# Inferential boundaries of age prediction: why prediction does not establish biological age measurement

**DOI:** 10.64898/2026.09.23.753690

**Authors:** Shiyu Song, Qiao Liang, Qian Gao

**Affiliations:** Nanjing Lupine Biotechnology; Medical School, Nanjing University

## Abstract

Chronological-age clocks reconstruct age from biological measurements, yet their outputs are interpreted as biological age, gaps as ageing acceleration and intervention-associated decreases as rejuvenation^1–5^. We show that age supervision identifies an age-task statistic, not a biological-age construct, and establish how this distinction changes biomarker construction and validation. Even at the population optimum, the same observable distribution and age-prediction performance admit incompatible biological-age interpretations. Resolving this ambiguity requires assumptions or evidence beyond the age task. Squared-error age loss penalizes within-age output dispersion without defining its biological direction. Given age and background, a gap re-expresses the compressed score; exact age recovery eliminates it even when heterogeneity remains in the measurements. Shared biological covariance permits genuine prognostic value without establishing construct identity. After allogeneic haematopoietic stem-cell transplantation, recipient-blood scores showed excess donor-lineage affiliation under a score-pairing null. In NHANES, age-trained scores improved held-out five-year mortality prediction beyond age and background, yet direct modelling of source measurements and mortality supervision at matched scalar capacity yielded further gains. The intended biological object must therefore guide study design, measurement selection and representation; validation must establish the claimed measurement relation rather than rely on age-prediction success alone.

## Main text

Chronological age is familiar, ordered and expressed in years. Age-trained molecular and clinical clocks reconstruct it from biological measurements and return a value on the same scale^1–5^. Subtracting known age produces an age gap. The common unit makes a further inference feel natural: the score becomes biological age, a positive gap becomes ageing acceleration, and an intervention-associated decrease becomes rejuvenation. These interpretations now shape research across molecular biology, clinical risk prediction and intervention evaluation (Table 1 and Supplementary Data 1)^3–5^. Yet sharing an age unit does not establish that the score measures the biological condition subsequently assigned to it.

**Table 1.**
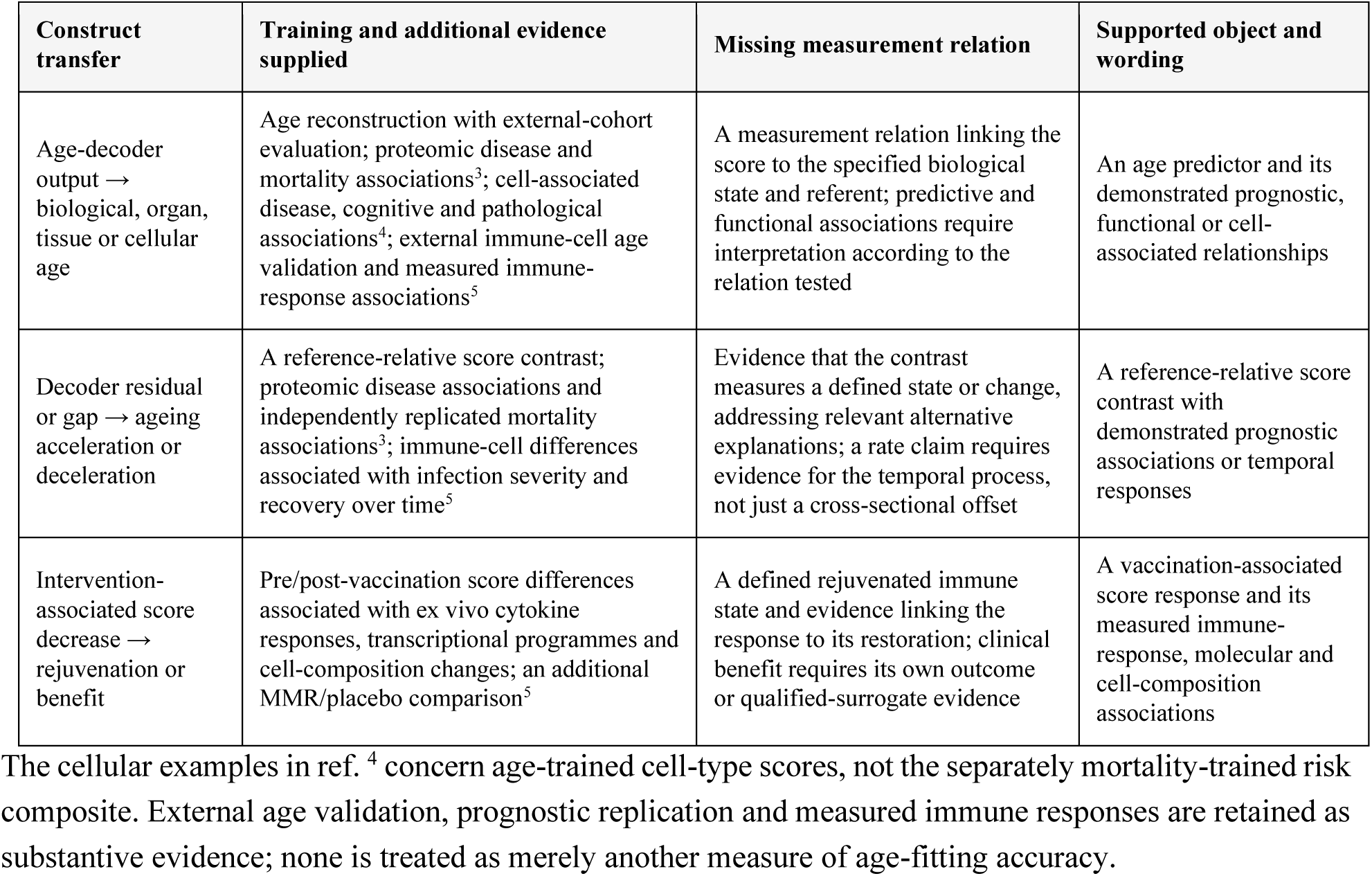
Contemporary construct transfer, available evidence and the missing identifying bridge. Rows locate three contemporary construct transfers and distinguish the evidence supplied by age training from the additional validation reported in the cited studies. These validations support the relations actually tested; the missing bridge concerns the further biological interpretation. Full source records and selection rules are provided in Supplementary Data 1.

| Construct transfer | Training and additional evidence supplied | Missing measurement relation | Supported object and wording |
| --- | --- | --- | --- |
| Age-decoder output → biological, organ, tissue or cellular age | Age reconstruction with external-cohort evaluation; proteomic disease and mortality associations<sup>3</sup>; cell-associated disease, cognitive and pathological associations<sup>4</sup>; external immune-cell age validation and measured immune-response associations<sup>5</sup> | A measurement relation linking the score to the specified biological state and referent; predictive and functional associations require interpretation according to the relation tested | An age predictor and its demonstrated prognostic, functional or cell-associated relationships |
| Decoder residual or gap → ageing acceleration or deceleration | A reference-relative score contrast; proteomic disease associations and independently replicated mortality associations<sup>3</sup>; immune-cell differences associated with infection severity and recovery over time<sup>5</sup> | Evidence that the contrast measures a defined state or change, addressing relevant alternative explanations; a rate claim requires evidence for the temporal process, not just a cross-sectional offset | A reference-relative score contrast with demonstrated prognostic associations or temporal responses |
| Intervention-associated score decrease → rejuvenation or benefit | Pre/post-vaccination score differences associated with ex vivo cytokine responses, transcriptional programmes and cell-composition changes; an additional MMR/placebo comparison<sup>5</sup> | A defined rejuvenated immune state and evidence linking the response to its restoration; clinical benefit requires its own outcome or qualified-surrogate evidence | A vaccination-associated score response and its measured immune-response, molecular and cell-composition associations |

The cellular examples in ref. <sup>4</sup> concern age-trained cell-type scores, not the separately mortality-trained risk composite. External age validation, prognostic replication and measured immune responses are retained as substantive evidence; none is treated as merely another measure of age-fitting accuracy.

Biological-age research seeks to resolve what chronological age leaves open: differences in physiological condition among people of the same age, changes in that condition over time and whether an intervention improves it. Age reconstruction asks a different question: which measurements, combined in what way, best predict chronological age? When its output is treated as biological age, age-prediction accuracy can become a criterion for selecting measurements, improving models and interpreting change. The inferential substitution occurs when success at the age-prediction task is accepted as evidence that the intended biological state has been measured.

Earlier work has questioned the construction and interpretation of clocks, established cross-sectional non-identifiability and challenged biological-age language^6–9^. Other studies have addressed calibration, uncertainty and validation^10–16^. Building on these contributions, we ask what chronological-age prediction can justify when it is used to select and validate a measure of biological ageing, including when the resulting score is accurate, reliable and genuinely prognostic.

Here we establish the consequences of using chronological-age prediction as the objective for developing an ageing measure. Age supervision identifies an age-task statistic, not the biological construct subsequently assigned to it; gap construction does not supply the missing measurement relation; and genuine prognostic value can arise through shared biological covariance without resolving construct identity. Human comparisons show that this distinction changes representation and measurement choice in practice. The resulting principle is constructive: the biological question, rather than age reconstruction, must determine the target, the representation and the evidence that counts as validation.

### Age supervision selects a statistic of chronological age

Let *A* denote chronological age, *W* background variables and *X* the measured profile (Fig. 1). Under squared-error loss, the unrestricted population-optimal age-prediction model, or decoder, is

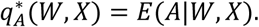

**Figure 1.**
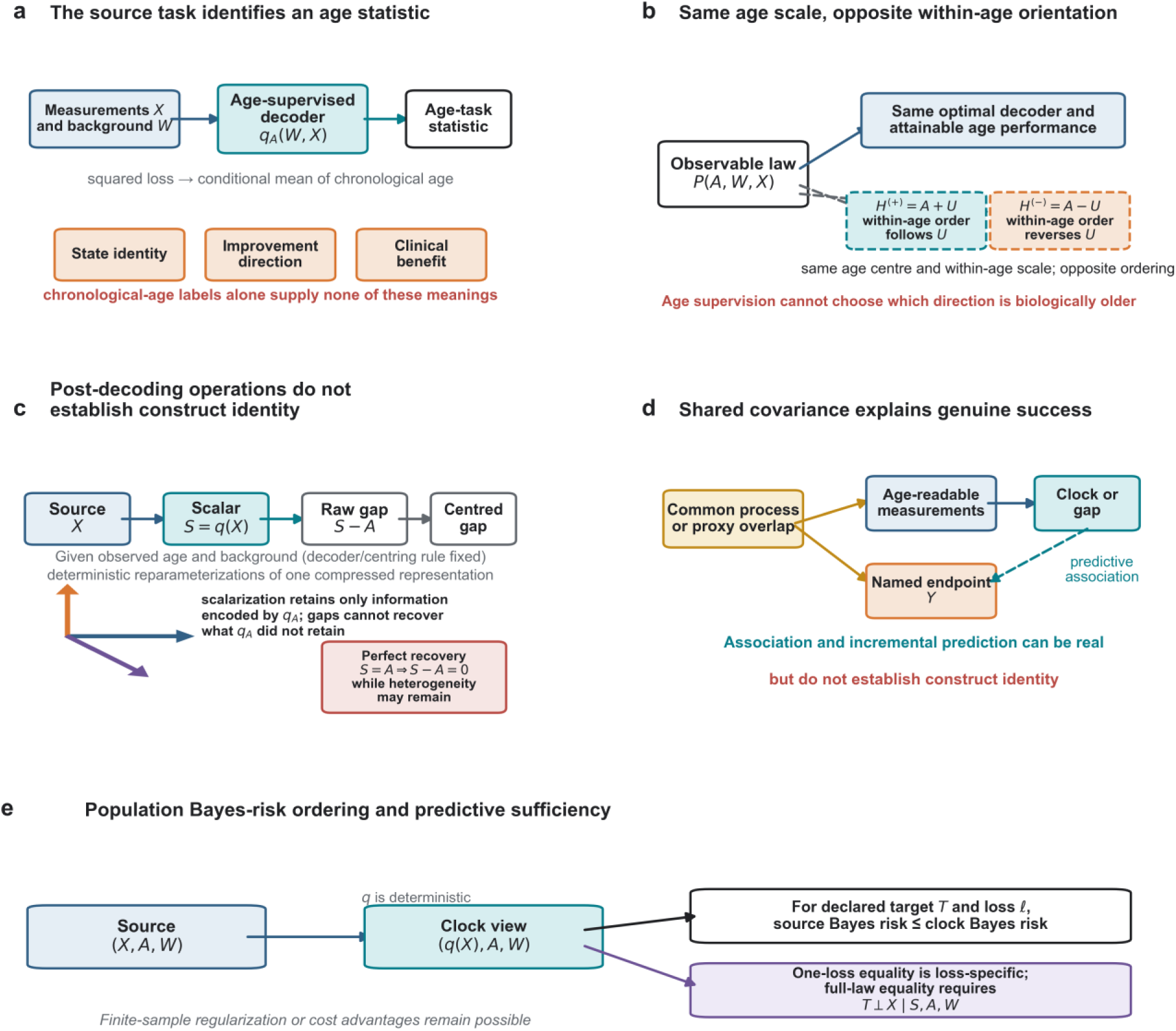
From age-task identification to genuine downstream success without construct identity a, Task–object mismatch. The supervised inverse task reconstructs ***A*** from ***X***, ***W***, whereas same-age state, future risk, longitudinal transition and intervention benefit require their own defining structures. b, Observable-law non-identification and within-age orientation. The same ***P***(***A***, ***W***, ***X***), population-optimal ***q_A_***(***W***, ***X***) and attainable age-prediction performance admit incompatible latent-state extensions; even age-centred, equal-scale candidates ***H***^(+)^ = ***A*** + ***U*** and ***H***^(−)^ = ***A*** − ***U*** can reverse the ordering of the same contrast. c, Post-decoding operations. For a fixed decoder and centring rule, the score, raw gap and conditionally centred gap are mutually recoverable given observed age and background; these operations cannot recover information discarded by the decoder. Perfect age recovery collapses the gap even when within-age heterogeneity remains. d, Shared covariance explains genuine downstream success: age-readable and endpoint-relevant directions can overlap, so a named association can be real without establishing construct identity. e, Source measurements cannot have greater population Bayes risk than their deterministic clock under the same declared target and loss. Equality for one loss is loss-specific; equality of the complete target law requires predictive sufficiency. Analytical results establish identification limits, human analyses establish materiality and endpoint-specific analyses motivate alternatives.

It returns the average chronological age among people with the same measured profile and background in the reference population. Within a restricted class, the population-optimal predictor gives the best squared-error approximation to this conditional mean; under absolute-error loss, the unrestricted optimum is a conditional median. The loss, model class and reference population therefore define an age-task statistic; they do not define what differences in that statistic mean biologically.

For same-age heterogeneity, the mismatch is visible in the loss itself:

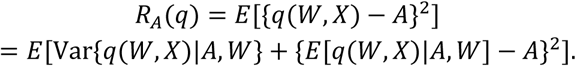

Within a fixed age-and-background stratum, the supervised target is the same for everyone. Dispersion in decoded age within that stratum therefore contributes directly to squared-error loss, whereas the second term measures whether the stratum mean is calibrated to chronological age. This is appropriate for age reconstruction. But the loss contains no label for whether a same-age difference reflects damage, reserve, adaptation, resilience or noise. Such output differences can remain because the same measurements can also improve reconstruction across age strata, but their biological orientation is not learned from the age objective. The loss acts on the scalar output, not on the existence of heterogeneity in *X*. Age-prediction accuracy therefore provides no criterion for deciding which same-age differences represent deterioration or improvement.

The loss decomposition describes how age supervision evaluates same-age output variation. A separate argument establishes the identification limit. Fix the complete observed distribution of age, background and measurements. The population-optimal age decoder and its attainable prediction performance are then fixed as well. Yet this same observable distribution is compatible with latent-state interpretations that support different biological-age values or same-age rankings. The age task therefore cannot determine which, if any, biological interpretation is correct. Building on the observational-equivalence construction of Sluiskes and colleagues^8^, Proposition 1 shows that this ambiguity persists at the population optimum, after finite-sample estimation error has disappeared.

Age anchoring does not resolve biological orientation. Let *U* be a measured contrast expressed in age units and centred within age and background. The two age-anchored quantities *A* + *U* and *A* − *U* have the same conditional centre, within-age variance and absolute departure from age, yet reverse every non-tied within-age ranking. Chronological-age information therefore cannot determine which direction, if either, corresponds to deterioration rather than improvement. That orientation must come from evidence outside the age task, such as an anchored measurement model or a separately defined endpoint.

### Age gaps preserve the score’s information and its identification limits

For a fixed decoder, the raw gap is *G*_raw_ = *q_A_* − *A*, so *q_A_* = *G*_raw_ + *A*. By definition, a positive raw gap means that the decoder assigns the measured profile an age-task value higher than the person’s observed chronological age; a negative gap means a lower one. The sign follows the orientation of the chronological-age coordinate and the chosen reporting convention, not an independently established biological direction. Likewise, a positive conditionally centred gap means that the score lies above the age-and background-specific reference mean.

Given chronological age, background and the centring rule, the score, raw gap and conditionally centred gap are one-to-one transformations of one another. Subtracting age or centring the score therefore re-expresses the fitted scalar; it adds neither a new observation nor a biological definition. Featurewise conditioning, *X* − E(*X*|*A*, *W*), is a different operation: it preserves feature-resolved same-age variation before scalar compression. Gap construction cannot recover profile distinctions that the decoder no longer represents. If those distinctions matter to the scientific question, they must remain represented and be evaluated directly.

The geometry of the decoder makes this compression explicit. For a linear decoder, profile differences in its null space leave the score unchanged. For a differentiable nonlinear decoder, changes orthogonal to the local gradient have no first-order effect on the score, although they may still affect it beyond first order (Supplementary Note 1).

Compression is only part of the problem: variation that remains in the gap has no privileged interpretation as biological-state information. Two limiting cases show both directions of the mismatch. Noisy age measurements can produce non-zero population-optimal gaps even when no additional state variable exists in the generating system. Conversely, a measured profile can contain chronological age exactly together with a separate state: age is then recovered perfectly and the gap is zero, while the state remains fully observed. Thus, non-zero gap variation does not establish state information, and a zero gap does not imply its absence from the source measurements. A frozen three-variable construction illustrates the same separation between age readability and state prediction (Extended Data Fig. 1).

The second limiting case extends to a general boundary. Under squared-error age prediction, the mean-squared raw gap equals the age-prediction risk. Subtracting the raw gap’s population conditional mean given age and background cannot increase its mean square in that population. As age-prediction error approaches zero, both gaps therefore collapse to zero in mean square, even though biologically relevant heterogeneity can remain in the source measurements. This is an amplitude result: it does not imply that endpoint information decreases monotonically as age accuracy improves. With imperfect prediction, an observed gap can reflect several statistically distinct sources, including conditional age ambiguity, reference-population differences, model restriction, estimation and calibration. Age-prediction accuracy alone cannot distinguish these contributions to an individual’s gap or establish what biological state the gap measures (Supplementary Note 1).

### Why age-trained clocks can predict outcomes beyond age without identifying biological age

Chronological age is itself informative about many health outcomes. Because age is already observed, the relevant question for an age-trained score is whether it contributes predictive information beyond chronological age and background. It can. Biological measurements contain age-associated variation arising from development, exposure, disease, adaptation, cell composition, cohort and selective survival. Some of this variation can also be associated with a named endpoint among people of the same chronological age and background. An age decoder can therefore retain variation that is useful both for reconstructing age across the population and for predicting an endpoint within age strata. We refer to this overlap as shared biological covariance. It provides a route to genuine incremental prognostic value without requiring the score to measure a biological-age construct.

A fixed clock is a deterministic compression of its source measurements and background. For any declared endpoint and loss, unrestricted population prediction using the source profile, age and background can achieve risk no greater than prediction using the clock, age and background^17–19^. This provides a direct benchmark for what the scalar compression preserves. Equality is possible when the scalar is predictively sufficient for that target and loss. Under log loss, this requires the source profile to add no endpoint information after conditioning on the clock, age and background. Chronological-age supervision neither imposes nor tests that condition. Finite-sample regularization, assay cost and transportability can still favour a scalar, but those are advantages to establish for its intended use rather than consequences of age reconstruction itself.

These distinctions separate three questions that are often conflated: what chronological age already predicts, what an age-trained score adds beyond age, and whether age reconstruction is the appropriate way to compress the source measurements for the intended task. None of these predictive comparisons, by itself, establishes the biological identity of the score. Validation must instead match the claim being made: prognosis, movement of a readout, restoration of a biological state and clinical benefit require different evidence^20^. We therefore use human data to examine the referent of an age-valued score, the consequences of scalar compression and the effect of changing the supervision target.

### Transplantation separates the candidate referents of a blood age score

Allogeneic haematopoietic stem-cell transplantation creates a natural referent split. The recipient remains the host organism, whereas the assayed circulating blood is produced largely by a donor-derived haematopoietic lineage (Fig. 2)^21,22^. A blood age score expressed in years can therefore be compared with two chronological coordinates: recipient age and donor age at transplantation plus elapsed follow-up. The latter indexes donor-lineage chronology, not the literal age of short-lived circulating cells.

**Figure 2.**
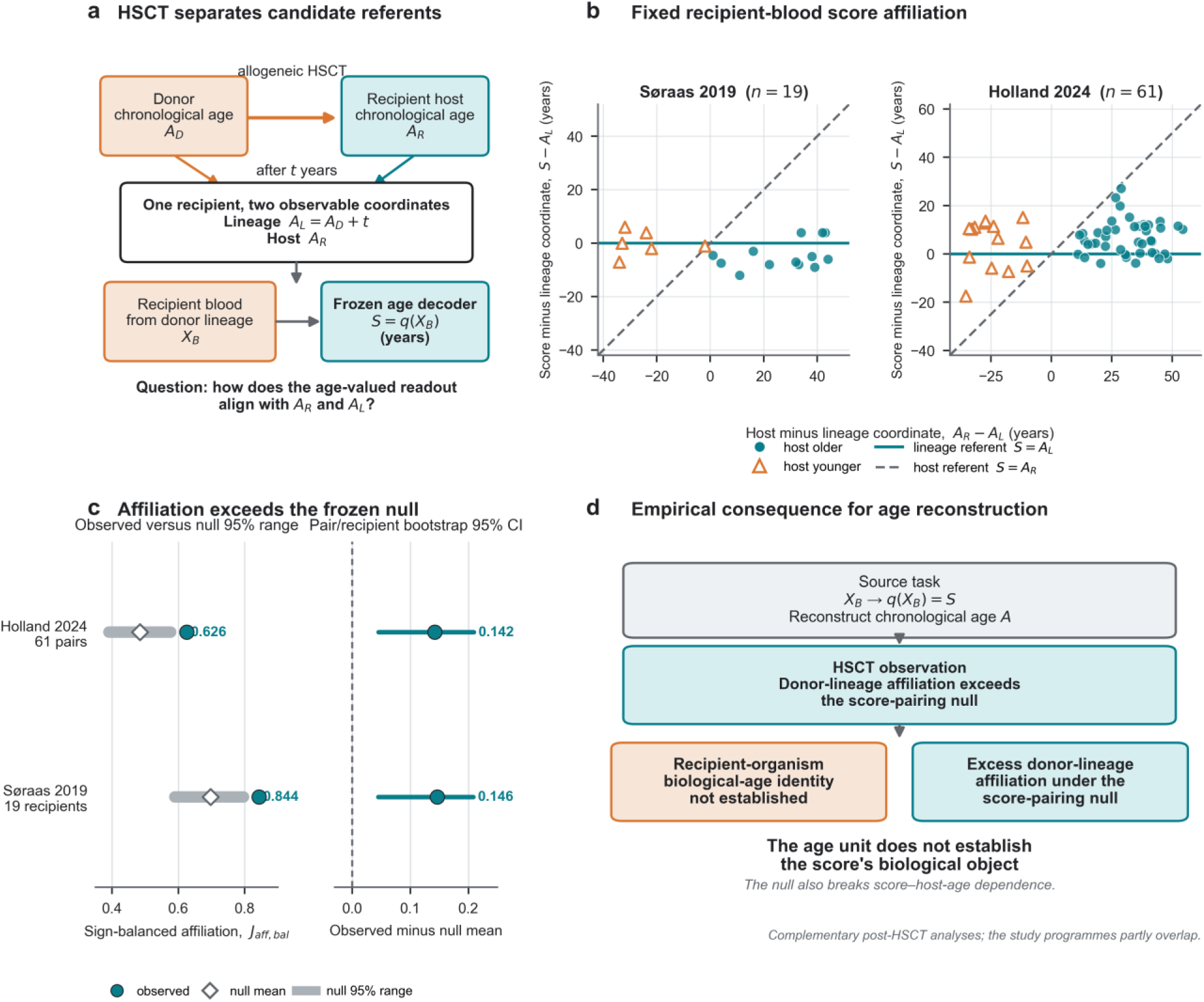
Haematopoietic stem-cell transplantation exposes referent ambiguity in an age-valued blood readout a, Allogeneic haematopoietic stem-cell transplantation separates recipient-host chronological age ***A_R_*** from the chronological coordinate of the sampled donor-derived haematopoietic lineage ***A_L_*** = ***A_D_*** + ***t***. A fixed recipient-blood score ***S*** = ***q***(***X_B_***) is expressed in years, but its numerical unit does not specify which referent it indexes. b, Recipient-or pair-level score geometry in Søraas 2019 and Holland 2024. The horizontal line denotes the lineage referent ***S*** = ***A_L_***; the dashed diagonal denotes the host referent ***S*** = ***A_R_***. Filled circles and open triangles distinguish host-older and host-younger directions. c, Sign-balanced affiliation ***J*_aff_**_,**bal**_: the affiliation index is evaluated after averaging scores and chronological coordinates within each pair (Holland) or recipient (Søraas), and the two age-difference directions are then equally weighted. These pair-summary indices, ***J*_pair_**, are computed before sign balancing and differ from the occasion-first indices underlying Extended Data Fig. 2a and the upper rows of 2b. Grey intervals show the frozen within-sign geometry-preserving score-pairing null, diamonds its mean and teal circles the observed statistic; the right subpanel shows observed-minus-null contrasts with stratified independent-unit bootstrap intervals. Holland provides the primary estimate and Søraas separately processed complementary evidence. The permutation also breaks the score–recipient-age relationship; it does not adjust for host-age-dependent shrinkage. d, The observed result is excess donor-lineage affiliation under that score-pairing null. Recipient-organism biological-age identity is not established, rather than empirically ruled out; lineage biological-age identity is also not established. Host influence remains compatible with this finding. The programmes partly overlap and are complementary rather than independent replications.

We tested whether recipient-blood scores showed greater donor-lineage affiliation than expected under a score-pairing null. We analysed the scores reported by Søraas and colleagues and applied fixed Horvath-2013 coefficients to the author-normalized Holland data using the declared external-mean imputation. The primary Holland analysis included 118 eligible occasions from 61 donor–recipient pairs. Scores and chronological coordinates were first averaged within each donor–recipient pair. An affiliation index quantified relative proximity to the two coordinates, with positive values indicating closer alignment with donor-lineage chronology. The two donor–recipient age-difference directions were weighted equally. Within each direction, the null held the donor–recipient age coordinates fixed and reassigned scores across pairs, preserving score marginals while breaking the observed score–referent pairing.

In Holland, observed affiliation was 0.626, exceeding the null mean of 0.484 and its 97.5th percentile of 0.577 (one-sided finite-permutation *P* = 8.50 × 10^−4^). The 95% bootstrap interval for observed-minus-null affiliation was 0.045–0.209, with positive contrasts in both age-difference directions. Separately processed Søraas scores showed the same direction. The programmes partly overlap and therefore provide complementary rather than independent evidence. The estimand is excess affiliation relative to a null in which the observed score–referent pairing is broken, not donor influence conditional on recipient age. Because the permutation also disrupts score–recipient-age dependence, it does not exclude host-age-dependent shrinkage.

The result makes the referent problem concrete. Recipient-blood scores showed structured affinity to donor-lineage chronology under the declared test. Neither chronological-age reconstruction nor expression in years establishes that the score measures the biological age of the recipient organism or the donor-derived lineage. Establishing either interpretation requires a measurement relation for the proposed referent.

### Mortality-directed modelling improves on useful age-trained scores

We next asked whether genuine prognostic value gives age reconstruction any privileged role as an intermediate representation. In continuous NHANES, we compared an age-trained score with direct modelling of its source measurements for held-out five-year mortality prediction (Fig. 3a,b). Among 13,724 participants aged 20–79 years, including 470 five-year deaths, a ridge model reconstructed age from background and 50 clinical measurements, with chronological age as its only supervised label. All mortality models included observed age and background. We compared this baseline with models adding either the cross-fitted age score or all 50 source measurements. Every comparison used the same held-out participants.

**Figure 3.**
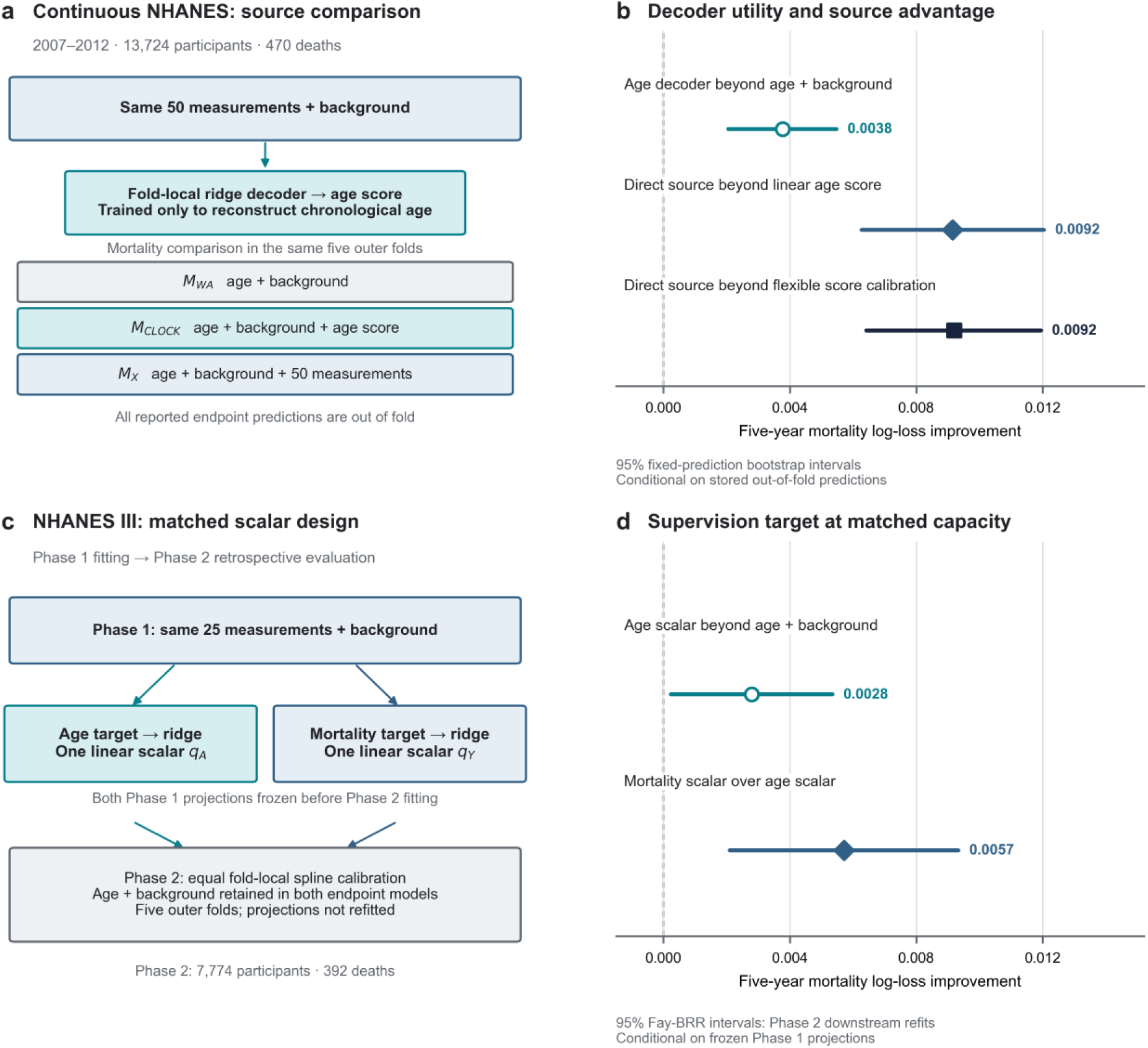
Mortality prediction from source measurements and matched scalar representations a, Nested comparison in continuous NHANES 2007–2012, using 13,724 participants with resolved five-year mortality status, including 470 deaths. A fold-local ridge decoder reconstructed chronological age from background and 50 measurements. Mortality models used age and background alone, added the cross-fitted age score, or directly modelled the same source measurements. b, Paired held-out log-loss contrasts show the clock’s gain beyond age and background and the further direct-source gain over linear and prespecified flexible clock calibration. Lines are 95% bootstrap intervals conditional on stored out-of-fold predictions. c, Separate NHANES III matched-scalar design. Age-and mortality-supervised linear projections used the same 25 measurements and background in Phase 1 (7,558 participants; 402 deaths). The fitted directions were frozen before application to Phase 2 (7,774 participants; 392 deaths), where they received identical fold-local endpoint calibration and held-out evaluation. Phase 2 outcomes did not enter projection fitting. d, Design-weighted Phase 2 log-loss contrasts. ***U_A_*** is the gain from the age-supervised scalar beyond observed age and background; ***Δ*** is the further gain from mortality supervision at matched scalar capacity. Lines are 95% Fay-BRR intervals incorporating Phase 2 downstream refitting conditional on the frozen Phase 1 directions. Positive values favour the representation named in each contrast. The NHANES III analysis is a retrospective, outcome-accessed protocol transfer, not independent confirmation. The separate age-substitution and measurement-removal comparisons are retained in Extended Data Fig. 3. Lower log loss means better probabilistic prediction, not clinical benefit or biological-age identification.

The age-trained score reduced mortality log loss by 0.003774 beyond age and background (95% fixed-prediction bootstrap interval, 0.002046–0.005474). Direct modelling of the same 50 measurements reduced log loss by a further 0.009152 compared with the clock model (0.006276–0.012034). At the point estimates, the additional improvement from the source measurements was more than twice the clock’s entire gain beyond age and background. Both contrasts had the same direction in all five outer folds, and prespecified nonlinear recalibration of the clock left the direct-source advantage essentially unchanged.

That comparison does not by itself isolate the effect of the supervision target: the source-measurement model has access to the full measurement profile rather than a single age-trained scalar. To test the supervision target at fixed scalar dimension, we used a distinct NHANES III design with two one-dimensional linear projections from the same 25-measurement panel (Fig. 3c,d). Both were trained in Phase 1 under otherwise matched conditions: one reconstructed chronological age and the other predicted five-year mortality. Their fitted directions were frozen and given identical downstream calibration opportunities in survey-weighted Phase 2 data comprising 7,774 participants and 392 deaths. Phase 2 outcomes did not enter projection fitting. The two projections therefore differed in supervision target, not source measurements, scalar dimension, linear capacity or downstream calibration opportunity.

Lower log loss indicates better probabilistic prediction. The age-supervised scalar remained incrementally predictive of mortality: adding it beyond observed age and background reduced Phase 2 log loss by 0.002792 (95% Fay-BRR interval, 0.000239–0.005345). Yet, with the same scalar capacity, mortality supervision reduced log loss by a further 0.005711 relative to the age-supervised representation (0.002093– 0.009329), with positive contrasts in all five outer folds. At the point estimates, the additional gain from mortality supervision was approximately twice the gain provided by adding the age-trained scalar itself.

Both analyses began with genuinely useful age-trained scores. Direct modelling of the source measurements nevertheless improved held-out mortality prediction, and mortality supervision yielded further gains at matched scalar capacity. The direct-outcome comparison was raised explicitly by Kriukov and colleagues^6^; our analyses make this comparison concrete through complementary source-profile and matched-scalar designs. The continuous-NHANES intervals condition on stored out-of-fold predictions. NHANES III is a retrospective protocol transfer with previously accessed outcomes; its Fay-BRR intervals incorporate Phase 2 downstream refitting conditional on frozen Phase 1 directions. Within the declared populations, measurements and model families, these results show that genuine prognostic value does not by itself justify choosing age reconstruction as the intermediate task.

### Similar AgeGaps can arise from different measurement profiles

The endpoint comparisons assess predictive performance for a named outcome. We next asked whether nearly identical age gaps corresponded to similar measurement-contribution profiles. Using the frozen continuous-NHANES age decoders, we decomposed each conditionally centred AgeGap into 50 exactly additive measurement contributions. We then formed 293 outcome-blind, non-reusing pairs matched exactly on fold, chronological age, sex, race or ethnicity and survey cycle, with conditionally centred gaps differing by no more than 0.25 decoder-output years (Extended Data Fig. 4).

The median absolute gap difference was 0.124 decoder-output years, whereas the median Euclidean distance between contribution profiles was 24.602 decoder-output years and median cosine similarity was 0.003. The distance is defined in the fitted decoder’s contribution space and is not an age difference between participants. Near-identical summed outputs therefore arose from markedly different signed measurement contributions. The scalar fixed only the sum of these contributions, not their configuration. These contributions exactly attribute the output of the frozen decoder under its specified centring and parameterization, but they are not causal components of ageing or components of a biological-age construct.

### The scientific target guides measurement and model selection

Target choice affects not only how measurements are compressed, but also which measurements are selected in the first place. Across five prespecified measurement-domain options in NHANES 2007–2008, mortality-directed selection chose hepatic, protein and injury measurements, whereas an outcome-blind inverse age decoder chose metabolic, adiposity and lipid measurements. We froze both choices and evaluated them in NHANES 2009–2010 (Fig. 4b; Extended Data Fig. 5). The endpoint-selected domain reduced five-year mortality log loss beyond age and background by 0.006418 (95% fixed-prediction bootstrap interval, 0.002297–0.010652), compared with 0.000646 (−0.001195–0.002481) for the age-selected domain. Their paired difference was 0.005772 (0.001380–0.010390), positive in four of five folds.

**Figure 4.**
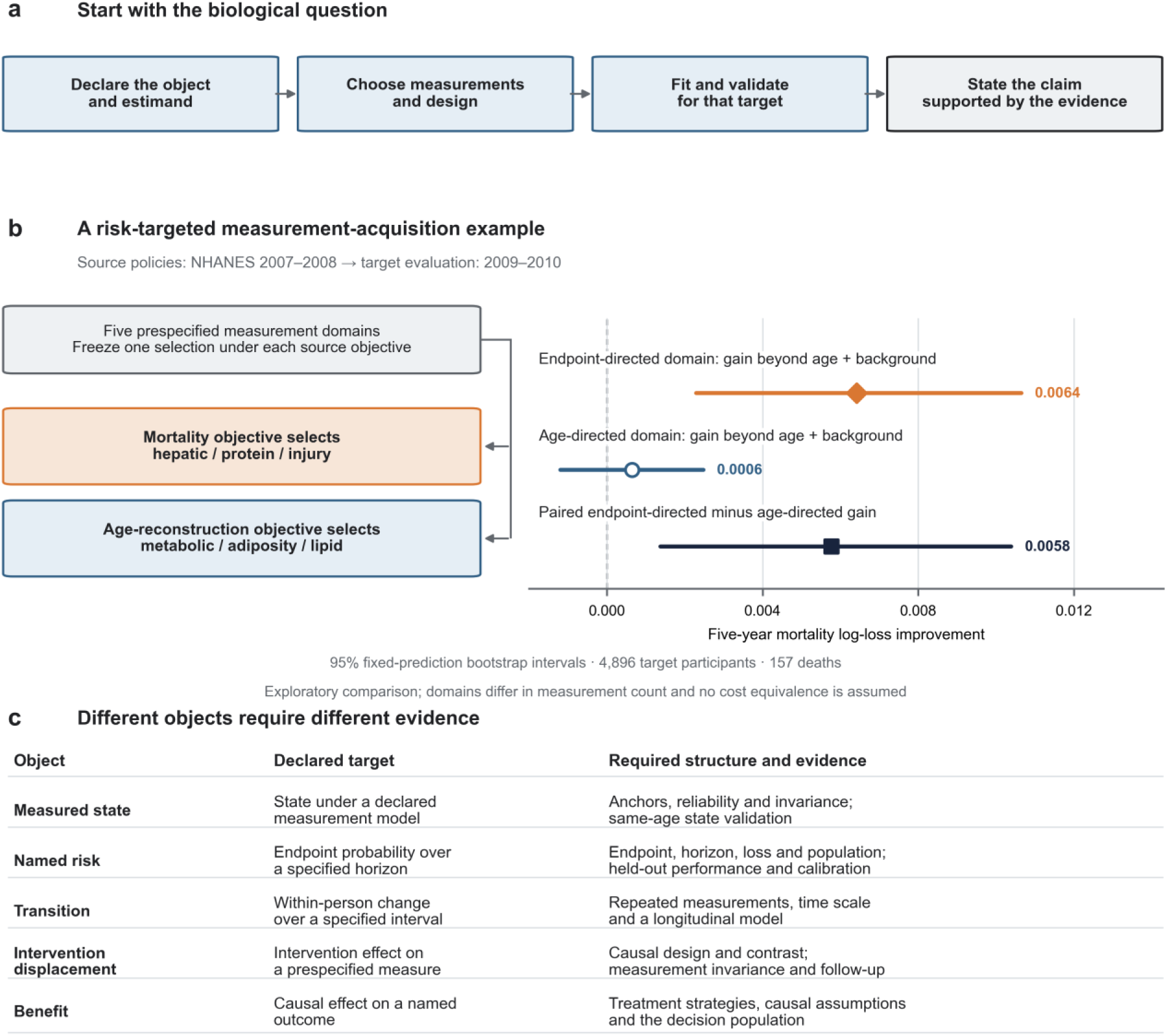
Target-first evidence replaces construct transfer from an age unit a, A target-first programme begins with the scientific question and declared object, then chooses measurements, design and supervision, evaluates the resulting statistic and makes the claim supported by that evidence. b, Exploratory acquisition comparison for five-year all-cause mortality. In NHANES 2007–2008 (4,599 participants; 174 deaths), endpoint-directed selection chose hepatic, protein and injury measurements from five prespecified domains, whereas inverse age decoding chose metabolic, adiposity and lipid measurements. In NHANES 2009– 2010 (4,896 participants; 157 deaths), their held-out log-loss gains beyond age and background were 0.006418 and 0.000646; the paired difference was 0.005772. Lines are 95% intervals from 5,000 event-stratified participant bootstraps conditional on fixed source selections and stored predictions. Domains differed in measurement count and were not matched for acquisition cost. The complete policy comparison is retained in Extended Data Fig. 5. c, Object-specific evidence map. A measured or latent state requires an anchored measurement model and state definition; named risk requires an endpoint, horizon, loss and validation population; transition requires longitudinal sampling and a transition model; intervention displacement requires a prespecified perturbation contrast and measurement response; and benefit requires a causal estimand, named outcome and decision context. The acquisition example illustrates the named-risk row, not empirical validation of every object in the map. Age-equivalent units may re-express a separately established quantity; the defining evidence supplies its meaning.

The two objectives therefore selected different measurements from the same prespecified menu, and the mortality-directed choice performed better for the mortality endpoint in the target cycle. This exploratory cross-cycle comparison involved domains with different numbers of measurements and did not assume equivalent acquisition costs. It illustrates how the scientific target can guide measurement selection as well as the representation subsequently constructed from those measurements.

A target-first programme therefore begins with the object to be measured and the decision it is intended to support (Fig. 4a,c). Different questions require different evidence. A same-age state requires a defined measurement model; named risk requires an endpoint, horizon and loss; transition requires repeated observations and a dynamic estimand; intervention displacement requires a causal contrast on a specified measurement; and clinical benefit requires a causal comparison on a clinical outcome or a surrogate qualified for the intended use. The measurement-selection comparison illustrates this programme for a named-risk task. Across these tasks, the intended object determines which differences matter, how performance should be evaluated and what evidence establishes progress.

## Discussion

The methodological error begins when success at reconstructing chronological age is treated as sufficient evidence that a representation measures a biological state. Biological-age measurement seeks to resolve differences that chronological age leaves open. Yet pure age-supervised clocks are constructed by optimizing prediction of that age label. This creates a tension between the reason for seeking a measure and the criterion used to develop it. Our analyses establish what the age objective supplies: a statistic for reconstructing age, not a relation defining the biological state it measures. Population-optimal prediction, gap construction and genuine prognostic value do not, by themselves, supply that missing relation. Progress in ageing measurement therefore requires evidence for the proposed biological object, rather than age-prediction accuracy as its substitute.

This argument builds on direct identification precedents and critiques of clock interpretation. Sluiskes and colleagues established observational non-identifiability and examined age-based marker selection^8^, while Ikram challenged prediction-error interpretation and clocks as disease explanations^7^. Kriukov and colleagues advocated direct outcome modelling and prospective target definition^6^; Johnson and Shokhirev addressed biological-age terminology^9^. Grødem and colleagues analysed the mismatch between age prediction and individual brain change and proposed reorienting prediction targets^23^. Hu’s House of Clocks demonstrated that different feature profiles can yield the same composite score and argued for prediction benchmarks using the source measurements^24^. Detailed comparisons are provided in Supplementary Note 2, with the structured review reported in Supplementary Table 1 and Supplementary Data 2. Here, we develop the consequences of these identification limits for measurement construction and validation. The analytical results distinguish what age supervision identifies, what gap operations preserve and what prognostic success can establish. The matched-scalar comparison evaluates supervision choice while holding source measurements, linear capacity and calibration opportunity fixed. Together, these results make the choice of training target part of the justification for a biological measurement, rather than merely a modelling decision.

The human comparisons make these consequences concrete. Transplantation separates the recipient organism and assayed lineage as candidate referents of an age-valued blood score. Near-identical gaps arise from different measurement-contribution profiles. Mortality-directed modelling improves on genuinely useful age-trained predictors, including at matched scalar capacity, while endpoint-directed selection changes which measurements are chosen. Shared biological covariance provides a route to the prognostic success of age-trained scores without requiring them first to measure a biological-age construct. Their usefulness therefore motivates evaluation for the intended task; it does not settle whether age reconstruction should be the intermediate objective.

The contemporary literature audit shows why this distinction remains consequential. Age-trained proteomic, cell-type and immune-age studies interpret age-valued scores, gaps and intervention-associated shifts as biological age, ageing acceleration or rejuvenation (Table 1 and Supplementary Data 1)^3–5^. These studies include external age validation, replicated prognostic associations and measured functional responses. The unresolved step is whether that evidence establishes the particular biological state or restoration claimed. The issue is therefore not an absence of validation, but what the validation establishes.

Field-scale studies report clock-specific disease associations, heterogeneous social gradients and discordant age-valued estimates^25–29^, while longitudinal and intervention studies establish prediction and response in particular settings^30–33^. These findings support the relations tested and can contribute to construct validation through an explicit measurement model. Their interpretation also shapes the phenotypes researchers investigate and the responses they regard as improvement. Our measurement-selection comparison demonstrates how target choice changes which measurements are prioritized for a named-risk task. More broadly, when age-valued scores guide clinical decisions or intervention development, movement of a readout and improvement in a defined state become different criteria of success. A claim of rejuvenation requires a defined biological state and evidence of its restoration; demonstrated functional or clinical benefit stands on its own outcome evidence.

Target-directed models already provide constructive starting points. Morbidity, mortality, lifespan and physiological-dynamics supervision select predictors for those targets^34–40^. These objectives differ from pure age reconstruction and must be assessed according to their own targets and evidence. Anchored deterioration models, causal constraints and compartment-specific perturbations can support biological constructs through added structure^41–43^. Such evidence can also validate a previously age-trained score for a defined object; its measurement validity then comes from that evidence. The population-optimal argument separates this identifying work from improvements in estimation, calibration or reliability. Chronological-age estimation remains a valid task, and fixed scores can have practical advantages in finite samples, cost or transportability. The human comparisons are retrospective evaluations of specified targets, measurement panels and estimators, not a universal ranking of age-and endpoint-directed models. They do not settle the existence of a low-dimensional ageing state.

A change of name leaves the age objective intact; a change of target changes the basis for measurement selection, model construction and validation. Chronological-age decoding does not identify biological age. Age-equivalent units can express a separately established quantity, but the defining evidence supplies its meaning. Measuring ageing requires the biological question, rather than the reconstruction of time, to define the target and the evidence that counts as success.

## Supporting information

Supplementary_Software_1

Supplementary_Software_1_verification_data

---

**Extended Data Figure 1.**
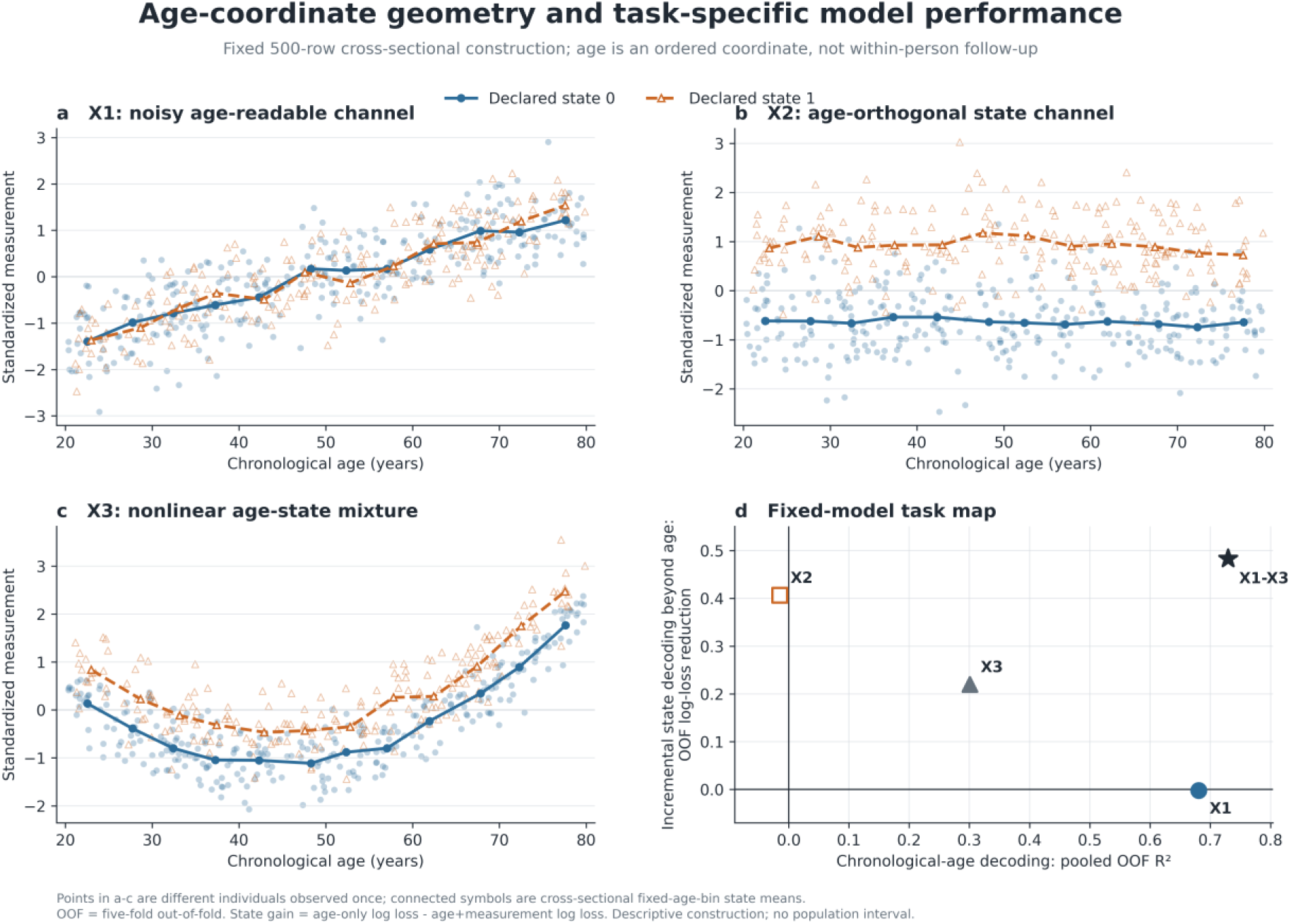
Age-coordinate geometry and task-specific model performance a–c, Frozen cross-sectional measurements X1–X3 plotted against chronological age. Each point is a different individual observed once; connected symbols are state-specific means within fixed age bins and are not longitudinal trajectories. X1 is a noisy age-readable channel without a designed state term; X2 is the declared state channel after exact projection from the realized intercept and linear age direction; X3 mixes linear and nonlinear age structure with state. d, Fixed five-fold out-of-fold task map. The horizontal axis is pooled chronological-age decoding ***R*^2^**. The vertical axis is the reduction in pooled state-decoding log loss obtained by adding the indicated measurement to the fixed age-only model. The display distinguishes age readability, same-age state value and their mixture under the declared models; it is explanatory rather than population inference, longitudinal evidence or a second proof of the analytical result.

**Extended Data Figure 2.**
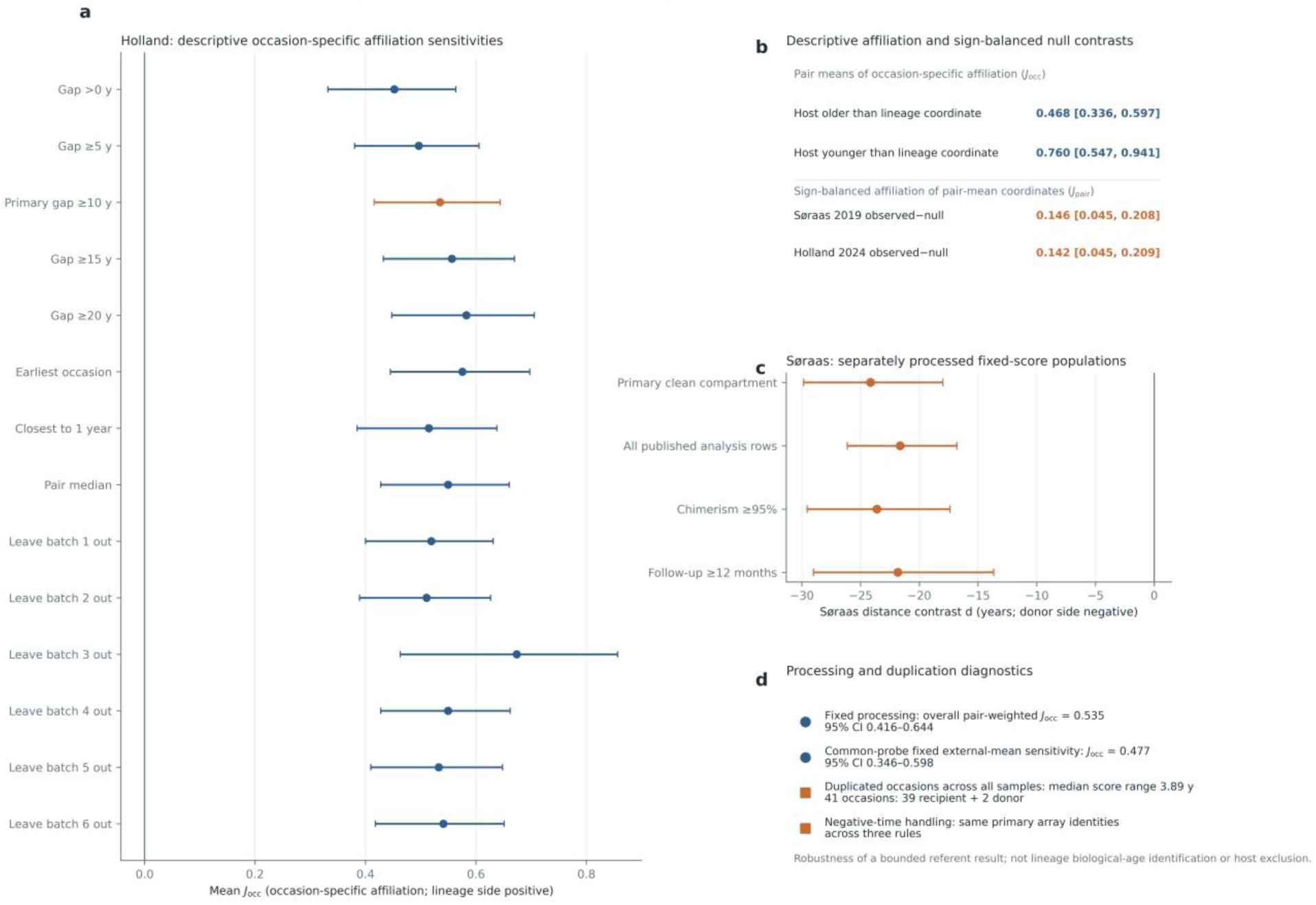
HSCT referent-null sensitivities preserve the interpretation boundary a, Holland occasion-first affiliation, ***J*_occ_**, across age-difference thresholds, follow-up selections and leave-one-batch-out analyses. Occasion indices are averaged within pair and then across pairs, except for the explicitly labelled within-pair median sensitivity. b, The upper rows show occasion-first host-older and host-younger means; the lower rows show the sign-balanced pair-summary observed-minus-null contrasts used in Fig. 2c. Aggregation order as well as final stratum weighting therefore differs between these summaries. c, Søraas clean-compartment and prespecified sensitivity populations after separate fixed-score processing. d, Primary and common-probe processing use the occasion-first pair-weighted statistic. The duplicated-occasion diagnostic covers all 41 duplicated donor or recipient occasions before primary-population screening (39 recipient and two donor occasions), not only the eligible recipient set. These analyses test robustness of the bounded referent result and do not identify lineage biological age or exclude host influence.

**Extended Data Figure 3.**
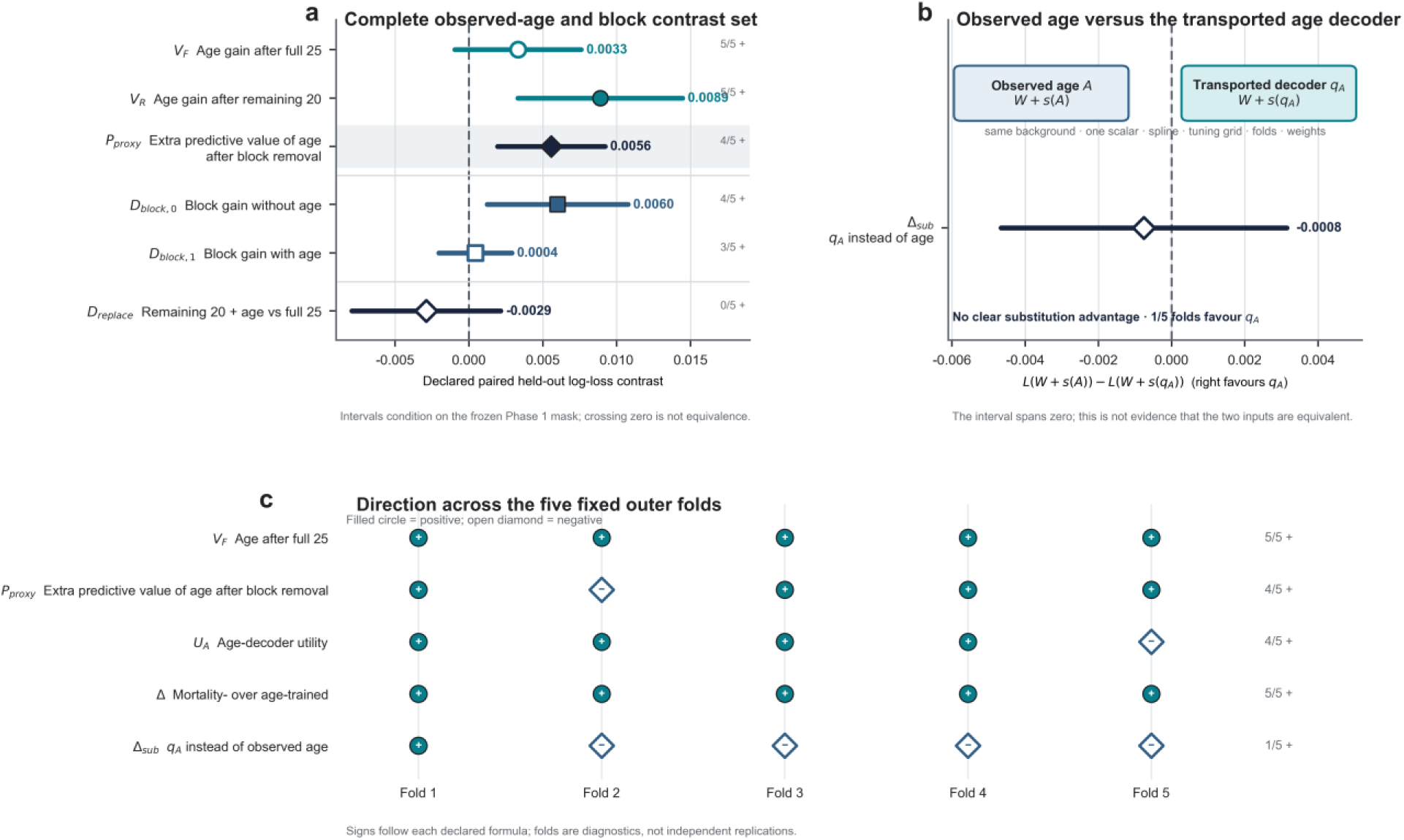
NHANES age-context, substitution and fold diagnostics a, Complete NHANES III Phase 2 contrast set for the four frozen endpoint models using all 25 measurements or the remaining 20 after the Phase 1 age-selected five-measurement block was withheld, without or with observed chronological age. Points are design-weighted paired held-out log-loss contrasts and lines are 95% Fay-BRR intervals. ***V_F_*** and ***V_R_*** are the increments from adding observed age; ***P*_proxy_** is their difference; ***D*_block_**_,**0**_ and ***D*_block_**_,**1**_ are the block contributions without and with age; ***D*_replace_** compares remaining 20 plus age with the full 25 without age. The intervals condition on the frozen Phase 1 mask. No assertion is made that the remaining 20 lack age-related information, and an interval spanning zero is not equivalence. b, Direct substitution boundary between observed age and the transported Phase 1 age decoder under the same background, one-scalar spline family, tuning grid, folds and weights. ***Δ*_sub_** was −0.000757 (95% Fay-BRR interval, −0.004661 to 0.003147), with one of five folds positive. There was no clear substitution advantage; this does not establish equality. c, Direction of five decision-facing paired contrasts in the five fixed primary-sampling-unit-atomic outer folds. Filled circles denote positive and open diamonds negative values under each declared formula. Fold directions are consistency diagnostics, not independent replications. Model-persistence and reload-parity records are retained in the reproducibility manifest rather than displayed as a scientific panel.

**Extended Data Figure 4.**
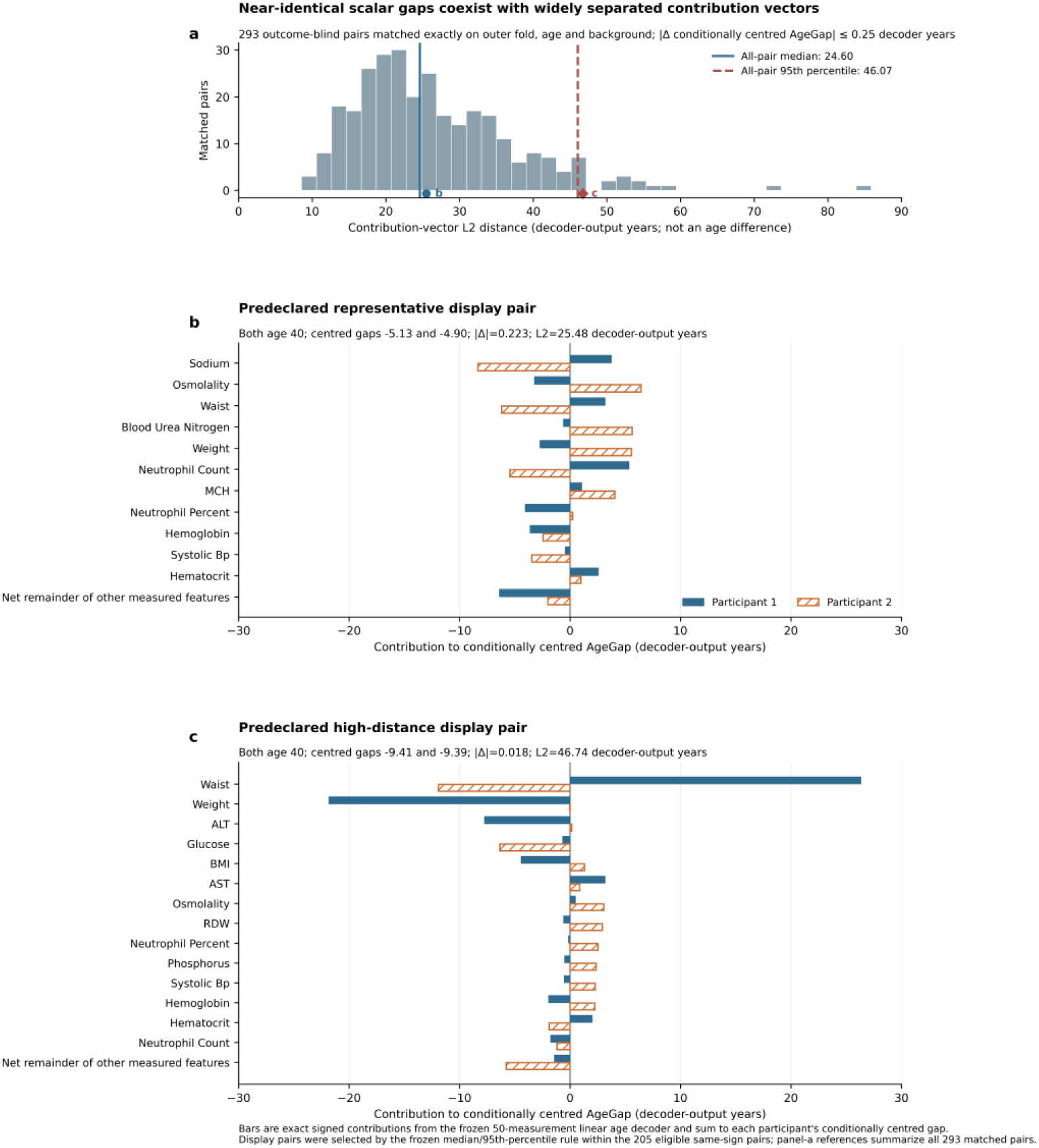
Near-identical conditionally centred AgeGaps arise from distinct measured-profile contributions a, Distribution of Euclidean distances between the 50-dimensional contribution vectors of 293 non-reusing, outcome-blind pairs matched exactly on outer fold, chronological age, sex, race or ethnicity and survey cycle and constrained to an absolute conditionally centred AgeGap difference of at most 0.25 decoder-output years. The solid and dashed lines mark the median (24.602) and 95th percentile (46.067) among all 293 pairs. The horizontal scale is distance in decoder-output contribution space, not an age difference between participants. b, Predeclared representative pair P00026, selected closest to the median contribution distance among the 205 display-eligible same-sign pairs with mean absolute centred AgeGap of at least two years. The centred deviations were −5.13 and −4.90 decoder-output years (absolute difference, 0.223); contribution-vector distance was 25.48 decoder-output years. c, Predeclared high-distance pair P00092, selected closest to the 95th percentile within the same display-eligible set. The centred deviations were −9.41 and −9.39 decoder-output years (absolute difference, 0.018); contribution-vector distance was 46.74 decoder-output years. In b and c, bars show signed contributions for the union of each participant’s eight largest absolute contributors; all remaining features form the exact net remainder, and displayed contributions sum to each participant’s conditionally centred AgeGap. Contributions attribute the output of the frozen fold-specific linear decoder after fold-local age and background centring; they are not causal, organ-specific or biological-age decompositions. This descriptive analysis uses the same unweighted complete-case interface as Fig. 3a,b, and attribution among correlated measurements depends on the frozen ridge parameterization.

**Extended Data Figure 5.**
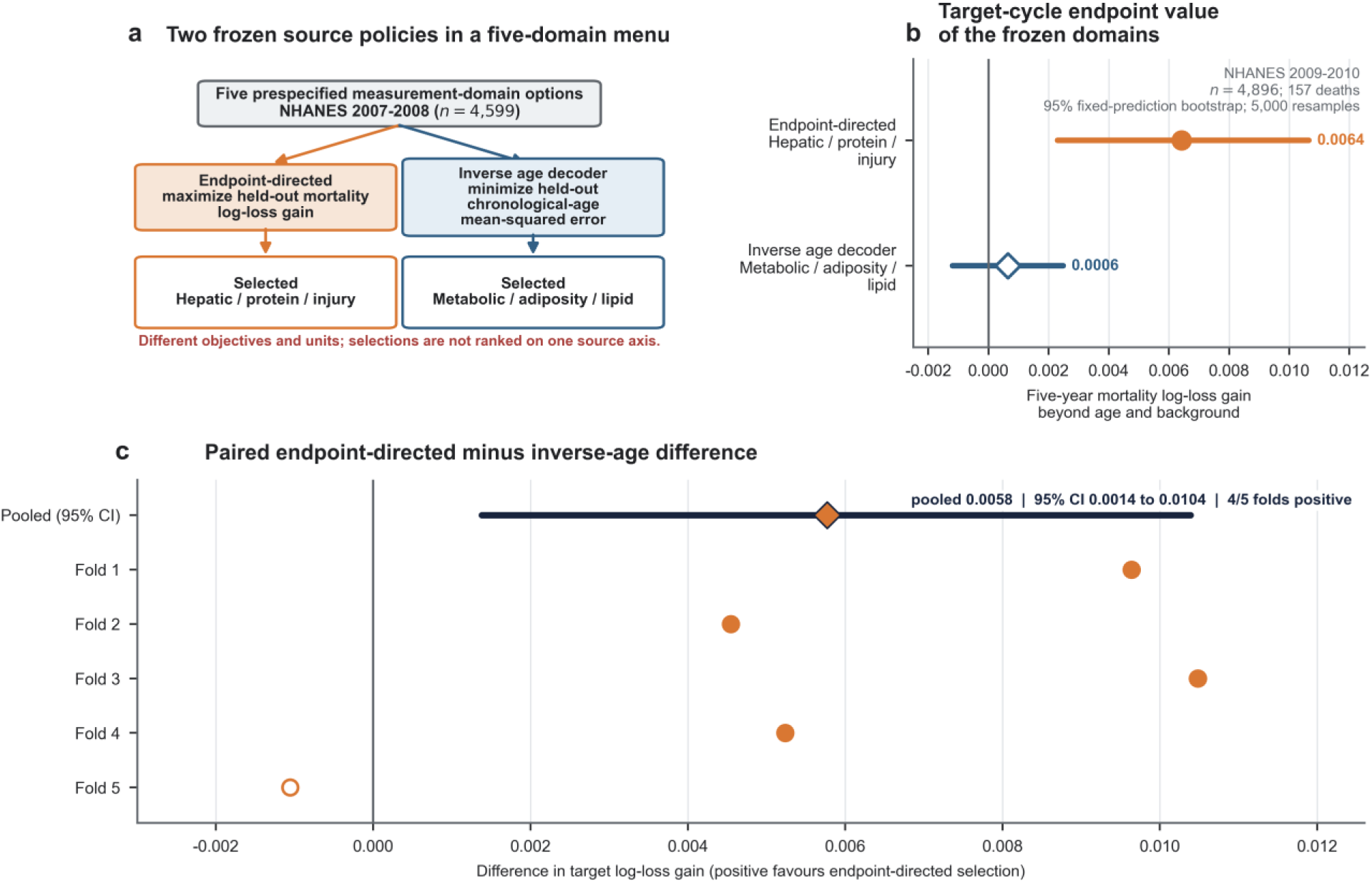
Endpoint-directed acquisition across five prespecified measurement-domain options a, Two policies were frozen in NHANES 2007–2008 from five prespecified measurement-domain options. The endpoint-directed policy selected hepatic, protein and injury measurements by maximizing held-out five-year mortality log-loss gain. The inverse chronological-age decoder selected metabolic, adiposity and lipid measurements by minimizing held-out chronological-age mean-squared error. The source criteria have different targets and units and are therefore shown as separate routes. b, In NHANES 2009–2010, the frozen endpoint-directed and inverse-age domains yielded log-loss gains beyond age and background of 0.006418 (95% fixed-prediction bootstrap interval, 0.002297–0.010652) and 0.000646 (−0.001195–0.002481), respectively. c, The paired endpoint-directed-minus-inverse-age difference was 0.005772 (0.001380–0.010390); four of five target-fold estimates were positive. The comparison is exploratory cross-cycle evidence within one declared five-domain menu. Each domain was treated as one acquisition option; domains contained different numbers of measurements and no equivalence of monetary resources or real-world costs was assumed. Intervals use 5,000 event-stratified participant bootstraps conditional on stored predictions and fixed source selections; they omit refitting and study-development uncertainty. The analysis is not clinical value of information, cost analysis, untouched confirmation, universal transportability or biological-age identification.

## Methods

### Study design

The study assigned distinct roles to analytical proof, empirical consequences and object-specific alternatives. The analytical sequence comprised loss-specific age-task identification, observable-law non-identification, meaning non-inheritance, post-decoding operator limits, a shared-covariance account of downstream success and population risk ordering under deterministic compression. Two literature modules served separate purposes: a contemporary construct-promotion audit documented current examples, whereas a structured review organized prior criticisms, proposed repairs and their identification ceilings. Neither literature module contributed to the proof. Haematopoietic stem-cell transplantation (HSCT) quantified donor-lineage affiliation of a fixed age-valued blood score relative to a score-pairing null preserving score marginals and donor–recipient age geometry. Continuous NHANES tested whether an age-supervised decoder could improve held-out mortality prediction while direct modelling of the same 50 source measurements performed better, and whether near-identical gaps from that decoder specified a unique measured-profile realization. NHANES III then held scalar dimension and model opportunity fixed while changing the training target from age to mortality, compared the age decoder with observed age itself, and tested how observed age’s predictive increment changed when a frozen age-selected measurement block was withheld. The estimand-specific replacement map supplied the constructive programme; endpoint-directed acquisition across five prespecified domains supplied an exploratory cross-cycle illustration.

A frozen X1–X3 subset of the earlier geometry package was retained as a transparent construction separating noisy age readability, linearly age-orthogonal declared state and their nonlinear mixture. Fixed five-fold out-of-fold models quantified chronological-age readability and incremental prediction of the generated state beyond age. The construction was not analysed as a human effect, repeated-simulation estimate, longitudinal trajectory or additional proof.

The contemporary examples in Table 1 were selected to represent distinct target and use classes, not to estimate the prevalence of any terminology or practice. The table is not a systematic review or a representative sample of all clocks. Its role is to establish that the target–interpretation promotions addressed by the analytical result remain current.

The central non-identification claim and meaning non-inheritance principle are analytical results. The X1– X3 construction and human analyses carry no proof responsibility for either result. The HSCT analyses were focused post-publication estimand audits rather than prospective preregistrations. All NHANES modules were retrospective and predictive or descriptive. For the continuous-NHANES pure-decoder comparison, the endpoint, population, panel, folds and model families were recorded in a versioned specification; the reported result was accepted without a confirmation refit or subsequent model, endpoint, panel or threshold search. The individual contribution analysis reused that frozen decoder, specified fold-local centring and outcome-blind matching before inspection, and introduced no decoder refit or endpoint-informed pair selection. The NHANES III modules were outcome-accessed fixed-protocol transfers from Phase 1 to Phase 2, not untouched confirmation or independent external validation. Phase 1 projections or measurement masks were frozen before the reported Phase 2 fitting and were not reselected. The cross-cycle acquisition rule was frozen in its source cycle before target-cycle evaluation, but the target cycle had been viewed during earlier reconnaissance and was classified as exploratory cross-cycle validation.

### Contemporary literature audits

The age-unit construct-promotion audit assembled source-verified contemporary examples from major general, medical, ageing and cell-biology journals. A paper was eligible as a main example when it supplied an individual-level predicted age, age gap or intervention-associated age-valued displacement; promoted that output to a biological, organ, cellular, immune or molecular ageing construct, rate or rejuvenation claim; and did not supply congruent named evidence for the promoted construct at the same conceptual level in its estimand, training target or validation. Boundary examples included clock applications constrained by a named endpoint, direct mechanism, longitudinal pace target or peripheral role; problem-diagnosing examples explicitly exposed clock heterogeneity or ambiguity. Downstream disease, mortality or frailty associations could establish narrower predictive uses but did not, by themselves, identify a broad latent ageing construct. Selection was designed to establish the existence and forms of the inferential operation, not a prevalence estimate, journal comparison or census. Supplementary Data 1 reports article title, journal and the relevant construct class.

The separate critical-literature review used local-first retrieval for 17 core critiques or methods papers published from 2024 through 2026 and three earlier context anchors, with targeted online supplementation where a local full text was unavailable. For each source we coded the problem targeted, the authors’ proposed repair, remedy family, earliest transition in the promotion chain, the narrow object supportable if the repair succeeded and the interpretation that still did not follow. The five transitions were features to age score, age score to gap or state, gap to faster or accelerated ageing, acceleration to intervention response, and response to rejuvenation or benefit. The map is a bounded interpretive synthesis, not a systematic review, ranking or claim that no prior work recognized identification. Supplementary Table 1 provides the structured synthesis; Supplementary Data 2 contains the complete source-level coding, evidence locators and codebook.

### Identification analysis

We treated the observable law *P*(*A*, *W*, *X*), declared loss and predictor class as the information available to an age-decoding task. Under unrestricted squared-error loss the target is *E*(*A*|*W*, *X*); under model restriction the fitted decoder approximates a loss-and class-specific projection. Latent-state non-identification was established by constructing distinct extensions *P*(*A*, *W*, *X*)*r_k_*(*H*|*A*, *W*, *X*) with identical observable laws and optimal age decoders. For any proposed biological property *ψ*, identification requires *ψ* to be constant over the resulting observational-equivalence class. If added measurement, endpoint, longitudinal or causal restrictions make *ψ* constant only in a narrower class, its identity arises from those restrictions. Prediction-error, transport, approximation, estimation and calibration contributions were separated algebraically in Supplementary Note 1; the decomposition is an identity, not an identifiable biological attribution of an observed individual gap.

Shared age-related covariance and proxy overlap among chronological age, measured biological processes, exposures, composition, cohort and survival selection were represented locally by Cov(*G_A_*, *Y*|*A*, *W*) ≈ *W_A_^T^ Σ_R_β_Y_*. This relation describes conditional association with the named endpoint under the stated local representation, not a causal decomposition or identification of ageing acceleration.

Equality of the complete endpoint law after scalar compression was represented by *Y_Δ_* ⊥ *X*|*A*, *W*, *q_A_*(*W*, *X*). This predictive-sufficiency condition is stronger than equality under one loss and is not inherited from age supervision. For an age-valued mapping *S* = *f*(*X*), intervention movement was represented exactly by *ΔS* = *f*(*X* + *ΔX*) − *f*(*X*). This identity supplies neither a causal outcome contrast nor a surrogate relationship.

For the age-and background-residualized score *G_A_* = *q_A_*(*W*, *X*) − *E*{*q_A_*(*W*, *X*)|*A*, *W*}, the conditional-expectation projection gives

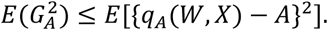

Full statements and regularity qualifications are assigned to Supplementary Note 1. The proposition concerns the identifying content of the chronological-age-decoding objective. Additional longitudinal, causal, measurement, endpoint or invariance restrictions may identify separately defined objects and may validate an already age-trained score for a specified use. That measurement validity derives from the added evidence, not from the age-decoding objective.

### Haematopoietic stem-cell transplantation analyses

We treated allogeneic haematopoietic stem-cell transplantation as an object-affiliation setting: the assayed circulating blood compartment is donor-derived, whereas the recipient remains the host organism. The analyses were not used to establish the analytical non-identification result.

For the Søraas *et al.* dataset^21^, we parsed Table 1 directly from the Europe PMC XML (PMCID: PMC6413751), without manually transcribing values or reprocessing methylation data. The table contained 41 rows. We reproduced the five published footnote-b exclusions—an outlying clock value or donor chimerism below 80%—leaving 36 published analysis samples from 30 recipients. The primary clean-compartment population retained samples whose latest recorded relapse status was “No”, because relapse can reintroduce recipient malignant cells into blood; this yielded 24 samples from 19 recipients. Let *S* denote the reported Horvath-2013 multi-tissue DNA-methylation-age score in recipient blood, *A_L_* the donor-lineage chronological coordinate at blood draw and *A_R_* recipient chronological age at blood draw. For each sample we calculated *d* = |*S* − *A_L_*| − |*S* − *A_R_*|; *d* < 0 indicates numerical proximity to the donor-lineage coordinate. Repeated observations were averaged within recipient, and recipients—not samples—were the independent units. We reported the recipient-weighted mean and median *d*, the numbers of negative, tied and positive recipient means, and an exact two-sided binomial test among non-tied recipients. A 95% percentile interval for the mean was obtained from 10,000 recipient bootstrap samples (seed 20260730). Sensitivities included all 36 published analysis samples, the non-relapse population with donor chimerism at least 95%, the non-relapse population sampled at least 12 months after transplantation and leave-one-recipient-out estimates. Five contemporaneous donor-blood comparisons were treated as descriptive only.

For Holland *et al.*^22^, we used the six author-normalized public beta-value matrices and comRand.csv metadata from Mendeley Data version 1 (doi:10.17632/j5krtjfj6y.1). The matrices comprised 334 arrays—303 recipient and 31 donor arrays. We implemented the 353-CpG Horvath-2013 coefficient formula and verified it against the bundled five-sample reference before scoring. Because the matrices had been processed separately by batch, we did not combine-renormalize them, apply ComBat or perform cohort-derived imputation. The six batches lacked 52, 28, 34, 18, 13 and 10 target probes, respectively. A probe absent from an entire batch was assigned the fixed overallMeanByCpGacross50data value from the local PC-Clocks resource; these values have not been established as the original Horvath training means. We therefore refer to the result as a Horvath-2013 coefficient implementation with fixed external-mean imputation, not as a paper-identical reconstruction of the original clock.

We parsed each array identifier into donor–recipient pair, role, batch and measurement. Arrays with the same pair, role and all four public age/time fields were averaged as one biological occasion, reducing 303 recipient arrays to 264 recipient occasions; distinct occasions were retained. For recipient samples, *A_L_* = *A_D_*_,TX_ + *t* denotes the donor-lineage chronological coordinate and *A_R_* = *A_R_*_,TX_ + *t* denotes recipient-host age. *A_L_* equalled the public AgeCells field to floating-point precision but was not interpreted as the literal age of the short-lived blood cells at sampling. The primary population retained recipient-role occasions at least 0.1 years after transplantation and with |*A_R_* − *A_L_*| ≥ 10 years, yielding 118 occasions from 61 donor– recipient pairs.

For each occasion we calculated

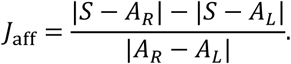

Positive values favour the lineage-coordinate side, negative values favour the host side and zero indicates equal distance. By the reverse triangle inequality, −1 ≤ *J*_aff_ ≤ 1. The index saturates at either boundary and is not a normalized measure of absolute accuracy. For the occasion-first summary, *J*_occ_, we averaged *J*_aff_ equally across occasions within pair and then equally across pairs. This gives the primary-processing value of 0.535 in Extended Data Fig. 2d; it is not the sign-balanced pair-summary statistic used in Fig. 2c. Uncertainty was quantified with 10,000 pair bootstrap samples (seed 20260730); signs of pair means were summarized with an exact two-sided binomial test among non-tied pairs. We also reported pair-weighted absolute errors to *A_L_* and *A_R_*.

Prespecified sensitivities used |*A_R_* − *A_L_*| > 0 and thresholds of 5, 15 and 20 years; the earliest eligible occasion; the occasion closest to one year; separate host-older and host-younger strata; the within-pair median; leave-one-batch-out reaggregation; and exclusion versus truncation of near-zero negative follow-up times (Extended Data Fig. 2). A post-specification processing sensitivity retained observed values only for the 253 probes present in all six batches and assigned the same fixed external mean to the other 100 probes in every batch. Across all samples before primary-population screening, 41 biological occasions had duplicate arrays: 39 recipient and two donor occasions. Their median within-occasion score range was 3.89 years and their maximum range was 11.502 years; duplicate arrays were not treated as independent observations. The pair bootstrap is conditional on the author-normalized matrices and does not incorporate preprocessing or imputation uncertainty. Holland *et al.* extended and partly overlapped the Søraas cohort and was not treated as an independent cohort replication.

Because numerical closeness to *A_L_* can arise from score marginals and donor/recipient age geometry alone, the prespecified geometry-preserving score-pairing null analysis used only the recipient-or pair-level summaries and did not rescore methylation. For this analysis, scores and chronological coordinates were first averaged within pair or recipient, and the same distance formula was then evaluated on those summaries to give *J*_pair_. Units were stratified by the sign of *A_R_* − *A_L_*, and the primary statistic *J*_aff,bal_ equally weighted the two stratum means. In Holland, the pair-summary stratum means were 0.488 and 0.763, giving 0.626. This differs from averaging occasion-specific indices before pair and stratum aggregation; the absolute-value operation makes the order consequential. Within each sign stratum, we permuted fixed scores across independent units while holding every (*A_R_*, *A_L_*) pair fixed, using 20,000 permutations (seed 20260823). This preserved score marginals, age geometry and stratum sizes while breaking observed score–referent pairing. It also broke the score–recipient-age relationship, so excess affiliation under this null does not exclude host-age-dependent shrinkage or estimate donor influence conditional on recipient age. The one-sided finite-permutation value used the add-one correction. Uncertainty for the observed statistic and observed-minus-null-mean contrast used 10,000 stratified independent-unit bootstrap samples (seed 20260824). Constant mean-and median-score controls and ordinary least-squares *S*-on-*A_L_* and *S*-on-*A_R_* fits were descriptive geometric and calibration diagnostics and were not used to recalibrate the scores. The prespecified analysis required the Holland observed statistic to exceed the null 97.5th percentile, a positive bootstrap interval for the observed-minus-null contrast, positive contrasts in both sign strata, the same direction in Søraas, positive Holland *S*-on-*A_L_* slopes in both strata and deterministic reconstruction.

As a conventional sensitivity, we fitted *S* − *A_R_* = *α* + *β*(*A_L_* − *A_R_*) + *ε* separately in Holland and Søraas at the frozen independent-unit level. Because the outcome and regressor share −*A_R_*, common-anchor covariance can contribute to *β*; we therefore treated it as a descriptive anchor-coordinate sensitivity, not an independent referent-identification parameter. We report ordinary least-squares estimates, HC3 intervals, 10,000-draw independent-unit bootstrap intervals (seed 20260826), *R*^2^, residual mean-squared error and constrained *β* = 0 and *β* = 1 anchor fits with a freely estimated intercept. The datasets were not pooled and the fitted scores were not recalibrated.

### NHANES data, endpoint and survey design

We used de-identified public-use National Health and Nutrition Examination Survey data and the 2019 public-use linked mortality file, with follow-up through 31 December 2019^44–46^. The originating protocols obtained ethics-review-board approval and informed consent^47^. NCHS reports that follow-up time or underlying cause was replaced by synthetic values for selected public-use records, whereas vital status was not perturbed^44^; released values were used without reconstruction. All modules used a fixed five-year all-cause mortality endpoint. We directly modelled five-year event probability and evaluated log loss.

The continuous-NHANES modules used participants aged 20–79 years from the 2007–2008, 2009–2010 and 2011–2012 cycles. Public-use ages reported as 80 were excluded because age is top-coded. These cycles supplied at least 60 months of administrative follow-up for every event-free included participant. The complete-case pure-decoder interface contained 13,724 participants and 470 deaths and was analysed without survey weights; it targets the declared predictive sample, not population prevalence. The endpoint-directed acquisition module used separately declared source-and target-cycle interfaces.

The NHANES III protocol-transfer modules used the canonical Phase 1 source interface (7,558 participants; 402 five-year deaths) and Phase 2 target interface (7,774 participants; 392 deaths), restricted to ages 20–79 years. The Phase 2 analysis used Mobile Examination Center weights, 23 released strata, 46 primary sampling units and fixed primary-sampling-unit-atomic folds. Missing measurements were retained and imputed by the training-fold weighted median. These modules estimate design-weighted predictive loss in the declared Phase 2 interface.

### Pure chronological-age decoder and same-source endpoint comparison

Background *W* comprised sex, race or ethnicity and survey cycle, observed chronological age was *A*, and *X* was the fixed ordered 50-measurement *S*_3_ panel inherited from the complete-support analysis. The sole endpoint was death by 60 months. We compared exactly three endpoint information sets: *M_WA_* = *P*(*Y*_5_|*W*, *A*); *M*_CLOCK_ = *P*(*Y*_5_|*W*, *A*, *S*), where *S* = *q_A_*(*W*, *X*) was trained only to reconstruct *A*; and *M_X_* = *P*(*Y*_5_|*W*, *A*, *X*). Given *A*, the raw gap *S* − *A* and *S* are deterministically equivalent, so no separate gap model was fitted.

Within every frozen outer training fold, categorical background was one-hot encoded and the 50 measurements were standardized using training data only. Ridge regression predicted chronological age. The penalty was selected by three-fold age-quintile-stratified root-mean-square error from the fixed grid {0.01,0.1,1,10,100,1000}. Cross-fitted scores were generated for the outer-training participants using the selected penalty and fixed inner folds; the decoder was then fitted on the full outer-training fold to score the held-out outer fold. No mortality label entered decoder preprocessing, tuning or fitting.

For *M*_CLOCK_, ridge logistic regression was fitted within each outer training fold using its cross-fitted clock scores, *W* and the same five-knot fold-local cubic age spline used in the pre-existing complete-support analysis. The endpoint penalty was selected by mean three-fold inner-validation log loss from the prespecified grid used in that analysis. Category encoding, score scaling, spline construction and tuning were training-fold local. *M_WA_* and *M_X_* were unweighted L2-logistic models with an intercept, reference-coded background and a cubic age basis with five uniformly spaced training-range knots and linear extrapolation; *M_X_* also included all 50 transformed measurements. Measurement and age-basis columns were standardized within each training partition, and three inner folds selected the penalty by mean fold-level validation log loss. Their stored predictions were reused after participant, fold and feature-order checks. SM2 identifies their generating script and model aliases; the software provides a combined ordered 50/25-variable dictionary with original units and transformations.

Let *L*(·) denote pooled participant-level out-of-fold log loss. The paired contrasts were

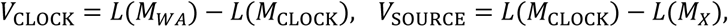

and *V_X_* = *L*(*M_WA_*) − *L*(*M_X_*). Positive *V*_SOURCE_ means that direct source-measurement modelling performed better for this endpoint under the declared estimators; it is not a population information estimate or a universal finite-sample theorem. We reused 2,000 paired participant-bootstrap indices from the pre-existing complete-support analysis, stratified jointly by endpoint and decade age band. Intervals condition on stored out-of-fold predictions and omit model refitting, tuning and study-development uncertainty.

### Individual measurement contributions to conditionally centred AgeGap

For participant *i* in outer fold *k*, we reloaded the frozen ridge age decoder and reproduced its stored held-out score *S_i_*. Using outer-training participants only, a multivariate ridge model estimated the conditional mean of each of the 50 measurements given chronological age and background. Age was represented by a five-knot cubic spline and background comprised sex, race or ethnicity and survey cycle. The penalty was selected by three-fold age-quintile-stratified cross-validation from {0.01,0.1,1,10,100,1000}, minimizing standardized measurement-reconstruction error. All tuning estimators and final residualizers were persisted, reloaded and parity-checked.

Let *β_kj_* and *σ_kj_* denote the frozen decoder coefficient and training-fold scale for measurement *j*, and let *μ*^*_kj_*(*A_i_*, *W_i_*) denote its fold-local conditional mean. The score centre was induced by passing the vector of these measurement means through the full frozen decoder:

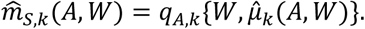

This evaluation retained the decoder’s background encoding, training-fold scaling and intercept; it was not a separately fitted score-on-age regression. We defined

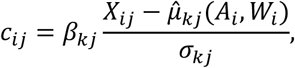

so that

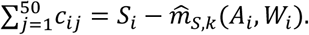

This sum is the conditionally centred AgeGap, not the raw gap *S_i_* − *A_i_*, which was retained separately. Because matched participants shared outer fold, age and background exactly, their raw-gap and centred-gap differences were equal to numerical precision.

Pairing was deterministic and outcome blind. Within exact strata of outer fold, chronological age, sex, race or ethnicity and survey cycle, participants were ordered by conditionally centred AgeGap; adjacent candidate edges were ordered by absolute gap difference and row identifiers and retained greedily without participant reuse. Pairs were retained when their centred gaps differed by no more than 0.25 decoder-output years. Contribution-profile distance was calculated before display selection. The two displayed pairs were chosen by the frozen median-and 95th-percentile-distance rules among same-sign pairs with mean absolute centred AgeGap of at least two decoder-output years. Distances are norms in model-contribution space, not participant age differences. The contributions explain how the frozen linear decoder generated its scalar and are not causal, organ-specific or biological-age components; their allocation among correlated measurements depends on the frozen ridge parameterization, scaling and centring reference.

### Flexible calibration of the continuous-NHANES age decoder

We reused the original five outer folds and frozen fold-local age scores. The endpoint model contained background, a five-knot cubic spline of chronological age and one prespecified five-knot cubic spline of *q_A_*; ridge-logistic regularization was selected inside each outer training set from the prespecified grid. No alternative spline or calibration search was performed. The contrast was the held-out log loss under this flexible clock model minus that under direct modelling of the frozen 50-measurement panel. The fully nested continuous-NHANES matched-scalar comparison is described in Supplementary Methods SM2b; the NHANES III Phase 1-to-Phase 2 protocol transfer is described below and in SM2c.

### NHANES III matched scalar target-supervision protocol transfer

Background comprised sex and race or ethnicity, and *X* comprised the same ordered 25 measurements for both one-dimensional projections. Chronological age was not an input. In NHANES III Phase 1, weighted-median imputation, one-hot background encoding, weighted measurement standardization and ridge regularization were identical for the two projections. The age-supervised projection *q_A_* used standardized chronological age as its target; the endpoint-supervised projection *q_Y_* used standardized five-year mortality. Target scaling and all preprocessing were estimated inside the relevant Phase 1 training folds. One final projection of each type was fitted on complete Phase 1 and applied unchanged to Phase 2; Phase 2 mortality outcomes did not enter projection preprocessing, tuning or fitting.

In Phase 2, both frozen scores received the same target-local opportunity: weighted L2-penalized logistic regression containing background, a cubic age spline with fixed knots at 20, 35, 50, 65 and 80 years and a training-local five-knot uniform cubic spline of the relevant score. Regularization was selected by pooled design-weighted log loss in three inner folds, and all reported probabilities were held out across five fixed outer folds. We compared the background-and-age model with the addition of *q_A_*, and directly compared otherwise identical models adding *q_A_* or *q_Y_*. A linear-score version was prespecified as a sensitivity.

The separate age-substitution comparison used the same Phase 2 participants, folds, weights and downstream model family. One arm contained background and a five-knot spline of observed age; the other replaced age with an identically constructed spline of the frozen *q_A_*. It tested superiority, not equivalence or non-inferiority.

For both modules, 32 Fay balanced repeated replication (Fay-BRR) weight sets with *ρ* = 0.3 repeated Phase 2 downstream preprocessing, inner tuning, fitting and held-out prediction; intervals used 23 design degrees of freedom. Phase 1 projection directions remained fixed. The intervals therefore quantify Phase 2 survey-design uncertainty conditional on the two frozen projections and do not include Phase 1 development uncertainty.

### NHANES III chronological-age increment and proxy-substitution comparison

We compared the incremental mortality-prediction value of observed chronological age after two information sets. Background *W* comprised sex and race or ethnicity. The full panel contained the ordered 25 measurements. The reduced panel removed exactly the five-measurement cardiorespiratory domain selected for age reconstruction in the earlier Phase 1 acquisition analysis: systolic blood pressure, diastolic blood pressure, radial pulse, forced expiratory volume in one second and forced vital capacity. The mask was frozen before the present Phase 2 fitting and was not reselected using Phase 2 outcomes. The remaining panel is described only as the remaining 20 measurements; it is not assumed to lack age-related information.

Four otherwise homologous weighted L2-logistic models predicted five-year death: full and reduced panels were each fitted without and with a training-local fixed-knot cubic spline of observed age. Encoding, weighted-median imputation, weighted standardization, spline construction, inner tuning and outer fitting were repeated separately for every model and weight set. No age-decoder score or endpoint-supervised scalar entered these models.

The full-panel age increment was the reduction in design-weighted held-out log loss from adding age to the 25-measurement model. The reduced-panel age increment was defined analogously after the frozen block was withheld. Their difference was the primary predictive proxy-substitution contrast. Secondary contrasts measured the block’s value without age, its value with age, and reduced-panel-plus-age versus full-panel-without-age. Thirty-two Fay-BRR replicates with *ρ* = 0.3 repeated the complete Phase 2 pipeline and used 23 design degrees of freedom. Intervals are conditional on the frozen Phase 1 mask and omit its development uncertainty. No contrast was interpreted as causal mediation, equivalence, biological-age identity or complete measured-state sufficiency.

### Endpoint-directed measurement acquisition

For measurement acquisition, the sole endpoint was five-year all-cause mortality. The source population was NHANES 2007–2008 (4,599 complete participants; 174 deaths) and the target was 2009–2010 (4,896; 157). Candidate acquisition categories were renal/mineral, metabolic/adiposity/lipid, hepatic/protein/injury, haematology and haemodynamic measurements. For domain *M_k_*, source measurement value was

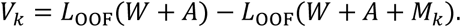

For the inverse chronological-age policy, each domain *M_k_* entered a ridge decoder *A* ∼ *W* + *M_k_*, where *W* comprised sex and race or ethnicity. Encoding, declared feature transformations and standardization were training-fold local. Five age-quintile-stratified outer folds (seed 20260826) supplied participant-level out-of-fold predictions. Within each outer training fold, the ridge penalty was selected from {0.01,0.1,1,10,100,1000} by pooled mean-squared error across three age-quintile-stratified inner folds (seed 20260827 + outer fold). The inverse-decoder policy selected the domain with the lowest pooled source out-of-fold age mean-squared error; the separately frozen endpoint policy selected argmax*_k_V_k_*. Source choices and the source-data signature were frozen before target evaluation, and target outcomes did not enter either source selection.

In the target cycle, we compared the held-out five-year mortality log-loss gains of the two source-frozen domains beyond age and background. Participant identity, fold assignment and ordered features matched the sealed predecessor. Uncertainty used 5,000 event-stratified paired participant bootstraps of fixed out-of-fold losses (seed 20260802), without policy reselection or model refitting. Each of the five domains was treated as one acquisition option. Domains differed in measurement count, and no equivalence of monetary resources or real-world costs was assumed.

### Transparent three-variable construction

We retained X1–X3 from the frozen cross-sectional geometry generated with seed 20260806. Age was sampled uniformly from 20 to 80 years and a binary separation state *D* with probability 0.40 (realized State 0, *n* = 299; State 1, *n* = 201). Let *A_z_* be standardized age, *D_c_* centred state, *Q_A_* the standardized quadratic age component after projection off the linear age direction, *ε_j_* independent standard-normal noise and *z*(·) sample standardization. The retained variables were

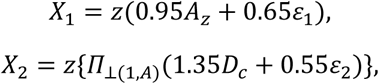

and

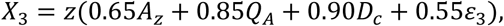

where *Π*_⊥(1,*A*)_ removes the realized intercept and linear age direction. Thus X1 supplied noisy age readability without a designed state term, X2 supplied declared same-age structure that was exactly linearly age-orthogonal in the realized construction, and X3 mixed linear and nonlinear age structure with state. The displayed limiting cases—noisy age without state and exact age alongside a separate state variable—are algebraic consequences rather than additional fitted analyses.

We assigned the 500 rows to five participant-disjoint folds stratified jointly by the declared binary state and five equal-frequency age groups (shuffle seed 20260827). Chronological-age decoding used four fixed ridge pipelines containing X1, X2, X3 or all three variables; each applied a five-knot cubic spline, training-fold standardization and Ridge(*α* = 1). Same-age state decoding used a fixed L2-logistic age-only model and four homologous age-plus-measurement models; age entered as a five-knot cubic spline and the measurements were standardized within training folds. The primary task-map quantities were pooled out-of-fold age *R*^2^ and the reduction in pooled state log loss relative to the age-only model. No model-family, penalty, split or threshold search was performed.

The pooled age *R*^2^/state log-loss gains beyond age were 0.681/−0.002 for X1, −0.015/0.407 for X2, 0.301/0.220 for X3 and 0.730/0.483 for X1–X3. These are fixed descriptive quantities for the declared construction and carry no population interval or human biological interpretation.

### Statistical analysis

Analysis records were constructed rows for X1–X3, recipient-or pair-level summaries for HSCT and participants for predictive NHANES analyses. Validation and resampling followed each module’s design; in NHANES III, validation folds kept primary sampling units (PSUs) intact. For the individual contribution summary, the descriptive unit was one retained non-reusing matched pair. The fixed X1–X3 construction was descriptive and supplied no population interval. The individual contribution analysis was also descriptive: matching used no endpoint information, all retained non-reusing pairs were reported and no hypothesis test or population interval was assigned to their empirical distance distribution. Continuous-NHANES uncertainty used the declared participant or event-stratified fixed-prediction bootstrap and therefore conditions on stored models, folds and policies. NHANES III uncertainty used full Phase 2 Fay-BRR downstream refits with 23 design degrees of freedom, conditional on frozen Phase 1 projection directions or masks. Exact two-sided binomial tests excluded ties. Lower log loss indicates better probabilistic prediction; reported reductions are paired held-out proper-score contrasts, not clinical effect sizes or percentage accuracy. An interval spanning zero was interpreted as an unresolved contrast in that design, not proof of no effect, equality or non-inferiority. Fold-construction rules and model grids are detailed in Supplementary Methods; persistence audits and sensitivity results are indexed in Supplementary Software 1 and Source Data. Repository-level version, artifact and reconstruction records are indexed in the reproducibility manifest included in Supplementary Software 1.

### Use of artificial intelligence

OpenAI Codex assisted with development of analysis and verification scripts, manuscript drafting and editorial revision. Analyses were executed using the versioned code and data described here; LLM-generated statements were not treated as data or bibliographic evidence.

## Data availability

The Søraas 2019 article table is available through Europe PMC (PMCID: PMC6413751). Holland 2024 individual-level metadata and six author-normalized methylation matrices are available from Mendeley Data version 1 (https://doi.org/10.17632/j5krtjfj6y.1); the article and supporting information are available through Europe PMC (PMCID: PMC11113269). Continuous NHANES, NHANES III and public linked-mortality data are available from the US Centers for Disease Control and Prevention/National Center for Health Statistics^44–46^. The Arivale and TwinsUK individual-level data underlying the published multi-omic BMI example are controlled access and were not reanalysed^48^. The source-verified construct-promotion audit and critical-literature remedy map are provided as Supplementary Data 1 and 2. Figure-level non-restricted derivatives, including aggregate matched-pair distributions and the predeclared display-pair contribution profiles underlying Extended Data Fig. 4, are provided in Source Data. Analysis specifications, code, ordered feature definitions, accession and module manifests, transformation definitions and hash/path references to internally retained reconstruction objects are provided in Supplementary Software 1. Supplementary Software 1 also provides frozen primary participant-level predictions, stored survey-replicate estimates, and the complete individual contribution profiles and matrix for deterministic result checks. Fitted-model binaries and large resampling-index arrays are retained internally and are not included in the submission archive. For peer review, the supplied code and derivative data are provided to editors and reviewers for evaluation. Third-party data remain subject to their source terms. Public deposition and reuse-licensing details will be supplied before publication.

## Code availability

Analysis and figure-generation code for the transparent age–state construction, HSCT object-affiliation and referent-null analyses, continuous-NHANES pure age decoder, individual conditionally centred AgeGap contribution decomposition and outcome-blind matching, NHANES III matched target-supervision, age-substitution and proxy-substitution comparisons, endpoint-directed acquisition and submission figures is provided as Supplementary Software 1. The snapshot includes the pinned environment specification, deterministic entry points, evidence-module registry, reproducibility manifest, source-data indices and non-restricted result tables used by the submission. The archive includes a portable stored-result verification entry point and installation and execution instructions. It is supplied for editorial and reviewer evaluation; no public reuse licence has been assigned.

## Acknowledgements

We thank the US National Center for Health Statistics for making NHANES survey and linked mortality data publicly available, and acknowledge the contributions of the survey participants and staff. We also thank the investigators of the transplantation studies by Søraas et al. and Holland et al. for sharing the data used in our analyses. We thank Dr Jianyue Ding for providing full-text articles and assisting with the review of the source literature.

## Author contributions

SS: Conceptualization, Methodology, Data Curation, and Writing - Original Draft. QL: Formal Analysis, Validation. QG: Conceptualization, Supervision, and Writing - Review & Editing,

## Additional information

Supplementary Information is available for this paper. Correspondence and requests for materials should be addressed to corresponding author.

