## Supplementary material for "Inferential boundaries of age prediction: why prediction does not establish biological age measurement": Source_Data: Figure_3_4_SOURCE_MAP.md

# Source and assembly map

The quantitative points and confidence endpoints are read without arithmetic
modification from byte-exact frozen source tables. Four-decimal point labels
are display rounding; interval geometry retains the full saved precision.

| New panel | Frozen figure evidence | Assembly change |
|---|---|---|
| Fig. 3a | Prior Fig. 3a design table | Reflow same 50-measurement, age-only training and three endpoint-model information sets |
| Fig. 3b | Prior Fig. 3b three-contrast table | Same estimates and confidence intervals; keep labels within panel |
| Fig. 3c | Current main Methods and HUM-06 contract | Diagram Phase 1 ridge projection fitting and freezing, then Phase 2 fold-local calibration; no new analysis |
| Fig. 3d | Prior Fig. 3c matched-scalar table | Same U_A and Delta estimates and confidence intervals, now paired with explicit design |
| Fig. 4a | Prior Fig. 4b target-first workflow | Reflow existing question → estimand → design/validation → claim structure |
| Fig. 4b | Current ED5 policy definitions, policy intervals and paired contrast | Compact source-selection diagram plus three existing target-cycle intervals |
| Fig. 4c | Prior Fig. 4c five-object map | Plain-language display condensation; full map copied with all fields |

Design and denominator statements are from
`manuscript/nature_first/MANUSCRIPT_NATURE_IDENTIFICATION_FIRST_v6_21_20260909.md`:
“NHANES data, endpoint and survey design”, “Pure chronological-age decoder and
same-source endpoint comparison”, “Flexible calibration of the continuous-NHANES
age decoder”, “NHANES III matched scalar target-supervision protocol transfer”,
and “Endpoint-directed measurement acquisition”. In particular:

- Continuous NHANES: 2007–2012; 13,724 participants; 470 five-year deaths;
  background includes sex, race/ethnicity and cycle; 50 measurements;
  five outer folds. Decoder training uses chronological-age labels only.
- NHANES III: the projection inputs are 25 measurements plus sex and
  race/ethnicity, not observed age. Standardized age or standardized five-year
  mortality supplies the ridge regression target. Phase 2 has 7,774 participants
  and 392 deaths; observed age enters the common downstream endpoint models.
  Projection directions remain frozen, but Phase 2 calibration is fitted,
  tuned and evaluated out of fold. Fay-BRR refits the Phase 2 downstream models.
- Acquisition: source 2007–2008; target 2009–2010 with 4,896 participants and
  157 deaths. Both domains were source-selected and then frozen. Their source
  criteria have different units and are not drawn on a common numeric axis.
  Target policy gains share the mortality log-loss scale. The source menu is
  not a matched resource budget.

Source_Data.zip routing: `source_data/Figure_3/*.tsv` maps to `Figure_3/`;
`source_data/Figure_4/*.tsv` maps to `Figure_4/`. Include this source map and the
source index as provenance. No old age-context evidence is retired: the complete
frozen HUM-03 contrast set is still supplied under Extended Data Fig. 3.
