## Supplementary material for "Inferential boundaries of age prediction: why prediction does not establish biological age measurement": Source_Data: README.md

# Source Data: V6.24 / SI V5.16

Figures 1–2 and Extended Data Figures 1–5 are unchanged. Figure 3 now pairs the continuous-NHANES design/results (a,b; 50 measurements) with the NHANES III matched-scalar design/results (c,d; 25 measurements). Former Figure 3d contextual-age contrasts remain complete in Extended Data Figure 3. Figure 4b reproduces the existing acquisition policy gains and paired contrast; Figure 4c retains the object/evidence map. The seven Figure 3/4 source tables are exact copies, not new estimates.

Figure_3_4_source_data_index.tsv maps these files to frozen sources. Figure_3_4_SOURCE_MAP.md explains the panel remapping. Main-manuscript legends are the canonical captions. Design diagrams use the frozen specifications and sample descriptions; they add no fitted result. The software archive contains the static builder.

V6.23 changes only the Figure 4b common-menu connector so both selection routes are explicitly fed. All Source Data entries other than this README are byte-identical to V6.22. Figure_3_4_SOURCE_MAP.md and the index retain the original aggregate-table provenance; the current Figure 4 export is built by analysis/plot_v6_23_figure4_connector_v0_1.py using those same four Figure 4 tables. No new estimates or data tables are introduced.

V6.24 changes Figure 1 conceptual labels only. All quantitative source table bytes remain identical. publication_labels_v6_24.tsv distinguishes HSCT occasion-first from pair-summary statistics; historical index field names retain original provenance. V6_24_SOURCE_REPLACEMENT_MAP.tsv supplies current routing.

V6.25 submission packaging: quantitative tables and figure definitions are unchanged from V6.24. Editorial/reviewer access only at this stage; no new public reuse licence.
