## Supplementary_Information for "Inferential boundaries of age prediction: why prediction does not establish biological age measurement"

#### Chronological-age decoding does not identify biological age

**Scope.** Supplementary Note 1 gives the derivations for loss-specific age-task identification, the conditional-state mismatch as one concrete case, the observable-law proposition, constrained within-age orientation, meaning non-inheritance, the unavoidable-residual argument, population Bayes-risk ordering under deterministic compression and the post-decoder operator chain. Supplementary Note 2 separates the contemporary construct-promotion audit from the structured review of critical and methodological responses, then states the Article's specific contribution relative to prior work. Supplementary Methods report the continuous-NHANES chronological-age decoder and supporting matched-scalar comparisons; the survey-weighted NHANES III Phase 1-to-Phase 2 matched-scalar protocol transfer, observed-age increment and proxy-substitution comparison, and observed-age-versus-age-decoder substitution boundary; the bounded endpoint-directed acquisition comparison; the fixed-model transparent age–state construction; and HSCT reproducibility pointers.

This Supplementary Information contains the analytical derivations; separate continuous-NHANES and NHANES III predictive analyses; a bounded endpoint-directed acquisition comparison; and one fixed cross-sectional age–state construction. The continuous-NHANES modules are within-interface supporting comparisons. The NHANES III analyses use survey weights and phase-frozen objects, but they are retrospective and outcome-accessed robustness analyses rather than untouched confirmations or independent external validations. HSCT referent methods are reported in the main Methods, with reproducibility pointers indexed here.

The logical and empirical responsibilities are separate. Supplementary Note 1 establishes the non-identification claim and its meaning non-inheritance consequence. The construct-promotion audit establishes that the inferential operation remains current, whereas the critical-literature review organizes proposed repairs and their identification ceilings. HSCT evaluates the referent of recipient-blood scores. The continuous-NHANES modules evaluate endpoint utility and additionally decompose each conditionally centred decoder deviation into exactly additive measurement contributions to test whether near-equal conditionally centred gaps can arise from dissimilar measurement-contribution profiles. The remaining NHANES modules evaluate whether supervision target changes one-dimensional endpoint performance, how observed age's increment changes when a pre-frozen measurement block is unavailable and whether an age decoder can substitute for observed age. The acquisition comparison is a bounded illustration of an endpoint-directed alternative. X1–X3 use fixed out-of-fold models to illustrate that age readability and declared same-age state value are distinct task properties. No construction or human analysis is required to prove the analytical results.

### Supplementary Note 1 | What chronological-age loss identifies

#### Observed task and latent extension

Let $A$ denote observed chronological age, $W$ declared background variables, $X$ a measured biological profile and $H$ a hypothetical scalar biological age. The observed law is

$$P_{\text{obs}}(A, W, X).$$

For a declared loss $\ell$ and predictor class $\mathcal{F}$, the population chronological-age task identifies, when the minimizer is unique,

$$q_A^* = \text{argmin}_{q \in \mathcal{F}} E_{P_{\text{obs}}}[\ell\{A, q(W, X)\}].$$

Under unrestricted squared-error loss,

$$q_A^*(W, X) = E(A|W, X).$$

This is a property of the observed task. It is not, without further restrictions, an identification result for $H$.

#### Biological claims and inverse label reconstruction are different estimands

One major claim made for biological-age measurement is same-age heterogeneity. That claim is naturally expressed through a declared conditional state law such as

$$P(X|A, W),$$

or, when the target is latent, through a separately defined state $Z$ with an anchored measurement model for $P(X|Z, A, W)$ and a declared conditional law $P(Z|A, W)$. These objects describe how measured or model-defined state varies among people sharing chronological age and background. Other proposed meanings of biological age concern accumulated damage, functional reserve, a named future risk, longitudinal pace or response to intervention. Chronological-age decoding targets none of these objects by virtue of its label. It targets $P(A|X, W)$ or a loss-specific functional of that inverse distribution. The age task contains no label for whether a same-age difference is favourable, predicts a named endpoint, represents a longitudinal rate or can be reversed beneficially.

Under squared-error loss, the mismatch is also visible directly in the population risk. For any scalar decoder $q(W, X)$,

$$\begin{aligned} R_A(q) &= E[\{q(W, X) - A\}^2] \\ &= E[\text{Var}\{q(W, X)|A, W\} + \{E[q(W, X)|A, W] - A\}^2]. \end{aligned}$$

The equality follows from conditioning on $A, W$, for which $A$ is fixed. The first term is a non-negative penalty on dispersion that the scalar decoder expresses within age-and-background strata; the second is its conditional calibration error. The loss does not suppress heterogeneity in the measured profile $X$ itself. Rather, any part of that heterogeneity expressed as different decoded ages among people with the same $A, W$ raises squared error. Such variation may remain because the same features improve reconstruction across age strata, but the objective supplies no supervision for its biological direction or meaning.

The decomposition establishes how the age objective evaluates same-age differences: output dispersion contributes to reconstruction error, whereas a biological state claim requires a criterion for deciding what those differences mean. This is appropriate for reconstructing chronological age; the methodological problem arises when this optimization is treated as identification of a different biological target. Proposition 1 supplies the separate identification argument. Additional labels, longitudinal observations, measurement restrictions or causal designs can identify declared biological objects and validate their measurements. Their identifying content belongs to those additions. Together, the two results make the choice of training target and validation evidence part of the scientific definition of the measurement.

#### Loss, model class and training population select the task statistic

The non-identification result is not specific to squared error. Different losses select different functionals of the same conditional chronological-age distribution. With an unrestricted predictor class, squared loss selects the conditional mean, absolute loss selects a conditional median and pinball loss at level $\tau$ selects a conditional quantile:

$$q_{\text{sq}}^*(W,X) = E(A|W,X), \quad q_{\text{abs}}^*(W,X) \in \text{Median}(A|W,X), \quad q_\tau^*(W,X) \in Q_\tau(A|W,X).$$

Here $Q_\tau$ denotes the set of conditional $\tau$-quantiles, which can be non-unique for rounded or discrete age. Under an unrestricted or correctly specified distributional class, a strictly proper probabilistic score can instead identify the complete conditional age distribution $P(A|W,X)$, up to almost-sure equivalence. These are different Bayes acts for a declared age-prediction task, not alternative proofs of a common biological-age entity. Squared loss is used in the main text because it makes the conditional-mean target and the gap-collapse boundary explicit.

The fitted score has additional dependencies. A general penalized empirical objective can be written as

$$\hat{q}_{\lambda,\omega} \in \text{argmin}_{q\in\mathcal{F}}[\frac{1}{n}\sum_{i=1}^{n}\omega_i \ell\{A_i, q(W_i,X_i)\} + \lambda J(q)], \quad \tilde{q} = c_{\hat{\eta}}(\hat{q}).$$

Here $\mathcal{F}$ encodes the model family and feature representation, $\omega_i$ the sampling or age weights, $J$ the penalty, $\lambda$ the selected regularization strength and $c_{\hat{\eta}}$ any post-fitting output-calibration map. Preprocessing, hyperparameter selection and finite-sample estimation add further dependence.

For example, after residualizing declared background variables and centring the features, a quadratically penalized linear population target has the form

$$\beta_\lambda = (\Sigma_{XX} + \lambda P)^{-1}\Sigma_{XA}.$$

Here $P \succcurlyeq 0$, the moments are taken under the declared weighted training law and the displayed matrix is assumed invertible. Feature scaling, covariance, collinearity, age sampling and the penalty matrix can therefore redistribute coefficients while leaving predictions similar. A fitted loading quantifies contribution to the declared prediction under the chosen parameterization; it is not by itself a contribution to ageing or evidence of a causal mechanism.

Three levels should therefore be distinguished:

1. different parameter vectors may implement the same or nearly the same prediction function;

2. different losses, model classes, penalties, training distributions and calibration rules may select different age-task functions;

3. even a unique, population-known age-task function does not identify a latent biological state.

The third level is the central proposition. The first two explain why practical scores and individual rankings can be even less stable than the proposition requires.

#### Inverse age prediction depends on the reference population

In a density representation for a declared training population, with the integral replaced by a sum for discrete or rounded age,

$$p_{\mathrm{tr}}(a|x,w) = \frac{p_{\mathrm{tr}}(x|a,w)p_{\mathrm{tr}}(a|w)}{\int p_{\mathrm{tr}}(x|a',w)p_{\mathrm{tr}}(a'|w)\,da'}.$$

Thus an inverse age decoder depends on both the age-conditional measurement law and the age distribution of the training population. Restricting the age range, reweighting younger or older participants, selecting unusually healthy survivors or changing cohort composition can alter $P_{\mathrm{tr}}(A|X,W)$ and its Bayes acts even if the measured person is unchanged. Let $\mu(a,w) = E(X|A=a,W=w)$. The map $a \mapsto \mu(a,w)$ can be multidimensional, noisy or non-injective, and an observed profile need not equal its age-conditioned mean. No inverse $\mu^{-1}$ is therefore defined in general, and $E(A|X,W)$ should not be interpreted as mathematical inversion of an individual biological trajectory.

#### Different objectives select different scientific quantities

Chronological-age prediction and endpoint prediction optimize different risks. For a named endpoint $Y_\Delta$, the corresponding population target is instead

$$q^*_{Y,\Delta} \in \mathrm{argmin}_{q\in\mathcal{G}} E[L_Y\{q(A,W,X),Y_\Delta\}].$$

To compare downstream use of two age decoders on a common domain, define

$$R_A(q) = E\{L_A[q(W,X),A]\},$$

and the best endpoint risk attainable from each scalar, chronological age and background,

$$R_{Y,\Delta}(q) = \inf_g E[L_Y\{g(A,W,q(W,X)),Y_\Delta\}].$$

An ordering by chronological-age risk need not be preserved by the endpoint risk:

$$R_A(q_1) < R_A(q_2) \;\not\Rightarrow\; R_{Y,\Delta}(q_1) < R_{Y,\Delta}(q_2).$$

An endpoint predictor may legitimately be converted to age-equivalent units through a declared reference-risk curve, for example $S_{\mathrm{age}} = g_{\mathrm{ref}}^{-1}\{q_{Y,\Delta}(A,W,X)\}$, when $g_{\mathrm{ref}}$ is strictly monotone on the relevant range. Otherwise a declared generalized inverse and tie convention are required. This is a reparameterization of a named endpoint statistic in a named reference population, not evidence that the statistic and a chronological-age decoder estimate the same latent object.

Similarly, a multitask objective that combines chronological age with one or more endpoints introduces labels absent from chronological-age loss. Any improvement in prediction of a named endpoint may come from those added targets. A shared internal representation remains dependent on task weights, architecture and structural assumptions and is not automatically identified as biological age. Ensembles and probabilistic decoders trained exclusively on chronological-age labels can improve chronological-age calibration or estimate $P(A|W,X)$, but remain functions of the observable age task unless separate identifying restrictions are supplied.

#### Proposition 1: chronological-age decoding does not identify $H$

Sluiskes *et al.* formalized the untestability of the identical-association assumption using observationally equivalent systems: the observed marker–age law can be preserved while the marker's relation to latent biological age changes (Main ref. 8). This is a direct mathematical predecessor of the argument below. We express the observational-equivalence result in terms of the population age task and then derive its consequences for within-age orientation, post-decoding gaps, downstream prediction and the evidence needed to validate a biological measurement.

Consider any fixed observed law $P_{\text{obs}}(A,W,X)$. Choose two distinct conditional laws

$$r_1(H|A,W,X) \text{ and } r_2(H|A,W,X),$$

and define two full-data extensions

$$P_k(A,W,X,H) = P_{\text{obs}}(A,W,X)r_k(H|A,W,X), \quad k \in \{1,2\}.$$

Both extensions induce exactly the same law for every observed variable. They therefore produce the same population-optimal age decoder and, under any fixed training and evaluation procedure, the same distribution of held-out performance and of every statistic computed only from $(A,W,X)$. Nevertheless, they can assign different values, rankings or biological meanings to $H$. Hence $H$ is not identified by the identifying content of the chronological-age objective.

**Proof.** Identification requires the target to take the same value under every admissible full-data law that induces the observed law. The two extensions above are observationally equivalent but differ in the conditional law of $H$. Therefore no functional that distinguishes these two versions of $H$ is identified by $P_{\text{obs}}$, the age-prediction loss or the achieved prediction accuracy. □

#### Constrained corollary: chronological-age loss does not identify within-age orientation

Fix the observed law and age task of Proposition 1. Let $U = u(A,W,X)$ be a square-integrable measured contrast expressed in age units, with

$$E(U|A,W) = 0, \quad P\{\text{Var}(U|A,W) > 0\} > 0.$$

Even if candidate constructs are restricted to have conditional centre $A$ and the same within-age scale, both

$$H^{(+)} = A + U, \quad H^{(-)} = A - U$$

are admissible because

$$E(H^{(\pm)}|A,W) = A, \quad \mathrm{Var}(H^{(\pm)}|A,W) = \mathrm{Var}(U|A,W),$$

and $(H^{(+)} - A)^2 = (H^{(-)} - A)^2 = U^2$ pointwise. Yet, for two conditional draws $i,j$ from the same $(A,W)$ stratum,

$$H_i^{(+)} - H_j^{(+)} = -\{H_i^{(-)} - H_j^{(-)}\},$$

so every non-tied within-stratum ordering is reversed. Equivalently,

$$\mathrm{Cov}(H^{(+)} - A, U|A,W) = \mathrm{Var}(U|A,W),$$

whereas

$$\mathrm{Cov}(H^{(-)} - A, U|A,W) = -\mathrm{Var}(U|A,W).$$

If $U|A,W \overset{d}{=} -U|A,W$, the two candidates additionally have the same complete conditional distribution given $A,W$.

**Proof.** Attach the point masses $H = A + U$ and $H = A - U$ to the same $P_{\mathrm{obs}}(A,W,X)$. Marginalizing $H$ gives the same observable law, so every chronological-age Bayes act, attainable risk and observable age-task statistic is identical. The displayed identities show that the age anchor and within-age scale agree while the orientation functional has opposite sign. It is therefore not constant across compatible extensions and is not identified by the age task. □

The two candidates make the missing requirement explicit: a same-age contrast needs an independently justified biological orientation. An endpoint, measurement relation or longitudinal anchor can supply that orientation. Age labels and age loss leave the two polarities observationally equivalent, and the construction assigns neither $U$ nor $-U$ to the fitted clock gap.

Perfect age recovery does not alter the argument. Even if $q_A^*(W,X) = A$ almost surely, the two extensions remain observationally equivalent and can retain different same-age latent states. Perfect prediction establishes recoverability of the age coordinate from $X$; it does not select a unique state that the prediction should be said to represent.

The proposition also does not require that a fitted score be unrelated to measured physiology or future outcomes. In a particular population, $q_A^*$ may correlate with a declared measured profile or improve prediction of a named endpoint. Such findings support those exact associations or predictive contrasts in the evaluated population and horizon. They do not show that chronological-age loss uniquely selected the latent entity to which the score was subsequently assigned, and they cannot retroactively change the object identified by the training objective.

#### Meaning non-inheritance across observationally equivalent systems

Let

$$\mathcal{E}(P_{\mathrm{obs}}) = \{P_H : P_H(A,W,X) = P_{\mathrm{obs}}(A,W,X)\}$$

denote the equivalence class of full-data laws that induce the observed law. Let $\psi(P_H)$ represent any claimed biological property of a candidate age construct: a same-age ordering, an accumulated-damage scale, functional reserve, a named future-risk meaning, a longitudinal pace or an intervention target. Identification by the observed age task requires

$$\psi(P_{H,1}) = \psi(P_{H,2}) \text{ for every } P_{H,1}, P_{H,2} \in \mathcal{E}(P_{\text{obs}}).$$

Proposition 1 constructs members of the same equivalence class for which this equality can fail. The age-task statistic may be uniquely defined and perfectly estimated while $\psi$ varies. Unique prediction is therefore not unique biological meaning.

Suppose additional structure $\mathcal{C}$ restricts the admissible laws to

$$\mathcal{E}_{\mathcal{C}}(P_{\text{obs}}) \subseteq \mathcal{E}(P_{\text{obs}}).$$

If a measurement model with anchors, endpoint observations, repeated-measurement dynamics or a causal design makes $\psi$ constant within this restricted class, then $\psi$ is identified under $\mathcal{C}$. Such evidence can validate an already-trained age clock as a measurement of a defined object when the required measurement relation is established. Its measurement identity then derives from $\mathcal{C}$, not from chronological-age loss. Likewise, a downstream association or intervention result establishes its named predictive or treatment contrast; a stronger measurement claim requires the corresponding identifying relation. We refer to the separation between these sources of meaning as the meaning non-inheritance principle.

The principle is agnostic to the substantive definition proposed for biological age. If biological age is defined tautologically as the output of a particular age decoder, the output is computable by definition, but no additional physiological, prognostic, longitudinal or causal meaning follows. If biological age is defined through any of those meanings, the corresponding measurements and restrictions carry the identification burden.

#### From a fitted score to raw and age-residualized gaps

For a predictor $q_A(W, X)$, define the raw age gap using the sign convention common in clock studies:

$$G_{\text{raw}} = q_A(W, X) - A.$$

This is the negative of the conventional regression residual $A - q_A(W, X)$. It is created after training by comparing the fitted age coordinate with the already observed chronological age; it is not itself the target optimized by the age loss.

Let $P_{AW}$ and $P_{XW}$ denote the conditional-expectation projections $P_{AW}Z = E(Z|A, W)$ and $P_{XW}Z = E(Z|X, W)$. Then the scalar age-residualization operator and the feature-wise same-age conditioning operator are

$$G_A = (I - P_{AW})q_A(W, X), \quad R_X = (I - P_{AW})X.$$

They are not interchangeable. The first residualizes only the information that survived age-supervised scalar compression; the second retains feature-resolved same-age variation. Conditional on a fixed fitted decoder and residualization rule, the data-processing inequality gives, for any named downstream variable $D$,

$$I(D; G_{\text{raw}}|A,W) = I(D; G_A|A,W) = I(D; q_A|A,W) \leq I(D; X|A,W).$$

Thus gap construction creates no additional individual information beyond the fitted scalar, chronological age and background; it cannot assign construct identity or recover information that $q_A$ discarded.

Only under squared-error loss in a common population, with a predictor class containing all square-integrable functions, is the unrestricted population target $q_A^* = P_{XW}A$. In that special case,

$$G_{\text{raw}} = (P_{XW} - I)A, \quad G_A = (I - P_{AW})P_{XW}A.$$

The second expression is a composition of conditional-expectation operators and is not generally an orthogonal projection because $P_{AW}$ and $P_{XW}$ need not commute. Restricted, penalized and finite-sample decoders must not be substituted into this population identity without qualification. If the decoder is trained under one law and centring is defined in another reference law, the correct notation is

$$G_A^{\text{ref}} = (I - P_{AW}^{\text{ref}})q_{\text{tr}}.$$

If $D$ is an observed state label, the conditional-mean state component is

$$S_D = E(X|A,W,D) - E(X|A,W) = (P_{AWD} - P_{AW})X.$$

Thus $G_A$, $R_X$ and $S_D$ are three different objects. For a fixed affine decoder $q_A = \beta^{\mathsf{T}}X + h(W)$, $G_A = \beta^{\mathsf{T}}R_X$, but even then the scalar projection does not equal $S_D$. Conditional-mean centring guarantees $E(G_A|A,W) = 0$ in the declared reference population; it does not guarantee independence from age or remove age dependence from conditional variance, tails or other distributional features.

The same identity makes scalar non-uniqueness explicit. For two observations with the same $A, W$ under a fixed affine decoder,

$$G_{A,i} - G_{A,\ell} = \beta^{\mathsf{T}}(R_{X,i} - R_{X,\ell}).$$

Equality of their centred gaps therefore imposes only one scalar constraint on their feature-resolved profiles. When the measured profile has more than one direction, differences in the null space of $\beta^{\mathsf{T}}$ remain invisible to the gap. Defining $c_{ij} = \beta_j R_{X,ij}$ gives the exact model-output attribution $G_{A,i} = \sum_j c_{ij}$. The components explain the fitted decoder calculation; they do not identify causal ageing mechanisms or biological-age components.

Now define the age- and background-residualized score

$$G_A = q_A(W,X) - m_q(A,W), \quad m_q(A,W) = E\{q_A(W,X)|A,W\},$$

where the expectation is taken in a declared reference population. This operation differs from predictive output calibration. One common output-calibration operation estimates a map from $(q_A, W)$ to $A$, involving $E(A|q_A,W)$; age residualization instead estimates $E(q_A|A,W)$.

The raw and age-residualized gaps are related by

$$G_A = G_{\text{raw}} - E(G_{\text{raw}}|A, W).$$

#### Corollary 1: age-gap construction adds no conditional individual information

For a fixed fitted decoder and residualization function, or conditional on the data used to estimate them, the score and both gaps are one-to-one transformations given $A, W$:

$$q_A = G_{\text{raw}} + A = G_A + m_q(A, W).$$

Consequently, for any named endpoint $Y$,

$$I(Y; G_{\text{raw}}|A, W) = I(Y; q_A|A, W) = I(Y; G_A|A, W).$$

Once chronological age and background are already known, subtracting age or centring within age strata adds no individual information beyond the score and those reference variables. It changes the reporting coordinate and, if chronological age is then omitted, can discard information contained jointly in $(A, q_A)$.

#### Population Bayes-risk ordering and predictive sufficiency

Let $T$ be an exactly declared target and let $\ell(d, T)$ be its declared decision loss. For an information set $\mathcal{J}$, define population Bayes risk

$$\mathcal{R}_\ell(\mathcal{J}) = \inf_{d\ \mathcal{J}\text{-measurable}} E\{\ell(d, T)\}.$$

For every deterministic score $S = q_A(X, W)$,

$$\sigma(S, A, W) \subseteq \sigma(X, A, W),$$

and therefore

$$\mathcal{R}_\ell\{\sigma(X, A, W)\} \leq \mathcal{R}_\ell\{\sigma(S, A, W)\}.$$

This classical data-processing and decision-theory ordering [Main refs. 17–19] is a population information-set result, not a claim that every finite-sample estimator fitted to a high-dimensional panel must outperform every frozen scalar. A scalar can improve finite-sample estimation through regularization, lower measurement cost or favourable transport. Each advantage is a claim about a specified target, population, sample size, estimator and resource constraint; none gives the age objective additional identifying content.

For probabilistic prediction of a named endpoint under log loss, the excess Bayes risk after scalar compression is

$$H(T|S, A, W) - H(T|X, A, W) = I(T; X|S, A, W).$$

It is zero exactly when the score retains all conditional predictive information in $X$ for that endpoint distribution. Under squared loss for a scalar target, the corresponding excess risk is

$$E[\{E(T|X, A, W) - E(T|S, A, W)\}^2].$$

Equality for one target–loss pair is only loss-specific Bayes-risk equivalence; under squared loss, for example, equality can require only equality of the relevant conditional means. Equality of the complete conditional target law is the stronger predictive-sufficiency condition $T \perp X|S, A, W$, which supports all decisions about that target. Equivalence across all decision problems of a statistical experiment is governed by Blackwell sufficiency or equivalence [Main refs. 17–19]. Chronological-age accuracy imposes none of these endpoint-specific equalities. Rank measures such as Harrell's C-index can quantify the declared discrimination comparison but are not proper-loss information quantities and should not be read as either side of the Bayes-risk bound.

#### Why unavoidable prediction error does not identify a state

For the unrestricted squared-loss target $q_A^*(W, X) = E(A|W, X)$, define $G^* = q_A^*(W, X) - A$. Then

$$E(G^*|W, X) = 0.$$

Non-zero $E\{(G^*)^2\}$ therefore states that chronological age is not a deterministic function of the observed profile and background. It does not identify the source of the conditional dispersion. In particular,

$$\mathrm{Var}(G^*) > 0 \ \not\Rightarrow \ I(G^*; H|A, W) > 0$$

for an unspecified latent state $H$. Distinct latent extensions can attach different, identical or independent versions of $H$ to the same observable law and the same residual distribution.

Two elementary constructions show that residual variation need not reflect state information, whereas source measurements can retain state information when the gap vanishes. Take $W$ to be constant. First, let $A \sim \mathrm{Uniform}(20,80)$ and let $\varepsilon \sim N(0, \sigma^2)$, with $\sigma^2 > 0$, independently of $A$. Set $X = A + \varepsilon$, with no additional state variable in the generator. The conditional age density is positive throughout (20,80) for every finite observed $X = x$, so $\mathrm{Var}(A|X) > 0$ almost surely and $\mathrm{Var}(G^*) = E\{\mathrm{Var}(A|X)\} > 0$, although no additional biological state was generated. Second, let $X = (A, D)$, where $D$ is a non-degenerate state independent of age. Then age is recovered exactly, $G^* = 0$, while $D$ remains completely observed in the source measurements.

The frozen X1–X3 construction retained in Supplementary Methods supplies a finite task map adjacent to the same geometry. X1 contains noisy age readability without a designed state term; X2 contains the declared state after exact projection off the realized intercept and linear age direction; and X3 contains both linear and nonlinear age structure together with state. Fixed out-of-fold models recover these deliberately separated task roles in the realized construction. This is explanatory rather than inferential: it does not estimate a human effect or add proof responsibility beyond the observable-law and limiting-case arguments.

#### Why a gap can contain genuine endpoint information without identifying ageing

The empirically most misleading case is not a false positive but a true association assigned the wrong construct. For two people with the same $A, W$, let $\delta$ be a local measured-profile displacement. With a fixed fitted decoder and centring rule,

$$G_A(X+\delta) - G_A(X) \simeq w_A^{\mathsf{T}}\delta, \quad w_A = \nabla_x q_A(W, X).$$

For an affine decoder the displayed relation is exact: two profiles differing by a null-space displacement $w_A^{\mathsf{T}}\delta = 0$ have the same gap. For a differentiable nonlinear decoder, this is a local first-order statement; higher-order terms can still change the score. If $w_A^{\mathsf{T}}\delta \neq 0$, the leading term describes the part of the local displacement expressed along the age-readable direction. Changes in the reference population, feature panel, loss or model can change this direction, and hence the sign or magnitude of a gap contrast, while leaving the measured-state contrast fixed. The calculation identifies how the decoder responds to a profile difference; a biological interpretation requires evidence defining that difference's meaning.

Write the measured profile as an age- and background-conditioned mean plus same-age residual variation,

$$X = \mu(A, W) + R, \quad E(R|A, W) = 0.$$

For a fixed linear decoder, or as a local approximation to a differentiable nonlinear decoder about $\mu(A, W)$,

$$G_A \simeq w_A^{\mathsf{T}}R, \quad w_A(A, W) = \nabla_x q_A\{W, \mu(A, W)\}.$$

If the residual variation of a named endpoint is represented locally by

$$Y - E(Y|A, W) = \beta_Y^{\mathsf{T}}R + \varepsilon, \quad E(\varepsilon R|A, W) = 0,$$

and $\Sigma_R(A, W) = \mathrm{Cov}(R|A, W)$, then

$$\mathrm{Cov}(G_A, Y|A, W) \simeq w_A^{\mathsf{T}}\Sigma_R\beta_Y.$$

This covariance expression is exact for the affine decoder under the stated endpoint decomposition and is a first-order approximation for the nonlinear decoder. It explains why a gap can improve endpoint prediction when age-readable variation and endpoint-relevant variation overlap. Such overlap may persist across specified samples and horizons. Linear null-space differences leave the scalar unchanged, but dependence among measured directions can still make the scalar informative about those directions; for nonlinear decoders, higher-order dependence also matters. The quantities $w_A$, $\beta_Y$ and $\Sigma_R$ can all vary with $A, W$. Thus shared biological covariance supplies a positive explanation of endpoint utility, while the identity and orientation of a biological construct require their own evidence.

The published multi-omic BMI analysis of Watanabe *et al.* (Main ref. 48) is a non-age analogue of an endpoint-associated decoder residual. Metabolomic, proteomic, clinical-chemistry and combined models reconstructed directly observed BMI, and their prediction–measurement discordances were associated with named cardiometabolic measurements and gut-microbiome features. Those results can validate the declared stratification contrasts without identifying a new BMI entity. At exact reconstruction the omics–BMI residuals would be zero although same-BMI physiological heterogeneity would remain. The different longitudinal trajectories of the modality-specific BMI scores during one lifestyle programme likewise demonstrate movement of different fitted mappings, not observation of one common latent metabolic quantity. We did not obtain or reanalyse the controlled-access Arivale or TwinsUK individual-level data; this example is based on the published article, Supplementary Data and public analysis code.

#### Corollary 2: perfect chronological-age decoding collapses the gap

Under squared-error loss,

$$E(G_{\mathrm{raw}}^2) = E[\{q_A(W,X) - A\}^2],$$

which is the population prediction risk. Thus, for any sequence of predictors whose squared-error risk converges to zero,

$$\|G_{\mathrm{raw}}\|_2 \to 0.$$

Conditional expectation is the $L^2$ projection of $q_A(W,X)$ onto functions of $(A,W)$. Because $A$ is itself an admissible function of $(A,W)$,

$$\begin{aligned} E(G_A^2) &= \min_{h(A,W)} E[\{q_A(W,X) - h(A,W)\}^2] \\ &\leq E[\{q_A(W,X) - A\}^2] = E(G_{\mathrm{raw}}^2). \end{aligned}$$

Therefore both gaps converge to zero in mean square as chronological age is recovered perfectly in mean square. This is an amplitude result: it does not imply that mutual information or endpoint utility decreases monotonically as age accuracy improves. A shrinking numerical scale can remain an invertible rescaling of the same variable at every non-zero scale. At exact recovery both gaps are identically zero. These statements establish the gap-collapse boundary without assigning a biological meaning to the residuals of imperfect predictors.

Even the population-optimal conditional-mean decoder can produce systematic age dependence in its raw gap. Assuming $E(A^2) < \infty$, omitting $W$ for clarity and taking $q_A^*(X) = E(A|X)$,

$$E(A|q_A^*) = q_A^*,$$

but generally $E(q_A^*|A) \neq A$, and

$$\mathrm{Cov}\{q_A^*(X) - A, A\} = -E\{\mathrm{Var}(A|X)\} \leq 0.$$

Thus positive raw gaps at younger ages and negative raw gaps at older ages can arise from conditional-mean shrinkage even when the population target is estimated without error. Age residualization can remove the chosen conditional mean in the reference population; it does not make the remaining quantity independent of age under all distributions or give it latent-state identity.

With imperfect prediction, a gap can combine several mathematically distinct sources. Let $m_{\mathrm{tr}} = E_{\mathrm{tr}}(A|X,W)$, let $m_{\mathrm{tar}} = E_{\mathrm{tar}}(A|X,W)$, let $q_{\mathcal{F},\lambda}^{\mathrm{tr}}$ be the penalized population target in the training law and let $\tilde{q}$ be the deployed, calibrated fitted score. Then the identity

$$\begin{aligned} \tilde{q} - A = \underbrace{(m_{\mathrm{tar}} - A)}_{\text{conditional age ambiguity}} + \underbrace{(m_{\mathrm{tr}} - m_{\mathrm{tar}})}_{\text{distribution shift}} + \underbrace{(q_{\mathcal{F},\lambda}^{\mathrm{tr}} - m_{\mathrm{tr}})}_{\text{model restriction and regularization}} \\ + \underbrace{(\hat{q} - q_{\mathcal{F},\lambda}^{\mathrm{tr}})}_{\text{estimation and tuning}} + \underbrace{(\tilde{q} - \hat{q})}_{\text{output calibration}} \end{aligned}$$

shows that an observed gap is compatible with multiple origins. Measurement error and omitted measurements enter through the observed profile and its conditional laws and need not form separately identifiable additive terms. Chronological-age accuracy alone cannot recover this decomposition or establish an intermediate accuracy regime in which the remainder acquires a privileged biological interpretation.

#### Prediction residuals are not longitudinal ageing changes

The gap decomposition concerns an inverse cross-sectional task. For a person observed at age $a$, a positive raw gap means only that the fitted reference system assigns the measured profile an age-task value above $a$; a negative gap means that it assigns a value below $a$. Because the assignment depends on $P_{\mathrm{tr}}(A|X,W)$, the same measured person can receive a different gap after the training-age distribution, sampled population, feature panel, model class, loss, regularization or calibration is changed. The terms “older-looking” and “younger-looking” are therefore shorthand for resemblance under a declared reference map, not observations of faster accumulation or reversal of a common biological process.

A longitudinal claim has a different information set and estimand. For repeated measurements, one may define a transition contrast

$$T_{\Delta} = X_{t+\Delta} - E(X_{t+\Delta}|X_t, A_t, W, U_t),$$

or a named trajectory parameter under an explicitly specified dynamic model. A risk claim requires a future endpoint $Y_{t+\Delta}$; an intervention claim requires a contrast under the assignment or causal design; clinical rejuvenation additionally requires functional, risk or benefit evidence in the favourable direction. These quantities may correlate with a clock gap, but none is identified by the age-prediction error.

The following terminology prevents three residual operations and two scientific changes from being conflated:

| Quantity | Definition | Information that generates it | What its sign can establish | What requires additional evidence |
|---|---|---|---|---|
| Conventional age-prediction residual | $A - \hat{q}(W, X)$ | chronological-age label, measured profile and fitted inverse decoder | over- or under-prediction of the known label | state, rate, risk or benefit |
| Raw clock gap | $\hat{q}(W, X) - A$ | the same information, with the opposite sign convention | reference-relative higher or lower age-task value | accelerated or decelerated ageing |
| Age-residualized clock score | $\hat{q} - E(\hat{q}|A, W)$ | fitted score plus a declared age/background reference law | position relative to the chosen conditional score mean | purification of model, measurement and biological contributions |
| Feature-resolved same-age state | $X - E(X|A, W)$ | direct measurements and an age/background reference model | signed measured deviation by feature | a latent cause or universally favourable direction |

| Quantity | Definition | Information that generates it | What its sign can establish | What requires additional evidence |
|---|---|---|---|---|
| Longitudinal transition or intervention displacement | repeated $X$, named window and, when relevant, assignment $U$ | new observations over time or under intervention | change in the declared measured object | clinical benefit or qualified surrogacy |

Chronological time is ordered, but age-prediction loss fitted to cross-sectional observations does not impose within-person monotonicity of the fitted score. If a score rises during an acute perturbation and falls during recovery, it may track that specified score response over the observed interval. Its reversibility does not, without an identified longitudinal model, make it elapsed biological time that advanced and then reversed. A non-decreasing output-calibration map cannot reverse the ordering of the uncalibrated scores, although it can introduce ties; it does not impose within-person monotonicity on the score or on the measured biological components.

#### How additional structure can identify an object and validate its measurement

The latent extension above is intentionally unrestricted because the claim under examination is whether chronological-age prediction loss itself identifies biological age. Additional restrictions may narrow the model class and identify an exact object. Examples include:

- a longitudinal transition model linking repeated measurements to a persistent latent state;
- a measurement model with declared invariances, anchors or multiple conditionally independent indicators;
- intervention restrictions that connect changes in the latent state to changes in measured outcomes;
- endpoint-specific sufficiency or transport restrictions;
- biological constraints that define the construct independently of chronological-age prediction.

Sufficient restrictions can identify the object defined by the longitudinal, measurement, intervention or endpoint model. They can also establish that an existing age-trained score is a valid measurement of that object. In either case, the added model and evidence supply the identifying relation; age loss and held-out age accuracy retain their original role as evidence for age reconstruction. The methodological requirement is to state that relation and evaluate the evidence supporting it.

A biological definition supplies identifying content when it restricts the admissible extensions and specifies the construct's relation to observations. Those restrictions must be stated and justified; identification can depend on assumptions that are not themselves testable from the age-task data. A semantic label alone leaves the observational-equivalence class unchanged.

Monotonicity alone does not close the identification problem. Chronological time is ordered, and a decoder can be required to preserve population age ordering, but this constrains the decoder rather than every biological measurement or same-age state. Even if a latent $H$ were assumed to be monotone in some declared sense, any admissible strictly increasing transformation of $H$ would preserve that ordering unless its scale and measurement relation were anchored separately. Monotonicity can therefore be one component of a narrower measurement model; it is not supplied by chronological-age accuracy and does not by itself identify the construct.

##### Scope of the proposition

The result does not assert that biological age cannot exist or that biological measurements cannot predict chronological age or a separately named endpoint. It establishes the non-implication

$$\text{chronological-age prediction} \nRightarrow \text{latent-state identity.}$$

The same boundary applies to other proposed meanings. Chronological-age prediction does not, by itself, identify a named future-risk construct, longitudinal pace, intervention response or clinical benefit. Those objects may be identified under their own data and assumptions; their meaning does not inherit from the age-decoding objective.

Chronological age records elapsed time from a declared origin. It can condition a specified population distribution, improve prediction of a named endpoint in a stated population and horizon, enter a declared effect-modification model or index endpoint-predictive variation unresolved by the current measurements. These task-specific roles do not rescue chronological-age decoding as an identification strategy for biological age.

#### Supplementary Note 2 | Contemporary construct promotion and critical responses

The two literature modules answer different questions. The construct-promotion audit asks whether the inferential operation remains present in contemporary research. The critical-literature review asks which problems have already been recognized, which repairs have been proposed and what each repair can support.

##### Contemporary construct-promotion audit

The existence audit asked whether the inferential pattern addressed by Proposition 1 remains observable in contemporary research. Source-verified examples were drawn from major general, medical, ageing and cell-biology journals to cover distinct age-unit operations and construct claims. The sampling frame was deliberately bounded and was not designed to estimate field-wide prevalence, compare journals or support a denominator-based claim.

An audit-positive main example had to satisfy three codebook conditions. First, it produced an individual-level predicted age, residual gap or intervention-associated age-valued displacement. Second, it promoted that unit to a biological, organ, cellular, immune or molecular ageing construct, an ageing rate or trajectory, or rejuvenation. Third, it lacked congruent named evidence at the same conceptual level in the estimand, training target or validation. A disease, mortality or frailty association could support a narrower endpoint prediction without identifying the broader ageing construct attached to the age unit. Boundary examples included a named endpoint, direct mechanism, longitudinal pace target or peripheral clock role; problem-diagnosing examples explicitly exposed clock heterogeneity or ambiguity. Directly named endpoint predictors, generative process parameters and longitudinal score-transition studies were therefore not treated as main examples merely because they used age-related language.

Supplementary Data 1 reports the submission examples by article title, journal, age-unit operation and highest construct-promotion level. The audit archive retains source locations and adjudication notes for reproducibility. The evidentiary statement used in the Article is existential: explicit contemporary examples instantiate the promotion chain. No ratio of selected examples is interpreted as the prevalence of the practice.

#### Critical literature and proposed remedies

The critical-literature review asked what recent critiques and methods propose in place of unqualified age-unit promotion. It comprised 17 core sources published from 2024 through 2026 and three earlier context anchors. Fifteen sources were reviewed from local full text; targeted online supplementation supplied three sources and two context anchors when local text was unavailable. For each source, we recorded the inferential or translational problem, the authors' proposed repair, remedy family, earliest promotion step addressed, the narrow object supportable if the repair succeeded and the claim that still did not follow.

The five ordered promotion steps were: biological features to an age-valued score; age score to gap or state; gap to faster or accelerated ageing; acceleration to intervention response; and response to rejuvenation or clinical benefit. The proposed remedies formed a qualification ladder rather than one replacement universal clock: target and estimand declaration; direct named-outcome prediction; reference or normative specification; repeated-measure trajectory modelling; mechanistic or generative decomposition; uncertainty and out-of-domain assessment; standardized external validation; referent and scale specificity; randomized displacement; and surrogate validation against named clinical outcomes.

The ladder is deliberately asymmetric. Repeated observations can establish a trajectory of the measured score but not its biological referent. Distribution-shift diagnostics can bound empirical support without creating ground truth. Direct outcome training can identify a predictor for the declared endpoint and horizon without identifying a common latent age. Randomization can identify treatment-induced score displacement without establishing mechanism, qualified surrogacy or benefit. Supplementary Data 2 contains the full source map, evidence locators, remedy ladder and codebook.

#### Supplementary Table 1 | Prior critiques and proposed repairs across the age-unit promotion chain

Source IDs refer to Supplementary Data 2, which contains complete citations, evidence locators and the coding fields used for this synthesis.

| Promotion link addressed | Problem recognized in prior work | Proposed repair | Evidentiary ceiling | Representative source IDs |
|---|---|---|---|---|
| Biological features → age-valued score | Chronological-age accuracy may arise from stochastic accumulation or mixtures of molecular process, composition, disease and nuisance variation; accuracy alone does not establish mechanism or health relevance. | Declare the target and context of use; separate forensic age prediction from molecular, mechanistic and clinical targets; use modality- or clock-specific names; decompose named processes, add explicit causal directionality or train directly on a named endpoint. | Supports a declared age decoder, model-dependent stochastic or molecular-process coordinate, or named endpoint predictor; does not identify a unique latent biological-age entity, causal ageing programme or clinical benefit. | R01, R02, R04, R06, R15, R18, R21 |
| Age score → gap or state | Cross-sectional marker–age association and residualization do not identify an unobserved ageing divergence; scores and gaps depend on reference population, calibration and model specification. | Specify a named endpoint, reference population, structural measurement model or normative estimand; disclose assumptions; benchmark against direct outcome models and use standardized external validation. | Supports a prespecified residual or normative deviation, a model-defined latent state, reference-dependent survival-age equivalent, or predictor for a named outcome and population; subtraction or centring does not confer construct identity. | R03, R08, R10, R12, R22 |
| Gap → faster or accelerated ageing | A single cross-sectional gap contains no within-person slope or acceleration and may have been established earlier or remain stable. | Obtain repeated measurements and model the trajectory directly, with sufficient observations, reliability and sampling cadence for the declared rate or acceleration. | Supports change or acceleration of the measured score under the declared longitudinal design; does not establish the biological referent, mechanism or beneficial meaning of that trajectory. | R13 |
| Acceleration → intervention response | Out-of-domain application, batch, composition, assay cadence, short-interval instability and cross-clock discordance can dominate intervention-associated score movement. | Quantify covariate shift, technical reproducibility, within-person stability and cross-clock disagreement; match sampling cadence to biomarker dynamics; use randomization, standardized preanalytics and orthogonal readouts. | Supports treatment-induced displacement of a declared readout, with uncertainty, within its supported domain; does not establish rejuvenation, state reversal, mechanism, qualified surrogacy or clinical benefit. | R07, R19, R23 |

| Promotion link addressed | Problem recognized in prior work | Proposed repair | Evidentiary ceiling | Representative source IDs |
|---|---|---|---|---|
| Response → rejuvenation or benefit | Responsiveness or prognostic association does not by itself establish surrogate validity, personal actionability or patient benefit. | Establish individual reliability and actionable thresholds, benchmark responsiveness, then validate the treatment → biomarker change → named clinical outcome chain across trials in a declared context of use. | Can support fit-for-purpose individual interpretation or context-specific surrogate validity for a named intervention, endpoint and population; does not establish generic rejuvenation, cross-organ reversal or benefit outside that context. | R09, R17, R20, R24 |

Main Table 1 and Supplementary Data 1 document contemporary construct promotion; Supplementary Table 1 and Supplementary Data 2 document the critical response and proposed remedy ladder. Repairs are link-specific: success at one link does not automatically authorize the next construct promotion.

Recent field-scale studies were retained as empirical context rather than treated as proof of the analytical result. In one cohort, associations between 14 DNA methylation clocks and 174 incident diseases varied across clock generations and diseases, with no clear best pan-disease clock [Main ref. 25]. A preregistered synthesis of 140 studies likewise found that associations between socioeconomic conditions and clock-derived ageing measures differed by clock generation [Main ref. 26]. OmniAge found substantial organization by biomarker class and reported that cell composition could account for selected cross-modality associations [Main ref. 27]; a same-cohort comparison of 16 ageing indicators found low-to-moderate agreement and marker-specific strengths [Main ref. 28]. A same-sample comparison of eight epigenetic clocks also documented substantial disagreement among their age-valued outputs and gaps [Main ref. 29]. These studies demonstrate heterogeneous, task- and construction-dependent association profiles. Their disagreement does not determine which, if any, corresponds to a biological-age entity.

Intervention evidence was separated by the object actually identified. TranslAGE harmonized responsiveness across 51 longitudinal intervention studies but explicitly left the relation between short-term biomarker movement and long-term disease, healthspan or lifespan unresolved [Main ref. 32]. In the randomized COSMOS ancillary study, multivitamin assignment altered two specified clock readouts, whereas cocoa extract altered none of the five tested; the clinical relevance of the clock changes remained unresolved [Main ref. 33]. These studies establish response properties or causal score displacement, not rejuvenation or clinical surrogacy. Conversely, explicit latent physiological-deterioration models with structural assumptions and mortality anchors [Main ref. 43], methylation scores supplied with putative causal directionality from Mendelian randomization [Main ref. 42], and multimodal clocks coupled to physiological capacity, compartment-specific measurements and targeted perturbations [Main ref. 41] illustrate how added information can define narrower objects. Their identifying content comes from that added information rather than the age unit.

Prior work supplies both direct identification precedents and critiques of particular uses. Sluiskes *et al.* established observational non-identifiability of the marker–biological-age relation while preserving the observed marker–age law [Main ref. 8]; Kriukov *et al.* analysed the age-clock layer and residual constructions [Main ref. 6]. Ikram and Johnson and Shokhirev challenged biological-age naming and whole-body interpretation [Main refs. 7,9], and Apsley *et al.* detailed the reliability, specificity, reference and actionability barriers to personal use [Main ref. 16]. Moqri *et al.* specified predictive, comparative and generalizability requirements for declared biomarkers [Main ref. 14]. Sehgal *et al.* separated technical reproducibility from short-interval within-person score stability [Main ref. 15], and later separated responsiveness benchmarking from still-unresolved surrogate qualification [Main ref. 32].

A closely related brain-age preprint by Grødem and colleagues combines theoretical analysis, simulations and longitudinal MRI to examine how chronological-age prediction can suppress differences in individual change [Main ref. 23]. It also proposes redirecting prediction targets towards individual change. The mismatch of training target and intended measurement is therefore a shared concern. The present Article addresses the relation among loss-specific identification, within-age orientation, gap operations, genuine prognostic success and target-specific comparison with source measurements.

Hu's preprint *House of Clocks* distinguished composite outputs from latent constructs, evaluated their utility by the intended task and argued that predictive evaluation should compare composites with their source measurements [Main ref. 24]. Its BLSA examples illustrated contrasting profiles at similar scores and age deviations, making scalar aliasing an empirical as well as a mathematical concern. Those insights are direct precedents for the present profile and source-comparison analyses. Here, the age objective is evaluated as a rule for constructing a biological measurement: loss-specific identification and within-age orientation specify what its label supplies; gap equivalence specifies what subsequent operations preserve; and target- and loss-specific risk ordering separates a population benchmark from finite-sample performance. The matched-scalar NHANES III comparison then changes the supervision target while holding source measurements, linear capacity and downstream calibration opportunity fixed. The comparisons with Hu and Grødem are targeted additions to the frozen 20-source structured review, not an expansion of its coded corpus.

The Article develops the methodological consequences of the identification problem for biomarker construction and validation. It specifies the statistical object selected by the training loss, shows how that objective evaluates same-age output differences, and makes the unresolved orientation of an age-anchored contrast explicit. The post-decoding analysis establishes what subtraction and centring preserve, while shared covariance explains why endpoint prediction can succeed. Population risk ordering then states the target- and loss-specific condition for the scalar to match the predictive risk attainable from its source measurements. Together these results require the biological question to govern representation choice and the evidence accepted as success. Structural, causal and endpoint-based alternatives provide routes to that requirement: their added evidence defines the object and can validate an existing score as its measurement. The distinction remains consequential even for accurate, reliable and predictively useful age decoders.

The human analyses make these distinctions observable in two empirical settings. HSCT separates recipient-host and donor-lineage chronological coordinates for a score-affiliation audit under a specified pairing null. In continuous NHANES, a pure age decoder added held-out mortality value while direct endpoint modelling from the same source measurements performed better; near-identical conditionally centred gaps from that decoder also arose from distinct 50-measurement contribution configurations. In survey-weighted NHANES III, a Phase 1-frozen age-supervised scalar added value beyond observed age and background, but the equally calibrated mortality-supervised scalar performed better; observed age became more useful when a pre-frozen age-selected measurement block was unavailable; and a matched direct substitution comparison found no clear advantage of the transported age decoder over known age. These results establish practical consequences for interpreting a score, selecting its supervision target and choosing measurements. Analytical proof carries the non-identification result; the human comparisons establish why the resulting methodological choices matter.

#### Supplementary Table 2 | Continuous HSCT referent-regression sensitivity

| **Study** | **Units** | $\alpha$ | $\beta$ | **HC3 95% CI** | **95% boot. CI** | $R^2$ | Residual MSE (free $\beta$) | $\beta = 0$ anchor MSE | $\beta = 1$ anchor MSE |
|---|---|---|---|---|---|---|---|---|---|
| Holland 2024 | 61 | 5.522 | 0.954 | 0.859–1.048 | 0.864–1.037 | 0.916 | 56.849 | 674.288 | 58.317 |
| Søraas 2019 | 19 | −2.622 | 1.039 | 0.930–1.148 | 0.940–1.129 | 0.969 | 26.509 | 856.982 | 27.655 |

The outcome is $S - A_R$ and the regressor is $A_L - A_R$. Because both contain the $-A_R$ anchor, common-anchor covariance can contribute to $\beta$; it is a descriptive anchor-coordinate sensitivity rather than an independent referent-identification coefficient. The `boot.` column reports the 95% independent-unit bootstrap interval. The analysis does not quantify lineage composition, identify a lineage biological age or exclude host influence. Anchor MSE values fit the intercept while fixing $\beta$ to 0 or 1. Holland provides the primary estimate; Søraas is separately processed complementary evidence. The analyses were not pooled, and the permutation null remains the primary referent-affiliation analysis.

The primary permutation holds each $(A_R, A_L)$ pair fixed and reassigns scores across independent units within the sign of $A_R - A_L$. It preserves score distributions and age-coordinate geometry while breaking the observed score pairing with both coordinates, including score–host-age dependence. Excess affiliation therefore refers to this score-pairing null. It does not estimate donor influence conditional on recipient age or exclude host-age-dependent shrinkage as a contributor. The result makes the assignment of an organismal referent to an age-valued blood readout an empirical question.

### Supplementary Methods

#### SM1. NHANES sources, ethics and cohorts

Data came from public National Health and Nutrition Examination Survey (NHANES) examination, laboratory and demographic files and the 2019 public-use linked mortality file, with follow-up through 31 December 2019 [Main Methods refs. 44,45]. NHANES protocols were approved by the relevant National Center for Health Statistics ethics review board, with informed consent obtained in the source study [Main Methods ref. 47]. The present work used de-identified public-use records and introduced no new participant contact or measurement. NCHS reports that follow-up time or underlying cause was replaced by synthetic values for selected public-use records, whereas vital status was not perturbed [Main Methods ref. 44]; the released values were used without reconstruction.

The continuous-NHANES pure-decoder and supporting matched-scalar analyses used one complete-support interface. The 2007–2008, 2009–2010 and 2011–2012 cycles supplied the fixed 50-measurement panel and at least 60 months of administrative follow-up for every event-free included participant. Participants were aged 20–79 years. Five-year death was death by 60 months. The three cycles contributed 4,599 participants and 174 deaths, 4,896 and 157, and 4,229 and 139, respectively. The pooled interface therefore contained 13,724 participants and 470 deaths, with no event-free participant censored before the horizon. These continuous-NHANES modules were complete-case and unweighted and target predictive performance in the declared analytical sample.

The survey-weighted NHANES III analyses used the canonical non-overlapping Phase 1 (1988–1991) and Phase 2 (1991–1994) interfaces frozen by the predecessor acquisition analysis. Phase 1 contained 7,558 participants and 402 resolved five-year deaths; Phase 2 contained 7,774 participants and 392. Both interfaces were restricted to examination ages 20–79 years and contained the same ordered 25-measurement clinical panel. Background comprised sex and race or ethnicity. The analysis weight was the MEC examination weight. Each phase contained 23 released strata and 46 PSU clusters; validation folds were PSU-atomic. Missing measurements were retained and handled by training-fold-local weighted-median imputation, so complete-case restriction was prohibited. The two phases support a phase-frozen protocol-transfer design within NHANES III, not an independent-cohort claim.

Phase 2 mortality had been accessed by an earlier, different panel-acquisition analysis before the three new NHANES III analyses were specified. They are therefore retrospective, outcome-accessed fixed-protocol-transfer robustness analyses. In the matched-scalar module, the two projection directions were nevertheless learned and frozen wholly in Phase 1 before being applied to Phase 2; the chronological-age-increment and observed-age-versus-decoder analyses reused fixed masks or projections but fitted their declared endpoint models in Phase 2. None is described as untouched confirmation or independent external validation.

The endpoint-directed acquisition module used its separately frozen source and target cycles, measurement menu and risk sets. Its populations are given with the corresponding methods below. Across modules, the estimands are predictive-performance contrasts for the declared analytical interfaces and estimators, not survey-population prevalence, a survey-weighted population hazard or a causal effect.

#### SM2. NHANES pure age-decoder source-information comparison

##### Frozen objects and endpoint information sets

The fixed interface contained 13,724 participants, 470 five-year deaths, ages 20–79 years and the 2007–2012 enrolment cycles. It used five prespecified outer folds, complete model inputs and the ordered 50-measurement $S_3$ panel. The endpoint, population, panel, folds and model families were fixed before this comparison.

Background $W$ comprised sex, race or ethnicity and survey cycle; $A$ was observed chronological age; $X$ was the fixed ordered 50-measurement panel; and $Y_5$ was death by 60 months. The clock $S = q_A(W, X)$ was a fold-local ridge decoder trained only to predict $A$. It received no mortality label. We compared exactly three endpoint information sets:

$$M_{WA} = P(Y_5|W, A), \quad M_{\text{CLOCK}} = P(Y_5|W, A, S), \quad M_X = P(Y_5|W, A, X).$$

Given observed $A$, the raw gap $S - A$ and $S$ are deterministic transforms of one another, so a separate gap endpoint model would not define a new information set and was not fitted.

##### Cross-fitted chronological-age decoder

Within each frozen outer training fold, categorical background was one-hot encoded and the 50 source measurements were standardized using training data only. Ridge regression predicted chronological age. Three inner folds stratified by age quintile selected the penalty by mean root-mean-square error from

$$\alpha \in \{0.01, 0.1, 1, 10, 100, 1000\}.$$

Using the selected penalty and the same inner folds, the analysis generated cross-fitted scores for every outer-training participant. A decoder fitted on the complete outer-training partition then produced scores for the held-out outer test fold. Thus the endpoint model never received an in-sample clock score, and the clock never received an endpoint label. Decoder diagnostics were held-out $R^2$, root-mean-square error, mean absolute error, Pearson correlation, observed-age-on-score calibration slope and intercept, residual summaries and age-band residuals. These quantify age readability and mean shrinkage, not biological validity.

For the pooled held-out scores, $R^2 = 0.5823$, root-mean-square error was 10.582 years and mean absolute error was 8.496 years. The observed-age-on-score slope was 0.9972 and the intercept was 0.127 years. These descriptive calibration summaries do not alter the decoder's target or establish a biological referent.

##### Endpoint models, contrasts and result boundary

For $M_{\text{CLOCK}}$, ridge logistic regression used the outer-training cross-fitted clock score, $W$ and the same five-knot fold-local cubic age spline used in the pre-existing complete-support analysis. Three inner folds selected

$$C \in \{0.001, 0.003, 0.01, 0.03, 0.1, 0.3, 1, 3, 10, 30, 100, 300\}$$

by mean validation log loss. Category encoding, score scaling, spline construction and tuning were confined to the relevant training partition. The held-out outer fold was scored by the full outer-training age decoder and then predicted by the fitted endpoint model. $M_{WA}$ and $M_X$ came from `analysis/analyze_nhanes_5y_complete_support_contextual_age_probe_v0_1.py`, where $M_X$ is stored as `M_S3A`. Both were unweighted L2-logistic models with an intercept, reference-coded sex, race or ethnicity and cycle, and a cubic age basis with five uniformly spaced training-range knots and linear extrapolation. $M_X$ additionally included all 50 transformed measurements. Measurement and age-basis columns were standardized within each training partition. Training medians were stored for preprocessing, although the accepted interface required complete observations. Within each outer fold, three stratified inner folds selected the same twelve-value inverse-penalty grid above by the arithmetic mean of fold-level validation log loss, followed by refitting on the outer-training set. Stored predictions were reused only after participant, fold and feature-order checks. The subsequent source-information entry is `analysis/analyze_nhanes_5y_age_decoder_source_information_probe_v0_1.py`, which writes the frozen v0.2 output directory. `reproducibility/ORDERED_VARIABLE_DICTIONARY.tsv` combines the ordered continuous-NHANES 50-variable and NHANES III 25-variable interfaces, preserving source variables, original units, transformations and missing-value rules.

For pooled out-of-fold log loss $L(\cdot)$, the declared contrasts were

$$V_{\text{CLOCK}} = L(M_{WA}) - L(M_{\text{CLOCK}}),$$

$$V_{\text{SOURCE}} = L(M_{\text{CLOCK}}) - L(M_X), \quad V_X = L(M_{WA}) - L(M_X).$$

The pooled endpoint results were:

| **Information set** | **Participants** | **Deaths** | **Log loss** | **Brier score** | **AUC** |
|---|---|---|---|---|---|
| $M_{WA}$ | 13,724 | 470 | 0.129791 | 0.031519 | 0.794128 |
| $M_{\text{CLOCK}}$ | 13,724 | 470 | 0.126017 | 0.030848 | 0.811423 |
| $M_X$ | 13,724 | 470 | 0.116865 | 0.029107 | 0.846111 |

Paired fixed-prediction bootstrap results were:

| **Contrast** | **Estimate** | **95% interval** |
|---|---|---|
| $V_{\text{CLOCK}}$ | 0.003774 | 0.002046–0.005474 |
| $V_{\text{SOURCE}}$ | 0.009152 | 0.006276–0.012034 |
| $V_X$ | 0.012926 | 0.009603–0.016113 |

All five fold-specific $V_{\text{CLOCK}}$ estimates were positive. All five $V_{\text{SOURCE}}$ estimates were also positive, ranging from 0.000535 to 0.014224. The paired intervals used 2,000 bootstrap resamples with indices reused from the pre-existing complete-support analysis and condition on stored out-of-fold predictions. They omit model refitting, tuning and study-development uncertainty.

The reported comparison was frozen as bounded materiality evidence without confirmation refitting or subsequent model, endpoint, panel or threshold search.

The result supports two bounded statements: one decoder trained only on chronological age added held-out predictive value for five-year all-cause mortality beyond age and background, and direct endpoint modelling from the same 50 measurements performed better under the declared finite-sample estimators. It does not estimate population information, prove universal conditional insufficiency, establish that direct models always outperform clocks, identify a true biological age or generalize beyond this decoder, panel, endpoint, horizon, population and model families.

#### SM2a. Individual contribution profiles for the frozen age decoder

This analysis reused the 13,724-participant complete-support interface, ordered 50-measurement panel, frozen outer folds, persisted ridge age decoders and stored held-out scores described in SM2. No age decoder was refitted. For each outer fold, a multivariate ridge model estimated $\hat{E}_k(X|A,W)$ from outer-training participants only, using a five-knot cubic age spline and one-hot sex, race or ethnicity and survey cycle. A single penalty was selected from $\{0.01, 0.1, 1, 10, 100, 1000\}$ by three-fold age-quintile-stratified cross-validation minimizing mean squared reconstruction error after scaling each outcome by its inner-training standard deviation. All 90 tuning estimators and five final residualizers were persisted and reloaded.

For the frozen fold-$k$ decoder, the contribution of measurement $j$ to participant $i$'s conditionally centred decoder deviation was

$$c_{ij} = \beta_{kj} \frac{X_{ij} - \hat{E}_k(X_j|A_i, W_i)}{\sigma_{kj}}.$$

Write $\hat{\mu}_k(A,W) = \hat{E}_k(X|A,W)$ for the vector of fitted measurement means. The score centre was induced by the full frozen linear decoder,

$$\widehat{m}_{S,k}(A,W) = q_{A,k}\{W, \hat{\mu}_k(A,W)\},$$

including its background encoding, training-fold scaling and intercept. It was not a separate score-on-age fit or a known population conditional expectation. The 50 contributions summed exactly to

$$G_{A,i}^{(k)} = S_i - \widehat{m}_{S,k}(A_i, W_i).$$

The field-standard raw AgeGap $G_{\mathrm{raw},i} = S_i - A_i$ was stored separately. Because matching was exact on outer fold, age and background, raw-gap and conditionally centred-gap differences were equal within each pair to numerical precision. The maximum frozen-decoder reproduction error was $1.42 \times 10^{-14}$ decoder-output years and the maximum contribution-sum error was $4.44 \times 10^{-14}$ decoder-output years.

Outcome-blind matching was performed within exact strata of outer fold, chronological age, sex, race or ethnicity and survey cycle. Participants were sorted by $G_A$; adjacent candidate edges were ordered by absolute gap difference and row identifier and retained greedily without reuse. Pairs with $|\Delta G_A| \leq 0.25$ decoder-output years were retained. The primary profile contrast was

$$d_2(i,\ell) = \{\textstyle\sum_{j=1}^{50}(c_{ij} - c_{\ell j})^2\}^{1/2}.$$

Cosine similarity and overlap between the five largest absolute contributors were secondary descriptive summaries. Mortality outcomes and endpoint-model outputs did not enter pair formation or display selection.

The algorithm retained 293 non-reusing pairs. Their median absolute conditionally centred-gap difference was 0.124 decoder-output years, whereas the median contribution-vector distance was 24.602 decoder-output years and its 95th percentile was 46.067. These distances are norms in model-contribution space, not age differences between participants. Median cosine similarity was 0.003 and median top-five-contributor Jaccard overlap was 0.429. Thus near-equality of the scalar did not imply similarity of the measured-feature contributions that produced it.

Display cases were selected before inspection of feature identities. Among 205 same-sign pairs with mean absolute conditionally centred AgeGap of at least two decoder-output years, one pair was selected closest to the median contribution distance and one closest to its 95th percentile. The union of each participant's eight largest absolute contributions was displayed; all remaining contributions were combined into an exact net remainder. This analysis makes the many-to-one geometry of one fitted age-decoder coordinate visible. Its attributions depend on the frozen model, scaling, centring reference and correlated-feature ridge parameterization; they do not assign causal ageing mechanisms, organ ages or biological-age components. The analysis used one unweighted complete-case NHANES interface and is descriptive rather than population representative.

#### SM2b. Supporting continuous-NHANES matched scalar target-supervision comparison

##### Role and frozen interface

This analysis is retained as a supporting within-interface continuous-NHANES comparison, not as the phase-frozen survey-weighted protocol-transfer result reported in SM2c. Its population, endpoint, ordered 50-measurement panel, background, model families, tuning grids, folds, seeds, metrics, bootstrap and decision rules were fixed; the fully nested construction below is the reported design.

The analysis used the same complete-support 13,724-participant interface and 470 five-year deaths as SM2. Background comprised sex, race or ethnicity and survey cycle. The source design contained background and the same ordered 50 measurements. Chronological age entered only the common downstream endpoint models in the matched comparison and was excluded from both scalar-projection inputs.

#### Flexible calibration of the age score

The original five outer folds and frozen fold-local age scores were reused. A single prespecified endpoint model added a five-knot cubic spline of the age-supervised score to background and the existing five-knot age spline. Ridge-logistic $C$ was selected inside each outer training set from $\{0.001,0.003,0.01,0.03,0.1,0.3,1,3,10,30,100,300\}$. No alternative spline or post-result calibration search was used. The fixed flexible-clock model had pooled out-of-fold log loss 0.126069, compared with 0.116865 for direct modelling of the same 50 measurements. Their difference was 0.009204 (95% fixed-prediction bootstrap interval, 0.006414–0.011941), with positive fold-specific differences in all five original folds.

#### Capacity-matched scalar construction and nesting

Two one-dimensional ridge projections were trained from identical fold-local encoded and standardized $(W, X)$ designs. The age-supervised projection $q_A$ targeted standardized chronological age; the mortality-supervised projection $q_Y$ targeted standardized five-year mortality. Both used squared-error loss, the same alpha grid $\{0.01,0.1,1,10,100,1000\}$, the same tuning folds, exclusion rules, preprocessing, linear capacity and scalar dimension. Each scalar then received the same two downstream opportunities: a linear scalar or a fixed five-knot scalar spline together with background and the same five-knot age spline. The spline contrast

$$D_{\text{TARGET}} = L(q_A \text{ spline}) - L(q_Y \text{ spline})$$

was primary; the linear contrast was diagnostic.

For every downstream validation fold, its complementary downstream-training set was defined first. Scores used to fit the downstream model were generated by three-fold projection cross-fitting wholly inside that training set; within each projection fold, alpha selection used only the projection-fold training participants. Validation scores were generated by a projection whose alpha was selected and whose coefficients were fitted only on the complete downstream-training side. Final outer-training scores were likewise projection-cross-fitted, and the outer-test scores came from projections selected and fitted on the complete outer-training set. Thus a downstream validation outcome could be used for the prespecified stratified fold assignment and then for downstream validation log loss, but it did not enter $q_Y$ preprocessing, alpha selection, fitting or training scores. Outer-test outcomes were used only for evaluation after probability generation.

Five prespecified five-fold outer partitions used seeds 20260823, 20260901, 20260902, 20260903 and 20260904, stratified by outcome and training-age quintile. The primary interval used 2,000 participant bootstraps (seed 20260905), stratified by outcome, age decade and cycle. Each participant's losses were averaged across repeats before resampling. Repeat- and cycle-specific summaries were consistency diagnostics and were not treated as independent samples or external confirmations.

##### Results, persistence and interpretation

The primary participant-averaged contrast was 0.009335 (95% participant-bootstrap interval, 0.007072–0.011666). All five repeat estimates were positive. The three enrolment-cycle estimates were also positive: 0.010361 for 2007–2008, 0.011114 for 2009–2010 and 0.006161 for 2011–2012. Linear-calibration repeat and cycle diagnostics had the same overall direction.

This supporting comparison shows that, for one continuous-NHANES analytical sample, endpoint, panel, fixed linear projection family and scalar dimension, supervision target materially changed held-out endpoint performance. It is not a protocol-transfer result, and it does not identify biological age, establish a universal mortality representation, validate a surrogate or imply causal benefit.

#### SM2c. NHANES III Phase 1-to-Phase 2 matched scalar target-supervision protocol transfer

##### Status, question and frozen interface

The reported protocol-transfer design, estimands, tuning grids, folds, survey inference and decision rules were frozen before the comparison reported below.

The fixed question was whether, given the same background, 25 measurements, one-dimensional linear capacity, Phase 1 fitting opportunity and Phase 2 endpoint calibration, a mortality-supervised scalar predicted five-year mortality better than a chronological-age-supervised scalar. This was a retrospective, outcome-accessed fixed-protocol-transfer robustness analysis because Phase 2 mortality had been accessed by an earlier panel-acquisition module. It is neither untouched confirmation nor independent external validation.

The source was the NHANES III Phase 1 interface of 7,558 participants and 402 five-year deaths; the target was the non-overlapping Phase 2 interface of 7,774 participants and 392 deaths. Both used ages 20–79 years, sex and race or ethnicity as background $W$, the same ordered 25-measurement panel $X$, MEC examination weights and the released survey design. Missing measurements were imputed by the training-local weighted median. Participant folds were PSU-atomic.

##### Phase 1 projections and operational age-decoder gate

Two ridge projections used identical Phase 1 $(W, X)$ inputs, encoding, weighted-median imputation, weighted standardization, normalized MEC fitting weights, squared-error loss, alpha grid $\{0.01, 0.1, 1, 10, 100, 1000\}$, folds and scalar dimension. Chronological age was not an input. The sole intended difference was the supervised label: $q_A$ targeted standardized chronological age and $q_Y$ targeted standardized five-year mortality. For the binary label, $q_Y$ is a mortality-supervised linear probability projection, not a latent mortality entity or a biological-age estimate.

Five fixed Phase 1 outer folds generated held-out scores. Final alphas were selected in the fixed complete-Phase 1 inner folds, after which one final $q_A$ and one final $q_Y$ object were fitted on complete Phase 1 and applied unchanged to Phase 2. Phase 2 outcomes did not enter projection preprocessing, tuning or fitting. Design-weighted Phase 1 out-of-fold age mean-squared error was 94.695287 for $q_A$, compared with 252.163340 for the otherwise matched $W$-only comparator. Transported Phase 2 $q_A$ age root-mean-squared error was 9.380644 years and $R^2$, relative to the frozen Phase 1 weighted-mean predictor, was 0.645207. These quantities establish age readability and transport diagnostics, not biological validity.

#### Equal Phase 2 endpoint calibration and estimands

The two frozen scores received identical target-local calibration in the fixed Phase 2 outer folds. We compared

$$M_0: Y \sim W + s(A), \quad M_A: Y \sim W + s(A) + s(q_A), \quad M_Y: Y \sim W + s(A) + s(q_Y).$$

The age basis used fixed knots at 20, 35, 50, 65 and 80 years. Each score used a training-local five-knot uniform cubic spline with linear extrapolation. Weighted L2-penalized logistic regression used the common grid $\{0.001, 0.003, 0.01, 0.03, 0.1, 0.3, 1, 3, 10, 30, 100, 300\}$, selected by pooled design-weighted inner-validation log loss. Every reported Phase 2 probability was outer-fold held out. Target-local fitting calibrated the two already frozen directions equally and did not relearn either projection.

For design-weighted participant-level outer-fold log loss $L(\cdot)$, the primary matched contrast was

$$\Delta = L(M_A) - L(M_Y),$$

where a positive value favours mortality supervision. The supporting age-decoder increment was

$$U_A = L(M_0) - L(M_A).$$

A positive $U_A$ distinguishes genuine downstream utility of $q_A$ from the stronger and unsupported claim that age supervision should have default representational privilege. A linear-score calibration was prespecified as a diagnostic; the spline-score comparison was primary.

#### Results, survey uncertainty and claim boundary

| Phase 2 information set | Design-weighted out-of-fold log loss |
|---|---|
| $M_0$: background and observed age | 0.124955173 |
| $M_A$: background, observed age and frozen $q_A$ | 0.122163092 |
| $M_Y$: background, observed age and frozen $q_Y$ | 0.116451991 |

The mortality-supervised scalar outperformed the age-supervised scalar: $\Delta = 0.005711101$, with a 95% Fay-BRR interval of 0.002093219–0.009328984 and positive fixed-fold contrasts in all five outer folds. The age-supervised scalar itself added held-out value beyond observed age and background, $U_A = 0.002792081$ (0.000239446–0.005344715). The prespecified linear-calibration diagnostic also favoured mortality supervision, with a point contrast of 0.004619814.

Each of 32 Phase 2 Fay-BRR replicates (ρ=0.3; 23 design degrees of freedom) repeated downstream inner tuning, outer fitting, held-out prediction and paired-loss evaluation under replicate weights; the Phase 1 projections remained fixed. The intervals therefore represent Phase 2 survey-design uncertainty conditional on those two Phase 1-fitted directions and omit source-model development uncertainty.

This result supports a bounded protocol-transfer claim: for this population, five-year endpoint, 25-measurement interface and one-dimensional linear projection family, chronological-age pretraining was unnecessary for constructing an endpoint-useful scalar and did not yield the best scalar direction for mortality. It does not identify biological age, establish universal mortality sufficiency, compare every clock, validate a surrogate, imply causal benefit or make the mortality-supervised score a general biological entity.

#### SM3. NHANES III observed chronological-age increment and predictive proxy substitution

##### Status, frozen block and four endpoint models

This analysis asked how much held-out predictive value observed chronological age added after the complete 25-measurement panel and whether that value increased when a previously age-selected measurement block was unavailable. It did not use an age-decoder score, endpoint-supervised scalar, age gap or any derived-age input. This was a retrospective, outcome-accessed fixed-protocol-transfer robustness analysis, not untouched confirmation or independent external validation.

The target was the canonical survey-weighted NHANES III Phase 2 interface of 7,774 participants and 392 resolved five-year deaths. Background $W$ comprised sex and race or ethnicity. The full panel $X_{25}$ comprised all 25 ordered measurements. The removed block was taken unchanged from the signed Phase 1 source-selection freeze, which selected the cardiorespiratory domain for age reconstruction before Phase 2 mortality was revealed in that predecessor selection stage. In manifest order the block was mean systolic blood pressure (`MEC_SBP_MEAN`), mean diastolic blood pressure (`MEC_DBP_MEAN`), radial pulse (`PEP6DR`), forced expiratory volume in one second (`SPPFEV1`) and forced vital capacity (`SPPFVC`). Removing exactly those five measurements yielded the ordered remaining-20 panel $X_{20R}$.

The mask was not revised in response to any Phase 2 outcome, coefficient, missingness pattern or result in this module. Selection occurred at the domain level: it does not establish that each removed variable is individually age-associated, and the remaining 20 measurements must not be described as lacking age-related information.

We fitted four target-local endpoint models:

$$F_0\colon Y \sim W + X_{25}, \quad F_1\colon Y \sim W + X_{25} + s(A),$$

$$R_0\colon Y \sim W + X_{20R}, \quad R_1\colon Y \sim W + X_{20R} + s(A).$$

Here $A$ was observed examination age and $s(A)$ was a cubic basis with fixed knots at ages 20, 35, 50, 65 and 80, no bias column and linear extrapolation. Within every training fold and weight set, background was encoded, measurements were weighted-median imputed and weighted-standardized, and the age basis for $F_1$ and $R_1$ was weighted-standardized. Weighted L2-penalized logistic regression used the common $C$ grid $\{0.001, 0.003, 0.01, 0.03, 0.1, 0.3, 1, 3, 10, 30, 100, 300\}$. Selection used pooled design-weighted log loss in the fixed three inner folds; every reported probability came from a held-out PSU-atomic outer fold.

#### Estimands and survey uncertainty

Let $L(M)$ denote the design-weighted mean participant-level outer-fold log loss. The two chronological-age increments were

$$V_F = L(F_0) - L(F_1), \quad V_R = L(R_0) - L(R_1).$$

The predictive proxy-substitution contrast was

$$P_{\text{proxy}} = V_R - V_F = \{L(R_0) - L(F_0)\} - \{L(R_1) - L(F_1)\}.$$

A positive value means that observed age became more predictively useful after the Phase 1 age-selected block was removed, equivalently that supplying age attenuated part of the held-out loss penalty from omitting that block under the declared estimators. This is an information-set-dependent predictive substitution pattern, not causal mediation or evidence that age is the biological state represented by the omitted measurements.

Three secondary contrasts were the block value without age, $D_{\text{block},0} = L(R_0) - L(F_0)$; the block value with age, $D_{\text{block},1} = L(R_1) - L(F_1)$; and the replacement comparison, $D_{\text{replace}} = L(R_1) - L(F_0)$. Failure of either latter interval to exclude zero cannot establish equality, equivalence, non-inferiority or complete replacement because no clinical margin was frozen.

For each of 32 Phase 2 Fay-BRR factors (ρ=0.3; 23 design degrees of freedom), the complete preprocessing, inner tuning, outer fitting, held-out prediction and paired-loss pipeline was rerun for all four arms. Intervals therefore include Phase 2 survey-design uncertainty conditional on the frozen mask, folds and estimator family. They do not include Phase 1 mask-development or broader study-development uncertainty.

#### Results, persistence and claim boundary

| Model | Inputs | Design-weighted out-of-fold log loss |
|---|---|---|
| $F_0$ | Background and all 25 measurements | 0.122980792 |
| $F_1$ | Background, all 25 measurements and observed age | 0.119664249 |
| $R_0$ | Background and remaining-20 panel | 0.128973446 |
| $R_1$ | Background, remaining-20 panel and observed age | 0.120087795 |

#### Paired predictive contrasts

| Contrast | Estimate | 95% Fay-BRR interval | Positive fixed outer folds |
|---|---|---|---|
| Full-panel age increment $V_F$ | 0.003316543 | −0.000966027 to 0.007599114 | 5 of 5 |
| Remaining-20 age increment $V_R$ | 0.008885651 | 0.003308686 to 0.014462615 | 5 of 5 |
| Predictive proxy contrast $P_{\text{proxy}}$ | 0.005569107 | 0.001925424 to 0.009212791 | 4 of 5 |
| Block value without age $D_{\text{block},0}$ | 0.005992654 | 0.001217282 to 0.010768026 | 4 of 5 |
| Block value with age $D_{\text{block},1}$ | 0.000423547 | −0.002053797 to 0.002900890 | 3 of 5 |
| Replacement comparison $D_{\text{replace}}$ | −0.002892997 | −0.007934182 to 0.002148189 | 0 of 5 |

The full-panel point estimate favoured adding observed age in all five folds, but its design interval crossed zero; incremental prediction after the complete 25-measurement panel was therefore not established under this comparison. After removal of the Phase 1 age-selected block, the age increment was larger and its interval excluded zero. The positive $P_{\text{proxy}}$ interval establishes the declared information-set-dependent predictive substitution pattern. The omitted block itself had positive predictive value without age, whereas its residual value after age and the reduced-plus-age versus full-without-age comparison were unresolved. In particular, neither crossing-zero secondary contrast establishes equivalence or replacement.

The result quantifies modest proper-score contrasts for one population, endpoint, mask and estimator family. It may be described as observed age attenuating part of the held-out penalty from omitting a pre-frozen age-selected block. It must not be described as a percentage or fraction of ageing, causal mediation, mechanism, biological-age identity, clinical equivalence, universal measured-state sufficiency or proof that the remaining-20 panel lacks age-related information. No post hoc recovery ratio was a frozen estimand or is reported.

#### SM3a. Observed chronological age versus a transported age decoder: substitution boundary

This analysis directly compared observed chronological age with the Phase 1-fitted $q_A$ from SM2c as alternative one-dimensional inputs for Phase 2 five-year mortality prediction after the same background adjustment. It did not ask whether either scalar added beyond the other, and neither model contained both. This was a retrospective, outcome-accessed fixed-protocol-transfer robustness analysis, not untouched confirmation. It is retained as a bounded diagnostic and carries no independent analytical proof responsibility.

The two target models were

$$M_{\mathrm{AGE}}: Y \sim W + s(A), \quad M_{Q_A}: Y \sim W + s(q_A).$$

Both arms used the same 7,774 participants and 392 deaths, MEC weights, PSU-atomic folds, one continuous scalar, training-local weighted standardization, five-knot uniform cubic spline with linear extrapolation, L2-logistic model, $C$ grid and design-weighted tuning rule. The Phase 1 $q_A$ direction remained frozen. The primary contrast was

$$\Delta_{\mathrm{sub}} = L(M_{\mathrm{AGE}}) - L(M_{Q_A}),$$

so positive values favour $q_A$ and negative values favour observed age. Each of the 32 Fay-BRR replicates repeated target-side tuning, fitting and held-out prediction for both arms without refitting $q_A$.

The observed-age arm is not numerically identical to $M_0$ in SM2c: this direct substitution comparison gave observed age and $q_A$ the same training-local uniform-spline opportunity, whereas SM2c retained its prespecified fixed chronological-age knot locations while testing incremental score utility.

Design-weighted out-of-fold log loss was 0.124777611 for observed age and 0.125534785 for $q_A$. The contrast was −0.000757174, with a 95% Fay-BRR interval of −0.004661445 to 0.003147097. One of five fixed-fold contrasts was positive and four were negative.

This boundary does not show equality, equivalence or non-inferiority, because no such margin was specified. It establishes neither that $q_A$ can replace known chronological age nor that observed age is generally superior to age decoders. It applies only to this transported Phase 1 decoder, Phase 2 population, endpoint and matched one-scalar model family. Together with SM2c, it shows that $q_A$ was an operational age decoder and could add endpoint value when supplied alongside age, while reconstructing age did not yield a clearly superior substitute for the known age itself.

#### SM4. Endpoint-directed versus inverse age-decoder acquisition analysis

##### Cross-cycle measurement acquisition

The sole endpoint was five-year all-cause mortality. The source cycle was NHANES 2007–2008 (4,599 complete participants; 174 deaths) and the target cycle was 2009–2010 (4,896; 157 deaths). The five candidate acquisition domains and feature manifests were fixed as renal/mineral, metabolic/adiposity/lipid, hepatic/protein/injury, haematology and haemodynamic measurements. Each policy selected one domain from the same five prespecified measurement-domain options. Domain measurement counts differed and no resource matching was imposed.

Within the source cycle, domain value was the reduction in pooled out-of-fold log loss when the actual domain was added to background plus age:

$$V_k = L_{\mathrm{OOF}}(W + A) - L_{\mathrm{OOF}}(W + A + M_k).$$

For the inverse chronological-age policy, each domain $M_k$ entered a ridge decoder $A \sim W + M_k$, where $W$ comprised sex and race or ethnicity. Encoding, declared feature transformations and standardization were training-fold local. Five age-quintile-stratified outer folds (seed 20260826) supplied participant-level out-of-fold predictions. Within each outer training fold, the ridge penalty was selected from $\{0.01, 0.1, 1, 10, 100, 1000\}$ by pooled mean-squared error across three age-quintile-stratified inner folds (seed $20260827 + \text{outer fold}$). The inverse-decoder policy selected the domain with the lowest pooled source out-of-fold age mean-squared error; the separately frozen endpoint policy selected $\text{argmax}_k V_k$. Source choices and the source-data signature were frozen before target evaluation, and target outcomes did not enter either source selection.

In the target cycle, we compared the held-out five-year mortality log-loss gains of the two source-frozen domains beyond age and background. Participant identity, fold assignment and ordered features matched the sealed predecessor. Uncertainty used 5,000 event-stratified paired participant bootstraps of fixed out-of-fold losses (seed 20260802), without policy reselection or model refitting. The target cycle had been viewed during earlier reconnaissance and is therefore exploratory cross-cycle evidence rather than historically untouched confirmation.

The endpoint-directed source policy selected hepatic, protein and injury measurements; the inverse age-decoder policy selected metabolic, adiposity and lipid measurements. The selected inverse decoder had pooled source out-of-fold age mean-squared error 177.923601, root-mean-squared error 13.338801 years, mean absolute error 10.889224 years and $R^2 = 0.342481$. In the target cycle, the endpoint-directed policy improved log loss beyond age and background by 0.006418 (95% fixed-prediction bootstrap interval, 0.002297–0.010652), whereas the inverse-decoder policy improved it by 0.000646 (−0.001195–0.002481). Their paired endpoint-minus-inverse difference was 0.005772 (0.001380–0.010390), with positive point estimates in four of five outer folds. These participant-averaged proper-score differences apply only to the declared menu, cycles and model families. They are not clinical value of information, net benefit, economic or resource-equivalence analysis, universal transportability or biological-age identification.

#### SM5. Transparent three-variable age–state task map

##### Evidential role and simulated population

This construction makes one task separation visible: noisy age readability, declared same-age state information and a mixture of the two are different properties of measured variables. It is not a model of human ageing, a disease-prevalence estimate, a prediction benchmark or a second proof of Proposition 1.

We retained X1–X3 from a frozen cross-sectional construction of 500 rows generated with seed 20260806. Chronological age was sampled uniformly from 20 to 80 years and a binary separation state $D$ with probability 0.40, yielding 299 State-0 and 201 State-1 rows. These observations are conditional cross-sectional draws; their conditional means are not within-person trajectories.

#### Declared feature geometry

Let $z(\cdot)$ denote standardization over the realized sample, $A_z$ standardized chronological age, $D_c$ centred state, $Q_A$ the standardized quadratic age component after projection off the linear age direction and $\varepsilon_j$ independent standard-normal noise. The retained variables were

$$X_1 = z(0.95A_z + 0.65\varepsilon_1),$$

$$X_2 = z\{\Pi_{\perp(1,A)}(1.35D_c + 0.55\varepsilon_2)\},$$

and

$$X_3 = z(0.65A_z + 0.85Q_A + 0.90D_c + 0.55\varepsilon_3),$$

where $\Pi_{\perp(1,A)}$ removes the realized intercept and linear age direction. X1 therefore supplied a noisy age-readable direction without a designed state term. X2 supplied declared state information while being exactly linearly age-orthogonal in the realized construction. X3 mixed linear and nonlinear age structure with state information. No biological interpretation was assigned to the generated state.

#### Fixed task-specific models

We assigned rows to five participant-disjoint outer folds using stratified random splitting with shuffle seed 20260827. The stratification variable crossed the declared binary state with five equal-frequency age groups. Fold assignments were fixed and persisted before model fitting. No alternative split, model-family search, tuning or threshold search was permitted.

Chronological-age decoding used four fixed ridge pipelines with X1, X2, X3 or X1–X3 as the measurement arm. Each measurement entered a five-knot cubic spline, followed by training-fold standardization and ridge regression with $\alpha = 1$. We report pooled five-fold out-of-fold $R^2$, root-mean-squared error and mean absolute error.

Declared-state decoding used five fixed L2-logistic pipelines: age alone, or age plus X1, X2, X3 or X1–X3. Chronological age entered through a five-knot cubic spline; measurements were standardized within each training fold. Logistic regression used $C = 1$, the `lbfgs` solver and a maximum of 2,000 iterations. The primary state quantity was the reduction in pooled out-of-fold log loss relative to the age-only model. Positive values mean that the declared measurement contributed held-out state prediction beyond the fixed age spline.

#### Fixed-model results

| **Measurement arm** | Age decoding: pooled OOF $R^2$ | **Age RMSE (years)** | **State log-loss gain beyond age** | **State ROC AUC** | **State log loss** |
|---|---|---|---|---|---|
| X1 | 0.680817 | 9.69424 | −0.002139 | 0.525117 | 0.674069 |
| X2 | −0.015023 | 17.2875 | 0.406866 | 0.955557 | 0.265064 |
| X3 | 0.300915 | 14.3469 | 0.219921 | 0.863093 | 0.452009 |
| X1–X3 | 0.729573 | 8.92316 | 0.483124 | 0.977138 | 0.188805 |

X1 was strongly age-readable but contributed no held-out state value beyond age under the declared models. X2 showed the converse task pattern: its age-decoding $R^2$ was slightly below zero, while its declared-state gain was large. X3 contributed to both tasks, and the joint X1–X3 arm supported both age decoding and state decoding (Extended Data Fig. 1). These quantities describe one frozen construction; they are not population estimates and are not interpreted as biological-information fractions.

The analysis reused the frozen 500-row source and did not regenerate any row.

##### Logical limiting cases and interpretation boundary

The finite construction is accompanied by the two limiting cases specified in Supplementary Note 1, with constant background. The uniform-age, independent-Gaussian-noise construction has positive population age-residual variance without an additional generated state. Conversely, when $X = (A, D)$ with non-degenerate $D$ independent of age, the gap is zero while the state remains completely observed in the source measurements. These cases—not the fitted task map—show that residual variation does not establish state information, and that a vanishing gap does not imply its absence from the source measurements. Panels a–c of Extended Data Fig. 1 plot different individuals once against a cross-sectional chronological-age coordinate; connected bin means are not within-person trajectories.

#### SM6. Leakage control and statistical reporting

The independent unit depended on the declared system. For X1–X3, one constructed row was the independent unit; fixed five-fold out-of-fold results supplied no population interval. HSCT used the recipient or donor–recipient pair as declared in its analysis. Predictive NHANES analyses contained one retained record per human participant. For the individual contribution summary, one retained non-reusing matched pair was the descriptive unit. Prespecified magnitude gates were decision rules rather than population-hypothesis tests.

For outer-fold evaluation, fitted operations excluded the corresponding outer-test participants. This applied to scaling, categorical encoding, imputation, spline construction, cross-fitted score generation, calibration and penalty selection. Resampling followed the module-specific fixed-prediction or refitting procedures described below. In continuous NHANES, the pure chronological-age decoder received no mortality labels, and its endpoint model received only training-side cross-fitted scores. The individual contribution module reloaded those frozen fold-local decoders and estimated $\hat{E}_k(X|A, W)$ on the complementary outer-training participants; pair formation and display selection used no mortality outcome or endpoint-model output. The supporting continuous-NHANES matched-scalar comparison used fully nested projection scores inside every downstream training partition.

For the NHANES III matched-scalar protocol transfer, the $q_A$ and $q_Y$ directions were learned and frozen in Phase 1; Phase 2 outcomes did not enter their preprocessing, tuning or fitting. Phase 2 endpoint calibration was identical across the two directions. The chronological-age-increment comparison used a Phase 1-selected domain mask fixed before any new Phase 2 model in that analysis, and no Phase 2 result could alter it. The observed-age-versus-$q_A$ comparison reused the same frozen Phase 1 $q_A$. These freezes prevent target-side relearning of the selected direction or mask, but the analyses remain retrospective and outcome-accessed because Phase 2 mortality had been accessed elsewhere before their specifications.

Uncertainty procedures followed the declared design. The continuous-NHANES pure-decoder and supporting matched-scalar intervals used 2,000 paired participant bootstraps of stored out-of-fold losses; those fixed-prediction intervals omit refitting, tuning and study-development uncertainty. The contribution-profile analysis was descriptive: all 293 retained non-reusing pairs entered its empirical summaries, and no hypothesis test or population interval was attached to that pair distribution. The cross-cycle acquisition analysis used 5,000 event-stratified fixed-prediction participant bootstraps. The survey-weighted NHANES III modules used the same 32 Phase 2 Fay-BRR factors with $\rho = 0.3$ and 23 design degrees of freedom. Every Fay-BRR replicate repeated the declared Phase 2 preprocessing, inner tuning, outer fitting, held-out prediction and paired-loss calculation. The matched-scalar module held the two Phase 1 projection directions fixed; the chronological-age-increment module refitted all four Phase 2 arms; and the observed-age-versus-$q_A$ module refitted both downstream arms while holding $q_A$ fixed. These intervals represent Phase 2 survey-design uncertainty conditional on the relevant Phase 1 objects, folds and estimator families. They do not include Phase 1 selection or projection-development uncertainty or broader study-development uncertainty.

No universal significance threshold was used to assign construct identity. Positive predictive contrasts were interpreted only for the declared model, cohort, endpoint and validation design. An interval crossing zero means that the corresponding increment or difference was not established under that design; it is not evidence that the associated biology is absent. Conversely, failure to distinguish two predictive losses does not establish equality, equivalence, non-inferiority, full substitution or replacement when no margin was prespecified. None of the reported proper-score differences is a clinical effect size, literal conditional mutual information, causal mediation estimand or fraction of ageing.

#### SM7. Source data and reproducibility

The two literature workbooks have distinct evidentiary responsibilities. Supplementary Data 1 and Main Table 1 contain the contemporary construct-promotion audit, including tier definitions, construct levels and source-level claim locators. Supplementary Data 2 and Supplementary Table 1 contain the critical-literature map, proposed remedies and their identification ceilings. These are interpretive literature Source Data rather than model outputs, and neither contributes to the mathematical proof.

The retained human analyses used frozen participant interfaces, ordered features, folds and declared estimators. Continuous-NHANES and NHANES III interfaces were analysed separately; Figure 3 labels their distinct populations and no estimates were pooled across them. For the individual contribution module, the internal project evidence directory retains participant-level raw and conditionally centred gaps, the complete participant-by-feature contribution matrix, all matched-pair metrics, ordered features, residualizer objects and reload-parity records; the source age decoders are referenced there by their frozen paths and hashes. The submission-facing Source Data contain aggregate pair metrics and the two de-identified display-pair contribution profiles. Supplementary Software 1 contains specifications, code, manifests and a stored-result audit layer comprising frozen primary participant-level predictions, survey-replicate estimates, all matched-pair profiles and the complete participant-by-feature contribution matrix. Fitted-model binaries and large resampling-index arrays remain internally retained rather than included in the submission archive. The HSCT analyses used public reported scores or author-processed beta matrices under the processing and imputation boundaries stated above. No raw-IDAT reprocessing was performed.

All reported fitted analyses were reconstructed after reload from persisted data, folds, preprocessing and model objects, with participant identity, fold assignment, ordered features and predictions matching the reported results at the prespecified tolerances. Fit-free figure builders read only frozen Source Data or conceptual inputs. Analyses ran in the project's pinned `lhstate` environment under the declared environment manifest and exact platform lock.

The software snapshot and figure-level Source Data described above are supplied for editorial and reviewer evaluation. Public deposition and reuse-licensing details remain to be finalized before publication. The portable verification entry point recalculates results from stored predictions, contributions and resampling summaries without refitting models. It also checks the flexible-calibration and repeated matched-scalar supporting contrasts from individual probabilities, averaging matched-scalar losses within participant across repeats. Their bootstrap intervals remain saved summaries because the original index archive is not included. For age-decoder substitution, the package supplies primary individual probabilities and 32 saved Fay-BRR replicate estimates for point-estimate and interval arithmetic; replicate-level prediction and fitting chains are not reexecuted. The snapshot includes the continuous-NHANES source-interface and baseline/direct-source model generators, with their execution order and input inventory in CORE_NHANES_REPRODUCTION.md. It also documents fold-construction rules and selected acquisition fold-assignment tables. Figure reconstruction, recalculation from stored predictions and source-to-model reexecution are distinct levels of reproducibility; end-to-end external reexecution has not been verified. Other modules can require the retained predecessor interfaces and fitted objects referenced by their manifests. NHANES participant records remain available from the originating public-use files under their terms. A prespecified Hannum-2013 sensitivity was not run because the complete-file transfer criterion was not met; no Hannum score or effect estimate was computed.

Exact script routes, output directories, internal decision records, artifact inventories, version histories and figure-resource mappings are indexed in `REPRODUCIBILITY_MANIFEST_v6_27_20260914.md`, which indexes the frozen scientific resources and their current publication mappings. These repository records support audit and rerun without adding scientific proof responsibility to the Supplementary Information.

### Supplementary reference note

References are keyed to the current main-manuscript sequence:

- `[Main ref. 36]` — published PhenoAge specification;
- `[Main ref. 8]` — Sluiskes observational-equivalence and identical-association predecessor;
- `[Main ref. 40]` — AccelerAge endpoint-first and reference-dependent age-equivalent framework;
- `[Main ref. 12]` — stochastic-accumulation clock construction;
- `[Main ref. 13]` — empirical stochastic CpG component analysis;
- `[Main ref. 20]` — BEST biomarker and surrogate framework;
- `[Main ref. 48]` — published multi-omic BMI analogue;
- `[Main ref. 21]` — Søraas post-transplant blood study;
- `[Main ref. 22]` — Holland post-transplant blood study;
- `[Main Methods ref. 44]` — public-use linked mortality documentation;
- `[Main Methods ref. 45]` — NHANES survey and public-use data documentation;
- `[Main Methods ref. 47]` — NHANES ethics-review-board approvals;
- `[Main ref. 17]` — Blackwell comparison and equivalence of experiments;
- `[Main ref. 18]` — data-processing inequality and information-theoretic sufficiency; and
- `[Main ref. 19]` — conditional independence in statistical theory.
