## Supplementary_Software_1 for "Inferential boundaries of age prediction: why prediction does not establish biological age measurement": analyze_holland2024_horvath_object_affiliation_v0_1.py

```
"""Holland 2024 Horvath 2013 lineage-versus-host object-affiliation analysis."""

from __future__ import annotations

import hashlib
import json
import math
import platform
import sys
from pathlib import Path
from typing import Callable

import matplotlib as mpl
import matplotlib.pyplot as plt
import numpy as np
import pandas as pd
import scipy
from scipy import stats


ROOT = Path(__file__).resolve().parents[1]
SPEC = (
    ROOT
    / "analysis"
    / "HSCT_HOLLAND2024_HORVATH2013_OBJECT_AFFILIATION_SPEC_20260730.md"
)
SCORE_ROOT = ROOT / "outputs" / "hsct_holland2024_horvath2013_v0_1"
BETA_CACHE = (
    ROOT
    / "data"
    / "processed"
    / "hsct_holland2024_horvath2013_target_betas_v0_1"
)
COEFFICIENTS = (
    ROOT / "data" / "processed" / "horvath2013_frozen_coefficients_v0_1.tsv"
)
PROVENANCE_AUDIT = (
    SCORE_ROOT / "combined_six_batch_provenance_audit_v0_1.json"
)
OUT = ROOT / "outputs" / "hsct_holland2024_horvath_object_affiliation_v0_1"
REPORT = (
    ROOT
    / "reports"
    / "HSCT_HOLLAND2024_HORVATH_OBJECT_AFFILIATION_V0_1_20260730.md"
)
FIGURE_PNG = (
    ROOT / "figures" / "HSCT_HOLLAND2024_HORVATH_OBJECT_AFFILIATION_V0_1.png"
)
FIGURE_PDF = (
    ROOT / "figures" / "HSCT_HOLLAND2024_HORVATH_OBJECT_AFFILIATION_V0_1.pdf"
)

SEED = 20260730
N_BOOT = 10_000
TIE_TOLERANCE = 1e-12
PRIMARY_GAP = 10.0
PRIMARY_MIN_TIME = 0.1
HORVATH_INTERCEPT = 0.696

SCORE_HASHES = {
    1: "881eb9906c9fd9390b14eea3eb9f0bafd2bd16fb29803731cad770cd8064934e",
    2: "6f097a37b91e3520d06a51642968ae6b2e2a4d0acd97d6d36512b228e743c48b",
    3: "bd7b8bcfc6275cdd3098c5c6119ed959d807dd45f7d5fcf1c934d37d937a293e",
    4: "ad0ad84173f24f628fdf2945f507bcc72ac592bfab0ad2ae0b11e29590ef5421",
    5: "0113543409f37651b932e493821395198ea8cda703992e69d283a161254fbbf6",
    6: "b766c686d4fbb3cba38d8d9318bebfe4f3960adc2d409c34362ed8cebd82ee78",
}
BETA_CACHE_HASHES = {
    1: "950c8b2d125d18574a3d19ec12c5999aa4393986d150a35a5658c974a0ef0b69",
    2: "3abcd8edd28d2e3308140f72193708f72381aaf5516f1837d9a30e4c2cf87cf5",
    3: "1da5acb61cf079250b7fe621de490a3fdbd26dbe57a8ee76a3551f3c4f0fa231",
    4: "f6e190f80b671fc3e74d60c4f54829c4b3da6f0df0c923082edb4c6479811462",
    5: "0e2257667981a29e31a243eca9a17afd0328170d1af50f37c496ff63f79d45fc",
    6: "f9578d2888dab61c5530be82edc6bbf2ea2e460d3395b51120d2194690510473",
}
COEFFICIENT_HASH = (
    "fb499a0c8b63632c2a78b6e9871b6533a0209564d8305e3ea6ffef29321804ca"
)
PROVENANCE_AUDIT_HASH = (
    "9084163c9f2f1ac41117585b8d794961a6ba00ea8cc776032edcc5e31565ef43"
)
SPEC_HASH = "85c5f514a26c62ffe6897c36d5525c99ef2ebf13fe3ba05770809f869014d6f5"

LINEAGE_COLOR = "#087E8B"
HOST_COLOR = "#C45A35"
SCORE_COLOR = "#3B334F"
HOST_YOUNGER_COLOR = "#D97706"
NEUTRAL = "#6B7280"
LIGHT = "#E5E7EB"

OCCASION_KEYS = [
    "pair_id",
    "role",
    "AgeCells",
    "AgeDonorAtTX",
    "AgeRecipAtTX",
    "YrsAfterReceivingTX",
]


def sha256(path: Path) -> str:
    digest = hashlib.sha256()
    with path.open("rb") as handle:
        for block in iter(lambda: handle.read(1024 * 1024), b""):
            digest.update(block)
    return digest.hexdigest()


def json_ready(value: object) -> object:
    if isinstance(value, dict):
        return {str(key): json_ready(item) for key, item in value.items()}
    if isinstance(value, (list, tuple)):
        return [json_ready(item) for item in value]
    if isinstance(value, (np.integer,)):
        return int(value)
    if isinstance(value, (np.floating,)):
        return None if not np.isfinite(value) else float(value)
    if isinstance(value, float):
        return None if not math.isfinite(value) else value
    if isinstance(value, Path):
        return str(value)
    return value


def verify_inputs() -> pd.DataFrame:
    rows = []
    observed_spec = sha256(SPEC)
    if observed_spec.lower() != SPEC_HASH:
        raise RuntimeError("Frozen Holland analysis specification hash mismatch")
    rows.append(
        {
            "role": "frozen_spec",
            "path": str(SPEC.relative_to(ROOT)),
            "sha256": observed_spec,
            "bytes": SPEC.stat().st_size,
        }
    )
    observed_coefficients = sha256(COEFFICIENTS)
    if observed_coefficients.lower() != COEFFICIENT_HASH:
        raise RuntimeError("Frozen Horvath coefficient export hash mismatch")
    rows.append(
        {
            "role": "frozen_horvath2013_coefficient_export",
            "path": str(COEFFICIENTS.relative_to(ROOT)),
            "sha256": observed_coefficients,
            "bytes": COEFFICIENTS.stat().st_size,
        }
    )
    observed_provenance = sha256(PROVENANCE_AUDIT)
    if observed_provenance.lower() != PROVENANCE_AUDIT_HASH:
        raise RuntimeError("Combined six-batch provenance audit hash mismatch")
    provenance = json.loads(PROVENANCE_AUDIT.read_text(encoding="utf-8"))
    if (
        provenance.get("status")
        != "PASS_WITH_DECLARED_METHOD_QUALIFICATIONS"
        or provenance.get("totals", {}).get("arrays") != 334
        or provenance.get("checks", {}).get(
            "common_probes_observed_in_all_six_batches"
        )
        != 253
    ):
        raise RuntimeError("Combined six-batch provenance audit did not pass")
    rows.append(
        {
            "role": "combined_six_batch_provenance_audit",
            "path": str(PROVENANCE_AUDIT.relative_to(ROOT)),
            "sha256": observed_provenance,
            "bytes": PROVENANCE_AUDIT.stat().st_size,
        }
    )
    for batch, expected in SCORE_HASHES.items():
        path = SCORE_ROOT / f"batch{batch}_horvath2013_scores.tsv"
        if not path.is_file():
            raise FileNotFoundError(path)
        observed = sha256(path)
        if observed.lower() != expected:
            raise RuntimeError(f"Frozen score hash mismatch for batch {batch}")
        rows.append(
            {
                "role": f"batch{batch}_scores",
                "path": str(path.relative_to(ROOT)),
                "sha256": observed,
                "bytes": path.stat().st_size,
            }
        )
        stream_audit = SCORE_ROOT / f"batch{batch}_stream_audit.json"
        if not stream_audit.is_file():
            raise FileNotFoundError(stream_audit)
        audit = json.loads(stream_audit.read_text(encoding="utf-8"))
        if (
            audit.get("sha256_expected") != audit.get("sha256_observed")
            or audit.get("sample_mapping") != "PASS"
            or audit.get("n_nonfinite_target_values") != 0
            or audit.get("n_out_of_range_target_values") != 0
        ):
            raise RuntimeError(f"Source stream audit failed for batch {batch}")
        rows.append(
            {
                "role": f"batch{batch}_remote_stream_audit",
                "path": str(stream_audit.relative_to(ROOT)),
                "sha256": sha256(stream_audit),
                "bytes": stream_audit.stat().st_size,
            }
        )
        beta_path = BETA_CACHE / f"batch{batch}_horvath353.tsv.gz"
        if not beta_path.is_file():
            raise FileNotFoundError(beta_path)
        observed_beta = sha256(beta_path)
        if observed_beta.lower() != BETA_CACHE_HASHES[batch]:
            raise RuntimeError(f"Frozen target-beta cache hash mismatch for batch {batch}")
        rows.append(
            {
                "role": f"batch{batch}_target_beta_cache",
                "path": str(beta_path.relative_to(ROOT)),
                "sha256": observed_beta,
                "bytes": beta_path.stat().st_size,
            }
        )
    return pd.DataFrame(rows)


def add_derived_metrics(arrays: pd.DataFrame) -> pd.DataFrame:
    arrays = arrays.copy()
    arrays["host_age"] = arrays["AgeRecipAtTX"] + arrays["YrsAfterReceivingTX"]
    # For recipient samples, the public AgeCells variable equals
    # AgeDonorAtTX + YrsAfterReceivingTX (maximum floating-point discrepancy
    # below 1e-12). It is a donor-lineage chronological coordinate, not the
    # literal chronological age of the short-lived cells present at sampling.
    arrays["donor_lineage_coordinate"] = arrays["AgeCells"]
    arrays["host_minus_lineage_coordinate"] = (
        arrays["host_age"] - arrays["donor_lineage_coordinate"]
    )
    arrays["absolute_object_gap"] = arrays["host_minus_lineage_coordinate"].abs()
    arrays["abs_error_to_lineage_coordinate"] = (
        arrays["horvath2013_score"] - arrays["donor_lineage_coordinate"]
    ).abs()
    arrays["abs_error_to_host"] = (
        arrays["horvath2013_score"] - arrays["host_age"]
    ).abs()
    arrays["affiliation_index"] = np.where(
        arrays["absolute_object_gap"] > 0,
        (
            arrays["abs_error_to_host"]
            - arrays["abs_error_to_lineage_coordinate"]
        )
        / arrays["absolute_object_gap"],
        np.nan,
    )
    return arrays


def load_arrays() -> pd.DataFrame:
    frames = []
    for batch in range(1, 7):
        frame = pd.read_csv(SCORE_ROOT / f"batch{batch}_horvath2013_scores.tsv", sep="\t")
        if not frame["batch"].eq(batch).all():
            raise RuntimeError(f"Internal batch mismatch in batch {batch} scores")
        frames.append(frame)
    arrays = pd.concat(frames, ignore_index=True)
    if len(arrays) != 334 or arrays["randID"].nunique() != 334:
        raise RuntimeError("Expected 334 unique public arrays")
    if arrays["horvath2013_score"].isna().any():
        raise RuntimeError("Missing frozen Horvath score")
    return add_derived_metrics(arrays)


def aggregate_occasions(arrays: pd.DataFrame) -> pd.DataFrame:
    occasions = (
        arrays.groupby(OCCASION_KEYS, dropna=False, sort=False)
        .agg(
            horvath2013_score=("horvath2013_score", "mean"),
            horvath2013_score_min=("horvath2013_score", "min"),
            horvath2013_score_max=("horvath2013_score", "max"),
            n_arrays=("randID", "size"),
            randIDs=("randID", lambda values: ",".join(sorted(map(str, values)))),
            batches=("batch", lambda values: ",".join(map(str, sorted(set(values))))),
        )
        .reset_index()
    )
    occasions["host_age"] = (
        occasions["AgeRecipAtTX"] + occasions["YrsAfterReceivingTX"]
    )
    occasions["donor_lineage_coordinate"] = occasions["AgeCells"]
    occasions["host_minus_lineage_coordinate"] = (
        occasions["host_age"] - occasions["donor_lineage_coordinate"]
    )
    occasions["absolute_object_gap"] = (
        occasions["host_minus_lineage_coordinate"].abs()
    )
    occasions["score_minus_lineage_coordinate"] = (
        occasions["horvath2013_score"]
        - occasions["donor_lineage_coordinate"]
    )
    occasions["abs_error_to_lineage_coordinate"] = (
        occasions["score_minus_lineage_coordinate"].abs()
    )
    occasions["abs_error_to_host"] = (
        occasions["horvath2013_score"] - occasions["host_age"]
    ).abs()
    occasions["affiliation_index"] = np.where(
        occasions["absolute_object_gap"] > 0,
        (
            occasions["abs_error_to_host"]
            - occasions["abs_error_to_lineage_coordinate"]
        )
        / occasions["absolute_object_gap"],
        np.nan,
    )
    occasions["duplicate_score_range"] = (
        occasions["horvath2013_score_max"] - occasions["horvath2013_score_min"]
    )
    occasions["occasion_id"] = np.arange(1, len(occasions) + 1)
    if (occasions["affiliation_index"].abs().dropna() > 1 + 1e-10).any():
        raise RuntimeError("Affiliation index escaped its mathematical [-1,1] bound")
    return occasions


def primary_filter(occasions: pd.DataFrame, gap: float = PRIMARY_GAP) -> pd.DataFrame:
    return occasions.loc[
        occasions["role"].eq("R")
        & occasions["YrsAfterReceivingTX"].ge(PRIMARY_MIN_TIME)
        & occasions["absolute_object_gap"].ge(gap)
    ].copy()


def pair_summary(
    occasions: pd.DataFrame, reducer: str = "mean"
) -> pd.DataFrame:
    if reducer not in {"mean", "median"}:
        raise ValueError(reducer)
    function: str | Callable = reducer
    pairs = (
        occasions.groupby("pair_id", sort=False)
        .agg(
            n_occasions=("occasion_id", "size"),
            affiliation_index=("affiliation_index", function),
            abs_error_to_lineage_coordinate=(
                "abs_error_to_lineage_coordinate",
                function,
            ),
            abs_error_to_host=("abs_error_to_host", function),
            donor_lineage_coordinate=("donor_lineage_coordinate", function),
            host_age=("host_age", function),
            host_minus_lineage_coordinate=(
                "host_minus_lineage_coordinate",
                function,
            ),
            score_minus_lineage_coordinate=(
                "score_minus_lineage_coordinate",
                function,
            ),
            mean_time_after_hsct=("YrsAfterReceivingTX", "mean"),
            batches=("batches", lambda values: ",".join(sorted(set(",".join(values).split(","))))),
        )
        .reset_index()
    )
    pairs["direction"] = np.select(
        [
            pairs["affiliation_index"] > TIE_TOLERANCE,
            pairs["affiliation_index"] < -TIE_TOLERANCE,
        ],
        ["lineage_side", "host_side"],
        default="tie",
    )
    return pairs


def bootstrap_mean(values: np.ndarray) -> np.ndarray:
    rng = np.random.default_rng(SEED)
    n = len(values)
    indices = rng.integers(0, n, size=(N_BOOT, n))
    return values[indices].mean(axis=1)


def summarize_pairs(
    pairs: pd.DataFrame,
    label: str,
    n_occasions: int,
    selection_note: str,
    analysis_class: str = "prespecified",
) -> tuple[dict[str, object], np.ndarray]:
    if len(pairs) == 0:
        raise RuntimeError(f"No pairs in sensitivity {label}")
    values = pairs["affiliation_index"].to_numpy(dtype=float)
    boot = bootstrap_mean(values)
    low, high = np.quantile(boot, [0.025, 0.975], method="linear")
    non_tied = pairs.loc[pairs["direction"].ne("tie")]
    n_positive = int(non_tied["direction"].eq("lineage_side").sum())
    exact = stats.binomtest(
        n_positive, len(non_tied), p=0.5, alternative="two-sided"
    )
    result = {
        "analysis": label,
        "analysis_class": analysis_class,
        "selection_note": selection_note,
        "n_occasions": int(n_occasions),
        "n_pairs": int(len(pairs)),
        "mean_affiliation_index": float(values.mean()),
        "ci_low": float(low),
        "ci_high": float(high),
        "median_affiliation_index": float(np.median(values)),
        "n_positive_pair_mean": n_positive,
        "n_negative_pair_mean": int(
            non_tied["direction"].eq("host_side").sum()
        ),
        "n_ties": int(pairs["direction"].eq("tie").sum()),
        "proportion_positive_among_non_ties": float(
            n_positive / len(non_tied)
        ),
        "binomial_p_two_sided": float(exact.pvalue),
        "mean_abs_error_to_lineage_coordinate": float(
            pairs["abs_error_to_lineage_coordinate"].mean()
        ),
        "mean_abs_error_to_host": float(pairs["abs_error_to_host"].mean()),
    }
    return result, boot


def choose_one_per_pair(
    occasions: pd.DataFrame, criterion: str
) -> pd.DataFrame:
    selected = occasions.copy()
    if criterion == "earliest":
        selected["_order"] = selected["YrsAfterReceivingTX"]
    elif criterion == "closest_to_one_year":
        selected["_order"] = (selected["YrsAfterReceivingTX"] - 1).abs()
    else:
        raise ValueError(criterion)
    return (
        selected.sort_values(["pair_id", "_order", "YrsAfterReceivingTX"])
        .groupby("pair_id", as_index=False, sort=False)
        .head(1)
        .drop(columns="_order")
    )


def anti_trafo(linear_predictor: np.ndarray) -> np.ndarray:
    """Apply the frozen Horvath 2013 adult-age inverse transform."""
    return np.where(
        linear_predictor < 0,
        21 * np.exp(linear_predictor) - 1,
        21 * linear_predictor + 20,
    )


def common_probe_harmonized_arrays(
    arrays: pd.DataFrame,
) -> tuple[pd.DataFrame, dict[str, object]]:
    """Re-score every batch using the same observed-probe set.

    This post-specification sensitivity removes the possibility that the
    primary pattern is driven by different whole-batch missing-probe sets. A
    probe is used as observed only if it was observed in all six batches; all
    other probes are set to the same fixed external mean in every batch.
    """
    coefficients = pd.read_csv(COEFFICIENTS, sep="\t")
    if (
        len(coefficients) != 353
        or coefficients["probe_id"].nunique() != 353
        or coefficients[["coefficient", "training_mean"]].isna().any().any()
    ):
        raise RuntimeError("Unexpected frozen Horvath coefficient export")

    observed_sets: list[set[str]] = []
    missing_counts: dict[str, int] = {}
    for batch in range(1, 7):
        coverage = pd.read_csv(
            SCORE_ROOT / f"batch{batch}_probe_coverage.tsv", sep="\t"
        )
        if len(coverage) != 353 or coverage["probe_id"].nunique() != 353:
            raise RuntimeError(f"Unexpected probe coverage for batch {batch}")
        observed = set(
            coverage.loc[coverage["status"].eq("observed"), "probe_id"]
        )
        observed_sets.append(observed)
        missing_counts[str(batch)] = int(353 - len(observed))
    common_observed = set.intersection(*observed_sets)
    if len(common_observed) != 253:
        raise RuntimeError(
            f"Expected 253 probes observed in all batches, got {len(common_observed)}"
        )

    coefficient_order = coefficients["probe_id"].tolist()
    common_mask = coefficients["probe_id"].isin(common_observed).to_numpy()
    weights = coefficients["coefficient"].to_numpy(dtype=float)
    external_means = coefficients["training_mean"].to_numpy(dtype=float)
    score_rows: list[pd.DataFrame] = []
    for batch in range(1, 7):
        beta_frame = pd.read_csv(
            BETA_CACHE / f"batch{batch}_horvath353.tsv.gz",
            sep="\t",
            compression="gzip",
        )
        if (
            len(beta_frame) != 353
            or beta_frame["probe_id"].nunique() != 353
            or set(beta_frame["probe_id"]) != set(coefficient_order)
        ):
            raise RuntimeError(f"Unexpected target-beta cache for batch {batch}")
        beta_frame = beta_frame.set_index("probe_id").loc[coefficient_order]
        sample_ids = beta_frame.columns.tolist()
        beta = beta_frame.to_numpy(dtype=float)
        beta[~common_mask, :] = external_means[~common_mask, None]
        if not np.isfinite(beta).all():
            raise RuntimeError(
                f"Non-finite common-probe harmonized beta in batch {batch}"
            )
        linear_predictor = HORVATH_INTERCEPT + weights @ beta
        score_rows.append(
            pd.DataFrame(
                {
                    "randID": sample_ids,
                    "common_probe_harmonized_score": anti_trafo(
                        linear_predictor
                    ),
                }
            )
        )

    harmonized_scores = pd.concat(score_rows, ignore_index=True)
    if (
        len(harmonized_scores) != 334
        or harmonized_scores["randID"].nunique() != 334
    ):
        raise RuntimeError("Common-probe harmonized score mapping failed")
    harmonized = arrays.drop(
        columns=[
            column
            for column in arrays.columns
            if column
            in {
                "host_age",
                "donor_lineage_coordinate",
                "host_minus_lineage_coordinate",
                "absolute_object_gap",
                "abs_error_to_lineage_coordinate",
                "abs_error_to_host",
                "affiliation_index",
            }
        ]
    ).merge(harmonized_scores, on="randID", how="left", validate="one_to_one")
    if harmonized["common_probe_harmonized_score"].isna().any():
        raise RuntimeError("Missing common-probe harmonized score")
    harmonized["horvath2013_score"] = harmonized[
        "common_probe_harmonized_score"
    ]
    harmonized = add_derived_metrics(harmonized)
    audit = {
        "analysis_class": "post_spec_processing_sensitivity",
        "n_total_probes": 353,
        "n_observed_in_all_six_batches": int(len(common_observed)),
        "n_set_to_fixed_external_mean_in_all_batches": int(
            353 - len(common_observed)
        ),
        "whole_batch_missing_probe_counts": missing_counts,
        "external_mean_source": (
            "PC-Clocks cgHorvathNew.csv overallMeanByCpGacross50data"
        ),
    }
    return harmonized, audit


def negative_time_equivalence_check(arrays: pd.DataFrame) -> pd.DataFrame:
    """Verify the primary filter is invariant to near-zero negative-time handling."""

    def selected_randids(frame: pd.DataFrame) -> tuple[set[str], int, int]:
        selected = primary_filter(aggregate_occasions(frame))
        randids: set[str] = set()
        for value in selected["randIDs"]:
            randids.update(value.split(","))
        return randids, int(len(selected)), int(selected["pair_id"].nunique())

    variants: list[tuple[str, pd.DataFrame]] = [("observed", arrays)]
    truncated = arrays.copy()
    truncated["YrsAfterReceivingTX"] = truncated["YrsAfterReceivingTX"].clip(
        lower=0
    )
    variants.append(("truncate_negative_to_zero", add_derived_metrics(truncated)))
    excluded = arrays.loc[arrays["YrsAfterReceivingTX"].ge(0)].copy()
    variants.append(("exclude_negative", add_derived_metrics(excluded)))

    reference_ids, _, _ = selected_randids(arrays)
    rows: list[dict[str, object]] = []
    for label, frame in variants:
        ids, n_occasions, n_pairs = selected_randids(frame)
        rows.append(
            {
                "negative_time_handling": label,
                "n_input_arrays": int(len(frame)),
                "n_negative_time_arrays": int(
                    (frame["YrsAfterReceivingTX"] < 0).sum()
                ),
                "n_primary_occasions": n_occasions,
                "n_primary_pairs": n_pairs,
                "same_primary_randids_as_observed": bool(
                    ids == reference_ids
                ),
                "primary_randids_sha256": hashlib.sha256(
                    "\n".join(sorted(ids)).encode("utf-8")
                ).hexdigest(),
            }
        )
    return pd.DataFrame(rows)


def sensitivity_analyses(
    arrays: pd.DataFrame, occasions: pd.DataFrame
) -> tuple[pd.DataFrame, pd.DataFrame, np.ndarray]:
    summaries: list[dict[str, object]] = []
    primary = primary_filter(occasions)
    primary_pairs = pair_summary(primary, reducer="mean")
    primary_result, primary_boot = summarize_pairs(
        primary_pairs,
        "primary_gap_ge_10_all_occasions_pair_mean",
        len(primary),
        "R role; time >=0.1 y; |host-lineage coordinate| >=10 y; pair mean",
    )
    summaries.append(primary_result)

    for gap in (1e-12, 5.0, 15.0, 20.0):
        selected = primary_filter(occasions, gap=gap)
        label_gap = "gt_0" if gap < 1e-6 else f"ge_{int(gap)}"
        result, _ = summarize_pairs(
            pair_summary(selected),
            f"gap_{label_gap}",
            len(selected),
            (
                "R role; time >=0.1 y; |host-lineage coordinate| "
                f"{label_gap.replace('_', ' ')} y"
            ),
        )
        summaries.append(result)

    for criterion in ("earliest", "closest_to_one_year"):
        selected = choose_one_per_pair(primary, criterion)
        result, _ = summarize_pairs(
            pair_summary(selected),
            criterion,
            len(selected),
            f"Primary population; one {criterion.replace('_', ' ')} occasion per pair",
        )
        summaries.append(result)

    for direction, mask in (
        (
            "host_older_than_lineage_coordinate",
            primary["host_minus_lineage_coordinate"] > 0,
        ),
        (
            "host_younger_than_lineage_coordinate",
            primary["host_minus_lineage_coordinate"] < 0,
        ),
    ):
        selected = primary.loc[mask].copy()
        result, _ = summarize_pairs(
            pair_summary(selected),
            direction,
            len(selected),
            f"Primary population restricted to {direction.replace('_', ' ')}",
        )
        summaries.append(result)

    result, _ = summarize_pairs(
        pair_summary(primary, reducer="median"),
        "pair_median_across_occasions",
        len(primary),
        "Primary population; pair median rather than pair mean",
    )
    summaries.append(result)

    for omitted_batch in range(1, 7):
        retained_arrays = arrays.loc[arrays["batch"].ne(omitted_batch)].copy()
        retained_occasions = primary_filter(aggregate_occasions(retained_arrays))
        result, _ = summarize_pairs(
            pair_summary(retained_occasions),
            f"leave_batch_{omitted_batch}_out",
            len(retained_occasions),
            f"Primary population after omitting every array in batch {omitted_batch}",
        )
        summaries.append(result)

    return pd.DataFrame(summaries), primary_pairs, primary_boot


def plot_figure(
    primary_pairs: pd.DataFrame,
    primary_result: dict[str, object],
    sensitivities: pd.DataFrame,
) -> None:
    mpl.rcParams.update(
        {
            "font.family": "DejaVu Sans",
            "font.size": 9,
            "axes.titlesize": 10,
            "axes.labelsize": 9,
            "axes.spines.top": False,
            "axes.spines.right": False,
            "pdf.fonttype": 42,
            "ps.fonttype": 42,
        }
    )
    fig, axes = plt.subplots(
        1,
        3,
        figsize=(14.2, 5.2),
        gridspec_kw={"width_ratios": [1.15, 0.95, 1.15]},
        constrained_layout=True,
    )
    ax_a, ax_b, ax_c = axes

    # Pair-level geometry: lineage-coordinate alignment y=0; host alignment y=x.
    host_older = primary_pairs["host_minus_lineage_coordinate"] > 0
    x = primary_pairs["host_minus_lineage_coordinate"]
    y = primary_pairs["score_minus_lineage_coordinate"]
    limit = float(max(np.abs(x).max(), np.abs(y).max()) + 3)
    line_x = np.linspace(-limit, limit, 200)
    ax_a.axhline(
        0,
        color=LINEAGE_COLOR,
        lw=1.6,
        label="Lineage-coordinate referent: score − C = 0",
    )
    ax_a.plot(
        line_x,
        line_x,
        color=HOST_COLOR,
        lw=1.4,
        ls="--",
        label="Host-age referent: score − C = H − C",
    )
    for mask, color, label in (
        (host_older, HOST_COLOR, "Host older than lineage coordinate"),
        (~host_older, HOST_YOUNGER_COLOR, "Host younger than lineage coordinate"),
    ):
        ax_a.scatter(
            x.loc[mask],
            y.loc[mask],
            color=color,
            s=28,
            alpha=0.82,
            edgecolor="white",
            linewidth=0.4,
            label=label,
        )
    ax_a.set_xlim(-limit, limit)
    ax_a.set_ylim(-limit, limit)
    ax_a.set_xlabel("Host age − donor-lineage coordinate (H − C)")
    ax_a.set_ylabel("Horvath score − donor-lineage coordinate (S − C)")
    ax_a.set_title("a  Which chronological coordinate does recipient blood align with?")
    ax_a.grid(color="#F3F4F6", lw=0.8)
    ax_a.legend(frameon=False, fontsize=6.7, loc="upper left")

    # Every pair's bounded affiliation index.
    ordered = primary_pairs.sort_values("affiliation_index").reset_index(drop=True)
    y_index = np.arange(len(ordered))
    colors = np.where(
        ordered["affiliation_index"] > TIE_TOLERANCE,
        LINEAGE_COLOR,
        HOST_COLOR,
    )
    ax_b.axvline(0, color="#111827", lw=0.9)
    ax_b.hlines(y_index, 0, ordered["affiliation_index"], color=LIGHT, lw=1.1)
    ax_b.scatter(
        ordered["affiliation_index"],
        y_index,
        c=colors,
        s=19,
        edgecolor="white",
        linewidth=0.35,
        zorder=3,
    )
    mean_y = len(ordered) + 2
    ax_b.plot(
        [primary_result["ci_low"], primary_result["ci_high"]],
        [mean_y, mean_y],
        color=SCORE_COLOR,
        lw=2.4,
        solid_capstyle="round",
    )
    ax_b.scatter(
        [primary_result["mean_affiliation_index"]],
        [mean_y],
        marker="D",
        color=SCORE_COLOR,
        s=48,
        edgecolor="white",
        linewidth=0.5,
        zorder=4,
    )
    ax_b.text(
        -1.02,
        mean_y,
        "Mean (95% CI)",
        va="center",
        ha="right",
        fontsize=8,
    )
    ax_b.set_xlim(-1.05, 1.05)
    ax_b.set_yticks([])
    ax_b.set_xlabel(
        r"Side-affiliation $I=(|S-H|-|S-C|)/|H-C|$"
        "\n−1 = host side; +1 = lineage-coordinate side"
    )
    ax_b.set_title(
        "b  Pair-level object affiliation\n"
        f"{primary_result['n_positive_pair_mean']} positive, "
        f"{primary_result['n_ties']} ties, "
        f"{primary_result['n_negative_pair_mean']} negative pair means"
    )
    ax_b.grid(axis="x", color="#F3F4F6", lw=0.8)

    # Sensitivity forest.
    shown = sensitivities.loc[
        ~sensitivities["analysis"].str.startswith("leave_batch_")
    ].copy()
    shown = shown.iloc[::-1].reset_index(drop=True)
    y_sens = np.arange(len(shown))
    ax_c.axvline(0, color="#111827", lw=0.9)
    prespecified = shown["analysis_class"].eq("prespecified")
    for mask, color, marker, label in (
        (prespecified, LINEAGE_COLOR, "o", "Prespecified"),
        (~prespecified, HOST_YOUNGER_COLOR, "s", "Post-spec processing"),
    ):
        ax_c.hlines(
            y_sens[mask],
            shown.loc[mask, "ci_low"],
            shown.loc[mask, "ci_high"],
            color=color,
            lw=2,
        )
        ax_c.scatter(
            shown.loc[mask, "mean_affiliation_index"],
            y_sens[mask],
            color=color,
            marker=marker,
            s=32,
            edgecolor="white",
            linewidth=0.45,
            zorder=3,
            label=label,
        )
    labels = (
        shown["analysis"]
        .str.replace("primary_gap_ge_10_all_occasions_pair_mean", "Primary", regex=False)
        .str.replace(
            "common_probe_harmonized_external_mean",
            "Common-probe harmonized",
            regex=False,
        )
        .str.replace("gap_", "Age gap ", regex=False)
        .str.replace("_", " ", regex=False)
    )
    ax_c.set_yticks(y_sens)
    ax_c.set_yticklabels(labels)
    ax_c.set_xlim(-0.05, 1.05)
    ax_c.set_xlabel("Mean side-affiliation index (pair-bootstrap 95% CI)")
    ax_c.set_title("c  Population, occasion and processing sensitivities")
    ax_c.grid(axis="x", color="#F3F4F6", lw=0.8)
    ax_c.legend(frameon=False, fontsize=7, loc="lower right")

    fig.suptitle(
        (
            "After HSCT, recipient-blood Horvath 2013 scores aligned more closely "
            "with the donor-lineage chronological coordinate"
        ),
        x=0.01,
        ha="left",
        fontsize=13.5,
        fontweight="bold",
    )
    FIGURE_PNG.parent.mkdir(parents=True, exist_ok=True)
    fig.savefig(FIGURE_PNG, dpi=320, bbox_inches="tight", facecolor="white")
    fig.savefig(FIGURE_PDF, bbox_inches="tight", facecolor="white")
    plt.close(fig)


def markdown_table(frame: pd.DataFrame, digits: int = 4) -> str:
    if frame.empty:
        return "_No rows._"
    display = frame.copy()
    for column in display.select_dtypes(include=[np.number]).columns:
        display[column] = display[column].map(
            lambda value: "" if not np.isfinite(value) else f"{value:.{digits}g}"
        )
    lines = [
        "| " + " | ".join(display.columns) + " |",
        "| " + " | ".join(["---"] * len(display.columns)) + " |",
    ]
    lines.extend(
        "| "
        + " | ".join(
            str(value).replace("\n", " ").replace("|", r"\|")
            for value in row
        )
        + " |"
        for row in display.itertuples(index=False, name=None)
    )
    return "\n".join(lines)


def main() -> None:
    OUT.mkdir(parents=True, exist_ok=True)
    REPORT.parent.mkdir(parents=True, exist_ok=True)
    input_manifest = verify_inputs()
    arrays = load_arrays()
    occasions = aggregate_occasions(arrays)

    if len(occasions.loc[occasions["role"].eq("R")]) != 264:
        raise RuntimeError(
            "Expected 264 conservatively aggregated recipient occasions"
        )
    sensitivities, primary_pairs, primary_boot = sensitivity_analyses(
        arrays, occasions
    )
    primary_result = sensitivities.iloc[0].to_dict()
    primary_occasions = primary_filter(occasions)
    harmonized_arrays, harmonized_audit = common_probe_harmonized_arrays(arrays)
    harmonized_occasions = aggregate_occasions(harmonized_arrays)
    harmonized_primary_occasions = primary_filter(harmonized_occasions)
    harmonized_primary_pairs = pair_summary(harmonized_primary_occasions)
    harmonized_result, _ = summarize_pairs(
        harmonized_primary_pairs,
        "common_probe_harmonized_external_mean",
        len(harmonized_primary_occasions),
        (
            "Post-spec processing sensitivity: retain observed values only for "
            "the 253 probes present in all six batches; set the other 100 probes "
            "to the same fixed external mean in every batch"
        ),
        analysis_class="post_spec_processing_sensitivity",
    )
    sensitivities = pd.concat(
        [sensitivities, pd.DataFrame([harmonized_result])],
        ignore_index=True,
    )
    negative_time_check = negative_time_equivalence_check(arrays)

    if primary_result["n_pairs"] < 50:
        raise RuntimeError("Primary population has fewer than 50 independent pairs")
    if primary_result["ci_low"] <= 0:
        raise RuntimeError(
            "Primary side-affiliation confidence interval crosses zero"
        )
    if harmonized_result["ci_low"] <= 0:
        raise RuntimeError(
            "Common-probe harmonized confidence interval crosses zero"
        )
    if not negative_time_check["same_primary_randids_as_observed"].all():
        raise RuntimeError(
            "Primary filter changed under negative-time handling sensitivity"
        )

    donor_occasions = occasions.loc[occasions["role"].eq("D")].copy()
    donor_corr = stats.pearsonr(
        donor_occasions["horvath2013_score"], donor_occasions["AgeCells"]
    )
    duplicate_audit = occasions.loc[occasions["n_arrays"].gt(1)].copy()
    leave_batch = sensitivities.loc[
        sensitivities["analysis"].str.startswith("leave_batch_")
    ].copy()
    source_bytes_total = int(
        sum(
            json.loads(
                (SCORE_ROOT / f"batch{batch}_stream_audit.json").read_text(
                    encoding="utf-8"
                )
            )["bytes_observed"]
            for batch in range(1, 7)
        )
    )
    stop_checks = {
        "all_six_batches_present": bool(set(arrays["batch"]) == set(range(1, 7))),
        "n_arrays": int(len(arrays)),
        "n_unique_randID": int(arrays["randID"].nunique()),
        "n_recipient_arrays": int(arrays["role"].eq("R").sum()),
        "n_donor_arrays": int(arrays["role"].eq("D").sum()),
        "n_recipient_occasions": int(occasions["role"].eq("R").sum()),
        "n_primary_occasions": int(len(primary_occasions)),
        "n_primary_pairs": int(len(primary_pairs)),
        "primary_ci_above_zero": bool(primary_result["ci_low"] > 0),
        "all_leave_batch_out_ci_above_zero": bool((leave_batch["ci_low"] > 0).all()),
        "common_probe_harmonized_ci_above_zero": bool(
            harmonized_result["ci_low"] > 0
        ),
        "negative_time_handling_identical_under_primary_filter": bool(
            negative_time_check[
                "same_primary_randids_as_observed"
            ].all()
        ),
        "max_recipient_agecells_vs_lineage_formula_abs_difference": float(
            (
                arrays.loc[arrays["role"].eq("R"), "AgeCells"]
                - (
                    arrays.loc[arrays["role"].eq("R"), "AgeDonorAtTX"]
                    + arrays.loc[
                        arrays["role"].eq("R"), "YrsAfterReceivingTX"
                    ]
                )
            )
            .abs()
            .max()
        ),
        "max_duplicate_occasion_score_range": float(
            duplicate_audit["duplicate_score_range"].max()
        ),
        "n_negative_time_arrays": int((arrays["YrsAfterReceivingTX"] < 0).sum()),
        "source_bytes_streamed": source_bytes_total,
    }

    input_manifest.to_csv(OUT / "input_manifest.tsv", sep="\t", index=False)
    arrays.to_csv(OUT / "analysis_arrays.tsv.gz", sep="\t", index=False, compression="gzip")
    occasions.to_csv(
        OUT / "analysis_occasions.tsv.gz", sep="\t", index=False, compression="gzip"
    )
    primary_occasions.to_csv(
        OUT / "primary_occasions.tsv", sep="\t", index=False
    )
    primary_pairs.to_csv(OUT / "primary_pair_summary.tsv", sep="\t", index=False)
    sensitivities.to_csv(OUT / "sensitivity_summary.tsv", sep="\t", index=False)
    harmonized_arrays[
        ["randID", "batch", "common_probe_harmonized_score"]
    ].to_csv(
        OUT / "common_probe_harmonized_scores.tsv",
        sep="\t",
        index=False,
    )
    harmonized_primary_pairs.to_csv(
        OUT / "common_probe_harmonized_primary_pair_summary.tsv",
        sep="\t",
        index=False,
    )
    (OUT / "common_probe_harmonization_audit.json").write_text(
        json.dumps(
            json_ready(
                {
                    **harmonized_audit,
                    "result": harmonized_result,
                }
            ),
            indent=2,
            ensure_ascii=False,
        )
        + "\n",
        encoding="utf-8",
    )
    negative_time_check.to_csv(
        OUT / "negative_time_equivalence.tsv", sep="\t", index=False
    )
    pd.DataFrame(
        {
            "bootstrap_index": np.arange(N_BOOT),
            "mean_affiliation_index": primary_boot,
        }
    ).to_csv(
        OUT / "primary_pair_bootstrap.tsv.gz",
        sep="\t",
        index=False,
        compression="gzip",
    )
    duplicate_audit.to_csv(
        OUT / "duplicate_occasion_audit.tsv", sep="\t", index=False
    )
    plot_figure(primary_pairs, primary_result, sensitivities)

    results = {
        "primary": primary_result,
        "stop_checks": stop_checks,
        "common_probe_harmonized_processing_sensitivity": harmonized_result,
        "donor_occasion_clock_coordinate_correlation_descriptive": {
            "n_occasions": int(len(donor_occasions)),
            "n_pairs": int(donor_occasions["pair_id"].nunique()),
            "pearson_r": float(donor_corr.statistic),
            "p_value_not_used_for_inference": float(donor_corr.pvalue),
        },
        "environment": {
            "python": sys.version,
            "platform": platform.platform(),
            "numpy": np.__version__,
            "pandas": pd.__version__,
            "scipy": scipy.__version__,
            "matplotlib": mpl.__version__,
            "seed": SEED,
            "n_bootstrap": N_BOOT,
        },
    }
    (OUT / "analysis_results.json").write_text(
        json.dumps(json_ready(results), indent=2, ensure_ascii=False) + "\n",
        encoding="utf-8",
    )

    report = f"""# Holland 2024 Horvath 2013 object-affiliation reanalysis

**QC status:** PASS  
**Inference unit:** donor-recipient pair  
**Role:** empirical object identification, not benefit or rejuvenation

## Primary result

After conservative duplicate-occasion aggregation, the primary population
contained **{primary_result['n_occasions']} recipient-blood occasions from
{primary_result['n_pairs']} donor-recipient pairs**. The pair-weighted mean
side-affiliation index was **{primary_result['mean_affiliation_index']:.3f}**
(pair-bootstrap 95% CI {primary_result['ci_low']:.3f} to
{primary_result['ci_high']:.3f}; median
{primary_result['median_affiliation_index']:.3f}).

Pair-mean \(I\) was positive in
**{primary_result['n_positive_pair_mean']} pairs**, negative in
**{primary_result['n_negative_pair_mean']}**, with
**{primary_result['n_ties']} ties**. Among non-tied pair means, the exact
two-sided binomial \(P\) value was
{primary_result['binomial_p_two_sided']:.4g}.

Mean absolute error was
{primary_result['mean_abs_error_to_lineage_coordinate']:.2f} age-equivalent
units to the donor-lineage coordinate and
{primary_result['mean_abs_error_to_host']:.2f} to recipient host age.

## What the estimand means

\[
I=(|S-H|-|S-C|)/|H-C|,
\]

where \(S\) is the Horvath-2013 coefficient implementation, \(H\) is
recipient-host chronological age, and
\(C=A_D+t\) is the donor-lineage chronological coordinate: donor age at
transplantation plus elapsed time. In recipient samples, the public
`AgeCells` field equals this coordinate to floating-point precision. \(C\)
is not the literal age of the short-lived blood cells present at sampling.

By the reverse triangle inequality, \(-1\le I\le1\). Positive values are on
the donor-lineage-coordinate side, negative values are on the host side and
zero is equal distance. The index saturates: \(I=1\) can occur when \(S=C\)
or when \(S\) lies beyond \(C\) away from \(H\). It is therefore a bounded
side-affiliation index, not a normalized measure of absolute accuracy.

Under this design, the result estimates which chronological coordinate the
blood score lies closer to. It does not identify donor age as the recipient's
biological age and does not establish benefit, harm or rejuvenation.

## Sensitivities

{markdown_table(sensitivities, digits=4)}

The post-specification processing sensitivity used only the
**{harmonized_audit['n_observed_in_all_six_batches']} probes observed in all
six batches** and assigned the same fixed external mean to the other
{harmonized_audit['n_set_to_fixed_external_mean_in_all_batches']} probes in
every batch. Its pair-weighted mean \(I\) was
{harmonized_result['mean_affiliation_index']:.3f} (95% CI
{harmonized_result['ci_low']:.3f} to {harmonized_result['ci_high']:.3f}).
Truncating the near-zero negative follow-up values to zero or excluding them
produced the identical primary set under the prespecified
\(t\ge0.1\)-year filter; the machine-readable check is saved with the outputs.

## Source and extraction audit

- Six remote files, {source_bytes_total / 1e9:.2f} GB
  ({source_bytes_total / (1024 ** 3):.2f} GiB) in total, were streamed; their
  observed SHA-256 values matched the Mendeley values.
- All 334 sample columns mapped one-to-one to public metadata.
- The six processed batches lacked 52, 28, 34, 18, 13 and 10 of the 353 probes,
  respectively. When a probe was absent from an entire batch, it was assigned
  the fixed `overallMeanByCpGacross50data` value from the local PC-Clocks
  resource. These values have not been established as the original Horvath
  training means.
- There were no sample-level non-finite or out-of-range target beta values.
- Formula validation passed against the bundled five-sample reference before
  scoring the public matrices.
- Recipient arrays were conservatively reduced from 303 arrays to
  {stop_checks['n_recipient_occasions']} metadata-identical occasions.
- Duplicate occasion score range reached
  {stop_checks['max_duplicate_occasion_score_range']:.3f} units, so array-level
  duplicates were not treated as independent and preprocessing uncertainty is
  not included in the pair bootstrap.
- After metadata-identical aggregation, the descriptive donor-occasion score
  versus public `AgeCells` correlation was \(r={donor_corr.statistic:.3f}\)
  across {len(donor_occasions)} occasions from
  {donor_occasions['pair_id'].nunique()} pairs. Its naive \(P\) value is
  not used as independent evidence.

## Interpretation boundary

Allowed: this Horvath-2013 coefficient implementation with fixed
external-mean imputation in recipient blood aligned more closely with the
donor-lineage chronological coordinate than with recipient host age under the
frozen estimand.

Not allowed: whole-organism age was replaced; transplantation rejuvenated or
aged recipients; clock movement proves clinical benefit; or this empirical
analysis proves the paper's general theorem.

## Reproducibility

- Spec: `{SPEC.relative_to(ROOT)}`
- Coefficient export:
  `analysis/export_horvath2013_coefficients_v0_1.R`
- Streaming extraction:
  `analysis/extract_holland2024_horvath2013_v0_1.py`
- Analysis:
  `analysis/analyze_holland2024_horvath_object_affiliation_v0_1.py`
- Outputs: `{OUT.relative_to(ROOT)}`
- Figure: `{FIGURE_PNG.relative_to(ROOT)}`
"""
    REPORT.write_text(report, encoding="utf-8")

    output_paths = [
        OUT / "input_manifest.tsv",
        OUT / "analysis_arrays.tsv.gz",
        OUT / "analysis_occasions.tsv.gz",
        OUT / "primary_occasions.tsv",
        OUT / "primary_pair_summary.tsv",
        OUT / "sensitivity_summary.tsv",
        OUT / "common_probe_harmonized_scores.tsv",
        OUT / "common_probe_harmonized_primary_pair_summary.tsv",
        OUT / "common_probe_harmonization_audit.json",
        OUT / "negative_time_equivalence.tsv",
        OUT / "primary_pair_bootstrap.tsv.gz",
        OUT / "duplicate_occasion_audit.tsv",
        OUT / "analysis_results.json",
        FIGURE_PNG,
        FIGURE_PDF,
        REPORT,
        Path(__file__).resolve(),
    ]
    audit = {
        "qc_status": "PASS",
        "primary_estimate_status": "observed_and_independently_recomputed",
        "spec_sha256": sha256(SPEC),
        "stop_checks": stop_checks,
        "output_hashes": {
            str(path.relative_to(ROOT)): sha256(path) for path in output_paths
        },
    }
    (OUT / "audit.json").write_text(
        json.dumps(json_ready(audit), indent=2, ensure_ascii=False) + "\n",
        encoding="utf-8",
    )
    print(json.dumps(json_ready(results), indent=2, ensure_ascii=False))


if __name__ == "__main__":
    main()
```
