## Supplementary_Software_1 for "Inferential boundaries of age prediction: why prediction does not establish biological age measurement": analyze_hsct_clock_object_replacement_v0_1.py

```
"""Focused HSCT clock object-replacement reanalysis.

Implements HSCT_CLOCK_OBJECT_REPLACEMENT_SPEC_20260730.md for the individual-
level Søraas et al. 2019 table embedded in Europe PMC XML.

The analysis asks whether an age-named score computed from post-HSCT recipient
blood is closer to the donor age (the source of the measured compartment) or
the recipient age (the host organism). It does not interpret score movement as
benefit, harm, rejuvenation, or organismal ageing.
"""

from __future__ import annotations

import hashlib
import json
import math
import platform
import re
import sys
from pathlib import Path
from typing import Callable
from xml.etree import ElementTree as ET

import matplotlib as mpl
import matplotlib.pyplot as plt
from matplotlib.lines import Line2D
import numpy as np
import pandas as pd
import scipy
from scipy import stats


ROOT = Path(__file__).resolve().parents[1]
SPEC = ROOT / "analysis" / "HSCT_CLOCK_OBJECT_REPLACEMENT_SPEC_20260730.md"
XML_2019 = ROOT / "data" / "raw" / "HSCT_Soraas2019" / "PMC6413751_fullText.xml"
PUBLISHED_2024 = (
    ROOT
    / "data"
    / "processed"
    / "hsct_holland2024_published_model1_recipient_status_v0_1.tsv"
)
ENVIRONMENT_POLICY = ROOT / "environment-policy.json"
ENVIRONMENT_MANIFEST = ROOT / "environment.yml"
ENVIRONMENT_LOCK = ROOT / "environment.win-64.lock.txt"
OUT = ROOT / "outputs" / "hsct_clock_object_replacement_v0_1"
PROCESSED = (
    ROOT / "data" / "processed" / "hsct_soraas2019_table1_extracted_v0_1.tsv"
)
REPORT = ROOT / "reports" / "HSCT_CLOCK_OBJECT_REPLACEMENT_V0_1_20260730.md"
FIGURE_PNG = ROOT / "figures" / "HSCT_CLOCK_OBJECT_REPLACEMENT_V0_1.png"
FIGURE_PDF = ROOT / "figures" / "HSCT_CLOCK_OBJECT_REPLACEMENT_V0_1.pdf"

SEED = 20260730
N_BOOT = 10_000

LOCKED_INPUTS = {
    SPEC: "6ac22a502f374276fd8b7e7017fc87d7eb11e206dab478398892b51dfb347498",
    XML_2019: "51ca18f969777d56caa275765e42a24452f072249831c7a39c97c962b99b0a74",
    PUBLISHED_2024: "f544ded0aa4038c6bb6a3d4c852ed6791a0b9d5d8c5a43511ff6aa4dcd995ede",
    ENVIRONMENT_POLICY: "b7ffa630ce7617f61eb3798f49604eba4dfa399a1d11fc7719e2971881322dd3",
    ENVIRONMENT_MANIFEST: "262e928e66e5d89a46429b409c513169d270fd74023b1ed30a55ebe2cb7c7773",
    ENVIRONMENT_LOCK: "85eae09d6fb170ff4616262d5263a7decfdac3e32e6242223d137054a483b5d6",
}

COLUMNS = (
    "sample_id_raw",
    "recipient_age",
    "donor_age",
    "dnam_age_recipient_blood",
    "repeated_sample",
    "months_after_hsct",
    "alive_last_followup",
    "relapsed",
    "recipient_minus_donor_age_reported",
    "dnam_minus_recipient_age_reported",
    "dnam_minus_donor_age_reported",
    "recipient_age_at_hsct",
    "donor_age_at_hsct",
    "donor_chimerism_raw",
    "dnam_age_donor_blood",
    "stem_cell_source",
)

NUMERIC_COLUMNS = (
    "recipient_age",
    "donor_age",
    "dnam_age_recipient_blood",
    "repeated_sample",
    "months_after_hsct",
    "recipient_minus_donor_age_reported",
    "dnam_minus_recipient_age_reported",
    "dnam_minus_donor_age_reported",
    "recipient_age_at_hsct",
    "donor_age_at_hsct",
    "dnam_age_donor_blood",
)

RECIPIENT_COLOR = "#C45A35"
DONOR_COLOR = "#087E8B"
SCORE_COLOR = "#3B334F"
NEUTRAL = "#6B7280"
LIGHT = "#E5E7EB"


def sha256(path: Path) -> str:
    digest = hashlib.sha256()
    with path.open("rb") as handle:
        for block in iter(lambda: handle.read(1024 * 1024), b""):
            digest.update(block)
    return digest.hexdigest()


def verify_inputs() -> pd.DataFrame:
    rows = []
    for path, expected in LOCKED_INPUTS.items():
        if not path.is_file():
            raise FileNotFoundError(path)
        observed = sha256(path)
        if observed.lower() != expected.lower():
            raise RuntimeError(
                f"Locked input hash mismatch for {path}: "
                f"expected {expected}, observed {observed}"
            )
        rows.append(
            {
                "path": str(path.relative_to(ROOT)),
                "sha256": observed,
                "bytes": path.stat().st_size,
            }
        )
    return pd.DataFrame(rows)


def clean_text(value: str | None) -> str:
    if value is None:
        return ""
    return (
        value.replace("\u2212", "-")
        .replace("\u2010", "-")
        .replace("\u2011", "-")
        .replace("\u00a0", " ")
        .strip()
    )


def node_text(node: ET.Element) -> str:
    return clean_text("".join(node.itertext()))


def parse_number(value: object) -> float:
    text = clean_text(str(value))
    if not text or text.upper() in {"NA", "N/A", "NAN"}:
        return float("nan")
    text = text.replace(">", "").replace("<", "")
    match = re.search(r"[-+]?\d+(?:\.\d+)?", text)
    return float(match.group(0)) if match else float("nan")


def parse_table1(path: Path) -> pd.DataFrame:
    root = ET.parse(path).getroot()
    table_wrap = None
    for node in root.iter():
        if node.attrib.get("id") == "acel12897-tbl-0001":
            table_wrap = node
            break
    if table_wrap is None:
        raise RuntimeError("Table 1 (acel12897-tbl-0001) not found")

    tbody = next((node for node in table_wrap.iter() if node.tag.endswith("tbody")), None)
    if tbody is None:
        raise RuntimeError("Table 1 body not found")

    parsed: list[dict[str, object]] = []
    for row_index, row in enumerate(
        [node for node in list(tbody) if node.tag.endswith("tr")], start=1
    ):
        cells = [node for node in list(row) if node.tag.endswith("td")]
        values = [node_text(cell) for cell in cells]
        if len(values) != len(COLUMNS):
            raise RuntimeError(
                f"Unexpected Table 1 width at row {row_index}: "
                f"{len(values)} instead of {len(COLUMNS)}"
            )
        item = dict(zip(COLUMNS, values))
        item["source_row"] = row_index
        item["published_exclusion_b"] = any(
            node.attrib.get("rid") == "acel12897-note-0005" for node in row.iter()
        )
        recipient_id = re.sub(r"\s*b\s*$", "", clean_text(values[0]), flags=re.I)
        if not recipient_id or re.fullmatch(r"\d+(?:/\d+)?", recipient_id) is None:
            raise RuntimeError(f"Could not parse sample ID from {values[0]!r}")
        item["recipient_id"] = recipient_id
        parsed.append(item)

    frame = pd.DataFrame(parsed)
    for column in NUMERIC_COLUMNS:
        frame[column] = frame[column].map(parse_number)
    frame["donor_chimerism_pct"] = frame["donor_chimerism_raw"].map(parse_number)
    frame["donor_chimerism_is_lower_bound"] = frame["donor_chimerism_raw"].str.contains(
        ">", regex=False, na=False
    )
    frame["published_analysis_row"] = ~frame["published_exclusion_b"]
    frame["primary_row"] = frame["published_analysis_row"] & frame["relapsed"].eq("No")

    frame["abs_error_to_donor"] = (
        frame["dnam_age_recipient_blood"] - frame["donor_age"]
    ).abs()
    frame["abs_error_to_recipient"] = (
        frame["dnam_age_recipient_blood"] - frame["recipient_age"]
    ).abs()
    frame["donor_minus_recipient_proximity"] = (
        frame["abs_error_to_donor"] - frame["abs_error_to_recipient"]
    )
    frame["recipient_minus_donor_age_calculated"] = (
        frame["recipient_age"] - frame["donor_age"]
    )
    frame["donor_blood_validation_contrast"] = (
        (frame["dnam_age_recipient_blood"] - frame["dnam_age_donor_blood"]).abs()
        - frame["abs_error_to_recipient"]
    )
    return frame


def validate_extraction(frame: pd.DataFrame) -> dict[str, object]:
    valid = frame.loc[frame["published_analysis_row"]].copy()
    primary = frame.loc[frame["primary_row"]].copy()
    checks = {
        "raw_rows": int(len(frame)),
        "published_analysis_rows": int(len(valid)),
        "published_analysis_recipients": int(valid["recipient_id"].nunique()),
        "primary_rows": int(len(primary)),
        "primary_recipients": int(primary["recipient_id"].nunique()),
        "excluded_rows": int(frame["published_exclusion_b"].sum()),
        "contemporaneous_donor_clock_rows": int(
            primary["dnam_age_donor_blood"].notna().sum()
        ),
        "reported_age_difference_max_abs_discrepancy": float(
            np.nanmax(
                np.abs(
                    frame["recipient_minus_donor_age_calculated"]
                    - frame["recipient_minus_donor_age_reported"]
                )
            )
        ),
    }
    expected = {
        "raw_rows": 41,
        "published_analysis_rows": 36,
        "published_analysis_recipients": 30,
        "primary_rows": 24,
        "primary_recipients": 19,
        "excluded_rows": 5,
        "contemporaneous_donor_clock_rows": 5,
    }
    for key, value in expected.items():
        if checks[key] != value:
            raise RuntimeError(f"Extraction check failed for {key}: {checks[key]} != {value}")
    if checks["reported_age_difference_max_abs_discrepancy"] > 0:
        raise RuntimeError("Calculated and published recipient-donor age differences disagree")

    all_donor_r = stats.pearsonr(valid["dnam_age_recipient_blood"], valid["donor_age"]).statistic
    all_recipient_r = stats.pearsonr(
        valid["dnam_age_recipient_blood"], valid["recipient_age"]
    ).statistic
    primary_donor_r = stats.pearsonr(
        primary["dnam_age_recipient_blood"], primary["donor_age"]
    ).statistic
    checks.update(
        {
            "published_reproduction_all_donor_r": float(all_donor_r),
            "published_reproduction_all_recipient_r": float(all_recipient_r),
            "published_reproduction_primary_donor_r": float(primary_donor_r),
            "published_reproduction_mean_abs_error_donor": float(
                valid["abs_error_to_donor"].mean()
            ),
            "published_reproduction_mean_abs_error_recipient": float(
                valid["abs_error_to_recipient"].mean()
            ),
        }
    )
    if abs(all_donor_r - 0.79) > 0.02:
        raise RuntimeError("Failed to reproduce published all-sample donor correlation")
    if abs(all_recipient_r - (-0.14)) > 0.02:
        raise RuntimeError("Failed to reproduce published all-sample recipient correlation")
    if abs(primary_donor_r - 0.89) > 0.02:
        raise RuntimeError("Failed to reproduce published non-relapse donor correlation")
    if abs(checks["published_reproduction_mean_abs_error_donor"] - 7.2) > 0.2:
        raise RuntimeError("Failed to reproduce published donor-age absolute difference")
    return checks


def recipient_level(frame: pd.DataFrame) -> pd.DataFrame:
    aggregations: dict[str, tuple[str, str | Callable]] = {
        "n_samples": ("source_row", "size"),
        "recipient_age": ("recipient_age", "mean"),
        "donor_age": ("donor_age", "mean"),
        "dnam_age_recipient_blood": ("dnam_age_recipient_blood", "mean"),
        "months_after_hsct": ("months_after_hsct", "mean"),
        "donor_chimerism_pct_min": ("donor_chimerism_pct", "min"),
        "abs_error_to_donor": ("abs_error_to_donor", "mean"),
        "abs_error_to_recipient": ("abs_error_to_recipient", "mean"),
        "d": ("donor_minus_recipient_proximity", "mean"),
        "stem_cell_source": (
            "stem_cell_source",
            lambda values: ",".join(sorted(set(map(str, values)))),
        ),
    }
    result = frame.groupby("recipient_id", sort=False).agg(**aggregations).reset_index()
    result["closer_to"] = np.select(
        [result["d"] < 0, result["d"] > 0],
        ["donor", "recipient"],
        default="tie",
    )
    result["donor_minus_recipient_age"] = (
        result["donor_age"] - result["recipient_age"]
    )
    return result


def bootstrap_subject_rows(
    subject_frame: pd.DataFrame,
    estimator: Callable[[pd.DataFrame], float],
    n_boot: int = N_BOOT,
) -> np.ndarray:
    rng = np.random.default_rng(SEED)
    n = len(subject_frame)
    draws = np.empty(n_boot, dtype=float)
    for index in range(n_boot):
        sample = subject_frame.iloc[rng.integers(0, n, size=n)]
        draws[index] = estimator(sample)
    return draws[np.isfinite(draws)]


def corr(frame: pd.DataFrame, x: str, y: str, method: str) -> float:
    if method == "pearson":
        return float(stats.pearsonr(frame[x], frame[y]).statistic)
    if method == "spearman":
        return float(stats.spearmanr(frame[x], frame[y]).statistic)
    raise ValueError(method)


def interval(values: np.ndarray) -> tuple[float, float]:
    low, high = np.quantile(values, [0.025, 0.975], method="linear")
    return float(low), float(high)


def bootstrap_correlations(subjects: pd.DataFrame) -> dict[str, np.ndarray]:
    """Bootstrap all correlation contrasts from the same recipient draws."""
    values = subjects[
        ["dnam_age_recipient_blood", "donor_age", "recipient_age"]
    ].to_numpy(dtype=float)
    rng = np.random.default_rng(SEED)
    n = len(values)
    draws = {
        "pearson_donor": np.full(N_BOOT, np.nan),
        "pearson_recipient": np.full(N_BOOT, np.nan),
        "pearson_difference": np.full(N_BOOT, np.nan),
        "spearman_donor": np.full(N_BOOT, np.nan),
        "spearman_recipient": np.full(N_BOOT, np.nan),
        "spearman_difference": np.full(N_BOOT, np.nan),
    }
    for index in range(N_BOOT):
        sample = values[rng.integers(0, n, size=n)]
        if np.all(np.std(sample, axis=0, ddof=0) > 0):
            matrix = np.corrcoef(sample, rowvar=False)
            donor_r = matrix[0, 1]
            recipient_r = matrix[0, 2]
            draws["pearson_donor"][index] = donor_r
            draws["pearson_recipient"][index] = recipient_r
            draws["pearson_difference"][index] = donor_r - recipient_r

            ranked = np.column_stack(
                [stats.rankdata(sample[:, column]) for column in range(3)]
            )
            rank_matrix = np.corrcoef(ranked, rowvar=False)
            donor_rank_r = rank_matrix[0, 1]
            recipient_rank_r = rank_matrix[0, 2]
            draws["spearman_donor"][index] = donor_rank_r
            draws["spearman_recipient"][index] = recipient_rank_r
            draws["spearman_difference"][index] = donor_rank_r - recipient_rank_r
    return {key: value[np.isfinite(value)] for key, value in draws.items()}


def sample_level_cluster_correlations(
    rows: pd.DataFrame, label: str
) -> pd.DataFrame:
    """Sample-level correlations with recipient-cluster bootstrap intervals."""
    value_columns = ["dnam_age_recipient_blood", "donor_age", "recipient_age"]
    clusters = [
        group[value_columns].to_numpy(dtype=float)
        for _, group in rows.groupby("recipient_id", sort=False)
    ]
    rng = np.random.default_rng(SEED)
    draws = {
        "pearson_donor": np.full(N_BOOT, np.nan),
        "pearson_recipient": np.full(N_BOOT, np.nan),
        "pearson_difference": np.full(N_BOOT, np.nan),
        "spearman_donor": np.full(N_BOOT, np.nan),
        "spearman_recipient": np.full(N_BOOT, np.nan),
        "spearman_difference": np.full(N_BOOT, np.nan),
    }
    for index in range(N_BOOT):
        selected = rng.integers(0, len(clusters), size=len(clusters))
        sample = np.concatenate([clusters[item] for item in selected], axis=0)
        if np.all(np.std(sample, axis=0, ddof=0) > 0):
            matrix = np.corrcoef(sample, rowvar=False)
            draws["pearson_donor"][index] = matrix[0, 1]
            draws["pearson_recipient"][index] = matrix[0, 2]
            draws["pearson_difference"][index] = matrix[0, 1] - matrix[0, 2]
            ranked = np.column_stack(
                [stats.rankdata(sample[:, column]) for column in range(3)]
            )
            rank_matrix = np.corrcoef(ranked, rowvar=False)
            draws["spearman_donor"][index] = rank_matrix[0, 1]
            draws["spearman_recipient"][index] = rank_matrix[0, 2]
            draws["spearman_difference"][index] = (
                rank_matrix[0, 1] - rank_matrix[0, 2]
            )

    result_rows: list[dict[str, object]] = []
    for method in ("pearson", "spearman"):
        point_donor = corr(
            rows, "dnam_age_recipient_blood", "donor_age", method=method
        )
        point_recipient = corr(
            rows, "dnam_age_recipient_blood", "recipient_age", method=method
        )
        for contrast, point, key in (
            ("corr(score, donor_age)", point_donor, f"{method}_donor"),
            ("corr(score, recipient_age)", point_recipient, f"{method}_recipient"),
            (
                "corr(score, donor_age) - corr(score, recipient_age)",
                point_donor - point_recipient,
                f"{method}_difference",
            ),
        ):
            finite = draws[key][np.isfinite(draws[key])]
            ci_low, ci_high = interval(finite)
            result_rows.append(
                {
                    "population": label,
                    "unit": "sample_level_with_recipient_cluster_bootstrap",
                    "method": method,
                    "contrast": contrast,
                    "estimate": point,
                    "ci_low": ci_low,
                    "ci_high": ci_high,
                    "n_samples": int(len(rows)),
                    "n_recipients": int(rows["recipient_id"].nunique()),
                }
            )
    return pd.DataFrame(result_rows)


def analyze_population(
    rows: pd.DataFrame, label: str
) -> tuple[pd.DataFrame, pd.DataFrame, dict[str, object], np.ndarray]:
    subjects = recipient_level(rows)
    non_ties = subjects.loc[subjects["d"].ne(0)]
    donor_closer = int((non_ties["d"] < 0).sum())
    exact = stats.binomtest(donor_closer, len(non_ties), p=0.5, alternative="two-sided")

    mean_boot = bootstrap_subject_rows(subjects, lambda data: float(data["d"].mean()))
    mean_ci = interval(mean_boot)
    wilcoxon = stats.wilcoxon(
        subjects["d"], alternative="two-sided", zero_method="wilcox", method="auto"
    )

    summary = {
        "population": label,
        "n_samples": int(len(rows)),
        "n_recipients": int(len(subjects)),
        "n_donor_closer": donor_closer,
        "n_recipient_closer": int((non_ties["d"] > 0).sum()),
        "n_ties": int((subjects["d"] == 0).sum()),
        "proportion_donor_closer": float(donor_closer / len(non_ties)),
        "binomial_p_two_sided": float(exact.pvalue),
        "mean_d": float(subjects["d"].mean()),
        "mean_d_ci_low": mean_ci[0],
        "mean_d_ci_high": mean_ci[1],
        "median_d": float(subjects["d"].median()),
        "wilcoxon_statistic": float(wilcoxon.statistic),
        "wilcoxon_p_two_sided": float(wilcoxon.pvalue),
        "mean_abs_error_to_donor": float(subjects["abs_error_to_donor"].mean()),
        "mean_abs_error_to_recipient": float(
            subjects["abs_error_to_recipient"].mean()
        ),
    }

    correlation_boot = bootstrap_correlations(subjects)
    correlation_rows: list[dict[str, object]] = []
    for method in ("pearson", "spearman"):
        donor_r = corr(
            subjects, "dnam_age_recipient_blood", "donor_age", method=method
        )
        recipient_r = corr(
            subjects, "dnam_age_recipient_blood", "recipient_age", method=method
        )
        donor_boot = correlation_boot[f"{method}_donor"]
        recipient_boot = correlation_boot[f"{method}_recipient"]
        difference_boot = correlation_boot[f"{method}_difference"]
        donor_ci = interval(donor_boot)
        recipient_ci = interval(recipient_boot)
        difference_ci = interval(difference_boot)
        correlation_rows.extend(
            [
                {
                    "population": label,
                    "method": method,
                    "contrast": "corr(score, donor_age)",
                    "estimate": donor_r,
                    "ci_low": donor_ci[0],
                    "ci_high": donor_ci[1],
                },
                {
                    "population": label,
                    "method": method,
                    "contrast": "corr(score, recipient_age)",
                    "estimate": recipient_r,
                    "ci_low": recipient_ci[0],
                    "ci_high": recipient_ci[1],
                },
                {
                    "population": label,
                    "method": method,
                    "contrast": "corr(score, donor_age) - corr(score, recipient_age)",
                    "estimate": donor_r - recipient_r,
                    "ci_low": difference_ci[0],
                    "ci_high": difference_ci[1],
                },
            ]
        )
    return subjects, pd.DataFrame(correlation_rows), summary, mean_boot


def leave_one_out(subjects: pd.DataFrame) -> pd.DataFrame:
    rows = []
    for recipient_id in subjects["recipient_id"]:
        retained = subjects.loc[subjects["recipient_id"].ne(recipient_id)]
        rows.append(
            {
                "left_out_recipient_id": recipient_id,
                "n_retained": int(len(retained)),
                "mean_d": float(retained["d"].mean()),
                "proportion_donor_closer": float((retained["d"] < 0).mean()),
            }
        )
    return pd.DataFrame(rows)


def same_donor_validation(primary_rows: pd.DataFrame) -> pd.DataFrame:
    columns = [
        "recipient_id",
        "recipient_age",
        "donor_age",
        "dnam_age_recipient_blood",
        "dnam_age_donor_blood",
        "abs_error_to_recipient",
        "donor_blood_validation_contrast",
    ]
    return primary_rows.loc[
        primary_rows["dnam_age_donor_blood"].notna(), columns
    ].copy()


def direction_reversal_cases(primary_rows: pd.DataFrame) -> pd.DataFrame:
    return primary_rows.loc[
        (primary_rows["recipient_age"] - primary_rows["donor_age"]).abs() >= 20,
        [
            "recipient_id",
            "recipient_age",
            "donor_age",
            "dnam_age_recipient_blood",
            "months_after_hsct",
            "donor_chimerism_pct",
            "donor_minus_recipient_proximity",
        ],
    ].copy()


def plot_figure(
    subjects: pd.DataFrame,
    summary: dict[str, object],
    same_donor: pd.DataFrame,
    cases: pd.DataFrame,
    published_2024: pd.DataFrame,
) -> None:
    mpl.rcParams.update(
        {
            "font.family": "DejaVu Sans",
            "font.size": 9,
            "axes.titlesize": 10,
            "axes.labelsize": 9,
            "axes.spines.top": False,
            "axes.spines.right": False,
            "pdf.fonttype": 42,
            "ps.fonttype": 42,
        }
    )
    fig = plt.figure(figsize=(12.4, 8.7), constrained_layout=True)
    grid = fig.add_gridspec(2, 2, width_ratios=(1.08, 1), height_ratios=(1, 1))
    ax_a = fig.add_subplot(grid[0, 0])
    ax_b = fig.add_subplot(grid[0, 1])
    ax_c = fig.add_subplot(grid[1, 0])
    ax_d = fig.add_subplot(grid[1, 1])

    # Panel a: all prespecified direction-reversal configurations.
    ordered_cases = cases.sort_values(
        ["recipient_age", "donor_age"], ascending=[True, True]
    ).reset_index(drop=True)
    y_positions = np.arange(len(ordered_cases))
    for y, row in ordered_cases.iterrows():
        lo = min(row["recipient_age"], row["donor_age"])
        hi = max(row["recipient_age"], row["donor_age"])
        ax_a.plot([lo, hi], [y, y], color=LIGHT, lw=2.2, zorder=1)
    ax_a.scatter(
        ordered_cases["recipient_age"],
        y_positions,
        s=28,
        facecolors="white",
        edgecolors=RECIPIENT_COLOR,
        linewidths=1.2,
        label="Recipient age",
        zorder=3,
    )
    ax_a.scatter(
        ordered_cases["donor_age"],
        y_positions,
        s=34,
        marker="D",
        color=DONOR_COLOR,
        label="Donor-lineage coordinate",
        zorder=3,
    )
    ax_a.scatter(
        ordered_cases["dnam_age_recipient_blood"],
        y_positions,
        s=31,
        color=SCORE_COLOR,
        label="Clock in recipient blood",
        zorder=4,
    )
    ax_a.axvline(0, color="none")
    ax_a.set_yticks(y_positions)
    ax_a.set_yticklabels(ordered_cases["recipient_id"].map(lambda value: f"ID {value}"))
    ax_a.invert_yaxis()
    ax_a.set_ylim(len(ordered_cases) - 0.5, -1.55)
    key_y = -0.92
    ax_a.scatter(
        [7],
        [key_y],
        s=28,
        facecolors="white",
        edgecolors=RECIPIENT_COLOR,
        linewidths=1.2,
        zorder=4,
    )
    ax_a.text(9, key_y, "Recipient age", va="center", fontsize=7.2)
    ax_a.scatter([25], [key_y], s=34, marker="D", color=DONOR_COLOR, zorder=4)
    ax_a.text(27, key_y, "Donor-lineage C", va="center", fontsize=7.2)
    ax_a.scatter([46], [key_y], s=31, color=SCORE_COLOR, zorder=4)
    ax_a.text(48, key_y, "Clock in recipient blood", va="center", fontsize=7.2)
    ax_a.set_xlabel("Age or age-equivalent clock value")
    ax_a.set_title(
        "a  Large donor-recipient age contrasts\n"
        "Clock values aligned with the donor-lineage coordinate"
    )

    # Panel b: all independent recipients in the primary population.
    ordered = subjects.sort_values("d").reset_index(drop=True)
    y = np.arange(len(ordered))
    ax_b.axvline(0, color="#111827", lw=0.9)
    ax_b.hlines(y, 0, ordered["d"], color=LIGHT, lw=2)
    point_colors = np.select(
        [ordered["d"] < 0, ordered["d"] > 0],
        [DONOR_COLOR, RECIPIENT_COLOR],
        default=NEUTRAL,
    )
    ax_b.scatter(
        ordered["d"],
        y,
        c=point_colors,
        s=33,
        edgecolor="white",
        linewidth=0.5,
        zorder=3,
    )
    mean_y = len(ordered)
    ax_b.plot(
        [summary["mean_d_ci_low"], summary["mean_d_ci_high"]],
        [mean_y, mean_y],
        color=SCORE_COLOR,
        lw=2.2,
        solid_capstyle="round",
        zorder=3,
    )
    ax_b.scatter(
        [summary["mean_d"]],
        [mean_y],
        marker="D",
        s=44,
        color=SCORE_COLOR,
        edgecolor="white",
        linewidth=0.5,
        zorder=4,
    )
    ax_b.set_yticks(np.concatenate([[mean_y], y]))
    ax_b.set_yticklabels(
        ["Mean (95% CI)"]
        + ordered["recipient_id"].map(lambda value: f"ID {value}").tolist()
    )
    ax_b.set_xlabel(
        r"$d=|S-A_D|-|S-A_R|$"
        "\nNegative values are on the donor-lineage-coordinate side"
    )
    ax_b.set_title(
        "b  Donor-lineage-versus-host proximity\n"
        f"{summary['n_donor_closer']} lineage-side, "
        f"{summary['n_ties']} tie, 0 host-side; "
        f"exact $P$={summary['binomial_p_two_sided']:.3g}"
    )
    ax_b.grid(axis="x", color="#F3F4F6", lw=0.8)

    # Panel c: contemporaneous donor-blood clock validation.
    for index, row in same_donor.reset_index(drop=True).iterrows():
        ax_c.plot(
            [0, 1, 2],
            [
                row["recipient_age"],
                row["dnam_age_recipient_blood"],
                row["dnam_age_donor_blood"],
            ],
            color=LIGHT,
            lw=1.5,
            zorder=1,
        )
        ax_c.text(
            2.06,
            row["dnam_age_donor_blood"],
            f"ID {row['recipient_id']}",
            va="center",
            fontsize=7.5,
            color=NEUTRAL,
        )
    ax_c.scatter(
        np.zeros(len(same_donor)),
        same_donor["recipient_age"],
        facecolors="white",
        edgecolors=RECIPIENT_COLOR,
        s=38,
        linewidth=1.2,
        zorder=3,
    )
    ax_c.scatter(
        np.ones(len(same_donor)),
        same_donor["dnam_age_recipient_blood"],
        color=SCORE_COLOR,
        s=38,
        zorder=3,
    )
    ax_c.scatter(
        np.full(len(same_donor), 2),
        same_donor["dnam_age_donor_blood"],
        marker="D",
        color=DONOR_COLOR,
        s=40,
        zorder=3,
    )
    ax_c.set_xticks([0, 1, 2])
    ax_c.set_xticklabels(
        ["Recipient\nchronological age", "Clock in\nrecipient blood", "Clock in\ndonor blood"]
    )
    ax_c.set_ylabel("Age or age-equivalent clock value")
    ax_c.set_xlim(-0.35, 2.65)
    ax_c.set_title(
        "c  Same donor lineage measured in two bodies\n"
        "Five contemporaneous donor-recipient pairs"
    )
    ax_c.grid(axis="y", color="#F3F4F6", lw=0.8)

    # Panel d: explicitly published 2024 Model 1 estimates.
    forest = published_2024.loc[published_2024["include_family_contrast"]].copy()
    forest["family_order"] = forest["clock_family"].map(
        {"chronological-target": 0, "endpoint-linked": 1}
    )
    forest = forest.sort_values(
        ["family_order", "estimate", "clock_label"], ascending=[True, True, True]
    ).reset_index(drop=True)
    y_forest = np.arange(len(forest))
    forest_colors = forest["clock_family"].map(
        {
            "chronological-target": DONOR_COLOR,
            "endpoint-linked": "#A23B72",
        }
    )
    ax_d.axvline(0, color="#111827", lw=0.9)
    for y_value, (_, row) in zip(y_forest, forest.iterrows()):
        ax_d.plot(
            [row["ci_low"], row["ci_high"]],
            [y_value, y_value],
            color=forest_colors.iloc[y_value],
            lw=1.8,
            solid_capstyle="round",
        )
    ax_d.scatter(
        forest["estimate"],
        y_forest,
        c=forest_colors,
        s=35,
        edgecolor="white",
        linewidth=0.5,
        zorder=3,
    )
    boundary = int((forest["clock_family"] == "chronological-target").sum()) - 0.5
    ax_d.axhline(boundary, color=LIGHT, lw=1)
    ax_d.set_yticks(y_forest)
    ax_d.set_yticklabels(forest["clock_label"])
    ax_d.invert_yaxis()
    ax_d.set_xlabel("Adjusted recipient-status displacement (age-equivalent units)")
    ax_d.set_title(
        "d  Different targets assign opposite directions\n"
        "Published Holland 2024 Model 1 estimates (95% CI)"
    )
    ax_d.grid(axis="x", color="#F3F4F6", lw=0.8)
    ax_d.text(
        0.98,
        0.96,
        "Chronological-age target",
        transform=ax_d.transAxes,
        color=DONOR_COLOR,
        fontsize=7.4,
        fontweight="bold",
        ha="right",
        va="top",
    )
    ax_d.text(
        0.02,
        0.44,
        "Endpoint-linked target",
        transform=ax_d.transAxes,
        color="#A23B72",
        fontsize=7.4,
        fontweight="bold",
        ha="left",
        va="top",
    )
    ax_d.text(
        0.01,
        -0.19,
        "Published evidence; overlapping/expanded HSCT cohorts, not an independent "
        "new reanalysis.",
        transform=ax_d.transAxes,
        fontsize=7.2,
        color=NEUTRAL,
        va="top",
    )

    fig.suptitle(
        (
            "Post-HSCT Horvath 2013 scores aligned with the donor-lineage "
            "chronological coordinate rather than host age"
        ),
        fontsize=13.5,
        fontweight="bold",
        x=0.01,
        ha="left",
    )
    FIGURE_PNG.parent.mkdir(parents=True, exist_ok=True)
    fig.savefig(FIGURE_PNG, dpi=320, bbox_inches="tight", facecolor="white")
    fig.savefig(FIGURE_PDF, bbox_inches="tight", facecolor="white")
    plt.close(fig)


def markdown_table(frame: pd.DataFrame, digits: int = 4) -> str:
    if frame.empty:
        return "_No rows._"
    display = frame.copy()
    for column in display.select_dtypes(include=[np.number]).columns:
        display[column] = display[column].map(
            lambda value: "" if not np.isfinite(value) else f"{value:.{digits}g}"
        )
    header = "| " + " | ".join(display.columns) + " |"
    rule = "| " + " | ".join(["---"] * len(display.columns)) + " |"
    rows = [
        "| " + " | ".join(str(value).replace("\n", " ") for value in row) + " |"
        for row in display.itertuples(index=False, name=None)
    ]
    return "\n".join([header, rule, *rows])


def json_ready(value: object) -> object:
    if isinstance(value, dict):
        return {str(key): json_ready(item) for key, item in value.items()}
    if isinstance(value, (list, tuple)):
        return [json_ready(item) for item in value]
    if isinstance(value, (np.integer,)):
        return int(value)
    if isinstance(value, (np.floating,)):
        return None if not np.isfinite(value) else float(value)
    if isinstance(value, float):
        return None if not math.isfinite(value) else value
    if isinstance(value, Path):
        return str(value)
    return value


def main() -> None:
    OUT.mkdir(parents=True, exist_ok=True)
    PROCESSED.parent.mkdir(parents=True, exist_ok=True)
    REPORT.parent.mkdir(parents=True, exist_ok=True)

    manifest = verify_inputs()
    frame = parse_table1(XML_2019)
    extraction_checks = validate_extraction(frame)
    frame.to_csv(PROCESSED, sep="\t", index=False, na_rep="")

    populations = {
        "primary_no_relapse": frame.loc[frame["primary_row"]].copy(),
        "all_published_analysis_rows": frame.loc[frame["published_analysis_row"]].copy(),
        "primary_chimerism_ge_95": frame.loc[
            frame["primary_row"] & frame["donor_chimerism_pct"].ge(95)
        ].copy(),
        "primary_at_least_12_months": frame.loc[
            frame["primary_row"] & frame["months_after_hsct"].ge(12)
        ].copy(),
    }

    subject_tables: dict[str, pd.DataFrame] = {}
    correlation_tables: list[pd.DataFrame] = []
    population_summaries: list[dict[str, object]] = []
    bootstrap_tables: list[pd.DataFrame] = []
    for label, rows in populations.items():
        subjects, correlations, summary, mean_boot = analyze_population(rows, label)
        subject_tables[label] = subjects
        correlation_tables.append(correlations)
        population_summaries.append(summary)
        bootstrap_tables.append(
            pd.DataFrame(
                {
                    "population": label,
                    "bootstrap_index": np.arange(len(mean_boot)),
                    "mean_d": mean_boot,
                }
            )
        )

    primary_subjects = subject_tables["primary_no_relapse"]
    population_summary = pd.DataFrame(population_summaries)
    correlation_summary = pd.concat(correlation_tables, ignore_index=True)
    bootstrap_summary = pd.concat(bootstrap_tables, ignore_index=True)
    loo = leave_one_out(primary_subjects)
    donor_validation = same_donor_validation(populations["primary_no_relapse"])
    cases = primary_subjects.loc[
        primary_subjects["donor_minus_recipient_age"].abs().ge(20)
    ].copy()
    published_2024 = pd.read_csv(PUBLISHED_2024, sep="\t")
    sample_level_correlations = pd.concat(
        [
            sample_level_cluster_correlations(
                populations["primary_no_relapse"], "primary_no_relapse"
            ),
            sample_level_cluster_correlations(
                populations["all_published_analysis_rows"],
                "all_published_analysis_rows",
            ),
        ],
        ignore_index=True,
    )
    source_quality = (
        populations["primary_no_relapse"]
        .groupby("recipient_id", sort=False)["stem_cell_source"]
        .agg(
            n_distinct_sources=lambda values: int(values.nunique()),
            sources=lambda values: ",".join(sorted(set(map(str, values)))),
        )
        .reset_index()
    )
    source_quality = source_quality.loc[source_quality["n_distinct_sources"].gt(1)]
    deviations = pd.DataFrame(
        [
            {
                "item": "stem_cell_source_sensitivity",
                "status": "not_performed",
                "reason": (
                    "The embedded source table assigns both BM and PB to recipient "
                    "754/812 across repeated rows; source stratification is not "
                    "interpretable without author clarification."
                ),
            },
            {
                "item": "holland2024_coefficient_reanalysis",
                "status": "performed_in_separate_module",
                "reason": (
                    "A six-batch Horvath-2013 coefficient implementation with fixed "
                    "external-mean imputation is reported separately. This figure "
                    "retains the published Model 1 estimates for the cross-clock "
                    "target comparison."
                ),
            },
            {
                "item": "wilcoxon_test",
                "status": "additional_secondary_analysis",
                "reason": (
                    "Reported only as a secondary magnitude-sensitive check; the "
                    "prespecified exact sign/binomial result remains primary."
                ),
            },
        ]
    )

    primary_summary = population_summaries[0]
    plot_figure(
        primary_subjects,
        primary_summary,
        donor_validation,
        cases,
        published_2024,
    )

    manifest.to_csv(OUT / "input_manifest.tsv", sep="\t", index=False)
    primary_subjects.to_csv(
        OUT / "primary_recipient_contrasts.tsv", sep="\t", index=False
    )
    population_summary.to_csv(OUT / "population_summary.tsv", sep="\t", index=False)
    correlation_summary.to_csv(OUT / "correlation_summary.tsv", sep="\t", index=False)
    sample_level_correlations.to_csv(
        OUT / "sample_level_cluster_correlation_summary.tsv",
        sep="\t",
        index=False,
    )
    bootstrap_summary.to_csv(
        OUT / "bootstrap_mean_d.tsv.gz",
        sep="\t",
        index=False,
        compression="gzip",
    )
    loo.to_csv(OUT / "leave_one_recipient_out.tsv", sep="\t", index=False)
    donor_validation.to_csv(
        OUT / "contemporaneous_donor_validation.tsv", sep="\t", index=False
    )
    cases.to_csv(OUT / "direction_reversal_cases.tsv", sep="\t", index=False)
    published_2024.to_csv(
        OUT / "holland2024_published_model1_recipient_status.tsv",
        sep="\t",
        index=False,
    )
    source_quality.to_csv(OUT / "data_quality_flags.tsv", sep="\t", index=False)
    deviations.to_csv(OUT / "deviations.tsv", sep="\t", index=False)

    environment = {
        "python": sys.version,
        "platform": platform.platform(),
        "numpy": np.__version__,
        "pandas": pd.__version__,
        "scipy": scipy.__version__,
        "matplotlib": mpl.__version__,
        "seed": SEED,
        "n_bootstrap": N_BOOT,
    }
    results = {
        "extraction_checks": extraction_checks,
        "primary": primary_summary,
        "correlations_primary": correlation_summary.loc[
            correlation_summary["population"].eq("primary_no_relapse")
        ].to_dict(orient="records"),
        "sample_level_cluster_correlations": sample_level_correlations.to_dict(
            orient="records"
        ),
        "contemporaneous_donor_validation": {
            "n": int(len(donor_validation)),
            "n_closer_to_donor_blood_clock_than_recipient_age": int(
                (donor_validation["donor_blood_validation_contrast"] < 0).sum()
            ),
            "median_contrast": float(
                donor_validation["donor_blood_validation_contrast"].median()
            ),
        },
        "leave_one_out": {
            "mean_d_min": float(loo["mean_d"].min()),
            "mean_d_max": float(loo["mean_d"].max()),
            "all_negative": bool((loo["mean_d"] < 0).all()),
        },
        "holland2024_published_model1": {
            "status": "published_estimates_not_new_reanalysis",
            "chronological_target_mean": float(
                published_2024.loc[
                    published_2024["clock_family"].eq("chronological-target")
                    & published_2024["include_family_contrast"],
                    "estimate",
                ].mean()
            ),
            "endpoint_linked_mean": float(
                published_2024.loc[
                    published_2024["clock_family"].eq("endpoint-linked")
                    & published_2024["include_family_contrast"],
                    "estimate",
                ].mean()
            ),
        },
        "environment": environment,
        "package_status": (
            "PASS_CORE_2019_REANALYSIS_WITH_DECLARED_SOURCE_DEVIATION"
            "_AND_SEPARATE_2024_MODULE"
        ),
    }
    (OUT / "analysis_results.json").write_text(
        json.dumps(json_ready(results), indent=2, ensure_ascii=False) + "\n",
        encoding="utf-8",
    )

    audit = {
        "status": "PASS_CORE_2019_REANALYSIS_WITH_DECLARED_DEVIATIONS",
        "spec_sha256": sha256(SPEC),
        "raw_xml_sha256": sha256(XML_2019),
        "checks": extraction_checks,
        "output_hashes": {},
    }
    output_paths = [
        PROCESSED,
        OUT / "input_manifest.tsv",
        OUT / "primary_recipient_contrasts.tsv",
        OUT / "population_summary.tsv",
        OUT / "correlation_summary.tsv",
        OUT / "sample_level_cluster_correlation_summary.tsv",
        OUT / "bootstrap_mean_d.tsv.gz",
        OUT / "leave_one_recipient_out.tsv",
        OUT / "contemporaneous_donor_validation.tsv",
        OUT / "direction_reversal_cases.tsv",
        OUT / "holland2024_published_model1_recipient_status.tsv",
        OUT / "data_quality_flags.tsv",
        OUT / "deviations.tsv",
        OUT / "analysis_results.json",
        FIGURE_PNG,
        FIGURE_PDF,
    ]
    for path in output_paths:
        audit["output_hashes"][str(path.relative_to(ROOT))] = sha256(path)

    primary_corr = correlation_summary.loc[
        correlation_summary["population"].eq("primary_no_relapse")
        & correlation_summary["method"].eq("pearson")
    ].copy()
    report = f"""# HSCT Horvath 2013 object-affiliation reanalysis

**Analysis version:** v0.1  
**QC status:** PASS for the 2019 core reanalysis, with declared deviations  
**Role in paper:** empirical materiality of object identification, not proof of
the central theorem

## Result

In the clean non-relapse population, the reported **Horvath 2013 multi-tissue
DNAm age estimator** in recipient blood was closer to the donor-lineage
chronological coordinate \(A_D\) than to recipient-host age for
**{primary_summary['n_donor_closer']} of
{primary_summary['n_recipients']} independent recipients**; one recipient was
tied and none was closer to recipient-host age. All 18 non-tied contrasts
favoured the donor-lineage coordinate (two-sided exact binomial
\(P={primary_summary['binomial_p_two_sided']:.4g}\)).
The recipient-weighted mean contrast
\(d=|S-A_D|-|S-A_R|\) was
**{primary_summary['mean_d']:.2f} age-equivalent units**
(recipient-bootstrap 95% CI
{primary_summary['mean_d_ci_low']:.2f} to
{primary_summary['mean_d_ci_high']:.2f}); negative values favour the
donor-lineage coordinate.

Here \(A_D\) is the donor's chronological age at blood draw: a temporal
coordinate associated with the donor-derived haematopoietic lineage, not the
literal age of the individual blood cells present at sampling. The result says
that the age-named score aligned more closely with that lineage coordinate than
with the host organism. It does not say that donor age is the recipient's true
biological age.

## Extraction audit

- Parsed 41 rows directly from the embedded XML table.
- Reproduced the five published footnote-`b` exclusions.
- Reproduced 36 published analysis samples from 30 recipients.
- The prespecified non-relapse object-replacement population contained 24
  samples from 19 recipients.
- Reproduced the published sample-level correlations:
  donor age \(r={extraction_checks['published_reproduction_all_donor_r']:.3f}\),
  recipient age \(r={extraction_checks['published_reproduction_all_recipient_r']:.3f}\),
  and non-relapse donor age
  \(r={extraction_checks['published_reproduction_primary_donor_r']:.3f}\).

## Population contrasts

{markdown_table(population_summary, digits=4)}

## Recipient-level correlations

{markdown_table(primary_corr, digits=4)}

## Published sample-level correlations with recipient-cluster intervals

{markdown_table(sample_level_correlations, digits=4)}

## Contemporaneous donor-blood validation

All five long-term pairs are shown without inferential overstatement. The
median contrast
\(c=|S_{{recipient\ blood}}-S_{{donor\ blood}}|-|S_{{recipient\ blood}}-A_R|\)
was **{donor_validation['donor_blood_validation_contrast'].median():.2f}**.

{markdown_table(donor_validation, digits=4)}

## Published 2024 cross-clock consequence

The later overlapping/expanded HSCT study reported negative adjusted
recipient-status displacement for all four chronological-age-target clocks and
positive displacement for all four endpoint-linked age-valued clocks. The
descriptive family means were
{results['holland2024_published_model1']['chronological_target_mean']:.2f} and
{results['holland2024_published_model1']['endpoint_linked_mean']:.2f}
age-equivalent units, respectively. These are published Model 1 estimates
transcribed from Supporting Information Tables 1-2; they are not a new
individual-level reanalysis, and no family-mean confidence interval is claimed.

{markdown_table(published_2024.loc[published_2024['include_family_contrast'], ['clock_label', 'clock_family', 'estimate', 'ci_low', 'ci_high', 'p_value']], digits=4)}

## Robustness

- Every leave-one-recipient-out mean contrast remained negative:
  range {loo['mean_d'].min():.2f} to {loo['mean_d'].max():.2f}.
- The chimerism and follow-up sensitivities are reported in the population table.
- Relapsed samples are retained only as a sensitivity because relapse can
  reintroduce recipient malignant cells into the measured compartment.
- The embedded table assigns both BM and PB to repeated rows for recipient
  `754/812`; the planned stem-cell-source stratification was therefore not
  interpreted.

## Declared deviations

{markdown_table(deviations, digits=4)}

## Interpretation boundary

Allowed conclusion: in this selected post-HSCT cohort, the reported Horvath
2013 score in recipient blood was numerically closer to the donor-lineage
chronological coordinate than to recipient-host chronological age. A valid
blood-compartment recorder can therefore fail to identify an organism-level
biological age; object specification is empirically consequential.

Not established: clinical benefit, harm, rejuvenation, a universal claim about
all clocks, or the paper's general mathematical non-identification proposition.

## Reproducibility

- Specification: `{SPEC.relative_to(ROOT)}`
- Script: `analysis/analyze_hsct_clock_object_replacement_v0_1.py`
- Extracted table: `{PROCESSED.relative_to(ROOT)}`
- Outputs: `{OUT.relative_to(ROOT)}`
- Figure: `{FIGURE_PNG.relative_to(ROOT)}`
- Random seed: {SEED}; recipient bootstrap replicates: {N_BOOT:,}
"""
    REPORT.write_text(report, encoding="utf-8")
    for path in (REPORT, Path(__file__).resolve()):
        audit["output_hashes"][str(path.relative_to(ROOT))] = sha256(path)
    audit["environment_lock_sha256"] = sha256(ENVIRONMENT_LOCK)
    (OUT / "audit.json").write_text(
        json.dumps(json_ready(audit), indent=2, ensure_ascii=False) + "\n",
        encoding="utf-8",
    )
    print(json.dumps(json_ready(results), indent=2, ensure_ascii=False))


if __name__ == "__main__":
    main()
```
