## Supplementary_Software_1 for "Inferential boundaries of age prediction: why prediction does not establish biological age measurement": analyze_hsct_fixed_score_referent_null_probe_v0_1.py

```
#!/usr/bin/env python
"""Author-authorized fixed-score HSCT referent-null placement probe.

This script is intentionally narrow. It reads two frozen independent-unit
summary tables, never rescoring methylation or fitting an AgeClock. It persists
all analysis-ready rows, random index arrays, descriptive OLS states, result
tables, figures, signatures and reload-parity checks required by the frozen
23 August 2026 contract.
"""

from __future__ import annotations

import hashlib
import json
import os
import platform
import shutil
import sys
from pathlib import Path

import matplotlib

matplotlib.use("Agg")
import matplotlib.pyplot as plt
import numpy as np
import pandas as pd


ANALYSIS_VERSION = "hsct_fixed_score_referent_null_probe_v0_1"
N_PERMUTATIONS = 20_000
PERMUTATION_SEED = 20_260_823
N_BOOTSTRAPS = 10_000
BOOTSTRAP_SEED = 20_260_824
TOLERANCE = 1e-12


def sha256_file(path: Path) -> str:
    digest = hashlib.sha256()
    with path.open("rb") as handle:
        for chunk in iter(lambda: handle.read(1024 * 1024), b""):
            digest.update(chunk)
    return digest.hexdigest()


def write_json(path: Path, obj: object) -> None:
    path.write_text(
        json.dumps(obj, indent=2, sort_keys=True, ensure_ascii=False) + "\n",
        encoding="utf-8",
    )


def assert_columns(frame: pd.DataFrame, required: list[str], label: str) -> None:
    missing = [column for column in required if column not in frame.columns]
    if missing:
        raise RuntimeError(f"{label} missing required columns: {missing}")


def affiliation(h: np.ndarray, c: np.ndarray, s: np.ndarray) -> np.ndarray:
    denominator = np.abs(h - c)
    if np.any(denominator <= 0):
        raise RuntimeError("H=C unit encountered in affiliation calculation")
    return (np.abs(s - h) - np.abs(s - c)) / denominator


def stable_quantile(values: np.ndarray, probability: float) -> float:
    return float(np.quantile(values, probability, method="linear"))


def load_analysis_ready(project_root: Path) -> tuple[pd.DataFrame, dict[str, Path]]:
    soraas_path = (
        project_root
        / "outputs"
        / "hsct_clock_object_replacement_v0_1"
        / "primary_recipient_contrasts.tsv"
    )
    holland_path = (
        project_root
        / "outputs"
        / "hsct_holland2024_horvath_object_affiliation_v0_1"
        / "primary_pair_summary.tsv"
    )

    soraas = pd.read_csv(soraas_path, sep="\t", dtype={"recipient_id": str})
    assert_columns(
        soraas,
        ["recipient_id", "recipient_age", "donor_age", "dnam_age_recipient_blood", "d"],
        "Soraas 2019",
    )
    soraas_ready = pd.DataFrame(
        {
            "source_study": "Soraas2019",
            "unit_id": soraas["recipient_id"].astype(str),
            "H": soraas["recipient_age"].astype(float),
            "C": soraas["donor_age"].astype(float),
            "S": soraas["dnam_age_recipient_blood"].astype(float),
            "source_affiliation": np.nan,
            "source_distance_d": soraas["d"].astype(float),
        }
    )

    holland = pd.read_csv(holland_path, sep="\t", dtype={"pair_id": str})
    assert_columns(
        holland,
        [
            "pair_id",
            "donor_lineage_coordinate",
            "host_age",
            "score_minus_lineage_coordinate",
            "affiliation_index",
            "host_minus_lineage_coordinate",
        ],
        "Holland 2024",
    )
    holland_c = holland["donor_lineage_coordinate"].astype(float)
    holland_ready = pd.DataFrame(
        {
            "source_study": "Holland2024",
            "unit_id": holland["pair_id"].astype(str),
            "H": holland["host_age"].astype(float),
            "C": holland_c,
            "S": holland_c + holland["score_minus_lineage_coordinate"].astype(float),
            "source_affiliation": holland["affiliation_index"].astype(float),
            "source_distance_d": np.nan,
        }
    )

    ready = pd.concat([soraas_ready, holland_ready], ignore_index=True)
    ready = ready.sort_values(["source_study", "unit_id"], kind="mergesort").reset_index(drop=True)
    ready["analysis_row"] = np.arange(len(ready), dtype=int)
    ready["H_minus_C"] = ready["H"] - ready["C"]
    if (ready["H_minus_C"] == 0).any():
        bad = ready.loc[ready["H_minus_C"] == 0, ["source_study", "unit_id"]]
        raise RuntimeError(f"Ineligible H=C units found: {bad.to_dict(orient='records')}")
    ready["stratum"] = np.where(ready["H_minus_C"] > 0, "host_older", "host_younger")
    ready["I"] = affiliation(
        ready["H"].to_numpy(float),
        ready["C"].to_numpy(float),
        ready["S"].to_numpy(float),
    )
    ready = ready[
        [
            "analysis_row",
            "source_study",
            "unit_id",
            "stratum",
            "H",
            "C",
            "S",
            "H_minus_C",
            "I",
            "source_affiliation",
            "source_distance_d",
        ]
    ]
    return ready, {"soraas2019": soraas_path, "holland2024": holland_path}


def build_affiliation_matrix(frame: pd.DataFrame) -> np.ndarray:
    h = frame["H"].to_numpy(float)[:, None]
    c = frame["C"].to_numpy(float)[:, None]
    s = frame["S"].to_numpy(float)[None, :]
    denominator = np.abs(h - c)
    return (np.abs(s - h) - np.abs(s - c)) / denominator


def make_permutation_maps(n: int, strata: np.ndarray, rng: np.random.Generator) -> np.ndarray:
    maps = np.tile(np.arange(n, dtype=np.int32), (N_PERMUTATIONS, 1))
    for label in ("host_older", "host_younger"):
        positions = np.flatnonzero(strata == label).astype(np.int32)
        for replicate in range(N_PERMUTATIONS):
            maps[replicate, positions] = rng.permutation(positions)
    return maps


def permutation_statistics(
    matrix: np.ndarray, maps: np.ndarray, strata: np.ndarray
) -> pd.DataFrame:
    row_positions = np.arange(matrix.shape[0], dtype=np.int32)[None, :]
    permuted_i = matrix[row_positions, maps]
    older = strata == "host_older"
    younger = strata == "host_younger"
    older_mean = permuted_i[:, older].mean(axis=1)
    younger_mean = permuted_i[:, younger].mean(axis=1)
    balanced = 0.5 * (older_mean + younger_mean)
    return pd.DataFrame(
        {
            "permutation_id": np.arange(N_PERMUTATIONS, dtype=int),
            "mean_I_host_older": older_mean,
            "mean_I_host_younger": younger_mean,
            "I_bal": balanced,
        }
    )


def make_bootstrap_maps(n: int, strata: np.ndarray, rng: np.random.Generator) -> np.ndarray:
    maps = np.empty((N_BOOTSTRAPS, n), dtype=np.int32)
    for label in ("host_older", "host_younger"):
        positions = np.flatnonzero(strata == label).astype(np.int32)
        maps[:, positions] = rng.choice(
            positions, size=(N_BOOTSTRAPS, len(positions)), replace=True
        )
    return maps


def bootstrap_statistics(
    frame: pd.DataFrame, matrix: np.ndarray, maps: np.ndarray
) -> pd.DataFrame:
    strata = frame["stratum"].to_numpy(str)
    observed_i = frame["I"].to_numpy(float)
    outputs: dict[str, np.ndarray] = {}
    exact_null_outputs: dict[str, np.ndarray] = {}

    for label in ("host_older", "host_younger"):
        positions = np.flatnonzero(strata == label)
        selected = maps[:, positions]
        outputs[label] = observed_i[selected].mean(axis=1)

        local_lookup = np.full(len(frame), -1, dtype=int)
        local_lookup[positions] = np.arange(len(positions), dtype=int)
        selected_local = local_lookup[selected]
        counts = np.zeros((N_BOOTSTRAPS, len(positions)), dtype=float)
        rows = np.repeat(np.arange(N_BOOTSTRAPS), len(positions))
        np.add.at(counts, (rows, selected_local.ravel()), 1.0)
        submatrix = matrix[np.ix_(positions, positions)]
        exact_null_outputs[label] = (
            np.einsum("bi,ij,bj->b", counts, submatrix, counts, optimize=True)
            / float(len(positions) ** 2)
        )

    observed_balanced = 0.5 * (outputs["host_older"] + outputs["host_younger"])
    exact_null_balanced = 0.5 * (
        exact_null_outputs["host_older"] + exact_null_outputs["host_younger"]
    )
    return pd.DataFrame(
        {
            "bootstrap_id": np.arange(N_BOOTSTRAPS, dtype=int),
            "observed_I_bal": observed_balanced,
            "exact_permutation_null_mean_I_bal": exact_null_balanced,
            "observed_minus_null_mean": observed_balanced - exact_null_balanced,
        }
    )


def ols_state(x: np.ndarray, y: np.ndarray) -> tuple[float, float, np.ndarray]:
    x_mean = float(x.mean())
    y_mean = float(y.mean())
    denominator = float(np.sum((x - x_mean) ** 2))
    if denominator <= 0:
        raise RuntimeError("OLS predictor has zero variance")
    slope = float(np.sum((x - x_mean) * (y - y_mean)) / denominator)
    intercept = float(y_mean - slope * x_mean)
    fitted = intercept + slope * x
    return slope, intercept, fitted


def create_calibration_artifacts(
    ready: pd.DataFrame, artifact_dir: Path, signature_core: dict[str, object]
) -> tuple[pd.DataFrame, dict[str, object]]:
    artifact_dir.mkdir(parents=True, exist_ok=False)
    rows: list[dict[str, object]] = []
    model_ids: list[str] = []
    slopes: list[float] = []
    intercepts: list[float] = []
    prediction_arrays: dict[str, np.ndarray] = {}
    row_index_arrays: dict[str, np.ndarray] = {}

    for study in ("Soraas2019", "Holland2024"):
        study_frame = ready.loc[ready["source_study"] == study].copy()
        for scope in ("overall", "host_older", "host_younger"):
            scope_frame = (
                study_frame
                if scope == "overall"
                else study_frame.loc[study_frame["stratum"] == scope]
            )
            for predictor in ("C", "H"):
                x = scope_frame[predictor].to_numpy(float)
                y = scope_frame["S"].to_numpy(float)
                slope, intercept, fitted = ols_state(x, y)
                model_id = f"{study}__{scope}__S_on_{predictor}"
                rows.append(
                    {
                        "model_id": model_id,
                        "source_study": study,
                        "scope": scope,
                        "outcome": "S",
                        "predictor": predictor,
                        "n_units": len(scope_frame),
                        "slope": slope,
                        "intercept": intercept,
                    }
                )
                model_ids.append(model_id)
                slopes.append(slope)
                intercepts.append(intercept)
                prediction_arrays[f"pred_{len(model_ids)-1:02d}"] = fitted
                row_index_arrays[f"rows_{len(model_ids)-1:02d}"] = scope_frame[
                    "analysis_row"
                ].to_numpy(np.int32)

    model_path = artifact_dir / "calibration_model_state.npz"
    prediction_path = artifact_dir / "calibration_fitted_values.npz"
    np.savez_compressed(
        model_path,
        model_id=np.asarray(model_ids, dtype="U80"),
        slope=np.asarray(slopes, dtype=float),
        intercept=np.asarray(intercepts, dtype=float),
    )
    np.savez_compressed(prediction_path, **prediction_arrays, **row_index_arrays)

    model_signature = {
        **signature_core,
        "artifact_type": "closed_form_ordinary_least_squares",
        "outcome": "S",
        "ordered_predictors": ["C", "H"],
        "scopes": ["overall", "host_older", "host_younger"],
        "preprocessing": "none",
        "selection": "none",
        "calibration_applied": False,
        "grouping_unit": {
            "Soraas2019": "recipient",
            "Holland2024": "donor-recipient pair",
        },
        "n_models": len(model_ids),
    }
    signature_path = artifact_dir / "calibration_model_signature.json"
    write_json(signature_path, model_signature)

    max_prediction_error = 0.0
    with (
        np.load(model_path, allow_pickle=False) as loaded_models,
        np.load(prediction_path, allow_pickle=False) as loaded_predictions,
    ):
        for index, model_id in enumerate(loaded_models["model_id"].tolist()):
            row_ids = loaded_predictions[f"rows_{index:02d}"].astype(int)
            predictor = "C" if str(model_id).endswith("S_on_C") else "H"
            x = ready.set_index("analysis_row").loc[row_ids, predictor].to_numpy(float)
            reloaded = (
                float(loaded_models["intercept"][index])
                + float(loaded_models["slope"][index]) * x
            )
            saved = loaded_predictions[f"pred_{index:02d}"]
            max_prediction_error = max(
                max_prediction_error, float(np.max(np.abs(reloaded - saved)))
            )
    if max_prediction_error > TOLERANCE:
        raise RuntimeError(
            f"Calibration model reload parity failed: {max_prediction_error}"
        )

    return pd.DataFrame(rows), {
        "calibration_model_count": len(model_ids),
        "calibration_reload_max_abs_prediction_error": max_prediction_error,
        "calibration_reload_parity": True,
    }


def summarize_constant_controls(frame: pd.DataFrame) -> list[dict[str, object]]:
    rows: list[dict[str, object]] = []
    h = frame["H"].to_numpy(float)
    c = frame["C"].to_numpy(float)
    strata = frame["stratum"].to_numpy(str)
    for control in ("marginal_mean_score", "marginal_median_score"):
        constant = (
            float(frame["S"].mean())
            if control == "marginal_mean_score"
            else float(frame["S"].median())
        )
        values = affiliation(h, c, np.full(len(frame), constant, dtype=float))
        older = float(values[strata == "host_older"].mean())
        younger = float(values[strata == "host_younger"].mean())
        rows.append(
            {
                "control": control,
                "constant_score": constant,
                "mean_I_overall": float(values.mean()),
                "mean_I_host_older": older,
                "mean_I_host_younger": younger,
                "I_bal": 0.5 * (older + younger),
            }
        )
    return rows


def summarize_study(
    study: str,
    frame: pd.DataFrame,
    matrix: np.ndarray,
    perm_stats: pd.DataFrame,
    boot_stats: pd.DataFrame,
) -> dict[str, object]:
    strata = frame["stratum"].to_numpy(str)
    values = frame["I"].to_numpy(float)
    observed_older = float(values[strata == "host_older"].mean())
    observed_younger = float(values[strata == "host_younger"].mean())
    observed_balanced = 0.5 * (observed_older + observed_younger)
    older_positions = np.flatnonzero(strata == "host_older")
    younger_positions = np.flatnonzero(strata == "host_younger")
    exact_null_older = float(matrix[np.ix_(older_positions, older_positions)].mean())
    exact_null_younger = float(matrix[np.ix_(younger_positions, younger_positions)].mean())
    exact_null_balanced = 0.5 * (exact_null_older + exact_null_younger)
    perm_balanced = perm_stats["I_bal"].to_numpy(float)
    perm_mean = float(perm_balanced.mean())
    exceedances = int(np.sum(perm_balanced >= observed_balanced))
    p_value = float((1 + exceedances) / (1 + N_PERMUTATIONS))
    observed_boot = boot_stats["observed_I_bal"].to_numpy(float)
    contrast_boot = boot_stats["observed_minus_null_mean"].to_numpy(float)
    return {
        "source_study": study,
        "n_overall": len(frame),
        "n_host_older": int(np.sum(strata == "host_older")),
        "n_host_younger": int(np.sum(strata == "host_younger")),
        "mean_I_overall": float(values.mean()),
        "mean_I_host_older": observed_older,
        "mean_I_host_younger": observed_younger,
        "I_bal_observed": observed_balanced,
        "permutation_null_mean_host_older": float(
            perm_stats["mean_I_host_older"].mean()
        ),
        "permutation_null_mean_host_younger": float(
            perm_stats["mean_I_host_younger"].mean()
        ),
        "I_bal_permutation_null_mean": perm_mean,
        "I_bal_permutation_q025": stable_quantile(perm_balanced, 0.025),
        "I_bal_permutation_q975": stable_quantile(perm_balanced, 0.975),
        "permutation_exceedances": exceedances,
        "permutation_p_one_sided": p_value,
        "exact_permutation_null_mean_host_older": exact_null_older,
        "exact_permutation_null_mean_host_younger": exact_null_younger,
        "exact_permutation_null_mean_I_bal": exact_null_balanced,
        "observed_minus_permutation_null_mean": observed_balanced - perm_mean,
        "observed_I_bal_bootstrap_ci_low": stable_quantile(observed_boot, 0.025),
        "observed_I_bal_bootstrap_ci_high": stable_quantile(observed_boot, 0.975),
        "observed_minus_null_bootstrap_ci_low": stable_quantile(contrast_boot, 0.025),
        "observed_minus_null_bootstrap_ci_high": stable_quantile(contrast_boot, 0.975),
    }


def draw_figure(
    ready: pd.DataFrame,
    permutation_frames: dict[str, pd.DataFrame],
    summaries: dict[str, dict[str, object]],
    png_path: Path,
    svg_path: Path,
) -> None:
    colours = {"host_older": "#C75D3B", "host_younger": "#D98B2B"}
    studies = ("Soraas2019", "Holland2024")
    titles = {"Soraas2019": "Søraas 2019", "Holland2024": "Holland 2024"}
    fig, axes = plt.subplots(2, 2, figsize=(11.2, 7.2), constrained_layout=True)
    for column, study in enumerate(studies):
        frame = ready.loc[ready["source_study"] == study]
        ax = axes[0, column]
        for stratum in ("host_older", "host_younger"):
            subset = frame.loc[frame["stratum"] == stratum]
            ax.scatter(
                subset["H_minus_C"],
                subset["S"] - subset["C"],
                s=34,
                alpha=0.85,
                color=colours[stratum],
                label=stratum.replace("_", " "),
            )
        limits = np.asarray(ax.get_xlim())
        low = min(float(frame["H_minus_C"].min()), float((frame["S"] - frame["C"]).min()))
        high = max(float(frame["H_minus_C"].max()), float((frame["S"] - frame["C"]).max()))
        span = high - low
        line = np.linspace(low - 0.08 * span, high + 0.08 * span, 100)
        ax.axhline(0, color="#008A92", linewidth=1.6, label="lineage referent")
        ax.plot(line, line, "--", color="#555555", linewidth=1.2, label="host referent")
        ax.set_title(titles[study])
        ax.set_xlabel("Host minus lineage coordinate (H − C)")
        ax.set_ylabel("Score minus lineage coordinate (S − C)")
        ax.grid(alpha=0.18)
        if column == 0:
            ax.legend(frameon=False, fontsize=8)

        ax_null = axes[1, column]
        perm_values = permutation_frames[study]["I_bal"].to_numpy(float)
        observed = float(summaries[study]["I_bal_observed"])
        q975 = float(summaries[study]["I_bal_permutation_q975"])
        ax_null.hist(perm_values, bins=45, color="#B8C7D1", edgecolor="none")
        ax_null.axvline(q975, color="#59636E", linestyle="--", linewidth=1.4, label="null 97.5%")
        ax_null.axvline(observed, color="#7A1F5C", linewidth=2.1, label="observed")
        ax_null.set_xlabel("Sign-balanced affiliation under within-sign permutation")
        ax_null.set_ylabel("Permutation count")
        ax_null.grid(alpha=0.15, axis="y")
        ax_null.legend(frameon=False, fontsize=8)
        ax_null.text(
            0.02,
            0.97,
            f"Observed={observed:.3f}\nNull 97.5%={q975:.3f}",
            transform=ax_null.transAxes,
            va="top",
            fontsize=9,
        )

    fig.suptitle(
        "Post-HSCT fixed-score affiliation exceeds sign-preserving marginal-score geometry",
        fontsize=14,
        fontweight="bold",
    )
    fig.savefig(png_path, dpi=240, metadata={"Software": "LHState"})
    fig.savefig(svg_path, metadata={"Date": None})
    plt.close(fig)


def create_input_manifest(paths: dict[str, Path], destination: Path) -> dict[str, str]:
    rows = []
    hashes: dict[str, str] = {}
    for role, path in paths.items():
        digest = sha256_file(path)
        hashes[role] = digest
        rows.append(
            {
                "role": role,
                "path": str(path),
                "sha256": digest,
                "bytes": path.stat().st_size,
            }
        )
    pd.DataFrame(rows).to_csv(destination, sep="\t", index=False)
    return hashes


def artifact_role(relative_path: str) -> str:
    if relative_path.startswith("model_artifacts/"):
        return "model_artifact"
    if relative_path.endswith((".png", ".svg")):
        return "figure"
    if relative_path.endswith(".json"):
        return "audit_or_decision"
    if relative_path.endswith((".tsv", ".tsv.gz", ".npz")):
        return "analysis_artifact"
    if relative_path.endswith(".md"):
        return "report"
    return "other"


def write_artifact_manifest(stage: Path) -> None:
    rows = []
    for path in sorted(stage.rglob("*")):
        if not path.is_file() or path.name == "artifact_manifest.tsv":
            continue
        relative = path.relative_to(stage).as_posix()
        rows.append(
            {
                "relative_path": relative,
                "role": artifact_role(relative),
                "bytes": path.stat().st_size,
                "sha256": sha256_file(path),
            }
        )
    pd.DataFrame(rows).to_csv(stage / "artifact_manifest.tsv", sep="\t", index=False)


def validate_cached(output_dir: Path) -> str:
    manifest = pd.read_csv(output_dir / "artifact_manifest.tsv", sep="\t")
    for row in manifest.itertuples(index=False):
        path = output_dir / row.relative_path
        if not path.is_file():
            raise RuntimeError(f"Cached artifact missing: {path}")
        if sha256_file(path) != row.sha256:
            raise RuntimeError(f"Cached artifact hash mismatch: {path}")
    decision = json.loads((output_dir / "decision.json").read_text(encoding="utf-8"))
    audit = json.loads((output_dir / "deterministic_audit.json").read_text(encoding="utf-8"))
    if audit.get("status") != "PASS":
        raise RuntimeError("Cached deterministic audit is not PASS")
    return str(decision["placement"])


def write_report(
    path: Path,
    placement: str,
    summaries: dict[str, dict[str, object]],
    gates: list[dict[str, object]],
) -> None:
    h = summaries["Holland2024"]
    s = summaries["Soraas2019"]
    gate_lines = "\n".join(
        f"- {'PASS' if gate['passed'] else 'FAIL'} — {gate['condition']}: {gate['detail']}"
        for gate in gates
    )
    report = f"""# HSCT fixed-score referent-null placement probe

Analysis version: `{ANALYSIS_VERSION}`  
Decision: **{placement}**

## Result

In Holland 2024, the observed sign-balanced affiliation was {h['I_bal_observed']:.3f},
compared with a within-sign permutation-null mean of
{h['I_bal_permutation_null_mean']:.3f} and 97.5th percentile of
{h['I_bal_permutation_q975']:.3f}. The one-sided finite-permutation p-value was
{h['permutation_p_one_sided']:.6g}. The observed-minus-null bootstrap interval
was {h['observed_minus_null_bootstrap_ci_low']:.3f} to
{h['observed_minus_null_bootstrap_ci_high']:.3f}.

Both Holland sign strata exceeded their permutation-null means. Søraas 2019
showed the same direction: observed sign-balanced affiliation
{s['I_bal_observed']:.3f} versus permutation-null mean
{s['I_bal_permutation_null_mean']:.3f}.

## Placement gate

{gate_lines}

## Allowed conclusion

> The evaluated post-HSCT recipient-blood readouts affiliated with the
> donor-derived haematopoietic lineage's chronological coordinate beyond the
> affiliation expected from the observed marginal score and donor/recipient age
> geometry.

This does not identify a unique lineage biological age, exclude host influence,
establish organismal age replacement, or validate rejuvenation. The two studies
partly overlap and were analysed separately.

## Scope

The probe used only the two frozen independent-unit summary tables. It did not
rescore methylation, fit an AgeClock, add a clock family, change a threshold,
use an external dataset or search an outcome. Ordinary least-squares fits were
descriptive calibration diagnostics and were not used to recalibrate the fixed
scores.
"""
    path.write_text(report, encoding="utf-8")


def main() -> int:
    project_root = Path(__file__).resolve().parents[1]
    output_dir = project_root / "outputs" / ANALYSIS_VERSION
    stage = project_root / "outputs" / f"{ANALYSIS_VERSION}.__staging__"
    spec_path = project_root / "analysis" / "HSCT_FIXED_SCORE_REFERENT_NULL_PROBE_SPEC_20260823.md"
    script_path = Path(__file__).resolve()
    lock_path = project_root / "environment.win-64.lock.txt"

    if output_dir.exists():
        placement = validate_cached(output_dir)
        print(json.dumps({"status": "REUSED_VERIFIED_ARTIFACTS", "placement": placement}))
        return 0
    if stage.exists():
        raise RuntimeError(
            f"Incomplete staging directory exists; fail closed instead of overwriting: {stage}"
        )
    stage.mkdir(parents=True, exist_ok=False)

    try:
        ready, frozen_inputs = load_analysis_ready(project_root)
        expected_counts = {"Soraas2019": 19, "Holland2024": 61}
        observed_counts = ready.groupby("source_study").size().to_dict()
        if observed_counts != expected_counts:
            raise RuntimeError(
                f"Independent-unit counts changed: expected {expected_counts}, got {observed_counts}"
            )
        if ready[["H", "C", "S", "I"]].isna().any().any():
            raise RuntimeError("Non-finite analysis-ready value")
        if not np.all(np.isfinite(ready[["H", "C", "S", "I"]].to_numpy(float))):
            raise RuntimeError("Non-finite analysis-ready value")
        if ready.duplicated(["source_study", "unit_id"]).any():
            raise RuntimeError("Duplicate independent unit")

        soraas_ready = ready.loc[ready["source_study"] == "Soraas2019"]
        reconstructed_d = np.abs(soraas_ready["S"] - soraas_ready["C"]) - np.abs(
            soraas_ready["S"] - soraas_ready["H"]
        )
        soraas_d_difference = np.abs(
            reconstructed_d - soraas_ready["source_distance_d"].to_numpy(float)
        )
        soraas_d_max_difference = float(np.max(soraas_d_difference))
        soraas_d_n_different = int(np.sum(soraas_d_difference > TOLERANCE))
        # The frozen recipient summary stores the mean of sample-level distance
        # contrasts, whereas this new contract requires I to be recomputed from
        # recipient-mean H, C and S. Absolute value and averaging do not commute;
        # a difference is therefore expected for a repeated recipient. Require
        # the directional classification, not the nonlinear magnitude, to agree.
        source_d = soraas_ready["source_distance_d"].to_numpy(float)
        soraas_d_direction_agreement = bool(
            np.all(
                (np.sign(reconstructed_d.to_numpy(float)) == np.sign(source_d))
                | (
                    (np.abs(reconstructed_d.to_numpy(float)) <= TOLERANCE)
                    & (np.abs(source_d) <= TOLERANCE)
                )
            )
        )
        if not soraas_d_direction_agreement:
            raise RuntimeError("Soraas reconstructed pair-level direction disagrees with source")

        holland_ready = ready.loc[ready["source_study"] == "Holland2024"]
        holland_hc_error = float(
            np.max(
                np.abs(
                    holland_ready["H_minus_C"]
                    - (holland_ready["H"] - holland_ready["C"])
                )
            )
        )
        if holland_hc_error > TOLERANCE:
            raise RuntimeError(f"Holland H-C reconstruction failed: {holland_hc_error}")

        ready.to_csv(stage / "analysis_ready_pairs.tsv", sep="\t", index=False)

        input_paths = {
            "frozen_spec": spec_path,
            "analysis_script": script_path,
            "environment_lock": lock_path,
            **frozen_inputs,
        }
        input_hashes = create_input_manifest(input_paths, stage / "input_manifest.tsv")
        signature_core: dict[str, object] = {
            "analysis_version": ANALYSIS_VERSION,
            "input_hashes": input_hashes,
            "ordered_analysis_rows_sha256": hashlib.sha256(
                ready[["source_study", "unit_id", "H", "C", "S"]]
                .to_csv(index=False)
                .encode("utf-8")
            ).hexdigest(),
            "permutation_seed": PERMUTATION_SEED,
            "bootstrap_seed": BOOTSTRAP_SEED,
            "n_permutations": N_PERMUTATIONS,
            "n_bootstraps": N_BOOTSTRAPS,
        }

        calibration, model_audit = create_calibration_artifacts(
            ready, stage / "model_artifacts", signature_core
        )
        calibration.to_csv(stage / "calibration_diagnostics.tsv", sep="\t", index=False)

        permutation_rng = np.random.default_rng(PERMUTATION_SEED)
        bootstrap_rng = np.random.default_rng(BOOTSTRAP_SEED)
        permutation_maps: dict[str, np.ndarray] = {}
        bootstrap_maps: dict[str, np.ndarray] = {}
        permutation_frames: dict[str, pd.DataFrame] = {}
        bootstrap_frames: dict[str, pd.DataFrame] = {}
        matrices: dict[str, np.ndarray] = {}
        constant_rows: list[dict[str, object]] = []

        for study in ("Soraas2019", "Holland2024"):
            frame = ready.loc[ready["source_study"] == study].reset_index(drop=True)
            strata = frame["stratum"].to_numpy(str)
            matrix = build_affiliation_matrix(frame)
            matrices[study] = matrix
            permutation_maps[study] = make_permutation_maps(len(frame), strata, permutation_rng)
            bootstrap_maps[study] = make_bootstrap_maps(len(frame), strata, bootstrap_rng)
            permutation_frames[study] = permutation_statistics(
                matrix, permutation_maps[study], strata
            )
            bootstrap_frames[study] = bootstrap_statistics(
                frame, matrix, bootstrap_maps[study]
            )
            for row in summarize_constant_controls(frame):
                constant_rows.append({"source_study": study, **row})

        np.savez_compressed(
            stage / "permutation_indices.npz",
            Soraas2019=permutation_maps["Soraas2019"],
            Holland2024=permutation_maps["Holland2024"],
        )
        np.savez_compressed(
            stage / "bootstrap_indices.npz",
            Soraas2019=bootstrap_maps["Soraas2019"],
            Holland2024=bootstrap_maps["Holland2024"],
        )

        permutation_output = pd.concat(
            [
                frame.assign(source_study=study)[
                    [
                        "source_study",
                        "permutation_id",
                        "mean_I_host_older",
                        "mean_I_host_younger",
                        "I_bal",
                    ]
                ]
                for study, frame in permutation_frames.items()
            ],
            ignore_index=True,
        )
        bootstrap_output = pd.concat(
            [
                frame.assign(source_study=study)[
                    [
                        "source_study",
                        "bootstrap_id",
                        "observed_I_bal",
                        "exact_permutation_null_mean_I_bal",
                        "observed_minus_null_mean",
                    ]
                ]
                for study, frame in bootstrap_frames.items()
            ],
            ignore_index=True,
        )
        compression = {"method": "gzip", "compresslevel": 9, "mtime": 0}
        permutation_output.to_csv(
            stage / "permutation_statistics.tsv.gz",
            sep="\t",
            index=False,
            compression=compression,
        )
        bootstrap_output.to_csv(
            stage / "bootstrap_statistics.tsv.gz",
            sep="\t",
            index=False,
            compression=compression,
        )
        pd.DataFrame(constant_rows).to_csv(
            stage / "constant_score_controls.tsv", sep="\t", index=False
        )

        max_perm_stat_error = 0.0
        max_boot_stat_error = 0.0
        index_checks: dict[str, bool] = {}
        with (
            np.load(stage / "permutation_indices.npz", allow_pickle=False) as loaded_perm,
            np.load(stage / "bootstrap_indices.npz", allow_pickle=False) as loaded_boot,
        ):
            for study in ("Soraas2019", "Holland2024"):
                frame = ready.loc[ready["source_study"] == study].reset_index(drop=True)
                strata = frame["stratum"].to_numpy(str)
                perm_equal = np.array_equal(loaded_perm[study], permutation_maps[study])
                boot_equal = np.array_equal(loaded_boot[study], bootstrap_maps[study])
                index_checks[f"{study}_permutation_index_reload_exact"] = bool(perm_equal)
                index_checks[f"{study}_bootstrap_index_reload_exact"] = bool(boot_equal)
                if not perm_equal or not boot_equal:
                    raise RuntimeError(f"Index-array reload parity failed for {study}")

                for label in ("host_older", "host_younger"):
                    positions = np.flatnonzero(strata == label)
                    expected = np.sort(positions)[None, :]
                    sorted_permutations = np.sort(
                        loaded_perm[study][:, positions], axis=1
                    )
                    if not np.array_equal(
                        sorted_permutations,
                        np.broadcast_to(expected, sorted_permutations.shape),
                    ):
                        raise RuntimeError(
                            f"Invalid within-stratum permutation for {study}, {label}"
                        )
                    if not np.all(np.isin(loaded_boot[study][:, positions], positions)):
                        raise RuntimeError(f"Bootstrap crossed strata for {study}, {label}")

                perm_recomputed = permutation_statistics(
                    matrices[study], loaded_perm[study], strata
                )
                boot_recomputed = bootstrap_statistics(
                    frame, matrices[study], loaded_boot[study]
                )
                perm_error = float(
                    np.max(
                        np.abs(
                            perm_recomputed[
                                ["mean_I_host_older", "mean_I_host_younger", "I_bal"]
                            ].to_numpy(float)
                            - permutation_frames[study][
                                ["mean_I_host_older", "mean_I_host_younger", "I_bal"]
                            ].to_numpy(float)
                        )
                    )
                )
                boot_error = float(
                    np.max(
                        np.abs(
                            boot_recomputed[
                                [
                                    "observed_I_bal",
                                    "exact_permutation_null_mean_I_bal",
                                    "observed_minus_null_mean",
                                ]
                            ].to_numpy(float)
                            - bootstrap_frames[study][
                                [
                                    "observed_I_bal",
                                    "exact_permutation_null_mean_I_bal",
                                    "observed_minus_null_mean",
                                ]
                            ].to_numpy(float)
                        )
                    )
                )
                max_perm_stat_error = max(max_perm_stat_error, perm_error)
                max_boot_stat_error = max(max_boot_stat_error, boot_error)
        if max_perm_stat_error > TOLERANCE or max_boot_stat_error > TOLERANCE:
            raise RuntimeError("Reloaded index arrays did not reproduce statistics")

        summaries: dict[str, dict[str, object]] = {}
        for study in ("Soraas2019", "Holland2024"):
            frame = ready.loc[ready["source_study"] == study].reset_index(drop=True)
            summaries[study] = summarize_study(
                study,
                frame,
                matrices[study],
                permutation_frames[study],
                bootstrap_frames[study],
            )
        observed_summary = pd.DataFrame(list(summaries.values()))
        observed_summary.to_csv(stage / "observed_summary.tsv", sep="\t", index=False)

        holland = summaries["Holland2024"]
        soraas = summaries["Soraas2019"]
        holland_c_slopes = calibration.loc[
            (calibration["source_study"] == "Holland2024")
            & (calibration["scope"].isin(["host_older", "host_younger"]))
            & (calibration["predictor"] == "C"),
            ["scope", "slope"],
        ].set_index("scope")["slope"]

        deterministic_checks_pass = (
            soraas_d_direction_agreement
            and holland_hc_error <= TOLERANCE
            and max_perm_stat_error <= TOLERANCE
            and max_boot_stat_error <= TOLERANCE
            and all(index_checks.values())
            and bool(model_audit["calibration_reload_parity"])
        )
        gates = [
            {
                "gate": 1,
                "condition": "Holland observed I_bal exceeds permutation-null 97.5th percentile",
                "passed": bool(
                    holland["I_bal_observed"] > holland["I_bal_permutation_q975"]
                ),
                "detail": f"{holland['I_bal_observed']:.6f} > {holland['I_bal_permutation_q975']:.6f}",
            },
            {
                "gate": 2,
                "condition": "Holland observed-minus-null bootstrap interval is above zero",
                "passed": bool(holland["observed_minus_null_bootstrap_ci_low"] > 0),
                "detail": (
                    f"95% interval {holland['observed_minus_null_bootstrap_ci_low']:.6f} "
                    f"to {holland['observed_minus_null_bootstrap_ci_high']:.6f}"
                ),
            },
            {
                "gate": 3,
                "condition": "Holland observed affiliation exceeds null mean in both sign strata",
                "passed": bool(
                    holland["mean_I_host_older"]
                    > holland["permutation_null_mean_host_older"]
                    and holland["mean_I_host_younger"]
                    > holland["permutation_null_mean_host_younger"]
                ),
                "detail": (
                    f"older {holland['mean_I_host_older']:.6f} vs "
                    f"{holland['permutation_null_mean_host_older']:.6f}; younger "
                    f"{holland['mean_I_host_younger']:.6f} vs "
                    f"{holland['permutation_null_mean_host_younger']:.6f}"
                ),
            },
            {
                "gate": 4,
                "condition": "Soraas observed-minus-null mean I_bal has the same positive sign",
                "passed": bool(soraas["observed_minus_permutation_null_mean"] > 0),
                "detail": f"contrast {soraas['observed_minus_permutation_null_mean']:.6f}",
            },
            {
                "gate": 5,
                "condition": "Holland score-on-C slopes are positive in both sign strata",
                "passed": bool(
                    holland_c_slopes["host_older"] > 0
                    and holland_c_slopes["host_younger"] > 0
                ),
                "detail": (
                    f"older {holland_c_slopes['host_older']:.6f}; "
                    f"younger {holland_c_slopes['host_younger']:.6f}"
                ),
            },
            {
                "gate": 6,
                "condition": "Source reconstruction and deterministic reload checks pass",
                "passed": bool(deterministic_checks_pass),
                "detail": (
                    f"Soraas direction agreement={soraas_d_direction_agreement}; "
                    f"Holland H-C error={holland_hc_error:.3g}; "
                    f"permutation parity error={max_perm_stat_error:.3g}; bootstrap parity error={max_boot_stat_error:.3g}"
                ),
            },
        ]
        placement = "MAIN" if all(gate["passed"] for gate in gates) else "EXTENDED_DATA"

        decision = {
            "analysis_version": ANALYSIS_VERSION,
            "placement": placement,
            "gate_status": gates,
            "allowed_claim": (
                "The evaluated post-HSCT recipient-blood readouts affiliated with the "
                "donor-derived haematopoietic lineage's chronological coordinate beyond "
                "the affiliation expected from the observed marginal score and donor/recipient "
                "age geometry."
            ),
            "forbidden_claims": [
                "unique lineage biological age",
                "absence of host influence",
                "organismal age replacement",
                "rejuvenation or clinical benefit",
                "independent replication of the two partly overlapping studies",
                "proof of the analytical non-identification proposition",
            ],
        }
        write_json(stage / "decision.json", decision)

        deterministic_audit = {
            "status": "PASS" if deterministic_checks_pass else "FAIL",
            "analysis_version": ANALYSIS_VERSION,
            "input_hashes": input_hashes,
            "independent_unit_counts": observed_counts,
            "stratum_counts": {
                f"{study}::{stratum}": int(count)
                for (study, stratum), count in ready.groupby(
                    ["source_study", "stratum"]
                ).size().items()
            },
            "soraas_source_sample_d_vs_recomputed_pair_mean_d_max_abs_difference": soraas_d_max_difference,
            "soraas_source_sample_d_vs_recomputed_pair_mean_d_n_different": soraas_d_n_different,
            "soraas_recomputed_direction_agrees_with_source": soraas_d_direction_agreement,
            "holland_H_minus_C_reconstruction_max_abs_error": holland_hc_error,
            "permutation_index_and_statistic_checks": index_checks,
            "permutation_statistic_reload_max_abs_error": max_perm_stat_error,
            "bootstrap_statistic_reload_max_abs_error": max_boot_stat_error,
            **model_audit,
            "numerical_tolerance": TOLERANCE,
            "randomization": {
                "permutation_seed": PERMUTATION_SEED,
                "n_permutations": N_PERMUTATIONS,
                "bootstrap_seed": BOOTSTRAP_SEED,
                "n_bootstraps": N_BOOTSTRAPS,
            },
            "environment": {
                "python": sys.version,
                "executable": sys.executable,
                "platform": platform.platform(),
                "numpy": np.__version__,
                "pandas": pd.__version__,
                "matplotlib": matplotlib.__version__,
                "environment_lock_sha256": sha256_file(lock_path),
            },
            "scope_checks": {
                "clock_rescored": False,
                "new_clock_added": False,
                "threshold_changed": False,
                "external_data_added": False,
                "alternative_null_added": False,
                "studies_pooled": False,
            },
        }
        write_json(stage / "deterministic_audit.json", deterministic_audit)

        draw_figure(
            ready,
            permutation_frames,
            summaries,
            stage / "HSCT_FIXED_SCORE_REFERENT_NULL_DECISION.png",
            stage / "HSCT_FIXED_SCORE_REFERENT_NULL_DECISION.svg",
        )
        write_report(stage / "REPORT.md", placement, summaries, gates)
        write_artifact_manifest(stage)
        stage.rename(output_dir)
        print(
            json.dumps(
                {
                    "status": "COMPLETE",
                    "placement": placement,
                    "output_dir": str(output_dir),
                },
                ensure_ascii=False,
            )
        )
        return 0
    except Exception:
        # Preserve the incomplete staging directory for diagnosis. Never promote
        # it or overwrite it silently on a later run.
        raise


if __name__ == "__main__":
    raise SystemExit(main())
```
