## Supplementary_Software_1 for "Inferential boundaries of age prediction: why prediction does not establish biological age measurement": analyze_hsct_referent_regression_sensitivity_v0_1.py

```
#!/usr/bin/env python
"""Conventional continuous-regression sensitivity for frozen HSCT units.

This analysis never rescales or recalibrates an age score.  It reads the
independent-unit interface frozen by the referent-null probe and fits, within
each study, S-A_R = alpha + beta (A_L-A_R) + epsilon.  All closed-form fitted
states, bootstrap maps, fitted values and verification records are persisted.
"""

from __future__ import annotations

import hashlib
import json
import platform
import sys
from pathlib import Path

import numpy as np
import pandas as pd
import scipy
from scipy import stats


ROOT = Path(__file__).resolve().parents[1]
ANALYSIS_VERSION = "hsct_referent_regression_clock_sensitivity_v0_1"
SPEC = ROOT / "analysis" / "HSCT_REFERENT_REGRESSION_CLOCK_SENSITIVITY_SPEC_20260826.md"
P2_LOCK = (
    ROOT
    / "manuscript"
    / "nature_first"
    / "next_round_20260826"
    / "P2_ANALYSIS_DIRECTION_LOCK_20260826.md"
)
FROZEN_READY = (
    ROOT
    / "outputs"
    / "hsct_fixed_score_referent_null_probe_v0_1"
    / "analysis_ready_pairs.tsv"
)
OUT_ROOT = ROOT / "outputs" / ANALYSIS_VERSION
OUT = OUT_ROOT / "regression"
MODEL_DIR = OUT / "model_artifacts"

N_BOOTSTRAPS = 10_000
BOOTSTRAP_SEED = 20_260_826
PARITY_TOLERANCE = 1e-12
STUDY_ORDER = ("Holland2024", "Soraas2019")

LOCKED_INPUTS = {
    SPEC: "9dbc15ea2bb7f50dacbd94d7f06256f79e13b36ab3f4d0c165fc25eef054711e",
    P2_LOCK: "af86c613d9fa77622370ebb8888f71dd712f084844d0cc215d0be2305e53b133",
    FROZEN_READY: "a1c911b205dd6838f30bf5e8b5f6e9b834acb4d122995bf474a74a15160e9b2d",
}


def sha256_file(path: Path) -> str:
    digest = hashlib.sha256()
    with path.open("rb") as handle:
        for block in iter(lambda: handle.read(1024 * 1024), b""):
            digest.update(block)
    return digest.hexdigest()


def write_json(path: Path, value: object) -> None:
    path.write_text(
        json.dumps(value, indent=2, sort_keys=True, ensure_ascii=False) + "\n",
        encoding="utf-8",
    )


def verify_locked_inputs() -> pd.DataFrame:
    rows: list[dict[str, object]] = []
    for path, expected in LOCKED_INPUTS.items():
        if not path.is_file():
            raise FileNotFoundError(path)
        observed = sha256_file(path)
        if observed.lower() != expected.lower():
            raise RuntimeError(f"Locked input hash mismatch: {path}")
        rows.append(
            {
                "relative_path": str(path.relative_to(ROOT)),
                "bytes": path.stat().st_size,
                "sha256": observed,
            }
        )
    return pd.DataFrame(rows)


def load_ready() -> pd.DataFrame:
    frame = pd.read_csv(FROZEN_READY, sep="\t", dtype={"unit_id": str})
    required = ["analysis_row", "source_study", "unit_id", "H", "C", "S"]
    missing = [column for column in required if column not in frame.columns]
    if missing:
        raise RuntimeError(f"Frozen interface missing columns: {missing}")
    if frame[required].isna().any().any():
        raise RuntimeError("Frozen regression inputs contain missing values")
    if frame["analysis_row"].duplicated().any():
        raise RuntimeError("analysis_row is not unique")
    observed_counts = frame["source_study"].value_counts().to_dict()
    expected_counts = {"Holland2024": 61, "Soraas2019": 19}
    if observed_counts != expected_counts:
        raise RuntimeError(
            f"Frozen independent-unit counts changed: {observed_counts}"
        )
    if set(frame["source_study"]) != set(STUDY_ORDER):
        raise RuntimeError("Unexpected study in frozen interface")
    frame = frame.sort_values(
        ["source_study", "unit_id"], kind="mergesort"
    ).reset_index(drop=True)
    frame["A_R"] = frame["H"].astype(float)
    frame["A_L"] = frame["C"].astype(float)
    frame["score_S"] = frame["S"].astype(float)
    frame["x_A_L_minus_A_R"] = frame["A_L"] - frame["A_R"]
    frame["y_S_minus_A_R"] = frame["score_S"] - frame["A_R"]
    if not np.isfinite(
        frame[["A_R", "A_L", "score_S", "x_A_L_minus_A_R", "y_S_minus_A_R"]]
        .to_numpy(float)
    ).all():
        raise RuntimeError("Non-finite value in regression interface")
    return frame[
        [
            "analysis_row",
            "source_study",
            "unit_id",
            "A_R",
            "A_L",
            "score_S",
            "x_A_L_minus_A_R",
            "y_S_minus_A_R",
        ]
    ]


def fit_ols(x: np.ndarray, y: np.ndarray) -> dict[str, object]:
    design = np.column_stack([np.ones(len(x), dtype=float), x])
    xtx = design.T @ design
    if np.linalg.matrix_rank(xtx) != 2:
        raise RuntimeError("Regression design is rank deficient")
    xtx_inv = np.linalg.inv(xtx)
    coefficients = xtx_inv @ design.T @ y
    fitted = design @ coefficients
    residual = y - fitted
    leverage = np.einsum("ij,jk,ik->i", design, xtx_inv, design)
    if np.any(leverage >= 1):
        raise RuntimeError("HC3 leverage is not below one")
    scaled = residual**2 / (1.0 - leverage) ** 2
    meat = design.T @ (scaled[:, None] * design)
    hc3_covariance = xtx_inv @ meat @ xtx_inv
    hc3_se = np.sqrt(np.diag(hc3_covariance))
    degrees_freedom = len(x) - 2
    critical = float(stats.t.ppf(0.975, degrees_freedom))
    ci_low = coefficients - critical * hc3_se
    ci_high = coefficients + critical * hc3_se
    sse = float(residual @ residual)
    sst = float(np.sum((y - y.mean()) ** 2))
    if sst <= 0:
        raise RuntimeError("Regression outcome has zero variance")
    return {
        "coefficients": coefficients,
        "fitted": fitted,
        "residual": residual,
        "hc3_covariance": hc3_covariance,
        "hc3_se": hc3_se,
        "ci_low": ci_low,
        "ci_high": ci_high,
        "r2": 1.0 - sse / sst,
        "residual_mse": sse / len(x),
        "degrees_freedom": degrees_freedom,
        "critical_value": critical,
    }


def fit_fixed_beta(x: np.ndarray, y: np.ndarray, beta: float) -> dict[str, object]:
    alpha = float(np.mean(y - beta * x))
    fitted = alpha + beta * x
    residual = y - fitted
    return {
        "coefficients": np.asarray([alpha, beta], dtype=float),
        "fitted": fitted,
        "residual": residual,
        "residual_mse": float(np.mean(residual**2)),
    }


def bootstrap_coefficients(
    x: np.ndarray, y: np.ndarray, maps: np.ndarray
) -> np.ndarray:
    x_selected = x[maps]
    y_selected = y[maps]
    x_mean = x_selected.mean(axis=1)
    y_mean = y_selected.mean(axis=1)
    x_centered = x_selected - x_mean[:, None]
    denominator = np.sum(x_centered**2, axis=1)
    if np.any(denominator <= 0):
        raise RuntimeError("A bootstrap resample has zero predictor variance")
    beta = np.sum(x_centered * (y_selected - y_mean[:, None]), axis=1) / denominator
    alpha = y_mean - beta * x_mean
    return np.column_stack([alpha, beta])


def quantile(values: np.ndarray, probability: float) -> float:
    return float(np.quantile(values, probability, method="linear"))


def main() -> None:
    if OUT_ROOT.exists():
        raise FileExistsError(
            f"New-version output already exists; refusing overwrite: {OUT_ROOT}"
        )
    OUT.mkdir(parents=True, exist_ok=False)
    MODEL_DIR.mkdir(parents=True, exist_ok=False)

    input_manifest = verify_locked_inputs()
    ready = load_ready()
    rng = np.random.default_rng(BOOTSTRAP_SEED)

    summaries: list[dict[str, object]] = []
    anchors: list[dict[str, object]] = []
    bootstrap_frames: list[pd.DataFrame] = []
    model_rows: list[dict[str, object]] = []
    model_ids: list[str] = []
    model_studies: list[str] = []
    model_kinds: list[str] = []
    model_coefficients: list[np.ndarray] = []
    fitted_arrays: dict[str, np.ndarray] = {}
    row_arrays: dict[str, np.ndarray] = {}
    covariance_arrays: dict[str, np.ndarray] = {}
    bootstrap_arrays: dict[str, np.ndarray] = {}

    for study in STUDY_ORDER:
        study_frame = ready.loc[ready["source_study"] == study].copy()
        x = study_frame["x_A_L_minus_A_R"].to_numpy(float)
        y = study_frame["y_S_minus_A_R"].to_numpy(float)
        row_ids = study_frame["analysis_row"].to_numpy(np.int32)
        ols = fit_ols(x, y)

        maps = rng.integers(
            0, len(study_frame), size=(N_BOOTSTRAPS, len(study_frame)), dtype=np.int32
        )
        boot = bootstrap_coefficients(x, y, maps)
        bootstrap_arrays[f"maps_{study}"] = maps
        bootstrap_frames.append(
            pd.DataFrame(
                {
                    "source_study": study,
                    "bootstrap_id": np.arange(N_BOOTSTRAPS, dtype=int),
                    "alpha": boot[:, 0],
                    "beta": boot[:, 1],
                }
            )
        )

        coefficients = np.asarray(ols["coefficients"], dtype=float)
        hc3_se = np.asarray(ols["hc3_se"], dtype=float)
        ci_low = np.asarray(ols["ci_low"], dtype=float)
        ci_high = np.asarray(ols["ci_high"], dtype=float)
        summaries.append(
            {
                "source_study": study,
                "n_independent_units": len(study_frame),
                "alpha": coefficients[0],
                "beta": coefficients[1],
                "alpha_hc3_se": hc3_se[0],
                "beta_hc3_se": hc3_se[1],
                "alpha_hc3_ci_low": ci_low[0],
                "alpha_hc3_ci_high": ci_high[0],
                "beta_hc3_ci_low": ci_low[1],
                "beta_hc3_ci_high": ci_high[1],
                "hc3_interval_reference": f"Student-t(df={ols['degrees_freedom']})",
                "r2": float(ols["r2"]),
                "residual_mse": float(ols["residual_mse"]),
                "alpha_bootstrap_ci_low": quantile(boot[:, 0], 0.025),
                "alpha_bootstrap_ci_high": quantile(boot[:, 0], 0.975),
                "beta_bootstrap_ci_low": quantile(boot[:, 1], 0.025),
                "beta_bootstrap_ci_high": quantile(boot[:, 1], 0.975),
                "bootstrap_draws": N_BOOTSTRAPS,
                "bootstrap_seed": BOOTSTRAP_SEED,
                "x_min": float(x.min()),
                "x_max": float(x.max()),
            }
        )

        model_index = len(model_ids)
        model_id = f"{study}__unconstrained_ols"
        model_ids.append(model_id)
        model_studies.append(study)
        model_kinds.append("unconstrained_ols")
        model_coefficients.append(coefficients)
        fitted_arrays[f"pred_{model_index:02d}"] = np.asarray(ols["fitted"], dtype=float)
        row_arrays[f"rows_{model_index:02d}"] = row_ids
        covariance_arrays[f"hc3_cov_{model_index:02d}"] = np.asarray(
            ols["hc3_covariance"], dtype=float
        )
        model_rows.append(
            {
                "model_id": model_id,
                "source_study": study,
                "model_kind": "unconstrained_ols",
                "n_independent_units": len(study_frame),
                "alpha": coefficients[0],
                "beta": coefficients[1],
                "residual_mse": float(ols["residual_mse"]),
            }
        )

        for label, fixed_beta in (("host_anchor", 0.0), ("lineage_anchor", 1.0)):
            fixed = fit_fixed_beta(x, y, fixed_beta)
            fixed_coefficients = np.asarray(fixed["coefficients"], dtype=float)
            anchors.append(
                {
                    "source_study": study,
                    "anchor": label,
                    "n_independent_units": len(study_frame),
                    "fixed_beta": fixed_beta,
                    "fitted_alpha": fixed_coefficients[0],
                    "residual_mse": float(fixed["residual_mse"]),
                }
            )
            model_index = len(model_ids)
            model_id = f"{study}__{label}"
            model_ids.append(model_id)
            model_studies.append(study)
            model_kinds.append(label)
            model_coefficients.append(fixed_coefficients)
            fitted_arrays[f"pred_{model_index:02d}"] = np.asarray(
                fixed["fitted"], dtype=float
            )
            row_arrays[f"rows_{model_index:02d}"] = row_ids
            model_rows.append(
                {
                    "model_id": model_id,
                    "source_study": study,
                    "model_kind": label,
                    "n_independent_units": len(study_frame),
                    "alpha": fixed_coefficients[0],
                    "beta": fixed_coefficients[1],
                    "residual_mse": float(fixed["residual_mse"]),
                }
            )

    summary_frame = pd.DataFrame(summaries)
    anchor_frame = pd.DataFrame(anchors)
    model_frame = pd.DataFrame(model_rows)
    bootstrap_frame = pd.concat(bootstrap_frames, ignore_index=True)

    ready.to_csv(OUT / "regression_analysis_ready.tsv", sep="\t", index=False)
    input_manifest.to_csv(OUT / "input_manifest.tsv", sep="\t", index=False)
    summary_frame.to_csv(OUT / "regression_summary.tsv", sep="\t", index=False)
    anchor_frame.to_csv(OUT / "constrained_anchor_summary.tsv", sep="\t", index=False)
    model_frame.to_csv(MODEL_DIR / "closed_form_model_coefficients.tsv", sep="\t", index=False)
    bootstrap_frame.to_csv(
        OUT / "bootstrap_coefficients.tsv.gz",
        sep="\t",
        index=False,
        compression="gzip",
    )
    np.savez_compressed(OUT / "bootstrap_indices.npz", **bootstrap_arrays)
    np.savez_compressed(
        MODEL_DIR / "closed_form_model_state.npz",
        model_id=np.asarray(model_ids, dtype="U80"),
        source_study=np.asarray(model_studies, dtype="U30"),
        model_kind=np.asarray(model_kinds, dtype="U40"),
        coefficient=np.vstack(model_coefficients),
        feature_order=np.asarray(["intercept", "A_L_minus_A_R"], dtype="U40"),
    )
    np.savez_compressed(
        MODEL_DIR / "closed_form_fitted_values.npz", **fitted_arrays, **row_arrays
    )
    np.savez_compressed(MODEL_DIR / "hc3_covariance.npz", **covariance_arrays)

    signature = {
        "analysis_version": ANALYSIS_VERSION,
        "artifact_type": "closed_form_ols_and_fixed_slope_anchor_models",
        "entry_point": str(Path(__file__).resolve().relative_to(ROOT)),
        "entry_point_sha256": sha256_file(Path(__file__).resolve()),
        "input_hashes": {
            str(path.relative_to(ROOT)): expected for path, expected in LOCKED_INPUTS.items()
        },
        "feature_order": ["intercept", "A_L_minus_A_R"],
        "outcome": "score_S_minus_A_R",
        "grouping_units": {
            "Holland2024": "donor-recipient pair",
            "Soraas2019": "recipient",
        },
        "study_order": list(STUDY_ORDER),
        "preprocessing": "none",
        "selection": "none",
        "weighting": "none",
        "calibration": "none",
        "bootstrap": {
            "draws": N_BOOTSTRAPS,
            "seed": BOOTSTRAP_SEED,
            "unit": "frozen independent unit within study",
        },
        "hc3_interval": "95% Student-t with n-2 degrees of freedom",
        "parity_tolerance": PARITY_TOLERANCE,
        "n_models": len(model_ids),
    }
    write_json(MODEL_DIR / "model_signature.json", signature)

    max_parity_error = 0.0
    with (
        np.load(MODEL_DIR / "closed_form_model_state.npz", allow_pickle=False) as state,
        np.load(MODEL_DIR / "closed_form_fitted_values.npz", allow_pickle=False) as fitted,
    ):
        ready_indexed = ready.set_index("analysis_row")
        for index, model_id in enumerate(state["model_id"].tolist()):
            rows = fitted[f"rows_{index:02d}"].astype(int)
            x = ready_indexed.loc[rows, "x_A_L_minus_A_R"].to_numpy(float)
            coefficients = state["coefficient"][index].astype(float)
            reloaded = coefficients[0] + coefficients[1] * x
            saved = fitted[f"pred_{index:02d}"].astype(float)
            error = float(np.max(np.abs(reloaded - saved)))
            max_parity_error = max(max_parity_error, error)
            if error > PARITY_TOLERANCE:
                raise RuntimeError(f"Reload parity failed for {model_id}: {error}")

    artifact_audit = {
        "status": "PASS",
        "n_models": len(model_ids),
        "model_ids": model_ids,
        "reload_parity_tolerance": PARITY_TOLERANCE,
        "reload_max_abs_prediction_error": max_parity_error,
        "reload_parity": True,
    }
    write_json(MODEL_DIR / "artifact_audit.json", artifact_audit)

    public_data_gate = {
        "status": "RAW_IDAT_UNAVAILABLE",
        "public_interface": "six separately ENmix-processed beta matrices plus comRand.csv",
        "raw_idat_present_in_public_interface": False,
        "raw_idat_reprocessing_performed": False,
    }
    write_json(OUT / "public_data_gate.json", public_data_gate)

    report_lines = [
        "# HSCT referent continuous-regression sensitivity",
        "",
        "**Status:** PASS",
        "",
        "The regression was fitted separately to the frozen 61 Holland donor-recipient pairs and 19 Søraas recipients. It is a conventional sensitivity to the existing geometry-preserving referent analysis, not a calibration or causal model.",
        "",
    ]
    for row in summaries:
        report_lines.extend(
            [
                f"## {row['source_study']}",
                "",
                f"- beta = {row['beta']:.6f}; HC3 95% CI {row['beta_hc3_ci_low']:.6f} to {row['beta_hc3_ci_high']:.6f}; independent-unit bootstrap 95% interval {row['beta_bootstrap_ci_low']:.6f} to {row['beta_bootstrap_ci_high']:.6f}.",
                f"- alpha = {row['alpha']:.6f}; R2 = {row['r2']:.6f}.",
                "",
            ]
        )
    report_lines.extend(
        [
            "## Boundary",
            "",
            "Beta is a linear score-coordinate alignment coefficient. It is not a biological mixture fraction, does not identify lineage biological age and cannot exclude host influence. Holland is placement-driving; Søraas is complementary and is not treated as independent replication.",
            "",
            "The public Holland interface contains author-processed ENmix beta matrices, not raw IDAT files; raw-IDAT reprocessing was not performed.",
            "",
        ]
    )
    (OUT / "REPORT.md").write_text("\n".join(report_lines), encoding="utf-8")

    audit_paths = [
        OUT / "regression_analysis_ready.tsv",
        OUT / "input_manifest.tsv",
        OUT / "regression_summary.tsv",
        OUT / "constrained_anchor_summary.tsv",
        OUT / "bootstrap_coefficients.tsv.gz",
        OUT / "bootstrap_indices.npz",
        OUT / "public_data_gate.json",
        OUT / "REPORT.md",
        MODEL_DIR / "closed_form_model_coefficients.tsv",
        MODEL_DIR / "closed_form_model_state.npz",
        MODEL_DIR / "closed_form_fitted_values.npz",
        MODEL_DIR / "hc3_covariance.npz",
        MODEL_DIR / "model_signature.json",
        MODEL_DIR / "artifact_audit.json",
    ]
    run_audit = {
        "analysis_version": ANALYSIS_VERSION,
        "status": "PASS",
        "n_holland_pairs": 61,
        "n_soraas_recipients": 19,
        "model_artifact_gate": artifact_audit,
        "public_data_gate": public_data_gate,
        "environment": {
            "python": sys.version,
            "platform": platform.platform(),
            "numpy": np.__version__,
            "pandas": pd.__version__,
            "scipy": scipy.__version__,
        },
        "output_hashes": {
            str(path.relative_to(ROOT)): sha256_file(path) for path in audit_paths
        },
    }
    write_json(OUT / "run_audit.json", run_audit)

    artifact_rows = [
        {
            "relative_path": str(path.relative_to(OUT_ROOT)),
            "role": "model_artifact" if MODEL_DIR in path.parents else "analysis_artifact",
            "bytes": path.stat().st_size,
            "sha256": sha256_file(path),
        }
        for path in audit_paths + [OUT / "run_audit.json"]
    ]
    pd.DataFrame(artifact_rows).to_csv(
        OUT / "artifact_manifest.tsv", sep="\t", index=False
    )
    print(summary_frame.to_string(index=False), flush=True)
    print(anchor_frame.to_string(index=False), flush=True)
    print(json.dumps(artifact_audit, indent=2), flush=True)


if __name__ == "__main__":
    main()
```
