## Supplementary_Software_1 for "Inferential boundaries of age prediction: why prediction does not establish biological age measurement": analyze_nhanes3_5y_age_decoder_substitution_comparison_v0_1.py

```
#!/usr/bin/env python
"""Directly compare known age with a frozen age-decoder scalar in NHANES III.

This fail-closed implementation follows
``NHANES3_5Y_AGE_DECODER_SUBSTITUTION_COMPARISON_SPEC_v0_1_20260827.md``.
It compares two otherwise identical one-scalar mortality models in the frozen
NHANES III Phase 2 interface: ``W + s(A)`` and ``W + s(q_A)``.  The age-decoder
direction was learned in Phase 1 and is never refitted here.

All 330 downstream fits (primary plus 32 Fay-BRR weight sets, two models and
five outer folds) are saved as signed joblib artifacts, immediately reloaded,
and checked for elementwise design and prediction parity.  Compatible reruns
reuse the artifacts; incompatible artifacts fail closed.
"""

from __future__ import annotations

import argparse
import json
import math
import sys
from pathlib import Path
from typing import Any, Sequence

import joblib
import numpy as np
import pandas as pd
from scipy.stats import t as student_t
from sklearn.preprocessing import OneHotEncoder, SplineTransformer, StandardScaler

import analyze_nhanes3_5y_matched_scalar_target_supervision_protocol_transfer_v0_2 as base


ROOT = Path(__file__).resolve().parents[1]
SCRIPT = Path(__file__).resolve()
SPEC = (
    ROOT
    / "analysis"
    / "NHANES3_5Y_AGE_DECODER_SUBSTITUTION_COMPARISON_SPEC_v0_1_20260827.md"
)
PARENT = (
    ROOT
    / "outputs"
    / "nhanes3_5y_matched_scalar_target_supervision_protocol_transfer_v0_2"
)
DEFAULT_OUT = (
    ROOT / "outputs" / "nhanes3_5y_age_decoder_substitution_comparison_v0_1"
)

ANALYSIS_VERSION = "nhanes3_5y_age_decoder_substitution_comparison_v0_1"
SCHEMA_VERSION = "1.0"
STATUS = "retrospective_outcome_accessed_fixed_protocol_transfer_robustness_analysis"
MODEL_SEED = 20260827

AGE = base.AGE
OUTCOME = base.OUTCOME
BACKGROUND = base.BACKGROUND
WEIGHT = base.WEIGHT
BASE_WEIGHT = base.BASE_WEIGHT
STRATUM = base.STRATUM
PSU = base.PSU
N_OUTER = base.N_OUTER
N_INNER = base.N_INNER
C_GRID = base.C_GRID
EPSILON = base.EPSILON
PARITY_TOLERANCE = base.PARITY_TOLERANCE
TIE_TOLERANCE = base.TIE_TOLERANCE
FAY_REPLICATES = base.FAY_REPLICATES
FAY_RHO = base.FAY_RHO
FAY_DF = base.FAY_DF

MODELS = {
    "M_AGE": AGE,
    "M_QA": "q_A",
}

PARENT_FILES = {
    "parent_analysis_signature": PARENT / "analysis_signature.json",
    "parent_decision": PARENT / "decision.json",
    "parent_source_model_audit": PARENT / "model_artifact_audit_source.json",
    "parent_frozen_phase2_q_A": PARENT / "frozen_phase2_scalar_scores.tsv.gz",
    "parent_phase1_projection_bundle": (
        PARENT / "models" / "source_final" / "phase1_final_projection_bundle.joblib"
    ),
}

FREEZE_CANONICAL_KEYS = (
    "phase2",
    "phase2_folds",
    "fay_registry",
    "fay_factors",
    "fay_design",
)


def configure_stdout() -> None:
    if hasattr(sys.stdout, "reconfigure"):
        sys.stdout.reconfigure(encoding="utf-8")


def input_record(role: str, path: Path) -> dict[str, Any]:
    base.require(path.is_file(), f"missing frozen input: {path}")
    return {
        "role": role,
        "path": str(path.resolve()),
        "sha256": base.sha256(path),
        "size_bytes": int(path.stat().st_size),
    }


def load_analysis_inputs() -> tuple[pd.DataFrame, np.ndarray, pd.DataFrame, list[dict[str, Any]]]:
    # The predecessor loader supplies the already-audited Phase 2 interface,
    # fixed folds and Fay-BRR factors.  Phase 1 is loaded only by that validator
    # and is not used by this substitution analysis.
    _, phase2, _, _, factors, _ = base.load_inputs()

    for path in PARENT_FILES.values():
        base.require(path.is_file(), f"parent evidence file is missing: {path}")
    parent_signature = json.loads(
        PARENT_FILES["parent_analysis_signature"].read_text(encoding="utf-8")
    )
    parent_decision = json.loads(
        PARENT_FILES["parent_decision"].read_text(encoding="utf-8")
    )
    parent_audit = json.loads(
        PARENT_FILES["parent_source_model_audit"].read_text(encoding="utf-8")
    )
    base.require(
        parent_signature.get("signature_payload", {}).get("analysis_version")
        == "nhanes3_5y_matched_scalar_target_supervision_protocol_transfer_v0_2",
        "parent analysis identity changed",
    )
    base.require(
        parent_decision.get("source_projection_decision", {}).get("age_decoder_gate")
        == "PASS",
        "parent age-decoder gate is not PASS",
    )
    base.require(parent_audit.get("status") == "PASS", "parent source-model audit is not PASS")

    frozen = base.read_table(PARENT_FILES["parent_frozen_phase2_q_A"])
    base.require({"SEQN", "q_A"}.issubset(frozen.columns), "frozen q_A table is malformed")
    frozen = frozen[["SEQN", "q_A"]].sort_values("SEQN", kind="mergesort").reset_index(drop=True)
    base.require(not frozen["SEQN"].duplicated().any(), "frozen q_A identities are duplicated")
    base.require(
        frozen["SEQN"].astype(int).tolist() == phase2["SEQN"].astype(int).tolist(),
        "frozen q_A identities or order differ from Phase 2",
    )
    base.require(np.isfinite(frozen["q_A"]).all(), "frozen q_A contains non-finite values")
    frame = phase2.merge(frozen, on="SEQN", how="left", validate="one_to_one")
    frame = frame.sort_values("SEQN", kind="mergesort").reset_index(drop=True)
    base.require(frame["q_A"].notna().all(), "frozen q_A merge is incomplete")

    records: list[dict[str, Any]] = []
    for key in FREEZE_CANONICAL_KEYS:
        records.append(input_record(f"canonical_{key}", base.CANONICAL / base.CANONICAL_FILES[key]))
    for role, path in PARENT_FILES.items():
        records.append(input_record(role, path))
    for role, path in (
        ("frozen_protocol", SPEC),
        ("analysis_script", SCRIPT),
        ("implementation_dependency_base_script", Path(base.__file__).resolve()),
        ("environment_policy", base.ENV_POLICY),
        ("environment_manifest", base.ENV_MANIFEST),
        ("environment_lock", base.ENV_LOCK),
    ):
        records.append(input_record(role, path))
    return frame, factors, frozen, records


def build_analysis_signature(input_records: list[dict[str, Any]]) -> tuple[str, dict[str, Any]]:
    payload = {
        "analysis_version": ANALYSIS_VERSION,
        "schema_version": SCHEMA_VERSION,
        "status": STATUS,
        "inputs": input_records,
        "population": "NHANES_III_Phase_2_age_20_to_79",
        "endpoint": "resolved_five_year_all_cause_mortality",
        "background": list(BACKGROUND),
        "models": {
            "M_AGE": "Y ~ W + one_uniform_five_knot_cubic_spline(A)",
            "M_QA": "Y ~ W + one_uniform_five_knot_cubic_spline(q_A)",
        },
        "scalar_preprocessing": (
            "training_fold_local_weighted_standardization_then_uniform_5_knot_"
            "cubic_spline_without_bias_linear_extrapolation"
        ),
        "C_grid": list(C_GRID),
        "outer_folds": N_OUTER,
        "inner_folds": N_INNER,
        "inner_loss_pooling": "sum_weighted_numerators_over_sum_weight_denominators",
        "tie_rule": "within_1e-12_choose_smaller_C",
        "fit_weight_normalization": "sum_to_n_within_fit",
        "Fay_BRR": {"replicates": FAY_REPLICATES, "rho": FAY_RHO, "df": FAY_DF},
        "model_seed": MODEL_SEED,
        "parity_tolerance": PARITY_TOLERANCE,
        "software": base.software_record(),
    }
    return base.stable_hash(payload), payload


def freeze_inputs(out: Path, input_records: list[dict[str, Any]]) -> None:
    frozen = out / "input_freeze"
    for key in FREEZE_CANONICAL_KEYS:
        source = base.CANONICAL / base.CANONICAL_FILES[key]
        base.copy_file_or_verify(source, frozen / source.name)
    for key in (
        "parent_analysis_signature",
        "parent_decision",
        "parent_source_model_audit",
        "parent_frozen_phase2_q_A",
    ):
        source = PARENT_FILES[key]
        base.copy_file_or_verify(source, frozen / source.name)
    base.write_table_or_verify(pd.DataFrame(input_records), out / "input_manifest.tsv")


def fit_scalar_preprocessor(
    frame: pd.DataFrame,
    scalar_column: str,
    weight: np.ndarray,
) -> tuple[dict[str, Any], np.ndarray]:
    fit_weight = base.normalized_weights(weight)
    encoder = OneHotEncoder(handle_unknown="ignore", sparse_output=False)
    categorical = encoder.fit_transform(frame.loc[:, list(BACKGROUND)].astype(str))

    scalar_raw = frame[[scalar_column]].to_numpy(float)
    base.require(np.isfinite(scalar_raw).all(), f"{scalar_column} contains non-finite values")
    scalar_scaler = StandardScaler().fit(scalar_raw, sample_weight=fit_weight)
    scalar_standardized = scalar_scaler.transform(scalar_raw)
    scalar_spline = SplineTransformer(
        n_knots=5,
        knots="uniform",
        degree=3,
        include_bias=False,
        extrapolation="linear",
    )
    scalar_design = scalar_spline.fit_transform(
        scalar_standardized, sample_weight=fit_weight
    )
    design = np.column_stack([categorical, scalar_design])
    base.require(np.isfinite(design).all(), "substitution design contains non-finite values")
    return {
        "background": list(BACKGROUND),
        "scalar_column": scalar_column,
        "encoder": encoder,
        "scalar_scaler": scalar_scaler,
        "scalar_spline": scalar_spline,
        "spline_contract": {
            "n_knots": 5,
            "knots": "uniform",
            "degree": 3,
            "include_bias": False,
            "extrapolation": "linear",
        },
        "design_feature_names": (
            [f"W__{name}" for name in encoder.get_feature_names_out(BACKGROUND)]
            + [f"scalar_spline_{index:02d}" for index in range(scalar_design.shape[1])]
        ),
    }, design


def transform_scalar_preprocessor(preprocessor: dict[str, Any], frame: pd.DataFrame) -> np.ndarray:
    categorical = preprocessor["encoder"].transform(
        frame.loc[:, list(preprocessor["background"])].astype(str)
    )
    scalar_raw = frame[[str(preprocessor["scalar_column"])]].to_numpy(float)
    scalar_standardized = preprocessor["scalar_scaler"].transform(scalar_raw)
    scalar_design = preprocessor["scalar_spline"].transform(scalar_standardized)
    design = np.column_stack([categorical, scalar_design])
    base.require(np.isfinite(design).all(), "reconstructed substitution design is non-finite")
    return design


def artifact_signature(
    stage_signature: str,
    model_name: str,
    outer_fold: int,
    training: pd.DataFrame,
    test: pd.DataFrame,
) -> str:
    scalar_column = MODELS[model_name]
    inner_column = f"inner_fold_outer_{outer_fold}"
    payload = {
        "stage_signature": stage_signature,
        "artifact_kind": "matched_single_scalar_target_local_mortality_model",
        "model_name": model_name,
        "scalar_column": scalar_column,
        "outer_fold": int(outer_fold),
        "training_seqn": training["SEQN"].astype(int).tolist(),
        "test_seqn": test["SEQN"].astype(int).tolist(),
        "training_weight": training[WEIGHT].astype(float).tolist(),
        "test_weight": test[WEIGHT].astype(float).tolist(),
        "training_scalar": training[scalar_column].astype(float).tolist(),
        "test_scalar": test[scalar_column].astype(float).tolist(),
        "inner_assignments": training[["SEQN", inner_column]].astype(
            {"SEQN": int, inner_column: int}
        ).to_dict(orient="records"),
        "background": list(BACKGROUND),
        "scalar_preprocessing": (
            "training_fold_local_weighted_standardization_then_uniform_5_knot_"
            "cubic_spline_without_bias_linear_extrapolation"
        ),
        "C_grid": list(C_GRID),
        "inner_loss_pooling": "sum_weighted_numerators_over_sum_weight_denominators",
        "tie_rule": "within_1e-12_choose_smaller_C",
        "fit_weight_normalization": "sum_to_n_within_fit",
        "log_loss_clip": EPSILON,
        "model_seed": MODEL_SEED,
        "parity_tolerance": PARITY_TOLERANCE,
    }
    return base.stable_hash(payload)


def fit_artifact(
    analysis_signature: str,
    stage_signature: str,
    model_name: str,
    outer_fold: int,
    frame: pd.DataFrame,
) -> dict[str, Any]:
    scalar_column = MODELS[model_name]
    training = frame.loc[frame["outer_fold"].ne(outer_fold)].sort_values(
        "SEQN", kind="mergesort"
    ).reset_index(drop=True)
    test = frame.loc[frame["outer_fold"].eq(outer_fold)].sort_values(
        "SEQN", kind="mergesort"
    ).reset_index(drop=True)
    inner_column = f"inner_fold_outer_{outer_fold}"
    inner_folds: list[dict[str, Any]] = []
    tuning_rows: list[dict[str, Any]] = []

    for inner_fold in range(N_INNER):
        inner_train = training.loc[training[inner_column].ne(inner_fold)].sort_values(
            "SEQN", kind="mergesort"
        ).reset_index(drop=True)
        inner_valid = training.loc[training[inner_column].eq(inner_fold)].sort_values(
            "SEQN", kind="mergesort"
        ).reset_index(drop=True)
        base.require(len(inner_train) > 0 and len(inner_valid) > 0, "empty inner fold")
        train_weight = inner_train[WEIGHT].to_numpy(float)
        valid_weight = inner_valid[WEIGHT].to_numpy(float)
        preprocessor, x_train = fit_scalar_preprocessor(
            inner_train, scalar_column, train_weight
        )
        x_valid = transform_scalar_preprocessor(preprocessor, inner_valid)
        y_train = inner_train[OUTCOME].to_numpy(int)
        y_valid = inner_valid[OUTCOME].to_numpy(int)
        candidate_fits: list[dict[str, Any]] = []
        for c_value in C_GRID:
            estimator = base.fit_logistic(x_train, y_train, train_weight, c_value)
            probability = base.predict_logistic(estimator, x_valid)
            losses = base.log_loss_vector(y_valid, probability)
            numerator = float(np.dot(valid_weight, losses))
            denominator = float(valid_weight.sum())
            candidate_fits.append(
                {
                    "C": float(c_value),
                    "estimator": estimator,
                    "valid_probability": probability,
                    "valid_loss_vector": losses,
                    "weighted_loss_numerator": numerator,
                    "weight_sum": denominator,
                }
            )
            tuning_rows.append(
                {
                    "model": model_name,
                    "outer_fold": int(outer_fold),
                    "inner_fold": int(inner_fold),
                    "C": float(c_value),
                    "n_valid": int(len(inner_valid)),
                    "events_valid": int(y_valid.sum()),
                    "weighted_loss_numerator": numerator,
                    "weight_sum": denominator,
                    "weighted_loss": numerator / denominator,
                }
            )
        inner_folds.append(
            {
                "inner_fold": int(inner_fold),
                "train_seqn": inner_train["SEQN"].to_numpy("int64"),
                "valid_seqn": inner_valid["SEQN"].to_numpy("int64"),
                "train_weight": train_weight,
                "valid_weight": valid_weight,
                "train_scalar": inner_train[scalar_column].to_numpy(float),
                "valid_scalar": inner_valid[scalar_column].to_numpy(float),
                "preprocessor": preprocessor,
                "x_train": x_train,
                "x_valid": x_valid,
                "train_outcome": y_train,
                "valid_outcome": y_valid,
                "candidate_fits": candidate_fits,
            }
        )

    selected_c = base.select_grid_value(tuning_rows, "C", C_GRID, prefer="small")
    final_weight = training[WEIGHT].to_numpy(float)
    final_preprocessor, x_training = fit_scalar_preprocessor(
        training, scalar_column, final_weight
    )
    x_test = transform_scalar_preprocessor(final_preprocessor, test)
    final_estimator = base.fit_logistic(
        x_training, training[OUTCOME].to_numpy(int), final_weight, selected_c
    )
    test_probability = base.predict_logistic(final_estimator, x_test)
    test_loss = base.log_loss_vector(test[OUTCOME].to_numpy(int), test_probability)
    return {
        "schema_version": SCHEMA_VERSION,
        "analysis_version": ANALYSIS_VERSION,
        "analysis_signature": analysis_signature,
        "stage_signature": stage_signature,
        "artifact_signature": artifact_signature(
            stage_signature, model_name, outer_fold, training, test
        ),
        "artifact_kind": "matched_single_scalar_target_local_mortality_model",
        "model_name": model_name,
        "scalar_column": scalar_column,
        "outer_fold": int(outer_fold),
        "inner_column": inner_column,
        "training_seqn": training["SEQN"].to_numpy("int64"),
        "test_seqn": test["SEQN"].to_numpy("int64"),
        "training_weight": final_weight,
        "test_weight": test[WEIGHT].to_numpy(float),
        "training_scalar": training[scalar_column].to_numpy(float),
        "test_scalar": test[scalar_column].to_numpy(float),
        "inner_folds": inner_folds,
        "tuning_rows": tuning_rows,
        "selected_C": float(selected_c),
        "final_preprocessor": final_preprocessor,
        "x_training": x_training,
        "x_test": x_test,
        "final_estimator": final_estimator,
        "test_probability": test_probability,
        "test_loss_vector": test_loss,
        "software": base.software_record(),
    }


def verify_artifact(
    artifact: dict[str, Any],
    analysis_signature: str,
    stage_signature: str,
    model_name: str,
    outer_fold: int,
    frame: pd.DataFrame,
) -> dict[str, float]:
    scalar_column = MODELS[model_name]
    training = frame.loc[frame["outer_fold"].ne(outer_fold)].sort_values(
        "SEQN", kind="mergesort"
    ).reset_index(drop=True)
    test = frame.loc[frame["outer_fold"].eq(outer_fold)].sort_values(
        "SEQN", kind="mergesort"
    ).reset_index(drop=True)
    expected_signature = artifact_signature(
        stage_signature, model_name, outer_fold, training, test
    )
    base.require(artifact.get("analysis_signature") == analysis_signature, "analysis signature changed")
    base.require(artifact.get("artifact_signature") == expected_signature, "artifact signature mismatch")
    base.require(artifact.get("model_name") == model_name, "artifact model identity changed")
    base.require(artifact.get("scalar_column") == scalar_column, "artifact scalar identity changed")
    base.require(
        np.array_equal(artifact["training_seqn"], training["SEQN"].to_numpy("int64")),
        "artifact training identities changed",
    )
    base.require(
        np.array_equal(artifact["test_seqn"], test["SEQN"].to_numpy("int64")),
        "artifact test identities changed",
    )
    base.require(
        base.maximum_error(artifact["training_weight"], training[WEIGHT])
        <= PARITY_TOLERANCE,
        "artifact training weights changed",
    )
    base.require(
        base.maximum_error(artifact["test_weight"], test[WEIGHT]) <= PARITY_TOLERANCE,
        "artifact test weights changed",
    )
    base.require(
        base.maximum_error(artifact["training_scalar"], training[scalar_column])
        <= PARITY_TOLERANCE,
        "artifact training scalar changed",
    )
    base.require(
        base.maximum_error(artifact["test_scalar"], test[scalar_column])
        <= PARITY_TOLERANCE,
        "artifact test scalar changed",
    )

    by_seqn = frame.set_index("SEQN", drop=False)
    reconstructed_tuning: list[dict[str, Any]] = []
    max_design_error = 0.0
    max_prediction_error = 0.0
    for stored in artifact["inner_folds"]:
        inner_fold = int(stored["inner_fold"])
        train_seqn = np.asarray(stored["train_seqn"], dtype="int64")
        valid_seqn = np.asarray(stored["valid_seqn"], dtype="int64")
        expected_train = training.loc[
            training[artifact["inner_column"]].ne(inner_fold), "SEQN"
        ].to_numpy("int64")
        expected_valid = training.loc[
            training[artifact["inner_column"]].eq(inner_fold), "SEQN"
        ].to_numpy("int64")
        base.require(np.array_equal(train_seqn, expected_train), "inner training identities changed")
        base.require(np.array_equal(valid_seqn, expected_valid), "inner validation identities changed")
        base.require(np.intersect1d(train_seqn, valid_seqn).size == 0, "inner-fold identity leakage")
        inner_train = by_seqn.loc[train_seqn].reset_index(drop=True)
        inner_valid = by_seqn.loc[valid_seqn].reset_index(drop=True)
        base.require(
            base.maximum_error(stored["train_weight"], inner_train[WEIGHT])
            <= PARITY_TOLERANCE,
            "inner training weights changed",
        )
        base.require(
            base.maximum_error(stored["valid_weight"], inner_valid[WEIGHT])
            <= PARITY_TOLERANCE,
            "inner validation weights changed",
        )
        base.require(
            base.maximum_error(stored["train_scalar"], inner_train[scalar_column])
            <= PARITY_TOLERANCE,
            "inner training scalar changed",
        )
        base.require(
            base.maximum_error(stored["valid_scalar"], inner_valid[scalar_column])
            <= PARITY_TOLERANCE,
            "inner validation scalar changed",
        )
        x_train = transform_scalar_preprocessor(stored["preprocessor"], inner_train)
        x_valid = transform_scalar_preprocessor(stored["preprocessor"], inner_valid)
        max_design_error = max(
            max_design_error,
            base.maximum_error(x_train, stored["x_train"]),
            base.maximum_error(x_valid, stored["x_valid"]),
        )
        y_valid = inner_valid[OUTCOME].to_numpy(int)
        valid_weight = inner_valid[WEIGHT].to_numpy(float)
        for candidate in stored["candidate_fits"]:
            probability = base.predict_logistic(candidate["estimator"], x_valid)
            losses = base.log_loss_vector(y_valid, probability)
            max_prediction_error = max(
                max_prediction_error,
                base.maximum_error(probability, candidate["valid_probability"]),
                base.maximum_error(losses, candidate["valid_loss_vector"]),
            )
            numerator = float(np.dot(valid_weight, losses))
            base.require(
                abs(numerator - float(candidate["weighted_loss_numerator"]))
                <= PARITY_TOLERANCE,
                "inner tuning loss changed",
            )
            reconstructed_tuning.append(
                {
                    "C": float(candidate["C"]),
                    "weighted_loss_numerator": numerator,
                    "weight_sum": float(valid_weight.sum()),
                }
            )
    selected_c = base.select_grid_value(
        reconstructed_tuning, "C", C_GRID, prefer="small"
    )
    base.require(selected_c == float(artifact["selected_C"]), "selected C changed")

    x_training = transform_scalar_preprocessor(artifact["final_preprocessor"], training)
    x_test = transform_scalar_preprocessor(artifact["final_preprocessor"], test)
    probability = base.predict_logistic(artifact["final_estimator"], x_test)
    losses = base.log_loss_vector(test[OUTCOME].to_numpy(int), probability)
    max_design_error = max(
        max_design_error,
        base.maximum_error(x_training, artifact["x_training"]),
        base.maximum_error(x_test, artifact["x_test"]),
    )
    max_prediction_error = max(
        max_prediction_error,
        base.maximum_error(probability, artifact["test_probability"]),
        base.maximum_error(losses, artifact["test_loss_vector"]),
    )
    base.require(
        max(max_design_error, max_prediction_error) <= PARITY_TOLERANCE,
        "reloaded model parity failed",
    )
    return {
        "max_design_error": max_design_error,
        "max_prediction_error": max_prediction_error,
    }


def fit_or_load_artifact(
    path: Path,
    analysis_signature: str,
    stage_signature: str,
    model_name: str,
    outer_fold: int,
    frame: pd.DataFrame,
) -> tuple[dict[str, Any], dict[str, Any]]:
    fitted = False
    if path.exists():
        artifact = joblib.load(path)
        training = frame.loc[frame["outer_fold"].ne(outer_fold)].sort_values(
            "SEQN", kind="mergesort"
        ).reset_index(drop=True)
        test = frame.loc[frame["outer_fold"].eq(outer_fold)].sort_values(
            "SEQN", kind="mergesort"
        ).reset_index(drop=True)
        expected = artifact_signature(
            stage_signature, model_name, outer_fold, training, test
        )
        base.require(
            artifact.get("analysis_signature") == analysis_signature,
            f"incompatible cached parent signature: {path}",
        )
        base.require(
            artifact.get("artifact_signature") == expected,
            f"incompatible cached artifact: {path}",
        )
    else:
        artifact = fit_artifact(
            analysis_signature, stage_signature, model_name, outer_fold, frame
        )
        base.save_joblib_atomic(artifact, path)
        fitted = True
    reloaded = joblib.load(path)
    parity = verify_artifact(
        reloaded, analysis_signature, stage_signature, model_name, outer_fold, frame
    )
    return reloaded, {
        "stage": "target_substitution_downstream",
        "target_refit_label": "",
        "model": model_name,
        "outer_fold": int(outer_fold),
        "artifact_path": str(path.resolve()),
        "artifact_sha256": base.sha256(path),
        "cache_action": "fitted" if fitted else "reused",
        "max_design_error": parity["max_design_error"],
        "max_prediction_error": parity["max_prediction_error"],
        "downstream_tuning_estimator_count": int(
            sum(len(fold["candidate_fits"]) for fold in reloaded["inner_folds"])
        ),
        "downstream_final_estimator_count": 1,
    }


def run_weight_set(
    out: Path,
    base_frame: pd.DataFrame,
    analysis_signature: str,
    label: str,
    weight_factor: np.ndarray,
) -> tuple[pd.DataFrame, pd.DataFrame, list[dict[str, Any]]]:
    base.require(len(weight_factor) == len(base_frame), "replicate weight length changed")
    frame = base_frame.copy()
    frame[WEIGHT] = frame[BASE_WEIGHT].to_numpy(float) * np.asarray(weight_factor, dtype=float)
    base.require(
        np.isfinite(frame[WEIGHT]).all() and frame[WEIGHT].gt(0).all(),
        "invalid target weights",
    )
    stage_signature = base.stable_hash(
        {
            "analysis_signature": analysis_signature,
            "stage": "phase2_matched_single_scalar_substitution",
            "label": label,
            "ordered_seqn": frame["SEQN"].astype(int).tolist(),
            "ordered_weight": frame[WEIGHT].astype(float).tolist(),
            "ordered_age": frame[AGE].astype(float).tolist(),
            "frozen_q_A": frame["q_A"].astype(float).tolist(),
            "models": list(MODELS),
            "q_A_refitted": False,
        }
    )
    prediction = frame[
        ["SEQN", "outer_fold", OUTCOME, BASE_WEIGHT, WEIGHT, STRATUM, PSU, AGE, "q_A"]
    ].copy()
    tuning_tables: list[pd.DataFrame] = []
    records: list[dict[str, Any]] = []
    model_root = (
        out / "models" / "primary"
        if label == "primary"
        else out / "models" / "fay_brr" / label
    )
    for model_name in MODELS:
        probability = np.full(len(frame), np.nan, dtype=float)
        for outer_fold in range(N_OUTER):
            artifact, record = fit_or_load_artifact(
                model_root / model_name / f"outer_fold_{outer_fold:02d}.joblib",
                analysis_signature,
                stage_signature,
                model_name,
                outer_fold,
                frame,
            )
            record["target_refit_label"] = label
            records.append(record)
            test_ids = np.asarray(artifact["test_seqn"], dtype="int64")
            positions = (
                frame.reset_index().set_index("SEQN").loc[test_ids, "index"].to_numpy(int)
            )
            probability[positions] = np.asarray(artifact["test_probability"], dtype=float)
            tuning = pd.DataFrame(artifact["tuning_rows"])
            tuning.insert(0, "target_refit_label", label)
            tuning_tables.append(tuning)
        base.require(np.isfinite(probability).all(), f"incomplete OOF predictions for {model_name}")
        prediction[f"probability_{model_name}"] = probability
        prediction[f"loss_{model_name}"] = base.log_loss_vector(
            frame[OUTCOME].to_numpy(int), probability
        )
    prediction["paired_delta_sub_loss"] = (
        prediction["loss_M_AGE"] - prediction["loss_M_QA"]
    )
    return prediction, pd.concat(tuning_tables, ignore_index=True), records


def summarize(prediction: pd.DataFrame) -> tuple[dict[str, float], pd.DataFrame]:
    estimates = {
        "loss_M_AGE": base.weighted_mean(prediction["loss_M_AGE"], prediction[WEIGHT]),
        "loss_M_QA": base.weighted_mean(prediction["loss_M_QA"], prediction[WEIGHT]),
        "Delta_sub": base.weighted_mean(
            prediction["paired_delta_sub_loss"], prediction[WEIGHT]
        ),
    }
    rows: list[dict[str, Any]] = []
    for outer_fold in range(N_OUTER):
        block = prediction.loc[prediction["outer_fold"].eq(outer_fold)]
        delta = base.weighted_mean(block["paired_delta_sub_loss"], block[WEIGHT])
        rows.append(
            {
                "outer_fold": int(outer_fold),
                "n": int(len(block)),
                "deaths_5y": int(block[OUTCOME].sum()),
                "Delta_sub": delta,
                "Delta_sub_positive": bool(delta > 0),
                "Delta_sub_negative": bool(delta < 0),
            }
        )
    return estimates, pd.DataFrame(rows)


def fay_interval(full_estimate: float, replicates: Sequence[float]) -> dict[str, Any]:
    values = np.asarray(replicates, dtype=float)
    base.require(
        values.shape == (FAY_REPLICATES,) and np.isfinite(values).all(),
        "Fay-BRR contrast estimates are incomplete",
    )
    variance = float(
        np.sum((values - float(full_estimate)) ** 2)
        / (FAY_REPLICATES * (1.0 - FAY_RHO) ** 2)
    )
    standard_error = float(math.sqrt(variance))
    critical = float(student_t.ppf(0.975, FAY_DF))
    return {
        "estimate": float(full_estimate),
        "variance": variance,
        "standard_error": standard_error,
        "critical_t": critical,
        "df": FAY_DF,
        "ci_lower": float(full_estimate - critical * standard_error),
        "ci_upper": float(full_estimate + critical * standard_error),
        "replicates": FAY_REPLICATES,
        "rho": FAY_RHO,
        "conditional_on": "the_frozen_Phase1_age_decoder_direction",
    }


def deterministic_model_index(records: list[dict[str, Any]]) -> pd.DataFrame:
    rows = [{key: value for key, value in record.items() if key != "cache_action"} for record in records]
    table = pd.DataFrame(rows).sort_values(
        ["target_refit_label", "model", "outer_fold"], kind="mergesort"
    ).reset_index(drop=True)
    base.require(not table["artifact_path"].duplicated().any(), "duplicate model artifact path")
    return table


def audit_models(records: list[dict[str, Any]]) -> dict[str, Any]:
    expected = (FAY_REPLICATES + 1) * len(MODELS) * N_OUTER
    base.require(len(records) == expected, f"expected {expected} artifacts, found {len(records)}")
    max_design = max(float(record["max_design_error"]) for record in records)
    max_prediction = max(float(record["max_prediction_error"]) for record in records)
    base.require(
        max(max_design, max_prediction) <= PARITY_TOLERANCE,
        "model artifact parity audit failed",
    )
    for record in records:
        path = Path(record["artifact_path"])
        base.require(path.is_file(), f"indexed model artifact is missing: {path}")
        base.require(
            base.sha256(path) == record["artifact_sha256"],
            f"indexed model artifact hash changed: {path}",
        )
    return {
        "status": "PASS",
        "analysis_version": ANALYSIS_VERSION,
        "artifact_count": int(len(records)),
        "expected_artifact_count": int(expected),
        "downstream_tuning_estimator_count": int(
            sum(record["downstream_tuning_estimator_count"] for record in records)
        ),
        "downstream_final_estimator_count": int(
            sum(record["downstream_final_estimator_count"] for record in records)
        ),
        "maximum_design_reconstruction_error": max_design,
        "maximum_prediction_reconstruction_error": max_prediction,
        "parity_tolerance": PARITY_TOLERANCE,
        "cache_policy": "compatible_artifacts_reused; incompatible_artifacts_fail_closed",
    }


def write_output_manifest(out: Path) -> None:
    rows: list[dict[str, Any]] = []
    manifest_path = out / "artifact_manifest.tsv"
    for path in sorted(out.rglob("*"), key=lambda item: str(item).lower()):
        if not path.is_file() or path == manifest_path:
            continue
        rows.append(
            {
                "relative_path": str(path.relative_to(out)).replace("\\", "/"),
                "size_bytes": int(path.stat().st_size),
                "sha256": base.sha256(path),
            }
        )
    base.write_table_or_verify(pd.DataFrame(rows), manifest_path)


def run(args: argparse.Namespace) -> int:
    configure_stdout()
    out = Path(args.output_dir).resolve()
    base.require(out.name == ANALYSIS_VERSION, "output basename must retain the frozen version")
    base.require(out != PARENT.resolve(), "new analysis may not write into its frozen parent")
    out.mkdir(parents=True, exist_ok=True)

    frame, factors, _, input_records = load_analysis_inputs()
    analysis_signature, signature_payload = build_analysis_signature(input_records)
    freeze_inputs(out, input_records)
    base.write_json_or_verify(
        out / "analysis_signature.json",
        {"analysis_signature": analysis_signature, "signature_payload": signature_payload},
    )

    all_records: list[dict[str, Any]] = []
    primary_prediction, primary_tuning, primary_records = run_weight_set(
        out,
        frame,
        analysis_signature,
        "primary",
        np.ones(len(frame), dtype=float),
    )
    all_records.extend(primary_records)
    primary_estimates, fold_contrasts = summarize(primary_prediction)
    base.write_table_or_verify(
        primary_prediction, out / "target_primary_oof_predictions_and_losses.tsv.gz"
    )
    base.write_table_or_verify(
        primary_tuning, out / "target_primary_inner_tuning.tsv.gz"
    )
    base.write_table_or_verify(
        fold_contrasts, out / "target_primary_fold_contrasts.tsv"
    )
    if args.stage == "primary":
        print(
            json.dumps(
                {
                    "analysis_version": ANALYSIS_VERSION,
                    "stage": "primary_probe",
                    "primary_estimates": primary_estimates,
                    "positive_outer_folds": int(
                        fold_contrasts["Delta_sub_positive"].sum()
                    ),
                    "negative_outer_folds": int(
                        fold_contrasts["Delta_sub_negative"].sum()
                    ),
                },
                indent=2,
                ensure_ascii=False,
            )
        )
        return 0

    replicate_rows: list[dict[str, Any]] = []
    replicate_tuning: list[pd.DataFrame] = []
    for replicate in range(FAY_REPLICATES):
        label = f"replicate_{replicate:02d}"
        prediction, tuning, records = run_weight_set(
            out, frame, analysis_signature, label, factors[replicate]
        )
        all_records.extend(records)
        estimates, folds = summarize(prediction)
        base.write_table_or_verify(
            prediction,
            out / "fay_brr_predictions" / f"{label}_oof_predictions_and_losses.tsv.gz",
        )
        replicate_tuning.append(tuning)
        replicate_rows.append(
            {
                "replicate": int(replicate),
                "loss_M_AGE": estimates["loss_M_AGE"],
                "loss_M_QA": estimates["loss_M_QA"],
                "Delta_sub": estimates["Delta_sub"],
                "Delta_sub_positive_outer_folds": int(folds["Delta_sub_positive"].sum()),
                "Delta_sub_negative_outer_folds": int(folds["Delta_sub_negative"].sum()),
            }
        )
    replicate_table = pd.DataFrame(replicate_rows)
    base.write_table_or_verify(
        replicate_table, out / "target_fay_brr_replicate_estimates.tsv"
    )
    base.write_table_or_verify(
        pd.concat(replicate_tuning, ignore_index=True),
        out / "target_fay_brr_inner_tuning.tsv.gz",
    )

    interval = fay_interval(primary_estimates["Delta_sub"], replicate_table["Delta_sub"])
    positive_folds = int(fold_contrasts["Delta_sub_positive"].sum())
    negative_folds = int(fold_contrasts["Delta_sub_negative"].sum())
    if (
        primary_estimates["Delta_sub"] > 0
        and interval["ci_lower"] > 0
        and positive_folds >= 4
    ):
        decision = "QA_SUBSTITUTION_ADVANTAGE"
    elif (
        primary_estimates["Delta_sub"] < 0
        and interval["ci_upper"] < 0
        and negative_folds >= 4
    ):
        decision = "AGE_SUBSTITUTION_ADVANTAGE"
    else:
        decision = "NO_CLEAR_SUBSTITUTION_ADVANTAGE"

    model_index = deterministic_model_index(all_records)
    model_audit = audit_models(all_records)
    base.write_table_or_verify(model_index, out / "model_artifact_index.tsv")
    base.write_json_or_verify(out / "model_artifact_audit.json", model_audit)
    result = {
        "analysis_version": ANALYSIS_VERSION,
        "status": STATUS,
        "decision": decision,
        "target_population_n": int(len(frame)),
        "target_deaths_5y": int(frame[OUTCOME].sum()),
        "models": {
            "M_AGE": "Y ~ W + training_local_uniform_five_knot_cubic_spline(A)",
            "M_QA": "Y ~ W + training_local_uniform_five_knot_cubic_spline(q_A)",
        },
        "primary_contrast": "Delta_sub = L(M_AGE) - L(M_QA); positive favours q_A",
        "primary_estimates": primary_estimates,
        "Delta_sub_Fay_BRR": interval,
        "Delta_sub_positive_outer_folds": positive_folds,
        "Delta_sub_negative_outer_folds": negative_folds,
        "Fay_BRR_full_refits_both_arms": True,
        "q_A_direction_refitted": False,
        "model_artifact_audit_status": model_audit["status"],
        "claim_ceiling": (
            "This comparison is limited to the frozen NHANES III Phase 2 population, "
            "five-year endpoint, transported q_A and matched one-scalar model family. "
            "It does not identify biological age, establish general superiority over "
            "chronological age, or imply causal or surrogate validity."
        ),
    }
    base.write_json_or_verify(out / "decision.json", result)
    report = f"""# NHANES III age-decoder substitution comparison v0.1

- Status: **{STATUS}**
- Decision: **{decision}**
- Phase 2 participants/deaths: {len(frame)} / {int(frame[OUTCOME].sum())}
- `M_AGE = W + s(A)` design-weighted OOF log loss: {primary_estimates['loss_M_AGE']:.9f}
- `M_QA = W + s(q_A)` design-weighted OOF log loss: {primary_estimates['loss_M_QA']:.9f}
- `Delta_sub = L(M_AGE)-L(M_QA)`: {primary_estimates['Delta_sub']:.9f}
- 95% Fay-BRR interval: {interval['ci_lower']:.9f} to {interval['ci_upper']:.9f}
- Positive fixed outer-fold contrasts: {positive_folds} of {N_OUTER}

Both arms used the same background variables, one continuous scalar, weighted
training-fold-local standardization, a uniform five-knot cubic spline, the same
regularization grid, folds and weights. Every Fay-BRR replicate repeated tuning,
fitting and held-out prediction for both arms. The Phase 1 age-decoder direction
was frozen throughout.
"""
    base.write_text_or_verify(out / "REPORT.md", report)
    write_output_manifest(out)
    print(json.dumps(result, indent=2, ensure_ascii=False))
    return 0


def build_parser() -> argparse.ArgumentParser:
    parser = argparse.ArgumentParser(description=__doc__)
    parser.add_argument(
        "--output-dir",
        default=str(DEFAULT_OUT),
        help="Versioned output directory; basename must equal the frozen analysis version.",
    )
    parser.add_argument(
        "--stage",
        choices=("primary", "all"),
        default="all",
        help="Run the minimal primary probe first, or complete all Fay-BRR refits.",
    )
    return parser


def main(argv: Sequence[str] | None = None) -> int:
    return run(build_parser().parse_args(argv))


if __name__ == "__main__":
    raise SystemExit(main())
```
