## Supplementary_Software_1 for "Inferential boundaries of age prediction: why prediction does not establish biological age measurement": analyze_nhanes3_5y_chronological_age_increment_and_proxy_comparison_v0_1.py

```
#!/usr/bin/env python
"""NHANES III chronological-age increment and proxy comparison.

This fail-closed implementation follows
``NHANES3_5Y_CHRONOLOGICAL_AGE_INCREMENT_AND_PROXY_COMPARISON_SPEC_v0_1_20260827.md``.
It fits four direct Phase 2 mortality models: the frozen full 25-measurement
panel with and without observed chronological age, and the ordered remaining-20
panel after removal of the frozen Phase 1 age-selected cardiorespiratory block,
again with and without observed age.

No age-decoder or endpoint-supervised scalar is an input. Every fitted fold
model, including all inner candidates and preprocessing objects, is saved,
immediately reloaded and checked for elementwise design and prediction parity.
Compatible reruns reuse artifacts; incompatible artifacts fail closed.
"""

from __future__ import annotations

import argparse
import json
import math
import sys
from pathlib import Path
from typing import Any, Sequence

import joblib
import numpy as np
import pandas as pd
from scipy.stats import t as student_t
from sklearn.preprocessing import OneHotEncoder, SplineTransformer, StandardScaler

import analyze_nhanes3_5y_matched_scalar_target_supervision_protocol_transfer_v0_2 as base


ROOT = Path(__file__).resolve().parents[1]
SCRIPT = Path(__file__).resolve()
SPEC = (
    ROOT
    / "analysis"
    / "NHANES3_5Y_CHRONOLOGICAL_AGE_INCREMENT_AND_PROXY_COMPARISON_SPEC_v0_1_20260827.md"
)
CANONICAL = base.CANONICAL
DEFAULT_OUT = (
    ROOT
    / "outputs"
    / "nhanes3_5y_chronological_age_increment_and_proxy_comparison_v0_1"
)

ANALYSIS_VERSION = (
    "nhanes3_5y_chronological_age_increment_and_proxy_comparison_v0_1"
)
SCHEMA_VERSION = "1.0"
STATUS = "retrospective_outcome_accessed_fixed_protocol_transfer_robustness_analysis"
MODEL_SEED = 20260827

AGE = base.AGE
OUTCOME = base.OUTCOME
BACKGROUND = base.BACKGROUND
WEIGHT = base.WEIGHT
BASE_WEIGHT = base.BASE_WEIGHT
STRATUM = base.STRATUM
PSU = base.PSU
N_OUTER = base.N_OUTER
N_INNER = base.N_INNER
C_GRID = base.C_GRID
EPSILON = base.EPSILON
PARITY_TOLERANCE = base.PARITY_TOLERANCE
TIE_TOLERANCE = base.TIE_TOLERANCE
FAY_REPLICATES = base.FAY_REPLICATES
FAY_RHO = base.FAY_RHO
FAY_DF = base.FAY_DF
AGE_KNOTS = (20.0, 35.0, 50.0, 65.0, 80.0)

MODEL_ORDER = ("F0", "F1", "R0", "R1")
EXPECTED_AGE_SELECTED_DOMAIN = "cardiorespiratory"
EXPECTED_AGE_SELECTED_FEATURES = (
    "MEC_SBP_MEAN",
    "MEC_DBP_MEAN",
    "PEP6DR",
    "SPPFEV1",
    "SPPFVC",
)
CANONICAL_FREEZE_KEYS = (
    "phase1",
    "phase2",
    "manifest",
    "phase1_folds",
    "phase2_folds",
    "fay_registry",
    "fay_factors",
    "fay_design",
)
SOURCE_MASK_FILES = {
    "source_selection_freeze": CANONICAL / "source_selection_freeze.json",
    "source_decision": CANONICAL / "source_decision.json",
    "source_model_artifact_audit": CANONICAL / "source_model_artifact_audit.json",
    "source_panel_policy_metrics": CANONICAL / "source_panel_policy_metrics.tsv",
}
CONTRAST_COLUMNS = {
    "V_F": "paired_V_F_loss",
    "V_R": "paired_V_R_loss",
    "P_proxy": "paired_P_proxy_loss",
    "D_block_0": "paired_D_block_0_loss",
    "D_block_1": "paired_D_block_1_loss",
    "D_replace": "paired_D_replace_loss",
}
EXPECTED_C_GRID = (
    0.001,
    0.003,
    0.01,
    0.03,
    0.1,
    0.3,
    1.0,
    3.0,
    10.0,
    30.0,
    100.0,
    300.0,
)


def configure_stdout() -> None:
    if hasattr(sys.stdout, "reconfigure"):
        sys.stdout.reconfigure(encoding="utf-8")


def validate_inherited_protocol_constants() -> None:
    base.require(AGE == "age_at_exam_years", "inherited age field changed")
    base.require(OUTCOME == "death_5y", "inherited outcome field changed")
    base.require(
        tuple(BACKGROUND) == ("sex", "race_ethnicity"),
        "inherited background fields changed",
    )
    base.require(
        (WEIGHT, BASE_WEIGHT, STRATUM, PSU)
        == ("analysis_weight", "mec_weight", "pseudo_stratum", "pseudo_psu"),
        "inherited survey interface changed",
    )
    base.require(N_OUTER == 5 and N_INNER == 3, "inherited fold counts changed")
    base.require(tuple(C_GRID) == EXPECTED_C_GRID, "inherited C grid changed")
    base.require(EPSILON == 1e-8, "inherited log-loss clipping changed")
    base.require(
        PARITY_TOLERANCE == 1e-12 and TIE_TOLERANCE == 1e-12,
        "inherited numerical tolerances changed",
    )
    base.require(
        FAY_REPLICATES == 32 and FAY_RHO == 0.3 and FAY_DF == 23,
        "inherited Fay-BRR protocol changed",
    )


def input_record(role: str, path: Path) -> dict[str, Any]:
    base.require(path.is_file(), f"missing frozen input: {path}")
    return {
        "role": role,
        "path": str(path.resolve()),
        "sha256": base.sha256(path),
        "size_bytes": int(path.stat().st_size),
    }


def validate_source_mask(
    ordered_features: list[str],
) -> tuple[dict[str, Any], list[str], list[str]]:
    for path in SOURCE_MASK_FILES.values():
        base.require(path.is_file(), f"missing source-mask evidence: {path}")

    freeze = json.loads(
        SOURCE_MASK_FILES["source_selection_freeze"].read_text(encoding="utf-8")
    )
    signature = str(freeze.get("freeze_signature", ""))
    payload = {key: value for key, value in freeze.items() if key != "freeze_signature"}
    base.require(signature == base.stable_hash(payload), "source selection freeze signature changed")
    base.require(
        freeze.get("analysis_version")
        == "nhanes3_phase1_to_phase2_endpoint_vs_age_acquisition_v0_2",
        "source selection analysis identity changed",
    )
    base.require(
        freeze.get("status") == "PASS_POLICY_CONTRAST",
        "source selection freeze is not PASS_POLICY_CONTRAST",
    )
    base.require(
        freeze.get("age_selected_domain") == EXPECTED_AGE_SELECTED_DOMAIN,
        "frozen age-selected domain changed",
    )
    base.require(
        freeze.get("target_outcomes_accessed") is False,
        "source freeze target-access flag changed",
    )
    domains = freeze.get("domains")
    base.require(isinstance(domains, dict), "source freeze domains are missing")
    selected = list(domains.get(EXPECTED_AGE_SELECTED_DOMAIN, []))
    base.require(
        selected == list(EXPECTED_AGE_SELECTED_FEATURES),
        "frozen age-selected feature block changed",
    )
    base.require(
        len(ordered_features) == len(set(ordered_features)) == 25,
        "canonical ordered full panel is not 25 unique measurements",
    )
    base.require(
        all(feature in ordered_features for feature in selected),
        "frozen age-selected block is not contained in the canonical panel",
    )
    remaining = [feature for feature in ordered_features if feature not in set(selected)]
    base.require(
        len(remaining) == len(set(remaining)) == 20,
        "ordered remaining panel is not 20 unique measurements",
    )

    source_decision = json.loads(
        SOURCE_MASK_FILES["source_decision"].read_text(encoding="utf-8")
    )
    base.require(
        source_decision.get("decision") == "PASS_POLICY_CONTRAST",
        "source decision did not pass",
    )
    base.require(
        source_decision.get("age_selected_domain") == EXPECTED_AGE_SELECTED_DOMAIN,
        "source decision age-selected domain differs from freeze",
    )
    source_audit = json.loads(
        SOURCE_MASK_FILES["source_model_artifact_audit"].read_text(encoding="utf-8")
    )
    base.require(source_audit.get("status") == "PASS", "source model audit is not PASS")

    source_metrics = base.read_table(SOURCE_MASK_FILES["source_panel_policy_metrics"])
    required = {"target", "domain", "design_weighted_loss"}
    base.require(required.issubset(source_metrics.columns), "source policy metrics are malformed")
    age_metrics = source_metrics.loc[source_metrics["target"].astype(str).eq("age")].copy()
    base.require(len(age_metrics) == 5, "source age-policy metric count changed")
    age_metrics["design_weighted_loss"] = pd.to_numeric(
        age_metrics["design_weighted_loss"], errors="raise"
    )
    minimum = float(age_metrics["design_weighted_loss"].min())
    minimum_domains = age_metrics.loc[
        age_metrics["design_weighted_loss"].le(minimum + TIE_TOLERANCE), "domain"
    ].astype(str).tolist()
    base.require(
        EXPECTED_AGE_SELECTED_DOMAIN in minimum_domains,
        "source age-selected domain no longer minimizes frozen age loss",
    )
    return freeze, selected, remaining


def build_models(
    full_features: list[str], remaining_features: list[str]
) -> dict[str, dict[str, Any]]:
    models = {
        "F0": {
            "panel": "full_25",
            "features": list(full_features),
            "include_age": False,
            "formula": "Y ~ W + X25",
        },
        "F1": {
            "panel": "full_25",
            "features": list(full_features),
            "include_age": True,
            "formula": "Y ~ W + X25 + fixed_knot_spline(A)",
        },
        "R0": {
            "panel": "remaining_20_after_age_selected_block",
            "features": list(remaining_features),
            "include_age": False,
            "formula": "Y ~ W + X20_remaining",
        },
        "R1": {
            "panel": "remaining_20_after_age_selected_block",
            "features": list(remaining_features),
            "include_age": True,
            "formula": "Y ~ W + X20_remaining + fixed_knot_spline(A)",
        },
    }
    base.require(tuple(models) == MODEL_ORDER, "model order changed")
    return models


def load_analysis_inputs() -> tuple[
    pd.DataFrame,
    np.ndarray,
    pd.DataFrame,
    list[str],
    list[str],
    dict[str, Any],
    list[dict[str, Any]],
]:
    # The predecessor validator loads both phases to verify the frozen canonical
    # contract. Only the returned Phase 2 interface is fitted in this analysis.
    _, phase2, manifest, features, factors, _ = base.load_inputs()
    forbidden = {"q_A", "q_Y"}.intersection(phase2.columns)
    base.require(not forbidden, f"derived scalar inputs are prohibited: {sorted(forbidden)}")
    freeze, selected, remaining = validate_source_mask(features)

    records: list[dict[str, Any]] = []
    for key in CANONICAL_FREEZE_KEYS:
        records.append(
            input_record(f"canonical_{key}", CANONICAL / base.CANONICAL_FILES[key])
        )
    for role, path in SOURCE_MASK_FILES.items():
        records.append(input_record(role, path))
    for role, path in (
        ("frozen_protocol", SPEC),
        ("analysis_script", SCRIPT),
        ("implementation_dependency_base_spec", base.SPEC),
        ("implementation_dependency_base_script", Path(base.__file__).resolve()),
        ("environment_policy", base.ENV_POLICY),
        ("environment_manifest", base.ENV_MANIFEST),
        ("environment_lock", base.ENV_LOCK),
    ):
        records.append(input_record(role, path))

    base.require(
        selected == list(EXPECTED_AGE_SELECTED_FEATURES),
        "validated selected feature block changed unexpectedly",
    )
    return phase2, factors, manifest, features, remaining, freeze, records


def build_analysis_signature(
    input_records: list[dict[str, Any]],
    models: dict[str, dict[str, Any]],
    freeze: dict[str, Any],
) -> tuple[str, dict[str, Any]]:
    payload = {
        "analysis_version": ANALYSIS_VERSION,
        "schema_version": SCHEMA_VERSION,
        "status": STATUS,
        "inputs": input_records,
        "population": "NHANES_III_Phase_2_age_20_to_79",
        "endpoint": "resolved_five_year_all_cause_mortality",
        "background": list(BACKGROUND),
        "models": models,
        "age_selected_domain": EXPECTED_AGE_SELECTED_DOMAIN,
        "age_selected_features": list(EXPECTED_AGE_SELECTED_FEATURES),
        "source_selection_freeze_signature": freeze["freeze_signature"],
        "remaining_panel_language": "remaining_20_after_age_selected_block",
        "age_knots": list(AGE_KNOTS),
        "age_spline": {
            "degree": 3,
            "include_bias": False,
            "extrapolation": "linear",
            "basis_standardization": "training_fold_MEC_weighted",
        },
        "measurement_preprocessing": (
            "training_fold_weighted_median_imputation_and_weighted_standardization"
        ),
        "C_grid": list(C_GRID),
        "outer_folds": N_OUTER,
        "inner_folds": N_INNER,
        "inner_loss_pooling": "sum_weighted_numerators_over_sum_weight_denominators",
        "tie_rule": "within_1e-12_choose_smaller_C",
        "fit_weight_normalization": "sum_to_n_within_fit",
        "Fay_BRR": {
            "replicates": FAY_REPLICATES,
            "rho": FAY_RHO,
            "df": FAY_DF,
        },
        "prohibited_inputs": ["q_A", "q_Y", "age_gap", "derived_age"],
        "model_seed": MODEL_SEED,
        "parity_tolerance": PARITY_TOLERANCE,
        "software": base.software_record(),
    }
    return base.stable_hash(payload), payload


def freeze_inputs(out: Path, input_records: list[dict[str, Any]]) -> None:
    frozen = out / "input_freeze"
    for key in CANONICAL_FREEZE_KEYS:
        source = CANONICAL / base.CANONICAL_FILES[key]
        base.copy_file_or_verify(source, frozen / source.name)
    for source in SOURCE_MASK_FILES.values():
        base.copy_file_or_verify(source, frozen / source.name)
    base.write_table_or_verify(pd.DataFrame(input_records), out / "input_manifest.tsv")


def fit_predictor_preprocessor(
    frame: pd.DataFrame,
    features: list[str],
    include_age: bool,
    weight: np.ndarray,
) -> tuple[dict[str, Any], np.ndarray]:
    base.require(features and len(features) == len(set(features)), "invalid feature panel")
    fit_weight = base.normalized_weights(weight)
    encoder = OneHotEncoder(handle_unknown="ignore", sparse_output=False)
    categorical = encoder.fit_transform(frame.loc[:, list(BACKGROUND)].astype(str))
    blocks = [categorical]
    names = [f"W__{name}" for name in encoder.get_feature_names_out(BACKGROUND)]

    raw = frame[features].to_numpy(float)
    base.require(
        np.isfinite(raw).sum(axis=0).min() > 0,
        "a measurement is all missing in a training fold",
    )
    imputer = base.WeightedMedianImputer().fit(raw, sample_weight=fit_weight)
    imputed = imputer.transform(raw)
    measurement_scaler = StandardScaler().fit(imputed, sample_weight=fit_weight)
    measured = measurement_scaler.transform(imputed)
    blocks.append(measured)
    names.extend(f"X__{feature}" for feature in features)

    age_spline: SplineTransformer | None = None
    age_scaler: StandardScaler | None = None
    if include_age:
        age_spline = SplineTransformer(
            degree=3,
            knots=np.asarray(AGE_KNOTS, dtype=float).reshape(-1, 1),
            include_bias=False,
            extrapolation="linear",
        )
        age_raw = age_spline.fit_transform(
            frame[[AGE]].to_numpy(float), sample_weight=fit_weight
        )
        age_scaler = StandardScaler().fit(age_raw, sample_weight=fit_weight)
        age_design = age_scaler.transform(age_raw)
        blocks.append(age_design)
        names.extend(f"A_spline_{index:02d}" for index in range(age_design.shape[1]))

    design = np.column_stack(blocks)
    base.require(np.isfinite(design).all(), "predictor design contains non-finite values")
    return {
        "background": list(BACKGROUND),
        "features": list(features),
        "include_age": bool(include_age),
        "encoder": encoder,
        "imputer": imputer,
        "measurement_scaler": measurement_scaler,
        "age_spline": age_spline,
        "age_scaler": age_scaler,
        "age_knots": list(AGE_KNOTS) if include_age else [],
        "design_feature_names": names,
        "fit_weight_normalization": "sum_to_n_within_fit",
    }, design


def transform_predictor_preprocessor(
    preprocessor: dict[str, Any], frame: pd.DataFrame
) -> np.ndarray:
    categorical = preprocessor["encoder"].transform(
        frame.loc[:, list(preprocessor["background"])].astype(str)
    )
    features = list(preprocessor["features"])
    measured = preprocessor["imputer"].transform(frame[features].to_numpy(float))
    measured = preprocessor["measurement_scaler"].transform(measured)
    blocks = [categorical, measured]
    if bool(preprocessor["include_age"]):
        age_raw = preprocessor["age_spline"].transform(frame[[AGE]].to_numpy(float))
        blocks.append(preprocessor["age_scaler"].transform(age_raw))
    design = np.column_stack(blocks)
    base.require(
        np.isfinite(design).all(),
        "reconstructed predictor design contains non-finite values",
    )
    return design


def verify_preprocessor_contract(
    preprocessor: dict[str, Any], model_spec: dict[str, Any], context: str
) -> None:
    base.require(
        list(preprocessor["background"]) == list(BACKGROUND),
        f"{context} preprocessor background order changed",
    )
    base.require(
        list(preprocessor["features"]) == list(model_spec["features"]),
        f"{context} preprocessor feature order changed",
    )
    include_age = bool(model_spec["include_age"])
    base.require(
        bool(preprocessor["include_age"]) == include_age,
        f"{context} preprocessor age inclusion changed",
    )
    encoder = preprocessor["encoder"]
    base.require(
        np.array_equal(
            np.asarray(encoder.feature_names_in_, dtype=str),
            np.asarray(BACKGROUND, dtype=str),
        ),
        f"{context} encoder input order changed",
    )
    expected_names = [
        f"W__{name}" for name in encoder.get_feature_names_out(BACKGROUND)
    ]
    expected_names.extend(f"X__{feature}" for feature in model_spec["features"])
    if include_age:
        age_spline = preprocessor["age_spline"]
        base.require(age_spline is not None, f"{context} age spline is missing")
        base.require(preprocessor["age_scaler"] is not None, f"{context} age scaler is missing")
        params = age_spline.get_params(deep=False)
        base.require(int(params["degree"]) == 3, f"{context} age spline degree changed")
        base.require(params["include_bias"] is False, f"{context} age spline bias changed")
        base.require(
            params["extrapolation"] == "linear",
            f"{context} age spline extrapolation changed",
        )
        knots = np.asarray(params["knots"], dtype=float).reshape(-1)
        base.require(
            knots.shape == (len(AGE_KNOTS),)
            and base.maximum_error(knots, AGE_KNOTS) <= PARITY_TOLERANCE,
            f"{context} age spline knots changed",
        )
        base.require(
            list(preprocessor["age_knots"]) == list(AGE_KNOTS),
            f"{context} stored age-knot metadata changed",
        )
        expected_names.extend(
            f"A_spline_{index:02d}"
            for index in range(int(age_spline.n_features_out_))
        )
    else:
        base.require(
            preprocessor["age_spline"] is None
            and preprocessor["age_scaler"] is None
            and list(preprocessor["age_knots"]) == [],
            f"{context} age-free preprocessor contains an age basis",
        )
    base.require(
        list(preprocessor["design_feature_names"]) == expected_names,
        f"{context} design feature names changed",
    )
    base.require(
        preprocessor["fit_weight_normalization"] == "sum_to_n_within_fit",
        f"{context} preprocessing weight normalization changed",
    )


def artifact_signature(
    analysis_signature: str,
    stage_signature: str,
    model_name: str,
    model_spec: dict[str, Any],
    outer_fold: int,
    training: pd.DataFrame,
    test: pd.DataFrame,
) -> str:
    inner_column = f"inner_fold_outer_{outer_fold}"
    payload = {
        "analysis_signature": analysis_signature,
        "stage_signature": stage_signature,
        "artifact_kind": "direct_measurement_chronological_age_endpoint_model",
        "model_name": model_name,
        "model_spec": model_spec,
        "outer_fold": int(outer_fold),
        "training_seqn": training["SEQN"].astype(int).tolist(),
        "test_seqn": test["SEQN"].astype(int).tolist(),
        "training_weight": training[WEIGHT].astype(float).tolist(),
        "test_weight": test[WEIGHT].astype(float).tolist(),
        "training_outcome": training[OUTCOME].astype(int).tolist(),
        "test_outcome": test[OUTCOME].astype(int).tolist(),
        "inner_assignments": training[["SEQN", inner_column]].astype(
            {"SEQN": int, inner_column: int}
        ).to_dict(orient="records"),
        "background": list(BACKGROUND),
        "age_knots": list(AGE_KNOTS),
        "measurement_preprocessing": "weighted_median_then_weighted_standardization",
        "C_grid": list(C_GRID),
        "inner_loss_pooling": "sum_weighted_numerators_over_sum_weight_denominators",
        "tie_rule": "within_1e-12_choose_smaller_C",
        "fit_weight_normalization": "sum_to_n_within_fit",
        "log_loss_clip": EPSILON,
        "model_seed": MODEL_SEED,
        "parity_tolerance": PARITY_TOLERANCE,
    }
    return base.stable_hash(payload)


def fit_artifact(
    analysis_signature: str,
    stage_signature: str,
    model_name: str,
    model_spec: dict[str, Any],
    outer_fold: int,
    frame: pd.DataFrame,
) -> dict[str, Any]:
    features = list(model_spec["features"])
    include_age = bool(model_spec["include_age"])
    training = frame.loc[frame["outer_fold"].ne(outer_fold)].sort_values(
        "SEQN", kind="mergesort"
    ).reset_index(drop=True)
    test = frame.loc[frame["outer_fold"].eq(outer_fold)].sort_values(
        "SEQN", kind="mergesort"
    ).reset_index(drop=True)
    inner_column = f"inner_fold_outer_{outer_fold}"
    inner_folds: list[dict[str, Any]] = []
    tuning_rows: list[dict[str, Any]] = []

    for inner_fold in range(N_INNER):
        inner_train = training.loc[training[inner_column].ne(inner_fold)].sort_values(
            "SEQN", kind="mergesort"
        ).reset_index(drop=True)
        inner_valid = training.loc[training[inner_column].eq(inner_fold)].sort_values(
            "SEQN", kind="mergesort"
        ).reset_index(drop=True)
        base.require(len(inner_train) > 0 and len(inner_valid) > 0, "empty inner fold")
        train_weight = inner_train[WEIGHT].to_numpy(float)
        valid_weight = inner_valid[WEIGHT].to_numpy(float)
        preprocessor, x_train = fit_predictor_preprocessor(
            inner_train, features, include_age, train_weight
        )
        x_valid = transform_predictor_preprocessor(preprocessor, inner_valid)
        y_train = inner_train[OUTCOME].to_numpy(int)
        y_valid = inner_valid[OUTCOME].to_numpy(int)
        candidate_fits: list[dict[str, Any]] = []
        for c_value in C_GRID:
            estimator = base.fit_logistic(x_train, y_train, train_weight, c_value)
            probability = base.predict_logistic(estimator, x_valid)
            losses = base.log_loss_vector(y_valid, probability)
            numerator = float(np.dot(valid_weight, losses))
            denominator = float(valid_weight.sum())
            candidate_fits.append(
                {
                    "C": float(c_value),
                    "estimator": estimator,
                    "valid_probability": probability,
                    "valid_loss_vector": losses,
                    "weighted_loss_numerator": numerator,
                    "weight_sum": denominator,
                }
            )
            tuning_rows.append(
                {
                    "model": model_name,
                    "outer_fold": int(outer_fold),
                    "inner_fold": int(inner_fold),
                    "C": float(c_value),
                    "n_valid": int(len(inner_valid)),
                    "events_valid": int(y_valid.sum()),
                    "weighted_loss_numerator": numerator,
                    "weight_sum": denominator,
                    "weighted_loss": numerator / denominator,
                }
            )
        inner_folds.append(
            {
                "inner_fold": int(inner_fold),
                "train_seqn": inner_train["SEQN"].to_numpy("int64"),
                "valid_seqn": inner_valid["SEQN"].to_numpy("int64"),
                "train_weight": train_weight,
                "valid_weight": valid_weight,
                "train_outcome": y_train,
                "valid_outcome": y_valid,
                "preprocessor": preprocessor,
                "x_train": x_train,
                "x_valid": x_valid,
                "candidate_fits": candidate_fits,
            }
        )

    selected_c = base.select_grid_value(tuning_rows, "C", C_GRID, prefer="small")
    final_weight = training[WEIGHT].to_numpy(float)
    final_preprocessor, x_training = fit_predictor_preprocessor(
        training, features, include_age, final_weight
    )
    x_test = transform_predictor_preprocessor(final_preprocessor, test)
    final_estimator = base.fit_logistic(
        x_training, training[OUTCOME].to_numpy(int), final_weight, selected_c
    )
    test_probability = base.predict_logistic(final_estimator, x_test)
    test_loss = base.log_loss_vector(test[OUTCOME].to_numpy(int), test_probability)
    return {
        "schema_version": SCHEMA_VERSION,
        "analysis_version": ANALYSIS_VERSION,
        "analysis_signature": analysis_signature,
        "stage_signature": stage_signature,
        "artifact_signature": artifact_signature(
            analysis_signature,
            stage_signature,
            model_name,
            model_spec,
            outer_fold,
            training,
            test,
        ),
        "artifact_kind": "direct_measurement_chronological_age_endpoint_model",
        "model_name": model_name,
        "model_spec": model_spec,
        "outer_fold": int(outer_fold),
        "inner_column": inner_column,
        "training_seqn": training["SEQN"].to_numpy("int64"),
        "test_seqn": test["SEQN"].to_numpy("int64"),
        "training_weight": final_weight,
        "test_weight": test[WEIGHT].to_numpy(float),
        "training_outcome": training[OUTCOME].to_numpy(int),
        "test_outcome": test[OUTCOME].to_numpy(int),
        "inner_folds": inner_folds,
        "tuning_rows": tuning_rows,
        "selected_C": float(selected_c),
        "final_preprocessor": final_preprocessor,
        "x_training": x_training,
        "x_test": x_test,
        "final_estimator": final_estimator,
        "test_probability": test_probability,
        "test_loss_vector": test_loss,
        "software": base.software_record(),
    }


def verify_artifact(
    artifact: dict[str, Any],
    analysis_signature: str,
    stage_signature: str,
    model_name: str,
    model_spec: dict[str, Any],
    outer_fold: int,
    frame: pd.DataFrame,
) -> dict[str, float]:
    training = frame.loc[frame["outer_fold"].ne(outer_fold)].sort_values(
        "SEQN", kind="mergesort"
    ).reset_index(drop=True)
    test = frame.loc[frame["outer_fold"].eq(outer_fold)].sort_values(
        "SEQN", kind="mergesort"
    ).reset_index(drop=True)
    expected_signature = artifact_signature(
        analysis_signature,
        stage_signature,
        model_name,
        model_spec,
        outer_fold,
        training,
        test,
    )
    base.require(artifact.get("schema_version") == SCHEMA_VERSION, "artifact schema changed")
    base.require(
        artifact.get("analysis_version") == ANALYSIS_VERSION,
        "artifact analysis version changed",
    )
    base.require(artifact.get("analysis_signature") == analysis_signature, "analysis signature changed")
    base.require(artifact.get("stage_signature") == stage_signature, "stage signature changed")
    base.require(artifact.get("artifact_signature") == expected_signature, "artifact signature mismatch")
    base.require(
        artifact.get("artifact_kind")
        == "direct_measurement_chronological_age_endpoint_model",
        "artifact kind changed",
    )
    base.require(artifact.get("model_name") == model_name, "artifact model identity changed")
    base.require(artifact.get("model_spec") == model_spec, "artifact model specification changed")
    base.require(int(artifact.get("outer_fold", -1)) == outer_fold, "outer-fold identity changed")
    base.require(
        artifact.get("inner_column") == f"inner_fold_outer_{outer_fold}",
        "inner-fold column changed",
    )
    base.require(
        np.array_equal(artifact["training_seqn"], training["SEQN"].to_numpy("int64")),
        "artifact training identities changed",
    )
    base.require(
        np.array_equal(artifact["test_seqn"], test["SEQN"].to_numpy("int64")),
        "artifact test identities changed",
    )
    for stored, current, message in (
        (artifact["training_weight"], training[WEIGHT], "artifact training weights changed"),
        (artifact["test_weight"], test[WEIGHT], "artifact test weights changed"),
        (artifact["training_outcome"], training[OUTCOME], "artifact training outcomes changed"),
        (artifact["test_outcome"], test[OUTCOME], "artifact test outcomes changed"),
    ):
        base.require(
            base.maximum_error(stored, current) <= PARITY_TOLERANCE,
            message,
        )

    by_seqn = frame.set_index("SEQN", drop=False)
    reconstructed_tuning: list[dict[str, Any]] = []
    reconstructed_tuning_rows: list[dict[str, Any]] = []
    max_design_error = 0.0
    max_prediction_error = 0.0
    base.require(
        [int(stored["inner_fold"]) for stored in artifact["inner_folds"]]
        == list(range(N_INNER)),
        "inner-fold artifact registry changed",
    )
    for stored in artifact["inner_folds"]:
        inner_fold = int(stored["inner_fold"])
        train_seqn = np.asarray(stored["train_seqn"], dtype="int64")
        valid_seqn = np.asarray(stored["valid_seqn"], dtype="int64")
        expected_train = training.loc[
            training[artifact["inner_column"]].ne(inner_fold), "SEQN"
        ].to_numpy("int64")
        expected_valid = training.loc[
            training[artifact["inner_column"]].eq(inner_fold), "SEQN"
        ].to_numpy("int64")
        base.require(np.array_equal(train_seqn, expected_train), "inner training identities changed")
        base.require(np.array_equal(valid_seqn, expected_valid), "inner validation identities changed")
        base.require(np.intersect1d(train_seqn, valid_seqn).size == 0, "inner-fold identity leakage")
        inner_train = by_seqn.loc[train_seqn].reset_index(drop=True)
        inner_valid = by_seqn.loc[valid_seqn].reset_index(drop=True)
        for stored_value, current, message in (
            (stored["train_weight"], inner_train[WEIGHT], "inner training weights changed"),
            (stored["valid_weight"], inner_valid[WEIGHT], "inner validation weights changed"),
            (stored["train_outcome"], inner_train[OUTCOME], "inner training outcomes changed"),
            (stored["valid_outcome"], inner_valid[OUTCOME], "inner validation outcomes changed"),
        ):
            base.require(
                base.maximum_error(stored_value, current) <= PARITY_TOLERANCE,
                message,
            )
        preprocessor = stored["preprocessor"]
        verify_preprocessor_contract(preprocessor, model_spec, "inner")
        x_train = transform_predictor_preprocessor(preprocessor, inner_train)
        x_valid = transform_predictor_preprocessor(preprocessor, inner_valid)
        max_design_error = max(
            max_design_error,
            base.maximum_error(x_train, stored["x_train"]),
            base.maximum_error(x_valid, stored["x_valid"]),
        )
        y_valid = inner_valid[OUTCOME].to_numpy(int)
        valid_weight = inner_valid[WEIGHT].to_numpy(float)
        base.require(
            [float(candidate["C"]) for candidate in stored["candidate_fits"]]
            == [float(value) for value in C_GRID],
            "inner candidate grid changed",
        )
        for candidate in stored["candidate_fits"]:
            base.require(
                float(candidate["estimator"].C) == float(candidate["C"]),
                "inner candidate estimator C changed",
            )
            probability = base.predict_logistic(candidate["estimator"], x_valid)
            losses = base.log_loss_vector(y_valid, probability)
            max_prediction_error = max(
                max_prediction_error,
                base.maximum_error(probability, candidate["valid_probability"]),
                base.maximum_error(losses, candidate["valid_loss_vector"]),
            )
            numerator = float(np.dot(valid_weight, losses))
            denominator = float(valid_weight.sum())
            base.require(
                abs(numerator - float(candidate["weighted_loss_numerator"]))
                <= PARITY_TOLERANCE,
                "inner tuning numerator changed",
            )
            base.require(
                abs(denominator - float(candidate["weight_sum"]))
                <= PARITY_TOLERANCE,
                "inner tuning denominator changed",
            )
            reconstructed_tuning.append(
                {
                    "C": float(candidate["C"]),
                    "weighted_loss_numerator": numerator,
                    "weight_sum": denominator,
                }
            )
            reconstructed_tuning_rows.append(
                {
                    "model": model_name,
                    "outer_fold": int(outer_fold),
                    "inner_fold": int(inner_fold),
                    "C": float(candidate["C"]),
                    "n_valid": int(len(inner_valid)),
                    "events_valid": int(y_valid.sum()),
                    "weighted_loss_numerator": numerator,
                    "weight_sum": denominator,
                    "weighted_loss": numerator / denominator,
                }
            )
    reconstructed_tuning_table = pd.DataFrame(reconstructed_tuning_rows)
    stored_tuning_table = pd.DataFrame(artifact["tuning_rows"])
    base.require(
        stored_tuning_table.columns.tolist()
        == reconstructed_tuning_table.columns.tolist(),
        "stored tuning-table schema changed",
    )
    try:
        pd.testing.assert_frame_equal(
            stored_tuning_table.reset_index(drop=True),
            reconstructed_tuning_table.reset_index(drop=True),
            check_dtype=False,
            check_exact=False,
            rtol=0.0,
            atol=PARITY_TOLERANCE,
        )
    except AssertionError as error:
        raise RuntimeError(
            "STOP_DATA_OR_ARTIFACT_FAIL: stored tuning rows changed"
        ) from error
    selected_c = base.select_grid_value(
        reconstructed_tuning, "C", C_GRID, prefer="small"
    )
    base.require(selected_c == float(artifact["selected_C"]), "selected C changed")

    final_preprocessor = artifact["final_preprocessor"]
    verify_preprocessor_contract(final_preprocessor, model_spec, "final")
    x_training = transform_predictor_preprocessor(final_preprocessor, training)
    x_test = transform_predictor_preprocessor(final_preprocessor, test)
    base.require(
        float(artifact["final_estimator"].C) == float(artifact["selected_C"]),
        "final estimator C changed",
    )
    probability = base.predict_logistic(artifact["final_estimator"], x_test)
    losses = base.log_loss_vector(test[OUTCOME].to_numpy(int), probability)
    max_design_error = max(
        max_design_error,
        base.maximum_error(x_training, artifact["x_training"]),
        base.maximum_error(x_test, artifact["x_test"]),
    )
    max_prediction_error = max(
        max_prediction_error,
        base.maximum_error(probability, artifact["test_probability"]),
        base.maximum_error(losses, artifact["test_loss_vector"]),
    )
    names = list(final_preprocessor["design_feature_names"])
    coefficients = np.asarray(artifact["final_estimator"].coef_, dtype=float)
    base.require(coefficients.shape == (1, len(names)), "final coefficient shape changed")
    base.require(
        max(max_design_error, max_prediction_error) <= PARITY_TOLERANCE,
        "reloaded model parity failed",
    )
    return {
        "max_design_error": max_design_error,
        "max_prediction_error": max_prediction_error,
    }


def fit_or_load_artifact(
    path: Path,
    analysis_signature: str,
    stage_signature: str,
    model_name: str,
    model_spec: dict[str, Any],
    outer_fold: int,
    frame: pd.DataFrame,
) -> tuple[dict[str, Any], dict[str, Any]]:
    fitted = False
    if path.exists():
        artifact = joblib.load(path)
        training = frame.loc[frame["outer_fold"].ne(outer_fold)].sort_values(
            "SEQN", kind="mergesort"
        ).reset_index(drop=True)
        test = frame.loc[frame["outer_fold"].eq(outer_fold)].sort_values(
            "SEQN", kind="mergesort"
        ).reset_index(drop=True)
        expected = artifact_signature(
            analysis_signature,
            stage_signature,
            model_name,
            model_spec,
            outer_fold,
            training,
            test,
        )
        base.require(
            artifact.get("analysis_signature") == analysis_signature,
            f"incompatible cached analysis signature: {path}",
        )
        base.require(
            artifact.get("artifact_signature") == expected,
            f"incompatible cached artifact: {path}",
        )
    else:
        artifact = fit_artifact(
            analysis_signature,
            stage_signature,
            model_name,
            model_spec,
            outer_fold,
            frame,
        )
        base.save_joblib_atomic(artifact, path)
        fitted = True
    reloaded = joblib.load(path)
    parity = verify_artifact(
        reloaded,
        analysis_signature,
        stage_signature,
        model_name,
        model_spec,
        outer_fold,
        frame,
    )
    return reloaded, {
        "stage": "target_direct_measurement_age_increment",
        "target_refit_label": "",
        "model": model_name,
        "panel": model_spec["panel"],
        "feature_count": int(len(model_spec["features"])),
        "include_age": bool(model_spec["include_age"]),
        "outer_fold": int(outer_fold),
        "selected_C": float(reloaded["selected_C"]),
        "design_feature_count": int(
            len(reloaded["final_preprocessor"]["design_feature_names"])
        ),
        "artifact_path": str(path.resolve()),
        "artifact_sha256": base.sha256(path),
        "cache_action": "fitted" if fitted else "reused",
        "max_design_error": parity["max_design_error"],
        "max_prediction_error": parity["max_prediction_error"],
        "downstream_tuning_estimator_count": int(
            sum(len(fold["candidate_fits"]) for fold in reloaded["inner_folds"])
        ),
        "downstream_final_estimator_count": 1,
    }


def final_coefficient_rows(
    artifact: dict[str, Any], label: str
) -> list[dict[str, Any]]:
    estimator = artifact["final_estimator"]
    names = list(artifact["final_preprocessor"]["design_feature_names"])
    coefficients = np.asarray(estimator.coef_, dtype=float).reshape(-1)
    base.require(len(names) == len(coefficients), "coefficient feature order changed")
    common = {
        "target_refit_label": label,
        "model": artifact["model_name"],
        "panel": artifact["model_spec"]["panel"],
        "include_age": bool(artifact["model_spec"]["include_age"]),
        "outer_fold": int(artifact["outer_fold"]),
        "selected_C": float(artifact["selected_C"]),
    }
    rows = [
        {
            **common,
            "term_order": 0,
            "term": "(Intercept)",
            "term_type": "intercept",
            "coefficient": float(np.asarray(estimator.intercept_, dtype=float)[0]),
        }
    ]
    for order, (name, coefficient) in enumerate(zip(names, coefficients), start=1):
        if name.startswith("W__"):
            term_type = "background"
        elif name.startswith("X__"):
            term_type = "measurement"
        elif name.startswith("A_spline_"):
            term_type = "chronological_age_spline"
        else:
            term_type = "unknown"
        rows.append(
            {
                **common,
                "term_order": order,
                "term": name,
                "term_type": term_type,
                "coefficient": float(coefficient),
            }
        )
    return rows


def add_contrast_vectors(prediction: pd.DataFrame) -> pd.DataFrame:
    output = prediction.copy()
    required = {f"loss_{model}" for model in MODEL_ORDER}
    base.require(required.issubset(output.columns), "model loss vectors are incomplete")
    output["paired_V_F_loss"] = output["loss_F0"] - output["loss_F1"]
    output["paired_V_R_loss"] = output["loss_R0"] - output["loss_R1"]
    output["paired_P_proxy_loss"] = (
        output["paired_V_R_loss"] - output["paired_V_F_loss"]
    )
    output["paired_D_block_0_loss"] = output["loss_R0"] - output["loss_F0"]
    output["paired_D_block_1_loss"] = output["loss_R1"] - output["loss_F1"]
    output["paired_D_replace_loss"] = output["loss_R1"] - output["loss_F0"]
    algebra = output["paired_D_block_0_loss"] - output["paired_D_block_1_loss"]
    base.require(
        base.maximum_error(algebra, output["paired_P_proxy_loss"])
        <= PARITY_TOLERANCE,
        "proxy contrast algebra changed",
    )
    return output


def run_weight_set(
    out: Path,
    base_frame: pd.DataFrame,
    analysis_signature: str,
    source_freeze_signature: str,
    models: dict[str, dict[str, Any]],
    label: str,
    weight_factor: np.ndarray,
) -> tuple[pd.DataFrame, pd.DataFrame, list[dict[str, Any]], list[dict[str, Any]]]:
    base.require(len(weight_factor) == len(base_frame), "replicate weight length changed")
    frame = base_frame.copy()
    frame[WEIGHT] = frame[BASE_WEIGHT].to_numpy(float) * np.asarray(
        weight_factor, dtype=float
    )
    base.require(
        np.isfinite(frame[WEIGHT]).all() and frame[WEIGHT].gt(0).all(),
        "invalid target weights",
    )
    stage_signature = base.stable_hash(
        {
            "analysis_signature": analysis_signature,
            "stage": "phase2_direct_measurement_chronological_age_increment",
            "label": label,
            "ordered_seqn": frame["SEQN"].astype(int).tolist(),
            "ordered_weight": frame[WEIGHT].astype(float).tolist(),
            "ordered_age": frame[AGE].astype(float).tolist(),
            "models": models,
            "source_selection_freeze_signature": source_freeze_signature,
            "derived_age_inputs_used": False,
        }
    )
    prediction = frame[
        ["SEQN", "outer_fold", OUTCOME, BASE_WEIGHT, WEIGHT, STRATUM, PSU, AGE]
    ].copy()
    tuning_tables: list[pd.DataFrame] = []
    records: list[dict[str, Any]] = []
    coefficient_rows: list[dict[str, Any]] = []
    model_root = (
        out / "models" / "primary"
        if label == "primary"
        else out / "models" / "fay_brr" / label
    )
    expected_artifact_paths = {
        model_root / model_name / f"outer_fold_{outer_fold:02d}.joblib"
        for model_name in MODEL_ORDER
        for outer_fold in range(N_OUTER)
    }
    existing_artifact_paths = (
        set(model_root.rglob("*.joblib")) if model_root.exists() else set()
    )
    base.require(
        not existing_artifact_paths
        or existing_artifact_paths == expected_artifact_paths,
        (
            f"partial or unexpected cache for {label}: found "
            f"{len(existing_artifact_paths)} of {len(expected_artifact_paths)} exact artifacts"
        ),
    )
    expected_cache_action = "reused" if existing_artifact_paths else "fitted"
    position_lookup = frame.reset_index().set_index("SEQN")["index"]
    for model_name in MODEL_ORDER:
        model_spec = models[model_name]
        probability = np.full(len(frame), np.nan, dtype=float)
        for outer_fold in range(N_OUTER):
            artifact, record = fit_or_load_artifact(
                model_root / model_name / f"outer_fold_{outer_fold:02d}.joblib",
                analysis_signature,
                stage_signature,
                model_name,
                model_spec,
                outer_fold,
                frame,
            )
            base.require(
                record["cache_action"] == expected_cache_action,
                f"cache state changed while processing {label}",
            )
            record["target_refit_label"] = label
            records.append(record)
            coefficient_rows.extend(final_coefficient_rows(artifact, label))
            test_ids = np.asarray(artifact["test_seqn"], dtype="int64")
            positions = position_lookup.loc[test_ids].to_numpy(int)
            probability[positions] = np.asarray(
                artifact["test_probability"], dtype=float
            )
            tuning = pd.DataFrame(artifact["tuning_rows"])
            tuning.insert(0, "target_refit_label", label)
            tuning_tables.append(tuning)
        base.require(
            np.isfinite(probability).all(),
            f"incomplete OOF predictions for {model_name}",
        )
        prediction[f"probability_{model_name}"] = probability
        prediction[f"loss_{model_name}"] = base.log_loss_vector(
            frame[OUTCOME].to_numpy(int), probability
        )
    prediction = add_contrast_vectors(prediction)
    return (
        prediction,
        pd.concat(tuning_tables, ignore_index=True),
        records,
        coefficient_rows,
    )


def summarize(
    prediction: pd.DataFrame,
) -> tuple[dict[str, float], pd.DataFrame]:
    estimates: dict[str, float] = {}
    for model_name in MODEL_ORDER:
        estimates[f"loss_{model_name}"] = base.weighted_mean(
            prediction[f"loss_{model_name}"], prediction[WEIGHT]
        )
    for contrast, column in CONTRAST_COLUMNS.items():
        estimates[contrast] = base.weighted_mean(prediction[column], prediction[WEIGHT])

    fold_rows: list[dict[str, Any]] = []
    for outer_fold in range(N_OUTER):
        block = prediction.loc[prediction["outer_fold"].eq(outer_fold)]
        row: dict[str, Any] = {
            "outer_fold": int(outer_fold),
            "n": int(len(block)),
            "deaths_5y": int(block[OUTCOME].sum()),
        }
        for model_name in MODEL_ORDER:
            row[f"loss_{model_name}"] = base.weighted_mean(
                block[f"loss_{model_name}"], block[WEIGHT]
            )
        for contrast, column in CONTRAST_COLUMNS.items():
            value = base.weighted_mean(block[column], block[WEIGHT])
            row[contrast] = value
            row[f"{contrast}_positive"] = bool(value > 0)
            row[f"{contrast}_negative"] = bool(value < 0)
        fold_rows.append(row)
    return estimates, pd.DataFrame(fold_rows)


def primary_probe_table(
    estimates: dict[str, float], fold_table: pd.DataFrame
) -> pd.DataFrame:
    rows: list[dict[str, Any]] = []
    for model_name in MODEL_ORDER:
        rows.append(
            {
                "metric_type": "model_loss",
                "metric": f"loss_{model_name}",
                "estimate": estimates[f"loss_{model_name}"],
                "positive_outer_folds": np.nan,
                "negative_outer_folds": np.nan,
                "formal_Fay_BRR_inference": False,
            }
        )
    for contrast in CONTRAST_COLUMNS:
        rows.append(
            {
                "metric_type": "paired_contrast",
                "metric": contrast,
                "estimate": estimates[contrast],
                "positive_outer_folds": int(fold_table[f"{contrast}_positive"].sum()),
                "negative_outer_folds": int(fold_table[f"{contrast}_negative"].sum()),
                "formal_Fay_BRR_inference": False,
            }
        )
    return pd.DataFrame(rows)


def classify_primary_promotion(
    estimates: dict[str, float], fold_table: pd.DataFrame
) -> dict[str, Any]:
    v_f_positive_outer_folds = int(fold_table["V_F_positive"].sum())
    p_proxy_positive_outer_folds = int(fold_table["P_proxy_positive"].sum())
    age_increment_pattern = bool(
        estimates["V_F"] > 0 and v_f_positive_outer_folds >= 4
    )
    proxy_substitution_pattern = bool(
        estimates["D_block_0"] > 0
        and estimates["P_proxy"] > 0
        and p_proxy_positive_outer_folds >= 4
    )
    promotion_permitted = bool(age_increment_pattern or proxy_substitution_pattern)
    return {
        "promotion_permitted": promotion_permitted,
        "classification": (
            "PROMOTE_TO_FORMAL_FAY_BRR"
            if promotion_permitted
            else "STOP_ABSENT_OR_AMBIGUOUS_PRIMARY_PATTERN"
        ),
        "age_increment_pattern": {
            "passed": age_increment_pattern,
            "requirements": "V_F > 0 and V_F positive in at least 4 of 5 outer folds",
            "V_F": float(estimates["V_F"]),
            "V_F_positive_outer_folds": v_f_positive_outer_folds,
        },
        "proxy_substitution_pattern": {
            "passed": proxy_substitution_pattern,
            "requirements": (
                "D_block_0 > 0, P_proxy > 0, and P_proxy positive in at least "
                "4 of 5 outer folds"
            ),
            "D_block_0": float(estimates["D_block_0"]),
            "P_proxy": float(estimates["P_proxy"]),
            "P_proxy_positive_outer_folds": p_proxy_positive_outer_folds,
        },
        "no_rescue_rule": (
            "If neither frozen primary pattern passes, formal Fay-BRR escalation stops; "
            "no post-hoc rescue analysis is attempted."
        ),
    }


def fay_interval(full_estimate: float, replicates: Sequence[float]) -> dict[str, Any]:
    values = np.asarray(replicates, dtype=float)
    base.require(
        values.shape == (FAY_REPLICATES,) and np.isfinite(values).all(),
        "Fay-BRR contrast estimates are incomplete",
    )
    variance = float(
        np.sum((values - float(full_estimate)) ** 2)
        / (FAY_REPLICATES * (1.0 - FAY_RHO) ** 2)
    )
    standard_error = float(math.sqrt(variance))
    critical = float(student_t.ppf(0.975, FAY_DF))
    return {
        "estimate": float(full_estimate),
        "variance": variance,
        "standard_error": standard_error,
        "critical_t": critical,
        "df": FAY_DF,
        "ci_lower": float(full_estimate - critical * standard_error),
        "ci_upper": float(full_estimate + critical * standard_error),
        "replicates": FAY_REPLICATES,
        "rho": FAY_RHO,
        "conditional_on": "the_frozen_Phase1_age_selected_cardiorespiratory_mask",
    }


def deterministic_model_index(records: list[dict[str, Any]]) -> pd.DataFrame:
    rows = [
        {key: value for key, value in record.items() if key != "cache_action"}
        for record in records
    ]
    table = pd.DataFrame(rows).sort_values(
        ["target_refit_label", "model", "outer_fold"], kind="mergesort"
    ).reset_index(drop=True)
    base.require(not table["artifact_path"].duplicated().any(), "duplicate model artifact path")
    return table


def deterministic_coefficients(rows: list[dict[str, Any]]) -> pd.DataFrame:
    return pd.DataFrame(rows).sort_values(
        ["target_refit_label", "model", "outer_fold", "term_order"],
        kind="mergesort",
    ).reset_index(drop=True)


def audit_models(
    records: list[dict[str, Any]], expected: int, inventory_root: Path
) -> dict[str, Any]:
    base.require(len(records) == expected, f"expected {expected} artifacts, found {len(records)}")
    paths = [str(record["artifact_path"]) for record in records]
    base.require(len(paths) == len(set(paths)), "model artifact paths are duplicated")
    indexed_paths = {Path(path).resolve() for path in paths}
    inventory_paths = (
        {path.resolve() for path in inventory_root.rglob("*.joblib")}
        if inventory_root.exists()
        else set()
    )
    base.require(
        inventory_paths == indexed_paths,
        (
            f"model artifact inventory mismatch under {inventory_root}: "
            f"indexed={len(indexed_paths)}, found={len(inventory_paths)}"
        ),
    )
    max_design = max(float(record["max_design_error"]) for record in records)
    max_prediction = max(float(record["max_prediction_error"]) for record in records)
    base.require(
        max(max_design, max_prediction) <= PARITY_TOLERANCE,
        "model artifact parity audit failed",
    )
    label_counts = pd.Series(
        [str(record["target_refit_label"]) for record in records], dtype="object"
    ).value_counts().sort_index()
    base.require(
        bool(label_counts.eq(len(MODEL_ORDER) * N_OUTER).all()),
        "a weight set does not contain exactly 20 model artifacts",
    )
    for record in records:
        path = Path(record["artifact_path"])
        base.require(path.is_file(), f"indexed model artifact is missing: {path}")
        base.require(
            base.sha256(path) == record["artifact_sha256"],
            f"indexed model artifact hash changed: {path}",
        )
    return {
        "status": "PASS",
        "analysis_version": ANALYSIS_VERSION,
        "artifact_count": int(len(records)),
        "expected_artifact_count": int(expected),
        "artifact_inventory_root": str(inventory_root.resolve()),
        "artifact_inventory_count": int(len(inventory_paths)),
        "weight_set_artifact_counts": {
            str(key): int(value) for key, value in label_counts.items()
        },
        "downstream_tuning_estimator_count": int(
            sum(record["downstream_tuning_estimator_count"] for record in records)
        ),
        "downstream_final_estimator_count": int(
            sum(record["downstream_final_estimator_count"] for record in records)
        ),
        "maximum_design_reconstruction_error": max_design,
        "maximum_prediction_reconstruction_error": max_prediction,
        "parity_tolerance": PARITY_TOLERANCE,
        "cache_policy": "compatible_artifacts_reused; incompatible_artifacts_fail_closed",
    }


def manifest_rows(paths: Sequence[Path], root: Path) -> pd.DataFrame:
    rows = []
    for path in sorted(set(paths), key=lambda item: str(item).lower()):
        if not path.is_file():
            continue
        rows.append(
            {
                "relative_path": str(path.relative_to(root)).replace("\\", "/"),
                "size_bytes": int(path.stat().st_size),
                "sha256": base.sha256(path),
            }
        )
    return pd.DataFrame(rows)


def write_primary_manifest(out: Path) -> None:
    target = out / "primary_artifact_manifest.tsv"
    explicit_names = {
        "analysis_signature.json",
        "input_manifest.tsv",
        "ordered_5x5_feature_manifest.tsv",
        "frozen_feature_mask.json",
        "frozen_feature_mask.tsv",
        "target_primary_oof_predictions_and_losses.tsv.gz",
        "target_primary_inner_tuning.tsv.gz",
        "target_primary_fold_contrasts.tsv",
        "target_primary_probe.tsv",
        "target_primary_probe.json",
        "model_artifact_index_primary.tsv",
        "model_artifact_audit_primary.json",
        "model_coefficients_primary.tsv.gz",
        "PRIMARY_REPORT.md",
    }
    paths: list[Path] = []
    for path in out.rglob("*"):
        if not path.is_file() or path == target:
            continue
        relative = str(path.relative_to(out)).replace("\\", "/")
        if (
            relative.startswith("input_freeze/")
            or relative.startswith("models/primary/")
            or path.name in explicit_names
        ):
            paths.append(path)
    base.write_table_or_verify(manifest_rows(paths, out), target)


def write_output_manifest(out: Path) -> None:
    target = out / "artifact_manifest.tsv"
    paths = [path for path in out.rglob("*") if path.is_file() and path != target]
    base.write_table_or_verify(manifest_rows(paths, out), target)


def run(args: argparse.Namespace) -> int:
    configure_stdout()
    validate_inherited_protocol_constants()
    out = Path(args.output_dir).resolve()
    base.require(out.name == ANALYSIS_VERSION, "output basename must retain the frozen version")
    base.require(out != CANONICAL.resolve(), "new analysis may not write into its canonical predecessor")
    out.mkdir(parents=True, exist_ok=True)

    phase2, factors, manifest, full_features, remaining_features, freeze, input_records = (
        load_analysis_inputs()
    )
    models = build_models(full_features, remaining_features)
    analysis_signature, signature_payload = build_analysis_signature(
        input_records, models, freeze
    )
    freeze_inputs(out, input_records)
    base.write_json_or_verify(
        out / "analysis_signature.json",
        {"analysis_signature": analysis_signature, "signature_payload": signature_payload},
    )
    base.write_table_or_verify(manifest, out / "ordered_5x5_feature_manifest.tsv")
    mask_payload = {
        "analysis_version": ANALYSIS_VERSION,
        "source_selection_freeze_signature": freeze["freeze_signature"],
        "age_selected_domain": EXPECTED_AGE_SELECTED_DOMAIN,
        "age_selected_features": list(EXPECTED_AGE_SELECTED_FEATURES),
        "full_features": list(full_features),
        "remaining_features": list(remaining_features),
        "full_feature_count": 25,
        "remaining_feature_count": 20,
        "remaining_panel_label": "remaining_20_after_age_selected_block",
        "language_guard": (
            "The remaining-20 panel is not asserted to be free of age-related information."
        ),
    }
    base.write_json_or_verify(out / "frozen_feature_mask.json", mask_payload)
    mask_rows = []
    selected_set = set(EXPECTED_AGE_SELECTED_FEATURES)
    for order, feature in enumerate(full_features, start=1):
        mask_rows.append(
            {
                "feature_order": order,
                "feature": feature,
                "in_phase1_age_selected_block": feature in selected_set,
                "in_remaining_20_panel": feature not in selected_set,
            }
        )
    base.write_table_or_verify(pd.DataFrame(mask_rows), out / "frozen_feature_mask.tsv")

    primary_prediction, primary_tuning, primary_records, primary_coefficients = (
        run_weight_set(
            out,
            phase2,
            analysis_signature,
            str(freeze["freeze_signature"]),
            models,
            "primary",
            np.ones(len(phase2), dtype=float),
        )
    )
    primary_estimates, primary_folds = summarize(primary_prediction)
    promotion_gate = classify_primary_promotion(primary_estimates, primary_folds)
    probe = primary_probe_table(primary_estimates, primary_folds)
    probe["promotion_permitted"] = promotion_gate["promotion_permitted"]
    probe["primary_classification"] = promotion_gate["classification"]
    probe["age_increment_pattern_passed"] = promotion_gate[
        "age_increment_pattern"
    ]["passed"]
    probe["proxy_substitution_pattern_passed"] = promotion_gate[
        "proxy_substitution_pattern"
    ]["passed"]
    base.write_table_or_verify(
        primary_prediction, out / "target_primary_oof_predictions_and_losses.tsv.gz"
    )
    base.write_table_or_verify(
        primary_tuning, out / "target_primary_inner_tuning.tsv.gz"
    )
    base.write_table_or_verify(
        primary_folds, out / "target_primary_fold_contrasts.tsv"
    )
    base.write_table_or_verify(probe, out / "target_primary_probe.tsv")
    primary_index = deterministic_model_index(primary_records)
    primary_audit = audit_models(
        primary_records, len(MODEL_ORDER) * N_OUTER, out / "models" / "primary"
    )
    base.write_table_or_verify(primary_index, out / "model_artifact_index_primary.tsv")
    base.write_json_or_verify(out / "model_artifact_audit_primary.json", primary_audit)
    base.write_table_or_verify(
        deterministic_coefficients(primary_coefficients),
        out / "model_coefficients_primary.tsv.gz",
    )
    primary_result = {
        "analysis_version": ANALYSIS_VERSION,
        "status": STATUS,
        "stage": "primary_probe",
        "decision": promotion_gate["classification"],
        "promotion_permitted": promotion_gate["promotion_permitted"],
        "promotion_gate": promotion_gate,
        "formal_Fay_BRR_inference": False,
        "target_population_n": int(len(phase2)),
        "target_deaths_5y": int(phase2[OUTCOME].sum()),
        "models": {name: models[name]["formula"] for name in MODEL_ORDER},
        "primary_estimates": primary_estimates,
        "outer_fold_signs": {
            contrast: {
                "positive": int(primary_folds[f"{contrast}_positive"].sum()),
                "negative": int(primary_folds[f"{contrast}_negative"].sum()),
            }
            for contrast in CONTRAST_COLUMNS
        },
        "primary_model_artifact_status": primary_audit["status"],
        "primary_model_artifact_count": primary_audit["artifact_count"],
        "claim_boundary": (
            "Primary point estimates only; no formal survey-design interval. The reduced "
            "panel is the remaining-20 after a frozen Phase 1 age-selected block, not a "
            "panel proven free of age-related information."
        ),
    }
    base.write_json_or_verify(out / "target_primary_probe.json", primary_result)
    primary_report = f"""# NHANES III chronological-age increment primary probe v0.1

- Status: **{STATUS}**
- Stage: **PRIMARY PROBE; NO FORMAL FAY-BRR INFERENCE**
- Phase 2 participants/deaths: {len(phase2)} / {int(phase2[OUTCOME].sum())}
- `V_F = L(F0)-L(F1)`: {primary_estimates['V_F']:.9f}
- `V_R = L(R0)-L(R1)`: {primary_estimates['V_R']:.9f}
- `P_proxy = V_R-V_F`: {primary_estimates['P_proxy']:.9f}
- Primary classification: **{promotion_gate['classification']}**
- `promotion_permitted`: **{str(promotion_gate['promotion_permitted']).lower()}**
- Persisted/reloaded primary models: {primary_audit['artifact_count']} / 20

The removed variables are the frozen Phase 1 age-selected cardiorespiratory
block. The retained variables are the remaining-20 panel; they are not asserted
to be free of age-related information. Formal inference requires both
`--stage all` and passage of the frozen primary promotion gate. Failure of both
primary patterns stops escalation without a post-hoc rescue analysis.
"""
    base.write_text_or_verify(out / "PRIMARY_REPORT.md", primary_report)
    write_primary_manifest(out)
    if args.stage == "primary":
        print(json.dumps(primary_result, indent=2, ensure_ascii=False))
        return 0

    if not promotion_gate["promotion_permitted"]:
        stopped_result = dict(primary_result)
        stopped_result.update(
            {
                "requested_stage": "all",
                "formal_Fay_BRR_executed": False,
                "stop_reason": (
                    "The frozen primary promotion gate was not met; formal Fay-BRR "
                    "escalation stopped without rescue analysis."
                ),
            }
        )
        print(json.dumps(stopped_result, indent=2, ensure_ascii=False))
        return 2

    all_records = list(primary_records)
    all_coefficients = list(primary_coefficients)
    replicate_rows: list[dict[str, Any]] = []
    replicate_tuning_tables: list[pd.DataFrame] = []
    for replicate in range(FAY_REPLICATES):
        label = f"replicate_{replicate:02d}"
        prediction, tuning, records, coefficients = run_weight_set(
            out,
            phase2,
            analysis_signature,
            str(freeze["freeze_signature"]),
            models,
            label,
            factors[replicate],
        )
        all_records.extend(records)
        all_coefficients.extend(coefficients)
        estimates, folds = summarize(prediction)
        base.write_table_or_verify(
            prediction,
            out / "fay_brr_predictions" / f"{label}_oof_predictions_and_losses.tsv.gz",
        )
        replicate_tuning_tables.append(tuning)
        row: dict[str, Any] = {"replicate": int(replicate)}
        row.update(estimates)
        for contrast in CONTRAST_COLUMNS:
            row[f"{contrast}_positive_outer_folds"] = int(
                folds[f"{contrast}_positive"].sum()
            )
            row[f"{contrast}_negative_outer_folds"] = int(
                folds[f"{contrast}_negative"].sum()
            )
        replicate_rows.append(row)

    replicate_table = pd.DataFrame(replicate_rows)
    base.write_table_or_verify(
        replicate_table, out / "target_fay_brr_replicate_estimates.tsv"
    )
    base.write_table_or_verify(
        pd.concat(replicate_tuning_tables, ignore_index=True),
        out / "target_fay_brr_inner_tuning.tsv.gz",
    )
    intervals = {
        contrast: fay_interval(primary_estimates[contrast], replicate_table[contrast])
        for contrast in CONTRAST_COLUMNS
    }
    formal_index = deterministic_model_index(all_records)
    expected_artifacts = (FAY_REPLICATES + 1) * len(MODEL_ORDER) * N_OUTER
    base.require(expected_artifacts == 660, "formal artifact-count formula changed")
    formal_audit = audit_models(all_records, expected_artifacts, out / "models")
    base.write_table_or_verify(formal_index, out / "model_artifact_index.tsv")
    base.write_json_or_verify(out / "model_artifact_audit.json", formal_audit)
    base.write_table_or_verify(
        deterministic_coefficients(all_coefficients), out / "model_coefficients.tsv.gz"
    )

    formal_result = {
        "analysis_version": ANALYSIS_VERSION,
        "status": STATUS,
        "stage": "formal_all_Fay_BRR",
        "decision": "FORMAL_ESTIMATES_COMPLETED",
        "formal_Fay_BRR_inference": True,
        "promotion_permitted": promotion_gate["promotion_permitted"],
        "promotion_gate": promotion_gate,
        "target_population_n": int(len(phase2)),
        "target_deaths_5y": int(phase2[OUTCOME].sum()),
        "models": {name: models[name]["formula"] for name in MODEL_ORDER},
        "primary_estimates": primary_estimates,
        "Fay_BRR_intervals": intervals,
        "outer_fold_signs": primary_result["outer_fold_signs"],
        "Fay_BRR_full_refits_all_four_arms": True,
        "Fay_BRR_replicates": FAY_REPLICATES,
        "Fay_BRR_rho": FAY_RHO,
        "source_selection_freeze_signature": freeze["freeze_signature"],
        "age_selected_domain": EXPECTED_AGE_SELECTED_DOMAIN,
        "age_selected_features": list(EXPECTED_AGE_SELECTED_FEATURES),
        "remaining_panel_label": "remaining_20_after_age_selected_block",
        "model_artifact_audit_status": formal_audit["status"],
        "model_artifact_count": formal_audit["artifact_count"],
        "interpretation": {
            "V_F_positive": bool(primary_estimates["V_F"] > 0),
            "V_R_positive": bool(primary_estimates["V_R"] > 0),
            "P_proxy_positive": bool(primary_estimates["P_proxy"] > 0),
            "V_F_interval_excludes_zero": bool(
                intervals["V_F"]["ci_lower"] > 0
                or intervals["V_F"]["ci_upper"] < 0
            ),
            "P_proxy_interval_excludes_zero": bool(
                intervals["P_proxy"]["ci_lower"] > 0
                or intervals["P_proxy"]["ci_upper"] < 0
            ),
            "non_significance_is_not_equivalence": True,
        },
        "claim_ceiling": (
            "Information-set-dependent incremental predictive value and proxy substitution "
            "under one frozen population, endpoint, mask and estimator family; not causal "
            "mediation, biological-age identity, clinical equivalence, universal measured-state "
            "sufficiency or proof that the remaining-20 lacks age-related information."
        ),
    }
    base.write_json_or_verify(out / "decision.json", formal_result)
    formal_report = f"""# NHANES III chronological-age increment and proxy comparison v0.1

- Status: **{STATUS}**
- Decision: **FORMAL_ESTIMATES_COMPLETED**
- Phase 2 participants/deaths: {len(phase2)} / {int(phase2[OUTCOME].sum())}
- `V_F = L(F0)-L(F1)`: {primary_estimates['V_F']:.9f}
  (95% Fay-BRR interval {intervals['V_F']['ci_lower']:.9f} to {intervals['V_F']['ci_upper']:.9f})
- `V_R = L(R0)-L(R1)`: {primary_estimates['V_R']:.9f}
  (95% Fay-BRR interval {intervals['V_R']['ci_lower']:.9f} to {intervals['V_R']['ci_upper']:.9f})
- `P_proxy = V_R-V_F`: {primary_estimates['P_proxy']:.9f}
  (95% Fay-BRR interval {intervals['P_proxy']['ci_lower']:.9f} to {intervals['P_proxy']['ci_upper']:.9f})
- Persisted/reloaded models: {formal_audit['artifact_count']} / 660

All 32 Fay-BRR weight sets repeated inner tuning, outer fitting and held-out
prediction for F0, F1, R0 and R1. A positive `P_proxy` is an
information-set-dependent predictive substitution pattern only. The
remaining-20 panel is not asserted to be free of age-related information, and
failure to reject a replacement contrast is not evidence of equivalence.
"""
    base.write_text_or_verify(out / "REPORT.md", formal_report)
    write_output_manifest(out)
    print(json.dumps(formal_result, indent=2, ensure_ascii=False))
    return 0


def build_parser() -> argparse.ArgumentParser:
    parser = argparse.ArgumentParser(description=__doc__)
    parser.add_argument(
        "--output-dir",
        default=str(DEFAULT_OUT),
        help="Versioned output directory; basename must equal the frozen analysis version.",
    )
    parser.add_argument(
        "--stage",
        choices=("primary", "all"),
        default="primary",
        help=(
            "Fit and audit the 20-model primary probe, or reuse it and complete "
            "all 640 Fay-BRR models."
        ),
    )
    return parser


def main(argv: Sequence[str] | None = None) -> int:
    return run(build_parser().parse_args(argv))


if __name__ == "__main__":
    raise SystemExit(main())
```
