## Supplementary_Software_1 for "Inferential boundaries of age prediction: why prediction does not establish biological age measurement": analyze_nhanes3_5y_matched_scalar_target_supervision_protocol_transfer_v0_2.py

```
#!/usr/bin/env python
"""NHANES III matched-scalar target-supervision protocol transfer.

This fail-closed implementation follows
``NHANES3_5Y_MATCHED_SCALAR_TARGET_SUPERVISION_PROTOCOL_TRANSFER_SPEC_v0_2_20260827.md``.
It learns two one-dimensional weighted ridge projections from the same Phase 1
background and 25-measurement design, differing only in whether chronological
age or five-year mortality supplies the supervised label.  The projections are
then frozen.  Phase 2 supplies equal target-local downstream calibration and
32 Fay-BRR full downstream refits; Phase 2 outcomes never enter either frozen
projection.

Every tuning fit, final preprocessor, estimator, derived design matrix and
prediction is saved in a signed joblib artifact.  Artifacts are reloaded and
their predictions reproduced elementwise with tolerance 1e-12.  A compatible
rerun reuses artifacts; an incompatible artifact fails closed and is never
overwritten.
"""

from __future__ import annotations

import argparse
import hashlib
import json
import math
import os
import platform
import shutil
import sys
import warnings
from pathlib import Path
from typing import Any, Iterable, Sequence

import joblib
import numpy as np
import pandas as pd
import scipy
import sklearn
from scipy.stats import t as student_t
from sklearn.exceptions import ConvergenceWarning
from sklearn.linear_model import LogisticRegression, Ridge
from sklearn.preprocessing import OneHotEncoder, SplineTransformer, StandardScaler


ROOT = Path(__file__).resolve().parents[1]
SCRIPT = Path(__file__).resolve()
SPEC = (
    ROOT
    / "analysis"
    / "NHANES3_5Y_MATCHED_SCALAR_TARGET_SUPERVISION_PROTOCOL_TRANSFER_SPEC_v0_2_20260827.md"
)
CANONICAL = (
    ROOT
    / "outputs"
    / "nhanes3_phase1_to_phase2_endpoint_vs_age_acquisition_v0_2_final_20260826"
)
DEFAULT_OUT = (
    ROOT
    / "outputs"
    / "nhanes3_5y_matched_scalar_target_supervision_protocol_transfer_v0_2"
)
ENV_POLICY = ROOT / "environment-policy.json"
ENV_MANIFEST = ROOT / "environment.yml"
ENV_LOCK = ROOT / "environment.win-64.lock.txt"

ANALYSIS_VERSION = "nhanes3_5y_matched_scalar_target_supervision_protocol_transfer_v0_2"
SCHEMA_VERSION = "1.0"
STATUS = "retrospective_outcome_accessed_fixed_protocol_transfer_robustness_analysis"
AGE = "age_at_exam_years"
OUTCOME = "death_5y"
BACKGROUND = ("sex", "race_ethnicity")
WEIGHT = "analysis_weight"
BASE_WEIGHT = "mec_weight"
STRATUM = "pseudo_stratum"
PSU = "pseudo_psu"
N_OUTER = 5
N_INNER = 3
ALPHA_GRID = (0.01, 0.1, 1.0, 10.0, 100.0, 1000.0)
C_GRID = (0.001, 0.003, 0.01, 0.03, 0.1, 0.3, 1.0, 3.0, 10.0, 30.0, 100.0, 300.0)
AGE_KNOTS = (20.0, 35.0, 50.0, 65.0, 80.0)
EPSILON = 1e-8
PARITY_TOLERANCE = 1e-12
TIE_TOLERANCE = 1e-12
MODEL_SEED = 20260827
FAY_REPLICATES = 32
FAY_RHO = 0.3
FAY_DF = 23

CANONICAL_FILES = {
    "phase1": "analysis_ready_phase1.tsv.gz",
    "phase2": "analysis_ready_phase2.tsv.gz",
    "manifest": "ordered_5x5_feature_manifest.tsv",
    "phase1_folds": "phase1_participant_folds_and_design.tsv",
    "phase2_folds": "phase2_participant_folds_and_design.tsv",
    "fay_registry": "fay_brr_factor_registry.tsv",
    "fay_factors": "fay_brr_participant_factors.npz",
    "fay_design": "fay_brr_design.json",
}


def configure_stdout() -> None:
    if hasattr(sys.stdout, "reconfigure"):
        sys.stdout.reconfigure(encoding="utf-8")


def require(condition: bool, message: str) -> None:
    if not condition:
        raise RuntimeError(f"STOP_DATA_OR_ARTIFACT_FAIL: {message}")


def json_ready(value: Any) -> Any:
    if isinstance(value, dict):
        return {str(key): json_ready(item) for key, item in value.items()}
    if isinstance(value, (list, tuple)):
        return [json_ready(item) for item in value]
    if isinstance(value, Path):
        return str(value)
    if isinstance(value, np.ndarray):
        return value.tolist()
    if isinstance(value, (np.integer,)):
        return int(value)
    if isinstance(value, (np.floating,)):
        return None if not np.isfinite(value) else float(value)
    if isinstance(value, float):
        return None if not math.isfinite(value) else value
    return value


def stable_hash(payload: Any) -> str:
    encoded = json.dumps(
        json_ready(payload), sort_keys=True, separators=(",", ":"), ensure_ascii=False
    ).encode("utf-8")
    return hashlib.sha256(encoded).hexdigest()


def sha256(path: Path) -> str:
    require(path.is_file(), f"missing file: {path}")
    digest = hashlib.sha256()
    with path.open("rb") as handle:
        for block in iter(lambda: handle.read(1024 * 1024), b""):
            digest.update(block)
    return digest.hexdigest()


def ordered_id_signature(values: Iterable[Any]) -> str:
    return stable_hash([int(value) for value in values])


def maximum_error(left: Any, right: Any) -> float:
    a = np.asarray(left, dtype=float)
    b = np.asarray(right, dtype=float)
    require(a.shape == b.shape, "parity arrays have different shapes")
    return 0.0 if a.size == 0 else float(np.max(np.abs(a - b)))


def normalized_weights(values: Sequence[float]) -> np.ndarray:
    weight = np.asarray(values, dtype=float)
    require(weight.ndim == 1, "fit weights must be one-dimensional")
    require(np.isfinite(weight).all() and (weight > 0).all(), "fit weights must be finite and positive")
    require(float(weight.sum()) > 0, "fit weights sum to zero")
    return weight * (len(weight) / float(weight.sum()))


def weighted_mean(values: Sequence[float], weights: Sequence[float]) -> float:
    value = np.asarray(values, dtype=float)
    weight = np.asarray(weights, dtype=float)
    require(value.shape == weight.shape, "weighted mean shape mismatch")
    require(np.isfinite(value).all() and np.isfinite(weight).all(), "non-finite weighted mean input")
    require((weight >= 0).all() and float(weight.sum()) > 0, "invalid weighted mean weights")
    return float(np.dot(value, weight) / float(weight.sum()))


def weighted_scale(values: Sequence[float], weights: Sequence[float]) -> dict[str, float]:
    value = np.asarray(values, dtype=float)
    mean = weighted_mean(value, weights)
    variance = weighted_mean((value - mean) ** 2, weights)
    require(variance > 0 and np.isfinite(variance), "target has zero or invalid weighted variance")
    return {"mean": mean, "scale": float(math.sqrt(variance))}


def standardize(values: Sequence[float], scaler: dict[str, float]) -> np.ndarray:
    return (np.asarray(values, dtype=float) - float(scaler["mean"])) / float(scaler["scale"])


def unstandardize(values: Sequence[float], scaler: dict[str, float]) -> np.ndarray:
    return np.asarray(values, dtype=float) * float(scaler["scale"]) + float(scaler["mean"])


def squared_loss(observed: np.ndarray, predicted: np.ndarray) -> np.ndarray:
    return (np.asarray(observed, dtype=float) - np.asarray(predicted, dtype=float)) ** 2


def log_loss_vector(observed: np.ndarray, probability: np.ndarray) -> np.ndarray:
    y = np.asarray(observed, dtype=float)
    p = np.clip(np.asarray(probability, dtype=float), EPSILON, 1.0 - EPSILON)
    require(y.shape == p.shape, "log-loss inputs have different shapes")
    return -(y * np.log(p) + (1.0 - y) * np.log(1.0 - p))


def software_record() -> dict[str, str]:
    return {
        "python": platform.python_version(),
        "python_executable": sys.executable,
        "platform": platform.platform(),
        "numpy": np.__version__,
        "pandas": pd.__version__,
        "scipy": scipy.__version__,
        "scikit_learn": sklearn.__version__,
        "joblib": joblib.__version__,
    }


def read_table(path: Path) -> pd.DataFrame:
    require(path.is_file(), f"missing table: {path}")
    suffixes = "".join(path.suffixes).lower()
    if suffixes.endswith(".parquet"):
        return pd.read_parquet(path)
    if suffixes.endswith(".csv") or suffixes.endswith(".csv.gz"):
        return pd.read_csv(path)
    return pd.read_csv(path, sep="\t")


def write_table_or_verify(frame: pd.DataFrame, path: Path) -> None:
    path.parent.mkdir(parents=True, exist_ok=True)
    if path.exists():
        existing = read_table(path)
        require(list(existing.columns) == list(frame.columns), f"cached table columns changed: {path}")
        try:
            pd.testing.assert_frame_equal(
                existing.reset_index(drop=True),
                frame.reset_index(drop=True),
                check_dtype=False,
                check_exact=False,
                rtol=1e-14,
                atol=PARITY_TOLERANCE,
            )
        except AssertionError as error:
            raise RuntimeError(f"STOP_DATA_OR_ARTIFACT_FAIL: cached table changed: {path}") from error
        return
    suffixes = "".join(path.suffixes).lower()
    if suffixes.endswith(".parquet"):
        frame.to_parquet(path, index=False)
    elif suffixes.endswith(".csv") or suffixes.endswith(".csv.gz"):
        frame.to_csv(path, index=False)
    else:
        frame.to_csv(path, sep="\t", index=False)


def write_json_or_verify(path: Path, payload: Any) -> None:
    ready = json_ready(payload)
    path.parent.mkdir(parents=True, exist_ok=True)
    if path.exists():
        existing = json.loads(path.read_text(encoding="utf-8"))
        require(stable_hash(existing) == stable_hash(ready), f"cached JSON changed: {path}")
        return
    path.write_text(json.dumps(ready, indent=2, ensure_ascii=False) + "\n", encoding="utf-8")


def write_text_or_verify(path: Path, text: str) -> None:
    path.parent.mkdir(parents=True, exist_ok=True)
    if path.exists():
        require(path.read_text(encoding="utf-8") == text, f"cached text changed: {path}")
        return
    path.write_text(text, encoding="utf-8")


def copy_file_or_verify(source: Path, target: Path) -> None:
    require(source.is_file(), f"canonical input is missing: {source}")
    target.parent.mkdir(parents=True, exist_ok=True)
    if target.exists():
        require(sha256(source) == sha256(target), f"frozen input copy changed: {target}")
        return
    shutil.copy2(source, target)
    require(sha256(source) == sha256(target), f"input copy parity failed: {target}")


def save_joblib_atomic(payload: Any, path: Path) -> None:
    path.parent.mkdir(parents=True, exist_ok=True)
    temporary = path.with_name(path.name + ".tmp")
    if temporary.exists():
        temporary.unlink()
    joblib.dump(payload, temporary, compress=3)
    os.replace(temporary, path)


def select_grid_value(
    rows: list[dict[str, Any]], key: str, grid: Sequence[float], prefer: str
) -> float:
    table = pd.DataFrame(rows)
    pooled = table.groupby(key, as_index=False).agg(
        numerator=("weighted_loss_numerator", "sum"), denominator=("weight_sum", "sum")
    )
    pooled["weighted_loss"] = pooled["numerator"] / pooled["denominator"]
    minimum = float(pooled["weighted_loss"].min())
    tied = set(pooled.loc[pooled["weighted_loss"].le(minimum + TIE_TOLERANCE), key].astype(float))
    require(prefer in {"small", "large"}, "unknown tuning tie rule")
    ordered = list(grid) if prefer == "small" else list(reversed(grid))
    for candidate in ordered:
        if float(candidate) in tied:
            return float(candidate)
    raise RuntimeError("STOP_DATA_OR_ARTIFACT_FAIL: no tuning candidate survived the fixed tie rule")


class WeightedMedianImputer:
    """Training-local weighted-median imputer with a sklearn-like interface."""

    def __init__(self) -> None:
        self.statistics_: np.ndarray | None = None
        self.n_features_in_: int | None = None

    def fit(self, values: np.ndarray, sample_weight: Sequence[float]) -> "WeightedMedianImputer":
        matrix = np.asarray(values, dtype=float)
        weight = np.asarray(sample_weight, dtype=float)
        require(matrix.ndim == 2 and len(matrix) == len(weight), "invalid imputer inputs")
        require(np.isfinite(weight).all() and (weight > 0).all(), "invalid imputer weights")
        statistics: list[float] = []
        for column in range(matrix.shape[1]):
            observed = np.isfinite(matrix[:, column])
            require(observed.any(), f"feature {column} is all missing in a training fold")
            x = matrix[observed, column]
            w = weight[observed]
            order = np.argsort(x, kind="mergesort")
            x = x[order]
            w = w[order]
            index = int(np.searchsorted(np.cumsum(w), 0.5 * float(w.sum()), side="left"))
            statistics.append(float(x[min(index, len(x) - 1)]))
        self.statistics_ = np.asarray(statistics, dtype=float)
        self.n_features_in_ = int(matrix.shape[1])
        return self

    def transform(self, values: np.ndarray) -> np.ndarray:
        require(self.statistics_ is not None, "imputer has not been fitted")
        matrix = np.asarray(values, dtype=float).copy()
        require(matrix.ndim == 2 and matrix.shape[1] == self.n_features_in_, "imputer feature order changed")
        missing = ~np.isfinite(matrix)
        if missing.any():
            matrix[missing] = np.take(self.statistics_, np.nonzero(missing)[1])
        return matrix


def validate_folds(frame: pd.DataFrame, phase: str) -> None:
    require(set(frame["outer_fold"].astype(int)) == set(range(N_OUTER)), f"{phase} outer folds malformed")
    require(set(frame["full_inner_fold"].astype(int)) == set(range(N_INNER)), f"{phase} full inner folds malformed")
    for outer in range(N_OUTER):
        column = f"inner_fold_outer_{outer}"
        training = frame.loc[frame["outer_fold"].ne(outer)]
        require(set(training[column].astype(int)) == set(range(N_INNER)), f"{phase} {column} malformed")
        test = frame.loc[frame["outer_fold"].eq(outer)]
        require(set(training[OUTCOME].astype(int)) == {0, 1}, f"{phase} outer training fold {outer} lacks an outcome class")
        require(set(test[OUTCOME].astype(int)) == {0, 1}, f"{phase} outer test fold {outer} lacks an outcome class")
        for inner in range(N_INNER):
            inner_train = training.loc[training[column].ne(inner)]
            inner_valid = training.loc[training[column].eq(inner)]
            require(set(inner_train[OUTCOME].astype(int)) == {0, 1}, f"{phase} {column} inner training fold {inner} lacks an outcome class")
            require(set(inner_valid[OUTCOME].astype(int)) == {0, 1}, f"{phase} {column} inner validation fold {inner} lacks an outcome class")
    cluster = frame.groupby([STRATUM, PSU], dropna=False).agg(
        outer_n=("outer_fold", "nunique"), full_inner_n=("full_inner_fold", "nunique")
    )
    require(int(cluster["outer_n"].max()) == 1, f"{phase} outer folds split a PSU")
    require(int(cluster["full_inner_n"].max()) == 1, f"{phase} full inner folds split a PSU")
    for outer in range(N_OUTER):
        column = f"inner_fold_outer_{outer}"
        training = frame.loc[frame["outer_fold"].ne(outer)]
        count = training.groupby([STRATUM, PSU], dropna=False)[column].nunique()
        require(int(count.max()) == 1, f"{phase} {column} splits a PSU")


def load_inputs() -> tuple[pd.DataFrame, pd.DataFrame, pd.DataFrame, list[str], np.ndarray, list[dict[str, Any]]]:
    paths = {key: CANONICAL / name for key, name in CANONICAL_FILES.items()}
    for path in paths.values():
        require(path.is_file(), f"canonical input is missing: {path}")
    phase1 = read_table(paths["phase1"]).sort_values("SEQN", kind="mergesort").reset_index(drop=True)
    phase2 = read_table(paths["phase2"]).sort_values("SEQN", kind="mergesort").reset_index(drop=True)
    manifest = read_table(paths["manifest"]).sort_values("feature_order", kind="mergesort").reset_index(drop=True)
    require(manifest["feature_order"].astype(int).tolist() == list(range(1, 26)), "feature manifest is not ordered 1..25")
    features = manifest["feature"].astype(str).tolist()
    require(len(features) == len(set(features)) == 25, "feature manifest must contain 25 unique measurements")
    required = {
        "row_id", "SEQN", "phase", AGE, *BACKGROUND, BASE_WEIGHT, WEIGHT, STRATUM, PSU,
        "outer_fold", "full_inner_fold", *[f"inner_fold_outer_{fold}" for fold in range(N_OUTER)],
        OUTCOME, *features,
    }
    for phase, frame, expected_n, expected_events in (
        ("phase1", phase1, 7558, 402), ("phase2", phase2, 7774, 392)
    ):
        require(required.issubset(frame.columns), f"{phase} analysis interface lacks required columns")
        require(len(frame) == expected_n, f"{phase} participant count changed")
        require(set(frame["phase"].astype(str)) == {phase}, f"{phase} label changed")
        require(int(frame[OUTCOME].sum()) == expected_events, f"{phase} five-year death count changed")
        require(not frame["SEQN"].duplicated().any(), f"{phase} duplicate participant identity")
        require(frame[AGE].ge(20).all() and frame[AGE].lt(80).all(), f"{phase} age range changed")
        require(set(frame[OUTCOME].astype(int)) == {0, 1}, f"{phase} endpoint support changed")
        require(np.isfinite(frame[WEIGHT]).all() and frame[WEIGHT].gt(0).all(), f"{phase} invalid analysis weights")
        require(maximum_error(frame[WEIGHT], frame[BASE_WEIGHT]) <= PARITY_TOLERANCE, f"{phase} base and analysis weights differ")
        require(frame[STRATUM].nunique() == 23 and frame[[STRATUM, PSU]].drop_duplicates().shape[0] == 46, f"{phase} survey design dimensions changed")
        validate_folds(frame, phase)
    fold_columns = [
        "SEQN", "phase", "outer_fold", "full_inner_fold",
        *[f"inner_fold_outer_{fold}" for fold in range(N_OUTER)],
        STRATUM, PSU, BASE_WEIGHT,
    ]
    for key, frame in (("phase1_folds", phase1), ("phase2_folds", phase2)):
        frozen_folds = read_table(paths[key]).sort_values("SEQN", kind="mergesort").reset_index(drop=True)
        try:
            pd.testing.assert_frame_equal(
                frozen_folds[fold_columns],
                frame[fold_columns],
                check_dtype=False,
                check_exact=False,
                rtol=0.0,
                atol=PARITY_TOLERANCE,
            )
        except AssertionError as error:
            raise RuntimeError(f"STOP_DATA_OR_ARTIFACT_FAIL: {key} differs from analysis interface") from error
    factor_file = np.load(paths["fay_factors"])
    factor_ids = np.asarray(factor_file["SEQN"], dtype="int64")
    factors = np.asarray(factor_file["factors"], dtype=float)
    require(factors.shape == (FAY_REPLICATES, len(factor_ids)), "Fay-BRR factor matrix shape changed")
    require(len(set(factor_ids.tolist())) == len(factor_ids), "duplicate Fay-BRR participant identity")
    require(set(factor_ids.tolist()) == set(phase2["SEQN"].astype(int)), "Fay-BRR identities differ from Phase 2")
    lookup = {int(seqn): index for index, seqn in enumerate(factor_ids)}
    factors = factors[:, [lookup[int(seqn)] for seqn in phase2["SEQN"]]]
    require(np.isfinite(factors).all() and (factors > 0).all(), "invalid Fay-BRR factors")
    require(set(np.round(np.unique(factors), 12).tolist()) == {0.3, 1.7}, "Fay-BRR factors are not the frozen 0.3/1.7 pair")
    registry = read_table(paths["fay_registry"])
    registry_required = {
        "replicate", "stratum_order", STRATUM, PSU, "hadamard_sign", "factor"
    }
    require(registry_required.issubset(registry.columns), "Fay-BRR registry columns changed")
    require(len(registry) == FAY_REPLICATES * 46, "Fay-BRR registry row count changed")
    require(set(registry["replicate"].astype(int)) == set(range(FAY_REPLICATES)), "Fay-BRR registry replicates changed")
    require(set(np.round(registry["factor"].astype(float).unique(), 12)) == {0.3, 1.7}, "Fay-BRR registry factors changed")
    participant_clusters = phase2[[STRATUM, PSU]].astype(str)
    for replicate in range(FAY_REPLICATES):
        block = registry.loc[registry["replicate"].astype(int).eq(replicate)].copy()
        require(len(block) == 46, f"Fay-BRR registry replicate {replicate} does not contain 46 PSU rows")
        block[STRATUM] = block[STRATUM].astype(str)
        block[PSU] = block[PSU].astype(str)
        require(not block[[STRATUM, PSU]].duplicated().any(), f"Fay-BRR registry replicate {replicate} duplicates a PSU")
        factor_lookup = {
            (str(row[STRATUM]), str(row[PSU])): float(row["factor"])
            for _, row in block.iterrows()
        }
        expected = np.asarray(
            [factor_lookup.get((str(row[STRATUM]), str(row[PSU])), np.nan) for _, row in participant_clusters.iterrows()],
            dtype=float,
        )
        require(np.isfinite(expected).all(), f"Fay-BRR registry replicate {replicate} does not cover every Phase 2 PSU")
        require(maximum_error(expected, factors[replicate]) <= PARITY_TOLERANCE, f"Fay-BRR registry/factor mismatch in replicate {replicate}")
    fay_design = json.loads(paths["fay_design"].read_text(encoding="utf-8"))
    require(int(fay_design.get("replicates", -1)) == FAY_REPLICATES, "Fay-BRR replicate count changed")
    require(abs(float(fay_design.get("rho", -1)) - FAY_RHO) <= PARITY_TOLERANCE, "Fay-BRR rho changed")
    require(int(fay_design.get("design_degrees_of_freedom", -1)) == FAY_DF, "Fay-BRR degrees of freedom changed")
    require(abs(float(fay_design.get("low_factor", -1)) - 0.3) <= PARITY_TOLERANCE, "Fay-BRR low factor changed")
    require(abs(float(fay_design.get("high_factor", -1)) - 1.7) <= PARITY_TOLERANCE, "Fay-BRR high factor changed")
    require(int(fay_design.get("Hadamard_order", -1)) == 32, "Fay-BRR Hadamard order changed")
    input_records = [
        {"role": key, "path": str(path.resolve()), "sha256": sha256(path), "size_bytes": path.stat().st_size}
        for key, path in paths.items()
    ]
    for role, path in (
        ("frozen_protocol", SPEC), ("analysis_script", SCRIPT), ("environment_policy", ENV_POLICY),
        ("environment_manifest", ENV_MANIFEST), ("environment_lock", ENV_LOCK),
    ):
        input_records.append({"role": role, "path": str(path.resolve()), "sha256": sha256(path), "size_bytes": path.stat().st_size})
    return phase1, phase2, manifest, features, factors, input_records


def fit_projection_preprocessor(
    frame: pd.DataFrame, features: list[str], weight: np.ndarray
) -> tuple[dict[str, Any], np.ndarray]:
    fit_weight = normalized_weights(weight)
    encoder = OneHotEncoder(handle_unknown="ignore", sparse_output=False)
    categorical = encoder.fit_transform(frame.loc[:, list(BACKGROUND)].astype(str))
    blocks = [categorical]
    names = [f"W__{name}" for name in encoder.get_feature_names_out(BACKGROUND)]
    imputer: WeightedMedianImputer | None = None
    scaler: StandardScaler | None = None
    if features:
        raw = frame[features].to_numpy(float)
        imputer = WeightedMedianImputer().fit(raw, sample_weight=fit_weight)
        imputed = imputer.transform(raw)
        scaler = StandardScaler().fit(imputed, sample_weight=fit_weight)
        measured = scaler.transform(imputed)
        blocks.append(measured)
        names.extend(f"X__{feature}" for feature in features)
    design = np.column_stack(blocks)
    require(np.isfinite(design).all(), "projection design contains non-finite values")
    return {
        "background": list(BACKGROUND),
        "features": list(features),
        "encoder": encoder,
        "imputer": imputer,
        "measurement_scaler": scaler,
        "design_feature_names": names,
        "fit_weight_normalization": "sum_to_n_within_fit",
    }, design


def transform_projection_preprocessor(preprocessor: dict[str, Any], frame: pd.DataFrame) -> np.ndarray:
    categorical = preprocessor["encoder"].transform(
        frame.loc[:, list(preprocessor["background"])].astype(str)
    )
    blocks = [categorical]
    features = list(preprocessor["features"])
    if features:
        measured = preprocessor["imputer"].transform(frame[features].to_numpy(float))
        blocks.append(preprocessor["measurement_scaler"].transform(measured))
    design = np.column_stack(blocks)
    require(np.isfinite(design).all(), "transformed projection design contains non-finite values")
    return design


def projection_targets(names: Sequence[str]) -> dict[str, str]:
    mapping = {"q_A": AGE, "q_Y": OUTCOME, "q_W": AGE}
    require(set(names).issubset(mapping), "unknown projection target")
    return {name: mapping[name] for name in names}


def fit_projection_subbundle(
    training: pd.DataFrame,
    holdout: pd.DataFrame,
    features: list[str],
    target_names: Sequence[str],
    inner_column: str,
    scope: str,
) -> dict[str, Any]:
    target_columns = projection_targets(target_names)
    tuning_rows: list[dict[str, Any]] = []
    inner_folds: list[dict[str, Any]] = []
    for inner_fold in range(N_INNER):
        inner_train = training.loc[training[inner_column].ne(inner_fold)].sort_values(
            "SEQN", kind="mergesort"
        ).reset_index(drop=True)
        inner_valid = training.loc[training[inner_column].eq(inner_fold)].sort_values(
            "SEQN", kind="mergesort"
        ).reset_index(drop=True)
        require(len(inner_train) > 0 and len(inner_valid) > 0, f"empty projection inner fold in {scope}")
        train_weight = inner_train[WEIGHT].to_numpy(float)
        valid_weight = inner_valid[WEIGHT].to_numpy(float)
        preprocessor, x_train = fit_projection_preprocessor(inner_train, features, train_weight)
        x_valid = transform_projection_preprocessor(preprocessor, inner_valid)
        target_fits: dict[str, Any] = {}
        for target_name, column in target_columns.items():
            target_scaler = weighted_scale(inner_train[column].to_numpy(float), train_weight)
            y_train = standardize(inner_train[column], target_scaler)
            y_valid = standardize(inner_valid[column], target_scaler)
            candidate_fits: list[dict[str, Any]] = []
            for alpha in ALPHA_GRID:
                estimator = Ridge(alpha=float(alpha), fit_intercept=True)
                estimator.fit(x_train, y_train, sample_weight=normalized_weights(train_weight))
                valid_score = np.asarray(estimator.predict(x_valid), dtype=float)
                losses = squared_loss(y_valid, valid_score)
                numerator = float(np.dot(valid_weight, losses))
                denominator = float(valid_weight.sum())
                candidate_fits.append(
                    {
                        "alpha": float(alpha),
                        "estimator": estimator,
                        "valid_score": valid_score,
                        "valid_loss_vector": losses,
                        "weighted_loss_numerator": numerator,
                        "weight_sum": denominator,
                    }
                )
                tuning_rows.append(
                    {
                        "scope": scope,
                        "target": target_name,
                        "inner_fold": int(inner_fold),
                        "alpha": float(alpha),
                        "n_valid": int(len(inner_valid)),
                        "weighted_loss_numerator": numerator,
                        "weight_sum": denominator,
                        "weighted_loss": numerator / denominator,
                    }
                )
            target_fits[target_name] = {
                "target_column": column,
                "target_scaler": target_scaler,
                "train_target_standardized": y_train,
                "valid_target_standardized": y_valid,
                "candidate_fits": candidate_fits,
            }
        inner_folds.append(
            {
                "inner_fold": int(inner_fold),
                "train_seqn": inner_train["SEQN"].to_numpy("int64"),
                "valid_seqn": inner_valid["SEQN"].to_numpy("int64"),
                "train_weight": train_weight,
                "valid_weight": valid_weight,
                "preprocessor": preprocessor,
                "x_train": x_train,
                "x_valid": x_valid,
                "targets": target_fits,
            }
        )

    selected_alpha: dict[str, float] = {}
    for target_name in target_columns:
        rows = [row for row in tuning_rows if row["target"] == target_name]
        selected_alpha[target_name] = select_grid_value(rows, "alpha", ALPHA_GRID, prefer="large")

    final_weight = training[WEIGHT].to_numpy(float)
    final_preprocessor, x_training = fit_projection_preprocessor(training, features, final_weight)
    x_holdout = transform_projection_preprocessor(final_preprocessor, holdout)
    final_targets: dict[str, Any] = {}
    for target_name, column in target_columns.items():
        target_scaler = weighted_scale(training[column].to_numpy(float), final_weight)
        y_training = standardize(training[column], target_scaler)
        estimator = Ridge(alpha=selected_alpha[target_name], fit_intercept=True)
        estimator.fit(x_training, y_training, sample_weight=normalized_weights(final_weight))
        training_score = np.asarray(estimator.predict(x_training), dtype=float)
        holdout_score = np.asarray(estimator.predict(x_holdout), dtype=float)
        final_targets[target_name] = {
            "target_column": column,
            "target_scaler": target_scaler,
            "selected_alpha": selected_alpha[target_name],
            "estimator": estimator,
            "training_score": training_score,
            "holdout_score": holdout_score,
        }
    return {
        "scope": scope,
        "features": list(features),
        "target_names": list(target_names),
        "inner_column": inner_column,
        "training_seqn": training["SEQN"].to_numpy("int64"),
        "holdout_seqn": holdout["SEQN"].to_numpy("int64"),
        "inner_folds": inner_folds,
        "tuning_rows": tuning_rows,
        "selected_alpha": selected_alpha,
        "final_preprocessor": final_preprocessor,
        "x_training": x_training,
        "x_holdout": x_holdout,
        "final_targets": final_targets,
    }


def verify_projection_subbundle(
    bundle: dict[str, Any], training: pd.DataFrame, holdout: pd.DataFrame
) -> dict[str, float]:
    features = list(bundle["features"])
    inner_column = str(bundle["inner_column"])
    require(
        np.array_equal(bundle["training_seqn"], training["SEQN"].to_numpy("int64")),
        "projection training identities changed",
    )
    require(
        np.array_equal(bundle["holdout_seqn"], holdout["SEQN"].to_numpy("int64")),
        "projection holdout identities changed",
    )
    by_seqn = training.set_index("SEQN", drop=False)
    max_design_error = 0.0
    max_prediction_error = 0.0
    reconstructed_rows: list[dict[str, Any]] = []
    for stored in bundle["inner_folds"]:
        inner_fold = int(stored["inner_fold"])
        train_seqn = np.asarray(stored["train_seqn"], dtype="int64")
        valid_seqn = np.asarray(stored["valid_seqn"], dtype="int64")
        expected_train = training.loc[training[inner_column].ne(inner_fold), "SEQN"].to_numpy("int64")
        expected_valid = training.loc[training[inner_column].eq(inner_fold), "SEQN"].to_numpy("int64")
        require(np.array_equal(train_seqn, expected_train), "projection inner training identities changed")
        require(np.array_equal(valid_seqn, expected_valid), "projection inner validation identities changed")
        require(np.intersect1d(train_seqn, valid_seqn).size == 0, "projection tuning identity leakage")
        inner_train = by_seqn.loc[train_seqn].reset_index(drop=True)
        inner_valid = by_seqn.loc[valid_seqn].reset_index(drop=True)
        x_train = transform_projection_preprocessor(stored["preprocessor"], inner_train)
        x_valid = transform_projection_preprocessor(stored["preprocessor"], inner_valid)
        max_design_error = max(
            max_design_error,
            maximum_error(x_train, stored["x_train"]),
            maximum_error(x_valid, stored["x_valid"]),
        )
        for target_name, target in stored["targets"].items():
            y_valid = standardize(inner_valid[target["target_column"]], target["target_scaler"])
            max_design_error = max(max_design_error, maximum_error(y_valid, target["valid_target_standardized"]))
            valid_weight = inner_valid[WEIGHT].to_numpy(float)
            for candidate in target["candidate_fits"]:
                prediction = np.asarray(candidate["estimator"].predict(x_valid), dtype=float)
                losses = squared_loss(y_valid, prediction)
                max_prediction_error = max(
                    max_prediction_error,
                    maximum_error(prediction, candidate["valid_score"]),
                    maximum_error(losses, candidate["valid_loss_vector"]),
                )
                numerator = float(np.dot(valid_weight, losses))
                denominator = float(valid_weight.sum())
                require(abs(numerator - float(candidate["weighted_loss_numerator"])) <= PARITY_TOLERANCE, "projection tuning loss changed")
                reconstructed_rows.append(
                    {
                        "target": target_name,
                        "alpha": float(candidate["alpha"]),
                        "weighted_loss_numerator": numerator,
                        "weight_sum": denominator,
                    }
                )
    for target_name in bundle["target_names"]:
        rows = [row for row in reconstructed_rows if row["target"] == target_name]
        expected = select_grid_value(rows, "alpha", ALPHA_GRID, prefer="large")
        require(expected == float(bundle["selected_alpha"][target_name]), "selected projection alpha changed")

    x_training = transform_projection_preprocessor(bundle["final_preprocessor"], training)
    x_holdout = transform_projection_preprocessor(bundle["final_preprocessor"], holdout)
    max_design_error = max(
        max_design_error,
        maximum_error(x_training, bundle["x_training"]),
        maximum_error(x_holdout, bundle["x_holdout"]),
    )
    for target in bundle["final_targets"].values():
        train_score = np.asarray(target["estimator"].predict(x_training), dtype=float)
        holdout_score = np.asarray(target["estimator"].predict(x_holdout), dtype=float)
        max_prediction_error = max(
            max_prediction_error,
            maximum_error(train_score, target["training_score"]),
            maximum_error(holdout_score, target["holdout_score"]),
        )
    require(max(max_design_error, max_prediction_error) <= PARITY_TOLERANCE, "projection reload parity failed")
    return {
        "max_design_error": max_design_error,
        "max_prediction_error": max_prediction_error,
    }


def projection_artifact_signature(
    analysis_signature: str,
    kind: str,
    training: pd.DataFrame,
    holdout: pd.DataFrame,
    features: list[str],
    inner_column: str,
) -> str:
    return stable_hash(
        {
            "analysis_signature": analysis_signature,
            "artifact_kind": kind,
            "training_seqn": training["SEQN"].astype(int).tolist(),
            "holdout_seqn": holdout["SEQN"].astype(int).tolist(),
            "training_weight": training[WEIGHT].astype(float).tolist(),
            "inner_assignments": training[["SEQN", inner_column]].astype(
                {"SEQN": int, inner_column: int}
            ).to_dict(orient="records"),
            "features": features,
            "background": BACKGROUND,
            "targets": {"q_A": AGE, "q_Y": OUTCOME, "q_W": AGE},
            "target_standardization": "training_fold_MEC_weighted_mean_and_population_sd",
            "alpha_grid": ALPHA_GRID,
            "loss": "MEC_weighted_squared_error_on_training_fold_standardized_target",
            "tie_rule": "within_1e-12_choose_larger_alpha",
            "fit_weight_normalization": "sum_to_n_within_fit",
            "model_seed": MODEL_SEED,
            "parity_tolerance": PARITY_TOLERANCE,
        }
    )


def fit_projection_artifact(
    analysis_signature: str,
    kind: str,
    training: pd.DataFrame,
    holdout: pd.DataFrame,
    features: list[str],
    inner_column: str,
    include_w_only: bool,
) -> dict[str, Any]:
    artifact_signature = projection_artifact_signature(
        analysis_signature, kind, training, holdout, features, inner_column
    )
    matched = fit_projection_subbundle(
        training, holdout, features, ("q_A", "q_Y"), inner_column, f"{kind}__matched"
    )
    w_only = None
    if include_w_only:
        w_only = fit_projection_subbundle(
            training, holdout, [], ("q_W",), inner_column, f"{kind}__w_only_age"
        )
    return {
        "schema_version": SCHEMA_VERSION,
        "analysis_version": ANALYSIS_VERSION,
        "analysis_signature": analysis_signature,
        "artifact_signature": artifact_signature,
        "artifact_kind": kind,
        "matched": matched,
        "w_only": w_only,
        "software": software_record(),
    }


def verify_projection_artifact(
    artifact: dict[str, Any],
    analysis_signature: str,
    kind: str,
    training: pd.DataFrame,
    holdout: pd.DataFrame,
    features: list[str],
    inner_column: str,
    include_w_only: bool,
) -> dict[str, Any]:
    expected = projection_artifact_signature(
        analysis_signature, kind, training, holdout, features, inner_column
    )
    require(artifact.get("artifact_signature") == expected, "projection artifact signature mismatch")
    matched = verify_projection_subbundle(artifact["matched"], training, holdout)
    w_only: dict[str, float] | None = None
    if include_w_only:
        require(artifact.get("w_only") is not None, "W-only age gate artifact is missing")
        w_only = verify_projection_subbundle(artifact["w_only"], training, holdout)
    else:
        require(artifact.get("w_only") is None, "unexpected W-only branch in final projection artifact")
    return {"matched": matched, "w_only": w_only}


def fit_or_load_projection_artifact(
    path: Path,
    analysis_signature: str,
    kind: str,
    training: pd.DataFrame,
    holdout: pd.DataFrame,
    features: list[str],
    inner_column: str,
    include_w_only: bool,
) -> tuple[dict[str, Any], dict[str, Any]]:
    newly_fitted = False
    if path.exists():
        artifact = joblib.load(path)
        expected = projection_artifact_signature(
            analysis_signature, kind, training, holdout, features, inner_column
        )
        require(artifact.get("artifact_signature") == expected, f"incompatible cached artifact: {path}")
    else:
        artifact = fit_projection_artifact(
            analysis_signature, kind, training, holdout, features, inner_column, include_w_only
        )
        save_joblib_atomic(artifact, path)
        newly_fitted = True
    reloaded = joblib.load(path)
    parity = verify_projection_artifact(
        reloaded,
        analysis_signature,
        kind,
        training,
        holdout,
        features,
        inner_column,
        include_w_only,
    )
    return reloaded, {
        "stage": "source_projection",
        "artifact_kind": kind,
        "artifact_path": str(path.resolve()),
        "artifact_sha256": sha256(path),
        "cache_action": "fitted" if newly_fitted else "reused",
        "max_design_error": max(
            parity["matched"]["max_design_error"],
            0.0 if parity["w_only"] is None else parity["w_only"]["max_design_error"],
        ),
        "max_prediction_error": max(
            parity["matched"]["max_prediction_error"],
            0.0 if parity["w_only"] is None else parity["w_only"]["max_prediction_error"],
        ),
        "projection_tuning_estimator_count": int(
            sum(len(target["candidate_fits"]) for fold in reloaded["matched"]["inner_folds"] for target in fold["targets"].values())
            + (
                0
                if reloaded["w_only"] is None
                else sum(len(target["candidate_fits"]) for fold in reloaded["w_only"]["inner_folds"] for target in fold["targets"].values())
            )
        ),
        "projection_final_estimator_count": int(
            len(reloaded["matched"]["final_targets"])
            + (0 if reloaded["w_only"] is None else len(reloaded["w_only"]["final_targets"]))
        ),
    }


DOWNSTREAM_MODELS = {
    "M0": {"score_column": None, "score_mode": "none"},
    "M_A": {"score_column": "q_A", "score_mode": "spline"},
    "M_Y": {"score_column": "q_Y", "score_mode": "spline"},
    "M_A_LINEAR": {"score_column": "q_A", "score_mode": "linear"},
    "M_Y_LINEAR": {"score_column": "q_Y", "score_mode": "linear"},
}


def fit_downstream_preprocessor(
    frame: pd.DataFrame,
    score: np.ndarray | None,
    score_mode: str,
    weight: np.ndarray,
) -> tuple[dict[str, Any], np.ndarray]:
    fit_weight = normalized_weights(weight)
    encoder = OneHotEncoder(handle_unknown="ignore", sparse_output=False)
    categorical = encoder.fit_transform(frame.loc[:, list(BACKGROUND)].astype(str))

    age_spline = SplineTransformer(
        degree=3,
        knots=np.asarray(AGE_KNOTS, dtype=float).reshape(-1, 1),
        include_bias=False,
        extrapolation="linear",
    )
    age_raw = age_spline.fit_transform(frame[[AGE]].to_numpy(float), sample_weight=fit_weight)
    age_scaler = StandardScaler().fit(age_raw, sample_weight=fit_weight)
    age_design = age_scaler.transform(age_raw)
    blocks = [categorical, age_design]
    names = (
        [f"W__{name}" for name in encoder.get_feature_names_out(BACKGROUND)]
        + [f"A_spline_{index:02d}" for index in range(age_design.shape[1])]
    )

    score_scaler: StandardScaler | None = None
    score_spline: SplineTransformer | None = None
    if score_mode != "none":
        require(score is not None, "downstream scalar is missing")
        score_values = np.asarray(score, dtype=float).reshape(-1, 1)
        require(len(score_values) == len(frame) and np.isfinite(score_values).all(), "invalid downstream scalar")
        score_scaler = StandardScaler().fit(score_values, sample_weight=fit_weight)
        score_scaled = score_scaler.transform(score_values)
        if score_mode == "linear":
            score_design = score_scaled
            score_names = ["scalar_linear"]
        elif score_mode == "spline":
            score_spline = SplineTransformer(
                n_knots=5,
                knots="uniform",
                degree=3,
                include_bias=False,
                extrapolation="linear",
            )
            score_design = score_spline.fit_transform(score_scaled, sample_weight=fit_weight)
            score_names = [f"scalar_spline_{index:02d}" for index in range(score_design.shape[1])]
        else:
            raise RuntimeError(f"STOP_DATA_OR_ARTIFACT_FAIL: unknown downstream score mode {score_mode}")
        blocks.append(score_design)
        names.extend(score_names)
    design = np.column_stack(blocks)
    require(np.isfinite(design).all(), "downstream design contains non-finite values")
    return {
        "background": list(BACKGROUND),
        "score_mode": score_mode,
        "encoder": encoder,
        "age_spline": age_spline,
        "age_scaler": age_scaler,
        "score_scaler": score_scaler,
        "score_spline": score_spline,
        "design_feature_names": names,
        "age_knots": list(AGE_KNOTS),
        "score_knots": "uniform_5_knots" if score_mode == "spline" else None,
    }, design


def transform_downstream_preprocessor(
    preprocessor: dict[str, Any], frame: pd.DataFrame, score: np.ndarray | None
) -> np.ndarray:
    categorical = preprocessor["encoder"].transform(
        frame.loc[:, list(preprocessor["background"])].astype(str)
    )
    age_raw = preprocessor["age_spline"].transform(frame[[AGE]].to_numpy(float))
    blocks = [categorical, preprocessor["age_scaler"].transform(age_raw)]
    mode = str(preprocessor["score_mode"])
    if mode != "none":
        require(score is not None, "stored downstream scalar is missing")
        score_scaled = preprocessor["score_scaler"].transform(
            np.asarray(score, dtype=float).reshape(-1, 1)
        )
        if mode == "linear":
            score_design = score_scaled
        elif mode == "spline":
            score_design = preprocessor["score_spline"].transform(score_scaled)
        else:
            raise RuntimeError("STOP_DATA_OR_ARTIFACT_FAIL: stored downstream mode is invalid")
        blocks.append(score_design)
    design = np.column_stack(blocks)
    require(np.isfinite(design).all(), "transformed downstream design contains non-finite values")
    return design


def fit_logistic(
    design: np.ndarray, outcome: np.ndarray, weight: np.ndarray, c_value: float
) -> LogisticRegression:
    require(set(np.asarray(outcome, dtype=int).tolist()) == {0, 1}, "downstream fit lacks an outcome class")
    estimator = LogisticRegression(
        C=float(c_value),
        penalty="l2",
        solver="lbfgs",
        max_iter=5000,
        tol=1e-10,
        random_state=MODEL_SEED,
    )
    with warnings.catch_warnings(record=True) as caught:
        warnings.simplefilter("always", ConvergenceWarning)
        warnings.filterwarnings("ignore", category=FutureWarning, message=".*penalty.*deprecated.*")
        estimator.fit(design, np.asarray(outcome, dtype=int), sample_weight=normalized_weights(weight))
    require(
        not any(issubclass(item.category, ConvergenceWarning) for item in caught),
        "weighted logistic estimator did not converge",
    )
    require(int(np.max(estimator.n_iter_)) < int(estimator.max_iter), "weighted logistic estimator reached max_iter")
    return estimator


def predict_logistic(estimator: LogisticRegression, design: np.ndarray) -> np.ndarray:
    return np.clip(np.asarray(estimator.predict_proba(design)[:, 1], dtype=float), EPSILON, 1.0 - EPSILON)


def downstream_artifact_signature(
    stage_signature: str,
    model_name: str,
    outer_fold: int,
    training: pd.DataFrame,
    test: pd.DataFrame,
    score_column: str | None,
    score_mode: str,
) -> str:
    inner_column = f"inner_fold_outer_{outer_fold}"
    payload: dict[str, Any] = {
        "stage_signature": stage_signature,
        "artifact_kind": "target_local_weighted_downstream_calibration",
        "model_name": model_name,
        "score_column": score_column,
        "score_mode": score_mode,
        "training_seqn": training["SEQN"].astype(int).tolist(),
        "test_seqn": test["SEQN"].astype(int).tolist(),
        "training_weight": training[WEIGHT].astype(float).tolist(),
        "test_weight": test[WEIGHT].astype(float).tolist(),
        "inner_assignments": training[["SEQN", inner_column]].astype(
            {"SEQN": int, inner_column: int}
        ).to_dict(orient="records"),
        "background": BACKGROUND,
        "fixed_age_knots": AGE_KNOTS,
        "score_spline": "uniform_5_knots_degree3_linear_extrapolation",
        "C_grid": C_GRID,
        "inner_loss_pooling": "sum_weighted_numerators_over_sum_weight_denominators",
        "tie_rule": "within_1e-12_choose_smaller_C",
        "fit_weight_normalization": "sum_to_n_within_fit",
        "log_loss_clip": EPSILON,
        "model_seed": MODEL_SEED,
        "parity_tolerance": PARITY_TOLERANCE,
    }
    if score_column is not None:
        payload["training_score"] = training[score_column].astype(float).tolist()
        payload["test_score"] = test[score_column].astype(float).tolist()
    return stable_hash(payload)


def fit_downstream_artifact(
    analysis_signature: str,
    stage_signature: str,
    model_name: str,
    outer_fold: int,
    frame: pd.DataFrame,
) -> dict[str, Any]:
    specification = DOWNSTREAM_MODELS[model_name]
    score_column = specification["score_column"]
    score_mode = specification["score_mode"]
    training = frame.loc[frame["outer_fold"].ne(outer_fold)].sort_values(
        "SEQN", kind="mergesort"
    ).reset_index(drop=True)
    test = frame.loc[frame["outer_fold"].eq(outer_fold)].sort_values(
        "SEQN", kind="mergesort"
    ).reset_index(drop=True)
    inner_column = f"inner_fold_outer_{outer_fold}"
    inner_folds: list[dict[str, Any]] = []
    tuning_rows: list[dict[str, Any]] = []
    for inner_fold in range(N_INNER):
        inner_train = training.loc[training[inner_column].ne(inner_fold)].sort_values(
            "SEQN", kind="mergesort"
        ).reset_index(drop=True)
        inner_valid = training.loc[training[inner_column].eq(inner_fold)].sort_values(
            "SEQN", kind="mergesort"
        ).reset_index(drop=True)
        require(len(inner_train) > 0 and len(inner_valid) > 0, "empty downstream inner fold")
        train_weight = inner_train[WEIGHT].to_numpy(float)
        valid_weight = inner_valid[WEIGHT].to_numpy(float)
        train_score = None if score_column is None else inner_train[score_column].to_numpy(float)
        valid_score = None if score_column is None else inner_valid[score_column].to_numpy(float)
        preprocessor, x_train = fit_downstream_preprocessor(
            inner_train, train_score, score_mode, train_weight
        )
        x_valid = transform_downstream_preprocessor(preprocessor, inner_valid, valid_score)
        y_train = inner_train[OUTCOME].to_numpy(int)
        y_valid = inner_valid[OUTCOME].to_numpy(int)
        candidate_fits: list[dict[str, Any]] = []
        for c_value in C_GRID:
            estimator = fit_logistic(x_train, y_train, train_weight, c_value)
            probability = predict_logistic(estimator, x_valid)
            losses = log_loss_vector(y_valid, probability)
            numerator = float(np.dot(valid_weight, losses))
            denominator = float(valid_weight.sum())
            candidate_fits.append(
                {
                    "C": float(c_value),
                    "estimator": estimator,
                    "valid_probability": probability,
                    "valid_loss_vector": losses,
                    "weighted_loss_numerator": numerator,
                    "weight_sum": denominator,
                }
            )
            tuning_rows.append(
                {
                    "model": model_name,
                    "outer_fold": int(outer_fold),
                    "inner_fold": int(inner_fold),
                    "C": float(c_value),
                    "n_valid": int(len(inner_valid)),
                    "events_valid": int(y_valid.sum()),
                    "weighted_loss_numerator": numerator,
                    "weight_sum": denominator,
                    "weighted_loss": numerator / denominator,
                }
            )
        inner_folds.append(
            {
                "inner_fold": int(inner_fold),
                "train_seqn": inner_train["SEQN"].to_numpy("int64"),
                "valid_seqn": inner_valid["SEQN"].to_numpy("int64"),
                "train_weight": train_weight,
                "valid_weight": valid_weight,
                "train_score": train_score,
                "valid_score": valid_score,
                "preprocessor": preprocessor,
                "x_train": x_train,
                "x_valid": x_valid,
                "train_outcome": y_train,
                "valid_outcome": y_valid,
                "candidate_fits": candidate_fits,
            }
        )
    selected_c = select_grid_value(tuning_rows, "C", C_GRID, prefer="small")
    final_weight = training[WEIGHT].to_numpy(float)
    final_train_score = None if score_column is None else training[score_column].to_numpy(float)
    final_test_score = None if score_column is None else test[score_column].to_numpy(float)
    final_preprocessor, x_training = fit_downstream_preprocessor(
        training, final_train_score, score_mode, final_weight
    )
    x_test = transform_downstream_preprocessor(final_preprocessor, test, final_test_score)
    final_estimator = fit_logistic(
        x_training, training[OUTCOME].to_numpy(int), final_weight, selected_c
    )
    test_probability = predict_logistic(final_estimator, x_test)
    test_losses = log_loss_vector(test[OUTCOME].to_numpy(int), test_probability)
    signature = downstream_artifact_signature(
        stage_signature,
        model_name,
        outer_fold,
        training,
        test,
        score_column,
        score_mode,
    )
    return {
        "schema_version": SCHEMA_VERSION,
        "analysis_version": ANALYSIS_VERSION,
        "analysis_signature": analysis_signature,
        "stage_signature": stage_signature,
        "artifact_signature": signature,
        "artifact_kind": "target_local_weighted_downstream_calibration",
        "model_name": model_name,
        "score_column": score_column,
        "score_mode": score_mode,
        "outer_fold": int(outer_fold),
        "inner_column": inner_column,
        "training_seqn": training["SEQN"].to_numpy("int64"),
        "test_seqn": test["SEQN"].to_numpy("int64"),
        "training_weight": final_weight,
        "test_weight": test[WEIGHT].to_numpy(float),
        "inner_folds": inner_folds,
        "tuning_rows": tuning_rows,
        "selected_C": selected_c,
        "final_train_score": final_train_score,
        "final_test_score": final_test_score,
        "final_preprocessor": final_preprocessor,
        "x_training": x_training,
        "x_test": x_test,
        "final_estimator": final_estimator,
        "test_probability": test_probability,
        "test_loss_vector": test_losses,
        "software": software_record(),
    }


def verify_downstream_artifact(
    artifact: dict[str, Any],
    analysis_signature: str,
    stage_signature: str,
    model_name: str,
    outer_fold: int,
    frame: pd.DataFrame,
) -> dict[str, float]:
    specification = DOWNSTREAM_MODELS[model_name]
    score_column = specification["score_column"]
    score_mode = specification["score_mode"]
    training = frame.loc[frame["outer_fold"].ne(outer_fold)].sort_values(
        "SEQN", kind="mergesort"
    ).reset_index(drop=True)
    test = frame.loc[frame["outer_fold"].eq(outer_fold)].sort_values(
        "SEQN", kind="mergesort"
    ).reset_index(drop=True)
    expected = downstream_artifact_signature(
        stage_signature, model_name, outer_fold, training, test, score_column, score_mode
    )
    require(artifact.get("artifact_signature") == expected, "downstream artifact signature mismatch")
    require(artifact.get("analysis_signature") == analysis_signature, "downstream parent analysis signature changed")
    require(artifact.get("model_name") == model_name, "downstream model identity changed")
    require(np.array_equal(artifact["training_seqn"], training["SEQN"].to_numpy("int64")), "downstream training identities changed")
    require(np.array_equal(artifact["test_seqn"], test["SEQN"].to_numpy("int64")), "downstream test identities changed")
    by_seqn = frame.set_index("SEQN", drop=False)
    reconstructed_rows: list[dict[str, Any]] = []
    max_design_error = 0.0
    max_prediction_error = 0.0
    for stored in artifact["inner_folds"]:
        inner_fold = int(stored["inner_fold"])
        train_seqn = np.asarray(stored["train_seqn"], dtype="int64")
        valid_seqn = np.asarray(stored["valid_seqn"], dtype="int64")
        expected_train = training.loc[training[artifact["inner_column"]].ne(inner_fold), "SEQN"].to_numpy("int64")
        expected_valid = training.loc[training[artifact["inner_column"]].eq(inner_fold), "SEQN"].to_numpy("int64")
        require(np.array_equal(train_seqn, expected_train), "downstream inner training identities changed")
        require(np.array_equal(valid_seqn, expected_valid), "downstream inner validation identities changed")
        require(np.intersect1d(train_seqn, valid_seqn).size == 0, "downstream tuning identity leakage")
        inner_train = by_seqn.loc[train_seqn].reset_index(drop=True)
        inner_valid = by_seqn.loc[valid_seqn].reset_index(drop=True)
        train_score = None if score_column is None else inner_train[score_column].to_numpy(float)
        valid_score = None if score_column is None else inner_valid[score_column].to_numpy(float)
        x_train = transform_downstream_preprocessor(stored["preprocessor"], inner_train, train_score)
        x_valid = transform_downstream_preprocessor(stored["preprocessor"], inner_valid, valid_score)
        max_design_error = max(
            max_design_error,
            maximum_error(x_train, stored["x_train"]),
            maximum_error(x_valid, stored["x_valid"]),
        )
        y_valid = inner_valid[OUTCOME].to_numpy(int)
        valid_weight = inner_valid[WEIGHT].to_numpy(float)
        for candidate in stored["candidate_fits"]:
            probability = predict_logistic(candidate["estimator"], x_valid)
            losses = log_loss_vector(y_valid, probability)
            max_prediction_error = max(
                max_prediction_error,
                maximum_error(probability, candidate["valid_probability"]),
                maximum_error(losses, candidate["valid_loss_vector"]),
            )
            numerator = float(np.dot(valid_weight, losses))
            require(abs(numerator - float(candidate["weighted_loss_numerator"])) <= PARITY_TOLERANCE, "downstream tuning loss changed")
            reconstructed_rows.append(
                {
                    "C": float(candidate["C"]),
                    "weighted_loss_numerator": numerator,
                    "weight_sum": float(valid_weight.sum()),
                }
            )
    selected_c = select_grid_value(reconstructed_rows, "C", C_GRID, prefer="small")
    require(selected_c == float(artifact["selected_C"]), "selected downstream C changed")
    final_train_score = None if score_column is None else training[score_column].to_numpy(float)
    final_test_score = None if score_column is None else test[score_column].to_numpy(float)
    x_training = transform_downstream_preprocessor(
        artifact["final_preprocessor"], training, final_train_score
    )
    x_test = transform_downstream_preprocessor(artifact["final_preprocessor"], test, final_test_score)
    probability = predict_logistic(artifact["final_estimator"], x_test)
    losses = log_loss_vector(test[OUTCOME].to_numpy(int), probability)
    max_design_error = max(
        max_design_error,
        maximum_error(x_training, artifact["x_training"]),
        maximum_error(x_test, artifact["x_test"]),
    )
    max_prediction_error = max(
        max_prediction_error,
        maximum_error(probability, artifact["test_probability"]),
        maximum_error(losses, artifact["test_loss_vector"]),
    )
    require(max(max_design_error, max_prediction_error) <= PARITY_TOLERANCE, "downstream reload parity failed")
    return {"max_design_error": max_design_error, "max_prediction_error": max_prediction_error}


def fit_or_load_downstream_artifact(
    path: Path,
    analysis_signature: str,
    stage_signature: str,
    model_name: str,
    outer_fold: int,
    frame: pd.DataFrame,
) -> tuple[dict[str, Any], dict[str, Any]]:
    newly_fitted = False
    if path.exists():
        artifact = joblib.load(path)
        require(artifact.get("analysis_signature") == analysis_signature, f"incompatible parent analysis signature: {path}")
        training = frame.loc[frame["outer_fold"].ne(outer_fold)].sort_values("SEQN", kind="mergesort").reset_index(drop=True)
        test = frame.loc[frame["outer_fold"].eq(outer_fold)].sort_values("SEQN", kind="mergesort").reset_index(drop=True)
        spec = DOWNSTREAM_MODELS[model_name]
        expected = downstream_artifact_signature(
            stage_signature,
            model_name,
            outer_fold,
            training,
            test,
            spec["score_column"],
            spec["score_mode"],
        )
        require(artifact.get("artifact_signature") == expected, f"incompatible cached artifact: {path}")
    else:
        artifact = fit_downstream_artifact(
            analysis_signature, stage_signature, model_name, outer_fold, frame
        )
        save_joblib_atomic(artifact, path)
        newly_fitted = True
    reloaded = joblib.load(path)
    parity = verify_downstream_artifact(
        reloaded, analysis_signature, stage_signature, model_name, outer_fold, frame
    )
    return reloaded, {
        "stage": "target_downstream",
        "model": model_name,
        "outer_fold": int(outer_fold),
        "artifact_path": str(path.resolve()),
        "artifact_sha256": sha256(path),
        "cache_action": "fitted" if newly_fitted else "reused",
        "max_design_error": parity["max_design_error"],
        "max_prediction_error": parity["max_prediction_error"],
        "downstream_tuning_estimator_count": int(
            sum(len(fold["candidate_fits"]) for fold in reloaded["inner_folds"])
        ),
        "downstream_final_estimator_count": 1,
    }


def phase1_source_analysis(
    out: Path,
    phase1: pd.DataFrame,
    phase2: pd.DataFrame,
    features: list[str],
    analysis_signature: str,
) -> tuple[pd.DataFrame, dict[str, Any], list[dict[str, Any]], list[dict[str, Any]]]:
    model_records: list[dict[str, Any]] = []
    tuning_rows: list[dict[str, Any]] = []
    score_rows: list[dict[str, Any]] = []
    for outer_fold in range(N_OUTER):
        training = phase1.loc[phase1["outer_fold"].ne(outer_fold)].sort_values(
            "SEQN", kind="mergesort"
        ).reset_index(drop=True)
        holdout = phase1.loc[phase1["outer_fold"].eq(outer_fold)].sort_values(
            "SEQN", kind="mergesort"
        ).reset_index(drop=True)
        inner_column = f"inner_fold_outer_{outer_fold}"
        kind = f"source_outer_fold_{outer_fold:02d}"
        artifact, record = fit_or_load_projection_artifact(
            out / "models" / "source_outer" / f"outer_fold_{outer_fold:02d}.joblib",
            analysis_signature,
            kind,
            training,
            holdout,
            features,
            inner_column,
            include_w_only=True,
        )
        model_records.append(record)
        for row in artifact["matched"]["tuning_rows"]:
            tuning_rows.append({"artifact_kind": kind, **row})
        for row in artifact["w_only"]["tuning_rows"]:
            tuning_rows.append({"artifact_kind": kind, **row})
        matched = artifact["matched"]["final_targets"]
        w_only = artifact["w_only"]["final_targets"]["q_W"]
        q_a = np.asarray(matched["q_A"]["holdout_score"], dtype=float)
        q_y = np.asarray(matched["q_Y"]["holdout_score"], dtype=float)
        q_w = np.asarray(w_only["holdout_score"], dtype=float)
        q_a_years = unstandardize(q_a, matched["q_A"]["target_scaler"])
        q_w_years = unstandardize(q_w, w_only["target_scaler"])
        for index, row in holdout.iterrows():
            score_rows.append(
                {
                    "SEQN": int(row["SEQN"]),
                    "outer_fold": int(outer_fold),
                    AGE: float(row[AGE]),
                    OUTCOME: int(row[OUTCOME]),
                    WEIGHT: float(row[WEIGHT]),
                    "q_A": float(q_a[index]),
                    "q_A_years": float(q_a_years[index]),
                    "q_Y": float(q_y[index]),
                    "q_W": float(q_w[index]),
                    "q_W_years": float(q_w_years[index]),
                    "q_A_selected_alpha": float(matched["q_A"]["selected_alpha"]),
                    "q_Y_selected_alpha": float(matched["q_Y"]["selected_alpha"]),
                    "q_W_selected_alpha": float(w_only["selected_alpha"]),
                }
            )
    source_scores = pd.DataFrame(score_rows).sort_values("SEQN", kind="mergesort").reset_index(drop=True)
    require(len(source_scores) == len(phase1), "Phase 1 OOF score coverage is incomplete")
    require(not source_scores["SEQN"].duplicated().any(), "Phase 1 OOF participant scored more than once")
    mse_q_a = weighted_mean(
        squared_loss(source_scores[AGE], source_scores["q_A_years"]), source_scores[WEIGHT]
    )
    mse_q_w = weighted_mean(
        squared_loss(source_scores[AGE], source_scores["q_W_years"]), source_scores[WEIGHT]
    )
    age_gate_pass = bool(mse_q_a < mse_q_w)

    phase2_projection = phase2[["SEQN", *BACKGROUND, *features]].copy()
    final_artifact, final_record = fit_or_load_projection_artifact(
        out / "models" / "source_final" / "phase1_final_projection_bundle.joblib",
        analysis_signature,
        "source_full_phase1_to_phase2_frozen_projection",
        phase1,
        phase2_projection,
        features,
        "full_inner_fold",
        include_w_only=False,
    )
    model_records.append(final_record)
    for row in final_artifact["matched"]["tuning_rows"]:
        tuning_rows.append({"artifact_kind": "source_full_phase1_to_phase2_frozen_projection", **row})
    final_targets = final_artifact["matched"]["final_targets"]
    phase2_q_a = np.asarray(final_targets["q_A"]["holdout_score"], dtype=float)
    phase2_q_y = np.asarray(final_targets["q_Y"]["holdout_score"], dtype=float)
    phase2_q_a_years = unstandardize(phase2_q_a, final_targets["q_A"]["target_scaler"])
    frozen_scores = pd.DataFrame(
        {
            "SEQN": phase2_projection["SEQN"].to_numpy("int64"),
            "q_A": phase2_q_a,
            "q_A_years": phase2_q_a_years,
            "q_Y": phase2_q_y,
            "q_A_selected_alpha": float(final_targets["q_A"]["selected_alpha"]),
            "q_Y_selected_alpha": float(final_targets["q_Y"]["selected_alpha"]),
        }
    )
    require(np.isfinite(frozen_scores.select_dtypes(include=[np.number])).all().all(), "frozen Phase 2 scores are non-finite")

    phase1_mean_age = weighted_mean(phase1[AGE], phase1[WEIGHT])
    phase2_weight = phase2[WEIGHT].to_numpy(float)
    phase2_age = phase2[AGE].to_numpy(float)
    phase2_mse = weighted_mean(squared_loss(phase2_age, phase2_q_a_years), phase2_weight)
    phase2_reference_mse = weighted_mean(
        squared_loss(phase2_age, np.repeat(phase1_mean_age, len(phase2))), phase2_weight
    )
    source_decision = {
        "analysis_version": ANALYSIS_VERSION,
        "status": STATUS,
        "source_population_n": int(len(phase1)),
        "source_deaths_5y": int(phase1[OUTCOME].sum()),
        "target_population_n": int(len(phase2)),
        "target_deaths_5y_not_used_in_projection_development": int(phase2[OUTCOME].sum()),
        "ordered_measurements_n": int(len(features)),
        "phase1_oof_q_A_age_mse": mse_q_a,
        "phase1_oof_W_only_age_mse": mse_q_w,
        "age_decoder_gate": "PASS" if age_gate_pass else "STOP_INVALID_AGE_DECODER",
        "phase2_q_A_age_rmse": float(math.sqrt(phase2_mse)),
        "phase2_q_A_R2_relative_to_frozen_phase1_weighted_mean": float(1.0 - phase2_mse / phase2_reference_mse),
        "phase1_weighted_mean_age": phase1_mean_age,
        "phase2_projection_outcomes_used": False,
        "claim_boundary": (
            "Projection development uses Phase 1 only. This is an outcome-accessed fixed "
            "protocol-transfer robustness analysis, not untouched confirmation."
        ),
    }
    write_table_or_verify(source_scores, out / "source_phase1_oof_projection_scores.tsv.gz")
    write_table_or_verify(pd.DataFrame(tuning_rows), out / "source_projection_inner_tuning.tsv.gz")
    write_table_or_verify(frozen_scores, out / "frozen_phase2_scalar_scores.tsv.gz")
    write_json_or_verify(out / "source_projection_decision.json", source_decision)
    source_report = f"""# NHANES III matched-scalar protocol transfer: Phase 1 projection freeze

- Status: **{STATUS}**
- Phase 1 participants/deaths: {len(phase1)} / {int(phase1[OUTCOME].sum())}
- Ordered source measurements: {len(features)}
- Phase 1 OOF age MSE, `q_A`: {mse_q_a:.9f}
- Phase 1 OOF age MSE, `W` only: {mse_q_w:.9f}
- Operational age-decoder gate: **{source_decision['age_decoder_gate']}**
- Phase 2 transported age RMSE: {math.sqrt(phase2_mse):.9f}

The two final scalar directions were trained and tuned only in Phase 1 from the
same background-plus-25-measurement design. Phase 2 mortality outcomes did not
enter imputation, standardization, tuning or projection fitting.
"""
    write_text_or_verify(out / "SOURCE_REPORT.md", source_report)
    return frozen_scores, source_decision, model_records, tuning_rows


def run_downstream_refits(
    out: Path,
    phase2: pd.DataFrame,
    frozen_scores: pd.DataFrame,
    analysis_signature: str,
    label: str,
    weight_factor: np.ndarray,
    model_names: Sequence[str],
) -> tuple[pd.DataFrame, pd.DataFrame, list[dict[str, Any]]]:
    require(len(weight_factor) == len(phase2), "target replicate weight length changed")
    frame = phase2.merge(frozen_scores, on="SEQN", how="left", validate="one_to_one")
    frame = frame.sort_values("SEQN", kind="mergesort").reset_index(drop=True)
    require(frame[["q_A", "q_Y"]].notna().all().all(), "frozen scalar merge is incomplete")
    frame[WEIGHT] = frame[BASE_WEIGHT].to_numpy(float) * np.asarray(weight_factor, dtype=float)
    require(np.isfinite(frame[WEIGHT]).all() and frame[WEIGHT].gt(0).all(), "invalid target replicate weights")
    stage_signature = stable_hash(
        {
            "analysis_signature": analysis_signature,
            "stage": "phase2_equal_target_local_calibration",
            "label": label,
            "ordered_seqn": frame["SEQN"].astype(int).tolist(),
            "ordered_weight": frame[WEIGHT].astype(float).tolist(),
            "frozen_score_signature": stable_hash(
                frozen_scores[["SEQN", "q_A", "q_Y"]].to_dict(orient="records")
            ),
            "models": list(model_names),
            "projections_refitted": False,
        }
    )
    prediction = frame[
        ["SEQN", "outer_fold", OUTCOME, BASE_WEIGHT, WEIGHT, STRATUM, PSU, "q_A", "q_Y"]
    ].copy()
    tuning_tables: list[pd.DataFrame] = []
    model_records: list[dict[str, Any]] = []
    if label == "primary":
        model_root = out / "models" / "target_primary"
    else:
        model_root = out / "models" / "target_fay_brr" / label
    for model_name in model_names:
        probability = np.full(len(frame), np.nan, dtype=float)
        for outer_fold in range(N_OUTER):
            artifact, record = fit_or_load_downstream_artifact(
                model_root / model_name / f"outer_fold_{outer_fold:02d}.joblib",
                analysis_signature,
                stage_signature,
                model_name,
                outer_fold,
                frame,
            )
            model_records.append({"target_refit_label": label, **record})
            test_ids = np.asarray(artifact["test_seqn"], dtype="int64")
            positions = frame.reset_index().set_index("SEQN").loc[test_ids, "index"].to_numpy(int)
            probability[positions] = np.asarray(artifact["test_probability"], dtype=float)
            tuning = pd.DataFrame(artifact["tuning_rows"])
            tuning.insert(0, "target_refit_label", label)
            tuning_tables.append(tuning)
        require(np.isfinite(probability).all(), f"incomplete OOF probability for {model_name}")
        prediction[f"probability_{model_name}"] = probability
        prediction[f"loss_{model_name}"] = log_loss_vector(frame[OUTCOME], probability)
    tuning_frame = pd.concat(tuning_tables, ignore_index=True)
    return prediction, tuning_frame, model_records


def summarize_contrasts(prediction: pd.DataFrame) -> tuple[dict[str, float], pd.DataFrame]:
    require({"loss_M0", "loss_M_A", "loss_M_Y"}.issubset(prediction.columns), "primary loss columns missing")
    prediction["paired_delta_loss"] = prediction["loss_M_A"] - prediction["loss_M_Y"]
    prediction["paired_U_A_loss"] = prediction["loss_M0"] - prediction["loss_M_A"]
    estimates = {
        "loss_M0": weighted_mean(prediction["loss_M0"], prediction[WEIGHT]),
        "loss_M_A": weighted_mean(prediction["loss_M_A"], prediction[WEIGHT]),
        "loss_M_Y": weighted_mean(prediction["loss_M_Y"], prediction[WEIGHT]),
        "Delta": weighted_mean(prediction["paired_delta_loss"], prediction[WEIGHT]),
        "U_A": weighted_mean(prediction["paired_U_A_loss"], prediction[WEIGHT]),
    }
    if {"loss_M_A_LINEAR", "loss_M_Y_LINEAR"}.issubset(prediction.columns):
        prediction["paired_delta_linear_loss"] = (
            prediction["loss_M_A_LINEAR"] - prediction["loss_M_Y_LINEAR"]
        )
        estimates.update(
            {
                "loss_M_A_LINEAR": weighted_mean(prediction["loss_M_A_LINEAR"], prediction[WEIGHT]),
                "loss_M_Y_LINEAR": weighted_mean(prediction["loss_M_Y_LINEAR"], prediction[WEIGHT]),
                "Delta_LINEAR": weighted_mean(prediction["paired_delta_linear_loss"], prediction[WEIGHT]),
            }
        )
    fold_rows: list[dict[str, Any]] = []
    for outer_fold in range(N_OUTER):
        block = prediction.loc[prediction["outer_fold"].eq(outer_fold)]
        delta = weighted_mean(block["paired_delta_loss"], block[WEIGHT])
        u_a = weighted_mean(block["paired_U_A_loss"], block[WEIGHT])
        fold_rows.append(
            {
                "outer_fold": int(outer_fold),
                "n": int(len(block)),
                "deaths_5y": int(block[OUTCOME].sum()),
                "Delta": delta,
                "Delta_positive": bool(delta > 0),
                "U_A": u_a,
                "U_A_positive": bool(u_a > 0),
            }
        )
    return estimates, pd.DataFrame(fold_rows)


def fay_interval(full_estimate: float, replicates: Sequence[float]) -> dict[str, float]:
    values = np.asarray(replicates, dtype=float)
    require(values.shape == (FAY_REPLICATES,) and np.isfinite(values).all(), "Fay-BRR estimates are incomplete")
    variance = float(np.sum((values - float(full_estimate)) ** 2) / (FAY_REPLICATES * (1.0 - FAY_RHO) ** 2))
    standard_error = float(math.sqrt(variance))
    critical = float(student_t.ppf(0.975, FAY_DF))
    return {
        "estimate": float(full_estimate),
        "variance": variance,
        "standard_error": standard_error,
        "critical_t": critical,
        "df": FAY_DF,
        "ci_lower": float(full_estimate - critical * standard_error),
        "ci_upper": float(full_estimate + critical * standard_error),
        "replicates": FAY_REPLICATES,
        "rho": FAY_RHO,
        "conditional_on": "two_fixed_Phase1_projection_directions",
    }


def phase2_target_analysis(
    out: Path,
    phase2: pd.DataFrame,
    frozen_scores: pd.DataFrame,
    factors: np.ndarray,
    analysis_signature: str,
) -> tuple[dict[str, Any], list[dict[str, Any]]]:
    all_records: list[dict[str, Any]] = []
    primary_models = ("M0", "M_A", "M_Y", "M_A_LINEAR", "M_Y_LINEAR")
    primary_prediction, primary_tuning, primary_records = run_downstream_refits(
        out,
        phase2,
        frozen_scores,
        analysis_signature,
        "primary",
        np.ones(len(phase2), dtype=float),
        primary_models,
    )
    all_records.extend(primary_records)
    primary_estimates, fold_contrasts = summarize_contrasts(primary_prediction)
    write_table_or_verify(primary_prediction, out / "target_primary_oof_predictions_and_losses.tsv.gz")
    write_table_or_verify(primary_tuning, out / "target_primary_inner_tuning.tsv.gz")
    write_table_or_verify(fold_contrasts, out / "target_primary_fold_contrasts.tsv")

    replicate_rows: list[dict[str, Any]] = []
    replicate_tuning_tables: list[pd.DataFrame] = []
    for replicate in range(FAY_REPLICATES):
        label = f"replicate_{replicate:02d}"
        prediction, tuning, records = run_downstream_refits(
            out,
            phase2,
            frozen_scores,
            analysis_signature,
            label,
            factors[replicate],
            ("M0", "M_A", "M_Y"),
        )
        all_records.extend(records)
        estimates, folds = summarize_contrasts(prediction)
        write_table_or_verify(
            prediction,
            out / "fay_brr_predictions" / f"{label}_oof_predictions_and_losses.tsv.gz",
        )
        replicate_tuning_tables.append(tuning)
        replicate_rows.append(
            {
                "replicate": int(replicate),
                "loss_M0": estimates["loss_M0"],
                "loss_M_A": estimates["loss_M_A"],
                "loss_M_Y": estimates["loss_M_Y"],
                "Delta": estimates["Delta"],
                "U_A": estimates["U_A"],
                "Delta_positive_outer_folds": int(folds["Delta_positive"].sum()),
                "U_A_positive_outer_folds": int(folds["U_A_positive"].sum()),
            }
        )
    replicate_table = pd.DataFrame(replicate_rows)
    write_table_or_verify(replicate_table, out / "target_fay_brr_replicate_estimates.tsv")
    write_table_or_verify(
        pd.concat(replicate_tuning_tables, ignore_index=True),
        out / "target_fay_brr_inner_tuning.tsv.gz",
    )
    delta_interval = fay_interval(primary_estimates["Delta"], replicate_table["Delta"])
    u_a_interval = fay_interval(primary_estimates["U_A"], replicate_table["U_A"])
    positive_folds = int(fold_contrasts["Delta_positive"].sum())
    delta_three_gates = bool(
        primary_estimates["Delta"] > 0
        and delta_interval["ci_lower"] > 0
        and positive_folds >= 4
    )
    u_a_support = bool(primary_estimates["U_A"] > 0 and u_a_interval["ci_lower"] > 0)
    if primary_estimates["Delta"] <= 0:
        decision = "NO_TRANSFERRED_TARGET_ADVANTAGE"
    elif delta_three_gates and u_a_support:
        decision = "UTILITY_WITHOUT_PRIVILEGE_PROTOCOL_TRANSFER"
    elif delta_three_gates:
        decision = "MATCHED_TARGET_SCALAR_ADVANTAGE_PROTOCOL_TRANSFER"
    else:
        decision = "DIRECTIONAL_PROTOCOL_TRANSFER"
    result = {
        "analysis_version": ANALYSIS_VERSION,
        "status": STATUS,
        "decision": decision,
        "target_population_n": int(len(phase2)),
        "target_deaths_5y": int(phase2[OUTCOME].sum()),
        "models": {
            "M0": "Y ~ W + fixed_five_knot_cubic_spline(A)",
            "M_A": "Y ~ W + fixed_five_knot_cubic_spline(A) + training_local_uniform_five_knot_cubic_spline(q_A)",
            "M_Y": "Y ~ W + fixed_five_knot_cubic_spline(A) + training_local_uniform_five_knot_cubic_spline(q_Y)",
        },
        "primary_estimates": primary_estimates,
        "Delta_Fay_BRR": delta_interval,
        "U_A_Fay_BRR": u_a_interval,
        "Delta_positive_outer_folds": positive_folds,
        "Delta_gate_pass": delta_three_gates,
        "U_A_support_gate_pass": u_a_support,
        "Fay_BRR_replicates": FAY_REPLICATES,
        "Fay_BRR_full_downstream_refits": True,
        "Fay_BRR_projection_refits": False,
        "claim_ceiling": (
            "For this population, five-year endpoint, 25-item interface and one-dimensional "
            "linear projection family, chronological-age pretraining was not necessary and "
            "did not provide the best scalar direction if the positive gates pass. This does "
            "not identify biological age, establish universal mortality sufficiency, compare "
            "all clocks, validate a surrogate or imply causal benefit."
        ),
    }
    write_json_or_verify(out / "target_decision.json", result)
    report = f"""# NHANES III matched-scalar target-supervision protocol transfer v0.2

- Status: **{STATUS}**
- Decision: **{decision}**
- Phase 2 participants/deaths: {len(phase2)} / {int(phase2[OUTCOME].sum())}
- `M0` design-weighted OOF log loss: {primary_estimates['loss_M0']:.9f}
- `M_A` design-weighted OOF log loss: {primary_estimates['loss_M_A']:.9f}
- `M_Y` design-weighted OOF log loss: {primary_estimates['loss_M_Y']:.9f}
- Primary `Delta = L(M_A)-L(M_Y)`: {primary_estimates['Delta']:.9f}
- 95% Fay-BRR interval for `Delta`: {delta_interval['ci_lower']:.9f} to {delta_interval['ci_upper']:.9f}
- `U_A = L(M0)-L(M_A)`: {primary_estimates['U_A']:.9f}
- 95% Fay-BRR interval for `U_A`: {u_a_interval['ci_lower']:.9f} to {u_a_interval['ci_upper']:.9f}
- Positive fixed outer-fold `Delta` contrasts: {positive_folds} of {N_OUTER}

The two one-dimensional directions were frozen from Phase 1. Every Fay-BRR
replicate repeated Phase 2 inner tuning, outer fitting, held-out prediction and
paired-loss evaluation under replicate weights; it did not relearn either
projection. The interval is conditional on the two Phase 1-fitted directions.
"""
    write_text_or_verify(out / "REPORT.md", report)
    return result, all_records


def deterministic_model_index(records: list[dict[str, Any]]) -> pd.DataFrame:
    rows = []
    for record in records:
        row = {key: value for key, value in record.items() if key != "cache_action"}
        rows.append(row)
    table = pd.DataFrame(rows).sort_values(["stage", "artifact_path"], kind="mergesort").reset_index(drop=True)
    require(not table["artifact_path"].duplicated().any(), "model artifact index has duplicate paths")
    return table


def audit_model_records(records: list[dict[str, Any]], expected_count: int) -> dict[str, Any]:
    require(len(records) > 0, "model artifact index is empty")
    require(len(records) == expected_count, f"expected {expected_count} model artifacts, found {len(records)}")
    max_design = max(float(record["max_design_error"]) for record in records)
    max_prediction = max(float(record["max_prediction_error"]) for record in records)
    require(max(max_design, max_prediction) <= PARITY_TOLERANCE, "model artifact parity audit failed")
    for record in records:
        path = Path(record["artifact_path"])
        require(path.is_file(), f"indexed model artifact is missing: {path}")
        require(sha256(path) == record["artifact_sha256"], f"indexed model artifact hash changed: {path}")
    return {
        "status": "PASS",
        "analysis_version": ANALYSIS_VERSION,
        "artifact_count": int(len(records)),
        "expected_artifact_count": int(expected_count),
        "projection_tuning_estimator_count": int(sum(record.get("projection_tuning_estimator_count", 0) for record in records)),
        "projection_final_estimator_count": int(sum(record.get("projection_final_estimator_count", 0) for record in records)),
        "downstream_tuning_estimator_count": int(sum(record.get("downstream_tuning_estimator_count", 0) for record in records)),
        "downstream_final_estimator_count": int(sum(record.get("downstream_final_estimator_count", 0) for record in records)),
        "maximum_design_reconstruction_error": max_design,
        "maximum_prediction_reconstruction_error": max_prediction,
        "parity_tolerance": PARITY_TOLERANCE,
        "cache_policy": "compatible_artifacts_reused; incompatible_artifacts_fail_closed",
    }


def write_output_manifest(out: Path) -> None:
    rows: list[dict[str, Any]] = []
    manifest_path = out / "artifact_manifest.tsv"
    for path in sorted(out.rglob("*"), key=lambda item: str(item).lower()):
        if not path.is_file() or path == manifest_path:
            continue
        rows.append(
            {
                "relative_path": str(path.relative_to(out)).replace("\\", "/"),
                "size_bytes": int(path.stat().st_size),
                "sha256": sha256(path),
            }
        )
    write_table_or_verify(pd.DataFrame(rows), manifest_path)


def build_analysis_signature(input_records: list[dict[str, Any]], features: list[str]) -> tuple[str, dict[str, Any]]:
    payload = {
        "analysis_version": ANALYSIS_VERSION,
        "schema_version": SCHEMA_VERSION,
        "status": STATUS,
        "canonical_input_directory": str(CANONICAL.resolve()),
        "inputs": input_records,
        "ordered_features": features,
        "background": BACKGROUND,
        "age_range": "[20,80)",
        "endpoint": "resolved_five_year_all_cause_mortality",
        "alpha_grid": ALPHA_GRID,
        "C_grid": C_GRID,
        "fixed_age_knots": AGE_KNOTS,
        "score_spline": "uniform_5_knots_degree3_linear_extrapolation",
        "outer_folds": N_OUTER,
        "inner_folds": N_INNER,
        "Fay_BRR": {"replicates": FAY_REPLICATES, "rho": FAY_RHO, "df": FAY_DF},
        "parity_tolerance": PARITY_TOLERANCE,
        "software": software_record(),
    }
    return stable_hash(payload), payload


def freeze_inputs(out: Path, input_records: list[dict[str, Any]]) -> None:
    frozen = out / "input_freeze"
    for key, name in CANONICAL_FILES.items():
        copy_file_or_verify(CANONICAL / name, frozen / name)
    write_table_or_verify(pd.DataFrame(input_records), out / "input_manifest.tsv")


def run(args: argparse.Namespace) -> int:
    configure_stdout()
    require(CANONICAL.resolve() == (ROOT / "outputs" / CANONICAL.name).resolve(), "canonical input directory changed")
    out = Path(args.output_dir).resolve()
    require(out.name == ANALYSIS_VERSION, "output directory must retain the frozen analysis-version name")
    require(out != CANONICAL.resolve(), "new analysis may not write into the canonical predecessor output")
    out.mkdir(parents=True, exist_ok=True)
    phase1, phase2, manifest, features, factors, input_records = load_inputs()
    analysis_signature, signature_payload = build_analysis_signature(input_records, features)
    freeze_inputs(out, input_records)
    write_json_or_verify(
        out / "analysis_signature.json",
        {"analysis_signature": analysis_signature, "signature_payload": signature_payload},
    )
    write_table_or_verify(manifest, out / "ordered_5x5_feature_manifest.tsv")

    frozen_scores, source_decision, source_records, _ = phase1_source_analysis(
        out, phase1, phase2, features, analysis_signature
    )
    source_index = deterministic_model_index(source_records)
    source_audit = audit_model_records(source_records, 6)
    write_table_or_verify(source_index, out / "model_artifact_index_source.tsv")
    write_json_or_verify(out / "model_artifact_audit_source.json", source_audit)
    if source_decision["age_decoder_gate"] != "PASS":
        write_json_or_verify(
            out / "decision.json",
            {
                "analysis_version": ANALYSIS_VERSION,
                "decision": "STOP_INVALID_AGE_DECODER",
                "source_projection_decision": source_decision,
                "target_analysis_executed": False,
            },
        )
        index = deterministic_model_index(source_records)
        write_table_or_verify(index, out / "model_artifact_index.tsv")
        write_json_or_verify(out / "model_artifact_audit.json", audit_model_records(source_records, 6))
        write_output_manifest(out)
        print(json.dumps({"decision": "STOP_INVALID_AGE_DECODER"}, indent=2))
        return 2
    if args.stage == "source":
        print(json.dumps(source_decision, indent=2, ensure_ascii=False))
        return 0

    target_decision, target_records = phase2_target_analysis(
        out, phase2, frozen_scores, factors, analysis_signature
    )
    all_records = [*source_records, *target_records]
    index = deterministic_model_index(all_records)
    write_table_or_verify(index, out / "model_artifact_index.tsv")
    audit = audit_model_records(all_records, 511)
    write_json_or_verify(out / "model_artifact_audit.json", audit)
    final = {
        "analysis_version": ANALYSIS_VERSION,
        "status": STATUS,
        "decision": target_decision["decision"],
        "source_projection_decision": source_decision,
        "target_decision": target_decision,
        "model_artifact_audit_status": audit["status"],
        "manuscript_or_frozen_predecessor_modified": False,
    }
    write_json_or_verify(out / "decision.json", final)
    write_output_manifest(out)
    print(json.dumps(final, indent=2, ensure_ascii=False))
    return 0


def build_parser() -> argparse.ArgumentParser:
    parser = argparse.ArgumentParser(description=__doc__)
    parser.add_argument(
        "--output-dir",
        default=str(DEFAULT_OUT),
        help="Versioned output directory; basename must equal the frozen analysis version.",
    )
    parser.add_argument(
        "--stage",
        choices=("source", "all"),
        default="all",
        help="Run and freeze Phase 1 only, or run the complete fixed protocol transfer.",
    )
    return parser


def main(argv: Sequence[str] | None = None) -> int:
    return run(build_parser().parse_args(argv))


if __name__ == "__main__":
    raise SystemExit(main())
```
