## Supplementary_Software_1 for "Inferential boundaries of age prediction: why prediction does not establish biological age measurement": analyze_nhanes_5y_age_decoder_source_information_probe_v0_1.py

````
#!/usr/bin/env python
"""Decision probe: pure age decoder versus its source measurements.

Implements NHANES_5Y_AGE_DECODER_SOURCE_INFORMATION_PROBE_SPEC_20260823.md.
The accepted complete-support NHANES interface, participant folds, baseline
outcome model and direct source-measurement outcome model are reused. Every new
age-decoder and clock-endpoint fit is persisted and reload-verified.
"""

from __future__ import annotations

import hashlib
import json
import math
import platform
import sys
import warnings
from pathlib import Path
from typing import Any

import joblib
import numpy as np
import pandas as pd
import sklearn
from sklearn.exceptions import ConvergenceWarning
from sklearn.linear_model import LogisticRegression, Ridge
from sklearn.metrics import brier_score_loss, log_loss, mean_absolute_error, mean_squared_error, r2_score, roc_auc_score
from sklearn.model_selection import StratifiedKFold
from sklearn.preprocessing import OneHotEncoder, SplineTransformer, StandardScaler


ROOT = Path(__file__).resolve().parents[1]
ANALYSIS_DIR = ROOT / "analysis"
PREDECESSOR = ROOT / "outputs" / "nhanes_5y_complete_support_contextual_age_probe_v0_1"
OUT = ROOT / "outputs" / "nhanes_5y_age_decoder_source_information_probe_v0_2"
MODEL_DIR = OUT / "models"

SOURCE_FRAME = PREDECESSOR / "analysis_ready_risk_set.tsv.gz"
SOURCE_FOLDS = PREDECESSOR / "participant_folds.tsv"
SOURCE_TIERS = PREDECESSOR / "ordered_feature_tiers.tsv"
SOURCE_OOF = PREDECESSOR / "oof_predictions.tsv.gz"
SOURCE_LOSSES = PREDECESSOR / "participant_loss_vectors.tsv.gz"
SOURCE_BOOTSTRAP = PREDECESSOR / "bootstrap_indices.npz"
SOURCE_SIGNATURE = PREDECESSOR / "analysis_signature.json"
SOURCE_MODEL_AUDIT = PREDECESSOR / "model_artifact_audit.json"
SPEC = ANALYSIS_DIR / "NHANES_5Y_AGE_DECODER_SOURCE_INFORMATION_PROBE_SPEC_20260823.md"
ENV_POLICY = ROOT / "environment-policy.json"
ENV_MANIFEST = ROOT / "environment.yml"
ENV_LOCK = ROOT / "environment.win-64.lock.txt"

ANALYSIS_VERSION = "nhanes_5y_age_decoder_source_information_probe_v0_2"
SCHEMA_VERSION = "1.0"
EXPECTED_N = 13_724
EXPECTED_EVENTS = 470
N_OUTER_FOLDS = 5
N_INNER_FOLDS = 3
OUTER_SEED = 20260823
AGE_INNER_SEED = 20260825
OUTCOME_INNER_SEED = 20260826
PARITY_TOLERANCE = 1e-12
EPSILON = 1e-7
BACKGROUND = ("sex", "race_ethnicity", "cycle")
ALPHA_GRID = (0.01, 0.1, 1.0, 10.0, 100.0, 1000.0)
C_GRID = (0.001, 0.003, 0.01, 0.03, 0.1, 0.3, 1.0, 3.0, 10.0, 30.0, 100.0, 300.0)


def configure_stdout() -> None:
    if hasattr(sys.stdout, "reconfigure"):
        sys.stdout.reconfigure(encoding="utf-8")


def sha256(path: Path) -> str:
    digest = hashlib.sha256()
    with path.open("rb") as handle:
        for block in iter(lambda: handle.read(1024 * 1024), b""):
            digest.update(block)
    return digest.hexdigest()


def stable_hash(payload: Any) -> str:
    encoded = json.dumps(payload, sort_keys=True, separators=(",", ":"), ensure_ascii=False).encode("utf-8")
    return hashlib.sha256(encoded).hexdigest()


def json_ready(value: Any) -> Any:
    if isinstance(value, dict):
        return {str(key): json_ready(item) for key, item in value.items()}
    if isinstance(value, (list, tuple)):
        return [json_ready(item) for item in value]
    if isinstance(value, Path):
        return str(value)
    if isinstance(value, np.ndarray):
        return value.tolist()
    if isinstance(value, (np.integer,)):
        return int(value)
    if isinstance(value, (np.floating,)):
        return None if not np.isfinite(value) else float(value)
    if isinstance(value, float):
        return None if not math.isfinite(value) else value
    return value


def write_json(path: Path, payload: Any) -> None:
    path.write_text(json.dumps(json_ready(payload), indent=2, ensure_ascii=False) + "\n", encoding="utf-8")


def clean_categories(frame: pd.DataFrame) -> pd.DataFrame:
    return frame.fillna("missing").astype(str).replace("", "missing")


def make_age_strata(age: np.ndarray, n_splits: int) -> np.ndarray:
    strata = pd.qcut(pd.Series(age), q=5, duplicates="drop", labels=False)
    strata = strata.fillna(-1).astype(int).to_numpy()
    counts = pd.Series(strata).value_counts()
    if int(counts.min()) < n_splits:
        raise RuntimeError("Age-quintile strata cannot support fixed inner folds")
    return strata


def make_outcome_strata(y: np.ndarray, age: np.ndarray, n_splits: int) -> np.ndarray:
    age_quintile = make_age_strata(age, n_splits)
    combined = np.char.add(np.char.add(y.astype(str), "_"), age_quintile.astype(str))
    counts = pd.Series(combined).value_counts()
    if int(counts.min()) >= n_splits:
        return combined
    if int(pd.Series(y).value_counts().min()) < n_splits:
        raise RuntimeError("Endpoint strata cannot support fixed inner folds")
    return y.astype(str)


def fit_age_preprocessor(frame: pd.DataFrame, features: list[str]) -> tuple[OneHotEncoder, StandardScaler, np.ndarray, list[str]]:
    encoder = OneHotEncoder(handle_unknown="ignore", sparse_output=False)
    categories = encoder.fit_transform(clean_categories(frame.loc[:, BACKGROUND]))
    scaler = StandardScaler()
    measurements = scaler.fit_transform(frame.loc[:, features].to_numpy(float))
    design = np.column_stack([categories, measurements])
    names = [f"W__{name}" for name in encoder.get_feature_names_out(BACKGROUND)] + [f"X__{name}" for name in features]
    return encoder, scaler, design, names


def transform_age_preprocessor(
    frame: pd.DataFrame,
    features: list[str],
    encoder: OneHotEncoder,
    scaler: StandardScaler,
) -> np.ndarray:
    categories = encoder.transform(clean_categories(frame.loc[:, BACKGROUND]))
    measurements = scaler.transform(frame.loc[:, features].to_numpy(float))
    return np.column_stack([categories, measurements])


def fit_outcome_preprocessor(
    frame: pd.DataFrame,
    score: np.ndarray,
) -> tuple[OneHotEncoder, SplineTransformer, StandardScaler, np.ndarray, list[str]]:
    encoder = OneHotEncoder(handle_unknown="ignore", sparse_output=False)
    categories = encoder.fit_transform(clean_categories(frame.loc[:, BACKGROUND]))
    spline = SplineTransformer(n_knots=5, degree=3, include_bias=False, extrapolation="linear")
    age_basis = spline.fit_transform(frame[["age_years"]].to_numpy(float))
    score_scaler = StandardScaler()
    score_design = score_scaler.fit_transform(np.asarray(score, dtype=float).reshape(-1, 1))
    design = np.column_stack([categories, age_basis, score_design])
    names = (
        [f"W__{name}" for name in encoder.get_feature_names_out(BACKGROUND)]
        + [f"A_spline_{index}" for index in range(age_basis.shape[1])]
        + ["clock_score"]
    )
    return encoder, spline, score_scaler, design, names


def transform_outcome_preprocessor(
    frame: pd.DataFrame,
    score: np.ndarray,
    encoder: OneHotEncoder,
    spline: SplineTransformer,
    score_scaler: StandardScaler,
) -> np.ndarray:
    categories = encoder.transform(clean_categories(frame.loc[:, BACKGROUND]))
    age_basis = spline.transform(frame[["age_years"]].to_numpy(float))
    score_design = score_scaler.transform(np.asarray(score, dtype=float).reshape(-1, 1))
    return np.column_stack([categories, age_basis, score_design])


def fit_ridge_age(frame: pd.DataFrame, features: list[str], alpha: float) -> dict[str, Any]:
    encoder, scaler, design, names = fit_age_preprocessor(frame, features)
    estimator = Ridge(alpha=float(alpha))
    estimator.fit(design, frame["age_years"].to_numpy(float))
    return {
        "encoder": encoder,
        "scaler": scaler,
        "estimator": estimator,
        "design_feature_names": names,
    }


def predict_ridge_age(model: dict[str, Any], frame: pd.DataFrame, features: list[str]) -> np.ndarray:
    design = transform_age_preprocessor(frame, features, model["encoder"], model["scaler"])
    return model["estimator"].predict(design)


def fit_clock_outcome(frame: pd.DataFrame, score: np.ndarray, c_value: float) -> dict[str, Any]:
    encoder, spline, score_scaler, design, names = fit_outcome_preprocessor(frame, score)
    estimator = LogisticRegression(C=float(c_value), penalty="l2", solver="lbfgs", max_iter=4000)
    with warnings.catch_warnings():
        warnings.filterwarnings("ignore", category=ConvergenceWarning)
        warnings.filterwarnings("ignore", category=FutureWarning, message=".*'penalty'.*deprecated.*")
        estimator.fit(design, frame["death_5y"].to_numpy(int))
    return {
        "encoder": encoder,
        "spline": spline,
        "score_scaler": score_scaler,
        "estimator": estimator,
        "design_feature_names": names,
    }


def predict_clock_outcome(model: dict[str, Any], frame: pd.DataFrame, score: np.ndarray) -> np.ndarray:
    design = transform_outcome_preprocessor(
        frame,
        score,
        model["encoder"],
        model["spline"],
        model["score_scaler"],
    )
    return model["estimator"].predict_proba(design)[:, 1]


def participant_loss(y: np.ndarray, probability: np.ndarray) -> np.ndarray:
    p = np.clip(np.asarray(probability, dtype=float), EPSILON, 1.0 - EPSILON)
    y_array = np.asarray(y, dtype=float)
    return -(y_array * np.log(p) + (1.0 - y_array) * np.log(1.0 - p))


def endpoint_metrics(y: np.ndarray, probability: np.ndarray) -> dict[str, float]:
    return {
        "log_loss": float(log_loss(y, probability, labels=[0, 1])),
        "brier": float(brier_score_loss(y, probability)),
        "auc": float(roc_auc_score(y, probability)),
    }


def load_and_gate() -> tuple[pd.DataFrame, list[str], pd.DataFrame, np.ndarray, dict[str, Any]]:
    source_audit = json.loads(SOURCE_MODEL_AUDIT.read_text(encoding="utf-8"))
    if source_audit.get("status") != "PASS":
        raise RuntimeError("STOP_DATA_OR_ARTIFACT_FAIL: predecessor model audit is not PASS")
    frame = pd.read_csv(SOURCE_FRAME, sep="\t")
    folds = pd.read_csv(SOURCE_FOLDS, sep="\t")
    tiers = pd.read_csv(SOURCE_TIERS, sep="\t")
    source_oof = pd.read_csv(SOURCE_OOF, sep="\t")
    bootstrap = np.load(SOURCE_BOOTSTRAP)["indices"]

    features = tiers.loc[tiers["tier"].eq("S3")].sort_values("order")["transformed_column"].tolist()
    required = ["row_id", "SEQN", "age_years", "death_5y", "outer_fold", *BACKGROUND, *features]
    missing = sorted(set(required) - set(frame.columns))
    if missing:
        raise RuntimeError(f"STOP_DATA_OR_ARTIFACT_FAIL: missing columns {missing}")
    if len(features) != 50 or len(set(features)) != 50:
        raise RuntimeError("STOP_DATA_OR_ARTIFACT_FAIL: S3 is not the ordered 50-measurement panel")
    if len(frame) != EXPECTED_N or int(frame["death_5y"].sum()) != EXPECTED_EVENTS:
        raise RuntimeError("STOP_DATA_OR_ARTIFACT_FAIL: participant or event count changed")
    if frame["SEQN"].nunique() != EXPECTED_N or sorted(frame["outer_fold"].unique()) != list(range(N_OUTER_FOLDS)):
        raise RuntimeError("STOP_DATA_OR_ARTIFACT_FAIL: participant identity or outer folds changed")
    if frame[required].isna().any().any():
        raise RuntimeError("STOP_DATA_OR_ARTIFACT_FAIL: missing analysis input")
    fold_check = frame[["row_id", "SEQN", "outer_fold"]].merge(
        folds[["row_id", "SEQN", "outer_fold"]],
        on=["row_id", "SEQN"],
        suffixes=("_frame", "_frozen"),
        validate="one_to_one",
    )
    if not np.array_equal(fold_check["outer_fold_frame"].to_numpy(), fold_check["outer_fold_frozen"].to_numpy()):
        raise RuntimeError("STOP_DATA_OR_ARTIFACT_FAIL: frozen fold mismatch")
    source_oof = frame[["row_id", "SEQN"]].merge(source_oof, on=["row_id", "SEQN"], validate="one_to_one")
    if not np.array_equal(source_oof["outer_fold"].to_numpy(int), frame["outer_fold"].to_numpy(int)):
        raise RuntimeError("STOP_DATA_OR_ARTIFACT_FAIL: predecessor prediction fold mismatch")
    if bootstrap.shape != (2000, EXPECTED_N):
        raise RuntimeError(f"STOP_DATA_OR_ARTIFACT_FAIL: bootstrap shape changed: {bootstrap.shape}")
    return frame.reset_index(drop=True), features, source_oof, bootstrap, source_audit


def fit_outer_artifact(
    frame: pd.DataFrame,
    features: list[str],
    outer_fold: int,
    analysis_signature: str,
) -> dict[str, Any]:
    train = frame.loc[frame["outer_fold"].ne(outer_fold)].copy().reset_index(drop=True)
    test = frame.loc[frame["outer_fold"].eq(outer_fold)].copy().reset_index(drop=True)
    age_strata = make_age_strata(train["age_years"].to_numpy(float), N_INNER_FOLDS)
    age_splitter = StratifiedKFold(n_splits=N_INNER_FOLDS, shuffle=True, random_state=AGE_INNER_SEED + outer_fold)
    age_splits = [(tr, va) for tr, va in age_splitter.split(np.zeros(len(train)), age_strata)]

    age_tuning_fits: list[dict[str, Any]] = []
    age_tuning_rows: list[dict[str, Any]] = []
    for alpha in ALPHA_GRID:
        for inner_fold, (inner_train, inner_valid) in enumerate(age_splits):
            model = fit_ridge_age(train.iloc[inner_train], features, alpha)
            prediction = predict_ridge_age(model, train.iloc[inner_valid], features)
            rmse = float(mean_squared_error(train.iloc[inner_valid]["age_years"], prediction) ** 0.5)
            fit = {
                "alpha": float(alpha),
                "inner_fold": inner_fold,
                "train_row_ids": train.iloc[inner_train]["row_id"].to_numpy(int),
                "valid_row_ids": train.iloc[inner_valid]["row_id"].to_numpy(int),
                "model": model,
                "valid_prediction": prediction,
            }
            age_tuning_fits.append(fit)
            age_tuning_rows.append({"alpha": float(alpha), "inner_fold": inner_fold, "rmse": rmse})
    age_tuning = pd.DataFrame(age_tuning_rows)
    mean_age_rmse = age_tuning.groupby("alpha", as_index=False)["rmse"].mean().sort_values(["rmse", "alpha"])
    selected_alpha = float(mean_age_rmse.iloc[0]["alpha"])
    train_clock = np.full(len(train), np.nan, dtype=float)
    for fit in age_tuning_fits:
        if fit["alpha"] == selected_alpha:
            positions = train.index[train["row_id"].isin(fit["valid_row_ids"])].to_numpy()
            expected_ids = train.iloc[positions]["row_id"].to_numpy(int)
            if not np.array_equal(expected_ids, fit["valid_row_ids"]):
                raise RuntimeError("Selected age cross-fit row order changed")
            train_clock[positions] = fit["valid_prediction"]
    if not np.isfinite(train_clock).all():
        raise RuntimeError("Incomplete cross-fitted training clock score")

    outer_age_model = fit_ridge_age(train, features, selected_alpha)
    test_clock = predict_ridge_age(outer_age_model, test, features)

    outcome_strata = make_outcome_strata(
        train["death_5y"].to_numpy(int),
        train["age_years"].to_numpy(float),
        N_INNER_FOLDS,
    )
    outcome_splitter = StratifiedKFold(
        n_splits=N_INNER_FOLDS,
        shuffle=True,
        random_state=OUTCOME_INNER_SEED + outer_fold,
    )
    outcome_splits = [(tr, va) for tr, va in outcome_splitter.split(np.zeros(len(train)), outcome_strata)]
    outcome_tuning_fits: list[dict[str, Any]] = []
    outcome_tuning_rows: list[dict[str, Any]] = []
    for c_value in C_GRID:
        for inner_fold, (inner_train, inner_valid) in enumerate(outcome_splits):
            model = fit_clock_outcome(train.iloc[inner_train], train_clock[inner_train], c_value)
            probability = predict_clock_outcome(model, train.iloc[inner_valid], train_clock[inner_valid])
            loss = float(log_loss(train.iloc[inner_valid]["death_5y"], probability, labels=[0, 1]))
            fit = {
                "C": float(c_value),
                "inner_fold": inner_fold,
                "train_row_ids": train.iloc[inner_train]["row_id"].to_numpy(int),
                "valid_row_ids": train.iloc[inner_valid]["row_id"].to_numpy(int),
                "train_clock": train_clock[inner_train],
                "valid_clock": train_clock[inner_valid],
                "model": model,
                "valid_probability": probability,
            }
            outcome_tuning_fits.append(fit)
            outcome_tuning_rows.append({"C": float(c_value), "inner_fold": inner_fold, "log_loss": loss})
    outcome_tuning = pd.DataFrame(outcome_tuning_rows)
    mean_outcome_loss = outcome_tuning.groupby("C", as_index=False)["log_loss"].mean().sort_values(["log_loss", "C"])
    selected_c = float(mean_outcome_loss.iloc[0]["C"])
    outer_outcome_model = fit_clock_outcome(train, train_clock, selected_c)
    test_probability = predict_clock_outcome(outer_outcome_model, test, test_clock)

    base_signature = stable_hash(
        {
            "analysis_signature": analysis_signature,
            "outer_fold": outer_fold,
            "train_row_ids": train["row_id"].tolist(),
            "test_row_ids": test["row_id"].tolist(),
            "features": features,
            "alpha_grid": ALPHA_GRID,
            "C_grid": C_GRID,
            "age_inner_seed": AGE_INNER_SEED + outer_fold,
            "outcome_inner_seed": OUTCOME_INNER_SEED + outer_fold,
        }
    )
    return {
        "schema_version": SCHEMA_VERSION,
        "analysis_signature": analysis_signature,
        "base_signature": base_signature,
        "outer_fold": outer_fold,
        "features": features,
        "train_row_ids": train["row_id"].to_numpy(int),
        "test_row_ids": test["row_id"].to_numpy(int),
        "selected_alpha": selected_alpha,
        "selected_C": selected_c,
        "age_tuning_fits": age_tuning_fits,
        "age_tuning_rows": age_tuning_rows,
        "train_clock": train_clock,
        "outer_age_model": outer_age_model,
        "test_clock": test_clock,
        "outcome_tuning_fits": outcome_tuning_fits,
        "outcome_tuning_rows": outcome_tuning_rows,
        "outer_outcome_model": outer_outcome_model,
        "test_probability": test_probability,
    }


def verify_outer_artifact(
    artifact: dict[str, Any],
    frame_by_row: pd.DataFrame,
    analysis_signature: str,
) -> tuple[float, float, np.ndarray, np.ndarray]:
    if artifact.get("analysis_signature") != analysis_signature:
        raise RuntimeError("Artifact analysis signature mismatch")
    features = list(artifact["features"])
    train = frame_by_row.loc[np.asarray(artifact["train_row_ids"], dtype=int)].copy()
    test = frame_by_row.loc[np.asarray(artifact["test_row_ids"], dtype=int)].copy()
    max_age_error = 0.0
    for fit in artifact["age_tuning_fits"]:
        valid = frame_by_row.loc[np.asarray(fit["valid_row_ids"], dtype=int)]
        prediction = predict_ridge_age(fit["model"], valid, features)
        error = float(np.max(np.abs(prediction - np.asarray(fit["valid_prediction"], dtype=float))))
        max_age_error = max(max_age_error, error)
    test_clock = predict_ridge_age(artifact["outer_age_model"], test, features)
    max_age_error = max(max_age_error, float(np.max(np.abs(test_clock - np.asarray(artifact["test_clock"], dtype=float)))))

    max_outcome_error = 0.0
    for fit in artifact["outcome_tuning_fits"]:
        valid = frame_by_row.loc[np.asarray(fit["valid_row_ids"], dtype=int)]
        probability = predict_clock_outcome(fit["model"], valid, np.asarray(fit["valid_clock"], dtype=float))
        error = float(np.max(np.abs(probability - np.asarray(fit["valid_probability"], dtype=float))))
        max_outcome_error = max(max_outcome_error, error)
    test_probability = predict_clock_outcome(artifact["outer_outcome_model"], test, test_clock)
    max_outcome_error = max(
        max_outcome_error,
        float(np.max(np.abs(test_probability - np.asarray(artifact["test_probability"], dtype=float)))),
    )
    if max_age_error > PARITY_TOLERANCE or max_outcome_error > PARITY_TOLERANCE:
        raise RuntimeError(
            f"STOP_DATA_OR_ARTIFACT_FAIL: reload parity failed age={max_age_error} outcome={max_outcome_error}"
        )
    if not np.array_equal(train["row_id"].to_numpy(int), np.asarray(artifact["train_row_ids"], dtype=int)):
        raise RuntimeError("Artifact train-row order mismatch")
    return max_age_error, max_outcome_error, test_clock, test_probability


def fit_or_load_outer_artifact(
    frame: pd.DataFrame,
    features: list[str],
    outer_fold: int,
    analysis_signature: str,
) -> tuple[dict[str, Any], bool, float, float, np.ndarray, np.ndarray]:
    path = MODEL_DIR / f"outer_fold_{outer_fold:02d}.joblib"
    if path.exists():
        artifact = joblib.load(path)
        reused = True
    else:
        artifact = fit_outer_artifact(frame, features, outer_fold, analysis_signature)
        path.parent.mkdir(parents=True, exist_ok=True)
        joblib.dump(artifact, path, compress=3)
        reused = False
    reloaded = joblib.load(path)
    frame_by_row = frame.set_index("row_id", drop=False)
    age_error, outcome_error, test_clock, test_probability = verify_outer_artifact(
        reloaded,
        frame_by_row,
        analysis_signature,
    )
    return reloaded, reused, age_error, outcome_error, test_clock, test_probability


def summarize(
    frame: pd.DataFrame,
    source_oof: pd.DataFrame,
    clock_score: np.ndarray,
    clock_probability: np.ndarray,
    bootstrap_indices: np.ndarray,
) -> tuple[pd.DataFrame, pd.DataFrame, pd.DataFrame, dict[str, Any]]:
    y = frame["death_5y"].to_numpy(int)
    baseline = source_oof["M_WA"].to_numpy(float)
    direct = source_oof["M_S3A"].to_numpy(float)
    probabilities = {"M_WA": baseline, "M_CLOCK": clock_probability, "M_X": direct}
    pooled = pd.DataFrame(
        [{"model": model, "n": len(frame), "events": int(y.sum()), **endpoint_metrics(y, probability)} for model, probability in probabilities.items()]
    )
    losses = {model: participant_loss(y, probability) for model, probability in probabilities.items()}
    contrast_vectors = {
        "V_CLOCK": losses["M_WA"] - losses["M_CLOCK"],
        "V_SOURCE": losses["M_CLOCK"] - losses["M_X"],
        "V_X": losses["M_WA"] - losses["M_X"],
    }
    boot = {name: vector[bootstrap_indices].mean(axis=1) for name, vector in contrast_vectors.items()}
    contrast_rows = []
    for name, vector in contrast_vectors.items():
        values = boot[name]
        contrast_rows.append(
            {
                "contrast": name,
                "estimate": float(vector.mean()),
                "q025": float(np.quantile(values, 0.025)),
                "q975": float(np.quantile(values, 0.975)),
            }
        )
    contrasts = pd.DataFrame(contrast_rows)

    fold_rows = []
    for outer_fold in range(N_OUTER_FOLDS):
        mask = frame["outer_fold"].eq(outer_fold).to_numpy()
        row = {"outer_fold": outer_fold, "n": int(mask.sum()), "events": int(y[mask].sum())}
        for model, probability in probabilities.items():
            row[f"{model}_log_loss"] = float(log_loss(y[mask], probability[mask], labels=[0, 1]))
        row["V_CLOCK"] = row["M_WA_log_loss"] - row["M_CLOCK_log_loss"]
        row["V_SOURCE"] = row["M_CLOCK_log_loss"] - row["M_X_log_loss"]
        row["V_X"] = row["M_WA_log_loss"] - row["M_X_log_loss"]
        fold_rows.append(row)
    fold_metrics = pd.DataFrame(fold_rows)

    age = frame["age_years"].to_numpy(float)
    age_residual = clock_score - age
    slope, intercept = np.polyfit(clock_score, age, 1)
    age_diagnostics = {
        "n": len(age),
        "r2": float(r2_score(age, clock_score)),
        "rmse": float(mean_squared_error(age, clock_score) ** 0.5),
        "mae": float(mean_absolute_error(age, clock_score)),
        "pearson_r": float(np.corrcoef(age, clock_score)[0, 1]),
        "observed_on_score_slope": float(slope),
        "observed_on_score_intercept": float(intercept),
        "residual_mean": float(age_residual.mean()),
        "residual_sd": float(age_residual.std(ddof=1)),
    }
    v_source = contrasts.set_index("contrast").loc["V_SOURCE"]
    positive_folds = int((fold_metrics["V_SOURCE"] > 0).sum())
    if float(v_source["estimate"]) > 0 and float(v_source["q025"]) > 0 and positive_folds >= 4:
        category = "PROMISING_SOURCE_INFORMATION"
    elif float(v_source["estimate"]) > 0:
        category = "WEAK_SOURCE_INFORMATION"
    elif float(v_source["q975"]) <= 0:
        category = "ABSENT_SOURCE_ADVANTAGE"
    else:
        category = "AMBIGUOUS_FINITE_SAMPLE_COMPARISON"
    decision = {
        "decision_category": category,
        "endpoint": "five_year_all_cause_mortality",
        "population": "NHANES 2007-2012 complete-support 50-measurement interface, ages 20-79",
        "n": len(frame),
        "events": int(y.sum()),
        "age_decoder": "fold-local ridge regression of age on W and fixed S3 measurements",
        "age_decoder_diagnostics": age_diagnostics,
        "V_CLOCK": float(contrasts.set_index("contrast").loc["V_CLOCK", "estimate"]),
        "V_SOURCE": float(v_source["estimate"]),
        "V_SOURCE_q025": float(v_source["q025"]),
        "V_SOURCE_q975": float(v_source["q975"]),
        "V_X": float(contrasts.set_index("contrast").loc["V_X", "estimate"]),
        "positive_V_SOURCE_folds": positive_folds,
        "n_bootstraps": int(bootstrap_indices.shape[0]),
        "claim_boundary": "Finite-sample target-specific comparison; not universal estimator dominance or biological-age identification.",
    }
    return pooled, contrasts, fold_metrics, decision


def markdown_table(frame: pd.DataFrame, digits: int = 6) -> str:
    lines = [
        "| " + " | ".join(map(str, frame.columns)) + " |",
        "| " + " | ".join(["---"] * len(frame.columns)) + " |",
    ]
    for row in frame.itertuples(index=False, name=None):
        values = []
        for value in row:
            if isinstance(value, (float, np.floating)):
                values.append("" if not np.isfinite(value) else f"{float(value):.{digits}g}")
            else:
                values.append(str(value))
        lines.append("| " + " | ".join(values) + " |")
    return "\n".join(lines)


def write_report(
    pooled: pd.DataFrame,
    contrasts: pd.DataFrame,
    fold_metrics: pd.DataFrame,
    decision: dict[str, Any],
    audit: dict[str, Any],
) -> None:
    age = decision["age_decoder_diagnostics"]
    source = contrasts.set_index("contrast").loc["V_SOURCE"]
    text = f"""# NHANES five-year age-decoder source-information probe

Date: 23 August 2026

## Decision

**{decision['decision_category']}**

The pure chronological-age decoder achieved held-out age R2={age['r2']:.4f}, RMSE={age['rmse']:.3f} years and MAE={age['mae']:.3f} years. For complete-support five-year mortality, direct use of age, background and the same 50 measurements improved pooled out-of-fold log loss beyond age, background and the clock by `V_SOURCE={source['estimate']:.6f}` (paired fixed-prediction bootstrap interval {source['q025']:.6f} to {source['q975']:.6f}); {decision['positive_V_SOURCE_folds']} of 5 fold contrasts were positive.

This is a finite-sample result for one linear age decoder, one endpoint, one population, one panel and one direct model family. It does not prove universal finite-sample superiority and does not identify biological age.

## Endpoint models

{markdown_table(pooled)}

## Paired log-loss contrasts

{markdown_table(contrasts)}

- `V_CLOCK = L(M_WA)-L(M_CLOCK)`.
- `V_SOURCE = L(M_CLOCK)-L(M_X)`.
- `V_X = L(M_WA)-L(M_X)`.

## Fold dispersion

{markdown_table(fold_metrics)}

## Age-decoder diagnostics

```json
{json.dumps(json_ready(age), indent=2, ensure_ascii=False)}
```

## Persistence and parity

- Outer artifacts: {audit['artifact_count']} of {audit['expected_artifact_count']}.
- Newly fitted: {audit['newly_fitted_artifact_count']}; reused: {audit['reused_artifact_count']}.
- Age tuning estimators: {audit['age_tuning_estimator_count']}.
- Outcome tuning estimators: {audit['outcome_tuning_estimator_count']}.
- Maximum reloaded age prediction error: {audit['max_age_prediction_error']:.3g}.
- Maximum reloaded endpoint probability error: {audit['max_outcome_prediction_error']:.3g}.

The 2,000 paired intervals reuse the accepted predecessor bootstrap indices and condition on stored out-of-fold predictions. They omit refitting and study-development uncertainty.
"""
    (OUT / "REPORT.md").write_text(text, encoding="utf-8")


def write_artifact_manifest() -> None:
    rows = []
    for path in sorted(OUT.rglob("*")):
        if path.is_file() and path.name != "artifact_manifest.tsv":
            rows.append(
                {
                    "path": str(path.relative_to(ROOT)),
                    "bytes": path.stat().st_size,
                    "sha256": sha256(path),
                }
            )
    pd.DataFrame(rows).to_csv(OUT / "artifact_manifest.tsv", sep="\t", index=False)


def main() -> int:
    configure_stdout()
    OUT.mkdir(parents=True, exist_ok=True)
    MODEL_DIR.mkdir(parents=True, exist_ok=True)
    required_paths = (
        SOURCE_FRAME,
        SOURCE_FOLDS,
        SOURCE_TIERS,
        SOURCE_OOF,
        SOURCE_LOSSES,
        SOURCE_BOOTSTRAP,
        SOURCE_SIGNATURE,
        SOURCE_MODEL_AUDIT,
        SPEC,
        ENV_POLICY,
        ENV_MANIFEST,
        ENV_LOCK,
        Path(__file__),
    )
    for path in required_paths:
        if not path.exists():
            raise RuntimeError(f"Required file missing: {path}")

    input_roles = {
        "analysis_ready_source": SOURCE_FRAME,
        "participant_folds": SOURCE_FOLDS,
        "ordered_feature_tiers": SOURCE_TIERS,
        "source_oof_predictions": SOURCE_OOF,
        "source_participant_losses": SOURCE_LOSSES,
        "source_bootstrap_indices": SOURCE_BOOTSTRAP,
        "source_analysis_signature": SOURCE_SIGNATURE,
        "source_model_audit": SOURCE_MODEL_AUDIT,
        "frozen_specification": SPEC,
        "analysis_script": Path(__file__),
        "environment_policy": ENV_POLICY,
        "environment_manifest": ENV_MANIFEST,
        "environment_lock": ENV_LOCK,
    }
    input_manifest = pd.DataFrame(
        [
            {"role": role, "path": str(path.relative_to(ROOT)), "bytes": path.stat().st_size, "sha256": sha256(path)}
            for role, path in input_roles.items()
        ]
    )
    signature_payload = {
        "analysis_version": ANALYSIS_VERSION,
        "schema_version": SCHEMA_VERSION,
        "inputs": input_manifest[["role", "sha256"]].to_dict(orient="records"),
        "models": ["M_WA", "M_CLOCK", "M_X"],
        "features": "frozen_S3_50_measurements",
        "age_decoder": "ridge",
        "alpha_grid": ALPHA_GRID,
        "clock_endpoint_model": "ridge_logistic",
        "C_grid": C_GRID,
        "outer_folds": N_OUTER_FOLDS,
        "age_inner_folds": N_INNER_FOLDS,
        "outcome_inner_folds": N_INNER_FOLDS,
        "age_inner_seed": AGE_INNER_SEED,
        "outcome_inner_seed": OUTCOME_INNER_SEED,
    }
    analysis_signature = stable_hash(signature_payload)
    signature_path = OUT / "analysis_signature.json"
    if signature_path.exists():
        existing = json.loads(signature_path.read_text(encoding="utf-8"))
        if existing.get("analysis_signature") != analysis_signature:
            raise RuntimeError("Existing output has an incompatible analysis signature")
    else:
        write_json(signature_path, {"analysis_signature": analysis_signature, "signature_payload": signature_payload})
    input_manifest.to_csv(OUT / "input_manifest.tsv", sep="\t", index=False)

    frame, features, source_oof, bootstrap_indices, source_audit = load_and_gate()
    frame[["row_id", "SEQN", "cycle", "age_years", "death_5y", "outer_fold"]].to_csv(
        OUT / "participant_folds.tsv", sep="\t", index=False
    )
    pd.DataFrame({"order": np.arange(1, len(features) + 1), "feature": features}).to_csv(
        OUT / "ordered_source_features.tsv", sep="\t", index=False
    )

    clock_score = np.full(len(frame), np.nan, dtype=float)
    clock_probability = np.full(len(frame), np.nan, dtype=float)
    index_rows = []
    tuning_rows = []
    max_age_error = 0.0
    max_outcome_error = 0.0
    reused_count = 0
    fitted_count = 0
    for outer_fold in range(N_OUTER_FOLDS):
        print(f"Outer fold {outer_fold + 1}/{N_OUTER_FOLDS}", flush=True)
        artifact, reused, age_error, outcome_error, test_score, test_probability = fit_or_load_outer_artifact(
            frame,
            features,
            outer_fold,
            analysis_signature,
        )
        mask = frame["outer_fold"].eq(outer_fold).to_numpy()
        clock_score[mask] = test_score
        clock_probability[mask] = test_probability
        max_age_error = max(max_age_error, age_error)
        max_outcome_error = max(max_outcome_error, outcome_error)
        reused_count += int(reused)
        fitted_count += int(not reused)
        path = MODEL_DIR / f"outer_fold_{outer_fold:02d}.joblib"
        index_rows.append(
            {
                "outer_fold": outer_fold,
                "selected_alpha": artifact["selected_alpha"],
                "selected_C": artifact["selected_C"],
                "n_train": len(artifact["train_row_ids"]),
                "n_test": len(artifact["test_row_ids"]),
                "artifact_path": str(path.relative_to(ROOT)),
                "artifact_sha256": sha256(path),
                "reused": reused,
                "max_age_prediction_error": age_error,
                "max_outcome_prediction_error": outcome_error,
            }
        )
        for row in artifact["age_tuning_rows"]:
            tuning_rows.append({"outer_fold": outer_fold, "stage": "age", **row})
        for row in artifact["outcome_tuning_rows"]:
            tuning_rows.append({"outer_fold": outer_fold, "stage": "outcome", **row})
        pd.DataFrame(
            {
                "row_id": artifact["train_row_ids"],
                "outer_fold": outer_fold,
                "crossfit_clock_score": artifact["train_clock"],
            }
        ).to_csv(OUT / f"outer_fold_{outer_fold:02d}_training_crossfit_scores.tsv.gz", sep="\t", index=False, compression="gzip")

    if not np.isfinite(clock_score).all() or not np.isfinite(clock_probability).all():
        raise RuntimeError("STOP_DATA_OR_ARTIFACT_FAIL: incomplete OOF output")
    model_index = pd.DataFrame(index_rows)
    model_index.to_csv(OUT / "model_artifact_index.tsv", sep="\t", index=False)
    pd.DataFrame(tuning_rows).to_csv(OUT / "inner_tuning.tsv.gz", sep="\t", index=False, compression="gzip")

    pooled, contrasts, fold_metrics, decision = summarize(
        frame,
        source_oof,
        clock_score,
        clock_probability,
        bootstrap_indices,
    )
    y = frame["death_5y"].to_numpy(int)
    baseline = source_oof["M_WA"].to_numpy(float)
    direct = source_oof["M_S3A"].to_numpy(float)
    oof = frame[["row_id", "SEQN", "cycle", "age_years", "death_5y", "outer_fold"]].copy()
    oof["clock_score"] = clock_score
    oof["clock_gap"] = clock_score - oof["age_years"].to_numpy(float)
    oof["M_WA"] = baseline
    oof["M_CLOCK"] = clock_probability
    oof["M_X"] = direct
    oof.to_csv(OUT / "oof_predictions.tsv.gz", sep="\t", index=False, compression="gzip")
    losses = oof[["row_id", "SEQN", "death_5y", "outer_fold"]].copy()
    losses["M_WA"] = participant_loss(y, baseline)
    losses["M_CLOCK"] = participant_loss(y, clock_probability)
    losses["M_X"] = participant_loss(y, direct)
    losses.to_csv(OUT / "participant_loss_vectors.tsv.gz", sep="\t", index=False, compression="gzip")
    pooled.to_csv(OUT / "pooled_metrics.tsv", sep="\t", index=False)
    contrasts.to_csv(OUT / "paired_contrasts.tsv", sep="\t", index=False)
    fold_metrics.to_csv(OUT / "fold_metrics.tsv", sep="\t", index=False)

    age_band = pd.cut(frame["age_years"], bins=[19, 29, 39, 49, 59, 69, 79], labels=["20-29", "30-39", "40-49", "50-59", "60-69", "70-79"])
    age_band_summary = (
        pd.DataFrame({"age_band": age_band, "age": frame["age_years"], "clock_score": clock_score, "gap": clock_score - frame["age_years"]})
        .groupby("age_band", observed=True)
        .agg(n=("age", "size"), mean_age=("age", "mean"), mean_clock=("clock_score", "mean"), mean_gap=("gap", "mean"), sd_gap=("gap", "std"))
        .reset_index()
    )
    age_band_summary.to_csv(OUT / "age_band_diagnostics.tsv", sep="\t", index=False)
    write_json(OUT / "age_decoder_diagnostics.json", decision["age_decoder_diagnostics"])

    audit = {
        "status": "PASS",
        "analysis_signature": analysis_signature,
        "source_analysis_signature": source_audit["analysis_signature"],
        "expected_artifact_count": N_OUTER_FOLDS,
        "artifact_count": int(len(model_index)),
        "newly_fitted_artifact_count": int(fitted_count),
        "reused_artifact_count": int(reused_count),
        "age_tuning_estimator_count": N_OUTER_FOLDS * len(ALPHA_GRID) * N_INNER_FOLDS,
        "outcome_tuning_estimator_count": N_OUTER_FOLDS * len(C_GRID) * N_INNER_FOLDS,
        "outer_final_estimator_count": N_OUTER_FOLDS * 2,
        "parity_tolerance": PARITY_TOLERANCE,
        "max_age_prediction_error": max_age_error,
        "max_outcome_prediction_error": max_outcome_error,
        "all_model_hashes_present": bool(model_index["artifact_sha256"].str.len().eq(64).all()),
        "software": {
            "python": sys.version,
            "python_executable": sys.executable,
            "platform": platform.platform(),
            "numpy": np.__version__,
            "pandas": pd.__version__,
            "scikit_learn": sklearn.__version__,
            "joblib": joblib.__version__,
        },
    }
    if (
        audit["artifact_count"] != audit["expected_artifact_count"]
        or max_age_error > PARITY_TOLERANCE
        or max_outcome_error > PARITY_TOLERANCE
        or not audit["all_model_hashes_present"]
    ):
        audit["status"] = "FAIL"
        decision["decision_category"] = "STOP_DATA_OR_ARTIFACT_FAIL"
    write_json(OUT / "model_artifact_audit.json", audit)
    write_json(OUT / "decision.json", decision)
    write_report(pooled, contrasts, fold_metrics, decision, audit)
    write_artifact_manifest()

    print(json.dumps(json_ready(decision), indent=2, ensure_ascii=False), flush=True)
    print(f"Wrote {OUT}", flush=True)
    return 0 if decision["decision_category"] != "STOP_DATA_OR_ARTIFACT_FAIL" else 2


if __name__ == "__main__":
    raise SystemExit(main())
````
