## Supplementary_Software_1 for "Inferential boundaries of age prediction: why prediction does not establish biological age measurement": analyze_nhanes_5y_complete_support_contextual_age_probe_v0_1.py

```
#!/usr/bin/env python
"""Five-year complete-support NHANES contextual-age decision probe.

Implements NHANES_5Y_COMPLETE_SUPPORT_CONTEXTUAL_AGE_PROBE_SPEC_20260823.md.
The script fits exactly eight ridge-logistic model specifications over one
five-year endpoint and persists every inner and outer fitted object used by
the analysis. Compatible artifacts are reloaded and reused on rerun.
"""

from __future__ import annotations

import hashlib
import json
import math
import os
import platform
import sys
import warnings
from dataclasses import asdict, dataclass
from pathlib import Path
from typing import Any

import joblib
import matplotlib.pyplot as plt
import numpy as np
import pandas as pd
import sklearn
from sklearn.exceptions import ConvergenceWarning
from sklearn.linear_model import LogisticRegression
from sklearn.metrics import brier_score_loss, log_loss, roc_auc_score
from sklearn.model_selection import StratifiedKFold
from sklearn.preprocessing import OneHotEncoder, SplineTransformer, StandardScaler


ROOT = Path(__file__).resolve().parents[1]
ANALYSIS = ROOT / "analysis"
PROCESSED = ROOT / "data" / "processed"
OUT = ROOT / "outputs" / "nhanes_5y_complete_support_contextual_age_probe_v0_1"
MODEL_DIR = OUT / "models"
REPORT = OUT / "REPORT.md"
PANEL = PROCESSED / "NHANES_broad_clinical_panel_v0_1.tsv.gz"
FEATURE_MANIFEST = PROCESSED / "NHANES_broad_clinical_feature_manifest_v0_1.tsv"
SPEC = ANALYSIS / "NHANES_5Y_COMPLETE_SUPPORT_CONTEXTUAL_AGE_PROBE_SPEC_20260823.md"
AUDIT_SCRIPT = ANALYSIS / "audit_nhanes_5y_complete_support_contextual_age_probe_v0_1.py"
ENV_POLICY = ROOT / "environment-policy.json"
ENV_MANIFEST = ROOT / "environment.yml"
ENV_LOCK = ROOT / "environment.win-64.lock.txt"

ANALYSIS_VERSION = "nhanes_5y_complete_support_contextual_age_probe_v0_1"
SCHEMA_VERSION = "1.0"
ALLOWED_CYCLES = ("2007_2008", "2009_2010", "2011_2012")
EXPECTED_N = 13_724
EXPECTED_EVENTS = 470
OUTER_SEED = 20260823
BOOTSTRAP_SEED = 20260824
N_OUTER_FOLDS = 5
N_INNER_FOLDS = 3
N_BOOTSTRAPS = 2000
PARITY_TOLERANCE = 1e-12
EPSILON = 1e-7
C_GRID = (0.001, 0.003, 0.01, 0.03, 0.1, 0.3, 1.0, 3.0, 10.0, 30.0, 100.0, 300.0)
BACKGROUND = ("sex", "race_ethnicity", "cycle")
AGE_BAND_LABELS = ("20-29", "30-39", "40-49", "50-59", "60-69", "70-79")


@dataclass(frozen=True)
class ModelSpec:
    name: str
    tier: str
    include_age: bool


MODEL_SPECS = (
    ModelSpec("M_W", "S0", False),
    ModelSpec("M_WA", "S0", True),
    ModelSpec("M_S1", "S1", False),
    ModelSpec("M_S1A", "S1", True),
    ModelSpec("M_S2c", "S2c", False),
    ModelSpec("M_S2cA", "S2c", True),
    ModelSpec("M_S3", "S3", False),
    ModelSpec("M_S3A", "S3", True),
)

AGE_VALUE_PAIRS = {
    "S0": ("M_W", "M_WA"),
    "S1": ("M_S1", "M_S1A"),
    "S2c": ("M_S2c", "M_S2cA"),
    "S3": ("M_S3", "M_S3A"),
}


def configure_stdout() -> None:
    if hasattr(sys.stdout, "reconfigure"):
        sys.stdout.reconfigure(encoding="utf-8")


def sha256(path: Path) -> str:
    digest = hashlib.sha256()
    with path.open("rb") as handle:
        for block in iter(lambda: handle.read(1024 * 1024), b""):
            digest.update(block)
    return digest.hexdigest()


def stable_hash(payload: Any) -> str:
    encoded = json.dumps(payload, sort_keys=True, separators=(",", ":"), ensure_ascii=False).encode("utf-8")
    return hashlib.sha256(encoded).hexdigest()


def json_ready(value: Any) -> Any:
    if isinstance(value, dict):
        return {str(key): json_ready(item) for key, item in value.items()}
    if isinstance(value, (list, tuple)):
        return [json_ready(item) for item in value]
    if isinstance(value, Path):
        return str(value)
    if isinstance(value, (np.integer,)):
        return int(value)
    if isinstance(value, (np.floating,)):
        return None if not np.isfinite(value) else float(value)
    if isinstance(value, np.ndarray):
        return value.tolist()
    if isinstance(value, float):
        return None if not math.isfinite(value) else value
    return value


def write_json(path: Path, payload: Any) -> None:
    path.write_text(json.dumps(json_ready(payload), indent=2, ensure_ascii=False) + "\n", encoding="utf-8")


def clean_categories(frame: pd.DataFrame) -> pd.DataFrame:
    return frame.fillna("missing").astype(str).replace("", "missing")


def age_band(values: pd.Series | np.ndarray) -> np.ndarray:
    age = np.asarray(values, dtype=float)
    return np.clip(((age - 20.0) // 10.0).astype(int), 0, 5)


def make_strata(frame: pd.DataFrame, y: np.ndarray, n_splits: int) -> tuple[np.ndarray, str]:
    age = pd.to_numeric(frame["age_years"], errors="raise").reset_index(drop=True)
    quintile = pd.qcut(age, q=5, duplicates="drop", labels=False)
    quintile = quintile.fillna(-1).astype(int).to_numpy()
    combined = np.char.add(np.char.add(np.asarray(y).astype(str), "_"), quintile.astype(str))
    counts = pd.Series(combined).value_counts()
    if len(counts) >= 2 and int(counts.min()) >= n_splits:
        return combined, "endpoint_plus_training_age_quintile"
    return np.asarray(y).astype(str), "endpoint_only_fallback"


def load_tiers(manifest: pd.DataFrame) -> dict[str, list[str]]:
    required = {"feature", "transformed_column", "source_module"}
    if not required.issubset(manifest.columns):
        raise RuntimeError(f"Feature manifest missing {sorted(required - set(manifest.columns))}")
    s1 = manifest["source_module"].isin(["BMX", "BPX"])
    s2c = manifest["source_module"].isin(["BMX", "BPX", "BIOPRO", "HDL"])
    tiers = {
        "S0": [],
        "S1": manifest.loc[s1, "transformed_column"].tolist(),
        "S2c": manifest.loc[s2c, "transformed_column"].tolist(),
        "S3": manifest["transformed_column"].tolist(),
    }
    observed = {key: len(value) for key, value in tiers.items()}
    expected = {"S0": 0, "S1": 6, "S2c": 30, "S3": 50}
    if observed != expected:
        raise RuntimeError(f"Unexpected ordered tier sizes: {observed}")
    if not set(tiers["S1"]).issubset(tiers["S2c"]):
        raise RuntimeError("S1 is not nested in S2c")
    if not set(tiers["S2c"]).issubset(tiers["S3"]):
        raise RuntimeError("S2c is not nested in S3")
    forbidden = ("age", "clock", "gap", "resid")
    bad = [column for column in tiers["S3"] if any(token in column.lower() for token in forbidden)]
    if bad:
        raise RuntimeError(f"Age-derived columns found: {bad}")
    return tiers


def load_and_gate() -> tuple[pd.DataFrame, pd.DataFrame, pd.DataFrame, dict[str, list[str]]]:
    frame = pd.read_csv(PANEL, sep="\t")
    manifest = pd.read_csv(FEATURE_MANIFEST, sep="\t")
    tiers = load_tiers(manifest)
    required = ["SEQN", "cycle", "age_years", *BACKGROUND, "event", "time_months", *tiers["S3"]]
    missing_columns = sorted(set(required) - set(frame.columns))
    if missing_columns:
        raise RuntimeError(f"Missing input columns: {missing_columns}")
    base = frame.loc[
        frame["cycle"].isin(ALLOWED_CYCLES) & frame["age_years"].between(20, 79, inclusive="both"),
        list(dict.fromkeys(required)),
    ].copy()
    base["death_5y"] = (base["event"].eq(1) & base["time_months"].le(60)).astype(int)
    base["event_free_censored_before_60"] = (base["event"].eq(0) & base["time_months"].lt(60)).astype(int)
    base["known_event_free_60"] = (base["time_months"].ge(60) & base["death_5y"].eq(0)).astype(int)
    support_rows = []
    for cycle in ALLOWED_CYCLES:
        chunk = base.loc[base["cycle"].eq(cycle)]
        event_free = chunk.loc[chunk["event"].eq(0), "time_months"]
        support_rows.append(
            {
                "cycle": cycle,
                "baseline_n": int(len(chunk)),
                "death_by_60m": int(chunk["death_5y"].sum()),
                "known_event_free_60m": int(chunk["known_event_free_60"].sum()),
                "event_free_censored_before_60m": int(chunk["event_free_censored_before_60"].sum()),
                "event_free_followup_min_months": float(event_free.min()),
                "event_free_followup_max_months": float(event_free.max()),
                "complete_support": bool(chunk["event_free_censored_before_60"].sum() == 0),
            }
        )
    support = pd.DataFrame(support_rows)
    if not support["complete_support"].all():
        raise RuntimeError("STOP_DATA_OR_AUDIT_FAIL: incomplete five-year cycle support")
    risk = base.loc[base["death_5y"].eq(1) | base["time_months"].ge(60)].copy().reset_index(drop=True)
    risk.insert(0, "row_id", np.arange(len(risk), dtype=int))
    model_columns = ["SEQN", "cycle", "age_years", *BACKGROUND, "death_5y", *tiers["S3"]]
    model_columns = list(dict.fromkeys(model_columns))
    if risk[model_columns].isna().any().any():
        raise RuntimeError("STOP_DATA_OR_AUDIT_FAIL: missing model inputs")
    if risk["SEQN"].duplicated().any():
        raise RuntimeError("STOP_DATA_OR_AUDIT_FAIL: duplicate SEQN")
    if len(risk) != EXPECTED_N or int(risk["death_5y"].sum()) != EXPECTED_EVENTS:
        raise RuntimeError(
            f"STOP_DATA_OR_AUDIT_FAIL: expected {EXPECTED_N}/{EXPECTED_EVENTS}, "
            f"observed {len(risk)}/{int(risk['death_5y'].sum())}"
        )
    if int(risk["event_free_censored_before_60"].sum()) != 0:
        raise RuntimeError("STOP_DATA_OR_AUDIT_FAIL: early event-free censoring remains")
    return risk, support, manifest, tiers


def assign_outer_folds(frame: pd.DataFrame) -> tuple[np.ndarray, str]:
    y = frame["death_5y"].to_numpy(int)
    strata, strategy = make_strata(frame, y, N_OUTER_FOLDS)
    splitter = StratifiedKFold(n_splits=N_OUTER_FOLDS, shuffle=True, random_state=OUTER_SEED)
    folds = np.full(len(frame), -1, dtype=int)
    for fold, (_, test_index) in enumerate(splitter.split(frame, strata)):
        folds[test_index] = fold
    if np.any(folds < 0):
        raise RuntimeError("Incomplete outer fold assignment")
    return folds, strategy


def inner_splits(outer_train: pd.DataFrame, outer_fold: int) -> tuple[list[tuple[np.ndarray, np.ndarray]], str]:
    y = outer_train["death_5y"].to_numpy(int)
    strata, strategy = make_strata(outer_train, y, N_INNER_FOLDS)
    splitter = StratifiedKFold(
        n_splits=N_INNER_FOLDS,
        shuffle=True,
        random_state=OUTER_SEED + 100 + outer_fold,
    )
    return list(splitter.split(outer_train, strata)), strategy


def fit_preprocessor(frame: pd.DataFrame, spec: ModelSpec, state_columns: list[str]) -> dict[str, Any]:
    encoder = OneHotEncoder(handle_unknown="ignore", drop="first", sparse_output=False)
    with warnings.catch_warnings():
        warnings.simplefilter("ignore")
        encoder.fit(clean_categories(frame[list(BACKGROUND)]))
    feature_names = encoder.get_feature_names_out(BACKGROUND).tolist()
    state_medians = None
    state_scaler = None
    if state_columns:
        raw = frame[state_columns].apply(pd.to_numeric, errors="coerce")
        state_medians = raw.median(axis=0)
        if state_medians.isna().any():
            raise RuntimeError("State median is non-finite")
        state_scaler = StandardScaler().fit(raw.fillna(state_medians))
        feature_names.extend(state_columns)
    age_median = None
    age_spline = None
    age_scaler = None
    if spec.include_age:
        age = pd.to_numeric(frame["age_years"], errors="coerce")
        age_median = float(age.median())
        values = age.fillna(age_median).to_numpy(float).reshape(-1, 1)
        age_spline = SplineTransformer(
            n_knots=5,
            degree=3,
            include_bias=False,
            extrapolation="linear",
        ).fit(values)
        age_raw = age_spline.transform(values)
        age_scaler = StandardScaler().fit(age_raw)
        feature_names.extend([f"age_spline_{index}" for index in range(age_raw.shape[1])])
    return {
        "background_columns": list(BACKGROUND),
        "encoder": encoder,
        "state_columns": list(state_columns),
        "state_medians": state_medians,
        "state_scaler": state_scaler,
        "include_age": bool(spec.include_age),
        "age_median": age_median,
        "age_spline": age_spline,
        "age_scaler": age_scaler,
        "feature_names": feature_names,
    }


def transform(preprocessor: dict[str, Any], frame: pd.DataFrame) -> np.ndarray:
    blocks = [preprocessor["encoder"].transform(clean_categories(frame[preprocessor["background_columns"]]))]
    state_columns = preprocessor["state_columns"]
    if state_columns:
        raw = frame[state_columns].apply(pd.to_numeric, errors="coerce")
        blocks.append(preprocessor["state_scaler"].transform(raw.fillna(preprocessor["state_medians"])))
    if preprocessor["include_age"]:
        age = pd.to_numeric(frame["age_years"], errors="coerce")
        values = age.fillna(preprocessor["age_median"]).to_numpy(float).reshape(-1, 1)
        age_raw = preprocessor["age_spline"].transform(values)
        blocks.append(preprocessor["age_scaler"].transform(age_raw))
    design = np.column_stack(blocks)
    if not np.isfinite(design).all():
        raise RuntimeError("Non-finite design matrix")
    if design.shape[1] != len(preprocessor["feature_names"]):
        raise RuntimeError("Design feature-order mismatch")
    return design


def fit_estimator(design: np.ndarray, y: np.ndarray, c_value: float) -> LogisticRegression:
    estimator = LogisticRegression(
        C=float(c_value),
        penalty="l2",
        solver="lbfgs",
        fit_intercept=True,
        max_iter=5000,
        random_state=OUTER_SEED,
    )
    with warnings.catch_warnings(record=True) as caught:
        warnings.simplefilter("always", ConvergenceWarning)
        estimator.fit(design, y)
    if any(issubclass(item.category, ConvergenceWarning) for item in caught):
        raise RuntimeError("STOP_DATA_OR_AUDIT_FAIL: logistic model did not converge")
    return estimator


def predict(estimator: LogisticRegression, design: np.ndarray) -> np.ndarray:
    return np.clip(estimator.predict_proba(design)[:, 1], EPSILON, 1.0 - EPSILON)


def artifact_signature(
    analysis_signature: str,
    spec: ModelSpec,
    outer_fold: int,
    train_ids: np.ndarray,
    test_ids: np.ndarray,
    state_columns: list[str],
) -> str:
    payload = {
        "analysis_signature": analysis_signature,
        "model_spec": asdict(spec),
        "outer_fold": outer_fold,
        "train_ids_sha256": stable_hash([int(value) for value in train_ids]),
        "test_ids_sha256": stable_hash([int(value) for value in test_ids]),
        "state_columns": state_columns,
        "C_grid": C_GRID,
        "outer_seed": OUTER_SEED,
        "inner_folds": N_INNER_FOLDS,
    }
    return stable_hash(payload)


def verify_artifact(
    artifact: dict[str, Any],
    frame_by_row: pd.DataFrame,
    expected_signature: str,
) -> tuple[np.ndarray, float, float, int]:
    if artifact.get("artifact_signature") != expected_signature:
        raise RuntimeError("STOP_DATA_OR_AUDIT_FAIL: model signature mismatch")
    test_rows = np.asarray(artifact["outer_test_row_ids"], dtype=int)
    test_frame = frame_by_row.loc[test_rows]
    outer_design = transform(artifact["outer_preprocessor"], test_frame)
    outer_prediction = predict(artifact["outer_estimator"], outer_design)
    stored_outer = np.asarray(artifact["outer_test_predictions"], dtype=float)
    outer_error = float(np.max(np.abs(outer_prediction - stored_outer)))
    inner_error = 0.0
    inner_models = 0
    for inner in artifact["inner_fits"]:
        validation_rows = np.asarray(inner["validation_row_ids"], dtype=int)
        validation_frame = frame_by_row.loc[validation_rows]
        validation_design = transform(inner["preprocessor"], validation_frame)
        for c_key, estimator in inner["estimators"].items():
            observed = predict(estimator, validation_design)
            stored = np.asarray(inner["validation_predictions"][c_key], dtype=float)
            inner_error = max(inner_error, float(np.max(np.abs(observed - stored))))
            inner_models += 1
    if outer_error > PARITY_TOLERANCE or inner_error > PARITY_TOLERANCE:
        raise RuntimeError(
            f"STOP_DATA_OR_AUDIT_FAIL: reload parity outer={outer_error}, inner={inner_error}"
        )
    return outer_prediction, outer_error, inner_error, inner_models


def fit_or_load_artifact(
    frame: pd.DataFrame,
    frame_by_row: pd.DataFrame,
    spec: ModelSpec,
    state_columns: list[str],
    outer_fold: int,
    analysis_signature: str,
) -> tuple[dict[str, Any], bool, float, float, int]:
    train = frame.loc[frame["outer_fold"].ne(outer_fold)].copy()
    test = frame.loc[frame["outer_fold"].eq(outer_fold)].copy()
    train_ids = train["SEQN"].to_numpy(int)
    test_ids = test["SEQN"].to_numpy(int)
    expected_signature = artifact_signature(
        analysis_signature, spec, outer_fold, train_ids, test_ids, state_columns
    )
    fold_dir = MODEL_DIR / f"outer_fold_{outer_fold:02d}"
    fold_dir.mkdir(parents=True, exist_ok=True)
    path = fold_dir / f"{spec.name}.joblib"
    if path.exists():
        artifact = joblib.load(path)
        _, outer_error, inner_error, inner_models = verify_artifact(
            artifact, frame_by_row, expected_signature
        )
        return artifact, True, outer_error, inner_error, inner_models

    split_list, inner_strategy = inner_splits(train, outer_fold)
    tuning_rows: list[dict[str, Any]] = []
    inner_fits: list[dict[str, Any]] = []
    for inner_fold, (inner_train_position, validation_position) in enumerate(split_list):
        inner_train = train.iloc[inner_train_position]
        validation = train.iloc[validation_position]
        preprocessor = fit_preprocessor(inner_train, spec, state_columns)
        inner_design = transform(preprocessor, inner_train)
        validation_design = transform(preprocessor, validation)
        estimators: dict[str, LogisticRegression] = {}
        validation_predictions: dict[str, np.ndarray] = {}
        y_inner = inner_train["death_5y"].to_numpy(int)
        y_validation = validation["death_5y"].to_numpy(int)
        for c_value in C_GRID:
            key = format(float(c_value), ".12g")
            estimator = fit_estimator(inner_design, y_inner, c_value)
            probability = predict(estimator, validation_design)
            estimators[key] = estimator
            validation_predictions[key] = probability
            tuning_rows.append(
                {
                    "outer_fold": outer_fold,
                    "inner_fold": inner_fold,
                    "inner_stratification": inner_strategy,
                    "model": spec.name,
                    "tier": spec.tier,
                    "include_age": spec.include_age,
                    "C": float(c_value),
                    "validation_log_loss": float(log_loss(y_validation, probability, labels=[0, 1])),
                    "n_train": int(len(inner_train)),
                    "n_validation": int(len(validation)),
                    "events_train": int(y_inner.sum()),
                    "events_validation": int(y_validation.sum()),
                    "n_design_features": int(inner_design.shape[1]),
                }
            )
        inner_fits.append(
            {
                "inner_fold": inner_fold,
                "stratification": inner_strategy,
                "train_row_ids": inner_train["row_id"].to_numpy(int),
                "validation_row_ids": validation["row_id"].to_numpy(int),
                "preprocessor": preprocessor,
                "estimators": estimators,
                "validation_predictions": validation_predictions,
            }
        )
    tuning_frame = pd.DataFrame(tuning_rows)
    mean_loss = tuning_frame.groupby("C", as_index=False)["validation_log_loss"].mean()
    mean_loss = mean_loss.sort_values("C", ascending=True).reset_index(drop=True)
    selected_c = float(mean_loss.loc[mean_loss["validation_log_loss"].idxmin(), "C"])
    tuning_frame = tuning_frame.merge(
        mean_loss.rename(columns={"validation_log_loss": "mean_validation_log_loss"}),
        on="C",
        how="left",
        validate="many_to_one",
    )
    tuning_frame["selected"] = tuning_frame["C"].eq(selected_c)

    outer_preprocessor = fit_preprocessor(train, spec, state_columns)
    outer_train_design = transform(outer_preprocessor, train)
    outer_test_design = transform(outer_preprocessor, test)
    outer_estimator = fit_estimator(
        outer_train_design, train["death_5y"].to_numpy(int), selected_c
    )
    outer_prediction = predict(outer_estimator, outer_test_design)
    artifact = {
        "schema_version": SCHEMA_VERSION,
        "analysis_version": ANALYSIS_VERSION,
        "analysis_signature": analysis_signature,
        "artifact_signature": expected_signature,
        "model_spec": asdict(spec),
        "outer_fold": outer_fold,
        "outer_train_row_ids": train["row_id"].to_numpy(int),
        "outer_test_row_ids": test["row_id"].to_numpy(int),
        "outer_train_SEQN": train_ids,
        "outer_test_SEQN": test_ids,
        "outer_stratification": str(frame["outer_stratification"].iloc[0]),
        "inner_stratification": inner_strategy,
        "selected_C": selected_c,
        "state_columns": state_columns,
        "inner_fits": inner_fits,
        "tuning_rows": tuning_frame.to_dict(orient="records"),
        "outer_preprocessor": outer_preprocessor,
        "outer_estimator": outer_estimator,
        "outer_test_predictions": outer_prediction,
        "outer_test_outcomes": test["death_5y"].to_numpy(int),
        "outer_design_feature_names": outer_preprocessor["feature_names"],
    }
    temporary = path.with_suffix(".tmp.joblib")
    joblib.dump(artifact, temporary, compress=3)
    os.replace(temporary, path)
    reloaded = joblib.load(path)
    _, outer_error, inner_error, inner_models = verify_artifact(
        reloaded, frame_by_row, expected_signature
    )
    return reloaded, False, outer_error, inner_error, inner_models


def participant_loss(y: np.ndarray, probability: np.ndarray) -> np.ndarray:
    probability = np.clip(np.asarray(probability, float), EPSILON, 1.0 - EPSILON)
    return -(y * np.log(probability) + (1 - y) * np.log(1 - probability))


def safe_auc(y: np.ndarray, probability: np.ndarray) -> float:
    return float(roc_auc_score(y, probability)) if len(np.unique(y)) == 2 else float("nan")


def metrics(y: np.ndarray, probability: np.ndarray) -> dict[str, float]:
    return {
        "log_loss": float(log_loss(y, probability, labels=[0, 1])),
        "brier": float(brier_score_loss(y, probability)),
        "auc": safe_auc(y, probability),
    }


def bootstrap_indices(frame: pd.DataFrame) -> np.ndarray:
    y = frame["death_5y"].to_numpy(int)
    bands = age_band(frame["age_years"])
    strata = np.char.add(np.char.add(y.astype(str), "_"), bands.astype(str))
    groups = [np.flatnonzero(strata == label) for label in sorted(np.unique(strata))]
    rng = np.random.default_rng(BOOTSTRAP_SEED)
    draws = np.empty((N_BOOTSTRAPS, len(frame)), dtype=np.int32)
    for bootstrap in range(N_BOOTSTRAPS):
        offset = 0
        for group in groups:
            sample = rng.choice(group, size=len(group), replace=True).astype(np.int32)
            draws[bootstrap, offset : offset + len(group)] = sample
            offset += len(group)
    return draws


def summarize_age_values(
    frame: pd.DataFrame,
    losses: pd.DataFrame,
    draws: np.ndarray,
) -> tuple[pd.DataFrame, pd.DataFrame, pd.DataFrame, dict[str, Any]]:
    model_names = [spec.name for spec in MODEL_SPECS]
    loss_matrix = losses[model_names].to_numpy(float)
    observed = loss_matrix.mean(axis=0)
    observed_map = dict(zip(model_names, observed))
    boot_values = np.empty((N_BOOTSTRAPS, len(AGE_VALUE_PAIRS)), dtype=float)
    tier_names = list(AGE_VALUE_PAIRS)
    for bootstrap in range(N_BOOTSTRAPS):
        means = loss_matrix[draws[bootstrap]].mean(axis=0)
        mean_map = dict(zip(model_names, means))
        for index, tier in enumerate(tier_names):
            without_age, with_age = AGE_VALUE_PAIRS[tier]
            boot_values[bootstrap, index] = mean_map[without_age] - mean_map[with_age]
    point_values = {
        tier: observed_map[without_age] - observed_map[with_age]
        for tier, (without_age, with_age) in AGE_VALUE_PAIRS.items()
    }
    d_boot = boot_values[:, 0] - boot_values[:, -1]
    d_point = point_values["S0"] - point_values["S3"]
    summary_rows = []
    for index, tier in enumerate(tier_names):
        values = boot_values[:, index]
        row = {
            "tier": tier,
            "without_age_model": AGE_VALUE_PAIRS[tier][0],
            "with_age_model": AGE_VALUE_PAIRS[tier][1],
            "n": len(frame),
            "events": int(frame["death_5y"].sum()),
            "age_value": point_values[tier],
            "q025": float(np.quantile(values, 0.025)),
            "q975": float(np.quantile(values, 0.975)),
            "bootstrap_probability_positive": float(np.mean(values > 0)),
        }
        if tier != "S0" and point_values["S0"] != 0:
            row["remaining_ratio_to_S0"] = point_values[tier] / point_values["S0"]
        else:
            row["remaining_ratio_to_S0"] = 1.0 if tier == "S0" else float("nan")
        summary_rows.append(row)
    d_summary = pd.DataFrame(
        [
            {
                "contrast": "D_0_to_3",
                "definition": "V_A(S0)-V_A(S3)",
                "estimate": d_point,
                "q025": float(np.quantile(d_boot, 0.025)),
                "q975": float(np.quantile(d_boot, 0.975)),
                "bootstrap_probability_positive": float(np.mean(d_boot > 0)),
                "n_bootstraps": N_BOOTSTRAPS,
            }
        ]
    )
    bootstrap_rows = []
    for bootstrap in range(N_BOOTSTRAPS):
        row: dict[str, Any] = {"bootstrap": bootstrap}
        for index, tier in enumerate(tier_names):
            row[f"V_A__{tier}"] = boot_values[bootstrap, index]
        row["D_0_to_3"] = d_boot[bootstrap]
        bootstrap_rows.append(row)
    v0_low = float(np.quantile(boot_values[:, 0], 0.025))
    d_low = float(np.quantile(d_boot, 0.025))
    d_high = float(np.quantile(d_boot, 0.975))
    if v0_low <= 0:
        category = "NO_VISIBLE_BASELINE_AGE_VALUE"
    elif d_low > 0:
        category = "CONTEXTUAL_SUBSTITUTION"
    elif d_high < 0:
        category = "CONTEXTUAL_SYNERGY"
    else:
        category = "NO_DETECTABLE_MIGRATION"
    decision = {
        "decision_category": category,
        "endpoint": "five_year_all_cause_mortality",
        "population": "NHANES 2007-2012 complete 50-measurement interface, ages 20-79",
        "n": len(frame),
        "events": int(frame["death_5y"].sum()),
        "V_A_S0": point_values["S0"],
        "V_A_S0_q025": v0_low,
        "V_A_S0_q975": float(np.quantile(boot_values[:, 0], 0.975)),
        "V_A_S3": point_values["S3"],
        "V_A_S3_q025": float(np.quantile(boot_values[:, -1], 0.025)),
        "V_A_S3_q975": float(np.quantile(boot_values[:, -1], 0.975)),
        "D_0_to_3": d_point,
        "D_0_to_3_q025": d_low,
        "D_0_to_3_q975": d_high,
        "n_bootstraps": N_BOOTSTRAPS,
        "bootstrap_seed": BOOTSTRAP_SEED,
        "interval_note": "Paired participant bootstrap of fixed OOF predictions; omits refitting and development uncertainty.",
        "claim_boundary": "Predictive substitution/synergy for this endpoint and representation; not causal mediation or a fraction of ageing.",
    }
    return pd.DataFrame(summary_rows), d_summary, pd.DataFrame(bootstrap_rows), decision


def make_figure(age_summary: pd.DataFrame, d_summary: pd.DataFrame, decision: dict[str, Any]) -> None:
    order = ["S0", "S1", "S2c", "S3"]
    view = age_summary.set_index("tier").loc[order]
    x = np.arange(len(order))
    y = view["age_value"].to_numpy(float)
    lower = y - view["q025"].to_numpy(float)
    upper = view["q975"].to_numpy(float) - y
    figure, axis = plt.subplots(figsize=(7.2, 4.8))
    axis.errorbar(x, y, yerr=np.vstack([lower, upper]), fmt="o-", capsize=4, color="#214e7a")
    axis.axhline(0, color="#666666", linewidth=0.8)
    axis.set_xticks(x, ["Background", "6 measures", "30 measures", "50 measures"])
    axis.set_ylabel("OOF log-loss gain from chronological age")
    axis.set_title("Contextual value of age under complete five-year support")
    d = d_summary.iloc[0]
    axis.text(
        0.98,
        0.96,
        f"D₀→₃ = {d['estimate']:.4f}\n95% interval {d['q025']:.4f} to {d['q975']:.4f}\n{decision['decision_category']}",
        transform=axis.transAxes,
        ha="right",
        va="top",
        fontsize=9,
        bbox={"boxstyle": "round", "facecolor": "white", "edgecolor": "#999999", "alpha": 0.9},
    )
    axis.grid(axis="y", alpha=0.2)
    figure.tight_layout()
    figure.savefig(OUT / "contextual_age_decision.png", dpi=220, bbox_inches="tight")
    figure.savefig(OUT / "contextual_age_decision.svg", bbox_inches="tight")
    plt.close(figure)


def markdown_table(frame: pd.DataFrame, digits: int = 6) -> str:
    display = frame.copy()
    lines = [
        "| " + " | ".join(str(column) for column in display.columns) + " |",
        "| " + " | ".join(["---"] * len(display.columns)) + " |",
    ]
    for row in display.itertuples(index=False, name=None):
        values = []
        for value in row:
            if isinstance(value, (float, np.floating)):
                values.append("" if not np.isfinite(value) else f"{float(value):.{digits}g}")
            else:
                values.append(str(value))
        lines.append("| " + " | ".join(values) + " |")
    return "\n".join(lines)


def write_report(
    support: pd.DataFrame,
    pooled: pd.DataFrame,
    age_summary: pd.DataFrame,
    d_summary: pd.DataFrame,
    decision: dict[str, Any],
    artifact_audit: dict[str, Any],
) -> None:
    display_age = age_summary[["tier", "age_value", "q025", "q975", "remaining_ratio_to_S0"]]
    selected = pooled[["model", "log_loss", "brier", "auc"]]
    text = f"""# NHANES five-year complete-support contextual-age probe

Date: 23 August 2026

## Decision

**{decision['decision_category']}**

For five-year all-cause mortality in the frozen 2007–2012 complete-support NHANES interface, the baseline OOF log-loss value of chronological age was {decision['V_A_S0']:.6f} (95% paired fixed-prediction bootstrap interval {decision['V_A_S0_q025']:.6f} to {decision['V_A_S0_q975']:.6f}). After all 50 age-blind measurements, it was {decision['V_A_S3']:.6f} ({decision['V_A_S3_q025']:.6f} to {decision['V_A_S3_q975']:.6f}). The prespecified change, `D_0_to_3 = V_A(S0)-V_A(S3)`, was {decision['D_0_to_3']:.6f} ({decision['D_0_to_3_q025']:.6f} to {decision['D_0_to_3_q975']:.6f}).

This is a predictive result for one endpoint, population, model family and fixed measurement ladder. It is not causal mediation, state sufficiency, biological-age identification or a fraction of ageing.

## Complete-support data gate

{markdown_table(support)}

Total: {decision['n']:,} participants and {decision['events']:,} deaths; zero event-free participants censored before 60 months.

## Age value across the fixed ladder

{markdown_table(display_age)}

The sequence was not required to be monotone. Ratios are secondary predictive quantities and are not percentages of ageing.

## Pooled model diagnostics

{markdown_table(selected)}

## Persistence and parity

- Serialized outer-fold/model artifacts: {artifact_audit['artifact_count']}.
- Persisted inner tuning estimators: {artifact_audit['inner_estimator_count']}.
- Maximum outer held-out reload error: {artifact_audit['max_outer_prediction_error']:.3g}.
- Maximum inner-validation reload error: {artifact_audit['max_inner_prediction_error']:.3g}.
- Compatible artifacts reused: {artifact_audit['reused_artifact_count']}.
- Newly fitted artifacts: {artifact_audit['newly_fitted_artifact_count']}.

Every compared model used the same participant folds. Category encoding, state scaling, five-knot cubic age spline construction, three-fold tuning and the final estimator were training-fold local. The independent numerical audit is written separately to `independent_audit.json`.

## Uncertainty boundary

The 2,000 paired bootstrap replicates reused identical endpoint-and-decade-stratified participant indices for all models. They quantify participant resampling conditional on stored OOF predictions and omit model refitting, tuning and study-development uncertainty.
"""
    REPORT.write_text(text, encoding="utf-8")


def main() -> int:
    configure_stdout()
    OUT.mkdir(parents=True, exist_ok=True)
    MODEL_DIR.mkdir(parents=True, exist_ok=True)
    for required in (PANEL, FEATURE_MANIFEST, SPEC, AUDIT_SCRIPT, ENV_POLICY, ENV_MANIFEST, ENV_LOCK, Path(__file__)):
        if not required.exists():
            raise RuntimeError(f"Required file missing: {required}")

    input_paths = {
        "analysis_ready_source": PANEL,
        "feature_manifest": FEATURE_MANIFEST,
        "frozen_specification": SPEC,
        "analysis_script": Path(__file__),
        "independent_audit_script": AUDIT_SCRIPT,
        "environment_policy": ENV_POLICY,
        "environment_manifest": ENV_MANIFEST,
        "environment_lock": ENV_LOCK,
    }
    input_manifest = pd.DataFrame(
        [
            {
                "role": role,
                "path": str(path.relative_to(ROOT)),
                "bytes": path.stat().st_size,
                "sha256": sha256(path),
            }
            for role, path in input_paths.items()
        ]
    )
    signature_payload = {
        "analysis_version": ANALYSIS_VERSION,
        "schema_version": SCHEMA_VERSION,
        "inputs": input_manifest[["role", "sha256"]].to_dict(orient="records"),
        "cycles": ALLOWED_CYCLES,
        "endpoint": "death_at_or_before_60_months",
        "model_specs": [asdict(spec) for spec in MODEL_SPECS],
        "C_grid": C_GRID,
        "outer_seed": OUTER_SEED,
        "bootstrap_seed": BOOTSTRAP_SEED,
        "outer_folds": N_OUTER_FOLDS,
        "inner_folds": N_INNER_FOLDS,
        "bootstraps": N_BOOTSTRAPS,
    }
    analysis_signature = stable_hash(signature_payload)
    signature_path = OUT / "analysis_signature.json"
    if signature_path.exists():
        existing = json.loads(signature_path.read_text(encoding="utf-8"))
        if existing.get("analysis_signature") != analysis_signature:
            raise RuntimeError("Existing output has an incompatible analysis signature")
    else:
        write_json(
            signature_path,
            {"analysis_signature": analysis_signature, "signature_payload": signature_payload},
        )
    input_manifest.to_csv(OUT / "input_manifest.tsv", sep="\t", index=False)

    frame, support, manifest, tiers = load_and_gate()
    folds, outer_strategy = assign_outer_folds(frame)
    frame["outer_fold"] = folds
    frame["outer_stratification"] = outer_strategy
    frame_by_row = frame.set_index("row_id", drop=False)

    frame.to_csv(OUT / "analysis_ready_risk_set.tsv.gz", sep="\t", index=False, compression="gzip")
    support.to_csv(OUT / "cycle_support_audit.tsv", sep="\t", index=False)
    frame[["row_id", "SEQN", "cycle", "age_years", "death_5y", "outer_fold", "outer_stratification"]].to_csv(
        OUT / "participant_folds.tsv", sep="\t", index=False
    )
    tier_rows = []
    for tier, columns in tiers.items():
        for order, column in enumerate(columns, start=1):
            source = manifest.loc[manifest["transformed_column"].eq(column)].iloc[0]
            tier_rows.append(
                {
                    "tier": tier,
                    "order": order,
                    "transformed_column": column,
                    "raw_feature": source["feature"],
                    "source_module": source["source_module"],
                    "clinical_label": source.get("clinical_label", ""),
                }
            )
    pd.DataFrame(tier_rows).to_csv(OUT / "ordered_feature_tiers.tsv", sep="\t", index=False)

    predictions = {spec.name: np.full(len(frame), np.nan, dtype=float) for spec in MODEL_SPECS}
    tuning_rows: list[dict[str, Any]] = []
    split_rows: list[dict[str, Any]] = []
    artifact_index_rows: list[dict[str, Any]] = []
    coefficient_rows: list[dict[str, Any]] = []
    fold_metric_rows: list[dict[str, Any]] = []
    max_outer_error = 0.0
    max_inner_error = 0.0
    inner_estimator_count = 0
    reused_count = 0
    fitted_count = 0

    for outer_fold in range(N_OUTER_FOLDS):
        print(f"Outer fold {outer_fold + 1}/{N_OUTER_FOLDS}", flush=True)
        test_mask = frame["outer_fold"].eq(outer_fold).to_numpy()
        y_test = frame.loc[test_mask, "death_5y"].to_numpy(int)
        for spec in MODEL_SPECS:
            print(f"  {spec.name}", flush=True)
            artifact, reused, outer_error, inner_error, inner_models = fit_or_load_artifact(
                frame,
                frame_by_row,
                spec,
                tiers[spec.tier],
                outer_fold,
                analysis_signature,
            )
            probability, _, _, _ = verify_artifact(
                artifact,
                frame_by_row,
                artifact["artifact_signature"],
            )
            predictions[spec.name][test_mask] = probability
            max_outer_error = max(max_outer_error, outer_error)
            max_inner_error = max(max_inner_error, inner_error)
            inner_estimator_count += inner_models
            reused_count += int(reused)
            fitted_count += int(not reused)
            tuning_rows.extend(artifact["tuning_rows"])
            for inner in artifact["inner_fits"]:
                for role, row_ids in (
                    ("train", inner["train_row_ids"]),
                    ("validation", inner["validation_row_ids"]),
                ):
                    for row_id in np.asarray(row_ids, dtype=int):
                        split_rows.append(
                            {
                                "outer_fold": outer_fold,
                                "inner_fold": int(inner["inner_fold"]),
                                "model": spec.name,
                                "role": role,
                                "row_id": int(row_id),
                                "SEQN": int(frame_by_row.at[int(row_id), "SEQN"]),
                            }
                        )
            artifact_path = MODEL_DIR / f"outer_fold_{outer_fold:02d}" / f"{spec.name}.joblib"
            artifact_index_rows.append(
                {
                    "outer_fold": outer_fold,
                    "model": spec.name,
                    "tier": spec.tier,
                    "include_age": spec.include_age,
                    "selected_C": artifact["selected_C"],
                    "n_train": len(artifact["outer_train_row_ids"]),
                    "n_test": len(artifact["outer_test_row_ids"]),
                    "n_design_features": len(artifact["outer_design_feature_names"]),
                    "artifact_path": str(artifact_path.relative_to(ROOT)),
                    "artifact_signature": artifact["artifact_signature"],
                    "artifact_sha256": sha256(artifact_path),
                    "reused": reused,
                    "outer_reload_max_abs_error": outer_error,
                    "inner_reload_max_abs_error": inner_error,
                }
            )
            estimator = artifact["outer_estimator"]
            for feature, coefficient in zip(
                artifact["outer_design_feature_names"], estimator.coef_[0]
            ):
                coefficient_rows.append(
                    {
                        "outer_fold": outer_fold,
                        "model": spec.name,
                        "selected_C": artifact["selected_C"],
                        "feature": feature,
                        "coefficient": float(coefficient),
                        "intercept": float(estimator.intercept_[0]),
                    }
                )
            fold_metric_rows.append(
                {
                    "outer_fold": outer_fold,
                    "model": spec.name,
                    "n": int(test_mask.sum()),
                    "events": int(y_test.sum()),
                    **metrics(y_test, probability),
                }
            )

    if any(not np.isfinite(value).all() for value in predictions.values()):
        raise RuntimeError("STOP_DATA_OR_AUDIT_FAIL: incomplete OOF predictions")
    oof = frame[["row_id", "SEQN", "cycle", "age_years", "death_5y", "outer_fold"]].copy()
    for spec in MODEL_SPECS:
        oof[spec.name] = predictions[spec.name]
    y = oof["death_5y"].to_numpy(int)
    losses = oof[["row_id", "SEQN", "death_5y", "outer_fold"]].copy()
    for spec in MODEL_SPECS:
        losses[spec.name] = participant_loss(y, oof[spec.name].to_numpy(float))
    oof.to_csv(OUT / "oof_predictions.tsv.gz", sep="\t", index=False, compression="gzip")
    losses.to_csv(OUT / "participant_loss_vectors.tsv.gz", sep="\t", index=False, compression="gzip")
    pd.DataFrame(tuning_rows).to_csv(OUT / "inner_tuning.tsv.gz", sep="\t", index=False, compression="gzip")
    pd.DataFrame(split_rows).to_csv(OUT / "inner_split_identities.tsv.gz", sep="\t", index=False, compression="gzip")
    artifact_index = pd.DataFrame(artifact_index_rows)
    artifact_index.to_csv(OUT / "model_artifact_index.tsv", sep="\t", index=False)
    pd.DataFrame(coefficient_rows).to_csv(OUT / "outer_model_coefficients.tsv.gz", sep="\t", index=False, compression="gzip")
    fold_metrics = pd.DataFrame(fold_metric_rows)
    fold_metrics.to_csv(OUT / "fold_metrics.tsv", sep="\t", index=False)

    pooled_rows = []
    calibration_rows = []
    bands = age_band(oof["age_years"])
    for spec in MODEL_SPECS:
        probability = oof[spec.name].to_numpy(float)
        pooled_rows.append(
            {
                "model": spec.name,
                "tier": spec.tier,
                "include_age": spec.include_age,
                "n": len(oof),
                "events": int(y.sum()),
                **metrics(y, probability),
            }
        )
        for band_index, label in enumerate(AGE_BAND_LABELS):
            mask = bands == band_index
            calibration_rows.append(
                {
                    "model": spec.name,
                    "age_band": label,
                    "n": int(mask.sum()),
                    "events": int(y[mask].sum()),
                    "observed_risk": float(y[mask].mean()),
                    "mean_predicted_risk": float(probability[mask].mean()),
                    "observed_minus_predicted": float(y[mask].mean() - probability[mask].mean()),
                }
            )
    pooled = pd.DataFrame(pooled_rows)
    pooled.to_csv(OUT / "pooled_metrics.tsv", sep="\t", index=False)
    pd.DataFrame(calibration_rows).to_csv(OUT / "age_band_calibration.tsv", sep="\t", index=False)

    draws_path = OUT / "bootstrap_indices.npz"
    if draws_path.exists():
        loaded = np.load(draws_path)
        draws = loaded["indices"]
        if draws.shape != (N_BOOTSTRAPS, len(frame)) or int(loaded["seed"][0]) != BOOTSTRAP_SEED:
            raise RuntimeError("STOP_DATA_OR_AUDIT_FAIL: incompatible bootstrap indices")
    else:
        draws = bootstrap_indices(frame)
        np.savez_compressed(draws_path, indices=draws, seed=np.asarray([BOOTSTRAP_SEED], dtype=np.int64))
    age_summary, d_summary, bootstrap_contrasts, decision = summarize_age_values(frame, losses, draws)
    age_summary.to_csv(OUT / "age_value_summary.tsv", sep="\t", index=False)
    d_summary.to_csv(OUT / "richest_change_summary.tsv", sep="\t", index=False)
    bootstrap_contrasts.to_csv(OUT / "bootstrap_contrasts.tsv.gz", sep="\t", index=False, compression="gzip")

    artifact_audit = {
        "status": "PASS",
        "analysis_signature": analysis_signature,
        "expected_artifact_count": N_OUTER_FOLDS * len(MODEL_SPECS),
        "artifact_count": int(len(artifact_index)),
        "inner_estimators_per_artifact": N_INNER_FOLDS * len(C_GRID),
        "inner_estimator_count": int(inner_estimator_count),
        "expected_inner_estimator_count": N_OUTER_FOLDS * len(MODEL_SPECS) * N_INNER_FOLDS * len(C_GRID),
        "reused_artifact_count": int(reused_count),
        "newly_fitted_artifact_count": int(fitted_count),
        "parity_tolerance": PARITY_TOLERANCE,
        "max_outer_prediction_error": max_outer_error,
        "max_inner_prediction_error": max_inner_error,
        "all_artifact_signatures_unique": bool(artifact_index["artifact_signature"].nunique() == len(artifact_index)),
        "all_model_hashes_present": bool(artifact_index["artifact_sha256"].str.len().eq(64).all()),
        "software": {
            "python": sys.version,
            "python_executable": sys.executable,
            "platform": platform.platform(),
            "numpy": np.__version__,
            "pandas": pd.__version__,
            "scikit_learn": sklearn.__version__,
            "joblib": joblib.__version__,
        },
    }
    if (
        artifact_audit["artifact_count"] != artifact_audit["expected_artifact_count"]
        or artifact_audit["inner_estimator_count"] != artifact_audit["expected_inner_estimator_count"]
        or max_outer_error > PARITY_TOLERANCE
        or max_inner_error > PARITY_TOLERANCE
        or not artifact_audit["all_artifact_signatures_unique"]
    ):
        artifact_audit["status"] = "FAIL"
        decision["decision_category"] = "STOP_DATA_OR_AUDIT_FAIL"
    write_json(OUT / "model_artifact_audit.json", artifact_audit)
    write_json(OUT / "decision.json", decision)
    make_figure(age_summary, d_summary, decision)
    write_report(support, pooled, age_summary, d_summary, decision, artifact_audit)

    print(json.dumps(json_ready(decision), indent=2, ensure_ascii=False), flush=True)
    print(f"Wrote {OUT}", flush=True)
    return 0 if decision["decision_category"] != "STOP_DATA_OR_AUDIT_FAIL" else 2


if __name__ == "__main__":
    raise SystemExit(main())
```
