## Supplementary_Software_1 for "Inferential boundaries of age prediction: why prediction does not establish biological age measurement": analyze_nhanes_5y_matched_scalar_target_supervision_probe_v0_1.py

```
from __future__ import annotations

import hashlib
import importlib.util
import json
import math
import os
import platform
import sys
import warnings
from pathlib import Path
from typing import Any

import joblib
import numpy as np
import pandas as pd
import sklearn
from sklearn.exceptions import ConvergenceWarning
from sklearn.linear_model import LogisticRegression, Ridge
from sklearn.metrics import brier_score_loss, log_loss, roc_auc_score
from sklearn.model_selection import StratifiedKFold
from sklearn.preprocessing import OneHotEncoder, SplineTransformer, StandardScaler


ROOT = Path(__file__).resolve().parents[1]
ANALYSIS = ROOT / "analysis"
PREDECESSOR = ROOT / "outputs" / "nhanes_5y_complete_support_contextual_age_probe_v0_1"
SOURCE = ROOT / "outputs" / "nhanes_5y_age_decoder_source_information_probe_v0_2"
OUT = ROOT / "outputs" / "nhanes_5y_matched_scalar_target_supervision_probe_v0_1"
MODEL_DIR = OUT / "models"
FLEX_MODEL_DIR = MODEL_DIR / "flexible_calibration"
MATCHED_MODEL_DIR = MODEL_DIR / "matched_scalar"

SOURCE_FRAME = PREDECESSOR / "analysis_ready_risk_set.tsv.gz"
SOURCE_FOLDS = PREDECESSOR / "participant_folds.tsv"
SOURCE_TIERS = PREDECESSOR / "ordered_feature_tiers.tsv"
SOURCE_BOOTSTRAP = PREDECESSOR / "bootstrap_indices.npz"
SOURCE_PREDECESSOR_AUDIT = PREDECESSOR / "model_artifact_audit.json"
SOURCE_OOF = SOURCE / "oof_predictions.tsv.gz"
SOURCE_AUDIT = SOURCE / "model_artifact_audit.json"
SOURCE_SIGNATURE = SOURCE / "analysis_signature.json"
SPEC = ANALYSIS / "NHANES_5Y_MATCHED_SCALAR_TARGET_SUPERVISION_SPEC_20260826.md"
BASE_SCRIPT = ANALYSIS / "analyze_nhanes_5y_age_decoder_source_information_probe_v0_1.py"
ENV_POLICY = ROOT / "environment-policy.json"
ENV_MANIFEST = ROOT / "environment.yml"
ENV_LOCK = ROOT / "environment.win-64.lock.txt"

ANALYSIS_VERSION = "nhanes_5y_matched_scalar_target_supervision_probe_v0_1"
SCHEMA_VERSION = "1.0"
EXPECTED_N = 13_724
EXPECTED_EVENTS = 470
EXPECTED_CYCLES = ("2007_2008", "2009_2010", "2011_2012")
BACKGROUND = ("sex", "race_ethnicity", "cycle")
OUTER_SEEDS = (20260823, 20260901, 20260902, 20260903, 20260904)
N_OUTER_FOLDS = 5
N_DOWNSTREAM_FOLDS = 3
N_PROJECTION_TUNING_FOLDS = 3
N_BOOTSTRAPS = 2_000
BOOTSTRAP_SEED = 20260905
ALPHA_GRID = (0.01, 0.1, 1.0, 10.0, 100.0, 1000.0)
C_GRID = (0.001, 0.003, 0.01, 0.03, 0.1, 0.3, 1.0, 3.0, 10.0, 30.0, 100.0, 300.0)
PARITY_TOLERANCE = 1e-12
EPSILON = 1e-7


def load_base_module() -> Any:
    module_spec = importlib.util.spec_from_file_location("nhanes_age_decoder_base", BASE_SCRIPT)
    if module_spec is None or module_spec.loader is None:
        raise RuntimeError("Could not load frozen NHANES age-decoder implementation")
    module = importlib.util.module_from_spec(module_spec)
    module_spec.loader.exec_module(module)
    return module


base = load_base_module()


def configure_stdout() -> None:
    if hasattr(sys.stdout, "reconfigure"):
        sys.stdout.reconfigure(encoding="utf-8")


def sha256(path: Path) -> str:
    digest = hashlib.sha256()
    with path.open("rb") as handle:
        for block in iter(lambda: handle.read(1024 * 1024), b""):
            digest.update(block)
    return digest.hexdigest()


def stable_hash(payload: Any) -> str:
    encoded = json.dumps(json_ready(payload), sort_keys=True, separators=(",", ":"), ensure_ascii=False).encode("utf-8")
    return hashlib.sha256(encoded).hexdigest()


def json_ready(value: Any) -> Any:
    if isinstance(value, dict):
        return {str(key): json_ready(item) for key, item in value.items()}
    if isinstance(value, (list, tuple)):
        return [json_ready(item) for item in value]
    if isinstance(value, np.ndarray):
        return value.tolist()
    if isinstance(value, (np.integer,)):
        return int(value)
    if isinstance(value, (np.floating,)):
        return float(value)
    if isinstance(value, (np.bool_,)):
        return bool(value)
    if isinstance(value, Path):
        return str(value)
    if pd.isna(value) if np.isscalar(value) and not isinstance(value, str) else False:
        return None
    return value


def write_json(path: Path, payload: Any) -> None:
    path.write_text(json.dumps(json_ready(payload), indent=2, ensure_ascii=False) + "\n", encoding="utf-8")


def save_joblib_atomic(payload: Any, path: Path) -> None:
    path.parent.mkdir(parents=True, exist_ok=True)
    temporary = path.with_suffix(path.suffix + ".tmp")
    if temporary.exists():
        temporary.unlink()
    joblib.dump(payload, temporary, compress=3)
    os.replace(temporary, path)


def clean_categories(frame: pd.DataFrame) -> pd.DataFrame:
    return frame.fillna("missing").astype(str).replace("", "missing")


def endpoint_metrics(y: np.ndarray, probability: np.ndarray) -> dict[str, float]:
    return {
        "log_loss": float(log_loss(y, probability, labels=[0, 1])),
        "brier": float(brier_score_loss(y, probability)),
        "auc": float(roc_auc_score(y, probability)),
    }


def participant_loss(y: np.ndarray, probability: np.ndarray) -> np.ndarray:
    p = np.clip(np.asarray(probability, dtype=float), EPSILON, 1.0 - EPSILON)
    target = np.asarray(y, dtype=float)
    return -(target * np.log(p) + (1.0 - target) * np.log(1.0 - p))


def age_decade(age: pd.Series | np.ndarray) -> np.ndarray:
    values = np.asarray(age, dtype=float)
    return np.clip(((values - 20.0) // 10.0).astype(int), 0, 5)


def make_joint_strata(frame: pd.DataFrame, n_splits: int) -> np.ndarray:
    age = pd.to_numeric(frame["age_years"], errors="raise").reset_index(drop=True)
    quintile = pd.qcut(age, q=5, duplicates="drop", labels=False).fillna(-1).astype(int).to_numpy()
    y = frame["death_5y"].to_numpy(int)
    combined = np.char.add(np.char.add(y.astype(str), "_"), quintile.astype(str))
    counts = pd.Series(combined).value_counts()
    if len(counts) < 2 or int(counts.min()) < n_splits:
        raise RuntimeError("STOP_DATA_OR_ARTIFACT_FAIL: joint outcome-age strata cannot support fixed folds")
    return combined


def make_splits(frame: pd.DataFrame, n_splits: int, seed: int) -> list[tuple[np.ndarray, np.ndarray]]:
    strata = make_joint_strata(frame, n_splits)
    splitter = StratifiedKFold(n_splits=n_splits, shuffle=True, random_state=int(seed))
    splits = [(np.asarray(train, dtype=int), np.asarray(valid, dtype=int)) for train, valid in splitter.split(np.zeros(len(frame)), strata)]
    for train, valid in splits:
        if np.intersect1d(train, valid).size or len(train) + len(valid) != len(frame):
            raise RuntimeError("STOP_DATA_OR_ARTIFACT_FAIL: invalid nested split identity")
        train_events = int(frame.iloc[train]["death_5y"].sum())
        valid_events = int(frame.iloc[valid]["death_5y"].sum())
        if train_events <= 0 or valid_events <= 0:
            raise RuntimeError("STOP_DATA_OR_ARTIFACT_FAIL: nested split lacks event support")
    return splits


def assign_outer_folds(frame: pd.DataFrame, seed: int) -> np.ndarray:
    strata = make_joint_strata(frame, N_OUTER_FOLDS)
    splitter = StratifiedKFold(n_splits=N_OUTER_FOLDS, shuffle=True, random_state=int(seed))
    assignment = np.full(len(frame), -1, dtype=int)
    for fold, (_, test) in enumerate(splitter.split(np.zeros(len(frame)), strata)):
        assignment[test] = fold
    if np.any(assignment < 0):
        raise RuntimeError("STOP_DATA_OR_ARTIFACT_FAIL: incomplete outer assignment")
    return assignment


def load_and_gate() -> tuple[pd.DataFrame, list[str], pd.DataFrame, list[dict[str, Any]]]:
    predecessor_audit = json.loads(SOURCE_PREDECESSOR_AUDIT.read_text(encoding="utf-8"))
    source_audit = json.loads(SOURCE_AUDIT.read_text(encoding="utf-8"))
    if predecessor_audit.get("status") != "PASS" or source_audit.get("status") != "PASS":
        raise RuntimeError("STOP_DATA_OR_ARTIFACT_FAIL: frozen source audit is not PASS")

    frame = pd.read_csv(SOURCE_FRAME, sep="\t").sort_values("row_id").reset_index(drop=True)
    tiers = pd.read_csv(SOURCE_TIERS, sep="\t")
    frozen_folds = pd.read_csv(SOURCE_FOLDS, sep="\t").sort_values("row_id").reset_index(drop=True)
    source_oof = pd.read_csv(SOURCE_OOF, sep="\t").sort_values("row_id").reset_index(drop=True)
    features = tiers.loc[tiers["tier"].eq("S3")].sort_values("order")["transformed_column"].tolist()

    required = ["row_id", "SEQN", "cycle", "age_years", "death_5y", "outer_fold", *BACKGROUND, *features]
    missing = sorted(set(required) - set(frame.columns))
    if missing:
        raise RuntimeError(f"STOP_DATA_OR_ARTIFACT_FAIL: missing input columns {missing}")
    if len(frame) != EXPECTED_N or int(frame["death_5y"].sum()) != EXPECTED_EVENTS:
        raise RuntimeError("STOP_DATA_OR_ARTIFACT_FAIL: participant or event gate changed")
    if frame["SEQN"].nunique() != EXPECTED_N or not frame["age_years"].between(20, 79).all():
        raise RuntimeError("STOP_DATA_OR_ARTIFACT_FAIL: participant identity or age gate changed")
    if tuple(sorted(frame["cycle"].unique())) != EXPECTED_CYCLES:
        raise RuntimeError("STOP_DATA_OR_ARTIFACT_FAIL: cycle gate changed")
    if len(features) != 50 or len(set(features)) != 50 or frame[required].isna().any().any():
        raise RuntimeError("STOP_DATA_OR_ARTIFACT_FAIL: feature-order or complete-interface gate failed")
    if not np.array_equal(frame["row_id"].to_numpy(int), np.arange(EXPECTED_N, dtype=int)):
        raise RuntimeError("STOP_DATA_OR_ARTIFACT_FAIL: row_id order changed")
    if not np.array_equal(frame[["row_id", "SEQN"]].to_numpy(), frozen_folds[["row_id", "SEQN"]].to_numpy()):
        raise RuntimeError("STOP_DATA_OR_ARTIFACT_FAIL: frozen fold participant identity changed")
    if not np.array_equal(frame[["row_id", "SEQN"]].to_numpy(), source_oof[["row_id", "SEQN"]].to_numpy()):
        raise RuntimeError("STOP_DATA_OR_ARTIFACT_FAIL: source OOF participant identity changed")
    if not {"M_WA", "M_CLOCK", "M_X", "clock_score"}.issubset(source_oof.columns):
        raise RuntimeError("STOP_DATA_OR_ARTIFACT_FAIL: required frozen predictions are missing")

    first_assignment = assign_outer_folds(frame, OUTER_SEEDS[0])
    if not np.array_equal(first_assignment, frozen_folds["outer_fold"].to_numpy(int)):
        raise RuntimeError("STOP_DATA_OR_ARTIFACT_FAIL: seed 20260823 no longer reproduces frozen folds")
    if not np.array_equal(first_assignment, frame["outer_fold"].to_numpy(int)):
        raise RuntimeError("STOP_DATA_OR_ARTIFACT_FAIL: analysis-ready frozen folds changed")

    source_artifacts: list[dict[str, Any]] = []
    for outer_fold in range(N_OUTER_FOLDS):
        artifact_path = SOURCE / "models" / f"outer_fold_{outer_fold:02d}.joblib"
        artifact = joblib.load(artifact_path)
        train = frame.loc[first_assignment != outer_fold].copy().reset_index(drop=True)
        test = frame.loc[first_assignment == outer_fold].copy().reset_index(drop=True)
        crossfit = pd.read_csv(SOURCE / f"outer_fold_{outer_fold:02d}_training_crossfit_scores.tsv.gz", sep="\t")
        if not np.array_equal(train["row_id"].to_numpy(int), artifact["train_row_ids"]):
            raise RuntimeError("STOP_DATA_OR_ARTIFACT_FAIL: frozen age artifact train identity changed")
        if not np.array_equal(test["row_id"].to_numpy(int), artifact["test_row_ids"]):
            raise RuntimeError("STOP_DATA_OR_ARTIFACT_FAIL: frozen age artifact test identity changed")
        if not np.array_equal(crossfit["row_id"].to_numpy(int), artifact["train_row_ids"]):
            raise RuntimeError("STOP_DATA_OR_ARTIFACT_FAIL: frozen age cross-fit identity changed")
        if np.max(np.abs(crossfit["crossfit_clock_score"].to_numpy(float) - artifact["train_clock"])) > PARITY_TOLERANCE:
            raise RuntimeError("STOP_DATA_OR_ARTIFACT_FAIL: frozen age cross-fit score parity failed")
        reproduced = base.predict_ridge_age(artifact["outer_age_model"], test, features)
        if np.max(np.abs(reproduced - artifact["test_clock"])) > PARITY_TOLERANCE:
            raise RuntimeError("STOP_DATA_OR_ARTIFACT_FAIL: frozen age outer score parity failed")
        source_artifacts.append(artifact)

    return frame, features, source_oof, source_artifacts


def fit_projection_preprocessor(frame: pd.DataFrame, features: list[str]) -> tuple[OneHotEncoder, StandardScaler, np.ndarray, list[str]]:
    encoder = OneHotEncoder(handle_unknown="ignore", sparse_output=False)
    categories = encoder.fit_transform(clean_categories(frame.loc[:, BACKGROUND]))
    scaler = StandardScaler()
    measurements = scaler.fit_transform(frame.loc[:, features].to_numpy(float))
    design = np.column_stack([categories, measurements])
    names = [f"W__{name}" for name in encoder.get_feature_names_out(BACKGROUND)] + [f"X__{name}" for name in features]
    return encoder, scaler, design, names


def transform_projection_preprocessor(
    frame: pd.DataFrame,
    features: list[str],
    encoder: OneHotEncoder,
    scaler: StandardScaler,
) -> np.ndarray:
    categories = encoder.transform(clean_categories(frame.loc[:, BACKGROUND]))
    measurements = scaler.transform(frame.loc[:, features].to_numpy(float))
    return np.column_stack([categories, measurements])


def fit_projection(frame: pd.DataFrame, features: list[str], target: str, alpha: float) -> dict[str, Any]:
    encoder, scaler, design, names = fit_projection_preprocessor(frame, features)
    raw_target = frame[target].to_numpy(float)
    target_mean = float(raw_target.mean())
    target_sd = float(raw_target.std(ddof=0))
    if not np.isfinite(target_sd) or target_sd <= 0:
        raise RuntimeError("STOP_DATA_OR_ARTIFACT_FAIL: projection target has zero variance")
    standardized_target = (raw_target - target_mean) / target_sd
    estimator = Ridge(alpha=float(alpha))
    estimator.fit(design, standardized_target)
    return {
        "target": target,
        "alpha": float(alpha),
        "encoder": encoder,
        "scaler": scaler,
        "target_mean": target_mean,
        "target_sd": target_sd,
        "estimator": estimator,
        "design_feature_names": names,
    }


def predict_projection(model: dict[str, Any], frame: pd.DataFrame, features: list[str]) -> np.ndarray:
    design = transform_projection_preprocessor(frame, features, model["encoder"], model["scaler"])
    return model["estimator"].predict(design)


def standardized_target_for_model(model: dict[str, Any], frame: pd.DataFrame) -> np.ndarray:
    raw = frame[model["target"]].to_numpy(float)
    return (raw - float(model["target_mean"])) / float(model["target_sd"])


def fit_projection_bundle(
    training: pd.DataFrame,
    holdout: pd.DataFrame,
    features: list[str],
    seed: int,
    bundle_name: str,
) -> dict[str, Any]:
    splits = make_splits(training, N_PROJECTION_TUNING_FOLDS, seed)
    targets: dict[str, Any] = {}
    for target_name, target_column in (("age", "age_years"), ("mortality", "death_5y")):
        tuning_fits = []
        tuning_rows = []
        for alpha in ALPHA_GRID:
            for inner_fold, (inner_train, inner_valid) in enumerate(splits):
                model = fit_projection(training.iloc[inner_train], features, target_column, alpha)
                prediction = predict_projection(model, training.iloc[inner_valid], features)
                expected = standardized_target_for_model(model, training.iloc[inner_valid])
                mse = float(np.mean((expected - prediction) ** 2))
                tuning_fits.append(
                    {
                        "alpha": float(alpha),
                        "inner_fold": int(inner_fold),
                        "train_row_ids": training.iloc[inner_train]["row_id"].to_numpy(int),
                        "valid_row_ids": training.iloc[inner_valid]["row_id"].to_numpy(int),
                        "model": model,
                        "valid_score": prediction,
                        "mse": mse,
                    }
                )
                tuning_rows.append({"alpha": float(alpha), "inner_fold": int(inner_fold), "mse": mse})
        tuning = pd.DataFrame(tuning_rows)
        selected_alpha = float(
            tuning.groupby("alpha", as_index=False)["mse"].mean().sort_values(["mse", "alpha"]).iloc[0]["alpha"]
        )
        final_model = fit_projection(training, features, target_column, selected_alpha)
        holdout_score = predict_projection(final_model, holdout, features)
        targets[target_name] = {
            "target_column": target_column,
            "selected_alpha": selected_alpha,
            "tuning_fits": tuning_fits,
            "tuning_rows": tuning_rows,
            "final_model": final_model,
            "holdout_score": holdout_score,
        }
    return {
        "bundle_name": bundle_name,
        "seed": int(seed),
        "training_row_ids": training["row_id"].to_numpy(int),
        "holdout_row_ids": holdout["row_id"].to_numpy(int),
        "targets": targets,
    }


def verify_projection_bundle(bundle: dict[str, Any], frame_by_row: pd.DataFrame, features: list[str]) -> float:
    training_ids = np.asarray(bundle["training_row_ids"], dtype=int)
    holdout_ids = np.asarray(bundle["holdout_row_ids"], dtype=int)
    if np.intersect1d(training_ids, holdout_ids).size:
        raise RuntimeError("STOP_DATA_OR_ARTIFACT_FAIL: projection bundle leaks holdout identity")
    max_error = 0.0
    for target in bundle["targets"].values():
        for fit in target["tuning_fits"]:
            fit_train = np.asarray(fit["train_row_ids"], dtype=int)
            fit_valid = np.asarray(fit["valid_row_ids"], dtype=int)
            if np.intersect1d(fit_train, fit_valid).size:
                raise RuntimeError("STOP_DATA_OR_ARTIFACT_FAIL: projection tuning leaks validation identity")
            valid = frame_by_row.loc[fit_valid]
            reproduced = predict_projection(fit["model"], valid, features)
            max_error = max(max_error, float(np.max(np.abs(reproduced - np.asarray(fit["valid_score"], dtype=float)))))
        holdout = frame_by_row.loc[holdout_ids]
        reproduced = predict_projection(target["final_model"], holdout, features)
        max_error = max(max_error, float(np.max(np.abs(reproduced - np.asarray(target["holdout_score"], dtype=float)))))
    if max_error > PARITY_TOLERANCE:
        raise RuntimeError(f"STOP_DATA_OR_ARTIFACT_FAIL: projection reload parity failed ({max_error})")
    return max_error


def fit_downstream_model(frame: pd.DataFrame, score: np.ndarray, c_value: float, mode: str) -> dict[str, Any]:
    encoder = OneHotEncoder(handle_unknown="ignore", sparse_output=False)
    categories = encoder.fit_transform(clean_categories(frame.loc[:, BACKGROUND]))
    age_spline = SplineTransformer(n_knots=5, degree=3, include_bias=False, extrapolation="linear")
    age_raw = age_spline.fit_transform(frame[["age_years"]].to_numpy(float))
    age_scaler = StandardScaler().fit(age_raw)
    age_design = age_scaler.transform(age_raw)
    score_scaler = StandardScaler().fit(np.asarray(score, dtype=float).reshape(-1, 1))
    scaled_score = score_scaler.transform(np.asarray(score, dtype=float).reshape(-1, 1))
    score_spline = None
    if mode == "linear":
        score_design = scaled_score
        score_names = ["scalar"]
    elif mode == "spline":
        score_spline = SplineTransformer(n_knots=5, degree=3, include_bias=False, extrapolation="linear")
        score_design = score_spline.fit_transform(scaled_score)
        score_names = [f"scalar_spline_{index}" for index in range(score_design.shape[1])]
    else:
        raise ValueError(f"Unknown downstream mode: {mode}")
    design = np.column_stack([categories, age_design, score_design])
    names = (
        [f"W__{name}" for name in encoder.get_feature_names_out(BACKGROUND)]
        + [f"A_spline_{index}" for index in range(age_design.shape[1])]
        + score_names
    )
    estimator = LogisticRegression(C=float(c_value), penalty="l2", solver="lbfgs", max_iter=5000)
    with warnings.catch_warnings(record=True) as caught:
        warnings.simplefilter("always", ConvergenceWarning)
        warnings.filterwarnings("ignore", category=FutureWarning, message=".*'penalty'.*deprecated.*")
        estimator.fit(design, frame["death_5y"].to_numpy(int))
    if any(issubclass(item.category, ConvergenceWarning) for item in caught):
        raise RuntimeError("STOP_DATA_OR_ARTIFACT_FAIL: downstream logistic model did not converge")
    return {
        "mode": mode,
        "C": float(c_value),
        "encoder": encoder,
        "age_spline": age_spline,
        "age_scaler": age_scaler,
        "score_scaler": score_scaler,
        "score_spline": score_spline,
        "estimator": estimator,
        "design_feature_names": names,
    }


def predict_downstream_model(model: dict[str, Any], frame: pd.DataFrame, score: np.ndarray) -> np.ndarray:
    categories = model["encoder"].transform(clean_categories(frame.loc[:, BACKGROUND]))
    age_raw = model["age_spline"].transform(frame[["age_years"]].to_numpy(float))
    age_design = model["age_scaler"].transform(age_raw)
    scaled_score = model["score_scaler"].transform(np.asarray(score, dtype=float).reshape(-1, 1))
    if model["mode"] == "linear":
        score_design = scaled_score
    elif model["mode"] == "spline":
        score_design = model["score_spline"].transform(scaled_score)
    else:
        raise RuntimeError("Stored downstream mode is invalid")
    design = np.column_stack([categories, age_design, score_design])
    return np.clip(model["estimator"].predict_proba(design)[:, 1], EPSILON, 1.0 - EPSILON)


def tune_downstream(
    training: pd.DataFrame,
    score: np.ndarray,
    splits: list[tuple[np.ndarray, np.ndarray]],
    mode: str,
) -> dict[str, Any]:
    tuning_fits = []
    tuning_rows = []
    for c_value in C_GRID:
        for downstream_fold, (inner_train, inner_valid) in enumerate(splits):
            model = fit_downstream_model(training.iloc[inner_train], score[inner_train], c_value, mode)
            probability = predict_downstream_model(model, training.iloc[inner_valid], score[inner_valid])
            loss = float(log_loss(training.iloc[inner_valid]["death_5y"], probability, labels=[0, 1]))
            tuning_fits.append(
                {
                    "C": float(c_value),
                    "downstream_fold": int(downstream_fold),
                    "train_row_ids": training.iloc[inner_train]["row_id"].to_numpy(int),
                    "valid_row_ids": training.iloc[inner_valid]["row_id"].to_numpy(int),
                    "train_score": np.asarray(score[inner_train], dtype=float),
                    "valid_score": np.asarray(score[inner_valid], dtype=float),
                    "model": model,
                    "valid_probability": probability,
                    "log_loss": loss,
                }
            )
            tuning_rows.append({"C": float(c_value), "downstream_fold": int(downstream_fold), "log_loss": loss})
    tuning = pd.DataFrame(tuning_rows)
    selected_c = float(
        tuning.groupby("C", as_index=False)["log_loss"].mean().sort_values(["log_loss", "C"]).iloc[0]["C"]
    )
    return {"selected_C": selected_c, "tuning_fits": tuning_fits, "tuning_rows": tuning_rows}


def verify_downstream_variant(variant: dict[str, Any], frame_by_row: pd.DataFrame) -> tuple[float, float]:
    max_validation_error = 0.0
    for fit in variant["tuning_fits"]:
        train_ids = np.asarray(fit["train_row_ids"], dtype=int)
        valid_ids = np.asarray(fit["valid_row_ids"], dtype=int)
        if np.intersect1d(train_ids, valid_ids).size:
            raise RuntimeError("STOP_DATA_OR_ARTIFACT_FAIL: downstream tuning leaks validation identity")
        valid = frame_by_row.loc[valid_ids]
        probability = predict_downstream_model(fit["model"], valid, np.asarray(fit["valid_score"], dtype=float))
        max_validation_error = max(
            max_validation_error,
            float(np.max(np.abs(probability - np.asarray(fit["valid_probability"], dtype=float)))),
        )
    test_ids = np.asarray(variant["test_row_ids"], dtype=int)
    test = frame_by_row.loc[test_ids]
    probability = predict_downstream_model(variant["final_model"], test, np.asarray(variant["test_score"], dtype=float))
    max_outer_error = float(np.max(np.abs(probability - np.asarray(variant["test_probability"], dtype=float))))
    if max(max_validation_error, max_outer_error) > PARITY_TOLERANCE:
        raise RuntimeError("STOP_DATA_OR_ARTIFACT_FAIL: downstream reload parity failed")
    return max_validation_error, max_outer_error


def flexible_signature(analysis_signature: str, outer_fold: int, train_ids: list[int], test_ids: list[int]) -> str:
    return stable_hash(
        {
            "analysis_signature": analysis_signature,
            "stage": "frozen_age_score_flexible_calibration",
            "outer_fold": int(outer_fold),
            "train_row_ids": train_ids,
            "test_row_ids": test_ids,
            "C_grid": C_GRID,
            "inner_seed": 20260826 + int(outer_fold),
            "mode": "five_knot_score_spline",
        }
    )


def fit_flexible_artifact(
    frame: pd.DataFrame,
    source_artifact: dict[str, Any],
    outer_fold: int,
    analysis_signature: str,
) -> dict[str, Any]:
    train = frame.loc[frame["outer_fold"].ne(outer_fold)].copy().reset_index(drop=True)
    test = frame.loc[frame["outer_fold"].eq(outer_fold)].copy().reset_index(drop=True)
    train_score = np.asarray(source_artifact["train_clock"], dtype=float)
    test_score = np.asarray(source_artifact["test_clock"], dtype=float)
    splits = make_splits(train, N_DOWNSTREAM_FOLDS, 20260826 + outer_fold)
    tuned = tune_downstream(train, train_score, splits, "spline")
    final_model = fit_downstream_model(train, train_score, tuned["selected_C"], "spline")
    test_probability = predict_downstream_model(final_model, test, test_score)
    train_ids = train["row_id"].astype(int).tolist()
    test_ids = test["row_id"].astype(int).tolist()
    return {
        "schema_version": SCHEMA_VERSION,
        "analysis_signature": analysis_signature,
        "artifact_signature": flexible_signature(analysis_signature, outer_fold, train_ids, test_ids),
        "outer_fold": int(outer_fold),
        "train_row_ids": np.asarray(train_ids, dtype=int),
        "test_row_ids": np.asarray(test_ids, dtype=int),
        "train_score": train_score,
        "test_score": test_score,
        "selected_C": tuned["selected_C"],
        "tuning_fits": tuned["tuning_fits"],
        "tuning_rows": tuned["tuning_rows"],
        "final_model": final_model,
        "test_probability": test_probability,
    }


def verify_flexible_artifact(
    artifact: dict[str, Any], frame: pd.DataFrame, outer_fold: int, analysis_signature: str
) -> tuple[np.ndarray, float, float]:
    train = frame.loc[frame["outer_fold"].ne(outer_fold)].copy().reset_index(drop=True)
    test = frame.loc[frame["outer_fold"].eq(outer_fold)].copy().reset_index(drop=True)
    train_ids = train["row_id"].astype(int).tolist()
    test_ids = test["row_id"].astype(int).tolist()
    expected = flexible_signature(analysis_signature, outer_fold, train_ids, test_ids)
    if artifact.get("artifact_signature") != expected:
        raise RuntimeError("STOP_DATA_OR_ARTIFACT_FAIL: flexible artifact signature mismatch")
    variant = {
        "tuning_fits": artifact["tuning_fits"],
        "test_row_ids": artifact["test_row_ids"],
        "test_score": artifact["test_score"],
        "final_model": artifact["final_model"],
        "test_probability": artifact["test_probability"],
    }
    validation_error, outer_error = verify_downstream_variant(variant, frame.set_index("row_id", drop=False))
    return np.asarray(artifact["test_probability"], dtype=float), validation_error, outer_error


def fit_or_load_flexible_artifact(
    frame: pd.DataFrame,
    source_artifact: dict[str, Any],
    outer_fold: int,
    analysis_signature: str,
) -> tuple[np.ndarray, dict[str, Any]]:
    path = FLEX_MODEL_DIR / f"outer_fold_{outer_fold:02d}.joblib"
    newly_fitted = False
    if path.exists():
        artifact = joblib.load(path)
    else:
        artifact = fit_flexible_artifact(frame, source_artifact, outer_fold, analysis_signature)
        save_joblib_atomic(artifact, path)
        newly_fitted = True
    reloaded = joblib.load(path)
    probability, validation_error, outer_error = verify_flexible_artifact(
        reloaded, frame, outer_fold, analysis_signature
    )
    metadata = {
        "stage": "flexible_calibration",
        "repeat": -1,
        "outer_seed": OUTER_SEEDS[0],
        "outer_fold": outer_fold,
        "artifact_path": str(path.relative_to(ROOT)),
        "artifact_sha256": sha256(path),
        "newly_fitted": newly_fitted,
        "projection_tuning_estimator_count": 0,
        "projection_final_estimator_count": 0,
        "downstream_tuning_estimator_count": len(reloaded["tuning_fits"]),
        "downstream_final_estimator_count": 1,
        "max_projection_score_error": 0.0,
        "max_downstream_validation_error": validation_error,
        "max_downstream_outer_error": outer_error,
    }
    return probability, metadata


def matched_signature(
    analysis_signature: str,
    repeat: int,
    outer_seed: int,
    outer_fold: int,
    train_ids: list[int],
    test_ids: list[int],
    features: list[str],
) -> str:
    return stable_hash(
        {
            "analysis_signature": analysis_signature,
            "stage": "matched_scalar_target_supervision",
            "repeat": int(repeat),
            "outer_seed": int(outer_seed),
            "outer_fold": int(outer_fold),
            "train_row_ids": train_ids,
            "test_row_ids": test_ids,
            "features": features,
            "alpha_grid": ALPHA_GRID,
            "C_grid": C_GRID,
            "downstream_seed": int(outer_seed + 1000 + outer_fold),
            "projection_seed_formula": "outer_seed+2000+10*outer_fold+downstream_fold",
        }
    )


def fit_matched_artifact(
    frame: pd.DataFrame,
    features: list[str],
    assignment: np.ndarray,
    repeat: int,
    outer_seed: int,
    outer_fold: int,
    analysis_signature: str,
) -> dict[str, Any]:
    train = frame.loc[assignment != outer_fold].copy().reset_index(drop=True)
    test = frame.loc[assignment == outer_fold].copy().reset_index(drop=True)
    downstream_seed = int(outer_seed + 1000 + outer_fold)
    downstream_splits = make_splits(train, N_DOWNSTREAM_FOLDS, downstream_seed)
    training_scores = {
        "age": np.full(len(train), np.nan, dtype=float),
        "mortality": np.full(len(train), np.nan, dtype=float),
    }
    projection_bundles = []
    for downstream_fold, (downstream_train, downstream_valid) in enumerate(downstream_splits):
        projection_seed = int(outer_seed + 2000 + 10 * outer_fold + downstream_fold)
        bundle = fit_projection_bundle(
            train.iloc[downstream_train].copy().reset_index(drop=True),
            train.iloc[downstream_valid].copy().reset_index(drop=True),
            features,
            projection_seed,
            f"downstream_fold_{downstream_fold}",
        )
        projection_bundles.append(bundle)
        for target in ("age", "mortality"):
            training_scores[target][downstream_valid] = bundle["targets"][target]["holdout_score"]
    if not all(np.isfinite(score).all() for score in training_scores.values()):
        raise RuntimeError("STOP_DATA_OR_ARTIFACT_FAIL: incomplete cross-fitted projection score")

    final_projection_seed = int(outer_seed + 2000 + 10 * outer_fold + N_DOWNSTREAM_FOLDS)
    final_bundle = fit_projection_bundle(
        train,
        test,
        features,
        final_projection_seed,
        "outer_test",
    )
    test_scores = {
        target: np.asarray(final_bundle["targets"][target]["holdout_score"], dtype=float)
        for target in ("age", "mortality")
    }

    downstream_variants: dict[str, Any] = {}
    for target in ("age", "mortality"):
        for mode in ("linear", "spline"):
            variant_name = f"{target}_{mode}"
            tuned = tune_downstream(train, training_scores[target], downstream_splits, mode)
            final_model = fit_downstream_model(train, training_scores[target], tuned["selected_C"], mode)
            test_probability = predict_downstream_model(final_model, test, test_scores[target])
            downstream_variants[variant_name] = {
                "target": target,
                "mode": mode,
                "selected_C": tuned["selected_C"],
                "tuning_fits": tuned["tuning_fits"],
                "tuning_rows": tuned["tuning_rows"],
                "final_model": final_model,
                "test_row_ids": test["row_id"].to_numpy(int),
                "test_score": test_scores[target],
                "test_probability": test_probability,
            }

    train_ids = train["row_id"].astype(int).tolist()
    test_ids = test["row_id"].astype(int).tolist()
    return {
        "schema_version": SCHEMA_VERSION,
        "analysis_signature": analysis_signature,
        "artifact_signature": matched_signature(
            analysis_signature, repeat, outer_seed, outer_fold, train_ids, test_ids, features
        ),
        "repeat": repeat,
        "outer_seed": outer_seed,
        "outer_fold": outer_fold,
        "features": features,
        "train_row_ids": np.asarray(train_ids, dtype=int),
        "test_row_ids": np.asarray(test_ids, dtype=int),
        "downstream_seed": downstream_seed,
        "downstream_splits": [
            {
                "downstream_fold": index,
                "train_row_ids": train.iloc[split[0]]["row_id"].to_numpy(int),
                "valid_row_ids": train.iloc[split[1]]["row_id"].to_numpy(int),
            }
            for index, split in enumerate(downstream_splits)
        ],
        "projection_bundles": projection_bundles,
        "final_projection_bundle": final_bundle,
        "training_scores": training_scores,
        "test_scores": test_scores,
        "downstream_variants": downstream_variants,
    }


def verify_matched_artifact(
    artifact: dict[str, Any],
    frame: pd.DataFrame,
    features: list[str],
    assignment: np.ndarray,
    repeat: int,
    outer_seed: int,
    outer_fold: int,
    analysis_signature: str,
) -> tuple[dict[str, np.ndarray], dict[str, np.ndarray], float, float, float, int, int, int, int]:
    train = frame.loc[assignment != outer_fold].copy().reset_index(drop=True)
    test = frame.loc[assignment == outer_fold].copy().reset_index(drop=True)
    train_ids = train["row_id"].astype(int).tolist()
    test_ids = test["row_id"].astype(int).tolist()
    expected = matched_signature(
        analysis_signature, repeat, outer_seed, outer_fold, train_ids, test_ids, features
    )
    if artifact.get("artifact_signature") != expected:
        raise RuntimeError("STOP_DATA_OR_ARTIFACT_FAIL: matched artifact signature mismatch")
    if not np.array_equal(np.asarray(artifact["train_row_ids"], dtype=int), np.asarray(train_ids, dtype=int)):
        raise RuntimeError("STOP_DATA_OR_ARTIFACT_FAIL: matched artifact train identity mismatch")
    if not np.array_equal(np.asarray(artifact["test_row_ids"], dtype=int), np.asarray(test_ids, dtype=int)):
        raise RuntimeError("STOP_DATA_OR_ARTIFACT_FAIL: matched artifact test identity mismatch")

    frame_by_row = frame.set_index("row_id", drop=False)
    max_projection_error = 0.0
    seen_holdouts: list[int] = []
    for bundle in artifact["projection_bundles"]:
        max_projection_error = max(max_projection_error, verify_projection_bundle(bundle, frame_by_row, features))
        seen_holdouts.extend(np.asarray(bundle["holdout_row_ids"], dtype=int).tolist())
    if sorted(seen_holdouts) != sorted(train_ids) or len(seen_holdouts) != len(set(seen_holdouts)):
        raise RuntimeError("STOP_DATA_OR_ARTIFACT_FAIL: cross-fitted training score coverage failed")
    max_projection_error = max(
        max_projection_error,
        verify_projection_bundle(artifact["final_projection_bundle"], frame_by_row, features),
    )

    max_validation_error = 0.0
    max_outer_error = 0.0
    probabilities: dict[str, np.ndarray] = {}
    for variant_name, variant in artifact["downstream_variants"].items():
        validation_error, outer_error = verify_downstream_variant(variant, frame_by_row)
        max_validation_error = max(max_validation_error, validation_error)
        max_outer_error = max(max_outer_error, outer_error)
        probabilities[variant_name] = np.asarray(variant["test_probability"], dtype=float)
    if max(max_projection_error, max_validation_error, max_outer_error) > PARITY_TOLERANCE:
        raise RuntimeError("STOP_DATA_OR_ARTIFACT_FAIL: matched artifact reload parity failed")

    projection_tuning_count = sum(
        len(target["tuning_fits"])
        for bundle in [*artifact["projection_bundles"], artifact["final_projection_bundle"]]
        for target in bundle["targets"].values()
    )
    projection_final_count = sum(
        1
        for bundle in [*artifact["projection_bundles"], artifact["final_projection_bundle"]]
        for _target in bundle["targets"].values()
    )
    downstream_tuning_count = sum(len(variant["tuning_fits"]) for variant in artifact["downstream_variants"].values())
    downstream_final_count = len(artifact["downstream_variants"])
    return (
        probabilities,
        {key: np.asarray(value, dtype=float) for key, value in artifact["test_scores"].items()},
        max_projection_error,
        max_validation_error,
        max_outer_error,
        projection_tuning_count,
        projection_final_count,
        downstream_tuning_count,
        downstream_final_count,
    )


def fit_or_load_matched_artifact(
    frame: pd.DataFrame,
    features: list[str],
    assignment: np.ndarray,
    repeat: int,
    outer_seed: int,
    outer_fold: int,
    analysis_signature: str,
) -> tuple[dict[str, np.ndarray], dict[str, np.ndarray], dict[str, Any]]:
    path = MATCHED_MODEL_DIR / f"repeat_{repeat:02d}" / f"outer_fold_{outer_fold:02d}.joblib"
    newly_fitted = False
    if path.exists():
        artifact = joblib.load(path)
    else:
        artifact = fit_matched_artifact(
            frame, features, assignment, repeat, outer_seed, outer_fold, analysis_signature
        )
        save_joblib_atomic(artifact, path)
        newly_fitted = True
    reloaded = joblib.load(path)
    (
        probabilities,
        scores,
        projection_error,
        validation_error,
        outer_error,
        projection_tuning_count,
        projection_final_count,
        downstream_tuning_count,
        downstream_final_count,
    ) = verify_matched_artifact(
        reloaded, frame, features, assignment, repeat, outer_seed, outer_fold, analysis_signature
    )
    metadata = {
        "stage": "matched_scalar",
        "repeat": repeat,
        "outer_seed": outer_seed,
        "outer_fold": outer_fold,
        "artifact_path": str(path.relative_to(ROOT)),
        "artifact_sha256": sha256(path),
        "newly_fitted": newly_fitted,
        "projection_tuning_estimator_count": projection_tuning_count,
        "projection_final_estimator_count": projection_final_count,
        "downstream_tuning_estimator_count": downstream_tuning_count,
        "downstream_final_estimator_count": downstream_final_count,
        "max_projection_score_error": projection_error,
        "max_downstream_validation_error": validation_error,
        "max_downstream_outer_error": outer_error,
    }
    return probabilities, scores, metadata


def build_bootstrap_indices(frame: pd.DataFrame) -> np.ndarray:
    strata = np.char.add(
        np.char.add(
            np.char.add(frame["death_5y"].to_numpy(int).astype(str), "_"),
            age_decade(frame["age_years"]).astype(str),
        ),
        np.char.add("_", frame["cycle"].astype(str).to_numpy()),
    )
    groups = [np.flatnonzero(strata == label) for label in sorted(np.unique(strata))]
    if min(len(group) for group in groups) <= 0:
        raise RuntimeError("STOP_DATA_OR_ARTIFACT_FAIL: bootstrap stratum is empty")
    rng = np.random.default_rng(BOOTSTRAP_SEED)
    indices = np.empty((N_BOOTSTRAPS, len(frame)), dtype=np.int32)
    for bootstrap in range(N_BOOTSTRAPS):
        offset = 0
        for group in groups:
            sample = rng.choice(group, size=len(group), replace=True).astype(np.int32)
            indices[bootstrap, offset : offset + len(group)] = sample
            offset += len(group)
    return indices


def interval_from_vector(vector: np.ndarray, indices: np.ndarray) -> tuple[float, float, float]:
    estimate = float(np.mean(vector))
    draws = np.asarray(vector, dtype=float)[indices].mean(axis=1)
    q025, q975 = np.quantile(draws, [0.025, 0.975])
    return estimate, float(q025), float(q975)


def markdown_table(frame: pd.DataFrame, digits: int = 6) -> str:
    display = frame.copy()
    for column in display.select_dtypes(include=[np.number]).columns:
        display[column] = display[column].map(lambda value: f"{value:.{digits}g}" if pd.notna(value) else "")
    headers = "| " + " | ".join(display.columns) + " |"
    divider = "| " + " | ".join(["---"] * len(display.columns)) + " |"
    rows = ["| " + " | ".join(map(str, row)) + " |" for row in display.itertuples(index=False, name=None)]
    return "\n".join([headers, divider, *rows])


def build_signature() -> tuple[str, pd.DataFrame, dict[str, Any]]:
    input_paths: list[tuple[str, Path]] = [
        ("analysis_ready_source", SOURCE_FRAME),
        ("participant_folds", SOURCE_FOLDS),
        ("ordered_feature_tiers", SOURCE_TIERS),
        ("source_bootstrap_indices", SOURCE_BOOTSTRAP),
        ("predecessor_model_audit", SOURCE_PREDECESSOR_AUDIT),
        ("source_oof_predictions", SOURCE_OOF),
        ("source_model_audit", SOURCE_AUDIT),
        ("source_analysis_signature", SOURCE_SIGNATURE),
        ("frozen_specification", SPEC),
        ("base_analysis_script", BASE_SCRIPT),
        ("analysis_script", Path(__file__)),
        ("environment_policy", ENV_POLICY),
        ("environment_manifest", ENV_MANIFEST),
        ("environment_lock", ENV_LOCK),
    ]
    for outer_fold in range(N_OUTER_FOLDS):
        input_paths.append((f"source_age_model_outer_{outer_fold}", SOURCE / "models" / f"outer_fold_{outer_fold:02d}.joblib"))
        input_paths.append(
            (
                f"source_age_crossfit_outer_{outer_fold}",
                SOURCE / f"outer_fold_{outer_fold:02d}_training_crossfit_scores.tsv.gz",
            )
        )
    for _, path in input_paths:
        if not path.exists():
            raise RuntimeError(f"Required input is missing: {path}")
    manifest = pd.DataFrame(
        [
            {"role": role, "path": str(path.relative_to(ROOT)), "bytes": path.stat().st_size, "sha256": sha256(path)}
            for role, path in input_paths
        ]
    )
    payload = {
        "analysis_version": ANALYSIS_VERSION,
        "schema_version": SCHEMA_VERSION,
        "inputs": manifest[["role", "sha256"]].to_dict(orient="records"),
        "population": "NHANES_2007_2012_complete_support_ages_20_79_S3_50",
        "outer_seeds": OUTER_SEEDS,
        "outer_folds": N_OUTER_FOLDS,
        "downstream_folds": N_DOWNSTREAM_FOLDS,
        "projection_tuning_folds": N_PROJECTION_TUNING_FOLDS,
        "projection_targets": ["standardized_chronological_age", "standardized_five_year_mortality"],
        "projection_model": "ridge_squared_error",
        "alpha_grid": ALPHA_GRID,
        "downstream_models": ["linear_scalar", "five_knot_scalar_spline"],
        "C_grid": C_GRID,
        "bootstrap_seed": BOOTSTRAP_SEED,
        "bootstraps": N_BOOTSTRAPS,
        "parity_tolerance": PARITY_TOLERANCE,
    }
    return stable_hash(payload), manifest, payload


def write_report(
    decision: dict[str, Any],
    flex_metrics: pd.DataFrame,
    flex_fold: pd.DataFrame,
    repeat_contrasts: pd.DataFrame,
    cycle_contrasts: pd.DataFrame,
    matched_metrics: pd.DataFrame,
    audit: dict[str, Any],
) -> None:
    flex = decision["flexible_calibration"]
    matched = decision["matched_scalar"]
    text = f"""# NHANES five-year matched scalar target-supervision probe

Date: 26 August 2026

## Decisions

- Flexible calibration: **{flex['decision_category']}**.
- Matched one-dimensional comparison: **{matched['decision_category']}**.

The fixed five-knot calibration of the frozen age score had log loss {flex['M_CLOCK_FLEX_log_loss']:.6f}. Its gap versus the frozen direct 50-measurement model was {flex['V_SOURCE_FLEX']:.6f} (participant-bootstrap interval {flex['q025']:.6f} to {flex['q975']:.6f}); {flex['positive_outer_folds']} of five original folds were positive.

Across five prespecified repeated outer partitions, the participant-averaged primary contrast `L(q_A spline)-L(q_Y spline)` was {matched['D_TARGET']:.6f} (participant-bootstrap interval {matched['q025']:.6f} to {matched['q975']:.6f}). {matched['positive_repeats']} of five repeat estimates were positive and {matched['nonreversing_cycles']} of three cycle estimates were non-negative.

This is a finite-sample comparison for one population, endpoint, 50-measurement panel, linear projection family and scalar dimension. Mortality supervision does not identify biological age, surrogate validity or causal benefit.

## Flexible-calibration models

{markdown_table(flex_metrics)}

## Flexible-calibration fold contrasts

{markdown_table(flex_fold)}

## Matched-scalar repeat contrasts

{markdown_table(repeat_contrasts)}

## Matched-scalar cycle contrasts

{markdown_table(cycle_contrasts)}

## Participant-averaged matched-scalar metrics

{markdown_table(matched_metrics)}

## Persistence and parity

- Artifact audit: **{audit['status']}**.
- Flexible artifacts: {audit['flexible_artifact_count']} of 5.
- Matched artifacts: {audit['matched_artifact_count']} of 25.
- Total persisted estimators: {audit['total_estimator_count']}.
- Newly fitted artifacts: {audit['newly_fitted_artifact_count']}; reused compatible artifacts: {audit['reused_artifact_count']}.
- Maximum reloaded projection-score error: {audit['max_projection_score_error']:.3g}.
- Maximum reloaded downstream validation-probability error: {audit['max_downstream_validation_error']:.3g}.
- Maximum reloaded outer-probability error: {audit['max_downstream_outer_error']:.3g}.

The paired interval averages each participant's loss across repeats before resampling. Repeated partitions are not treated as independent samples. Repeat- and cycle-specific results are consistency diagnostics, not replication tests.
"""
    (OUT / "REPORT.md").write_text(text, encoding="utf-8")


def write_artifact_manifest() -> None:
    rows = []
    for path in sorted(item for item in OUT.rglob("*") if item.is_file() and item.name != "artifact_manifest.tsv"):
        rows.append(
            {
                "path": str(path.relative_to(ROOT)),
                "bytes": path.stat().st_size,
                "sha256": sha256(path),
            }
        )
    pd.DataFrame(rows).to_csv(OUT / "artifact_manifest.tsv", sep="\t", index=False)


def main() -> int:
    configure_stdout()
    OUT.mkdir(parents=True, exist_ok=True)
    FLEX_MODEL_DIR.mkdir(parents=True, exist_ok=True)
    MATCHED_MODEL_DIR.mkdir(parents=True, exist_ok=True)
    analysis_signature, input_manifest, signature_payload = build_signature()
    signature_path = OUT / "analysis_signature.json"
    if signature_path.exists():
        existing = json.loads(signature_path.read_text(encoding="utf-8"))
        if existing.get("analysis_signature") != analysis_signature:
            raise RuntimeError("STOP_DATA_OR_ARTIFACT_FAIL: incompatible output analysis signature")
    else:
        write_json(signature_path, {"analysis_signature": analysis_signature, "signature_payload": signature_payload})
    input_manifest.to_csv(OUT / "input_manifest.tsv", sep="\t", index=False)

    frame, features, source_oof, source_artifacts = load_and_gate()
    pd.DataFrame({"order": np.arange(1, len(features) + 1), "feature": features}).to_csv(
        OUT / "ordered_source_features.tsv", sep="\t", index=False
    )
    bootstrap_indices = build_bootstrap_indices(frame)
    np.savez_compressed(
        OUT / "bootstrap_indices.npz",
        indices=bootstrap_indices,
        seed=np.asarray([BOOTSTRAP_SEED], dtype=np.int64),
    )

    artifact_rows: list[dict[str, Any]] = []

    # Stage 1: fixed flexible calibration of the frozen fold-local age score.
    print("Stage 1/2: flexible calibration of frozen age scores", flush=True)
    flexible_probability = np.full(EXPECTED_N, np.nan, dtype=float)
    for outer_fold in range(N_OUTER_FOLDS):
        print(f"  flexible outer fold {outer_fold + 1}/{N_OUTER_FOLDS}", flush=True)
        probability, metadata = fit_or_load_flexible_artifact(
            frame, source_artifacts[outer_fold], outer_fold, analysis_signature
        )
        mask = frame["outer_fold"].eq(outer_fold).to_numpy()
        flexible_probability[mask] = probability
        artifact_rows.append(metadata)
    if not np.isfinite(flexible_probability).all():
        raise RuntimeError("STOP_DATA_OR_ARTIFACT_FAIL: incomplete flexible-calibration prediction")

    y = frame["death_5y"].to_numpy(int)
    flex_predictions = frame[["row_id", "SEQN", "cycle", "age_years", "death_5y", "outer_fold"]].copy()
    flex_predictions["M_CLOCK_LINEAR"] = source_oof["M_CLOCK"].to_numpy(float)
    flex_predictions["M_CLOCK_FLEX"] = flexible_probability
    flex_predictions["M_X"] = source_oof["M_X"].to_numpy(float)
    flex_predictions.to_csv(OUT / "flexible_calibration_oof_predictions.tsv.gz", sep="\t", index=False, compression="gzip")
    flex_losses = flex_predictions[["row_id", "SEQN", "cycle", "age_years", "death_5y", "outer_fold"]].copy()
    flex_metric_rows = []
    for model in ("M_CLOCK_LINEAR", "M_CLOCK_FLEX", "M_X"):
        flex_losses[model] = participant_loss(y, flex_predictions[model].to_numpy(float))
        flex_metric_rows.append({"model": model, **endpoint_metrics(y, flex_predictions[model].to_numpy(float))})
    flex_losses.to_csv(OUT / "flexible_calibration_participant_losses.tsv.gz", sep="\t", index=False, compression="gzip")
    flex_metrics = pd.DataFrame(flex_metric_rows)
    flex_metrics.to_csv(OUT / "flexible_calibration_metrics.tsv", sep="\t", index=False)
    flex_vector = flex_losses["M_CLOCK_FLEX"].to_numpy(float) - flex_losses["M_X"].to_numpy(float)
    flex_estimate, flex_q025, flex_q975 = interval_from_vector(flex_vector, bootstrap_indices)
    flex_fold_rows = []
    for outer_fold, subset in flex_losses.groupby("outer_fold", sort=True):
        contrast = float((subset["M_CLOCK_FLEX"] - subset["M_X"]).mean())
        flex_fold_rows.append(
            {
                "outer_fold": int(outer_fold),
                "n": int(len(subset)),
                "events": int(subset["death_5y"].sum()),
                "V_SOURCE_FLEX": contrast,
            }
        )
    flex_fold = pd.DataFrame(flex_fold_rows)
    flex_fold.to_csv(OUT / "flexible_calibration_fold_contrasts.tsv", sep="\t", index=False)
    positive_flex_folds = int((flex_fold["V_SOURCE_FLEX"] > 0).sum())
    flex_decision = (
        "HUM04_ROBUST_TO_FLEX_CALIBRATION"
        if flex_estimate > 0 and flex_q025 > 0 and positive_flex_folds >= 4
        else "CALIBRATION_SENSITIVE"
    )
    flex_metric_lookup = flex_metrics.set_index("model")

    # Stage 2: matched scalar target-supervision comparison.
    print("Stage 2/2: matched scalar target-supervision repeats", flush=True)
    repeat_fold_rows = []
    prediction_rows = []
    fold_identity_rows = []
    nested_identity_rows = []
    for repeat, outer_seed in enumerate(OUTER_SEEDS):
        assignment = assign_outer_folds(frame, outer_seed)
        if repeat == 0 and not np.array_equal(assignment, frame["outer_fold"].to_numpy(int)):
            raise RuntimeError("STOP_DATA_OR_ARTIFACT_FAIL: original repeated partition changed")
        for position, row in frame.iterrows():
            fold_identity_rows.append(
                {
                    "repeat": repeat,
                    "outer_seed": outer_seed,
                    "row_id": int(row["row_id"]),
                    "SEQN": int(row["SEQN"]),
                    "outer_fold": int(assignment[position]),
                }
            )
        for outer_fold in range(N_OUTER_FOLDS):
            print(f"  repeat {repeat + 1}/{len(OUTER_SEEDS)}, outer fold {outer_fold + 1}/{N_OUTER_FOLDS}", flush=True)
            probabilities, scores, metadata = fit_or_load_matched_artifact(
                frame, features, assignment, repeat, outer_seed, outer_fold, analysis_signature
            )
            artifact_rows.append(metadata)
            mask = assignment == outer_fold
            test = frame.loc[mask].copy().reset_index(drop=True)
            if len(test) != len(next(iter(probabilities.values()))):
                raise RuntimeError("STOP_DATA_OR_ARTIFACT_FAIL: matched held-out length changed")
            for index, row in test.iterrows():
                prediction_rows.append(
                    {
                        "repeat": repeat,
                        "outer_seed": outer_seed,
                        "outer_fold": outer_fold,
                        "row_id": int(row["row_id"]),
                        "SEQN": int(row["SEQN"]),
                        "cycle": row["cycle"],
                        "age_years": float(row["age_years"]),
                        "death_5y": int(row["death_5y"]),
                        "q_A": float(scores["age"][index]),
                        "q_Y": float(scores["mortality"][index]),
                        "M_QA_LINEAR": float(probabilities["age_linear"][index]),
                        "M_QA_SPLINE": float(probabilities["age_spline"][index]),
                        "M_QY_LINEAR": float(probabilities["mortality_linear"][index]),
                        "M_QY_SPLINE": float(probabilities["mortality_spline"][index]),
                    }
                )
            artifact = joblib.load(ROOT / metadata["artifact_path"])
            for split in artifact["downstream_splits"]:
                for row_id in split["valid_row_ids"]:
                    nested_identity_rows.append(
                        {
                            "repeat": repeat,
                            "outer_seed": outer_seed,
                            "outer_fold": outer_fold,
                            "stage": "downstream",
                            "downstream_fold": int(split["downstream_fold"]),
                            "projection_inner_fold": -1,
                            "row_id": int(row_id),
                            "role": "valid",
                        }
                    )
            for downstream_fold, bundle in enumerate(artifact["projection_bundles"]):
                for target_name, target in bundle["targets"].items():
                    selected_alpha = float(target["selected_alpha"])
                    for fit in target["tuning_fits"]:
                        if fit["alpha"] != selected_alpha:
                            continue
                        for row_id in fit["valid_row_ids"]:
                            nested_identity_rows.append(
                                {
                                    "repeat": repeat,
                                    "outer_seed": outer_seed,
                                    "outer_fold": outer_fold,
                                    "stage": f"projection_{target_name}",
                                    "downstream_fold": downstream_fold,
                                    "projection_inner_fold": int(fit["inner_fold"]),
                                    "row_id": int(row_id),
                                    "role": "valid",
                                }
                            )

    folds_table = pd.DataFrame(fold_identity_rows)
    folds_table.to_csv(OUT / "participant_repeat_folds.tsv.gz", sep="\t", index=False, compression="gzip")
    pd.DataFrame(nested_identity_rows).to_csv(
        OUT / "nested_split_identities.tsv.gz", sep="\t", index=False, compression="gzip"
    )
    predictions = pd.DataFrame(prediction_rows).sort_values(["repeat", "row_id"]).reset_index(drop=True)
    if len(predictions) != EXPECTED_N * len(OUTER_SEEDS):
        raise RuntimeError("STOP_DATA_OR_ARTIFACT_FAIL: repeated OOF prediction row count failed")
    if not predictions.groupby("repeat")["row_id"].nunique().eq(EXPECTED_N).all():
        raise RuntimeError("STOP_DATA_OR_ARTIFACT_FAIL: repeated OOF participant coverage failed")
    predictions.to_csv(OUT / "matched_scalar_oof_predictions.tsv.gz", sep="\t", index=False, compression="gzip")

    model_columns = ("M_QA_LINEAR", "M_QA_SPLINE", "M_QY_LINEAR", "M_QY_SPLINE")
    losses = predictions[["repeat", "outer_seed", "outer_fold", "row_id", "SEQN", "cycle", "age_years", "death_5y"]].copy()
    for model in model_columns:
        losses[model] = participant_loss(predictions["death_5y"].to_numpy(int), predictions[model].to_numpy(float))
    losses.to_csv(OUT / "matched_scalar_participant_losses.tsv.gz", sep="\t", index=False, compression="gzip")

    repeat_metric_rows = []
    repeat_contrast_rows = []
    for repeat, subset in predictions.groupby("repeat", sort=True):
        repeat_losses = losses.loc[losses["repeat"].eq(repeat)]
        for model in model_columns:
            repeat_metric_rows.append(
                {
                    "repeat": int(repeat),
                    "outer_seed": int(subset["outer_seed"].iloc[0]),
                    "model": model,
                    **endpoint_metrics(subset["death_5y"].to_numpy(int), subset[model].to_numpy(float)),
                }
            )
        repeat_contrast_rows.append(
            {
                "repeat": int(repeat),
                "outer_seed": int(subset["outer_seed"].iloc[0]),
                "D_TARGET_SPLINE": float((repeat_losses["M_QA_SPLINE"] - repeat_losses["M_QY_SPLINE"]).mean()),
                "D_TARGET_LINEAR": float((repeat_losses["M_QA_LINEAR"] - repeat_losses["M_QY_LINEAR"]).mean()),
            }
        )
    repeat_metrics = pd.DataFrame(repeat_metric_rows)
    repeat_metrics.to_csv(OUT / "matched_scalar_repeat_metrics.tsv", sep="\t", index=False)
    repeat_contrasts = pd.DataFrame(repeat_contrast_rows)
    repeat_contrasts.to_csv(OUT / "matched_scalar_repeat_contrasts.tsv", sep="\t", index=False)

    average_losses = losses.groupby(["row_id", "SEQN", "cycle", "age_years", "death_5y"], as_index=False)[list(model_columns)].mean()
    if len(average_losses) != EXPECTED_N:
        raise RuntimeError("STOP_DATA_OR_ARTIFACT_FAIL: participant-average loss identity failed")
    average_losses.to_csv(OUT / "matched_scalar_participant_average_losses.tsv.gz", sep="\t", index=False, compression="gzip")
    primary_vector = average_losses["M_QA_SPLINE"].to_numpy(float) - average_losses["M_QY_SPLINE"].to_numpy(float)
    target_estimate, target_q025, target_q975 = interval_from_vector(primary_vector, bootstrap_indices)
    cycle_rows = []
    for cycle, subset in average_losses.groupby("cycle", sort=True):
        cycle_rows.append(
            {
                "cycle": cycle,
                "n": int(len(subset)),
                "events": int(subset["death_5y"].sum()),
                "D_TARGET_SPLINE": float((subset["M_QA_SPLINE"] - subset["M_QY_SPLINE"]).mean()),
                "D_TARGET_LINEAR": float((subset["M_QA_LINEAR"] - subset["M_QY_LINEAR"]).mean()),
            }
        )
    cycle_contrasts = pd.DataFrame(cycle_rows)
    cycle_contrasts.to_csv(OUT / "matched_scalar_cycle_contrasts.tsv", sep="\t", index=False)
    positive_repeats = int((repeat_contrasts["D_TARGET_SPLINE"] > 0).sum())
    nonreversing_cycles = int((cycle_contrasts["D_TARGET_SPLINE"] >= 0).sum())
    if target_estimate <= 0 or target_q025 <= 0:
        matched_decision = "NO_MATCHED_TARGET_ADVANTAGE"
    elif positive_repeats >= 4 and nonreversing_cycles == len(EXPECTED_CYCLES):
        matched_decision = "MATCHED_TARGET_SCALAR_ADVANTAGE"
    else:
        matched_decision = "SPLIT_OR_CYCLE_SENSITIVE"

    average_predictions = predictions.groupby(["row_id", "SEQN", "cycle", "age_years", "death_5y"], as_index=False)[list(model_columns)].mean()
    matched_metric_rows = []
    for model in model_columns:
        matched_metric_rows.append(
            {
                "model": model,
                **endpoint_metrics(average_predictions["death_5y"].to_numpy(int), average_predictions[model].to_numpy(float)),
            }
        )
    matched_metrics = pd.DataFrame(matched_metric_rows)
    matched_metrics.to_csv(OUT / "matched_scalar_participant_average_metrics.tsv", sep="\t", index=False)

    artifact_index = pd.DataFrame(artifact_rows)
    artifact_index.to_csv(OUT / "model_artifact_index.tsv", sep="\t", index=False)
    audit = {
        "status": "PASS",
        "analysis_signature": analysis_signature,
        "expected_flexible_artifact_count": 5,
        "flexible_artifact_count": int(artifact_index["stage"].eq("flexible_calibration").sum()),
        "expected_matched_artifact_count": 25,
        "matched_artifact_count": int(artifact_index["stage"].eq("matched_scalar").sum()),
        "newly_fitted_artifact_count": int(artifact_index["newly_fitted"].sum()),
        "reused_artifact_count": int((~artifact_index["newly_fitted"]).sum()),
        "projection_tuning_estimator_count": int(artifact_index["projection_tuning_estimator_count"].sum()),
        "projection_final_estimator_count": int(artifact_index["projection_final_estimator_count"].sum()),
        "downstream_tuning_estimator_count": int(artifact_index["downstream_tuning_estimator_count"].sum()),
        "downstream_final_estimator_count": int(artifact_index["downstream_final_estimator_count"].sum()),
        "total_estimator_count": int(
            artifact_index[
                [
                    "projection_tuning_estimator_count",
                    "projection_final_estimator_count",
                    "downstream_tuning_estimator_count",
                    "downstream_final_estimator_count",
                ]
            ].to_numpy().sum()
        ),
        "parity_tolerance": PARITY_TOLERANCE,
        "max_projection_score_error": float(artifact_index["max_projection_score_error"].max()),
        "max_downstream_validation_error": float(artifact_index["max_downstream_validation_error"].max()),
        "max_downstream_outer_error": float(artifact_index["max_downstream_outer_error"].max()),
        "all_model_hashes_present": bool(artifact_index["artifact_sha256"].str.len().eq(64).all()),
        "software": {
            "python": sys.version,
            "python_executable": sys.executable,
            "platform": platform.platform(),
            "numpy": np.__version__,
            "pandas": pd.__version__,
            "scikit_learn": sklearn.__version__,
            "joblib": joblib.__version__,
        },
    }
    if (
        audit["flexible_artifact_count"] != audit["expected_flexible_artifact_count"]
        or audit["matched_artifact_count"] != audit["expected_matched_artifact_count"]
        or audit["max_projection_score_error"] > PARITY_TOLERANCE
        or audit["max_downstream_validation_error"] > PARITY_TOLERANCE
        or audit["max_downstream_outer_error"] > PARITY_TOLERANCE
        or not audit["all_model_hashes_present"]
    ):
        audit["status"] = "FAIL"
    write_json(OUT / "model_artifact_audit.json", audit)

    decision = {
        "analysis_version": ANALYSIS_VERSION,
        "population": "NHANES 2007-2012 complete-support 50-measurement interface, ages 20-79",
        "n": EXPECTED_N,
        "events": EXPECTED_EVENTS,
        "flexible_calibration": {
            "decision_category": flex_decision,
            "M_CLOCK_LINEAR_log_loss": float(flex_metric_lookup.loc["M_CLOCK_LINEAR", "log_loss"]),
            "M_CLOCK_FLEX_log_loss": float(flex_metric_lookup.loc["M_CLOCK_FLEX", "log_loss"]),
            "M_X_log_loss": float(flex_metric_lookup.loc["M_X", "log_loss"]),
            "V_SOURCE_FLEX": flex_estimate,
            "q025": flex_q025,
            "q975": flex_q975,
            "positive_outer_folds": positive_flex_folds,
        },
        "matched_scalar": {
            "decision_category": matched_decision,
            "D_TARGET": target_estimate,
            "q025": target_q025,
            "q975": target_q975,
            "positive_repeats": positive_repeats,
            "nonreversing_cycles": nonreversing_cycles,
            "outer_seeds": OUTER_SEEDS,
        },
        "artifact_status": audit["status"],
        "claim_boundary": "One population, endpoint, 50-measurement panel, fixed linear projection family and scalar dimension; not biological-age identification, surrogate validity or causal benefit.",
    }
    if audit["status"] != "PASS":
        decision["flexible_calibration"]["decision_category"] = "STOP_DATA_OR_ARTIFACT_FAIL"
        decision["matched_scalar"]["decision_category"] = "STOP_DATA_OR_ARTIFACT_FAIL"
    write_json(OUT / "decision.json", decision)
    write_report(decision, flex_metrics, flex_fold, repeat_contrasts, cycle_contrasts, matched_metrics, audit)
    write_artifact_manifest()
    print(json.dumps(json_ready(decision), indent=2, ensure_ascii=False), flush=True)
    print(f"Wrote {OUT}", flush=True)
    return 0 if audit["status"] == "PASS" else 2


if __name__ == "__main__":
    raise SystemExit(main())
```
