## Supplementary_Software_1 for "Inferential boundaries of age prediction: why prediction does not establish biological age measurement": analyze_nhanes_5y_matched_scalar_target_supervision_probe_v0_2.py

```
from __future__ import annotations

import importlib.util
import json
import sys
from pathlib import Path
from typing import Any

import joblib
import numpy as np
import pandas as pd
from sklearn.metrics import log_loss


ROOT = Path(__file__).resolve().parents[1]
CORE_SCRIPT = ROOT / "analysis" / "analyze_nhanes_5y_matched_scalar_target_supervision_probe_v0_1.py"
SPEC = ROOT / "analysis" / "NHANES_5Y_MATCHED_SCALAR_TARGET_SUPERVISION_SPEC_v0_2_20260826.md"
OUT = ROOT / "outputs" / "nhanes_5y_matched_scalar_target_supervision_probe_v0_2"
ANALYSIS_VERSION = "nhanes_5y_matched_scalar_target_supervision_probe_v0_2"
SCHEMA_VERSION = "2.0"
NESTING_VERSION = "fully_nested_downstream_projection_v0_2"


def load_core() -> Any:
    module_spec = importlib.util.spec_from_file_location("nhanes_matched_scalar_v01_core", CORE_SCRIPT)
    if module_spec is None or module_spec.loader is None:
        raise RuntimeError("Could not load frozen HUM-06 v0.1 implementation")
    module = importlib.util.module_from_spec(module_spec)
    module_spec.loader.exec_module(module)
    return module


core = load_core()
core.ANALYSIS_VERSION = ANALYSIS_VERSION
core.SCHEMA_VERSION = SCHEMA_VERSION
core.OUT = OUT
core.MODEL_DIR = OUT / "models"
core.FLEX_MODEL_DIR = core.MODEL_DIR / "flexible_calibration"
core.MATCHED_MODEL_DIR = core.MODEL_DIR / "matched_scalar"
core.SPEC = SPEC


def require(condition: bool, message: str) -> None:
    if not condition:
        raise RuntimeError(f"STOP_DATA_OR_ARTIFACT_FAIL: {message}")


def as_ids(values: Any) -> np.ndarray:
    return np.asarray(values, dtype=int)


def maximum_error(left: Any, right: Any) -> float:
    a = np.asarray(left, dtype=float)
    b = np.asarray(right, dtype=float)
    require(a.shape == b.shape, "score or prediction shape mismatch")
    return 0.0 if a.size == 0 else float(np.max(np.abs(a - b)))


def matched_signature_v2(
    analysis_signature: str,
    repeat: int,
    outer_seed: int,
    outer_fold: int,
    train_ids: list[int],
    test_ids: list[int],
    features: list[str],
) -> str:
    return core.stable_hash(
        {
            "analysis_signature": analysis_signature,
            "stage": "matched_scalar_target_supervision",
            "nesting_version": NESTING_VERSION,
            "repeat": int(repeat),
            "outer_seed": int(outer_seed),
            "outer_fold": int(outer_fold),
            "train_row_ids": train_ids,
            "test_row_ids": test_ids,
            "features": features,
            "alpha_grid": core.ALPHA_GRID,
            "C_grid": core.C_GRID,
            "downstream_seed": int(outer_seed + 1000 + outer_fold),
            "projection_seed_formula": "outer_seed+2000+10*outer_fold+downstream_fold",
            "inner_training_score_rule": "three_fold_projection_crossfit_wholly_within_downstream_training",
            "inner_validation_score_rule": "projection_selected_and_fitted_on_complete_downstream_training",
        }
    )


def build_fully_nested_score_chain(
    downstream_training: pd.DataFrame,
    downstream_validation: pd.DataFrame,
    features: list[str],
    seed: int,
    downstream_fold: int,
) -> dict[str, Any]:
    score_splits = core.make_splits(downstream_training, core.N_PROJECTION_TUNING_FOLDS, seed)
    training_scores = {
        "age": np.full(len(downstream_training), np.nan, dtype=float),
        "mortality": np.full(len(downstream_training), np.nan, dtype=float),
    }
    training_projection_bundles = []
    split_records = []
    for projection_fold, (projection_train, projection_valid) in enumerate(score_splits):
        bundle = core.fit_projection_bundle(
            downstream_training.iloc[projection_train].copy().reset_index(drop=True),
            downstream_training.iloc[projection_valid].copy().reset_index(drop=True),
            features,
            seed,
            f"downstream_{downstream_fold}_training_projection_fold_{projection_fold}",
        )
        training_projection_bundles.append(bundle)
        split_records.append(
            {
                "projection_fold": int(projection_fold),
                "train_row_ids": downstream_training.iloc[projection_train]["row_id"].to_numpy(int),
                "valid_row_ids": downstream_training.iloc[projection_valid]["row_id"].to_numpy(int),
            }
        )
        for target in ("age", "mortality"):
            training_scores[target][projection_valid] = bundle["targets"][target]["holdout_score"]
    require(all(np.isfinite(values).all() for values in training_scores.values()), "incomplete nested training score")

    validation_bundle = core.fit_projection_bundle(
        downstream_training,
        downstream_validation,
        features,
        seed,
        f"downstream_{downstream_fold}_validation_projection",
    )
    validation_scores = {
        target: np.asarray(validation_bundle["targets"][target]["holdout_score"], dtype=float)
        for target in ("age", "mortality")
    }
    return {
        "downstream_fold": int(downstream_fold),
        "seed": int(seed),
        "training_row_ids": downstream_training["row_id"].to_numpy(int),
        "validation_row_ids": downstream_validation["row_id"].to_numpy(int),
        "training_projection_splits": split_records,
        "training_projection_bundles": training_projection_bundles,
        "validation_projection_bundle": validation_bundle,
        "training_scores": training_scores,
        "validation_scores": validation_scores,
    }


def tune_downstream_fully_nested(
    frame_by_row: pd.DataFrame,
    chains: list[dict[str, Any]],
    target: str,
    mode: str,
) -> dict[str, Any]:
    tuning_fits = []
    tuning_rows = []
    for c_value in core.C_GRID:
        for chain in chains:
            train_ids = as_ids(chain["training_row_ids"])
            valid_ids = as_ids(chain["validation_row_ids"])
            training = frame_by_row.loc[train_ids].reset_index(drop=True)
            validation = frame_by_row.loc[valid_ids].reset_index(drop=True)
            train_score = np.asarray(chain["training_scores"][target], dtype=float)
            valid_score = np.asarray(chain["validation_scores"][target], dtype=float)
            model = core.fit_downstream_model(training, train_score, c_value, mode)
            probability = core.predict_downstream_model(model, validation, valid_score)
            loss = float(log_loss(validation["death_5y"], probability, labels=[0, 1]))
            row = {
                "C": float(c_value),
                "downstream_fold": int(chain["downstream_fold"]),
                "train_row_ids": train_ids,
                "valid_row_ids": valid_ids,
                "train_score": train_score,
                "valid_score": valid_score,
                "model": model,
                "valid_probability": probability,
                "log_loss": loss,
            }
            tuning_fits.append(row)
            tuning_rows.append(
                {
                    "C": float(c_value),
                    "downstream_fold": int(chain["downstream_fold"]),
                    "log_loss": loss,
                }
            )
    tuning = pd.DataFrame(tuning_rows)
    selected_c = float(
        tuning.groupby("C", as_index=False)["log_loss"].mean().sort_values(["log_loss", "C"]).iloc[0]["C"]
    )
    return {"selected_C": selected_c, "tuning_fits": tuning_fits, "tuning_rows": tuning_rows}


def fit_matched_artifact_v2(
    frame: pd.DataFrame,
    features: list[str],
    assignment: np.ndarray,
    repeat: int,
    outer_seed: int,
    outer_fold: int,
    analysis_signature: str,
) -> dict[str, Any]:
    train = frame.loc[assignment != outer_fold].copy().reset_index(drop=True)
    test = frame.loc[assignment == outer_fold].copy().reset_index(drop=True)
    downstream_seed = int(outer_seed + 1000 + outer_fold)
    downstream_splits = core.make_splits(train, core.N_DOWNSTREAM_FOLDS, downstream_seed)

    # Scores used by the final outer-training calibration remain participant-level
    # cross-fitted scores; every participant is excluded from alpha selection and fit.
    training_scores = {
        "age": np.full(len(train), np.nan, dtype=float),
        "mortality": np.full(len(train), np.nan, dtype=float),
    }
    projection_bundles = []
    downstream_score_chains = []
    for downstream_fold, (downstream_train, downstream_valid) in enumerate(downstream_splits):
        projection_seed = int(outer_seed + 2000 + 10 * outer_fold + downstream_fold)
        outer_training_bundle = core.fit_projection_bundle(
            train.iloc[downstream_train].copy().reset_index(drop=True),
            train.iloc[downstream_valid].copy().reset_index(drop=True),
            features,
            projection_seed,
            f"outer_training_projection_fold_{downstream_fold}",
        )
        projection_bundles.append(outer_training_bundle)
        for target in ("age", "mortality"):
            training_scores[target][downstream_valid] = outer_training_bundle["targets"][target]["holdout_score"]

        downstream_score_chains.append(
            build_fully_nested_score_chain(
                train.iloc[downstream_train].copy().reset_index(drop=True),
                train.iloc[downstream_valid].copy().reset_index(drop=True),
                features,
                projection_seed,
                downstream_fold,
            )
        )
    require(all(np.isfinite(values).all() for values in training_scores.values()), "incomplete outer-training score")

    final_projection_seed = int(outer_seed + 2000 + 10 * outer_fold + core.N_DOWNSTREAM_FOLDS)
    final_bundle = core.fit_projection_bundle(train, test, features, final_projection_seed, "outer_test")
    test_scores = {
        target: np.asarray(final_bundle["targets"][target]["holdout_score"], dtype=float)
        for target in ("age", "mortality")
    }

    frame_by_row = frame.set_index("row_id", drop=False)
    downstream_variants: dict[str, Any] = {}
    for target in ("age", "mortality"):
        for mode in ("linear", "spline"):
            variant_name = f"{target}_{mode}"
            tuned = tune_downstream_fully_nested(frame_by_row, downstream_score_chains, target, mode)
            final_model = core.fit_downstream_model(train, training_scores[target], tuned["selected_C"], mode)
            test_probability = core.predict_downstream_model(final_model, test, test_scores[target])
            downstream_variants[variant_name] = {
                "target": target,
                "mode": mode,
                "selected_C": tuned["selected_C"],
                "tuning_fits": tuned["tuning_fits"],
                "tuning_rows": tuned["tuning_rows"],
                "final_model": final_model,
                "final_train_row_ids": train["row_id"].to_numpy(int),
                "final_train_score": np.asarray(training_scores[target], dtype=float),
                "test_row_ids": test["row_id"].to_numpy(int),
                "test_score": np.asarray(test_scores[target], dtype=float),
                "test_probability": test_probability,
            }

    train_ids = train["row_id"].astype(int).tolist()
    test_ids = test["row_id"].astype(int).tolist()
    return {
        "schema_version": SCHEMA_VERSION,
        "analysis_signature": analysis_signature,
        "artifact_signature": matched_signature_v2(
            analysis_signature, repeat, outer_seed, outer_fold, train_ids, test_ids, features
        ),
        "nesting_version": NESTING_VERSION,
        "repeat": int(repeat),
        "outer_seed": int(outer_seed),
        "outer_fold": int(outer_fold),
        "features": list(features),
        "train_row_ids": np.asarray(train_ids, dtype=int),
        "test_row_ids": np.asarray(test_ids, dtype=int),
        "downstream_seed": downstream_seed,
        "downstream_splits": [
            {
                "downstream_fold": int(index),
                "train_row_ids": train.iloc[split[0]]["row_id"].to_numpy(int),
                "valid_row_ids": train.iloc[split[1]]["row_id"].to_numpy(int),
            }
            for index, split in enumerate(downstream_splits)
        ],
        "projection_bundles": projection_bundles,
        "downstream_score_chains": downstream_score_chains,
        "final_projection_bundle": final_bundle,
        "training_scores": training_scores,
        "test_scores": test_scores,
        "downstream_variants": downstream_variants,
    }


def verify_matching_preprocessors(age_model: dict[str, Any], mortality_model: dict[str, Any]) -> None:
    require(age_model["design_feature_names"] == mortality_model["design_feature_names"], "qA/qY feature order differs")
    require(len(age_model["encoder"].categories_) == len(mortality_model["encoder"].categories_), "qA/qY encoder differs")
    for left, right in zip(age_model["encoder"].categories_, mortality_model["encoder"].categories_):
        require(np.array_equal(left, right), "qA/qY encoded categories differ")
    require(maximum_error(age_model["scaler"].mean_, mortality_model["scaler"].mean_) <= core.PARITY_TOLERANCE, "qA/qY scaler mean differs")
    require(maximum_error(age_model["scaler"].scale_, mortality_model["scaler"].scale_) <= core.PARITY_TOLERANCE, "qA/qY scaler scale differs")


def verify_projection_bundle_v2(
    bundle: dict[str, Any],
    frame_by_row: pd.DataFrame,
    features: list[str],
    expected_training_ids: np.ndarray,
    expected_holdout_ids: np.ndarray,
) -> tuple[float, float]:
    training_ids = as_ids(bundle["training_row_ids"])
    holdout_ids = as_ids(bundle["holdout_row_ids"])
    require(np.array_equal(training_ids, as_ids(expected_training_ids)), "projection training identity mismatch")
    require(np.array_equal(holdout_ids, as_ids(expected_holdout_ids)), "projection holdout identity mismatch")
    require(np.intersect1d(training_ids, holdout_ids).size == 0, "projection holdout identity leaked")
    require(set(bundle["targets"]) == {"age", "mortality"}, "projection target set changed")

    max_score_error = 0.0
    max_metric_error = 0.0
    target_columns = {"age": "age_years", "mortality": "death_5y"}
    for target_name, target_column in target_columns.items():
        target = bundle["targets"][target_name]
        require(target["target_column"] == target_column, "projection target column changed")
        require(len(target["tuning_fits"]) == len(core.ALPHA_GRID) * core.N_PROJECTION_TUNING_FOLDS, "projection tuning fit count changed")
        mean_rows = []
        for alpha in core.ALPHA_GRID:
            fits = [fit for fit in target["tuning_fits"] if float(fit["alpha"]) == float(alpha)]
            require(len(fits) == core.N_PROJECTION_TUNING_FOLDS, "projection alpha fold count changed")
            seen_valid: list[int] = []
            losses = []
            for fit in sorted(fits, key=lambda item: int(item["inner_fold"])):
                fit_train = as_ids(fit["train_row_ids"])
                fit_valid = as_ids(fit["valid_row_ids"])
                require(np.intersect1d(fit_train, fit_valid).size == 0, "projection tuning train/valid overlap")
                require(set(fit_train).union(set(fit_valid)) == set(training_ids), "projection tuning is not contained in bundle training")
                require(np.intersect1d(fit_valid, holdout_ids).size == 0, "projection tuning uses bundle holdout")
                model = fit["model"]
                require(model["target"] == target_column and float(model["alpha"]) == float(alpha), "stored projection model metadata changed")
                valid = frame_by_row.loc[fit_valid]
                reproduced = core.predict_projection(model, valid, features)
                score_error = maximum_error(reproduced, fit["valid_score"])
                max_score_error = max(max_score_error, score_error)
                expected_target = core.standardized_target_for_model(model, valid)
                reproduced_mse = float(np.mean((expected_target - reproduced) ** 2))
                metric_error = abs(reproduced_mse - float(fit["mse"]))
                max_metric_error = max(max_metric_error, metric_error)
                seen_valid.extend(fit_valid.tolist())
                losses.append(reproduced_mse)
            require(sorted(seen_valid) == sorted(training_ids.tolist()) and len(seen_valid) == len(set(seen_valid)), "projection tuning validation coverage failed")
            mean_rows.append({"alpha": float(alpha), "mse": float(np.mean(losses))})
        selected = float(pd.DataFrame(mean_rows).sort_values(["mse", "alpha"]).iloc[0]["alpha"])
        require(selected == float(target["selected_alpha"]), "selected projection alpha does not reconstruct")
        require(float(target["final_model"]["alpha"]) == selected, "final projection alpha differs from selected alpha")
        require(target["final_model"]["target"] == target_column, "final projection target changed")
        holdout = frame_by_row.loc[holdout_ids]
        reproduced = core.predict_projection(target["final_model"], holdout, features)
        max_score_error = max(max_score_error, maximum_error(reproduced, target["holdout_score"]))

    age_fits = bundle["targets"]["age"]["tuning_fits"]
    mortality_fits = bundle["targets"]["mortality"]["tuning_fits"]
    require(len(age_fits) == len(mortality_fits), "qA/qY tuning opportunity differs")
    for age_fit, mortality_fit in zip(age_fits, mortality_fits):
        require(float(age_fit["alpha"]) == float(mortality_fit["alpha"]), "qA/qY alpha grid order differs")
        require(int(age_fit["inner_fold"]) == int(mortality_fit["inner_fold"]), "qA/qY tuning fold differs")
        require(np.array_equal(as_ids(age_fit["train_row_ids"]), as_ids(mortality_fit["train_row_ids"])), "qA/qY tuning train identity differs")
        require(np.array_equal(as_ids(age_fit["valid_row_ids"]), as_ids(mortality_fit["valid_row_ids"])), "qA/qY tuning valid identity differs")
        verify_matching_preprocessors(age_fit["model"], mortality_fit["model"])
    verify_matching_preprocessors(
        bundle["targets"]["age"]["final_model"], bundle["targets"]["mortality"]["final_model"]
    )
    require(max_score_error <= core.PARITY_TOLERANCE, "projection reload score parity failed")
    require(max_metric_error <= core.PARITY_TOLERANCE, "projection tuning metric parity failed")
    return max_score_error, max_metric_error


def verify_matched_artifact_v2(
    artifact: dict[str, Any],
    frame: pd.DataFrame,
    features: list[str],
    assignment: np.ndarray,
    repeat: int,
    outer_seed: int,
    outer_fold: int,
    analysis_signature: str,
) -> dict[str, Any]:
    train = frame.loc[assignment != outer_fold].copy().reset_index(drop=True)
    test = frame.loc[assignment == outer_fold].copy().reset_index(drop=True)
    train_ids = train["row_id"].to_numpy(int)
    test_ids = test["row_id"].to_numpy(int)
    expected_signature = matched_signature_v2(
        analysis_signature,
        repeat,
        outer_seed,
        outer_fold,
        train_ids.tolist(),
        test_ids.tolist(),
        features,
    )
    require(artifact.get("artifact_signature") == expected_signature, "matched artifact signature mismatch")
    require(artifact.get("nesting_version") == NESTING_VERSION, "matched nesting version mismatch")
    require(artifact.get("features") == features, "matched feature order mismatch")
    require(np.array_equal(as_ids(artifact["train_row_ids"]), train_ids), "matched outer-train identity mismatch")
    require(np.array_equal(as_ids(artifact["test_row_ids"]), test_ids), "matched outer-test identity mismatch")
    require(np.intersect1d(train_ids, test_ids).size == 0, "matched outer identity overlap")

    downstream_splits = core.make_splits(train, core.N_DOWNSTREAM_FOLDS, int(outer_seed + 1000 + outer_fold))
    require(len(artifact["downstream_splits"]) == core.N_DOWNSTREAM_FOLDS, "downstream split count changed")
    require(len(artifact["projection_bundles"]) == core.N_DOWNSTREAM_FOLDS, "outer-training projection bundle count changed")
    require(len(artifact["downstream_score_chains"]) == core.N_DOWNSTREAM_FOLDS, "nested score chain count changed")
    frame_by_row = frame.set_index("row_id", drop=False)
    train_position = {int(row_id): index for index, row_id in enumerate(train_ids)}
    reconstructed_training_scores = {
        "age": np.full(len(train), np.nan, dtype=float),
        "mortality": np.full(len(train), np.nan, dtype=float),
    }
    max_projection_error = 0.0
    max_metric_error = 0.0
    max_chain_error = 0.0
    seen_outer_training_holdouts: list[int] = []

    for downstream_fold, (split_train, split_valid) in enumerate(downstream_splits):
        expected_train_ids = train.iloc[split_train]["row_id"].to_numpy(int)
        expected_valid_ids = train.iloc[split_valid]["row_id"].to_numpy(int)
        split_record = artifact["downstream_splits"][downstream_fold]
        require(np.array_equal(as_ids(split_record["train_row_ids"]), expected_train_ids), "stored downstream train identity changed")
        require(np.array_equal(as_ids(split_record["valid_row_ids"]), expected_valid_ids), "stored downstream valid identity changed")
        require(np.intersect1d(expected_train_ids, test_ids).size == 0, "downstream train uses outer test")
        require(np.intersect1d(expected_valid_ids, test_ids).size == 0, "downstream valid uses outer test")

        outer_bundle = artifact["projection_bundles"][downstream_fold]
        score_error, metric_error = verify_projection_bundle_v2(
            outer_bundle, frame_by_row, features, expected_train_ids, expected_valid_ids
        )
        max_projection_error = max(max_projection_error, score_error)
        max_metric_error = max(max_metric_error, metric_error)
        seen_outer_training_holdouts.extend(expected_valid_ids.tolist())
        for target in ("age", "mortality"):
            values = np.asarray(outer_bundle["targets"][target]["holdout_score"], dtype=float)
            positions = [train_position[int(row_id)] for row_id in expected_valid_ids]
            reconstructed_training_scores[target][positions] = values

        chain = artifact["downstream_score_chains"][downstream_fold]
        require(int(chain["downstream_fold"]) == downstream_fold, "score chain downstream fold changed")
        expected_projection_seed = int(outer_seed + 2000 + 10 * outer_fold + downstream_fold)
        require(int(chain["seed"]) == expected_projection_seed, "score chain seed changed")
        require(np.array_equal(as_ids(chain["training_row_ids"]), expected_train_ids), "score chain train identity changed")
        require(np.array_equal(as_ids(chain["validation_row_ids"]), expected_valid_ids), "score chain valid identity changed")

        downstream_training = frame_by_row.loc[expected_train_ids].reset_index(drop=True)
        score_splits = core.make_splits(
            downstream_training, core.N_PROJECTION_TUNING_FOLDS, expected_projection_seed
        )
        require(len(chain["training_projection_bundles"]) == core.N_PROJECTION_TUNING_FOLDS, "nested training bundle count changed")
        reconstructed_chain_scores = {
            "age": np.full(len(downstream_training), np.nan, dtype=float),
            "mortality": np.full(len(downstream_training), np.nan, dtype=float),
        }
        chain_position = {int(row_id): index for index, row_id in enumerate(expected_train_ids)}
        seen_chain_holdouts: list[int] = []
        for projection_fold, (projection_train, projection_valid) in enumerate(score_splits):
            projection_train_ids = downstream_training.iloc[projection_train]["row_id"].to_numpy(int)
            projection_valid_ids = downstream_training.iloc[projection_valid]["row_id"].to_numpy(int)
            split_state = chain["training_projection_splits"][projection_fold]
            require(np.array_equal(as_ids(split_state["train_row_ids"]), projection_train_ids), "nested projection train identity changed")
            require(np.array_equal(as_ids(split_state["valid_row_ids"]), projection_valid_ids), "nested projection valid identity changed")
            require(np.intersect1d(projection_train_ids, expected_valid_ids).size == 0, "downstream validation enters nested projection training")
            bundle = chain["training_projection_bundles"][projection_fold]
            score_error, metric_error = verify_projection_bundle_v2(
                bundle, frame_by_row, features, projection_train_ids, projection_valid_ids
            )
            max_projection_error = max(max_projection_error, score_error)
            max_metric_error = max(max_metric_error, metric_error)
            seen_chain_holdouts.extend(projection_valid_ids.tolist())
            for target in ("age", "mortality"):
                values = np.asarray(bundle["targets"][target]["holdout_score"], dtype=float)
                positions = [chain_position[int(row_id)] for row_id in projection_valid_ids]
                reconstructed_chain_scores[target][positions] = values
        require(sorted(seen_chain_holdouts) == sorted(expected_train_ids.tolist()) and len(seen_chain_holdouts) == len(set(seen_chain_holdouts)), "nested projection holdout coverage failed")

        validation_bundle = chain["validation_projection_bundle"]
        score_error, metric_error = verify_projection_bundle_v2(
            validation_bundle, frame_by_row, features, expected_train_ids, expected_valid_ids
        )
        max_projection_error = max(max_projection_error, score_error)
        max_metric_error = max(max_metric_error, metric_error)
        for target in ("age", "mortality"):
            max_chain_error = max(
                max_chain_error,
                maximum_error(reconstructed_chain_scores[target], chain["training_scores"][target]),
                maximum_error(
                    validation_bundle["targets"][target]["holdout_score"], chain["validation_scores"][target]
                ),
            )

    require(sorted(seen_outer_training_holdouts) == sorted(train_ids.tolist()) and len(seen_outer_training_holdouts) == len(set(seen_outer_training_holdouts)), "outer-training projection coverage failed")
    for target in ("age", "mortality"):
        max_chain_error = max(
            max_chain_error,
            maximum_error(reconstructed_training_scores[target], artifact["training_scores"][target]),
        )

    final_projection_seed = int(outer_seed + 2000 + 10 * outer_fold + core.N_DOWNSTREAM_FOLDS)
    require(int(artifact["final_projection_bundle"]["seed"]) == final_projection_seed, "outer-test projection seed changed")
    score_error, metric_error = verify_projection_bundle_v2(
        artifact["final_projection_bundle"], frame_by_row, features, train_ids, test_ids
    )
    max_projection_error = max(max_projection_error, score_error)
    max_metric_error = max(max_metric_error, metric_error)
    for target in ("age", "mortality"):
        max_chain_error = max(
            max_chain_error,
            maximum_error(
                artifact["final_projection_bundle"]["targets"][target]["holdout_score"],
                artifact["test_scores"][target],
            ),
        )

    max_validation_error = 0.0
    max_outer_error = 0.0
    probabilities: dict[str, np.ndarray] = {}
    expected_variants = {f"{target}_{mode}" for target in ("age", "mortality") for mode in ("linear", "spline")}
    require(set(artifact["downstream_variants"]) == expected_variants, "downstream variant set changed")
    chains_by_fold = {int(chain["downstream_fold"]): chain for chain in artifact["downstream_score_chains"]}
    for variant_name, variant in artifact["downstream_variants"].items():
        target = variant["target"]
        mode = variant["mode"]
        require(variant_name == f"{target}_{mode}", "downstream variant metadata changed")
        require(len(variant["tuning_fits"]) == len(core.C_GRID) * core.N_DOWNSTREAM_FOLDS, "downstream tuning fit count changed")
        mean_rows = []
        for c_value in core.C_GRID:
            fits = [fit for fit in variant["tuning_fits"] if float(fit["C"]) == float(c_value)]
            require(len(fits) == core.N_DOWNSTREAM_FOLDS, "downstream C fold count changed")
            losses = []
            seen_valid: list[int] = []
            for fit in sorted(fits, key=lambda item: int(item["downstream_fold"])):
                chain = chains_by_fold[int(fit["downstream_fold"])]
                expected_train_ids = as_ids(chain["training_row_ids"])
                expected_valid_ids = as_ids(chain["validation_row_ids"])
                require(np.array_equal(as_ids(fit["train_row_ids"]), expected_train_ids), "downstream fit train identity breaks score chain")
                require(np.array_equal(as_ids(fit["valid_row_ids"]), expected_valid_ids), "downstream fit valid identity breaks score chain")
                require(np.intersect1d(expected_train_ids, expected_valid_ids).size == 0, "downstream fit train/valid overlap")
                max_chain_error = max(
                    max_chain_error,
                    maximum_error(fit["train_score"], chain["training_scores"][target]),
                    maximum_error(fit["valid_score"], chain["validation_scores"][target]),
                )
                valid = frame_by_row.loc[expected_valid_ids]
                probability = core.predict_downstream_model(fit["model"], valid, fit["valid_score"])
                max_validation_error = max(
                    max_validation_error, maximum_error(probability, fit["valid_probability"])
                )
                reproduced_loss = float(log_loss(valid["death_5y"], probability, labels=[0, 1]))
                max_metric_error = max(max_metric_error, abs(reproduced_loss - float(fit["log_loss"])))
                require(float(fit["model"]["C"]) == float(c_value), "stored downstream tuning C changed")
                require(fit["model"]["mode"] == mode, "stored downstream tuning mode changed")
                losses.append(reproduced_loss)
                seen_valid.extend(expected_valid_ids.tolist())
            require(sorted(seen_valid) == sorted(train_ids.tolist()) and len(seen_valid) == len(set(seen_valid)), "downstream tuning validation coverage failed")
            mean_rows.append({"C": float(c_value), "log_loss": float(np.mean(losses))})
        selected_c = float(pd.DataFrame(mean_rows).sort_values(["log_loss", "C"]).iloc[0]["C"])
        require(selected_c == float(variant["selected_C"]), "selected downstream C does not reconstruct")
        require(float(variant["final_model"]["C"]) == selected_c, "final downstream C differs from selected C")
        require(variant["final_model"]["mode"] == mode, "final downstream mode changed")
        require(np.array_equal(as_ids(variant["final_train_row_ids"]), train_ids), "final downstream train identity changed")
        require(np.array_equal(as_ids(variant["test_row_ids"]), test_ids), "final downstream test identity changed")
        max_chain_error = max(
            max_chain_error,
            maximum_error(variant["final_train_score"], artifact["training_scores"][target]),
            maximum_error(variant["test_score"], artifact["test_scores"][target]),
        )
        probability = core.predict_downstream_model(
            variant["final_model"], test, np.asarray(variant["test_score"], dtype=float)
        )
        max_outer_error = max(max_outer_error, maximum_error(probability, variant["test_probability"]))
        probabilities[variant_name] = np.asarray(variant["test_probability"], dtype=float)

    require(max_projection_error <= core.PARITY_TOLERANCE, "projection parity exceeds tolerance")
    require(max_chain_error <= core.PARITY_TOLERANCE, "end-to-end score-chain parity exceeds tolerance")
    require(max_metric_error <= core.PARITY_TOLERANCE, "tuning metric reconstruction exceeds tolerance")
    require(max_validation_error <= core.PARITY_TOLERANCE, "downstream validation parity exceeds tolerance")
    require(max_outer_error <= core.PARITY_TOLERANCE, "downstream outer parity exceeds tolerance")

    bundles = [*artifact["projection_bundles"], artifact["final_projection_bundle"]]
    for chain in artifact["downstream_score_chains"]:
        bundles.extend(chain["training_projection_bundles"])
        bundles.append(chain["validation_projection_bundle"])
    projection_tuning_count = sum(
        len(target["tuning_fits"]) for bundle in bundles for target in bundle["targets"].values()
    )
    projection_final_count = sum(1 for bundle in bundles for _ in bundle["targets"].values())
    downstream_tuning_count = sum(
        len(variant["tuning_fits"]) for variant in artifact["downstream_variants"].values()
    )
    downstream_final_count = len(artifact["downstream_variants"])
    return {
        "probabilities": probabilities,
        "scores": {key: np.asarray(value, dtype=float) for key, value in artifact["test_scores"].items()},
        "max_projection_score_error": max_projection_error,
        "max_end_to_end_score_error": max_chain_error,
        "max_tuning_metric_error": max_metric_error,
        "max_downstream_validation_error": max_validation_error,
        "max_downstream_outer_error": max_outer_error,
        "projection_tuning_estimator_count": projection_tuning_count,
        "projection_final_estimator_count": projection_final_count,
        "downstream_tuning_estimator_count": downstream_tuning_count,
        "downstream_final_estimator_count": downstream_final_count,
        "nested_identity_gate_pass": True,
    }


def fit_or_load_matched_artifact_v2(
    frame: pd.DataFrame,
    features: list[str],
    assignment: np.ndarray,
    repeat: int,
    outer_seed: int,
    outer_fold: int,
    analysis_signature: str,
) -> tuple[dict[str, np.ndarray], dict[str, np.ndarray], dict[str, Any]]:
    path = core.MATCHED_MODEL_DIR / f"repeat_{repeat:02d}" / f"outer_fold_{outer_fold:02d}.joblib"
    newly_fitted = False
    if path.exists():
        artifact = joblib.load(path)
    else:
        artifact = fit_matched_artifact_v2(
            frame, features, assignment, repeat, outer_seed, outer_fold, analysis_signature
        )
        core.save_joblib_atomic(artifact, path)
        newly_fitted = True
    reloaded = joblib.load(path)
    verified = verify_matched_artifact_v2(
        reloaded, frame, features, assignment, repeat, outer_seed, outer_fold, analysis_signature
    )
    metadata = {
        "stage": "matched_scalar",
        "repeat": int(repeat),
        "outer_seed": int(outer_seed),
        "outer_fold": int(outer_fold),
        "artifact_path": str(path.relative_to(ROOT)),
        "artifact_sha256": core.sha256(path),
        "newly_fitted": newly_fitted,
        "projection_tuning_estimator_count": verified["projection_tuning_estimator_count"],
        "projection_final_estimator_count": verified["projection_final_estimator_count"],
        "downstream_tuning_estimator_count": verified["downstream_tuning_estimator_count"],
        "downstream_final_estimator_count": verified["downstream_final_estimator_count"],
        "max_projection_score_error": verified["max_projection_score_error"],
        "max_end_to_end_score_error": verified["max_end_to_end_score_error"],
        "max_tuning_metric_error": verified["max_tuning_metric_error"],
        "max_downstream_validation_error": verified["max_downstream_validation_error"],
        "max_downstream_outer_error": verified["max_downstream_outer_error"],
        "nested_identity_gate_pass": verified["nested_identity_gate_pass"],
    }
    return verified["probabilities"], verified["scores"], metadata


def build_signature_v2() -> tuple[str, pd.DataFrame, dict[str, Any]]:
    input_paths: list[tuple[str, Path]] = [
        ("analysis_ready_source", core.SOURCE_FRAME),
        ("participant_folds", core.SOURCE_FOLDS),
        ("ordered_feature_tiers", core.SOURCE_TIERS),
        ("source_bootstrap_indices", core.SOURCE_BOOTSTRAP),
        ("predecessor_model_audit", core.SOURCE_PREDECESSOR_AUDIT),
        ("source_oof_predictions", core.SOURCE_OOF),
        ("source_model_audit", core.SOURCE_AUDIT),
        ("source_analysis_signature", core.SOURCE_SIGNATURE),
        ("frozen_v0_2_specification", SPEC),
        ("HUM06_v0_1_core_script", CORE_SCRIPT),
        ("base_age_decoder_script", core.BASE_SCRIPT),
        ("analysis_script", Path(__file__)),
        ("environment_policy", core.ENV_POLICY),
        ("environment_manifest", core.ENV_MANIFEST),
        ("environment_lock", core.ENV_LOCK),
    ]
    for outer_fold in range(core.N_OUTER_FOLDS):
        input_paths.append(
            (f"source_age_model_outer_{outer_fold}", core.SOURCE / "models" / f"outer_fold_{outer_fold:02d}.joblib")
        )
        input_paths.append(
            (
                f"source_age_crossfit_outer_{outer_fold}",
                core.SOURCE / f"outer_fold_{outer_fold:02d}_training_crossfit_scores.tsv.gz",
            )
        )
    for _, path in input_paths:
        require(path.exists(), f"required input is missing: {path}")
    manifest = pd.DataFrame(
        [
            {
                "role": role,
                "path": str(path.relative_to(ROOT)),
                "bytes": path.stat().st_size,
                "sha256": core.sha256(path),
            }
            for role, path in input_paths
        ]
    )
    payload = {
        "analysis_version": ANALYSIS_VERSION,
        "schema_version": SCHEMA_VERSION,
        "nesting_version": NESTING_VERSION,
        "inputs": manifest[["role", "sha256"]].to_dict(orient="records"),
        "population": "NHANES_2007_2012_complete_support_ages_20_79_S3_50",
        "outer_seeds": core.OUTER_SEEDS,
        "outer_folds": core.N_OUTER_FOLDS,
        "downstream_folds": core.N_DOWNSTREAM_FOLDS,
        "projection_tuning_folds": core.N_PROJECTION_TUNING_FOLDS,
        "projection_targets": ["standardized_chronological_age", "standardized_five_year_mortality"],
        "projection_model": "ridge_squared_error",
        "alpha_grid": core.ALPHA_GRID,
        "downstream_models": ["linear_scalar", "five_knot_scalar_spline"],
        "C_grid": core.C_GRID,
        "bootstrap_seed": core.BOOTSTRAP_SEED,
        "bootstraps": core.N_BOOTSTRAPS,
        "parity_tolerance": core.PARITY_TOLERANCE,
    }
    return core.stable_hash(payload), manifest, payload


def id_signature(values: Any) -> str:
    return core.stable_hash(as_ids(values).tolist())


def persist_v2_review_tables() -> None:
    identity_rows = []
    tuning_rows = []
    for path in sorted(core.MATCHED_MODEL_DIR.rglob("outer_fold_*.joblib")):
        artifact = joblib.load(path)
        base = {
            "repeat": int(artifact["repeat"]),
            "outer_seed": int(artifact["outer_seed"]),
            "outer_fold": int(artifact["outer_fold"]),
        }

        def add_identity(stage: str, downstream_fold: int, projection_fold: int, bundle: dict[str, Any]) -> None:
            train_ids = as_ids(bundle["training_row_ids"])
            valid_ids = as_ids(bundle["holdout_row_ids"])
            identity_rows.append(
                {
                    **base,
                    "stage": stage,
                    "downstream_fold": downstream_fold,
                    "projection_fold": projection_fold,
                    "train_n": len(train_ids),
                    "valid_n": len(valid_ids),
                    "train_id_signature": id_signature(train_ids),
                    "valid_id_signature": id_signature(valid_ids),
                    "disjoint": np.intersect1d(train_ids, valid_ids).size == 0,
                }
            )
            for target_name, target in bundle["targets"].items():
                tuning_rows.append(
                    {
                        **base,
                        "stage": stage,
                        "downstream_fold": downstream_fold,
                        "projection_fold": projection_fold,
                        "target": target_name,
                        "mode": "ridge_projection",
                        "selected_parameter": float(target["selected_alpha"]),
                    }
                )

        for downstream_fold, bundle in enumerate(artifact["projection_bundles"]):
            add_identity("outer_training_projection", downstream_fold, downstream_fold, bundle)
        add_identity("outer_test_projection", -1, -1, artifact["final_projection_bundle"])
        for chain in artifact["downstream_score_chains"]:
            downstream_fold = int(chain["downstream_fold"])
            add_identity("downstream_validation_projection", downstream_fold, -1, chain["validation_projection_bundle"])
            for projection_fold, bundle in enumerate(chain["training_projection_bundles"]):
                add_identity("downstream_training_crossfit_projection", downstream_fold, projection_fold, bundle)
        for variant_name, variant in artifact["downstream_variants"].items():
            tuning_rows.append(
                {
                    **base,
                    "stage": "downstream_calibration",
                    "downstream_fold": -1,
                    "projection_fold": -1,
                    "target": variant["target"],
                    "mode": variant["mode"],
                    "selected_parameter": float(variant["selected_C"]),
                }
            )
    pd.DataFrame(identity_rows).to_csv(OUT / "fully_nested_identity_audit.tsv.gz", sep="\t", index=False, compression="gzip")
    pd.DataFrame(tuning_rows).to_csv(OUT / "fully_nested_tuning_summary.tsv.gz", sep="\t", index=False, compression="gzip")


def finalize_v2_audit() -> None:
    artifact_index = pd.read_csv(OUT / "model_artifact_index.tsv", sep="\t")
    matched = artifact_index.loc[artifact_index["stage"].eq("matched_scalar")].copy()
    audit_path = OUT / "model_artifact_audit.json"
    audit = json.loads(audit_path.read_text(encoding="utf-8"))
    audit.update(
        {
            "schema_version": SCHEMA_VERSION,
            "nesting_version": NESTING_VERSION,
            "v0_1_audit_disposition": "retained_provenance_inner_validation_not_fully_nested",
            "max_end_to_end_score_error": float(matched["max_end_to_end_score_error"].max()),
            "max_tuning_metric_error": float(matched["max_tuning_metric_error"].max()),
            "all_nested_identity_gates_pass": bool(matched["nested_identity_gate_pass"].fillna(False).all()),
        }
    )
    if (
        audit["max_end_to_end_score_error"] > core.PARITY_TOLERANCE
        or audit["max_tuning_metric_error"] > core.PARITY_TOLERANCE
        or not audit["all_nested_identity_gates_pass"]
    ):
        audit["status"] = "FAIL"
    core.write_json(audit_path, audit)

    decision_path = OUT / "decision.json"
    decision = json.loads(decision_path.read_text(encoding="utf-8"))
    decision["analysis_version"] = ANALYSIS_VERSION
    decision["nesting_version"] = NESTING_VERSION
    decision["supersedes_for_matched_verdict"] = "nhanes_5y_matched_scalar_target_supervision_probe_v0_1"
    decision["artifact_status"] = audit["status"]
    if audit["status"] != "PASS":
        decision["flexible_calibration"]["decision_category"] = "STOP_DATA_OR_ARTIFACT_FAIL"
        decision["matched_scalar"]["decision_category"] = "STOP_DATA_OR_ARTIFACT_FAIL"
    core.write_json(decision_path, decision)

    core.write_json(
        OUT / "version_correction.json",
        {
            "analysis_version": ANALYSIS_VERSION,
            "retained_prior_version": "nhanes_5y_matched_scalar_target_supervision_probe_v0_1",
            "prior_audit_finding": "outer test clean; downstream C-tuning validation not fully nested",
            "correction": NESTING_VERSION,
            "unchanged": [
                "population",
                "endpoint",
                "ordered_50_measurement_panel",
                "outer_and_downstream_folds",
                "seeds",
                "model_families",
                "alpha_and_C_grids",
                "bootstrap",
                "decision_gates",
            ],
            "HUM04_status": "frozen_read_only",
        },
    )
    persist_v2_review_tables()

    report_path = OUT / "REPORT.md"
    report = report_path.read_text(encoding="utf-8")
    report += f"""

## V0.2 nesting correction

V0.1 is retained as audit provenance because its downstream C-tuning folds did
not fully isolate mortality-supervised training scores from the corresponding
downstream validation outcomes. V0.2 generated each downstream-training score
by projection cross-fitting wholly inside that training set and generated each
validation score from a projection selected and fitted only on the complete
training side.

- Fully nested identity gates: **{audit['all_nested_identity_gates_pass']}**.
- Maximum end-to-end score-chain error: {audit['max_end_to_end_score_error']:.3g}.
- Maximum tuning-metric reconstruction error: {audit['max_tuning_metric_error']:.3g}.
- HUM-04 remained frozen and read-only.
"""
    report_path.write_text(report, encoding="utf-8")
    core.write_artifact_manifest()


core.build_signature = build_signature_v2
core.fit_or_load_matched_artifact = fit_or_load_matched_artifact_v2


def main() -> int:
    result = core.main()
    if result != 0:
        return int(result)
    finalize_v2_audit()
    audit = json.loads((OUT / "model_artifact_audit.json").read_text(encoding="utf-8"))
    decision = json.loads((OUT / "decision.json").read_text(encoding="utf-8"))
    print(json.dumps({"audit": audit, "decision": decision}, indent=2, ensure_ascii=False), flush=True)
    return 0 if audit["status"] == "PASS" else 2


if __name__ == "__main__":
    raise SystemExit(main())
```
