## Supplementary_Software_1 for "Inferential boundaries of age prediction: why prediction does not establish biological age measurement": analyze_nhanes_endpoint_vs_inverse_age_decoder_acquisition_v0_2.py

```
#!/usr/bin/env python
"""Branch A: endpoint-selected versus inverse-age-decoder acquisition.

This analysis implements
NHANES_ENDPOINT_VS_INVERSE_AGE_DECODER_ACQUISITION_SPEC_20260826.md.
It fits source-cycle age decoders only. The endpoint-selected source policy and
target mortality predictions are sealed upstream inputs. Every fitted
preprocessor and ridge estimator is persisted and reload-verified.
"""

from __future__ import annotations

import hashlib
import json
import math
import platform
import runpy
import sys
from pathlib import Path
from typing import Any

import joblib
import numpy as np
import pandas as pd
import sklearn
from sklearn.linear_model import LogisticRegression, Ridge
from sklearn.metrics import mean_absolute_error, mean_squared_error, r2_score
from sklearn.model_selection import StratifiedKFold
from sklearn.preprocessing import OneHotEncoder, StandardScaler


ROOT = Path(__file__).resolve().parents[1]
SCRIPT = Path(__file__).resolve()
ANALYSIS_DIR = ROOT / "analysis"
MANUSCRIPT_DIR = ROOT / "manuscript" / "nature_first"
PREDECESSOR = ROOT / "outputs" / "nhanes_age_prior_guided_risk_acquisition_probe_v0_1"
OUT = ROOT / "outputs" / "nhanes_endpoint_vs_inverse_age_decoder_acquisition_v0_2"
MODEL_DIR = OUT / "models"

PANEL = ROOT / "data" / "processed" / "NHANES_broad_clinical_panel_v0_1.tsv.gz"
FEATURE_MANIFEST = (
    ROOT / "data" / "processed" / "NHANES_broad_clinical_feature_manifest_v0_1.tsv"
)
DOMAIN_SOURCE = ANALYSIS_DIR / "probe_nhanes_age_prior_guided_risk_acquisition_v0_1.py"
SOURCE_FREEZE = PREDECESSOR / "source_recommendation_freeze.json"
SOURCE_SUMMARY = PREDECESSOR / "source_domain_summary.tsv"
SOURCE_OOF = PREDECESSOR / "source_oof_predictions.tsv.gz"
TARGET_OOF = PREDECESSOR / "target_oof_predictions.tsv.gz"
SPEC = ANALYSIS_DIR / "NHANES_ENDPOINT_VS_INVERSE_AGE_DECODER_ACQUISITION_SPEC_20260826.md"
DIRECTION_LOCK = (
    MANUSCRIPT_DIR
    / "next_round_20260826"
    / "P2_ANALYSIS_DIRECTION_LOCK_20260826.md"
)
ENV_POLICY = ROOT / "environment-policy.json"
ENV_MANIFEST = ROOT / "environment.yml"
ENV_LOCK = ROOT / "environment.win-64.lock.txt"

ANALYSIS_VERSION = "nhanes_endpoint_vs_inverse_age_decoder_acquisition_v0_2"
SCHEMA_VERSION = "1.0"
SOURCE_CYCLE = "2007_2008"
TARGET_CYCLE = "2009_2010"
EXPECTED_SOURCE_N = 4599
EXPECTED_SOURCE_EVENTS = 174
EXPECTED_TARGET_N = 4896
EXPECTED_TARGET_EVENTS = 157
BACKGROUND = ("sex", "race_ethnicity")
N_OUTER_FOLDS = 5
N_INNER_FOLDS = 3
OUTER_SEED = 20260826
INNER_SEED_BASE = 20260827
BOOTSTRAP_SEED = 20260802
N_BOOTSTRAP = 5000
ALPHA_GRID = (0.01, 0.1, 1.0, 10.0, 100.0, 1000.0)
PARITY_TOLERANCE = 1e-12
LOG_LOSS_EPSILON = 1e-8


def configure_stdout() -> None:
    if hasattr(sys.stdout, "reconfigure"):
        sys.stdout.reconfigure(encoding="utf-8")


def sha256(path: Path) -> str:
    digest = hashlib.sha256()
    with path.open("rb") as handle:
        for block in iter(lambda: handle.read(1024 * 1024), b""):
            digest.update(block)
    return digest.hexdigest()


def stable_hash(payload: Any) -> str:
    encoded = json.dumps(
        payload,
        sort_keys=True,
        separators=(",", ":"),
        ensure_ascii=False,
    ).encode("utf-8")
    return hashlib.sha256(encoded).hexdigest()


def json_ready(value: Any) -> Any:
    if isinstance(value, dict):
        return {str(key): json_ready(item) for key, item in value.items()}
    if isinstance(value, (list, tuple)):
        return [json_ready(item) for item in value]
    if isinstance(value, Path):
        return str(value)
    if isinstance(value, np.ndarray):
        return value.tolist()
    if isinstance(value, (np.integer,)):
        return int(value)
    if isinstance(value, (np.floating,)):
        return None if not np.isfinite(value) else float(value)
    if isinstance(value, float):
        return None if not math.isfinite(value) else value
    return value


def write_json(path: Path, payload: Any) -> None:
    path.write_text(
        json.dumps(json_ready(payload), indent=2, ensure_ascii=False) + "\n",
        encoding="utf-8",
    )


def relative(path: Path) -> str:
    return str(path.relative_to(ROOT))


def load_domains() -> dict[str, list[str]]:
    namespace = runpy.run_path(str(DOMAIN_SOURCE), run_name="mth01_domain_source")
    raw = namespace.get("DOMAINS")
    if not isinstance(raw, dict) or len(raw) != 5:
        raise RuntimeError("STOP_DATA_OR_ARTIFACT_FAIL: frozen DOMAINS not recovered")
    domains = {str(key): [str(item) for item in value] for key, value in raw.items()}
    expected_counts = [10, 7, 10, 20, 3]
    if list(map(len, domains.values())) != expected_counts:
        raise RuntimeError("STOP_DATA_OR_ARTIFACT_FAIL: frozen domain sizes changed")
    flattened = [feature for features in domains.values() for feature in features]
    if len(flattened) != 50 or len(set(flattened)) != 50:
        raise RuntimeError("STOP_DATA_OR_ARTIFACT_FAIL: domain feature partition changed")
    return domains


def load_cycle(
    panel: pd.DataFrame,
    cycle: str,
    domains: dict[str, list[str]],
) -> pd.DataFrame:
    transformed = [
        f"T__{feature}" for features in domains.values() for feature in features
    ]
    required = [
        "SEQN",
        "cycle",
        "age_years",
        "sex",
        "race_ethnicity",
        "event",
        "time_months",
        *transformed,
    ]
    missing = sorted(set(required) - set(panel.columns))
    if missing:
        raise RuntimeError(
            f"STOP_DATA_OR_ARTIFACT_FAIL: panel missing required columns {missing}"
        )
    died = panel["event"].eq(1) & panel["time_months"].le(60)
    resolved = died | panel["time_months"].ge(60)
    keep = (
        panel["cycle"].eq(cycle)
        & panel["age_years"].between(20, 79)
        & resolved
        & panel[required].notna().all(axis=1)
    )
    frame = (
        panel.loc[keep, required]
        .copy()
        .sort_values("SEQN", kind="mergesort")
        .reset_index(drop=True)
    )
    frame.insert(0, "row_id", np.arange(len(frame), dtype=int))
    frame["death_5y"] = (
        frame["event"].eq(1) & frame["time_months"].le(60)
    ).astype(int)
    if frame["SEQN"].duplicated().any():
        raise RuntimeError("STOP_DATA_OR_ARTIFACT_FAIL: duplicate participant")
    return frame


def make_age_strata(age: np.ndarray, n_splits: int) -> np.ndarray:
    strata = pd.qcut(
        pd.Series(np.asarray(age, dtype=float)),
        q=5,
        labels=False,
        duplicates="drop",
    )
    if strata.isna().any():
        raise RuntimeError("STOP_DATA_OR_ARTIFACT_FAIL: age strata contain missing")
    values = strata.astype(int).to_numpy()
    counts = pd.Series(values).value_counts()
    if len(counts) != 5 or int(counts.min()) < n_splits:
        raise RuntimeError("STOP_DATA_OR_ARTIFACT_FAIL: age strata cannot support folds")
    return values


def assign_outer_folds(frame: pd.DataFrame) -> pd.DataFrame:
    age = frame["age_years"].to_numpy(float)
    strata = make_age_strata(age, N_OUTER_FOLDS)
    splitter = StratifiedKFold(
        n_splits=N_OUTER_FOLDS,
        shuffle=True,
        random_state=OUTER_SEED,
    )
    fold = np.full(len(frame), -1, dtype=int)
    for outer_fold, (_, test_index) in enumerate(
        splitter.split(np.zeros(len(frame)), strata)
    ):
        fold[test_index] = outer_fold
    if sorted(np.unique(fold).tolist()) != list(range(N_OUTER_FOLDS)):
        raise RuntimeError("STOP_DATA_OR_ARTIFACT_FAIL: incomplete source outer folds")
    result = frame.copy()
    result["age_stratum"] = strata
    result["outer_fold"] = fold
    return result


def clean_categories(frame: pd.DataFrame) -> pd.DataFrame:
    return frame.fillna("missing").astype(str).replace("", "missing")


def fit_age_model(
    frame: pd.DataFrame,
    features: list[str],
    alpha: float,
) -> dict[str, Any]:
    encoder = OneHotEncoder(handle_unknown="ignore", sparse_output=False)
    categories = encoder.fit_transform(
        clean_categories(frame.loc[:, BACKGROUND])
    )
    scaler: StandardScaler | None = None
    blocks = [categories]
    names = [
        f"W__{name}" for name in encoder.get_feature_names_out(BACKGROUND)
    ]
    if features:
        scaler = StandardScaler()
        measurements = scaler.fit_transform(frame.loc[:, features].to_numpy(float))
        blocks.append(measurements)
        names.extend(f"M__{name}" for name in features)
    design = np.column_stack(blocks)
    estimator = Ridge(alpha=float(alpha), fit_intercept=True)
    estimator.fit(design, frame["age_years"].to_numpy(float))
    return {
        "encoder": encoder,
        "scaler": scaler,
        "estimator": estimator,
        "feature_names": list(features),
        "design_feature_names": names,
    }


def predict_age_model(model: dict[str, Any], frame: pd.DataFrame) -> np.ndarray:
    categories = model["encoder"].transform(
        clean_categories(frame.loc[:, BACKGROUND])
    )
    blocks = [categories]
    features = list(model["feature_names"])
    if features:
        measurements = model["scaler"].transform(
            frame.loc[:, features].to_numpy(float)
        )
        blocks.append(measurements)
    return model["estimator"].predict(np.column_stack(blocks))


def model_specs(domains: dict[str, list[str]]) -> dict[str, list[str]]:
    return {
        "W_only": [],
        **{
            f"domain__{domain}": [f"T__{feature}" for feature in features]
            for domain, features in domains.items()
        },
    }


def fit_outer_artifact(
    frame: pd.DataFrame,
    domains: dict[str, list[str]],
    outer_fold: int,
    analysis_signature: str,
) -> dict[str, Any]:
    train = (
        frame.loc[frame["outer_fold"].ne(outer_fold)]
        .copy()
        .sort_values("row_id")
        .reset_index(drop=True)
    )
    test = (
        frame.loc[frame["outer_fold"].eq(outer_fold)]
        .copy()
        .sort_values("row_id")
        .reset_index(drop=True)
    )
    inner_strata = make_age_strata(
        train["age_years"].to_numpy(float), N_INNER_FOLDS
    )
    inner_seed = INNER_SEED_BASE + outer_fold
    splitter = StratifiedKFold(
        n_splits=N_INNER_FOLDS,
        shuffle=True,
        random_state=inner_seed,
    )
    inner_splits = [
        (inner_fold, train_index, valid_index)
        for inner_fold, (train_index, valid_index) in enumerate(
            splitter.split(np.zeros(len(train)), inner_strata)
        )
    ]

    fitted_models: dict[str, Any] = {}
    tuning_rows: list[dict[str, Any]] = []
    inner_assignment_rows: list[dict[str, Any]] = []
    for inner_fold, _, valid_index in inner_splits:
        inner_assignment_rows.extend(
            {
                "outer_fold": outer_fold,
                "inner_fold": inner_fold,
                "row_id": int(row_id),
                "SEQN": int(seqn),
            }
            for row_id, seqn in train.iloc[valid_index][["row_id", "SEQN"]]
            .itertuples(index=False, name=None)
        )

    for name, features in model_specs(domains).items():
        candidate_fits: list[dict[str, Any]] = []
        for alpha in ALPHA_GRID:
            for inner_fold, inner_train, inner_valid in inner_splits:
                fit_frame = train.iloc[inner_train].copy()
                valid_frame = train.iloc[inner_valid].copy()
                fitted = fit_age_model(fit_frame, features, alpha)
                prediction = predict_age_model(fitted, valid_frame)
                residual = (
                    valid_frame["age_years"].to_numpy(float) - prediction
                )
                squared_error_sum = float(np.dot(residual, residual))
                row = {
                    "outer_fold": outer_fold,
                    "model": name,
                    "alpha": float(alpha),
                    "inner_fold": inner_fold,
                    "n": int(len(valid_frame)),
                    "squared_error_sum": squared_error_sum,
                    "mse": squared_error_sum / len(valid_frame),
                }
                tuning_rows.append(row)
                candidate_fits.append(
                    {
                        "alpha": float(alpha),
                        "inner_fold": inner_fold,
                        "train_row_ids": fit_frame["row_id"].to_numpy(int),
                        "valid_row_ids": valid_frame["row_id"].to_numpy(int),
                        "model": fitted,
                        "valid_prediction": prediction,
                        "squared_error_sum": squared_error_sum,
                    }
                )
        tuning = pd.DataFrame(
            [row for row in tuning_rows if row["model"] == name]
        )
        pooled = (
            tuning.groupby("alpha", as_index=False)
            .agg(
                squared_error_sum=("squared_error_sum", "sum"),
                n=("n", "sum"),
            )
        )
        pooled["mse"] = pooled["squared_error_sum"] / pooled["n"]
        selected_alpha = float(
            pooled.sort_values(["mse", "alpha"], kind="mergesort")
            .iloc[0]["alpha"]
        )
        outer_model = fit_age_model(train, features, selected_alpha)
        test_prediction = predict_age_model(outer_model, test)
        fitted_models[name] = {
            "features": list(features),
            "selected_alpha": selected_alpha,
            "tuning_fits": candidate_fits,
            "outer_model": outer_model,
            "test_prediction": test_prediction,
        }

    base_signature = stable_hash(
        {
            "analysis_signature": analysis_signature,
            "outer_fold": outer_fold,
            "train_row_ids": train["row_id"].tolist(),
            "test_row_ids": test["row_id"].tolist(),
            "domain_order": list(domains),
            "domains": domains,
            "background": BACKGROUND,
            "alpha_grid": ALPHA_GRID,
            "inner_seed": inner_seed,
        }
    )
    return {
        "schema_version": SCHEMA_VERSION,
        "analysis_signature": analysis_signature,
        "base_signature": base_signature,
        "outer_fold": outer_fold,
        "inner_seed": inner_seed,
        "train_row_ids": train["row_id"].to_numpy(int),
        "test_row_ids": test["row_id"].to_numpy(int),
        "models": fitted_models,
        "tuning_rows": tuning_rows,
        "inner_assignment_rows": inner_assignment_rows,
    }


def verify_outer_artifact(
    artifact: dict[str, Any],
    frame_by_row: pd.DataFrame,
    analysis_signature: str,
) -> tuple[float, dict[str, np.ndarray]]:
    if artifact.get("analysis_signature") != analysis_signature:
        raise RuntimeError(
            "STOP_DATA_OR_ARTIFACT_FAIL: artifact analysis signature mismatch"
        )
    test_ids = np.asarray(artifact["test_row_ids"], dtype=int)
    test = frame_by_row.loc[test_ids].copy()
    max_error = 0.0
    predictions: dict[str, np.ndarray] = {}
    for name, payload in artifact["models"].items():
        for fit in payload["tuning_fits"]:
            valid_ids = np.asarray(fit["valid_row_ids"], dtype=int)
            valid = frame_by_row.loc[valid_ids].copy()
            prediction = predict_age_model(fit["model"], valid)
            error = float(
                np.max(
                    np.abs(
                        prediction
                        - np.asarray(fit["valid_prediction"], dtype=float)
                    )
                )
            )
            max_error = max(max_error, error)
        prediction = predict_age_model(payload["outer_model"], test)
        error = float(
            np.max(
                np.abs(
                    prediction
                    - np.asarray(payload["test_prediction"], dtype=float)
                )
            )
        )
        max_error = max(max_error, error)
        predictions[name] = prediction
    if max_error > PARITY_TOLERANCE:
        raise RuntimeError(
            f"STOP_DATA_OR_ARTIFACT_FAIL: reload parity error {max_error}"
        )
    return max_error, predictions


def fit_or_load_outer_artifact(
    frame: pd.DataFrame,
    domains: dict[str, list[str]],
    outer_fold: int,
    analysis_signature: str,
) -> tuple[dict[str, Any], bool, float, dict[str, np.ndarray]]:
    path = MODEL_DIR / f"source_outer_fold_{outer_fold:02d}.joblib"
    if path.exists():
        artifact = joblib.load(path)
        reused = True
    else:
        artifact = fit_outer_artifact(
            frame, domains, outer_fold, analysis_signature
        )
        path.parent.mkdir(parents=True, exist_ok=True)
        joblib.dump(artifact, path, compress=3)
        reused = False
    reloaded = joblib.load(path)
    frame_by_row = frame.set_index("row_id", drop=False)
    max_error, predictions = verify_outer_artifact(
        reloaded, frame_by_row, analysis_signature
    )
    return reloaded, reused, max_error, predictions


def regression_metrics(age: np.ndarray, prediction: np.ndarray) -> dict[str, float]:
    mse = float(mean_squared_error(age, prediction))
    return {
        "mse": mse,
        "rmse": float(math.sqrt(mse)),
        "mae": float(mean_absolute_error(age, prediction)),
        "r2": float(r2_score(age, prediction)),
    }


def binary_log_loss_vector(y: np.ndarray, p: np.ndarray) -> np.ndarray:
    probability = np.clip(
        np.asarray(p, dtype=float),
        LOG_LOSS_EPSILON,
        1.0 - LOG_LOSS_EPSILON,
    )
    outcome = np.asarray(y, dtype=float)
    return -(
        outcome * np.log(probability)
        + (1.0 - outcome) * np.log(1.0 - probability)
    )


def stratified_bootstrap_indices(
    outcome: np.ndarray,
    n_bootstrap: int,
    seed: int,
) -> np.ndarray:
    rng = np.random.default_rng(seed)
    strata = [
        np.flatnonzero(outcome == value) for value in np.unique(outcome)
    ]
    return np.stack(
        [
            np.concatenate(
                [
                    rng.choice(index, size=len(index), replace=True)
                    for index in strata
                ]
            )
            for _ in range(n_bootstrap)
        ]
    )


def endpoint_categorical_matrix(frame: pd.DataFrame) -> np.ndarray:
    sex = frame["sex"].to_numpy(float)
    race = frame["race_ethnicity"].to_numpy(float)
    if not set(np.unique(sex)).issubset({1.0, 2.0}):
        raise RuntimeError(
            "STOP_DATA_OR_ARTIFACT_FAIL: unexpected endpoint sex category"
        )
    if not set(np.unique(race)).issubset({1.0, 2.0, 3.0, 4.0, 5.0}):
        raise RuntimeError(
            "STOP_DATA_OR_ARTIFACT_FAIL: unexpected endpoint race category"
        )
    return np.column_stack(
        [
            (sex == 2.0).astype(float),
            *[(race == value).astype(float) for value in (2, 3, 4, 5)],
        ]
    )


def endpoint_model_specs(
    domains: dict[str, list[str]],
) -> dict[str, str | None]:
    return {
        "CA": None,
        **{f"CA_M__{domain}": domain for domain in domains},
    }


def fit_endpoint_design(
    frame: pd.DataFrame,
    categories: np.ndarray,
    train_index: np.ndarray,
    domain: str | None,
    domains: dict[str, list[str]],
) -> tuple[np.ndarray, dict[str, Any]]:
    blocks = [categories[train_index]]
    names = [
        "W__female",
        "W__race_2",
        "W__race_3",
        "W__race_4",
        "W__race_5",
    ]
    age = frame["age_years"].to_numpy(float).reshape(-1, 1)
    train_age = age[train_index]
    age_mean = train_age.mean(axis=0)
    age_scale = train_age.std(axis=0, ddof=0)
    age_scale[age_scale < 1e-12] = 1.0
    age_z = (train_age - age_mean) / age_scale
    blocks.append(np.column_stack([age_z, age_z**2, age_z**3]))
    names.extend(["A_z", "A_z2", "A_z3"])

    measurement_mean: np.ndarray | None = None
    measurement_scale: np.ndarray | None = None
    transformed_columns: list[str] = []
    if domain is not None:
        transformed_columns = [
            f"T__{feature}" for feature in domains[domain]
        ]
        values = frame[transformed_columns].to_numpy(float)
        train_values = values[train_index]
        measurement_mean = train_values.mean(axis=0)
        measurement_scale = train_values.std(axis=0, ddof=0)
        measurement_scale[measurement_scale < 1e-12] = 1.0
        blocks.append(
            (train_values - measurement_mean) / measurement_scale
        )
        names.extend(f"M__{column}" for column in transformed_columns)
    return np.column_stack(blocks), {
        "domain": domain,
        "age_mean": age_mean,
        "age_scale": age_scale,
        "transformed_columns": transformed_columns,
        "measurement_mean": measurement_mean,
        "measurement_scale": measurement_scale,
        "design_feature_names": names,
    }


def transform_endpoint_design(
    frame: pd.DataFrame,
    categories: np.ndarray,
    row_index: np.ndarray,
    preprocessor: dict[str, Any],
) -> np.ndarray:
    blocks = [categories[row_index]]
    age = frame["age_years"].to_numpy(float).reshape(-1, 1)[row_index]
    age_z = (
        age - np.asarray(preprocessor["age_mean"], dtype=float)
    ) / np.asarray(preprocessor["age_scale"], dtype=float)
    blocks.append(np.column_stack([age_z, age_z**2, age_z**3]))
    columns = list(preprocessor["transformed_columns"])
    if columns:
        values = frame[columns].to_numpy(float)[row_index]
        blocks.append(
            (
                values
                - np.asarray(preprocessor["measurement_mean"], dtype=float)
            )
            / np.asarray(preprocessor["measurement_scale"], dtype=float)
        )
    return np.column_stack(blocks)


def fit_endpoint_outer_artifact(
    frame: pd.DataFrame,
    domains: dict[str, list[str]],
    population: str,
    outer_fold: int,
    promotion_signature: str,
) -> dict[str, Any]:
    fold = frame["legacy_endpoint_fold"].to_numpy(int)
    train_index = np.flatnonzero(fold != outer_fold)
    test_index = np.flatnonzero(fold == outer_fold)
    categories = endpoint_categorical_matrix(frame)
    outcome = frame["death_5y"].to_numpy(int)
    models: dict[str, Any] = {}
    for name, domain in endpoint_model_specs(domains).items():
        x_train, preprocessor = fit_endpoint_design(
            frame, categories, train_index, domain, domains
        )
        estimator = LogisticRegression(
            C=1.0,
            l1_ratio=0.0,
            solver="lbfgs",
            max_iter=3000,
        )
        estimator.fit(x_train, outcome[train_index])
        x_test = transform_endpoint_design(
            frame, categories, test_index, preprocessor
        )
        prediction = estimator.predict_proba(x_test)[:, 1]
        models[name] = {
            "domain": domain,
            "preprocessor": preprocessor,
            "estimator": estimator,
            "test_prediction": prediction,
        }
    base_signature = stable_hash(
        {
            "promotion_signature": promotion_signature,
            "population": population,
            "outer_fold": outer_fold,
            "train_row_ids": frame.iloc[train_index]["row_id"].tolist(),
            "test_row_ids": frame.iloc[test_index]["row_id"].tolist(),
            "model_specs": endpoint_model_specs(domains),
            "C": 1.0,
            "solver": "lbfgs",
            "max_iter": 3000,
        }
    )
    return {
        "schema_version": SCHEMA_VERSION,
        "promotion_signature": promotion_signature,
        "base_signature": base_signature,
        "population": population,
        "outer_fold": outer_fold,
        "train_row_ids": frame.iloc[train_index]["row_id"].to_numpy(int),
        "test_row_ids": frame.iloc[test_index]["row_id"].to_numpy(int),
        "models": models,
    }


def verify_endpoint_outer_artifact(
    artifact: dict[str, Any],
    frame: pd.DataFrame,
    promotion_signature: str,
) -> tuple[float, dict[str, np.ndarray]]:
    if artifact.get("promotion_signature") != promotion_signature:
        raise RuntimeError(
            "STOP_DATA_OR_ARTIFACT_FAIL: endpoint artifact signature mismatch"
        )
    frame_by_row = frame.set_index("row_id", drop=False)
    test_ids = np.asarray(artifact["test_row_ids"], dtype=int)
    test = frame_by_row.loc[test_ids].copy().reset_index(drop=True)
    categories = endpoint_categorical_matrix(test)
    row_index = np.arange(len(test), dtype=int)
    predictions: dict[str, np.ndarray] = {}
    max_error = 0.0
    for name, payload in artifact["models"].items():
        design = transform_endpoint_design(
            test, categories, row_index, payload["preprocessor"]
        )
        prediction = payload["estimator"].predict_proba(design)[:, 1]
        error = float(
            np.max(
                np.abs(
                    prediction
                    - np.asarray(payload["test_prediction"], dtype=float)
                )
            )
        )
        max_error = max(max_error, error)
        predictions[name] = prediction
    if max_error > PARITY_TOLERANCE:
        raise RuntimeError(
            "STOP_DATA_OR_ARTIFACT_FAIL: endpoint reload parity failed "
            f"at {max_error}"
        )
    return max_error, predictions


def reconstruct_endpoint_population(
    frame: pd.DataFrame,
    sealed_oof: pd.DataFrame,
    domains: dict[str, list[str]],
    population: str,
    promotion_signature: str,
) -> dict[str, Any]:
    required = {
        "SEQN",
        "outcome",
        "fold",
        *endpoint_model_specs(domains).keys(),
    }
    missing = sorted(required - set(sealed_oof.columns))
    if missing:
        raise RuntimeError(
            f"STOP_DATA_OR_ARTIFACT_FAIL: {population} sealed OOF missing {missing}"
        )
    analysis = frame.merge(
        sealed_oof[sorted(required)],
        on="SEQN",
        how="inner",
        validate="one_to_one",
        sort=False,
    ).sort_values("row_id").reset_index(drop=True)
    if len(analysis) != len(frame):
        raise RuntimeError(
            f"STOP_DATA_OR_ARTIFACT_FAIL: {population} endpoint identity changed"
        )
    if not np.array_equal(
        analysis["death_5y"].to_numpy(int),
        analysis["outcome"].to_numpy(int),
    ):
        raise RuntimeError(
            f"STOP_DATA_OR_ARTIFACT_FAIL: {population} endpoint outcome changed"
        )
    if sorted(analysis["fold"].unique().tolist()) != list(range(N_OUTER_FOLDS)):
        raise RuntimeError(
            f"STOP_DATA_OR_ARTIFACT_FAIL: {population} endpoint folds changed"
        )
    analysis["legacy_endpoint_fold"] = analysis["fold"].astype(int)
    analysis[
        ["row_id", "SEQN", "death_5y", "legacy_endpoint_fold"]
    ].to_csv(
        OUT / f"legacy_{population}_endpoint_folds.tsv",
        sep="\t",
        index=False,
    )

    reconstructed = analysis[
        ["row_id", "SEQN", "death_5y", "legacy_endpoint_fold"]
    ].copy()
    for name in endpoint_model_specs(domains):
        reconstructed[name] = np.nan
    index_rows: list[dict[str, Any]] = []
    coefficient_rows: list[dict[str, Any]] = []
    max_reload_error = 0.0
    newly_fitted = 0
    reused = 0

    for outer_fold in range(N_OUTER_FOLDS):
        path = (
            MODEL_DIR
            / f"endpoint_{population}_outer_fold_{outer_fold:02d}.joblib"
        )
        if path.exists():
            artifact = joblib.load(path)
            reused += 1
        else:
            artifact = fit_endpoint_outer_artifact(
                analysis,
                domains,
                population,
                outer_fold,
                promotion_signature,
            )
            joblib.dump(artifact, path, compress=3)
            newly_fitted += 1
        reloaded = joblib.load(path)
        reload_error, predictions = verify_endpoint_outer_artifact(
            reloaded, analysis, promotion_signature
        )
        max_reload_error = max(max_reload_error, reload_error)
        test_ids = np.asarray(reloaded["test_row_ids"], dtype=int)
        expected_ids = (
            analysis.loc[
                analysis["legacy_endpoint_fold"].eq(outer_fold), "row_id"
            ]
            .sort_values()
            .to_numpy(int)
        )
        if not np.array_equal(test_ids, expected_ids):
            raise RuntimeError(
                f"STOP_DATA_OR_ARTIFACT_FAIL: {population} endpoint test IDs changed"
            )
        for name, prediction in predictions.items():
            reconstructed.loc[
                reconstructed["row_id"].isin(test_ids), name
            ] = prediction
            payload = reloaded["models"][name]
            estimator = payload["estimator"]
            coefficient_rows.append(
                {
                    "population": population,
                    "outer_fold": outer_fold,
                    "model": name,
                    "term": "(intercept)",
                    "coefficient": float(estimator.intercept_[0]),
                }
            )
            coefficient_rows.extend(
                {
                    "population": population,
                    "outer_fold": outer_fold,
                    "model": name,
                    "term": term,
                    "coefficient": float(coefficient),
                }
                for term, coefficient in zip(
                    payload["preprocessor"]["design_feature_names"],
                    estimator.coef_[0],
                )
            )
        index_rows.append(
            {
                "population": population,
                "outer_fold": outer_fold,
                "n_train": len(reloaded["train_row_ids"]),
                "n_test": len(reloaded["test_row_ids"]),
                "artifact_path": relative(path),
                "artifact_sha256": sha256(path),
                "reload_max_abs_prediction_error": reload_error,
            }
        )

    model_columns = list(endpoint_model_specs(domains))
    if reconstructed[model_columns].isna().any().any():
        raise RuntimeError(
            f"STOP_DATA_OR_ARTIFACT_FAIL: incomplete {population} endpoint OOF"
        )
    reconstructed.to_csv(
        OUT / f"reconstructed_{population}_endpoint_oof_predictions.tsv.gz",
        sep="\t",
        index=False,
        compression="gzip",
    )
    sealed_aligned = analysis.set_index("row_id")
    parity_rows = []
    max_upstream_error = 0.0
    for name in model_columns:
        upstream = sealed_aligned[name].to_numpy(float)
        regenerated = reconstructed[name].to_numpy(float)
        error = float(np.max(np.abs(regenerated - upstream)))
        max_upstream_error = max(max_upstream_error, error)
        parity_rows.append(
            {
                "population": population,
                "model": name,
                "n": len(regenerated),
                "max_abs_upstream_prediction_error": error,
                "exact_elementwise_equal": bool(
                    np.array_equal(regenerated, upstream)
                ),
                "within_tolerance_1e_12": bool(
                    error <= PARITY_TOLERANCE
                ),
            }
        )
    if max_upstream_error > PARITY_TOLERANCE:
        raise RuntimeError(
            f"STOP_DATA_OR_ARTIFACT_FAIL: {population} upstream OOF parity "
            f"{max_upstream_error} exceeds {PARITY_TOLERANCE}"
        )
    return {
        "index_rows": index_rows,
        "coefficient_rows": coefficient_rows,
        "parity_rows": parity_rows,
        "max_reload_error": max_reload_error,
        "max_upstream_error": max_upstream_error,
        "newly_fitted": newly_fitted,
        "reused": reused,
        "estimator_count": N_OUTER_FOLDS * len(model_columns),
    }


def input_manifest(paths: dict[str, Path]) -> pd.DataFrame:
    rows = []
    for role, path in paths.items():
        if not path.is_file():
            raise RuntimeError(
                f"STOP_DATA_OR_ARTIFACT_FAIL: missing input {role}: {path}"
            )
        rows.append(
            {
                "role": role,
                "path": relative(path),
                "bytes": int(path.stat().st_size),
                "sha256": sha256(path),
            }
        )
    return pd.DataFrame(rows)


def write_artifact_manifest() -> None:
    rows = []
    for path in sorted(OUT.rglob("*")):
        if path.is_file() and path.name != "artifact_manifest.tsv":
            rows.append(
                {
                    "path": relative(path),
                    "bytes": int(path.stat().st_size),
                    "sha256": sha256(path),
                }
            )
    pd.DataFrame(rows).to_csv(
        OUT / "artifact_manifest.tsv", sep="\t", index=False
    )


def main() -> int:
    configure_stdout()
    domains = load_domains()
    input_paths = {
        "analysis_ready_panel": PANEL,
        "feature_manifest": FEATURE_MANIFEST,
        "frozen_domain_source": DOMAIN_SOURCE,
        "source_endpoint_selection_freeze": SOURCE_FREEZE,
        "source_endpoint_summary": SOURCE_SUMMARY,
        "sealed_source_oof_predictions": SOURCE_OOF,
        "sealed_target_oof_predictions": TARGET_OOF,
        "frozen_specification": SPEC,
        "p2_direction_lock": DIRECTION_LOCK,
        "analysis_script": SCRIPT,
        "environment_policy": ENV_POLICY,
        "environment_manifest": ENV_MANIFEST,
        "environment_lock": ENV_LOCK,
    }
    inputs = input_manifest(input_paths)
    input_hashes = dict(zip(inputs["role"], inputs["sha256"]))
    signature_payload = {
        "analysis_version": ANALYSIS_VERSION,
        "schema_version": SCHEMA_VERSION,
        "inputs": inputs[["role", "sha256"]].to_dict(orient="records"),
        "source_cycle": SOURCE_CYCLE,
        "target_cycle": TARGET_CYCLE,
        "expected_source_n": EXPECTED_SOURCE_N,
        "expected_target_n": EXPECTED_TARGET_N,
        "background": BACKGROUND,
        "domains": domains,
        "outer_folds": N_OUTER_FOLDS,
        "inner_folds": N_INNER_FOLDS,
        "outer_seed": OUTER_SEED,
        "inner_seed_rule": "20260827 + outer_fold",
        "alpha_grid": ALPHA_GRID,
        "source_selection_loss": "pooled_participant_oof_age_mse",
        "bootstrap_seed": BOOTSTRAP_SEED,
        "bootstrap_draws": N_BOOTSTRAP,
        "parity_tolerance": PARITY_TOLERANCE,
        "target_endpoint_predictions": "sealed_upstream",
        "evidence_status": "probe",
    }
    candidate_analysis_signature = stable_hash(signature_payload)

    OUT.mkdir(parents=True, exist_ok=True)
    MODEL_DIR.mkdir(parents=True, exist_ok=True)
    signature_path = OUT / "analysis_signature.json"
    if signature_path.exists():
        existing = json.loads(signature_path.read_text(encoding="utf-8"))
        existing_payload = existing.get("signature_payload", {})
        locked_fields = [
            "analysis_version",
            "schema_version",
            "source_cycle",
            "target_cycle",
            "background",
            "domains",
            "outer_folds",
            "inner_folds",
            "outer_seed",
            "inner_seed_rule",
            "alpha_grid",
            "source_selection_loss",
            "bootstrap_seed",
            "bootstrap_draws",
            "parity_tolerance",
        ]
        if any(
            existing_payload.get(field)
            != json_ready(signature_payload.get(field))
            for field in locked_fields
        ):
            raise RuntimeError(
                "STOP_DATA_OR_ARTIFACT_FAIL: locked source analysis contract changed"
            )
        analysis_signature = str(existing["analysis_signature"])
    else:
        analysis_signature = candidate_analysis_signature
        write_json(
            signature_path,
            {
                "analysis_signature": analysis_signature,
                "signature_payload": signature_payload,
            },
        )
    promotion_signature = stable_hash(
        {
            "promotion_kind": "legacy_endpoint_fitted_object_reconstruction",
            "source_analysis_signature": analysis_signature,
            "current_extension_payload": signature_payload,
            "endpoint_models": endpoint_model_specs(domains),
            "endpoint_C": 1.0,
            "endpoint_solver": "lbfgs",
            "endpoint_max_iter": 3000,
            "upstream_oof_parity_tolerance": PARITY_TOLERANCE,
        }
    )
    write_json(
        OUT / "promotion_signature.json",
        {
            "promotion_signature": promotion_signature,
            "source_analysis_signature": analysis_signature,
            "candidate_current_script_signature": candidate_analysis_signature,
            "contract": {
                "source_and_target_models": list(
                    endpoint_model_specs(domains)
                ),
                "folds": "sealed legacy source and target folds",
                "C": 1.0,
                "solver": "lbfgs",
                "max_iter": 3000,
                "upstream_oof_parity_tolerance": PARITY_TOLERANCE,
            },
        },
    )
    inputs.to_csv(OUT / "input_manifest.tsv", sep="\t", index=False)

    feature_manifest = pd.read_csv(FEATURE_MANIFEST, sep="\t")
    manifest_features = set(feature_manifest["feature"].astype(str))
    domain_features = {
        feature for features in domains.values() for feature in features
    }
    if domain_features != manifest_features:
        raise RuntimeError(
            "STOP_DATA_OR_ARTIFACT_FAIL: domain and feature manifests disagree"
        )
    domain_rows = []
    for domain_order, (domain, features) in enumerate(domains.items(), start=1):
        for feature_order, feature in enumerate(features, start=1):
            row = feature_manifest.loc[
                feature_manifest["feature"].eq(feature)
            ].iloc[0]
            domain_rows.append(
                {
                    "domain_order": domain_order,
                    "domain": domain,
                    "feature_order": feature_order,
                    "feature": feature,
                    "transformed_column": f"T__{feature}",
                    "source_module": row["source_module"],
                    "source_column": row["source_column"],
                    "transform": row["transform"],
                }
            )
    pd.DataFrame(domain_rows).to_csv(
        OUT / "ordered_domain_features.tsv", sep="\t", index=False
    )

    source_freeze = json.loads(SOURCE_FREEZE.read_text(encoding="utf-8"))
    if source_freeze.get("source_summary_sha256") != sha256(SOURCE_SUMMARY):
        raise RuntimeError(
            "STOP_DATA_OR_ARTIFACT_FAIL: source endpoint summary hash mismatch"
        )
    endpoint_domain = str(
        source_freeze.get("source_recommended_measurement_domain")
    )
    if endpoint_domain not in domains:
        raise RuntimeError(
            "STOP_DATA_OR_ARTIFACT_FAIL: frozen endpoint domain is invalid"
        )
    if source_freeze.get("target_outcomes_used_in_source_recommendation") is not False:
        raise RuntimeError(
            "STOP_DATA_OR_ARTIFACT_FAIL: endpoint source selection provenance changed"
        )

    panel = pd.read_csv(PANEL, sep="\t")
    source = load_cycle(panel, SOURCE_CYCLE, domains)
    if (
        len(source) != EXPECTED_SOURCE_N
        or int(source["death_5y"].sum()) != EXPECTED_SOURCE_EVENTS
    ):
        raise RuntimeError(
            "STOP_DATA_OR_ARTIFACT_FAIL: source population changed"
        )
    source = assign_outer_folds(source)
    source.to_csv(
        OUT / "analysis_ready_source.tsv.gz",
        sep="\t",
        index=False,
        compression="gzip",
    )
    source[
        ["row_id", "SEQN", "age_years", "age_stratum", "outer_fold"]
    ].to_csv(OUT / "source_participant_folds.tsv", sep="\t", index=False)

    specs = model_specs(domains)
    oof = source[
        ["row_id", "SEQN", "age_years", "age_stratum", "outer_fold"]
    ].copy()
    for name in specs:
        oof[f"predicted_age__{name}"] = np.nan
    model_index_rows = []
    tuning_rows = []
    inner_assignment_rows = []
    coefficient_rows = []
    max_parity_error = 0.0
    fitted_count = 0
    reused_count = 0
    tuning_estimator_count = 0
    outer_estimator_count = 0

    for outer_fold in range(N_OUTER_FOLDS):
        print(
            f"Source inverse-age decoder outer fold {outer_fold + 1}/{N_OUTER_FOLDS}",
            flush=True,
        )
        artifact, reused, parity_error, fold_predictions = (
            fit_or_load_outer_artifact(
                source, domains, outer_fold, analysis_signature
            )
        )
        max_parity_error = max(max_parity_error, parity_error)
        reused_count += int(reused)
        fitted_count += int(not reused)
        test_ids = np.asarray(artifact["test_row_ids"], dtype=int)
        expected_ids = (
            source.loc[source["outer_fold"].eq(outer_fold), "row_id"]
            .sort_values()
            .to_numpy(int)
        )
        if not np.array_equal(test_ids, expected_ids):
            raise RuntimeError(
                "STOP_DATA_OR_ARTIFACT_FAIL: artifact test identity mismatch"
            )
        for name, prediction in fold_predictions.items():
            oof.loc[
                oof["row_id"].isin(test_ids), f"predicted_age__{name}"
            ] = prediction
        tuning_rows.extend(artifact["tuning_rows"])
        inner_assignment_rows.extend(artifact["inner_assignment_rows"])
        tuning_estimator_count += sum(
            len(payload["tuning_fits"])
            for payload in artifact["models"].values()
        )
        outer_estimator_count += len(artifact["models"])

        artifact_path = MODEL_DIR / f"source_outer_fold_{outer_fold:02d}.joblib"
        index_row = {
            "outer_fold": outer_fold,
            "n_train": len(artifact["train_row_ids"]),
            "n_test": len(artifact["test_row_ids"]),
            "inner_seed": artifact["inner_seed"],
            "artifact_path": relative(artifact_path),
            "artifact_sha256": sha256(artifact_path),
            "reload_max_abs_prediction_error": parity_error,
        }
        for name, payload in artifact["models"].items():
            index_row[f"selected_alpha__{name}"] = payload["selected_alpha"]
            model = payload["outer_model"]
            estimator = model["estimator"]
            coefficient_rows.append(
                {
                    "outer_fold": outer_fold,
                    "model": name,
                    "selected_alpha": payload["selected_alpha"],
                    "term": "(intercept)",
                    "coefficient": float(estimator.intercept_),
                }
            )
            coefficient_rows.extend(
                {
                    "outer_fold": outer_fold,
                    "model": name,
                    "selected_alpha": payload["selected_alpha"],
                    "term": term,
                    "coefficient": float(coefficient),
                }
                for term, coefficient in zip(
                    model["design_feature_names"], estimator.coef_
                )
            )
        model_index_rows.append(index_row)

    prediction_columns = [f"predicted_age__{name}" for name in specs]
    if oof[prediction_columns].isna().any().any():
        raise RuntimeError(
            "STOP_DATA_OR_ARTIFACT_FAIL: incomplete source age OOF predictions"
        )
    pd.DataFrame(model_index_rows).to_csv(
        OUT / "model_artifact_index.tsv", sep="\t", index=False
    )
    pd.DataFrame(tuning_rows).to_csv(
        OUT / "source_age_decoder_tuning.tsv.gz",
        sep="\t",
        index=False,
        compression="gzip",
    )
    pd.DataFrame(inner_assignment_rows).to_csv(
        OUT / "source_inner_fold_assignments.tsv.gz",
        sep="\t",
        index=False,
        compression="gzip",
    )
    pd.DataFrame(coefficient_rows).to_csv(
        OUT / "source_outer_model_coefficients.tsv.gz",
        sep="\t",
        index=False,
        compression="gzip",
    )
    oof.to_csv(
        OUT / "source_age_decoder_oof_predictions.tsv.gz",
        sep="\t",
        index=False,
        compression="gzip",
    )

    age = oof["age_years"].to_numpy(float)
    loss_vectors = oof[
        ["row_id", "SEQN", "age_years", "outer_fold"]
    ].copy()
    pooled_rows = []
    for model_order, name in enumerate(specs):
        prediction = oof[f"predicted_age__{name}"].to_numpy(float)
        loss_vectors[f"squared_error__{name}"] = (age - prediction) ** 2
        pooled_rows.append(
            {
                "model_order": model_order,
                "model": name,
                "domain": (
                    "W_only"
                    if name == "W_only"
                    else name.replace("domain__", "", 1)
                ),
                **regression_metrics(age, prediction),
            }
        )
    loss_vectors.to_csv(
        OUT / "source_age_decoder_participant_losses.tsv.gz",
        sep="\t",
        index=False,
        compression="gzip",
    )
    pooled = pd.DataFrame(pooled_rows)
    baseline_mse = float(
        pooled.loc[pooled["model"].eq("W_only"), "mse"].iloc[0]
    )
    pooled["delta_mse_vs_W_only"] = baseline_mse - pooled["mse"]
    domain_pooled = pooled.loc[pooled["model"].ne("W_only")].copy()
    domain_order_map = {domain: index for index, domain in enumerate(domains)}
    domain_pooled["domain_order"] = domain_pooled["domain"].map(domain_order_map)
    domain_pooled = domain_pooled.sort_values(
        ["mse", "domain_order"], kind="mergesort"
    ).reset_index(drop=True)
    domain_pooled["source_age_decoder_rank"] = (
        np.arange(len(domain_pooled), dtype=int) + 1
    )
    inverse_domain = str(domain_pooled.iloc[0]["domain"])

    fold_rows = []
    for outer_fold in range(N_OUTER_FOLDS):
        mask = oof["outer_fold"].eq(outer_fold).to_numpy()
        fold_age = age[mask]
        for model_order, name in enumerate(specs):
            fold_prediction = oof.loc[
                mask, f"predicted_age__{name}"
            ].to_numpy(float)
            fold_rows.append(
                {
                    "outer_fold": outer_fold,
                    "model_order": model_order,
                    "model": name,
                    "domain": (
                        "W_only"
                        if name == "W_only"
                        else name.replace("domain__", "", 1)
                    ),
                    "n": int(mask.sum()),
                    **regression_metrics(fold_age, fold_prediction),
                }
            )
    fold_metrics = pd.DataFrame(fold_rows)
    fold_metrics["delta_mse_vs_W_only"] = np.nan
    fold_metrics["domain_rank"] = np.nan
    for outer_fold in range(N_OUTER_FOLDS):
        base = float(
            fold_metrics.loc[
                fold_metrics["outer_fold"].eq(outer_fold)
                & fold_metrics["model"].eq("W_only"),
                "mse",
            ].iloc[0]
        )
        fold_mask = fold_metrics["outer_fold"].eq(outer_fold)
        fold_metrics.loc[fold_mask, "delta_mse_vs_W_only"] = (
            base - fold_metrics.loc[fold_mask, "mse"]
        )
        domain_mask = fold_mask & fold_metrics["model"].ne("W_only")
        ranked = (
            fold_metrics.loc[domain_mask]
            .assign(
                domain_order=lambda x: x["domain"].map(domain_order_map)
            )
            .sort_values(["mse", "domain_order"], kind="mergesort")
        )
        fold_metrics.loc[ranked.index, "domain_rank"] = np.arange(
            1, len(ranked) + 1
        )
    pooled.to_csv(
        OUT / "source_all_age_decoder_metrics.tsv", sep="\t", index=False
    )
    domain_pooled.to_csv(
        OUT / "source_domain_age_decoder_metrics.tsv", sep="\t", index=False
    )
    fold_metrics.to_csv(
        OUT / "source_fold_age_decoder_metrics.tsv", sep="\t", index=False
    )

    selection_metrics_path = OUT / "source_domain_age_decoder_metrics.tsv"
    source_selection = {
        "analysis_version": ANALYSIS_VERSION,
        "analysis_signature": analysis_signature,
        "source_cycle": SOURCE_CYCLE,
        "selection_target": "chronological_age",
        "selection_rule": "argmin_source_pooled_participant_oof_age_mse",
        "source_inverse_age_decoder_domain": inverse_domain,
        "source_endpoint_selected_domain": endpoint_domain,
        "domain_order": list(domains),
        "selected_domain_mse": float(domain_pooled.iloc[0]["mse"]),
        "selected_domain_rmse": float(domain_pooled.iloc[0]["rmse"]),
        "selected_domain_delta_mse_vs_W_only": float(
            domain_pooled.iloc[0]["delta_mse_vs_W_only"]
        ),
        "source_selection_metrics_file": selection_metrics_path.name,
        "source_selection_metrics_sha256": sha256(selection_metrics_path),
        "outer_fold_rule": "five age-quintile-stratified folds",
        "outer_seed": OUTER_SEED,
        "target_outcomes_used_in_inverse_age_selection": False,
        "target_oof_loaded_after_source_freeze": True,
        "historically_unseen_target": False,
        "evidence_status": "probe",
    }
    write_json(OUT / "source_selection_freeze.json", source_selection)

    # Target outcome data and sealed target OOF predictions are loaded only
    # after the source age-policy selection has been frozen above.
    target = load_cycle(panel, TARGET_CYCLE, domains)
    if (
        len(target) != EXPECTED_TARGET_N
        or int(target["death_5y"].sum()) != EXPECTED_TARGET_EVENTS
    ):
        raise RuntimeError(
            "STOP_DATA_OR_ARTIFACT_FAIL: target population changed"
        )
    target.to_csv(
        OUT / "analysis_ready_target.tsv.gz",
        sep="\t",
        index=False,
        compression="gzip",
    )
    sealed_oof = pd.read_csv(TARGET_OOF, sep="\t")
    required_target_columns = {
        "SEQN",
        "outcome",
        "fold",
        "CA",
        f"CA_M__{endpoint_domain}",
        f"CA_M__{inverse_domain}",
    }
    missing_target = sorted(required_target_columns - set(sealed_oof.columns))
    if missing_target:
        raise RuntimeError(
            f"STOP_DATA_OR_ARTIFACT_FAIL: target OOF missing {missing_target}"
        )
    target_eval = target[
        ["row_id", "SEQN", "age_years", "death_5y"]
    ].merge(
        sealed_oof,
        on="SEQN",
        how="inner",
        validate="one_to_one",
        sort=False,
    )
    target_eval = target_eval.sort_values("row_id").reset_index(drop=True)
    if (
        len(target_eval) != EXPECTED_TARGET_N
        or not np.array_equal(
            target_eval["death_5y"].to_numpy(int),
            target_eval["outcome"].to_numpy(int),
        )
        or sorted(target_eval["fold"].unique().tolist())
        != list(range(N_OUTER_FOLDS))
    ):
        raise RuntimeError(
            "STOP_DATA_OR_ARTIFACT_FAIL: sealed target OOF identity changed"
        )

    outcome = target_eval["outcome"].to_numpy(int)
    losses = {
        "baseline_CA": binary_log_loss_vector(outcome, target_eval["CA"]),
        "source_endpoint_policy": binary_log_loss_vector(
            outcome, target_eval[f"CA_M__{endpoint_domain}"]
        ),
        "source_inverse_age_decoder_policy": binary_log_loss_vector(
            outcome, target_eval[f"CA_M__{inverse_domain}"]
        ),
    }
    target_policy = target_eval[
        ["row_id", "SEQN", "outcome", "fold", "CA"]
    ].copy()
    target_policy["endpoint_domain"] = endpoint_domain
    target_policy["inverse_age_decoder_domain"] = inverse_domain
    target_policy["probability_endpoint_policy"] = target_eval[
        f"CA_M__{endpoint_domain}"
    ]
    target_policy["probability_inverse_age_decoder_policy"] = target_eval[
        f"CA_M__{inverse_domain}"
    ]
    for name, values in losses.items():
        target_policy[f"log_loss__{name}"] = values
    target_policy["gain__source_endpoint_policy"] = (
        losses["baseline_CA"] - losses["source_endpoint_policy"]
    )
    target_policy["gain__source_inverse_age_decoder_policy"] = (
        losses["baseline_CA"] - losses["source_inverse_age_decoder_policy"]
    )
    target_policy["contrast__endpoint_minus_inverse"] = (
        losses["source_inverse_age_decoder_policy"]
        - losses["source_endpoint_policy"]
    )
    target_policy.to_csv(
        OUT / "target_oof_policy_predictions_and_losses.tsv.gz",
        sep="\t",
        index=False,
        compression="gzip",
    )

    policy_rows = []
    baseline_loss = float(losses["baseline_CA"].mean())
    for policy, domain, loss in [
        ("source_endpoint_policy", endpoint_domain, losses["source_endpoint_policy"]),
        (
            "source_inverse_age_decoder_policy",
            inverse_domain,
            losses["source_inverse_age_decoder_policy"],
        ),
    ]:
        policy_rows.append(
            {
                "policy": policy,
                "selected_domain": domain,
                "log_loss": float(loss.mean()),
                "gain_vs_age_and_background": baseline_loss - float(loss.mean()),
            }
        )
    policy_summary = pd.DataFrame(policy_rows)
    policy_summary.to_csv(
        OUT / "target_policy_comparison.tsv", sep="\t", index=False
    )

    target_fold_rows = []
    for outer_fold in range(N_OUTER_FOLDS):
        mask = target_eval["fold"].eq(outer_fold).to_numpy()
        base = float(losses["baseline_CA"][mask].mean())
        endpoint_loss = float(
            losses["source_endpoint_policy"][mask].mean()
        )
        inverse_loss = float(
            losses["source_inverse_age_decoder_policy"][mask].mean()
        )
        target_fold_rows.append(
            {
                "outer_fold": outer_fold,
                "n": int(mask.sum()),
                "events": int(outcome[mask].sum()),
                "endpoint_domain": endpoint_domain,
                "inverse_age_decoder_domain": inverse_domain,
                "gain_endpoint": base - endpoint_loss,
                "gain_inverse_age_decoder": base - inverse_loss,
                "endpoint_minus_inverse": inverse_loss - endpoint_loss,
            }
        )
    pd.DataFrame(target_fold_rows).to_csv(
        OUT / "target_fold_policy_metrics.tsv", sep="\t", index=False
    )

    bootstrap = stratified_bootstrap_indices(
        outcome, N_BOOTSTRAP, BOOTSTRAP_SEED
    )
    np.savez_compressed(OUT / "bootstrap_indices.npz", indices=bootstrap)
    policy_interval_rows = []
    for policy, participant_gain in [
        (
            "source_endpoint_policy",
            losses["baseline_CA"] - losses["source_endpoint_policy"],
        ),
        (
            "source_inverse_age_decoder_policy",
            losses["baseline_CA"]
            - losses["source_inverse_age_decoder_policy"],
        ),
    ]:
        replicates = participant_gain[bootstrap].mean(axis=1)
        q025, q975 = np.quantile(replicates, [0.025, 0.975])
        policy_interval_rows.append(
            {
                "policy": policy,
                "estimate": float(participant_gain.mean()),
                "ci_low": float(q025),
                "ci_high": float(q975),
                "bootstrap_probability_positive": float(
                    np.mean(replicates > 0)
                ),
                "n_bootstrap": N_BOOTSTRAP,
            }
        )
    pd.DataFrame(policy_interval_rows).to_csv(
        OUT / "target_policy_bootstrap.tsv", sep="\t", index=False
    )

    participant_difference = (
        losses["source_inverse_age_decoder_policy"]
        - losses["source_endpoint_policy"]
    )
    contrast_replicates = participant_difference[bootstrap].mean(axis=1)
    contrast_estimate = float(participant_difference.mean())
    contrast_low, contrast_high = np.quantile(
        contrast_replicates, [0.025, 0.975]
    )
    contrast = pd.DataFrame(
        [
            {
                "contrast": "source_endpoint_policy_minus_source_inverse_age_decoder_policy",
                "estimate": contrast_estimate,
                "ci_low": float(contrast_low),
                "ci_high": float(contrast_high),
                "bootstrap_probability_positive": float(
                    np.mean(contrast_replicates > 0)
                ),
                "n_bootstrap": N_BOOTSTRAP,
            }
        ]
    )
    contrast.to_csv(
        OUT / "target_policy_pairwise_bootstrap.tsv",
        sep="\t",
        index=False,
    )

    if endpoint_domain == inverse_domain or contrast_estimate <= 0:
        classification = "NO_ENDPOINT_ADVANTAGE"
    elif float(contrast_low) > 0:
        classification = "ENDPOINT_OUTPERFORMS_INVERSE_DECODER"
    else:
        classification = "DIRECTIONAL_ONLY"

    model_index = pd.DataFrame(model_index_rows)
    source_model_gate_pass = bool(
        len(model_index) == N_OUTER_FOLDS
        and max_parity_error <= PARITY_TOLERANCE
        and model_index["artifact_sha256"].str.len().eq(64).all()
    )
    if not source_model_gate_pass:
        raise RuntimeError(
            "STOP_DATA_OR_ARTIFACT_FAIL: source age-model persistence audit failed"
        )

    promotion_succeeded = False
    promotion_failure_reason: str | None = None
    endpoint_audit: dict[str, Any]
    try:
        source_endpoint_reconstruction = reconstruct_endpoint_population(
            source.drop(columns=["age_stratum", "outer_fold"]),
            pd.read_csv(SOURCE_OOF, sep="\t"),
            domains,
            "source",
            promotion_signature,
        )
        target_endpoint_reconstruction = reconstruct_endpoint_population(
            target,
            sealed_oof,
            domains,
            "target",
            promotion_signature,
        )
        endpoint_results = [
            source_endpoint_reconstruction,
            target_endpoint_reconstruction,
        ]
        endpoint_index = pd.DataFrame(
            [
                row
                for result in endpoint_results
                for row in result["index_rows"]
            ]
        )
        endpoint_coefficients = pd.DataFrame(
            [
                row
                for result in endpoint_results
                for row in result["coefficient_rows"]
            ]
        )
        endpoint_parity = pd.DataFrame(
            [
                row
                for result in endpoint_results
                for row in result["parity_rows"]
            ]
        )
        endpoint_index.to_csv(
            OUT / "endpoint_model_artifact_index.tsv",
            sep="\t",
            index=False,
        )
        endpoint_coefficients.to_csv(
            OUT / "endpoint_model_coefficients.tsv.gz",
            sep="\t",
            index=False,
            compression="gzip",
        )
        endpoint_parity.to_csv(
            OUT / "endpoint_oof_parity.tsv",
            sep="\t",
            index=False,
        )
        max_endpoint_reload_error = max(
            result["max_reload_error"] for result in endpoint_results
        )
        max_endpoint_upstream_error = max(
            result["max_upstream_error"] for result in endpoint_results
        )
        endpoint_artifact_count = int(len(endpoint_index))
        endpoint_hash_gate = bool(
            endpoint_index["artifact_sha256"].str.len().eq(64).all()
        )
        endpoint_parity_gate = bool(
            endpoint_parity["within_tolerance_1e_12"].all()
        )
        promotion_succeeded = bool(
            endpoint_artifact_count == 2 * N_OUTER_FOLDS
            and endpoint_hash_gate
            and endpoint_parity_gate
            and max_endpoint_reload_error <= PARITY_TOLERANCE
            and max_endpoint_upstream_error <= PARITY_TOLERANCE
        )
        if not promotion_succeeded:
            raise RuntimeError(
                "STOP_DATA_OR_ARTIFACT_FAIL: endpoint reconstruction gate failed"
            )
        endpoint_audit = {
            "status": "PASS",
            "promotion_signature": promotion_signature,
            "source_analysis_signature": analysis_signature,
            "expected_outer_artifact_count": 2 * N_OUTER_FOLDS,
            "outer_artifact_count": endpoint_artifact_count,
            "newly_fitted_outer_artifact_count": int(
                sum(result["newly_fitted"] for result in endpoint_results)
            ),
            "reused_outer_artifact_count": int(
                sum(result["reused"] for result in endpoint_results)
            ),
            "fitted_endpoint_estimator_count": int(
                sum(result["estimator_count"] for result in endpoint_results)
            ),
            "model_columns": list(endpoint_model_specs(domains)),
            "reload_parity_tolerance": PARITY_TOLERANCE,
            "upstream_oof_parity_tolerance": PARITY_TOLERANCE,
            "max_reload_prediction_error": max_endpoint_reload_error,
            "max_upstream_oof_prediction_error": max_endpoint_upstream_error,
            "all_model_hashes_present": endpoint_hash_gate,
            "all_upstream_oof_columns_within_tolerance": endpoint_parity_gate,
            "source_and_target_folds": "sealed legacy folds",
            "model_specification": {
                "C": 1.0,
                "l1_ratio": 0.0,
                "solver": "lbfgs",
                "max_iter": 3000,
                "background": list(BACKGROUND),
                "age_basis": "training-fold standardized linear, square and cube",
                "measurement_scaling": "training-fold mean and population SD",
            },
        }
    except Exception as error:
        promotion_succeeded = False
        promotion_failure_reason = f"{type(error).__name__}: {error}"
        endpoint_audit = {
            "status": "FAIL",
            "promotion_signature": promotion_signature,
            "source_analysis_signature": analysis_signature,
            "evidence_status_retained": "probe",
            "failure_reason": promotion_failure_reason,
            "required_upstream_oof_parity_tolerance": PARITY_TOLERANCE,
        }
    write_json(
        OUT / "endpoint_model_artifact_audit.json", endpoint_audit
    )
    evidence_status = (
        "validated_candidate" if promotion_succeeded else "probe"
    )

    audit = {
        "status": (
            "PASS" if promotion_succeeded else "FAIL_ENDPOINT_RECONSTRUCTION"
        ),
        "analysis_version": ANALYSIS_VERSION,
        "analysis_signature": analysis_signature,
        "promotion_signature": promotion_signature,
        "evidence_status": evidence_status,
        "target_endpoint_predictions": "sealed_upstream",
        "legacy_source_and_target_endpoint_models_persisted_here": (
            promotion_succeeded
        ),
        "endpoint_reconstruction_failure_reason": promotion_failure_reason,
        "expected_outer_artifact_count": N_OUTER_FOLDS,
        "outer_artifact_count": int(len(model_index)),
        "newly_fitted_outer_artifact_count": fitted_count,
        "reused_outer_artifact_count": reused_count,
        "tuning_estimator_count": tuning_estimator_count,
        "outer_final_estimator_count": outer_estimator_count,
        "parity_tolerance": PARITY_TOLERANCE,
        "max_reload_prediction_error": max_parity_error,
        "all_model_hashes_present": bool(
            model_index["artifact_sha256"].str.len().eq(64).all()
        ),
        "endpoint_model_artifact_audit_status": endpoint_audit["status"],
        "endpoint_outer_artifact_count": int(
            endpoint_audit.get("outer_artifact_count", 0)
        ),
        "endpoint_estimator_count": int(
            endpoint_audit.get("fitted_endpoint_estimator_count", 0)
        ),
        "endpoint_max_reload_prediction_error": endpoint_audit.get(
            "max_reload_prediction_error"
        ),
        "endpoint_max_upstream_oof_prediction_error": endpoint_audit.get(
            "max_upstream_oof_prediction_error"
        ),
        "source_participants": int(len(source)),
        "source_events": int(source["death_5y"].sum()),
        "target_participants": int(len(target_eval)),
        "target_events": int(outcome.sum()),
        "software": {
            "python": sys.version,
            "python_executable": sys.executable,
            "platform": platform.platform(),
            "numpy": np.__version__,
            "pandas": pd.__version__,
            "scikit_learn": sklearn.__version__,
            "joblib": joblib.__version__,
        },
    }
    write_json(OUT / "model_artifact_audit.json", audit)

    decision = {
        "classification": classification,
        "evidence_status": evidence_status,
        "analysis_version": ANALYSIS_VERSION,
        "analysis_signature": analysis_signature,
        "promotion_signature": promotion_signature,
        "source_cycle": SOURCE_CYCLE,
        "target_cycle": TARGET_CYCLE,
        "source_endpoint_selected_domain": endpoint_domain,
        "source_inverse_age_decoder_selected_domain": inverse_domain,
        "target_endpoint_gain": float(
            policy_summary.loc[
                policy_summary["policy"].eq("source_endpoint_policy"),
                "gain_vs_age_and_background",
            ].iloc[0]
        ),
        "target_inverse_age_decoder_gain": float(
            policy_summary.loc[
                policy_summary["policy"].eq(
                    "source_inverse_age_decoder_policy"
                ),
                "gain_vs_age_and_background",
            ].iloc[0]
        ),
        "endpoint_minus_inverse_estimate": contrast_estimate,
        "endpoint_minus_inverse_ci_low": float(contrast_low),
        "endpoint_minus_inverse_ci_high": float(contrast_high),
        "endpoint_minus_inverse_bootstrap_probability_positive": float(
            np.mean(contrast_replicates > 0)
        ),
        "target_predictions_are_sealed_upstream": True,
        "legacy_source_and_target_endpoint_fitted_objects_reconstructed": (
            promotion_succeeded
        ),
        "endpoint_reconstruction_audit_status": endpoint_audit["status"],
        "endpoint_reconstruction_failure_reason": promotion_failure_reason,
        "claim_ceiling": (
            "One exploratory source-target cycle pair, five frozen "
            "equal-category-cost domains, declared ridge age decoders and "
            "exactly parity-checked source and target endpoint predictions."
        ),
    }
    write_json(OUT / "decision.json", decision)

    selected_row = domain_pooled.set_index("domain").loc[inverse_domain]
    report = f"""# NHANES endpoint versus inverse-age-decoder acquisition v0.2

Date: 26 August 2026

## Verdict

**{classification}**

Evidence status: **{evidence_status}**. The endpoint fitted-object
reconstruction audit is **{endpoint_audit['status']}**.

## Source inverse chronological-age decoder

The source cycle contained {len(source):,} participants. Five prespecified
domains were compared with nested outcome-blind ridge decoders of chronological
age. The selected domain was **{inverse_domain}** with pooled OOF age
MSE {selected_row['mse']:.6f}, RMSE {selected_row['rmse']:.6f} years,
MAE {selected_row['mae']:.6f} years and R2 {selected_row['r2']:.6f}.
The delta MSE versus W-only was {selected_row['delta_mse_vs_W_only']:.6f}.

The separately frozen endpoint-informed source policy selected
**{endpoint_domain}**.

## Target mortality policy comparison

In the 2009--2010 target cycle ({len(target_eval):,} participants;
{int(outcome.sum()):,} five-year deaths), the endpoint-selected policy gained
{decision['target_endpoint_gain']:.9f} log-loss units beyond age and
background. The inverse-age-decoder-selected policy gained
{decision['target_inverse_age_decoder_gain']:.9f}.

The paired endpoint-minus-inverse contrast was
{contrast_estimate:.9f}, with a fixed-prediction event-stratified bootstrap
interval from {float(contrast_low):.9f} to {float(contrast_high):.9f}.

## Persistence and boundary

All {N_OUTER_FOLDS} outer artifacts, {tuning_estimator_count} inner tuning
estimators and {outer_estimator_count} outer final estimators were persisted,
reloaded and reproduced at maximum absolute prediction error
{max_parity_error:.3g} (tolerance {PARITY_TOLERANCE:.1e}).

The legacy source and target endpoint models for CA and all five CA_M domains
were {'reconstructed, persisted and reload-verified' if promotion_succeeded else 'not promoted because reconstruction failed'}.
The endpoint reconstruction used
{endpoint_audit.get('outer_artifact_count', 0)} outer artifacts and
{endpoint_audit.get('fitted_endpoint_estimator_count', 0)} fitted endpoint
estimators. The maximum reload error was
{endpoint_audit.get('max_reload_prediction_error', float('nan'))} and the
maximum difference from sealed upstream OOF predictions was
{endpoint_audit.get('max_upstream_oof_prediction_error', float('nan'))},
against tolerance {PARITY_TOLERANCE:.1e}.
{'Failure reason: ' + promotion_failure_reason if promotion_failure_reason else ''}

This result compares two frozen acquisition policies across five
equal-category-cost domains in one exploratory source/target cycle pair. It is
not clinical value of information, a cost analysis, untouched confirmation,
universal transportability or biological-age identification. Bootstrap
intervals condition on stored predictions and fixed source selections and omit
refitting and study-development uncertainty.
"""
    (OUT / "REPORT.md").write_text(report, encoding="utf-8")
    write_artifact_manifest()

    print(json.dumps(json_ready(decision), indent=2, ensure_ascii=False))
    print()
    print(domain_pooled.to_string(index=False))
    print()
    print(contrast.to_string(index=False))
    print()
    print(json.dumps(json_ready(audit), indent=2, ensure_ascii=False))
    return 0


if __name__ == "__main__":
    raise SystemExit(main())
```
