## Supplementary_Software_1 for "Inferential boundaries of age prediction: why prediction does not establish biological age measurement": analyze_nhanes_individual_agegap_contributions_v0_1.py

```
#!/usr/bin/env python
"""Individual additive AgeGap contributions under the frozen NHANES age decoder."""

from __future__ import annotations

import hashlib
import json
import math
import platform
import sys
from pathlib import Path
from typing import Any

import joblib
import matplotlib
import numpy as np
import pandas as pd
import sklearn
from sklearn.linear_model import Ridge
from sklearn.model_selection import StratifiedKFold
from sklearn.preprocessing import OneHotEncoder, SplineTransformer


matplotlib.use("Agg")
import matplotlib.pyplot as plt


ROOT = Path(__file__).resolve().parents[1]
ANALYSIS = ROOT / "analysis"
BASE = ROOT / "outputs" / "nhanes_5y_complete_support_contextual_age_probe_v0_1"
DECODER = ROOT / "outputs" / "nhanes_5y_age_decoder_source_information_probe_v0_2"
GAP_COMPARISON = ROOT / "outputs" / "nhanes_5y_age_clock_gap_comparison_v0_1"
OUT = ROOT / "outputs" / "nhanes_individual_agegap_contributions_v0_1"
MODELS = OUT / "models"
FIGURES = OUT / "figures"

FRAME_PATH = BASE / "analysis_ready_risk_set.tsv.gz"
FOLDS_PATH = BASE / "participant_folds.tsv"
TIERS_PATH = BASE / "ordered_feature_tiers.tsv"
DECODER_OOF_PATH = DECODER / "oof_predictions.tsv.gz"
DECODER_AUDIT_PATH = DECODER / "model_artifact_audit.json"
DECODER_SIGNATURE_PATH = DECODER / "analysis_signature.json"
GAP_OOF_PATH = GAP_COMPARISON / "oof_predictions.tsv.gz"
GAP_AUDIT_PATH = GAP_COMPARISON / "model_artifact_audit.json"
SPEC_PATH = ANALYSIS / "NHANES_INDIVIDUAL_AGEGAP_CONTRIBUTIONS_SPEC_20260831.md"
ENV_POLICY = ROOT / "environment-policy.json"
ENV_MANIFEST = ROOT / "environment.yml"
ENV_LOCK = ROOT / "environment.win-64.lock.txt"

ANALYSIS_VERSION = "nhanes_individual_agegap_contributions_v0_1"
SCHEMA_VERSION = "1.0"
EXPECTED_N = 13_724
N_OUTER_FOLDS = 5
N_INNER_FOLDS = 3
INNER_SEED = 20260831
BACKGROUND = ("sex", "race_ethnicity", "cycle")
ALPHA_GRID = (0.01, 0.1, 1.0, 10.0, 100.0, 1000.0)
PARITY_TOLERANCE = 1e-12
ADDITIVITY_TOLERANCE = 1e-10
MATCH_CALIPER_YEARS = 0.25


DOMAIN_MAP = {
    "albumin": "protein/liver",
    "alt": "liver",
    "ast": "liver",
    "alkaline_phosphatase": "liver/bone",
    "blood_urea_nitrogen": "renal",
    "calcium": "mineral",
    "total_cholesterol": "lipid/metabolic",
    "bicarbonate": "electrolyte",
    "creatinine": "renal",
    "ggt": "liver",
    "glucose": "metabolic",
    "iron": "haematologic/mineral",
    "ldh": "tissue-turnover",
    "phosphorus": "mineral/renal",
    "total_bilirubin": "liver/haematologic",
    "total_protein": "protein/liver",
    "triglycerides": "lipid/metabolic",
    "uric_acid": "renal/metabolic",
    "sodium": "electrolyte",
    "potassium": "electrolyte",
    "chloride": "electrolyte",
    "osmolality": "electrolyte",
    "globulin": "protein/immune",
    "wbc": "haematologic/immune",
    "lymphocyte_percent": "haematologic/immune",
    "monocyte_percent": "haematologic/immune",
    "neutrophil_percent": "haematologic/immune",
    "eosinophil_percent": "haematologic/immune",
    "basophil_percent": "haematologic/immune",
    "lymphocyte_count": "haematologic/immune",
    "monocyte_count": "haematologic/immune",
    "neutrophil_count": "haematologic/immune",
    "eosinophil_count": "haematologic/immune",
    "basophil_count": "haematologic/immune",
    "rbc": "haematologic",
    "hemoglobin": "haematologic",
    "hematocrit": "haematologic",
    "mcv": "haematologic",
    "mch": "haematologic",
    "mchc": "haematologic",
    "rdw": "haematologic",
    "platelets": "haematologic",
    "mean_platelet_volume": "haematologic",
    "weight": "anthropometric",
    "bmi": "anthropometric/metabolic",
    "waist": "anthropometric/metabolic",
    "systolic_bp": "cardiovascular",
    "diastolic_bp": "cardiovascular",
    "pulse": "cardiovascular",
    "hdl": "lipid/metabolic",
}


def configure_stdout() -> None:
    if hasattr(sys.stdout, "reconfigure"):
        sys.stdout.reconfigure(encoding="utf-8")


def sha256(path: Path) -> str:
    digest = hashlib.sha256()
    with path.open("rb") as handle:
        for block in iter(lambda: handle.read(1024 * 1024), b""):
            digest.update(block)
    return digest.hexdigest()


def stable_hash(payload: Any) -> str:
    encoded = json.dumps(payload, sort_keys=True, separators=(",", ":"), ensure_ascii=False).encode("utf-8")
    return hashlib.sha256(encoded).hexdigest()


def json_ready(value: Any) -> Any:
    if isinstance(value, dict):
        return {str(key): json_ready(item) for key, item in value.items()}
    if isinstance(value, (list, tuple)):
        return [json_ready(item) for item in value]
    if isinstance(value, Path):
        return str(value)
    if isinstance(value, np.ndarray):
        return value.tolist()
    if isinstance(value, np.integer):
        return int(value)
    if isinstance(value, np.floating):
        return None if not np.isfinite(value) else float(value)
    if isinstance(value, float):
        return None if not math.isfinite(value) else value
    return value


def write_json(path: Path, payload: Any) -> None:
    path.write_text(json.dumps(json_ready(payload), indent=2, ensure_ascii=False) + "\n", encoding="utf-8")


def clean_categories(frame: pd.DataFrame) -> pd.DataFrame:
    return frame.loc[:, BACKGROUND].fillna("missing").astype(str).replace("", "missing")


def display_name(feature: str) -> str:
    plain = feature.removeprefix("T__")
    replacements = {
        "alt": "ALT",
        "ast": "AST",
        "ggt": "GGT",
        "ldh": "LDH",
        "wbc": "WBC",
        "rbc": "RBC",
        "mcv": "MCV",
        "mch": "MCH",
        "mchc": "MCHC",
        "rdw": "RDW",
        "bmi": "BMI",
        "hdl": "HDL",
    }
    if plain in replacements:
        return replacements[plain]
    return plain.replace("_", " ").title()


def make_age_strata(age: np.ndarray) -> np.ndarray:
    strata = pd.qcut(pd.Series(age), q=5, duplicates="drop", labels=False).fillna(-1).astype(int).to_numpy()
    if int(pd.Series(strata).value_counts().min()) < N_INNER_FOLDS:
        raise RuntimeError("Age strata cannot support inner folds")
    return strata


def fit_conditional_model(frame: pd.DataFrame, features: list[str], alpha: float) -> dict[str, Any]:
    encoder = OneHotEncoder(handle_unknown="ignore", sparse_output=False)
    w = encoder.fit_transform(clean_categories(frame))
    spline = SplineTransformer(n_knots=5, degree=3, include_bias=False, extrapolation="linear")
    a = spline.fit_transform(frame[["age_years"]].to_numpy(float))
    design = np.column_stack([w, a])
    estimator = Ridge(alpha=float(alpha))
    estimator.fit(design, frame.loc[:, features].to_numpy(float))
    return {
        "encoder": encoder,
        "spline": spline,
        "estimator": estimator,
        "design_feature_names": [f"W__{x}" for x in encoder.get_feature_names_out(BACKGROUND)]
        + [f"A_spline_{i}" for i in range(a.shape[1])],
        "features": features,
    }


def predict_conditional_model(model: dict[str, Any], frame: pd.DataFrame) -> np.ndarray:
    w = model["encoder"].transform(clean_categories(frame))
    a = model["spline"].transform(frame[["age_years"]].to_numpy(float))
    return model["estimator"].predict(np.column_stack([w, a]))


def standardized_mse(train: pd.DataFrame, valid: pd.DataFrame, features: list[str], prediction: np.ndarray) -> float:
    scale = train.loc[:, features].std(axis=0, ddof=0).to_numpy(float)
    scale[scale < 1e-12] = 1.0
    error = (valid.loc[:, features].to_numpy(float) - prediction) / scale
    return float(np.mean(error**2))


def predict_decoder(model: dict[str, Any], frame: pd.DataFrame, features: list[str]) -> np.ndarray:
    w = model["encoder"].transform(clean_categories(frame))
    x = model["scaler"].transform(frame.loc[:, features].to_numpy(float))
    return model["estimator"].predict(np.column_stack([w, x]))


def load_and_gate() -> tuple[pd.DataFrame, list[str], pd.DataFrame, dict[str, Any]]:
    decoder_audit = json.loads(DECODER_AUDIT_PATH.read_text(encoding="utf-8"))
    gap_audit = json.loads(GAP_AUDIT_PATH.read_text(encoding="utf-8"))
    if decoder_audit.get("status") != "PASS" or gap_audit.get("status") != "PASS":
        raise RuntimeError("STOP_DATA_OR_ARTIFACT_FAIL: predecessor artifact audit is not PASS")
    frame = pd.read_csv(FRAME_PATH, sep="\t")
    folds = pd.read_csv(FOLDS_PATH, sep="\t")
    tiers = pd.read_csv(TIERS_PATH, sep="\t")
    decoder_oof = pd.read_csv(DECODER_OOF_PATH, sep="\t")
    gap_oof = pd.read_csv(GAP_OOF_PATH, sep="\t")
    features = tiers.loc[tiers["tier"].eq("S3")].sort_values("order")["transformed_column"].tolist()
    if len(features) != 50 or len(set(features)) != 50:
        raise RuntimeError("Frozen S3 panel is not 50 ordered unique features")
    required = ["row_id", "SEQN", "age_years", "outer_fold", *BACKGROUND, *features]
    if sorted(set(required) - set(frame.columns)):
        raise RuntimeError(f"Missing analysis columns: {sorted(set(required) - set(frame.columns))}")
    if len(frame) != EXPECTED_N or frame["row_id"].nunique() != EXPECTED_N or frame["SEQN"].nunique() != EXPECTED_N:
        raise RuntimeError("Participant identity/count changed")
    if frame[required].isna().any().any():
        raise RuntimeError("Missing analysis input")
    fold_check = frame[["row_id", "SEQN", "outer_fold"]].merge(
        folds[["row_id", "SEQN", "outer_fold"]], on=["row_id", "SEQN"], suffixes=("_frame", "_frozen"), validate="one_to_one"
    )
    if not np.array_equal(fold_check["outer_fold_frame"].to_numpy(), fold_check["outer_fold_frozen"].to_numpy()):
        raise RuntimeError("Frozen fold mismatch")
    keep_oof = ["row_id", "SEQN", "outer_fold", "clock_score", "clock_gap"]
    decoder_oof = frame[["row_id", "SEQN"]].merge(decoder_oof[keep_oof], on=["row_id", "SEQN"], validate="one_to_one")
    gap_keep = ["row_id", "SEQN", "outer_fold", "M_X", "M_AGE_GAP"]
    gap_oof = frame[["row_id", "SEQN"]].merge(gap_oof[gap_keep], on=["row_id", "SEQN"], validate="one_to_one")
    for joined in (decoder_oof, gap_oof):
        if not np.array_equal(joined["outer_fold"].to_numpy(int), frame["outer_fold"].to_numpy(int)):
            raise RuntimeError("OOF participant/fold order mismatch")
    if np.max(np.abs(decoder_oof["clock_gap"].to_numpy(float) - (decoder_oof["clock_score"] - frame["age_years"]))) > PARITY_TOLERANCE:
        raise RuntimeError("Stored raw AgeGap identity failed")
    endpoint = gap_oof[["row_id", "M_X", "M_AGE_GAP"]].copy()
    return frame.reset_index(drop=True), features, decoder_oof, endpoint


def fit_residualizer_artifact(
    frame: pd.DataFrame,
    features: list[str],
    outer_fold: int,
    analysis_signature: str,
) -> dict[str, Any]:
    train = frame.loc[frame["outer_fold"].ne(outer_fold)].copy().reset_index(drop=True)
    test = frame.loc[frame["outer_fold"].eq(outer_fold)].copy().reset_index(drop=True)
    splitter = StratifiedKFold(
        n_splits=N_INNER_FOLDS,
        shuffle=True,
        random_state=INNER_SEED + outer_fold,
    )
    splits = list(splitter.split(np.zeros(len(train)), make_age_strata(train["age_years"].to_numpy(float))))
    tuning_fits: list[dict[str, Any]] = []
    tuning_rows: list[dict[str, Any]] = []
    for alpha in ALPHA_GRID:
        for inner_fold, (tr, va) in enumerate(splits):
            model = fit_conditional_model(train.iloc[tr], features, alpha)
            prediction = predict_conditional_model(model, train.iloc[va])
            mse = standardized_mse(train.iloc[tr], train.iloc[va], features, prediction)
            tuning_fits.append(
                {
                    "alpha": float(alpha),
                    "inner_fold": inner_fold,
                    "train_row_ids": train.iloc[tr]["row_id"].to_numpy(int),
                    "valid_row_ids": train.iloc[va]["row_id"].to_numpy(int),
                    "model": model,
                    "valid_standardized_mse": mse,
                }
            )
            tuning_rows.append({"alpha": float(alpha), "inner_fold": inner_fold, "standardized_mse": mse})
    tuning = pd.DataFrame(tuning_rows)
    selected_alpha = float(tuning.groupby("alpha", as_index=False)["standardized_mse"].mean().sort_values(["standardized_mse", "alpha"]).iloc[0]["alpha"])
    final_model = fit_conditional_model(train, features, selected_alpha)
    test_conditional_mean = predict_conditional_model(final_model, test)
    signature = stable_hash(
        {
            "analysis_signature": analysis_signature,
            "outer_fold": outer_fold,
            "train_row_ids": train["row_id"].tolist(),
            "test_row_ids": test["row_id"].tolist(),
            "features": features,
            "alpha_grid": ALPHA_GRID,
            "inner_seed": INNER_SEED + outer_fold,
        }
    )
    return {
        "schema_version": SCHEMA_VERSION,
        "analysis_signature": analysis_signature,
        "artifact_signature": signature,
        "outer_fold": outer_fold,
        "features": features,
        "train_row_ids": train["row_id"].to_numpy(int),
        "test_row_ids": test["row_id"].to_numpy(int),
        "tuning_fits": tuning_fits,
        "tuning_rows": tuning_rows,
        "selected_alpha": selected_alpha,
        "final_model": final_model,
        "test_conditional_mean": test_conditional_mean,
    }


def verify_residualizer_artifact(
    artifact: dict[str, Any], frame_by_row: pd.DataFrame, analysis_signature: str
) -> tuple[float, np.ndarray]:
    if artifact.get("analysis_signature") != analysis_signature:
        raise RuntimeError("Residualizer analysis signature mismatch")
    features = list(artifact["features"])
    max_error = 0.0
    for fit in artifact["tuning_fits"]:
        train = frame_by_row.loc[np.asarray(fit["train_row_ids"], dtype=int)]
        valid = frame_by_row.loc[np.asarray(fit["valid_row_ids"], dtype=int)]
        prediction = predict_conditional_model(fit["model"], valid)
        mse = standardized_mse(train, valid, features, prediction)
        max_error = max(max_error, abs(mse - float(fit["valid_standardized_mse"])))
    test = frame_by_row.loc[np.asarray(artifact["test_row_ids"], dtype=int)]
    prediction = predict_conditional_model(artifact["final_model"], test)
    max_error = max(max_error, float(np.max(np.abs(prediction - np.asarray(artifact["test_conditional_mean"], dtype=float)))))
    if max_error > PARITY_TOLERANCE:
        raise RuntimeError(f"STOP_DATA_OR_ARTIFACT_FAIL: residualizer reload parity error={max_error}")
    return max_error, prediction


def fit_or_load_residualizer(
    frame: pd.DataFrame, features: list[str], outer_fold: int, analysis_signature: str
) -> tuple[dict[str, Any], bool, float, np.ndarray]:
    path = MODELS / f"outer_fold_{outer_fold:02d}_conditional_x.joblib"
    if path.exists():
        artifact = joblib.load(path)
        reused = True
    else:
        artifact = fit_residualizer_artifact(frame, features, outer_fold, analysis_signature)
        joblib.dump(artifact, path, compress=3)
        reused = False
    reloaded = joblib.load(path)
    error, prediction = verify_residualizer_artifact(reloaded, frame.set_index("row_id", drop=False), analysis_signature)
    return reloaded, reused, error, prediction


def calculate_contributions(
    frame: pd.DataFrame,
    features: list[str],
    decoder_oof: pd.DataFrame,
    endpoint: pd.DataFrame,
    analysis_signature: str,
) -> tuple[pd.DataFrame, pd.DataFrame, pd.DataFrame, pd.DataFrame, dict[str, Any]]:
    n, p = len(frame), len(features)
    conditional_x = np.full((n, p), np.nan)
    contributions = np.full((n, p), np.nan)
    expected_score = np.full(n, np.nan)
    reconstructed_score = np.full(n, np.nan)
    index_rows: list[dict[str, Any]] = []
    tuning_rows: list[dict[str, Any]] = []
    coefficient_rows: list[dict[str, Any]] = []
    max_decoder_error = 0.0
    max_residualizer_error = 0.0
    reused_count = 0
    fitted_count = 0
    decoder_index = pd.read_csv(DECODER / "model_artifact_index.tsv", sep="\t").set_index("outer_fold")
    for outer_fold in range(N_OUTER_FOLDS):
        print(f"Outer fold {outer_fold + 1}/{N_OUTER_FOLDS}", flush=True)
        mask = frame["outer_fold"].eq(outer_fold).to_numpy()
        positions = np.where(mask)[0]
        test = frame.iloc[positions].copy()
        decoder_path = DECODER / "models" / f"outer_fold_{outer_fold:02d}.joblib"
        if sha256(decoder_path) != str(decoder_index.loc[outer_fold, "artifact_sha256"]):
            raise RuntimeError("Frozen decoder artifact hash mismatch")
        decoder_artifact = joblib.load(decoder_path)
        decoder_model = decoder_artifact["outer_age_model"]
        if list(decoder_artifact["features"]) != features:
            raise RuntimeError("Frozen decoder feature order mismatch")
        if not np.array_equal(test["row_id"].to_numpy(int), np.asarray(decoder_artifact["test_row_ids"], dtype=int)):
            raise RuntimeError("Frozen decoder test order mismatch")
        decoder_prediction = predict_decoder(decoder_model, test, features)
        stored = decoder_oof.loc[mask, "clock_score"].to_numpy(float)
        decoder_error = float(np.max(np.abs(decoder_prediction - stored)))
        max_decoder_error = max(max_decoder_error, decoder_error)
        if decoder_error > PARITY_TOLERANCE:
            raise RuntimeError("Frozen decoder prediction parity failed")

        residualizer, reused, residualizer_error, conditional = fit_or_load_residualizer(
            frame, features, outer_fold, analysis_signature
        )
        reused_count += int(reused)
        fitted_count += int(not reused)
        max_residualizer_error = max(max_residualizer_error, residualizer_error)
        if not np.array_equal(test["row_id"].to_numpy(int), np.asarray(residualizer["test_row_ids"], dtype=int)):
            raise RuntimeError("Residualizer test order mismatch")
        conditional_x[positions] = conditional

        n_w = sum(len(categories) for categories in decoder_model["encoder"].categories_)
        names = list(decoder_model["design_feature_names"])
        expected_measure_names = [f"X__{feature}" for feature in features]
        if names[n_w:] != expected_measure_names:
            raise RuntimeError("Decoder measurement coefficient order mismatch")
        beta = np.asarray(decoder_model["estimator"].coef_, dtype=float)[n_w:]
        scale = np.asarray(decoder_model["scaler"].scale_, dtype=float)
        mean = np.asarray(decoder_model["scaler"].mean_, dtype=float)
        contributions[positions] = (test.loc[:, features].to_numpy(float) - conditional) / scale * beta
        w = decoder_model["encoder"].transform(clean_categories(test))
        expected_design = np.column_stack([w, (conditional - mean) / scale])
        expected_score[positions] = decoder_model["estimator"].predict(expected_design)
        reconstructed_score[positions] = expected_score[positions] + contributions[positions].sum(axis=1)

        residualizer_path = MODELS / f"outer_fold_{outer_fold:02d}_conditional_x.joblib"
        index_rows.append(
            {
                "outer_fold": outer_fold,
                "selected_alpha": residualizer["selected_alpha"],
                "n_train": len(residualizer["train_row_ids"]),
                "n_test": len(residualizer["test_row_ids"]),
                "artifact_path": str(residualizer_path.relative_to(ROOT)),
                "artifact_sha256": sha256(residualizer_path),
                "source_decoder_path": str(decoder_path.relative_to(ROOT)),
                "source_decoder_sha256": sha256(decoder_path),
                "reused": reused,
                "max_reload_error": residualizer_error,
                "decoder_prediction_error": decoder_error,
            }
        )
        for row in residualizer["tuning_rows"]:
            tuning_rows.append({"outer_fold": outer_fold, **row})
        for j, feature in enumerate(features):
            plain = feature.removeprefix("T__")
            coefficient_rows.append(
                {
                    "outer_fold": outer_fold,
                    "feature_order": j + 1,
                    "feature": feature,
                    "display_feature": display_name(feature),
                    "domain": DOMAIN_MAP.get(plain, "other"),
                    "decoder_beta_per_standardized_unit": beta[j],
                    "decoder_beta_per_raw_unit": beta[j] / scale[j],
                    "decoder_scaler_mean": mean[j],
                    "decoder_scaler_scale": scale[j],
                    "residualizer_alpha": residualizer["selected_alpha"],
                }
            )

    if not np.isfinite(contributions).all() or not np.isfinite(conditional_x).all():
        raise RuntimeError("Incomplete participant contribution output")
    score = decoder_oof["clock_score"].to_numpy(float)
    raw_gap = decoder_oof["clock_gap"].to_numpy(float)
    centered_gap = contributions.sum(axis=1)
    score_error = float(np.max(np.abs(reconstructed_score - score)))
    centered_identity_error = float(np.max(np.abs(centered_gap - (score - expected_score))))
    if score_error > ADDITIVITY_TOLERANCE or centered_identity_error > ADDITIVITY_TOLERANCE:
        raise RuntimeError(f"Contribution additivity failed score={score_error} centered={centered_identity_error}")
    participants = frame[["row_id", "SEQN", "outer_fold", "age_years", *BACKGROUND]].copy()
    participants["clock_score"] = score
    participants["raw_agegap_years"] = raw_gap
    participants["conditional_expected_clock_score"] = expected_score
    participants["conditional_expected_raw_agegap"] = expected_score - participants["age_years"].to_numpy(float)
    participants["centered_agegap_years"] = centered_gap
    participants["sum_feature_contributions_years"] = centered_gap
    participants = participants.merge(endpoint, on="row_id", validate="one_to_one")

    dictionary = pd.DataFrame(
        [
            {
                "feature_order": j + 1,
                "feature": feature,
                "display_feature": display_name(feature),
                "domain": DOMAIN_MAP.get(feature.removeprefix("T__"), "other"),
            }
            for j, feature in enumerate(features)
        ]
    )
    long = pd.DataFrame(
        {
            "row_id": np.repeat(participants["row_id"].to_numpy(int), p),
            "SEQN": np.repeat(participants["SEQN"].to_numpy(int), p),
            "outer_fold": np.repeat(participants["outer_fold"].to_numpy(int), p),
            "feature_order": np.tile(np.arange(1, p + 1), n),
            "feature": np.tile(features, n),
            "observed_value": frame.loc[:, features].to_numpy(float).ravel(),
            "conditional_expected_value": conditional_x.ravel(),
            "conditional_residual": (frame.loc[:, features].to_numpy(float) - conditional_x).ravel(),
            "contribution_years": contributions.ravel(),
        }
    ).merge(dictionary, on=["feature_order", "feature"], validate="many_to_one")
    audit = {
        "status": "PASS",
        "analysis_signature": analysis_signature,
        "expected_residualizer_artifact_count": N_OUTER_FOLDS,
        "residualizer_artifact_count": len(index_rows),
        "newly_fitted_residualizer_count": fitted_count,
        "reused_residualizer_count": reused_count,
        "tuning_estimator_count": N_OUTER_FOLDS * len(ALPHA_GRID) * N_INNER_FOLDS,
        "final_residualizer_count": N_OUTER_FOLDS,
        "source_decoder_count": N_OUTER_FOLDS,
        "parity_tolerance": PARITY_TOLERANCE,
        "additivity_tolerance": ADDITIVITY_TOLERANCE,
        "max_residualizer_reload_error": max_residualizer_error,
        "max_decoder_prediction_error": max_decoder_error,
        "max_score_reconstruction_error": score_error,
        "max_centered_gap_identity_error": centered_identity_error,
        "all_residualizer_hashes_present": all(len(row["artifact_sha256"]) == 64 for row in index_rows),
    }
    return participants, long, pd.DataFrame(coefficient_rows), pd.DataFrame(index_rows), pd.DataFrame(tuning_rows), audit


def logit(probability: np.ndarray) -> np.ndarray:
    p = np.clip(np.asarray(probability, dtype=float), 1e-7, 1.0 - 1e-7)
    return np.log(p / (1.0 - p))


def build_pairs(participants: pd.DataFrame, contribution_matrix: np.ndarray) -> pd.DataFrame:
    by_row = {int(row_id): i for i, row_id in enumerate(participants["row_id"].to_numpy(int))}
    rows: list[dict[str, Any]] = []
    strata = ["outer_fold", "age_years", *BACKGROUND]
    for _, group in participants.groupby(strata, sort=True, dropna=False):
        ordered = group.sort_values(["centered_agegap_years", "row_id"]).reset_index(drop=True)
        if len(ordered) < 2:
            continue
        edges = []
        for j in range(len(ordered) - 1):
            left = ordered.iloc[j]
            right = ordered.iloc[j + 1]
            difference = abs(float(right["centered_agegap_years"] - left["centered_agegap_years"]))
            edges.append((difference, int(left["row_id"]), int(right["row_id"]), left, right))
        used: set[int] = set()
        for difference, left_id, right_id, left, right in sorted(edges, key=lambda item: (item[0], item[1], item[2])):
            if difference > MATCH_CALIPER_YEARS or left_id in used or right_id in used:
                continue
            used.update((left_id, right_id))
            c1 = contribution_matrix[by_row[left_id]]
            c2 = contribution_matrix[by_row[right_id]]
            delta = c1 - c2
            norm_product = float(np.linalg.norm(c1) * np.linalg.norm(c2))
            cosine = float(c1 @ c2 / norm_product) if norm_product > 0 else np.nan
            top1 = set(np.argsort(np.abs(c1))[-5:].tolist())
            top2 = set(np.argsort(np.abs(c2))[-5:].tolist())
            top_jaccard = len(top1 & top2) / len(top1 | top2)
            rows.append(
                {
                    "row_id_1": left_id,
                    "row_id_2": right_id,
                    "SEQN_1": int(left["SEQN"]),
                    "SEQN_2": int(right["SEQN"]),
                    "outer_fold": int(left["outer_fold"]),
                    "age_years": float(left["age_years"]),
                    "sex": left["sex"],
                    "race_ethnicity": left["race_ethnicity"],
                    "cycle": left["cycle"],
                    "raw_agegap_1": float(left["raw_agegap_years"]),
                    "raw_agegap_2": float(right["raw_agegap_years"]),
                    "raw_agegap_mean": float((left["raw_agegap_years"] + right["raw_agegap_years"]) / 2),
                    "raw_agegap_abs_difference": abs(float(left["raw_agegap_years"] - right["raw_agegap_years"])),
                    "centered_agegap_1": float(left["centered_agegap_years"]),
                    "centered_agegap_2": float(right["centered_agegap_years"]),
                    "centered_agegap_mean": float((left["centered_agegap_years"] + right["centered_agegap_years"]) / 2),
                    "centered_agegap_abs_difference": difference,
                    "contribution_l2_distance_years": float(np.linalg.norm(delta)),
                    "contribution_l1_distance_years": float(np.abs(delta).sum()),
                    "contribution_cosine_similarity": cosine,
                    "top5_absolute_contributor_jaccard": top_jaccard,
                    "M_X_1": float(left["M_X"]),
                    "M_X_2": float(right["M_X"]),
                    "M_X_abs_logit_difference": float(abs(logit(np.array([left["M_X"]]))[0] - logit(np.array([right["M_X"]]))[0])),
                    "M_AGE_GAP_1": float(left["M_AGE_GAP"]),
                    "M_AGE_GAP_2": float(right["M_AGE_GAP"]),
                    "M_AGE_GAP_abs_logit_difference": float(abs(logit(np.array([left["M_AGE_GAP"]]))[0] - logit(np.array([right["M_AGE_GAP"]]))[0])),
                }
            )
    pairs = pd.DataFrame(rows).sort_values(["outer_fold", "age_years", "row_id_1", "row_id_2"]).reset_index(drop=True)
    pairs.insert(0, "pair_id", [f"P{i + 1:05d}" for i in range(len(pairs))])
    if pairs.empty:
        raise RuntimeError("No matched pairs passed the frozen caliper")
    if pairs[["row_id_1", "row_id_2"]].stack().duplicated().any():
        raise RuntimeError("Matched-pair participant reuse detected")
    identity_error = float(np.max(np.abs(pairs["raw_agegap_abs_difference"] - pairs["centered_agegap_abs_difference"])))
    if identity_error > ADDITIVITY_TOLERANCE:
        raise RuntimeError(f"Raw/centered within-stratum gap-difference identity failed: {identity_error}")
    return pairs


def summarize_pairs(pairs: pd.DataFrame) -> pd.DataFrame:
    metrics = [
        "centered_agegap_abs_difference",
        "contribution_l2_distance_years",
        "contribution_l1_distance_years",
        "contribution_cosine_similarity",
        "top5_absolute_contributor_jaccard",
        "M_X_abs_logit_difference",
        "M_AGE_GAP_abs_logit_difference",
    ]
    rows = []
    for metric in metrics:
        values = pairs[metric].dropna().to_numpy(float)
        rows.append(
            {
                "metric": metric,
                "n": len(values),
                "mean": float(np.mean(values)),
                "sd": float(np.std(values, ddof=1)),
                "q025": float(np.quantile(values, 0.025)),
                "q25": float(np.quantile(values, 0.25)),
                "median": float(np.quantile(values, 0.5)),
                "q75": float(np.quantile(values, 0.75)),
                "q95": float(np.quantile(values, 0.95)),
                "q975": float(np.quantile(values, 0.975)),
                "max": float(np.max(values)),
            }
        )
    return pd.DataFrame(rows)


def select_display_pairs(pairs: pd.DataFrame, contributions: pd.DataFrame) -> tuple[pd.DataFrame, pd.DataFrame]:
    eligible = pairs.loc[
        np.sign(pairs["centered_agegap_1"]).eq(np.sign(pairs["centered_agegap_2"]))
        & pairs["centered_agegap_mean"].abs().ge(2.0)
    ].copy()
    if len(eligible) < 2:
        raise RuntimeError("Insufficient display-eligible matched pairs")
    selected_rows = []
    for label, quantile in (("median_distance", 0.50), ("p95_distance", 0.95)):
        target = float(eligible["contribution_l2_distance_years"].quantile(quantile))
        choice = eligible.assign(distance_to_target=(eligible["contribution_l2_distance_years"] - target).abs()).sort_values(
            ["distance_to_target", "pair_id"]
        ).iloc[0].drop(labels="distance_to_target")
        selected_rows.append({"case": label, "target_quantile": quantile, "target_distance": target, **choice.to_dict()})
    selected = pd.DataFrame(selected_rows)
    by = contributions.set_index(["row_id", "feature_order"])
    case_rows = []
    for case in selected.itertuples(index=False):
        ids = [int(case.row_id_1), int(case.row_id_2)]
        series = {row_id: by.loc[row_id]["contribution_years"].sort_index() for row_id in ids}
        top_union: set[int] = set()
        for row_id in ids:
            top_union.update((series[row_id].abs().nlargest(8).index.astype(int)).tolist())
        ordered_features = sorted(top_union, key=lambda j: -max(abs(series[ids[0]].loc[j]), abs(series[ids[1]].loc[j])))
        for person_number, row_id in enumerate(ids, start=1):
            person = contributions.loc[contributions["row_id"].eq(row_id)].set_index("feature_order")
            for feature_order in ordered_features:
                row = person.loc[feature_order]
                case_rows.append(
                    {
                        "case": case.case,
                        "pair_id": case.pair_id,
                        "person": person_number,
                        "row_id": row_id,
                        "feature_order": int(feature_order),
                        "feature": row["feature"],
                        "display_feature": row["display_feature"],
                        "domain": row["domain"],
                        "contribution_years": float(row["contribution_years"]),
                        "is_remainder": False,
                    }
                )
            remainder = float(person.loc[~person.index.isin(ordered_features), "contribution_years"].sum())
            case_rows.append(
                {
                    "case": case.case,
                    "pair_id": case.pair_id,
                    "person": person_number,
                    "row_id": row_id,
                    "feature_order": 999,
                    "feature": "OTHER",
                    "display_feature": "Other measured features",
                    "domain": "remainder",
                    "contribution_years": remainder,
                    "is_remainder": True,
                }
            )
    case_contributions = pd.DataFrame(case_rows)
    check = case_contributions.groupby(["case", "person"])["contribution_years"].sum().reset_index()
    expected = selected.set_index("case")
    for row in check.itertuples(index=False):
        target = float(expected.loc[row.case, f"centered_agegap_{row.person}"])
        if abs(float(row.contribution_years) - target) > ADDITIVITY_TOLERANCE:
            raise RuntimeError("Selected-case remainder additivity failed")
    return selected, case_contributions


def plot_case(ax: plt.Axes, selected: pd.DataFrame, case_data: pd.DataFrame, case: str, title: str) -> None:
    subset = case_data.loc[case_data["case"].eq(case)].copy()
    order = (
        subset.groupby("display_feature")["contribution_years"].apply(lambda x: float(np.max(np.abs(x)))).sort_values().index.tolist()
    )
    pivot = subset.pivot(index="display_feature", columns="person", values="contribution_years").reindex(order).fillna(0.0)
    y = np.arange(len(pivot))
    ax.barh(y - 0.18, pivot[1], height=0.34, color="#32688E", label="Participant A")
    ax.barh(y + 0.18, pivot[2], height=0.34, color="#D07A40", label="Participant B")
    ax.axvline(0, color="#777777", linewidth=0.8)
    ax.set_yticks(y, pivot.index, fontsize=7)
    ax.set_xlabel("Contribution to centred AgeGap (years)")
    row = selected.set_index("case").loc[case]
    ax.set_title(
        f"{title}\ncentred gaps {row['centered_agegap_1']:.2f}, {row['centered_agegap_2']:.2f}; profile L2={row['contribution_l2_distance_years']:.2f}",
        loc="left",
        fontsize=9,
        fontweight="bold",
    )


def make_figure(pairs: pd.DataFrame, selected: pd.DataFrame, case_data: pd.DataFrame) -> None:
    plt.rcParams.update({"font.family": "DejaVu Sans", "font.size": 8.5, "axes.spines.top": False, "axes.spines.right": False})
    fig, axes = plt.subplots(2, 2, figsize=(11.2, 8.4), constrained_layout=True)
    ax = axes[0, 0]
    ax.hist(pairs["contribution_l2_distance_years"], bins=45, color="#8498A5", edgecolor="white", linewidth=0.3)
    for case, color in (("median_distance", "#32688E"), ("p95_distance", "#B54C4C")):
        value = float(selected.set_index("case").loc[case, "contribution_l2_distance_years"])
        ax.axvline(value, color=color, linewidth=1.5, label=case.replace("_", " "))
    ax.set_xlabel("Matched-pair contribution-profile L2 distance (years)")
    ax.set_ylabel("Pairs")
    ax.set_title("a  Near-equal AgeGaps retain diverse internal profiles", loc="left", fontweight="bold")
    ax.legend(frameon=False, fontsize=7)
    plot_case(axes[0, 1], selected, case_data, "median_distance", "b  Predeclared median-distance case")
    plot_case(axes[1, 0], selected, case_data, "p95_distance", "c  Predeclared high-distance case")
    axes[0, 1].legend(frameon=False, fontsize=7)
    ax = axes[1, 1]
    for column, label, color in (
        ("M_AGE_GAP_abs_logit_difference", "Age + AgeGap model", "#7C8790"),
        ("M_X_abs_logit_difference", "Direct age + measurements model", "#B54C4C"),
    ):
        values = np.sort(pairs[column].to_numpy(float))
        cdf = np.arange(1, len(values) + 1) / len(values)
        ax.plot(values, cdf, color=color, linewidth=1.8, label=label)
    ax.set_xlabel("Absolute within-pair difference in mortality logit")
    ax.set_ylabel("Empirical cumulative proportion")
    ax.set_title("d  Endpoint-directed scores vary within AgeGap matches", loc="left", fontweight="bold")
    ax.legend(frameon=False, fontsize=7)
    fig.savefig(FIGURES / "nhanes_individual_agegap_contributions_v0_1.png", dpi=300, bbox_inches="tight")
    fig.savefig(FIGURES / "nhanes_individual_agegap_contributions_v0_1.svg", bbox_inches="tight")
    plt.close(fig)


def markdown_table(frame: pd.DataFrame, digits: int = 5) -> str:
    lines = ["| " + " | ".join(map(str, frame.columns)) + " |", "| " + " | ".join(["---"] * len(frame.columns)) + " |"]
    for values in frame.itertuples(index=False, name=None):
        rendered = []
        for value in values:
            if isinstance(value, (float, np.floating)):
                rendered.append("" if not np.isfinite(value) else f"{float(value):.{digits}g}")
            else:
                rendered.append(str(value))
        lines.append("| " + " | ".join(rendered) + " |")
    return "\n".join(lines)


def write_report(
    participants: pd.DataFrame,
    pairs: pd.DataFrame,
    summary: pd.DataFrame,
    selected: pd.DataFrame,
    audit: dict[str, Any],
) -> None:
    metrics = summary.set_index("metric")
    selected_view = selected[
        [
            "case",
            "pair_id",
            "age_years",
            "centered_agegap_1",
            "centered_agegap_2",
            "centered_agegap_abs_difference",
            "contribution_l2_distance_years",
            "contribution_cosine_similarity",
            "M_X_abs_logit_difference",
        ]
    ]
    text = f"""# NHANES individual AgeGap contribution analysis v0.1

Date: 31 August 2026

## Result

All **{len(participants):,}** participants received an out-of-fold raw AgeGap and
an exactly additive 50-feature decomposition of the conditional-centred AgeGap.
The maximum score reconstruction error was
`{audit['max_score_reconstruction_error']:.3g}` years.

The frozen outcome-blind algorithm retained **{len(pairs):,}** non-reusing pairs
with identical outer fold, chronological age, sex, race/ethnicity and survey
cycle and centred-AgeGap differences no greater than 0.25 years. Their median
AgeGap difference was
**{metrics.loc['centered_agegap_abs_difference', 'median']:.3f} years**, while
the median 50-dimensional contribution-profile L2 distance was
**{metrics.loc['contribution_l2_distance_years', 'median']:.3f} years**
(95th percentile **{metrics.loc['contribution_l2_distance_years', 'q95']:.3f}**).
The median cosine similarity was
**{metrics.loc['contribution_cosine_similarity', 'median']:.3f}**, and the median
top-five absolute-contributor Jaccard overlap was
**{metrics.loc['top5_absolute_contributor_jaccard', 'median']:.3f}**.

Within these near-equal AgeGap pairs, the median absolute difference in the
direct age-plus-measurements mortality logit was
**{metrics.loc['M_X_abs_logit_difference', 'median']:.3f}**, versus
**{metrics.loc['M_AGE_GAP_abs_logit_difference', 'median']:.3f}** for the
age-plus-AgeGap endpoint model. These are cross-fitted model-output contrasts,
not calibrated individual effects.

## Predeclared display cases

{markdown_table(selected_view)}

## Pair-distribution summary

{markdown_table(summary)}

## Methods

For each existing outer fold, the frozen Ridge age decoder was reloaded and its
stored held-out scores were reproduced. A fold-local multivariate Ridge model
estimated `E(X|A,W)` from the outer-training participants using a five-knot cubic
age spline and one-hot sex, race/ethnicity and survey cycle. Its penalty was
selected inside the outer training set by three-fold age-stratified measurement
reconstruction error. Every tuning model and final residualizer was persisted,
reloaded and verified.

For decoder measurement coefficient `beta_kj` and decoder scaler `sigma_kj`, the
participant contribution was

`c_ij = beta_kj (X_ij - E_hat_k[X_j|A_i,W_i]) / sigma_kj`.

Thus `sum_j c_ij = S_i-E_hat_k(S|A_i,W_i)`, which is the conditional-centred
AgeGap. The raw field-standard AgeGap `S_i-A_i` is stored separately. Because
pairs are exact on age, background and outer fold, their raw-gap and centred-gap
differences are equal up to numerical precision.

Pairing was outcome blind and did not use contribution distance or endpoint
scores. Within exact strata, adjacent centred-gap edges were ordered by gap
difference and greedily retained without participant reuse. The two display
cases follow the frozen median/95th-percentile rule among same-sign pairs with
mean absolute centred AgeGap of at least two years.

## Interpretation boundary

The result demonstrates the many-to-one geometry of this fitted age-decoder
coordinate: near-identical scalar AgeGaps can be generated by different measured
profiles. Contributions explain the decoder calculation; they do not identify
causal ageing mechanisms, organ ages, or biological-age components. The endpoint
comparison is specific to five-year all-cause mortality, this population, this
measurement panel and the existing fitted models.

## Persistence audit

- analysis signature: `{audit['analysis_signature']}`
- residualizer artifacts: {audit['residualizer_artifact_count']}/{audit['expected_residualizer_artifact_count']}
- tuning estimators persisted: {audit['tuning_estimator_count']}
- newly fitted/reused residualizers: {audit['newly_fitted_residualizer_count']}/{audit['reused_residualizer_count']}
- maximum residualizer reload error: `{audit['max_residualizer_reload_error']:.3g}`
- maximum frozen-decoder prediction error: `{audit['max_decoder_prediction_error']:.3g}`
- maximum contribution additivity error: `{audit['max_centered_gap_identity_error']:.3g}`

The complete participant table, long contribution matrix, matched pairs,
selected-case source data, fold coefficients, model artifacts and manifests are
in this output directory.
"""
    (OUT / "REPORT.md").write_text(text, encoding="utf-8")


def write_artifact_manifest() -> None:
    rows = []
    for path in sorted(OUT.rglob("*")):
        if path.is_file() and path.name != "artifact_manifest.tsv":
            rows.append({"path": str(path.relative_to(ROOT)), "bytes": path.stat().st_size, "sha256": sha256(path)})
    pd.DataFrame(rows).to_csv(OUT / "artifact_manifest.tsv", sep="\t", index=False)


def main() -> int:
    configure_stdout()
    OUT.mkdir(parents=True, exist_ok=True)
    MODELS.mkdir(parents=True, exist_ok=True)
    FIGURES.mkdir(parents=True, exist_ok=True)
    required = {
        "analysis_ready_source": FRAME_PATH,
        "participant_folds": FOLDS_PATH,
        "ordered_feature_tiers": TIERS_PATH,
        "decoder_oof_predictions": DECODER_OOF_PATH,
        "decoder_model_audit": DECODER_AUDIT_PATH,
        "decoder_analysis_signature": DECODER_SIGNATURE_PATH,
        "gap_endpoint_predictions": GAP_OOF_PATH,
        "gap_model_audit": GAP_AUDIT_PATH,
        "frozen_specification": SPEC_PATH,
        "analysis_script": Path(__file__),
        "environment_policy": ENV_POLICY,
        "environment_manifest": ENV_MANIFEST,
        "environment_lock": ENV_LOCK,
    }
    for path in required.values():
        if not path.exists():
            raise RuntimeError(f"Required input missing: {path}")
    for outer_fold in range(N_OUTER_FOLDS):
        required[f"decoder_outer_fold_{outer_fold}"] = DECODER / "models" / f"outer_fold_{outer_fold:02d}.joblib"
    input_manifest = pd.DataFrame(
        [
            {"role": role, "path": str(path.relative_to(ROOT)), "bytes": path.stat().st_size, "sha256": sha256(path)}
            for role, path in required.items()
        ]
    )
    signature_payload = {
        "analysis_version": ANALYSIS_VERSION,
        "schema_version": SCHEMA_VERSION,
        "inputs": input_manifest[["role", "sha256"]].to_dict(orient="records"),
        "features": "frozen_S3_50_measurements",
        "outer_folds": N_OUTER_FOLDS,
        "conditional_model": "multioutput_ridge_on_onehot_W_plus_5knot_cubic_age_spline",
        "alpha_grid": ALPHA_GRID,
        "inner_folds": N_INNER_FOLDS,
        "inner_seed": INNER_SEED,
        "match_caliper_years": MATCH_CALIPER_YEARS,
        "matching": "exact_strata_adjacent_edge_greedy_without_reuse",
    }
    analysis_signature = stable_hash(signature_payload)
    signature_path = OUT / "analysis_signature.json"
    if signature_path.exists():
        existing = json.loads(signature_path.read_text(encoding="utf-8"))
        if existing.get("analysis_signature") != analysis_signature:
            raise RuntimeError("Existing output has incompatible analysis signature")
    else:
        write_json(signature_path, {"analysis_signature": analysis_signature, "signature_payload": signature_payload})
    input_manifest.to_csv(OUT / "input_manifest.tsv", sep="\t", index=False)

    frame, features, decoder_oof, endpoint = load_and_gate()
    dictionary = pd.DataFrame(
        [
            {
                "feature_order": i + 1,
                "feature": feature,
                "display_feature": display_name(feature),
                "domain": DOMAIN_MAP.get(feature.removeprefix("T__"), "other"),
            }
            for i, feature in enumerate(features)
        ]
    )
    dictionary.to_csv(OUT / "feature_dictionary.tsv", sep="\t", index=False)
    frame[["row_id", "SEQN", "outer_fold", "age_years", *BACKGROUND, *features]].to_csv(
        OUT / "analysis_ready_interface.tsv.gz", sep="\t", index=False, compression="gzip"
    )

    participants, contributions, coefficients, model_index, tuning, audit = calculate_contributions(
        frame, features, decoder_oof, endpoint, analysis_signature
    )
    participants.to_csv(OUT / "participant_agegap_profiles.tsv.gz", sep="\t", index=False, compression="gzip")
    contributions.to_csv(OUT / "participant_feature_contributions.tsv.gz", sep="\t", index=False, compression="gzip")
    coefficients.to_csv(OUT / "fold_decoder_coefficients.tsv", sep="\t", index=False)
    model_index.to_csv(OUT / "model_artifact_index.tsv", sep="\t", index=False)
    tuning.to_csv(OUT / "conditional_model_inner_tuning.tsv", sep="\t", index=False)

    matrix = contributions.pivot(index="row_id", columns="feature_order", values="contribution_years").reindex(participants["row_id"]).to_numpy(float)
    pairs = build_pairs(participants, matrix)
    pair_summary = summarize_pairs(pairs)
    selected, selected_contributions = select_display_pairs(pairs, contributions)
    pairs.to_csv(OUT / "matched_pairs.tsv.gz", sep="\t", index=False, compression="gzip")
    pair_summary.to_csv(OUT / "matched_pair_distribution_summary.tsv", sep="\t", index=False)
    selected.to_csv(OUT / "selected_display_pairs.tsv", sep="\t", index=False)
    selected_contributions.to_csv(OUT / "selected_pair_contributions.tsv", sep="\t", index=False)

    audit.update(
        {
            "matched_pair_count": len(pairs),
            "matched_unique_participant_count": int(pd.unique(pairs[["row_id_1", "row_id_2"]].to_numpy().ravel()).size),
            "match_caliper_years": MATCH_CALIPER_YEARS,
            "max_matched_raw_vs_centered_gap_difference_error": float(
                np.max(np.abs(pairs["raw_agegap_abs_difference"] - pairs["centered_agegap_abs_difference"]))
            ),
            "software": {
                "python": sys.version,
                "python_executable": sys.executable,
                "platform": platform.platform(),
                "numpy": np.__version__,
                "pandas": pd.__version__,
                "scikit_learn": sklearn.__version__,
                "joblib": joblib.__version__,
                "matplotlib": matplotlib.__version__,
            },
        }
    )
    if (
        audit["residualizer_artifact_count"] != audit["expected_residualizer_artifact_count"]
        or audit["max_residualizer_reload_error"] > PARITY_TOLERANCE
        or audit["max_decoder_prediction_error"] > PARITY_TOLERANCE
        or audit["max_score_reconstruction_error"] > ADDITIVITY_TOLERANCE
        or not audit["all_residualizer_hashes_present"]
    ):
        audit["status"] = "FAIL"
    write_json(OUT / "model_artifact_audit.json", audit)
    if audit["status"] != "PASS":
        raise RuntimeError("STOP_DATA_OR_ARTIFACT_FAIL: final audit is not PASS")
    make_figure(pairs, selected, selected_contributions)
    write_report(participants, pairs, pair_summary, selected, audit)
    write_artifact_manifest()
    print(pair_summary.to_string(index=False), flush=True)
    print(selected[["case", "pair_id", "centered_agegap_1", "centered_agegap_2", "contribution_l2_distance_years", "M_X_abs_logit_difference"]].to_string(index=False), flush=True)
    print(f"Wrote {OUT}", flush=True)
    return 0


if __name__ == "__main__":
    raise SystemExit(main())
```
