## Supplementary_Software_1 for "Inferential boundaries of age prediction: why prediction does not establish biological age measurement": analyze_three_variable_age_state_task_map_probe_v0_1.py

```
"""Model-backed task map for the frozen three-variable age/state construction.

This script reuses the 500 frozen cross-sectional rows. It does not regenerate
people or interpret the ordered age coordinate as within-person follow-up.
Every fitted outer-fold pipeline is persisted and immediately reloaded for
elementwise held-out prediction parity.
"""

from __future__ import annotations

import gzip
import hashlib
import json
import math
import platform
import shutil
import sys
from dataclasses import asdict, dataclass
from pathlib import Path
from typing import Any

import joblib
import matplotlib
import matplotlib.pyplot as plt
import numpy as np
import pandas as pd
import sklearn
from sklearn.compose import ColumnTransformer
from sklearn.linear_model import LogisticRegression, Ridge
from sklearn.metrics import (
    brier_score_loss,
    log_loss,
    mean_absolute_error,
    mean_squared_error,
    r2_score,
    roc_auc_score,
)
from sklearn.model_selection import StratifiedKFold
from sklearn.pipeline import Pipeline
from sklearn.preprocessing import SplineTransformer, StandardScaler


ROOT = Path(__file__).resolve().parents[1]
SOURCE_PATH = ROOT / "outputs" / "three_variable_age_state_geometry_v0_1" / "simulated_people.tsv.gz"
PUBLIC_SOURCE_PATH = (
    ROOT
    / "outputs"
    / "v6_2_extended_data_figure_sync_v0_1"
    / "extended_data_figure_1"
    / "source_data"
    / "panel_a_x1_x3_geometry.tsv"
)
OUTPUT_DIR = ROOT / "outputs" / "three_variable_age_state_task_map_probe_v0_1"
MODEL_DIR = OUTPUT_DIR / "model_artifacts"
FIGURE_DIR = OUTPUT_DIR / "figures"
SOURCE_DATA_DIR = OUTPUT_DIR / "source_data"

EXPECTED_SOURCE_SHA256 = "55cf8290cbd3acf9e827428df0751361c4110eaa5f2650188ac946ebc84cae49"
ANALYSIS_VERSION = "three_variable_age_state_task_map_probe_v0_1"
FOLD_SEED = 20260827
N_FOLDS = 5
PARITY_TOLERANCE = 1e-12

ID_COL = "person_id"
AGE_COL = "age"
STATE_COL = "separation_state"
FEATURES = {
    "X1": "x_age_linear",
    "X2": "x_age_orthogonal_state",
    "X3": "x_age_nonlinear_state",
}
AGE_ARMS = {
    "X1": [FEATURES["X1"]],
    "X2": [FEATURES["X2"]],
    "X3": [FEATURES["X3"]],
    "X1-X3": list(FEATURES.values()),
}
STATE_ARMS = {
    "Age only": [],
    "X1": [FEATURES["X1"]],
    "X2": [FEATURES["X2"]],
    "X3": [FEATURES["X3"]],
    "X1-X3": list(FEATURES.values()),
}

BLUE = "#2F6690"
ORANGE = "#C96B2C"
INK = "#1F2933"
MUTED = "#66727D"
GRID = "#D7DDE2"


@dataclass(frozen=True)
class ModelRecord:
    task: str
    arm: str
    fold: int
    model_path: str
    signature_path: str
    model_sha256: str
    signature_sha256: str
    model_bytes: int
    parity_max_abs_error: float
    reused: bool


def sha256_file(path: Path) -> str:
    digest = hashlib.sha256()
    with path.open("rb") as handle:
        for block in iter(lambda: handle.read(1024 * 1024), b""):
            digest.update(block)
    return digest.hexdigest()


def stable_hash(values: list[str]) -> str:
    return hashlib.sha256("\n".join(values).encode("utf-8")).hexdigest()


def write_json(path: Path, payload: Any) -> None:
    path.parent.mkdir(parents=True, exist_ok=True)
    path.write_text(json.dumps(payload, indent=2, ensure_ascii=False), encoding="utf-8")


def canonical_json_hash(payload: Any) -> str:
    encoded = json.dumps(payload, sort_keys=True, separators=(",", ":"), ensure_ascii=False).encode("utf-8")
    return hashlib.sha256(encoded).hexdigest()


def load_source() -> pd.DataFrame:
    if sha256_file(SOURCE_PATH) != EXPECTED_SOURCE_SHA256:
        raise RuntimeError("Frozen X1-X3 source hash does not match the declared contract")
    frame = pd.read_csv(SOURCE_PATH, sep="\t", compression="gzip")
    required = [ID_COL, AGE_COL, STATE_COL, *FEATURES.values()]
    if list(frame.columns) != required:
        raise RuntimeError(f"Unexpected source schema/order: {list(frame.columns)}")
    if len(frame) != 500 or frame[ID_COL].duplicated().any():
        raise RuntimeError("Expected exactly 500 unique synthetic people")
    if not frame[STATE_COL].isin([0, 1]).all():
        raise RuntimeError("Declared state must be binary")
    if frame.isna().any().any():
        raise RuntimeError("Frozen interface unexpectedly contains missing values")

    public = pd.read_csv(PUBLIC_SOURCE_PATH, sep="\t").rename(
        columns={
            "chronological_age_years": AGE_COL,
            "declared_state": STATE_COL,
            "X1_age_readable": FEATURES["X1"],
            "X2_age_orthogonal_state": FEATURES["X2"],
            "X3_age_nonlinear_state": FEATURES["X3"],
        }
    )
    public = public[required]
    if public[ID_COL].tolist() != frame[ID_COL].tolist():
        raise RuntimeError("Public Source Data participant order differs from the frozen source")
    numeric = [AGE_COL, STATE_COL, *FEATURES.values()]
    max_source_error = float(np.max(np.abs(public[numeric].to_numpy(float) - frame[numeric].to_numpy(float))))
    if max_source_error > 1e-8:
        raise RuntimeError(f"Public Source Data parity failed: {max_source_error}")
    return frame


def make_folds(frame: pd.DataFrame) -> pd.DataFrame:
    age_group = pd.qcut(frame[AGE_COL], q=5, labels=False, duplicates="raise").astype(int)
    stratum = frame[STATE_COL].astype(str) + "_" + age_group.astype(str)
    splitter = StratifiedKFold(n_splits=N_FOLDS, shuffle=True, random_state=FOLD_SEED)
    fold_id = np.zeros(len(frame), dtype=int)
    for fold, (_, test_index) in enumerate(splitter.split(np.zeros(len(frame)), stratum), start=1):
        fold_id[test_index] = fold
    if sorted(np.unique(fold_id).tolist()) != list(range(1, N_FOLDS + 1)):
        raise RuntimeError("Fold assignment failed")
    return pd.DataFrame(
        {
            ID_COL: frame[ID_COL],
            "fold_id": fold_id,
            "age_quintile": age_group,
            "stratum": stratum,
        }
    )


def age_pipeline() -> Pipeline:
    return Pipeline(
        steps=[
            (
                "spline",
                SplineTransformer(
                    n_knots=5,
                    degree=3,
                    include_bias=False,
                    knots="uniform",
                    extrapolation="linear",
                ),
            ),
            ("scale", StandardScaler()),
            ("ridge", Ridge(alpha=1.0)),
        ]
    )


def state_pipeline(feature_columns: list[str]) -> Pipeline:
    transformers: list[tuple[str, Any, list[str]]] = [
        (
            "age",
            Pipeline(
                steps=[
                    (
                        "spline",
                        SplineTransformer(
                            n_knots=5,
                            degree=3,
                            include_bias=False,
                            knots="uniform",
                            extrapolation="linear",
                        ),
                    ),
                    ("scale", StandardScaler()),
                ]
            ),
            [AGE_COL],
        )
    ]
    if feature_columns:
        transformers.append(("measurements", StandardScaler(), feature_columns))
    preprocessor = ColumnTransformer(transformers=transformers, remainder="drop", verbose_feature_names_out=True)
    return Pipeline(
        steps=[
            ("preprocess", preprocessor),
            (
                "logistic",
                LogisticRegression(
                    C=1.0,
                    penalty="l2",
                    solver="lbfgs",
                    max_iter=2000,
                    random_state=FOLD_SEED,
                ),
            ),
        ]
    )


def model_signature(
    *,
    analysis_signature_hash: str,
    task: str,
    arm: str,
    fold: int,
    features: list[str],
    train_ids: list[str],
    test_ids: list[str],
) -> dict[str, Any]:
    settings: dict[str, Any]
    if task == "age":
        settings = {
            "spline": {"n_knots": 5, "degree": 3, "knots": "uniform", "extrapolation": "linear"},
            "scaler": "StandardScaler",
            "estimator": "Ridge",
            "alpha": 1.0,
        }
    else:
        settings = {
            "age_spline": {"n_knots": 5, "degree": 3, "knots": "uniform", "extrapolation": "linear"},
            "measurement_scaler": "StandardScaler",
            "estimator": "LogisticRegression",
            "C": 1.0,
            "penalty": "l2",
            "solver": "lbfgs",
            "max_iter": 2000,
        }
    return {
        "analysis_version": ANALYSIS_VERSION,
        "analysis_signature_hash": analysis_signature_hash,
        "source_sha256": EXPECTED_SOURCE_SHA256,
        "task": task,
        "arm": arm,
        "fold": fold,
        "features": features,
        "train_ids_sha256": stable_hash(train_ids),
        "test_ids_sha256": stable_hash(test_ids),
        "n_train": len(train_ids),
        "n_test": len(test_ids),
        "settings": settings,
        "fold_seed": FOLD_SEED,
        "sklearn_version": sklearn.__version__,
        "python_version": platform.python_version(),
    }


def extract_coefficients(model: Pipeline, task: str, arm: str, fold: int) -> pd.DataFrame:
    if task == "age":
        transformer = model.named_steps["spline"]
        terms = transformer.get_feature_names_out().tolist()
        estimator = model.named_steps["ridge"]
        coefficients = np.asarray(estimator.coef_).reshape(-1)
        intercept = float(estimator.intercept_)
    else:
        transformer = model.named_steps["preprocess"]
        terms = transformer.get_feature_names_out().tolist()
        estimator = model.named_steps["logistic"]
        coefficients = np.asarray(estimator.coef_).reshape(-1)
        intercept = float(np.asarray(estimator.intercept_).reshape(-1)[0])
    rows = pd.DataFrame(
        {
            "task": task,
            "arm": arm,
            "fold": fold,
            "term": terms,
            "coefficient": coefficients,
            "intercept": intercept,
        }
    )
    return rows


def fit_or_load_model(
    *,
    task: str,
    arm: str,
    fold: int,
    features: list[str],
    train: pd.DataFrame,
    test: pd.DataFrame,
    analysis_signature_hash: str,
    reuse: bool,
) -> tuple[np.ndarray, ModelRecord, pd.DataFrame]:
    arm_slug = arm.lower().replace(" ", "_").replace("-", "_")
    artifact_dir = MODEL_DIR / task / arm_slug
    artifact_dir.mkdir(parents=True, exist_ok=True)
    model_path = artifact_dir / f"fold_{fold}.joblib"
    signature_path = artifact_dir / f"fold_{fold}.signature.json"
    expected_signature = model_signature(
        analysis_signature_hash=analysis_signature_hash,
        task=task,
        arm=arm,
        fold=fold,
        features=features,
        train_ids=train[ID_COL].tolist(),
        test_ids=test[ID_COL].tolist(),
    )

    if reuse:
        if not model_path.exists() or not signature_path.exists():
            raise RuntimeError(f"Missing persisted model during reuse: {task}/{arm}/fold {fold}")
        stored_signature = json.loads(signature_path.read_text(encoding="utf-8"))
        if stored_signature != expected_signature:
            raise RuntimeError(f"Model signature mismatch: {task}/{arm}/fold {fold}")
        model = joblib.load(model_path)
        fitted_prediction: np.ndarray | None = None
    else:
        if model_path.exists() or signature_path.exists():
            raise RuntimeError(f"Refusing to overwrite partial model artifact: {model_path}")
        if task == "age":
            model = age_pipeline()
            model.fit(train[features], train[AGE_COL])
            fitted_prediction = np.asarray(model.predict(test[features]), dtype=float)
        else:
            model = state_pipeline(features)
            model.fit(train[[AGE_COL, *features]], train[STATE_COL])
            fitted_prediction = np.asarray(model.predict_proba(test[[AGE_COL, *features]])[:, 1], dtype=float)
        joblib.dump(model, model_path, compress=3)
        write_json(signature_path, expected_signature)

    reloaded = joblib.load(model_path)
    if task == "age":
        reloaded_prediction = np.asarray(reloaded.predict(test[features]), dtype=float)
    else:
        reloaded_prediction = np.asarray(reloaded.predict_proba(test[[AGE_COL, *features]])[:, 1], dtype=float)
    if fitted_prediction is None:
        fitted_prediction = reloaded_prediction.copy()
    parity = float(np.max(np.abs(fitted_prediction - reloaded_prediction)))
    if not math.isfinite(parity) or parity > PARITY_TOLERANCE:
        raise RuntimeError(f"Reload prediction parity failed: {task}/{arm}/fold {fold}: {parity}")
    record = ModelRecord(
        task=task,
        arm=arm,
        fold=fold,
        model_path=str(model_path.relative_to(ROOT)),
        signature_path=str(signature_path.relative_to(ROOT)),
        model_sha256=sha256_file(model_path),
        signature_sha256=sha256_file(signature_path),
        model_bytes=model_path.stat().st_size,
        parity_max_abs_error=parity,
        reused=reuse,
    )
    return reloaded_prediction, record, extract_coefficients(reloaded, task, arm, fold)


def age_metrics(y_true: np.ndarray, y_pred: np.ndarray) -> dict[str, float]:
    return {
        "r2": float(r2_score(y_true, y_pred)),
        "rmse_years": float(np.sqrt(mean_squared_error(y_true, y_pred))),
        "mae_years": float(mean_absolute_error(y_true, y_pred)),
    }


def state_metrics(y_true: np.ndarray, probability: np.ndarray) -> dict[str, float]:
    return {
        "log_loss": float(log_loss(y_true, probability, labels=[0, 1])),
        "roc_auc": float(roc_auc_score(y_true, probability)),
        "brier_score": float(brier_score_loss(y_true, probability)),
    }


def run_models(
    frame: pd.DataFrame,
    folds: pd.DataFrame,
    analysis_signature_hash: str,
    reuse: bool,
) -> tuple[pd.DataFrame, pd.DataFrame, pd.DataFrame, pd.DataFrame, list[ModelRecord], pd.DataFrame]:
    merged = frame.merge(folds[[ID_COL, "fold_id"]], on=ID_COL, validate="one_to_one")
    predictions = merged[[ID_COL, AGE_COL, STATE_COL, "fold_id"]].copy()
    model_records: list[ModelRecord] = []
    coefficient_tables: list[pd.DataFrame] = []
    fold_rows: list[dict[str, Any]] = []

    for arm, features in AGE_ARMS.items():
        prediction = np.full(len(merged), np.nan)
        for fold in range(1, N_FOLDS + 1):
            train = merged.loc[merged["fold_id"] != fold].copy()
            test = merged.loc[merged["fold_id"] == fold].copy()
            values, record, coefficients = fit_or_load_model(
                task="age",
                arm=arm,
                fold=fold,
                features=features,
                train=train,
                test=test,
                analysis_signature_hash=analysis_signature_hash,
                reuse=reuse,
            )
            prediction[test.index] = values
            model_records.append(record)
            coefficient_tables.append(coefficients)
            fold_rows.append({"task": "age", "arm": arm, "fold": fold, **age_metrics(test[AGE_COL], values)})
        if np.isnan(prediction).any():
            raise RuntimeError(f"Incomplete age predictions for {arm}")
        predictions[f"age_pred_{arm}"] = prediction

    for arm, features in STATE_ARMS.items():
        probability = np.full(len(merged), np.nan)
        for fold in range(1, N_FOLDS + 1):
            train = merged.loc[merged["fold_id"] != fold].copy()
            test = merged.loc[merged["fold_id"] == fold].copy()
            values, record, coefficients = fit_or_load_model(
                task="state",
                arm=arm,
                fold=fold,
                features=features,
                train=train,
                test=test,
                analysis_signature_hash=analysis_signature_hash,
                reuse=reuse,
            )
            probability[test.index] = values
            model_records.append(record)
            coefficient_tables.append(coefficients)
            fold_rows.append({"task": "state", "arm": arm, "fold": fold, **state_metrics(test[STATE_COL], values)})
        if np.isnan(probability).any():
            raise RuntimeError(f"Incomplete state predictions for {arm}")
        predictions[f"state_prob_{arm}"] = probability

    pooled_rows: list[dict[str, Any]] = []
    for arm in AGE_ARMS:
        pooled_rows.append(
            {
                "task": "age",
                "arm": arm,
                **age_metrics(merged[AGE_COL].to_numpy(), predictions[f"age_pred_{arm}"].to_numpy()),
            }
        )
    state_pooled: dict[str, dict[str, float]] = {}
    for arm in STATE_ARMS:
        state_pooled[arm] = state_metrics(
            merged[STATE_COL].to_numpy(), predictions[f"state_prob_{arm}"].to_numpy()
        )
    baseline_loss = state_pooled["Age only"]["log_loss"]
    for arm, metrics in state_pooled.items():
        pooled_rows.append(
            {
                "task": "state",
                "arm": arm,
                **metrics,
                "log_loss_gain_beyond_age": float(baseline_loss - metrics["log_loss"]),
            }
        )

    fold_metrics = pd.DataFrame(fold_rows)
    state_baseline_by_fold = (
        fold_metrics.query("task == 'state' and arm == 'Age only'")
        .set_index("fold")["log_loss"]
        .to_dict()
    )
    state_mask = fold_metrics["task"].eq("state")
    fold_metrics.loc[state_mask, "log_loss_gain_beyond_age"] = fold_metrics.loc[state_mask].apply(
        lambda row: state_baseline_by_fold[int(row["fold"])] - float(row["log_loss"]), axis=1
    )
    pooled_metrics = pd.DataFrame(pooled_rows)
    coefficients = pd.concat(coefficient_tables, ignore_index=True)

    task_rows = []
    for arm in AGE_ARMS:
        age_row = pooled_metrics.query("task == 'age' and arm == @arm").iloc[0]
        state_row = pooled_metrics.query("task == 'state' and arm == @arm").iloc[0]
        task_rows.append(
            {
                "arm": arm,
                "age_oof_r2": float(age_row["r2"]),
                "age_oof_rmse_years": float(age_row["rmse_years"]),
                "state_oof_log_loss_gain_beyond_age": float(state_row["log_loss_gain_beyond_age"]),
                "state_oof_roc_auc": float(state_row["roc_auc"]),
                "state_oof_log_loss": float(state_row["log_loss"]),
            }
        )
    task_map = pd.DataFrame(task_rows)
    return predictions, pooled_metrics, fold_metrics, task_map, model_records, coefficients


def make_age_coordinate_source(frame: pd.DataFrame) -> tuple[pd.DataFrame, pd.DataFrame]:
    long_rows = []
    for label, column in FEATURES.items():
        part = frame[[ID_COL, AGE_COL, STATE_COL, column]].rename(columns={column: "value"})
        part.insert(3, "feature", label)
        long_rows.append(part)
    points = pd.concat(long_rows, ignore_index=True)
    edges = np.linspace(20.0, 80.0, 13)
    points["age_bin"] = pd.cut(points[AGE_COL], bins=edges, include_lowest=True, right=False)
    binned = (
        points.groupby(["feature", STATE_COL, "age_bin"], observed=True)
        .agg(
            n=("value", "size"),
            age_mean=(AGE_COL, "mean"),
            value_mean=("value", "mean"),
            value_sd=("value", "std"),
        )
        .reset_index()
    )
    binned["age_bin_left"] = binned["age_bin"].map(lambda interval: float(interval.left))
    binned["age_bin_right"] = binned["age_bin"].map(lambda interval: float(interval.right))
    binned = binned.drop(columns="age_bin")
    return points, binned


def plot_figure(points: pd.DataFrame, binned: pd.DataFrame, task_map: pd.DataFrame) -> None:
    matplotlib.rcParams.update(
        {
            "font.family": "DejaVu Sans",
            "axes.edgecolor": INK,
            "axes.labelcolor": INK,
            "xtick.color": INK,
            "ytick.color": INK,
            "text.color": INK,
            "axes.titleweight": "bold",
        }
    )
    fig, axes = plt.subplots(2, 2, figsize=(12.4, 9.2), constrained_layout=False)
    plt.subplots_adjust(left=0.08, right=0.97, top=0.84, bottom=0.12, hspace=0.34, wspace=0.26)
    feature_titles = {
        "X1": "X1: noisy age-readable channel",
        "X2": "X2: age-orthogonal state channel",
        "X3": "X3: nonlinear age–state mixture",
    }
    state_style = {
        0: {"color": BLUE, "marker": "o", "linestyle": "-", "label": "Declared state 0"},
        1: {"color": ORANGE, "marker": "^", "linestyle": "--", "label": "Declared state 1"},
    }
    for panel, (axis, feature) in enumerate(zip(axes.flat[:3], ["X1", "X2", "X3"], strict=True)):
        feature_points = points.loc[points["feature"] == feature]
        feature_bins = binned.loc[binned["feature"] == feature]
        for state, style in state_style.items():
            subset = feature_points.loc[feature_points[STATE_COL] == state]
            if state == 0:
                axis.scatter(
                    subset[AGE_COL],
                    subset["value"],
                    s=17,
                    alpha=0.30,
                    marker=style["marker"],
                    facecolors=style["color"],
                    edgecolors="none",
                    rasterized=True,
                )
            else:
                axis.scatter(
                    subset[AGE_COL],
                    subset["value"],
                    s=20,
                    alpha=0.42,
                    marker=style["marker"],
                    facecolors="none",
                    edgecolors=style["color"],
                    linewidths=0.65,
                    rasterized=True,
                )
            means = feature_bins.loc[feature_bins[STATE_COL] == state].sort_values("age_mean")
            axis.plot(
                means["age_mean"],
                means["value_mean"],
                color=style["color"],
                linestyle=style["linestyle"],
                marker=style["marker"],
                markersize=4.5,
                linewidth=2.0,
                markerfacecolor=style["color"] if state == 0 else "white",
                markeredgecolor=style["color"],
                label=style["label"],
            )
        axis.axhline(0, color=GRID, linewidth=0.9, zorder=0)
        axis.grid(axis="y", color=GRID, linewidth=0.7, alpha=0.65)
        axis.set_xlim(19, 81)
        axis.set_xlabel("Chronological age (years)")
        axis.set_ylabel("Standardized measurement")
        axis.set_title(f"{chr(97 + panel)}   {feature_titles[feature]}", loc="left", fontsize=12)
        axis.spines[["top", "right"]].set_visible(False)

    axis = axes.flat[3]
    marker_map = {"X1": "o", "X2": "s", "X3": "^", "X1-X3": "*"}
    color_map = {"X1": BLUE, "X2": ORANGE, "X3": MUTED, "X1-X3": INK}
    for _, row in task_map.iterrows():
        arm = str(row["arm"])
        axis.scatter(
            row["age_oof_r2"],
            row["state_oof_log_loss_gain_beyond_age"],
            s=145 if arm == "X1-X3" else 90,
            marker=marker_map[arm],
            facecolor=color_map[arm] if arm != "X2" else "white",
            edgecolor=color_map[arm],
            linewidth=1.6,
            zorder=3,
        )
        offset = {"X1": (7, -15), "X2": (7, 8), "X3": (7, 8), "X1-X3": (7, -16)}[arm]
        axis.annotate(arm, (row["age_oof_r2"], row["state_oof_log_loss_gain_beyond_age"]), xytext=offset,
                      textcoords="offset points", fontsize=10, fontweight="bold")
    axis.axhline(0, color=INK, linewidth=0.9)
    axis.axvline(0, color=INK, linewidth=0.9)
    x_values = task_map["age_oof_r2"].to_numpy(float)
    y_values = task_map["state_oof_log_loss_gain_beyond_age"].to_numpy(float)
    x_pad = max(0.05, 0.10 * (x_values.max() - x_values.min()))
    y_pad = max(0.004, 0.14 * (y_values.max() - y_values.min()))
    axis.set_xlim(min(-0.05, x_values.min() - x_pad), x_values.max() + x_pad)
    axis.set_ylim(min(-0.005, y_values.min() - y_pad), y_values.max() + y_pad)
    axis.grid(color=GRID, linewidth=0.7, alpha=0.65)
    axis.set_xlabel("Chronological-age decoding: pooled OOF R²")
    axis.set_ylabel("Same-age state decoding:\nOOF log-loss gain beyond age")
    axis.set_title("d   Fixed-model task map", loc="left", fontsize=12)
    axis.spines[["top", "right"]].set_visible(False)

    handles, labels = axes.flat[0].get_legend_handles_labels()
    fig.legend(handles[:2], labels[:2], loc="upper center", ncol=2, frameon=False, bbox_to_anchor=(0.5, 0.89))
    fig.suptitle("Age-coordinate geometry and task-specific model performance", fontsize=18, fontweight="bold", y=0.965)
    fig.text(
        0.5,
        0.918,
        "Fixed 500-row cross-sectional construction; age is an ordered coordinate, not within-person follow-up",
        ha="center",
        va="center",
        fontsize=10.5,
        color=MUTED,
    )
    fig.text(
        0.5,
        0.035,
        "Points in a–c are individuals; lines are fixed-bin state-specific means. OOF = five-fold out-of-fold. "
        "State gain = age-only log loss − age+measurement log loss. Descriptive construction; no population interval.",
        ha="center",
        va="center",
        fontsize=8.8,
        color=MUTED,
    )
    FIGURE_DIR.mkdir(parents=True, exist_ok=True)
    stem = FIGURE_DIR / "Figure_X123_age_coordinate_and_task_map"
    fig.savefig(stem.with_suffix(".png"), dpi=600, facecolor="white")
    fig.savefig(stem.with_suffix(".svg"), facecolor="white")
    fig.savefig(stem.with_suffix(".pdf"), facecolor="white")
    plt.close(fig)


def evaluate_gate(task_map: pd.DataFrame, model_records: list[ModelRecord]) -> tuple[dict[str, bool], str, str]:
    lookup = task_map.set_index("arm")
    gates = {
        "x1_age_r2_exceeds_x2": bool(lookup.loc["X1", "age_oof_r2"] > lookup.loc["X2", "age_oof_r2"]),
        "x2_state_gain_exceeds_x1": bool(
            lookup.loc["X2", "state_oof_log_loss_gain_beyond_age"]
            > lookup.loc["X1", "state_oof_log_loss_gain_beyond_age"]
        ),
        "x3_positive_age_r2": bool(lookup.loc["X3", "age_oof_r2"] > 0),
        "x3_positive_state_gain": bool(lookup.loc["X3", "state_oof_log_loss_gain_beyond_age"] > 0),
        "joint_positive_age_r2": bool(lookup.loc["X1-X3", "age_oof_r2"] > 0),
        "joint_positive_state_gain": bool(lookup.loc["X1-X3", "state_oof_log_loss_gain_beyond_age"] > 0),
        "all_45_models_persisted": len(model_records) == 45,
        "all_reload_parity_within_tolerance": all(
            record.parity_max_abs_error <= PARITY_TOLERANCE for record in model_records
        ),
    }
    if all(gates.values()):
        return gates, "PROMISING", "PROMOTE"
    return gates, "AMBIGUOUS", "STOP"


def write_reports(
    *,
    frame: pd.DataFrame,
    task_map: pd.DataFrame,
    pooled_metrics: pd.DataFrame,
    model_records: list[ModelRecord],
    gates: dict[str, bool],
    category: str,
    decision: str,
    reuse: bool,
) -> None:
    table = task_map.copy()
    for column in table.columns[1:]:
        table[column] = table[column].map(lambda value: f"{float(value):.6f}")
    report = f"""# Three-variable age–state task-map probe v0.1

## Verdict

**{category} / {decision}** under the single frozen comparison. The script ran in
`{'reuse' if reuse else 'fit-once'}` mode and did not regenerate the 500 source rows.

## Model-backed task map

{table.to_markdown(index=False)}

The horizontal model quantity is pooled five-fold out-of-fold chronological-age
R2. The vertical model quantity is pooled five-fold out-of-fold state log-loss
reduction relative to an age-only model. Positive state gain means that the
measurement contributed held-out information about the declared binary state
after the fixed age spline.

## Gate

"""
    for name, passed in gates.items():
        report += f"- `{name}`: `{'PASS' if passed else 'FAIL'}`\n"
    report += f"""

## Interpretation boundary

This fixed cross-sectional construction separates age readability, same-age
state information and their mixture under the declared models. It is not a
longitudinal analysis, human biological-age model, disease prediction,
population estimate or proof of the analytical theorem.

## Persistence

- Persisted fold-local pipelines: {len(model_records)}.
- Maximum immediate reload prediction error:
  {max(record.parity_max_abs_error for record in model_records):.3e}.
- Source rows: {len(frame)} unique synthetic people.
"""
    (OUTPUT_DIR / "RESULTS_REPORT.md").write_text(report, encoding="utf-8")

    legend = """# Figure legend

## English

**Figure X | Age-coordinate geometry and task-specific model performance in a transparent three-variable construction.**
**a–c,** Individual standardized values of X1, X2 and X3 across chronological age in the fixed 500-row cross-sectional construction. Colour and marker/fill distinguish the declared binary states; connected points are fixed-bin state-specific means and are not within-person trajectories. X1 was generated as a noisy age-readable channel without a designed state term, X2 as a state channel projected exactly off the realized linear age direction and X3 as a nonlinear age–state mixture. **d,** Pooled five-fold out-of-fold chronological-age R2 versus pooled out-of-fold state log-loss gain beyond an age-only model for fixed spline-ridge and spline-logistic pipelines. The construction is descriptive, supplies no population interval and is not a model of human ageing.

## 中文

**图 X｜透明三变量构造中的年龄坐标几何与任务特异模型表现。**
**a–c，** 固定 500 行横截面构造中，X1、X2 和 X3 的标准化数值沿时间年龄坐标的分布。颜色及点形/填充区分预先声明的二元状态；连线为固定年龄分箱内的状态特异均值，并非同一人的纵向轨迹。X1 被构造为不含预设状态项的噪声年龄可读通道，X2 为在已实现样本内精确投影掉线性年龄方向的状态通道，X3 为非线性年龄–状态混合通道。**d，** 固定 spline–ridge 与 spline–logistic pipeline 中，合并五折样本外时间年龄 R2 与相对 age-only 模型的样本外状态 log-loss 改善。该构造是描述性的，不提供总体区间，也不是人类衰老模型。
"""
    (OUTPUT_DIR / "FIGURE_LEGEND_BILINGUAL.md").write_text(legend, encoding="utf-8")

    readme = f"""# Three-variable age–state task-map probe v0.1

- Decision: `{category} / {decision}`
- Frozen input: `outputs/three_variable_age_state_geometry_v0_1/simulated_people.tsv.gz`
- Models: 45 persisted outer-fold pipelines with immediate reload parity
- Figure: `figures/Figure_X123_age_coordinate_and_task_map` in PNG, SVG and PDF
- Source Data: `source_data/`
- Canonical report: `RESULTS_REPORT.md`
- Current manuscript v6.7/v5.2 was not edited by this probe.
"""
    (OUTPUT_DIR / "README.md").write_text(readme, encoding="utf-8")


def artifact_manifest() -> pd.DataFrame:
    excluded = {"artifact_manifest.tsv"}
    rows = []
    for path in sorted(OUTPUT_DIR.rglob("*")):
        if not path.is_file() or path.name in excluded:
            continue
        rows.append(
            {
                "path": str(path.relative_to(ROOT)),
                "bytes": path.stat().st_size,
                "sha256": sha256_file(path),
            }
        )
    return pd.DataFrame(rows)


def main() -> None:
    OUTPUT_DIR.mkdir(parents=True, exist_ok=True)
    MODEL_DIR.mkdir(parents=True, exist_ok=True)
    FIGURE_DIR.mkdir(parents=True, exist_ok=True)
    SOURCE_DATA_DIR.mkdir(parents=True, exist_ok=True)

    frame = load_source()
    folds = make_folds(frame)
    script_sha = sha256_file(Path(__file__))
    analysis_signature = {
        "analysis_version": ANALYSIS_VERSION,
        "source_path": str(SOURCE_PATH.relative_to(ROOT)),
        "source_sha256": EXPECTED_SOURCE_SHA256,
        "public_source_path": str(PUBLIC_SOURCE_PATH.relative_to(ROOT)),
        "public_source_sha256": sha256_file(PUBLIC_SOURCE_PATH),
        "script_path": str(Path(__file__).relative_to(ROOT)),
        "script_sha256": script_sha,
        "n_people": len(frame),
        "fold_seed": FOLD_SEED,
        "n_folds": N_FOLDS,
        "age_arms": AGE_ARMS,
        "state_arms": STATE_ARMS,
        "parity_tolerance": PARITY_TOLERANCE,
        "python_version": platform.python_version(),
        "numpy_version": np.__version__,
        "pandas_version": pd.__version__,
        "sklearn_version": sklearn.__version__,
        "matplotlib_version": matplotlib.__version__,
    }
    signature_hash = canonical_json_hash(analysis_signature)
    signature_path = OUTPUT_DIR / "analysis_signature.json"
    reuse = signature_path.exists()
    if reuse:
        stored = json.loads(signature_path.read_text(encoding="utf-8"))
        if stored != analysis_signature:
            raise RuntimeError("Existing analysis signature is incompatible; use a new versioned output directory")
    elif any(MODEL_DIR.rglob("*.joblib")):
        raise RuntimeError("Partial model artifacts exist without an analysis signature; refusing to overwrite")

    predictions, pooled_metrics, fold_metrics, task_map, model_records, coefficients = run_models(
        frame, folds, signature_hash, reuse
    )
    points, binned = make_age_coordinate_source(frame)
    plot_figure(points, binned, task_map)
    gates, category, decision = evaluate_gate(task_map, model_records)

    analysis_ready = frame.merge(folds, on=ID_COL, validate="one_to_one")
    with gzip.open(OUTPUT_DIR / "analysis_ready_interface.tsv.gz", "wt", encoding="utf-8", newline="") as handle:
        analysis_ready.to_csv(handle, sep="\t", index=False, float_format="%.12g")
    analysis_ready[[ID_COL, AGE_COL, STATE_COL, "fold_id", "age_quintile", "stratum"]].to_csv(
        OUTPUT_DIR / "participant_metadata.tsv", sep="\t", index=False, float_format="%.12g"
    )
    folds.to_csv(OUTPUT_DIR / "fold_assignments.tsv", sep="\t", index=False)
    pd.DataFrame(
        [
            {"publication_name": label, "source_column": column, "ordered_position": index + 1}
            for index, (label, column) in enumerate(FEATURES.items())
        ]
    ).to_csv(OUTPUT_DIR / "feature_manifest.tsv", sep="\t", index=False)
    with gzip.open(OUTPUT_DIR / "oof_predictions.tsv.gz", "wt", encoding="utf-8", newline="") as handle:
        predictions.to_csv(handle, sep="\t", index=False, float_format="%.12g")
    pooled_metrics.to_csv(OUTPUT_DIR / "pooled_metrics.tsv", sep="\t", index=False, float_format="%.12g")
    fold_metrics.to_csv(OUTPUT_DIR / "fold_metrics.tsv", sep="\t", index=False, float_format="%.12g")
    task_map.to_csv(OUTPUT_DIR / "task_map.tsv", sep="\t", index=False, float_format="%.12g")
    coefficients.to_csv(OUTPUT_DIR / "model_coefficients.tsv", sep="\t", index=False, float_format="%.12g")
    pd.DataFrame([asdict(record) for record in model_records]).to_csv(
        OUTPUT_DIR / "model_artifact_manifest.tsv", sep="\t", index=False, float_format="%.12g"
    )

    with gzip.open(SOURCE_DATA_DIR / "panels_a_c_age_coordinate_points.tsv.gz", "wt", encoding="utf-8", newline="") as handle:
        points.to_csv(handle, sep="\t", index=False, float_format="%.12g")
    binned.to_csv(SOURCE_DATA_DIR / "panels_a_c_binned_means.tsv", sep="\t", index=False, float_format="%.12g")
    task_map.to_csv(SOURCE_DATA_DIR / "panel_d_model_task_map.tsv", sep="\t", index=False, float_format="%.12g")

    environment = {
        "python_executable": sys.executable,
        "python_version": platform.python_version(),
        "platform": platform.platform(),
        "numpy": np.__version__,
        "pandas": pd.__version__,
        "scikit_learn": sklearn.__version__,
        "matplotlib": matplotlib.__version__,
        "joblib": joblib.__version__,
    }
    write_json(OUTPUT_DIR / "environment.json", environment)
    write_json(
        OUTPUT_DIR / "decision.json",
        {
            "analysis_version": ANALYSIS_VERSION,
            "category": category,
            "decision": decision,
            "gates": gates,
            "claim": "Fixed models separate age readability, same-age state information and their mixture in the frozen construction.",
            "forbidden_claims": [
                "longitudinal change",
                "human biological age",
                "disease prediction",
                "population inference",
                "proof of the analytical theorem",
            ],
        },
    )
    write_json(
        OUTPUT_DIR / "audit.json",
        {
            "status": "PASS" if all(gates.values()) else "FAIL_CLOSED",
            "source_sha256": sha256_file(SOURCE_PATH),
            "public_source_sha256": sha256_file(PUBLIC_SOURCE_PATH),
            "n_rows": len(frame),
            "n_unique_people": int(frame[ID_COL].nunique()),
            "n_models": len(model_records),
            "max_reload_prediction_error": max(record.parity_max_abs_error for record in model_records),
            "parity_tolerance": PARITY_TOLERANCE,
            "run_mode": "reuse" if reuse else "fit-once",
            "all_gates_passed": all(gates.values()),
        },
    )
    write_reports(
        frame=frame,
        task_map=task_map,
        pooled_metrics=pooled_metrics,
        model_records=model_records,
        gates=gates,
        category=category,
        decision=decision,
        reuse=reuse,
    )
    if not reuse:
        write_json(signature_path, analysis_signature)
    artifact_manifest().to_csv(OUTPUT_DIR / "artifact_manifest.tsv", sep="\t", index=False)

    payload = {
        "analysis_version": ANALYSIS_VERSION,
        "category": category,
        "decision": decision,
        "n_models": len(model_records),
        "max_reload_prediction_error": max(record.parity_max_abs_error for record in model_records),
        "task_map": task_map.to_dict(orient="records"),
        "figure": str((FIGURE_DIR / "Figure_X123_age_coordinate_and_task_map.png").relative_to(ROOT)),
        "run_mode": "reuse" if reuse else "fit-once",
    }
    print(json.dumps(payload, indent=2, ensure_ascii=False))


if __name__ == "__main__":
    main()
```
