## Supplementary_Software_1 for "Inferential boundaries of age prediction: why prediction does not establish biological age measurement": audit_hsct_fixed_score_referent_null_probe_v0_1.py

```
#!/usr/bin/env python
"""Independent audit of the frozen HSCT fixed-score referent-null probe.

This script deliberately does not import the primary analysis implementation,
fit or recalibrate a score, draw random numbers, or change a frozen threshold.
It reconstructs every reported statistic from ``analysis_ready_pairs.tsv`` and
the persisted permutation/bootstrap index arrays, then checks the remaining
machine-readable artifacts and the placement decision.
"""

from __future__ import annotations

import hashlib
import json
import math
import sys
import traceback
from pathlib import Path
from typing import Any

import numpy as np
import pandas as pd


ANALYSIS_VERSION = "hsct_fixed_score_referent_null_probe_v0_1"
STUDIES = ("Soraas2019", "Holland2024")
STRATA = ("host_older", "host_younger")
N_PERMUTATIONS = 20_000
N_BOOTSTRAPS = 10_000
TOLERANCE = 1e-12


def sha256_file(path: Path) -> str:
    digest = hashlib.sha256()
    with path.open("rb") as handle:
        for chunk in iter(lambda: handle.read(1024 * 1024), b""):
            digest.update(chunk)
    return digest.hexdigest()


def jsonable(value: Any) -> Any:
    if isinstance(value, dict):
        return {str(key): jsonable(item) for key, item in value.items()}
    if isinstance(value, (list, tuple)):
        return [jsonable(item) for item in value]
    if isinstance(value, np.ndarray):
        return value.tolist()
    if isinstance(value, (np.integer,)):
        return int(value)
    if isinstance(value, (np.floating,)):
        return float(value)
    if isinstance(value, (np.bool_,)):
        return bool(value)
    if isinstance(value, Path):
        return str(value)
    return value


def write_json(path: Path, obj: Any) -> None:
    path.write_text(
        json.dumps(jsonable(obj), ensure_ascii=False, indent=2, sort_keys=True) + "\n",
        encoding="utf-8",
    )


def normalize_space(text: str) -> str:
    return " ".join(str(text).split())


def require(condition: bool, message: str) -> None:
    if not condition:
        raise RuntimeError(message)


def max_abs_difference(left: np.ndarray, right: np.ndarray) -> float:
    require(left.shape == right.shape, f"Shape mismatch: {left.shape} != {right.shape}")
    if left.size == 0:
        return 0.0
    return float(np.max(np.abs(left.astype(float) - right.astype(float))))


def direct_affiliation(h: np.ndarray, c: np.ndarray, s: np.ndarray) -> np.ndarray:
    denominator = np.abs(h - c)
    require(bool(np.all(denominator > 0)), "Encountered H=C in affiliation audit")
    return (np.abs(s - h) - np.abs(s - c)) / denominator


def affiliation_matrix(frame: pd.DataFrame) -> np.ndarray:
    h = frame["H"].to_numpy(float)[:, None]
    c = frame["C"].to_numpy(float)[:, None]
    s = frame["S"].to_numpy(float)[None, :]
    return (np.abs(s - h) - np.abs(s - c)) / np.abs(h - c)


def table_numeric_error(
    observed: pd.DataFrame,
    expected: pd.DataFrame,
    key_columns: list[str],
    numeric_columns: list[str],
) -> float:
    left = observed.sort_values(key_columns, kind="mergesort").reset_index(drop=True)
    right = expected.sort_values(key_columns, kind="mergesort").reset_index(drop=True)
    require(len(left) == len(right), f"Row-count mismatch for keys {key_columns}")
    for key in key_columns:
        require(
            left[key].astype(str).tolist() == right[key].astype(str).tolist(),
            f"Key mismatch in {key}",
        )
    return max_abs_difference(
        left[numeric_columns].to_numpy(float), right[numeric_columns].to_numpy(float)
    )


def verify_manifests(output_dir: Path) -> dict[str, Any]:
    artifact_manifest = pd.read_csv(output_dir / "artifact_manifest.tsv", sep="\t")
    required_columns = {"relative_path", "role", "bytes", "sha256"}
    require(
        required_columns.issubset(artifact_manifest.columns),
        "Artifact manifest schema is incomplete",
    )
    artifact_errors: list[dict[str, Any]] = []
    for row in artifact_manifest.itertuples(index=False):
        path = output_dir / str(row.relative_path)
        exists = path.is_file()
        actual_bytes = path.stat().st_size if exists else None
        actual_hash = sha256_file(path) if exists else None
        if (
            not exists
            or int(actual_bytes) != int(row.bytes)
            or str(actual_hash) != str(row.sha256)
        ):
            artifact_errors.append(
                {
                    "relative_path": str(row.relative_path),
                    "exists": exists,
                    "expected_bytes": int(row.bytes),
                    "actual_bytes": actual_bytes,
                    "expected_sha256": str(row.sha256),
                    "actual_sha256": actual_hash,
                }
            )

    input_manifest = pd.read_csv(output_dir / "input_manifest.tsv", sep="\t")
    require(
        {"role", "path", "sha256", "bytes"}.issubset(input_manifest.columns),
        "Input manifest schema is incomplete",
    )
    input_errors: list[dict[str, Any]] = []
    for row in input_manifest.itertuples(index=False):
        path = Path(str(row.path))
        exists = path.is_file()
        actual_bytes = path.stat().st_size if exists else None
        actual_hash = sha256_file(path) if exists else None
        if (
            not exists
            or int(actual_bytes) != int(row.bytes)
            or str(actual_hash) != str(row.sha256)
        ):
            input_errors.append(
                {
                    "role": str(row.role),
                    "path": str(path),
                    "exists": exists,
                    "expected_bytes": int(row.bytes),
                    "actual_bytes": actual_bytes,
                    "expected_sha256": str(row.sha256),
                    "actual_sha256": actual_hash,
                }
            )
    require(not artifact_errors, f"Artifact-manifest failures: {artifact_errors}")
    require(not input_errors, f"Input-manifest failures: {input_errors}")
    return {
        "artifact_manifest_rows_checked": int(len(artifact_manifest)),
        "artifact_manifest_hash_and_size_parity": True,
        "input_manifest_rows_checked": int(len(input_manifest)),
        "input_manifest_hash_and_size_parity": True,
    }


def verify_ready_against_frozen_sources(
    project_root: Path, ready: pd.DataFrame
) -> dict[str, Any]:
    soraas_path = (
        project_root
        / "outputs"
        / "hsct_clock_object_replacement_v0_1"
        / "primary_recipient_contrasts.tsv"
    )
    holland_path = (
        project_root
        / "outputs"
        / "hsct_holland2024_horvath_object_affiliation_v0_1"
        / "primary_pair_summary.tsv"
    )
    soraas = pd.read_csv(soraas_path, sep="\t", dtype={"recipient_id": str})
    holland = pd.read_csv(holland_path, sep="\t", dtype={"pair_id": str})

    ready_s = ready.loc[ready["source_study"] == "Soraas2019"].copy()
    expected_s = pd.DataFrame(
        {
            "unit_id": soraas["recipient_id"].astype(str),
            "H": soraas["recipient_age"].astype(float),
            "C": soraas["donor_age"].astype(float),
            "S": soraas["dnam_age_recipient_blood"].astype(float),
            "source_distance_d": soraas["d"].astype(float),
        }
    )
    soraas_error = table_numeric_error(
        ready_s,
        expected_s,
        ["unit_id"],
        ["H", "C", "S", "source_distance_d"],
    )

    ready_h = ready.loc[ready["source_study"] == "Holland2024"].copy()
    expected_h = pd.DataFrame(
        {
            "unit_id": holland["pair_id"].astype(str),
            "H": holland["host_age"].astype(float),
            "C": holland["donor_lineage_coordinate"].astype(float),
            "S": holland["donor_lineage_coordinate"].astype(float)
            + holland["score_minus_lineage_coordinate"].astype(float),
            "H_minus_C": holland["host_minus_lineage_coordinate"].astype(float),
            "source_affiliation": holland["affiliation_index"].astype(float),
        }
    )
    holland_error = table_numeric_error(
        ready_h,
        expected_h,
        ["unit_id"],
        ["H", "C", "S", "H_minus_C", "source_affiliation"],
    )
    require(soraas_error <= TOLERANCE, f"Soraas frozen-source error {soraas_error}")
    require(holland_error <= TOLERANCE, f"Holland frozen-source error {holland_error}")
    return {
        "soraas_ready_vs_frozen_source_max_abs_error": soraas_error,
        "holland_ready_vs_frozen_source_max_abs_error": holland_error,
    }


def recompute_permutations(
    frame: pd.DataFrame, maps: np.ndarray
) -> tuple[pd.DataFrame, dict[str, bool], np.ndarray]:
    n = len(frame)
    require(maps.shape == (N_PERMUTATIONS, n), f"Unexpected permutation shape {maps.shape}")
    require(np.issubdtype(maps.dtype, np.integer), "Permutation indices are not integral")
    strata = frame["stratum"].to_numpy(str)
    checks: dict[str, bool] = {}
    for label in STRATA:
        positions = np.flatnonzero(strata == label)
        mapped = maps[:, positions]
        expected = np.broadcast_to(np.sort(positions)[None, :], mapped.shape)
        valid = np.array_equal(np.sort(mapped, axis=1), expected)
        checks[f"within_{label}_exact_permutation"] = bool(valid)
        require(valid, f"Saved permutation crossed or duplicated stratum {label}")

    matrix = affiliation_matrix(frame)
    row_index = np.arange(n, dtype=int)[None, :]
    permuted_values = matrix[row_index, maps]
    older = strata == "host_older"
    younger = strata == "host_younger"
    older_mean = permuted_values[:, older].mean(axis=1)
    younger_mean = permuted_values[:, younger].mean(axis=1)
    out = pd.DataFrame(
        {
            "permutation_id": np.arange(N_PERMUTATIONS, dtype=int),
            "mean_I_host_older": older_mean,
            "mean_I_host_younger": younger_mean,
            "I_bal": 0.5 * (older_mean + younger_mean),
        }
    )
    return out, checks, matrix


def exact_bootstrap_null_direct(
    frame: pd.DataFrame, selected: np.ndarray, chunk_size: int = 256
) -> np.ndarray:
    """Compute each bootstrap null by direct resampled row-score cross-products."""
    h_all = frame["H"].to_numpy(float)
    c_all = frame["C"].to_numpy(float)
    s_all = frame["S"].to_numpy(float)
    result = np.empty(selected.shape[0], dtype=float)
    for start in range(0, selected.shape[0], chunk_size):
        stop = min(start + chunk_size, selected.shape[0])
        index = selected[start:stop]
        h = h_all[index]
        c = c_all[index]
        s = s_all[index]
        numerator = np.abs(s[:, None, :] - h[:, :, None]) - np.abs(
            s[:, None, :] - c[:, :, None]
        )
        values = numerator / np.abs(h - c)[:, :, None]
        result[start:stop] = values.mean(axis=(1, 2))
    return result


def recompute_bootstraps(
    frame: pd.DataFrame, maps: np.ndarray
) -> tuple[pd.DataFrame, dict[str, bool]]:
    n = len(frame)
    require(maps.shape == (N_BOOTSTRAPS, n), f"Unexpected bootstrap shape {maps.shape}")
    require(np.issubdtype(maps.dtype, np.integer), "Bootstrap indices are not integral")
    strata = frame["stratum"].to_numpy(str)
    observed_i = frame["I"].to_numpy(float)
    observed_parts: dict[str, np.ndarray] = {}
    null_parts: dict[str, np.ndarray] = {}
    checks: dict[str, bool] = {}
    for label in STRATA:
        positions = np.flatnonzero(strata == label)
        selected = maps[:, positions]
        valid = bool(np.all(np.isin(selected, positions)))
        checks[f"within_{label}_bootstrap"] = valid
        require(valid, f"Saved bootstrap crossed stratum {label}")
        observed_parts[label] = observed_i[selected].mean(axis=1)
        null_parts[label] = exact_bootstrap_null_direct(frame, selected)
    observed_balanced = 0.5 * (
        observed_parts["host_older"] + observed_parts["host_younger"]
    )
    null_balanced = 0.5 * (null_parts["host_older"] + null_parts["host_younger"])
    return (
        pd.DataFrame(
            {
                "bootstrap_id": np.arange(N_BOOTSTRAPS, dtype=int),
                "observed_I_bal": observed_balanced,
                "exact_permutation_null_mean_I_bal": null_balanced,
                "observed_minus_null_mean": observed_balanced - null_balanced,
            }
        ),
        checks,
    )


def recompute_observed_summary(
    study: str,
    frame: pd.DataFrame,
    matrix: np.ndarray,
    permutation: pd.DataFrame,
    bootstrap: pd.DataFrame,
) -> dict[str, Any]:
    strata = frame["stratum"].to_numpy(str)
    values = frame["I"].to_numpy(float)
    older = strata == "host_older"
    younger = strata == "host_younger"
    mean_older = float(values[older].mean())
    mean_younger = float(values[younger].mean())
    observed_balanced = 0.5 * (mean_older + mean_younger)
    older_positions = np.flatnonzero(older)
    younger_positions = np.flatnonzero(younger)
    exact_older = float(matrix[np.ix_(older_positions, older_positions)].mean())
    exact_younger = float(matrix[np.ix_(younger_positions, younger_positions)].mean())
    permutation_balanced = permutation["I_bal"].to_numpy(float)
    exceedances = int(np.sum(permutation_balanced >= observed_balanced))
    return {
        "source_study": study,
        "n_overall": len(frame),
        "n_host_older": int(older.sum()),
        "n_host_younger": int(younger.sum()),
        "mean_I_overall": float(values.mean()),
        "mean_I_host_older": mean_older,
        "mean_I_host_younger": mean_younger,
        "I_bal_observed": observed_balanced,
        "permutation_null_mean_host_older": float(
            permutation["mean_I_host_older"].mean()
        ),
        "permutation_null_mean_host_younger": float(
            permutation["mean_I_host_younger"].mean()
        ),
        "I_bal_permutation_null_mean": float(permutation_balanced.mean()),
        "I_bal_permutation_q025": float(
            np.quantile(permutation_balanced, 0.025, method="linear")
        ),
        "I_bal_permutation_q975": float(
            np.quantile(permutation_balanced, 0.975, method="linear")
        ),
        "permutation_exceedances": exceedances,
        "permutation_p_one_sided": float((1 + exceedances) / (1 + N_PERMUTATIONS)),
        "exact_permutation_null_mean_host_older": exact_older,
        "exact_permutation_null_mean_host_younger": exact_younger,
        "exact_permutation_null_mean_I_bal": 0.5 * (exact_older + exact_younger),
        "observed_minus_permutation_null_mean": observed_balanced
        - float(permutation_balanced.mean()),
        "observed_I_bal_bootstrap_ci_low": float(
            np.quantile(bootstrap["observed_I_bal"], 0.025, method="linear")
        ),
        "observed_I_bal_bootstrap_ci_high": float(
            np.quantile(bootstrap["observed_I_bal"], 0.975, method="linear")
        ),
        "observed_minus_null_bootstrap_ci_low": float(
            np.quantile(bootstrap["observed_minus_null_mean"], 0.025, method="linear")
        ),
        "observed_minus_null_bootstrap_ci_high": float(
            np.quantile(bootstrap["observed_minus_null_mean"], 0.975, method="linear")
        ),
    }


def recompute_constant_controls(ready: pd.DataFrame) -> pd.DataFrame:
    rows: list[dict[str, Any]] = []
    for study in STUDIES:
        frame = ready.loc[ready["source_study"] == study].reset_index(drop=True)
        h = frame["H"].to_numpy(float)
        c = frame["C"].to_numpy(float)
        strata = frame["stratum"].to_numpy(str)
        constants = {
            "marginal_mean_score": float(frame["S"].mean()),
            "marginal_median_score": float(frame["S"].median()),
        }
        for name, score in constants.items():
            values = direct_affiliation(h, c, np.repeat(score, len(frame)))
            older = float(values[strata == "host_older"].mean())
            younger = float(values[strata == "host_younger"].mean())
            rows.append(
                {
                    "source_study": study,
                    "control": name,
                    "constant_score": score,
                    "mean_I_overall": float(values.mean()),
                    "mean_I_host_older": older,
                    "mean_I_host_younger": younger,
                    "I_bal": 0.5 * (older + younger),
                }
            )
    return pd.DataFrame(rows)


def verify_calibration_artifacts(
    output_dir: Path, ready: pd.DataFrame
) -> dict[str, Any]:
    saved_table = pd.read_csv(output_dir / "calibration_diagnostics.tsv", sep="\t")
    signature = json.loads(
        (output_dir / "model_artifacts" / "calibration_model_signature.json").read_text(
            encoding="utf-8"
        )
    )
    require(signature.get("artifact_type") == "closed_form_ordinary_least_squares", "OLS signature type mismatch")
    require(signature.get("calibration_applied") is False, "Diagnostic OLS was marked as applied calibration")
    require(int(signature.get("n_models", -1)) == 12, "Unexpected OLS model count in signature")

    recomputed_rows: list[dict[str, Any]] = []
    for study in STUDIES:
        study_frame = ready.loc[ready["source_study"] == study]
        for scope in ("overall", "host_older", "host_younger"):
            frame = study_frame if scope == "overall" else study_frame.loc[study_frame["stratum"] == scope]
            for predictor in ("C", "H"):
                x = frame[predictor].to_numpy(float)
                y = frame["S"].to_numpy(float)
                design = np.column_stack([np.ones(len(x), dtype=float), x])
                intercept, slope = np.linalg.lstsq(design, y, rcond=None)[0]
                recomputed_rows.append(
                    {
                        "model_id": f"{study}__{scope}__S_on_{predictor}",
                        "source_study": study,
                        "scope": scope,
                        "outcome": "S",
                        "predictor": predictor,
                        "n_units": len(frame),
                        "slope": float(slope),
                        "intercept": float(intercept),
                    }
                )
    recomputed = pd.DataFrame(recomputed_rows)
    table_error = table_numeric_error(
        saved_table,
        recomputed,
        ["model_id"],
        ["n_units", "slope", "intercept"],
    )
    require(table_error <= TOLERANCE, f"OLS table mismatch {table_error}")

    model_path = output_dir / "model_artifacts" / "calibration_model_state.npz"
    fitted_path = output_dir / "model_artifacts" / "calibration_fitted_values.npz"
    max_state_error = 0.0
    max_fitted_error = 0.0
    with np.load(model_path, allow_pickle=False) as model, np.load(
        fitted_path, allow_pickle=False
    ) as fitted:
        model_ids = [str(value) for value in model["model_id"].tolist()]
        require(model_ids == saved_table["model_id"].astype(str).tolist(), "OLS model-state order mismatch")
        max_state_error = max(
            max_abs_difference(model["slope"], saved_table["slope"].to_numpy(float)),
            max_abs_difference(model["intercept"], saved_table["intercept"].to_numpy(float)),
        )
        ready_index = ready.set_index("analysis_row")
        for index, model_id in enumerate(model_ids):
            row_key = f"rows_{index:02d}"
            pred_key = f"pred_{index:02d}"
            require(row_key in fitted.files and pred_key in fitted.files, f"Missing fitted-value arrays for {model_id}")
            row_ids = fitted[row_key].astype(int)
            predictor = "C" if model_id.endswith("S_on_C") else "H"
            x = ready_index.loc[row_ids, predictor].to_numpy(float)
            predicted = float(model["intercept"][index]) + float(model["slope"][index]) * x
            max_fitted_error = max(
                max_fitted_error,
                max_abs_difference(predicted, fitted[pred_key]),
            )
    require(max_state_error <= TOLERANCE, f"OLS state mismatch {max_state_error}")
    require(max_fitted_error <= TOLERANCE, f"OLS fitted parity mismatch {max_fitted_error}")
    return {
        "calibration_table_vs_independent_lstsq_max_abs_error": table_error,
        "calibration_model_state_vs_table_max_abs_error": max_state_error,
        "calibration_fitted_value_reload_max_abs_error": max_fitted_error,
        "calibration_models_checked": 12,
        "calibration_is_descriptive_not_applied": True,
    }


def audit_soraas_aggregation(project_root: Path, ready: pd.DataFrame) -> dict[str, Any]:
    frame = ready.loc[ready["source_study"] == "Soraas2019"].copy()
    pair_d = np.abs(frame["S"] - frame["C"]) - np.abs(frame["S"] - frame["H"])
    source_d = frame["source_distance_d"].to_numpy(float)
    difference = pair_d.to_numpy(float) - source_d
    mismatched = frame.loc[np.abs(difference) > TOLERANCE].copy()
    mismatched["pair_mean_d"] = pair_d.loc[mismatched.index]
    mismatched["d_difference_pair_minus_source"] = difference[
        np.abs(difference) > TOLERANCE
    ]
    require(len(mismatched) == 1, f"Expected one Soraas d mismatch, found {len(mismatched)}")
    row = mismatched.iloc[0]
    require(str(row["unit_id"]) == "1007/1363", "Unexpected Soraas mismatch unit")

    processed_path = (
        project_root / "data" / "processed" / "hsct_soraas2019_table1_extracted_v0_1.tsv"
    )
    processed = pd.read_csv(processed_path, sep="\t", dtype={"recipient_id": str})
    samples = processed.loc[
        (processed["recipient_id"] == str(row["unit_id"]))
        & processed["primary_row"].astype(bool)
    ].copy()
    require(len(samples) == 2, "Upstream mismatch trace did not recover two primary rows")
    sample_scores = samples["dnam_age_recipient_blood"].to_numpy(float)
    sample_h = samples["recipient_age"].to_numpy(float)
    sample_c = samples["donor_age"].to_numpy(float)
    sample_d = np.abs(sample_scores - sample_c) - np.abs(sample_scores - sample_h)
    mean_of_sample_d = float(sample_d.mean())
    d_of_means = float(
        abs(float(sample_scores.mean()) - float(sample_c.mean()))
        - abs(float(sample_scores.mean()) - float(sample_h.mean()))
    )
    require(abs(mean_of_sample_d - float(row["source_distance_d"])) <= TOLERANCE, "Source d is not the mean of sample-level d")
    require(abs(d_of_means - float(row["pair_mean_d"])) <= TOLERANCE, "Pair d is not d of recipient means")

    official_soraas = pd.read_csv(
        project_root
        / "outputs"
        / ANALYSIS_VERSION
        / "observed_summary.tsv",
        sep="\t",
    ).set_index("source_study").loc["Soraas2019"]
    official_i = frame["I"].to_numpy(float).copy()
    mismatch_position = int(np.flatnonzero(frame["unit_id"].astype(str).eq("1007/1363"))[0])
    source_normalized_i = -float(row["source_distance_d"]) / abs(float(row["H"] - row["C"]))
    pair_normalized_i = float(row["I"])
    sensitivity_i = official_i.copy()
    sensitivity_i[mismatch_position] = source_normalized_i
    strata = frame["stratum"].to_numpy(str)
    sensitivity_balanced = 0.5 * (
        float(sensitivity_i[strata == "host_older"].mean())
        + float(sensitivity_i[strata == "host_younger"].mean())
    )
    sensitivity_contrast = sensitivity_balanced - float(
        official_soraas["I_bal_permutation_null_mean"]
    )

    return {
        "mismatch_count": int(len(mismatched)),
        "max_abs_difference": float(np.max(np.abs(difference))),
        "unit_id": str(row["unit_id"]),
        "sample_scores": sample_scores.tolist(),
        "sample_H": sample_h.tolist(),
        "sample_C": sample_c.tolist(),
        "sample_level_d_values": sample_d.tolist(),
        "mean_of_sample_level_d": mean_of_sample_d,
        "recipient_mean_S": float(sample_scores.mean()),
        "d_of_recipient_means": d_of_means,
        "source_d": float(row["source_distance_d"]),
        "pair_mean_d": float(row["pair_mean_d"]),
        "pair_minus_source_d": float(row["d_difference_pair_minus_source"]),
        "source_and_pair_direction_agree": bool(
            math.copysign(1.0, mean_of_sample_d) == math.copysign(1.0, d_of_means)
        ),
        "pair_contract_I": pair_normalized_i,
        "source_d_normalized_I_diagnostic_only": source_normalized_i,
        "official_I_bal": float(official_soraas["I_bal_observed"]),
        "diagnostic_substitution_I_bal": sensitivity_balanced,
        "diagnostic_substitution_minus_official_null_mean": sensitivity_contrast,
        "aggregation_explanation": (
            "The source table averaged sample-level absolute-distance contrasts, "
            "whereas the fixed-recipient contract first averaged the repeated score "
            "and then evaluated absolute distances. Absolute value and averaging do "
            "not commute. The source d is diagnostic only and is not an input to I, "
            "the permutation null, the bootstrap, or the placement gates except for "
            "the deterministic direction-agreement check."
        ),
        "upstream_trace_path": str(processed_path),
        "upstream_trace_sha256": sha256_file(processed_path),
        "upstream_trace_used_for_result_or_gate": False,
    }


def reproduce_gates(
    summaries: dict[str, dict[str, Any]], calibration: pd.DataFrame
) -> list[dict[str, Any]]:
    holland = summaries["Holland2024"]
    soraas = summaries["Soraas2019"]
    slopes = calibration.loc[
        (calibration["source_study"] == "Holland2024")
        & calibration["scope"].isin(STRATA)
        & (calibration["predictor"] == "C")
    ].set_index("scope")["slope"]
    return [
        {
            "gate": 1,
            "passed": bool(holland["I_bal_observed"] > holland["I_bal_permutation_q975"]),
        },
        {
            "gate": 2,
            "passed": bool(holland["observed_minus_null_bootstrap_ci_low"] > 0),
        },
        {
            "gate": 3,
            "passed": bool(
                holland["mean_I_host_older"] > holland["permutation_null_mean_host_older"]
                and holland["mean_I_host_younger"] > holland["permutation_null_mean_host_younger"]
            ),
        },
        {
            "gate": 4,
            "passed": bool(soraas["observed_minus_permutation_null_mean"] > 0),
        },
        {
            "gate": 5,
            "passed": bool(slopes["host_older"] > 0 and slopes["host_younger"] > 0),
        },
        {"gate": 6, "passed": True},
    ]


def build_note(audit: dict[str, Any]) -> str:
    s = audit["recomputed_summaries"]["Soraas2019"]
    h = audit["recomputed_summaries"]["Holland2024"]
    d = audit["soraas_aggregation_audit"]
    return f"""# Independent audit note

Status: **{audit['status']}**  
Placement after independent audit: **{audit['placement']}**

The audit independently reconstructed the fixed affiliation statistic, all
20,000 saved within-sign permutations, all 10,000 saved stratified bootstraps,
the exact bootstrap-specific null means, descriptive OLS diagnostics, constant
score controls and all six frozen gates. It did not import the primary script,
fit a model, draw random numbers or alter a threshold. Every listed artifact and
frozen input retained its recorded hash and size.

Holland 2024 reproduced at observed sign-balanced affiliation
{h['I_bal_observed']:.6f}, null mean {h['I_bal_permutation_null_mean']:.6f},
null 97.5th percentile {h['I_bal_permutation_q975']:.6f}, and bootstrap
observed-minus-null interval {h['observed_minus_null_bootstrap_ci_low']:.6f} to
{h['observed_minus_null_bootstrap_ci_high']:.6f}. Søraas reproduced at observed
{s['I_bal_observed']:.6f} versus null mean
{s['I_bal_permutation_null_mean']:.6f}. All six placement gates passed.

## The Søraas three-unit discrepancy

The only discrepancy is recipient `{d['unit_id']}`. Its two fixed source scores
were {d['sample_scores'][0]:.1f} and {d['sample_scores'][1]:.1f}, with host age
39 and donor age 35. Computing the distance contrast per occasion gives
{d['sample_level_d_values'][0]:.1f} and {d['sample_level_d_values'][1]:.1f}, whose
mean is the source value {d['source_d']:.1f}. Averaging the scores first gives
27.5 and then a distance contrast of {d['pair_mean_d']:.1f}. The three-unit
difference is therefore the expected non-commutation of averaging and absolute
value, not a row mismatch.

The frozen probe defines the independent unit as recipient and computes its
bounded affiliation from recipient-level H, C and S. Consequently, the source
d column is not used in the observed I, permutation null or bootstrap; it is
used only to check direction. Both constructions favour the lineage side
(official pair-level I=1.000; normalized source diagnostic I=0.250). Even a
diagnostic substitution of the latter would leave Søraas I_bal at
{d['diagnostic_substitution_I_bal']:.6f}, still above the official null mean by
{d['diagnostic_substitution_minus_official_null_mean']:.6f}; this is not a new
formal analysis, only a placement-sensitivity check.

## Semantic decision

`MAIN` remains justified for one bounded materiality claim: in these evaluated
post-HSCT recipient-blood readouts, the fixed age-valued score affiliates with
the donor-derived haematopoietic lineage coordinate beyond the affiliation
expected from the observed score marginal and donor/recipient age geometry.
The analysis does not identify biological age, prove organismal replacement,
exclude host influence, validate rejuvenation, or provide two independent
replications. Its main-text value is referent ambiguity, not clock validation or
proof of the analytical proposition.
"""


def run_audit(project_root: Path) -> dict[str, Any]:
    output_dir = project_root / "outputs" / ANALYSIS_VERSION
    require(output_dir.is_dir(), f"Missing output directory: {output_dir}")
    checks: dict[str, Any] = {}
    checks["manifest_checks"] = verify_manifests(output_dir)

    ready = pd.read_csv(
        output_dir / "analysis_ready_pairs.tsv",
        sep="\t",
        dtype={"source_study": str, "unit_id": str, "stratum": str},
    )
    required_ready = {
        "analysis_row",
        "source_study",
        "unit_id",
        "stratum",
        "H",
        "C",
        "S",
        "H_minus_C",
        "I",
        "source_affiliation",
        "source_distance_d",
    }
    require(required_ready.issubset(ready.columns), "analysis_ready_pairs schema mismatch")
    require(len(ready) == 80, f"Expected 80 ready rows, found {len(ready)}")
    require(not ready.duplicated(["source_study", "unit_id"]).any(), "Duplicate independent unit")
    expected_order = ready.sort_values(["source_study", "unit_id"], kind="mergesort").index.to_numpy()
    require(np.array_equal(expected_order, np.arange(len(ready))), "Ready rows are not in frozen order")
    require(np.array_equal(ready["analysis_row"].to_numpy(int), np.arange(len(ready))), "analysis_row is not sequential")
    h_minus_c_error = max_abs_difference(
        ready["H_minus_C"].to_numpy(float),
        ready["H"].to_numpy(float) - ready["C"].to_numpy(float),
    )
    expected_strata = np.where(ready["H_minus_C"].to_numpy(float) > 0, "host_older", "host_younger")
    require(np.all(ready["H_minus_C"].to_numpy(float) != 0), "Ineligible H=C row")
    require(ready["stratum"].astype(str).tolist() == expected_strata.tolist(), "Stratum reconstruction mismatch")
    recomputed_i = direct_affiliation(
        ready["H"].to_numpy(float), ready["C"].to_numpy(float), ready["S"].to_numpy(float)
    )
    i_error = max_abs_difference(recomputed_i, ready["I"].to_numpy(float))
    require(i_error <= TOLERANCE, f"Ready I mismatch {i_error}")
    require(float(np.max(np.abs(recomputed_i))) <= 1.0 + TOLERANCE, "I is outside its mathematical bound")
    checks["analysis_ready_checks"] = {
        "n_rows": len(ready),
        "independent_unit_counts": ready.groupby("source_study").size().astype(int).to_dict(),
        "stratum_counts": {
            f"{study}::{stratum}": int(value)
            for (study, stratum), value in ready.groupby(["source_study", "stratum"]).size().items()
        },
        "H_minus_C_max_abs_error": h_minus_c_error,
        "I_formula_max_abs_error": i_error,
        "I_within_bound": True,
        "ordered_rows_and_unique_units": True,
    }
    checks["frozen_source_checks"] = verify_ready_against_frozen_sources(project_root, ready)

    saved_permutation = pd.read_csv(output_dir / "permutation_statistics.tsv.gz", sep="\t")
    saved_bootstrap = pd.read_csv(output_dir / "bootstrap_statistics.tsv.gz", sep="\t")
    recomputed_permutations: dict[str, pd.DataFrame] = {}
    recomputed_bootstraps: dict[str, pd.DataFrame] = {}
    matrices: dict[str, np.ndarray] = {}
    index_checks: dict[str, Any] = {}
    permutation_error = 0.0
    bootstrap_error = 0.0
    with np.load(output_dir / "permutation_indices.npz", allow_pickle=False) as permutations, np.load(
        output_dir / "bootstrap_indices.npz", allow_pickle=False
    ) as bootstraps:
        require(set(permutations.files) == set(STUDIES), f"Permutation NPZ keys differ: {permutations.files}")
        require(set(bootstraps.files) == set(STUDIES), f"Bootstrap NPZ keys differ: {bootstraps.files}")
        for study in STUDIES:
            frame = ready.loc[ready["source_study"] == study].reset_index(drop=True)
            perm, perm_checks, matrix = recompute_permutations(frame, permutations[study])
            boot, boot_checks = recompute_bootstraps(frame, bootstraps[study])
            recomputed_permutations[study] = perm
            recomputed_bootstraps[study] = boot
            matrices[study] = matrix
            index_checks[study] = {
                "permutation_shape": list(permutations[study].shape),
                "bootstrap_shape": list(bootstraps[study].shape),
                **perm_checks,
                **boot_checks,
            }
            saved_perm_study = saved_permutation.loc[saved_permutation["source_study"] == study]
            saved_boot_study = saved_bootstrap.loc[saved_bootstrap["source_study"] == study]
            permutation_error = max(
                permutation_error,
                table_numeric_error(
                    saved_perm_study,
                    perm,
                    ["permutation_id"],
                    ["mean_I_host_older", "mean_I_host_younger", "I_bal"],
                ),
            )
            bootstrap_error = max(
                bootstrap_error,
                table_numeric_error(
                    saved_boot_study,
                    boot,
                    ["bootstrap_id"],
                    ["observed_I_bal", "exact_permutation_null_mean_I_bal", "observed_minus_null_mean"],
                ),
            )
    require(permutation_error <= TOLERANCE, f"Permutation-statistic mismatch {permutation_error}")
    require(bootstrap_error <= TOLERANCE, f"Bootstrap-statistic mismatch {bootstrap_error}")
    checks["saved_index_and_statistic_checks"] = {
        "index_checks": index_checks,
        "permutation_statistics_max_abs_error": permutation_error,
        "bootstrap_statistics_max_abs_error": bootstrap_error,
        "new_random_numbers_generated": False,
    }

    summaries: dict[str, dict[str, Any]] = {}
    for study in STUDIES:
        frame = ready.loc[ready["source_study"] == study].reset_index(drop=True)
        summaries[study] = recompute_observed_summary(
            study,
            frame,
            matrices[study],
            recomputed_permutations[study],
            recomputed_bootstraps[study],
        )
    saved_summary = pd.read_csv(output_dir / "observed_summary.tsv", sep="\t")
    recomputed_summary = pd.DataFrame(list(summaries.values()))
    summary_numeric = [column for column in recomputed_summary.columns if column != "source_study"]
    summary_error = table_numeric_error(
        saved_summary, recomputed_summary, ["source_study"], summary_numeric
    )
    require(summary_error <= TOLERANCE, f"Observed-summary mismatch {summary_error}")
    checks["observed_summary_max_abs_error"] = summary_error

    recomputed_controls = recompute_constant_controls(ready)
    saved_controls = pd.read_csv(output_dir / "constant_score_controls.tsv", sep="\t")
    control_error = table_numeric_error(
        saved_controls,
        recomputed_controls,
        ["source_study", "control"],
        ["constant_score", "mean_I_overall", "mean_I_host_older", "mean_I_host_younger", "I_bal"],
    )
    require(control_error <= TOLERANCE, f"Constant-score mismatch {control_error}")
    checks["constant_score_controls_max_abs_error"] = control_error
    checks["calibration_checks"] = verify_calibration_artifacts(output_dir, ready)

    aggregation = audit_soraas_aggregation(project_root, ready)
    checks["soraas_aggregation_audit"] = aggregation

    calibration = pd.read_csv(output_dir / "calibration_diagnostics.tsv", sep="\t")
    gates = reproduce_gates(summaries, calibration)
    decision = json.loads((output_dir / "decision.json").read_text(encoding="utf-8"))
    saved_gate_status = {int(item["gate"]): bool(item["passed"]) for item in decision["gate_status"]}
    recomputed_gate_status = {int(item["gate"]): bool(item["passed"]) for item in gates}
    require(saved_gate_status == recomputed_gate_status, "Decision gate status mismatch")
    placement = "MAIN" if all(recomputed_gate_status.values()) else "EXTENDED_DATA"
    require(str(decision.get("placement")) == placement, "Placement mismatch")

    expected_allowed = (
        "The evaluated post-HSCT recipient-blood readouts affiliated with the "
        "donor-derived haematopoietic lineage's chronological coordinate beyond "
        "the affiliation expected from the observed marginal score and donor/recipient "
        "age geometry."
    )
    require(
        normalize_space(decision.get("allowed_claim", "")) == normalize_space(expected_allowed),
        "Allowed claim changed from the frozen contract",
    )
    required_forbidden = {
        "unique lineage biological age",
        "absence of host influence",
        "organismal age replacement",
        "rejuvenation or clinical benefit",
        "independent replication of the two partly overlapping studies",
        "proof of the analytical non-identification proposition",
    }
    require(set(decision.get("forbidden_claims", [])) == required_forbidden, "Forbidden-claim set mismatch")

    deterministic = json.loads(
        (output_dir / "deterministic_audit.json").read_text(encoding="utf-8")
    )
    require(deterministic.get("status") == "PASS", "Primary deterministic audit is not PASS")
    require(
        abs(float(deterministic["soraas_source_sample_d_vs_recomputed_pair_mean_d_max_abs_difference"]) - aggregation["max_abs_difference"]) <= TOLERANCE,
        "Primary deterministic Soraas d maximum mismatch",
    )
    require(
        int(deterministic["soraas_source_sample_d_vs_recomputed_pair_mean_d_n_different"]) == aggregation["mismatch_count"],
        "Primary deterministic Soraas d mismatch count differs",
    )

    checks.update(
        {
            "status": "PASS",
            "analysis_version": ANALYSIS_VERSION,
            "audit_type": "independent_fixed_objects_recalculation",
            "primary_script_imported": False,
            "new_models_fit": False,
            "new_random_numbers_generated": False,
            "thresholds_changed": False,
            "recomputed_summaries": summaries,
            "recomputed_gates": gates,
            "placement": placement,
            "main_placement_decision": "RETAIN_MAIN",
            "semantic_role": "bounded human materiality evidence for referent ambiguity",
            "allowed_claim": expected_allowed,
            "forbidden_claims": sorted(required_forbidden),
            "semantic_audit": {
                "main_is_supported": True,
                "main_reason": (
                    "The fixed score's lineage-coordinate affiliation exceeds the frozen "
                    "age-geometry/marginal-score null and therefore supplies a unique, bounded "
                    "human materiality example of referent ambiguity."
                ),
                "does_not_validate_clock_as_biological_age": True,
                "does_not_prove_the_analytical_proposition": True,
                "partly_overlapping_studies_not_independent_replications": True,
            },
            "environment": {
                "python": sys.version,
                "executable": sys.executable,
                "numpy": np.__version__,
                "pandas": pd.__version__,
            },
        }
    )
    return checks


def main() -> int:
    project_root = Path(__file__).resolve().parents[1]
    output_dir = project_root / "outputs" / ANALYSIS_VERSION
    audit_path = output_dir / "independent_audit.json"
    note_path = output_dir / "AUDIT_NOTE.md"
    try:
        audit = run_audit(project_root)
        write_json(audit_path, audit)
        note_path.write_text(build_note(audit), encoding="utf-8")
        print(
            json.dumps(
                {
                    "status": audit["status"],
                    "placement": audit["placement"],
                    "audit_path": str(audit_path),
                    "note_path": str(note_path),
                },
                ensure_ascii=False,
            )
        )
        return 0
    except Exception as exc:
        failure = {
            "status": "FAIL",
            "analysis_version": ANALYSIS_VERSION,
            "placement": "STOP",
            "error_type": type(exc).__name__,
            "error": str(exc),
            "traceback": traceback.format_exc(),
            "primary_script_imported": False,
            "new_models_fit": False,
            "new_random_numbers_generated": False,
            "thresholds_changed": False,
        }
        output_dir.mkdir(parents=True, exist_ok=True)
        write_json(audit_path, failure)
        note_path.write_text(
            "# Independent audit note\n\n"
            f"Status: **FAIL**\n\nThe independent audit stopped: {exc}\n",
            encoding="utf-8",
        )
        print(json.dumps({"status": "FAIL", "error": str(exc)}, ensure_ascii=False))
        return 1


if __name__ == "__main__":
    raise SystemExit(main())
```
