## Supplementary_Software_1 for "Inferential boundaries of age prediction: why prediction does not establish biological age measurement": audit_hsct_referent_regression_sensitivity_v0_1.py

```
#!/usr/bin/env python
"""Independent end-to-end audit of the frozen HSCT referent regression.

The audit does not import the primary analysis implementation and does not
generate bootstrap samples.  It reconstructs the closed-form regressions,
HC3 covariance matrices, fixed-slope anchors, bootstrap coefficients and
quantiles from frozen rows and persisted resampling maps.  It then checks the
frozen model objects, all 15 manifest hashes, the direction-lock provenance
and the separately persisted Hannum/network boundary.
"""

from __future__ import annotations

import hashlib
import json
import platform
import sys
from pathlib import Path
from typing import Any

import numpy as np
import pandas as pd
import scipy
from scipy import stats


ROOT = Path(__file__).resolve().parents[1]
AUDIT_VERSION = "hsct_referent_regression_clock_sensitivity_v0_1_independent_audit_v0_1"
OUT = ROOT / "outputs" / AUDIT_VERSION
FROZEN_ROOT = ROOT / "outputs" / "hsct_referent_regression_clock_sensitivity_v0_1"
REGRESSION = FROZEN_ROOT / "regression"
MODEL_DIR = REGRESSION / "model_artifacts"
HANNUM_DIR = FROZEN_ROOT / "hannum2013"
PRIMARY_SCRIPT = ROOT / "analysis" / "analyze_hsct_referent_regression_sensitivity_v0_1.py"
SPEC = ROOT / "analysis" / "HSCT_REFERENT_REGRESSION_CLOCK_SENSITIVITY_SPEC_20260826.md"
P2_LOCK = (
    ROOT
    / "manuscript"
    / "nature_first"
    / "next_round_20260826"
    / "P2_ANALYSIS_DIRECTION_LOCK_20260826.md"
)
FROZEN_READY = (
    ROOT
    / "outputs"
    / "hsct_fixed_score_referent_null_probe_v0_1"
    / "analysis_ready_pairs.tsv"
)

STUDY_ORDER = ("Holland2024", "Soraas2019")
EXPECTED_COUNTS = {"Holland2024": 61, "Soraas2019": 19}
N_BOOTSTRAPS = 10_000
BOOTSTRAP_SEED = 20_260_826
TOLERANCE = 1e-12
FEATURE_ORDER = ["intercept", "A_L_minus_A_R"]

BRANCH_C_START = "## Branch C: HUM-06 matched scalar target supervision\n"
SHARED_CONTRACT_START = "## Shared execution contract\n"
ORIGINAL_BRANCH_C = """## Branch C: HUM-06 matched scalar target supervision

- New module and analysis version: `HUM-06`,
  `nhanes_5y_matched_scalar_target_supervision_probe_v0_1`.
- `HUM-04` remains frozen and is not silently rewritten.
- Flexible calibration uses the existing fold-local age scores and compares a
  fixed five-knot score spline with the frozen direct 50-measurement mortality
  model.
- The matched comparison constructs one age-supervised and one
  mortality-supervised ridge projection from the same 50 measurements and
  background, with identical nested fold exclusions, preprocessing, ridge
  grid, squared-error tuning rule and scalar dimension. Chronological age enters
  only the common downstream endpoint model.
- Both scalars receive the same linear and fixed spline downstream calibration
  opportunities. No learner library or post-result calibration search is used.
- Five prespecified five-fold outer partitions use seeds `20260823`,
  `20260901`, `20260902`, `20260903` and `20260904`. Repeats are not treated as
  independent samples.
- The primary metric is participant-level OOF binary log loss. A paired
  participant bootstrap averages each participant's losses across repeats.
- Failure of identity, feature-order, event-support, leakage, persistence or
  reload-parity gates stops the branch.

"""


def require(condition: bool, message: str) -> None:
    if not condition:
        raise RuntimeError(message)


def sha256_bytes(value: bytes) -> str:
    return hashlib.sha256(value).hexdigest()


def sha256_file(path: Path) -> str:
    digest = hashlib.sha256()
    with path.open("rb") as handle:
        for block in iter(lambda: handle.read(1024 * 1024), b""):
            digest.update(block)
    return digest.hexdigest()


def jsonable(value: Any) -> Any:
    if isinstance(value, dict):
        return {str(key): jsonable(item) for key, item in value.items()}
    if isinstance(value, (list, tuple)):
        return [jsonable(item) for item in value]
    if isinstance(value, np.ndarray):
        return value.tolist()
    if isinstance(value, (np.integer,)):
        return int(value)
    if isinstance(value, (np.floating,)):
        return float(value)
    if isinstance(value, (np.bool_,)):
        return bool(value)
    if isinstance(value, Path):
        return str(value)
    return value


def write_json(path: Path, value: Any) -> None:
    path.write_text(
        json.dumps(jsonable(value), indent=2, sort_keys=True, ensure_ascii=False) + "\n",
        encoding="utf-8",
    )


def max_abs_difference(left: np.ndarray, right: np.ndarray) -> float:
    require(left.shape == right.shape, f"Shape mismatch: {left.shape} != {right.shape}")
    if left.size == 0:
        return 0.0
    return float(np.max(np.abs(left.astype(float) - right.astype(float))))


def add_parity(
    rows: list[dict[str, Any]],
    check_id: str,
    artifact: str,
    error: float,
    detail: str,
    tolerance: float = TOLERANCE,
) -> None:
    require(np.isfinite(error), f"Non-finite parity error for {check_id}")
    passed = bool(error <= tolerance)
    rows.append(
        {
            "check_id": check_id,
            "artifact": artifact,
            "max_abs_error": float(error),
            "tolerance": float(tolerance),
            "status": "PASS" if passed else "FAIL",
            "detail": detail,
        }
    )
    require(passed, f"Parity failed for {check_id}: {error} > {tolerance}")


def compare_table(
    observed: pd.DataFrame,
    expected: pd.DataFrame,
    keys: list[str],
    exact_columns: list[str],
    numeric_columns: list[str],
) -> float:
    left = observed.sort_values(keys, kind="mergesort").reset_index(drop=True)
    right = expected.sort_values(keys, kind="mergesort").reset_index(drop=True)
    require(list(left.columns) == list(right.columns), f"Column mismatch for {keys}")
    require(len(left) == len(right), f"Row-count mismatch for {keys}")
    for column in keys + exact_columns:
        require(
            left[column].astype(str).tolist() == right[column].astype(str).tolist(),
            f"Exact-value mismatch in {column}",
        )
    return max_abs_difference(
        left[numeric_columns].to_numpy(float), right[numeric_columns].to_numpy(float)
    )


def verify_frozen_manifest() -> tuple[pd.DataFrame, pd.DataFrame]:
    manifest_path = REGRESSION / "artifact_manifest.tsv"
    manifest = pd.read_csv(manifest_path, sep="\t")
    require(
        list(manifest.columns) == ["relative_path", "role", "bytes", "sha256"],
        "Frozen artifact-manifest schema changed",
    )
    require(len(manifest) == 15, f"Expected 15 frozen manifest rows, found {len(manifest)}")
    require(not manifest["relative_path"].duplicated().any(), "Duplicate frozen manifest path")
    checks: list[dict[str, Any]] = []
    for row in manifest.itertuples(index=False):
        path = FROZEN_ROOT / Path(str(row.relative_path))
        exists = path.is_file()
        actual_bytes = path.stat().st_size if exists else -1
        actual_hash = sha256_file(path) if exists else ""
        passed = (
            exists
            and int(actual_bytes) == int(row.bytes)
            and actual_hash.lower() == str(row.sha256).lower()
        )
        checks.append(
            {
                "relative_path": str(row.relative_path),
                "role": str(row.role),
                "expected_bytes": int(row.bytes),
                "observed_bytes": int(actual_bytes),
                "expected_sha256": str(row.sha256),
                "observed_sha256": actual_hash,
                "status": "PASS" if passed else "FAIL",
            }
        )
        require(passed, f"Frozen manifest mismatch: {row.relative_path}")
    return manifest, pd.DataFrame(checks)


def audit_direction_lock(input_manifest: pd.DataFrame) -> dict[str, Any]:
    lock_rows = input_manifest.loc[
        input_manifest["relative_path"].astype(str).str.endswith(
            "P2_ANALYSIS_DIRECTION_LOCK_20260826.md"
        )
    ]
    require(len(lock_rows) == 1, "Frozen input manifest does not identify one direction lock")
    lock_row = lock_rows.iloc[0]
    expected_hash = str(lock_row["sha256"]).lower()
    expected_bytes = int(lock_row["bytes"])

    current_raw = P2_LOCK.read_bytes()
    require(not current_raw.startswith(b"\xef\xbb\xbf"), "Current direction lock has a BOM")
    require(b"\r\n" not in current_raw, "Current direction lock is not LF-normalized")
    current_text = current_raw.decode("utf-8")
    require(current_text.count(BRANCH_C_START) == 1, "Branch C heading is not unique")
    require(
        current_text.count(SHARED_CONTRACT_START) == 1,
        "Shared execution-contract heading is not unique",
    )
    start = current_text.index(BRANCH_C_START)
    end = current_text.index(SHARED_CONTRACT_START, start)
    current_branch = current_text[start:end]
    prefix = current_text[:start]
    suffix = current_text[end:]
    reconstructed_text = prefix + ORIGINAL_BRANCH_C + suffix
    reconstructed_without_eof_normalization = reconstructed_text.encode("utf-8")
    require(
        current_raw.endswith(b"\n") and not current_raw.endswith(b"\n\n"),
        "Current direction-lock EOF normalization is not the audited one-LF state",
    )
    reconstructed_raw = reconstructed_without_eof_normalization + b"\n"
    reconstructed_hash = sha256_bytes(reconstructed_raw)

    require(
        len(reconstructed_raw) == expected_bytes,
        "Branch C plus terminal-LF reconstruction has the wrong byte count: "
        f"{len(reconstructed_raw)} != {expected_bytes}",
    )
    require(
        reconstructed_hash == expected_hash,
        "Cannot account exactly for direction-lock drift: "
        f"{reconstructed_hash} != {expected_hash}",
    )
    required_later_governance = [
        "Independent code audit found",
        "v0.1 cannot carry the matched-target verdict",
        "nhanes_5y_matched_scalar_target_supervision_probe_v0_2",
        "Within every downstream C-tuning fold in v0.2",
        "V0.2 reload validation must connect every persisted score end to end",
    ]
    for phrase in required_later_governance:
        require(phrase in current_branch, f"Later Branch C governance marker missing: {phrase}")
    require(
        "Holland is placement-driving; Soeraas is complementary and is not an\n"
        "  independent replication." in prefix,
        "Branch B interpretation boundary changed",
    )
    return {
        "status": "PASS",
        "recorded_pre_update_sha256": expected_hash,
        "recorded_pre_update_bytes": expected_bytes,
        "reconstructed_pre_update_sha256": reconstructed_hash,
        "reconstructed_pre_update_bytes": len(reconstructed_raw),
        "current_sha256": sha256_bytes(current_raw),
        "current_bytes": len(current_raw),
        "unchanged_prefix_sha256": sha256_bytes(prefix.encode("utf-8")),
        "unchanged_suffix_sha256": sha256_bytes(suffix.encode("utf-8")),
        "recorded_branch_c_sha256": sha256_bytes(ORIGINAL_BRANCH_C.encode("utf-8")),
        "current_branch_c_sha256": sha256_bytes(current_branch.encode("utf-8")),
        "semantic_content_difference_confined_to": (
            "the exact bytes from the unique Branch C heading up to, but not "
            "including, the Shared execution contract heading"
        ),
        "additional_nonsemantic_byte_difference": (
            "the recorded pre-update file ended with two LF bytes; the current "
            "file ends with one LF byte"
        ),
        "outside_branch_c_equal_after_terminal_lf_normalization": True,
        "later_update_classification": (
            "Branch C governance update recording the v0.1 leakage audit and the "
            "fully nested v0.2 correction"
        ),
        "strict_proof": (
            "Replacing the current Branch C region with the contemporaneously "
            "recorded block and restoring the single removed terminal LF reproduces "
            "both the frozen byte count and SHA-256. No other textual difference exists."
        ),
        "reconstructed_file_written": False,
    }


def verify_frozen_inputs(input_manifest: pd.DataFrame) -> dict[str, Any]:
    require(
        list(input_manifest.columns) == ["relative_path", "bytes", "sha256"],
        "Frozen input-manifest schema changed",
    )
    require(len(input_manifest) == 3, "Frozen input manifest no longer has three rows")
    results: list[dict[str, Any]] = []
    direction_audit = audit_direction_lock(input_manifest)
    for row in input_manifest.itertuples(index=False):
        relative = str(row.relative_path)
        path = ROOT / Path(relative)
        if relative.endswith("P2_ANALYSIS_DIRECTION_LOCK_20260826.md"):
            results.append(
                {
                    "relative_path": relative,
                    "status": "PASS_VIA_BRANCH_C_RECONSTRUCTION",
                    "expected_sha256": str(row.sha256),
                    "observed_or_reconstructed_sha256": direction_audit[
                        "reconstructed_pre_update_sha256"
                    ],
                }
            )
            continue
        require(path.is_file(), f"Frozen input is missing: {relative}")
        actual_hash = sha256_file(path)
        require(path.stat().st_size == int(row.bytes), f"Frozen input size mismatch: {relative}")
        require(actual_hash == str(row.sha256), f"Frozen input hash mismatch: {relative}")
        results.append(
            {
                "relative_path": relative,
                "status": "PASS",
                "expected_sha256": str(row.sha256),
                "observed_or_reconstructed_sha256": actual_hash,
            }
        )
    return {"status": "PASS", "inputs": results, "direction_lock": direction_audit}


def load_and_validate_ready(parity_rows: list[dict[str, Any]]) -> pd.DataFrame:
    saved = pd.read_csv(
        REGRESSION / "regression_analysis_ready.tsv",
        sep="\t",
        dtype={"unit_id": str, "source_study": str},
    )
    required = [
        "analysis_row",
        "source_study",
        "unit_id",
        "A_R",
        "A_L",
        "score_S",
        "x_A_L_minus_A_R",
        "y_S_minus_A_R",
    ]
    require(list(saved.columns) == required, "Frozen regression-ready schema changed")
    require(len(saved) == 80, f"Expected 80 frozen units, found {len(saved)}")
    require(not saved["analysis_row"].duplicated().any(), "analysis_row is not unique")
    require(
        not saved.duplicated(["source_study", "unit_id"]).any(),
        "Frozen independent unit is duplicated",
    )
    require(saved[required].notna().all().all(), "Missing value in frozen regression-ready data")
    require(
        np.isfinite(saved[required[3:]].to_numpy(float)).all(),
        "Non-finite regression value",
    )
    counts = saved.groupby("source_study").size().astype(int).to_dict()
    require(counts == EXPECTED_COUNTS, f"Independent-unit counts changed: {counts}")

    x_error = max_abs_difference(
        saved["x_A_L_minus_A_R"].to_numpy(float),
        saved["A_L"].to_numpy(float) - saved["A_R"].to_numpy(float),
    )
    y_error = max_abs_difference(
        saved["y_S_minus_A_R"].to_numpy(float),
        saved["score_S"].to_numpy(float) - saved["A_R"].to_numpy(float),
    )
    add_parity(parity_rows, "ready_x_identity", "regression_analysis_ready.tsv", x_error, "x=A_L-A_R")
    add_parity(parity_rows, "ready_y_identity", "regression_analysis_ready.tsv", y_error, "y=S-A_R")

    source = pd.read_csv(
        FROZEN_READY,
        sep="\t",
        dtype={"unit_id": str, "source_study": str},
    )
    source_required = ["analysis_row", "source_study", "unit_id", "H", "C", "S"]
    require(set(source_required).issubset(source.columns), "Frozen source-ready schema changed")
    source = source.sort_values(["source_study", "unit_id"], kind="mergesort").reset_index(drop=True)
    expected = pd.DataFrame(
        {
            "analysis_row": source["analysis_row"].astype(int),
            "source_study": source["source_study"].astype(str),
            "unit_id": source["unit_id"].astype(str),
            "A_R": source["H"].astype(float),
            "A_L": source["C"].astype(float),
            "score_S": source["S"].astype(float),
        }
    )
    expected["x_A_L_minus_A_R"] = expected["A_L"] - expected["A_R"]
    expected["y_S_minus_A_R"] = expected["score_S"] - expected["A_R"]
    ready_source_error = compare_table(
        saved,
        expected,
        ["source_study", "unit_id"],
        ["analysis_row"],
        ["A_R", "A_L", "score_S", "x_A_L_minus_A_R", "y_S_minus_A_R"],
    )
    add_parity(
        parity_rows,
        "ready_vs_frozen_source",
        "analysis_ready_pairs.tsv",
        ready_source_error,
        "saved regression interface independently derived from H,C,S",
    )
    return saved


def fit_ols(x: np.ndarray, y: np.ndarray) -> dict[str, Any]:
    design = np.column_stack([np.ones(len(x), dtype=float), x])
    xtx = design.T @ design
    require(np.linalg.matrix_rank(xtx) == 2, "OLS design is rank deficient")
    xtx_inv = np.linalg.inv(xtx)
    coefficients = xtx_inv @ design.T @ y
    fitted = design @ coefficients
    residual = y - fitted
    leverage = np.einsum("ij,jk,ik->i", design, xtx_inv, design)
    require(bool(np.all(leverage < 1)), "HC3 leverage is not below one")
    scaled = residual**2 / (1.0 - leverage) ** 2
    meat = design.T @ (scaled[:, None] * design)
    covariance = xtx_inv @ meat @ xtx_inv
    se = np.sqrt(np.diag(covariance))
    df = len(x) - 2
    critical = float(stats.t.ppf(0.975, df))
    ci_low = coefficients - critical * se
    ci_high = coefficients + critical * se
    sse = float(residual @ residual)
    sst = float(np.sum((y - y.mean()) ** 2))
    require(sst > 0, "Regression outcome has zero variance")
    return {
        "coefficients": coefficients,
        "fitted": fitted,
        "residual": residual,
        "covariance": covariance,
        "se": se,
        "ci_low": ci_low,
        "ci_high": ci_high,
        "df": df,
        "r2": 1.0 - sse / sst,
        "residual_mse": sse / len(x),
    }


def fit_fixed_beta(x: np.ndarray, y: np.ndarray, beta: float) -> dict[str, Any]:
    alpha = float(np.mean(y - beta * x))
    fitted = alpha + beta * x
    return {
        "coefficients": np.asarray([alpha, beta], dtype=float),
        "fitted": fitted,
        "residual_mse": float(np.mean((y - fitted) ** 2)),
    }


def bootstrap_coefficients(x: np.ndarray, y: np.ndarray, maps: np.ndarray) -> np.ndarray:
    selected_x = x[maps]
    selected_y = y[maps]
    x_mean = selected_x.mean(axis=1)
    y_mean = selected_y.mean(axis=1)
    centered_x = selected_x - x_mean[:, None]
    denominator = np.sum(centered_x**2, axis=1)
    require(bool(np.all(denominator > 0)), "A persisted bootstrap draw has zero x variance")
    beta = np.sum(centered_x * (selected_y - y_mean[:, None]), axis=1) / denominator
    alpha = y_mean - beta * x_mean
    return np.column_stack([alpha, beta])


def linear_quantile(values: np.ndarray, probability: float) -> float:
    return float(np.quantile(values, probability, method="linear"))


def recompute_objects(
    ready: pd.DataFrame,
) -> tuple[
    pd.DataFrame,
    pd.DataFrame,
    pd.DataFrame,
    pd.DataFrame,
    pd.DataFrame,
    pd.DataFrame,
    dict[str, dict[str, Any]],
]:
    summaries: list[dict[str, Any]] = []
    anchors: list[dict[str, Any]] = []
    boot_rows: list[pd.DataFrame] = []
    model_rows: list[dict[str, Any]] = []
    fitted_rows: list[pd.DataFrame] = []
    covariance_rows: list[dict[str, Any]] = []
    objects: dict[str, dict[str, Any]] = {}

    with np.load(REGRESSION / "bootstrap_indices.npz", allow_pickle=False) as maps_file:
        require(
            set(maps_file.files) == {f"maps_{study}" for study in STUDY_ORDER},
            f"Unexpected bootstrap map keys: {maps_file.files}",
        )
        for study in STUDY_ORDER:
            frame = ready.loc[ready["source_study"] == study].copy()
            x = frame["x_A_L_minus_A_R"].to_numpy(float)
            y = frame["y_S_minus_A_R"].to_numpy(float)
            row_ids = frame["analysis_row"].to_numpy(int)
            ols = fit_ols(x, y)
            maps = maps_file[f"maps_{study}"]
            require(
                maps.shape == (N_BOOTSTRAPS, len(frame)),
                f"Unexpected bootstrap shape for {study}: {maps.shape}",
            )
            require(np.issubdtype(maps.dtype, np.integer), "Bootstrap maps are not integers")
            require(int(maps.min()) >= 0 and int(maps.max()) < len(frame), "Bootstrap index out of range")
            boot = bootstrap_coefficients(x, y, maps)
            boot_rows.append(
                pd.DataFrame(
                    {
                        "source_study": study,
                        "bootstrap_id": np.arange(N_BOOTSTRAPS, dtype=int),
                        "alpha": boot[:, 0],
                        "beta": boot[:, 1],
                    }
                )
            )
            coefficients = np.asarray(ols["coefficients"], dtype=float)
            se = np.asarray(ols["se"], dtype=float)
            ci_low = np.asarray(ols["ci_low"], dtype=float)
            ci_high = np.asarray(ols["ci_high"], dtype=float)
            summaries.append(
                {
                    "source_study": study,
                    "n_independent_units": len(frame),
                    "alpha": coefficients[0],
                    "beta": coefficients[1],
                    "alpha_hc3_se": se[0],
                    "beta_hc3_se": se[1],
                    "alpha_hc3_ci_low": ci_low[0],
                    "alpha_hc3_ci_high": ci_high[0],
                    "beta_hc3_ci_low": ci_low[1],
                    "beta_hc3_ci_high": ci_high[1],
                    "hc3_interval_reference": f"Student-t(df={ols['df']})",
                    "r2": float(ols["r2"]),
                    "residual_mse": float(ols["residual_mse"]),
                    "alpha_bootstrap_ci_low": linear_quantile(boot[:, 0], 0.025),
                    "alpha_bootstrap_ci_high": linear_quantile(boot[:, 0], 0.975),
                    "beta_bootstrap_ci_low": linear_quantile(boot[:, 1], 0.025),
                    "beta_bootstrap_ci_high": linear_quantile(boot[:, 1], 0.975),
                    "bootstrap_draws": N_BOOTSTRAPS,
                    "bootstrap_seed": BOOTSTRAP_SEED,
                    "x_min": float(x.min()),
                    "x_max": float(x.max()),
                }
            )

            ordered_models: list[tuple[str, str, dict[str, Any]]] = [
                ("unconstrained_ols", f"{study}__unconstrained_ols", ols)
            ]
            for anchor, beta in (("host_anchor", 0.0), ("lineage_anchor", 1.0)):
                fixed = fit_fixed_beta(x, y, beta)
                anchors.append(
                    {
                        "source_study": study,
                        "anchor": anchor,
                        "n_independent_units": len(frame),
                        "fixed_beta": beta,
                        "fitted_alpha": float(fixed["coefficients"][0]),
                        "residual_mse": float(fixed["residual_mse"]),
                    }
                )
                ordered_models.append((anchor, f"{study}__{anchor}", fixed))

            for model_kind, model_id, model in ordered_models:
                model_coefficients = np.asarray(model["coefficients"], dtype=float)
                model_rows.append(
                    {
                        "model_id": model_id,
                        "source_study": study,
                        "model_kind": model_kind,
                        "n_independent_units": len(frame),
                        "alpha": model_coefficients[0],
                        "beta": model_coefficients[1],
                        "residual_mse": float(model["residual_mse"]),
                    }
                )
                fitted = np.asarray(model["fitted"], dtype=float)
                fitted_rows.append(
                    pd.DataFrame(
                        {
                            "model_id": model_id,
                            "source_study": study,
                            "analysis_row": row_ids,
                            "fitted": fitted,
                        }
                    )
                )
                objects[model_id] = {
                    "source_study": study,
                    "model_kind": model_kind,
                    "coefficients": model_coefficients,
                    "rows": row_ids,
                    "fitted": fitted,
                }
                if model_kind == "unconstrained_ols":
                    covariance = np.asarray(model["covariance"], dtype=float)
                    objects[model_id]["covariance"] = covariance
                    for row_index in range(2):
                        for column_index in range(2):
                            covariance_rows.append(
                                {
                                    "model_id": model_id,
                                    "row": row_index,
                                    "column": column_index,
                                    "value": covariance[row_index, column_index],
                                }
                            )

    return (
        pd.DataFrame(summaries),
        pd.DataFrame(anchors),
        pd.concat(boot_rows, ignore_index=True),
        pd.DataFrame(model_rows),
        pd.concat(fitted_rows, ignore_index=True),
        pd.DataFrame(covariance_rows),
        objects,
    )


def verify_recomputed_tables(
    summary: pd.DataFrame,
    anchors: pd.DataFrame,
    bootstrap: pd.DataFrame,
    models: pd.DataFrame,
    parity_rows: list[dict[str, Any]],
) -> None:
    saved_summary = pd.read_csv(REGRESSION / "regression_summary.tsv", sep="\t")
    summary_numeric = [
        column
        for column in summary.columns
        if column not in {"source_study", "hc3_interval_reference", "n_independent_units", "bootstrap_draws", "bootstrap_seed"}
    ]
    summary_error = compare_table(
        saved_summary,
        summary,
        ["source_study"],
        ["hc3_interval_reference", "n_independent_units", "bootstrap_draws", "bootstrap_seed"],
        summary_numeric,
    )
    add_parity(parity_rows, "regression_summary", "regression_summary.tsv", summary_error, "all reported OLS, HC3, bootstrap and range values")

    saved_anchors = pd.read_csv(REGRESSION / "constrained_anchor_summary.tsv", sep="\t")
    anchor_error = compare_table(
        saved_anchors,
        anchors,
        ["source_study", "anchor"],
        ["n_independent_units"],
        ["fixed_beta", "fitted_alpha", "residual_mse"],
    )
    add_parity(parity_rows, "anchor_summary", "constrained_anchor_summary.tsv", anchor_error, "host and lineage fixed-beta anchors")

    saved_models = pd.read_csv(MODEL_DIR / "closed_form_model_coefficients.tsv", sep="\t")
    model_error = compare_table(
        saved_models,
        models,
        ["model_id"],
        ["source_study", "model_kind", "n_independent_units"],
        ["alpha", "beta", "residual_mse"],
    )
    add_parity(parity_rows, "model_coefficient_table", "closed_form_model_coefficients.tsv", model_error, "all six persisted closed-form model rows")

    saved_bootstrap = pd.read_csv(REGRESSION / "bootstrap_coefficients.tsv.gz", sep="\t")
    bootstrap_error = compare_table(
        saved_bootstrap,
        bootstrap,
        ["source_study", "bootstrap_id"],
        [],
        ["alpha", "beta"],
    )
    add_parity(parity_rows, "bootstrap_coefficients", "bootstrap_coefficients.tsv.gz", bootstrap_error, "20,000 coefficient pairs reconstructed from persisted maps")


def verify_persisted_models(
    ready: pd.DataFrame,
    models: pd.DataFrame,
    objects: dict[str, dict[str, Any]],
    parity_rows: list[dict[str, Any]],
) -> dict[str, Any]:
    signature = json.loads((MODEL_DIR / "model_signature.json").read_text(encoding="utf-8"))
    expected_entry = ROOT / Path(str(signature["entry_point"]))
    require(expected_entry.resolve() == PRIMARY_SCRIPT.resolve(), "Persisted entry point changed")
    actual_entry_hash = sha256_file(PRIMARY_SCRIPT)
    require(actual_entry_hash == signature["entry_point_sha256"], "Primary entry-point hash mismatch")
    require(signature["feature_order"] == FEATURE_ORDER, "Persisted feature order changed")
    require(signature["study_order"] == list(STUDY_ORDER), "Persisted study order changed")
    require(int(signature["n_models"]) == 6, "Persisted model count changed")
    require(float(signature["parity_tolerance"]) == TOLERANCE, "Persisted parity tolerance changed")
    require(signature["preprocessing"] == "none", "Unexpected preprocessing")
    require(signature["selection"] == "none", "Unexpected selection")
    require(signature["weighting"] == "none", "Unexpected weighting")
    require(signature["calibration"] == "none", "Unexpected calibration")
    require(int(signature["bootstrap"]["draws"]) == N_BOOTSTRAPS, "Bootstrap draw count changed")
    require(int(signature["bootstrap"]["seed"]) == BOOTSTRAP_SEED, "Bootstrap seed changed")

    input_manifest = pd.read_csv(REGRESSION / "input_manifest.tsv", sep="\t")
    manifest_hashes = {
        str(row.relative_path): str(row.sha256) for row in input_manifest.itertuples(index=False)
    }
    require(signature["input_hashes"] == manifest_hashes, "Model signature input hashes differ")

    expected_order = models["model_id"].astype(str).tolist()
    with np.load(MODEL_DIR / "closed_form_model_state.npz", allow_pickle=False) as state:
        require(
            set(state.files) == {"model_id", "source_study", "model_kind", "coefficient", "feature_order"},
            f"Unexpected model-state keys: {state.files}",
        )
        model_ids = state["model_id"].astype(str).tolist()
        require(model_ids == expected_order, "Persisted model order changed")
        require(state["feature_order"].astype(str).tolist() == FEATURE_ORDER, "State feature order changed")
        require(
            state["source_study"].astype(str).tolist() == models["source_study"].astype(str).tolist(),
            "State study order differs",
        )
        require(
            state["model_kind"].astype(str).tolist() == models["model_kind"].astype(str).tolist(),
            "State model-kind order differs",
        )
        expected_coefficients = np.vstack([objects[model_id]["coefficients"] for model_id in model_ids])
        state_error = max_abs_difference(state["coefficient"], expected_coefficients)
        add_parity(parity_rows, "persisted_model_state", "closed_form_model_state.npz", state_error, "six coefficient vectors and exact model metadata")

    fitted_max = 0.0
    row_match = True
    with np.load(MODEL_DIR / "closed_form_fitted_values.npz", allow_pickle=False) as fitted_file:
        expected_keys = {f"pred_{i:02d}" for i in range(6)} | {f"rows_{i:02d}" for i in range(6)}
        require(set(fitted_file.files) == expected_keys, "Unexpected fitted-value NPZ keys")
        for index, model_id in enumerate(expected_order):
            rows = fitted_file[f"rows_{index:02d}"].astype(int)
            saved = fitted_file[f"pred_{index:02d}"].astype(float)
            expected_rows = objects[model_id]["rows"].astype(int)
            row_match = row_match and np.array_equal(rows, expected_rows)
            require(np.array_equal(rows, expected_rows), f"Persisted row identity changed for {model_id}")
            ready_indexed = ready.set_index("analysis_row")
            x = ready_indexed.loc[rows, "x_A_L_minus_A_R"].to_numpy(float)
            coefficient = objects[model_id]["coefficients"]
            independently_fitted = coefficient[0] + coefficient[1] * x
            fitted_max = max(fitted_max, max_abs_difference(saved, independently_fitted))
    add_parity(parity_rows, "persisted_fitted_predictions", "closed_form_fitted_values.npz", fitted_max, "all six models at exact persisted analysis-row identities")

    covariance_max = 0.0
    ols_indices = {"Holland2024__unconstrained_ols": "hc3_cov_00", "Soraas2019__unconstrained_ols": "hc3_cov_03"}
    with np.load(MODEL_DIR / "hc3_covariance.npz", allow_pickle=False) as covariance_file:
        require(set(covariance_file.files) == set(ols_indices.values()), "Unexpected HC3 covariance keys")
        for model_id, key in ols_indices.items():
            covariance_max = max(
                covariance_max,
                max_abs_difference(covariance_file[key], objects[model_id]["covariance"]),
            )
    add_parity(parity_rows, "persisted_hc3_covariance", "hc3_covariance.npz", covariance_max, "both independently reconstructed 2x2 HC3 covariance matrices")

    primary_artifact_audit = json.loads(
        (MODEL_DIR / "artifact_audit.json").read_text(encoding="utf-8")
    )
    require(primary_artifact_audit["status"] == "PASS", "Primary artifact audit is not PASS")
    require(primary_artifact_audit["reload_parity"] is True, "Primary reload parity not true")
    require(primary_artifact_audit["model_ids"] == expected_order, "Primary audit model IDs differ")

    run_audit = json.loads((REGRESSION / "run_audit.json").read_text(encoding="utf-8"))
    require(run_audit["status"] == "PASS", "Frozen regression run audit is not PASS")
    require(int(run_audit["n_holland_pairs"]) == 61, "Frozen Holland count differs")
    require(int(run_audit["n_soraas_recipients"]) == 19, "Frozen Soraas count differs")
    for relative, recorded_hash in run_audit["output_hashes"].items():
        path = ROOT / Path(str(relative))
        require(path.is_file(), f"Run-audit output missing: {relative}")
        require(sha256_file(path) == recorded_hash, f"Run-audit hash mismatch: {relative}")

    return {
        "status": "PASS",
        "primary_entry_point_sha256": actual_entry_hash,
        "n_models": 6,
        "model_order": expected_order,
        "feature_order": FEATURE_ORDER,
        "row_identity_exact": row_match,
        "max_abs_fitted_prediction_error": fitted_max,
        "max_abs_hc3_covariance_error": covariance_max,
        "max_abs_model_state_error": state_error,
    }


def verify_hannum_boundary() -> dict[str, Any]:
    decision = json.loads((HANNUM_DIR / "decision.json").read_text(encoding="utf-8"))
    expected = {
        "status": "NETWORK_TRANSFER_BLOCKED",
        "hannum_remote_manifest": "VERIFIED",
        "hannum_coverage_gate": "NOT_EVALUATED",
        "hannum_score": "NOT_CALCULATED",
        "hannum_regression": "NOT_RUN",
        "hannum_affiliation": "NOT_RUN",
        "full_file_hash_gate_relaxed": False,
        "third_clock_searched": False,
        "third_source_searched": False,
        "regression_branch": "PASS",
    }
    for key, value in expected.items():
        require(decision.get(key) == value, f"Hannum boundary field changed: {key}")
    run_audit = json.loads((HANNUM_DIR / "run_audit.json").read_text(encoding="utf-8"))
    require(run_audit["status"] == "NETWORK_TRANSFER_BLOCKED", "Hannum run status changed")
    require(run_audit["hannum_branch"] == decision, "Hannum run decision differs")
    network = json.loads(
        (HANNUM_DIR / "network_range_probe_audit.json").read_text(encoding="utf-8")
    )
    require(network["status"] == "NETWORK_TRANSFER_BLOCKED", "Network audit status changed")
    require(network["probe_is_formal_analysis_input"] is False, "Network probe became an analysis input")
    require(network["temporary_probe_bytes_persisted"] is False, "Temporary range bytes were persisted")
    require(network["validation"]["all_byte_counts_exact"] is True, "Range byte counts not exact")
    require(network["validation"]["all_content_ranges_exact"] is True, "Content ranges not exact")
    require(network["validation"]["all_status_206"] is True, "Range responses were not all 206")
    return {
        "status": "PASS",
        "decision": decision,
        "interpretation": (
            "NETWORK_TRANSFER_BLOCKED is solely a public-data transfer/throughput boundary. "
            "It is not a negative second-score result, not a coverage failure, and not "
            "evidence of absent score affiliation."
        ),
        "third_clock_or_source_search": False,
    }


def build_input_manifest() -> pd.DataFrame:
    paths = [
        Path(__file__).resolve(),
        PRIMARY_SCRIPT,
        SPEC,
        P2_LOCK,
        FROZEN_READY,
        REGRESSION / "artifact_manifest.tsv",
        REGRESSION / "regression_analysis_ready.tsv",
        REGRESSION / "bootstrap_indices.npz",
        HANNUM_DIR / "decision.json",
        HANNUM_DIR / "run_audit.json",
        HANNUM_DIR / "network_range_probe_audit.json",
    ]
    rows = []
    for path in paths:
        require(path.is_file(), f"Audit input missing: {path}")
        rows.append(
            {
                "relative_path": str(path.relative_to(ROOT)),
                "bytes": path.stat().st_size,
                "sha256": sha256_file(path),
            }
        )
    return pd.DataFrame(rows)


def write_tsv(frame: pd.DataFrame, path: Path, compressed: bool = False) -> None:
    compression: Any = {"method": "gzip", "mtime": 0} if compressed else None
    frame.to_csv(
        path,
        sep="\t",
        index=False,
        float_format="%.17g",
        compression=compression,
    )


def persist_and_reload(
    ready: pd.DataFrame,
    summary: pd.DataFrame,
    anchors: pd.DataFrame,
    bootstrap: pd.DataFrame,
    models: pd.DataFrame,
    fitted: pd.DataFrame,
    covariance: pd.DataFrame,
    parity: pd.DataFrame,
    hashes: pd.DataFrame,
    input_manifest: pd.DataFrame,
) -> dict[str, Any]:
    write_tsv(ready, OUT / "audited_regression_analysis_ready.tsv")
    write_tsv(summary, OUT / "recomputed_regression_summary.tsv")
    write_tsv(anchors, OUT / "recomputed_constrained_anchor_summary.tsv")
    write_tsv(bootstrap, OUT / "recomputed_bootstrap_coefficients.tsv.gz", compressed=True)
    write_tsv(models, OUT / "recomputed_model_coefficients.tsv")
    write_tsv(fitted, OUT / "recomputed_fitted_values.tsv.gz", compressed=True)
    write_tsv(covariance, OUT / "recomputed_hc3_covariance.tsv")
    write_tsv(parity, OUT / "parity_checks.tsv")
    write_tsv(hashes, OUT / "hash_verification.tsv")
    write_tsv(input_manifest, OUT / "input_manifest.tsv")

    checks: list[dict[str, Any]] = []

    def reload_table(
        filename: str,
        expected: pd.DataFrame,
        keys: list[str],
        exact: list[str],
        numeric: list[str],
    ) -> None:
        observed = pd.read_csv(OUT / filename, sep="\t")
        error = compare_table(observed, expected, keys, exact, numeric)
        require(error <= TOLERANCE, f"Own artifact reload failed: {filename}: {error}")
        checks.append({"artifact": filename, "max_abs_error": error, "status": "PASS"})

    reload_table(
        "audited_regression_analysis_ready.tsv",
        ready,
        ["source_study", "unit_id"],
        ["analysis_row"],
        ["A_R", "A_L", "score_S", "x_A_L_minus_A_R", "y_S_minus_A_R"],
    )
    reload_table(
        "recomputed_regression_summary.tsv",
        summary,
        ["source_study"],
        ["n_independent_units", "hc3_interval_reference", "bootstrap_draws", "bootstrap_seed"],
        [column for column in summary.columns if column not in {"source_study", "n_independent_units", "hc3_interval_reference", "bootstrap_draws", "bootstrap_seed"}],
    )
    reload_table(
        "recomputed_constrained_anchor_summary.tsv",
        anchors,
        ["source_study", "anchor"],
        ["n_independent_units"],
        ["fixed_beta", "fitted_alpha", "residual_mse"],
    )
    reload_table(
        "recomputed_bootstrap_coefficients.tsv.gz",
        bootstrap,
        ["source_study", "bootstrap_id"],
        [],
        ["alpha", "beta"],
    )
    reload_table(
        "recomputed_model_coefficients.tsv",
        models,
        ["model_id"],
        ["source_study", "model_kind", "n_independent_units"],
        ["alpha", "beta", "residual_mse"],
    )
    reload_table(
        "recomputed_fitted_values.tsv.gz",
        fitted,
        ["model_id", "analysis_row"],
        ["source_study"],
        ["fitted"],
    )
    reload_table(
        "recomputed_hc3_covariance.tsv",
        covariance,
        ["model_id", "row", "column"],
        [],
        ["value"],
    )
    require((pd.read_csv(OUT / "parity_checks.tsv", sep="\t")["status"] == "PASS").all(), "Reloaded parity table contains failure")
    require((pd.read_csv(OUT / "hash_verification.tsv", sep="\t")["status"] == "PASS").all(), "Reloaded hash table contains failure")
    reloaded_manifest = pd.read_csv(OUT / "input_manifest.tsv", sep="\t")
    require(reloaded_manifest.astype(str).equals(input_manifest.astype(str)), "Own input manifest reload differs")
    return {
        "status": "PASS",
        "tolerance": TOLERANCE,
        "artifact_checks": checks,
        "maximum_abs_reload_error": max(float(row["max_abs_error"]) for row in checks),
    }


def build_report(audit: dict[str, Any], summary: pd.DataFrame, anchors: pd.DataFrame) -> str:
    holland = summary.loc[summary["source_study"] == "Holland2024"].iloc[0]
    soraas = summary.loc[summary["source_study"] == "Soraas2019"].iloc[0]
    anchor_lookup = anchors.set_index(["source_study", "anchor"])
    return f"""# Independent audit: HSCT continuous referent regression

**Status:** PASS

This independent script did not import the primary analysis and generated no new random numbers. It reconstructed both regressions from the frozen 80-unit interface and the persisted 10,000-draw bootstrap maps. All 15 frozen artifact hashes passed, and all reported coefficients, HC3 covariance terms and intervals, bootstrap coefficients and linear quantiles, fixed-slope anchor mean squared errors, model states and fitted predictions matched within `1e-12`.

## Recomputed results

- Holland (61 donor-recipient pairs): beta = {holland['beta']:.6f}; HC3 95% CI {holland['beta_hc3_ci_low']:.6f} to {holland['beta_hc3_ci_high']:.6f}; bootstrap 95% interval {holland['beta_bootstrap_ci_low']:.6f} to {holland['beta_bootstrap_ci_high']:.6f}. Host-anchor MSE = {anchor_lookup.loc[('Holland2024', 'host_anchor'), 'residual_mse']:.6f}; lineage-anchor MSE = {anchor_lookup.loc[('Holland2024', 'lineage_anchor'), 'residual_mse']:.6f}.
- Søraas (19 recipients): beta = {soraas['beta']:.6f}; HC3 95% CI {soraas['beta_hc3_ci_low']:.6f} to {soraas['beta_hc3_ci_high']:.6f}; bootstrap 95% interval {soraas['beta_bootstrap_ci_low']:.6f} to {soraas['beta_bootstrap_ci_high']:.6f}. Host-anchor MSE = {anchor_lookup.loc[('Soraas2019', 'host_anchor'), 'residual_mse']:.6f}; lineage-anchor MSE = {anchor_lookup.loc[('Soraas2019', 'lineage_anchor'), 'residual_mse']:.6f}.

## Provenance and interpretation boundary

The frozen direction-lock hash drift is fully accounted for by the later Branch C governance update: replacing only that exact region reproduces the frozen byte count and SHA-256. No reconstructed lock was written.

Beta is a linear score-coordinate alignment coefficient. It is **not** a donor fraction or biological mixture fraction, does not identify lineage biological age, and cannot exclude host influence. Holland remains placement-driving; Søraas is separately processed complementary evidence and is not an independent replication. The geometry-preserving permutation analysis remains the primary non-parametric context; this regression is a sensitivity analysis.

`NETWORK_TRANSFER_BLOCKED` in the Hannum branch is a data-transfer/throughput boundary. Coverage was not evaluated, no Hannum score was calculated, and no second-score regression or affiliation analysis was run. It is not a negative result.

Maximum frozen-object parity error: {audit['parity']['maximum_abs_error']:.3g}. Maximum audit-artifact reload error: {audit['persistence']['maximum_abs_reload_error']:.3g}.
"""


def main() -> None:
    if OUT.exists():
        raise FileExistsError(f"Refusing to overwrite independent audit output: {OUT}")

    _, hash_checks = verify_frozen_manifest()
    frozen_input_manifest = pd.read_csv(REGRESSION / "input_manifest.tsv", sep="\t")
    provenance = verify_frozen_inputs(frozen_input_manifest)
    parity_rows: list[dict[str, Any]] = []
    ready = load_and_validate_ready(parity_rows)
    summary, anchors, bootstrap, models, fitted, covariance, objects = recompute_objects(ready)
    verify_recomputed_tables(summary, anchors, bootstrap, models, parity_rows)
    model_audit = verify_persisted_models(ready, models, objects, parity_rows)
    hannum_audit = verify_hannum_boundary()

    parity = pd.DataFrame(parity_rows)
    require((parity["status"] == "PASS").all(), "At least one parity check failed")
    maximum_parity = float(parity["max_abs_error"].max())
    require(maximum_parity <= TOLERANCE, "Maximum parity error exceeds tolerance")
    input_manifest = build_input_manifest()

    OUT.mkdir(parents=True, exist_ok=False)
    persistence = persist_and_reload(
        ready,
        summary,
        anchors,
        bootstrap,
        models,
        fitted,
        covariance,
        parity,
        hash_checks,
        input_manifest,
    )
    write_json(OUT / "provenance_lock_audit.json", provenance["direction_lock"])
    write_json(OUT / "artifact_reload_audit.json", persistence)

    audit = {
        "status": "PASS",
        "audit_version": AUDIT_VERSION,
        "audit_type": "independent_end_to_end_fixed_object_recalculation",
        "primary_script_imported": False,
        "new_random_numbers_generated": False,
        "new_models_or_scores_searched": False,
        "frozen_regression_modified": False,
        "tolerance": TOLERANCE,
        "independent_units": EXPECTED_COUNTS,
        "formula": "S-A_R = alpha + beta(A_L-A_R) + epsilon, separately by study",
        "bootstrap": {
            "draws_per_study": N_BOOTSTRAPS,
            "seed_recorded_by_primary_analysis": BOOTSTRAP_SEED,
            "persisted_maps_reused": True,
            "new_maps_generated": False,
            "quantile_method": "linear",
        },
        "frozen_manifest": {
            "status": "PASS",
            "verified_hashes": int(len(hash_checks)),
            "expected_hashes": 15,
        },
        "parity": {
            "status": "PASS",
            "n_checks": int(len(parity)),
            "maximum_abs_error": maximum_parity,
            "tolerance": TOLERANCE,
        },
        "model_artifacts": model_audit,
        "persistence": persistence,
        "direction_lock_provenance": provenance["direction_lock"],
        "hannum_boundary": hannum_audit,
        "recomputed_results": summary.to_dict(orient="records"),
        "recomputed_anchor_results": anchors.to_dict(orient="records"),
        "claim_boundaries": {
            "beta_is": "a linear score-coordinate alignment coefficient",
            "beta_is_not": [
                "a donor fraction",
                "a biological mixture fraction",
                "identification of lineage biological age",
                "evidence excluding host influence",
                "a causal or calibration parameter",
            ],
            "study_relationship": (
                "Holland is placement-driving; Soraas is separately processed "
                "complementary evidence and is not an independent replication."
            ),
            "primary_context": (
                "The geometry-preserving permutation analysis remains primary; "
                "the continuous regression is a conventional sensitivity."
            ),
        },
        "environment": {
            "python": sys.version,
            "executable": sys.executable,
            "platform": platform.platform(),
            "numpy": np.__version__,
            "pandas": pd.__version__,
            "scipy": scipy.__version__,
        },
        "artifact_manifest_excludes_self": True,
    }
    write_json(OUT / "independent_audit.json", audit)
    (OUT / "REPORT.md").write_text(build_report(audit, summary, anchors), encoding="utf-8")

    artifact_paths = sorted(
        [path for path in OUT.iterdir() if path.is_file() and path.name != "artifact_manifest.tsv"],
        key=lambda path: path.name,
    )
    manifest = pd.DataFrame(
        [
            {
                "relative_path": path.name,
                "role": "independent_audit_artifact",
                "bytes": path.stat().st_size,
                "sha256": sha256_file(path),
            }
            for path in artifact_paths
        ]
    )
    write_tsv(manifest, OUT / "artifact_manifest.tsv")
    reloaded_manifest = pd.read_csv(OUT / "artifact_manifest.tsv", sep="\t")
    require(len(reloaded_manifest) == len(artifact_paths), "Audit manifest row count differs")
    for row in reloaded_manifest.itertuples(index=False):
        path = OUT / str(row.relative_path)
        require(path.is_file(), f"Audit artifact missing after manifest write: {row.relative_path}")
        require(path.stat().st_size == int(row.bytes), f"Audit artifact size differs: {row.relative_path}")
        require(sha256_file(path) == str(row.sha256), f"Audit artifact hash differs: {row.relative_path}")

    print(
        json.dumps(
            {
                "status": "PASS",
                "output": str(OUT),
                "frozen_hashes_verified": len(hash_checks),
                "parity_checks": len(parity),
                "maximum_abs_parity_error": maximum_parity,
                "maximum_abs_reload_error": persistence["maximum_abs_reload_error"],
                "direction_lock_branch_c_only": True,
                "hannum_status": hannum_audit["decision"]["status"],
            },
            indent=2,
        ),
        flush=True,
    )


if __name__ == "__main__":
    main()
```
