## Supplementary_Software_1 for "Inferential boundaries of age prediction: why prediction does not establish biological age measurement": audit_nhanes_5y_complete_support_contextual_age_probe_v0_1.py

```
#!/usr/bin/env python
"""Independent numerical audit for the NHANES five-year successor probe.

This script fits no model. It recomputes data gates, participant losses,
pooled metrics, paired-bootstrap estimands and the frozen decision from stored
artifacts, then writes the final artifact manifest.
"""

from __future__ import annotations

import hashlib
import json
from pathlib import Path
from typing import Any

import numpy as np
import pandas as pd
from sklearn.metrics import brier_score_loss, log_loss, roc_auc_score


ROOT = Path(__file__).resolve().parents[1]
OUT = ROOT / "outputs" / "nhanes_5y_complete_support_contextual_age_probe_v0_1"
EXPECTED_N = 13_724
EXPECTED_EVENTS = 470
EXPECTED_CYCLES = {"2007_2008", "2009_2010", "2011_2012"}
EXPECTED_MODELS = ("M_W", "M_WA", "M_S1", "M_S1A", "M_S2c", "M_S2cA", "M_S3", "M_S3A")
PAIRS = {
    "S0": ("M_W", "M_WA"),
    "S1": ("M_S1", "M_S1A"),
    "S2c": ("M_S2c", "M_S2cA"),
    "S3": ("M_S3", "M_S3A"),
}
TOLERANCE = 1e-12


def sha256(path: Path) -> str:
    digest = hashlib.sha256()
    with path.open("rb") as handle:
        for block in iter(lambda: handle.read(1024 * 1024), b""):
            digest.update(block)
    return digest.hexdigest()


def write_json(path: Path, payload: Any) -> None:
    path.write_text(json.dumps(payload, indent=2, ensure_ascii=False) + "\n", encoding="utf-8")


def participant_loss(y: np.ndarray, probability: np.ndarray) -> np.ndarray:
    probability = np.clip(np.asarray(probability, float), 1e-7, 1 - 1e-7)
    return -(y * np.log(probability) + (1 - y) * np.log(1 - probability))


def maximum_error(values: list[float]) -> float:
    return float(max(values)) if values else 0.0


def main() -> int:
    required = [
        "analysis_ready_risk_set.tsv.gz",
        "cycle_support_audit.tsv",
        "participant_folds.tsv",
        "ordered_feature_tiers.tsv",
        "analysis_signature.json",
        "input_manifest.tsv",
        "model_artifact_index.tsv",
        "model_artifact_audit.json",
        "inner_tuning.tsv.gz",
        "inner_split_identities.tsv.gz",
        "oof_predictions.tsv.gz",
        "participant_loss_vectors.tsv.gz",
        "pooled_metrics.tsv",
        "fold_metrics.tsv",
        "age_band_calibration.tsv",
        "bootstrap_indices.npz",
        "bootstrap_contrasts.tsv.gz",
        "age_value_summary.tsv",
        "richest_change_summary.tsv",
        "decision.json",
        "contextual_age_decision.png",
        "contextual_age_decision.svg",
        "REPORT.md",
    ]
    missing = [name for name in required if not (OUT / name).exists()]
    if missing:
        raise RuntimeError(f"Missing audit inputs: {missing}")

    risk = pd.read_csv(OUT / "analysis_ready_risk_set.tsv.gz", sep="\t")
    support = pd.read_csv(OUT / "cycle_support_audit.tsv", sep="\t")
    folds = pd.read_csv(OUT / "participant_folds.tsv", sep="\t")
    tiers = pd.read_csv(OUT / "ordered_feature_tiers.tsv", sep="\t")
    oof = pd.read_csv(OUT / "oof_predictions.tsv.gz", sep="\t")
    losses = pd.read_csv(OUT / "participant_loss_vectors.tsv.gz", sep="\t")
    pooled = pd.read_csv(OUT / "pooled_metrics.tsv", sep="\t")
    age_summary = pd.read_csv(OUT / "age_value_summary.tsv", sep="\t")
    d_summary = pd.read_csv(OUT / "richest_change_summary.tsv", sep="\t")
    tuning = pd.read_csv(OUT / "inner_tuning.tsv.gz", sep="\t")
    artifact_index = pd.read_csv(OUT / "model_artifact_index.tsv", sep="\t")
    artifact_audit = json.loads((OUT / "model_artifact_audit.json").read_text(encoding="utf-8"))
    decision = json.loads((OUT / "decision.json").read_text(encoding="utf-8"))
    signature = json.loads((OUT / "analysis_signature.json").read_text(encoding="utf-8"))
    draws_file = np.load(OUT / "bootstrap_indices.npz")
    draws = draws_file["indices"]

    checks: dict[str, bool] = {}
    checks["risk_n"] = len(risk) == EXPECTED_N
    checks["risk_events"] = int(risk["death_5y"].sum()) == EXPECTED_EVENTS
    checks["risk_cycles"] = set(risk["cycle"].astype(str)) == EXPECTED_CYCLES
    checks["unique_SEQN"] = not risk["SEQN"].duplicated().any()
    checks["zero_early_event_free_censoring"] = int(risk["event_free_censored_before_60"].sum()) == 0
    checks["cycle_support_complete"] = bool(support["complete_support"].astype(bool).all())
    checks["fold_count"] = folds["outer_fold"].nunique() == 5 and set(folds["outer_fold"]) == set(range(5))
    checks["fold_identity"] = bool(
        folds.sort_values("row_id")["outer_fold"].to_numpy().tolist()
        == oof.sort_values("row_id")["outer_fold"].to_numpy().tolist()
    )
    checks["model_columns_exact"] = set(EXPECTED_MODELS).issubset(oof.columns)
    checks["tier_sizes"] = tiers.groupby("tier").size().to_dict() == {"S1": 6, "S2c": 30, "S3": 50}
    checks["tuning_models_exact"] = set(tuning["model"]) == set(EXPECTED_MODELS)
    checks["tuning_row_count"] = len(tuning) == 5 * 8 * 3 * 12
    checks["artifact_count"] = len(artifact_index) == 40
    checks["artifact_audit_pass"] = artifact_audit.get("status") == "PASS"
    checks["signature_present"] = len(signature.get("analysis_signature", "")) == 64
    checks["bootstrap_shape"] = draws.shape == (2000, EXPECTED_N)
    checks["bootstrap_bounds"] = bool(draws.min() >= 0 and draws.max() < EXPECTED_N)

    probability_bounds = []
    loss_errors = []
    pooled_errors = []
    y = oof["death_5y"].to_numpy(int)
    for model in EXPECTED_MODELS:
        probability = oof[model].to_numpy(float)
        probability_bounds.append(bool(np.isfinite(probability).all() and np.all((probability > 0) & (probability < 1))))
        recalculated_loss = participant_loss(y, probability)
        loss_errors.append(float(np.max(np.abs(recalculated_loss - losses[model].to_numpy(float)))))
        row = pooled.loc[pooled["model"].eq(model)].iloc[0]
        pooled_errors.extend(
            [
                abs(float(log_loss(y, probability, labels=[0, 1])) - float(row["log_loss"])),
                abs(float(brier_score_loss(y, probability)) - float(row["brier"])),
                abs(float(roc_auc_score(y, probability)) - float(row["auc"])),
            ]
        )
    checks["probabilities_valid"] = all(probability_bounds)
    checks["participant_loss_parity"] = maximum_error(loss_errors) <= TOLERANCE
    checks["pooled_metric_parity"] = maximum_error(pooled_errors) <= TOLERANCE

    loss_matrix = losses[list(EXPECTED_MODELS)].to_numpy(float)
    observed_means = loss_matrix.mean(axis=0)
    observed = dict(zip(EXPECTED_MODELS, observed_means))
    tier_names = list(PAIRS)
    point = {
        tier: observed[without_age] - observed[with_age]
        for tier, (without_age, with_age) in PAIRS.items()
    }
    boot = np.empty((draws.shape[0], len(tier_names)), float)
    for bootstrap in range(draws.shape[0]):
        means = loss_matrix[draws[bootstrap]].mean(axis=0)
        mapped = dict(zip(EXPECTED_MODELS, means))
        for index, tier in enumerate(tier_names):
            without_age, with_age = PAIRS[tier]
            boot[bootstrap, index] = mapped[without_age] - mapped[with_age]
    d_boot = boot[:, 0] - boot[:, -1]
    d_point = point["S0"] - point["S3"]

    summary_errors = []
    for index, tier in enumerate(tier_names):
        stored = age_summary.loc[age_summary["tier"].eq(tier)].iloc[0]
        summary_errors.extend(
            [
                abs(point[tier] - float(stored["age_value"])),
                abs(float(np.quantile(boot[:, index], 0.025)) - float(stored["q025"])),
                abs(float(np.quantile(boot[:, index], 0.975)) - float(stored["q975"])),
            ]
        )
    stored_d = d_summary.iloc[0]
    d_errors = [
        abs(d_point - float(stored_d["estimate"])),
        abs(float(np.quantile(d_boot, 0.025)) - float(stored_d["q025"])),
        abs(float(np.quantile(d_boot, 0.975)) - float(stored_d["q975"])),
    ]
    checks["age_value_summary_parity"] = maximum_error(summary_errors) <= TOLERANCE
    checks["richest_change_parity"] = maximum_error(d_errors) <= TOLERANCE

    v0_low = float(np.quantile(boot[:, 0], 0.025))
    d_low = float(np.quantile(d_boot, 0.025))
    d_high = float(np.quantile(d_boot, 0.975))
    if v0_low <= 0:
        independent_decision = "NO_VISIBLE_BASELINE_AGE_VALUE"
    elif d_low > 0:
        independent_decision = "CONTEXTUAL_SUBSTITUTION"
    elif d_high < 0:
        independent_decision = "CONTEXTUAL_SYNERGY"
    else:
        independent_decision = "NO_DETECTABLE_MIGRATION"
    checks["decision_parity"] = independent_decision == decision.get("decision_category")

    artifact_hash_errors = []
    artifact_missing = []
    for row in artifact_index.itertuples(index=False):
        path = ROOT / str(row.artifact_path)
        if not path.exists():
            artifact_missing.append(str(path))
            continue
        artifact_hash_errors.append(sha256(path) != str(row.artifact_sha256))
    checks["artifact_files_present"] = not artifact_missing
    checks["artifact_hash_parity"] = not any(artifact_hash_errors)
    checks["artifact_reload_parity_recorded"] = bool(
        artifact_audit.get("max_outer_prediction_error", 1) <= TOLERANCE
        and artifact_audit.get("max_inner_prediction_error", 1) <= TOLERANCE
    )

    passed = all(checks.values())
    audit = {
        "status": "PASS" if passed else "FAIL",
        "checks": checks,
        "independent_decision_category": independent_decision,
        "stored_decision_category": decision.get("decision_category"),
        "n": len(risk),
        "events": int(risk["death_5y"].sum()),
        "maximum_participant_loss_error": maximum_error(loss_errors),
        "maximum_pooled_metric_error": maximum_error(pooled_errors),
        "maximum_age_value_summary_error": maximum_error(summary_errors),
        "maximum_richest_change_error": maximum_error(d_errors),
        "artifact_missing": artifact_missing,
        "artifact_manifest_excludes_self": True,
        "note": "Independent audit fits no model and recomputes the decision from stored OOF losses and stored bootstrap indices.",
    }
    write_json(OUT / "independent_audit.json", audit)

    manifest_rows = []
    for path in sorted(OUT.rglob("*"), key=lambda item: str(item)):
        if not path.is_file() or path.name == "artifact_manifest.tsv":
            continue
        manifest_rows.append(
            {
                "path": str(path.relative_to(ROOT)),
                "bytes": path.stat().st_size,
                "sha256": sha256(path),
            }
        )
    pd.DataFrame(manifest_rows).to_csv(OUT / "artifact_manifest.tsv", sep="\t", index=False)

    print(json.dumps(audit, indent=2, ensure_ascii=False))
    return 0 if passed else 2


if __name__ == "__main__":
    raise SystemExit(main())
```
