## Supplementary_Software_1 for "Inferential boundaries of age prediction: why prediction does not establish biological age measurement": audit_nhanes_individual_agegap_contributions_v0_1.py

```
#!/usr/bin/env python
"""Independent saved-output audit for NHANES individual AgeGap contributions."""

from __future__ import annotations

import hashlib
import json
from pathlib import Path

import numpy as np
import pandas as pd


ROOT = Path(__file__).resolve().parents[1]
OUT = ROOT / "outputs" / "nhanes_individual_agegap_contributions_v0_1"
SOURCE_OOF = ROOT / "outputs" / "nhanes_5y_age_decoder_source_information_probe_v0_2" / "oof_predictions.tsv.gz"
TOL = 1e-10


def sha256(path: Path) -> str:
    digest = hashlib.sha256()
    with path.open("rb") as handle:
        for block in iter(lambda: handle.read(1024 * 1024), b""):
            digest.update(block)
    return digest.hexdigest()


def main() -> int:
    participants = pd.read_csv(OUT / "participant_agegap_profiles.tsv.gz", sep="\t")
    contributions = pd.read_csv(OUT / "participant_feature_contributions.tsv.gz", sep="\t")
    pairs = pd.read_csv(OUT / "matched_pairs.tsv.gz", sep="\t")
    summary = pd.read_csv(OUT / "matched_pair_distribution_summary.tsv", sep="\t").set_index("metric")
    selected = pd.read_csv(OUT / "selected_display_pairs.tsv", sep="\t")
    selected_contributions = pd.read_csv(OUT / "selected_pair_contributions.tsv", sep="\t")
    model_audit = json.loads((OUT / "model_artifact_audit.json").read_text(encoding="utf-8"))
    model_index = pd.read_csv(OUT / "model_artifact_index.tsv", sep="\t")
    source_oof = pd.read_csv(SOURCE_OOF, sep="\t")

    checks: dict[str, object] = {}
    checks["participant_count"] = int(len(participants))
    checks["participant_identity_unique"] = bool(participants["row_id"].is_unique and participants["SEQN"].is_unique)
    counts = contributions.groupby("row_id").size()
    checks["contribution_row_count"] = int(len(contributions))
    checks["exactly_50_contributions_per_participant"] = bool(counts.eq(50).all() and len(counts) == len(participants))
    sums = contributions.groupby("row_id")["contribution_years"].sum()
    expected = participants.set_index("row_id")["centered_agegap_years"].reindex(sums.index)
    checks["max_participant_contribution_sum_error"] = float(np.max(np.abs(sums.to_numpy() - expected.to_numpy())))

    source = participants[["row_id", "clock_score", "raw_agegap_years", "age_years"]].merge(
        source_oof[["row_id", "clock_score", "clock_gap"]], on="row_id", suffixes=("_saved", "_source"), validate="one_to_one"
    )
    checks["max_source_clock_score_error"] = float(np.max(np.abs(source["clock_score_saved"] - source["clock_score_source"])))
    checks["max_source_raw_agegap_error"] = float(np.max(np.abs(source["raw_agegap_years"] - source["clock_gap"])))
    checks["max_raw_agegap_identity_error"] = float(
        np.max(np.abs(source["raw_agegap_years"] - (source["clock_score_saved"] - source["age_years"])))
    )

    stacked_ids = pairs[["row_id_1", "row_id_2"]].to_numpy().ravel()
    checks["matched_pair_count"] = int(len(pairs))
    checks["matched_unique_participant_count"] = int(pd.unique(stacked_ids).size)
    checks["no_participant_reuse"] = bool(pd.Series(stacked_ids).is_unique)
    checks["max_pair_caliper"] = float(pairs["centered_agegap_abs_difference"].max())
    checks["max_pair_raw_centered_difference_error"] = float(
        np.max(np.abs(pairs["raw_agegap_abs_difference"] - pairs["centered_agegap_abs_difference"]))
    )

    p = participants.set_index("row_id")
    exact_columns = ["outer_fold", "age_years", "sex", "race_ethnicity", "cycle"]
    exact_ok = True
    for pair in pairs.itertuples(index=False):
        left = p.loc[int(pair.row_id_1)]
        right = p.loc[int(pair.row_id_2)]
        exact_ok = exact_ok and all(left[column] == right[column] for column in exact_columns)
    checks["all_pairs_exact_on_frozen_strata"] = bool(exact_ok)

    matrix = contributions.pivot(index="row_id", columns="feature_order", values="contribution_years")
    l2_errors = []
    l1_errors = []
    for pair in pairs.itertuples(index=False):
        delta = matrix.loc[int(pair.row_id_1)].to_numpy(float) - matrix.loc[int(pair.row_id_2)].to_numpy(float)
        l2_errors.append(abs(float(np.linalg.norm(delta)) - float(pair.contribution_l2_distance_years)))
        l1_errors.append(abs(float(np.abs(delta).sum()) - float(pair.contribution_l1_distance_years)))
    checks["max_recomputed_l2_error"] = float(max(l2_errors))
    checks["max_recomputed_l1_error"] = float(max(l1_errors))

    headline = {
        "median_gap_difference": float(pairs["centered_agegap_abs_difference"].median()),
        "median_profile_l2": float(pairs["contribution_l2_distance_years"].median()),
        "p95_profile_l2": float(pairs["contribution_l2_distance_years"].quantile(0.95)),
        "median_profile_cosine": float(pairs["contribution_cosine_similarity"].median()),
        "median_top5_jaccard": float(pairs["top5_absolute_contributor_jaccard"].median()),
        "median_direct_endpoint_logit_difference": float(pairs["M_X_abs_logit_difference"].median()),
        "median_agegap_endpoint_logit_difference": float(pairs["M_AGE_GAP_abs_logit_difference"].median()),
    }
    summary_map = {
        "median_gap_difference": ("centered_agegap_abs_difference", "median"),
        "median_profile_l2": ("contribution_l2_distance_years", "median"),
        "p95_profile_l2": ("contribution_l2_distance_years", "q95"),
        "median_profile_cosine": ("contribution_cosine_similarity", "median"),
        "median_top5_jaccard": ("top5_absolute_contributor_jaccard", "median"),
        "median_direct_endpoint_logit_difference": ("M_X_abs_logit_difference", "median"),
        "median_agegap_endpoint_logit_difference": ("M_AGE_GAP_abs_logit_difference", "median"),
    }
    checks["max_headline_summary_error"] = float(
        max(abs(value - float(summary.loc[metric, column])) for key, value in headline.items() for metric, column in [summary_map[key]])
    )

    eligible = pairs.loc[
        np.sign(pairs["centered_agegap_1"]).eq(np.sign(pairs["centered_agegap_2"]))
        & pairs["centered_agegap_mean"].abs().ge(2.0)
    ].copy()
    selection_ok = True
    for label, quantile in (("median_distance", 0.5), ("p95_distance", 0.95)):
        target = float(eligible["contribution_l2_distance_years"].quantile(quantile))
        expected_pair = eligible.assign(
            distance_to_target=(eligible["contribution_l2_distance_years"] - target).abs()
        ).sort_values(["distance_to_target", "pair_id"]).iloc[0]["pair_id"]
        actual_pair = selected.set_index("case").loc[label, "pair_id"]
        selection_ok = selection_ok and expected_pair == actual_pair
    checks["display_pair_selection_rule_reproduced"] = bool(selection_ok)
    case_sum = selected_contributions.groupby(["case", "person"])["contribution_years"].sum()
    case_errors = []
    selected_by_case = selected.set_index("case")
    for (case, person), value in case_sum.items():
        case_errors.append(abs(float(value) - float(selected_by_case.loc[case, f"centered_agegap_{int(person)}"])))
    checks["max_selected_case_remainder_error"] = float(max(case_errors))

    artifact_hash_ok = True
    for row in model_index.itertuples(index=False):
        path = ROOT / Path(row.artifact_path)
        artifact_hash_ok = artifact_hash_ok and path.exists() and sha256(path) == row.artifact_sha256
    checks["all_model_artifact_hashes_reproduced"] = bool(artifact_hash_ok)
    checks["model_artifact_audit_status"] = model_audit.get("status")
    checks["rerun_reused_all_residualizers"] = bool(
        model_audit.get("reused_residualizer_count") == 5 and model_audit.get("newly_fitted_residualizer_count") == 0
    )
    checks["figure_png_exists"] = (OUT / "figures" / "nhanes_individual_agegap_contributions_v0_1.png").exists()
    checks["figure_svg_exists"] = (OUT / "figures" / "nhanes_individual_agegap_contributions_v0_1.svg").exists()

    numerical_keys = [key for key in checks if key.startswith("max_")]
    pass_status = (
        checks["participant_count"] == 13_724
        and checks["contribution_row_count"] == 13_724 * 50
        and all(bool(checks[key]) for key in checks if isinstance(checks[key], bool))
        and all(float(checks[key]) <= TOL for key in numerical_keys if key != "max_pair_caliper")
        and float(checks["max_pair_caliper"]) <= 0.25 + TOL
        and checks["model_artifact_audit_status"] == "PASS"
    )
    payload = {
        "status": "PASS" if pass_status else "FAIL",
        "confidence_assessment": "Ready to share with explicit non-causal and target-specific caveats" if pass_status else "Needs revision",
        "audit_script": str(Path(__file__).relative_to(ROOT)),
        "audit_script_sha256": sha256(Path(__file__)),
        "checks": checks,
        "headline_recomputation": headline,
        "required_caveats": [
            "Contributions explain the fitted decoder and are not causal decompositions of biological ageing.",
            "The conditional expectation is a fold-local linear Ridge estimate, not a known population conditional mean.",
            "Feature-specific magnitudes belong to this frozen correlated-feature Ridge parameterization and are not stable measures of biological importance.",
            "Endpoint-score contrasts are model-specific five-year mortality predictions, not causal or calibrated individual effects.",
            "The matched-pair analysis is descriptive for the complete-support NHANES cohort and does not use survey weights for population prevalence.",
        ],
    }
    (OUT / "validation_audit.json").write_text(json.dumps(payload, indent=2, ensure_ascii=False) + "\n", encoding="utf-8")
    report = f"""# Validation audit: NHANES individual AgeGap contributions

**Overall assessment: {payload['confidence_assessment']}**

- participant profiles: {checks['participant_count']:,}
- long contribution rows: {checks['contribution_row_count']:,}
- matched pairs / unique participants: {checks['matched_pair_count']:,} / {checks['matched_unique_participant_count']:,}
- maximum participant contribution-sum error: {checks['max_participant_contribution_sum_error']:.3g}
- maximum independently recomputed L2-distance error: {checks['max_recomputed_l2_error']:.3g}
- maximum headline-summary error: {checks['max_headline_summary_error']:.3g}
- frozen decoder and residualizer audit: {checks['model_artifact_audit_status']}
- routine rerun reused all five residualizers: {checks['rerun_reused_all_residualizers']}
- display selection rule reproduced: {checks['display_pair_selection_rule_reproduced']}

No calculation or persistence blocker was found. The required interpretation
caveats are recorded in `validation_audit.json` and the main report.
"""
    (OUT / "VALIDATION_REPORT.md").write_text(report, encoding="utf-8")

    manifest_rows = []
    for path in sorted(OUT.rglob("*")):
        if path.is_file() and path.name != "artifact_manifest.tsv":
            manifest_rows.append(
                {"path": str(path.relative_to(ROOT)), "bytes": path.stat().st_size, "sha256": sha256(path)}
            )
    pd.DataFrame(manifest_rows).to_csv(OUT / "artifact_manifest.tsv", sep="\t", index=False)
    print(json.dumps(payload, indent=2, ensure_ascii=False))
    return 0 if pass_status else 2


if __name__ == "__main__":
    raise SystemExit(main())
```
