## Supplementary_Software_1 for "Inferential boundaries of age prediction: why prediction does not establish biological age measurement": HSCT_FIXED_SCORE_REFERENT_NULL_PROBE_SPEC_20260823.md

# HSCT fixed-score referent-null probe

Date frozen: 23 August 2026  
Status: author-authorized decision probe  
Module: HUM-01 placement revision  
Primary role: materiality placement gate  
Analysis version: `hsct_fixed_score_referent_null_probe_v0_1`

## 1. Decision served

The sole decision is whether the existing HSCT fixed-score referent result can
remain a main human result or must move to Extended Data after accounting for
the non-zero lineage-side baseline created by donor/recipient age geometry and
score shrinkage.

The probe does not rescore methylation, fit an AgeClock, add clock families,
estimate biological age, test rejuvenation or prove the analytical proposition.

## 2. Frozen inputs

Only these persisted pair/recipient summaries may be used:

- `outputs/hsct_clock_object_replacement_v0_1/primary_recipient_contrasts.tsv`;
- `outputs/hsct_holland2024_horvath_object_affiliation_v0_1/primary_pair_summary.tsv`.

The two studies partly overlap and are analysed separately. They are not
pooled as independent replications.

## 3. Objects and independent unit

For every independent recipient or donor-recipient pair:

- (H): recipient-host chronological age at sampling;
- (C=A_D+t): donor-lineage chronological coordinate;
- (S): fixed age-valued recipient-blood readout.

For both studies compute the common bounded affiliation statistic

\[
I=\frac{|S-H|-|S-C|}{|H-C|}.
\]

Positive values favour the lineage-coordinate side. The statistic is bounded
and saturating; it is not normalized age accuracy.

The independent unit is recipient for Søraas 2019 and donor-recipient pair for
Holland 2024. No sample or occasion is treated as independent after the frozen
aggregation.

Define two strata before calculation:

- `host_older`: (H-C>0);
- `host_younger`: (H-C<0).

Pairs with (H=C) are ineligible; none may be assigned to a stratum by an
arbitrary rule.

## 4. Primary observed statistic

The primary statistic in each study is sign-balanced affiliation:

\[
I_{bal}=\frac{1}{2}
\left\{
\overline I_{host\ older}+
\overline I_{host\ younger}
\right\}.
\]

This gives equal weight to the two age-difference directions and prevents the
larger stratum from determining the result. Also report pair-weighted overall
mean (I) and both stratum means as secondary summaries.

## 5. Frozen nulls and diagnostics

### Primary permutation null

Within each age-difference stratum, permute the fixed score (S) across
independent units while holding each unit's (H,C) pair fixed. This breaks
score–referent pairing while preserving:

- the observed score marginal distribution within sign;
- the observed host and lineage-coordinate distributions;
- stratum sizes;
- the frozen independent unit.

Use 20,000 permutations with NumPy generator seed `20260823`. Apply the same
permutation indices to every recomputation saved for audit. For each permutation
compute stratum means and (I_{bal}).

### Constant-score shrinkage controls

Within each study, replace every score by:

1. the observed marginal mean score;
2. the observed marginal median score.

Compute overall, stratum-specific and sign-balanced affiliation. These are
geometric controls, not null distributions.

### Calibration diagnostics

Report ordinary least-squares slopes and intercepts for:

- (S\sim C);
- (S\sim H);

overall and within each age-difference stratum. These are descriptive
fixed-score diagnostics. They are not used to recalibrate scores and do not
carry a causal interpretation.

## 6. Uncertainty and p-values

Use 10,000 stratified independent-unit bootstraps with seed `20260824` for the
observed sign-balanced statistic and observed-minus-permutation-null-mean
contrast. Resample independently within the two sign strata and preserve the
original stratum sizes.

Permutation p-values use the finite-sample correction

\[
p=\frac{1+\#\{I_{bal}^{perm}\ge I_{bal}^{obs}\}}{1+20{,}000}.
\]

P-values are placement diagnostics rather than evidence for a biological
entity. Do not select a one- versus two-sided convention after seeing the
result; the directional one-sided test is frozen because the existing claim is
lineage-side affiliation.

## 7. Placement gate

Classify `MAIN` only if all conditions hold:

1. Holland 2024 observed (I_{bal}) exceeds the 97.5th percentile of its
   within-sign permutation null;
2. the observed-minus-null-mean bootstrap interval is above zero in Holland;
3. observed affiliation exceeds the permutation-null mean in both Holland sign
   strata;
4. Søraas 2019 has the same sign for observed-minus-null-mean (I_{bal});
5. score-on-(C) slopes are positive in both Holland sign strata;
6. all source reconstruction and deterministic checks pass.

Classify `EXTENDED_DATA` if the inputs and computations pass but any placement
condition fails. Classify `STOP` only if input identity, independent-unit
construction or deterministic reproduction fails.

No threshold, stratum, permutation rule or statistic may be changed to rescue
placement.

## 8. Required artifacts

Write to `outputs/hsct_fixed_score_referent_null_probe_v0_1/`:

- `analysis_ready_pairs.tsv` with source-study identifier and reconstructed
  (H,C,S,I);
- `input_manifest.tsv` with paths, sizes and hashes;
- `permutation_indices.npz`;
- `permutation_statistics.tsv.gz`;
- `bootstrap_indices.npz`;
- `bootstrap_statistics.tsv.gz`;
- `observed_summary.tsv`;
- `constant_score_controls.tsv`;
- `calibration_diagnostics.tsv`;
- `decision.json`;
- `deterministic_audit.json`;
- `artifact_manifest.tsv`;
- one compact decision figure in PNG and SVG;
- one decision-ready report.

Reload the saved index arrays and reproduce all reported permutation and
bootstrap summaries elementwise or to a declared tolerance. A parity failure is
`STOP`.

## 9. Claim boundary

If `MAIN`, the allowed conclusion is:

> The evaluated post-HSCT recipient-blood readouts affiliated with the
> donor-derived haematopoietic lineage's chronological coordinate beyond the
> affiliation expected from the observed marginal score and donor/recipient age
> geometry.

This does not identify a unique lineage biological age, exclude host influence,
establish organismal age replacement, or validate rejuvenation.

## 10. Stop rule

Stop after the two frozen inputs, two sign strata, two constant-score controls,
the declared permutation/bootstrap procedures and audit. Do not add clocks,
alternative age-gap thresholds, calibration functions, external cohorts or
outcomes.
