## Supplementary_Software_1 for "Inferential boundaries of age prediction: why prediction does not establish biological age measurement": HSCT_REFERENT_REGRESSION_CLOCK_SENSITIVITY_SPEC_20260826.md

# HSCT referent regression and second-score sensitivity specification

Date frozen: 26 August 2026  
Analysis version: `hsct_referent_regression_clock_sensitivity_v0_1`

## Decision

Test whether the existing HSCT referent result remains visible under a
conventional continuous regression estimand and, if a fixed cross-batch probe
panel passes a coverage gate, a second chronological-age coefficient score.

## Primary regression

At the existing independent-unit level define recipient age `A_R`, donor-lineage
chronology `A_L` and fixed recipient-blood score `S`. Fit separately by study:

`S - A_R = alpha + beta (A_L - A_R) + epsilon`.

- Holland: 61 donor-recipient pairs from the frozen primary pair summary.
- Soeraas: 19 recipients from the frozen primary recipient summary.
- Report OLS beta and alpha, HC3 intervals, R2 and a 10,000-draw
  independent-unit bootstrap interval with seed `20260826`.
- Report constrained residual MSE for host-side `beta=0` and lineage-side
  `beta=1`, each with its own fitted intercept, as descriptive anchors.
- Do not pool studies or treat them as independent replication.

## Public-data gate

The Mendeley release contains six separately ENmix-processed beta matrices and
`comRand.csv`; it does not contain IDAT files. Raw-IDAT reprocessing is therefore
recorded as unavailable in the public interface.

## Second-score gate

- Candidate fixed in advance: Hannum 2013 chronological-age coefficients,
  71 probes.
- Stream only the 71 requested rows while byte-counting and validating each of
  the six public beta matrices.
- Prefer the complete 71-probe formula. If it is not common to all batches, one
  common-probe coefficient implementation is allowed only when the identical
  intersection is present in all six batches, contains at least 65 probes and
  uses zero cohort-derived imputation.
- If the gate fails, classify `SECOND_SCORE_COVERAGE_STOP`; do not search a
  third clock.
- If it passes, compute the fixed score for every array, aggregate to the same
  primary Holland pair interface and rerun the regression and frozen
  geometry-preserving affiliation summaries. Call it a coefficient-score
  sensitivity, not a paper-identical reconstruction unless the complete
  formula is available.

## Controls and boundaries

Donor arrays and the predeclared stable post-transplant window may be used as
diagnostics. Pre-transplant recipient blood and Holland chimerism are absent.
No result can establish lineage biological age, exclude host influence, prove
the analytical theorem or generalize to all clocks.
