## Supplementary_Software_1 for "Inferential boundaries of age prediction: why prediction does not establish biological age measurement": NHANES3_5Y_AGE_DECODER_SUBSTITUTION_COMPARISON_SPEC_v0_1_20260827.md

# NHANES III five-year mortality: chronological age versus an age decoder

## Frozen status

- Analysis version: `nhanes3_5y_age_decoder_substitution_comparison_v0_1`
- Status: retrospective, outcome-accessed, fixed protocol-transfer robustness analysis
- Source representation: the Phase 1-fitted `q_A` frozen in
  `nhanes3_5y_matched_scalar_target_supervision_protocol_transfer_v0_2`
- Target evaluation population: the already frozen NHANES III Phase 2 five-year
  mortality interface, participant folds, survey weights and Fay-BRR design
- This analysis is separate from v0.2. It must not modify v0.2, a manuscript,
  or any other frozen evidence module.

The five-year endpoint has already been accessed in this project. This is
therefore not an untouched confirmation. No endpoint, horizon, population,
feature, model-family or tuning-grid search is allowed.

## Question and estimand

Does the frozen age-decoder scalar replace known chronological age as a better
one-dimensional input for predicting five-year all-cause mortality, after the
same background variables are supplied?

Fit and compare only:

- `M_AGE`: `Y ~ W + s(A)`
- `M_QA`: `Y ~ W + s(q_A)`

Here `Y` is resolved five-year all-cause mortality, `W` is sex and
race/ethnicity, `A` is observed chronological age at examination, and `q_A` is
the Phase 1-fitted age decoder applied unchanged to Phase 2.

The primary contrast is

`Delta_sub = L(M_AGE) - L(M_QA)`,

where `L` is design-weighted held-out log loss. Positive values favour `q_A`;
negative values favour observed age.

This is a direct substitution comparison. Neither model may contain both `A`
and `q_A`. No `q_Y`, source-measurement model or additional performance metric
belongs in this analysis.

## Equal modelling opportunity

Both arms use exactly the same target-local pipeline:

1. one-hot encode `W` within the training fold;
2. weighted-standardize the single continuous scalar within the training fold;
3. transform that scalar with a five-knot uniform cubic spline, without a bias
   column and with linear extrapolation;
4. fit weighted L2-penalized logistic regression;
5. tune `C` by the same three-fold inner-validation log loss over
   `{0.001, 0.003, 0.01, 0.03, 0.1, 0.3, 1, 3, 10, 30, 100, 300}`;
6. pool inner-fold weighted loss numerators and denominators, and break ties
   within `1e-12` in favour of the smaller `C`.

The fixed five outer folds provide out-of-fold predictions. All preprocessing,
spline knots, tuning and fitting are training-fold local. Both arms use the
same participants, outcomes, folds and weights.

## Survey-design uncertainty and decision rule

Run the primary fit with the MEC examination weights. For each of the 32 frozen
Fay-BRR replicates (`rho=0.3`, 23 design degrees of freedom), repeat inner
tuning, outer fitting and held-out prediction for both arms using the replicate
weights. Do not refit or retune the frozen Phase 1 age-decoder direction.

Report the full-sample contrast, its 95% Fay-BRR interval and the number of the
five fixed outer-fold contrasts that are positive.

- `QA_SUBSTITUTION_ADVANTAGE` requires `Delta_sub > 0`, a positive lower
  interval bound and at least four of five positive outer-fold contrasts.
- `AGE_SUBSTITUTION_ADVANTAGE` requires `Delta_sub < 0`, a negative upper
  interval bound and at least four of five negative outer-fold contrasts.
- Otherwise report `NO_CLEAR_SUBSTITUTION_ADVANTAGE`.
- Any schema, identity, weight, fold, cache-signature or reload-parity failure
  is `STOP_DATA_OR_ARTIFACT_FAIL`; no rescue analysis follows.

## Persistence contract

Persist one signed joblib artifact for every model, outer fold and weight set:
`33 x 2 x 5 = 330` downstream artifacts. Each artifact must contain its
training-local encoder, scalar standardizer, spline transformer, all inner
candidate estimators and predictions, selected `C`, final estimator, ordered
identities, weights, derived designs and held-out predictions.

Immediately reload every artifact and verify design and prediction parity
elementwise to tolerance `1e-12`. A compatible rerun must reuse all 330
artifacts. An incompatible artifact must fail closed rather than be overwritten.

## Claim ceiling

A positive result would show only that, for this frozen NHANES III population,
five-year endpoint, transported age decoder and matched one-scalar model family,
the age-decoder scalar predicted the endpoint better than observed age after the
same background adjustment. It would not identify biological age, establish
general superiority over chronological age, prove causal or surrogate validity,
or apply to every clock or endpoint.
