## Supplementary_Software_1 for "Inferential boundaries of age prediction: why prediction does not establish biological age measurement": NHANES3_5Y_CHRONOLOGICAL_AGE_INCREMENT_AND_PROXY_COMPARISON_SPEC_v0_1_20260827.md

# NHANES III five-year chronological-age increment and proxy comparison v0.1

Date frozen: 27 August 2026  
Analysis version: `nhanes3_5y_chronological_age_increment_and_proxy_comparison_v0_1`

## Frozen status and question

This is a retrospective, outcome-accessed, fixed protocol-transfer robustness
analysis. NHANES III Phase 2 five-year mortality outcomes have already been
accessed by predecessor analyses. This module is therefore not an untouched
confirmation and must not be described as an independent external validation.

The fixed questions are:

1. How much held-out predictive value does observed chronological age add after
   sex, race/ethnicity and the complete frozen 25-measurement clinical panel are
   already supplied?
2. Does chronological age become more predictively valuable after removing the
   five-measurement block that an earlier Phase 1 analysis selected for age
   reconstruction?

This analysis does not use an age-decoder score, an endpoint-supervised scalar,
an age gap or any derived-age input. In particular, `q_A` and `q_Y` are
prohibited as inputs.

## Frozen data interface

- Target population: the canonical NHANES III Phase 2 analysis-ready interface,
  7,774 participants and 392 resolved five-year all-cause deaths.
- Age range: examination age `[20,80)` years.
- Outcome `Y`: resolved five-year all-cause mortality.
- Background `W`: sex and race/ethnicity.
- Full measurement panel `X25`: all 25 measurements in the canonical ordered
  `ordered_5x5_feature_manifest.tsv`, in `feature_order=1,...,25`.
- Survey design: `analysis_weight=mec_weight`, 23 released strata and 46 PSU
  clusters.
- Validation design: the existing PSU-atomic five outer folds and the
  outer-fold-specific three inner folds.
- Survey uncertainty: the existing 32 Phase 2 Fay-BRR factor vectors with
  `rho=0.3` and 23 design degrees of freedom.
- Missing measurements: training-fold-local weighted-median imputation.
  Complete-case restriction is prohibited.

The canonical inputs are read only from
`outputs/nhanes3_phase1_to_phase2_endpoint_vs_age_acquisition_v0_2_final_20260826`.
The new analysis must not modify that directory or any predecessor module.

## Frozen Phase 1 age-selected block

The removed block is taken unchanged from the signed
`source_selection_freeze.json` in the canonical directory. The freeze must:

- have analysis identity
  `nhanes3_phase1_to_phase2_endpoint_vs_age_acquisition_v0_2`;
- have status `PASS_POLICY_CONTRAST`;
- identify `cardiorespiratory` as `age_selected_domain`;
- reproduce its own `freeze_signature`; and
- map that domain, in this exact order, to:

  1. `MEC_SBP_MEAN`
  2. `MEC_DBP_MEAN`
  3. `PEP6DR`
  4. `SPPFEV1`
  5. `SPPFVC`

The reduced panel `X20_remaining` is the ordered full panel after removing
exactly those five variables. Its 20 measurements are retained in their
original manifest order.

The five variables are called the **Phase 1 age-selected block**. The remaining
20 variables are called the **remaining-20** or **reduced panel**. Neither this
selection nor this analysis establishes that every removed variable is
individually age-associated, and the remaining 20 must never be called
"non-age-related".

The mask is frozen before any new Phase 2 model is fitted. No Phase 2 outcome,
coefficient, association, missingness pattern or performance result may alter
the mask.

## Four locked endpoint models

Fit exactly four target-local models:

- `F0`: `Y ~ W + X25`
- `F1`: `Y ~ W + X25 + s(A)`
- `R0`: `Y ~ W + X20_remaining`
- `R1`: `Y ~ W + X20_remaining + s(A)`

Here `A` is observed chronological age at examination. `s(A)` is a cubic spline
with fixed knots at ages `20,35,50,65,80`, no bias column and linear
extrapolation. The spline basis is standardized with training-fold normalized
MEC weights. It is absent from `F0` and `R0`.

Within every training fold and independently for every model and weight set:

1. one-hot encode `W`, with unknown held-out levels ignored;
2. weighted-median impute the model's ordered measurement panel;
3. weighted-standardize the imputed measurements;
4. for `F1` and `R1`, construct and weighted-standardize the fixed-knot age
   basis;
5. concatenate the ordered design as background, measurements, then age basis;
6. fit weighted L2-penalized logistic regression; and
7. tune `C` by pooled design-weighted inner-validation log loss over
   `{0.001,0.003,0.01,0.03,0.1,0.3,1,3,10,30,100,300}`.

Inner losses are pooled as the sum of weighted loss numerators divided by the
sum of weight denominators. Within a `1e-12` loss tolerance, choose the smaller
`C`. Fitting weights are normalized to sum to the number of observations in the
corresponding fit. Probabilities are clipped to `[1e-8,1-1e-8]` for log-loss
evaluation. Every reported probability is outer-fold held out.

No interaction, feature selection, nonlinear measurement transformation,
alternative age basis, endpoint, horizon, age range, tuning grid or model
family may be added after this freeze.

## Locked estimands

Let `L(M)` be the design-weighted mean participant-level outer-fold log loss for
model `M`; positive contrasts below favour the richer or substituting input.

### Incremental chronological-age value

- Full panel: `V_F = L(F0) - L(F1)`.
- Remaining-20 panel: `V_R = L(R0) - L(R1)`.

`V_F` is the primary answer to how much predictive value chronological age adds
after the complete measured interface. `V_R` is the corresponding value after
the frozen age-selected block is unavailable.

### Predictive proxy-substitution contrast

The proxy contrast is

`P_proxy = V_R - V_F`

which is algebraically identical to

`[L(R0)-L(F0)] - [L(R1)-L(F1)]`.

A positive value means that chronological age becomes more useful after the
Phase 1 age-selected block is removed, or equivalently that supplying age
attenuates part of the held-out predictive value of that omitted block under the
declared estimators. This is an information-set-dependent predictive
substitution pattern, not causal mediation and not evidence that age is the
biological state represented by the omitted measurements.

Also report:

- block value without age: `D_block_0 = L(R0)-L(F0)`;
- block value with age: `D_block_1 = L(R1)-L(F1)`; and
- replacement comparison: `D_replace = L(R1)-L(F0)`.

Ordinary failure to reject `D_replace=0` does not establish equivalence or
non-inferiority. No equivalence claim is permitted because no clinical margin
is frozen in this specification.

All estimands are formed first as participant-level paired loss vectors and
then design-weighted. Calling these finite-estimator contrasts "incremental
predictive value" is permitted; calling them literal conditional mutual
information or a fraction of ageing is not.

## Staged execution

### `--stage primary`

Fit the four models in the five fixed outer folds using the base MEC weights:
`4 x 5 = 20` artifacts. Persist the OOF probabilities, participant losses,
paired contrast vectors, tuning results, fold contrasts, coefficient ledger,
artifact index and artifact parity audit. Write a primary probe table and exit.

The primary probe has no Fay-BRR interval and cannot be labelled formal survey
inference. Its purpose is to expose the fixed point estimates and validate the
implementation before the expensive branch.

Promote the probe to `--stage all` when at least one declared question has a
coherent point-estimate pattern:

- chronological-age increment: `V_F > 0` and at least four of five fixed
  outer-fold `V_F` contrasts are positive; or
- predictive substitution: `D_block_0 > 0`, `P_proxy > 0` and at least four of
  five fixed outer-fold `P_proxy` contrasts are positive.

Otherwise classify the probe as absent or ambiguous and stop rather than
changing the mask, endpoint, spline, model family or folds to rescue it. This
promotion rule governs computational escalation only; it is not a formal
significance claim. The primary TSV, JSON and report must record both branch
decisions, the resulting classification and a machine-readable
`promotion_permitted` Boolean. A requested `--stage all` must exit before any
Fay-BRR fit when that Boolean is false.

### `--stage all`

First validate and reuse all 20 compatible primary artifacts. Then, for each of
the 32 frozen Fay-BRR weight sets, repeat inner tuning, outer fitting, held-out
prediction and all paired-loss calculations for all four arms:

`32 x 4 x 5 = 640` additional artifacts.

The completed formal run therefore contains exactly `660` model artifacts.
For each locked contrast, report the full-sample estimate, Fay-BRR standard
error and two-sided 95% interval using

`sum_r (theta_r-theta)^2 / [32*(1-0.3)^2]`

and a `t_23` critical value. Also report the sign of each of the five fixed
outer-fold contrasts. The intervals represent Phase 2 survey-design uncertainty
conditional on the frozen mask, endpoint, folds and estimator family; they do
not include Phase 1 mask-development uncertainty.

No scientific result is gated solely on statistical significance. The formal
decision object records estimates, intervals and prespecified sign diagnostics
without turning a non-significant difference into proof of equality.

## Persistence and fail-closed contract

Persist one signed joblib artifact per model, outer fold and weight set. Every
artifact must contain:

- the complete ordered model specification and signature;
- training and held-out participant identities, outcomes and weights;
- every inner-fold identity assignment;
- every fitted encoder, imputer, scaler and age spline;
- every inner candidate logistic estimator and its validation predictions and
  loss vector;
- the selected `C`;
- the final fitted estimator;
- fitted design matrices, feature names and held-out predictions; and
- environment and software provenance.

Immediately reload every artifact and reconstruct all stored designs, tuning
losses and predictions elementwise to tolerance `1e-12`. Persist a
human-readable coefficient ledger for every final fold estimator. A compatible
rerun must reuse the artifact. A missing object, changed signature, changed
feature order or parity failure stops the analysis. An incompatible cache is
never overwritten and cannot be rescued within the same versioned directory.

Persist the exact canonical inputs used, the frozen mask, analysis-ready target
interface, folds, Fay factors, all result tables, decision objects and
machine-readable manifests. The primary and all-stage audits must state the
expected and observed artifact counts explicitly.

## Claim ceiling

This analysis can quantify whether observed chronological age adds held-out
predictive value after a declared clinical representation and whether that
value increases when a previously age-selected measurement block is omitted.
It cannot establish biological-age identity, causal mediation, mechanism,
surrogate validity, clinical utility, universal measured-state sufficiency or
that the remaining-20 panel is free of age-related information.
