## Supplementary_Software_1 for "Inferential boundaries of age prediction: why prediction does not establish biological age measurement": NHANES3_5Y_MATCHED_SCALAR_TARGET_SUPERVISION_PROTOCOL_TRANSFER_SPEC_v0_2_20260827.md

# NHANES III five-year matched-scalar target-supervision protocol transfer v0.2

Date frozen: 27 August 2026  
Analysis version: `nhanes3_5y_matched_scalar_target_supervision_protocol_transfer_v0_2`

## Revision provenance

The v0.1 source models and numerical results passed all model-object reload and
prediction-parity checks, but its first compatible cache rerun stopped when a
compressed TSV round trip changed large tuning-loss numerators by roughly one
machine-precision unit. V0.1 remains unchanged as audit provenance and is not a
complete formal run. V0.2 changes only the persistence comparison for
human-readable numeric tables: it retains absolute tolerance `1e-12` and adds
relative tolerance `1e-14` for serialized numeric cells. Model designs,
predictions, losses reconstructed directly from joblib objects, estimands,
grids, folds, survey inference and decision gates are unchanged and remain
subject to the absolute `1e-12` parity requirement.

## Status and question

This is a retrospective, outcome-accessed **fixed protocol-transfer robustness
analysis**. Phase 2 mortality outcomes were accessed in an earlier, different
panel-acquisition analysis. The present analysis is therefore not an untouched
confirmation and cannot be described as an independent external validation.

The fixed question is:

> Using the same measurements, background inputs, one-dimensional linear
> capacity, fitting opportunity and downstream calibration, is a scalar learned
> under five-year mortality supervision more useful for five-year mortality than
> a scalar learned under chronological-age supervision?

No panel, feature, endpoint, horizon, age range, fold, model family or score
dimension may be selected after this specification.

## Frozen data interface

- Source: NHANES III Phase 1, 7,558 participants and 402 five-year deaths.
- Target: NHANES III Phase 2, 7,774 participants and 392 five-year deaths.
- Age range: examination age `[20,80)` years.
- Endpoint: resolved five-year all-cause mortality indicator.
- Measurements `X`: all 25 measurements in
  `ordered_5x5_feature_manifest.tsv`, exactly in `feature_order=1,...,25`.
- Background `W`: sex and race/ethnicity.
- Survey design: `analysis_weight=mec_weight`, 23 released strata and 46 PSU
  clusters per phase.
- Folds: the existing PSU-atomic five outer folds, three full inner folds and
  the outer-fold-specific three inner folds.
- Missing measurements: training-local weighted-median imputation. Complete-case
  restriction is prohibited.

Canonical inputs are read only from
`outputs/nhanes3_phase1_to_phase2_endpoint_vs_age_acquisition_v0_2_final_20260826`.
The prior five-domain acquisition models and decisions are not reused.

## Matched one-dimensional projections

Both projections use the same Phase 1 participants, ordered `(W,X)` design,
weighted-median imputation, one-hot background encoding, weighted measurement
standardization, normalized MEC fitting weights, ridge regression, alpha grid
`{0.01,0.1,1,10,100,1000}`, squared-error tuning loss and scalar output.
Chronological age is not an input to either projection. The only intended
difference is the supervised label:

- `q_A`: standardized chronological age;
- `q_Y`: standardized five-year mortality.

For the binary mortality label, `q_Y` is a mortality-supervised linear
probability projection. It is not a latent mortality entity or a biological-age
estimate.

Target means and standard deviations are MEC-weighted and estimated separately
inside every projection training fold; they are stored with the projection.
The `W`-only operational comparator uses the identical folds, encoding, target
standardization, ridge family, alpha grid and tuning rule, with only `X` removed.

### Phase 1 fitting and freezing

1. Use the five fixed outer folds and their corresponding three fixed inner
   folds to generate fully held-out Phase 1 scores and select alpha inside each
   outer training set.
2. Use `full_inner_fold` to select one final alpha separately for `q_A` and
   `q_Y` on complete Phase 1.
3. Fit one final preprocessor and projection for each target on complete Phase
   1, persist them, reload them and require elementwise prediction parity.
4. Apply these two frozen projections to every Phase 2 participant. Phase 2
   outcomes may not enter projection preprocessing, tuning or fitting.

The operational age-decoder gate requires the Phase 1 OOF `q_A`, transformed
back to years, to improve design-weighted age MSE over a Phase 1 OOF model using
`W` alone. Phase 2 age RMSE, R-squared and performance relative to the frozen
Phase 1 weighted-mean age predictor are reported as transport diagnostics. A
failed source age-decoder gate stops the substantive branch.

## Equal Phase 2 endpoint calibration

The frozen Phase 1 scores receive identical target-local calibration in the
fixed Phase 2 outer folds:

- `M0: Y ~ W + s(A)`;
- `M_A: Y ~ W + s(A) + s(q_A)`;
- `M_Y: Y ~ W + s(A) + s(q_Y)`.

`s(A)` is the fixed five-knot cubic basis at ages 20, 35, 50, 65 and 80.
`s(q)` is a five-knot uniform cubic spline with linear extrapolation, fitted
only in the corresponding training set. All models use weighted L2-logistic
regression and the same C
grid `{0.001,0.003,0.01,0.03,0.1,0.3,1,3,10,30,100,300}`. C is selected by
design-weighted log loss in the fixed three inner folds. Every reported Phase 2
probability is outer-fold held out.

For both alpha and C, inner validation losses are pooled through their weighted
loss numerators and denominators, not averaged equally over folds. Within a
`1e-12` loss tolerance, select the more strongly regularized candidate (larger
alpha or smaller C). Binary probabilities are clipped to
`[1e-8,1-1e-8]` before log-loss evaluation.

The spline-scalar comparison is primary. The otherwise identical linear-scalar
comparison is a prespecified diagnostic, not a second decision analysis.

Target-local calibration is permitted because it calibrates two already frozen
directions equally; it may not relearn either Phase 1 projection.

## Estimands and decision gates

The sole primary estimand is the Phase 2 design-weighted paired OOF log-loss
contrast

`Delta = L(M_A) - L(M_Y)`.

Positive `Delta` favours mortality supervision. Also report

`U_A = L(M0) - L(M_A)`

to distinguish an uninformative age decoder from the scientifically important
pattern `L(M_Y) < L(M_A) < L(M0)`: genuine downstream utility without default
representational privilege.

Decision labels:

- `UTILITY_WITHOUT_PRIVILEGE_PROTOCOL_TRANSFER`: `Delta>0`, its Fay-BRR 95%
  interval has lower bound above zero, at least four of five fixed outer-fold
  contrasts are positive, and `U_A>0` with a Fay-BRR lower bound above zero.
- `MATCHED_TARGET_SCALAR_ADVANTAGE_PROTOCOL_TRANSFER`: the three `Delta` gates
  pass but the `U_A` support gate does not.
- `DIRECTIONAL_PROTOCOL_TRANSFER`: `Delta>0` but an interval or fold-agreement
  gate fails.
- `NO_TRANSFERRED_TARGET_ADVANTAGE`: `Delta<=0`.
- `STOP_INVALID_AGE_DECODER`: the Phase 1 OOF age-decoder gate fails.
- `STOP_DATA_OR_ARTIFACT_FAIL`: any identity, design, outcome-support, leakage,
  persistence or parity gate fails.

## Design-aware uncertainty

Use the existing 32 Phase 2 Fay-BRR participant factor vectors with `rho=0.3`
and `df=23`. In each replicate, repeat all Phase 2 downstream inner tuning,
outer fitting, held-out prediction and paired-loss calculation under replicate
weights. Phase 1 projections remain fixed. The resulting interval represents
Phase 2 design uncertainty conditional on the two Phase 1-fitted projections;
it does not include source-model development uncertainty.

## Artifact contract

Persist the analysis-ready Phase 1 and Phase 2 interfaces, ordered feature
manifest, fold and design identities, Phase 1 OOF predictions, final Phase 1
projection artifacts, frozen Phase 2 scores, every Phase 2 primary and Fay-BRR
downstream artifact, tuning tables, OOF probabilities, participant losses,
fold contrasts, design estimates, decision object and machine-readable
manifests. Each artifact records input signatures, ordered features,
participant/fold identities, targets, grids, weights, software and analysis
signature.

Immediately reload every fitted object and reconstruct its relevant tuning
metric and prediction. Maximum elementwise error must not exceed `1e-12`.
Compatible reruns must reuse the stored objects; incompatible artifacts fail
closed and must be written to a new versioned directory.

## Claim ceiling

A positive result would show that, for this population, endpoint, 25-item
measurement interface and one-dimensional linear model family, chronological-age
pretraining was not necessary and did not provide the best scalar direction for
five-year mortality. It would not identify biological age, establish universal
mortality sufficiency, show that all clocks are inferior, validate a surrogate,
or imply causal benefit.
