## Supplementary_Software_1 for "Inferential boundaries of age prediction: why prediction does not establish biological age measurement": NHANES_5Y_AGE_DECODER_SOURCE_INFORMATION_PROBE_SPEC_20260823.md

# NHANES five-year age-decoder source-information probe

Date frozen: 23 August 2026  
Status: author-authorized decision probe  
Primary role: empirical materiality for target-specific clock compression  
Analysis version: `nhanes_5y_age_decoder_source_information_probe_v0_2`

Implementation provenance: the v0.1 implementation completed and persisted all
five fold fits but stopped before scientific summarization because an invalid
report-stage `DataFrame` argument was supplied. No v0.1 verdict was produced.
The scientific contract is unchanged in v0.2; it is rerun into a new output
directory with a new signature rather than reusing artifacts created under the
incomplete implementation.

## 1. Decision served

The sole question is:

> For complete-support five-year all-cause mortality in NHANES, does a pure
> chronological-age decoder retain all of the held-out endpoint information
> available in the same 50 source measurements once chronological age and
> background are already included?

The result decides whether the manuscript should add one direct human example
of chronological-age-supervised compression being insufficient, adequate or
advantageous in finite samples for a named endpoint. It does not test or define
biological age.

## 2. Frozen inputs

Reuse only the persisted outputs from
`nhanes_5y_complete_support_contextual_age_probe_v0_1`:

- `analysis_ready_risk_set.tsv.gz`;
- `participant_folds.tsv`;
- `ordered_feature_tiers.tsv`;
- `oof_predictions.tsv.gz`;
- `participant_loss_vectors.tsv.gz`;
- `bootstrap_indices.npz`;
- `analysis_signature.json`;
- `model_artifact_audit.json`.

The data gate requires exactly 13,724 participants, 470 deaths, five existing
outer folds, ages 20--79 years, cycles 2007--2012, no missing model input and
the ordered 50-measurement `S3` panel. Existing baseline `M_WA` and direct
measurement `M_S3A` predictions are reused only if the predecessor artifact
audit is `PASS` and participant/fold identities match exactly.

## 3. Fixed objects and information sets

- Endpoint: death by 60 months.
- Background `W`: sex, race/ethnicity and survey cycle.
- Observed age: `A`.
- Source measurements: the fixed ordered 50-measurement `S3` panel, `X`.
- Clock: a fold-local ridge decoder `S=q_A(W,X)` trained only to predict `A`.
- Primary loss: pooled participant-level out-of-fold binary log loss.

Compare exactly three endpoint information sets:

1. `M_WA = P(Y5 | W,A)` from the accepted predecessor;
2. `M_CLOCK = P(Y5 | W,A,S)` fitted in this probe;
3. `M_X = P(Y5 | W,A,X)`, the predecessor `M_S3A`.

Given `A`, raw AgeGap `S-A` and `S` are deterministically equivalent, so no
separate AgeGap model is fitted.

## 4. Age decoder

Within every frozen outer training fold:

- one-hot encode `W` and standardize the 50 measurements using training data
  only;
- fit ridge regression to chronological age;
- select `alpha` by three-fold age-quintile-stratified RMSE from the fixed grid
  `{0.01,0.1,1,10,100,1000}`;
- generate cross-fitted scores for all outer-training participants using the
  selected `alpha` and the same inner folds;
- refit on the full outer-training fold and predict the outer test fold.

The decoder receives no mortality labels. Report pooled held-out age `R2`,
RMSE, MAE, calibration slope/intercept and age-band mean residuals as
descriptions of age readability and mean shrinkage, not biological validity.

## 5. Clock endpoint model

For each outer fold, fit ridge logistic regression for `M_CLOCK` on the outer-
training cross-fitted clock score, `W` and the same five-knot fold-local cubic
age spline used by the predecessor. Choose `C` by minimum mean three-fold inner
validation log loss from the predecessor grid
`{0.001,0.003,0.01,0.03,0.1,0.3,1,3,10,30,100,300}`. Category encoding,
clock-score scaling, spline construction and tuning are training-fold local.

The frozen outer test fold is predicted using its score from the full outer-
training age decoder. Every age-decoder and endpoint-model fit is persisted and
reload-verified.

## 6. Primary contrasts

Let `L(.)` denote pooled out-of-fold log loss.

- `V_CLOCK = L(M_WA) - L(M_CLOCK)`: clock value beyond age and background.
- `V_SOURCE = L(M_CLOCK) - L(M_X)`: source-measurement value beyond the clock.
- `V_X = L(M_WA) - L(M_X)`: direct source-measurement value beyond age and
  background.

Positive `V_SOURCE` means that the direct source-measurement model performed
better for this endpoint; it is not a universal theorem about finite-sample
estimators. Reuse the predecessor's 2,000 paired participant bootstrap indices
to obtain fixed-prediction percentile intervals, and report the sign of
`V_SOURCE` in each outer fold.

## 7. Decision gate

- `PROMISING_SOURCE_INFORMATION`: `V_SOURCE>0`, its bootstrap lower bound is
  above zero and at least four of five fold contrasts are positive.
- `WEAK_SOURCE_INFORMATION`: `V_SOURCE>0` but either the interval includes zero
  or fewer than four folds are positive.
- `ABSENT_SOURCE_ADVANTAGE`: `V_SOURCE<=0` and its upper interval is at or below
  zero.
- `AMBIGUOUS_FINITE_SAMPLE_COMPARISON`: all other valid outcomes.
- `STOP_DATA_OR_ARTIFACT_FAIL`: any data, signature, persistence or parity gate
  fails.

The verdict is a probe classification. `PROMISING_SOURCE_INFORMATION` permits
a separately frozen confirmatory analysis; no other category permits model,
endpoint, feature-panel or threshold search.

## 8. Persistence

Write to
`outputs/nhanes_5y_age_decoder_source_information_probe_v0_2/` and persist:

- copied participant/fold and ordered-feature interfaces;
- input manifest and deterministic analysis signature;
- one complete artifact per outer fold containing all age-decoder tuning fits,
  selected cross-fit models, full outer age decoder, endpoint tuning fits and
  final endpoint estimator;
- cross-fitted training clock scores and outer-test scores;
- pooled and fold metrics, participant losses, bootstrap contrasts and
  age-decoder diagnostics;
- model index and reload-parity audit;
- `decision.json` and a compact decision report.

Compatible artifacts are reloaded rather than refitted. Elementwise prediction
parity tolerance is `1e-12`; any failure stops the probe.

## 9. Claim boundary and stop rule

The result concerns one fixed linear age decoder, one endpoint, one population,
one source panel, one direct model family and one finite sample. Population
information containment is analytical; this probe asks whether its practical
consequence is visible under the declared estimators. A clock advantage would
be a finite-sample engineering result, not biological-age identity.

Stop after the three information sets, one age decoder, one endpoint, one
paired uncertainty analysis and persistence audit. Do not add clocks,
nonlinear decoders, alternative panels, horizons, endpoints or cohorts.
