## Supplementary_Software_1 for "Inferential boundaries of age prediction: why prediction does not establish biological age measurement": NHANES_5Y_COMPLETE_SUPPORT_CONTEXTUAL_AGE_PROBE_SPEC_20260823.md

# NHANES five-year complete-support contextual-age probe

Date frozen: 23 August 2026  
Status: author-authorized decision probe  
Module: HUM-03 successor candidate  
Primary role: positive-boundary  
Analysis version: `nhanes_5y_complete_support_contextual_age_probe_v0_1`

## 1. Decision served

The sole question is:

> For five-year all-cause mortality in NHANES cycles with complete
> administrative support, how does the held-out predictive contribution of
> chronological age change as a prespecified age-blind clinical measurement
> representation expands from background only to 6, 30 and 50 measurements?

The result decides whether HUM-03 returns to the main text as a quantitative
contextual-age result. It does not estimate causal mediation, biological age or
a percentage of ageing.

## 2. Frozen input and data gate

Use only:

- `data/processed/NHANES_broad_clinical_panel_v0_1.tsv.gz`;
- `data/processed/NHANES_broad_clinical_feature_manifest_v0_1.tsv`.

Restrict before model fitting to:

- cycles 2007–2008, 2009–2010 and 2011–2012;
- ages 20–79 years;
- the existing complete 50-measurement interface;
- five-year death defined by death at or before 60 months;
- event-free participants observed for at least 60 months.

The pre-fit gate requires exactly:

- (N=13{,}724);
- 470 five-year deaths;
- zero event-free participants censored before 60 months;
- one row per `SEQN`;
- the frozen ordered feature manifest and no missing model inputs.

Fail closed if any gate differs. Do not revise the cycle set, horizon or feature
panel after the gate.

## 3. Fixed information sets

Background (W): sex, race/ethnicity and survey cycle.

The age-blind measurement tiers are inherited without reordering:

- (S_1): six examination measurements;
- (S_{2c}): (S_1) plus BIOPRO and HDL, 30 measurements total;
- (S_3): all 50 measurements after adding CBC.

No feature is selected by age association, mortality association, clock
loading or observed performance. Do not include the historical haematology-first
order sensitivity, age interactions or common-support model in this decision
probe.

## 4. Model ladder

For (k\in\{0,1,2c,3\}), fit:

\[
M_{W+S_k}
\quad\text{and}\quad
M_{W+S_k+A},
\]

where (S_0) is empty and chronological age is represented by the same
fold-local cubic spline basis with five knots used in the predecessor analysis.

Use ridge logistic regression and the frozen penalty grid:

\[
C\in\{0.001,0.003,0.01,0.03,0.1,0.3,1,3,10,30,100,300\}.
\]

Within each outer training fold, choose (C) by minimum mean three-fold inner
validation log loss; deterministic ties choose the smaller (C). All category
encoding, scaling, spline construction and tuning are fold-local.

## 5. Validation

- Five outer participant-disjoint folds.
- Seed `20260823`.
- Stratify by five-year endpoint and training-population age quintile when all
  cells permit; endpoint-only fallback must be declared before fitting.
- Use the same outer folds for every model.
- Primary score: pooled participant-level out-of-fold binary log loss.
- Secondary diagnostics: Brier score, AUC and age-band calibration.

## 6. Primary estimands

For each tier:

\[
V_A(S_k)=
L_{OOF}(W,S_k)-L_{OOF}(W,S_k,A).
\]

Primary reporting uses the absolute paired loss improvement. Define the
richest-panel change:

\[
D_{0\rightarrow3}=V_A(S_0)-V_A(S_3).
\]

Positive values indicate predictive substitution by the measured panel;
negative values indicate age–measurement synergy; zero indicates no detectable
change. Ratios to (V_A(S_0)) are secondary and may be reported only if the
baseline value is clearly positive. No ratio is a fraction of ageing.

## 7. Uncertainty

Use 2,000 paired participant bootstraps of fixed out-of-fold predictions with
seed `20260824`, stratified by endpoint and decade age band. Apply identical
indices to all models in each replicate. Report percentile intervals for every
(V_A(S_k)) and (D_{0\rightarrow3}).

Fixed-prediction bootstrap intervals omit model-development and refitting
uncertainty; state this explicitly.

## 8. Decision categories

- `CONTEXTUAL_SUBSTITUTION`: lower interval bound for (V_A(S_0)) is above
  zero and lower interval bound for (D_{0\rightarrow3}) is above zero.
- `CONTEXTUAL_SYNERGY`: lower interval bound for (V_A(S_0)) is above zero and
  upper interval bound for (D_{0\rightarrow3}) is below zero.
- `NO_DETECTABLE_MIGRATION`: baseline age value is positive but the interval for
  (D_{0\rightarrow3}) includes zero.
- `NO_VISIBLE_BASELINE_AGE_VALUE`: the interval for (V_A(S_0)) includes or
  crosses zero.
- `STOP_DATA_OR_AUDIT_FAIL`: any data gate, artifact or reload parity check
  fails.

The sequence need not be monotone. No decision category authorizes changing
the horizon, tiers, age form, model family or threshold.

Main-text quantitative placement is permitted for
`CONTEXTUAL_SUBSTITUTION` or `CONTEXTUAL_SYNERGY`. Other valid outcomes remain
framework or Extended Data boundaries.

## 9. Persistence contract

Write to `outputs/nhanes_5y_complete_support_contextual_age_probe_v0_1/` and
persist:

- `analysis_ready_risk_set.tsv.gz`;
- `cycle_support_audit.tsv`;
- `participant_folds.tsv`;
- `ordered_feature_tiers.tsv`;
- `input_manifest.tsv` and deterministic analysis signature;
- one serialized fold-local object for every outer-fold model, including
  encoder, scaler, age spline, chosen penalty and fitted estimator;
- inner tuning tables and split identities;
- out-of-fold probabilities and participant loss vectors;
- pooled/fold metrics and age-band calibration;
- bootstrap indices and contrast tables;
- `decision.json`;
- `model_artifact_audit.json` with object counts and reload parity;
- `artifact_manifest.tsv`;
- one compact decision figure in PNG and SVG;
- one decision-ready report.

If a compatible artifact exists, validate and reuse it. Do not silently refit.
Immediately reload every saved fold object and reproduce held-out probabilities
elementwise; record maximum absolute error and fail closed if it exceeds
(10^{-12}).

## 10. Runtime

Run the environment gate before Python work and use only:

```powershell
& .\LifeHistoryStateDecomposition\scripts\Invoke-ProjectPython.ps1 -- <args>
```

No bare Python, pip or environment substitution is permitted.

## 11. Claim boundary

The result concerns predictive value for five-year mortality in the declared
complete-case NHANES population, measurements and model. Remaining age value
does not identify what is missing; a reduction does not establish causal
mediation or state sufficiency; a synergy does not establish an age mechanism.

## 12. Stop rule

Stop after the single five-year endpoint, three fixed measurement tiers, eight
paired models, one bootstrap analysis, persistence audit and independent
numerical audit. Do not add ten-year mortality, alternative orders,
interactions, clocks, causes of death or new feature panels.
