## Supplementary_Software_1 for "Inferential boundaries of age prediction: why prediction does not establish biological age measurement": NHANES_5Y_MATCHED_SCALAR_TARGET_SUPERVISION_SPEC_v0_2_20260826.md

# NHANES five-year matched scalar target-supervision specification v0.2

Date frozen: 26 August 2026  
Module: `HUM-06`  
Analysis version: `nhanes_5y_matched_scalar_target_supervision_probe_v0_2`

## Revision provenance

The frozen v0.1 implementation kept every outer test fold fully held out, but
used one common outer-training OOF score vector when tuning the downstream
calibration. When a downstream fold was held out for C selection, mortality
outcomes in that fold could have contributed to projection scores used on the
downstream-training side. V0.1 is retained unchanged as audit provenance and
does not carry the matched-target verdict.

V0.2 changes only this nesting and the corresponding identity/parity audit. It
does not change the population, measurements, endpoint, model families, grids,
fold counts, seeds, metrics, bootstrap or decision gates.

## Question

For the frozen complete-support NHANES population and 50-measurement panel:

1. does the direct-source advantage in `HUM-04` survive a fixed flexible
   calibration of its age-supervised score; and
2. when inputs, linear capacity, scalar dimension and validation opportunity
   are matched, does mortality supervision produce a scalar better suited to
   five-year mortality than chronological-age supervision?

## Frozen interface

- 13,724 participants, 470 five-year deaths, cycles 2007--2012, ages 20--79.
- Independent unit: participant.
- Source measurements: ordered S3 panel of 50 measurements.
- Background `W`: sex, race/ethnicity and cycle.
- Chronological age enters only the common downstream endpoint model in the
  matched comparison.
- Primary metric: participant-level OOF binary log loss.

## Flexible calibration check

Reuse the original `20260823` outer folds and fold-local/cross-fitted age scores.
Compare the frozen linear clock model with a single prespecified model using
`W + five-knot cubic spline(A) + five-knot cubic spline(q_A)` and ridge
logistic regression. The fixed C grid is
`{0.001,0.003,0.01,0.03,0.1,0.3,1,3,10,30,100,300}`.

The primary robustness contrast is the flexible clock model's log loss minus
the frozen direct 50-measurement model's log loss. No second calibration search
is permitted.

## Matched one-dimensional comparison

Construct two scalars from the same fold-local encoded and standardized
`(W,X)` design:

- `q_A`: ridge projection trained on standardized chronological age;
- `q_Y`: ridge projection trained on standardized five-year mortality.

Both projections use squared-error loss, the same alpha grid
`{0.01,0.1,1,10,100,1000}`, identical exclusion rules and one scalar output.
Both scalars receive the same two prespecified downstream models:

- linear scalar plus `W` and a five-knot age spline;
- five-knot scalar spline plus `W` and the same age spline.

The spline comparison is primary; the linear comparison is diagnostic.

## Fully nested downstream tuning

For each outer-training set and each downstream validation fold:

1. define the complementary downstream-training set;
2. generate scores for that training set by three-fold projection
   cross-fitting wholly inside it; within each projection fold, alpha is
   selected using only the projection-fold training participants;
3. separately select alpha using only the complete downstream-training set,
   fit the projection on that complete set and score the downstream validation
   fold; and
4. fit and evaluate every candidate downstream C using only these paired
   training and validation scores.

Thus neither a downstream validation participant nor that participant's
mortality outcome enters any mortality-supervised score used to fit the model
evaluated on that validation fold. The final outer-test score remains generated
from the complete outer-training set, and the final downstream model is fitted
to projection-cross-fitted scores for the complete outer-training set.

## Repeated partitions

Use five five-fold outer partitions with seeds `20260823`, `20260901`,
`20260902`, `20260903` and `20260904`, stratified by five-year outcome and
training-age quintile. Downstream inner seed is
`outer_seed + 1000 + outer_fold`; projection-tuning seed is
`outer_seed + 2000 + 10*outer_fold + downstream_fold`. The same declared
projection seed is reused inside that downstream fold's corrective nesting; no
additional seed or split search is introduced.

Use 2,000 participant bootstraps with seed `20260905`, stratified by outcome,
age decade and cycle. Average each participant's loss across repeats before the
paired bootstrap. Repeats are not independent samples. Report repeat- and
cycle-specific point estimates without treating them as replication tests.

## Persistence and fail-closed audit

Persist every projection tuning fit, fold-local preprocessor, final projection,
downstream tuning fit and final downstream model. After reload:

- verify every inner identity is a subset of its declared outer and downstream
  training set and that held-out identities form exact non-overlapping covers;
- reconstruct projection tuning metrics and selected alpha values;
- connect projection holdout scores to assembled training, validation and test
  score vectors by participant identity;
- connect those vectors to every downstream stored train, validation and test
  score;
- reconstruct downstream tuning losses, selected C values and all held-out
  probabilities; and
- require elementwise parity at tolerance `1e-12`.

Any failure stops the branch.

## Gates

- `MATCHED_TARGET_SCALAR_ADVANTAGE`: primary log-loss contrast is positive,
  participant-bootstrap lower bound is above zero, at least four of five
  repeat estimates are positive and none of the three cycle estimates reverses.
- `SPLIT_OR_CYCLE_SENSITIVE`: pooled contrast is positive but an agreement gate
  fails.
- `NO_MATCHED_TARGET_ADVANTAGE`: estimate is non-positive or its interval
  includes zero.
- `HUM04_ROBUST_TO_FLEX_CALIBRATION`: flexible-clock versus direct-source gap
  is positive with lower interval above zero and at least four of five original
  fold contrasts positive.
- `CALIBRATION_SENSITIVE`: the prespecified flexible calibration removes that
  gap.
- `STOP_DATA_OR_ARTIFACT_FAIL`: any input, identity, event-support, leakage,
  persistence or parity gate fails.

## Claim ceiling

Success would show that, for one population, endpoint, panel and fixed linear
capacity, supervision target matters even when the learned representation is
one-dimensional. It would not identify biological age, establish universal
mortality representation, prove surrogate validity or imply causal benefit.
