## Supplementary_Software_1 for "Inferential boundaries of age prediction: why prediction does not establish biological age measurement": NHANES_HUM03_HUM04_HUM06_EXTENDED_DATA_FIGURE3_SUBMISSION_CONTRACT_v0_3_20260828.md

# Extended Data Figure 3 submission contract v0.3

## Purpose

Create a fit-free three-panel submission successor for Extended Data Figure 3.
The figure retains the complete observed-age/block contrast set, the matched
observed-age-versus-decoder substitution boundary and the fixed-fold direction
diagnostics. The former engineering persistence panel is removed from the
scientific display and remains documented in the reproducibility manifest.

## Frozen inputs and claim boundary

- Reuse the validated v0.2 TSV/JSON input gate and all frozen estimands.
- Do not load a fitted model, refit or tune an estimator, resample participants
  or generate random numbers.
- Do not change any estimate, interval, fold sign or interpretation boundary.
- An interval crossing zero is unresolved in the declared design; it is not
  equivalence or non-inferiority.
- Fold directions are diagnostics, not independent replications.

## Output

- Three panels labelled a-c.
- 600-dpi PNG, editable SVG and vector PDF.
- Panel-level Source Data and a fit-free visualization audit.
