## Supplementary_Software_1 for "Inferential boundaries of age prediction: why prediction does not establish biological age measurement": NHANES_INDIVIDUAL_AGEGAP_CONTRIBUTIONS_SPEC_20260831.md

# NHANES individual AgeGap contribution analysis: frozen specification

Date frozen: 31 August 2026  
Analysis version: `nhanes_individual_agegap_contributions_v0_1`  
Status: author-authorized descriptive analysis

## Question

Can near-identical individual AgeGaps arise from materially different measured-
profile contributions under the already frozen NHANES chronological-age decoder?

The analysis describes how the fitted decoder generated its score. It is not a
causal decomposition of biological ageing and does not assign mechanisms.

## Frozen population and decoder

Reuse without refitting:

- the 13,724 complete-support NHANES 2007--2012 participants aged 20--79;
- the fixed 50-measurement S3 panel and five outer folds;
- each participant's existing out-of-fold Ridge chronological-age score;
- the five persisted fold-local age-decoder artifacts; and
- the existing out-of-fold five-year mortality predictions from the direct
  age-plus-measurement model (`M_X`) and adjusted AgeGap model (`M_AGE_GAP`).

All predecessor artifact audits must be `PASS`, participant/fold identities must
match exactly, and reloaded decoder predictions must reproduce the stored scores
within `1e-12`.

## Fold-local conditional centering

For each outer fold, fit a multivariate linear conditional-mean model on outer-
training participants only:

`E_hat(X | A,W) = Ridge([one-hot(W), cubic-spline(A)])`,

where `W={sex, race/ethnicity, survey cycle}`. Use five knots, degree three,
linear extrapolation, and the fixed alpha grid
`{0.01,0.1,1,10,100,1000}`. Select alpha by three-fold age-quintile-stratified
inner cross-validation minimizing mean squared error after standardizing each of
the 50 outcomes by its inner-training standard deviation. The inner seed is
`20260831 + outer_fold`.

Persist every tuning fit, the selected final residualizer, preprocessing,
ordered features, row identities, and predictions. Reload immediately and
verify held-out conditional means elementwise within `1e-12`. Existing decoder
objects are referenced by signature and hash rather than duplicated.

## Exact additive decomposition

For participant `i` in outer fold `k`, let the frozen decoder use measurement
scaler mean `mu_k`, scale `sigma_k`, and measurement coefficient `beta_kj`.
Define

`c_ij = beta_kj * (X_ij - E_hat_k[X_j | A_i,W_i]) / sigma_kj`.

Then define the conditional-centred AgeGap

`G_centered_i = sum_j c_ij = S_i - E_hat_k[S | A_i,W_i]`.

Because chronological age is fixed inside the conditioning set, this is also

`(S_i-A_i) - E_hat_k[S_i-A_i | A_i,W_i]`.

The field-standard raw AgeGap `G_raw_i=S_i-A_i` is retained separately for
every participant. Contributions are in decoder-score years. The maximum
absolute difference between `sum_j c_ij` and `G_centered_i` must not exceed
`1e-10`.

## Outcome-blind matching

Create deterministic, non-reusing pairs within exact strata of:

- outer fold;
- chronological age;
- sex;
- race/ethnicity; and
- survey cycle.

Within each stratum, sort participants by `G_centered`. Form candidate edges
between adjacent sorted participants, order edges by absolute gap difference
then row identifiers, and greedily retain an edge only if neither participant
has already been used. Retain pairs with absolute centred-gap difference no
greater than 0.25 decoder years. Outcomes, endpoint predictions, contribution
distance and feature extremes are not used to form pairs.

For every retained pair report:

- raw and centred AgeGap differences;
- Euclidean and L1 distance between 50-dimensional contribution vectors;
- cosine similarity;
- Jaccard overlap of the five largest absolute contributors;
- absolute logit difference for `M_X` and `M_AGE_GAP`.

The primary profile-distance quantity is Euclidean distance in decoder years.
Endpoint logit differences are descriptive model-output contrasts, not causal
hazard or risk effects.

## Display-case rule

Among matched pairs whose two centred AgeGaps have the same sign and whose mean
absolute centred AgeGap is at least 2 years, select:

1. the pair closest to the median Euclidean contribution distance; and
2. the pair closest to the 95th percentile Euclidean contribution distance.

Ties are resolved by pair identifier. Display the union of each pair's eight
largest absolute feature contributions; aggregate all other features into an
exact `Other measured features` remainder so displayed bars still sum to the
participant's centred AgeGap. These rules are frozen before pair results are
inspected.

## Required persistence

Write to `outputs/nhanes_individual_agegap_contributions_v0_1/`:

- signed input and artifact manifests;
- the analysis-ready participant interface;
- all fold-local residualizer artifacts and parity audit;
- ordered feature/domain dictionary and fold-local decoder coefficients;
- participant-level raw/centred AgeGaps and conditional expected scores;
- the complete long-form participant-by-feature contribution matrix;
- all matched-pair metrics, distribution summaries and selected-case data;
- a publication-oriented PNG/SVG figure; and
- a methods/results report with explicit interpretation boundaries.

Any missing object, incompatible signature, participant-order discrepancy,
feature-order discrepancy, or failed numerical parity check stops the run.
