## Supplementary_Software_1 for "Inferential boundaries of age prediction: why prediction does not establish biological age measurement": plot_hsct_age_reconstruction_semantic_counterexample_v0_4.py

```
#!/usr/bin/env python
"""Build the publication figure for the HSCT semantic-counterexample synthesis.

This is a fit-free visualization of frozen HUM-01 artifacts. It does not draw
new random numbers, rescore methylation, refit a clock, or change the referent
estimand.
"""

from __future__ import annotations

import hashlib
import json
import platform
import sys
from pathlib import Path

import matplotlib
import matplotlib.pyplot as plt
from matplotlib.lines import Line2D
from matplotlib.patches import FancyArrowPatch, FancyBboxPatch
import numpy as np
import pandas as pd


ROOT = Path(__file__).resolve().parents[1]
ANALYSIS_VERSION = "hsct_age_reconstruction_semantic_counterexample_figure_v0_4"
FROZEN = ROOT / "outputs" / "hsct_fixed_score_referent_null_probe_v0_1"
READY_PATH = FROZEN / "analysis_ready_pairs.tsv"
SUMMARY_PATH = FROZEN / "observed_summary.tsv"
DECISION_PATH = FROZEN / "decision.json"
AUDIT_PATH = FROZEN / "independent_audit.json"
SYNTHESIS_PATH = ROOT / "reports" / "HSCT_AGE_RECONSTRUCTION_SEMANTIC_COUNTEREXAMPLE_SYNTHESIS_V0_1_20260824.md"
SCRIPT_PATH = Path(__file__).resolve()
OUT = ROOT / "outputs" / ANALYSIS_VERSION
STAGE = ROOT / "outputs" / f".{ANALYSIS_VERSION}.staging"
FIGURE_STEM = "Figure_HSCT_age_reconstruction_semantic_counterexample"
TOL = 1e-12

INK = "#202124"
MUTED = "#62676D"
GRID = "#D9DDE1"
LIGHT = "#EEF1F3"
TEAL = "#087E8B"
TEAL_LIGHT = "#D7EEF0"
ORANGE = "#D97732"
ORANGE_LIGHT = "#F7E6D8"
WHITE = "#FFFFFF"


def sha256(path: Path) -> str:
    digest = hashlib.sha256()
    with path.open("rb") as handle:
        for block in iter(lambda: handle.read(1024 * 1024), b""):
            digest.update(block)
    return digest.hexdigest()


def write_json(path: Path, payload: object) -> None:
    path.write_text(json.dumps(payload, indent=2, ensure_ascii=False), encoding="utf-8")


def require(condition: bool, message: str) -> None:
    if not condition:
        raise RuntimeError(message)


def box(
    ax: plt.Axes,
    xy: tuple[float, float],
    width: float,
    height: float,
    text: str,
    *,
    facecolor: str = WHITE,
    edgecolor: str = MUTED,
    fontsize: float = 7.0,
    weight: str = "normal",
    color: str = INK,
    linewidth: float = 1.0,
) -> FancyBboxPatch:
    patch = FancyBboxPatch(
        xy,
        width,
        height,
        boxstyle="round,pad=0.018,rounding_size=0.018",
        facecolor=facecolor,
        edgecolor=edgecolor,
        linewidth=linewidth,
        transform=ax.transAxes,
        clip_on=False,
    )
    ax.add_patch(patch)
    ax.text(
        xy[0] + width / 2,
        xy[1] + height / 2,
        text,
        transform=ax.transAxes,
        ha="center",
        va="center",
        fontsize=fontsize,
        fontweight=weight,
        color=color,
        linespacing=1.16,
    )
    return patch


def arrow(
    ax: plt.Axes,
    start: tuple[float, float],
    end: tuple[float, float],
    *,
    color: str = MUTED,
    linewidth: float = 1.1,
    style: str = "-|>",
) -> None:
    ax.add_patch(
        FancyArrowPatch(
            start,
            end,
            transform=ax.transAxes,
            arrowstyle=style,
            mutation_scale=9,
            linewidth=linewidth,
            color=color,
            shrinkA=0,
            shrinkB=0,
            clip_on=False,
        )
    )


def style_axis(ax: plt.Axes) -> None:
    ax.spines["top"].set_visible(False)
    ax.spines["right"].set_visible(False)
    ax.spines["left"].set_color(MUTED)
    ax.spines["bottom"].set_color(MUTED)
    ax.tick_params(colors=INK, labelsize=6.3, width=0.7, length=3)
    ax.grid(color=GRID, linewidth=0.55, alpha=0.72)
    ax.set_axisbelow(True)


def panel_label(ax: plt.Axes, label: str, title: str) -> None:
    ax.text(
        -0.02,
        1.065,
        label,
        transform=ax.transAxes,
        ha="left",
        va="bottom",
        fontsize=10,
        fontweight="bold",
        color=INK,
        clip_on=False,
    )
    ax.text(
        0.055,
        1.065,
        title,
        transform=ax.transAxes,
        ha="left",
        va="bottom",
        fontsize=8.2,
        fontweight="bold",
        color=INK,
        clip_on=False,
    )


def draw_panel_a(ax: plt.Axes) -> None:
    ax.set_axis_off()
    panel_label(ax, "a", "HSCT separates candidate referents")

    box(ax, (0.03, 0.73), 0.33, 0.15, "Donor\nchronological age  $A_D$", facecolor=ORANGE_LIGHT, edgecolor=ORANGE)
    box(ax, (0.63, 0.73), 0.33, 0.15, "Recipient host\nchronological age  $A_R$", facecolor=TEAL_LIGHT, edgecolor=TEAL)
    arrow(ax, (0.37, 0.805), (0.61, 0.805), color=ORANGE, linewidth=1.5)
    ax.text(0.49, 0.845, "allogeneic HSCT", transform=ax.transAxes, ha="center", va="bottom", fontsize=6.3, color=MUTED)

    ax.text(0.49, 0.655, "after $t$ years", transform=ax.transAxes, ha="center", va="center", fontsize=6.5, color=MUTED)
    arrow(ax, (0.20, 0.71), (0.38, 0.59), color=ORANGE)
    arrow(ax, (0.79, 0.71), (0.62, 0.59), color=TEAL)
    box(
        ax,
        (0.27, 0.45),
        0.46,
        0.14,
        "One recipient, two observable coordinates\nlineage  $A_L = A_D + t$     ≠     host  $A_R$",
        facecolor=WHITE,
        edgecolor=INK,
        fontsize=7.0,
        weight="bold",
    )

    arrow(ax, (0.50, 0.43), (0.50, 0.35), color=MUTED)
    box(ax, (0.08, 0.19), 0.36, 0.15, "Recipient blood\n$X_B$ from donor-derived lineage", facecolor=ORANGE_LIGHT, edgecolor=ORANGE, fontsize=6.8)
    arrow(ax, (0.45, 0.265), (0.58, 0.265), color=MUTED)
    box(ax, (0.59, 0.19), 0.32, 0.15, "Frozen age decoder\n$S=q(X_B)$  (years)", facecolor=TEAL_LIGHT, edgecolor=TEAL, fontsize=7.0, weight="bold")

    ax.text(
        0.50,
        0.075,
        "Question: does the age-valued readout affiliate with $A_R$ or $A_L$?",
        transform=ax.transAxes,
        ha="center",
        va="center",
        fontsize=7.0,
        color=INK,
        fontweight="bold",
    )


def draw_geometry_axis(ax: plt.Axes, frame: pd.DataFrame, title: str) -> None:
    style_axis(ax)
    older = frame.loc[frame["stratum"].eq("host_older")]
    younger = frame.loc[frame["stratum"].eq("host_younger")]
    ax.scatter(
        older["H_minus_C"],
        older["S"] - older["C"],
        s=19,
        marker="o",
        facecolor=TEAL,
        edgecolor=WHITE,
        linewidth=0.35,
        alpha=0.82,
        zorder=3,
    )
    ax.scatter(
        younger["H_minus_C"],
        younger["S"] - younger["C"],
        s=23,
        marker="^",
        facecolor=WHITE,
        edgecolor=ORANGE,
        linewidth=1.0,
        alpha=0.95,
        zorder=4,
    )
    all_values = np.concatenate(
        [frame["H_minus_C"].to_numpy(float), (frame["S"] - frame["C"]).to_numpy(float)]
    )
    lo = min(-5.0, float(np.floor(all_values.min() / 10) * 10 - 2))
    hi = max(5.0, float(np.ceil(all_values.max() / 10) * 10 + 2))
    ax.plot([lo, hi], [0, 0], color=TEAL, linewidth=1.25, zorder=2)
    ax.plot([lo, hi], [lo, hi], color=MUTED, linewidth=1.0, linestyle=(0, (4, 2)), zorder=2)
    ax.set_xlim(lo, hi)
    ax.set_ylim(lo, hi)
    ax.set_aspect("equal", adjustable="box")
    ax.set_title(f"{title}  ($n={len(frame)}$)", fontsize=7.2, color=INK, pad=3.0, fontweight="bold")
    ax.set_xlabel("")
    ax.set_ylabel("Score minus lineage coordinate,  $S-A_L$ (years)", fontsize=6.1, color=INK, labelpad=2)


def draw_panel_b(container: plt.Axes, ready: pd.DataFrame) -> None:
    container.set_axis_off()
    panel_label(container, "b", "Fixed recipient-blood score affiliation")
    sub = container.get_subplotspec().subgridspec(1, 2, wspace=0.40)
    ax_s = container.figure.add_subplot(sub[0, 0])
    ax_h = container.figure.add_subplot(sub[0, 1])
    draw_geometry_axis(ax_s, ready.loc[ready["source_study"].eq("Soraas2019")], "Søraas 2019")
    draw_geometry_axis(ax_h, ready.loc[ready["source_study"].eq("Holland2024")], "Holland 2024")
    container.text(
        0.5,
        -0.035,
        "Host minus lineage coordinate,  $A_R-A_L$ (years)",
        transform=container.transAxes,
        ha="center",
        va="top",
        fontsize=6.3,
        color=INK,
    )
    handles = [
        Line2D([0], [0], marker="o", color="none", markerfacecolor=TEAL, markeredgecolor=WHITE, markersize=5.3, label="host older"),
        Line2D([0], [0], marker="^", color="none", markerfacecolor=WHITE, markeredgecolor=ORANGE, markersize=5.5, label="host younger"),
        Line2D([0], [0], color=TEAL, linewidth=1.3, label="lineage referent  $S=A_L$"),
        Line2D([0], [0], color=MUTED, linewidth=1.0, linestyle=(0, (4, 2)), label="host referent  $S=A_R$"),
    ]
    container.legend(
        handles=handles,
        loc="lower center",
        bbox_to_anchor=(0.5, -0.20),
        ncol=2,
        frameon=False,
        fontsize=5.8,
        handlelength=2.0,
        columnspacing=1.1,
    )


def draw_panel_c(container: plt.Axes, summary: pd.DataFrame) -> None:
    container.set_axis_off()
    panel_label(container, "c", "Affiliation exceeds the frozen null")
    sub = container.get_subplotspec().subgridspec(1, 2, width_ratios=[1.12, 1.0], wspace=0.33)
    ax_level = container.figure.add_subplot(sub[0, 0])
    ax_delta = container.figure.add_subplot(sub[0, 1])

    order = ["Holland2024", "Soraas2019"]
    labels = ["Holland 2024\n61 pairs", "Søraas 2019\n19 recipients"]
    y = np.array([1.0, 0.0])
    summary_by_study = summary.set_index("source_study")

    for yi, study in zip(y, order):
        row = summary_by_study.loc[study]
        ax_level.plot(
            [row["I_bal_permutation_q025"], row["I_bal_permutation_q975"]],
            [yi, yi],
            color="#AAB1B7",
            linewidth=5.0,
            solid_capstyle="round",
            zorder=1,
        )
        ax_level.scatter(row["I_bal_permutation_null_mean"], yi, marker="D", s=24, facecolor=WHITE, edgecolor=MUTED, linewidth=0.9, zorder=3)
        ax_level.scatter(row["I_bal_observed"], yi, marker="o", s=38, facecolor=TEAL, edgecolor=INK, linewidth=0.45, zorder=4)
        ax_level.text(row["I_bal_observed"] + 0.012, yi, f"{row['I_bal_observed']:.3f}", va="center", ha="left", fontsize=6.2, color=TEAL, fontweight="bold")

        delta = row["observed_minus_permutation_null_mean"]
        low = row["observed_minus_null_bootstrap_ci_low"]
        high = row["observed_minus_null_bootstrap_ci_high"]
        ax_delta.plot([low, high], [yi, yi], color=TEAL, linewidth=2.0, solid_capstyle="round", zorder=2)
        ax_delta.scatter(delta, yi, marker="o", s=38, facecolor=TEAL, edgecolor=INK, linewidth=0.45, zorder=3)
        ax_delta.text(high + 0.008, yi, f"{delta:.3f}", va="center", ha="left", fontsize=6.2, color=TEAL, fontweight="bold")

    for ax in (ax_level, ax_delta):
        ax.spines["top"].set_visible(False)
        ax.spines["right"].set_visible(False)
        ax.spines["left"].set_visible(False)
        ax.spines["bottom"].set_color(MUTED)
        ax.set_ylim(-0.55, 1.55)
        ax.tick_params(axis="x", labelsize=6.3, colors=INK, width=0.7, length=3)
        ax.tick_params(axis="y", length=0)
        ax.grid(axis="x", color=GRID, linewidth=0.55)
        ax.set_axisbelow(True)

    ax_level.set_xlim(0.35, 0.90)
    ax_level.set_yticks(y, labels, fontsize=6.4, color=INK)
    ax_level.set_xlabel("Sign-balanced affiliation,  $J_{aff,bal}$", fontsize=6.3, color=INK)
    ax_level.set_title("Observed versus null 95% range", fontsize=6.7, color=INK, pad=5)

    ax_delta.axvline(0, color=MUTED, linewidth=0.9, linestyle=(0, (3, 2)))
    ax_delta.set_xlim(-0.03, 0.25)
    ax_delta.set_yticks(y, ["", ""])
    ax_delta.set_xlabel("Observed minus null mean", fontsize=6.3, color=INK)
    ax_delta.set_title("Pair/recipient bootstrap 95% CI", fontsize=6.7, color=INK, pad=5)

    handles = [
        Line2D([0], [0], marker="o", color="none", markerfacecolor=TEAL, markeredgecolor=INK, markeredgewidth=0.45, markersize=5.3, label="observed"),
        Line2D([0], [0], marker="D", color="none", markerfacecolor=WHITE, markeredgecolor=MUTED, markersize=4.5, label="null mean"),
        Line2D([0], [0], color="#AAB1B7", linewidth=4.0, label="null 95% range"),
    ]
    ax_level.legend(handles=handles, loc="upper left", bbox_to_anchor=(0.0, -0.25), ncol=3, frameon=False, fontsize=5.6, handlelength=1.5, columnspacing=0.8)


def draw_panel_d(ax: plt.Axes) -> None:
    ax.set_axis_off()
    panel_label(ax, "d", "Empirical consequence for age reconstruction")

    box(ax, (0.06, 0.76), 0.88, 0.12, "Source task\nX_B  →  q(X_B) = S is trained to reconstruct chronological age A", facecolor=LIGHT, edgecolor=MUTED, fontsize=6.7)
    arrow(ax, (0.50, 0.74), (0.50, 0.66), color=MUTED)
    box(ax, (0.06, 0.52), 0.88, 0.13, "HSCT observation\n$A_R \\ne A_L$ and the fixed blood score affiliates with $A_L$ beyond the frozen null", facecolor=TEAL_LIGHT, edgecolor=TEAL, fontsize=6.7, weight="bold")
    arrow(ax, (0.50, 0.50), (0.50, 0.43), color=MUTED)

    box(ax, (0.04, 0.23), 0.42, 0.17, "Not identified here\n$S$ = recipient-wide biological age", facecolor=ORANGE_LIGHT, edgecolor=ORANGE, fontsize=7.0, weight="bold")
    ax.plot([0.08, 0.42], [0.25, 0.38], transform=ax.transAxes, color=ORANGE, linewidth=1.5, clip_on=False)
    box(ax, (0.54, 0.23), 0.42, 0.17, "Supported here\nSampled-lineage referent affiliation", facecolor=TEAL_LIGHT, edgecolor=TEAL, fontsize=7.0, weight="bold")

    ax.text(
        0.50,
        0.075,
        "Age reconstruction does not fix a unique organismal referent",
        transform=ax.transAxes,
        ha="center",
        va="center",
        fontsize=8.0,
        color=INK,
        fontweight="bold",
    )
    ax.text(
        0.50,
        0.015,
        "Materiality witness, not empirical proof of the analytical theorem",
        transform=ax.transAxes,
        ha="center",
        va="center",
        fontsize=6.1,
        color=MUTED,
        style="italic",
    )


def build_figure(ready: pd.DataFrame, summary: pd.DataFrame, destination: Path) -> None:
    matplotlib.rcParams.update(
        {
            "font.family": "Arial",
            "font.size": 7,
            "axes.linewidth": 0.75,
            "pdf.fonttype": 42,
            "ps.fonttype": 42,
            "svg.fonttype": "none",
            "savefig.facecolor": "white",
        }
    )
    fig = plt.figure(figsize=(7.15, 5.75), facecolor=WHITE)
    grid = fig.add_gridspec(
        2,
        2,
        width_ratios=[0.90, 1.35],
        height_ratios=[1.03, 0.97],
        left=0.055,
        right=0.985,
        top=0.955,
        bottom=0.075,
        wspace=0.20,
        hspace=0.36,
    )
    ax_a = fig.add_subplot(grid[0, 0])
    ax_b = fig.add_subplot(grid[0, 1])
    ax_c = fig.add_subplot(grid[1, 0])
    ax_d = fig.add_subplot(grid[1, 1])

    draw_panel_a(ax_a)
    draw_panel_b(ax_b, ready)
    draw_panel_c(ax_c, summary)
    draw_panel_d(ax_d)

    fig.text(
        0.985,
        0.018,
        "Complementary post-HSCT analyses; the study programmes partly overlap.",
        ha="right",
        va="bottom",
        fontsize=5.6,
        color=MUTED,
        style="italic",
    )
    fig.savefig(destination.with_suffix(".png"), dpi=600, bbox_inches="tight", pad_inches=0.035)
    fig.savefig(destination.with_suffix(".svg"), bbox_inches="tight", pad_inches=0.035)
    fig.savefig(destination.with_suffix(".pdf"), bbox_inches="tight", pad_inches=0.035)
    plt.close(fig)


def build_legend(path: Path) -> None:
    text = r"""# Figure legend | 图注

## English

**Figure | HSCT exposes the referent ambiguity of an age-valued blood readout.**
**a,** Allogeneic haematopoietic stem-cell transplantation separates recipient-host chronological age \(A_R\) from the chronological coordinate of the sampled donor-derived haematopoietic lineage \(A_L=A_D+t\). A fixed recipient-blood score \(S=q(X_B)\) is expressed in years, but its numerical unit does not specify which referent it indexes. **b,** Recipient- or pair-level score geometry in Søraas 2019 (19 recipients) and Holland 2024 (61 donor-recipient pairs). The horizontal line denotes the lineage referent \(S=A_L\); the dashed diagonal denotes the host referent \(S=A_R\). Filled circles and open triangles distinguish host-older and host-younger directions. **c,** Sign-balanced affiliation \(J_{\mathrm{aff,bal}}\), which equally weights the two age-difference directions. Grey intervals show the 2.5th–97.5th percentiles of the frozen within-sign geometry-preserving score-pairing null, diamonds its mean and teal circles the observed statistic. The right subpanel shows the observed-minus-null contrast with stratified independent-unit bootstrap 95% intervals. Holland provided the primary estimate (0.626 versus null mean 0.484; contrast interval 0.045–0.209); Søraas supplied separately processed same-direction evidence (0.844 versus 0.697; contrast interval 0.045–0.208). **d,** The empirical counterexample targets semantic inheritance, not age reconstruction as an engineering task: an age-valued score can affiliate with the sampled lineage rather than uniquely indexing the recipient organism. The result does not identify a unique lineage biological age, exclude host influence, establish organismal age replacement or prove rejuvenation. The two programmes partly overlap and are complementary rather than independent replications.

## 中文

**图｜HSCT 暴露年龄值血液读数的参照物歧义。****a，**异基因造血干细胞移植将受者宿主实际年龄 \(A_R\) 与被取样的供者来源造血谱系时间坐标 \(A_L=A_D+t\) 分离。固定的受者血液分数 \(S=q(X_B)\) 以年龄单位表达，但该单位不能确定其参照物。**b，**Søraas 2019（19 名受者）和 Holland 2024（61 对供受者）的受者/配对层级分数几何。水平线表示谱系参照 \(S=A_L\)，虚线对角线表示宿主参照 \(S=A_R\)；实心圆和空心三角分别表示宿主较老和宿主较年轻。**c，**对两个年龄差方向等权的 sign-balanced affiliation \(J_{\mathrm{aff,bal}}\)。灰色区间为冻结的 sign 内、保留年龄几何的分数配对置换零分布 2.5%–97.5% 分位数，菱形为零均值，teal 圆点为观察统计量；右侧显示观察值减零均值及分层独立单位 bootstrap 95% 区间。Holland 提供主估计（0.626 对零均值 0.484；差值区间 0.045–0.209），Søraas 提供独立处理但同方向的互补证据（0.844 对 0.697；差值区间 0.045–0.208）。**d，**该人体反例针对的是语义继承，而非年龄预测这一工程任务：年龄值分数可以指向被取样的谱系，而非唯一指向受者整体。该结果不识别唯一谱系 biological age，不排除宿主影响，不建立受者整体年龄替换，也不证明 rejuvenation。两项研究计划部分重叠，只能作为互补分析而非独立重复验证。

Source Data are supplied as panel-level TSV files in the adjacent `source_data` directory.
"""
    path.write_text(text, encoding="utf-8")


def main() -> int:
    required = [READY_PATH, SUMMARY_PATH, DECISION_PATH, AUDIT_PATH, SYNTHESIS_PATH, SCRIPT_PATH]
    missing = [str(path) for path in required if not path.exists()]
    require(not missing, f"Missing required inputs: {missing}")
    require(not OUT.exists(), f"Output already exists; use a new analysis version: {OUT}")
    require(not STAGE.exists(), f"Staging directory already exists: {STAGE}")

    ready = pd.read_csv(READY_PATH, sep="\t")
    summary = pd.read_csv(SUMMARY_PATH, sep="\t")
    decision = json.loads(DECISION_PATH.read_text(encoding="utf-8"))
    independent_audit = json.loads(AUDIT_PATH.read_text(encoding="utf-8"))

    require(len(ready) == 80, "Expected 80 independent units")
    require(not ready.duplicated(["source_study", "unit_id"]).any(), "Independent-unit duplication")
    require(np.isfinite(ready[["H", "C", "S", "I"]].to_numpy(float)).all(), "Non-finite analysis-ready values")
    require(np.max(np.abs(ready["I"].to_numpy(float))) <= 1 + TOL, "Affiliation index outside bounds")
    require(np.min(np.abs(ready["H_minus_C"].to_numpy(float))) > TOL, "Ineligible H=C unit")
    require(decision.get("placement") == "MAIN", "Frozen placement is not MAIN")
    require(independent_audit.get("status") == "PASS", "Independent audit is not PASS")

    expected_counts = {
        ("Holland2024", "host_older"): 47,
        ("Holland2024", "host_younger"): 14,
        ("Soraas2019", "host_older"): 13,
        ("Soraas2019", "host_younger"): 6,
    }
    observed_counts = ready.groupby(["source_study", "stratum"]).size().to_dict()
    require(observed_counts == expected_counts, f"Stratum counts changed: {observed_counts}")

    for study, group in ready.groupby("source_study", sort=False):
        calculated = 0.5 * (
            group.loc[group["stratum"].eq("host_older"), "I"].mean()
            + group.loc[group["stratum"].eq("host_younger"), "I"].mean()
        )
        stored = float(summary.loc[summary["source_study"].eq(study), "I_bal_observed"].iloc[0])
        require(abs(calculated - stored) <= TOL, f"I_bal mismatch for {study}")

    STAGE.mkdir(parents=True)
    (STAGE / "figures").mkdir()
    (STAGE / "source_data").mkdir()
    (STAGE / "visualization").mkdir()

    panel_b = ready[["analysis_row", "source_study", "unit_id", "stratum", "H", "C", "S", "H_minus_C", "I"]].copy()
    panel_b["S_minus_C"] = panel_b["S"] - panel_b["C"]
    panel_b.to_csv(STAGE / "source_data" / "panel_b_affiliation_geometry.tsv", sep="\t", index=False)

    panel_c_columns = [
        "source_study", "n_overall", "n_host_older", "n_host_younger",
        "I_bal_observed", "I_bal_permutation_null_mean", "I_bal_permutation_q025",
        "I_bal_permutation_q975", "observed_minus_permutation_null_mean",
        "observed_minus_null_bootstrap_ci_low", "observed_minus_null_bootstrap_ci_high",
        "permutation_p_one_sided",
    ]
    summary[panel_c_columns].to_csv(STAGE / "source_data" / "panel_c_referent_null_summary.tsv", sep="\t", index=False)
    pd.DataFrame(
        [
            {"panel": "a", "object": "recipient_host_coordinate", "symbol": "A_R", "definition": "recipient-host chronological age at sampling"},
            {"panel": "a", "object": "donor_lineage_coordinate", "symbol": "A_L=A_D+t", "definition": "donor age plus elapsed time since transplantation"},
            {"panel": "a", "object": "fixed_recipient_blood_score", "symbol": "S=q(X_B)", "definition": "fixed age-valued score from recipient blood"},
            {"panel": "d", "object": "not_identified_interpretation", "symbol": "S=recipient-wide biological age", "definition": "not identified by chronological-age reconstruction"},
            {"panel": "d", "object": "supported_result", "symbol": "sampled-lineage referent affiliation", "definition": "identified in the evaluated HSCT contrast"},
        ]
    ).to_csv(STAGE / "source_data" / "panel_a_d_concept_map.tsv", sep="\t", index=False)

    pd.DataFrame(
        [
            {"panel": "a", "source_file": str(READY_PATH.relative_to(ROOT)), "row_grain": "concept definitions from frozen HSCT estimand"},
            {"panel": "b", "source_file": "source_data/panel_b_affiliation_geometry.tsv", "row_grain": "recipient or donor-recipient pair"},
            {"panel": "c", "source_file": "source_data/panel_c_referent_null_summary.tsv", "row_grain": "study summary"},
            {"panel": "d", "source_file": "source_data/panel_a_d_concept_map.tsv", "row_grain": "claim-boundary object"},
        ]
    ).to_csv(STAGE / "source_data" / "source_data_index.tsv", sep="\t", index=False)

    figure_path = STAGE / "figures" / FIGURE_STEM
    build_figure(ready, summary, figure_path)
    build_legend(STAGE / "FIGURE_LEGEND_BILINGUAL.md")

    chart_contract = {
        "analysis_version": ANALYSIS_VERSION,
        "analytical_question": "Does a fixed age-valued recipient-blood score affiliate with the sampled donor-derived lineage beyond marginal-score and donor/recipient age geometry, thereby exposing failure of organismal semantic inheritance from age reconstruction?",
        "takeaway": "The evaluated score showed greater donor-lineage affiliation than expected under a geometry-preserving score-pairing null; age reconstruction did not fix a unique recipient-organism referent.",
        "surface": "standalone static publication figure",
        "footprint": "180 mm wide, four panels, 600 dpi PNG plus editable SVG and vector PDF",
        "panels": {
            "a": "conceptual HSCT referent-separation diagram",
            "b": "faceted recipient/pair-level scatter with lineage and host reference lines",
            "c": "dot-and-interval comparison of observed affiliation, permutation null and bootstrap contrast",
            "d": "claim-boundary flow diagram",
        },
        "palette_policy": "hard two-root cap",
        "palette": {"teal": TEAL, "orange": ORANGE, "ink": INK, "neutral": MUTED},
        "non_color_distinction": "filled circles versus open triangles; solid versus dashed reference lines; direct labels",
        "data_grain": {"Soraas2019": "19 recipients", "Holland2024": "61 donor-recipient pairs"},
    }
    write_json(STAGE / "visualization" / "chart_contract.json", chart_contract)

    input_manifest = []
    for path in required:
        input_manifest.append(
            {
                "path": str(path.relative_to(ROOT)),
                "sha256": sha256(path),
                "bytes": path.stat().st_size,
            }
        )
    pd.DataFrame(input_manifest).to_csv(STAGE / "visualization" / "input_manifest.tsv", sep="\t", index=False)

    audit = {
        "status": "PASS",
        "analysis_version": ANALYSIS_VERSION,
        "frozen_analysis_version": independent_audit.get("analysis_version"),
        "placement": decision.get("placement"),
        "new_models_fit": False,
        "new_random_numbers_generated": False,
        "n_independent_units": len(ready),
        "study_counts": ready.groupby("source_study").size().to_dict(),
        "stratum_counts": {f"{key[0]}::{key[1]}": value for key, value in observed_counts.items()},
        "affiliation_index_within_bounds": True,
        "i_bal_recomputed_within_tolerance": True,
        "numerical_tolerance": TOL,
        "formats": ["png_600dpi", "svg_editable_text", "pdf_vector"],
        "environment": {
            "python": platform.python_version(),
            "executable": sys.executable,
            "numpy": np.__version__,
            "pandas": pd.__version__,
            "matplotlib": matplotlib.__version__,
        },
    }
    write_json(STAGE / "visualization" / "visualization_audit.json", audit)

    readme = f"""# HSCT age-reconstruction semantic-counterexample figure v0.4

Publication-ready four-panel figure using only frozen `HUM-01` artifacts.

- Figure: `figures/{FIGURE_STEM}.png`, `.svg`, `.pdf`
- Bilingual legend: `FIGURE_LEGEND_BILINGUAL.md`
- Panel-level Source Data: `source_data/`
- Chart contract and audit: `visualization/`
- Reproducible entry point: `analysis/{SCRIPT_PATH.name}`

The figure frames HSCT as an empirical counterexample to semantic inheritance
from chronological-age reconstruction. It does not refit a clock, generate new
random numbers, or change the frozen `HUM-01` estimand, role or placement.
"""
    (STAGE / "README.md").write_text(readme, encoding="utf-8")

    manifest_rows = []
    for path in sorted(STAGE.rglob("*")):
        if path.is_file() and path.name != "figure_manifest.tsv":
            manifest_rows.append(
                {
                    "path": str(path.relative_to(STAGE)),
                    "role": "figure" if path.parent.name == "figures" else "source_or_audit",
                    "bytes": path.stat().st_size,
                    "sha256": sha256(path),
                }
            )
    pd.DataFrame(manifest_rows).to_csv(STAGE / "visualization" / "figure_manifest.tsv", sep="\t", index=False)

    STAGE.rename(OUT)
    print(json.dumps({"status": "COMPLETE", "analysis_version": ANALYSIS_VERSION, "output_dir": str(OUT)}, ensure_ascii=False))
    return 0


if __name__ == "__main__":
    raise SystemExit(main())
```
