## Supplementary_Software_1 for "Inferential boundaries of age prediction: why prediction does not establish biological age measurement": plot_hsct_age_reconstruction_semantic_counterexample_v0_5.py

```
#!/usr/bin/env python
"""Text/layout-only HSCT figure derivative; no fitting or statistical estimation.

Use frozen v0.4 numerical plotting functions and figure assembly.  Only panels
a/d are replaced. Quantitative source tables are copied without serialization.
"""

from __future__ import annotations

import importlib.util
import json
import shutil
from pathlib import Path

import pandas as pd

ROOT = Path(__file__).resolve().parents[1]
OLD_SCRIPT = ROOT / "analysis/plot_hsct_age_reconstruction_semantic_counterexample_v0_4.py"
OLD_OUT = ROOT / "outputs/hsct_age_reconstruction_semantic_counterexample_figure_v0_4"
OUT = ROOT / "outputs/hsct_age_reconstruction_semantic_counterexample_figure_v0_5"

spec = importlib.util.spec_from_file_location("frozen_hsct_figure_v0_4", OLD_SCRIPT)
if spec is None or spec.loader is None:
    raise RuntimeError("Cannot load frozen plotting module")
old = importlib.util.module_from_spec(spec)
spec.loader.exec_module(old)

BOX_ITEMS = []


def checked_box(ax, xy, width, height, text, **kwargs):
    patch = old.box(ax, xy, width, height, text, **kwargs)
    BOX_ITEMS.append((patch, ax.texts[-1]))
    return patch


def draw_panel_a(ax):
    ax.set_axis_off()
    old.panel_label(ax, "a", "HSCT separates candidate referents")
    checked_box(ax, (0.03, 0.72), 0.33, 0.18,
                "Donor\nchronological age\n$A_D$",
                facecolor=old.ORANGE_LIGHT, edgecolor=old.ORANGE, fontsize=6.6)
    checked_box(ax, (0.63, 0.72), 0.33, 0.18,
                "Recipient host\nchronological age\n$A_R$",
                facecolor=old.TEAL_LIGHT, edgecolor=old.TEAL, fontsize=6.6)
    old.arrow(ax, (0.38, 0.805), (0.60, 0.805), color=old.ORANGE, linewidth=1.5)
    ax.text(0.49, 0.93, "allogeneic HSCT", transform=ax.transAxes,
            ha="center", va="center", fontsize=6.3, color=old.INK)
    ax.text(0.50, 0.66, "after $t$ years", transform=ax.transAxes,
            ha="center", va="center", fontsize=6.5, color=old.INK)
    old.arrow(ax, (0.20, 0.70), (0.31, 0.62), color=old.ORANGE)
    old.arrow(ax, (0.79, 0.70), (0.69, 0.62), color=old.TEAL)
    checked_box(ax, (0.08, 0.43), 0.84, 0.18,
                "One recipient, two observable coordinates\n"
                "Lineage  $A_L = A_D + t$\nHost  $A_R$",
                facecolor=old.WHITE, edgecolor=old.INK, fontsize=6.6, weight="bold")
    old.arrow(ax, (0.50, 0.41), (0.50, 0.35), color=old.MUTED)
    checked_box(ax, (0.04, 0.16), 0.39, 0.18,
                "Recipient blood\nfrom donor lineage\n$X_B$",
                facecolor=old.ORANGE_LIGHT, edgecolor=old.ORANGE, fontsize=6.6)
    old.arrow(ax, (0.45, 0.25), (0.56, 0.25), color=old.MUTED)
    checked_box(ax, (0.58, 0.16), 0.38, 0.18,
                "Frozen age decoder\n$S=q(X_B)$\n(years)",
                facecolor=old.TEAL_LIGHT, edgecolor=old.TEAL,
                fontsize=6.6, weight="bold")
    ax.text(0.50, 0.045,
            "Question: how does the age-valued readout\nalign with $A_R$ and $A_L$?",
            transform=ax.transAxes, ha="center", va="center", fontsize=6.7,
            color=old.INK, fontweight="bold", linespacing=1.12)


def draw_panel_d(ax):
    ax.set_axis_off()
    old.panel_label(ax, "d", "Empirical consequence for age reconstruction")
    checked_box(ax, (0.06, 0.74), 0.88, 0.18,
                "Source task\n$X_B$ → $q(X_B)=S$\nReconstruct chronological age $A$",
                facecolor=old.LIGHT, edgecolor=old.MUTED, fontsize=6.7)
    old.arrow(ax, (0.50, 0.72), (0.50, 0.70), color=old.MUTED)
    checked_box(ax, (0.06, 0.49), 0.88, 0.19,
                "HSCT observation\nDonor-lineage affiliation exceeds\nthe score-pairing null",
                facecolor=old.TEAL_LIGHT, edgecolor=old.TEAL,
                fontsize=6.7, weight="bold")
    old.arrow(ax, (0.50, 0.47), (0.50, 0.43), color=old.MUTED)
    checked_box(ax, (0.04, 0.21), 0.42, 0.19,
                "Recipient-organism\nbiological-age identity\nnot established",
                facecolor=old.ORANGE_LIGHT, edgecolor=old.ORANGE,
                fontsize=7.0, weight="bold")
    checked_box(ax, (0.54, 0.21), 0.42, 0.19,
                "Excess donor-lineage\naffiliation under the\nscore-pairing null",
                facecolor=old.TEAL_LIGHT, edgecolor=old.TEAL,
                fontsize=7.0, weight="bold")
    ax.text(0.50, 0.09,
            "The age unit does not establish\nthe score's biological object",
            transform=ax.transAxes, ha="center", va="center", fontsize=7.7,
            color=old.INK, fontweight="bold", linespacing=1.12)
    ax.text(0.50, 0.005,
            "The null also breaks score–host-age dependence.",
            transform=ax.transAxes, ha="center", va="center", fontsize=6.1,
            color=old.MUTED, style="italic")


def main():
    for path in (OLD_SCRIPT, OLD_OUT, OUT / "README.md", OUT / "FIGURE_LEGEND_BILINGUAL.md",
                 old.READY_PATH, old.SUMMARY_PATH):
        old.require(path.exists(), f"Required source missing: {path}")
    old.require(not (OUT / "DATA_PARITY_VALIDATION.json").exists(),
                "Validated output already exists; use a new version")
    frozen_paths = [OLD_SCRIPT, old.READY_PATH, old.SUMMARY_PATH]
    frozen_paths += list((OLD_OUT / "source_data").glob("*.tsv"))
    before = {str(p): old.sha256(p) for p in frozen_paths}

    (OUT / "figures").mkdir(exist_ok=True)
    (OUT / "source_data").mkdir(exist_ok=True)
    tables = []
    for filename in ("panel_b_affiliation_geometry.tsv", "panel_c_referent_null_summary.tsv",
                     "source_data_index.tsv"):
        source = OLD_OUT / "source_data" / filename
        target = OUT / "source_data" / filename
        shutil.copyfile(source, target)
        old.require(source.read_bytes() == target.read_bytes(), f"Byte parity failed: {filename}")
        left, right = (pd.read_csv(p, sep="\t") for p in (source, target))
        pd.testing.assert_frame_equal(left, right, check_exact=True)
        tables.append({"file": filename, "rows": len(left), "columns": len(left.columns),
                       "sha256": old.sha256(source), "byte_parity": "PASS",
                       "exact_table_parity": "PASS"})
    concept_map = pd.DataFrame([
        {"panel": "a", "object": "recipient_host_coordinate", "symbol": "A_R",
         "definition": "recipient-host chronological age at sampling"},
        {"panel": "a", "object": "donor_lineage_coordinate", "symbol": "A_L=A_D+t",
         "definition": "donor age plus elapsed time since transplantation"},
        {"panel": "a", "object": "fixed_recipient_blood_score", "symbol": "S=q(X_B)",
         "definition": "fixed age-valued score from recipient blood"},
        {"panel": "d", "object": "not_established_identity", "symbol": "recipient-organism biological-age identity",
         "definition": "not established by the chronological-age reconstruction task; not empirically refuted by the contrast"},
        {"panel": "d", "object": "observed_result", "symbol": "excess donor-lineage affiliation under the score-pairing null",
         "definition": "the null also breaks score-host-age dependence; not donor influence conditional on host age"},
    ])
    concept_map.to_csv(OUT / "source_data/panel_a_d_concept_map.tsv", sep="\t", index=False)

    # Same persisted numerical inputs, same b/c functions, same figure gridspec.
    ready = pd.read_csv(old.READY_PATH, sep="\t")
    summary = pd.read_csv(old.SUMMARY_PATH, sep="\t")
    frozen_b, frozen_c = old.draw_panel_b, old.draw_panel_c
    old.draw_panel_a, old.draw_panel_d = draw_panel_a, draw_panel_d
    old.require(old.draw_panel_b is frozen_b and old.draw_panel_c is frozen_c,
                "Quantitative plotting functions changed")
    old.build_figure(ready, summary, OUT / "figures" / old.FIGURE_STEM)

    # Layout-only containment check; no analysis estimand is calculated here.
    figure = BOX_ITEMS[0][0].figure
    figure.canvas.draw()
    renderer = figure.canvas.get_renderer()
    containment = []
    for patch, label in BOX_ITEMS:
        pb, tb = patch.get_window_extent(renderer), label.get_window_extent(renderer)
        contained = tb.x0 >= pb.x0 and tb.x1 <= pb.x1 and tb.y0 >= pb.y0 and tb.y1 <= pb.y1
        containment.append({"text": label.get_text(), "contained": bool(contained)})
        old.require(contained, f"Schematic label exceeds its box: {label.get_text()}")
    after = {str(p): old.sha256(p) for p in frozen_paths}
    old.require(before == after, "A frozen input changed")

    report = {
        "status": "PASS", "derivative": OUT.name, "source_figure": OLD_OUT.name,
        "scope": "a/d semantics and layout only; b/c numerical source and plotting unchanged",
        "fitting": False, "random_draws": False, "statistical_estimation": False,
        "frozen_input_identity": "PASS", "unchanged_b_c_plotting_functions": "PASS",
        "tables": tables, "schematic_label_containment": containment,
        "frozen_input_sha256": before,
        "outputs": [str(OUT / "figures" / (old.FIGURE_STEM + suffix))
                    for suffix in (".png", ".svg", ".pdf")],
    }
    old.write_json(OUT / "DATA_PARITY_VALIDATION.json", report)
    lines = ["# Data and layout parity validation", "", "Status: PASS", "",
             "- Quantitative panel b/c TSVs: byte-identical and exact table equality with v0.4.",
             "- Frozen analysis inputs and all original source TSVs: unchanged.",
             "- Original b/c plotting functions and figure assembly: reused unchanged.",
             "- Schematic box labels: contained within their boxes.",
             "- No fitted objects, random numbers or new statistical estimates.", "",
             "See DATA_PARITY_VALIDATION.json for per-file hashes and exact checks."]
    (OUT / "VALIDATION_REPORT.md").write_text("\n".join(lines) + "\n", encoding="utf-8")
    print(json.dumps({"status": "PASS", "output": str(OUT), "tables": tables}, indent=2))


if __name__ == "__main__":
    main()
```
