## Supplementary_Software_1 for "Inferential boundaries of age prediction: why prediction does not establish biological age measurement": plot_nhanes_hum03_hum04_hum06_extended_data_figure3_submission_v0_3.py

```
#!/usr/bin/env python
"""Build the fit-free three-panel submission successor for Extended Data Fig. 3.

The script reuses the frozen v0.2 evidence gate and changes only the
submission-facing layout. It reads tabular and JSON evidence; it does not load
model objects, fit or tune estimators, resample participants or generate random
numbers.
"""

from __future__ import annotations

import argparse
import importlib.util
import json
import platform
import sys
from pathlib import Path
from typing import Any

import matplotlib.pyplot as plt
import pandas as pd


ROOT = Path(__file__).resolve().parents[1]
SCRIPT = Path(__file__).resolve()
LEGACY_SCRIPT = (
    ROOT
    / "analysis"
    / "plot_nhanes_hum03_hum04_hum06_figure3_successor_v0_2.py"
)
CONTRACT = (
    ROOT
    / "analysis"
    / "NHANES_HUM03_HUM04_HUM06_EXTENDED_DATA_FIGURE3_SUBMISSION_CONTRACT_v0_3_20260828.md"
)
OUT = (
    ROOT
    / "outputs"
    / "nhanes_hum03_hum04_hum06_extended_data_figure3_submission_v0_3"
)


def load_legacy_module() -> Any:
    spec = importlib.util.spec_from_file_location("nhanes_ed3_v0_2", LEGACY_SCRIPT)
    if spec is None or spec.loader is None:
        raise RuntimeError(f"Could not load frozen evidence gate: {LEGACY_SCRIPT}")
    module = importlib.util.module_from_spec(spec)
    spec.loader.exec_module(module)
    return module


legacy = load_legacy_module()

INK = legacy.INK
NAVY = legacy.NAVY
MUTED = legacy.MUTED
GRID = legacy.GRID
LIGHT = legacy.LIGHT
WHITE = legacy.WHITE
BLUE = legacy.BLUE
BLUE_LIGHT = legacy.BLUE_LIGHT
TEAL = legacy.TEAL
TEAL_LIGHT = legacy.TEAL_LIGHT

INPUTS = [
    ("submission_contract_v0_3", CONTRACT),
    ("submission_plot_script_v0_3", SCRIPT),
    ("frozen_evidence_gate_v0_2", LEGACY_SCRIPT),
] + [
    (role, path)
    for role, path in legacy.INPUTS
    if role not in {"chart_contract", "plot_script"}
]


def require(condition: bool, message: str) -> None:
    if not condition:
        raise RuntimeError(message)


def write_json(path: Path, payload: Any) -> None:
    path.write_text(
        json.dumps(payload, indent=2, ensure_ascii=False) + "\n",
        encoding="utf-8",
    )


def ensure_package() -> None:
    require(OUT.name.endswith("_v0_3"), f"Output is not versioned v0.3: {OUT}")
    (OUT / "figures").mkdir(parents=True, exist_ok=True)
    (OUT / "source_data").mkdir(parents=True, exist_ok=True)


def write_input_manifest() -> Path:
    rows: list[dict[str, Any]] = []
    for role, path in INPUTS:
        require(path.exists(), f"Required frozen input is missing: {path}")
        require(
            path.suffix.lower() not in {".joblib", ".pkl", ".pickle"},
            f"Fitted-model input is forbidden: {path}",
        )
        rows.append(
            {
                "role": role,
                "path": str(path.relative_to(ROOT)),
                "bytes": path.stat().st_size,
                "sha256": legacy.sha256(path),
            }
        )
    output = OUT / "INPUT_MANIFEST.tsv"
    pd.DataFrame(rows).to_csv(output, sep="\t", index=False)
    return output


def write_output_manifest() -> Path:
    rows: list[dict[str, Any]] = []
    for path in sorted(
        item
        for item in OUT.rglob("*")
        if item.is_file() and item.name != "OUTPUT_MANIFEST.tsv"
    ):
        rows.append(
            {
                "path": str(path.relative_to(ROOT)),
                "bytes": path.stat().st_size,
                "sha256": legacy.sha256(path),
            }
        )
    output = OUT / "OUTPUT_MANIFEST.tsv"
    pd.DataFrame(rows).to_csv(output, sep="\t", index=False)
    return output


def draw_complete_contrasts(ax: plt.Axes, data: dict[str, Any]) -> None:
    age_rows = data["age_rows"].set_index("contrast_id")
    legacy.panel_label(ax, "a", "Complete observed-age and block contrast set")
    legacy.style_axis(ax, grid_axis="x")
    ax.axvline(0, color=MUTED, linewidth=1.0, linestyle=(0, (4, 2)))
    order = ["V_F", "V_R", "P_proxy", "D_block_0", "D_block_1", "D_replace"]
    labels = [
        "$V_F$  Age gain after full 25",
        "$V_R$  Age gain after remaining 20",
        "$P_{proxy}$  Extra predictive value of age\nafter block removal",
        "$D_{block,0}$  Block gain without age",
        "$D_{block,1}$  Block gain with age",
        "$D_{replace}$  Remaining 20 + age vs full 25",
    ]
    y_values = [5.0, 4.0, 3.0, 1.8, 0.8, -0.4]
    styles = [
        (TEAL, "o", False),
        (TEAL, "o", True),
        (NAVY, "D", True),
        (BLUE, "s", True),
        (BLUE, "s", False),
        (NAVY, "D", False),
    ]
    ax.axhspan(2.58, 3.42, color=LIGHT, zorder=-1)
    ax.axhline(2.4, color=GRID, linewidth=0.75)
    ax.axhline(0.2, color=GRID, linewidth=0.75)
    for contrast_id, y, (color, marker, filled) in zip(
        order, y_values, styles, strict=True
    ):
        row = age_rows.loc[contrast_id]
        legacy.plot_interval(
            ax,
            y,
            row,
            color=color,
            marker=marker,
            filled=filled,
            label_offset=0.00035,
        )
        ax.text(
            0.99,
            y,
            f"{int(row['positive_outer_folds'])}/5 +",
            transform=ax.get_yaxis_transform(),
            ha="right",
            va="bottom",
            fontsize=5.0,
            color=MUTED,
        )
    ax.set_yticks(y_values, labels)
    ax.set_ylim(-1.0, 5.62)
    ax.set_xlim(-0.009, 0.019)
    ax.set_xlabel("Declared paired held-out log-loss contrast", fontsize=5.85)
    ax.text(
        0.01,
        -0.25,
        "Intervals condition on the frozen Phase 1 mask; crossing zero is not equivalence.",
        transform=ax.transAxes,
        fontsize=5.15,
        color=MUTED,
    )


def draw_substitution(ax: plt.Axes, data: dict[str, Any]) -> None:
    row = data["substitution_row"].iloc[0]
    legacy.panel_label(ax, "b", "Observed age versus the transported age decoder")
    legacy.style_axis(ax, grid_axis="x")
    ax.axvline(0, color=MUTED, linewidth=1.0, linestyle=(0, (4, 2)))
    legacy.box(
        ax,
        (0.03, 0.70),
        0.39,
        0.14,
        "Observed age $A$\n$W+s(A)$",
        face=BLUE_LIGHT,
        edge=BLUE,
        fontsize=6.0,
        weight="bold",
    )
    legacy.box(
        ax,
        (0.58, 0.70),
        0.39,
        0.14,
        "Transported decoder $q_A$\n$W+s(q_A)$",
        face=TEAL_LIGHT,
        edge=TEAL,
        fontsize=6.0,
        weight="bold",
    )
    ax.text(
        0.50,
        0.64,
        "same background · one scalar · spline · tuning grid · folds · weights",
        transform=ax.transAxes,
        ha="center",
        fontsize=5.15,
        color=MUTED,
    )
    y_value = 0.30
    legacy.plot_interval(
        ax,
        y_value,
        row,
        color=NAVY,
        marker="D",
        filled=False,
        label_offset=0.00025,
    )
    ax.set_yticks([y_value], ["$\\Delta_{sub}$\n$q_A$ instead of age"])
    ax.set_ylim(-0.12, 1.08)
    ax.set_xlim(-0.0061, 0.0052)
    ax.set_xlabel("$L(W+s(A))-L(W+s(q_A))$  (right favours $q_A$)", fontsize=5.7)
    ax.text(
        0.01,
        0.06,
        "No clear substitution advantage · 1/5 folds favour $q_A$",
        transform=ax.transAxes,
        fontsize=5.55,
        color=NAVY,
        fontweight="bold",
    )
    ax.text(
        0.01,
        -0.25,
        "The interval spans zero; this is not evidence that the two inputs are equivalent.",
        transform=ax.transAxes,
        fontsize=5.2,
        color=MUTED,
    )


def draw_fold_directions(ax: plt.Axes, data: dict[str, Any]) -> None:
    fold_diagnostics = data["fold_diagnostics"]
    legacy.panel_label(ax, "c", "Direction across the five fixed outer folds")
    ax.spines[["top", "right", "left", "bottom"]].set_visible(False)
    ax.tick_params(axis="both", colors=INK, labelsize=6.0, length=0)
    order = ["V_F", "P_proxy", "U_A", "Delta", "Delta_sub"]
    labels = [
        "$V_F$  Age after full 25",
        "$P_{proxy}$  Extra predictive value of age after block removal",
        "$U_A$  Age-decoder utility",
        "$\\Delta$  Mortality- over age-trained",
        "$\\Delta_{sub}$  $q_A$ instead of observed age",
    ]
    y_values = list(range(len(order)))[::-1]
    for y, contrast_id in zip(y_values, order, strict=True):
        subset = fold_diagnostics.loc[
            fold_diagnostics["contrast_id"] == contrast_id
        ].sort_values("outer_fold")
        require(len(subset) == 5, f"Fold diagnostic rows changed for {contrast_id}")
        for _, row in subset.iterrows():
            x = int(row["outer_fold"]) + 1
            positive = bool(row["positive"])
            ax.scatter(
                x,
                y,
                s=58,
                marker="o" if positive else "D",
                facecolor=TEAL if positive else WHITE,
                edgecolor=INK if positive else BLUE,
                linewidth=0.6 if positive else 1.25,
                zorder=2,
            )
            ax.text(
                x,
                y,
                "+" if positive else "−",
                ha="center",
                va="center",
                fontsize=5.0,
                color=WHITE if positive else BLUE,
                fontweight="bold",
                zorder=3,
            )
        ax.text(
            5.62,
            y,
            f"{int(subset['positive'].sum())}/5 +",
            ha="left",
            va="center",
            fontsize=5.55,
            color=MUTED,
        )
    ax.set_xticks(range(1, 6), [f"Fold {value}" for value in range(1, 6)])
    ax.set_yticks(y_values, labels)
    ax.set_xlim(0.55, 6.15)
    ax.set_ylim(-0.65, 4.65)
    for x in range(1, 6):
        ax.axvline(x, color=GRID, linewidth=0.45, zorder=0)
    ax.text(
        0.01,
        0.93,
        "Filled circle = positive; open diamond = negative",
        transform=ax.transAxes,
        fontsize=5.4,
        color=MUTED,
    )
    ax.text(
        0.01,
        -0.20,
        "Signs follow each declared formula; folds are diagnostics, not independent replications.",
        transform=ax.transAxes,
        fontsize=5.25,
        color=MUTED,
    )


def build(data: dict[str, Any], visual_qa_status: str) -> dict[str, Path]:
    ensure_package()
    write_input_manifest()
    fig = plt.figure(figsize=(7.15, 5.75), facecolor=WHITE)
    grid = fig.add_gridspec(
        2,
        2,
        left=0.09,
        right=0.985,
        bottom=0.075,
        top=0.90,
        height_ratios=[1.0, 0.86],
        hspace=0.55,
        wspace=0.48,
    )
    ax_a = fig.add_subplot(grid[0, 0])
    ax_b = fig.add_subplot(grid[0, 1])
    ax_c = fig.add_subplot(grid[1, :])
    fig.suptitle(
        "Extended Data Figure 3 | NHANES age-context, substitution and fold diagnostics",
        x=0.09,
        y=0.975,
        ha="left",
        fontsize=9.75,
        fontweight="bold",
        color=INK,
    )
    draw_complete_contrasts(ax_a, data)
    draw_substitution(ax_b, data)
    draw_fold_directions(ax_c, data)
    stem = "Extended_Data_Figure_3_NHANES_age_context_substitution_and_fold_diagnostics"
    figures = legacy.save_figure(fig, OUT, stem)

    panel_a = OUT / "source_data" / "panel_a_six_contrast_forest.tsv"
    panel_b = OUT / "source_data" / "panel_b_actual_age_vs_qA_substitution.tsv"
    panel_c = OUT / "source_data" / "panel_c_fold_direction_diagnostics.tsv"
    data["age_rows"].to_csv(panel_a, sep="\t", index=False)
    data["substitution_row"].to_csv(panel_b, sep="\t", index=False)
    data["fold_diagnostics"].to_csv(panel_c, sep="\t", index=False)
    pd.DataFrame(
        [
            ("a", panel_a.name, "complete six-contrast observed-age and block forest with Fay-BRR intervals"),
            ("b", panel_b.name, "matched observed-age versus transported-q_A substitution boundary"),
            ("c", panel_c.name, "five fixed outer-fold sign diagnostics for decision-facing contrasts"),
        ],
        columns=["panel", "file", "content"],
    ).to_csv(OUT / "source_data" / "source_data_index.tsv", sep="\t", index=False)

    legend = """# Extended Data Figure 3 | Observed-age context, direct substitution and fold direction

**a**, Complete NHANES III Phase 2 contrast set for the four frozen endpoint models using all 25 measurements or the remaining 20 after the Phase 1 age-selected five-measurement block was withheld, without or with observed chronological age. Points are design-weighted paired held-out log-loss contrasts and lines are 95% Fay-BRR intervals. V_F and V_R are the increments from adding observed age; P_proxy is their difference; D_block,0 and D_block,1 are the block contributions without and with age; D_replace compares remaining 20 plus age with the full 25 without age. The intervals condition on the frozen Phase 1 mask. The remaining 20 are not asserted to be non-age-related, and an interval spanning zero is not equivalence. **b**, Direct substitution boundary between observed age and the transported Phase 1 age decoder under the same background, one-scalar spline family, tuning grid, folds and weights. Delta_sub was −0.000757 (95% Fay-BRR interval, −0.004661 to 0.003147), with one of five folds positive. There was no clear substitution advantage; this does not establish equality. **c**, Direction of five decision-facing paired contrasts in the five fixed primary-sampling-unit-atomic outer folds. Filled circles denote positive and open diamonds negative values under each declared formula. Fold directions are consistency diagnostics, not independent replications. Model-persistence and reload-parity records are retained in the reproducibility manifest rather than displayed as a scientific panel.

# 扩展数据图 3｜真实年龄背景效用、直接替代与逐折方向

**a**，NHANES III Phase 2 的完整六对比集合。四个冻结终点模型使用全部 25 项测量，或使用移除 Phase 1 年龄选择五项 block 后的 remaining-20，并分别不加入或加入真实时序年龄。点为设计加权的配对留出 log-loss 对比，线为 95% Fay-BRR 区间。V_F 与 V_R 为加入真实年龄的增量；P_proxy 为二者之差；D_block,0 与 D_block,1 分别为无年龄和有年龄时五项 block 的贡献；D_replace 比较 remaining-20 加年龄与完整 25 项但不加年龄。区间条件于冻结的 Phase 1 mask。remaining-20 不被称为“非年龄相关”，区间跨零也不代表等价。**b**，在相同背景、单标量 spline 模型族、调参网格、折和权重下，直接比较真实年龄与从 Phase 1 转移的年龄解码器。Delta_sub 为 −0.000757（95% Fay-BRR 区间 −0.004661 至 0.003147），五折中一折为正；没有明确的替代优势，但这不建立相等。**c**，五个决策相关配对对比在五个固定 PSU-atomic 外层折中的方向。实心圆表示按各自公式为正，空心菱形表示为负；逐折方向是一致性诊断，不是独立重复。模型持久化和重载一致性记录保留在可重复性清单中，不再作为科学 panel 展示。
"""
    (OUT / "FIGURE_LEGEND_BILINGUAL.md").write_text(legend, encoding="utf-8")
    (OUT / "README.md").write_text(
        "# NHANES Extended Data Figure 3 submission successor v0.3\n\n"
        "Fit-free three-panel presentation successor. The v0.2 engineering "
        "persistence panel is retained in repository provenance and omitted from "
        "the submission-facing scientific figure. No estimate or estimand changed.\n",
        encoding="utf-8",
    )
    write_json(
        OUT / "CHART_CONTRACT.json",
        {
            "figure": "Extended Data Figure 3 submission successor v0.3",
            "surface": "standalone 180-mm static Extended Data figure",
            "renderer": "Matplotlib via pinned lhstate environment",
            "layout": "two upper panels and one full-width lower panel",
            "panels": {
                "a": "complete six-contrast Fay-BRR forest",
                "b": "observed age versus transported q_A substitution boundary",
                "c": "five fixed outer-fold direction matrix",
            },
            "removed_from_scientific_display": "engineering persistence panel; retained in reproducibility manifest",
            "metric_boundary": "paired held-out log-loss contrasts; interval overlap with zero is not equivalence",
            "exports": ["600 dpi PNG", "editable SVG", "vector PDF"],
            "fit_free": True,
        },
    )
    audit_status = "PASS" if visual_qa_status == "pass" else "PENDING_MANUAL_VISUAL_QA"
    write_json(
        OUT / "VISUAL_AUDIT.json",
        {
            "status": audit_status,
            "automated_source_schema_checks": "PASS",
            "manual_visual_qa": visual_qa_status.upper(),
            "manual_visual_qa_checks": {
                "all_six_intervals_and_zero_line_legible": visual_qa_status == "pass",
                "actual_age_vs_qA_direction_unambiguous": visual_qa_status == "pass",
                "fold_sign_matrix_readable_without_colour": visual_qa_status == "pass",
                "no_clipping_collision_or_detached_annotation": visual_qa_status == "pass",
            },
            "fit_free": True,
            "fitted_models_loaded": False,
            "random_numbers_generated": False,
            "exports_created": sorted(figures),
        },
    )
    write_output_manifest()
    return figures


def parse_args() -> argparse.Namespace:
    parser = argparse.ArgumentParser(description=__doc__)
    parser.add_argument(
        "--visual-qa-status",
        choices=["pending", "pass"],
        default="pending",
    )
    return parser.parse_args()


def main() -> int:
    args = parse_args()
    data = legacy.load_and_gate()
    figures = build(data, args.visual_qa_status)
    summary = {
        "status": (
            "PASS_AUTOMATED_BUILD_AND_MANUAL_VISUAL_QA"
            if args.visual_qa_status == "pass"
            else "PASS_AUTOMATED_BUILD_VISUAL_QA_PENDING"
        ),
        "output": str(OUT),
        "figures": {key: str(path) for key, path in figures.items()},
        "fit_free": True,
        "models_loaded": False,
        "random_numbers_generated": False,
        "visual_qa_status": args.visual_qa_status,
        "python": sys.version,
        "python_executable": sys.executable,
        "platform": platform.platform(),
    }
    write_json(OUT / "BUILD_SUMMARY.json", summary)
    write_output_manifest()
    print(json.dumps(summary, indent=2, ensure_ascii=False))
    return 0


if __name__ == "__main__":
    raise SystemExit(main())
```
