## Supplementary_Software_1 for "Inferential boundaries of age prediction: why prediction does not establish biological age measurement": plot_nhanes_hum03_hum04_hum06_figure3_successor_v0_3.py

```
#!/usr/bin/env python
"""Build fit-free NHANES Figure 3 successor v0.3.

The builder reuses the v0.2 frozen-input integrity gate, then changes only the
reader-facing panel hierarchy. It reads TSV/JSON evidence only; it does not
load a fitted model, fit or tune an estimator, resample participants, or
generate random numbers.
"""

from __future__ import annotations

import argparse
import importlib.util
import json
import platform
import sys
from pathlib import Path
from typing import Any

import matplotlib.pyplot as plt
import pandas as pd


ROOT = Path(__file__).resolve().parents[1]
SCRIPT = Path(__file__).resolve()
LEGACY_SCRIPT = (
    ROOT
    / "analysis"
    / "plot_nhanes_hum03_hum04_hum06_figure3_successor_v0_2.py"
)
CONTRACT = (
    ROOT
    / "analysis"
    / "NHANES_HUM03_HUM04_HUM06_FIGURE3_SUCCESSOR_CHART_CONTRACT_v0_3_20260827.md"
)
OUT = ROOT / "outputs" / "nhanes_hum03_hum04_hum06_figure3_successor_v0_3"


def load_legacy_module() -> Any:
    spec = importlib.util.spec_from_file_location("nhanes_figure3_v0_2", LEGACY_SCRIPT)
    if spec is None or spec.loader is None:
        raise RuntimeError(f"Could not load frozen gate helper: {LEGACY_SCRIPT}")
    module = importlib.util.module_from_spec(spec)
    spec.loader.exec_module(module)
    return module


legacy = load_legacy_module()

INK = legacy.INK
NAVY = legacy.NAVY
MUTED = legacy.MUTED
GRID = legacy.GRID
LIGHT = legacy.LIGHT
WHITE = legacy.WHITE
BLUE = legacy.BLUE
BLUE_LIGHT = legacy.BLUE_LIGHT
TEAL = legacy.TEAL
TEAL_LIGHT = legacy.TEAL_LIGHT
RED = legacy.RED
RED_LIGHT = legacy.RED_LIGHT

INPUTS = [
    ("chart_contract_v0_3", CONTRACT),
    ("plot_script_v0_3", SCRIPT),
    ("frozen_gate_and_render_helpers_v0_2", LEGACY_SCRIPT),
] + [
    (role, path)
    for role, path in legacy.INPUTS
    if role not in {"chart_contract", "plot_script"}
]


def require(condition: bool, message: str) -> None:
    if not condition:
        raise RuntimeError(message)


def ensure_package() -> None:
    require(OUT.name.endswith("_v0_3"), f"Output is not versioned v0.3: {OUT}")
    (OUT / "figures").mkdir(parents=True, exist_ok=True)
    (OUT / "source_data").mkdir(parents=True, exist_ok=True)


def write_json(path: Path, payload: Any) -> None:
    path.write_text(
        json.dumps(payload, indent=2, ensure_ascii=False) + "\n",
        encoding="utf-8",
    )


def write_input_manifest() -> Path:
    rows: list[dict[str, Any]] = []
    for role, path in INPUTS:
        require(path.exists(), f"Required frozen input is missing: {path}")
        require(
            path.suffix.lower() not in {".joblib", ".pkl", ".pickle"},
            f"Fitted-model input is forbidden: {path}",
        )
        rows.append(
            {
                "role": role,
                "path": str(path.relative_to(ROOT)),
                "bytes": path.stat().st_size,
                "sha256": legacy.sha256(path),
            }
        )
    output = OUT / "INPUT_MANIFEST.tsv"
    pd.DataFrame(rows).to_csv(output, sep="\t", index=False)
    return output


def write_output_manifest() -> Path:
    rows: list[dict[str, Any]] = []
    for path in sorted(
        item
        for item in OUT.rglob("*")
        if item.is_file() and item.name != "OUTPUT_MANIFEST.tsv"
    ):
        rows.append(
            {
                "path": str(path.relative_to(ROOT)),
                "bytes": path.stat().st_size,
                "sha256": legacy.sha256(path),
            }
        )
    output = OUT / "OUTPUT_MANIFEST.tsv"
    pd.DataFrame(rows).to_csv(output, sep="\t", index=False)
    return output


def draw_design(ax: plt.Axes, diagnostics: dict[str, Any]) -> pd.DataFrame:
    ax.set_axis_off()
    legacy.panel_label(ax, "a", "Age-supervised decoder and direct-source design")
    legacy.box(
        ax,
        (-0.06, 0.67),
        0.415,
        0.16,
        "Same 50 measurements $X$\n+ background $W$",
        face=BLUE_LIGHT,
        edge=BLUE,
        fontsize=6.35,
        weight="bold",
    )
    legacy.arrow(ax, (0.365, 0.75), (0.40, 0.75), color=BLUE)
    legacy.box(
        ax,
        (0.41, 0.65),
        0.27,
        0.20,
        "Fold-local ridge\nage decoder\ntrained only on $A$",
        face=TEAL_LIGHT,
        edge=TEAL,
        fontsize=6.45,
        weight="bold",
    )
    legacy.arrow(ax, (0.68, 0.75), (0.79, 0.75), color=TEAL)
    legacy.box(
        ax,
        (0.80, 0.67),
        0.17,
        0.16,
        "OOF decoder\n$q_A(W,X)$",
        face=WHITE,
        edge=INK,
        fontsize=6.25,
        weight="bold",
    )
    ax.text(
        0.53,
        0.56,
        f"held-out age $R^2$ = {diagnostics['r2']:.2f}; 5 outer folds",
        transform=ax.transAxes,
        ha="center",
        fontsize=6.0,
        color=MUTED,
    )
    for y, text_value, face, edge in zip(
        [0.36, 0.20, 0.04],
        [
            "$M_{WA}$: age + background",
            "$M_{CLOCK}$: age + background + OOF decoder",
            "$M_X$: age + background + the same 50 measurements",
        ],
        [LIGHT, TEAL_LIGHT, BLUE_LIGHT],
        [MUTED, TEAL, BLUE],
    ):
        legacy.box(
            ax,
            (0.10, y),
            0.80,
            0.105,
            text_value,
            face=face,
            edge=edge,
            fontsize=6.0,
            weight="bold",
        )
    ax.text(
        0.50,
        -0.055,
        "NHANES 2007–2012 · $n=13{,}724$ · 470 deaths",
        transform=ax.transAxes,
        ha="center",
        fontsize=5.75,
        color=MUTED,
        fontstyle="italic",
    )
    return pd.DataFrame(
        [
            ("input", "W + ordered 50-measurement panel", "same source panel for decoder and direct endpoint model"),
            ("decoder", "fold-local ridge q_A(W,X)", "trained only to reconstruct chronological age"),
            ("endpoint model", "M_WA", "age + background"),
            ("endpoint model", "M_CLOCK", "age + background + OOF q_A"),
            ("endpoint model", "M_X", "age + background + direct 50 measurements"),
            ("validation", "five outer folds", "all reported predictions are participant-level OOF"),
            ("boundary", "construct identity", "not identified by age reconstruction or endpoint utility"),
        ],
        columns=["stage", "object", "detail"],
    )


def draw_continuous_contrasts(ax: plt.Axes, rows: pd.DataFrame) -> None:
    legacy.panel_label(ax, "b", "Held-out utility of decoder and source measurements")
    legacy.style_axis(ax, grid_axis="x")
    ax.axvline(0, color=MUTED, linewidth=1.0, linestyle=(0, (4, 2)))
    labels = [
        "Age-supervised decoder beyond\nage and background",
        "Direct source beyond\nlinear decoder",
        "Direct source beyond fixed\nflexible decoder calibration",
    ]
    y_values = [2.0, 1.0, 0.0]
    colors = [TEAL, BLUE, NAVY]
    for y, (_, row), color in zip(y_values, rows.iterrows(), colors):
        legacy.plot_interval(
            ax,
            y,
            row,
            color=color,
            marker="o",
            filled=True,
            estimate_digits=3,
            label_offset=0.00040,
        )
    ax.set_yticks(y_values, labels)
    ax.set_ylim(-0.55, 2.48)
    ax.set_xlim(-0.001, 0.0156)
    ax.set_xlabel(
        "Paired held-out log-loss improvement (right is better)",
        fontsize=5.9,
        color=INK,
    )
    ax.text(
        0.01,
        -0.24,
        "NHANES 2007–2012; intervals condition on stored OOF predictions.",
        transform=ax.transAxes,
        ha="left",
        fontsize=5.4,
        color=MUTED,
    )


def draw_matched_scalar(ax: plt.Axes, rows: pd.DataFrame) -> None:
    indexed = rows.set_index("contrast_id")
    legacy.panel_label(ax, "c", "Matched scalar capacity under two supervision targets")
    legacy.style_axis(ax, grid_axis="x")
    ax.axvline(0, color=MUTED, linewidth=1.0, linestyle=(0, (4, 2)))
    order = ["U_A", "Delta"]
    labels = [
        "Age-supervised scalar beyond\nage + background",
        "Mortality-supervised over\nage-supervised scalar",
    ]
    y_values = [1.0, 0.0]
    styles = [(TEAL, "o", False), (BLUE, "D", True)]
    for contrast_id, y, (color, marker, filled) in zip(order, y_values, styles):
        legacy.plot_interval(
            ax,
            y,
            indexed.loc[contrast_id],
            color=color,
            marker=marker,
            filled=filled,
        )
    ax.set_yticks(y_values, labels)
    ax.set_ylim(-0.67, 1.58)
    ax.set_xlim(-0.0012, 0.0114)
    ax.set_xlabel(
        "Paired held-out log-loss improvement (right is better)",
        fontsize=5.85,
        color=INK,
    )
    ax.text(
        0.01,
        0.89,
        "Same 25 inputs · one-dimensional linear capacity · equal Phase 2 calibration",
        transform=ax.transAxes,
        ha="left",
        fontsize=5.1,
        color=MUTED,
    )
    ax.text(
        0.98,
        0.04,
        "Utility can be genuine without construct identity",
        transform=ax.transAxes,
        ha="right",
        va="bottom",
        fontsize=5.15,
        color=RED,
        fontweight="bold",
        bbox={
            "boxstyle": "round,pad=0.22",
            "facecolor": RED_LIGHT,
            "edgecolor": RED,
            "linewidth": 0.75,
        },
    )
    ax.text(
        0.01,
        -0.24,
        "NHANES III Phase 1→2 frozen protocol transfer; Phase 2 did not refit either direction.",
        transform=ax.transAxes,
        ha="left",
        fontsize=5.15,
        color=MUTED,
    )


def draw_observed_age_context(ax: plt.Axes, rows: pd.DataFrame) -> None:
    indexed = rows.set_index("contrast_id")
    legacy.panel_label(ax, "d", "Observed-age value by measured information set")
    legacy.style_axis(ax, grid_axis="x")
    ax.axvline(0, color=MUTED, linewidth=1.0, linestyle=(0, (4, 2)))
    ax.axhspan(-0.34, 0.34, color=LIGHT, zorder=-1)
    order = ["V_F", "V_R", "P_proxy"]
    labels = [
        "Age after all 25\nclinical measurements",
        "Age after remaining 20\n(5-item block withheld)",
        "Extra predictive value of age\nafter block removal",
    ]
    y_values = [2.0, 1.0, 0.0]
    styles = [(TEAL, "o", False), (BLUE, "o", True), (NAVY, "D", True)]
    for contrast_id, y, (color, marker, filled) in zip(order, y_values, styles):
        legacy.plot_interval(
            ax,
            y,
            indexed.loc[contrast_id],
            color=color,
            marker=marker,
            filled=filled,
        )
    ax.axhline(0.5, color=GRID, linewidth=0.75)
    ax.set_yticks(y_values, labels)
    ax.set_ylim(-0.62, 2.55)
    ax.set_xlim(-0.0025, 0.0172)
    ax.set_xlabel(
        "Paired held-out log-loss improvement (right is better)",
        fontsize=5.85,
        color=INK,
    )
    ax.text(
        0.01,
        0.91,
        "NHANES III Phase 2 · $n=7{,}774$ · 392 deaths · 95% Fay-BRR CI",
        transform=ax.transAxes,
        ha="left",
        fontsize=5.35,
        color=MUTED,
    )
    ax.text(
        0.01,
        -0.25,
        "The remaining 20 are not asserted to be free of age-related information.",
        transform=ax.transAxes,
        ha="left",
        fontsize=5.25,
        color=MUTED,
    )


def build_figure(data: dict[str, Any], visual_qa_status: str) -> dict[str, Path]:
    ensure_package()
    write_input_manifest()

    fig = plt.figure(figsize=(7.15, 6.05), facecolor=WHITE)
    grid = fig.add_gridspec(
        2,
        2,
        left=0.085,
        right=0.985,
        bottom=0.085,
        top=0.96,
        hspace=0.48,
        wspace=0.47,
    )
    ax_a = fig.add_subplot(grid[0, 0])
    ax_b = fig.add_subplot(grid[0, 1])
    ax_c = fig.add_subplot(grid[1, 0])
    ax_d = fig.add_subplot(grid[1, 1])

    design_source = draw_design(ax_a, data["h04_diagnostics"])
    draw_continuous_contrasts(ax_b, data["h04_plot_rows"])
    draw_matched_scalar(ax_c, data["matched_rows"])
    draw_observed_age_context(ax_d, data["age_rows"])

    stem = "Figure_3_NHANES_endpoint_utility_information_set_and_supervision_target"
    figures = legacy.save_figure(fig, OUT, stem)

    design_path = OUT / "source_data" / "panel_a_continuous_nhanes_design.tsv"
    design_source.to_csv(design_path, sep="\t", index=False)
    continuous_path = OUT / "source_data" / "panel_b_continuous_nhanes_predictive_contrasts.tsv"
    data["h04_plot_rows"].to_csv(continuous_path, sep="\t", index=False)
    matched_path = OUT / "source_data" / "panel_c_nhanes3_matched_scalar_target_supervision.tsv"
    data["matched_rows"].to_csv(matched_path, sep="\t", index=False)
    age_path = OUT / "source_data" / "panel_d_nhanes3_observed_age_information_set.tsv"
    data["age_rows"].query("contrast_id in ['V_F', 'V_R', 'P_proxy']").to_csv(
        age_path,
        sep="\t",
        index=False,
    )
    source_index = OUT / "source_data" / "source_data_index.tsv"
    pd.DataFrame(
        [
            ("a", design_path.name, "frozen continuous-NHANES age-decoder/direct-source design"),
            ("b", continuous_path.name, "frozen HUM-04 decoder-utility and direct-source contrasts"),
            ("c", matched_path.name, "NHANES III U_A and q_Y-over-q_A matched-scalar contrasts"),
            ("d", age_path.name, "NHANES III V_F, V_R and P_proxy observed-age contrasts"),
        ],
        columns=["panel", "file", "content"],
    ).to_csv(source_index, sep="\t", index=False)

    legend = """# Figure 3 | Endpoint utility depends on information set and supervision target, not on the age unit alone

**a**, Frozen nested comparison in the complete-support continuous NHANES 2007–2012 interface. A fold-local ridge decoder was trained only to reconstruct chronological age from background and the same 50 measurements used by the direct endpoint model. Mortality models used age and background alone, added the out-of-fold age-supervised decoder, or directly modelled the same 50 source measurements. **b**, Paired held-out contrasts from that interface show the age-supervised decoder's predictive utility beyond age and background and the advantage of direct source-measurement modelling over the linear decoder and over one prespecified flexible calibration of that decoder. Intervals condition on stored out-of-fold predictions. **c**, Fixed NHANES III Phase 1-to-Phase 2 matched-scalar protocol transfer. The age-supervised and mortality-supervised one-dimensional directions used the same 25 measurements, background, linear capacity and Phase 1 fitting opportunity, then received equal Phase 2 calibration opportunity. U_A measures endpoint utility of the transported age-supervised scalar beyond observed age and background; Delta measures the additional advantage of the mortality-supervised direction over the age-supervised direction. **d**, Separate NHANES III Phase 2 comparison of the paired held-out mortality log-loss reduction from adding observed chronological age after all 25 measurements or after the frozen Phase 1 age-selected five-measurement block was withheld. Their difference quantifies the increase in age's predictive contribution after block removal. Points in c and d are design-weighted estimates and lines are 95% Fay-BRR intervals. The full-panel interval in d spans zero; this is not evidence of zero value or equivalence, and the remaining 20 measurements are not claimed to be free of age-related information. The NHANES III analyses are retrospective, outcome-accessed protocol-transfer robustness analyses rather than independent confirmation. These results do not identify biological age, surrogate validity or causal benefit.

# 图 3｜终点效用取决于信息集和监督目标，而非年龄单位本身

**a**，在具有完整五年随访支持的 continuous NHANES 2007–2012 接口中，保留冻结的嵌套比较。每个折内 ridge 解码器仅以时序年龄为监督目标，使用背景和与直接终点模型相同的 50 项测量；死亡模型依次使用年龄加背景、加入折外年龄监督解码器，或直接建模同源 50 项测量。**b**，该接口的配对留出对比显示年龄监督解码器相对年龄和背景的预测效用，以及直接源测量相对线性解码器和一次预设柔性校准解码器的优势；区间以已保存的折外预测为条件。**c**，冻结的 NHANES III Phase 1→Phase 2 匹配标量协议转移。年龄监督与死亡监督的一维方向使用相同 25 项测量、背景、线性容量和 Phase 1 拟合机会，并获得相同的 Phase 2 校准机会。U_A 衡量转移后的年龄监督标量在真实年龄和背景之外的终点预测效用；Delta 衡量死亡监督方向相对年龄监督方向的额外优势。**d**，在 NHANES III Phase 2 中，分别比较在全部 25 项测量后加入真实时序年龄，以及在冻结的 Phase 1 年龄选择五项 block 被移除后加入真实年龄所带来的配对留出死亡 log-loss 降低；二者之差衡量删块后年龄预测贡献的增加。c、d 中的点为设计加权估计，线为 95% Fay-BRR 区间。d 中完整面板的区间跨零不等于年龄价值为零或模型等价；remaining-20 也不被称为“非年龄相关”。NHANES III 分析是回顾性、结局已访问的固定协议转移稳健性分析，而非独立确认。上述结果不识别 biological age、surrogate validity 或 causal benefit。
"""
    (OUT / "FIGURE_LEGEND_BILINGUAL.md").write_text(legend, encoding="utf-8")
    (OUT / "README.md").write_text(
        "# NHANES Figure 3 successor v0.3\n\n"
        "Fit-free hierarchy revision of Figure 3. The panel order is now continuous "
        "NHANES design, continuous NHANES contrasts, NHANES III matched scalar "
        "target supervision, then observed-age information-set dependence. Only "
        "frozen TSV/JSON evidence is read; no fitted model is loaded, no estimator "
        "is fit or tuned, and no random number is generated. The v0.2 package is "
        "unchanged.\n",
        encoding="utf-8",
    )
    write_json(
        OUT / "CHART_CONTRACT.json",
        {
            "figure": "Figure 3 successor v0.3",
            "title": "Endpoint utility depends on information set and supervision target, not on the age unit alone",
            "question": "How do information set and supervision target affect held-out value for one named mortality endpoint?",
            "surface": "standalone 180-mm static publication figure",
            "renderer": "Matplotlib via pinned lhstate environment",
            "layout": "2 x 2; continuous NHANES in row 1 and NHANES III in row 2",
            "panels": {
                "a": "continuous NHANES age-supervised decoder/direct-source design",
                "b": "continuous NHANES decoder-utility and direct-source contrasts",
                "c": "NHANES III matched q_A/q_Y protocol-transfer U_A and Delta",
                "d": "NHANES III observed-age increment V_F/V_R and P_proxy",
            },
            "family": "task schematic plus uncertainty-and-benchmark dot-and-interval comparisons",
            "metric": "paired held-out mortality log-loss contrasts; positive direction defined per row",
            "palette": "hard two-root cap (blue and teal) plus neutrals; marker fill and shape provide non-colour distinction",
            "exports": ["600 dpi PNG", "editable SVG", "vector PDF"],
            "fit_free": True,
        },
    )
    audit_status = "PASS" if visual_qa_status == "pass" else "PENDING_MANUAL_VISUAL_QA"
    write_json(
        OUT / "VISUAL_AUDIT.json",
        {
            "status": audit_status,
            "automated_source_schema_checks": "PASS",
            "manual_visual_qa": visual_qa_status.upper(),
            "manual_visual_qa_checks": {
                "panel_order_matches_contract": visual_qa_status == "pass",
                "panel_titles_and_labels_legible": visual_qa_status == "pass",
                "intervals_and_zero_lines_visible": visual_qa_status == "pass",
                "no_clipping_collision_or_detached_annotation": visual_qa_status == "pass",
                "information_sets_and_supervision_targets_clear": visual_qa_status == "pass",
            },
            "fit_free": True,
            "fitted_models_loaded": False,
            "random_numbers_generated": False,
            "exports_created": sorted(figures),
        },
    )
    return figures


def parse_args() -> argparse.Namespace:
    parser = argparse.ArgumentParser(
        description="Build fit-free NHANES Figure 3 successor v0.3."
    )
    parser.add_argument(
        "--visual-qa-status",
        choices=["pending", "pass"],
        default="pending",
        help="Record manual visual QA as pending or pass after PNG inspection.",
    )
    return parser.parse_args()


def main() -> int:
    args = parse_args()
    data = legacy.load_and_gate()
    figures = build_figure(data, args.visual_qa_status)
    status = (
        "PASS_AUTOMATED_BUILD_AND_MANUAL_VISUAL_QA"
        if args.visual_qa_status == "pass"
        else "PASS_AUTOMATED_BUILD_VISUAL_QA_PENDING"
    )
    summary = {
        "status": status,
        "output": str(OUT),
        "figures": {key: str(path) for key, path in figures.items()},
        "panel_order": [
            "a continuous NHANES design",
            "b continuous NHANES contrasts",
            "c NHANES III capacity-matched supervision targets",
            "d NHANES III observed-age information-set dependence",
        ],
        "fit_free": True,
        "models_loaded": False,
        "random_numbers_generated": False,
        "visual_qa_status": args.visual_qa_status,
        "python": sys.version,
        "python_executable": sys.executable,
        "platform": platform.platform(),
    }
    write_json(OUT / "BUILD_SUMMARY.json", summary)
    write_output_manifest()
    print(json.dumps(summary, indent=2, ensure_ascii=False))
    return 0


if __name__ == "__main__":
    raise SystemExit(main())
```
