## Supplementary_Software_1 for "Inferential boundaries of age prediction: why prediction does not establish biological age measurement": plot_nhanes_individual_agegap_contributions_extended_data_v0_2.py

```
from __future__ import annotations

from pathlib import Path

import matplotlib.pyplot as plt
import numpy as np
import pandas as pd


PROJECT = Path(__file__).resolve().parents[1]
SOURCE = PROJECT / "outputs" / "nhanes_individual_agegap_contributions_v0_1"
OUT = PROJECT / "outputs" / "nhanes_individual_agegap_contributions_figure_v0_2"
FIGURES = OUT / "figures"
SOURCE_DATA = OUT / "source_data"


def panel_label(ax: plt.Axes, label: str) -> None:
    ax.text(
        -0.075,
        1.10,
        label,
        transform=ax.transAxes,
        fontsize=12,
        fontweight="bold",
        va="top",
        ha="left",
    )


def plot_pair(
    ax: plt.Axes,
    selected: pd.DataFrame,
    contributions: pd.DataFrame,
    case: str,
    title: str,
) -> None:
    row = selected.set_index("case").loc[case]
    subset = contributions.loc[contributions["case"].eq(case)].copy()
    subset["display_feature"] = subset["display_feature"].replace(
        {"Other measured features": "Net remainder of other measured features"}
    )
    feature_order = (
        subset.loc[~subset["is_remainder"].astype(bool)]
        .groupby("display_feature")["contribution_years"]
        .apply(lambda values: float(np.max(np.abs(values))))
        .sort_values(ascending=False)
        .index.tolist()
    )
    feature_order.append("Net remainder of other measured features")
    pivot = (
        subset.pivot(index="display_feature", columns="person", values="contribution_years")
        .reindex(feature_order)
        .fillna(0.0)
    )

    y = np.arange(len(pivot))
    ax.barh(
        y - 0.19,
        pivot[1],
        height=0.34,
        color="#2F6B8E",
        edgecolor="#2F6B8E",
        linewidth=0.8,
        label="Participant 1",
        zorder=3,
    )
    ax.barh(
        y + 0.19,
        pivot[2],
        height=0.34,
        color="white",
        edgecolor="#C96F32",
        linewidth=1.2,
        hatch="///",
        label="Participant 2",
        zorder=3,
    )
    ax.axvline(0, color="#5E666B", linewidth=0.9, zorder=2)
    ax.grid(axis="x", color="#D9DEE1", linewidth=0.55, alpha=0.75, zorder=1)
    ax.set_yticks(y, pivot.index, fontsize=8.5)
    ax.invert_yaxis()
    ax.set_xlim(-30, 30)
    ax.set_xlabel("Contribution to conditionally centred AgeGap (decoder-output years)")
    ax.set_title(title, loc="left", fontsize=10.5, fontweight="bold", pad=24)
    subtitle = (
        f"Both age {row['age_years']:.0f}; centred gaps {row['centered_agegap_1']:.2f} and "
        f"{row['centered_agegap_2']:.2f}; |Δ|={row['centered_agegap_abs_difference']:.3f}; "
        f"L2={row['contribution_l2_distance_years']:.2f} decoder-output years"
    )
    ax.text(0, 1.012, subtitle, transform=ax.transAxes, fontsize=8.5, ha="left", va="bottom")


def main() -> int:
    FIGURES.mkdir(parents=True, exist_ok=True)
    SOURCE_DATA.mkdir(parents=True, exist_ok=True)

    pairs = pd.read_csv(SOURCE / "matched_pairs.tsv.gz", sep="\t")
    summary = pd.read_csv(SOURCE / "matched_pair_distribution_summary.tsv", sep="\t")
    selected = pd.read_csv(SOURCE / "selected_display_pairs.tsv", sep="\t")
    contributions = pd.read_csv(SOURCE / "selected_pair_contributions.tsv", sep="\t")

    if len(pairs) != 293:
        raise RuntimeError(f"Expected 293 frozen pairs, found {len(pairs)}")
    if set(selected["case"]) != {"median_distance", "p95_distance"}:
        raise RuntimeError("Frozen display cases changed")

    l2_summary = summary.set_index("metric").loc["contribution_l2_distance_years"]
    median_l2 = float(l2_summary["median"])
    p95_l2 = float(l2_summary["q95"])

    plt.rcParams.update(
        {
            "font.family": "DejaVu Sans",
            "font.size": 9,
            "axes.spines.top": False,
            "axes.spines.right": False,
            "axes.titleweight": "bold",
            "pdf.fonttype": 42,
            "ps.fonttype": 42,
        }
    )
    fig, axes = plt.subplots(
        3,
        1,
        figsize=(10.8, 12.8),
        gridspec_kw={"height_ratios": [0.78, 1.38, 1.62]},
    )
    fig.subplots_adjust(left=0.31, right=0.985, top=0.965, bottom=0.07, hspace=0.58)

    ax = axes[0]
    values = pairs["contribution_l2_distance_years"].to_numpy(float)
    ax.hist(values, bins=38, color="#8CA0AC", edgecolor="white", linewidth=0.45)
    ax.axvline(median_l2, color="#2F6B8E", linewidth=1.8, label=f"All-pair median: {median_l2:.2f}")
    ax.axvline(
        p95_l2,
        color="#A64B4B",
        linewidth=1.8,
        linestyle="--",
        label=f"All-pair 95th percentile: {p95_l2:.2f}",
    )
    case_rows = selected.set_index("case")
    display_b_x = float(case_rows.loc["median_distance", "contribution_l2_distance_years"])
    display_c_x = float(case_rows.loc["p95_distance", "contribution_l2_distance_years"])
    ax.scatter(
        [display_b_x],
        [-0.7],
        marker="o",
        s=34,
        color="#2F6B8E",
        clip_on=False,
    )
    ax.scatter(
        [display_c_x],
        [-0.7],
        marker="D",
        s=30,
        color="#A64B4B",
        clip_on=False,
    )
    ax.text(display_b_x + 1.2, -0.7, "b", color="#2F6B8E", fontsize=8.5, fontweight="bold", va="center")
    ax.text(display_c_x + 1.2, -0.7, "c", color="#A64B4B", fontsize=8.5, fontweight="bold", va="center")
    ax.set_xlim(0, 90)
    ax.set_ylim(-1.6, None)
    ax.set_xlabel("Contribution-vector L2 distance (decoder-output years; not an age difference)")
    ax.set_ylabel("Matched pairs")
    ax.set_title(
        "Near-identical scalar gaps coexist with widely separated contribution vectors",
        loc="left",
        fontsize=10.5,
        pad=24,
    )
    ax.text(
        0,
        1.015,
        "293 outcome-blind pairs matched exactly on outer fold, age and background; |Δ conditionally centred AgeGap| ≤ 0.25 decoder years",
        transform=ax.transAxes,
        fontsize=8.5,
        ha="left",
        va="bottom",
    )
    ax.legend(frameon=False, fontsize=8.0, ncol=1, loc="upper right")
    panel_label(ax, "a")

    plot_pair(
        axes[1],
        selected,
        contributions,
        "median_distance",
        "Predeclared representative display pair",
    )
    panel_label(axes[1], "b")
    axes[1].legend(frameon=False, fontsize=8.2, ncol=2, loc="lower right")

    plot_pair(
        axes[2],
        selected,
        contributions,
        "p95_distance",
        "Predeclared high-distance display pair",
    )
    panel_label(axes[2], "c")

    note = (
        "Bars are exact signed contributions from the frozen 50-measurement linear age decoder and sum to each participant's conditionally centred gap.\n"
        "Display pairs were selected by the frozen median/95th-percentile rule within the 205 eligible same-sign pairs; panel-a references summarize all 293 matched pairs."
    )
    fig.text(0.31, 0.018, note, fontsize=7.7, ha="left", va="bottom")

    stem = "Extended_Data_Figure_4_near_identical_AgeGaps_distinct_measured_contributions"
    fig.savefig(FIGURES / f"{stem}.png", dpi=600, bbox_inches="tight")
    fig.savefig(FIGURES / f"{stem}.svg", bbox_inches="tight")
    fig.savefig(FIGURES / f"{stem}.pdf", bbox_inches="tight")
    plt.close(fig)

    pair_source = pairs[
        [
            "centered_agegap_abs_difference",
            "contribution_l2_distance_years",
            "contribution_cosine_similarity",
            "top5_absolute_contributor_jaccard",
        ]
    ].copy()
    pair_source.insert(0, "pair_index", np.arange(1, len(pair_source) + 1))
    pair_source.to_csv(SOURCE_DATA / "panel_a_matched_pair_profile_distances.tsv", sep="\t", index=False)

    summary.loc[
        summary["metric"].isin(
            [
                "centered_agegap_abs_difference",
                "contribution_l2_distance_years",
                "contribution_cosine_similarity",
                "top5_absolute_contributor_jaccard",
            ]
        )
    ].to_csv(SOURCE_DATA / "panel_a_distribution_summary.tsv", sep="\t", index=False)

    display_source = selected[
        [
            "case",
            "pair_id",
            "age_years",
            "centered_agegap_1",
            "centered_agegap_2",
            "centered_agegap_abs_difference",
            "contribution_l2_distance_years",
            "contribution_cosine_similarity",
            "top5_absolute_contributor_jaccard",
        ]
    ].rename(columns={"pair_id": "display_pair"})
    display_source.to_csv(SOURCE_DATA / "panel_b_c_display_pairs.tsv", sep="\t", index=False)

    contribution_source = contributions[
        ["case", "person", "feature_order", "display_feature", "contribution_years", "is_remainder"]
    ].copy()
    contribution_source["display_feature"] = contribution_source["display_feature"].replace(
        {"Other measured features": "Net remainder of other measured features"}
    )
    contribution_source.to_csv(SOURCE_DATA / "panel_b_c_contributions.tsv", sep="\t", index=False)
    return 0


if __name__ == "__main__":
    raise SystemExit(main())
```
