## Supplementary_Software_1 for "Inferential boundaries of age prediction: why prediction does not establish biological age measurement": plot_three_variable_age_state_task_map_figure_v0_2.py

```
"""Render the submission-facing X1-X3 age-coordinate/task-map figure.

This script is deliberately plotting-only. It consumes the frozen source-data
tables produced by three_variable_age_state_task_map_probe_v0_1 and never fits
or applies a model.
"""

from __future__ import annotations

import hashlib
import json
import shutil
from pathlib import Path

import matplotlib
import matplotlib.pyplot as plt
import pandas as pd


ROOT = Path(__file__).resolve().parents[1]
UPSTREAM = ROOT / "outputs" / "three_variable_age_state_task_map_probe_v0_1"
OUTPUT = ROOT / "outputs" / "three_variable_age_state_task_map_figure_v0_2"
FIGURES = OUTPUT / "figures"
SOURCE_DATA = OUTPUT / "source_data"

POINTS_NAME = "panels_a_c_age_coordinate_points.tsv.gz"
BINS_NAME = "panels_a_c_binned_means.tsv"
TASK_MAP_NAME = "panel_d_model_task_map.tsv"

INK = "#202933"
MUTED = "#66727D"
BLUE = "#2C6C99"
ORANGE = "#CB6724"
GRID = "#D9DEE3"


def sha256(path: Path) -> str:
    digest = hashlib.sha256()
    with path.open("rb") as handle:
        for chunk in iter(lambda: handle.read(1024 * 1024), b""):
            digest.update(chunk)
    return digest.hexdigest()


def copy_source_data() -> list[dict[str, object]]:
    SOURCE_DATA.mkdir(parents=True, exist_ok=True)
    records: list[dict[str, object]] = []
    for name in (POINTS_NAME, BINS_NAME, TASK_MAP_NAME):
        source = UPSTREAM / "source_data" / name
        target = SOURCE_DATA / name
        if not source.exists():
            raise FileNotFoundError(source)
        shutil.copy2(source, target)
        if sha256(source) != sha256(target):
            raise RuntimeError(f"Source-data copy parity failed: {name}")
        records.append(
            {
                "file": str(target.relative_to(ROOT)),
                "upstream_file": str(source.relative_to(ROOT)),
                "sha256": sha256(target),
                "bytes": target.stat().st_size,
            }
        )
    return records


def render(points: pd.DataFrame, binned: pd.DataFrame, task_map: pd.DataFrame) -> list[Path]:
    matplotlib.rcParams.update(
        {
            "font.family": "DejaVu Sans",
            "axes.edgecolor": INK,
            "axes.labelcolor": INK,
            "xtick.color": INK,
            "ytick.color": INK,
            "text.color": INK,
            "axes.titleweight": "bold",
        }
    )
    fig, axes = plt.subplots(2, 2, figsize=(12.4, 9.2), constrained_layout=False)
    plt.subplots_adjust(left=0.08, right=0.97, top=0.84, bottom=0.155, hspace=0.34, wspace=0.26)

    feature_titles = {
        "X1": "X1: noisy age-readable channel",
        "X2": "X2: age-orthogonal state channel",
        "X3": "X3: nonlinear age-state mixture",
    }
    state_style = {
        0: {"color": BLUE, "marker": "o", "linestyle": "-", "label": "Declared state 0"},
        1: {"color": ORANGE, "marker": "^", "linestyle": "--", "label": "Declared state 1"},
    }

    for panel, (axis, feature) in enumerate(zip(axes.flat[:3], ["X1", "X2", "X3"], strict=True)):
        feature_points = points.loc[points["feature"] == feature]
        feature_bins = binned.loc[binned["feature"] == feature]
        for state, style in state_style.items():
            subset = feature_points.loc[feature_points["separation_state"] == state]
            if state == 0:
                axis.scatter(
                    subset["age"], subset["value"], s=17, alpha=0.30,
                    marker=style["marker"], facecolors=style["color"], edgecolors="none",
                    rasterized=True,
                )
            else:
                axis.scatter(
                    subset["age"], subset["value"], s=20, alpha=0.42,
                    marker=style["marker"], facecolors="none", edgecolors=style["color"],
                    linewidths=0.65, rasterized=True,
                )
            means = feature_bins.loc[feature_bins["separation_state"] == state].sort_values("age_mean")
            axis.plot(
                means["age_mean"], means["value_mean"], color=style["color"],
                linestyle=style["linestyle"], marker=style["marker"], markersize=4.5,
                linewidth=2.0, markerfacecolor=style["color"] if state == 0 else "white",
                markeredgecolor=style["color"], label=style["label"],
            )
        axis.axhline(0, color=GRID, linewidth=0.9, zorder=0)
        axis.grid(axis="y", color=GRID, linewidth=0.7, alpha=0.65)
        axis.set_xlim(19, 81)
        axis.set_xlabel("Chronological age (years)")
        axis.set_ylabel("Standardized measurement")
        axis.set_title(f"{chr(97 + panel)}   {feature_titles[feature]}", loc="left", fontsize=12)
        axis.spines[["top", "right"]].set_visible(False)

    axis = axes.flat[3]
    marker_map = {"X1": "o", "X2": "s", "X3": "^", "X1-X3": "*"}
    color_map = {"X1": BLUE, "X2": ORANGE, "X3": MUTED, "X1-X3": INK}
    for _, row in task_map.iterrows():
        arm = str(row["arm"])
        axis.scatter(
            row["age_oof_r2"], row["state_oof_log_loss_gain_beyond_age"],
            s=145 if arm == "X1-X3" else 90, marker=marker_map[arm],
            facecolor=color_map[arm] if arm != "X2" else "white",
            edgecolor=color_map[arm], linewidth=1.6, zorder=3,
        )
        offset = {"X1": (7, -15), "X2": (7, 8), "X3": (7, 8), "X1-X3": (7, -16)}[arm]
        axis.annotate(
            arm,
            (row["age_oof_r2"], row["state_oof_log_loss_gain_beyond_age"]),
            xytext=offset, textcoords="offset points", fontsize=10, fontweight="bold",
        )
    axis.axhline(0, color=INK, linewidth=0.9)
    axis.axvline(0, color=INK, linewidth=0.9)
    x_values = task_map["age_oof_r2"].to_numpy(float)
    y_values = task_map["state_oof_log_loss_gain_beyond_age"].to_numpy(float)
    x_pad = max(0.05, 0.10 * (x_values.max() - x_values.min()))
    y_pad = max(0.004, 0.14 * (y_values.max() - y_values.min()))
    axis.set_xlim(min(-0.05, x_values.min() - x_pad), x_values.max() + x_pad)
    axis.set_ylim(min(-0.005, y_values.min() - y_pad), y_values.max() + y_pad)
    axis.grid(color=GRID, linewidth=0.7, alpha=0.65)
    axis.set_xlabel("Chronological-age decoding: pooled OOF R²")
    axis.set_ylabel("Incremental state decoding beyond age:\nOOF log-loss reduction")
    axis.set_title("d   Fixed-model task map", loc="left", fontsize=12)
    axis.spines[["top", "right"]].set_visible(False)

    handles, labels = axes.flat[0].get_legend_handles_labels()
    fig.legend(handles[:2], labels[:2], loc="upper center", ncol=2, frameon=False, bbox_to_anchor=(0.5, 0.89))
    fig.suptitle("Age-coordinate geometry and task-specific model performance", fontsize=18, fontweight="bold", y=0.965)
    fig.text(
        0.5, 0.918,
        "Fixed 500-row cross-sectional construction; age is an ordered coordinate, not within-person follow-up",
        ha="center", va="center", fontsize=10.5, color=MUTED,
    )
    fig.text(
        0.08, 0.072,
        "Points in a-c are different individuals observed once; connected symbols are cross-sectional fixed-age-bin state means.\n"
        "OOF = five-fold out-of-fold. State gain = age-only log loss - age+measurement log loss. Descriptive construction; no population interval.",
        ha="left", va="center", fontsize=8.2, color=MUTED, linespacing=1.35,
    )

    FIGURES.mkdir(parents=True, exist_ok=True)
    stem = FIGURES / "Figure_X123_age_coordinate_and_task_map_v0_2"
    paths = [stem.with_suffix(ext) for ext in (".png", ".svg", ".pdf")]
    fig.savefig(paths[0], dpi=600, facecolor="white")
    fig.savefig(paths[1], facecolor="white")
    fig.savefig(paths[2], facecolor="white")
    plt.close(fig)
    return paths


def write_text_artifacts(source_records: list[dict[str, object]], figure_paths: list[Path]) -> None:
    OUTPUT.mkdir(parents=True, exist_ok=True)
    manifest = source_records + [
        {
            "file": str(path.relative_to(ROOT)),
            "upstream_file": "",
            "sha256": sha256(path),
            "bytes": path.stat().st_size,
        }
        for path in figure_paths
    ]
    pd.DataFrame(manifest).to_csv(OUTPUT / "artifact_manifest.tsv", sep="\t", index=False)

    provenance = {
        "figure_version": "three_variable_age_state_task_map_figure_v0_2",
        "analysis_source": "three_variable_age_state_task_map_probe_v0_1",
        "plotting_only": True,
        "models_refit": False,
        "source_data_copied_with_sha256_parity": True,
        "interpretation_boundary": "Cross-sectional age coordinate, not within-person time or change.",
    }
    (OUTPUT / "provenance.json").write_text(json.dumps(provenance, indent=2) + "\n", encoding="utf-8")

    (OUTPUT / "README.md").write_text(
        "# X1-X3 age-coordinate and task-map figure v0.2\n\n"
        "Submission-facing plotting-only successor to the v0.1 probe figure. The underlying 500 rows, "
        "folds, predictions and model metrics are unchanged; no model was fitted or applied here. "
        "The footer was wrapped and moved inside the canvas after visual QA identified clipping.\n\n"
        "This display is cross-sectional. Its connected bin means are not within-person trajectories.\n",
        encoding="utf-8",
    )
    (OUTPUT / "FIGURE_LEGEND_BILINGUAL.md").write_text(
        "# Figure legend\n\n"
        "**English.** **Age-coordinate geometry and task-specific model performance.** "
        "a-c, Frozen cross-sectional measurements X1-X3 plotted against chronological age. Each point is a different individual observed once; "
        "connected symbols are state-specific means within fixed age bins and are not longitudinal trajectories. "
        "X1 is a noisy age-readable channel without a designed state term; X2 is the declared state channel after exact projection from the realized intercept and linear age direction; "
        "X3 mixes linear and nonlinear age structure with state. d, Fixed five-fold out-of-fold task map. "
        "The horizontal axis is pooled chronological-age decoding R². The vertical axis is the reduction in pooled state-decoding log loss obtained by adding the indicated measurement to the fixed age-only model. "
        "The display is explanatory: it distinguishes age readability, same-age state value and their mixture under the declared models; it is not population inference or longitudinal evidence.\n\n"
        "**中文。** **年龄坐标几何与任务特异的模型表现。** "
        "a-c，将冻结的横断面测量 X1-X3 对实际年龄作图。每个点代表一名仅观察一次的不同个体；连线是固定年龄分箱内按状态计算的均值，不是个体纵向轨迹。"
        "X1 是含噪声的年龄可读通道且生成式中没有设计状态项；X2 是从实际样本的截距和线性年龄方向精确投影后得到的声明状态通道；X3 同时包含线性及非线性年龄结构和状态。"
        "d，固定五折折外任务图。横轴为合并的实际年龄解码 R²；纵轴为在固定的仅年龄模型上加入相应测量后，合并状态解码 log loss 的下降。"
        "该图只解释在声明模型下年龄可读性、同龄状态价值及二者混合可以彼此分离；它不是总体推断或纵向证据。\n",
        encoding="utf-8",
    )


def main() -> None:
    source_records = copy_source_data()
    points = pd.read_csv(SOURCE_DATA / POINTS_NAME, sep="\t")
    binned = pd.read_csv(SOURCE_DATA / BINS_NAME, sep="\t")
    task_map = pd.read_csv(SOURCE_DATA / TASK_MAP_NAME, sep="\t")
    if len(points) != 1500 or set(points["feature"]) != {"X1", "X2", "X3"}:
        raise RuntimeError("Unexpected panels a-c source-data schema or row count")
    if set(task_map["arm"]) != {"X1", "X2", "X3", "X1-X3"}:
        raise RuntimeError("Unexpected panel d task-map arms")
    figure_paths = render(points, binned, task_map)
    write_text_artifacts(source_records, figure_paths)
    print(json.dumps({"figure_paths": [str(path.relative_to(ROOT)) for path in figure_paths], "models_refit": False}, indent=2))


if __name__ == "__main__":
    main()
```
