## Supplementary_Software_1 for "Inferential boundaries of age prediction: why prediction does not establish biological age measurement": plot_v6_2_submission_extended_data_v0_1.py

```
from __future__ import annotations

import argparse
import hashlib
import json
from pathlib import Path

import matplotlib

matplotlib.use("Agg")
import matplotlib.pyplot as plt
from matplotlib.gridspec import GridSpec
from matplotlib.patches import FancyArrowPatch, FancyBboxPatch
import numpy as np
import pandas as pd
from PIL import Image


ROOT = Path(__file__).resolve().parents[1]
OUTPUTS = ROOT / "outputs"
SOURCE_PACKAGE = OUTPUTS / "v6_2_extended_data_figure_sync_v0_1"
PACKAGE = OUTPUTS / "v6_2_submission_extended_data_v0_1"

BLUE = "#2F5D8C"
BLUE_LIGHT = "#DCE8F2"
ORANGE = "#C66B32"
ORANGE_LIGHT = "#F4E1D5"
INK = "#1E2933"
MID = "#65727E"
LIGHT = "#E7EBEF"
PALE = "#F7F8FA"
WHITE = "#FFFFFF"

SPECS = {
    1: {
        "title": "Residual and operator boundaries do not confer construct identity",
        "legend_en": (
            "a, Two logical limiting cases separate age residuals from same-age state information. A noisy age-only "
            "channel can produce a non-zero residual without any generated state, whereas exact observation of age "
            "together with a separate state channel yields a zero age gap while same-age heterogeneity remains observed. "
            "b, Feature-wise conditioning X−E(X|A,W) retains a multivariate same-age representation, whereas fitting an "
            "age decoder and subtracting age returns one decoder-specific scalar. Post-decoding gaps cannot recover "
            "directions discarded by scalarization, and neither operator confers biological-age or ageing-acceleration "
            "identity. These are analytical boundaries, not a second simulation result or a human effect estimate."
        ),
        "legend_zh": (
            "a，两个逻辑极限情形将年龄残差与同龄状态信息分开。带噪声的纯年龄通道可以在未生成任何状态时产生非零残差；而精确观测年龄并同时观测独立状态通道时，年龄 gap 为零，同龄异质性仍可被观测。"
            "b，逐特征条件化 X−E(X|A,W) 保留多变量同龄表示；先拟合年龄解码器再减去年龄只返回一个解码器特异标量。解码后的 gap 无法恢复标量化丢失的方向，两种算子都不赋予 biological-age 或 ageing-acceleration identity。"
            "这些是解析边界，不是第二个 simulation result 或人体效应估计。"
        ),
    },
    2: {
        "title": "HSCT referent-null sensitivities preserve the interpretation boundary",
        "legend_en": (
            "a, Holland affiliation estimates across donor–recipient age-difference thresholds, occasion summaries, "
            "follow-up selections and leave-one-batch-out analyses. b, Host-older and host-younger strata and "
            "observed-minus-null contrasts. c, Søraas clean-compartment and prespecified sensitivity populations after "
            "separate fixed-score processing. d, Processing, duplicated-occasion and fixed-imputation diagnostics. "
            "These analyses test robustness of the bounded referent result and do not identify lineage biological age "
            "or exclude host influence."
        ),
        "legend_zh": (
            "a，Holland 数据中跨供者–受者年龄差阈值、测量时点汇总、随访选择及逐批留出分析的归属估计。"
            "b，宿主年龄较大与较小分层，以及观察值减置换零分布均值的对比。"
            "c，Søraas 数据经独立固定分数处理后的清洁 compartment 主分析和预设敏感性人群。"
            "d，处理方式、重复时点和固定插补诊断。这些分析检验有界 referent 结果的稳健性，但不识别谱系 biological age，也不排除宿主影响。"
        ),
    },
    3: {
        "title": "Age-decoder and endpoint-model diagnostics for the NHANES comparison",
        "legend_en": (
            "a, Held-out chronological-age calibration and residual summaries for the fold-local ridge decoder. "
            "b, Fold-specific log loss for M_WA, M_CLOCK and M_X. c, Paired fold-specific clock and source-measurement "
            "gains for five-year all-cause mortality. Intervals condition on stored out-of-fold predictions and omit "
            "refitting and study-development uncertainty. Artifact reload parity and participant-loss checks are reported "
            "in Source Data and the resource-package README rather than as a scientific panel."
        ),
        "legend_zh": (
            "a，折内局部 ridge 年龄解码器的留出实际年龄校准与残差汇总。b，M_WA、M_CLOCK 和 M_X 的逐折 log loss。"
            "c，五年全因死亡中 clock 增益与源测量增益的逐折配对结果。区间条件于已存储折外预测，不包含重拟合或研究开发不确定性。"
            "模型产物重载一致性和参与者 loss 检查记录在 Source Data 与资源包 README 中，不作为科学 panel。"
        ),
    },
}


def sha256(path: Path) -> str:
    digest = hashlib.sha256()
    with path.open("rb") as handle:
        for chunk in iter(lambda: handle.read(1024 * 1024), b""):
            digest.update(chunk)
    return digest.hexdigest()


def read_tsv(path: Path) -> pd.DataFrame:
    return pd.read_csv(path, sep="\t")


def read_json(path: Path) -> dict:
    return json.loads(path.read_text(encoding="utf-8"))


def write_tsv(frame: pd.DataFrame, path: Path) -> None:
    path.parent.mkdir(parents=True, exist_ok=True)
    frame.to_csv(path, sep="\t", index=False, lineterminator="\n")


def configure() -> None:
    plt.rcParams.update(
        {
            "font.family": "DejaVu Sans",
            "font.size": 9,
            "axes.titlesize": 10,
            "axes.labelsize": 9,
            "axes.edgecolor": MID,
            "axes.linewidth": 0.8,
            "xtick.color": MID,
            "ytick.color": MID,
            "text.color": INK,
            "axes.labelcolor": INK,
            "figure.facecolor": WHITE,
            "axes.facecolor": WHITE,
            "savefig.facecolor": WHITE,
            "svg.fonttype": "none",
            "pdf.fonttype": 42,
        }
    )


def panel_label(ax, label: str) -> None:
    ax.text(-0.08, 1.06, label, transform=ax.transAxes, fontsize=14, fontweight="bold", va="top")


def clean(ax, grid="y") -> None:
    ax.spines["top"].set_visible(False)
    ax.spines["right"].set_visible(False)
    if grid:
        ax.grid(axis=grid, color=LIGHT, linewidth=0.7, zorder=0)


def box(ax, xy, width, height, text, *, face=PALE, edge=MID, weight="normal", fontsize=9):
    patch = FancyBboxPatch(
        xy,
        width,
        height,
        boxstyle="round,pad=0.02,rounding_size=0.025",
        transform=ax.transAxes,
        facecolor=face,
        edgecolor=edge,
        linewidth=1.0,
    )
    ax.add_patch(patch)
    ax.text(xy[0] + width / 2, xy[1] + height / 2, text, transform=ax.transAxes,
            ha="center", va="center", fontsize=fontsize, fontweight=weight)


def arrow(ax, start, end, color=MID):
    ax.add_patch(FancyArrowPatch(start, end, transform=ax.transAxes, arrowstyle="-|>", mutation_scale=12,
                                linewidth=1.2, color=color))


def add_header(fig, number: int) -> None:
    fig.suptitle(f"Extended Data Figure {number} | {SPECS[number]['title']}", x=0.055, y=0.985,
                 ha="left", fontsize=15, fontweight="bold")


def export(fig, package_dir: Path, filename: str) -> list[Path]:
    figures = package_dir / "figures"
    figures.mkdir(parents=True, exist_ok=True)
    paths = [figures / f"{filename}.png", figures / f"{filename}.svg", figures / f"{filename}.pdf"]
    fig.savefig(paths[0], dpi=600, bbox_inches="tight", pad_inches=0.12)
    fig.savefig(paths[1], bbox_inches="tight", pad_inches=0.12)
    fig.savefig(paths[2], bbox_inches="tight", pad_inches=0.12)
    plt.close(fig)
    return paths


def write_docs(number: int, package_dir: Path, input_paths: list[Path], source_index: pd.DataFrame, extra_readme: str = "") -> None:
    spec = SPECS[number]
    (package_dir / "FIGURE_LEGEND_EN.md").write_text(
        f"# Extended Data Figure {number} | {spec['title']}\n\n{spec['legend_en']}\n", encoding="utf-8"
    )
    (package_dir / "FIGURE_LEGEND_BILINGUAL.md").write_text(
        f"# Extended Data Figure {number} | {spec['title']}\n\n## English\n\n{spec['legend_en']}\n\n"
        f"## 中文\n\n{spec['legend_zh']}\n", encoding="utf-8"
    )
    manifest_rows = []
    for path in input_paths:
        manifest_rows.append({
            "role": "existing_frozen_or_audited_input",
            "path": str(path.relative_to(ROOT)).replace("/", "\\"),
            "bytes": path.stat().st_size,
            "sha256": sha256(path),
        })
    write_tsv(pd.DataFrame(manifest_rows), package_dir / "INPUT_MANIFEST.tsv")
    write_tsv(source_index, package_dir / "source_data" / "source_data_index.tsv")
    contract = {
        "figure": f"Extended Data Figure {number}",
        "question": spec["title"],
        "surface": "submission-facing static Extended Data",
        "renderer": "Matplotlib via pinned lhstate environment",
        "palette": "restrained blue/orange roots plus neutrals; non-colour marker/fill distinctions",
        "exports": ["600 dpi PNG", "editable SVG", "vector PDF"],
        "evidence": "deterministic fit-free redraw from frozen or previously audited source data",
    }
    (package_dir / "CHART_CONTRACT.json").write_text(json.dumps(contract, indent=2), encoding="utf-8")
    (package_dir / "README.md").write_text(
        f"# Extended Data Figure {number} submission resource\n\n"
        f"**Title:** {spec['title']}\n\n"
        "This package is a deterministic, fit-free submission redraw. It does not fit, tune, calibrate, bootstrap, "
        "permute, resample or generate analytical random numbers. The prior five-figure audit package remains unchanged.\n\n"
        "## Contents\n\n"
        "- `figures/`: 600 dpi PNG, editable SVG and vector PDF.\n"
        "- `source_data/`: panel Source Data, scientific/engineering checks where applicable, and an index.\n"
        "- English and bilingual legends, chart contract, input/output manifests and visual audit.\n\n"
        f"## Claim boundary\n\n{spec['legend_en']}\n\n{extra_readme}",
        encoding="utf-8",
    )


def build_ed1() -> None:
    number = 1
    package_dir = PACKAGE / "extended_data_figure_1"
    source_dir = package_dir / "source_data"
    source_dir.mkdir(parents=True, exist_ok=True)
    cases_path = SOURCE_PACKAGE / "extended_data_figure_1" / "source_data" / "panel_b_logical_limiting_cases.tsv"
    operators_path = SOURCE_PACKAGE / "extended_data_figure_1" / "source_data" / "panel_c_operator_boundary.tsv"
    cases = read_tsv(cases_path)
    operators = read_tsv(operators_path)
    out_cases = source_dir / "panel_a_logical_limiting_cases.tsv"
    out_operators = source_dir / "panel_b_operator_boundary.tsv"
    write_tsv(cases, out_cases)
    write_tsv(operators, out_operators)

    fig = plt.figure(figsize=(12.6, 6.5))
    gs = GridSpec(1, 2, figure=fig, wspace=0.25, left=0.07, right=0.97, bottom=0.14, top=0.86)
    axa = fig.add_subplot(gs[0, 0])
    axa.axis("off")
    panel_label(axa, "a")
    axa.set_title("Residual existence and state observability are logically distinct", loc="left")
    box(axa, (0.03, 0.58), 0.28, 0.20, "Noisy age-only\nchannel\nX=A+ε", face=BLUE_LIGHT, edge=BLUE, weight="bold")
    arrow(axa, (0.32, 0.68), (0.45, 0.68), BLUE)
    box(axa, (0.46, 0.56), 0.48, 0.24, "Age residual ≠ 0\nNo state was generated", face=WHITE, edge=BLUE)
    box(axa, (0.03, 0.20), 0.28, 0.20, "Exact age +\nseparate state\nX=(A,D)", face=ORANGE_LIGHT, edge=ORANGE, weight="bold")
    arrow(axa, (0.32, 0.30), (0.45, 0.30), ORANGE)
    box(axa, (0.46, 0.18), 0.48, 0.24, "Age gap = 0\nSame-age D remains observed", face=WHITE, edge=ORANGE)
    axa.text(0.49, 0.04, "Gap magnitude alone does not determine whether same-age state exists.",
             transform=axa.transAxes, ha="center", va="center", fontsize=8.5, color=MID)

    axb = fig.add_subplot(gs[0, 1])
    axb.axis("off")
    panel_label(axb, "b")
    axb.set_title("Feature-wise conditioning and a post-decoding gap are different operators", loc="left")
    box(axb, (0.04, 0.62), 0.22, 0.18, "Measurements\nX", face=PALE, edge=MID, weight="bold")
    arrow(axb, (0.27, 0.71), (0.41, 0.71), BLUE)
    box(axb, (0.42, 0.58), 0.52, 0.26, "Feature-wise conditioning\nX−E(X|A,W)\nmultivariate same-age representation",
        face=BLUE_LIGHT, edge=BLUE)
    arrow(axb, (0.15, 0.62), (0.15, 0.41), ORANGE)
    box(axb, (0.04, 0.19), 0.22, 0.18, "Age decoder\nq=f(X,A,W)", face=ORANGE_LIGHT, edge=ORANGE, weight="bold")
    arrow(axb, (0.27, 0.28), (0.41, 0.28), ORANGE)
    box(axb, (0.42, 0.15), 0.52, 0.26, "Post-decoding gap\nq−A\none decoder-specific scalar",
        face=WHITE, edge=ORANGE)
    axb.text(0.49, 0.04, "Scalarization can discard directions that gap construction cannot restore.",
             transform=axb.transAxes, ha="center", va="center", fontsize=8.5, color=MID)

    fig.text(0.50, 0.055, "Neither residualization outcome nor operator choice confers biological-age or ageing-acceleration identity.",
             ha="center", va="center", fontsize=9.2, color=INK,
             bbox=dict(boxstyle="round,pad=0.35", facecolor=PALE, edgecolor=LIGHT))
    add_header(fig, number)
    export(fig, package_dir, "Extended_Data_Figure_1_theory_boundaries")
    source_index = pd.DataFrame([
        {"panel": "a", "file": out_cases.name, "source": str(cases_path.relative_to(ROOT)), "role": "audited logical boundary"},
        {"panel": "b", "file": out_operators.name, "source": str(operators_path.relative_to(ROOT)), "role": "audited operator definitions"},
    ])
    write_docs(number, package_dir, [cases_path, operators_path], source_index,
               "No X1–X3 participant scatter or finite simulated sample is presented in this submission figure.\n")


def build_ed2() -> None:
    number = 2
    package_dir = PACKAGE / "extended_data_figure_2"
    source_dir = package_dir / "source_data"
    source_dir.mkdir(parents=True, exist_ok=True)
    old = SOURCE_PACKAGE / "extended_data_figure_3" / "source_data"
    input_paths = [
        old / "panel_a_holland_sensitivities.tsv",
        old / "panel_b_strata_and_null.tsv",
        old / "panel_c_soraas_populations.tsv",
        old / "panel_d_processing_diagnostics.tsv",
    ]
    panel_a, panel_b, panel_c, panel_d = [read_tsv(path) for path in input_paths]
    out_paths = [source_dir / path.name for path in input_paths]
    for frame, path in zip([panel_a, panel_b, panel_c, panel_d], out_paths):
        write_tsv(frame, path)

    fig = plt.figure(figsize=(13.5, 10.5))
    gs = GridSpec(3, 2, figure=fig, width_ratios=[1.35, 1.0], height_ratios=[1.05, 1.0, 0.85],
                  hspace=0.50, wspace=0.32, left=0.12, right=0.97, bottom=0.07, top=0.91)
    axa = fig.add_subplot(gs[:, 0])
    panel_label(axa, "a")
    y = np.arange(len(panel_a))[::-1]
    est, lo, hi = [panel_a[col].to_numpy() for col in ["mean_affiliation_index", "ci_low", "ci_high"]]
    colors = [ORANGE if name == "primary_gap_ge_10_all_occasions_pair_mean" else BLUE for name in panel_a["analysis"]]
    for yi, e, l, h, color in zip(y, est, lo, hi, colors):
        axa.errorbar(e, yi, xerr=[[e-l], [h-e]], fmt="o", color=color, ecolor=color, capsize=2.5, markersize=5.5)
    axa.axvline(0, color=MID, linewidth=1)
    axa.set_yticks(y, panel_a["display_label"])
    axa.set_xlabel("Mean bounded affiliation index (lineage side positive)")
    axa.set_title("Holland: prespecified thresholds, occasion summaries and batch leave-outs", loc="left")
    clean(axa, "x")

    axb = fig.add_subplot(gs[0, 1])
    panel_label(axb, "b")
    axb.axis("off")
    axb.set_title("Sign strata and observed-minus-null contrasts", loc="left")
    strata = panel_b[panel_b["metric"] == "affiliation_index"].reset_index(drop=True)
    contrasts = panel_b[panel_b["metric"] == "observed_minus_null"].reset_index(drop=True)
    for i, row in strata.iterrows():
        axb.text(0.02, 0.72-i*0.18, row["display_label"], transform=axb.transAxes, ha="left", va="center", fontsize=8.5)
        axb.text(0.97, 0.72-i*0.18, f"{row['estimate']:.3f} [{row['q025']:.3f}, {row['q975']:.3f}]",
                 transform=axb.transAxes, ha="right", va="center", fontsize=8.5, color=BLUE, fontweight="bold")
    axb.plot([0.02, 0.97], [0.42, 0.42], transform=axb.transAxes, color=LIGHT, linewidth=1)
    for i, row in contrasts.iterrows():
        axb.text(0.02, 0.31-i*0.17, f"{row['display_label']} observed−null", transform=axb.transAxes, ha="left", va="center", fontsize=8.5)
        axb.text(0.97, 0.31-i*0.17, f"{row['estimate']:.3f} [{row['q025']:.3f}, {row['q975']:.3f}]",
                 transform=axb.transAxes, ha="right", va="center", fontsize=8.5, color=ORANGE, fontweight="bold")

    axc = fig.add_subplot(gs[1, 1])
    panel_label(axc, "c")
    y = np.arange(len(panel_c))[::-1]
    est, lo, hi = [panel_c[col].to_numpy() for col in ["mean_d", "mean_d_ci_low", "mean_d_ci_high"]]
    axc.errorbar(est, y, xerr=np.vstack([est-lo, hi-est]), fmt="o", color=ORANGE, ecolor=ORANGE, capsize=3, markersize=6)
    axc.axvline(0, color=MID, linewidth=1)
    axc.set_yticks(y, panel_c["display_label"])
    axc.set_xlabel("Søraas distance contrast d (years; donor side negative)")
    axc.set_title("Søraas: separately processed fixed-score populations", loc="left")
    clean(axc, "x")

    axd = fig.add_subplot(gs[2, 1])
    axd.axis("off")
    panel_label(axd, "d")
    axd.set_title("Processing and duplication diagnostics", loc="left")
    for i, row in panel_d.iterrows():
        ytxt = 0.78-i*0.20
        color = BLUE if i < 2 else ORANGE
        marker = "o" if i < 2 else "s"
        axd.scatter([0.05], [ytxt], transform=axd.transAxes, s=55, c=color, marker=marker)
        if i < 2:
            detail = f"{row['diagnostic']}: I={row['value']:.3f}; {row['detail']}"
        elif i == 2:
            detail = f"{row['diagnostic']}: {row['value']:.2f} y; {row['detail']}"
        else:
            detail = f"{row['diagnostic']}: {row['detail']}"
        axd.text(0.10, ytxt, detail, transform=axd.transAxes, ha="left", va="center", fontsize=8.5)
    axd.text(0.02, -0.02, "Robustness of a bounded referent result; not lineage biological-age identification or host exclusion.",
             transform=axd.transAxes, ha="left", va="bottom", fontsize=8.2, color=MID)

    add_header(fig, number)
    export(fig, package_dir, "Extended_Data_Figure_2_hsct_referent_sensitivities")
    source_index = pd.DataFrame([
        {"panel": panel, "file": output.name, "source": str(source.relative_to(ROOT)), "role": role}
        for panel, output, source, role in zip(
            ["a", "b", "c", "d"], out_paths, input_paths,
            ["Holland sensitivity summaries", "sign strata and frozen referent-null contrasts",
             "separately processed Søraas populations", "processing and duplication diagnostics"]
        )
    ])
    write_docs(number, package_dir, input_paths, source_index)


def build_ed3() -> None:
    number = 3
    package_dir = PACKAGE / "extended_data_figure_3"
    source_dir = package_dir / "source_data"
    source_dir.mkdir(parents=True, exist_ok=True)
    base = OUTPUTS / "nhanes_5y_age_decoder_source_information_probe_v0_2"
    age_path = base / "age_band_diagnostics.tsv"
    diagnostics_path = base / "age_decoder_diagnostics.json"
    fold_path = base / "fold_metrics.tsv"
    contrast_path = base / "paired_contrasts.tsv"
    audit_path = base / "model_artifact_audit.json"
    loss_path = base / "participant_loss_vectors.tsv.gz"
    pooled_path = base / "pooled_metrics.tsv"
    age = read_tsv(age_path)
    diagnostics = read_json(diagnostics_path)
    folds = read_tsv(fold_path)
    contrasts = read_tsv(contrast_path)
    artifact_audit = read_json(audit_path)
    losses = read_tsv(loss_path)
    pooled = read_tsv(pooled_path)

    panel_a_path = source_dir / "panel_a_age_calibration.tsv"
    panel_b_path = source_dir / "panel_b_fold_performance.tsv"
    panel_c_path = source_dir / "panel_c_fold_gains.tsv"
    engineering_path = source_dir / "engineering_qa_not_plotted.tsv"
    write_tsv(age, panel_a_path)
    write_tsv(folds[["outer_fold", "n", "events", "M_WA_log_loss", "M_CLOCK_log_loss", "M_X_log_loss"]], panel_b_path)
    write_tsv(folds[["outer_fold", "V_CLOCK", "V_SOURCE", "V_X"]], panel_c_path)
    expected = pooled.set_index("model")["log_loss"].to_dict()
    participant_parity = max(abs(float(losses[model].mean()) - float(expected[model])) for model in ["M_WA", "M_CLOCK", "M_X"])
    engineering = pd.DataFrame([
        {"check": "participant-loss pooled parity", "status": "PASS" if participant_parity < 1e-12 else "FAIL", "value": participant_parity,
         "detail": "maximum absolute difference between stored participant-loss mean and pooled log loss"},
        {"check": "artifact count", "status": artifact_audit["status"], "value": artifact_audit["artifact_count"],
         "detail": f"expected {artifact_audit['expected_artifact_count']}; reused {artifact_audit['reused_artifact_count']}; newly fitted {artifact_audit['newly_fitted_artifact_count']}"},
        {"check": "age-prediction reload parity", "status": artifact_audit["status"], "value": artifact_audit["max_age_prediction_error"],
         "detail": f"tolerance {artifact_audit['parity_tolerance']}"},
        {"check": "endpoint-prediction reload parity", "status": artifact_audit["status"], "value": artifact_audit["max_outcome_prediction_error"],
         "detail": f"tolerance {artifact_audit['parity_tolerance']}"},
    ])
    write_tsv(engineering, engineering_path)

    fig = plt.figure(figsize=(14.2, 5.4))
    gs = GridSpec(1, 3, figure=fig, wspace=0.30, left=0.065, right=0.98, bottom=0.17, top=0.83)
    axa = fig.add_subplot(gs[0, 0])
    panel_label(axa, "a")
    axa.plot([20, 80], [20, 80], color=MID, linestyle="--", linewidth=1.2, label="Identity")
    axa.plot(age["mean_age"], age["mean_clock"], color=BLUE, marker="o", markersize=6, linewidth=1.6, label="Age-band mean")
    for _, row in age.iterrows():
        axa.text(row["mean_age"], row["mean_clock"]+1.5, row["age_band"], ha="center", fontsize=7.2, color=MID)
    axa.set_xlim(19, 80); axa.set_ylim(19, 80)
    axa.set_xlabel("Mean chronological age (years)")
    axa.set_ylabel("Mean held-out decoder score (years)")
    axa.set_title("Held-out chronological-age calibration", loc="left")
    axa.text(0.03, 0.95, f"R²={diagnostics['r2']:.3f}; RMSE={diagnostics['rmse']:.2f} y\nresidual mean={diagnostics['residual_mean']:.3f} y; SD={diagnostics['residual_sd']:.2f} y",
             transform=axa.transAxes, ha="left", va="top", fontsize=8,
             bbox=dict(boxstyle="round,pad=0.3", facecolor=WHITE, edgecolor=LIGHT))
    axa.legend(frameon=False, fontsize=7.5, loc="lower right")
    clean(axa, "both")

    axb = fig.add_subplot(gs[0, 1])
    panel_label(axb, "b")
    for column, label, color, marker in [
        ("M_WA_log_loss", "M_WA", MID, "o"),
        ("M_CLOCK_log_loss", "M_CLOCK", ORANGE, "s"),
        ("M_X_log_loss", "M_X", BLUE, "^"),
    ]:
        axb.plot(folds["outer_fold"]+1, folds[column], label=label, color=color, marker=marker, linewidth=1.5, markersize=6)
    axb.set_xticks(np.arange(1, 6))
    axb.set_xlabel("Outer fold")
    axb.set_ylabel("Held-out log loss (lower is better)")
    axb.set_title("Fold-specific mortality performance", loc="left")
    axb.legend(frameon=False, fontsize=7.5, ncol=3, loc="upper center")
    clean(axb)

    axc = fig.add_subplot(gs[0, 2])
    panel_label(axc, "c")
    x = folds["outer_fold"].to_numpy()+1
    axc.plot(x, folds["V_CLOCK"], color=ORANGE, marker="s", linewidth=1.5, label="Clock gain")
    axc.plot(x, folds["V_SOURCE"], color=BLUE, marker="o", linewidth=1.5, label="Source gain")
    axc.axhline(0, color=MID, linewidth=0.9)
    for _, row in contrasts[contrasts["contrast"].isin(["V_CLOCK", "V_SOURCE"])].iterrows():
        color = ORANGE if row["contrast"] == "V_CLOCK" else BLUE
        label = "pooled clock" if row["contrast"] == "V_CLOCK" else "pooled source"
        axc.axhspan(row["q025"], row["q975"], color=color, alpha=0.08)
        axc.axhline(row["estimate"], color=color, linestyle="--", linewidth=1, label=label)
    axc.set_xticks(x)
    axc.set_xlabel("Outer fold")
    axc.set_ylabel("Paired held-out log-loss gain")
    axc.set_title("Clock utility and source value coexist", loc="left")
    axc.legend(frameon=False, fontsize=7.2, loc="upper right")
    clean(axc)

    add_header(fig, number)
    export(fig, package_dir, "Extended_Data_Figure_3_nhanes_decoder_diagnostics")
    source_index = pd.DataFrame([
        {"panel": "a", "file": panel_a_path.name, "source": f"{age_path.relative_to(ROOT)}; {diagnostics_path.relative_to(ROOT)}", "role": "scientific panel: frozen held-out age diagnostics"},
        {"panel": "b", "file": panel_b_path.name, "source": str(fold_path.relative_to(ROOT)), "role": "scientific panel: frozen fold endpoint performance"},
        {"panel": "c", "file": panel_c_path.name, "source": f"{fold_path.relative_to(ROOT)}; {contrast_path.relative_to(ROOT)}", "role": "scientific panel: frozen paired gains"},
        {"panel": "not plotted", "file": engineering_path.name, "source": f"{audit_path.relative_to(ROOT)}; {loss_path.relative_to(ROOT)}; {pooled_path.relative_to(ROOT)}", "role": "engineering QA retained outside scientific panels"},
    ])
    extra = (
        "## Engineering QA retained outside the scientific panels\n\n"
        f"- Participant-loss pooled parity: {'PASS' if participant_parity < 1e-12 else 'FAIL'}; maximum absolute difference {participant_parity:.3g}.\n"
        f"- Persisted model artifacts: {artifact_audit['artifact_count']}/{artifact_audit['expected_artifact_count']}; reused {artifact_audit['reused_artifact_count']}; newly fitted {artifact_audit['newly_fitted_artifact_count']}.\n"
        f"- Reload parity: maximum age-prediction error {artifact_audit['max_age_prediction_error']}; maximum endpoint-probability error {artifact_audit['max_outcome_prediction_error']}.\n"
        "- These checks support reproducibility but are not presented as a scientific result panel.\n"
    )
    write_docs(number, package_dir, [age_path, diagnostics_path, fold_path, contrast_path, audit_path, loss_path, pooled_path], source_index, extra)


def validate_science_contract() -> None:
    ed1a = read_tsv(PACKAGE / "extended_data_figure_1" / "source_data" / "panel_a_logical_limiting_cases.tsv")
    ed1b = read_tsv(PACKAGE / "extended_data_figure_1" / "source_data" / "panel_b_operator_boundary.tsv")
    if len(ed1a) != 2 or len(ed1b) != 2:
        raise RuntimeError("ED1 theory-boundary source contract failed")
    ed2a = read_tsv(PACKAGE / "extended_data_figure_2" / "source_data" / "panel_a_holland_sensitivities.tsv")
    primary = ed2a.loc[ed2a["analysis"] == "primary_gap_ge_10_all_occasions_pair_mean", "mean_affiliation_index"].iloc[0]
    if not np.isclose(primary, 0.5349228774016916):
        raise RuntimeError("ED2 Holland primary estimate mismatch")
    ed3c = read_tsv(PACKAGE / "extended_data_figure_3" / "source_data" / "panel_c_fold_gains.tsv")
    if len(ed3c) != 5 or not (ed3c["V_SOURCE"] > 0).all():
        raise RuntimeError("ED3 source-gain contract failed")
    qa = read_tsv(PACKAGE / "extended_data_figure_3" / "source_data" / "engineering_qa_not_plotted.tsv")
    if not (qa["status"] == "PASS").all():
        raise RuntimeError("ED3 engineering QA failed")


def write_audits_and_manifests(manual_pass: bool) -> None:
    index_rows = []
    for number in SPECS:
        package_dir = PACKAGE / f"extended_data_figure_{number}"
        pngs = list((package_dir / "figures").glob("*.png"))
        svgs = list((package_dir / "figures").glob("*.svg"))
        pdfs = list((package_dir / "figures").glob("*.pdf"))
        if len(pngs) != 1 or len(svgs) != 1 or len(pdfs) != 1:
            raise RuntimeError(f"ED{number}: expected one PNG, SVG and PDF")
        with Image.open(pngs[0]) as image:
            width, height = image.size
        programmatic = width >= 6000 and height >= 3000 and svgs[0].stat().st_size > 1000 and pdfs[0].stat().st_size > 1000
        overall = "PASS" if programmatic and manual_pass else ("PENDING_MANUAL" if programmatic else "FAIL")
        audit = {
            "figure": number,
            "programmatic_status": "PASS" if programmatic else "FAIL",
            "manual_visual_status": "PASS" if manual_pass else "PENDING",
            "overall_status": overall,
            "png_pixels": [width, height],
            "required_exports_present": True,
            "fit_free": True,
            "manual_review_scope": "panel labels, legibility, clipping, overlap, scale honesty, theme dilution and boundary wording",
        }
        (package_dir / "VISUAL_AUDIT.json").write_text(json.dumps(audit, indent=2), encoding="utf-8")
        rows = []
        for path in sorted(package_dir.rglob("*")):
            if path.is_file() and path.name != "OUTPUT_MANIFEST.tsv":
                rows.append({"path": str(path.relative_to(package_dir)).replace("/", "\\"),
                             "bytes": path.stat().st_size, "sha256": sha256(path)})
        write_tsv(pd.DataFrame(rows), package_dir / "OUTPUT_MANIFEST.tsv")
        index_rows.append({
            "figure": number,
            "title": SPECS[number]["title"],
            "package": str(package_dir.relative_to(ROOT)).replace("/", "\\"),
            "png": str(pngs[0].relative_to(ROOT)).replace("/", "\\"),
            "svg": str(svgs[0].relative_to(ROOT)).replace("/", "\\"),
            "pdf": str(pdfs[0].relative_to(ROOT)).replace("/", "\\"),
            "audit_status": overall,
        })
    write_tsv(pd.DataFrame(index_rows), PACKAGE / "FIGURE_INDEX.tsv")
    lines = ["# v6.2 submission Extended Data v0.1", "",
             "Theme-diluted submission set: three deterministic fit-free figures. The prior five-figure audit package is preserved unchanged.", "",
             "| Final figure | Responsibility | Audit |", "|---|---|---|"]
    for row in index_rows:
        lines.append(f"| ED{row['figure']} | {row['title']} | {row['audit_status']} |")
    lines.extend(["", "Unified script: `analysis/plot_v6_2_submission_extended_data_v0_1.py`.", ""])
    (PACKAGE / "FIGURE_INDEX.md").write_text("\n".join(lines), encoding="utf-8")
    qa_status = "PASS" if manual_pass and all(row["audit_status"] == "PASS" for row in index_rows) else "PENDING_MANUAL"
    (PACKAGE / "QA_REPORT.md").write_text(
        "# v6.2 submission Extended Data v0.1 — QA\n\n"
        f"**Overall status: {qa_status}.**\n\n"
        "- Pinned `lhstate` environment and project Python wrapper used.\n"
        "- Three figures exported as 600 dpi PNG, editable SVG and vector PDF.\n"
        "- Scientific source contracts and ED3 engineering QA: PASS.\n"
        "- No fitting, tuning, resampling, bootstrap/permutation execution or analytical RNG.\n"
        "- Final ED1 contains no n=500 scatter or simulation-result presentation.\n"
        "- Final ED2 retains the audited HSCT scientific content.\n"
        "- Final ED3 contains three scientific panels; reload parity and participant-loss checks are Source Data/README only.\n"
        "- Prior `v6_2_extended_data_figure_sync_v0_1` package was not overwritten or deleted.\n"
        f"- Manual visual inspection: {'PASS for ED1–ED3' if manual_pass else 'pending'}.\n",
        encoding="utf-8",
    )
    root_rows = []
    for path in sorted(PACKAGE.rglob("*")):
        if path.is_file() and path.name != "PACKAGE_MANIFEST.tsv":
            root_rows.append({"path": str(path.relative_to(PACKAGE)).replace("/", "\\"),
                              "bytes": path.stat().st_size, "sha256": sha256(path)})
    write_tsv(pd.DataFrame(root_rows), PACKAGE / "PACKAGE_MANIFEST.tsv")


def main() -> None:
    parser = argparse.ArgumentParser()
    parser.add_argument("--audit-only", action="store_true")
    parser.add_argument("--manual-pass", action="store_true")
    args = parser.parse_args()
    configure()
    PACKAGE.mkdir(parents=True, exist_ok=True)
    if not args.audit_only:
        build_ed1()
        build_ed2()
        build_ed3()
    validate_science_contract()
    write_audits_and_manifests(args.manual_pass)
    print(f"{'PASS' if args.manual_pass else 'PENDING_MANUAL'}: v6.2 submission Extended Data at {PACKAGE}")


if __name__ == "__main__":
    main()
```
