## Supplementary_Software_1 for "Inferential boundaries of age prediction: why prediction does not establish biological age measurement": plot_v6_8_figure4_target_first_and_ed_acquisition_v0_1.py

```
#!/usr/bin/env python
"""Build the v6.8 target-first Figure 4 and bounded acquisition ED figure.

This entry point is deliberately fit-free. It reads only the source-data tables
already frozen in ``v6_6_figure4_inverse_decoder_v0_1``, verifies those tables
against that package's figure manifest, and performs deterministic rendering.
It does not load or fit models, regenerate resampling indices, or mutate any
predecessor output.
"""

from __future__ import annotations

import argparse
import hashlib
import json
import platform
import sys
from pathlib import Path

import matplotlib
import matplotlib.pyplot as plt
from matplotlib.patches import FancyArrowPatch, FancyBboxPatch
import numpy as np
import pandas as pd


ROOT = Path(__file__).resolve().parents[1]
SCRIPT = Path(__file__).resolve()
SOURCE = ROOT / "outputs" / "v6_6_figure4_inverse_decoder_v0_1"
MAIN_PACKAGE = ROOT / "outputs" / "v6_8_figure4_target_first_framework_v0_1"
ED_PACKAGE = ROOT / "outputs" / "v6_8_extended_data_figure4_bounded_acquisition_v0_1"
MAIN_STEM = "Figure_4_target_first_estimand_specific_replacement_framework"
ED_STEM = "Extended_Data_Figure_4_bounded_equal_category_cost_acquisition_comparison"
VERSION = "v6_8_figure4_target_first_and_ed_acquisition_v0_1"

INK = "#202124"
NAVY = "#18243D"
MUTED = "#62676D"
GRID = "#D9DDE1"
LIGHT = "#EEF1F3"
WHITE = "#FFFFFF"
BLUE = "#315F85"
BLUE_LIGHT = "#E5EEF5"
TEAL = "#087E8B"
TEAL_LIGHT = "#D7EEF0"
ORANGE = "#D97732"
ORANGE_LIGHT = "#F7E6D8"
GOLD = "#C08A18"
GOLD_LIGHT = "#F8EED3"
PURPLE = "#6D5396"
PURPLE_LIGHT = "#EEEAF7"
RED = "#A83F3B"

matplotlib.rcParams.update(
    {
        "font.family": "sans-serif",
        "font.sans-serif": ["Arial", "DejaVu Sans"],
        "font.size": 7.2,
        "axes.unicode_minus": False,
        "svg.fonttype": "none",
        "pdf.fonttype": 42,
        "ps.fonttype": 42,
    }
)


def require(condition: bool, message: str) -> None:
    if not condition:
        raise RuntimeError(message)


def sha256(path: Path) -> str:
    digest = hashlib.sha256()
    with path.open("rb") as handle:
        for block in iter(lambda: handle.read(1024 * 1024), b""):
            digest.update(block)
    return digest.hexdigest()


def write_json(path: Path, payload: object) -> None:
    path.write_text(json.dumps(payload, indent=2, ensure_ascii=False) + "\n", encoding="utf-8")


def ensure_packages() -> None:
    for package in (MAIN_PACKAGE, ED_PACKAGE):
        (package / "figures").mkdir(parents=True, exist_ok=True)
        (package / "source_data").mkdir(parents=True, exist_ok=True)
        (package / "visualization").mkdir(parents=True, exist_ok=True)


def box(
    ax: plt.Axes,
    xy: tuple[float, float],
    width: float,
    height: float,
    text: str,
    *,
    face: str = WHITE,
    edge: str = MUTED,
    fontsize: float = 6.5,
    weight: str = "normal",
    color: str = INK,
    linewidth: float = 1.0,
    linestyle: str = "solid",
) -> FancyBboxPatch:
    patch = FancyBboxPatch(
        xy,
        width,
        height,
        boxstyle="round,pad=0.014,rounding_size=0.015",
        transform=ax.transAxes,
        facecolor=face,
        edgecolor=edge,
        linewidth=linewidth,
        linestyle=linestyle,
        clip_on=False,
    )
    ax.add_patch(patch)
    ax.text(
        xy[0] + width / 2,
        xy[1] + height / 2,
        text,
        transform=ax.transAxes,
        ha="center",
        va="center",
        fontsize=fontsize,
        fontweight=weight,
        color=color,
        linespacing=1.12,
    )
    return patch


def arrow(
    ax: plt.Axes,
    start: tuple[float, float],
    end: tuple[float, float],
    *,
    color: str = MUTED,
    linewidth: float = 1.1,
    linestyle: str = "solid",
) -> None:
    ax.add_patch(
        FancyArrowPatch(
            start,
            end,
            transform=ax.transAxes,
            arrowstyle="-|>",
            mutation_scale=8.5,
            color=color,
            linewidth=linewidth,
            linestyle=linestyle,
            shrinkA=0,
            shrinkB=0,
            clip_on=False,
        )
    )


def panel_label(ax: plt.Axes, label: str, title: str) -> None:
    ax.text(-0.018, 1.045, label, transform=ax.transAxes, fontsize=10, fontweight="bold", color=INK, va="bottom")
    ax.text(0.055, 1.045, title, transform=ax.transAxes, fontsize=8.25, fontweight="bold", color=INK, va="bottom")


def style_axis(ax: plt.Axes, *, grid_axis: str = "both") -> None:
    ax.spines["top"].set_visible(False)
    ax.spines["right"].set_visible(False)
    ax.spines["left"].set_color(MUTED)
    ax.spines["bottom"].set_color(MUTED)
    ax.tick_params(colors=INK, labelsize=6.3, width=0.7, length=3)
    ax.grid(axis=grid_axis, color=GRID, linewidth=0.55, alpha=0.78)
    ax.set_axisbelow(True)


def load_frozen_source() -> dict[str, object]:
    paths = {
        "manifest": SOURCE / "visualization" / "figure_manifest.tsv",
        "policy": SOURCE / "source_data" / "panel_a_frozen_policy_definitions.tsv",
        "target": SOURCE / "source_data" / "panel_b_target_policy_intervals.tsv",
        "pair": SOURCE / "source_data" / "panel_c_pooled_policy_difference.tsv",
        "folds": SOURCE / "source_data" / "panel_c_fold_policy_differences.tsv",
        "replacement": SOURCE / "source_data" / "panel_d_estimand_replacement_map.tsv",
    }
    for label, path in paths.items():
        require(path.exists(), f"Required frozen {label} input is missing: {path}")

    manifest = pd.read_csv(paths["manifest"], sep="\t")
    manifest_by_path = manifest.set_index("path")
    for label, path in paths.items():
        if label == "manifest":
            continue
        relative = path.relative_to(ROOT).as_posix()
        require(relative in manifest_by_path.index, f"Frozen manifest does not list {relative}")
        require(sha256(path) == manifest_by_path.loc[relative, "sha256"], f"Frozen source-data hash changed: {relative}")

    policy = pd.read_csv(paths["policy"], sep="\t")
    target = pd.read_csv(paths["target"], sep="\t")
    pair = pd.read_csv(paths["pair"], sep="\t")
    folds = pd.read_csv(paths["folds"], sep="\t")
    replacement = pd.read_csv(paths["replacement"], sep="\t")

    require(len(policy) == 2, "Expected two frozen acquisition policies.")
    require(len(target) == 2, "Expected two target-cycle policy estimates.")
    require(len(pair) == 1, "Expected one pooled paired policy contrast.")
    require(len(folds) == 5, "Expected five frozen target folds.")
    require(len(replacement) == 5, "Expected five target-first objects.")
    require(int((folds["endpoint_minus_inverse"] > 0).sum()) == 4, "Expected four of five positive fold contrasts.")
    require((target["n_bootstrap"] == 5000).all() and int(pair.iloc[0]["n_bootstrap"]) == 5000, "Bootstrap count changed.")
    require(
        set(replacement["object"])
        == {"Measured state", "Named risk", "Transition", "Intervention displacement", "Benefit"},
        "Target-first object set changed.",
    )
    return {"paths": paths, "policy": policy, "target": target, "pair": pair, "folds": folds, "replacement": replacement}


def main_framework_rows() -> pd.DataFrame:
    return pd.DataFrame(
        [
            {
                "object": "Measured state",
                "target_estimand": "Z or P(Z | A,W,X) under a declared state model",
                "identifying_structure_and_evidence": "Anchored measurement model; reliability, invariance and same-age validation",
                "permissible_claim": "Measured or model-defined state under the declared reference",
            },
            {
                "object": "Named risk",
                "target_estimand": "P(Y at horizon Delta | A,W,X) or its declared loss-specific Bayes act",
                "identifying_structure_and_evidence": "Named endpoint, horizon and loss; held-out validation and calibration in the target population",
                "permissible_claim": "Risk of the named endpoint over the declared horizon",
            },
            {
                "object": "Transition",
                "target_estimand": "P(Z at t+Delta | Z at t,A,W) or a declared within-person change",
                "identifying_structure_and_evidence": "Repeated measurements, time scale and longitudinal transition model",
                "permissible_claim": "Within-person dynamics over the declared interval",
            },
            {
                "object": "Intervention displacement",
                "target_estimand": "Contrast in a prespecified measure S under intervention strategies",
                "identifying_structure_and_evidence": "Randomized or causal design, prespecified contrast, follow-up and measurement invariance",
                "permissible_claim": "The intervention changes the declared measure",
            },
            {
                "object": "Benefit",
                "target_estimand": "E[Y(1)-Y(0)] or another declared decision-specific causal estimand",
                "identifying_structure_and_evidence": "Treatment strategies, outcome, causal assumptions, follow-up and decision population",
                "permissible_claim": "Effect on the named outcome in the declared decision context",
            },
        ]
    )


def write_source_data(data: dict[str, object]) -> tuple[list[Path], list[Path]]:
    main_dir = MAIN_PACKAGE / "source_data"
    ed_dir = ED_PACKAGE / "source_data"

    main_rows = main_framework_rows()
    main_map = main_dir / "panel_c_estimand_specific_replacement_map.tsv"
    main_route = main_dir / "panels_a_b_target_first_routes.tsv"
    main_index = main_dir / "source_data_index.tsv"
    main_rows.to_csv(main_map, sep="\t", index=False)
    pd.DataFrame(
        [
            ("a", "chronological-age estimation", "age-supervised decoder", "conditional chronological-age statistic", "chronological-age estimation"),
            ("a", "biological state/risk/transition/displacement/benefit", "age unit alone", "no identifying bridge", "no automatic construct transfer"),
            ("b", "target-first programme", "declare object and estimand", "choose design/supervision and validate", "restrict claim to identified object"),
        ],
        columns=["panel", "question_or_claim", "input_or_start", "required_bridge", "permissible_output"],
    ).to_csv(main_route, sep="\t", index=False)
    pd.DataFrame(
        [
            ("a-b", main_route.name, "task-object route and target-first claim gate"),
            ("c", main_map.name, "estimand-specific replacement map for five declared objects"),
        ],
        columns=["panel", "file", "content"],
    ).to_csv(main_index, sep="\t", index=False)

    ed_files: list[Path] = []
    mapping = {
        "panel_a_frozen_policy_definitions.tsv": data["policy"],
        "panel_b_target_policy_intervals.tsv": data["target"],
        "panel_c_pooled_policy_difference.tsv": data["pair"],
        "panel_c_fold_policy_differences.tsv": data["folds"],
    }
    for filename, frame in mapping.items():
        path = ed_dir / filename
        frame.to_csv(path, sep="\t", index=False)
        ed_files.append(path)
    ed_index = ed_dir / "source_data_index.tsv"
    pd.DataFrame(
        [
            ("a", "panel_a_frozen_policy_definitions.tsv", "two source-frozen supervision targets, rules and selected domains"),
            ("b", "panel_b_target_policy_intervals.tsv", "target-cycle endpoint gains and fixed-prediction bootstrap intervals"),
            ("c", "panel_c_pooled_policy_difference.tsv", "pooled paired endpoint-directed minus inverse-age contrast"),
            ("c", "panel_c_fold_policy_differences.tsv", "five target-fold paired contrasts"),
        ],
        columns=["panel", "file", "content"],
    ).to_csv(ed_index, sep="\t", index=False)
    ed_files.append(ed_index)
    return [main_route, main_map, main_index], ed_files


def build_main_figure() -> dict[str, Path]:
    fig = plt.figure(figsize=(7.15, 5.55), facecolor=WHITE)
    grid = fig.add_gridspec(2, 2, left=0.045, right=0.985, bottom=0.055, top=0.965, height_ratios=[1.0, 3.45], hspace=0.28, wspace=0.22)
    ax_a = fig.add_subplot(grid[0, 0])
    ax_b = fig.add_subplot(grid[0, 1])
    ax_c = fig.add_subplot(grid[1, :])

    # Panel a: identify the legitimate endpoint of the age task and stop the shortcut.
    ax_a.set_axis_off()
    panel_label(ax_a, "a", "The age task identifies an age statistic")
    box(ax_a, (0.00, 0.50), 0.25, 0.28, "Measurements\n$X$ and background $W$", face=LIGHT, edge=MUTED, fontsize=6.2, weight="bold")
    arrow(ax_a, (0.265, 0.64), (0.355, 0.64), color=BLUE)
    box(ax_a, (0.37, 0.50), 0.25, 0.28, "Age-supervised\ndecoder", face=BLUE_LIGHT, edge=BLUE, fontsize=6.4, weight="bold")
    arrow(ax_a, (0.635, 0.64), (0.725, 0.64), color=BLUE)
    box(ax_a, (0.74, 0.50), 0.25, 0.28, "Chronological-age\ntask statistic", face=WHITE, edge=BLUE, fontsize=6.25, weight="bold")
    ax_a.plot([0.865, 0.865], [0.49, 0.35], transform=ax_a.transAxes, color=RED, linewidth=1.0, linestyle=(0, (3, 2)))
    ax_a.text(0.865, 0.29, "no automatic identity transfer", transform=ax_a.transAxes, ha="center", va="center", fontsize=5.7, color=RED, fontweight="bold")
    ax_a.text(0.865, 0.10, "state   risk   transition\ndisplacement   benefit", transform=ax_a.transAxes, ha="center", va="center", fontsize=5.7, color=INK, linespacing=1.2)

    # Panel b: a compact target-first gate.
    ax_b.set_axis_off()
    panel_label(ax_b, "b", "A target-first claim gate")
    labels = ["Question", "Object +\nestimand", "Design +\nsupervision", "Validation", "Bounded\nclaim"]
    faces = [LIGHT, BLUE_LIGHT, TEAL_LIGHT, ORANGE_LIGHT, GOLD_LIGHT]
    edges = [MUTED, BLUE, TEAL, ORANGE, GOLD]
    x_positions = [0.00, 0.205, 0.41, 0.615, 0.82]
    for i, (x, label, face, edge) in enumerate(zip(x_positions, labels, faces, edges)):
        box(ax_b, (x, 0.49), 0.16, 0.29, label, face=face, edge=edge, fontsize=6.0, weight="bold")
        if i < len(labels) - 1:
            arrow(ax_b, (x + 0.165, 0.635), (x_positions[i + 1] - 0.01, 0.635), color=MUTED, linewidth=1.0)
    ax_b.text(0.50, 0.20, "Meaning comes from the declared object and evidence, not from expressing a score in years.", transform=ax_b.transAxes, ha="center", va="center", fontsize=5.8, color=NAVY, fontweight="bold")

    # Panel c: the dominant estimand-specific replacement map.
    ax_c.set_axis_off()
    panel_label(ax_c, "c", "Estimand-specific replacements for biological claims")
    columns = [
        (0.00, 0.18, "Declared object"),
        (0.19, 0.27, "Target estimand"),
        (0.47, 0.32, "Identifying structure and evidence"),
        (0.80, 0.20, "Permissible conclusion"),
    ]
    for x, width, label in columns:
        box(ax_c, (x, 0.88), width, 0.095, label, face=NAVY, edge=NAVY, fontsize=6.25, weight="bold", color=WHITE, linewidth=0.9)

    rows = [
        ("Measured state", "$Z$ or $P(Z\mid A,W,X)$\nunder a declared state model", "Anchored measurement model; reliability,\ninvariance and same-age validation", "State under the\ndeclared reference", BLUE_LIGHT, BLUE),
        ("Named risk", "$P(Y_{\Delta}=1\mid A,W,X)$ or a\ndeclared loss-specific Bayes act", "Named endpoint, horizon and loss; held-out\nvalidation and calibration in the target population", "Risk for the named endpoint\nover the declared horizon", TEAL_LIGHT, TEAL),
        ("Transition", "$P(Z_{t+\Delta}\mid Z_t,A,W)$ or a\ndeclared within-person change", "Repeated measurements, time scale and\nlongitudinal transition model", "Within-person dynamics\nover the declared interval", PURPLE_LIGHT, PURPLE),
        ("Intervention\ndisplacement", "Contrast in a prespecified measure $S$\nunder intervention strategies", "Randomized or causal design; prespecified contrast,\nfollow-up and measurement invariance", "Change in the declared\nmeasure under intervention", ORANGE_LIGHT, ORANGE),
        ("Benefit", "$E\{Y(1)-Y(0)\}$ or another\ndecision-specific causal estimand", "Treatment strategies, outcome, causal assumptions,\nfollow-up and decision population", "Effect on the named outcome\nin the decision context", GOLD_LIGHT, GOLD),
    ]
    y_positions = [0.715, 0.555, 0.395, 0.235, 0.075]
    for y, (obj, estimand, evidence, claim, face, edge) in zip(y_positions, rows):
        box(ax_c, (0.00, y), 0.18, 0.12, obj, face=face, edge=edge, fontsize=6.1, weight="bold")
        arrow(ax_c, (0.185, y + 0.06), (0.205, y + 0.06), color=edge, linewidth=0.9)
        box(ax_c, (0.21, y), 0.25, 0.12, estimand, face=WHITE, edge=edge, fontsize=5.8, weight="bold")
        arrow(ax_c, (0.465, y + 0.06), (0.485, y + 0.06), color=edge, linewidth=0.9)
        box(ax_c, (0.49, y), 0.29, 0.12, evidence, face=WHITE, edge=edge, fontsize=5.55)
        arrow(ax_c, (0.785, y + 0.06), (0.805, y + 0.06), color=edge, linewidth=0.9)
        box(ax_c, (0.81, y), 0.19, 0.12, claim, face=face, edge=edge, fontsize=5.6, weight="bold")
    ax_c.text(0.50, 0.008, "Age-equivalent units may re-express a separately identified target; they do not supply object identity or causal meaning.", transform=ax_c.transAxes, ha="center", va="bottom", fontsize=6.05, color=RED, fontweight="bold")

    outputs = {
        "png": MAIN_PACKAGE / "figures" / f"{MAIN_STEM}.png",
        "svg": MAIN_PACKAGE / "figures" / f"{MAIN_STEM}.svg",
        "pdf": MAIN_PACKAGE / "figures" / f"{MAIN_STEM}.pdf",
    }
    fig.savefig(outputs["png"], dpi=600, bbox_inches="tight", pad_inches=0.035, facecolor=WHITE)
    fig.savefig(outputs["svg"], bbox_inches="tight", pad_inches=0.035, facecolor=WHITE)
    fig.savefig(outputs["pdf"], bbox_inches="tight", pad_inches=0.035, facecolor=WHITE)
    plt.close(fig)
    return outputs


def build_ed_figure(data: dict[str, object]) -> dict[str, Path]:
    policy = data["policy"].set_index("policy")
    target = data["target"].set_index("policy")
    pair = data["pair"].iloc[0]
    folds = data["folds"]

    endpoint_policy = policy.loc["source_endpoint_policy"]
    inverse_policy = policy.loc["source_inverse_age_decoder_policy"]
    fig = plt.figure(figsize=(7.15, 4.65), facecolor=WHITE)
    grid = fig.add_gridspec(2, 2, left=0.065, right=0.985, bottom=0.075, top=0.955, height_ratios=[1.05, 1.25], hspace=0.42, wspace=0.34)
    ax_a = fig.add_subplot(grid[0, 0])
    ax_b = fig.add_subplot(grid[0, 1])
    ax_c = fig.add_subplot(grid[1, :])

    ax_a.set_axis_off()
    panel_label(ax_a, "a", "Two frozen source policies in a five-domain menu")
    box(ax_a, (0.13, 0.78), 0.74, 0.13, "Five prespecified equal-category-cost domains\nNHANES 2007-2008 ($n=4{,}599$)", face=LIGHT, edge=MUTED, fontsize=6.25, weight="bold")
    arrow(ax_a, (0.50, 0.77), (0.26, 0.64), color=ORANGE)
    arrow(ax_a, (0.50, 0.77), (0.74, 0.64), color=BLUE)
    box(ax_a, (0.01, 0.42), 0.47, 0.20, "Endpoint-directed\nmaximize held-out mortality\nlog-loss gain", face=ORANGE_LIGHT, edge=ORANGE, fontsize=6.05, weight="bold")
    box(ax_a, (0.52, 0.42), 0.47, 0.20, "Inverse age decoder\nminimize held-out chronological-age\nmean-squared error", face=BLUE_LIGHT, edge=BLUE, fontsize=5.95, weight="bold")
    arrow(ax_a, (0.245, 0.41), (0.245, 0.32), color=ORANGE)
    arrow(ax_a, (0.755, 0.41), (0.755, 0.32), color=BLUE)
    box(ax_a, (0.01, 0.08), 0.47, 0.22, "Selected\nHepatic / protein / injury", face=WHITE, edge=ORANGE, fontsize=6.1, weight="bold")
    box(ax_a, (0.52, 0.08), 0.47, 0.22, "Selected\nMetabolic / adiposity / lipid", face=WHITE, edge=BLUE, fontsize=6.1, weight="bold")
    ax_a.text(0.50, 0.005, "Different objectives and units; selections are not ranked on one source axis.", transform=ax_a.transAxes, ha="center", fontsize=5.55, color=RED, fontweight="bold")

    panel_label(ax_b, "b", "Target-cycle endpoint value of the frozen domains")
    style_axis(ax_b, grid_axis="x")
    order = ["source_endpoint_policy", "source_inverse_age_decoder_policy"]
    y = np.array([1, 0])
    labels = ["Endpoint-directed\nHepatic / protein / injury", "Inverse age decoder\nMetabolic / adiposity / lipid"]
    colors = [ORANGE, BLUE]
    markers = ["o", "D"]
    for yi, key, color, marker in zip(y, order, colors, markers):
        row = target.loc[key]
        ax_b.plot([row["ci_low"], row["ci_high"]], [yi, yi], color=color, linewidth=2.2, solid_capstyle="round")
        ax_b.scatter(row["estimate"], yi, s=45, marker=marker, facecolor=color if key == order[0] else WHITE, edgecolor=color, linewidth=1.2, zorder=3)
        ax_b.text(row["ci_high"] + 0.00028, yi, f"{row['estimate']:.4f}", va="center", fontsize=6.0, color=color, fontweight="bold")
    ax_b.axvline(0, color=MUTED, linewidth=0.9)
    ax_b.set_yticks(y, labels)
    ax_b.set_ylim(-0.65, 1.65)
    ax_b.set_xlim(-0.0025, 0.0125)
    ax_b.set_xlabel("Five-year mortality log-loss gain\nbeyond age and background", fontsize=6.15, color=INK)
    ax_b.text(
        0.99,
        0.97,
        "NHANES 2009-2010\n$n=4{,}896$; 157 deaths\n95% fixed-prediction bootstrap; 5,000 resamples",
        transform=ax_b.transAxes,
        ha="right",
        va="top",
        fontsize=5.7,
        color=MUTED,
        linespacing=1.2,
    )

    panel_label(ax_c, "c", "Paired endpoint-directed minus inverse-age difference")
    style_axis(ax_c, grid_axis="x")
    fold_values = folds["endpoint_minus_inverse"].to_numpy(float)
    y_rows = np.arange(6)[::-1]
    labels_c = ["Pooled (95% CI)"] + [f"Fold {i}" for i in range(1, 6)]
    ax_c.plot([pair["ci_low"], pair["ci_high"]], [y_rows[0], y_rows[0]], color=NAVY, linewidth=2.5, solid_capstyle="round")
    ax_c.scatter(pair["estimate"], y_rows[0], s=52, marker="D", facecolor=ORANGE, edgecolor=NAVY, linewidth=0.7, zorder=4)
    for yi, value in zip(y_rows[1:], fold_values):
        ax_c.scatter(value, yi, s=36, marker="o", facecolor=ORANGE if value > 0 else WHITE, edgecolor=ORANGE, linewidth=1.1, zorder=3)
    ax_c.axvline(0, color=MUTED, linewidth=0.9)
    ax_c.set_yticks(y_rows, labels_c)
    ax_c.set_ylim(-0.65, 5.65)
    ax_c.set_xlim(-0.0030, 0.0126)
    ax_c.set_xlabel("Difference in target log-loss gain (positive favours endpoint-directed selection)", fontsize=6.15, color=INK)
    ax_c.text(0.99, 0.96, f"pooled {pair['estimate']:.4f}  |  95% CI {pair['ci_low']:.4f} to {pair['ci_high']:.4f}  |  4/5 folds positive", transform=ax_c.transAxes, ha="right", va="top", fontsize=5.95, color=NAVY, fontweight="bold", bbox=dict(facecolor=WHITE, edgecolor="none", pad=1.2))

    outputs = {
        "png": ED_PACKAGE / "figures" / f"{ED_STEM}.png",
        "svg": ED_PACKAGE / "figures" / f"{ED_STEM}.svg",
        "pdf": ED_PACKAGE / "figures" / f"{ED_STEM}.pdf",
    }
    fig.savefig(outputs["png"], dpi=600, bbox_inches="tight", pad_inches=0.04, facecolor=WHITE)
    fig.savefig(outputs["svg"], bbox_inches="tight", pad_inches=0.04, facecolor=WHITE)
    fig.savefig(outputs["pdf"], bbox_inches="tight", pad_inches=0.04, facecolor=WHITE)
    plt.close(fig)
    return outputs


def write_docs() -> tuple[list[Path], list[Path]]:
    main_legend = MAIN_PACKAGE / "FIGURE_LEGEND_BILINGUAL.md"
    main_readme = MAIN_PACKAGE / "README.md"
    ed_legend = ED_PACKAGE / "FIGURE_LEGEND_BILINGUAL.md"
    ed_readme = ED_PACKAGE / "README.md"

    main_legend.write_text(
        """# Figure 4 | Target-first evidence replaces construct transfer from an age unit

**a**, Chronological-age supervision maps measurements and background to a statistic for reconstructing chronological age. This is a valid age-estimation route, but the resulting age unit does not automatically identify biological state, named risk, transition, intervention displacement or benefit. **b**, A target-first claim gate begins with the scientific question, declares the object and estimand, chooses the corresponding design and supervision, validates the resulting statistic, and limits the conclusion to the object supported by that evidence. **c**, Estimand-specific replacement map. A measured or latent state requires an anchored measurement model and state definition; named risk requires an endpoint, horizon, loss and validation population; transition requires longitudinal sampling and a transition model; intervention displacement requires a prespecified perturbation contrast and measurement response; and benefit requires a causal estimand, named outcome and decision context. Age-equivalent units may re-express a separately identified target, but do not themselves supply object identity or causal meaning.

# 图 4｜Target-first 证据取代由年龄单位直接转移构念含义

**a**，时序年龄监督把测量与背景映射为重建时序年龄的统计量。这是一条合法的年龄估计路径，但所得年龄单位不会自动识别生物状态、命名风险、转变、干预位移或获益。**b**，target-first 主张闸门从科学问题出发，先声明对象与 estimand，再选择相应的设计和监督，验证所得统计量，并把结论限制在证据真正支持的对象上。**c**，estimand-specific 替代图。测量或潜在状态需要有锚定的测量模型与状态定义；命名风险需要终点、时间窗、损失函数与验证人群；转变需要纵向采样与转变模型；干预位移需要预设的扰动对比与测量反应；获益需要因果 estimand、命名结局与决策语境。年龄等价单位可以重表达一个已被单独识别的目标，但不能自行赋予对象身份或因果含义。
""",
        encoding="utf-8",
    )
    main_readme.write_text(
        """# v6.8 Figure 4 target-first framework v0.1

Submission-facing, fit-free successor that makes the estimand-specific replacement programme the whole main figure. It reworks only the conceptual target-first map from the frozen v6.6 Figure 4 package. No empirical acquisition value is presented as a main-figure result.

The figure separates the valid chronological-age task from unsupported construct transfer, then shows the target-first claim gate and five object-specific replacements: measured state, named risk, transition, intervention displacement and benefit. Every row names its estimand, identifying structure and permissible conclusion.

The companion acquisition result has been moved to `outputs/v6_8_extended_data_figure4_bounded_acquisition_v0_1/`. This package does not edit the manuscript, evidence registry, project state or frozen predecessor outputs.
""",
        encoding="utf-8",
    )
    ed_legend.write_text(
        """# Extended Data Figure 4 | Endpoint-directed acquisition in a declared five-domain equal-category-cost comparison

**a**, Two policies were frozen in NHANES 2007-2008 from five prespecified equal-category-cost measurement domains. The endpoint-directed policy selected hepatic, protein and injury measurements by maximizing held-out five-year mortality log-loss gain. The inverse chronological-age decoder selected metabolic, adiposity and lipid measurements by minimizing held-out chronological-age mean-squared error. The source criteria have different targets and units and are therefore shown as separate routes. **b**, In NHANES 2009-2010, the frozen endpoint-directed and inverse-age domains yielded log-loss gains beyond age and background of 0.006418 (95% fixed-prediction bootstrap interval, 0.002297 to 0.010652) and 0.000646 (-0.001195 to 0.002481), respectively. **c**, The paired endpoint-directed minus inverse-age difference was 0.005772 (0.001380 to 0.010390); four of five target-fold estimates were positive. The comparison is exploratory cross-cycle evidence within one declared five-domain menu. Equal category status is not equal variable count, monetary budget or real-world cost. Intervals use 5,000 event-stratified participant bootstraps conditional on stored predictions and fixed source selections; they omit refitting and study-development uncertainty. The analysis is not clinical value of information, cost analysis, untouched confirmation, universal transportability or biological-age identification.

# 扩展数据图 4｜在一个预设的五域等类别成本比较中进行终点导向测量选择

**a**，在 NHANES 2007-2008 的五个预设等类别成本测量域中冻结两项策略。终点导向策略按折外五年死亡 log-loss 增量最大选择肝脏／蛋白／损伤测量；逆向时序年龄解码策略按折外年龄均方误差最小选择代谢／脂肪／血脂测量。两种来源判据的目标和单位不同，因此以分离路径呈现。**b**，在 NHANES 2009-2010 中，冻结的终点导向域和逆年龄域相对年龄与背景的 log-loss 增量分别为 0.006418（95% 固定预测 bootstrap 区间 0.002297 至 0.010652）和 0.000646（-0.001195 至 0.002481）。**c**，终点导向减逆年龄策略的配对差值为 0.005772（0.001380 至 0.010390）；五个目标折中四折点估计为正。该结果只是一个预设五域菜单内的探索性跨周期证据。“等类别成本”不等于等变量数、等货币预算或真实成本。区间来自 5,000 次事件分层参与者 bootstrap，并以已存储预测和冻结来源选择为条件，不包含重拟合和研究开发不确定性。该分析不是临床信息价值、成本分析、未查看确认、普遍可迁移性或生物年龄识别。
""",
        encoding="utf-8",
    )
    ed_readme.write_text(
        """# v6.8 Extended Data Figure 4 bounded acquisition v0.1

Fit-free Extended Data successor containing the empirical acquisition panels a-c from the frozen v6.6 Figure 4 package. It reads only the predecessor source-data tables, verifies their hashes against the predecessor manifest, and preserves all stored estimates and intervals.

The title and figure language explicitly bound the result to five prespecified equal-category-cost domains in one exploratory NHANES source-target cycle comparison. Equal-category-cost is not equal budget, equal variable count, monetary cost equivalence or universal measurement selection.

No models were loaded or fit, no random numbers were generated, and no frozen output or canonical manuscript file was modified.
""",
        encoding="utf-8",
    )
    return [main_legend, main_readme], [ed_legend, ed_readme]


def write_package_manifests(
    package: Path,
    package_kind: str,
    figures: dict[str, Path],
    source_files: list[Path],
    docs: list[Path],
    input_paths: list[Path],
    *,
    manual_pass: bool,
) -> None:
    if package_kind == "main":
        contract = {
            "analytical_question": "What object-specific evidence should replace semantic promotion from a chronological-age unit?",
            "takeaway": "Start from the scientific object and estimand; state, risk, transition, intervention displacement and benefit require different identifying structures and support different conclusions.",
            "canonical_families": ["decomposition and progression", "matrix and cohort"],
            "renderer": "static Matplotlib",
            "surface": "standalone static publication figure",
            "footprint": "180 mm wide, three panels, 600 dpi PNG plus editable SVG and vector PDF",
            "panels": {"a": "valid age-task route and blocked identity shortcut", "b": "target-first claim gate", "c": "five-row estimand-specific replacement map"},
            "palette_policy": "relaxed multi-category, five approved roots with direct row labels and repeated geometry",
            "non_color_distinction": "fixed row order, direct labels, column headers and identical left-to-right claim-gate geometry",
            "claim_boundary": "conceptual replacement framework; no empirical effect estimate and no claim that one universal low-dimensional ageing construct is impossible",
        }
    else:
        contract = {
            "analytical_question": "Within one declared five-domain menu, which source-frozen selection policy carried more five-year mortality value into the target NHANES cycle?",
            "takeaway": "The endpoint-directed frozen domain had a positive paired target-cycle advantage over the inverse-age-decoder frozen domain; four of five fold estimates were positive.",
            "canonical_families": ["decomposition and progression", "uncertainty and benchmark", "comparison"],
            "renderer": "static Matplotlib",
            "surface": "standalone static Extended Data figure",
            "footprint": "180 mm wide, three panels, 600 dpi PNG plus editable SVG and vector PDF",
            "panels": {"a": "two source-frozen policy routes with distinct targets and units", "b": "target-cycle policy gains and intervals", "c": "pooled and fold-level paired contrasts"},
            "data_sufficiency": "five prespecified domains; 4,599 source participants; 4,896 target participants with 157 deaths; five folds and 5,000 fixed-prediction bootstrap resamples",
            "palette_policy": "hard two-root cap plus neutrals",
            "non_color_distinction": "separate routes, direct labels, circle versus diamond, filled versus open markers, zero-line context",
            "claim_boundary": "one exploratory source-target cycle pair and five equal-category-cost domains; not equal budget, equal variable count, clinical value of information, cost analysis, untouched confirmation, universal transportability or biological-age identification",
        }
    write_json(package / "visualization" / "chart_contract.json", contract)

    pd.DataFrame(
        [{"path": path.relative_to(ROOT).as_posix(), "bytes": path.stat().st_size, "sha256": sha256(path)} for path in input_paths]
    ).to_csv(package / "visualization" / "input_manifest.tsv", sep="\t", index=False)

    rows = []
    for kind, path in figures.items():
        rows.append({"artifact": kind, "path": path.relative_to(ROOT).as_posix(), "bytes": path.stat().st_size, "sha256": sha256(path)})
    for path in source_files:
        rows.append({"artifact": "source_data", "path": path.relative_to(ROOT).as_posix(), "bytes": path.stat().st_size, "sha256": sha256(path)})
    for path in docs:
        rows.append({"artifact": "documentation", "path": path.relative_to(ROOT).as_posix(), "bytes": path.stat().st_size, "sha256": sha256(path)})
    pd.DataFrame(rows).to_csv(package / "visualization" / "figure_manifest.tsv", sep="\t", index=False)

    write_json(
        package / "visualization" / "visualization_audit.json",
        {
            "status": "PASS" if manual_pass else "PENDING_MANUAL_VISUAL_QA",
            "analysis_version": VERSION,
            "package_kind": package_kind,
            "source_package": "v6_6_figure4_inverse_decoder_v0_1",
            "fit_free": True,
            "models_loaded": False,
            "models_refit": False,
            "random_numbers_generated": False,
            "frozen_inputs_mutated": False,
            "manual_visual_qa": manual_pass,
            "manual_visual_qa_scope": "PNG inspected at exported resolution; PDF rendered and inspected; SVG structure and declared colors checked" if manual_pass else "pending",
            "python": sys.version,
            "platform": platform.platform(),
            "matplotlib": matplotlib.__version__,
            "source_data_files": len(source_files),
            "figure_formats": sorted(figures),
        },
    )


def parse_args() -> argparse.Namespace:
    parser = argparse.ArgumentParser(description=__doc__)
    parser.add_argument("--manual-pass", action="store_true", help="Record manual PNG/PDF/SVG visual QA as complete after inspection.")
    return parser.parse_args()


def main() -> None:
    args = parse_args()
    ensure_packages()
    data = load_frozen_source()
    main_source, ed_source = write_source_data(data)
    main_figures = build_main_figure()
    ed_figures = build_ed_figure(data)
    main_docs, ed_docs = write_docs()
    inputs = [SCRIPT] + list(data["paths"].values())
    write_package_manifests(MAIN_PACKAGE, "main", main_figures, main_source, main_docs, inputs, manual_pass=args.manual_pass)
    write_package_manifests(ED_PACKAGE, "extended_data", ed_figures, ed_source, ed_docs, inputs, manual_pass=args.manual_pass)
    status = "PASS" if args.manual_pass else "PENDING_MANUAL_VISUAL_QA"
    print(f"{status}: built main Figure 4 and bounded Extended Data acquisition figure from frozen source data; no models loaded or fitted.")


if __name__ == "__main__":
    main()
```
