## Supplementary_Software_1 for "Inferential boundaries of age prediction: why prediction does not establish biological age measurement": plot_v6_9_figure1_precision_revision_v0_1.py

```
#!/usr/bin/env python
"""Build the v6.9 precision-revised Figure 1 resource.

This entry point is deliberately fit-free. It reads only frozen analytical
outputs, performs deterministic tabular reshaping for figure Source Data and
renders publication graphics. It never loads a fitted estimator, refits a
model, draws random numbers or mutates the frozen analysis directories.
"""

from __future__ import annotations

import hashlib
import json
import platform
import shutil
import sys
from pathlib import Path

import matplotlib
import matplotlib.pyplot as plt
from matplotlib.lines import Line2D
from matplotlib.patches import FancyArrowPatch, FancyBboxPatch
import numpy as np
import pandas as pd


ROOT = Path(__file__).resolve().parents[1]
SCRIPT = Path(__file__).resolve()

NHANES = ROOT / "outputs" / "nhanes_5y_age_decoder_source_information_probe_v0_2"
ACQ = ROOT / "outputs" / "nature_v4_3_result_figure_packages_v0_1" / "prior_positive_results" / "source_data"
HSCT_PACKAGE = ROOT / "outputs" / "hsct_age_reconstruction_semantic_counterexample_figure_v0_3"
OLD_FIG1 = ROOT / "figures" / "nature_v4_1_draft_v0_1" / "Figure_1_Chronological_age_loss_does_not_identify_biological_age.png"

PACKAGES = {
    "figure1": ROOT / "outputs" / "v6_9_figure1_theory_chain_v0_1",
    "figure3": ROOT / "outputs" / "v6_5_figure3_nhanes_age_decoder_v0_2",
    "figure4": ROOT / "outputs" / "v6_4_figure4_target_first_v0_1",
}
INDEX_PACKAGE = ROOT / "outputs" / "v6_5_main_figure_sync_v0_2"

INK = "#202124"
NAVY = "#18243D"
MUTED = "#62676D"
GRID = "#D9DDE1"
LIGHT = "#EEF1F3"
WHITE = "#FFFFFF"
BLUE = "#315F85"
BLUE_LIGHT = "#E5EEF5"
TEAL = "#087E8B"
TEAL_LIGHT = "#D7EEF0"
ORANGE = "#D97732"
ORANGE_LIGHT = "#F7E6D8"
GOLD = "#C08A18"
GOLD_LIGHT = "#F8EED3"
PURPLE = "#6D5396"
PURPLE_LIGHT = "#EEEAF7"
RED = "#B94B45"
RED_LIGHT = "#F6E6E4"

matplotlib.rcParams.update(
    {
        "font.family": "sans-serif",
        "font.sans-serif": ["Arial", "DejaVu Sans"],
        "font.size": 7.2,
        "axes.unicode_minus": False,
        "svg.fonttype": "none",
        "pdf.fonttype": 42,
        "ps.fonttype": 42,
    }
)


def require(condition: bool, message: str) -> None:
    if not condition:
        raise RuntimeError(message)


def sha256(path: Path) -> str:
    digest = hashlib.sha256()
    with path.open("rb") as handle:
        for block in iter(lambda: handle.read(1024 * 1024), b""):
            digest.update(block)
    return digest.hexdigest()


def write_json(path: Path, payload: object) -> None:
    path.write_text(json.dumps(payload, indent=2, ensure_ascii=False) + "\n", encoding="utf-8")


def ensure_package(path: Path) -> None:
    (path / "figures").mkdir(parents=True, exist_ok=True)
    (path / "source_data").mkdir(parents=True, exist_ok=True)
    (path / "visualization").mkdir(parents=True, exist_ok=True)


def save_figure(fig: plt.Figure, package: Path, stem: str) -> dict[str, Path]:
    outputs = {
        "png": package / "figures" / f"{stem}.png",
        "svg": package / "figures" / f"{stem}.svg",
        "pdf": package / "figures" / f"{stem}.pdf",
    }
    fig.savefig(outputs["png"], dpi=600, bbox_inches="tight", pad_inches=0.045, facecolor=WHITE)
    fig.savefig(outputs["svg"], bbox_inches="tight", pad_inches=0.045, facecolor=WHITE)
    fig.savefig(outputs["pdf"], bbox_inches="tight", pad_inches=0.045, facecolor=WHITE)
    plt.close(fig)
    return outputs


def box(
    ax: plt.Axes,
    xy: tuple[float, float],
    width: float,
    height: float,
    text: str,
    *,
    face: str = WHITE,
    edge: str = MUTED,
    fontsize: float = 6.5,
    weight: str = "normal",
    color: str = INK,
    linewidth: float = 1.0,
    linestyle: str = "-",
) -> FancyBboxPatch:
    patch = FancyBboxPatch(
        xy,
        width,
        height,
        boxstyle="round,pad=0.016,rounding_size=0.017",
        transform=ax.transAxes,
        facecolor=face,
        edgecolor=edge,
        linewidth=linewidth,
        linestyle=linestyle,
        clip_on=False,
    )
    ax.add_patch(patch)
    ax.text(
        xy[0] + width / 2,
        xy[1] + height / 2,
        text,
        transform=ax.transAxes,
        ha="center",
        va="center",
        fontsize=fontsize,
        fontweight=weight,
        color=color,
        linespacing=1.14,
    )
    return patch


def arrow(
    ax: plt.Axes,
    start: tuple[float, float],
    end: tuple[float, float],
    *,
    color: str = MUTED,
    linewidth: float = 1.1,
    linestyle: str = "-",
    style: str = "-|>",
) -> None:
    ax.add_patch(
        FancyArrowPatch(
            start,
            end,
            transform=ax.transAxes,
            arrowstyle=style,
            mutation_scale=8.5,
            color=color,
            linewidth=linewidth,
            linestyle=linestyle,
            shrinkA=0,
            shrinkB=0,
            clip_on=False,
        )
    )


def panel_label(ax: plt.Axes, label: str, title: str) -> None:
    ax.text(-0.02, 1.055, label, transform=ax.transAxes, fontsize=10, fontweight="bold", color=INK, va="bottom")
    ax.text(0.055, 1.055, title, transform=ax.transAxes, fontsize=8.2, fontweight="bold", color=INK, va="bottom")


def style_axis(ax: plt.Axes, *, grid_axis: str = "both") -> None:
    ax.spines["top"].set_visible(False)
    ax.spines["right"].set_visible(False)
    ax.spines["left"].set_color(MUTED)
    ax.spines["bottom"].set_color(MUTED)
    ax.tick_params(colors=INK, labelsize=6.3, width=0.7, length=3)
    ax.grid(axis=grid_axis, color=GRID, linewidth=0.55, alpha=0.78)
    ax.set_axisbelow(True)


def write_common_visualization_files(
    package: Path,
    analysis_version: str,
    chart_contract: dict,
    inputs: list[Path],
    figures: dict[str, Path],
    source_files: list[Path],
) -> None:
    write_json(package / "visualization" / "chart_contract.json", chart_contract)
    input_rows = []
    for path in inputs:
        input_rows.append(
            {
                "path": path.relative_to(ROOT).as_posix(),
                "bytes": path.stat().st_size,
                "sha256": sha256(path),
            }
        )
    pd.DataFrame(input_rows).to_csv(package / "visualization" / "input_manifest.tsv", sep="\t", index=False)

    output_rows = []
    for kind, path in figures.items():
        output_rows.append(
            {
                "artifact": kind,
                "path": path.relative_to(ROOT).as_posix(),
                "bytes": path.stat().st_size,
                "sha256": sha256(path),
            }
        )
    for path in source_files:
        output_rows.append(
            {
                "artifact": "source_data",
                "path": path.relative_to(ROOT).as_posix(),
                "bytes": path.stat().st_size,
                "sha256": sha256(path),
            }
        )
    pd.DataFrame(output_rows).to_csv(package / "visualization" / "figure_manifest.tsv", sep="\t", index=False)
    write_json(
        package / "visualization" / "visualization_audit.json",
        {
            "status": "PASS",
            "analysis_version": analysis_version,
            "fit_free": True,
            "models_loaded": False,
            "models_refit": False,
            "random_numbers_generated": False,
            "frozen_inputs_mutated": False,
            "python": sys.version,
            "platform": platform.platform(),
            "matplotlib": matplotlib.__version__,
            "source_data_files": len(source_files),
            "figure_formats": sorted(figures),
        },
    )


def build_figure1() -> dict[str, Path]:
    package = PACKAGES["figure1"]
    ensure_package(package)
    stem = "Figure_1_age_decoding_identifies_an_age_task_not_biological_age"

    fig = plt.figure(figsize=(7.15, 6.6), facecolor=WHITE)
    grid = fig.add_gridspec(3, 2, left=0.055, right=0.985, bottom=0.045, top=0.965, hspace=0.43, wspace=0.27, height_ratios=[1.0, 1.0, 0.82])
    ax_a = fig.add_subplot(grid[0, 0])
    ax_b = fig.add_subplot(grid[0, 1])
    ax_c = fig.add_subplot(grid[1, 0])
    ax_d = fig.add_subplot(grid[1, 1])
    ax_e = fig.add_subplot(grid[2, :])
    for ax in [ax_a, ax_b, ax_c, ax_d, ax_e]:
        ax.set_axis_off()

    panel_label(ax_a, "a", "The source task identifies an age statistic")
    box(ax_a, (0.03, 0.63), 0.27, 0.17, "Measurements $X$\nand background $W$", face=BLUE_LIGHT, edge=BLUE, fontsize=6.4, weight="bold")
    arrow(ax_a, (0.30, 0.715), (0.40, 0.715), color=BLUE)
    box(ax_a, (0.41, 0.61), 0.27, 0.21, "Age-supervised\ndecoder\n$q_A(W,X)$", face=TEAL_LIGHT, edge=TEAL, fontsize=7.1, weight="bold")
    arrow(ax_a, (0.69, 0.715), (0.78, 0.715), color=TEAL)
    box(ax_a, (0.79, 0.63), 0.18, 0.17, "Age-task\nstatistic", face=WHITE, edge=INK, fontsize=6.8, weight="bold")
    ax_a.text(0.50, 0.50, "squared loss → conditional mean of chronological age", transform=ax_a.transAxes, ha="center", va="center", fontsize=6.5, color=MUTED)
    box(ax_a, (0.08, 0.16), 0.23, 0.15, "State identity", face=ORANGE_LIGHT, edge=ORANGE, fontsize=6.5, weight="bold")
    box(ax_a, (0.385, 0.16), 0.23, 0.15, "Improvement\ndirection", face=ORANGE_LIGHT, edge=ORANGE, fontsize=6.5, weight="bold")
    box(ax_a, (0.69, 0.16), 0.23, 0.15, "Clinical\nbenefit", face=ORANGE_LIGHT, edge=ORANGE, fontsize=6.5, weight="bold")
    ax_a.text(0.50, 0.075, "same-age labels supply none of these meanings", transform=ax_a.transAxes, ha="center", fontsize=6.5, color=RED, fontweight="bold")

    panel_label(ax_b, "b", "Same age scale, opposite within-age orientation")
    box(ax_b, (0.03, 0.42), 0.27, 0.20, "Observable law\n$P(A,W,X)$", face=WHITE, edge=INK, fontsize=7.3, weight="bold")
    arrow(ax_b, (0.30, 0.54), (0.44, 0.69), color=BLUE)
    box(ax_b, (0.44, 0.63), 0.50, 0.16, "Same optimal decoder and\nattainable age performance", face=BLUE_LIGHT, edge=BLUE, fontsize=6.8, weight="bold")
    arrow(ax_b, (0.30, 0.48), (0.45, 0.38), color=MUTED, linestyle="--")
    arrow(ax_b, (0.30, 0.45), (0.72, 0.38), color=MUTED, linestyle="--")
    box(ax_b, (0.41, 0.27), 0.25, 0.17, "$H^{(+)}=A+U$\nwithin-age order\nfollows $U$", face=TEAL_LIGHT, edge=TEAL, fontsize=5.9, weight="bold", linestyle="--")
    box(ax_b, (0.70, 0.27), 0.25, 0.17, "$H^{(-)}=A-U$\nwithin-age order\nreverses $U$", face=ORANGE_LIGHT, edge=ORANGE, fontsize=5.9, weight="bold", linestyle="--")
    ax_b.text(0.68, 0.17, "same age centre and within-age scale; opposite ordering", transform=ax_b.transAxes, ha="center", fontsize=6.35, color=INK)
    ax_b.text(0.50, 0.055, "Age supervision cannot choose which direction is biologically older", transform=ax_b.transAxes, ha="center", fontsize=6.25, color=RED, fontweight="bold")

    panel_label(ax_c, "c", "Post-decoding operations do not restore identity")
    box(ax_c, (0.02, 0.63), 0.18, 0.15, "Source\n$X$", face=BLUE_LIGHT, edge=BLUE, fontsize=7.2, weight="bold")
    arrow(ax_c, (0.20, 0.705), (0.29, 0.705), color=BLUE)
    box(ax_c, (0.30, 0.63), 0.18, 0.15, "Scalar\n$S=q(X)$", face=TEAL_LIGHT, edge=TEAL, fontsize=7.0, weight="bold")
    arrow(ax_c, (0.48, 0.705), (0.57, 0.705), color=TEAL)
    box(ax_c, (0.58, 0.63), 0.18, 0.15, "Raw gap\n$S-A$", face=WHITE, edge=MUTED, fontsize=7.0, weight="bold")
    arrow(ax_c, (0.76, 0.705), (0.83, 0.705), color=MUTED)
    box(ax_c, (0.84, 0.63), 0.14, 0.15, "Centred\ngap", face=WHITE, edge=MUTED, fontsize=6.7, weight="bold")
    ax_c.text(0.50, 0.49, "deterministic reparameterizations of one compressed representation", transform=ax_c.transAxes, ha="center", fontsize=6.3, color=MUTED)
    ax_c.arrow(0.09, 0.29, 0.25, 0.0, transform=ax_c.transAxes, width=0.006, head_width=0.035, head_length=0.028, color=BLUE, length_includes_head=True)
    ax_c.arrow(0.09, 0.29, 0.0, 0.18, transform=ax_c.transAxes, width=0.006, head_width=0.035, head_length=0.028, color=ORANGE, length_includes_head=True)
    ax_c.arrow(0.09, 0.29, 0.16, -0.14, transform=ax_c.transAxes, width=0.006, head_width=0.035, head_length=0.028, color=PURPLE, length_includes_head=True)
    ax_c.text(0.36, 0.36, "scalarization retains only information\nencoded by $q_A$; gaps cannot recover\nwhat $q_A$ did not retain", transform=ax_c.transAxes, ha="left", va="center", fontsize=5.95, color=INK, fontweight="bold")
    box(ax_c, (0.62, 0.035), 0.34, 0.22, "Perfect recovery\n$S=A \Rightarrow S-A=0$\nwhile heterogeneity\nmay remain", face=RED_LIGHT, edge=RED, fontsize=5.7, weight="bold")

    panel_label(ax_d, "d", "Shared covariance explains genuine success")
    box(ax_d, (0.04, 0.62), 0.25, 0.17, "Common process\nor proxy overlap", face=GOLD_LIGHT, edge=GOLD, fontsize=6.7, weight="bold")
    arrow(ax_d, (0.29, 0.69), (0.44, 0.77), color=GOLD)
    arrow(ax_d, (0.29, 0.68), (0.44, 0.48), color=GOLD)
    box(ax_d, (0.45, 0.68), 0.23, 0.16, "Age-readable\nmeasurements", face=BLUE_LIGHT, edge=BLUE, fontsize=6.5, weight="bold")
    box(ax_d, (0.45, 0.37), 0.23, 0.16, "Named endpoint\n$Y$", face=ORANGE_LIGHT, edge=ORANGE, fontsize=6.5, weight="bold")
    arrow(ax_d, (0.68, 0.76), (0.78, 0.76), color=BLUE)
    box(ax_d, (0.79, 0.68), 0.18, 0.16, "Clock or\ngap", face=TEAL_LIGHT, edge=TEAL, fontsize=6.6, weight="bold")
    arrow(ax_d, (0.88, 0.67), (0.68, 0.48), color=TEAL)
    ax_d.text(0.53, 0.25, "Association and incremental prediction can be real", transform=ax_d.transAxes, ha="center", fontsize=6.7, color=TEAL, fontweight="bold")
    ax_d.text(0.53, 0.12, "but do not establish construct identity", transform=ax_d.transAxes, ha="center", fontsize=6.8, color=RED, fontweight="bold")

    panel_label(ax_e, "e", "Population Bayes-risk ordering and predictive sufficiency")
    box(ax_e, (0.02, 0.43), 0.18, 0.24, "Source\n$(X,A,W)$", face=BLUE_LIGHT, edge=BLUE, fontsize=7.4, weight="bold")
    arrow(ax_e, (0.20, 0.55), (0.31, 0.55), color=BLUE)
    box(ax_e, (0.32, 0.43), 0.18, 0.24, "Clock view\n$(q(X),A,W)$", face=TEAL_LIGHT, edge=TEAL, fontsize=7.1, weight="bold")
    ax_e.text(0.35, 0.70, "$q$ is deterministic", transform=ax_e.transAxes, ha="center", fontsize=6.2, color=MUTED)
    box(ax_e, (0.61, 0.54), 0.36, 0.20, "For declared target $T$ and loss $\\ell$,\nsource Bayes risk ≤ clock Bayes risk", face=WHITE, edge=INK, fontsize=6.7, weight="bold")
    box(ax_e, (0.61, 0.22), 0.36, 0.20, "One-loss equality is loss-specific;\nfull-law equality requires\n$T\\perp X\\mid S,A,W$", face=PURPLE_LIGHT, edge=PURPLE, fontsize=6.25, weight="bold")
    arrow(ax_e, (0.50, 0.58), (0.60, 0.64), color=INK)
    arrow(ax_e, (0.50, 0.51), (0.60, 0.34), color=PURPLE)
    ax_e.text(0.26, 0.17, "Finite-sample regularization or cost advantages remain possible", transform=ax_e.transAxes, ha="center", fontsize=6.5, color=MUTED, fontstyle="italic")

    figures = save_figure(fig, package, stem)

    concept_rows = [
        ("a", "edge", "measurements", "age_decoder", "age supervision"),
        ("a", "edge", "age_decoder", "age_statistic", "identified task output"),
        ("a", "missing_bridge", "age_statistic", "biological_age_identity", "not supplied by same-age labels"),
        ("b", "equivalence", "H_plus=A+U", "H_minus=A-U", "same observable law, age centre and scale; opposite within-age orientation"),
        ("c", "operator", "X", "clock_to_raw_gap_to_centred_gap", "deterministic scalar chain"),
        ("c", "boundary", "perfect_age_recovery", "zero_gap", "biological heterogeneity may remain"),
        ("d", "mechanism", "shared_covariance", "endpoint_association", "genuine association without construct identity"),
        ("e", "comparison", "clock_view", "source_view", "source view contains clock view"),
        ("e", "ordering", "source_bayes_risk", "clock_bayes_risk", "source risk cannot exceed deterministic-clock risk"),
        ("e", "equality_condition", "one_loss", "equal_risk", "loss-specific Bayes-risk equivalence"),
        ("e", "equality_condition", "full_target_law", "predictive_sufficiency", "T independent of X conditional on S,A,W"),
    ]
    source = package / "source_data" / "figure1_theory_chain.tsv"
    pd.DataFrame(concept_rows, columns=["panel", "relation", "from", "to", "interpretation"]).to_csv(source, sep="\t", index=False)
    source_index = package / "source_data" / "source_data_index.tsv"
    pd.DataFrame([{"panel": "a-e", "file": source.name, "content": "conceptual analytical map; no empirical estimates"}]).to_csv(source_index, sep="\t", index=False)

    legend = """# Figure 1 | Chronological-age decoding identifies an age task, not a biological-age construct

**a**, Chronological-age supervision identifies an age-task statistic; same-age labels do not supply state identity, an improvement direction or clinical benefit. **b**, Even after fixing the age centre and within-age scale, $H^{(+)}=A+U$ and $H^{(-)}=A-U$ reverse the orientation of the same non-degenerate contrast while leaving the observable age task unchanged; conditional symmetry additionally gives the same conditional distribution. **c**, A clock, its raw gap and its conditionally centred gap are deterministic scalar operations; post-decoding gaps cannot recover information not retained in the scalar, and perfect age recovery collapses the gap even when within-age heterogeneity remains. **d**, Shared age-related covariance and proxy overlap can yield genuine endpoint associations without establishing construct identity. **e**, For a declared target and loss, source measurements cannot have greater population Bayes risk than their deterministic clock. Equality under one loss is loss-specific; equality of the complete conditional target law requires predictive sufficiency. Finite-sample regularization, measurement-cost and implementation advantages remain possible.

# 图 1｜时序年龄解码识别的是年龄任务，而不是生物年龄构念

**a**，时序年龄监督识别一个年龄任务统计量；同龄标签本身不提供状态身份、改善方向或临床获益。**b**，即使固定年龄中心与同龄尺度，$H^{(+)}=A+U$ 与 $H^{(-)}=A-U$ 仍会反转同一非退化对比的方向，而不改变可观测年龄任务；若条件分布对称，两者还具有相同的条件分布。**c**，时钟、原始 gap 与条件中心化 gap 都是确定性的标量操作；解码后的 gap 不能恢复标量中未保留的信息，而完美年龄复原会令 gap 归零，即使同龄异质性仍存在。**d**，共享年龄相关协方差与代理重叠可以产生真实终点关联，却不建立构念身份。**e**，对已声明的目标和损失，源测量的总体 Bayes 风险不会高于其确定性时钟；单一损失下的风险相等只是损失特异的，完整目标条件分布相等需要预测充分性。有限样本正则化、测量成本和实施优势仍可能存在。
"""
    (package / "FIGURE_LEGEND_BILINGUAL.md").write_text(legend, encoding="utf-8")
    (package / "README.md").write_text(
        "# v6.9 Figure 1 theory chain v0.1\n\nFit-free precision revision of the analytical figure. Scalar compression is described through information retained in the deterministic representation rather than a linear-direction metaphor. No empirical analysis or model fitting is performed.\n",
        encoding="utf-8",
    )
    write_common_visualization_files(
        package,
        "v6_9_figure1_theory_chain_v0_1",
        {
            "analytical_question": "What does chronological-age decoding identify, and which stronger meanings do post-decoding operations or downstream associations fail to confer?",
            "takeaway": "Age decoding identifies an age-task statistic; it does not orient a same-age contrast, and full target-law equivalence requires additional predictive sufficiency.",
            "surface": "standalone static publication figure",
            "footprint": "180 mm wide, five panels, 600 dpi PNG plus editable SVG and vector PDF",
            "panels": {"a": "task-object mismatch", "b": "constrained within-age orientation non-identification", "c": "gap and collapse boundary", "d": "shared covariance", "e": "Bayes-risk ordering and predictive sufficiency"},
            "palette_policy": "hard two-root cap per panel with shared neutral system",
            "non_color_distinction": "box hierarchy, line style, direct text and spatial grouping",
        },
        [SCRIPT],
        figures,
        [source, source_index],
    )
    return figures


def build_figure3() -> dict[str, Path]:
    package = PACKAGES["figure3"]
    ensure_package(package)
    stem = "Figure_3_NHANES_age_decoder_utility_and_source_measurement_performance"
    pooled_path = NHANES / "pooled_metrics.tsv"
    contrast_path = NHANES / "paired_contrasts.tsv"
    folds_path = NHANES / "fold_metrics.tsv"
    diagnostics_path = NHANES / "age_decoder_diagnostics.json"
    decision_path = NHANES / "decision.json"
    pooled = pd.read_csv(pooled_path, sep="\t")
    contrasts = pd.read_csv(contrast_path, sep="\t")
    folds = pd.read_csv(folds_path, sep="\t")
    diagnostics = json.loads(diagnostics_path.read_text(encoding="utf-8"))
    decision = json.loads(decision_path.read_text(encoding="utf-8"))

    require(list(pooled["model"]) == ["M_WA", "M_CLOCK", "M_X"], "Unexpected pooled model ordering.")
    require(len(folds) == 5 and int(pooled.loc[0, "n"]) == 13724 and int(pooled.loc[0, "events"]) == 470, "Frozen NHANES dimensions changed.")
    require((folds["V_CLOCK"] > 0).all() and (folds["V_SOURCE"] > 0).all(), "Expected positive fold contrasts were not recovered.")
    require(decision.get("decision_category") == "PROMISING_SOURCE_INFORMATION", "Frozen decision changed; stop for governance review.")

    fig = plt.figure(figsize=(7.15, 5.85), facecolor=WHITE)
    grid = fig.add_gridspec(2, 2, left=0.07, right=0.985, bottom=0.07, top=0.96, hspace=0.39, wspace=0.34)
    ax_a = fig.add_subplot(grid[0, 0])
    ax_b = fig.add_subplot(grid[0, 1])
    ax_c = fig.add_subplot(grid[1, 0])
    ax_d = fig.add_subplot(grid[1, 1])

    ax_a.set_axis_off()
    panel_label(ax_a, "a", "Nested cross-fitting separates source and endpoint tasks")
    box(ax_a, (0.03, 0.67), 0.26, 0.16, "Same 50 measurements $X$\n+ background $W$", face=BLUE_LIGHT, edge=BLUE, fontsize=6.5, weight="bold")
    arrow(ax_a, (0.29, 0.75), (0.40, 0.75), color=BLUE)
    box(ax_a, (0.41, 0.65), 0.27, 0.20, "Fold-local ridge\nage decoder\ntrained only on $A$", face=TEAL_LIGHT, edge=TEAL, fontsize=6.6, weight="bold")
    arrow(ax_a, (0.68, 0.75), (0.79, 0.75), color=TEAL)
    box(ax_a, (0.80, 0.67), 0.17, 0.16, "OOF clock\n$q_A(W,X)$", face=WHITE, edge=INK, fontsize=6.5, weight="bold")
    ax_a.text(0.53, 0.56, f"held-out age $R^2$ = {diagnostics['r2']:.2f}; 5 outer folds", transform=ax_a.transAxes, ha="center", fontsize=6.2, color=MUTED)
    model_y = [0.36, 0.20, 0.04]
    model_text = ["$M_{WA}$: age + background", "$M_{CLOCK}$: age + background + OOF clock", "$M_X$: age + background + the same 50 measurements"]
    model_face = [LIGHT, TEAL_LIGHT, BLUE_LIGHT]
    model_edge = [MUTED, TEAL, BLUE]
    for y, text_value, face, edge in zip(model_y, model_text, model_face, model_edge):
        box(ax_a, (0.12, y), 0.76, 0.105, text_value, face=face, edge=edge, fontsize=6.35, weight="bold")
    ax_a.text(0.50, -0.055, "Complete five-year administrative support · one endpoint · one declared model family", transform=ax_a.transAxes, ha="center", fontsize=5.9, color=MUTED, fontstyle="italic")

    panel_label(ax_b, "b", "Out-of-fold mortality performance")
    style_axis(ax_b, grid_axis="x")
    labels = ["Age +\nbackground", "+ age-only\ndecoder", "+ same 50\nmeasurements"]
    colors = ["#AAB1B7", TEAL, BLUE]
    y = np.arange(3)[::-1]
    vals = pooled["log_loss"].to_numpy(float)
    ax_b.barh(y, vals, height=0.56, color=colors, edgecolor=INK, linewidth=0.45)
    ax_b.set_yticks(y, labels)
    ax_b.set_xlim(0, max(vals) * 1.12)
    ax_b.set_xlabel("Five-year all-cause mortality log loss (lower is better)", fontsize=6.3, color=INK)
    for yi, val in zip(y, vals):
        ax_b.text(val + 0.0013, yi, f"{val:.3f}", va="center", fontsize=6.5, color=INK, fontweight="bold")
    ax_b.text(0.99, 0.02, "$n=13{,}724$; 470 deaths", transform=ax_b.transAxes, ha="right", va="bottom", fontsize=6.0, color=MUTED)

    panel_label(ax_c, "c", "Paired held-out predictive value")
    style_axis(ax_c, grid_axis="x")
    order = ["V_CLOCK", "V_SOURCE", "V_X"]
    c = contrasts.set_index("contrast").loc[order]
    labels_c = ["Clock beyond age\nand background", "Source measurements\nbeyond the clock", "Source measurements\nbeyond age and background"]
    y_c = np.arange(3)[::-1]
    col_c = [TEAL, BLUE, NAVY]
    for yi, (_, row), color in zip(y_c, c.iterrows(), col_c):
        ax_c.plot([row["q025"], row["q975"]], [yi, yi], color=color, linewidth=2.2, solid_capstyle="round")
        ax_c.scatter(row["estimate"], yi, s=42, facecolor=color, edgecolor=INK, linewidth=0.5, zorder=3)
        ax_c.text(row["q975"] + 0.00045, yi, f"{row['estimate']:.3f}", va="center", fontsize=6.3, color=color, fontweight="bold")
    ax_c.axvline(0, color=MUTED, linewidth=0.9, linestyle=(0, (4, 2)))
    ax_c.set_yticks(y_c, labels_c)
    ax_c.set_xlim(-0.001, 0.0183)
    ax_c.set_xlabel("Paired out-of-fold log-loss improvement", fontsize=6.3, color=INK)
    ax_c.text(0.02, -0.21, "Intervals condition on stored OOF predictions; they omit refitting and study-development uncertainty.", transform=ax_c.transAxes, ha="left", fontsize=5.7, color=MUTED)

    ax_d.set_axis_off()
    panel_label(ax_d, "d", "Utility, source performance and construct identity are distinct")
    box(ax_d, (0.05, 0.68), 0.90, 0.17, "Observed here\nThe age-only decoder adds held-out mortality value\nbeyond age and background.", face=TEAL_LIGHT, edge=TEAL, fontsize=6.25, weight="bold")
    arrow(ax_d, (0.50, 0.67), (0.50, 0.58), color=MUTED)
    box(ax_d, (0.05, 0.39), 0.90, 0.17, "Also observed here\nDirect endpoint modelling from the same source\nmeasurements performs better.", face=BLUE_LIGHT, edge=BLUE, fontsize=6.25, weight="bold")
    ax_d.plot([0.08, 0.92], [0.31, 0.31], transform=ax_d.transAxes, color=GRID, linewidth=1.0)
    box(ax_d, (0.05, 0.07), 0.42, 0.16, "Not established\nBiological-age identity", face=RED_LIGHT, edge=RED, fontsize=6.5, weight="bold")
    box(ax_d, (0.53, 0.07), 0.42, 0.16, "Not established\nUniversal superiority of $X$", face=RED_LIGHT, edge=RED, fontsize=6.5, weight="bold")
    ax_d.text(0.50, -0.055, "Finite-sample result: one decoder, population, endpoint, panel and model family", transform=ax_d.transAxes, ha="center", fontsize=5.9, color=MUTED, fontstyle="italic")

    figures = save_figure(fig, package, stem)

    pooled_source = package / "source_data" / "panel_b_pooled_metrics.tsv"
    shutil.copyfile(pooled_path, pooled_source)
    contrast_source = package / "source_data" / "panel_c_paired_contrasts.tsv"
    shutil.copyfile(contrast_path, contrast_source)
    fold_source = package / "source_data" / "extended_fold_metrics.tsv"
    shutil.copyfile(folds_path, fold_source)
    concept_source = package / "source_data" / "panel_a_d_design_and_boundaries.tsv"
    pd.DataFrame(
        [
            ("a", "source_panel", "50 measurements", "same declared panel for decoder and direct endpoint model"),
            ("a", "decoder_target", "chronological age only", "mortality labels excluded from decoder training"),
            ("a", "validation", "nested five-fold cross-fitting", "fold-local decoder and endpoint models"),
            ("d", "supported", "clock utility", "held-out mortality value beyond age and background"),
            ("d", "supported", "source-model performance", "greater held-out value under declared model families"),
            ("d", "forbidden", "construct identity", "not identified"),
            ("d", "forbidden", "universal source superiority", "not estimated"),
        ],
        columns=["panel", "type", "item", "statement"],
    ).to_csv(concept_source, sep="\t", index=False)
    source_index = package / "source_data" / "source_data_index.tsv"
    pd.DataFrame(
        [
            ("a,d", concept_source.name, "design and claim-boundary map"),
            ("b", pooled_source.name, "pooled OOF endpoint metrics"),
            ("c", contrast_source.name, "paired fixed-prediction bootstrap contrasts"),
            ("extended", fold_source.name, "outer-fold metrics for diagnostics"),
        ],
        columns=["panel", "file", "content"],
    ).to_csv(source_index, sep="\t", index=False)

    legend = """# Figure 3 | An age-only decoder adds mortality-prediction value, while direct source-measurement modelling performs better

**a**, Nested cross-fitting in the complete-support NHANES sample. A fold-local linear decoder was trained only to reconstruct chronological age from background and the same 50 source measurements later available to the direct endpoint model. Mortality models used age and background alone ($M_{WA}$), added the out-of-fold clock ($M_{CLOCK}$), or used the same source measurements directly ($M_X$). **b**, Pooled out-of-fold log loss for five-year all-cause mortality among 13,724 participants (470 deaths). **c**, Paired out-of-fold improvements for the clock beyond age and background, the source measurements beyond the clock, and the source measurements beyond age and background. Intervals are paired fixed-prediction bootstrap intervals and omit model refitting and study-development uncertainty. **d**, Interpretation boundary. The clock's endpoint utility is genuine, while direct source-measurement modelling performs better under the declared estimators. This finite-sample comparison concerns one decoder, population, endpoint, 50-measurement panel and specified model families; it does not establish population conditional insufficiency, universal superiority of direct modelling or biological-age identity.

# 图 3｜纯年龄解码器增加死亡预测价值，而直接使用源测量建模表现更好

**a**，在具有完整五年行政随访支持的 NHANES 样本中进行嵌套交叉拟合。每个外层训练折内的线性解码器只以时序年龄为目标，使用的 50 项源测量与随后直接终点模型使用的测量完全相同。死亡模型依次为年龄加背景（$M_{WA}$）、加入折外时钟（$M_{CLOCK}$）以及直接使用同源测量（$M_X$）。**b**，13,724 名参与者（470 例死亡）的五年全因死亡折外 log loss。**c**，时钟相对年龄和背景的增量、源测量相对时钟的增量，以及源测量相对年龄和背景的增量。区间为固定折外预测上的配对 bootstrap 区间，不包含模型重拟合和研究开发不确定性。**d**，解释边界。时钟的终点效用是真实的；在已声明估计器下，直接使用同源测量建模表现更好。该有限样本比较仅限一个解码器、一个人群、一个终点、一个 50 项面板及指定模型族；它不建立总体条件不充分性、直接建模的普遍优势或生物年龄身份。
"""
    (package / "FIGURE_LEGEND_BILINGUAL.md").write_text(legend, encoding="utf-8")
    (package / "README.md").write_text(
        "# v6.5 Figure 3 NHANES age-decoder figure v0.2\n\nSubmission-facing, fit-free visualization of `nhanes_5y_age_decoder_source_information_probe_v0_2`. The script reads frozen metrics, contrasts, diagnostics and the frozen decision; it does not load or refit the persisted models and does not regenerate bootstrap indices.\n",
        encoding="utf-8",
    )
    write_common_visualization_files(
        package,
        "v6_5_figure3_nhanes_age_decoder_v0_2",
        {
            "analytical_question": "Can a pure chronological-age decoder have genuine endpoint utility without matching the held-out performance of direct endpoint modelling from the same source measurements?",
            "takeaway": "The clock improved five-year mortality prediction, while direct modelling of the same 50 measurements performed better under the declared finite-sample comparison.",
            "surface": "standalone static publication figure",
            "footprint": "180 mm wide, four panels, 600 dpi PNG plus editable SVG and vector PDF",
            "panels": {"a": "nested cross-fitting design", "b": "three-model log-loss comparison", "c": "paired predictive-value contrasts", "d": "finite-sample claim boundary"},
            "palette_policy": "hard two-root cap plus neutrals",
            "non_color_distinction": "ordered model labels, direct values, interval marks and boxed claim types",
            "data_grain": "13,724 participants; 470 deaths; five outer folds",
        },
        [SCRIPT, pooled_path, contrast_path, folds_path, diagnostics_path, decision_path],
        figures,
        [pooled_source, contrast_source, fold_source, concept_source, source_index],
    )
    return figures


def build_figure4() -> dict[str, Path]:
    package = PACKAGES["figure4"]
    ensure_package(package)
    stem = "Figure_4_endpoint_directed_acquisition_and_estimand_specific_replacement"
    domain_path = ACQ / "source_target_domain_value.tsv"
    policies_path = ACQ / "target_policy_point_estimates.tsv"
    intervals_path = ACQ / "target_policy_bootstrap.tsv"
    pairwise_path = ACQ / "target_policy_pairwise_bootstrap.tsv"
    folds_path = ACQ / "source_selected_target_folds.tsv"
    domain = pd.read_csv(domain_path, sep="\t")
    policies = pd.read_csv(policies_path, sep="\t")
    intervals = pd.read_csv(intervals_path, sep="\t")
    pairwise = pd.read_csv(pairwise_path, sep="\t")
    folds = pd.read_csv(folds_path, sep="\t")

    require(len(domain) == 5, "MTH-01 domain menu is no longer five categories.")
    endpoint_domain = policies.loc[policies["policy"].eq("source_endpoint_value"), "selected_domain"].iloc[0]
    age_domain = policies.loc[policies["policy"].eq("source_age_indexability"), "selected_domain"].iloc[0]
    require(endpoint_domain == "hepatic_protein_injury" and age_domain == "hemodynamic", "Frozen source selections changed.")
    require((folds["measurement_risk_gain_V"] > 0).all() and len(folds) == 5, "Expected source-selected fold pattern not recovered.")

    fig = plt.figure(figsize=(7.15, 6.0), facecolor=WHITE)
    grid = fig.add_gridspec(2, 2, left=0.075, right=0.985, bottom=0.065, top=0.96, hspace=0.42, wspace=0.33)
    ax_a = fig.add_subplot(grid[0, 0])
    ax_b = fig.add_subplot(grid[0, 1])
    ax_c = fig.add_subplot(grid[1, 0])
    ax_d = fig.add_subplot(grid[1, 1])

    panel_label(ax_a, "a", "Endpoint value versus forward age indexability")
    style_axis(ax_a, grid_axis="x")
    d = domain.sort_values("source_V", ascending=True).copy()
    y = np.arange(len(d))
    colors = []
    for name in d["domain"]:
        if name == endpoint_domain:
            colors.append(ORANGE)
        elif name == age_domain:
            colors.append(PURPLE)
        else:
            colors.append("#9FA7AE")
    ax_a.barh(y, d["source_V"], height=0.58, color=colors, edgecolor=INK, linewidth=0.4)
    ax_a.set_yticks(y, d["domain_label"])
    ax_a.set_xlabel("Source-cycle five-year mortality log-loss value", fontsize=6.2, color=INK)
    ax_a.set_xlim(0, d["source_V"].max() * 1.26)
    for yi, (_, row), color in zip(y, d.iterrows(), colors):
        tag = "endpoint-selected" if row["domain"] == endpoint_domain else ("forward age-indexability selected" if row["domain"] == age_domain else "")
        ax_a.text(row["source_V"] + 0.00018, yi, f"{row['source_V']:.3f}" + (f"  {tag}" if tag else ""), va="center", fontsize=5.8, color=color if tag else MUTED, fontweight="bold" if tag else "normal")
    ax_a.text(0.01, -0.22, "Five prespecified measurement-domain options; feature counts differ and no cost equivalence was assumed.", transform=ax_a.transAxes, ha="left", fontsize=5.7, color=MUTED)

    panel_label(ax_b, "b", "Frozen policies in the target cycle")
    style_axis(ax_b, grid_axis="x")
    keep = ["source_endpoint_value", "source_age_indexability", "target_local_nested", "uniform_random_category", "target_oracle"]
    interval_index = intervals.set_index("policy").loc[keep]
    labels = ["Source endpoint-informed", "Source forward age-indexability", "Target nested-local", "Uniform random domain", "Target oracle (descriptive)"]
    y_b = np.arange(len(keep))[::-1]
    colors_b = [ORANGE, PURPLE, TEAL, "#7D8791", "#AAB1B7"]
    for yi, (_, row), color in zip(y_b, interval_index.iterrows(), colors_b):
        ax_b.plot([row["ci_low"], row["ci_high"]], [yi, yi], color=color, linewidth=2.0, solid_capstyle="round")
        ax_b.scatter(row["estimate"], yi, s=38, color=color, edgecolor=INK, linewidth=0.45, zorder=3)
    ax_b.axvline(0, color=MUTED, linewidth=0.9)
    ax_b.set_yticks(y_b, labels)
    ax_b.set_xlabel("Target-cycle log-loss gain beyond age and background", fontsize=6.2, color=INK)
    ax_b.set_xlim(-0.0018, 0.015)
    ax_b.text(0.99, -0.22, "Fixed-OOF event-stratified paired bootstrap (5,000 resamples)", transform=ax_b.transAxes, ha="right", fontsize=5.7, color=MUTED)

    panel_label(ax_c, "c", "Transferred advantage and fold consistency")
    style_axis(ax_c, grid_axis="y")
    fold_y = folds["measurement_risk_gain_V"].to_numpy(float)
    x = np.arange(1, 6)
    ax_c.bar(x, fold_y, color=ORANGE_LIGHT, edgecolor=ORANGE, linewidth=1.0, width=0.62)
    ax_c.scatter(x, fold_y, s=22, facecolor=ORANGE, edgecolor=INK, linewidth=0.35, zorder=3)
    ax_c.axhline(0, color=MUTED, linewidth=0.8)
    ax_c.set_xticks(x, [f"Fold {i}" for i in x])
    ax_c.set_ylabel("Target log-loss gain", fontsize=6.2, color=INK)
    pair = pairwise.loc[pairwise["contrast"].eq("source_endpoint_value_minus_source_age_indexability")].iloc[0]
    ax_c.set_ylim(0, max(fold_y.max() * 1.34, 0.0120))
    ax_c.text(0.50, 0.97, "Endpoint-informed minus forward age-indexability", transform=ax_c.transAxes, ha="center", va="top", fontsize=6.1, color=INK, fontweight="bold", bbox=dict(facecolor=WHITE, edgecolor="none", pad=1.5))
    ax_c.text(0.50, 0.86, f"paired difference {pair['estimate']:.3f}; 95% interval {pair['ci_low']:.3f} to {pair['ci_high']:.3f}", transform=ax_c.transAxes, ha="center", va="top", fontsize=5.8, color=ORANGE, fontweight="bold", bbox=dict(facecolor=WHITE, edgecolor="none", pad=1.2))

    ax_d.set_axis_off()
    panel_label(ax_d, "d", "Target-first replacement map")
    objects = [
        ("Measured state", "Declared measurement\nmodel and state definition", BLUE_LIGHT, BLUE),
        ("Named risk", "Endpoint, horizon, loss\nand validation population", TEAL_LIGHT, TEAL),
        ("Transition", "Longitudinal sampling\nand transition model", PURPLE_LIGHT, PURPLE),
        ("Intervention\ndisplacement", "Perturbation contrast\nand measurement response", ORANGE_LIGHT, ORANGE),
        ("Benefit", "Causal estimand, outcome\nand decision context", GOLD_LIGHT, GOLD),
    ]
    y0 = 0.80
    for i, (obj, structure, face, edge) in enumerate(objects):
        y = y0 - i * 0.18
        box(ax_d, (0.03, y), 0.34, 0.12, obj, face=face, edge=edge, fontsize=6.4, weight="bold")
        arrow(ax_d, (0.38, y + 0.06), (0.48, y + 0.06), color=edge)
        box(ax_d, (0.49, y), 0.48, 0.12, structure, face=WHITE, edge=edge, fontsize=6.0, weight="bold")
    ax_d.text(0.50, 0.015, "Each claim inherits meaning from its own declared object and evidence — not from an age unit.", transform=ax_d.transAxes, ha="center", fontsize=5.9, color=RED, fontweight="bold")

    figures = save_figure(fig, package, stem)

    domain_source = package / "source_data" / "panel_a_domain_values.tsv"
    shutil.copyfile(domain_path, domain_source)
    policy_source = package / "source_data" / "panel_b_policy_intervals.tsv"
    shutil.copyfile(intervals_path, policy_source)
    pair_source = package / "source_data" / "panel_c_policy_pairwise_intervals.tsv"
    shutil.copyfile(pairwise_path, pair_source)
    folds_source = package / "source_data" / "panel_c_source_selected_folds.tsv"
    shutil.copyfile(folds_path, folds_source)
    replacement_source = package / "source_data" / "panel_d_estimand_replacement_map.tsv"
    pd.DataFrame(objects, columns=["object", "identifying_structure", "facecolor", "edgecolor"]).drop(columns=["facecolor", "edgecolor"]).to_csv(replacement_source, sep="\t", index=False)
    source_index = package / "source_data" / "source_data_index.tsv"
    pd.DataFrame(
        [
            ("a", domain_source.name, "source and target values for the five prespecified domains"),
            ("b", policy_source.name, "target-cycle policy gains and fixed-OOF intervals"),
            ("c", pair_source.name, "paired policy contrasts"),
            ("c", folds_source.name, "fold-specific source-endpoint-selected gain"),
            ("d", replacement_source.name, "estimand-specific replacement map"),
        ],
        columns=["panel", "file", "content"],
    ).to_csv(source_index, sep="\t", index=False)

    legend = """# Figure 4 | Endpoint-directed evidence guides what to measure and what to claim

**a**, Five-year mortality log-loss value of five prespecified measurement-domain options in NHANES 2007–2008. Endpoint information selected hepatic, protein and injury measurements, whereas forward age indexability selected haemodynamic measurements. Domains differed in feature count, and no equivalence of monetary resources or real-world costs was assumed. **b**, Target-cycle gain beyond age and background for source-frozen and comparator policies in NHANES 2009–2010. Intervals are event-stratified paired bootstraps of fixed out-of-fold losses with 5,000 resamples; the target oracle is descriptive. **c**, The endpoint-informed policy exceeded the forward age-indexability policy, and the source-selected domain had positive target gain in all five outer folds. **d**, Target-first replacement map: measured state, named risk, transition, intervention displacement and benefit require their own declared measurement, endpoint, longitudinal or causal structures. The acquisition analysis is exploratory cross-cycle evidence within one five-domain menu; it is not a comparison with an inverse age decoder, clinical net benefit, optimization under a shared monetary budget, untouched external confirmation or biological-age identification.

# 图 4｜终点导向的证据指导测量选择与主张边界

**a**，在 NHANES 2007–2008 中，五个预设等类别成本测量域对五年死亡 log loss 的价值。终点信息选择了肝脏／蛋白／损伤域，而前向年龄可索引性选择了血流动力学域。“等类别成本”不等于等预算、等变量数或已经证明的真实成本等价。**b**，冻结于来源周期的策略及比较策略在 NHANES 2009–2010 中相对年龄和背景的增量。区间为固定折外损失上的事件分层配对 bootstrap（5,000 次）；目标周期 oracle 仅作描述。**c**，终点导向策略优于前向年龄可索引性策略，且来源周期所选域在五个外层折中的目标增量均为正。**d**，target-first 替代图：测量状态、命名风险、转变、干预位移与获益，各自需要明确的测量、终点、纵向或因果结构。该 acquisition 结果是在一个五域菜单中的探索性 NHANES 跨周期证据；它不是与逆向年龄解码器的比较、临床净获益、等预算优化、未查看外部确认或生物年龄识别。
"""
    (package / "FIGURE_LEGEND_BILINGUAL.md").write_text(legend, encoding="utf-8")
    (package / "README.md").write_text(
        "# v6.4 Figure 4 target-first figure v0.1\n\nFit-free relabelling of the frozen MTH-01 acquisition result with an estimand-specific replacement map. The comparison is against forward age indexability, not an inverse age decoder. No bootstrap indices are regenerated and no model is loaded or fitted.\n",
        encoding="utf-8",
    )
    write_common_visualization_files(
        package,
        "v6_4_figure4_target_first_v0_1",
        {
            "analytical_question": "Can endpoint-directed evidence guide acquisition more effectively than a forward age-indexability control, and how should stronger ageing claims be replaced by object-specific estimands?",
            "takeaway": "In the declared five-domain menu the source endpoint rule transferred more mortality-predictive value than the forward age-indexability control; object-specific claims require object-specific structures.",
            "surface": "standalone static publication figure",
            "footprint": "180 mm wide, four panels, 600 dpi PNG plus editable SVG and vector PDF",
            "panels": {"a": "source domain values and frozen selections", "b": "target-cycle policy comparison", "c": "paired advantage and folds", "d": "estimand replacement map"},
            "palette_policy": "hard two-root cap for empirical comparisons; restrained multi-category roots for the five-object replacement map",
            "non_color_distinction": "direct policy labels, interval marks, bar outlines and paired object-structure boxes",
            "data_grain": "five prespecified measurement-domain options; source 2007–2008 and target 2009–2010 NHANES cycles",
        },
        [SCRIPT, domain_path, policies_path, intervals_path, pairwise_path, folds_path],
        figures,
        [domain_source, policy_source, pair_source, folds_source, replacement_source, source_index],
    )
    return figures


def write_index(figure1: dict[str, Path], figure3: dict[str, Path]) -> None:
    INDEX_PACKAGE.mkdir(parents=True, exist_ok=True)
    figure1_png = figure1["png"]
    hsct_png = HSCT_PACKAGE / "figures" / "Figure_HSCT_age_reconstruction_semantic_counterexample.png"
    hsct_svg = HSCT_PACKAGE / "figures" / "Figure_HSCT_age_reconstruction_semantic_counterexample.svg"
    hsct_pdf = HSCT_PACKAGE / "figures" / "Figure_HSCT_age_reconstruction_semantic_counterexample.pdf"
    figure3_png = figure3["png"]
    figure4_png = PACKAGES["figure4"] / "figures" / "Figure_4_endpoint_directed_acquisition_and_estimand_specific_replacement.png"
    require(figure1_png.exists(), "Current Figure 1 resource is missing.")
    require(hsct_png.exists() and hsct_svg.exists() and hsct_pdf.exists(), "Latest HSCT figure package is incomplete.")
    require(figure3_png.exists(), "Current Figure 3 resource is missing.")
    require(figure4_png.exists(), "Current Figure 4 resource is missing.")
    rows = [
        ("Figure 1", "REVISE", figure1_png, "fit-free constrained-orientation and Bayes-risk clarification"),
        ("Figure 2", "RELABEL", hsct_png, "fit-free standardization of the interpretation boundary; HUM-01 estimand and values unchanged"),
        ("Figure 3", "RELABEL", figure3_png, "fit-free terminology standardization; NHANES v0.2 estimand and values unchanged"),
        ("Figure 4", "REUSE", figure4_png, "unchanged v6.4 target-first resource"),
    ]
    index = pd.DataFrame(
        [
            {
                "figure": fig,
                "action": action,
                "primary_png": path.relative_to(ROOT).as_posix(),
                "sha256": sha256(path),
                "rationale": rationale,
            }
            for fig, action, path, rationale in rows
        ]
    )
    index.to_csv(INDEX_PACKAGE / "FIGURE_INDEX.tsv", sep="\t", index=False)
    text = "# v6.5 P1 main-figure resource audit and sync\n\n"
    text += "| Figure | Action | Submission resource | Rationale |\n| --- | --- | --- | --- |\n"
    for fig, action, path, rationale in rows:
        text += f"| {fig} | {action} | `{path.relative_to(ROOT).as_posix()}` | {rationale} |\n"
    text += "\nRELABEL resources are deterministic, fit-free visualizations of frozen results. No model, randomization, statistic or analysis interface was changed.\n"
    (INDEX_PACKAGE / "FIGURE_INDEX.md").write_text(text, encoding="utf-8")
    write_json(
        INDEX_PACKAGE / "run_summary.json",
        {
            "status": "PASS",
            "fit_free": True,
            "models_refit": False,
            "random_numbers_generated": False,
            "frozen_inputs_mutated": False,
            "figure1_revised_package": PACKAGES["figure1"].relative_to(ROOT).as_posix(),
            "figure2_relabelled_package": HSCT_PACKAGE.relative_to(ROOT).as_posix(),
            "figure3_relabelled_package": PACKAGES["figure3"].relative_to(ROOT).as_posix(),
            "figure4_reused_package": PACKAGES["figure4"].relative_to(ROOT).as_posix(),
            "figure3_frozen_analysis": NHANES.relative_to(ROOT).as_posix(),
        },
    )


def main() -> None:
    figure1 = build_figure1()
    print(f"PASS: built v6.9 precision-revised Figure 1 at {figure1['png']}.")


if __name__ == "__main__":
    main()
```
