## Supplementary_Software_1 for "Inferential boundaries of age prediction: why prediction does not establish biological age measurement": plot_v6_16_extended_data_figure5_layout_v0_1.py

```
#!/usr/bin/env python
"""Re-render frozen Extended Data Figure 5 with corrected label clearance."""

from __future__ import annotations

import importlib.util
import json
import shutil
import sys
from pathlib import Path

sys.dont_write_bytecode = True
ROOT = Path(__file__).resolve().parents[1]
PREDECESSOR_SCRIPT = ROOT / "analysis" / "plot_v6_9_extended_data_figure4_domain_options_v0_1.py"
PREDECESSOR = ROOT / "outputs" / "v6_9_extended_data_figure4_domain_options_v0_1"
OUTPUT = ROOT / "outputs" / "v6_16_extended_data_figure5_layout_v0_1"
STEM = "Extended_Data_Figure_5_five_domain_acquisition_comparison"

spec = importlib.util.spec_from_file_location("frozen_ed5_plot", PREDECESSOR_SCRIPT)
assert spec is not None and spec.loader is not None
old = importlib.util.module_from_spec(spec)
spec.loader.exec_module(old)


def main() -> None:
    """Reuse only frozen tabular source and deterministic plotting primitives."""
    for name in ("figures", "source_data", "visualization"):
        (OUTPUT / name).mkdir(parents=True, exist_ok=True)
    old.require((OUTPUT / "README.md").exists(), "Chart contract must precede rendering.")
    data = old.load_frozen_source()
    source_rows = []
    for source in sorted((PREDECESSOR / "source_data").glob("*.tsv")):
        target_file = OUTPUT / "source_data" / source.name
        shutil.copy2(source, target_file)
        old.require(old.sha256(source) == old.sha256(target_file), f"Source copy changed: {source.name}")
        old.pd.testing.assert_frame_equal(
            old.pd.read_csv(source, sep="\t"),
            old.pd.read_csv(target_file, sep="\t"),
            check_exact=True,
        )
        source_rows.append({"file": source.name, "sha256": old.sha256(source), "exact_copy": True})
    old.require(len(source_rows) == 5, "Expected four source tables and their index.")
    for key, filename in {
        "policy": "panel_a_frozen_policy_definitions.tsv",
        "target": "panel_b_target_policy_intervals.tsv",
        "pair": "panel_c_pooled_policy_difference.tsv",
        "folds": "panel_c_fold_policy_differences.tsv",
    }.items():
        old.pd.testing.assert_frame_equal(
            data[key], old.pd.read_csv(OUTPUT / "source_data" / filename, sep="\t"), check_exact=True
        )

    target = data["target"].set_index("policy")
    pair = data["pair"].iloc[0]
    folds = data["folds"]
    fig = old.plt.figure(figsize=(7.15, 4.65), facecolor=old.WHITE)
    grid = fig.add_gridspec(
        2, 2, left=0.065, right=0.985, bottom=0.075, top=0.955,
        height_ratios=[1.05, 1.25], hspace=0.42, wspace=0.34,
    )
    ax_a = fig.add_subplot(grid[0, 0])
    ax_b = fig.add_subplot(grid[0, 1])
    ax_c = fig.add_subplot(grid[1, :])
    # Keep panel a and the entire lower panel fixed; reserve extra space for b labels.
    pos_b = ax_b.get_position()
    ax_b.set_position([pos_b.x0 + 0.055, pos_b.y0, pos_b.width - 0.055, pos_b.height])

    ax_a.set_axis_off()
    old.panel_label(ax_a, "a", "Two frozen source policies in a five-domain menu")
    old.box(ax_a, (0.13, 0.78), 0.74, 0.13, "Five prespecified measurement-domain options\nNHANES 2007-2008 ($n=4{,}599$)", face=old.LIGHT, edge=old.MUTED, fontsize=6.25, weight="bold")
    old.arrow(ax_a, (0.50, 0.77), (0.26, 0.64), color=old.ORANGE)
    old.arrow(ax_a, (0.50, 0.77), (0.74, 0.64), color=old.BLUE)
    old.box(ax_a, (0.01, 0.42), 0.47, 0.20, "Endpoint-directed\nmaximize held-out mortality\nlog-loss gain", face=old.ORANGE_LIGHT, edge=old.ORANGE, fontsize=6.05, weight="bold")
    # Four-line wrapping receives local vertical padding; no words or font size change.
    inverse_box = old.box(ax_a, (0.52, 0.40), 0.47, 0.24, "Inverse age decoder\nminimize held-out\nchronological-age\nmean-squared error", face=old.BLUE_LIGHT, edge=old.BLUE, fontsize=5.95, weight="bold")
    inverse_text = ax_a.texts[-1]
    old.arrow(ax_a, (0.245, 0.41), (0.245, 0.32), color=old.ORANGE)
    old.arrow(ax_a, (0.755, 0.39), (0.755, 0.32), color=old.BLUE)
    old.box(ax_a, (0.01, 0.08), 0.47, 0.22, "Selected\nHepatic / protein / injury", face=old.WHITE, edge=old.ORANGE, fontsize=6.1, weight="bold")
    selected_box = old.box(ax_a, (0.52, 0.08), 0.47, 0.22, "Selected\nMetabolic / adiposity / lipid", face=old.WHITE, edge=old.BLUE, fontsize=6.1, weight="bold")
    ax_a.text(0.50, 0.005, "Different objectives and units; selections are not ranked on one source axis.", transform=ax_a.transAxes, ha="center", fontsize=5.55, color=old.RED, fontweight="bold")

    old.panel_label(ax_b, "b", "Target-cycle endpoint value\nof the frozen domains")
    old.style_axis(ax_b, grid_axis="x")
    order = ["source_endpoint_policy", "source_inverse_age_decoder_policy"]
    y = old.np.array([1, 0])
    labels = ["Endpoint-directed\nHepatic / protein /\ninjury", "Inverse age decoder\nMetabolic / adiposity /\nlipid"]
    for yi, key, color, marker in zip(y, order, [old.ORANGE, old.BLUE], ["o", "D"]):
        row = target.loc[key]
        ax_b.plot([row["ci_low"], row["ci_high"]], [yi, yi], color=color, linewidth=2.2, solid_capstyle="round")
        ax_b.scatter(row["estimate"], yi, s=45, marker=marker, facecolor=color if key == order[0] else old.WHITE, edgecolor=color, linewidth=1.2, zorder=3)
        ax_b.text(row["ci_high"] + 0.00028, yi, f"{row['estimate']:.4f}", va="center", fontsize=6.0, color=color, fontweight="bold")
    ax_b.axvline(0, color=old.MUTED, linewidth=0.9)
    ax_b.set_yticks(y, labels)
    ax_b.set_ylim(-0.65, 1.65)
    ax_b.set_xlim(-0.0025, 0.0125)
    ax_b.set_xlabel("Five-year mortality log-loss gain\nbeyond age and background", fontsize=6.15, color=old.INK)
    ax_b.text(0.99, 0.97, "NHANES 2009-2010\n$n=4{,}896$; 157 deaths\n95% fixed-prediction bootstrap; 5,000 resamples", transform=ax_b.transAxes, ha="right", va="top", fontsize=5.7, color=old.MUTED, linespacing=1.2)

    # Panel c is inherited without geometric or semantic changes.
    old.panel_label(ax_c, "c", "Paired endpoint-directed minus inverse-age difference")
    old.style_axis(ax_c, grid_axis="x")
    fold_values = folds["endpoint_minus_inverse"].to_numpy(float)
    y_rows = old.np.arange(6)[::-1]
    labels_c = ["Pooled (95% CI)"] + [f"Fold {i}" for i in range(1, 6)]
    ax_c.plot([pair["ci_low"], pair["ci_high"]], [y_rows[0], y_rows[0]], color=old.NAVY, linewidth=2.5, solid_capstyle="round")
    ax_c.scatter(pair["estimate"], y_rows[0], s=52, marker="D", facecolor=old.ORANGE, edgecolor=old.NAVY, linewidth=0.7, zorder=4)
    for yi, value in zip(y_rows[1:], fold_values):
        ax_c.scatter(value, yi, s=36, marker="o", facecolor=old.ORANGE if value > 0 else old.WHITE, edgecolor=old.ORANGE, linewidth=1.1, zorder=3)
    ax_c.axvline(0, color=old.MUTED, linewidth=0.9)
    ax_c.set_yticks(y_rows, labels_c)
    ax_c.set_ylim(-0.65, 5.65)
    ax_c.set_xlim(-0.0030, 0.0126)
    ax_c.set_xlabel("Difference in target log-loss gain (positive favours endpoint-directed selection)", fontsize=6.15, color=old.INK)
    ax_c.text(0.99, 0.96, f"pooled {pair['estimate']:.4f}  |  95% CI {pair['ci_low']:.4f} to {pair['ci_high']:.4f}  |  4/5 folds positive", transform=ax_c.transAxes, ha="right", va="top", fontsize=5.95, color=old.NAVY, fontweight="bold", bbox=dict(facecolor=old.WHITE, edgecolor="none", pad=1.2))

    fig.canvas.draw()
    renderer = fig.canvas.get_renderer()
    leftmost_b_label = min(label.get_window_extent(renderer).x0 for label in ax_b.get_yticklabels())
    panel_a_right = selected_box.get_window_extent(renderer).x1
    clearance = leftmost_b_label - panel_a_right
    old.require(clearance > 5, f"Panel a/b labels still collide: {clearance:.3f}px")
    text_bounds = inverse_text.get_window_extent(renderer)
    box_bounds = inverse_box.get_window_extent(renderer)
    old.require(text_bounds.x0 >= box_bounds.x0 and text_bounds.x1 <= box_bounds.x1, "Wrapped objective exceeds box horizontally.")
    old.require(text_bounds.y0 >= box_bounds.y0 and text_bounds.y1 <= box_bounds.y1, "Wrapped objective exceeds box vertically.")

    figures = {}
    for extension in ("png", "svg", "pdf"):
        path = OUTPUT / "figures" / f"{STEM}.{extension}"
        extra = {"dpi": 600} if extension == "png" else {}
        fig.savefig(path, bbox_inches="tight", pad_inches=0.04, facecolor=old.WHITE, **extra)
        figures[extension] = str(path)
    old.plt.close(fig)
    report = {
        "status": "PENDING_VISUAL_QA",
        "source_data_exact": source_rows,
        "all_four_plotted_tables_equal_predecessor": True,
        "label_clearance_at_100dpi_px": clearance,
        "wrapped_objective_within_box": True,
        "models_loaded": False,
        "models_fitted": False,
        "random_numbers_generated": False,
        "new_analysis": False,
        "predecessor_script": str(PREDECESSOR_SCRIPT),
        "figures": figures,
    }
    old.write_json(OUTPUT / "visualization" / "VALIDATION_REPORT.json", report)
    print(json.dumps(report, indent=2))


if __name__ == "__main__":
    main()
```
