## Supplementary_Software_1 for "Inferential boundaries of age prediction: why prediction does not establish biological age measurement": plot_v6_22_main_figure_assembly_v0_1.py

```
#!/usr/bin/env python
"""Fit-free Figure 3/4 assembly from frozen aggregate figure source tables."""

from __future__ import annotations

import hashlib
import argparse
import json
import shutil
import sys
from pathlib import Path

import matplotlib
matplotlib.use("Agg")
import matplotlib.pyplot as plt
from matplotlib.patches import FancyArrowPatch, FancyBboxPatch
from matplotlib.ticker import FixedLocator, FormatStrFormatter
import pandas as pd


ROOT = Path(__file__).resolve().parents[1]
OUT = ROOT / "outputs" / "v6_22_main_figure_assembly_v0_1"
OLD3 = ROOT / "outputs" / "nhanes_hum03_hum04_hum06_figure3_successor_v0_3" / "source_data"
OLD4 = ROOT / "outputs" / "v6_8_figure4_target_first_framework_v0_1" / "source_data"
ED5 = ROOT / "outputs" / "v6_16_extended_data_figure5_layout_v0_1" / "source_data"

INK = "#202124"
MUTED = "#62676D"
GRID = "#D9DDE1"
LIGHT = "#EEF1F3"
WHITE = "#FFFFFF"
NAVY = "#18243D"
BLUE = "#315F85"
BLUE_LIGHT = "#E5EEF5"
TEAL = "#087E8B"
TEAL_LIGHT = "#D7EEF0"
ORANGE = "#D97732"
ORANGE_LIGHT = "#F7E6D8"

matplotlib.rcParams.update({
    "font.family": "sans-serif", "font.sans-serif": ["Arial", "DejaVu Sans"],
    "font.size": 7, "axes.unicode_minus": False, "svg.fonttype": "none",
    "pdf.fonttype": 42, "ps.fonttype": 42,
})

INPUTS = [
    ("fig3_a_design", "3a", OLD3 / "panel_a_continuous_nhanes_design.tsv"),
    ("fig3_b_continuous_contrasts", "3b", OLD3 / "panel_b_continuous_nhanes_predictive_contrasts.tsv"),
    ("fig3_d_matched_scalar_contrasts", "3c,d", OLD3 / "panel_c_nhanes3_matched_scalar_target_supervision.tsv"),
    ("fig4_b_source_policies", "4b", ED5 / "panel_a_frozen_policy_definitions.tsv"),
    ("fig4_b_policy_intervals", "4b", ED5 / "panel_b_target_policy_intervals.tsv"),
    ("fig4_b_paired_difference", "4b", ED5 / "panel_c_pooled_policy_difference.tsv"),
    ("fig4_c_replacement_map", "4c", OLD4 / "panel_c_estimand_specific_replacement_map.tsv"),
]

BOX_RECORDS: list[tuple[plt.Figure, object, object]] = []


def require(value: bool, message: str) -> None:
    if not value:
        raise RuntimeError(message)


def sha256(path: Path) -> str:
    return hashlib.sha256(path.read_bytes()).hexdigest()


def load_inputs(supplied_only: bool = False) -> tuple[dict[str, pd.DataFrame], list[dict[str, object]]]:
    require((OUT / "README.md").exists(), "Chart contract must precede rendering")
    (OUT / "source_data").mkdir(parents=True, exist_ok=True)
    (OUT / "figures").mkdir(exist_ok=True)
    index_path = OUT / "source_data" / "source_data_index.tsv"
    saved_index = pd.read_csv(index_path, sep="\t").set_index("file") if index_path.exists() else None
    result, records = {}, []
    for key, panel, path in INPUTS:
        figure_dir = "Figure_3" if key.startswith("fig3_") else "Figure_4"
        target = OUT / "source_data" / figure_dir / f"{key}.tsv"
        target.parent.mkdir(exist_ok=True)
        relative_target = target.relative_to(OUT / "source_data").as_posix()
        source_available = path.exists() and not supplied_only
        if not target.exists():
            require(source_available, f"Missing source and supplied copy: {target}")
            shutil.copy2(path, target)
        copied_frame = pd.read_csv(target, sep="\t")
        if source_available:
            source_frame = pd.read_csv(path, sep="\t")
            pd.testing.assert_frame_equal(source_frame, copied_frame, check_exact=True)
            require(sha256(path) == sha256(target), f"Copy changed bytes: {key}")
        else:
            require(saved_index is not None and relative_target in saved_index.index,
                    f"Frozen-copy manifest missing: {relative_target}")
            require(sha256(target) == saved_index.loc[relative_target, "sha256"],
                    f"Supplied aggregate copy differs from frozen index: {relative_target}")
        result[key] = copied_frame
        records.append({"panels": panel, "file": relative_target,
                        "frozen_source": path.relative_to(ROOT).as_posix(),
                        "rows": len(copied_frame), "sha256": sha256(target), "byte_exact": True})
    pd.DataFrame(records).to_csv(OUT / "source_data" / "source_data_index.tsv", sep="\t", index=False)
    require(list(result["fig3_b_continuous_contrasts"].contrast_id) == ["V_CLOCK", "V_SOURCE", "V_SOURCE_FLEX"], "Continuous contrast order changed")
    require(list(result["fig3_d_matched_scalar_contrasts"].contrast_id) == ["U_A", "Delta"], "Matched scalar order changed")
    require(len(result["fig4_b_policy_intervals"]) == 2, "Expected two source-frozen policies")
    require(len(result["fig4_b_paired_difference"]) == 1, "Expected one paired difference")
    require(len(result["fig4_c_replacement_map"]) == 5, "Expected five target-first objects")
    for key, columns in [("fig3_b_continuous_contrasts", ("ci_lower", "estimate", "ci_upper")),
                         ("fig3_d_matched_scalar_contrasts", ("ci_lower", "estimate", "ci_upper")),
                         ("fig4_b_policy_intervals", ("ci_low", "estimate", "ci_high")),
                         ("fig4_b_paired_difference", ("ci_low", "estimate", "ci_high"))]:
        lo, est, hi = columns
        require((result[key][lo] <= result[key][est]).all() and (result[key][est] <= result[key][hi]).all(), f"Invalid interval in {key}")
    return result, records


def text(ax, x, y, value, size=7, **kw):
    return ax.text(x, y, value, transform=ax.transAxes, color=kw.pop("color", INK), fontsize=size, **kw)


def panel(fig, rect, letter, title, subtitle=None):
    ax = fig.add_axes(rect)
    ax.set_axis_off()
    text(ax, 0, 1.02, letter, size=10, weight="bold", va="bottom")
    text(ax, 0.06, 1.025, title, size=8.1, weight="bold", va="bottom")
    if subtitle:
        text(ax, 0.06, .975, subtitle, size=6.35, color=MUTED, va="top", linespacing=1.2)
    return ax


def box(ax, x, y, width, height, label, edge=MUTED, face=WHITE, size=6.6, bold=False):
    patch = FancyBboxPatch((x, y), width, height, boxstyle="round,pad=0.009,rounding_size=0.008",
                          transform=ax.transAxes, facecolor=face, edgecolor=edge, linewidth=.9, clip_on=False)
    ax.add_patch(patch)
    label_obj = text(ax, x + width / 2, y + height / 2, label, size=size, ha="center", va="center",
                     weight="bold" if bold else "normal", linespacing=1.17)
    BOX_RECORDS.append((ax.figure, patch, label_obj))
    return patch


def arrow(ax, xy0, xy1, color=MUTED):
    ax.add_patch(FancyArrowPatch(xy0, xy1, transform=ax.transAxes, arrowstyle="-|>",
                                mutation_scale=7, color=color, linewidth=.8, shrinkA=1, shrinkB=1))


def plot_rows(ax, rows, labels, colors, markers, *, low="ci_lower", high="ci_upper", xmax=.014, footer=None):
    n = len(rows)
    ax.set_axis_on()
    ax.set_xlim(-.0015, xmax)
    ax.set_ylim(-.55, n - .02)
    ax.spines[["left", "right", "top"]].set_visible(False)
    ax.spines["bottom"].set_color(MUTED)
    ax.set_yticks([])
    ax.xaxis.set_major_locator(FixedLocator([0, .004, .008, .012]))
    ax.xaxis.set_major_formatter(FormatStrFormatter("%.3f"))
    ax.tick_params(axis="x", labelsize=6, length=3, color=MUTED)
    ax.grid(axis="x", color=GRID, linewidth=.55, zorder=-1)
    ax.axvline(0, color=MUTED, linewidth=.8, linestyle=(0, (3, 2)), zorder=0)
    for k, ((_, row), label, color, marker) in enumerate(zip(rows.iterrows(), labels, colors, markers)):
        y = n - 1 - k
        # Row labels are kept inside this panel rather than protruding into its neighbour.
        ax.text(.015, (y + .44 + .55) / (n + .53), label, transform=ax.transAxes,
                fontsize=6.35, color=INK, va="center", linespacing=1.13,
                bbox={"facecolor": WHITE, "edgecolor": "none", "pad": .5})
        ax.plot([row[low], row[high]], [y, y], color=color, linewidth=1.65, solid_capstyle="round")
        ax.scatter(row.estimate, y, s=30, marker=marker, facecolor=WHITE if marker == "o" else color,
                   edgecolor=color, linewidth=1.15, zorder=3)
        ax.text(row[high] + .00035, y, f"{row.estimate:.4f}", fontsize=6.1, color=color, weight="bold", va="center")
    ax.set_xlabel("Five-year mortality log-loss improvement", fontsize=6.5, labelpad=4)
    if footer:
        text(ax, 0, -.205, footer, size=5.8, color=MUTED, va="top", linespacing=1.2)


def figure3(data):
    fig = plt.figure(figsize=(7.15, 6.4), facecolor=WHITE)
    a = panel(fig, [.04, .60, .425, .32], "a", "Continuous NHANES: source comparison",
              "2007–2012 · 13,724 participants · 470 deaths")
    box(a, .025, .70, .95, .16, "Same 50 measurements + background", BLUE, BLUE_LIGHT, bold=True)
    arrow(a, (.50, .69), (.50, .605), TEAL)
    box(a, .13, .435, .74, .16, "Fold-local ridge decoder → age score\nTrained only to reconstruct chronological age", TEAL, TEAL_LIGHT, 6.4, True)
    text(a, .5, .374, "Mortality comparison in the same five outer folds", size=6.1, color=MUTED, ha="center")
    for y, label, color, face in [(.245, "$M_{WA}$   age + background", MUTED, LIGHT),
                                  (.115, "$M_{CLOCK}$   age + background + age score", TEAL, TEAL_LIGHT),
                                  (-.015, "$M_X$   age + background + 50 measurements", BLUE, BLUE_LIGHT)]:
        box(a, .04, y, .92, .095, label, color, face, 6.3)
    text(a, .5, -.10, "All reported endpoint predictions are out of fold", size=6, color=MUTED, ha="center")

    b = panel(fig, [.535, .60, .425, .32], "b", "Decoder utility and source advantage")
    plot_rows(b, data["fig3_b_continuous_contrasts"],
              ["Age decoder beyond age + background", "Direct source beyond linear age score", "Direct source beyond flexible score calibration"],
              [TEAL, BLUE, NAVY], ["o", "D", "s"], xmax=.0152,
              footer="95% fixed-prediction bootstrap intervals\nConditional on stored out-of-fold predictions")

    c = panel(fig, [.04, .105, .425, .355], "c", "NHANES III: matched scalar design",
              "Phase 1 fitting → Phase 2 retrospective evaluation")
    box(c, .025, .70, .95, .15, "Phase 1: same 25 measurements + background", BLUE, BLUE_LIGHT, 6.35, True)
    arrow(c, (.33, .69), (.245, .58), TEAL)
    arrow(c, (.67, .69), (.755, .58), BLUE)
    box(c, .015, .40, .46, .175, "Age target → ridge\nOne linear scalar $q_A$", TEAL, TEAL_LIGHT, 6.65, True)
    box(c, .525, .40, .46, .175, "Mortality target → ridge\nOne linear scalar $q_Y$", BLUE, BLUE_LIGHT, 6.65, True)
    text(c, .5, .338, "Both Phase 1 projections frozen before Phase 2 fitting", size=6, color=MUTED, ha="center")
    arrow(c, (.25, .29), (.35, .22), TEAL)
    arrow(c, (.75, .29), (.65, .22), BLUE)
    box(c, .025, -.005, .95, .22, "Phase 2: equal fold-local spline calibration\nAge + background retained in both endpoint models\nFive outer folds; projections not refitted", MUTED, LIGHT, 6.25)
    text(c, .5, -.10, "Phase 2: 7,774 participants · 392 deaths", size=6, color=MUTED, ha="center")

    d = panel(fig, [.535, .105, .425, .355], "d", "Supervision target at matched capacity")
    plot_rows(d, data["fig3_d_matched_scalar_contrasts"],
              ["Age scalar beyond age + background", "Mortality scalar over age scalar"],
              [TEAL, BLUE], ["o", "D"], xmax=.0152,
              footer="95% Fay-BRR intervals: Phase 2 downstream refits\nConditional on frozen Phase 1 projections")
    return fig


def figure4(data):
    fig = plt.figure(figsize=(7.15, 6.7), facecolor=WHITE)
    a = panel(fig, [.04, .80, .92, .12], "a", "Start with the biological question")
    labels = ["Declare the object\nand estimand", "Choose measurements\nand design", "Fit and validate\nfor that target", "State the claim\nsupported by the evidence"]
    xs = [.005, .26, .515, .77]
    for i, (x, label) in enumerate(zip(xs, labels)):
        box(a, x, .25, .22, .51, label, BLUE if i < 3 else INK,
            BLUE_LIGHT if i < 3 else LIGHT, 6.65, True)
        if i < 3:
            arrow(a, (x + .225, .505), (xs[i + 1] - .008, .505))

    b = panel(fig, [.04, .44, .92, .29], "b", "A risk-targeted measurement-acquisition example",
              "Source policies: NHANES 2007–2008 → target evaluation: 2009–2010")
    box(b, .005, .64, .395, .18, "Five prespecified measurement domains\nFreeze one selection under each source objective", MUTED, LIGHT, 6.3)
    arrow(b, (.20, .63), (.20, .535))
    box(b, .005, .315, .395, .205, "Mortality objective selects\nhepatic / protein / injury", ORANGE, ORANGE_LIGHT, 6.5, True)
    box(b, .005, .045, .395, .205, "Age-reconstruction objective selects\nmetabolic / adiposity / lipid", BLUE, BLUE_LIGHT, 6.5, True)
    p = fig.add_axes([.455, .44, .505, .23])
    target = data["fig4_b_policy_intervals"]
    pair = data["fig4_b_paired_difference"]
    plotted = pd.concat([target[["estimate", "ci_low", "ci_high"]], pair[["estimate", "ci_low", "ci_high"]]], ignore_index=True)
    plot_rows(p, plotted, ["Endpoint-directed domain: gain beyond age + background",
                           "Age-directed domain: gain beyond age + background",
                           "Paired endpoint-directed minus age-directed gain"],
              [ORANGE, BLUE, NAVY], ["D", "o", "s"], low="ci_low", high="ci_high", xmax=.0143)
    p.set_xlim(-.002, .0143)
    p.set_xlabel("Five-year mortality log-loss improvement", fontsize=6.3, labelpad=4)
    text(b, .5, -.23, "95% fixed-prediction bootstrap intervals · 4,896 target participants · 157 deaths", size=6, color=MUTED, ha="center")
    text(b, .5, -.315, "Exploratory comparison; domains differ in measurement count and no cost equivalence is assumed", size=6, color=MUTED, ha="center")

    c = panel(fig, [.04, .06, .92, .255], "c", "Different objects require different evidence")
    c.set_xlim(0, 1)
    c.set_ylim(0, 1)
    headers = [(0.005, "Object"), (.23, "Declared target"), (.52, "Required structure and evidence")]
    for x, label in headers:
        text(c, x, .90, label, size=6.8, weight="bold")
    short = [
        ("Measured state", "State under a declared\nmeasurement model", "Anchors, reliability and invariance;\nsame-age state validation"),
        ("Named risk", "Endpoint probability over\na specified horizon", "Endpoint, horizon, loss and population;\nheld-out performance and calibration"),
        ("Transition", "Within-person change\nover a specified interval", "Repeated measurements, time scale\nand a longitudinal model"),
        ("Intervention\ndisplacement", "Intervention effect on\na prespecified measure", "Causal design and contrast;\nmeasurement invariance and follow-up"),
        ("Benefit", "Causal effect on a named\noutcome", "Treatment strategies, causal assumptions\nand the decision population"),
    ]
    for i, (obj, target, evidence) in enumerate(short):
        y = .75 - i * .168
        c.plot([0, 1], [y + .115, y + .115], transform=c.transAxes, color=GRID, linewidth=.55)
        text(c, .005, y, obj, size=6.6, weight="bold", va="center", linespacing=1.14)
        text(c, .23, y, target, size=6.6, va="center", linespacing=1.14)
        text(c, .52, y, evidence, size=6.6, va="center", linespacing=1.14)
    c.plot([0, 1], [.012, .012], transform=c.transAxes, color=GRID, linewidth=.55)
    return fig


def export(fig, stem):
    fig.canvas.draw()
    renderer = fig.canvas.get_renderer()
    checked = 0
    for owner, patch, label in BOX_RECORDS:
        if owner is not fig:
            continue
        pb, tb = patch.get_window_extent(renderer), label.get_window_extent(renderer)
        require(tb.x0 >= pb.x0 and tb.x1 <= pb.x1 and tb.y0 >= pb.y0 and tb.y1 <= pb.y1,
                f"Text exceeds box in {stem}: {label.get_text()}")
        checked += 1
    files = []
    for suffix in ("png", "svg", "pdf"):
        dest = OUT / "figures" / f"{stem}.{suffix}"
        kwargs = {"dpi": 600} if suffix == "png" else {}
        fig.savefig(dest, bbox_inches="tight", pad_inches=.04, facecolor=WHITE, **kwargs)
        files.append({"path": dest.relative_to(ROOT).as_posix(), "bytes": dest.stat().st_size})
    plt.close(fig)
    return {"stem": stem, "contained_box_labels": checked, "exports": files}


def main():
    parser = argparse.ArgumentParser(description=__doc__)
    parser.add_argument("--validate-supplied-only", action="store_true",
                        help="Validate supplied aggregate copies without reading predecessor paths or rendering")
    args = parser.parse_args()
    data, sources = load_inputs(supplied_only=args.validate_supplied_only)
    if args.validate_supplied_only:
        print(json.dumps({"status": "PASS_SUPPLIED_AGGREGATE_COPIES_ONLY", "tables": len(sources),
                          "predecessor_files_read": False, "models_loaded": False,
                          "models_fitted": False, "new_estimates_computed": False}, indent=2))
        return
    figures = [export(figure3(data), "Figure_3_NHANES_source_representation_and_supervision_target"),
               export(figure4(data), "Figure_4_target_first_measurement_and_evidence")]
    report = {"automated_status": "PASS", "manual_visual_status": "SEE_VISUAL_QA_MD",
              "source_copy_checks": sources, "figures": figures,
              "models_loaded": False, "models_fitted": False, "random_numbers_generated": False,
              "participant_data_read": False, "new_estimates_computed": False,
              "python_executable": sys.executable,
              "scope": "Static reassembly of frozen aggregate figure evidence only"}
    (OUT / "BUILD_REPORT.json").write_text(json.dumps(report, indent=2) + "\n", encoding="utf-8")
    print(json.dumps(report, indent=2))


if __name__ == "__main__":
    main()
```
