## Supplementary_Software_1 for "Inferential boundaries of age prediction: why prediction does not establish biological age measurement": plot_v6_23_figure4_connector_v0_1.py

```
#!/usr/bin/env python
"""Static Figure 4b connector correction using frozen v6.22 aggregate evidence.

Reuses the v6.22 Figure 4 renderer without invoking its input-copy or main paths.
No Figure 3, participant rows, fitted models, or new estimates are accessed.
"""

from __future__ import annotations

from io import BytesIO
import json
from pathlib import Path
import sys

import matplotlib.pyplot as plt
from matplotlib.lines import Line2D
from matplotlib.patches import FancyArrowPatch
import numpy as np
import pandas as pd
from PIL import Image

import plot_v6_22_main_figure_assembly_v0_1 as baseline


ROOT = Path(__file__).resolve().parents[1]
OUT = ROOT / "outputs" / "v6_23_figure4_connector_v0_1"
FROZEN = ROOT / "outputs" / "v6_22_main_figure_assembly_v0_1"
STEM = "Figure_4_target_first_measurement_and_evidence"
DPI = 600
PAD = .04


def render_png(fig) -> np.ndarray:
    stream = BytesIO()
    fig.savefig(stream, format="png", dpi=DPI, bbox_inches="tight",
                pad_inches=PAD, facecolor=baseline.WHITE)
    stream.seek(0)
    return np.asarray(Image.open(stream).convert("RGBA")).copy()


def text_state(fig):
    return [(text.get_text(), text.get_position(), text.get_fontsize(),
             text.get_color(), text.get_fontweight())
            for ax in fig.axes for text in ax.texts]


def load_frozen_figure4():
    index_path = FROZEN / "source_data" / "source_data_index.tsv"
    index = pd.read_csv(index_path, sep="\t")
    rows = index[index.file.str.startswith("Figure_4/")]
    baseline.require(len(rows) == 4, "Expected exactly four frozen Figure 4 tables")
    data, records, original_bytes = {}, [], {}
    for row in rows.itertuples(index=False):
        path = FROZEN / "source_data" / row.file
        payload = path.read_bytes()
        baseline.require(baseline.sha256(path) == row.sha256,
                         f"Frozen aggregate differs from v6.22 index: {row.file}")
        key = path.stem
        data[key] = pd.read_csv(BytesIO(payload), sep="\t")
        baseline.require(len(data[key]) == row.rows, f"Row count changed: {key}")
        original_bytes[path] = payload
        records.append({"path": path.relative_to(ROOT).as_posix(),
                        "rows": int(row.rows), "sha256": row.sha256,
                        "matches_frozen_index": True, "copied": False})
    return data, records, original_bytes


def replace_connector(fig):
    b = fig.axes[1]
    candidates = [patch for patch in b.patches if isinstance(patch, FancyArrowPatch)]
    baseline.require(len(candidates) == 1, "Unexpected Figure 4b connector structure")
    fig.canvas.draw()
    removed_extent = candidates[0].get_window_extent(fig.canvas.get_renderer()).frozen()
    candidates[0].remove()
    # The common candidate box feeds both objectives via one shared right-hand
    # branch. All boxes, their labels, and all quantitative marks remain in place.
    trunk = Line2D([.409, .435, .435], [.730, .730, .1475],
                   transform=b.transAxes, color=baseline.MUTED, linewidth=.8,
                   solid_capstyle="butt", solid_joinstyle="miter")
    b.add_line(trunk)
    added = [trunk]
    for y in (.4175, .1475):
        baseline.arrow(b, (.435, y), (.410, y))
        added.append(b.patches[-1])
    fig.canvas.draw()
    extents = [removed_extent] + [artist.get_window_extent(fig.canvas.get_renderer()).frozen()
                                  for artist in added]
    return extents


def compare_pixels(before, after, fig, extents):
    baseline.require(before.shape == after.shape, "Connector edit changed image dimensions")
    changed = np.any(before != after, axis=2)
    allowed = np.zeros(changed.shape, dtype=bool)
    bounds = fig.get_tightbbox(fig.canvas.get_renderer())
    height, width = changed.shape
    for extent in extents:
        # Artist extents are display pixels at figure DPI; exports use a tight
        # inch-space bounding box plus PAD. Include antialiasing and stroke width.
        x0 = int(np.floor((extent.x0 / fig.dpi - bounds.x0 + PAD) * DPI)) - 8
        x1 = int(np.ceil((extent.x1 / fig.dpi - bounds.x0 + PAD) * DPI)) + 8
        y0 = int(np.floor((bounds.y1 + PAD - extent.y1 / fig.dpi) * DPI)) - 8
        y1 = int(np.ceil((bounds.y1 + PAD - extent.y0 / fig.dpi) * DPI)) + 8
        allowed[max(0, y0):min(height, y1 + 1), max(0, x0):min(width, x1 + 1)] = True
    outside = int(np.count_nonzero(changed & ~allowed))
    baseline.require(outside == 0, f"Changed {outside} pixels outside connector extents")
    baseline.require(np.any(changed), "No connector correction appeared in the export")
    ys, xs = np.nonzero(changed)
    return {"image_size_pixels": [width, height],
            "changed_pixels": int(np.count_nonzero(changed)),
            "changed_pixel_bbox_xyxy": [int(xs.min()), int(ys.min()), int(xs.max()), int(ys.max())],
            "changed_pixels_outside_connector_extents": outside}


def main():
    baseline.require((OUT / "README.md").exists(), "README must precede rendering")
    (OUT / "figures").mkdir(exist_ok=True)
    data, source_records, original_bytes = load_frozen_figure4()
    fig = baseline.figure4(data)
    before_text = text_state(fig)
    before = render_png(fig)
    with Image.open(FROZEN / "figures" / f"{STEM}.png") as frozen_png:
        frozen_pixels = np.asarray(frozen_png.convert("RGBA"))
    baseline.require(np.array_equal(before, frozen_pixels),
                     "Current baseline renderer does not match frozen v6.22 PNG pixels")
    extents = replace_connector(fig)
    baseline.require(before_text == text_state(fig), "Text or text styling changed")
    renderer = fig.canvas.get_renderer()
    contained_labels = 0
    for owner, patch, label in baseline.BOX_RECORDS:
        if owner is fig:
            pb, tb = patch.get_window_extent(renderer), label.get_window_extent(renderer)
            baseline.require(tb.x0 >= pb.x0 and tb.x1 <= pb.x1 and
                             tb.y0 >= pb.y0 and tb.y1 <= pb.y1,
                             f"Text exceeds box: {label.get_text()}")
            contained_labels += 1
    exports = []
    for suffix in ("png", "svg", "pdf"):
        path = OUT / "figures" / f"{STEM}.{suffix}"
        fig.savefig(path, bbox_inches="tight", pad_inches=PAD, facecolor=baseline.WHITE,
                    **({"dpi": DPI} if suffix == "png" else {}))
        exports.append({"path": path.relative_to(ROOT).as_posix(), "bytes": path.stat().st_size})
    with Image.open(OUT / "figures" / f"{STEM}.png") as final_png:
        after = np.asarray(final_png.convert("RGBA"))
        actual_dpi = list(final_png.info["dpi"])
    pixel_check = compare_pixels(before, after, fig, extents)
    for path, payload in original_bytes.items():
        baseline.require(path.read_bytes() == payload, f"Frozen table changed during render: {path}")
    plt.close(fig)
    report = {"automated_status": "PASS", "manual_visual_status": "SEE_VISUAL_QA_MD",
              "scope": "Figure 4b connector only; one common source branches to both objective boxes",
              "baseline_png_pixel_exact_before_connector_edit": True,
              "text_labels_and_styling_unchanged": True,
              "contained_box_labels": contained_labels,
              "pixel_comparison": pixel_check, "png_dpi": actual_dpi,
              "frozen_aggregate_sources": source_records,
              "all_source_table_bytes_unchanged_after_render": True,
              "source_data_copied": False, "figure3_rendered": False,
              "participant_data_read": False, "models_loaded": False, "models_fitted": False,
              "random_numbers_generated": False, "new_estimates_computed": False,
              "python_executable": sys.executable, "exports": exports}
    (OUT / "BUILD_REPORT.json").write_text(json.dumps(report, indent=2) + "\n", encoding="utf-8")
    print(json.dumps(report, indent=2))


if __name__ == "__main__":
    main()
```
