## Supplementary_Software_1 for "Inferential boundaries of age prediction: why prediction does not establish biological age measurement": plot_v6_24_figure1_ed2_labels_v0_1.py

```
#!/usr/bin/env python
"""Fit-free Fig.1 and ED2 label synchronization from frozen v6.23 assets.

Only the named renderer entry points are called. Their export hooks stop each
builder before its downstream documentation work; source writes are intercepted.
No other figures, participant data, statistical objects or random draws are used.
"""
from __future__ import annotations

from io import BytesIO
import json
from pathlib import Path
import sys
from unittest.mock import patch

import matplotlib
matplotlib.use("Agg")
import matplotlib.pyplot as plt
from matplotlib.patches import FancyArrowPatch
import numpy as np
import pandas as pd
from PIL import Image

import plot_v6_9_figure1_precision_revision_v0_1 as f1
import plot_v6_2_submission_extended_data_v0_1 as ed2

ROOT = Path(__file__).resolve().parents[1]
OUT = ROOT / "outputs" / "v6_24_figure1_ed2_labels_v0_1"
PARENT = ROOT / "manuscript/nature_first/submission_20260911_v6_23_candidate/upload"
OLD1 = ROOT / "outputs/v6_9_figure1_theory_chain_v0_1"
OLDE = ROOT / "outputs/v6_2_submission_extended_data_v0_1/extended_data_figure_2"
STEM1 = "Figure_1_age_decoding_identifies_an_age_task_not_biological_age"
STEME = "Extended_Data_Figure_2_hsct_referent_sensitivities"
FIG1_RC = dict(matplotlib.rcParams)


def require(ok, message):
    if not ok:
        raise RuntimeError(message)


class Captured(Exception):
    def __init__(self, fig):
        self.fig = fig


def capture(fig, *args, **kwargs):
    raise Captured(fig)


def capture_figure1():
    with patch.object(f1, "ensure_package", lambda *args: None), patch.object(f1, "save_figure", capture):
        try:
            f1.build_figure1()
        except Captured as result:
            return result.fig
    raise RuntimeError("Figure 1 capture failed")


def capture_ed2(tables):
    def frozen_read(path):
        require(path.name in tables, f"Unexpected source read: {path}")
        return tables[path.name].copy(deep=True)
    with patch.object(ed2, "PACKAGE", OUT / "renderer_scratch"), \
         patch.object(ed2, "read_tsv", frozen_read), \
         patch.object(ed2, "write_tsv", lambda *args: None), \
         patch.object(ed2, "export", capture):
        try:
            ed2.build_ed2()
        except Captured as result:
            return result.fig
    raise RuntimeError("ED2 capture failed")


def image_array(fig, pad):
    stream = BytesIO()
    fig.savefig(stream, format="png", dpi=600, bbox_inches="tight", pad_inches=pad, facecolor="white")
    stream.seek(0)
    return np.asarray(Image.open(stream).convert("RGBA")).copy()


def parent_parity(fig, reference, pad):
    actual = image_array(fig, pad)
    with Image.open(reference) as image:
        expected = np.asarray(image.convert("RGBA"))
    same_shape = actual.shape == expected.shape
    result = {"reference": str(reference.relative_to(ROOT)), "same_dimensions": same_shape,
              "pixel_exact": bool(same_shape and np.array_equal(actual, expected))}
    if same_shape:
        result["changed_pixels"] = int(np.count_nonzero(np.any(actual != expected, axis=2)))
    return result


def find_text(ax, old):
    found = [item for item in ax.texts if item.get_text() == old]
    require(len(found) == 1, f"Expected one text match: {old}")
    return found[0]


def change_text(ax, old, new, changes, panel):
    obj = find_text(ax, old)
    obj.set_text(new)
    changes.append({"figure": "Figure_1", "panel": panel, "old_label": old, "new_label": new})
    return obj


def numeric_artists(fig):
    state = []
    for ax in fig.axes:
        state.append({"position": list(ax.get_position().bounds),
                      "xlim": list(ax.get_xlim()), "ylim": list(ax.get_ylim()),
                      "lines": [(np.asarray(line.get_xdata()).tolist(), np.asarray(line.get_ydata()).tolist())
                                for line in ax.lines],
                      "collections": [(collection.get_offsets().tolist(),
                                        [path.vertices.tolist() for path in collection.get_paths()])
                                       for collection in ax.collections]})
    return state


def edit_figure1(fig, changes):
    a, b, c, d, e = fig.axes
    change_text(a, "same-age labels supply none of these meanings",
                "chronological-age labels alone supply none of these meanings", changes, "a")
    change_text(c, "Post-decoding operations do not restore identity",
                "Post-decoding operations do not\nestablish construct identity", changes, "c")
    condition = "Given observed age and background (decoder/centring rule fixed)"
    c.text(.5, .555, condition, transform=c.transAxes, ha="center", fontsize=6.3, color=f1.MUTED)
    changes.append({"figure": "Figure_1", "panel": "c", "old_label": "", "new_label": condition})
    candidates = [p for p in d.patches if isinstance(p, FancyArrowPatch)]
    require(len(candidates) == 4, "Unexpected Figure 1d link count")
    candidates[-1].set_linestyle("--")
    # Place a short association label beside the dashed link, clear of its nodes.
    d.text(.91, .525, "predictive\nassociation", transform=d.transAxes, ha="center", va="center",
           fontsize=6.3, color=f1.TEAL, linespacing=1.12)
    changes.append({"figure": "Figure_1", "panel": "d", "old_label": "unlabelled solid score-to-endpoint arrow",
                    "new_label": "dashed arrow labelled predictive association"})


def edit_ed2(fig, tables, changes):
    a, b, c, d = fig.axes
    a.set_xlabel("Mean $J_{\\mathrm{occ}}$ (occasion-specific affiliation; lineage side positive)")
    a.set_title("Holland: descriptive occasion-specific affiliation sensitivities", loc="left")
    b.set_title("Descriptive affiliation and sign-balanced null contrasts", loc="left")
    b.text(.02, .90, "Pair means of occasion-specific affiliation ($J_{\\mathrm{occ}}$)",
           transform=b.transAxes, fontsize=8.5, color=ed2.MID, va="center")
    b.text(.02, .39, "Sign-balanced affiliation of pair-mean coordinates ($J_{\\mathrm{pair}}$)",
           transform=b.transAxes, fontsize=8.5, color=ed2.MID, va="top")
    # Give the lower heading room by moving its two complete text rows together.
    for text in b.texts:
        x, y = text.get_position()
        if np.isclose(y, .31) or np.isclose(y, .14):
            text.set_position((x, y - .055))
        if "observed−null" in text.get_text():
            text.set_text(text.get_text().replace("Soraas2019", "Søraas 2019").replace("Holland2024", "Holland 2024"))
    details = [
        "Fixed processing: overall pair-weighted $J_{\\mathrm{occ}}$ = 0.535\n95% CI 0.416–0.644",
        "Common-probe fixed external-mean sensitivity: $J_{\\mathrm{occ}}$ = 0.477\n95% CI 0.346–0.598",
        "Duplicated occasions across all samples: median score range 3.89 y\n41 occasions: 39 recipient + 2 donor",
        "Negative-time handling: same primary array identities\nacross three rules",
    ]
    detail_texts = [t for t in d.texts if t.get_position()[0] == .10]
    require(len(detail_texts) == 4, "Unexpected ED2 diagnostic labels")
    for obj, label in zip(detail_texts, details):
        changes.append({"figure": "Extended_Data_Figure_2", "panel": "d", "old_label": obj.get_text(), "new_label": label})
        obj.set_text(label)
        obj.set_linespacing(1.12)
    changes.extend([
        {"figure": "Extended_Data_Figure_2", "panel": "a", "old_label": "Mean bounded affiliation index",
         "new_label": "Mean J_occ; pair summaries of occasion-specific affiliation (pair-median row retained)"},
        {"figure": "Extended_Data_Figure_2", "panel": "b", "old_label": "Sign strata and observed-minus-null contrasts",
         "new_label": "Upper: pair means of J_occ; lower: sign-balanced J_pair observed-minus-null"},
    ])


def export(fig, stem, pad):
    paths = []
    for suffix in ("png", "svg", "pdf"):
        path = OUT / "figures" / f"{stem}.{suffix}"
        fig.savefig(path, dpi=600 if suffix == "png" else None, bbox_inches="tight", pad_inches=pad, facecolor="white")
        paths.append(str(path.relative_to(ROOT)))
    plt.close(fig)
    return paths


def source_data(changes, payloads):
    concept_path = OLD1 / "source_data/figure1_theory_chain.tsv"
    concepts = pd.read_csv(concept_path, sep="\t")
    concepts.loc[concepts.relation == "missing_bridge", "interpretation"] = "chronological-age labels alone supply none of these meanings"
    concepts.loc[concepts.relation == "operator", "interpretation"] = "Given observed age and background, with decoder and centring rule fixed: deterministic scalar chain; post-decoding operations do not establish construct identity"
    concepts.loc[concepts.panel == "d", "interpretation"] = "predictive association under shared covariance; dashed score-endpoint link does not assert causation"
    concepts.loc[concepts.panel == "d", "relation"] = "association"
    dest1 = OUT / "source_data/Figure_1/figure1_theory_chain.tsv"
    dest1.parent.mkdir(parents=True, exist_ok=True)
    concepts.to_csv(dest1, sep="\t", index=False)
    records = [{"publication": "Figure_1", "source": str(concept_path.relative_to(ROOT)),
                "successor": str(dest1.relative_to(ROOT)), "action": "replace conceptual table; no numerical estimates"}]
    for path, payload in payloads.items():
        dest = OUT / "source_data/Extended_Data_Figure_2" / path.name
        dest.parent.mkdir(parents=True, exist_ok=True)
        dest.write_bytes(payload)
        require(dest.read_bytes() == payload, f"Source byte parity failed: {dest}")
        records.append({"publication": "Extended_Data_Figure_2", "source": str(path.relative_to(ROOT)),
                        "successor": str(dest.relative_to(ROOT)), "action": "byte-exact frozen aggregate copy; labels interpreted by publication_labels.tsv"})
    labels = OUT / "source_data/publication_labels.tsv"
    pd.DataFrame(changes).to_csv(labels, sep="\t", index=False)
    records.append({"publication": "Figure_1;Extended_Data_Figure_2", "source": "approved v6.24 label correction",
                    "successor": str(labels.relative_to(ROOT)), "action": "add explicit publication-label and estimand map"})
    pd.DataFrame(records).to_csv(OUT / "SOURCE_REPLACEMENT_MAP.tsv", sep="\t", index=False)
    return records


def main():
    (OUT / "figures").mkdir(parents=True, exist_ok=True)
    sources = [OLDE / "source_data" / name for name in
               ("panel_a_holland_sensitivities.tsv", "panel_b_strata_and_null.tsv", "panel_c_soraas_populations.tsv", "panel_d_processing_diagnostics.tsv")]
    payloads = {path: path.read_bytes() for path in sources}
    tables = {path.name: pd.read_csv(BytesIO(payload), sep="\t") for path, payload in payloads.items()}
    changes, outputs = [], []
    with matplotlib.rc_context(FIG1_RC):
        fig = capture_figure1()
        p1 = parent_parity(fig, PARENT / "Main_Figures/Figure_1.png", .045)
        require(p1["pixel_exact"], "Figure 1 parent reproduction drift")
        edit_figure1(fig, changes)
        outputs.extend(export(fig, STEM1, .045))
    with matplotlib.rc_context(matplotlib.rcParamsDefault):
        ed2.configure()
        fig = capture_ed2(tables)
        pe = parent_parity(fig, PARENT / "Extended_Data_Figures/Extended_Data_Figure_2.png", .12)
        require(pe["pixel_exact"], "ED2 parent reproduction drift")
        before = numeric_artists(fig)
        edit_ed2(fig, tables, changes)
        require(before == numeric_artists(fig), "ED2 numerical marks, intervals, axes or layout changed")
        outputs.extend(export(fig, STEME, .12))
    records = source_data(changes, payloads)
    require(all(path.read_bytes() == payload for path, payload in payloads.items()), "Frozen source changed")
    report = {"status": "PASS", "manual_visual_status": "SEE_VISUAL_QA_MD", "parent_reproduction": {"Figure_1": p1, "ED2": pe},
              "ed2_numeric_artists_intervals_axes_and_positions_exact": True,
              "ed2_aggregate_source_copies_byte_exact": True, "ed2_frozen_sources_unchanged": True,
              "models_loaded": False, "models_fitted": False, "random_numbers_generated": False,
              "participant_data_read": False, "new_estimates_computed": False,
              "only_figures_rendered": ["Figure_1", "Extended_Data_Figure_2"],
              "python_executable": sys.executable, "outputs": outputs, "source_map": records}
    (OUT / "BUILD_REPORT.json").write_text(json.dumps(report, indent=2, ensure_ascii=False) + "\n", encoding="utf-8")
    print(json.dumps(report, indent=2, ensure_ascii=False))


if __name__ == "__main__":
    main()
```
