## Supplementary_Software_1 for "Inferential boundaries of age prediction: why prediction does not establish biological age measurement": prepare_nhanes_broad_clinical_panel_v0_1.py

```
#!/usr/bin/env python
"""Assemble a broad, cross-cycle NHANES clinical panel with linked mortality.

The panel deliberately expands beyond the nine PhenoAge biomarkers. Candidate
features are required to have the same NHANES variable code in every included
cycle (2007-2018), and the primary table uses complete cases so feature-level
age and endpoint tests share an identical risk set.
"""

from __future__ import annotations

import json
from pathlib import Path

import numpy as np
import pandas as pd


ROOT = Path(__file__).resolve().parents[1]
RAW = ROOT / "data" / "raw" / "NHANES_mortality_mvp"
PROCESSED = ROOT / "data" / "processed"
OUT_TABLE = PROCESSED / "NHANES_broad_clinical_panel_v0_1.tsv.gz"
OUT_MANIFEST = PROCESSED / "NHANES_broad_clinical_feature_manifest_v0_1.tsv"
OUT_SUMMARY = PROCESSED / "NHANES_broad_clinical_panel_summary_v0_1.json"

CYCLES = [
    ("2007_2008", "E"),
    ("2009_2010", "F"),
    ("2011_2012", "G"),
    ("2013_2014", "H"),
    ("2015_2016", "I"),
    ("2017_2018", "J"),
]

# feature, source module, source column, label, unit, transform
FEATURES = [
    ("albumin", "BIOPRO", "LBDSALSI", "Albumin", "g/L", "identity"),
    ("alt", "BIOPRO", "LBXSATSI", "Alanine aminotransferase", "U/L", "log1p"),
    ("ast", "BIOPRO", "LBXSASSI", "Aspartate aminotransferase", "U/L", "log1p"),
    ("alkaline_phosphatase", "BIOPRO", "LBXSAPSI", "Alkaline phosphatase", "U/L", "log1p"),
    ("blood_urea_nitrogen", "BIOPRO", "LBDSBUSI", "Blood urea nitrogen", "mmol/L", "log1p"),
    ("calcium", "BIOPRO", "LBDSCASI", "Calcium", "mmol/L", "identity"),
    ("total_cholesterol", "BIOPRO", "LBDSCHSI", "Total cholesterol", "mmol/L", "identity"),
    ("bicarbonate", "BIOPRO", "LBXSC3SI", "Bicarbonate", "mmol/L", "identity"),
    ("creatinine", "BIOPRO", "LBDSCRSI", "Creatinine", "umol/L", "log1p"),
    ("ggt", "BIOPRO", "LBXSGTSI", "Gamma-glutamyl transferase", "U/L", "log1p"),
    ("glucose", "BIOPRO", "LBDSGLSI", "Glucose", "mmol/L", "log1p"),
    ("iron", "BIOPRO", "LBDSIRSI", "Iron", "umol/L", "log1p"),
    ("ldh", "BIOPRO", "LBXSLDSI", "Lactate dehydrogenase", "U/L", "log1p"),
    ("phosphorus", "BIOPRO", "LBDSPHSI", "Phosphorus", "mmol/L", "identity"),
    ("total_bilirubin", "BIOPRO", "LBDSTBSI", "Total bilirubin", "umol/L", "log1p"),
    ("total_protein", "BIOPRO", "LBDSTPSI", "Total protein", "g/L", "identity"),
    ("triglycerides", "BIOPRO", "LBDSTRSI", "Triglycerides", "mmol/L", "log1p"),
    ("uric_acid", "BIOPRO", "LBDSUASI", "Uric acid", "umol/L", "log1p"),
    ("sodium", "BIOPRO", "LBXSNASI", "Sodium", "mmol/L", "identity"),
    ("potassium", "BIOPRO", "LBXSKSI", "Potassium", "mmol/L", "identity"),
    ("chloride", "BIOPRO", "LBXSCLSI", "Chloride", "mmol/L", "identity"),
    ("osmolality", "BIOPRO", "LBXSOSSI", "Osmolality", "mmol/kg", "identity"),
    ("globulin", "BIOPRO", "LBDSGBSI", "Globulin", "g/L", "identity"),
    ("wbc", "CBC", "LBXWBCSI", "White-blood-cell count", "10^9/L", "log1p"),
    ("lymphocyte_percent", "CBC", "LBXLYPCT", "Lymphocyte percentage", "%", "identity"),
    ("monocyte_percent", "CBC", "LBXMOPCT", "Monocyte percentage", "%", "identity"),
    ("neutrophil_percent", "CBC", "LBXNEPCT", "Neutrophil percentage", "%", "identity"),
    ("eosinophil_percent", "CBC", "LBXEOPCT", "Eosinophil percentage", "%", "log1p"),
    ("basophil_percent", "CBC", "LBXBAPCT", "Basophil percentage", "%", "log1p"),
    ("lymphocyte_count", "CBC", "LBDLYMNO", "Lymphocyte count", "10^9/L", "log1p"),
    ("monocyte_count", "CBC", "LBDMONO", "Monocyte count", "10^9/L", "log1p"),
    ("neutrophil_count", "CBC", "LBDNENO", "Neutrophil count", "10^9/L", "log1p"),
    ("eosinophil_count", "CBC", "LBDEONO", "Eosinophil count", "10^9/L", "log1p"),
    ("basophil_count", "CBC", "LBDBANO", "Basophil count", "10^9/L", "log1p"),
    ("rbc", "CBC", "LBXRBCSI", "Red-blood-cell count", "10^12/L", "identity"),
    ("hemoglobin", "CBC", "LBXHGB", "Hemoglobin", "g/dL", "identity"),
    ("hematocrit", "CBC", "LBXHCT", "Hematocrit", "%", "identity"),
    ("mcv", "CBC", "LBXMCVSI", "Mean corpuscular volume", "fL", "identity"),
    ("mch", "CBC", "LBXMCHSI", "Mean corpuscular hemoglobin", "pg", "identity"),
    ("mchc", "CBC", "LBXMC", "Mean corpuscular hemoglobin concentration", "g/dL", "identity"),
    ("rdw", "CBC", "LBXRDW", "Red-cell distribution width", "%", "log1p"),
    ("platelets", "CBC", "LBXPLTSI", "Platelet count", "10^9/L", "log1p"),
    ("mean_platelet_volume", "CBC", "LBXMPSI", "Mean platelet volume", "fL", "identity"),
    ("weight", "BMX", "BMXWT", "Weight", "kg", "log1p"),
    ("bmi", "BMX", "BMXBMI", "Body-mass index", "kg/m2", "log1p"),
    ("waist", "BMX", "BMXWAIST", "Waist circumference", "cm", "log1p"),
    ("systolic_bp", "BPX", "__MEAN_SYS__", "Systolic blood pressure", "mmHg", "identity"),
    ("diastolic_bp", "BPX", "__MEAN_DIA__", "Diastolic blood pressure", "mmHg", "identity"),
    ("pulse", "BPX", "BPXPLS", "Pulse", "beats/min", "log1p"),
    ("hdl", "HDL", "LBDHDDSI", "HDL cholesterol", "mmol/L", "log1p"),
]


def read_xpt(path: Path) -> pd.DataFrame:
    frame = pd.read_sas(path, format="xport", encoding="latin1")
    frame.columns = [str(column).upper() for column in frame.columns]
    return frame


def row_mean(frame: pd.DataFrame, prefix: str) -> pd.Series:
    columns = [f"{prefix}{index}" for index in range(1, 5)]
    return frame[columns].apply(pd.to_numeric, errors="coerce").mean(axis=1, skipna=True)


def build_cycle(base: pd.DataFrame, cycle: str, suffix: str) -> pd.DataFrame:
    directory = RAW / cycle
    modules = {
        module: read_xpt(directory / f"{module}_{suffix}.XPT")
        for module in ["BIOPRO", "CBC", "BMX", "BPX", "HDL"]
    }
    frame = base.loc[base["cycle"].eq(cycle)].copy()
    for module, table in modules.items():
        required = {"SEQN"}
        required.update(
            source
            for _, source_module, source, _, _, _ in FEATURES
            if source_module == module and not source.startswith("__")
        )
        if module == "BPX":
            required.update(f"BPXSY{index}" for index in range(1, 5))
            required.update(f"BPXDI{index}" for index in range(1, 5))
        missing = sorted(required - set(table.columns))
        if missing:
            raise ValueError(f"{cycle} {module} missing required columns: {missing}")
        frame = frame.merge(table[sorted(required)], on="SEQN", how="left", validate="one_to_one")

    frame["__MEAN_SYS__"] = row_mean(frame, "BPXSY")
    frame["__MEAN_DIA__"] = row_mean(frame, "BPXDI")
    for feature, _, source, _, _, transform in FEATURES:
        values = pd.to_numeric(frame[source], errors="coerce")
        frame[feature] = values
        frame[f"T__{feature}"] = (
            np.log1p(values.clip(lower=0)) if transform == "log1p" else values
        )
    return frame


def main() -> int:
    base = pd.read_csv(PROCESSED / "NHANES_mortality_mvp_biomarkers.tsv", sep="\t")
    combined = pd.concat(
        [build_cycle(base, cycle, suffix) for cycle, suffix in CYCLES],
        ignore_index=True,
        sort=False,
    )
    combined["event"] = combined["mortality_status"].eq(1).astype(int)
    combined["time_months"] = pd.to_numeric(
        combined["followup_months_exam"], errors="coerce"
    )
    transformed = [f"T__{feature}" for feature, *_ in FEATURES]
    eligibility = (
        combined["age_years"].between(20, 79, inclusive="both")
        & combined["eligible_mortality"].eq(1)
        & combined["time_months"].gt(0)
        & combined["mortality_status"].notna()
    )
    complete = eligibility & combined[transformed].notna().all(axis=1)
    metadata = [
        "SEQN",
        "cycle",
        "age_years",
        "sex",
        "race_ethnicity",
        "exam_weight",
        "event",
        "time_months",
    ]
    raw_features = [feature for feature, *_ in FEATURES]
    analysis = combined.loc[complete, metadata + raw_features + transformed].copy()
    if analysis["SEQN"].duplicated().any():
        raise ValueError("SEQN must be unique")
    if len(raw_features) != 50:
        raise ValueError(f"Expected 50 features, found {len(raw_features)}")
    analysis.to_csv(OUT_TABLE, sep="\t", index=False, compression="gzip")

    manifest = pd.DataFrame(
        [
            {
                "feature": feature,
                "transformed_column": f"T__{feature}",
                "source_module": module,
                "source_column": source,
                "clinical_label": label,
                "unit": unit,
                "transform": transform,
            }
            for feature, module, source, label, unit, transform in FEATURES
        ]
    )
    manifest.to_csv(OUT_MANIFEST, sep="\t", index=False)
    by_cycle = (
        analysis.groupby("cycle", observed=True)
        .agg(n=("SEQN", "size"), events=("event", "sum"))
        .reset_index()
    )
    summary = {
        "n_raw_rows": int(len(combined)),
        "n_eligible_before_complete_case": int(eligibility.sum()),
        "n_complete_50_feature_panel": int(len(analysis)),
        "n_events": int(analysis["event"].sum()),
        "age_min": float(analysis["age_years"].min()),
        "age_max": float(analysis["age_years"].max()),
        "n_features": len(raw_features),
        "cycles": by_cycle.to_dict(orient="records"),
        "features": raw_features,
        "selection_rule": (
            "Same NHANES variable code present in all 2007-2018 cycles; "
            "complete cases across all 50 features."
        ),
        "crp_excluded_reason": (
            "No common CRP assay file across all six cycles in the local public extract."
        ),
        "output_table": str(OUT_TABLE.relative_to(ROOT)),
    }
    OUT_SUMMARY.write_text(
        json.dumps(summary, indent=2, ensure_ascii=False),
        encoding="utf-8",
    )
    print(json.dumps(summary, indent=2, ensure_ascii=False))
    return 0


if __name__ == "__main__":
    raise SystemExit(main())
```
