## Supplementary_Software_1 for "Inferential boundaries of age prediction: why prediction does not establish biological age measurement": prepare_nhanes_mortality_mvp.py

```
"""Prepare NHANES clinical biomarker + linked mortality MVP table.

This first pass uses public NHANES 2007-2018 files downloaded under
data/raw/NHANES_mortality_mvp.  It builds a sample-level table for testing
clinical biological-age scalars as lossy quotient coordinates.
"""

from __future__ import annotations

from pathlib import Path

import numpy as np
import pandas as pd


ROOT = Path(__file__).resolve().parents[1]
RAW = ROOT / "data/raw/NHANES_mortality_mvp"
PROCESSED = ROOT / "data/processed"
REPORTS = ROOT / "reports"

OUT_TABLE = PROCESSED / "NHANES_mortality_mvp_biomarkers.tsv"
OUT_READINESS = PROCESSED / "NHANES_mortality_mvp_readiness.tsv"
OUT_REPORT = REPORTS / "NHANES_MORTALITY_MVP_READINESS.md"


CYCLES = [
    ("2007_2008", "E"),
    ("2009_2010", "F"),
    ("2011_2012", "G"),
    ("2013_2014", "H"),
    ("2015_2016", "I"),
    ("2017_2018", "J"),
]


def markdown_table(df: pd.DataFrame) -> str:
    if df.empty:
        return "_No rows._"
    cols = list(df.columns)
    lines = [
        "| " + " | ".join(cols) + " |",
        "| " + " | ".join(["---"] * len(cols)) + " |",
    ]
    for _, row in df.iterrows():
        vals = [str(row[c]).replace("\n", " ") for c in cols]
        lines.append("| " + " | ".join(vals) + " |")
    return "\n".join(lines)


def read_xpt(path: Path) -> pd.DataFrame:
    df = pd.read_sas(path, format="xport", encoding="latin1")
    df.columns = [str(c).upper() for c in df.columns]
    return df


def coalesce_columns(df: pd.DataFrame, candidates: list[str]) -> pd.Series:
    out = pd.Series(np.nan, index=df.index, dtype="float64")
    for col in candidates:
        if col in df.columns:
            out = out.fillna(pd.to_numeric(df[col], errors="coerce"))
    return out


def read_mortality(path: Path) -> pd.DataFrame:
    rows: list[dict[str, object]] = []
    with path.open("r", encoding="ascii", errors="ignore") as handle:
        for line in handle:
            if len(line) < 16:
                continue
            seqn = pd.to_numeric(line[0:6].strip(), errors="coerce")
            if pd.isna(seqn):
                continue

            def field(start_1: int, end_1: int) -> str:
                return line[start_1 - 1 : end_1].strip()

            def num(start_1: int, end_1: int) -> float:
                return pd.to_numeric(field(start_1, end_1), errors="coerce")

            rows.append(
                {
                    "SEQN": int(seqn),
                    "ELIGSTAT": num(15, 15),
                    "MORTSTAT": num(16, 16),
                    "UCOD_LEADING": field(17, 19),
                    "DIABETES_DEATH": num(20, 20),
                    "HYPERTEN_DEATH": num(21, 21),
                    "PERMTH_INT": num(43, 45),
                    "PERMTH_EXM": num(46, 48),
                }
            )
    return pd.DataFrame(rows)


def phenoage_linear_score(df: pd.DataFrame) -> pd.Series:
    """Levine Phenotypic Age xb using common published units.

    Inputs are expected in:
    albumin g/L, creatinine umol/L, glucose mmol/L, CRP mg/L,
    lymphocyte %, MCV fL, RDW %, alkaline phosphatase U/L, WBC 1000 cells/uL,
    age years.
    """

    crp = pd.to_numeric(df["crp_mg_l"], errors="coerce").clip(lower=1e-4)
    return (
        -19.9067
        - 0.0336 * df["albumin_g_l"]
        + 0.0095 * df["creatinine_umol_l"]
        + 0.1953 * df["glucose_mmol_l"]
        + 0.0954 * np.log(crp)
        - 0.0120 * df["lymphocyte_percent"]
        + 0.0268 * df["mcv_f_l"]
        + 0.3306 * df["rdw_percent"]
        + 0.00188 * df["alkaline_phosphatase_u_l"]
        + 0.0554 * df["wbc_1000_ul"]
        + 0.0804 * df["age_years"]
    )


def load_cycle(cycle: str, suffix: str) -> tuple[pd.DataFrame, dict[str, object]]:
    cdir = RAW / cycle
    demo = read_xpt(cdir / f"DEMO_{suffix}.XPT")
    cbc = read_xpt(cdir / f"CBC_{suffix}.XPT")
    biopro = read_xpt(cdir / f"BIOPRO_{suffix}.XPT")
    crp_files = list(cdir.glob("CRP_*.XPT")) + list(cdir.glob("HSCRP_*.XPT"))
    crp = read_xpt(crp_files[0]) if crp_files else pd.DataFrame({"SEQN": []})
    mort_path = cdir / f"NHANES_{cycle}_MORT_2019_PUBLIC.dat"
    mort = read_mortality(mort_path)

    df = demo.merge(cbc, on="SEQN", how="left").merge(biopro, on="SEQN", how="left")
    if not crp.empty:
        df = df.merge(crp, on="SEQN", how="left", suffixes=("", "_CRP"))
    df = df.merge(mort, on="SEQN", how="left")
    df["cycle"] = cycle

    out = pd.DataFrame(
        {
            "SEQN": pd.to_numeric(df["SEQN"], errors="coerce").astype("Int64"),
            "cycle": cycle,
            "age_years": coalesce_columns(df, ["RIDAGEYR"]),
            "sex": coalesce_columns(df, ["RIAGENDR"]),
            "race_ethnicity": coalesce_columns(df, ["RIDRETH1", "RIDRETH3"]),
            "exam_weight": coalesce_columns(df, ["WTMEC2YR"]),
            "albumin_g_l": coalesce_columns(df, ["LBDSALSI"]),
            "creatinine_umol_l": coalesce_columns(df, ["LBDSCRSI"]),
            "glucose_mmol_l": coalesce_columns(df, ["LBDSGLSI", "LBXGLUSI"]),
            "alkaline_phosphatase_u_l": coalesce_columns(df, ["LBXSAPSI", "LBXSAP"]),
            "wbc_1000_ul": coalesce_columns(df, ["LBXWBCSI", "LBXWBC"]),
            "lymphocyte_percent": coalesce_columns(df, ["LBXLYPCT", "LBDLYMNO"]),
            "mcv_f_l": coalesce_columns(df, ["LBXMCVSI", "LBXMCV"]),
            "rdw_percent": coalesce_columns(df, ["LBXRDW", "LBXRDWCV"]),
            "crp_mg_l": coalesce_columns(df, ["LBXHSCRP", "LBXCRP"]),
            "eligible_mortality": coalesce_columns(df, ["ELIGSTAT"]),
            "mortality_status": coalesce_columns(df, ["MORTSTAT"]),
            "followup_months_exam": coalesce_columns(df, ["PERMTH_EXM"]),
            "followup_months_interview": coalesce_columns(df, ["PERMTH_INT"]),
        }
    )
    # Older CRP files are commonly mg/dL; HSCRP files are commonly mg/L.
    if crp_files and crp_files[0].name.startswith("CRP_"):
        out["crp_mg_l"] = out["crp_mg_l"] * 10.0
    out["phenoage_xb"] = phenoage_linear_score(out)

    required = [
        "age_years",
        "albumin_g_l",
        "creatinine_umol_l",
        "glucose_mmol_l",
        "crp_mg_l",
        "lymphocyte_percent",
        "mcv_f_l",
        "rdw_percent",
        "alkaline_phosphatase_u_l",
        "wbc_1000_ul",
        "mortality_status",
        "followup_months_exam",
    ]
    summary = {
        "cycle": cycle,
        "n_demo": len(demo),
        "n_rows": len(out),
        "n_adult_20plus": int(out["age_years"].ge(20).sum()),
        "has_crp_file": bool(crp_files),
        "crp_file": crp_files[0].name if crp_files else "",
        "n_complete_phenoage_mortality_adult": int((out[required].notna().all(axis=1) & out["age_years"].ge(20)).sum()),
        "n_deaths_complete_adult": int(((out[required].notna().all(axis=1)) & out["age_years"].ge(20) & out["mortality_status"].eq(1)).sum()),
        "n_mortality_eligible": int(out["eligible_mortality"].eq(1).sum()),
        "n_deaths_total": int(out["mortality_status"].eq(1).sum()),
    }
    return out, summary


def main() -> int:
    tables: list[pd.DataFrame] = []
    summaries: list[dict[str, object]] = []
    for cycle, suffix in CYCLES:
        table, summary = load_cycle(cycle, suffix)
        tables.append(table)
        summaries.append(summary)

    combined = pd.concat(tables, ignore_index=True)
    readiness = pd.DataFrame(summaries)
    combined.to_csv(OUT_TABLE, sep="\t", index=False)
    readiness.to_csv(OUT_READINESS, sep="\t", index=False)

    complete = combined[
        combined[
            [
                "age_years",
                "albumin_g_l",
                "creatinine_umol_l",
                "glucose_mmol_l",
                "crp_mg_l",
                "lymphocyte_percent",
                "mcv_f_l",
                "rdw_percent",
                "alkaline_phosphatase_u_l",
                "wbc_1000_ul",
                "mortality_status",
                "followup_months_exam",
            ]
        ]
        .notna()
        .all(axis=1)
        & combined["age_years"].ge(20)
    ]
    lines = [
        "# NHANES Mortality MVP Readiness",
        "",
        "## Purpose",
        "",
        "Prepare a public NHANES clinical-biomarker + linked mortality table for testing whether biological-age scalar information is absorbed by explicit physiological state.",
        "",
        "## Cycle Coverage",
        "",
        markdown_table(readiness),
        "",
        "## Complete Adult Analysis Set",
        "",
        f"- rows: {len(complete)}",
        f"- deaths: {int(complete['mortality_status'].eq(1).sum())}",
        f"- cycles with CRP available in this first pass: {', '.join(readiness.loc[readiness['has_crp_file'], 'cycle'])}",
        "",
        "## Interpretation",
        "",
        "This first pass has a complete PhenoAge-style biomarker set for cycles with CRP/HSCRP files.  2011-2014 were downloaded for demographics/CBC/BIOPRO/mortality but are incomplete for CRP until the correct inflammatory marker files are added or the model is run without CRP.",
        "",
        "## Outputs",
        "",
        f"- `{OUT_TABLE.relative_to(ROOT)}`",
        f"- `{OUT_READINESS.relative_to(ROOT)}`",
    ]
    OUT_REPORT.write_text("\n".join(lines) + "\n", encoding="utf-8")
    print(readiness.to_string(index=False))
    print()
    print(f"complete adult rows={len(complete)} deaths={int(complete['mortality_status'].eq(1).sum())}")
    print(f"wrote {OUT_REPORT}")
    return 0


if __name__ == "__main__":
    raise SystemExit(main())
```
