## Supplementary_Software_1 for "Inferential boundaries of age prediction: why prediction does not establish biological age measurement": simulate_three_variable_age_state_geometry_v0_1.py

```
"""Simulate a minimal three-variable age/state geometry for 500 people.

The construction is deliberately transparent:

1. ``x_age_linear`` follows age plus noise.
2. ``x_age_orthogonal_state`` carries a binary separation-state signal but is
   projected to be exactly orthogonal to age in this realized sample.
3. ``x_age_nonlinear_state`` contains a curved age component and a separation-
   state offset.

This is a descriptive simulation. It fits no predictive or inferential model.
"""

from __future__ import annotations

import gzip
import hashlib
import json
from pathlib import Path

import matplotlib.pyplot as plt
import numpy as np
import pandas as pd


ROOT = Path(__file__).resolve().parents[1]
OUTPUT_DIR = ROOT / "outputs" / "three_variable_age_state_geometry_v0_1"
FIGURE_DIR = ROOT / "figures" / "three_variable_age_state_geometry_v0_1"

N_PEOPLE = 500
SEED = 20260806


def zscore(values: np.ndarray) -> np.ndarray:
    centered = values - np.mean(values)
    return centered / np.std(centered, ddof=1)


def remove_linear_age_direction(values: np.ndarray, age_z: np.ndarray) -> np.ndarray:
    """Remove intercept and the realized sample age direction exactly."""
    centered = values - np.mean(values)
    residual = centered - np.dot(centered, age_z) / np.dot(age_z, age_z) * age_z
    residual -= np.mean(residual)
    return residual


def pearson(left: np.ndarray, right: np.ndarray) -> float:
    return float(np.corrcoef(left, right)[0, 1])


def sha256(path: Path) -> str:
    digest = hashlib.sha256()
    with path.open("rb") as handle:
        for block in iter(lambda: handle.read(1024 * 1024), b""):
            digest.update(block)
    return digest.hexdigest()


def simulate() -> tuple[pd.DataFrame, dict[str, object], dict[str, np.ndarray | float]]:
    rng = np.random.default_rng(SEED)

    age = rng.uniform(20.0, 80.0, size=N_PEOPLE)
    age_z = zscore(age)
    separation_state = rng.binomial(1, 0.40, size=N_PEOPLE)
    state_centered = separation_state - np.mean(separation_state)

    x1 = zscore(0.95 * age_z + rng.normal(0.0, 0.65, size=N_PEOPLE))

    x2_candidate = 1.35 * state_centered + rng.normal(0.0, 0.55, size=N_PEOPLE)
    x2 = zscore(remove_linear_age_direction(x2_candidate, age_z))

    age_quadratic = remove_linear_age_direction(age_z**2, age_z)
    age_quadratic = zscore(age_quadratic)
    x3_raw = (
        0.65 * age_z
        + 0.85 * age_quadratic
        + 0.90 * state_centered
        + rng.normal(0.0, 0.55, size=N_PEOPLE)
    )
    x3_mean = float(np.mean(x3_raw))
    x3_sd = float(np.std(x3_raw, ddof=1))
    x3 = (x3_raw - x3_mean) / x3_sd

    frame = pd.DataFrame(
        {
            "person_id": [f"P{i:04d}" for i in range(1, N_PEOPLE + 1)],
            "age": age,
            "separation_state": separation_state,
            "x_age_linear": x1,
            "x_age_orthogonal_state": x2,
            "x_age_nonlinear_state": x3,
        }
    )

    correlations = {
        "corr_age_x_age_linear": pearson(age, x1),
        "corr_age_x_age_orthogonal_state": pearson(age, x2),
        "corr_state_x_age_orthogonal_state": pearson(separation_state, x2),
        "corr_age_x_age_nonlinear_state": pearson(age, x3),
        "corr_age_quadratic_x_age_nonlinear_state": pearson(age_quadratic, x3),
        "corr_state_x_age_nonlinear_state": pearson(separation_state, x3),
    }
    checks = {
        "n_people_is_500": len(frame) == 500,
        "x1_has_age_signal": abs(correlations["corr_age_x_age_linear"]) > 0.60,
        "x2_is_age_orthogonal": abs(correlations["corr_age_x_age_orthogonal_state"]) < 1e-12,
        "x2_has_state_signal": abs(correlations["corr_state_x_age_orthogonal_state"]) > 0.50,
        "x3_has_linear_age_signal": abs(correlations["corr_age_x_age_nonlinear_state"]) > 0.25,
        "x3_has_quadratic_age_signal": abs(correlations["corr_age_quadratic_x_age_nonlinear_state"]) > 0.40,
        "x3_has_state_signal": abs(correlations["corr_state_x_age_nonlinear_state"]) > 0.20,
    }
    if not all(checks.values()):
        failed = [name for name, passed in checks.items() if not passed]
        raise RuntimeError(f"Simulation checks failed: {failed}")

    summary: dict[str, object] = {
        "simulation": "three_variable_age_state_geometry_v0_1",
        "n_people": N_PEOPLE,
        "seed": SEED,
        "age_range": [float(np.min(age)), float(np.max(age))],
        "separation_state_prevalence": float(np.mean(separation_state)),
        "correlations": correlations,
        "checks": checks,
        "construction": {
            "x_age_linear": "z(0.95 * z(age) + Normal(0, 0.65))",
            "x_age_orthogonal_state": (
                "z(project_off_age(1.35 * centered(separation_state) + Normal(0, 0.55)))"
            ),
            "x_age_nonlinear_state": (
                "z(0.65 * z(age) + 0.85 * orthogonal_quadratic_age "
                "+ 0.90 * centered(separation_state) + Normal(0, 0.55))"
            ),
        },
    }
    plotting = {
        "age_z": age_z,
        "age_quadratic": age_quadratic,
        "state_centered": state_centered,
        "x3_mean": x3_mean,
        "x3_sd": x3_sd,
    }
    return frame, summary, plotting


def make_figure(
    frame: pd.DataFrame,
    summary: dict[str, object],
    plotting: dict[str, np.ndarray | float],
) -> plt.Figure:
    colors = np.where(frame["separation_state"].to_numpy() == 1, "#D55E00", "#0072B2")
    correlations = summary["correlations"]
    assert isinstance(correlations, dict)

    fig = plt.figure(figsize=(12.8, 9.0), constrained_layout=True)
    grid = fig.add_gridspec(2, 2)
    axes = [fig.add_subplot(grid[0, 0]), fig.add_subplot(grid[0, 1]), fig.add_subplot(grid[1, 0])]
    ax3d = fig.add_subplot(grid[1, 1], projection="3d")

    panels = [
        ("x_age_linear", "A  Linear age signal + noise", "corr(Age, X1)"),
        ("x_age_orthogonal_state", "B  Age-orthogonal state signal", "corr(Age, X2)"),
        ("x_age_nonlinear_state", "C  Nonlinear age + state signal", "corr(Age, X3)"),
    ]
    corr_keys = [
        "corr_age_x_age_linear",
        "corr_age_x_age_orthogonal_state",
        "corr_age_x_age_nonlinear_state",
    ]
    for axis, (column, title, corr_label), corr_key in zip(axes, panels, corr_keys, strict=True):
        axis.scatter(frame["age"], frame[column], c=colors, s=22, alpha=0.66, linewidths=0)
        axis.set_title(title, loc="left", fontweight="bold")
        axis.set_xlabel("Age (years)")
        axis.set_ylabel(column)
        axis.grid(alpha=0.18)
        axis.text(
            0.03,
            0.96,
            f"{corr_label} = {float(correlations[corr_key]):.3f}",
            transform=axis.transAxes,
            va="top",
            bbox={"boxstyle": "round,pad=0.25", "facecolor": "white", "alpha": 0.88, "edgecolor": "#cccccc"},
        )

    axes[1].text(
        0.03,
        0.84,
        f"corr(State, X2) = {float(correlations['corr_state_x_age_orthogonal_state']):.3f}",
        transform=axes[1].transAxes,
        va="top",
    )
    axes[2].text(
        0.03,
        0.84,
        (
            f"corr(quadratic Age, X3) = "
            f"{float(correlations['corr_age_quadratic_x_age_nonlinear_state']):.3f}\n"
            f"corr(State, X3) = {float(correlations['corr_state_x_age_nonlinear_state']):.3f}"
        ),
        transform=axes[2].transAxes,
        va="top",
    )

    ax3d.scatter(
        frame["x_age_linear"],
        frame["x_age_orthogonal_state"],
        frame["x_age_nonlinear_state"],
        c=colors,
        s=18,
        alpha=0.62,
        linewidths=0,
    )
    ax3d.set_title("D  The three-variable object", loc="left", fontweight="bold")
    ax3d.set_xlabel("X1: age-linear")
    ax3d.set_ylabel("X2: age-orthogonal state")
    ax3d.set_zlabel("X3: nonlinear age + state")
    ax3d.view_init(elev=22, azim=-54)

    from matplotlib.lines import Line2D

    legend = [
        Line2D([0], [0], marker="o", color="w", markerfacecolor="#0072B2", label="State 0", markersize=7),
        Line2D([0], [0], marker="o", color="w", markerfacecolor="#D55E00", label="State 1", markersize=7),
    ]
    fig.legend(handles=legend, loc="upper center", ncol=2, frameon=False, bbox_to_anchor=(0.5, 1.02))
    fig.suptitle("A minimal 500-person age/state simulation", fontsize=16, fontweight="bold", y=1.055)
    return fig


def write_report(summary: dict[str, object], data_path: Path, figure_path: Path) -> str:
    corr = summary["correlations"]
    assert isinstance(corr, dict)
    return f"""# 三变量 Age–State 模拟（v0.1）

## 最小探针问题

在 500 个模拟个体中，能否同时构造：线性年龄方向、与年龄严格正交但携带分离状态的方向，以及同时携带非线性年龄与分离状态的混合方向？

## 构造

- `X1 = Age线性项 + 噪声`。
- `X2 = 分离状态项 + 噪声`，随后在本样本内投影掉 Age 方向。
- `X3 = Age线性项 + Age正交二次项 + 分离状态项 + 噪声`。

## 决定性诊断

- `corr(Age, X1) = {float(corr['corr_age_x_age_linear']):.6f}`
- `corr(Age, X2) = {float(corr['corr_age_x_age_orthogonal_state']):.3e}`
- `corr(State, X2) = {float(corr['corr_state_x_age_orthogonal_state']):.6f}`
- `corr(Age, X3) = {float(corr['corr_age_x_age_nonlinear_state']):.6f}`
- `corr(Age二次正交项, X3) = {float(corr['corr_age_quadratic_x_age_nonlinear_state']):.6f}`
- `corr(State, X3) = {float(corr['corr_state_x_age_nonlinear_state']):.6f}`

## 探针结论

`PROMISING`：三个预定几何角色可以同时实现。这里的“严格正交”是这 500 人样本内的代数约束，不是对新样本总体性质的推断。

## 产物

- 数据：`{data_path.relative_to(ROOT)}`
- 图：`{figure_path.relative_to(ROOT)}`
- 随机种子：`{SEED}`
"""


def main() -> None:
    OUTPUT_DIR.mkdir(parents=True, exist_ok=True)
    FIGURE_DIR.mkdir(parents=True, exist_ok=True)

    frame, summary, plotting = simulate()
    data_path = OUTPUT_DIR / "simulated_people.tsv.gz"
    summary_path = OUTPUT_DIR / "summary.json"
    report_path = OUTPUT_DIR / "PROBE_REPORT_CN.md"
    png_path = FIGURE_DIR / "three_variable_age_state_geometry.png"
    svg_path = FIGURE_DIR / "three_variable_age_state_geometry.svg"

    with gzip.open(data_path, "wt", encoding="utf-8", newline="") as handle:
        frame.to_csv(handle, sep="\t", index=False, float_format="%.10g")

    figure = make_figure(frame, summary, plotting)
    figure.savefig(png_path, dpi=220, bbox_inches="tight", facecolor="white")
    figure.savefig(svg_path, bbox_inches="tight", facecolor="white")
    plt.close(figure)

    report_path.write_text(write_report(summary, data_path, png_path), encoding="utf-8")

    summary["artifacts"] = {
        str(path.relative_to(ROOT)): {"sha256": sha256(path), "bytes": path.stat().st_size}
        for path in [data_path, report_path, png_path, svg_path]
    }
    summary_path.write_text(json.dumps(summary, indent=2, ensure_ascii=False), encoding="utf-8")
    print(json.dumps(summary, indent=2, ensure_ascii=False))


if __name__ == "__main__":
    main()
```
