## Supplementary_Software_1 for "Inferential boundaries of age prediction: why prediction does not establish biological age measurement": THREE_VARIABLE_AGE_STATE_TASK_MAP_PROBE_SPEC_v0_1_20260827.md

# Three-variable age–state task-map probe v0.1

## Decision

Decide whether the frozen X1–X3 construction can support one concise,
model-backed Results paragraph and a publication-facing explanatory figure
without changing the manuscript's analytical proof responsibility.

## Fixed source

- Source rows: `outputs/three_variable_age_state_geometry_v0_1/simulated_people.tsv.gz`.
- Required source SHA-256:
  `55cf8290cbd3acf9e827428df0751361c4110eaa5f2650188ac946ebc84cae49`.
- Independent unit: one synthetic row/person.
- Population: the existing 500 cross-sectional rows generated with seed
  `20260806`; no row is regenerated.
- Age is an ordered cross-sectional coordinate, not within-person follow-up.

## One empirical question

Can fixed models recover the deliberately separated roles of X1, X2 and X3:
age readability, same-age state information and their mixture?

## Fixed folds

Create five participant-disjoint outer folds with `StratifiedKFold`, shuffle
enabled and seed `20260827`. Stratify jointly by the declared binary state and
five equal-frequency age groups. Persist every row's fold assignment.

## Fixed model arms

No tuning, model-family search or alternative split is permitted.

### Chronological-age decoding

Fit four fixed ridge pipelines using X1, X2, X3 or X1–X3. Each pipeline applies
a five-knot cubic `SplineTransformer`, fold-local standardization and
`Ridge(alpha=1.0)`. Report pooled out-of-fold R2, RMSE and MAE.

### Same-age state decoding

Fit five fixed L2-logistic pipelines: age spline only; age spline plus X1, X2,
X3 or X1–X3. Age uses a five-knot cubic spline; X variables are standardized
inside the training fold. Use `LogisticRegression(C=1.0, solver='lbfgs',
max_iter=2000)`. Report pooled out-of-fold log loss, ROC AUC and Brier score.
The primary state estimand is the log-loss reduction relative to the age-only
arm; positive values mean incremental held-out state prediction beyond age.

## Probe gate

Classify the fixed result as `PROMISING` only if all qualitative roles are
recovered:

1. X1 has greater OOF age R2 than X2.
2. X2 has greater state log-loss gain beyond age than X1.
3. X3 has positive OOF age R2 and positive state log-loss gain beyond age.
4. X1–X3 jointly have positive OOF age R2 and positive state log-loss gain.
5. Every persisted model reloads and reproduces its held-out predictions with
   maximum absolute error at most `1e-12`.

Otherwise classify `WEAK`, `ABSENT` or `AMBIGUOUS` and stop. Do not rescue the
probe by changing features, folds, splines, penalties or metrics.

## Figure contract

- Surface: standalone static research figure in PNG, SVG and vector PDF.
- Panels a–c: individual X1, X2 and X3 values against chronological age, with
  the two declared states distinguished by both colour and marker/fill. Overlay
  fixed-bin state-specific means only; do not draw or call them longitudinal
  trajectories.
- Panel d: task map with pooled OOF age R2 on the horizontal axis and pooled
  state log-loss gain beyond age on the vertical axis for X1, X2, X3 and X1–X3.
- Palette: hard two-root cap (blue and orange) plus neutrals.
- Visible subtitle: fixed 500-row cross-sectional construction; age is an
  ordered coordinate, not within-person follow-up.
- No population confidence interval is reported for this fixed descriptive
  construction.

## Required persisted objects

Persist the analysis-ready interface, participant metadata, ordered feature
manifest, folds, all 45 fold-local fitted pipelines, predictions, pooled and
fold metrics, figure Source Data, signatures, artifact manifest, environment
record, reload-parity audit and decision file. A missing object or parity
failure makes the run incomplete.

## Claim boundary

Allowed: in this frozen construction, fixed models separate age readability,
same-age state information and their mixture.

Forbidden: longitudinal change; a human biological-age model; disease
prediction; clinical validation; a universal model ranking; proof of the
analytical non-identification theorem; or a population estimate inferred from
the 500 constructed rows.
