## Supplementary_Software_1 for "Inferential boundaries of age prediction: why prediction does not establish biological age measurement": CORE_NHANES_REPRODUCTION.md

# Continuous NHANES core comparison: inputs and execution order

Version: V6.21, 9 September 2026. Scope: the main continuous-NHANES
age-decoder versus direct-source comparison (SM2). This document does not
certify the other NHANES modules as independently rerunnable.

## What has been checked

The two unchanged public-data preparation scripts were executed in an isolated
directory using the locally retained public NHANES files. The complete-support
loader and fold constructor then reproduced all 13,724 rows, 470 events, the
ordered 50 features and five outer-fold assignments exactly, including every
numeric and categorical value after serialization and reload. The public input
identities are listed in `CORE_NHANES_INPUT_INVENTORY.tsv`; the check report is
`CORE_NHANES_INTERFACE_QA.json`. The analysis-ready 50-feature participant interface is not supplied; the selected frozen participant predictions used by the verifier are supplied.

No estimator was fitted or tuned in that check. Public downloads on a fresh
machine, the source-to-prediction rerun, and numerical parity of refitted
predictions remain unverified. The included software now exposes the missing
generators for the baseline and direct-source predictions; the archive is not
being labelled a verified end-to-end reproduction.

## Inputs and environment

Unpack Supplementary Software 1 into a new directory. Its root contains
`analysis/`, `scripts/` and the environment contract. Do not run these entry
points inside an author checkout containing frozen `outputs/` directories.

Use the documented mamba environment in `ENVIRONMENT.md`, `environment.yml`,
`environment.win-64.lock.txt` and `environment-policy.json`. The lock targets
Windows 64-bit. Run `scripts/Test-ProjectEnvironment.ps1` before Python work.
All Python execution goes through `scripts/Invoke-ProjectPython.ps1`; the
required verifier `scripts/verify_project_environment.py` is included.

Obtain the CDC/NCHS public files named in the input inventory and place them at
the listed relative paths. The unchanged upstream builders iterate over all
six 2007–2018 cycles; the analytical gate subsequently selects 2007–2012.
For each cycle they require DEMO, CBC, BIOPRO, BMX, BPX and HDL XPT files and
the 2019 public linked-mortality DAT file. The four listed CRP/HSCRP files feed
legacy readiness columns only; CRP is excluded from the 50-feature panel.
Their presence matches the original preparation inputs. No WHQ, glucose,
insulin or triglyceride module is needed for this core route.

The inventory records the retained input hashes and source locations. Live
availability of every download was not retested. Mortality files and layouts
are obtainable through the [NCHS linked-mortality page](https://www.cdc.gov/nchs/linked-data/mortality-files/index.html);
survey modules through the [NHANES data catalogue](https://wwwn.cdc.gov/nchs/nhanes/continuousnhanes/).
Use the 2019 mortality release; a later release changes the study input.

## Execution order

From the unpacked software root, create `data/processed/` and `reports/`.
The commands below use a fresh PowerShell child process for each Python call
because the project wrapper isolates that process's runtime PATH.

```powershell
$taskShell = (Get-Process -Id $PID).Path
New-Item -ItemType Directory -Force -Path '.\data\processed', '.\reports' | Out-Null
& $taskShell -NoProfile -File '.\scripts\Test-ProjectEnvironment.ps1'
& $taskShell -NoProfile -File '.\scripts\Invoke-ProjectPython.ps1' -- '.\analysis\prepare_nhanes_mortality_mvp.py'
& $taskShell -NoProfile -File '.\scripts\Invoke-ProjectPython.ps1' -- '.\analysis\prepare_nhanes_broad_clinical_panel_v0_1.py'
```

These preparation commands generate the broad panel and feature manifest from
public sources. The following commands fit the original, specified models and
should be run only in that fresh reproduction directory. They were not executed
during V6.21 manuscript cleanup.

```powershell
& $taskShell -NoProfile -File '.\scripts\Invoke-ProjectPython.ps1' -- '.\analysis\analyze_nhanes_5y_complete_support_contextual_age_probe_v0_1.py'
& $taskShell -NoProfile -File '.\scripts\Invoke-ProjectPython.ps1' -- '.\analysis\audit_nhanes_5y_complete_support_contextual_age_probe_v0_1.py'
& $taskShell -NoProfile -File '.\scripts\Invoke-ProjectPython.ps1' -- '.\analysis\analyze_nhanes_5y_age_decoder_source_information_probe_v0_1.py'
```

The complete-support predecessor creates the analysis-ready risk set, ordered
feature tiers, participant folds, bootstrap indices, all eight specified
endpoint models, their out-of-fold predictions, participant losses and model
reload checks. In the main comparison, **M_WA** supplies the baseline and
**M_S3A** supplies the direct-source model (**M_X** downstream). These are
generated predictions, not unexplained external constants.

The last script learns the pure age decoder and its endpoint calibration using
the same participants and folds. Its filename ends in `v0_1`, but the inspected
source writes the accepted `outputs/nhanes_5y_age_decoder_source_information_probe_v0_2/`
directory. It consumes eight predecessor files: `analysis_ready_risk_set.tsv.gz`,
`participant_folds.tsv`, `ordered_feature_tiers.tsv`, `oof_predictions.tsv.gz`,
`participant_loss_vectors.tsv.gz`, `bootstrap_indices.npz`,
`analysis_signature.json` and `model_artifact_audit.json`.

The scripts persist fitted objects, check signatures and reload outputs. Do not
copy old fitted objects into a newly reconstructed input tree: gzip metadata and
other byte-level provenance may differ even when table values agree. Preserve
new signatures and compare participant identity, folds, ordered features,
predictions and reported contrasts, rather than forcing old hash identities.
The audit-only predecessor's additional contrasts are provenance for generating
the required inputs, not new manuscript results.

## Remaining verification and other modules

Three achievements must remain separate: rebuilding figures from aggregate
Source Data; recalculating statistics from predictions, labels, weights and
folds; and rebuilding predictions from original data. The current bundle provides the selected participant inputs used by the verifier, but not every participant-level input for all supplementary analyses.
Internal model reload checks and the interface check above do not establish the
third route on an external machine.

For NHANES III, the current matched-target script consumes the canonical
`nhanes3_phase1_to_phase2_endpoint_vs_age_acquisition_v0_2_final_20260826`
interfaces, including phase-specific analysis tables, folds/design, feature
order and Fay-BRR factors. Local upstream preparation and mortality-truth
scripts exist, but their historical stage/signature dependencies have not yet
been converted into and tested as a stand-alone public reconstruction chain.
That path remains an explicit delivery item; it is not evidence that the frozen
result is wrong, nor is a path/hash reference an external reconstruction test.

The acceptance check still needed is a fresh extraction with publicly obtained
inputs, followed by the specified model execution and comparison against frozen
results. This check must preserve the existing estimands, seeds, fold rules,
models and inclusion criteria. It requires no new scientific analysis design.


V6.26 delivery note: the historical interface check above was not repeated in this repair. The current model definitions are clarified in main Methods and SM2, and the ordered 50/25-variable dictionary is supplied under reproducibility/. Stored-result checks do not execute the training commands above.


## Dependency closure, 14 September 2026

Ten previously omitted frozen dependencies are now supplied at their original
paths. See reproducibility/DEPENDENCY_CLOSURE_20260914.md for exact scope,
identity checks and the read-only preflight. This does not certify training
or public-source-to-model reconstruction.
