## Supplementary_Software_1 for "Inferential boundaries of age prediction: why prediction does not establish biological age measurement": CURRENT_MANUSCRIPT_VERSION.md

# Review texts for this software package — 22 September 2026

The authoritative review texts are the separately supplied submission files:

| Role | Submission file |
|---|---|
| Main manuscript | `Manuscript_Zotero_20260921.docx` |
| Supplementary Information | `Supplementary_Information.pdf` |

Use the Zotero-labelled DOCX designated by the authors on 22 September 2026.
Its filename includes 21 September; that filename is not a content-freeze date.
These two review documents accompany the submission and are not embedded in
this software archive. Their byte identities at this navigation update are
recorded in [the review-file record](reproducibility/REVIEW_TEXT_FILES_20260922.tsv).

The Markdown manuscript and SI files dated 14 September 2026 under
`manuscript/nature_first/` are historical analysis-era snapshots. They retain
their original text, including any author placeholders, and are not alternative
current review texts. Other dated manuscript and reproducibility documents in
that directory likewise describe their original snapshots. V6.27 / SI V5.19
labels and earlier version identifiers are retained for provenance.

Start with [the software guide](README.md) for verification commands and
[the manuscript index](manuscript/nature_first/README.md) for snapshot roles.
The inventory for this software archive is
[ARCHIVE_MEMBERS_20260922.tsv](reproducibility/ARCHIVE_MEMBERS_20260922.tsv).
`V6_27_ARCHIVE_MEMBERS.tsv` and earlier inventories are unchanged historical
records; they do not inventory the revised navigation files in this archive.

This update changes document navigation only. It does not revise the review
documents, analysis files, numerical results, historical versions or historical
hashes. Access and reuse terms remain as stated in the existing software guide.
