## Supplementary_Software_1 for "Inferential boundaries of age prediction: why prediction does not establish biological age measurement": ENVIRONMENT.md

# Environment / 环境

## Contract

All Python work in this project uses the mamba-managed `lhstate` environment.
The runtime is fail-closed: a missing environment, wrong version, missing
package, or lock drift is an error and must never trigger a fallback to base,
`py`, bare `python`, or the Codex bundled runtime.

The environment has three complementary sources:

- `environment-policy.json`: manager, environment name, Python version, platform,
  and deterministic runtime settings.
- `environment.yml`: exact pins for direct project dependencies.
- `environment.win-64.lock.txt`: exact URLs, builds, and hashes for the complete
  transitive win-64 environment.

The current contract is mamba 2.6.0, environment `lhstate`, Python 3.11.15,
and platform `win-64`.

## Verify and run

From the Aging workspace root:

```powershell
& .\LifeHistoryStateDecomposition\scripts\Test-ProjectEnvironment.ps1
& .\LifeHistoryStateDecomposition\scripts\Invoke-ProjectPython.ps1 -- .\LifeHistoryStateDecomposition\analysis\SCRIPT.py ARGUMENTS
```

For single-cell/Scanpy work, run the slower full import gate once first:

```powershell
& .\LifeHistoryStateDecomposition\scripts\Test-ProjectEnvironment.ps1 -Full
```

The wrapper resolves `lhstate` from `mamba env list`, verifies the interpreter,
all exact direct dependency versions, import availability, and the full explicit
lock, then invokes the resolved `lhstate\python.exe` directly. Direct invocation
avoids fragile shell activation while still using only the mamba-managed runtime.
It also supplies the Windows Conda DLL search paths and directs Numba,
Matplotlib, and joblib caches to the project-local ignored `.runtime-cache`
directory; this prevents native-library crashes and protected-prefix cache hangs.

Do not use `pip install` in this project. Do not work around a missing package by
switching interpreters.

For Codex tasks, `.codex/hooks.json` adds a mechanical pre-command guard that
denies bare Python, alternate Python package managers, Conda, and
`mamba run ... python`. Review and trust this project-local hook when Codex asks;
the approved entry point remains `Invoke-ProjectPython.ps1`.

`gseapy=0.12.0` is intentionally excluded from the win-64 contract: the
Bioconda `noarch` artifact contains a Linux-only compiled extension and cannot
import on Windows. The one enrichment script that can use it already has a
documented Enrichr REST fallback.

## Create or restore

For a human-readable solve using the direct dependency manifest:

```powershell
mamba env create -f .\LifeHistoryStateDecomposition\environment.yml
```

For the exact current win-64 environment:

```powershell
mamba create -n lhstate --file .\LifeHistoryStateDecomposition\environment.win-64.lock.txt
```

## Deliberate dependency updates

1. Edit `environment.yml` and use exact versions.
2. Apply it with mamba:

   ```powershell
   mamba env update -n lhstate -f .\LifeHistoryStateDecomposition\environment.yml
   ```

3. Regenerate the exact lock:

   ```powershell
   & .\LifeHistoryStateDecomposition\scripts\Update-ProjectEnvironmentLock.ps1
   ```

4. Run `Test-ProjectEnvironment.ps1` before any analysis.

If the mamba, Python, environment name, or platform contract changes, update
`environment-policy.json` in the same change.

## Notebook kernel

Register the verified interpreter only through the wrapper:

```powershell
& .\LifeHistoryStateDecomposition\scripts\Invoke-ProjectPython.ps1 -- -m ipykernel install --user --name lhstate --display-name "Python (lhstate)"
```

Always select `Python (lhstate)` and verify the notebook kernel executable before
running cells.

## R / Bioconductor entry

R work remains separate from the Python environment and uses:

```text
D:\APP\R-4.5.1\bin\x64\Rscript.exe
```

Use it only for workflows that require R/Bioconductor (for example GEOquery,
Biobase, limma, edgeR, or DESeq2). Python downstream analysis still uses the
verified `lhstate` wrapper.
