## Supplementary_Software_1 for "Inferential boundaries of age prediction: why prediction does not establish biological age measurement": environment.yml

```
name: lhstate
channels:
  - conda-forge
  - bioconda
  - nodefaults
dependencies:
  - python=3.11.15
  - numpy=2.4.4
  - pandas=2.3.3
  - tabulate=0.9.0
  - scipy=1.17.1
  - scikit-learn=1.8.0
  - statsmodels=0.14.6
  - matplotlib=3.10.9
  - seaborn=0.13.2
  - requests=2.34.1
  - pyyaml=6.0.3
  - tqdm=4.67.3
  - ipykernel=7.2.0
  - lifelines=0.30.3
  - pyarrow=24.0.0
  - polars=1.40.1
  - h5py=3.16.0
  - pytables=3.11.1
  - anndata=0.12.14
  - scanpy=1.11.5
  - mygene=3.2.2
  - goatools=1.6.4
  - openpyxl=3.1.5
  - xlsxwriter=3.2.9
  - python-docx=1.2.0
  - pillow=12.2.0
  - pdf2image=1.17.0
  - pdfplumber=0.11.10
  - pypdf=6.14.2
  - pypdfium2=5.12.1
  - pymupdf=1.27.2.2
  - setuptools=80.9.0
```
