## Supplementary_Software_1 for "Inferential boundaries of age prediction: why prediction does not establish biological age measurement": README.md

# Extended Data figure: individual AgeGap contribution geometry v0.2

Date: 31 August 2026

This is a fit-free submission figure assembled from the frozen and independently
validated `nhanes_individual_agegap_contributions_v0_1` tables. No decoder,
conditional-centring model, endpoint model or matching rule was refitted or
reselected.

## Scientific responsibility

The figure shows that, in one frozen continuous-NHANES linear chronological-age
decoder, near-identical conditionally centred AgeGaps can arise from markedly
different signed contributions from the same 50 measured clinical features.
The scalar therefore does not specify a unique measured-profile realization.

The contributions are exact attributions of this fitted decoder output. They
are conditional on the frozen model, scaling and fold-local centring reference;
they are not causal ageing mechanisms, organ ages, biological-age components
or biological burden. Contribution-vector L2 distances use decoder-output-year
units and are not age differences between participants.

## Panels

- **a** uses all 293 outcome-blind, non-reusing pairs and displays the frozen
  distribution of 50-dimensional contribution-vector distances. The solid and
  dashed lines are the all-pair median and 95th percentile.
- **b** displays frozen pair `P00026`, selected by the prespecified
  representative-distance rule within the 205 display-eligible pairs.
- **c** displays frozen pair `P00092`, selected by the prespecified
  high-distance rule within the same eligible set.

## Files

- `figures/Extended_Data_Figure_4_near_identical_AgeGaps_distinct_measured_contributions.png`
- `figures/Extended_Data_Figure_4_near_identical_AgeGaps_distinct_measured_contributions.svg`
- `figures/Extended_Data_Figure_4_near_identical_AgeGaps_distinct_measured_contributions.pdf`
- `source_data/` contains sanitized aggregate pair metrics, frozen display-pair
  values and signed plotted contributions. Participant identifiers,
  matching-stratum values and endpoint-model outputs are excluded.

Reproducible plotting entry:
`analysis/plot_nhanes_individual_agegap_contributions_extended_data_v0_2.py`.
