## Supplementary_Software_1 for "Inferential boundaries of age prediction: why prediction does not establish biological age measurement": README.md

# Submission Extended Data Figure 5: frozen domain-options comparison

Fit-free precision successor containing the empirical acquisition panels from the frozen v6.6 Figure 4 source package. It verifies and re-renders only predecessor Source Data; no model is loaded or fitted and no random number is generated.

The comparison is limited to five prespecified measurement-domain options in one exploratory NHANES source-target cycle pair. Each domain is one acquisition option; domains differ in measurement count, and no monetary or real-world cost equivalence is assumed. The v6.8 main Figure 4 package and all frozen analytical predecessors remain unchanged.

The underlying frozen resource is named `v6_9_extended_data_figure4_domain_options_v0_1` because it was originally assigned Extended Data Figure 4. In v6.13 it is renumbered only, without analytical, numerical or graphical change, as Extended Data Figure 5.
