## Supplementary_Software_1 for "Inferential boundaries of age prediction: why prediction does not establish biological age measurement": FIGURE_LEGEND_BILINGUAL.md

# Figure legend | 图注

## English

**Figure | HSCT separates chronological coordinates without establishing biological-age identity.**
**a,** Allogeneic haematopoietic stem-cell transplantation separates recipient-host chronological age \(A_R\) from the chronological coordinate of the sampled donor-derived haematopoietic lineage \(A_L=A_D+t\). A fixed recipient-blood score \(S=q(X_B)\) is expressed in years, but that unit does not establish the biological object represented by the score. **b,** Recipient- or pair-level score geometry in Søraas 2019 (19 recipients) and Holland 2024 (61 donor–recipient pairs). The horizontal line denotes \(S=A_L\); the dashed diagonal denotes \(S=A_R\). Filled circles and open triangles distinguish host-older and host-younger directions. **c,** Sign-balanced affiliation \(J_{\mathrm{aff,bal}}\), which equally weights the two age-difference directions. Grey intervals show the 2.5th–97.5th percentiles of the frozen within-sign score-pairing null, diamonds its mean and teal circles the observed statistic. The null preserves score marginals and donor–recipient age geometry, but also breaks score–host-age dependence. The right subpanel shows the observed-minus-null contrast with stratified independent-unit bootstrap 95% intervals. Holland provided the primary estimate (0.626 versus null mean 0.484; contrast interval 0.045–0.209); Søraas supplied separately processed same-direction evidence (0.844 versus 0.697; contrast interval 0.045–0.208). **d,** The observed result is excess donor-lineage affiliation under this score-pairing null, not identification of the sampled lineage as the score's biological-age referent. Chronological-age reconstruction does not by itself establish recipient-organism biological-age identity; the empirical contrast neither identifies nor refutes that identity. It does not rule out host-age-dependent shrinkage or estimate donor influence conditional on host age, and it does not establish a unique lineage biological age, exclude host influence, imply organismal age replacement or prove rejuvenation. The two study programmes partly overlap and are complementary rather than independent replications.

## 中文

**图｜HSCT 分离时间年龄坐标，但不确立 biological-age identity。**
**a，**异基因造血干细胞移植将受者宿主实际年龄 \(A_R\) 与被取样的供者来源造血谱系时间坐标 \(A_L=A_D+t\) 分离。固定的受者血液分数 \(S=q(X_B)\) 以年龄单位表达，但该单位并未确立分数所表征的生物学对象。**b，**Søraas 2019（19 名受者）和 Holland 2024（61 对供受者）的受者/配对层级分数几何。水平线表示 \(S=A_L\)，虚线对角线表示 \(S=A_R\)；实心圆和空心三角分别表示宿主较老和宿主较年轻。**c，**对两个年龄差方向等权的 sign-balanced affiliation \(J_{\mathrm{aff,bal}}\)。灰色区间为冻结的 sign 内分数配对置换零分布 2.5%–97.5% 分位数，菱形为零均值，teal 圆点为观察统计量。该零分布保留分数边际分布及供受者年龄几何，但也破坏分数与宿主年龄之间的依赖。右侧显示观察值减零均值及分层独立单位 bootstrap 95% 区间。Holland 提供主估计（0.626 对零均值 0.484；差值区间 0.045–0.209），Søraas 提供独立处理、同方向的互补证据（0.844 对 0.697；差值区间 0.045–0.208）。**d，**观察结果是该分数配对零分布下的超额供者谱系归属，而不是识别被取样谱系为分数的 biological-age 参照对象。时间年龄重建本身并未确立受者整体的 biological-age identity；该经验对比既不识别也不否定这一 identity。该对比不排除宿主年龄依赖的收缩，也不估计给定宿主年龄后的条件供者影响；它不建立唯一的谱系 biological age、不排除宿主影响、不意味着受者整体年龄被替换，也不证明 rejuvenation。两项研究计划部分重叠，只能作为互补分析而非独立重复验证。

Source Data are supplied as panel-level TSV files in the adjacent `source_data` directory. Panels b/c are byte-identical to v0.4; only a/d wording and layout are revised.
