## Supplementary_Software_1 for "Inferential boundaries of age prediction: why prediction does not establish biological age measurement": README.md

# HSCT age-reconstruction figure v0.5

## Bounded chart contract

- **Question:** what does the frozen HSCT score-pairing result establish, and what biological-age interpretation is not established by age reconstruction?
- **Takeaway:** excess donor-lineage affiliation under the score-pairing null is a measured result; recipient-organism biological-age identity is not established by the age unit. The null also breaks score–host-age dependence, so this is not a conditional donor-attribution test.
- **Chart families and roles:** a, coordinate schematic; b, frozen paired-coordinate scatterplots; c, frozen observed/null estimates and bootstrap intervals; d, task–result–interpretation diagram.
- **Frozen grain:** the same 80 independent units (19 Søraas recipients; 61 Holland donor–recipient pairs), two study summaries and four age-difference strata used by v0.4. The study programmes partly overlap; these are complementary analyses.
- **Permitted changes:** a/d labels and wrapping/box placement, and a matching bilingual legend. No new analysis, fitted object, score, random draw or statistical estimate.
- **Invariant b/c:** original plotting functions, coordinates, axes, ranges, markers, colours, point positions, numerical labels and interval encoding are reused. Quantitative panel TSVs are copied byte-for-byte.
- **Surface:** reproducible static matplotlib figure, 600-dpi PNG plus editable SVG and vector PDF. Keep the v0.4 teal/orange/neutral palette and its shape/line redundancies.
- **QA:** verify copied data byte identity and exact table equality; verify frozen input identity; visually inspect schematic text containment, arrows, quantitative panels and the final PDF. Save the parity report in this directory.

## Reproduction

Run `analysis/plot_hsct_age_reconstruction_semantic_counterexample_v0_5.py` only through the pinned LHState gate/wrapper. The script imports the frozen v0.4 plotting module but never calls its analysis/audit `main()`; only its plotting helpers and unchanged figure assembly are used.

The authoritative analysis remains `hsct_fixed_score_referent_null_probe_v0_1`. This derivative does not change HUM-01's estimand, frozen result, role or placement.
