## Supplementary_Software_1 for "Inferential boundaries of age prediction: why prediction does not establish biological age measurement": VALIDATION_REPORT_v0_5.md

# Data and layout parity validation

Status: PASS

- Quantitative panel b/c TSVs: byte-identical and exact table equality with v0.4.
- Frozen analysis inputs and all original source TSVs: unchanged.
- Original b/c plotting functions and figure assembly: reused unchanged.
- Schematic box labels: contained within their boxes.
- No fitted objects, random numbers or new statistical estimates.

See DATA_PARITY_VALIDATION.json for per-file hashes and exact checks.

## Visual inspection

- Inspected the 600-dpi PNG and the one-page vector PDF rendered with bundled Poppler.
- Panels a/d: all node text is inside its box; no crossed-out identity, clipped label or arrow/text collision. The biological object is not presented as empirically identified or refuted.
- Panels b/c retain their original plot layout and numerical annotations. No point, axis, interval, marker or number was changed.
- The a/d concept table is intentionally revised to match the new labels; it is not included in the quantitative byte-parity claim.
- Final a/d semantic and layout QA: PASS.
