## Supplementary_Software_1 for "Inferential boundaries of age prediction: why prediction does not establish biological age measurement": COVER_LETTER_NATURE_v6_27_20260914.md

# Cover letter

14 September 2026

Dear Editors,

Please consider our manuscript, “Chronological-age decoding does not identify biological age”, as an Article in Nature.

When an age-clock output is interpreted as biological age, success at reconstructing chronological age can become the criterion for selecting measurements, improving models and judging rejuvenation. A biological measurement question is then replaced by an age-prediction task. Our Article establishes why success at that substituted task does not establish that the original biological question has been answered.

We show that chronological-age supervision identifies an age-task statistic, not a biological-age construct. Population-optimal age prediction, gap construction and genuine prognostic value do not supply the missing measurement relation. Shared biological covariance explains how an age-trained score can carry real endpoint information without first having measured a defined biological-age object. The methodological finding concerns the basis for choosing and evaluating a biological representation, not a demand for greater age-prediction accuracy.

The human comparisons begin with useful age-trained scores. In continuous NHANES, an age decoder improved held-out five-year mortality prediction beyond age and background, yet direct modelling of the same source measurements performed better. A separate NHANES III comparison held the measurement panel, scalar dimension, linear capacity and downstream calibration opportunity constant: changing supervision from age to mortality improved prediction. These results distinguish a score's genuine utility from the justification for choosing age reconstruction as the intermediate task. Transplantation provides a complementary referent example: recipient-blood scores showed excess donor-lineage affiliation under the declared score-pairing null.

The manuscript credits the direct precedents on non-identification, composite-score ambiguity, source-measurement comparison, outcome-first modelling and measurement terminology. Its central judgement is that age-task success cannot serve as the criterion for progress in a biological measurement task that age supervision does not define. This changes what researchers should select, optimize and accept as validation. Additional measurement or causal evidence can validate a defined biological construct, including one measured by an already age-trained score; that evidence establishes the relation the age objective leaves open.

We believe this finding warrants Nature’s broad readership because the same inferential step shapes biomarker development across molecular, cellular, organ and clinical settings. Measuring ageing requires the biological question, rather than the reconstruction of time, to define the target and the evidence that counts as success. The Article supplies analytical grounds, human-data comparisons and an estimand-specific research programme for that change.

Code and derivative data are supplied for editorial and reviewer evaluation, with a portable stored-result verification entry point. OpenAI Codex assisted with code development and manuscript preparation, as disclosed in Methods.

Sincerely,

[CORRESPONDING AUTHOR]
