## Supplementary_Software_1 for "Inferential boundaries of age prediction: why prediction does not establish biological age measurement": MANUSCRIPT_NATURE_IDENTIFICATION_FIRST_v6_27_20260914.md

# Chronological-age decoding does not identify biological age

Authors: **[AUTHOR LIST]**  
Affiliations: **[AFFILIATIONS]**  
Correspondence: **[CORRESPONDING AUTHOR AND EMAIL]**

## Summary

Chronological-age clocks reconstruct age from biological measurements, yet their outputs are interpreted as biological age, gaps as ageing acceleration and intervention-associated decreases as rejuvenation<sup>1–5</sup>. We show that age supervision identifies an age-task statistic, not a biological-age construct, and establish how this distinction changes biomarker construction and validation. Even at the population optimum, the same observable distribution and age-prediction performance admit incompatible biological-age interpretations. Squared-error age loss penalizes within-age output dispersion without defining its biological direction. Given age, a gap re-expresses the compressed score; exact age recovery eliminates it despite persistent heterogeneity. Shared biological covariance explains how genuine endpoint prediction can arise without establishing construct identity. In human analyses, recipient-blood scores after allogeneic haematopoietic stem-cell transplantation showed excess donor-lineage affiliation under a score-pairing null. In complementary NHANES analyses, age-trained scores improved held-out five-year mortality prediction beyond age and background, yet direct modelling of source measurements and mortality supervision at matched scalar capacity yielded further gains. These findings place the biological question at the start of biomarker development: the intended object must determine the training target, the representation selected and the evidence that counts as success.

## Main text

Chronological age is familiar, ordered and expressed in years. Molecular and clinical clocks reconstruct it from biological measurements and return a value on the same scale<sup>1–5</sup>. Subtracting known age produces a gap. The common unit makes a further inference feel natural: the score becomes biological age, a positive gap becomes ageing acceleration and an intervention-associated decrease becomes rejuvenation. These quantities are computationally connected. Establishing their biological meaning requires a relation between the number and the condition it is claimed to measure.

That relation matters from the start of model development. Biological-age research seeks to resolve differences that chronological age leaves open: physiological condition among same-age people, change in that condition and improvement under intervention. Age reconstruction answers a different question: which combination of measurements best predicts the age label? Treating its output as a biological coordinate lets age-prediction accuracy govern measurement selection and model improvement, while the resulting ordering determines who appears biologically older and which changes appear beneficial. The inferential substitution occurs when success at reconstructing age is accepted as success at measuring the proposed state.

Contemporary proteomic and single-cell studies continue to assign biological-age, acceleration and rejuvenation meanings to age-valued outputs, gaps and score movements (Table 1 and Supplementary Data 1)<sup>3–5</sup>. Their prognostic and functional findings can be real and useful. Such successes make the biological interpretation compelling, while leaving a further question: what relation establishes that this particular score measures the proposed ageing state? A clock that predicts both age and mortality makes that question especially consequential.

Here we establish how the age objective constrains the construction and validation of an ageing measure. It selects a statistic for reconstructing age, evaluates same-age differences in its output as prediction error and leaves their biological interpretation unidentified even at the population optimum. Gap construction re-expresses that statistic. Shared biology explains how it can predict a health endpoint without acquiring biological-age identity. The methodological consequence is that the biological question must determine which representation is selected and which evidence establishes its meaning.

Earlier work questioned the construction and interpretation of clocks, established cross-sectional non-identifiability and challenged biological-age language<sup>6–9</sup>. Calibration, uncertainty and validation studies address further aspects of their use<sup>10–16</sup>. The question here concerns the justification for selecting a biological measurement, including when an age-trained score has genuine prognostic value. Our analytical results establish what the age objective selects and what its success can mean; human comparisons make the consequences for score interpretation and representation choice concrete.

### Age supervision selects a statistic of chronological age

Let \(A\) denote chronological age, \(W\) background variables and \(X\) the measured profile (Fig. 1). Under squared-error loss, the unrestricted population-optimal age-prediction model, or decoder, is

\[
q_A^*(W,X)=E(A\mid W,X).
\]

It returns the average chronological age among people with the same measured profile and background in the reference population. A restricted model supplies the best approximation within its class; another loss selects a different summary, such as a conditional median. The loss, model class and reference population therefore define an age-task statistic. Biological ranking, future risk and improvement require evidence for those objects.

For same-age heterogeneity, the mismatch is visible in the loss itself:

\[
\begin{aligned}
R_A(q)&=E[\{q(W,X)-A\}^2]\\
&=E[\operatorname{Var}\{q(W,X)\mid A,W\}
+\{E[q(W,X)\mid A,W]-A\}^2].
\end{aligned}
\]

The first term penalizes differences in decoded age among people of the same age and background; the second measures conditional calibration error. This is appropriate for reconstructing age. The methodological problem arises when that criterion is used to judge measurement of a different biological object. Same-age output differences can remain because the same measurements improve reconstruction across ages, but their biological direction receives no supervision: variation interpreted as greater damage, reserve or resilience enters the objective as age-prediction error. This is a property of the scalar output, not elimination of heterogeneity from \(X\). Selecting a model by age accuracy therefore supplies no criterion for deciding which same-age differences represent deterioration or improvement.

The identification problem extends beyond this loss decomposition. Hold fixed the entire observed distribution of ages, backgrounds and measurements. Different latent biological states can be attached to that distribution while preserving the optimal age decoder and attainable prediction performance, yet assigning different same-age rankings and mechanisms to the observations. Age decoding cannot select among these compatible interpretations (Proposition 1; Supplementary Note 1). This argument builds on the observational-equivalence construction of Sluiskes and colleagues<sup>8</sup>. Its consequence holds at the population optimum, after finite-sample estimation error has disappeared.

Age anchoring also leaves biological orientation open. For a measured contrast \(U\) centred within age and background, \(A+U\) and \(A-U\) have the same age centre, within-age variance and absolute departure from age, with opposite within-age rankings. Both numbers are computable. What remains to be established is which, if either, has the measurement relation claimed for a biological state. An independently defined endpoint or anchored measurement model can supply orientation through its additional evidence. The same question applies to cellular, organ and organismal ages: specifying the compartment locates a proposed state; a measurement relation establishes what the score measures there.

### Age gaps preserve the score's information and its identification limits

For a fixed decoder, the raw gap is \(G=q_A-A\), so \(q_A=G+A\). Once age and background are known, the score, raw gap and conditionally centred gap carry the same individual information. Subtraction changes the reference point. It introduces neither a new observation nor a biological definition.

Featurewise conditioning, \(X-E(X\mid A,W)\), instead retains a profile of same-age deviations before scalar compression. This distinction matters because post-decoding gaps cannot recover directions discarded by scalarization. If the scientific question concerns those differences, they must be preserved and evaluated in the representation used to answer it.

A linear decoder assigns the same score to profiles that differ along its null space. For a differentiable nonlinear decoder, a gradient describes local first-order sensitivity; orthogonality to that gradient does not rule out higher-order dependence. These geometries concern which profile differences the output distinguishes, not whether the score has a biological identity (Supplementary Note 1).

Two limiting cases separate gap variation from biological-state information. Noisy age measurements can produce non-zero optimal gaps with no additional state in the generating system. Conversely, a profile containing exact age and a separate state yields perfect age recovery and a zero gap, although the state remains fully observed. Gap variation does not establish state information, and a zero gap does not imply its absence from the source measurements. The algebraic limits establish these non-implications; a frozen three-variable construction illustrates age readability and state prediction as distinct tasks (Extended Data Fig. 1).

As squared age-prediction error approaches zero, raw and conditionally centred gaps approach zero in mean square. This is an amplitude result, not a claim that endpoint information decreases monotonically as accuracy improves. Under imperfect prediction, gaps combine conditional age ambiguity, reference-population differences, model restriction, estimation and calibration. The decomposition separates these statistical terms algebraically; age-prediction accuracy does not identify their individual contributions or biological meaning (Supplementary Note 1).

### Shared biology explains genuine prediction without construct identity

An age-trained score can predict mortality because age-readable and mortality-relevant biology covary. Development, exposure, disease, adaptation, cell composition, cohort and selective survival shape the measured profile. The combination that reconstructs age can overlap the combination that predicts a named endpoint, including within age strata. This explains genuine prognostic value while leaving the identity of a proposed biological-age construct unresolved.

A fixed clock is a deterministic function of its source measurements and background. For any declared endpoint and loss, optimal population risk using the source profile, age and background is no greater than optimal risk using the clock, age and background<sup>17–19</sup>. This gives a direct benchmark for the compressed representation. A scalar can attain equality when it is predictively sufficient for that target and loss. Under log loss, equality requires the source profile to add no endpoint information after conditioning on the clock, age and background. Age supervision neither imposes nor tests that condition. Finite-sample regularization, assay cost and transport can still favour a scalar, but those advantages are properties to establish for its intended use.

Validation must establish the relation claimed for the score: evidence for prognosis, change in a readout and restoration of a biological state answer different questions<sup>20</sup>. With the identification boundary established analytically, we use human data to examine its consequences for the referent of an age-valued score, the choice of a predictive representation and the interpretation of an individual gap.

### Transplantation separates the candidate referents of a blood age score

Allogeneic haematopoietic stem-cell transplantation separates the recipient organism from the donor-derived lineage producing the assayed blood (Fig. 2)<sup>21,22</sup>. The recipient retains the host's chronological history, while the lineage carries another age coordinate. We compared recipient age with donor age at transplantation plus elapsed follow-up. The latter describes lineage chronology, not the literal age of short-lived circulating cells.

We reproduced the reported-score direction in Søraas and colleagues and applied fixed Horvath-2013 coefficients with external-mean imputation to author-normalized Holland data. The primary Holland population contained 118 eligible occasions from 61 donor–recipient pairs. A sign-balanced affiliation index compared the score's distances to recipient and donor-lineage coordinates. Within each age-difference direction, a score-pairing permutation preserved score marginals and the observed age-coordinate pairs while breaking their pairing with the scores.

Holland affiliation was 0.626, above the null mean of 0.484 and its 97.5th percentile of 0.577 (one-sided finite-permutation \(P=8.50\times10^{-4}\)). The bootstrap interval for observed-minus-null affiliation was 0.045–0.209, with positive contrasts in both age-difference directions. Separately processed Søraas scores showed the same direction. The programmes partly overlap and provide complementary evidence. The estimand is excess affiliation relative to broken score pairing. Because this permutation also breaks the score–recipient-age relationship, it does not exclude host-age-dependent shrinkage or estimate donor influence conditional on recipient age.

The comparison makes the referent problem concrete: recipient-blood scores showed structured affinity to donor-lineage chronology under the declared test. The year unit itself supplies no rule assigning that readout to the recipient organism or to a lineage biological age. Establishing either interpretation requires a measurement relation for the proposed referent.

### Mortality-directed modelling improves on useful age-trained scores

We next compared representations for a named endpoint, beginning with an age decoder that improved held-out five-year mortality prediction beyond age and background (Fig. 3a,b). Among 13,724 participants aged 20–79 years in NHANES 2007–2012, including 470 five-year deaths, a ridge model reconstructed age from background and 50 clinical measurements. Age was its only supervised label. Mortality models used age and background, added the cross-fitted age score, or directly modelled the same source measurements. Every contrast used the same held-out participants.

The clock reduced mortality log loss by 0.003774 beyond age and background (95% fixed-prediction bootstrap interval, 0.002046–0.005474). Direct modelling reduced loss by a further 0.009152 beyond the clock (0.006276–0.012034). At the point estimates, the additional direct-source gain was more than twice the clock's entire improvement over age and background. Both contrasts had the same direction in all five outer folds, and prespecified nonlinear recalibration of the clock left the direct-source advantage essentially unchanged.

Lower log loss means better probabilistic prediction. The direct-source comparison changes both the information supplied to the endpoint model and its dimension. To separate dimension from supervision target, we used a separate NHANES III comparison of two one-dimensional linear projections from the same 25-measurement panel (Fig. 3c,d). Both were trained in Phase 1: one reconstructed age and the other predicted five-year mortality. We froze their directions and gave them identical downstream calibration opportunities in survey-weighted Phase 2 data comprising 7,774 participants and 392 deaths. Phase 2 outcomes did not enter projection fitting. Inputs, scalar dimension, linear capacity and downstream opportunity were matched.

The age-supervised scalar again added mortality-prediction value beyond age and background: the Phase 2 log-loss gain was 0.002792 (95% Fay-BRR interval, 0.000239–0.005345). Mortality supervision added a further 0.005711 over the age-supervised scalar (0.002093–0.009329), positive in all five outer folds. Redirecting the same scalar capacity added approximately twice the age-trained scalar's baseline gain at the point estimates. A separate matched substitution comparison found no clear advantage from replacing observed age with the transported age decoder (Extended Data Fig. 3). A supporting analysis shows that observed age's own predictive increment increased when an age-selected measurement block was withheld (Extended Data Fig. 3; Supplementary Methods SM3).

Both comparisons began with useful age-trained scores, yet direct mortality modelling improved prediction, including at matched scalar capacity. The designs have different evaluation boundaries: NHANES III was a retrospective protocol transfer with previously accessed outcomes, and its Fay-BRR intervals incorporate Phase 2 survey-design refitting conditional on frozen Phase 1 directions; the continuous-NHANES intervals condition on stored out-of-fold predictions. The direct-modelling question was raised explicitly by Kriukov and colleagues<sup>6</sup>. In the declared populations and model families, these results show why prognostic success does not settle whether age reconstruction should be the intermediate task.

### Near-identical AgeGaps have different measurement contributions

The frozen age decoders from continuous NHANES also allowed individual outputs to be resolved into their measurement contributions. We decomposed each conditionally centred AgeGap into 50 exactly additive terms and formed 293 outcome-blind, non-reusing pairs matched exactly on fold, chronological age, sex, race or ethnicity and survey cycle. Within each pair, gaps differed by at most 0.25 decoder-output years (Extended Data Fig. 4).

The median gap difference was 0.124, whereas the median Euclidean distance between contribution vectors was 24.602 decoder-output years and median cosine similarity was 0.003. The distance describes separation in the fitted model's contribution space, not an age difference between people. Near-identical gaps arose from markedly different signed measurement contributions. The scalar specified their sum while leaving the contributing profile unresolved. These are exact attributions of the frozen model output, rather than causal components of ageing.

### The biological target guides measurement and model selection

The choice of target changed measurement selection before a score was constructed. Across five prespecified measurement-domain options in NHANES 2007–2008, mortality-directed selection chose hepatic, protein and injury measurements, whereas an outcome-blind inverse age decoder chose metabolic, adiposity and lipid measurements. We evaluated both frozen choices in NHANES 2009–2010 (Fig. 4b; Extended Data Fig. 5). The endpoint-selected domain improved mortality log loss by 0.006418 (95% fixed-prediction bootstrap interval, 0.002297–0.010652); the age-selected domain improved it by 0.000646 (−0.001195–0.002481). Their paired difference was 0.005772 (0.001380–0.010390), positive in four of five folds.

Selecting measurements for age reconstruction and selecting them for mortality were therefore materially different decisions in this menu. The comparison is exploratory cross-cycle evidence; domains differed in measurement count and no equivalence of acquisition cost was assumed. The result illustrates how a scientific target can guide both measurement acquisition and subsequent modelling directly.

A target-first programme begins with the object to be measured and the decision to be supported (Fig. 4a,c). Same-age state calls for a defined measurement model. Named risk calls for an endpoint, horizon and loss. Transition calls for repeated observations and a dynamic estimand. Intervention displacement calls for a causal contrast on a specified measurement. Benefit calls for a clinical outcome or qualified surrogate. These definitions specify which differences matter and which comparisons can establish progress; the acquisition example demonstrates this approach for the named risk task.

### Discussion

The methodological error begins when age-reconstruction success is used to justify a representation as a measure of a biological state that the age task does not define. We show why population-optimal prediction, gap construction and genuine prognostic value do not establish the missing measurement relation. Shared biological covariance explains how the prognostic success can arise, while human comparisons show that changing the target changes the representations and measurements selected. The consequence is a change in the criterion for progress in ageing measurement: age-prediction accuracy cannot stand in for evidence that the proposed biological object has been measured.

Sluiskes and colleagues established observational non-identifiability and examined age-based marker selection<sup>8</sup>, while Ikram challenged prediction-error interpretation and clocks as disease explanations<sup>7</sup>. Kriukov and colleagues advocated direct outcome modelling and prospective target definition<sup>6</sup>; Johnson and Shokhirev addressed biological-age terminology<sup>9</sup>. Grødem and colleagues analysed the mismatch between age prediction and individual brain change and proposed reorienting prediction targets<sup>23</sup>. Hu's *House of Clocks* showed that different feature profiles can yield the same composite score and argued for prediction benchmarks using the source measurements<sup>24</sup>. Detailed source comparisons are provided in Supplementary Note 2, with the 20-source structured review reported in Supplementary Table 1 and Supplementary Data 2. We specify what the age objective identifies, what gap operations preserve and what genuine prognostic success can establish, and test representation choice with source measurements and scalar capacity held fixed. These results make the training target and validation criterion part of the measurement claim itself.

The human comparisons show why this distinction matters in practice. Transplantation separates host and assayed lineage as candidate referents; near-identical gaps summarize different measurement contributions; and mortality-directed representations outperformed useful age-trained predictors, including at matched scalar capacity. Genuine predictive success makes the biological interpretation persuasive without supplying its measurement relation. Field-scale studies report clock-specific disease associations, heterogeneous social gradients and discordant age-valued estimates<sup>25–29</sup>, while longitudinal and intervention studies establish prediction and response in particular settings<sup>30–33</sup>. Those findings support the relations tested and can contribute to construct validation when a measurement model specifies that relation. A claim of rejuvenation calls for a defined biological state and evidence of its restoration; demonstrated functional or clinical benefit stands on its own outcome evidence.

Target-directed models already provide constructive starting points. Morbidity, mortality, lifespan and physiological-dynamics supervision select predictors for those outcomes<sup>34–40</sup>. These objectives differ from pure age reconstruction and must be assessed according to their own targets and evidence. Anchored deterioration models, causal constraints and compartment-specific perturbations can support biological constructs through added structure<sup>41–43</sup>. Such evidence can also validate a previously age-trained score for a defined object; its measurement validity then comes from that evidence. The population-optimal argument separates this identifying work from improvements in estimation, calibration or reliability. Chronological-age estimation remains a valid task, and fixed scores can have practical advantages in finite samples, cost or transport. The scope of the human comparisons lies in their declared targets, populations and estimators; they do not settle the existence of a low-dimensional ageing state.

A change of name leaves the age objective intact; a change of target alters which variation is selected, which measurements are acquired and which outcomes validate the result. Chronological-age decoding does not identify biological age. Its output scale can express a separately established quantity. Measuring ageing requires the biological question, rather than the reconstruction of time, to define the target and the evidence that counts as success.

## References

1. Horvath, S. DNA methylation age of human tissues and cell types. *Genome Biol.* **14**, R115 (2013). https://doi.org/10.1186/gb-2013-14-10-r115
2. Hannum, G. *et al.* Genome-wide methylation profiles reveal quantitative views of human aging rates. *Mol. Cell* **49**, 359–367 (2013). https://doi.org/10.1016/j.molcel.2012.10.016
3. Robinson, O. *et al.* Associations of proteomic age clocks with lifestyle risk factors, incident chronic diseases and mortality in two European cohorts. *Nat. Aging* **6**, 1437–1451 (2026). https://doi.org/10.1038/s43587-026-01163-6
4. Ding, D. Y. *et al.* Plasma proteomic signatures of cellular aging predict human disease. *Nat. Med.* **32**, 2060–2072 (2026). https://doi.org/10.1038/s41591-026-04446-y
5. Li, W. *et al.* Single-cell immune aging clocks reveal inter-individual heterogeneity during infection and vaccination. *Nat. Aging* **5**, 607–621 (2025). https://doi.org/10.1038/s43587-025-00819-z
6. Kriukov, D. *et al.* Do we actually need aging clocks? *npj Aging* **12**, 15 (2026). https://doi.org/10.1038/s41514-025-00312-2
7. Ikram, M. A. The use and misuse of ‘biological aging’ in health research. *Nat. Med.* **30**, 3045 (2024). https://doi.org/10.1038/s41591-024-03297-9
8. Sluiskes, M. H. *et al.* Clarifying the biological and statistical assumptions of cross-sectional biological age predictors: an elaborate illustration using synthetic and real data. *BMC Med. Res. Methodol.* **24**, 58 (2024). https://doi.org/10.1186/s12874-024-02181-x
9. Johnson, A. A. & Shokhirev, M. N. Contextualizing aging clocks and properly describing biological age. *Aging Cell* **23**, e14377 (2024). https://doi.org/10.1111/acel.14377
10. Dabrowski, J. K. *et al.* Probabilistic inference of epigenetic age acceleration from cellular dynamics. *Nat. Aging* **4**, 1493–1507 (2024). https://doi.org/10.1038/s43587-024-00700-5
11. Wyss-Coray, T. & Topol, E. J. Biological aging clocks in health and disease. *Nat. Med.* **32**, 2383–2394 (2026). https://doi.org/10.1038/s41591-026-04495-3
12. Meyer, D. H. & Schumacher, B. Aging clocks based on accumulating stochastic variation. *Nat. Aging* **4**, 871–885 (2024). https://doi.org/10.1038/s43587-024-00619-x
13. Tong, H. *et al.* Quantifying the stochastic component of epigenetic aging. *Nat. Aging* **4**, 886–901 (2024). https://doi.org/10.1038/s43587-024-00600-8
14. Moqri, M. *et al.* Validation of biomarkers of aging. *Nat. Med.* **30**, 360–372 (2024). https://doi.org/10.1038/s41591-023-02784-9
15. Sehgal, R. *et al.* Biological versus technical reliability of epigenetic clocks and implications for disease prognosis and intervention response. *Aging Cell* **25**, e70635 (2026). https://doi.org/10.1111/acel.70635
16. Apsley, A. T., Etzel, L., Ye, Q. & Shalev, I. From population science to the clinic? Limits of epigenetic clocks as personal biomarkers. *Epigenomics* **17**, 1447–1461 (2025). https://doi.org/10.1080/17501911.2025.2603880
17. Blackwell, D. Equivalent comparisons of experiments. *Ann. Math. Stat.* **24**, 265–272 (1953). https://doi.org/10.1214/aoms/1177729032
18. Cover, T. M. & Thomas, J. A. *Elements of Information Theory*, 2nd edn (Wiley, 2006). https://doi.org/10.1002/047174882X
19. Dawid, A. P. Conditional independence in statistical theory. *J. R. Stat. Soc. B* **41**, 1–31 (1979). https://doi.org/10.1111/j.2517-6161.1979.tb01052.x
20. FDA–NIH Biomarker Working Group. *BEST (Biomarkers, EndpointS, and other Tools) Resource* (National Center for Biotechnology Information, 2021). https://www.ncbi.nlm.nih.gov/books/NBK338448/
21. Søraas, A. *et al.* Epigenetic age is a cell-intrinsic property in transplanted human hematopoietic cells. *Aging Cell* **18**, e12897 (2019). https://doi.org/10.1111/acel.12897
22. Holland, P. *et al.* Epigenetic aging of human blood cells is influenced by the age of the host body. *Aging Cell* **23**, e14112 (2024). https://doi.org/10.1111/acel.14112
23. Grødem, E. O. S. *et al.* Reconsidering Brain Age: Why Age-Prediction Models Fail as Measures of Brain Aging. *medRxiv* (2026). https://doi.org/10.64898/2026.05.07.26352620
24. Hu, I. House of Clocks: On the Misuse of Ageing Composite Measures. *bioRxiv* (2025). https://doi.org/10.1101/2025.05.24.655934
25. Mavrommatis, C. *et al.* An unbiased comparison of 14 epigenetic clocks in relation to 174 incident disease outcomes. *Nat. Commun.* **16**, 11164 (2025). https://doi.org/10.1038/s41467-025-66106-y
26. Willems, Y. E. *et al.* Social determinants of health and epigenetic clocks: a systematic review and meta-analysis of 140 studies. *Nat. Hum. Behav.* (2026). https://doi.org/10.1038/s41562-026-02477-6
27. Du, Z. *et al.* The OmniAge compendium of aging omic biomarkers links mitotic clocks to clonal hematopoiesis and causality. *Nat. Commun.* **17**, 9130 (2026). https://doi.org/10.1038/s41467-026-76038-w
28. Vetter, V. M. *et al.* Comprehensive cross-sectional and longitudinal comparison of sixteen markers of biological aging from the Berlin Aging Study II. *Commun. Med.* **6**, 168 (2026). https://doi.org/10.1038/s43856-025-01233-7
29. Johnson, A. A. & Shokhirev, M. N. Understanding and making sense of epigenetic age misalignment across different aging clocks. *Epigenomics* **18**, 965–972 (2026). https://doi.org/10.1080/17501911.2026.2708567
30. Kuo, P.-L. *et al.* Longitudinal changes in epigenetic clocks predict survival in the InCHIANTI cohort. *Nat. Aging* **6**, 534–540 (2026). https://doi.org/10.1038/s43587-026-01066-6
31. Mishra, M. *et al.* Exoproteome of calorie-restricted humans identifies complement deactivation as an immunometabolic checkpoint reducing inflammaging. *Nat. Aging* **6**, 1064–1078 (2026). https://doi.org/10.1038/s43587-026-01107-0
32. Sehgal, R. *et al.* Responsiveness of epigenetic aging biomarkers to longevity interventions in humans. *Nat. Med.* (2026). https://doi.org/10.1038/s41591-026-04562-9
33. Li, S. *et al.* Effects of daily multivitamin–multimineral and cocoa extract supplementation on epigenetic aging clocks in the COSMOS randomized clinical trial. *Nat. Med.* **32**, 1012–1022 (2026). https://doi.org/10.1038/s41591-026-04239-3
34. Chen, Q. *et al.* OMICmAge quantifies biological age by integrating multi-omics with electronic medical records. *Nat. Aging* **6**, 722–737 (2026). https://doi.org/10.1038/s43587-026-01073-7
35. Tyshkovskiy, A. *et al.* Universal transcriptomic hallmarks of mammalian ageing and mortality. *Nature* **654**, 173–188 (2026). https://doi.org/10.1038/s41586-026-10542-3
36. Levine, M. E. *et al.* An epigenetic biomarker of aging for lifespan and healthspan. *Aging* **10**, 573–591 (2018). https://doi.org/10.18632/aging.101414
37. Lu, A. T. *et al.* DNA methylation GrimAge strongly predicts lifespan and healthspan. *Aging* **11**, 303–327 (2019). https://doi.org/10.18632/aging.101684
38. Lu, A. T. *et al.* DNA methylation GrimAge version 2. *Aging* **14**, 9484–9549 (2022). https://doi.org/10.18632/aging.204434
39. Belsky, D. W. *et al.* DunedinPACE, a DNA methylation biomarker of the pace of aging. *eLife* **11**, e73420 (2022). https://doi.org/10.7554/eLife.73420
40. Sluiskes, M. H. *et al.* The AccelerAge framework: a new statistical approach to predict biological age based on time-to-event data. *Eur. J. Epidemiol.* **39**, 623–641 (2024). https://doi.org/10.1007/s10654-024-01114-8
41. Li, J. *et al.* Multimodal clocks of human aging. *Cell* **189**, 4489–4511.e31 (2026). https://doi.org/10.1016/j.cell.2026.04.025
42. Ying, K. *et al.* Causality-enriched epigenetic age uncouples damage and adaptation. *Nat. Aging* **4**, 231–246 (2024). https://doi.org/10.1038/s43587-023-00557-0
43. Beltrán-Sánchez, H. *et al.* Modeling biological age and its link with the aging process. *PNAS Nexus* **1**, pgac135 (2022). https://doi.org/10.1093/pnasnexus/pgac135
44. CDC/NCHS. National Death Index linked mortality files: 2019 public-use files. https://www.cdc.gov/nchs/linked-data/mortality-files/index.html
45. CDC/NCHS. NHANES questionnaires, datasets and related documentation. https://wwwn.cdc.gov/nchs/nhanes/continuousnhanes/
46. CDC/NCHS. Third National Health and Nutrition Examination Survey (NHANES III), 1988–1994: data and documentation. https://wwwn.cdc.gov/nchs/nhanes/nhanes3/default.aspx
47. CDC/NCHS. Ethics Review Board approval for the National Health and Nutrition Examination Survey. https://www.cdc.gov/nchs/nhanes/about/erb.html
48. Watanabe, K. *et al.* Multiomic signatures of body mass index identify heterogeneous health phenotypes and responses to a lifestyle intervention. *Nat. Med.* **29**, 996–1008 (2023). https://doi.org/10.1038/s41591-023-02248-0

## Table

### Table 1 | Contemporary construct transfer, available evidence and the missing identifying bridge

Rows locate three contemporary construct transfers and distinguish the evidence supplied by age training from the additional validation reported in the cited studies. These validations support the relations actually tested; the missing bridge concerns the further biological interpretation. Full source records and selection rules are provided in Supplementary Data 1.

| Construct transfer | Training and additional evidence supplied | Missing measurement relation | Supported object and wording |
| --- | --- | --- | --- |
| Age-decoder output → biological, organ, tissue or cellular age | Age reconstruction with external-cohort evaluation; proteomic disease and mortality associations<sup>3</sup>; cell-associated disease, cognitive and pathological associations<sup>4</sup>; external immune-cell age validation and measured immune-response associations<sup>5</sup> | A measurement relation linking the score to the specified biological state and referent; predictive and functional associations require interpretation according to the relation tested | An age predictor and its demonstrated prognostic, functional or cell-associated relationships |
| Decoder residual or gap → ageing acceleration or deceleration | A reference-relative score contrast; proteomic disease associations and independently replicated mortality associations<sup>3</sup>; immune-cell differences associated with infection severity and recovery over time<sup>5</sup> | Evidence that the contrast measures a defined state or change, addressing relevant alternative explanations; a rate claim requires evidence for the temporal process, not just a cross-sectional offset | A reference-relative score contrast with demonstrated prognostic associations or temporal responses |
| Intervention-associated score decrease → rejuvenation or benefit | Pre/post-vaccination score differences associated with ex vivo cytokine responses, transcriptional programmes and cell-composition changes; an additional MMR/placebo comparison<sup>5</sup> | A defined rejuvenated immune state and evidence linking the response to its restoration; clinical benefit requires its own outcome or qualified-surrogate evidence | A vaccination-associated score response and its measured immune-response, molecular and cell-composition associations |

The cellular examples in ref. 4 concern age-trained cell-type scores, not the separately mortality-trained risk composite. External age validation, prognostic replication and measured immune responses are retained as substantive evidence; none is treated as merely another measure of age-fitting accuracy.

## Figure legends

### Figure 1 | From age-task identification to genuine downstream success without construct identity

**a**, Task–object mismatch. The supervised inverse task reconstructs \(A\) from \(X,W\), whereas same-age state, future risk, longitudinal transition and intervention benefit require their own defining structures. **b**, Observable-law non-identification and within-age orientation. The same \(P(A,W,X)\), population-optimal \(q_A(W,X)\) and attainable age-prediction performance admit incompatible latent-state extensions; even age-centred, equal-scale candidates \(H^{(+)}=A+U\) and \(H^{(-)}=A-U\) can reverse the ordering of the same contrast. **c**, Post-decoding operations. For a fixed decoder and centring rule, the score, raw gap and conditionally centred gap are mutually recoverable given observed age and background; these operations cannot recover information discarded by the decoder. Perfect age recovery collapses the gap even when within-age heterogeneity remains. **d**, Shared covariance explains genuine downstream success: age-readable and endpoint-relevant directions can overlap, so a named association can be real without establishing construct identity. **e**, Source measurements cannot have greater population Bayes risk than their deterministic clock under the same declared target and loss. Equality for one loss is loss-specific; equality of the complete target law requires predictive sufficiency. Analytical results establish identification limits, human analyses establish materiality and endpoint-specific analyses motivate alternatives.

### Figure 2 | Haematopoietic stem-cell transplantation exposes referent ambiguity in an age-valued blood readout

**a**, Allogeneic haematopoietic stem-cell transplantation separates recipient-host chronological age \(A_R\) from the chronological coordinate of the sampled donor-derived haematopoietic lineage \(A_L=A_D+t\). A fixed recipient-blood score \(S=q(X_B)\) is expressed in years, but its numerical unit does not specify which referent it indexes. **b**, Recipient- or pair-level score geometry in Søraas 2019 and Holland 2024. The horizontal line denotes the lineage referent \(S=A_L\); the dashed diagonal denotes the host referent \(S=A_R\). Filled circles and open triangles distinguish host-older and host-younger directions. **c**, Sign-balanced affiliation \(J_{\mathrm{aff,bal}}\): the affiliation index is evaluated after averaging scores and chronological coordinates within each pair (Holland) or recipient (Søraas), and the two age-difference directions are then equally weighted. These pair-summary indices, \(J_{\mathrm{pair}}\), are computed before sign balancing and differ from the occasion-first indices underlying Extended Data Fig. 2a and the upper rows of 2b. Grey intervals show the frozen within-sign geometry-preserving score-pairing null, diamonds its mean and teal circles the observed statistic; the right subpanel shows observed-minus-null contrasts with stratified independent-unit bootstrap intervals. Holland provides the primary estimate and Søraas separately processed complementary evidence. The permutation also breaks the score–recipient-age relationship; it does not adjust for host-age-dependent shrinkage. **d**, The observed result is excess donor-lineage affiliation under that score-pairing null. Recipient-organism biological-age identity is not established, rather than empirically ruled out; lineage biological-age identity is also not established. Host influence remains compatible with this finding. The programmes partly overlap and are complementary rather than independent replications.

### Figure 3 | Mortality prediction from source measurements and matched scalar representations

**a**, Nested comparison in continuous NHANES 2007–2012, using 13,724 participants with resolved five-year mortality status, including 470 deaths. A fold-local ridge decoder reconstructed chronological age from background and 50 measurements. Mortality models used age and background alone, added the cross-fitted age score, or directly modelled the same source measurements. **b**, Paired held-out log-loss contrasts show the clock's gain beyond age and background and the further direct-source gain over linear and prespecified flexible clock calibration. Lines are 95% bootstrap intervals conditional on stored out-of-fold predictions. **c**, Separate NHANES III matched-scalar design. Age- and mortality-supervised linear projections used the same 25 measurements and background in Phase 1 (7,558 participants; 402 deaths). The fitted directions were frozen before application to Phase 2 (7,774 participants; 392 deaths), where they received identical fold-local endpoint calibration and held-out evaluation. Phase 2 outcomes did not enter projection fitting. **d**, Design-weighted Phase 2 log-loss contrasts. \(U_A\) is the gain from the age-supervised scalar beyond observed age and background; \(\Delta\) is the further gain from mortality supervision at matched scalar capacity. Lines are 95% Fay-BRR intervals incorporating Phase 2 downstream refitting conditional on the frozen Phase 1 directions. Positive values favour the representation named in each contrast. The NHANES III analysis is a retrospective, outcome-accessed protocol transfer, not independent confirmation. The separate age-substitution and measurement-removal comparisons are retained in Extended Data Fig. 3. Lower log loss means better probabilistic prediction, not clinical benefit or biological-age identification.

### Figure 4 | Target-first evidence replaces construct transfer from an age unit

**a**, A target-first programme begins with the scientific question and declared object, then chooses measurements, design and supervision, evaluates the resulting statistic and makes the claim supported by that evidence. **b**, Exploratory acquisition comparison for five-year all-cause mortality. In NHANES 2007–2008 (4,599 participants; 174 deaths), endpoint-directed selection chose hepatic, protein and injury measurements from five prespecified domains, whereas inverse age decoding chose metabolic, adiposity and lipid measurements. In NHANES 2009–2010 (4,896 participants; 157 deaths), their held-out log-loss gains beyond age and background were 0.006418 and 0.000646; the paired difference was 0.005772. Lines are 95% intervals from 5,000 event-stratified participant bootstraps conditional on fixed source selections and stored predictions. Domains differed in measurement count and were not matched for acquisition cost. The complete policy comparison is retained in Extended Data Fig. 5. **c**, Object-specific evidence map. A measured or latent state requires an anchored measurement model and state definition; named risk requires an endpoint, horizon, loss and validation population; transition requires longitudinal sampling and a transition model; intervention displacement requires a prespecified perturbation contrast and measurement response; and benefit requires a causal estimand, named outcome and decision context. The acquisition example illustrates the named-risk row, not empirical validation of every object in the map. Age-equivalent units may re-express a separately established quantity; the defining evidence supplies its meaning.

### Extended Data Figure 1 | Age-coordinate geometry and task-specific model performance

**a–c**, Frozen cross-sectional measurements X1–X3 plotted against chronological age. Each point is a different individual observed once; connected symbols are state-specific means within fixed age bins and are not longitudinal trajectories. X1 is a noisy age-readable channel without a designed state term; X2 is the declared state channel after exact projection from the realized intercept and linear age direction; X3 mixes linear and nonlinear age structure with state. **d**, Fixed five-fold out-of-fold task map. The horizontal axis is pooled chronological-age decoding \(R^2\). The vertical axis is the reduction in pooled state-decoding log loss obtained by adding the indicated measurement to the fixed age-only model. The display distinguishes age readability, same-age state value and their mixture under the declared models; it is explanatory rather than population inference, longitudinal evidence or a second proof of the analytical result.

### Extended Data Figure 2 | HSCT referent-null sensitivities preserve the interpretation boundary

**a**, Holland occasion-first affiliation, \(J_{\mathrm{occ}}\), across age-difference thresholds, follow-up selections and leave-one-batch-out analyses. Occasion indices are averaged within pair and then across pairs, except for the explicitly labelled within-pair median sensitivity. **b**, The upper rows show occasion-first host-older and host-younger means; the lower rows show the sign-balanced pair-summary observed-minus-null contrasts used in Fig. 2c. Aggregation order as well as final stratum weighting therefore differs between these summaries. **c**, Søraas clean-compartment and prespecified sensitivity populations after separate fixed-score processing. **d**, Primary and common-probe processing use the occasion-first pair-weighted statistic. The duplicated-occasion diagnostic covers all 41 duplicated donor or recipient occasions before primary-population screening (39 recipient and two donor occasions), not only the eligible recipient set. These analyses test robustness of the bounded referent result and do not identify lineage biological age or exclude host influence.

### Extended Data Figure 3 | NHANES age-context, substitution and fold diagnostics

**a**, Complete NHANES III Phase 2 contrast set for the four frozen endpoint models using all 25 measurements or the remaining 20 after the Phase 1 age-selected five-measurement block was withheld, without or with observed chronological age. Points are design-weighted paired held-out log-loss contrasts and lines are 95% Fay-BRR intervals. \(V_F\) and \(V_R\) are the increments from adding observed age; \(P_{\mathrm{proxy}}\) is their difference; \(D_{\mathrm{block},0}\) and \(D_{\mathrm{block},1}\) are the block contributions without and with age; \(D_{\mathrm{replace}}\) compares remaining 20 plus age with the full 25 without age. The intervals condition on the frozen Phase 1 mask. No assertion is made that the remaining 20 lack age-related information, and an interval spanning zero is not equivalence. **b**, Direct substitution boundary between observed age and the transported Phase 1 age decoder under the same background, one-scalar spline family, tuning grid, folds and weights. \(\Delta_{\mathrm{sub}}\) was −0.000757 (95% Fay-BRR interval, −0.004661 to 0.003147), with one of five folds positive. There was no clear substitution advantage; this does not establish equality. **c**, Direction of five decision-facing paired contrasts in the five fixed primary-sampling-unit-atomic outer folds. Filled circles denote positive and open diamonds negative values under each declared formula. Fold directions are consistency diagnostics, not independent replications. Model-persistence and reload-parity records are retained in the reproducibility manifest rather than displayed as a scientific panel.

### Extended Data Figure 4 | Near-identical conditionally centred AgeGaps arise from distinct measured-profile contributions

**a**, Distribution of Euclidean distances between the 50-dimensional contribution vectors of 293 non-reusing, outcome-blind pairs matched exactly on outer fold, chronological age, sex, race or ethnicity and survey cycle and constrained to an absolute conditionally centred AgeGap difference of at most 0.25 decoder-output years. The solid and dashed lines mark the median (24.602) and 95th percentile (46.067) among all 293 pairs. The horizontal scale is distance in decoder-output contribution space, not an age difference between participants. **b**, Predeclared representative pair P00026, selected closest to the median contribution distance among the 205 display-eligible same-sign pairs with mean absolute centred AgeGap of at least two years. The centred deviations were −5.13 and −4.90 decoder-output years (absolute difference, 0.223); contribution-vector distance was 25.48 decoder-output years. **c**, Predeclared high-distance pair P00092, selected closest to the 95th percentile within the same display-eligible set. The centred deviations were −9.41 and −9.39 decoder-output years (absolute difference, 0.018); contribution-vector distance was 46.74 decoder-output years. In **b** and **c**, bars show signed contributions for the union of each participant's eight largest absolute contributors; all remaining features form the exact net remainder, and displayed contributions sum to each participant's conditionally centred AgeGap. Contributions attribute the output of the frozen fold-specific linear decoder after fold-local age and background centring; they are not causal, organ-specific or biological-age decompositions. This descriptive analysis uses the same unweighted complete-case interface as Fig. 3a,b, and attribution among correlated measurements depends on the frozen ridge parameterization.

### Extended Data Figure 5 | Endpoint-directed acquisition across five prespecified measurement-domain options

**a**, Two policies were frozen in NHANES 2007–2008 from five prespecified measurement-domain options. The endpoint-directed policy selected hepatic, protein and injury measurements by maximizing held-out five-year mortality log-loss gain. The inverse chronological-age decoder selected metabolic, adiposity and lipid measurements by minimizing held-out chronological-age mean-squared error. The source criteria have different targets and units and are therefore shown as separate routes. **b**, In NHANES 2009–2010, the frozen endpoint-directed and inverse-age domains yielded log-loss gains beyond age and background of 0.006418 (95% fixed-prediction bootstrap interval, 0.002297–0.010652) and 0.000646 (−0.001195–0.002481), respectively. **c**, The paired endpoint-directed-minus-inverse-age difference was 0.005772 (0.001380–0.010390); four of five target-fold estimates were positive. The comparison is exploratory cross-cycle evidence within one declared five-domain menu. Each domain was treated as one acquisition option; domains contained different numbers of measurements and no equivalence of monetary resources or real-world costs was assumed. Intervals use 5,000 event-stratified participant bootstraps conditional on stored predictions and fixed source selections; they omit refitting and study-development uncertainty. The analysis is not clinical value of information, cost analysis, untouched confirmation, universal transportability or biological-age identification.

## Methods

### Study design

The study assigned distinct roles to analytical proof, empirical consequences and object-specific alternatives. The analytical sequence comprised loss-specific age-task identification, observable-law non-identification, meaning non-inheritance, post-decoding operator limits, a shared-covariance account of downstream success and population risk ordering under deterministic compression. Two literature modules served separate purposes: a contemporary construct-promotion audit documented current examples, whereas a structured review organized prior criticisms, proposed repairs and their identification ceilings. Neither literature module contributed to the proof. Haematopoietic stem-cell transplantation (HSCT) quantified donor-lineage affiliation of a fixed age-valued blood score relative to a score-pairing null preserving score marginals and donor–recipient age geometry. Continuous NHANES tested whether an age-supervised decoder could improve held-out mortality prediction while direct modelling of the same 50 source measurements performed better, and whether near-identical gaps from that decoder specified a unique measured-profile realization. NHANES III then held scalar dimension and model opportunity fixed while changing the training target from age to mortality, compared the age decoder with observed age itself, and tested how observed age's predictive increment changed when a frozen age-selected measurement block was withheld. The estimand-specific replacement map supplied the constructive programme; endpoint-directed acquisition across five prespecified domains supplied an exploratory cross-cycle illustration.

A frozen X1–X3 subset of the earlier geometry package was retained as a transparent construction separating noisy age readability, linearly age-orthogonal declared state and their nonlinear mixture. Fixed five-fold out-of-fold models quantified chronological-age readability and incremental prediction of the generated state beyond age. The construction was not analysed as a human effect, repeated-simulation estimate, longitudinal trajectory or additional proof.

The contemporary examples in Table 1 were selected to represent distinct target and use classes, not to estimate the prevalence of any terminology or practice. The table is not a systematic review or a representative sample of all clocks. Its role is to establish that the target–interpretation promotions addressed by the analytical result remain current.

The central non-identification claim and meaning non-inheritance principle are analytical results. The X1–X3 construction and human analyses carry no proof responsibility for either result. The HSCT analyses were focused post-publication estimand audits rather than prospective preregistrations. All NHANES modules were retrospective and predictive or descriptive. For the continuous-NHANES pure-decoder comparison, the endpoint, population, panel, folds and model families were recorded in a versioned specification; the reported result was accepted without a confirmation refit or subsequent model, endpoint, panel or threshold search. The individual contribution analysis reused that frozen decoder, specified fold-local centring and outcome-blind matching before inspection, and introduced no decoder refit or endpoint-informed pair selection. The NHANES III modules were outcome-accessed fixed-protocol transfers from Phase 1 to Phase 2, not untouched confirmation or independent external validation. Phase 1 projections or measurement masks were frozen before the reported Phase 2 fitting and were not reselected. The cross-cycle acquisition rule was frozen in its source cycle before target-cycle evaluation, but the target cycle had been viewed during earlier reconnaissance and was classified as exploratory cross-cycle validation.

### Contemporary literature audits

The age-unit construct-promotion audit assembled source-verified contemporary examples from major general, medical, ageing and cell-biology journals. A paper was eligible as a main example when it supplied an individual-level predicted age, age gap or intervention-associated age-valued displacement; promoted that output to a biological, organ, cellular, immune or molecular ageing construct, rate or rejuvenation claim; and did not supply congruent named evidence for the promoted construct at the same conceptual level in its estimand, training target or validation. Boundary examples included clock applications constrained by a named endpoint, direct mechanism, longitudinal pace target or peripheral role; problem-diagnosing examples explicitly exposed clock heterogeneity or ambiguity. Downstream disease, mortality or frailty associations could establish narrower predictive uses but did not, by themselves, identify a broad latent ageing construct. Selection was designed to establish the existence and forms of the inferential operation, not a prevalence estimate, journal comparison or census. Supplementary Data 1 reports article title, journal and the relevant construct class.

The separate critical-literature review used local-first retrieval for 17 core critiques or methods papers published from 2024 through 2026 and three earlier context anchors, with targeted online supplementation where a local full text was unavailable. For each source we coded the problem targeted, the authors' proposed repair, remedy family, earliest transition in the promotion chain, the narrow object supportable if the repair succeeded and the interpretation that still did not follow. The five transitions were features to age score, age score to gap or state, gap to faster or accelerated ageing, acceleration to intervention response, and response to rejuvenation or benefit. The map is a bounded interpretive synthesis, not a systematic review, ranking or claim that no prior work recognized identification. Supplementary Table 1 provides the structured synthesis; Supplementary Data 2 contains the complete source-level coding, evidence locators and codebook.

### Identification analysis

We treated the observable law \(P(A,W,X)\), declared loss and predictor class as the information available to an age-decoding task. Under unrestricted squared-error loss the target is \(E(A\mid W,X)\); under model restriction the fitted decoder approximates a loss- and class-specific projection. Latent-state non-identification was established by constructing distinct extensions \(P(A,W,X)r_k(H\mid A,W,X)\) with identical observable laws and optimal age decoders. For any proposed biological property \(\psi\), identification requires \(\psi\) to be constant over the resulting observational-equivalence class. If added measurement, endpoint, longitudinal or causal restrictions make \(\psi\) constant only in a narrower class, its identity arises from those restrictions. Prediction-error, transport, approximation, estimation and calibration contributions were separated algebraically in Supplementary Note 1; the decomposition is an identity, not an identifiable biological attribution of an observed individual gap.

Shared age-related covariance and proxy overlap among chronological age, measured biological processes, exposures, composition, cohort and survival selection were represented locally by \(\operatorname{Cov}(G_A,Y\mid A,W)\approx w_A^{\mathsf T}\Sigma_R\beta_Y\). This relation describes conditional association with the named endpoint under the stated local representation, not a causal decomposition or identification of ageing acceleration.

Equality of the complete endpoint law after scalar compression was represented by \(Y_\Delta\perp X\mid A,W,q_A(W,X)\). This predictive-sufficiency condition is stronger than equality under one loss and is not inherited from age supervision. For an age-valued mapping \(S=f(X)\), intervention movement was represented exactly by \(\Delta S=f(X+\Delta X)-f(X)\). This identity supplies neither a causal outcome contrast nor a surrogate relationship.

For the age- and background-residualized score \(G_A=q_A(W,X)-E\{q_A(W,X)\mid A,W\}\), the conditional-expectation projection gives

\[
E(G_A^2)
\leq
E[\{q_A(W,X)-A\}^2].
\]

Full statements and regularity qualifications are assigned to Supplementary Note 1. The proposition concerns the identifying content of the chronological-age-decoding objective. Additional longitudinal, causal, measurement, endpoint or invariance restrictions may identify separately defined objects and may validate an already age-trained score for a specified use. That measurement validity derives from the added evidence, not from the age-decoding objective.

### Haematopoietic stem-cell transplantation analyses

We treated allogeneic haematopoietic stem-cell transplantation as an object-affiliation setting: the assayed circulating blood compartment is donor-derived, whereas the recipient remains the host organism. The analyses were not used to establish the analytical non-identification result.

For the Søraas *et al.* dataset<sup>21</sup>, we parsed Table 1 directly from the Europe PMC XML (PMCID: PMC6413751), without manually transcribing values or reprocessing methylation data. The table contained 41 rows. We reproduced the five published footnote-b exclusions—an outlying clock value or donor chimerism below 80%—leaving 36 published analysis samples from 30 recipients. The primary clean-compartment population retained samples whose latest recorded relapse status was “No”, because relapse can reintroduce recipient malignant cells into blood; this yielded 24 samples from 19 recipients. Let \(S\) denote the reported Horvath-2013 multi-tissue DNA-methylation-age score in recipient blood, \(A_L\) the donor-lineage chronological coordinate at blood draw and \(A_R\) recipient chronological age at blood draw. For each sample we calculated \(d=|S-A_L|-|S-A_R|\); \(d<0\) indicates numerical proximity to the donor-lineage coordinate. Repeated observations were averaged within recipient, and recipients—not samples—were the independent units. We reported the recipient-weighted mean and median \(d\), the numbers of negative, tied and positive recipient means, and an exact two-sided binomial test among non-tied recipients. A 95% percentile interval for the mean was obtained from 10,000 recipient bootstrap samples (seed 20260730). Sensitivities included all 36 published analysis samples, the non-relapse population with donor chimerism at least 95%, the non-relapse population sampled at least 12 months after transplantation and leave-one-recipient-out estimates. Five contemporaneous donor-blood comparisons were treated as descriptive only.

For Holland *et al.*<sup>22</sup>, we used the six author-normalized public beta-value matrices and `comRand.csv` metadata from Mendeley Data version 1 (doi:10.17632/j5krtjfj6y.1). The matrices comprised 334 arrays—303 recipient and 31 donor arrays. We implemented the 353-CpG Horvath-2013 coefficient formula and verified it against the bundled five-sample reference before scoring. Because the matrices had been processed separately by batch, we did not combine-renormalize them, apply ComBat or perform cohort-derived imputation. The six batches lacked 52, 28, 34, 18, 13 and 10 target probes, respectively. A probe absent from an entire batch was assigned the fixed `overallMeanByCpGacross50data` value from the local PC-Clocks resource; these values have not been established as the original Horvath training means. We therefore refer to the result as a Horvath-2013 coefficient implementation with fixed external-mean imputation, not as a paper-identical reconstruction of the original clock.

We parsed each array identifier into donor–recipient pair, role, batch and measurement. Arrays with the same pair, role and all four public age/time fields were averaged as one biological occasion, reducing 303 recipient arrays to 264 recipient occasions; distinct occasions were retained. For recipient samples, \(A_L=A_{D,\mathrm{TX}}+t\) denotes the donor-lineage chronological coordinate and \(A_R=A_{R,\mathrm{TX}}+t\) denotes recipient-host age. \(A_L\) equalled the public `AgeCells` field to floating-point precision but was not interpreted as the literal age of the short-lived blood cells at sampling. The primary population retained recipient-role occasions at least 0.1 years after transplantation and with \(|A_R-A_L|\geq10\) years, yielding 118 occasions from 61 donor–recipient pairs.

For each occasion we calculated

\[
J_{\mathrm{aff}}=\frac{|S-A_R|-|S-A_L|}{|A_R-A_L|}.
\]

Positive values favour the lineage-coordinate side, negative values favour the host side and zero indicates equal distance. By the reverse triangle inequality, \(-1\leq J_{\mathrm{aff}}\leq1\). The index saturates at either boundary and is not a normalized measure of absolute accuracy. For the occasion-first summary, \(J_{\mathrm{occ}}\), we averaged \(J_{\mathrm{aff}}\) equally across occasions within pair and then equally across pairs. This gives the primary-processing value of 0.535 in Extended Data Fig. 2d; it is not the sign-balanced pair-summary statistic used in Fig. 2c. Uncertainty was quantified with 10,000 pair bootstrap samples (seed 20260730); signs of pair means were summarized with an exact two-sided binomial test among non-tied pairs. We also reported pair-weighted absolute errors to \(A_L\) and \(A_R\).

Prespecified sensitivities used \(|A_R-A_L|>0\) and thresholds of 5, 15 and 20 years; the earliest eligible occasion; the occasion closest to one year; separate host-older and host-younger strata; the within-pair median; leave-one-batch-out reaggregation; and exclusion versus truncation of near-zero negative follow-up times (Extended Data Fig. 2). A post-specification processing sensitivity retained observed values only for the 253 probes present in all six batches and assigned the same fixed external mean to the other 100 probes in every batch. Across all samples before primary-population screening, 41 biological occasions had duplicate arrays: 39 recipient and two donor occasions. Their median within-occasion score range was 3.89 years and their maximum range was 11.502 years; duplicate arrays were not treated as independent observations. The pair bootstrap is conditional on the author-normalized matrices and does not incorporate preprocessing or imputation uncertainty. Holland *et al.* extended and partly overlapped the Søraas cohort and was not treated as an independent cohort replication.

Because numerical closeness to \(A_L\) can arise from score marginals and donor/recipient age geometry alone, the prespecified geometry-preserving score-pairing null analysis used only the recipient- or pair-level summaries and did not rescore methylation. For this analysis, scores and chronological coordinates were first averaged within pair or recipient, and the same distance formula was then evaluated on those summaries to give \(J_{\mathrm{pair}}\). Units were stratified by the sign of \(A_R-A_L\), and the primary statistic \(J_{\mathrm{aff,bal}}\) equally weighted the two stratum means. In Holland, the pair-summary stratum means were 0.488 and 0.763, giving 0.626. This differs from averaging occasion-specific indices before pair and stratum aggregation; the absolute-value operation makes the order consequential. Within each sign stratum, we permuted fixed scores across independent units while holding every \((A_R,A_L)\) pair fixed, using 20,000 permutations (seed 20260823). This preserved score marginals, age geometry and stratum sizes while breaking observed score–referent pairing. It also broke the score–recipient-age relationship, so excess affiliation under this null does not exclude host-age-dependent shrinkage or estimate donor influence conditional on recipient age. The one-sided finite-permutation value used the add-one correction. Uncertainty for the observed statistic and observed-minus-null-mean contrast used 10,000 stratified independent-unit bootstrap samples (seed 20260824). Constant mean- and median-score controls and ordinary least-squares \(S\)-on-\(A_L\) and \(S\)-on-\(A_R\) fits were descriptive geometric and calibration diagnostics and were not used to recalibrate the scores. The prespecified analysis required the Holland observed statistic to exceed the null 97.5th percentile, a positive bootstrap interval for the observed-minus-null contrast, positive contrasts in both sign strata, the same direction in Søraas, positive Holland \(S\)-on-\(A_L\) slopes in both strata and deterministic reconstruction.

As a conventional sensitivity, we fitted \(S-A_R=\alpha+\beta(A_L-A_R)+\varepsilon\) separately in Holland and Søraas at the frozen independent-unit level. Because the outcome and regressor share \(-A_R\), common-anchor covariance can contribute to \(\beta\); we therefore treated it as a descriptive anchor-coordinate sensitivity, not an independent referent-identification parameter. We report ordinary least-squares estimates, HC3 intervals, 10,000-draw independent-unit bootstrap intervals (seed 20260826), \(R^2\), residual mean-squared error and constrained \(\beta=0\) and \(\beta=1\) anchor fits with a freely estimated intercept. The datasets were not pooled and the fitted scores were not recalibrated.

### NHANES data, endpoint and survey design

We used de-identified public-use National Health and Nutrition Examination Survey data and the 2019 public-use linked mortality file, with follow-up through 31 December 2019<sup>44–46</sup>. The originating protocols obtained ethics-review-board approval and informed consent<sup>47</sup>. NCHS reports that follow-up time or underlying cause was replaced by synthetic values for selected public-use records, whereas vital status was not perturbed<sup>44</sup>; released values were used without reconstruction. All modules used a fixed five-year all-cause mortality endpoint. We directly modelled five-year event probability and evaluated log loss.

The continuous-NHANES modules used participants aged 20–79 years from the 2007–2008, 2009–2010 and 2011–2012 cycles. Public-use ages reported as 80 were excluded because age is top-coded. These cycles supplied at least 60 months of administrative follow-up for every event-free included participant. The complete-case pure-decoder interface contained 13,724 participants and 470 deaths and was analysed without survey weights; it targets the declared predictive sample, not population prevalence. The endpoint-directed acquisition module used separately declared source- and target-cycle interfaces.

The NHANES III protocol-transfer modules used the canonical Phase 1 source interface (7,558 participants; 402 five-year deaths) and Phase 2 target interface (7,774 participants; 392 deaths), restricted to ages 20–79 years. The Phase 2 analysis used Mobile Examination Center weights, 23 released strata, 46 primary sampling units and fixed primary-sampling-unit-atomic folds. Missing measurements were retained and imputed by the training-fold weighted median. These modules estimate design-weighted predictive loss in the declared Phase 2 interface.

### Pure chronological-age decoder and same-source endpoint comparison

Background \(W\) comprised sex, race or ethnicity and survey cycle, observed chronological age was \(A\), and \(X\) was the fixed ordered 50-measurement \(S_3\) panel inherited from the complete-support analysis. The sole endpoint was death by 60 months. We compared exactly three endpoint information sets: \(M_{WA}=P(Y_5\mid W,A)\); \(M_{\mathrm{CLOCK}}=P(Y_5\mid W,A,S)\), where \(S=q_A(W,X)\) was trained only to reconstruct \(A\); and \(M_X=P(Y_5\mid W,A,X)\). Given \(A\), the raw gap \(S-A\) and \(S\) are deterministically equivalent, so no separate gap model was fitted.

Within every frozen outer training fold, categorical background was one-hot encoded and the 50 measurements were standardized using training data only. Ridge regression predicted chronological age. The penalty was selected by three-fold age-quintile-stratified root-mean-square error from the fixed grid \(\{0.01,0.1,1,10,100,1000\}\). Cross-fitted scores were generated for the outer-training participants using the selected penalty and fixed inner folds; the decoder was then fitted on the full outer-training fold to score the held-out outer fold. No mortality label entered decoder preprocessing, tuning or fitting.

For \(M_{\mathrm{CLOCK}}\), ridge logistic regression was fitted within each outer training fold using its cross-fitted clock scores, \(W\) and the same five-knot fold-local cubic age spline used in the pre-existing complete-support analysis. The endpoint penalty was selected by mean three-fold inner-validation log loss from the prespecified grid used in that analysis. Category encoding, score scaling, spline construction and tuning were training-fold local. \(M_{WA}\) and \(M_X\) were unweighted L2-logistic models with an intercept, reference-coded background and a cubic age basis with five uniformly spaced training-range knots and linear extrapolation; \(M_X\) also included all 50 transformed measurements. Measurement and age-basis columns were standardized within each training partition, and three inner folds selected the penalty by mean fold-level validation log loss. Their stored predictions were reused after participant, fold and feature-order checks. SM2 identifies their generating script and model aliases; the software provides a combined ordered 50/25-variable dictionary with original units and transformations.

Let \(L(\cdot)\) denote pooled participant-level out-of-fold log loss. The paired contrasts were

\[
V_{\mathrm{CLOCK}}=L(M_{WA})-L(M_{\mathrm{CLOCK}}),
\qquad
V_{\mathrm{SOURCE}}=L(M_{\mathrm{CLOCK}})-L(M_X),
\]

and \(V_X=L(M_{WA})-L(M_X)\). Positive \(V_{\mathrm{SOURCE}}\) means that direct source-measurement modelling performed better for this endpoint under the declared estimators; it is not a population information estimate or a universal finite-sample theorem. We reused 2,000 paired participant-bootstrap indices from the pre-existing complete-support analysis, stratified jointly by endpoint and decade age band. Intervals condition on stored out-of-fold predictions and omit model refitting, tuning and study-development uncertainty.

### Individual measurement contributions to conditionally centred AgeGap

For participant \(i\) in outer fold \(k\), we reloaded the frozen ridge age decoder and reproduced its stored held-out score \(S_i\). Using outer-training participants only, a multivariate ridge model estimated the conditional mean of each of the 50 measurements given chronological age and background. Age was represented by a five-knot cubic spline and background comprised sex, race or ethnicity and survey cycle. The penalty was selected by three-fold age-quintile-stratified cross-validation from \(\{0.01,0.1,1,10,100,1000\}\), minimizing standardized measurement-reconstruction error. All tuning estimators and final residualizers were persisted, reloaded and parity-checked.

Let \(\beta_{kj}\) and \(\sigma_{kj}\) denote the frozen decoder coefficient and training-fold scale for measurement \(j\), and let \(\widehat{\mu}_{kj}(A_i,W_i)\) denote its fold-local conditional mean. The score centre was induced by passing the vector of these measurement means through the full frozen decoder:
\[
\widehat m_{S,k}(A,W)=q_{A,k}\{W,\widehat{\mu}_k(A,W)\}.
\]
This evaluation retained the decoder's background encoding, training-fold scaling and intercept; it was not a separately fitted score-on-age regression. We defined

\[
c_{ij}=\beta_{kj}\frac{X_{ij}-\widehat{\mu}_{kj}(A_i,W_i)}{\sigma_{kj}},
\]

so that

\[
\sum_{j=1}^{50}c_{ij}=S_i-\widehat m_{S,k}(A_i,W_i).
\]

This sum is the conditionally centred AgeGap, not the raw gap \(S_i-A_i\), which was retained separately. Because matched participants shared outer fold, age and background exactly, their raw-gap and centred-gap differences were equal to numerical precision.

Pairing was deterministic and outcome blind. Within exact strata of outer fold, chronological age, sex, race or ethnicity and survey cycle, participants were ordered by conditionally centred AgeGap; adjacent candidate edges were ordered by absolute gap difference and row identifiers and retained greedily without participant reuse. Pairs were retained when their centred gaps differed by no more than 0.25 decoder-output years. Contribution-profile distance was calculated before display selection. The two displayed pairs were chosen by the frozen median- and 95th-percentile-distance rules among same-sign pairs with mean absolute centred AgeGap of at least two decoder-output years. Distances are norms in model-contribution space, not participant age differences. The contributions explain how the frozen linear decoder generated its scalar and are not causal, organ-specific or biological-age components; their allocation among correlated measurements depends on the frozen ridge parameterization, scaling and centring reference.

### Flexible calibration of the continuous-NHANES age decoder

We reused the original five outer folds and frozen fold-local age scores. The endpoint model contained background, a five-knot cubic spline of chronological age and one prespecified five-knot cubic spline of \(q_A\); ridge-logistic regularization was selected inside each outer training set from the prespecified grid. No alternative spline or calibration search was performed. The contrast was the held-out log loss under this flexible clock model minus that under direct modelling of the frozen 50-measurement panel. The fully nested continuous-NHANES matched-scalar comparison is described in Supplementary Methods SM2b; the NHANES III Phase 1-to-Phase 2 protocol transfer is described below and in SM2c.

### NHANES III matched scalar target-supervision protocol transfer

Background comprised sex and race or ethnicity, and \(X\) comprised the same ordered 25 measurements for both one-dimensional projections. Chronological age was not an input. In NHANES III Phase 1, weighted-median imputation, one-hot background encoding, weighted measurement standardization and ridge regularization were identical for the two projections. The age-supervised projection \(q_A\) used standardized chronological age as its target; the endpoint-supervised projection \(q_Y\) used standardized five-year mortality. Target scaling and all preprocessing were estimated inside the relevant Phase 1 training folds. One final projection of each type was fitted on complete Phase 1 and applied unchanged to Phase 2; Phase 2 mortality outcomes did not enter projection preprocessing, tuning or fitting.

In Phase 2, both frozen scores received the same target-local opportunity: weighted L2-penalized logistic regression containing background, a cubic age spline with fixed knots at 20, 35, 50, 65 and 80 years and a training-local five-knot uniform cubic spline of the relevant score. Regularization was selected by pooled design-weighted log loss in three inner folds, and all reported probabilities were held out across five fixed outer folds. We compared the background-and-age model with the addition of \(q_A\), and directly compared otherwise identical models adding \(q_A\) or \(q_Y\). A linear-score version was prespecified as a sensitivity.

The separate age-substitution comparison used the same Phase 2 participants, folds, weights and downstream model family. One arm contained background and a five-knot spline of observed age; the other replaced age with an identically constructed spline of the frozen \(q_A\). It tested superiority, not equivalence or non-inferiority.

For both modules, 32 Fay balanced repeated replication (Fay-BRR) weight sets with \(\rho=0.3\) repeated Phase 2 downstream preprocessing, inner tuning, fitting and held-out prediction; intervals used 23 design degrees of freedom. Phase 1 projection directions remained fixed. The intervals therefore quantify Phase 2 survey-design uncertainty conditional on the two frozen projections and do not include Phase 1 development uncertainty.

### NHANES III chronological-age increment and proxy-substitution comparison

We compared the incremental mortality-prediction value of observed chronological age after two information sets. Background \(W\) comprised sex and race or ethnicity. The full panel contained the ordered 25 measurements. The reduced panel removed exactly the five-measurement cardiorespiratory domain selected for age reconstruction in the earlier Phase 1 acquisition analysis: systolic blood pressure, diastolic blood pressure, radial pulse, forced expiratory volume in one second and forced vital capacity. The mask was frozen before the present Phase 2 fitting and was not reselected using Phase 2 outcomes. The remaining panel is described only as the remaining 20 measurements; it is not assumed to lack age-related information.

Four otherwise homologous weighted L2-logistic models predicted five-year death: full and reduced panels were each fitted without and with a training-local fixed-knot cubic spline of observed age. Encoding, weighted-median imputation, weighted standardization, spline construction, inner tuning and outer fitting were repeated separately for every model and weight set. No age-decoder score or endpoint-supervised scalar entered these models.

The full-panel age increment was the reduction in design-weighted held-out log loss from adding age to the 25-measurement model. The reduced-panel age increment was defined analogously after the frozen block was withheld. Their difference was the primary predictive proxy-substitution contrast. Secondary contrasts measured the block's value without age, its value with age, and reduced-panel-plus-age versus full-panel-without-age. Thirty-two Fay-BRR replicates with \(\rho=0.3\) repeated the complete Phase 2 pipeline and used 23 design degrees of freedom. Intervals are conditional on the frozen Phase 1 mask and omit its development uncertainty. No contrast was interpreted as causal mediation, equivalence, biological-age identity or complete measured-state sufficiency.

### Endpoint-directed measurement acquisition

For measurement acquisition, the sole endpoint was five-year all-cause mortality. The source population was NHANES 2007–2008 (4,599 complete participants; 174 deaths) and the target was 2009–2010 (4,896; 157). Candidate acquisition categories were renal/mineral, metabolic/adiposity/lipid, hepatic/protein/injury, haematology and haemodynamic measurements. For domain \(M_k\), source measurement value was

\[
V_k
=
L_{\mathrm{OOF}}(W+A)
-
L_{\mathrm{OOF}}(W+A+M_k).
\]

For the inverse chronological-age policy, each domain \(M_k\) entered a ridge decoder \(A\sim W+M_k\), where \(W\) comprised sex and race or ethnicity. Encoding, declared feature transformations and standardization were training-fold local. Five age-quintile-stratified outer folds (seed 20260826) supplied participant-level out-of-fold predictions. Within each outer training fold, the ridge penalty was selected from \(\{0.01,0.1,1,10,100,1000\}\) by pooled mean-squared error across three age-quintile-stratified inner folds (seed \(20260827+\text{outer fold}\)). The inverse-decoder policy selected the domain with the lowest pooled source out-of-fold age mean-squared error; the separately frozen endpoint policy selected \(\arg\max_k V_k\). Source choices and the source-data signature were frozen before target evaluation, and target outcomes did not enter either source selection.

In the target cycle, we compared the held-out five-year mortality log-loss gains of the two source-frozen domains beyond age and background. Participant identity, fold assignment and ordered features matched the sealed predecessor. Uncertainty used 5,000 event-stratified paired participant bootstraps of fixed out-of-fold losses (seed 20260802), without policy reselection or model refitting. Each of the five domains was treated as one acquisition option. Domains differed in measurement count, and no equivalence of monetary resources or real-world costs was assumed.

### Transparent three-variable construction

We retained X1–X3 from the frozen cross-sectional geometry generated with seed 20260806. Age was sampled uniformly from 20 to 80 years and a binary separation state \(D\) with probability 0.40 (realized State 0, \(n=299\); State 1, \(n=201\)). Let \(A_z\) be standardized age, \(D_c\) centred state, \(Q_A\) the standardized quadratic age component after projection off the linear age direction, \(\varepsilon_j\) independent standard-normal noise and \(z(\cdot)\) sample standardization. The retained variables were

\[
X_1=z(0.95A_z+0.65\varepsilon_1),
\]

\[
X_2=z\{\Pi_{\perp(1,A)}(1.35D_c+0.55\varepsilon_2)\},
\]

and

\[
X_3=z(0.65A_z+0.85Q_A+0.90D_c+0.55\varepsilon_3),
\]

where \(\Pi_{\perp(1,A)}\) removes the realized intercept and linear age direction. Thus X1 supplied noisy age readability without a designed state term, X2 supplied declared same-age structure that was exactly linearly age-orthogonal in the realized construction, and X3 mixed linear and nonlinear age structure with state. The displayed limiting cases—noisy age without state and exact age alongside a separate state variable—are algebraic consequences rather than additional fitted analyses.

We assigned the 500 rows to five participant-disjoint folds stratified jointly by the declared binary state and five equal-frequency age groups (shuffle seed 20260827). Chronological-age decoding used four fixed ridge pipelines containing X1, X2, X3 or all three variables; each applied a five-knot cubic spline, training-fold standardization and \(\mathrm{Ridge}(\alpha=1)\). Same-age state decoding used a fixed L2-logistic age-only model and four homologous age-plus-measurement models; age entered as a five-knot cubic spline and the measurements were standardized within training folds. The primary task-map quantities were pooled out-of-fold age \(R^2\) and the reduction in pooled state log loss relative to the age-only model. No model-family, penalty, split or threshold search was performed.

The pooled age \(R^2\)/state log-loss gains beyond age were 0.681/−0.002 for X1, −0.015/0.407 for X2, 0.301/0.220 for X3 and 0.730/0.483 for X1–X3. These are fixed descriptive quantities for the declared construction and carry no population interval or human biological interpretation.

### Statistical analysis

Analysis records were constructed rows for X1–X3, recipient- or pair-level summaries for HSCT and participants for predictive NHANES analyses. Validation and resampling followed each module's design; in NHANES III, validation folds kept primary sampling units (PSUs) intact. For the individual contribution summary, the descriptive unit was one retained non-reusing matched pair. The fixed X1–X3 construction was descriptive and supplied no population interval. The individual contribution analysis was also descriptive: matching used no endpoint information, all retained non-reusing pairs were reported and no hypothesis test or population interval was assigned to their empirical distance distribution. Continuous-NHANES uncertainty used the declared participant or event-stratified fixed-prediction bootstrap and therefore conditions on stored models, folds and policies. NHANES III uncertainty used full Phase 2 Fay-BRR downstream refits with 23 design degrees of freedom, conditional on frozen Phase 1 projection directions or masks. Exact two-sided binomial tests excluded ties. Lower log loss indicates better probabilistic prediction; reported reductions are paired held-out proper-score contrasts, not clinical effect sizes or percentage accuracy. An interval spanning zero was interpreted as an unresolved contrast in that design, not proof of no effect, equality or non-inferiority. Fold-construction rules and model grids are detailed in Supplementary Methods; persistence audits and sensitivity results are indexed in Supplementary Software 1 and Source Data. Repository-level version, artifact and reconstruction records are indexed in the reproducibility manifest included in Supplementary Software 1.

### Use of artificial intelligence

OpenAI Codex assisted with development of analysis and verification scripts, manuscript drafting and editorial revision. Analyses were executed using the versioned code and data described here; LLM-generated statements were not treated as data or bibliographic evidence.

## Data availability

The Søraas 2019 article table is available through Europe PMC (PMCID: PMC6413751). Holland 2024 individual-level metadata and six author-normalized methylation matrices are available from Mendeley Data version 1 (https://doi.org/10.17632/j5krtjfj6y.1); the article and supporting information are available through Europe PMC (PMCID: PMC11113269). Continuous NHANES, NHANES III and public linked-mortality data are available from the US Centers for Disease Control and Prevention/National Center for Health Statistics<sup>44–46</sup>. The Arivale and TwinsUK individual-level data underlying the published multi-omic BMI example are controlled access and were not reanalysed<sup>48</sup>. The source-verified construct-promotion audit and critical-literature remedy map are provided as Supplementary Data 1 and 2. Figure-level non-restricted derivatives, including aggregate matched-pair distributions and the predeclared display-pair contribution profiles underlying Extended Data Fig. 4, are provided in Source Data. Analysis specifications, code, ordered feature definitions, accession and module manifests, transformation definitions and hash/path references to internally retained reconstruction objects are provided in Supplementary Software 1. Supplementary Software 1 also provides frozen primary participant-level predictions, stored survey-replicate estimates, and the complete individual contribution profiles and matrix for deterministic result checks. Fitted-model binaries and large resampling-index arrays are retained internally and are not included in the submission archive. For peer review, the supplied code and derivative data are provided to editors and reviewers for evaluation. Third-party data remain subject to their source terms. Public deposition and reuse-licensing details will be supplied before publication.

## Code availability

Analysis and figure-generation code for the transparent age–state construction, HSCT object-affiliation and referent-null analyses, continuous-NHANES pure age decoder, individual conditionally centred AgeGap contribution decomposition and outcome-blind matching, NHANES III matched target-supervision, age-substitution and proxy-substitution comparisons, endpoint-directed acquisition and submission figures is provided as Supplementary Software 1. The snapshot includes the pinned environment specification, deterministic entry points, evidence-module registry, reproducibility manifest, source-data indices and non-restricted result tables used by the submission. The archive includes a portable stored-result verification entry point and installation and execution instructions. It is supplied for editorial and reviewer evaluation; no public reuse licence has been assigned.

## Acknowledgements

**[DATASET ACKNOWLEDGEMENTS, INFRASTRUCTURE AND FUNDING TO BE INSERTED]**

## Funding

**[FUNDING SOURCES AND GRANT NUMBERS TO BE INSERTED]**

## Author contributions

**[CRediT CONTRIBUTIONS BY AUTHOR INITIALS TO BE INSERTED]**

## Competing interests

The authors declare **[NO COMPETING INTERESTS / DETAILS TO BE INSERTED]**.

## Additional information

Supplementary Information is available for this paper. Correspondence and requests for materials should be addressed to **[CORRESPONDING AUTHOR]**.
