## Supplementary_Software_1 for "Inferential boundaries of age prediction: why prediction does not establish biological age measurement": P2_ANALYSIS_DIRECTION_LOCK_20260826.md

# P2 analysis direction lock

Date: 26 August 2026

## Authorization and decision served

The author explicitly authorized the previously deferred P2 empirical work on
26 August 2026. P2 closes three avoidable reviewer exits without changing the
Article's north-star claim:

1. replace the forward-age-indexability control in `MTH-01` with a true
   domain-level inverse chronological-age decoder;
2. test whether the HSCT referent result survives a conventional continuous
   regression estimand and, only after a frozen coverage gate, a second fixed
   chronological-age coefficient score;
3. test whether the NHANES source-measurement advantage survives flexible
   downstream calibration and whether mortality supervision outperforms age
   supervision when input, linear capacity and scalar dimension are matched.

P2 is an analysis phase. It does not authorize endpoint, population, clock or
model-family search beyond the prespecified single revisions below.

## Branch A: MTH-01 inverse-age comparator

- New analysis version:
  `nhanes_endpoint_vs_inverse_age_decoder_acquisition_v0_2`.
- Source and target remain NHANES 2007--2008 and 2009--2010, respectively,
  within the frozen 20--79-year complete-case 50-measurement interface.
- The five candidate domains and their order are unchanged.
- The source age policy selects the domain minimizing pooled OOF MSE from
  `A ~ W + M_k`, using outcome-blind nested folds and a fixed ridge grid.
- The source endpoint policy remains frozen from v0.1. Target outcomes do not
  enter either source selection.
- The target comparison uses the frozen participant-level mortality OOF
  predictions. It is a decision probe until the legacy endpoint fitted-object
  state is separately reconstructed and parity-audited.
- No domain, seed, loss, endpoint or model rescue is permitted after inspection.

## Branch B: HUM-01 HSCT robustness

- New analysis version:
  `hsct_referent_regression_clock_sensitivity_v0_1`.
- Primary estimand, separately in Holland and Soeraas, is
  `S-A_R = alpha + beta(A_L-A_R) + epsilon` at the frozen independent-unit
  level. Holland is placement-driving; Soeraas is complementary and is not an
  independent replication.
- The existing geometry-preserving score-pairing permutation remains the
  non-parametric primary context. The regression is a conventional sensitivity,
  not a calibration of the score and not a causal model.
- Public Holland data contain six author-processed ENmix beta matrices, not raw
  IDAT. P2 must report raw-IDAT reprocessing as unavailable rather than relabel
  beta values as raw data.
- The single prespecified second-score attempt is a Hannum-2013 coefficient
  score. All 71 coefficient probes are streamed for a coverage audit. If the
  complete 71-probe formula is unavailable, one common-probe implementation may
  be used only when the same fixed probe intersection applies to all six
  batches, contains at least 65 probes and uses no cohort-derived imputation.
  Otherwise the second-score branch stops. No third clock is searched.
- Pre-transplant recipient controls are unavailable. Donor arrays and stable
  post-transplant windows may be used only as declared sensitivities.

## Branch C: HUM-06 matched scalar target supervision

- New module: `HUM-06`.
- The first implementation,
  `nhanes_5y_matched_scalar_target_supervision_probe_v0_1`, is retained as
  audit provenance. Independent code audit found that its outer held-out
  predictions were clean, but its downstream C-tuning folds reused one common
  outer-training OOF score vector. For mortality supervision, scores on a
  downstream-training fold could therefore have been fitted using outcomes in
  that downstream validation fold. Its inner validation was not fully nested,
  and v0.1 cannot carry the matched-target verdict.
- The corrective analysis version is
  `nhanes_5y_matched_scalar_target_supervision_probe_v0_2`. It preserves the
  population, endpoint, feature panel, outer and inner folds, seeds, model
  families, tuning grids, metrics and bootstrap declared for v0.1.
- `HUM-04` remains frozen and is not silently rewritten.
- Flexible calibration uses the existing fold-local age scores and compares a
  fixed five-knot score spline with the frozen direct 50-measurement mortality
  model.
- The matched comparison constructs one age-supervised and one
  mortality-supervised ridge projection from the same 50 measurements and
  background, with identical nested fold exclusions, preprocessing, ridge
  grid, squared-error tuning rule and scalar dimension. Chronological age enters
  only the common downstream endpoint model.
- Both scalars receive the same linear and fixed spline downstream calibration
  opportunities. No learner library or post-result calibration search is used.
- Within every downstream C-tuning fold in v0.2, projection alpha selection and
  the validation projection are restricted to that fold's training side. The
  downstream-training scores are generated by a further projection
  cross-fitting layer wholly inside that training side; the downstream
  validation scores are generated by a projection fitted on the complete
  training side. No validation outcome may enter a mortality-supervised score
  used to fit the downstream model for that validation fold.
- Five prespecified five-fold outer partitions use seeds `20260823`,
  `20260901`, `20260902`, `20260903` and `20260904`. Repeats are not treated as
  independent samples.
- The primary metric is participant-level OOF binary log loss. A paired
  participant bootstrap averages each participant's losses across repeats.
- Failure of identity, feature-order, event-support, leakage, persistence or
  reload-parity gates stops the branch.
- V0.2 reload validation must connect every persisted score end to end:
  projection-bundle holdouts to assembled training/validation/test scores, and
  those scores to every downstream tuning and final fitted object. Inner
  identities must be verified as subsets of the declared outer and downstream
  training sets.

## Shared execution contract

- Use only the pinned `lhstate` Python environment and the documented R entry
  when R is genuinely required.
- Every analysis-ready interface, feature order, fold identity, derived score,
  prediction, loss vector, tuning result and final table is persisted.
- Every fitted preprocessor, projection, calibration object and endpoint model
  is serialized, reloaded and checked for elementwise prediction parity at
  tolerance `1e-12`.
- Existing frozen output directories are never overwritten. Incompatible
  reruns go to a new versioned directory and fail closed.
- Negative, weak or split-sensitive results are retained as audit evidence;
  there is no rescue search.
- Manuscript, figures and registry are updated only after a branch passes its
  declared scientific and artifact gates.

## Non-goals

- No universal biological-age estimator or clock leaderboard.
- No new endpoint, horizon, subgroup or measurement-domain search.
- No claim that mortality supervision identifies biological age.
- No claim that one HSCT score excludes host influence or identifies lineage
  biological age.
- No use of repeated folds as independent replications.
