## Supplementary_Software_1 for "Inferential boundaries of age prediction: why prediction does not establish biological age measurement": HUM_08_NHANES_INDIVIDUAL_AGEGAP_REALIZATION_CONTRACT_20260831.md

# Evidence module contract — HUM-08

Date: 31 August 2026

## Identity

- **Module ID:** HUM-08
- **Semantic name:** NHANES non-unique measured-profile realizations of individual AgeGap
- **Analysis version:** nhanes_individual_agegap_contributions_v0_1
- **Manuscript pair:** v6.13 / Supplementary Information v5.5
- **Evidence role:** materiality
- **Status:** current
- **Placement:** main+extended-data+supplement
- **Supersedes:** none

HUM-08 is a new semantic module. It does not revise HUM-04, whose responsibility
is endpoint utility and same-source mortality performance. HUM-07 remains a
historical rejected identifier and is not reused.

## Scientific contract

- **Question:** Under one frozen chronological-age decoder, can near-identical
  individual AgeGaps arise from materially different configurations of
  contributions from the same measured biological panel?
- **Estimand:** The outcome-blind distribution of 50-dimensional
  decoder-contribution distances among non-reusing pairs matched exactly on
  outer fold, chronological age, sex, race or ethnicity and survey cycle and
  restricted to an absolute conditionally centred AgeGap difference no greater
  than 0.25 decoder-output years. Euclidean contribution-vector distance is the
  primary profile-distance quantity.
- **Independent unit:** one non-reusing matched participant pair.
- **Decision served:** establish one bounded human instance in which a fitted
  AgeGap scalar does not specify a unique measured-profile realization.

## Claim contract

- **Allowed main claim:** In one frozen continuous-NHANES linear age decoder,
  near-identical conditionally centred AgeGaps among participants matched on
  age, background and fold arose from markedly different 50-measurement
  contribution configurations; the scalar therefore did not specify a unique
  measured-profile realization.
- **Forbidden claims:** causal ageing mechanisms, organ ages, biological-age
  components, biological burden, damage, benefit or a unique latent state;
  contribution-space distance as an age difference; model-invariant or
  parameterization-invariant feature attribution; universal transport to every
  clock, cohort or panel; endpoint-logit differences as individual effects; or
  proof of ANA-01–ANA-03.
- **Proof dependency:** none. ANA-02 establishes the analytical scalarization
  boundary; HUM-08 shows that the many-to-one consequence is visible in one
  fitted human model.
- **Known limitations:** one unweighted complete-case NHANES 2007–2012
  interface, one ordered 50-measurement panel, one linear ridge age decoder and
  one fold-local conditional-centring model. Contributions depend on the
  frozen model, scaling, reference law and correlated-feature parameterization.

## Frozen evidence

- 13,724 participant profiles with exact 50-feature additivity;
- 293 outcome-blind, non-reusing matched pairs;
- median absolute conditionally centred-gap difference, 0.124 decoder-output
  years;
- median contribution-vector L2 distance, 24.602 decoder-output years;
- median cosine similarity, 0.003;
- median top-five-contributor Jaccard overlap, 0.429;
- maximum decoder reproduction error, 1.42 × 10⁻¹⁴;
- maximum contribution-sum error, 4.44 × 10⁻¹⁴.

No confirmatory population-effect threshold was prespecified. Artifact identity,
fold identity, feature order, residualizer reload and contribution additivity
were fail-closed requirements and all passed. Independent validation reproduced
the pairing, display selection and numerical summaries.

## Placement contract

- no Summary sentence;
- no main figure;
- one bounded Results paragraph;
- one Extended Data figure;
- full Methods and Supplementary Methods;
- at most one Discussion sentence.

The submission figure is a fit-free three-panel successor assembled only from
frozen tables. It excludes the endpoint-logit panel and does not depend on the
older adjusted-AgeGap endpoint comparator.

## Provenance

- specification:
  \`analysis/NHANES_INDIVIDUAL_AGEGAP_CONTRIBUTIONS_SPEC_20260831.md\`
- analysis:
  \`analysis/analyze_nhanes_individual_agegap_contributions_v0_1.py\`
- report and validation:
  \`outputs/nhanes_individual_agegap_contributions_v0_1/\`
- fit-free submission figure:
  \`outputs/nhanes_individual_agegap_contributions_figure_v0_2/\`
- plotting entry:
  \`analysis/plot_nhanes_individual_agegap_contributions_extended_data_v0_2.py\`
