## Supplementary_Software_1 for "Inferential boundaries of age prediction: why prediction does not establish biological age measurement": PROJECT_STATE.md

# Review package navigation state — 22 September 2026

Use [the review-text index](README.md). The separately supplied
`Manuscript_Zotero_20260921.docx` and `Supplementary_Information.pdf` are the
review texts designated for this submission. This software directory contains
historical manuscript snapshots, including V6.27 / SI V5.19 dated 14 September
2026; their text and original version labels are unchanged.

See [the software guide](../../README.md) for access terms and stored-result
verification, and [the current archive inventory](../../reproducibility/ARCHIVE_MEMBERS_20260922.tsv)
for this navigation update. Existing dated reproducibility manifests remain
historical records. This update does not establish author declarations,
public licensing, or complete source-to-model reconstruction.
