## Supplementary_Software_1 for "Inferential boundaries of age prediction: why prediction does not establish biological age measurement": README.md

# Review texts and historical manuscript snapshots

The current review texts are the separately supplied
`Manuscript_Zotero_20260921.docx` and `Supplementary_Information.pdf`, as designated
for this submission on 22 September 2026. They are outside this software archive.
See [review-text selection and identities](../../CURRENT_MANUSCRIPT_VERSION.md).

The files below are unchanged historical analysis-era snapshots, not the current
review manuscript or SI:

- [Main manuscript snapshot, 14 September 2026](MANUSCRIPT_NATURE_IDENTIFICATION_FIRST_v6_27_20260914.md)
- [Supplementary Information snapshot, 14 September 2026](SUPPLEMENTARY_INFORMATION_IDENTIFICATION_FIRST_v5_19_20260914.md)
- [Reproducibility scope recorded on 14 September 2026](REPRODUCIBILITY_MANIFEST_v6_27_20260914.md)
- [Evidence registry supplied with that analysis snapshot](EVIDENCE_MODULE_REGISTRY.tsv)

The old Markdown text, author placeholders, version labels and dated manifests
are retained without rewriting. Statements such as "current" inside those dated
records apply to their original snapshot, not to the review-text selection above.
For this archive's member identities, use
[the 22 September inventory](../../reproducibility/ARCHIVE_MEMBERS_20260922.tsv).
Earlier member inventories remain historical records.

Start with [the software guide](../../README.md) and the
[stored-result verification instructions](../../reproducibility/README.md).
Frozen analysis scripts and historical input manifests retain their original
identities. Complete raw-source-to-model reexecution is not established.
Editorial/reviewer access does not grant a new public reuse licence.
