## Supplementary_Software_1 for "Inferential boundaries of age prediction: why prediction does not establish biological age measurement": REPRODUCIBILITY_MANIFEST_v6_9_20260827.md

# Reproducibility manifest for manuscript v6.9 / Supplementary Information v5.4

Date: 2026-08-28  
Scope: repository-level provenance for `Chronological-age decoding does not identify biological age`

## Purpose and boundary

This manifest indexes the executable contracts, frozen interfaces, fitted-object archives, figure resources and reconstruction records used by manuscript v6.9 and Supplementary Information v5.4. It is an engineering provenance map, not part of the scientific argument. The Article and Supplementary Information retain the populations, estimands, models, leakage controls, uncertainty procedures and interpretation boundaries required to assess the work; version histories, object inventories, internal decisions and file-by-file routes are kept here so that they do not obscure those methods.

No analysis was fitted or refitted for the v6.9/v5.4 precision revision. Its new Figure 1, Figure 2 and Extended Data Figure 4 packages are deterministic renderings of frozen analytical or conceptual inputs. The v6.8/v5.3 submission pair and all predecessor outputs remain unchanged.

## Environment contract

- Human-maintained environment: `environment.yml`
- Exact win-64 lock: `environment.win-64.lock.txt`
- Machine-readable policy: `environment-policy.json`
- Environment gate: `scripts/Test-ProjectEnvironment.ps1`
- Python wrapper: `scripts/Invoke-ProjectPython.ps1`

All LHState Python work is run in the pinned mamba-managed `lhstate` environment. The output-level manifests record interpreter and package versions where applicable.

## Literature evidence

The two workbooks have different evidentiary roles and neither contributes to the mathematical proof.

- Contemporary construct-promotion audit: `Papers/Audit/outputs/age-unit-construct-typical-examples-2024-2026/age_unit_construct_audit_2024_2026.xlsx`
- Critical-literature and remedy map: `Papers/Audit/outputs/age-clock-problem-remedy-2024-2026/age_clock_problem_remedy_2024_2026.xlsx`

Main Table 1 uses only promotion-positive records supported by source-level claim locators. Named endpoint, pace, mechanism and peripheral-clock statements are boundary cases unless the source explicitly promotes a broader construct. The v6.9 audit removed the endpoint-trained aggregate row because OMICmAge, the audited transcriptomic mortality model, DunedinPACE and AccelerAge were coded as bounded or boundary cases under the project codebook; an age unit alone was not treated as proof of a broader author claim.

## Frozen human modules

### HSCT referent-affiliation module (`HUM-01`)

Primary frozen analysis:

- `analysis/analyze_hsct_fixed_score_referent_null_probe_v0_1.py`
- `analysis/audit_hsct_fixed_score_referent_null_probe_v0_1.py`
- `outputs/hsct_fixed_score_referent_null_probe_v0_1/`

Public-source reconstruction and score implementation:

- `analysis/analyze_hsct_clock_object_replacement_v0_1.py`
- `analysis/extract_holland2024_horvath2013_v0_1.py`
- `analysis/analyze_holland2024_horvath_object_affiliation_v0_1.py`
- `analysis/freeze_holland2024_horvath2013_provenance_v0_1.py`

Continuous anchor-coordinate sensitivity:

- `analysis/analyze_hsct_referent_regression_sensitivity_v0_1.py`
- `analysis/audit_hsct_referent_regression_sensitivity_v0_1.py`
- `outputs/hsct_referent_regression_clock_sensitivity_v0_1/regression/`
- `outputs/hsct_referent_regression_clock_sensitivity_v0_1_independent_audit_v0_1/`

The primary statistic is the sign-balanced affiliation index under a within-sign geometry-preserving score-pairing null. The continuous model shares the `-A_R` anchor on both sides and is retained only as an anchor-coordinate compatibility sensitivity. Holland supplies the primary estimate; Søraas is separately processed complementary evidence. The two programmes are not independent cohort replications.

The prespecified Hannum-2013 fixed-coefficient sensitivity stopped before scoring because the complete-file transfer criterion was not met. Coverage was not evaluated and no Hannum score, regression or affiliation estimate was computed. The stopped implementation record is in `outputs/hsct_referent_regression_clock_sensitivity_v0_1/hannum2013/`; no raw-IDAT reprocessing was performed.

### Continuous-NHANES pure age decoder (`HUM-04`)

- Analysis: `analysis/analyze_nhanes_5y_age_decoder_source_information_probe_v0_1.py`
- Frozen output: `outputs/nhanes_5y_age_decoder_source_information_probe_v0_2/`
- Numerical report: `outputs/nhanes_5y_age_decoder_source_information_probe_v0_2/REPORT.md`

Key machine-readable objects include `ordered_source_features.tsv`, `pooled_metrics.tsv`, `paired_contrasts.tsv`, `fold_metrics.tsv`, `age_decoder_diagnostics.json`, `model_artifact_audit.json` and `decision.json`. Five outer artifacts contain fold-local preprocessing, chronological-age decoder fitting and endpoint models. The archived audit indexes 90 age-tuning and 180 endpoint-tuning estimators; reloaded score and probability outputs matched the reported predictions exactly. The author accepted the bounded v0.2 result without confirmation refitting or subsequent model, endpoint, panel or threshold search.

### Supporting continuous-NHANES matched scalar comparison

- Analysis: `analysis/analyze_nhanes_5y_matched_scalar_target_supervision_probe_v0_2.py`
- Frozen output: `outputs/nhanes_5y_matched_scalar_target_supervision_probe_v0_2/`
- Numerical report: `outputs/nhanes_5y_matched_scalar_target_supervision_probe_v0_2/REPORT.md`

The formal run persisted 25 matched-comparison and five flexible-calibration artifacts containing 19,085 estimators. The audit reconstructed the fully nested projection and downstream score chain; maximum score and probability discrepancies were at floating-point tolerance. The earlier non-nested version is retained only as versioned audit provenance and does not support the reported result.

### NHANES III matched-scalar protocol transfer (`HUM-06`)

- Contract: `analysis/NHANES3_5Y_MATCHED_SCALAR_TARGET_SUPERVISION_PROTOCOL_TRANSFER_SPEC_v0_2_20260827.md`
- Analysis: `analysis/analyze_nhanes3_5y_matched_scalar_target_supervision_protocol_transfer_v0_2.py`
- Frozen output: `outputs/nhanes3_5y_matched_scalar_target_supervision_protocol_transfer_v0_2/`
- Numerical report: `outputs/nhanes3_5y_matched_scalar_target_supervision_protocol_transfer_v0_2/REPORT.md`

The current run preserves the v0.1 scientific design and changes only the tolerance used when comparing serialized human-readable numeric cells. Its internal decision label is `UTILITY_WITHOUT_PRIVILEGE_PROTOCOL_TRANSFER`. The archive contains the two Phase 1 projections and 505 Phase 2 downstream artifacts, including all Fay-BRR refits; design and prediction reconstruction errors were zero at the declared tolerance.

### NHANES III observed-age contextual module (`HUM-03`)

- Contract: `analysis/NHANES3_5Y_CHRONOLOGICAL_AGE_INCREMENT_AND_PROXY_COMPARISON_SPEC_v0_1_20260827.md`
- Analysis: `analysis/analyze_nhanes3_5y_chronological_age_increment_and_proxy_comparison_v0_1.py`
- Frozen output: `outputs/nhanes3_5y_chronological_age_increment_and_proxy_comparison_v0_1/`
- Numerical report: `outputs/nhanes3_5y_chronological_age_increment_and_proxy_comparison_v0_1/REPORT.md`

The archive contains 20 primary models and 640 full-pipeline Fay-BRR models. All persisted designs, tuning losses and probabilities reconstructed at the declared absolute tolerance of `1e-12`.

### NHANES III observed-age versus transported-decoder boundary

- Contract: `analysis/NHANES3_5Y_AGE_DECODER_SUBSTITUTION_COMPARISON_SPEC_v0_1_20260827.md`
- Analysis: `analysis/analyze_nhanes3_5y_age_decoder_substitution_comparison_v0_1.py`
- Frozen output: `outputs/nhanes3_5y_age_decoder_substitution_comparison_v0_1/`
- Numerical report: `outputs/nhanes3_5y_age_decoder_substitution_comparison_v0_1/REPORT.md`

The internal decision label is `NO_CLEAR_SUBSTITUTION_ADVANTAGE`. All 330 expected downstream artifacts were present and reconstructed with zero design and prediction discrepancy at the declared tolerance.

The three NHANES III modules inherit the canonical Phase 1/Phase 2 interfaces, ordered 25-measurement manifest, survey-design objects, folds and source-selection freeze from `outputs/nhanes3_phase1_to_phase2_endpoint_vs_age_acquisition_v0_2_final_20260826/`. They do not modify that directory.

### Endpoint-directed acquisition (`MTH-01`)

- Analysis: `analysis/analyze_nhanes_endpoint_vs_inverse_age_decoder_acquisition_v0_2.py`
- Frozen output: `outputs/nhanes_endpoint_vs_inverse_age_decoder_acquisition_v0_2/`
- Numerical report: `outputs/nhanes_endpoint_vs_inverse_age_decoder_acquisition_v0_2/REPORT.md`

The persisted audit indexes 540 inverse-decoder tuning estimators, 30 inverse-decoder final estimators and 60 endpoint estimators. Reloaded predictions matched the sealed predictions at the recorded tolerances. The scientific comparison treats each policy as selecting one domain from the same five prespecified measurement-domain options. Domains differ in measurement count and no monetary, economic or real-world resource equivalence is asserted.

## Transparent X1–X3 construction (`SYN-02`)

- Contract: `analysis/THREE_VARIABLE_AGE_STATE_TASK_MAP_PROBE_SPEC_v0_1_20260827.md`
- Generator: `analysis/simulate_three_variable_age_state_geometry_v0_1.py`
- Analysis: `analysis/analyze_three_variable_age_state_task_map_probe_v0_1.py`
- Figure builder: `analysis/plot_three_variable_age_state_task_map_figure_v0_2.py`
- Frozen analysis: `outputs/three_variable_age_state_task_map_probe_v0_1/`
- Figure package: `outputs/three_variable_age_state_task_map_figure_v0_2/`

The frozen 500-row interface, participant metadata, fold assignments, ordered features, predictions, result tables and 45 fitted pipelines are persisted. Immediate reload reproduced all held-out predictions at tolerance `1e-12`. The construction is explanatory; the algebraic limiting cases, not these fitted models, establish the strict non-implications.

## Submission figure routes

Main figures:

1. `outputs/v6_9_figure1_theory_chain_v0_1/` — generated by `analysis/plot_v6_9_figure1_precision_revision_v0_1.py`; fit-free.
2. `outputs/hsct_age_reconstruction_semantic_counterexample_figure_v0_4/` — generated by `analysis/plot_hsct_age_reconstruction_semantic_counterexample_v0_4.py` from frozen `HUM-01` inputs; fit-free.
3. `outputs/nhanes_hum03_hum04_hum06_figure3_successor_v0_3/` — combines separately labelled continuous-NHANES and NHANES III interfaces without pooling.
4. `outputs/v6_8_figure4_target_first_framework_v0_1/` — conceptual target-first framework; reused unchanged.

Extended Data figures:

1. `outputs/three_variable_age_state_task_map_figure_v0_2/`
2. `outputs/v6_2_submission_extended_data_v0_1/extended_data_figure_2/`
3. `outputs/nhanes_hum03_hum04_hum06_extended_data_figure3_successor_v0_2/`
4. `outputs/v6_9_extended_data_figure4_domain_options_v0_1/` — generated by `analysis/plot_v6_9_extended_data_figure4_domain_options_v0_1.py`; fit-free.

Every figure package contains its image files, panel Source Data, legend, chart contract, input manifest and visualization audit. The package-level manifests carry exact file digests.

## Submission build and QA

- Manuscript source: `manuscript/nature_first/MANUSCRIPT_NATURE_IDENTIFICATION_FIRST_v6_9_20260827.md`
- Supplementary source: `manuscript/nature_first/SUPPLEMENTARY_INFORMATION_IDENTIFICATION_FIRST_v5_4_20260827.md`
- DOCX builder: `manuscript/nature_first/scripts/build_v6_9_docx.py`
- Main figure resource index: `manuscript/nature_first/V6_9_MAIN_FIGURE_RESOURCE_INDEX_20260827.md`
- Extended Data resource index: `manuscript/nature_first/V6_9_EXTENDED_DATA_RESOURCE_INDEX_20260827.md`

The v6.9 build forces a page break before Figure 1. Final QA records document summary length, forbidden-phrase checks, table classification, figure routing, mathematical and citation cross-references, DOCX/PDF rendering and page-by-page visual inspection.

## Consolidated reconstruction statement

All reported fitted analyses were reconstructed after reload from persisted data, folds, preprocessing and model objects, with participant identity, fold assignment, ordered features and predictions matching the reported results at the prespecified tolerances. Routine reruns validate and reuse compatible objects; they do not silently refit frozen evidence.
