## Supplementary_Software_1 for "Inferential boundaries of age prediction: why prediction does not establish biological age measurement": REPRODUCIBILITY_MANIFEST_v6_13_20260831.md

# Reproducibility manifest v6.13

Date: 31 August 2026
Scope: bounded HUM-08 evidence integration

## Scientific pair

- Main: \`MANUSCRIPT_NATURE_IDENTIFICATION_FIRST_v6_13_20260831.md\`
- Supplement: \`SUPPLEMENTARY_INFORMATION_IDENTIFICATION_FIRST_v5_5_20260831.md\`

V6.13 inherits the complete v6.12 / v5.4 evidence package and adds one
materiality module: HUM-08. The new module reuses the frozen continuous-NHANES
linear age decoder and validated
\`nhanes_individual_agegap_contributions_v0_1\` analysis. No decoder or endpoint
model was refitted. The submission Extended Data Figure 4 is a fit-free
three-panel successor assembled only from frozen pair and contribution tables.
The former acquisition Extended Data Figure 4 is scientifically unchanged and
is routed as Extended Data Figure 5.

## HUM-08 analysis and persistence

- frozen specification:
  \`analysis/NHANES_INDIVIDUAL_AGEGAP_CONTRIBUTIONS_SPEC_20260831.md\`
- analysis entry:
  \`analysis/analyze_nhanes_individual_agegap_contributions_v0_1.py\`
- frozen output:
  \`outputs/nhanes_individual_agegap_contributions_v0_1/\`
- independent validation:
  \`outputs/nhanes_individual_agegap_contributions_v0_1/VALIDATION_REPORT.md\`
- fit-free plot entry:
  \`analysis/plot_nhanes_individual_agegap_contributions_extended_data_v0_2.py\`
- fit-free figure and Source Data:
  \`outputs/nhanes_individual_agegap_contributions_figure_v0_2/\`
- module contract:
  \`manuscript/nature_first/next_round_20260831/HUM_08_NHANES_INDIVIDUAL_AGEGAP_REALIZATION_CONTRACT_20260831.md\`

The frozen audit reports 13,724 participant profiles, 686,200
participant-by-feature contribution rows, five persisted residualizers, 90
persisted tuning estimators, exact decoder reproduction and exact additive
parity. The independent validation reproduced all 293 non-reusing matched
pairs, both display selections and the reported summaries.

## Governance and checks

- direction lock:
  \`next_round_20260831/V6_13_DIRECTION_LOCK_20260831.md\`
- version map:
  \`next_round_20260831/V6_13_VERSION_MAP_20260831.md\`
- main-figure index:
  \`V6_13_MAIN_FIGURE_RESOURCE_INDEX_20260831.md\`
- Extended Data index:
  \`V6_13_EXTENDED_DATA_RESOURCE_INDEX_20260831.md\`
- evidence QA:
  \`qa/Test-V6_13ManuscriptEvidence.ps1\`
- DOCX structure QA:
  \`qa/Test-V6_13DocxStructure.ps1\`
- submission DOCX builder:
  \`scripts/build_submission_v6_13_docx.py\`
- registry validator:
  \`../../../.agents/skills/lhstate-integrate-manuscript-evidence/scripts/Test-EvidenceRegistry.ps1\`

All v6.9–v6.12 resources not named above are inherited unchanged. The v6.12
candidate and every earlier frozen output remain untouched.
