## Supplementary_Software_1 for "Inferential boundaries of age prediction: why prediction does not establish biological age measurement": REPRODUCIBILITY_MANIFEST_v6_16_20260907.md

# Reproducibility manifest v6.16

Date: 7 September 2026
Scope: full publication-writing successor to v6.15 / SI v5.7

## Current documents

- Main: MANUSCRIPT_NATURE_IDENTIFICATION_FIRST_v6_16_20260907.md
- SI: SUPPLEMENTARY_INFORMATION_IDENTIFICATION_FIRST_v5_8_20260907.md
- Cover: COVER_LETTER_NATURE_v6_16_20260907.md
- Writing contract: next_round_20260907/V6_16_WRITING_DIRECTION_AND_VERSION_MAP_20260907.md
- Reference map: qa/V6_16_REFERENCE_RENUMBERING_20260907.json
- Content QA: qa/V6_16_CONTENT_QA_20260907.md
- Candidate: submission_20260907_v6_16_candidate/

## Frozen evidence and the scope of revision

All analyses, populations, endpoints, fitted objects, predictions, panels,
folds, thresholds and numerical results are inherited unchanged. The main
text is rewritten around the methodological consequences of the
identification result. The SI synchronizes six local mathematical,
prior-art and empirical-interpretation clarifications. The 46 source
references are unchanged; their display numbers now follow first citation.
All nine SI tables retain their scientific content.

Main Figures 1–4 and Extended Data Figures 1–4 retain the prior submission
images. Extended Data Figure 5 has a versioned label-spacing derivative,
with identical source data and plotted values. The v6.13 analytical
manifest continues to locate underlying analyses and model objects.
The v6.15 figure indexes supply inherited resource paths except for the
explicit ED5 routing below.

- ED5 layout derivative: outputs/v6_16_extended_data_figure5_layout_v0_1/
- Layout-only plotting script: analysis/plot_v6_16_extended_data_figure5_layout_v0_1.py
- Historical figure indexes: V6_15_MAIN_FIGURE_RESOURCE_INDEX_20260904.md and V6_15_EXTENDED_DATA_RESOURCE_INDEX_20260904.md

## Literature supplements

Supplementary Data 1 and 2 are unchanged. A targeted read-only check of the
Sluiskes source-map row found no erroneous restriction to an additive-only
theorem or unsupported priority claim. Neither workbook embeds manuscript
reference numbers, so renumbering requires no workbook edit.

## Reproduction and release

- DOCX builder: scripts/build_submission_v6_16_docx.py (inherits v6.15/v6.9)
- PDF export: scripts/Export-V6_16WordPdf.ps1, using a hidden Word instance
  and read-only document opening.
- PDF rendering: bundled Poppler; manuscript and SI page inspection is
  recorded in the candidate QA report.
- Runtime: pinned LHState mamba environment through Invoke-ProjectPython.ps1.
- Registry validator: .agents/skills/lhstate-integrate-manuscript-evidence/scripts/Test-EvidenceRegistry.ps1

Source_Data.zip retains the frozen aggregate data. The new
Supplementary_Software_1.zip preserves all inherited analysis entries;
only current governance metadata and the fit-free ED5 layout script are
added or synchronized. The package records entry-level integrity checks.
No participant-level derived contributions or fitted-model binaries are
added to the submission archive.

Author metadata, acknowledgements, funding, CRediT, conflict declarations
and a public repository persistent identifier remain author-side items.
