## Supplementary_Software_1 for "Inferential boundaries of age prediction: why prediction does not establish biological age measurement": REPRODUCIBILITY_MANIFEST_v6_17_20260908.md

# Reproducibility manifest v6.17

Date: 8 September 2026
Scope: publication-writing successor to v6.16 / SI v5.8

## Current documents

- Main: MANUSCRIPT_NATURE_IDENTIFICATION_FIRST_v6_17_20260908.md
- SI: SUPPLEMENTARY_INFORMATION_IDENTIFICATION_FIRST_v5_9_20260908.md
- Cover: COVER_LETTER_NATURE_v6_17_20260908.md
- Writing contract: next_round_20260908/V6_17_WRITING_DIRECTION_AND_VERSION_MAP_20260908.md
- Reference numbering: qa/V6_16_REFERENCE_RENUMBERING_20260907.json (unchanged)
- Content QA: qa/V6_17_CONTENT_QA_20260908.md
- Candidate: submission_20260908_v6_17_candidate/

## Frozen scientific resources

This revision changes prose, not analyses. All equations, fitted objects,
populations, panels, folds, endpoints, predictions, numerical results, Methods,
Table 1 and the nine SI tables are retained. The same 46 references retain
their numbering. Four main and five Extended Data figures are inherited
byte-for-byte from the v6.16 release, including its ED5 label-spacing derivative.

The v6.13 analytical manifest continues to locate underlying analyses and
persisted model objects. The v6.15 figure indexes locate inherited resources,
with the ED5 exception recorded in the v6.16 manifest:

- REPRODUCIBILITY_MANIFEST_v6_13_20260831.md
- REPRODUCIBILITY_MANIFEST_v6_16_20260907.md
- V6_15_MAIN_FIGURE_RESOURCE_INDEX_20260904.md
- V6_15_EXTENDED_DATA_RESOURCE_INDEX_20260904.md
- ED5: outputs/v6_16_extended_data_figure5_layout_v0_1/

## Reproduction and packaging

- DOCX: scripts/build_submission_v6_16_docx.py, using the unchanged v6.15/v6.9
  inherited builders and frozen figure resources.
- PDF: scripts/Export-V6_16WordPdf.ps1, hidden Word instance, read-only opening.
- Page rendering: bundled Poppler; visual checks recorded in candidate QA.
- Runtime: pinned LHState mamba environment via Invoke-ProjectPython.ps1.
- Registry validator: .agents/skills/lhstate-integrate-manuscript-evidence/scripts/Test-EvidenceRegistry.ps1

Source_Data.zip and both literature workbooks are unchanged. The software ZIP
preserves all inherited analysis entries; only current publication metadata is
synchronized, with old metadata retained as provenance. The release records
document-copy parity, unchanged-figure/data parity and ZIP-entry verification.
No participant-level derived contributions or fitted-model binaries are added.

Original manuscripts and frozen analysis artifacts remain intact. Author
metadata, acknowledgements, funding, CRediT, conflicts and public repository
deposition remain author-side items.
