## Supplementary_Software_1 for "Inferential boundaries of age prediction: why prediction does not establish biological age measurement": REPRODUCIBILITY_MANIFEST_v6_18_20260908.md

# Reproducibility manifest v6.18

Date: 8 September 2026
Scope: local writing and Figure 2 semantic synchronization from v6.17 / SI v5.9.

## Current documents

- Main: MANUSCRIPT_NATURE_IDENTIFICATION_FIRST_v6_18_20260908.md
- SI: SUPPLEMENTARY_INFORMATION_IDENTIFICATION_FIRST_v5_10_20260908.md
- Cover: COVER_LETTER_NATURE_v6_18_20260908.md
- Contract: next_round_20260908/V6_18_LOCAL_SYNCHRONIZATION_CONTRACT_20260908.md
- Reference numbering: qa/V6_16_REFERENCE_RENUMBERING_20260907.json (unchanged)
- Content QA: qa/V6_18_CONTENT_QA_20260908.md
- Candidate: submission_20260908_v6_18_candidate/

## Scientific resources and changes

All fitted objects, populations, panels, folds, endpoints, predictions,
numerical estimates and equations are inherited unchanged. Main Methods and
all nine SI tables are unchanged. The same 46 references retain numbering.
Main Table 1 now records substantive external and downstream validation already
present in its three source papers and Supplementary Data 1. It adds no new
study or empirical result. The gap-decomposition language is corrected from
identifying components to separating them algebraically.

Figure 2 uses a fit-free v0.5 successor that changes conceptual labels and node
layout. The b/c quantitative Source Data are unchanged; the a/d conceptual
Source Data and bilingual legend are synchronized to the pairing-null result.
This revision adds no conditional host/donor attribution analysis.

- Figure 2: outputs/hsct_age_reconstruction_semantic_counterexample_figure_v0_5/
- Figure 2 script: analysis/plot_hsct_age_reconstruction_semantic_counterexample_v0_5.py
- Underlying analytical objects: REPRODUCIBILITY_MANIFEST_v6_13_20260831.md
- Inherited main/ED resource indexes: V6_15_MAIN_FIGURE_RESOURCE_INDEX_20260904.md
  and V6_15_EXTENDED_DATA_RESOURCE_INDEX_20260904.md
- Inherited ED5 label-layout exception: REPRODUCIBILITY_MANIFEST_v6_16_20260907.md
- Prior writing release: REPRODUCIBILITY_MANIFEST_v6_17_20260908.md

All figures other than Figure 2 are copied unchanged from v6.17.

## Build and release

- DOCX: scripts/build_submission_v6_18_docx.py, inheriting the v6.16/v6.15/v6.9
  builders; routes the new Figure 2 and gives Table 1 deliberate column widths.
- PDF: hidden Word read-only export via scripts/Export-V6_16WordPdf.ps1.
- Rendering: bundled Poppler, followed by page inspection.
- Runtime: pinned LHState mamba environment through Invoke-ProjectPython.ps1.
- Registry: 13 current modules retain roles, estimands and analysis versions;
  publication mappings change, with HUM-01's figure source redirected to v0.5.

The data archive preserves all quantitative entries and synchronizes only
Figure 2 conceptual metadata. Literature workbooks are unchanged. The software
archive preserves inherited analytical entries, retains superseded metadata as
provenance and adds the fit-free figure script and current governance metadata.
Package verification records the exact changed and preserved entries.
Participant-level contributions and fitted-model binaries are not added.

Original manuscripts, original figures and frozen analysis outputs remain
intact. Author administrative fields and public repository deposition remain
open; this revision does not submit the manuscript.
