## Supplementary_Software_1 for "Inferential boundaries of age prediction: why prediction does not establish biological age measurement": REPRODUCIBILITY_MANIFEST_v6_19_20260908.md

# Reproducibility manifest v6.19

Date: 8 September 2026
Scope: final local text and limiting-construction correction from v6.18 / SI v5.10.

## Current documents

- Main: MANUSCRIPT_NATURE_IDENTIFICATION_FIRST_v6_19_20260908.md
- SI: SUPPLEMENTARY_INFORMATION_IDENTIFICATION_FIRST_v5_11_20260908.md
- Cover retained unchanged: COVER_LETTER_NATURE_v6_18_20260908.md
- Contract: next_round_20260908/V6_19_FINAL_LOCAL_CORRECTION_20260908.md
- Content QA: qa/V6_19_CONTENT_QA_20260908.md
- Candidate: submission_20260908_v6_19_candidate/

## Scientific scope

The first mathematical limiting example now specifies constant background,
uniform continuous age and independent Gaussian noise of positive variance.
Its positive conditional age density guarantees positive population residual
variance. SM5 refers to that example; main and SI distinguish variation in the
gap from state information present in the source measurements. One Introduction
sentence is synchronized to the established methodological contribution.

No fitted object, data row, panel, fold, endpoint, prediction, numerical estimate,
figure, table or citation number changes. The theoretical example is not a revised
generator for the frozen 500-row X1–X3 construction. The title, 187-word Summary,
Results order, Discussion and scientific Methods remain unchanged. One Methods
availability sentence now names fold-construction rules rather than implying
that participant-level fold assignments are included in the supplied Source Data.

## Inherited resources

- Immediate resource and figure authority: REPRODUCIBILITY_MANIFEST_v6_18_20260908.md
- Underlying analytical resources: REPRODUCIBILITY_MANIFEST_v6_13_20260831.md
- Figure 2 remains the v0.5 semantic derivative; all nine figure resources unchanged.
- Source_Data.zip and both literature workbooks are inherited in full from v6.18.
- Software archive retains all scientific entries; current entry-point metadata
  is synchronized and superseded metadata retained as provenance.
- The pre-existing `analysis/NHANES_5Y_AGE_DECODER_SOURCE_INFORMATION_PROBE_SPEC_20260823.md`
  is added unchanged after the availability check found it absent from the prior ZIP.
  Archive entry points explicitly describe their dependence on retained predecessor
  data interfaces and the original project layout; path/hash records are locators,
  not the underlying data or fitted objects.

## Build and verification

- DOCX: unchanged scripts/build_submission_v6_18_docx.py.
- PDF: hidden read-only Word export via scripts/Export-V6_16WordPdf.ps1.
- Rendering: bundled Poppler; final page inspection.
- Runtime: pinned LHState mamba environment through Invoke-ProjectPython.ps1.
- Registry: thirteen current modules retain estimands, roles, placement and analysis
  versions; only manuscript/SI publication mappings change.

Local checks cover the repaired example, document/source consistency, unchanged
empirical content and archive availability/integrity. They are not a clean-machine
reproduction of fitted models, P values or confidence intervals. Public DOI
deposition and author administrative fields remain open. No upload or submission
is performed by this revision.
