## Supplementary_Software_1 for "Inferential boundaries of age prediction: why prediction does not establish biological age measurement": REPRODUCIBILITY_MANIFEST_v6_20_20260909.md

# Reproducibility manifest V6.20

Date: 9 September 2026.
Scope: minimal final manuscript cleanup from main V6.19 / SI V5.11.

## Current documents

- Main: MANUSCRIPT_NATURE_IDENTIFICATION_FIRST_v6_20_20260909.md
- SI: SUPPLEMENTARY_INFORMATION_IDENTIFICATION_FIRST_v5_12_20260909.md
- Cover unchanged: COVER_LETTER_NATURE_v6_18_20260908.md
- Cleanup note: next_round_20260908/V6_20_FINAL_CLEANUP_20260909.md
- Content QA: qa/V6_20_CONTENT_QA_20260909.json
- Candidate: submission_20260909_v6_20_candidate/

## Scope and retained resources

The main Methods replaces one editorial-selection sentence with method
cross-references. SI corrects monotone-calibration wording and the bootstrap
resample count's referent, removes one duplicated qualification and one
production instruction, and matches planned archive wording to supplied files.
The SI manifest pointer advances. All equations, empirical results, tables,
reference numbers, figure resources/legends and the cover remain unchanged.

The immediate resource authority is REPRODUCIBILITY_MANIFEST_v6_19_20260908.md;
its inherited manifests retain exact scientific routes and historical versions.
Both literature workbooks, Source_Data.zip and all figure PNG/PDF resources are
copied unchanged from the V6.19 candidate. Software scientific entries remain
unchanged; publication metadata advance, with displaced metadata retained under
provenance/v6_19/. No model or participant record is changed or refitted.

The software remains a code, specification and provenance snapshot, dependent
where indicated on retained predecessor interfaces and fitted objects. It does
not supply every participant-level fold assignment or fitted-model binary.
Paths and hashes are locators, not the corresponding objects. Public deposition,
a persistent identifier and clean-machine reproduction remain incomplete.

## Build and verification boundary

Use the existing scripts/build_submission_v6_18_docx.py builder and hidden
read-only Word PDF export; rasterize the final PDFs with bundled Poppler.
Python operations use the pinned LHState wrapper after the environment gate.
Verify the exact baseline-to-successor delta, source/Word agreement, unchanged
mathematics/tables/figures, and final page images. Thirteen current registry
rows change publication mappings only. No analytical estimates are recomputed.
