## Supplementary_Software_1 for "Inferential boundaries of age prediction: why prediction does not establish biological age measurement": REPRODUCIBILITY_MANIFEST_v6_21_20260909.md

# Reproducibility manifest V6.21

Date: 9 September 2026. Parent: V6.20 / SI V5.12.

Current main: `MANUSCRIPT_NATURE_IDENTIFICATION_FIRST_v6_21_20260909.md`.
Current SI: `SUPPLEMENTARY_INFORMATION_IDENTIFICATION_FIRST_v5_13_20260909.md`.
Retained cover: `COVER_LETTER_NATURE_v6_18_20260908.md`.
Candidate: `submission_20260909_v6_21_candidate/`.
Review response: `next_round_20260909/V6_21_REVIEW_ABSORPTION_20260909.md`.

The Methods states direct five-year event-probability modelling. One main
sentence and one SI paragraph acknowledge the related Grødem brain-age
preprint. Reference 23 is inserted; previous 23–46 advance by one in the main
and SI. All previous bibliography entries, scientific equations, numerical
results, tables, figure assets and scientific legend content are retained.
The supplementary 20-source review and both workbooks are unchanged.

The scientific resource authority remains
`REPRODUCIBILITY_MANIFEST_v6_20_20260909.md` and its predecessor manifests.
No model is fitted and no scientific estimate is added or updated.

The software successor adds the existing continuous-NHANES raw-interface and
complete-support baseline/direct-source model generators, their required
specification, the environment verifier and execution/input documentation.
`CORE_NHANES_REPRODUCTION.md` names each predecessor interface and its generator.
The `CORE_NHANES_INPUT_INVENTORY.tsv` and `CORE_NHANES_INTERFACE_QA.json` files
record a local public-source-to-interface check: all 13,724 participant rows,
470 events, 50 ordered features and five folds exactly match frozen objects
after write/reload. The private reconstructed rows are not included in the
submission. Existing scientific software entries remain byte-identical.

This validates interface reconstruction using retained public files, not an
external source-download-to-prediction rerun. No prediction model was refitted.
The NHANES III stand-alone upstream reconstruction route remains unverified.
Figure reconstruction, recalculation from stored predictions and source-to-model
reexecution are separate verification levels. Public deposition and the
persistent archive identifier remain incomplete.

Document validation: `qa/V6_21_CONTENT_QA_20260909.json` and
`qa/v6_21_render/PAGE_DELTA.json`. Use the inherited V6.18 builder and hidden
read-only native Word export, followed by the document skill rasterizer.
Registry changes are limited to current publication mappings and review date.
Displaced software metadata are retained under `provenance/v6_20/`.
