## Supplementary_Software_1 for "Inferential boundaries of age prediction: why prediction does not establish biological age measurement": REPRODUCIBILITY_MANIFEST_v6_22_20260910.md

# Reproducibility manifest V6.22

Date: 10 September 2026. Parent: V6.21 / SI V5.13.

Current main: `MANUSCRIPT_NATURE_IDENTIFICATION_FIRST_v6_22_20260910.md`.
Current SI: `SUPPLEMENTARY_INFORMATION_IDENTIFICATION_FIRST_v5_14_20260910.md`.
Retained cover: `COVER_LETTER_NATURE_v6_18_20260908.md`.
Candidate: `submission_20260910_v6_22_candidate/`.
Change map: `next_round_20260910/V6_22_STRUCTURAL_EDIT_20260910.md`.

## Scientific authority and editorial scope

The scientific resource authority remains
`REPRODUCIBILITY_MANIFEST_v6_21_20260909.md` and its predecessor manifests.
This successor changes presentation, not estimates, models, estimands or
analytical proof responsibility. No participant data are analysed, no model is
loaded or fitted, no random numbers are generated and no new estimate is
computed for this revision.

The main has seven Results sections, pairing the two distinct NHANES predictive
comparisons without pooling them. Contextual-age contrasts are retained in
Extended Data Figure 3 and Supplementary Methods 3, with a substantive main
pointer. Individual AgeGap contributions retain a separate main subsection.
The title, 187-word Summary, displayed equations, scientific Methods and
availability statements, Table 1 and all 47 references are unchanged. SI
changes only the manifest pointer. Cover V6.18 and both literature workbooks
are unchanged.

## Figure and Source Data map

Figures 1–2 and Extended Data Figures 1–5 are inherited unchanged. New static
Figures 3–4 are built by
`analysis/plot_v6_22_main_figure_assembly_v0_1.py`, with resources under
`outputs/v6_22_main_figure_assembly_v0_1/`, both relative to the project root.

- Figure 3a,b: continuous NHANES design and its three frozen predictive
  contrasts; 50-measurement interface.
- Figure 3c,d: NHANES III Phase 1 projection/Phase 2 calibration design and
  the two frozen matched-scalar contrasts; 25-measurement interface. The new
  design diagram adds no statistical estimate.
- Former Figure 3d contextual-age results: complete in unchanged ED3/SM3.
- Figure 4a: target-first workflow; Figure 4b: the frozen exploratory
  acquisition gains and paired difference, with all original intervals;
  Figure 4c: the five-object evidence map. ED5 retains the full acquisition
  selection and fold diagnostics. No cost equivalence is implied.

The seven quantitative/conceptual source TSVs are byte-exact copies of their
frozen predecessors. Their index, source map, builder and render QA accompany
the software; Source Data uses the new panel paths. Main-manuscript legends
are the canonical submission captions. Figure assembly documentation may
retain proposed working legends as provenance.

The supplied aggregate tables support static reconstruction of Figures 3–4.
They do not turn the whole software snapshot into an independently verified
source-to-model execution bundle. The V6.21 local continuous-NHANES interface
check remains the authority for that distinct reconstruction level; it is not
rerun here. See `CORE_NHANES_REPRODUCTION.md`.

## Verification and remaining items

Content: `qa/V6_22_CONTENT_QA_20260910.json`.
Rendering: `qa/v6_22_render/PAGE_DELTA.json` and
`qa/V6_22_VISUAL_QA_20260910.md`.
Package: `submission_20260910_v6_22_candidate/PACKAGE_INTEGRITY.json`.
Build: `scripts/build_submission_v6_22_docx.py`, inherited Word builder and
hidden read-only native Word PDF export. All changed pages are visually
inspected; pixel-identical pages inherit the prior inspection. The existing
Cambria Math text-run equation representation is retained, not converted to
native Word equation objects.

Registry changes are limited to current publication mappings, review date,
HUM-03 placement and Figure 3/4 routing. Estimands, evidence roles, scientific
claims, forbidden claims and analysis versions are unchanged. Displaced
software metadata remain under `provenance/v6_21/`; older scientific software
and prior candidates remain recoverable.

Still open: full derivation-level comparison with Grødem; external end-to-end
model reexecution; an independently verified NHANES III upstream public route;
author administrative fields; and public deposition/DOI. This editorial
successor does not claim to complete any of those tasks.
