## Supplementary_Software_1 for "Inferential boundaries of age prediction: why prediction does not establish biological age measurement": REPRODUCIBILITY_MANIFEST_v6_23_20260911.md

# Reproducibility manifest V6.23

Date: 11 September 2026. Parent: V6.22 / SI V5.14.

Current main: `MANUSCRIPT_NATURE_IDENTIFICATION_FIRST_v6_23_20260911.md`.
Current SI: `SUPPLEMENTARY_INFORMATION_IDENTIFICATION_FIRST_v5_15_20260911.md`.
Cover: unchanged `COVER_LETTER_NATURE_v6_18_20260908.md`.
Candidate: `submission_20260911_v6_23_candidate/`.
Change note: `next_round_20260911/V6_23_LOCAL_REVISION_20260911.md`.

This is a bounded editorial successor. The Summary combines the complementary
NHANES comparisons and omits the separate individual-contribution headline.
That result remains unchanged in its own Results section, ED4 and SI. The ED3
information-set sentence moves before the NHANES section's concluding judgment.
The Figure 4 legend omits the version-relative word Existing. All scientific
results, equations, Methods, references, tables, section order and Discussion
remain unchanged. SI updates only this manifest locator.

Figure 4 changes only the common-menu connector: both selection objectives
are explicitly connected to the same candidate menu. The new static builder is
`analysis/plot_v6_23_figure4_connector_v0_1.py`; its output is
`outputs/v6_23_figure4_connector_v0_1/`, relative to the project root. The
V6.22 figure builder and aggregate source tables remain dependencies and are
retained in Supplementary Software 1. Main Figures 1–3, ED1–5, both literature
workbooks and the cover are inherited unchanged. Main-manuscript captions
remain canonical.

The parent package's standalone Figures 3 and 4 already used the axis title
Five-year mortality log-loss improvement, whereas its DOCX/PDF still embedded
earlier axis titles. The V6.23 build synchronizes those embedded images with
the existing standalone assets; Figure 4 additionally receives the authorized
connector change. No frozen image is overwritten. The nine current embedded
images are checked against current standalone exports in
`qa/V6_23_FIGURE_INHERITANCE_20260911.json`.

The scientific resource authority remains
`REPRODUCIBILITY_MANIFEST_v6_22_20260910.md` and its predecessors. Existing
aggregate Source Data are preserved; no new participant analysis, model fit,
estimate, resampling or random-number generation is performed. The static
figure correction is not a source-to-model reproduction.

The affected artifacts are checked with `qa/check_v6_23_content.py`, native
Word PDF export and the document-skill rasterizer. Changed pages are inspected
visually; pixel-identical pages inherit V6.22/V5.14 inspection. See
`qa/V6_23_CONTENT_QA_20260911.json`, `qa/V6_23_VISUAL_QA_20260911.md`,
`qa/v6_23_render/PAGE_DELTA.json` and the candidate's `PACKAGE_INTEGRITY.json`.
The existing Cambria Math text-run representation is retained; equations are
not newly converted to OMML. Registry edits are limited to publication
versions/dates and current Figure 4 resource paths.

Unresolved items are unchanged: full derivation-level comparison with Grødem;
external source-to-model execution; an independently verified NHANES III
upstream public route; author administrative fields; and public deposition/DOI.
The software remains a source/provenance snapshot rather than a certified
self-contained full-study execution bundle. Historical metadata displaced by
packaging are retained under `provenance/v6_22/`.
