## Supplementary_Software_1 for "Inferential boundaries of age prediction: why prediction does not establish biological age measurement": REPRODUCIBILITY_MANIFEST_v6_24_20260911.md

# Reproducibility manifest V6.24 / SI V5.16

Date: 11 September 2026. Parent V6.23 / SI V5.15 remains intact.

Current sources: `MANUSCRIPT_NATURE_IDENTIFICATION_FIRST_v6_24_20260911.md`
and `SUPPLEMENTARY_INFORMATION_IDENTIFICATION_FIRST_v5_16_20260911.md`.
Cover: unchanged `COVER_LETTER_NATURE_v6_18_20260908.md`.
Candidate: `submission_20260911_v6_24_candidate/`.
Local contract: `next_round_20260911/V6_24_LOCAL_SYNC_AND_TRACEABILITY_20260911.md`.

Scientific resources remain those indexed by the V6.23/V6.22 manifests and
their frozen predecessor manifests. Current Fig.1 and ED2 are label-only
successors in `outputs/v6_24_figure1_ed2_labels_v0_1/`, built by
`analysis/plot_v6_24_figure1_ed2_labels_v0_1.py`, relative to the project root.
The four ED2 aggregate tables are byte-identical; historical field names are
interpreted by the new `source_data/publication_labels.tsv`. Fig.1's
conceptual table is synchronized. Fig.2, Fig.3, Fig.4 and ED1/ED3–5 remain
the V6.23 package resources. Both literature workbooks and cover are unchanged.

## What was checked

The current audit records are in `qa/v6_24_traceability/`. They distinguish
participant-prediction recalculation, stored-draw reduction, fitted-object
integrity, public mortality-record linkage and actual ZIP-member inclusion.
The report `TRACEABILITY_REVIEW.md` summarizes the evidence and remaining
limits; detailed machine-readable records retain file and result locators.
The exact editorial delta is `qa/V6_24_EXACT_EDIT_LEDGER.json`.

The software snapshot adds a compact stored-result audit layer: primary
NHANES probabilities, stored BRR replicate estimates, participant contribution
profiles and matrix, synthetic OOF predictions/folds, and small HSCT inputs
and scripts. `V6_24_ADDED_FILES.tsv` in that ZIP is the actual inclusion and
integrity record. Large model binaries, resampling-index arrays and full
replicate-prediction collections remain local. Their existence and hashes
do not make the ZIP a self-contained source-to-model execution bundle.

Historical environment manifests matching eight frozen input references
were recovered from a retained predecessor snapshot; they are supplied in
their original directory, not substituted for the current environment.
One historical P2 direction-lock hash has not been recovered. Its current
file is retained with an explicit mismatch record; no frozen signature is
rewritten. Primary data, models and probabilities matched their recorded
integrity checks. Forty-five NHANES III primary estimators were reloaded and
replayed against stored design matrices; this was not a clean source-to-model
execution or a replay of every fitted object.

## Publication checks and limits

See `qa/V6_24_CONTENT_QA.json`, `qa/V6_24_COLD_READ.md`, the rendered-page
records and the candidate's `PACKAGE_INTEGRITY.json`. Existing Cambria Math
text-run equation representation is retained; this is not an OMML conversion.
All source and historical package originals remain unchanged.

Public deposition/DOI and a clean external full-study reexecution remain
open. Original mortality linkages were checked locally; this does not
constitute an independent rebuild of the entire NHANES III clinical interface.
Literature workbook mappings and targeted original passages were audited,
not every original paper or every Grødem derivation. User-managed author
administration remains outside this revision. These boundaries belong in
the audit and availability documentation, not as new defensive main-text claims.
