## Supplementary_Software_1 for "Inferential boundaries of age prediction: why prediction does not establish biological age measurement": REPRODUCIBILITY_MANIFEST_v6_25_20260911.md

# Reproducibility manifest V6.25

This initial-submission assembly retains all V6.24 scientific estimates, source tables, model contracts, four main figures and five Extended Data figures. Main V6.25 and SI V5.17 add review-access and verification instructions; the Summary gains one existing-reference callout and Methods/cover disclose Codex assistance.

Current scientific module contracts are recorded in EVIDENCE_MODULE_REGISTRY.tsv. The prior detailed resource map is REPRODUCIBILITY_MANIFEST_v6_24_20260911.md, retained unchanged. Historical script paths remain dependencies/provenance and are not competing submission versions.

The supplied executable is reproducibility/verify_stored_results.py. Extract both Supplementary Software 1 archives into one root. It recalculates implemented principal point results for SYN-02, HUM-01, HUM-03, HUM-04, HUM-06, HUM-08 and MTH-01; saved HSCT permutation/bootstrap statistics and NHANES III BRR estimates support the stated uncertainty checks. It preserves the full contribution matrix and all matched pairs. Read its output for exact coverage.

Verification levels are distinct: preservation of file identity, figure Source Data consistency, recalculation from stored predictions/statistics, fitted-object reload parity and raw-source-to-model reexecution. The final isolated extraction test establishes only the executable checks actually run. Binary models and large NHANES bootstrap-index arrays are retained in the author project and are not in these archives. A complete external source-to-model rebuild has not been verified.

The V6.24 traceability review also retains one historical provenance exception: the exact original P2 direction-lock snapshot associated with one frozen manifest could not be recovered. Neither the manifest nor the historical file was altered to conceal the discrepancy. Reported quantitative outputs passed the scoped recalculations. The detailed internal audit is retained by the authors.

Review access is authorized. Public upload, licensing, persistent identifiers and publication-stage release are not completed or authorized by this assembly.
