## Supplementary_Software_1 for "Inferential boundaries of age prediction: why prediction does not establish biological age measurement": REPRODUCIBILITY_MANIFEST_v6_26_20260914.md

# Reproducibility manifest V6.26 / SI V5.18

This bounded delivery repair preserves the title, analytical definitions, scientific estimates, four main figures and five Extended Data figures of V6.25 / SI V5.17. Native Word equations replace the previous flattened equation rendering. Methods now expose the existing M_WA and M_X model definitions, the M_X = M_S3A source alias and the ordered variable dictionary. No model, feature set, participant, split, seed or scientific result was newly selected.

Supplementary Data 1 retains 62 selected records and their Tier/Level/rank codes. Source-author claims, actual reported validation, training targets and the present Article's interpretation are separate fields. Public DOI and published section/figure locators replace private paths. Nine role labels formerly occupying the terminal-claim field have been replaced with scientific claim paraphrases. Mixed, endpoint, longitudinal and non-CA models remain distinct. Source-grounded corrections include the pregnancy cell-adjustment evidence, mortality supervision of PCAge and Systems Age, fMRI/EEG model modalities and the primary clinical PhenoAge score in the cross-population multiomics study. Supplementary Data 2 remains the separate prior-critique table. This documentary review neither estimates prevalence nor independently reproduces the source papers.

## Executable scope

Extract both Supplementary Software 1 ZIPs into the same empty directory. The verification entry is `reproducibility/verify_stored_results.py`; the runtime and invocation are in `reproducibility/README.md`. The first ZIP includes 17 newly packaged frozen files whose original manifest identities were checked, including the three ED4 plotting inputs at their original paths (8,094 bytes total). The second ZIP retains the complete participant-by-feature contribution matrix. Each ZIP is below 30 MB.

The verifier covers the seven existing principal result modules (SYN-02, HUM-01, HUM-03, HUM-04, HUM-06, HUM-08 and MTH-01) and three supporting comparisons. SUP-FLEX, SUP-MATCHED and SUP-SUBSTITUTION are verification labels, not new evidence-registry modules.

| Supporting comparison | Recalculated layer | Interval boundary |
|---|---|---|
| Flexible calibration | Individual probability-derived losses, primary contrast and fold contrasts; 13,724 participants and 470 events | The published bootstrap interval remains a frozen summary. Its index archive is not supplied. |
| Repeated matched scalar | Five fixed repeat/fold assignments, individual losses, within-person mean losses across repeats, overall/repeat/cycle contrasts | Repeats are not independent people. Losses are averaged before the final participant mean; probabilities are not averaged before loss. The frozen bootstrap interval is not recalculated. |
| NHANES III age-decoder substitution | Primary probabilities, losses, weights, participant/design/fold identities and 32 stored Fay-BRR replicate contrasts | The rho = 0.3, t_23 interval is recomputed from saved replicate estimates. Replicate-level fitting and predictions are not regenerated. |

The flexible/matched-scalar frozen bootstrap-index archive is 59,391,693 bytes and remains in the author project. Other large NHANES index arrays and fitted objects also remain local. No replacement random draws, fitting, tuning, raw-data download or source-to-model reconstruction were performed. HSCT uncertainty checks summarize saved permutation/bootstrap statistics; NHANES III intervals summarize saved BRR estimates. These layers do not establish a clean-machine end-to-end rerun.

The final two-ZIP extraction test and executed plot commands are recorded in the submission handoff directory's author-only `internal_qa/`, outside the software ZIPs. The verifier produces the same report types for a reviewer: input identities, runtime, recovered estimates, individual checks and exact limitations. A successful exit means all implemented checks passed against supplied inputs, not that every supplementary analysis or model-generation chain was rerun.

## Figure checks and methods

The ED4 entry reads the original three selected-pair/source tables without choosing new participants. The main-figure assembly entry offers `--validate-supplied-only`. The Fig.1/ED2 wrapper offers `fixed-source` mode to test numerical artists, intervals, labels and source preservation, and `historical-pixel-audit` mode to additionally require historical pixel identity. Redraw outputs are for inspection and do not replace the unchanged publication figures.

`reproducibility/ORDERED_VARIABLE_DICTIONARY.tsv` supplies the frozen feature order, source variable codes, labels, units, transformations and missing-data/derivation notes for the 50 continuous-NHANES and 25 NHANES III variables. `CORE_NHANES_REPRODUCTION.md` retains the original preparation and training routes. The source-information script's filename ends in v0_1 while its accepted output directory ends in v0_2; these are not competing model versions.

## Preserved limits and access

All 13 current scientific module contracts remain unchanged. Earlier detailed maps and frozen manifests remain provenance. One historical P2 direction-lock snapshot still differs from the exact original identity recorded in its frozen manifest; that original was not recovered, and the manifest was not rewritten. P2 denotes the historical analysis stage, not NHANES III Phase 2.

The V6.25 upload manuscript was open in another Word process during repair. Its unsaved state could not be compared. This version was built from the preserved named mother documents without closing or changing that open file.

Materials remain for editorial and reviewer evaluation. Public release, a new reuse licence, DOI creation, submission, author signatures and dynamic XFA/portal acceptance are not established by this repair. The existing three author forms are preserved as drafts with their original completion boundaries.
