## Supplementary_Software_1 for "Inferential boundaries of age prediction: why prediction does not establish biological age measurement": V6_9_EXTENDED_DATA_RESOURCE_INDEX_20260828.md

# V6.9 Extended Data governance resource index

Date: 28 August 2026

| Figure | Resource | Registered responsibility | Boundary |
|---|---|---|---|
| Extended Data Fig. 1 | `outputs/three_variable_age_state_task_map_figure_v0_2/figures/Figure_X123_age_coordinate_and_task_map_v0_2.png` | `SYN-02` cross-sectional age/state task map | one frozen construction and fixed models; bin means are not trajectories and do not prove the theorem |
| Extended Data Fig. 2 | `outputs/v6_2_submission_extended_data_v0_1/extended_data_figure_2/figures/Extended_Data_Figure_2_hsct_referent_sensitivities.png` | `HUM-01` referent sensitivities | reused unchanged |
| Extended Data Fig. 3 | `outputs/nhanes_hum03_hum04_hum06_extended_data_figure3_successor_v0_2/figures/Extended_Data_Figure_3_NHANES_age_substitution_fold_and_persistence_diagnostics.png` | `HUM-03`, `HUM-04` and `HUM-06` diagnostics | reused unchanged; crossing-zero intervals are not equivalence |
| Extended Data Fig. 4 | `outputs/v6_9_extended_data_figure4_domain_options_v0_1/figures/Extended_Data_Figure_4_five_domain_acquisition_comparison.png` | `MTH-01` bounded acquisition illustration | five prespecified measurement-domain options; no equality of variable count, budget, monetary cost or real-world resources |

Editable SVG and vector PDF companions are stored beside each listed PNG.
`HUM-02`, `HUM-05` and `MTH-02` remain audit-only and absent from the
submission-facing chain.
