## Supplementary_Software_1 for "Inferential boundaries of age prediction: why prediction does not establish biological age measurement": V6_9_MAIN_FIGURE_RESOURCE_INDEX_20260828.md

# V6.9 main-figure governance resource index

Date: 28 August 2026

| Figure | Resource | Registered responsibility | Boundary |
|---|---|---|---|
| Figure 1 | `outputs/v6_9_figure1_theory_chain_v0_1/figures/Figure_1_age_decoding_identifies_an_age_task_not_biological_age.png` | `ANA-01`--`ANA-03` analytical chain | wording-only successor; scalarization is stated as information retention, not a universal one-direction claim |
| Figure 2 | `outputs/hsct_age_reconstruction_semantic_counterexample_figure_v0_4/figures/Figure_HSCT_age_reconstruction_semantic_counterexample.png` | `HUM-01` referent ambiguity | geometry-preserving score-pairing null is primary; regression is a coordinate-compatibility sensitivity |
| Figure 3 | `outputs/nhanes_hum03_hum04_hum06_figure3_successor_v0_3/figures/Figure_3_NHANES_endpoint_utility_information_set_and_supervision_target.png` | panel a design; panel b `HUM-04`; panel c `HUM-06`; panel d `HUM-03` | reused unchanged; distinct populations, interfaces and uncertainty procedures are not pooled |
| Figure 4 | `outputs/v6_8_figure4_target_first_framework_v0_1/figures/Figure_4_target_first_estimand_specific_replacement_framework.png` | `FRM-01` target-first replacement | reused unchanged; conceptual estimand map, not a new empirical result |

Editable SVG and vector PDF companions are stored beside each listed PNG.
The v6.9 Figure 1 and v0.4 Figure 2 packages are fit-free presentation
successors. No frozen model or estimand changed.
