## Supplementary_Software_1 for "Inferential boundaries of age prediction: why prediction does not establish biological age measurement": V6_13_EXTENDED_DATA_RESOURCE_INDEX_20260831.md

# V6.13 Extended Data governance resource index

Date: 31 August 2026

| Figure | Resource | Registered responsibility | Boundary |
|---|---|---|---|
| Extended Data Fig. 1 | \`outputs/three_variable_age_state_task_map_figure_v0_2/figures/Figure_X123_age_coordinate_and_task_map_v0_2.png\` | SYN-02 cross-sectional age/state task map | unchanged |
| Extended Data Fig. 2 | \`outputs/v6_2_submission_extended_data_v0_1/extended_data_figure_2/figures/Extended_Data_Figure_2_hsct_referent_sensitivities.png\` | HUM-01 referent sensitivities | unchanged |
| Extended Data Fig. 3 | \`outputs/nhanes_hum03_hum04_hum06_extended_data_figure3_submission_v0_3/figures/Extended_Data_Figure_3_NHANES_age_context_substitution_and_fold_diagnostics.png\` | HUM-03, HUM-04 and HUM-06 diagnostics | unchanged; crossing-zero intervals are not equivalence |
| Extended Data Fig. 4 | \`outputs/nhanes_individual_agegap_contributions_figure_v0_2/figures/Extended_Data_Figure_4_near_identical_AgeGaps_distinct_measured_contributions.png\` | HUM-08 non-unique measured-profile realization | exact fitted-model contributions; no causal, organ-age or biological-age-component interpretation |
| Extended Data Fig. 5 | \`outputs/v6_9_extended_data_figure4_domain_options_v0_1/figures/Extended_Data_Figure_4_five_domain_acquisition_comparison.png\` | MTH-01 bounded acquisition illustration | same frozen resource, renumbered only; five prespecified domains and no resource equivalence |

Editable SVG and vector PDF companions are stored beside each listed PNG. The
submission-facing HUM-08 figure omits endpoint-logit comparisons.
