## Supplementary_Software_1 for "Inferential boundaries of age prediction: why prediction does not establish biological age measurement": V6_13_MAIN_FIGURE_RESOURCE_INDEX_20260831.md

# V6.13 main-figure governance resource index

Date: 31 August 2026

| Figure | Resource | Registered responsibility | Boundary |
|---|---|---|---|
| Figure 1 | \`outputs/v6_9_figure1_theory_chain_v0_1/figures/Figure_1_age_decoding_identifies_an_age_task_not_biological_age.png\` | ANA-01–ANA-03 analytical chain | unchanged |
| Figure 2 | \`outputs/hsct_age_reconstruction_semantic_counterexample_figure_v0_4/figures/Figure_HSCT_age_reconstruction_semantic_counterexample.png\` | HUM-01 referent ambiguity | unchanged; geometry-preserving score-pairing null remains primary |
| Figure 3 | \`outputs/nhanes_hum03_hum04_hum06_figure3_successor_v0_3/figures/Figure_3_NHANES_endpoint_utility_information_set_and_supervision_target.png\` | HUM-03, HUM-04 and HUM-06 | unchanged; distinct populations and uncertainty procedures are not pooled |
| Figure 4 | \`outputs/v6_8_figure4_target_first_framework_v0_1/figures/Figure_4_target_first_estimand_specific_replacement_framework.png\` | FRM-01 target-first replacement | unchanged conceptual estimand map |

Editable SVG and vector PDF companions remain beside each listed PNG. HUM-08
does not alter or enter a main figure.
