## Supplementary_Software_1 for "Inferential boundaries of age prediction: why prediction does not establish biological age measurement": FIGURE_LEGENDS.md

# Figure 3 | Mortality prediction depends on source representation and supervision target

**a**, Continuous NHANES 2007–2012 comparison in participants whose survey cycles supplied at least five years of administrative follow-up. A fold-local ridge decoder reconstructed chronological age from background and the same 50 measurements used by the direct endpoint model. Mortality models used age and background alone, added the age-supervised score, or directly modelled the 50 measurements; all endpoint predictions were out of fold. **b**, Paired held-out five-year all-cause mortality log-loss improvements: the age decoder beyond age and background, direct source modelling beyond the linear score, and direct source modelling beyond one prespecified flexible calibration of that score. Lines show 95% fixed-prediction bootstrap intervals conditional on stored out-of-fold predictions. **c**, NHANES III matched-scalar workflow. The same 25 measurements and background were inputs to two Phase 1 ridge projections trained on standardized chronological age or five-year mortality. Both one-dimensional linear projections were frozen and applied unchanged to Phase 2. Phase 2 endpoint models retained chronological age and background and fitted the same fold-local spline calibration and regularization procedures to each score. **d**, Paired design-weighted held-out contrasts in Phase 2: age-score utility beyond age and background, and mortality-score advantage over the age score. Lines show 95% Fay-BRR intervals from Phase 2 downstream refits, conditional on the frozen Phase 1 projections. NHANES III was a retrospective, outcome-accessed protocol transfer, not untouched confirmation. Positive values favour the added or substituted representation named in each row. The two populations, measurement panels and uncertainty procedures are distinct; these comparisons establish performance in the declared designs, not a general ordering of finite-sample estimators.

# Figure 4 | A target-first programme for measurement and evidence

**a**, The biological question defines the object and estimand, the measurement and study design, the fitting and validation target, and the evidence-supported claim. **b**, An exploratory risk-prediction example. In NHANES 2007–2008, two policies selected one domain from the same five prespecified measurement-domain options. Mortality-directed selection maximized held-out five-year all-cause mortality log-loss gain and selected hepatic, protein and injury measurements; inverse age-decoder selection minimized held-out chronological-age mean-squared error and selected metabolic, adiposity and lipid measurements. The frozen selections were evaluated in NHANES 2009–2010. Points show each policy's held-out mortality log-loss gain beyond age and background and their paired difference; lines show the original 95% intervals from 5,000 event-stratified participant bootstraps conditional on fixed source selections and stored predictions. Domains differed in measurement count, and no equivalence of acquisition costs was assumed. Full selection and fold diagnostics remain in Extended Data Fig. 5. **c**, Estimand-specific replacement map: measured state, named risk, transition, intervention displacement and benefit require their respective measurement, endpoint, longitudinal or causal evidence. The risk-acquisition example illustrates one route through this programme; it does not validate the other four objects. Age-equivalent units may re-express a separately identified target but do not supply its identity or causal meaning.
