## Supplementary_Software_1 for "Inferential boundaries of age prediction: why prediction does not establish biological age measurement": README.md

# V6.22 main-figure assembly v0.1

Authorized static writing/assembly update, 10 September 2026. This package
changes figure presentation, not frozen analyses, estimands, estimates or
uncertainty calculations. It does not read participant data, load fitted models,
fit or tune estimators, resample observations or generate random numbers.

## Chart contract

- Surface: static 180-mm manuscript figures; editable SVG, vector PDF and
  600-dpi PNG. Renderer: Matplotlib in the pinned project environment.
- Figure 3 question: how do source representation and supervision target affect
  held-out five-year mortality prediction? Two rows distinguish the continuous
  NHANES comparison (50 measurements) from the NHANES III matched-scalar
  comparison (25 measurements). Each row pairs a design diagram with its frozen
  dot-and-interval contrasts. Three continuous-NHANES contrasts and two NHANES
  III contrasts are the complete specified comparisons, not a sparse trend.
- Figure 3 retains actual age plus background in each endpoint comparison;
  distinguishes age-only decoder training from mortality modelling; and
  distinguishes frozen Phase 1 projections from Phase 2 fold-local fitted
  calibration. No direct Cox or population-information estimate is implied.
- Figure 4 question: how can a declared biological target guide measurement,
  model selection and the claim? A four-stage workflow, a compact acquisition
  example, and a five-object evidence map connect the practical programme.
  The example displays both frozen policy gains and their paired difference,
  all with their original intervals. It is an exploratory risk example, not
  validation of all five objects or equivalence of acquisition costs.
- Styling: white background; Arial; blue/teal for Figure 3 and blue/orange for
  Figure 4, with dark neutral text and reference lines. Marker shape, fill,
  direct row labels and order preserve meaning without colour. No branding.
- Full observed-age information-set evidence stays in existing Extended Data
  Figure 3. Full acquisition selection and fold diagnostics stay in existing
  Extended Data Figure 5. Those files are not modified.

## Rebuild

From `E:/Papers/Aging`:

```powershell
& .\LifeHistoryStateDecomposition\scripts\Test-ProjectEnvironment.ps1
& .\LifeHistoryStateDecomposition\scripts\Invoke-ProjectPython.ps1 -- .\LifeHistoryStateDecomposition\analysis\plot_v6_22_main_figure_assembly_v0_1.py
```

The builder reads the seven frozen aggregate source-table copies identified in
`source_data/source_data_index.tsv`, preserving all columns and exact bytes.
When predecessor files are available, copies must match them. Without those
directories, the supplied copies are checked against the frozen source index.
Existing copies are reused and mismatches fail closed.
`BUILD_REPORT.json` records source-copy parity and static-render checks;
`VISUAL_QA.md` records the separate manual export inspection. Figure legends
are supplied in `FIGURE_LEGENDS.md` for the main writer to integrate.

The source tables and builder are sufficient for this static figure assembly;
this is not an end-to-end reproduction of the underlying scientific analyses.
