## Supplementary_Software_1 for "Inferential boundaries of age prediction: why prediction does not establish biological age measurement": VISUAL_QA.md

# Static figure export QA — 10 September 2026

Status: PASS for the exported PNG figures inspected by the figure-assembly
agent. Final placement in Word/PDF is a separate manuscript-build check.

- Both final 600-dpi PNG exports were opened and visually inspected after the
  final axis-label update. Titles, panel letters, axes, row labels and footnotes
  are legible; no clipped boxes, title/footnote collisions or interval clipping
  remain. The first draft's clipped box edges and Fig. 3 inter-row collision
  were repaired before this pass.
- Figure 3 keeps the continuous 50-measurement comparison in a–b and the
  NHANES III 25-measurement comparison in c–d. Age/mortality are Phase 1 ridge
  targets rather than extra inputs. The Phase 1 directions are frozen; Phase 2
  calibration is explicitly fold-local and fitted. Both endpoint comparisons
  retain actual age and background.
- All five Figure 3 point estimates and full confidence intervals match their
  frozen source rows. Identical axes permit within-figure scale reading without
  implying identical samples, panels or uncertainty procedures. Fixed-prediction
  bootstrap and conditional Phase 2 Fay-BRR uncertainty are labelled separately.
- Figure 4 shows both policy gains and their paired contrast, including the
  inverse-age policy's interval across zero. All three intervals are complete.
  Source criteria are shown as distinct objectives, not compared on one scale.
  The named-risk example is separated from the five-object evidence map.
- Shape, open/filled markers, direct labels and fixed row ordering supplement
  colour. The figures have no decorative branding.
- Automated box-containment checks passed for 9 labels in Figure 3 and 7 in
  Figure 4. All 7 source tables were byte-exact and DataFrame-exact copies of
  their frozen predecessors. Supplied-copy-only validation separately passed
  without reading predecessor directories.
- SVG and PDF exports were produced from the same checked figure objects;
  editable SVG text and PDF TrueType embedding are configured. This record does
  not claim separate raster inspection of the vector PDFs or an independent
  end-to-end scientific reproduction.

No participants, fitted objects or bootstrap draws were read. No estimate,
confidence interval, model, threshold or outcome was recomputed or selected.
Only new versioned figure-assembly files were written; frozen predecessors,
main/SI text, registry and existing Extended Data files were not modified.
