## Supplementary_Software_1 for "Inferential boundaries of age prediction: why prediction does not establish biological age measurement": Figure_4_target_first_measurement_and_evidence.pdf

**a** Start with the biological question

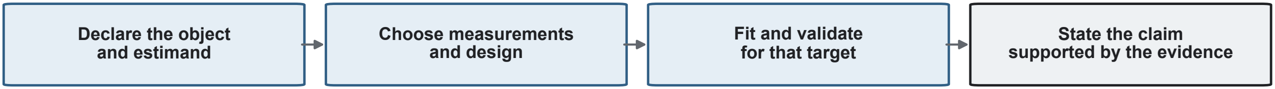


**b** A risk-targeted measurement-acquisition example

Source policies: NHANES 2007–2008 → target evaluation: 2009–2010

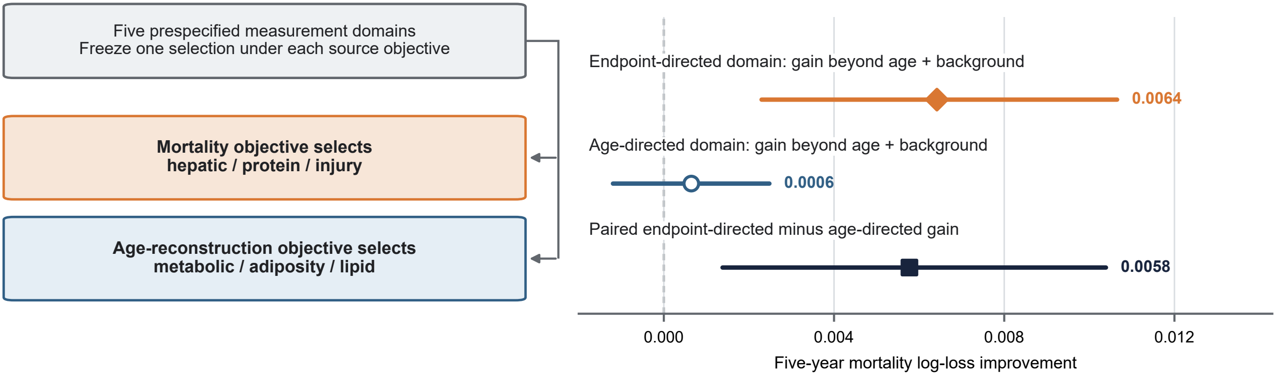


95% fixed-prediction bootstrap intervals · 4,896 target participants · 157 deaths

Exploratory comparison; domains differ in measurement count and no cost equivalence is assumed

**c** Different objects require different evidence

| Object | Declared target | Required structure and evidence |
|---|---|---|
| **Measured state** | State under a declared measurement model | Anchors, reliability and invariance; same-age state validation |
| **Named risk** | Endpoint probability over a specified horizon | Endpoint, horizon, loss and population; held-out performance and calibration |
| **Transition** | Within-person change over a specified interval | Repeated measurements, time scale and a longitudinal model |
| **Intervention displacement** | Intervention effect on a prespecified measure | Causal design and contrast; measurement invariance and follow-up |
| **Benefit** | Causal effect on a named outcome | Treatment strategies, causal assumptions and the decision population |
