## Supplementary_Software_1 for "Inferential boundaries of age prediction: why prediction does not establish biological age measurement": README.md

# Figure 4b shared-candidate connector correction (v6.23)

This static, fit-free correction starts from the frozen v6.22 Figure 4. Its only
visual change replaces the single source-to-mortality arrow with a shared branch
from the five prespecified measurement domains to both the mortality-objective
and age-reconstruction-objective selection boxes. Box positions, labels, colors,
all estimates and intervals, and panels 4a and 4c are preserved.

## Outputs and verification

`figures/Figure_4_target_first_measurement_and_evidence` is exported as 600 dpi
PNG, editable-text SVG, and vector PDF. `BUILD_REPORT.json` records the exact
pixel comparison with the frozen v6.22 PNG, unchanged text and source-table bytes,
box-label containment, and the only permitted connector extents. See
`VISUAL_QA.md` for visual review of the exported PNG and rendered PDF.

No Figure 3 is rendered. No participant data, model fitting/loading, random
numbers, or new estimates are involved. Frozen v6.22 artifacts are read only.

## Reproduction and software-package dependencies

From the Aging workspace root, run the project environment gate and then:

```powershell
& .\LifeHistoryStateDecomposition\scripts\Invoke-ProjectPython.ps1 -- .\LifeHistoryStateDecomposition\analysis\plot_v6_23_figure4_connector_v0_1.py
```

The software ZIP must retain these paths relative to `LifeHistoryStateDecomposition`:

- `analysis/plot_v6_23_figure4_connector_v0_1.py` (this bounded correction).
- `analysis/plot_v6_22_main_figure_assembly_v0_1.py` (imported Figure 4 renderer).
- `outputs/v6_23_figure4_connector_v0_1/README.md` (this contract).
- `outputs/v6_22_main_figure_assembly_v0_1/source_data/source_data_index.tsv`.
- The four `.tsv` files under `outputs/v6_22_main_figure_assembly_v0_1/source_data/Figure_4/`.
- `outputs/v6_22_main_figure_assembly_v0_1/figures/Figure_4_target_first_measurement_and_evidence.png` (frozen pixel-comparison reference).
- The pinned project environment contract, manifests, and runtime wrappers.

The four original aggregate TSVs are reused at their frozen paths; no new data
copies are created. The v6.22 module's main routine and input-copy routines are
not called, so its Figure 3 data and older predecessor source locations are not
runtime dependencies of this correction.
