## Supplementary_Software_1 for "Inferential boundaries of age prediction: why prediction does not establish biological age measurement": VISUAL_QA.md

# Figure 4 connector QA

Status: PASS, reviewed 2026-09-11.

- Inspected the frozen v6.22 PNG, the new 600 dpi PNG, and the new one-page PDF
  rendered with Poppler at 180 dpi. Both objective boxes now receive an arrow
  from the common candidate-domain branch. The branch stays within the gap
  between the boxes and the quantitative panel, with no text overlap or clipping.
- Before the connector edit, the imported renderer reproduced the frozen v6.22
  PNG pixel for pixel. After the edit, the image remains 4010 x 3589 pixels.
  Exactly 8,584 pixels changed, all within the removed-arrow and new-branch
  extents; zero pixels changed elsewhere. Thus all unchanged boxes, labels,
  numerical marks, intervals, colors, and panels 4a/4c retain the frozen image.
- All 7 boxed labels pass geometric containment. All 63 SVG text nodes retain
  identical content, position, and styling compared with v6.22.
- The four original aggregate TSVs match their frozen v6.22 manifest checksums
  and remain byte-identical after rendering. They are reused in place; no
  participant data, Figure 3 data, models, or new estimates are involved.
- PDF inspection reports one page, 481.270 x 430.907 pt. PNG metadata reports
  599.9988 dpi in each dimension (the PNG resolution-field representation of
  the requested 600 dpi). SVG retains editable text; PDF uses vector output.

Machine-readable results and exact data dependencies are in `BUILD_REPORT.json`;
reproduction and software-ZIP requirements are in `README.md`.
