## Supplementary_Software_1 for "Inferential boundaries of age prediction: why prediction does not establish biological age measurement": Extended_Data_Figure_2_hsct_referent_sensitivities.pdf

### Extended Data Figure 2 | HSCT referent-null sensitivities preserve the interpretation boundary

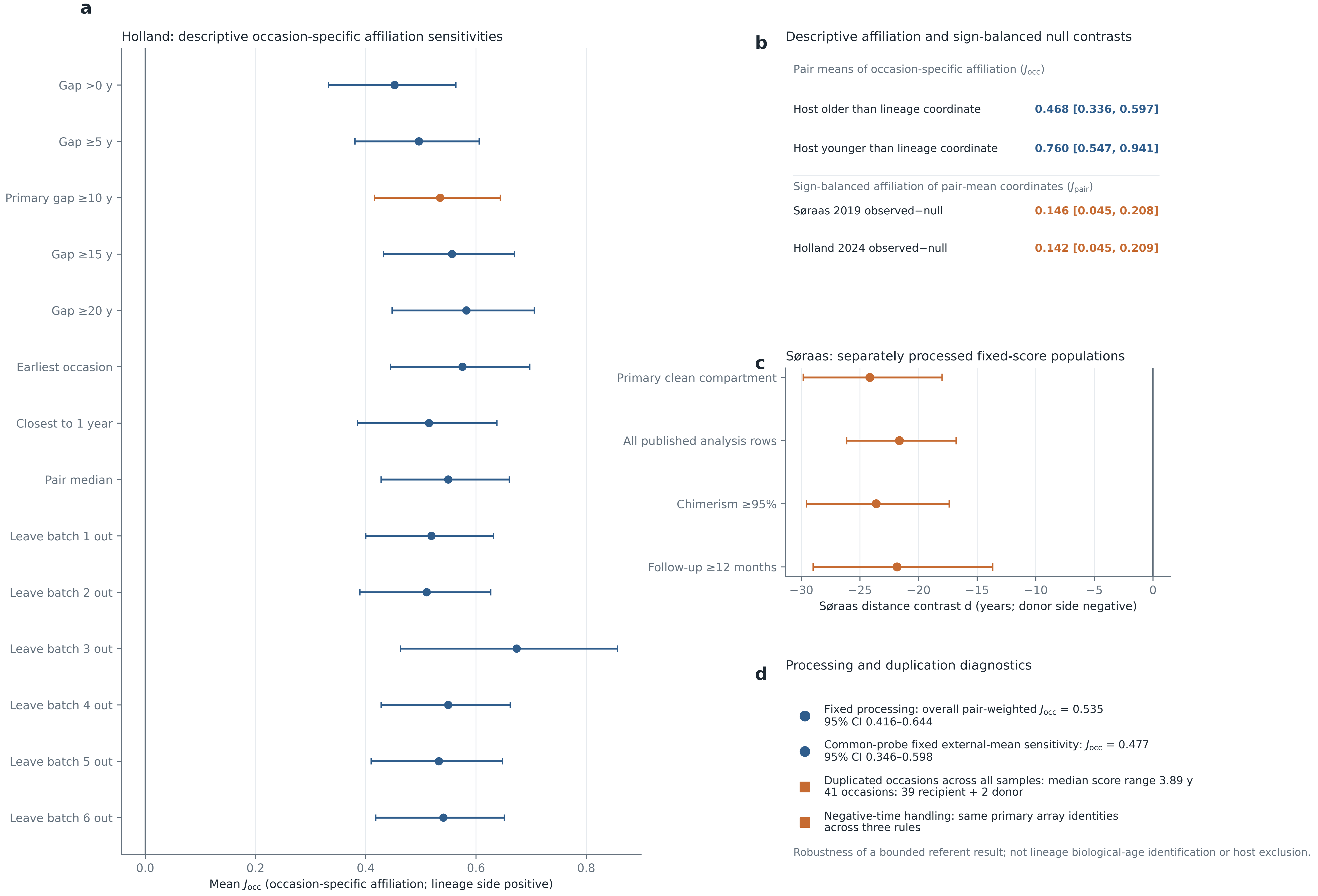
