## Supplementary_Software_1 for "Inferential boundaries of age prediction: why prediction does not establish biological age measurement": README.md

# Figure 1 and Extended Data Figure 2 label synchronization, v6.24

This fit-free successor changes only the author-approved labels and one Figure
1d link style. Figure 1a specifies chronological-age labels alone; Figure 1c
states the fixed-decoder/fixed-centring, observed-age/background condition and
the construct-identity boundary; Figure 1d labels its dashed link as predictive
association. No other Figure 1 content changes.

ED2 distinguishes the older descriptive occasion-specific affiliation index
`J_occ` from the sign-balanced affiliation of pair-mean coordinates `J_pair`.
Panel a averages the declared pair-level summaries: the ordinary rows summarize
occasion-specific affiliation within pairs, while the explicitly named Pair
median sensitivity retains its within-pair median rule. Panel b's upper strata
use pair means of occasion-specific affiliation; its lower observed-minus-null
rows use sign-balanced `J_pair`. Panel d reports overall pair-weighted `J_occ`
under fixed processing and the same-statistic common-probe/fixed-external-mean
sensitivity. Its duplication scope covers all 41 duplicated occasions: 39
recipient and 2 donor. These are label clarifications, not new estimates.

All four ED2 aggregate tables are copied byte for byte. Their historic field
names are retained for reproducibility; `source_data/publication_labels.tsv`
provides the explicit current publication labels. The conceptual Figure 1
table is versioned to match the figure. `SOURCE_REPLACEMENT_MAP.tsv` identifies
the exact replacement/addition route for the submission Source Data package.

## Reproduction and software ZIP dependencies

From the Aging workspace root, pass `Test-ProjectEnvironment.ps1`, then run:

```powershell
& .\LifeHistoryStateDecomposition\scripts\Invoke-ProjectPython.ps1 -- .\LifeHistoryStateDecomposition\analysis\plot_v6_24_figure1_ed2_labels_v0_1.py
```

Retain these paths relative to `LifeHistoryStateDecomposition` in the software ZIP:

- `analysis/plot_v6_24_figure1_ed2_labels_v0_1.py`.
- `analysis/plot_v6_9_figure1_precision_revision_v0_1.py`.
- `analysis/plot_v6_2_submission_extended_data_v0_1.py`.
- `outputs/v6_9_figure1_theory_chain_v0_1/source_data/figure1_theory_chain.tsv`.
- The four original `panel_*.tsv` files in
  `outputs/v6_2_submission_extended_data_v0_1/extended_data_figure_2/source_data/`.
- Frozen reference images
  `manuscript/nature_first/submission_20260911_v6_23_candidate/upload/Main_Figures/Figure_1.png`
  and `manuscript/nature_first/submission_20260911_v6_23_candidate/upload/Extended_Data_Figures/Extended_Data_Figure_2.png`.
- The project environment contract, pinned manifests and runtime wrappers.

Only the named Figure 1 and ED2 builder functions are invoked. Figure 1's
filesystem setup and both builders' export/source-write hooks are intercepted;
the ED2 reads are routed to the four frozen aggregate copies. Neither original
module's `main` routine or any other figure builder runs. The original ED2
predecessor source directory is therefore not a dependency of this entry point.

Exports are 600 dpi PNG, editable-text SVG and vector PDF. `BUILD_REPORT.json`
records exact baseline-pixel reproduction, unchanged ED2 numerical artist state
and aggregate byte parity. `VISUAL_QA.md` records the exported-PNG/PDF inspection.
