## Supplementary_Software_1 for "Inferential boundaries of age prediction: why prediction does not establish biological age measurement": VISUAL_QA.md

# Figure 1 and ED2 visual QA

Status: PASS. Reviewed 2026-09-11.

- Both original renderer entry points independently reproduced their v6.23
  submission PNG pixel for pixel before the edits: identical image dimensions,
  zero changed pixels for both Figure 1 and ED2.
- Inspected both new 600 dpi PNGs and both PDFs rendered by Poppler at normal
  reading resolution (Figure 1: 150 dpi; ED2: 120 dpi). The added labels are
  legible, with no clipping, text collision or ambiguous attachment. The ED2b
  lower text rows were moved together slightly to give their estimand heading
  clear space; neither their values nor any numerical plot position changed.
- Figure 1c's longer title wraps at its original font size. The conditioning
  statement sits immediately below the scalar chain. The Figure 1d endpoint
  link is dashed and explicitly labelled predictive association. Its nodes,
  equations, other panels and scientific content remain in place.
- ED2a retains every threshold, occasion-summary and batch-leave-out row. ED2c
  is unchanged. ED2 numerical marks, error intervals, axes, limits and axes
  positions pass exact artist-state parity before/after label edits. All four
  aggregate TSVs remain byte-identical to the frozen sources.
- ED2b values and intervals remain 0.468 [0.336, 0.597], 0.760 [0.547, 0.941],
  0.146 [0.045, 0.208], and 0.142 [0.045, 0.209]. ED2d retains 0.535 with
  95% CI 0.416-0.644, 0.477 with 95% CI 0.346-0.598, and the 3.89-year
  duplicated-occasion median score range. The authorized scope clarification
  specifies 41 occasions across all samples: 39 recipient and 2 donor.
- Historical source column names and diagnostic names are retained in the
  byte-exact tables. The versioned publication-label map explicitly maps them
  to J_occ versus J_pair, fixed processing versus sensitivity, and the complete
  duplicated-occasion scope. No new estimate or interval was calculated.
- Only Figure 1 and ED2 were rendered. No participant record, fitted model,
  resampling procedure, random stream or other plot was accessed.

Machine checks: `BUILD_REPORT.json`. Source Data replacement/addition routing:
`SOURCE_REPLACEMENT_MAP.tsv`. Exact software dependencies: `README.md`.
