## Supplementary_Software_1 for "Inferential boundaries of age prediction: why prediction does not establish biological age measurement": README.md

# Supplementary Software 1 — frozen V6.27 analysis bundle

Review navigation updated on 22 September 2026; original analysis versions are unchanged.

For editorial and reviewer evaluation. No new public reuse licence is assigned.
Third-party source terms continue to apply. Extract both software ZIPs into the
same empty directory. The second ZIP supplies the full contribution matrix.

## Review texts and historical snapshots

The authoritative review texts are the separately supplied
`Manuscript_Zotero_20260921.docx` and `Supplementary_Information.pdf`.
They accompany this submission and are not stored in this software archive.
See [the review-text selection](CURRENT_MANUSCRIPT_VERSION.md), including the
saved review-file identities for this update.

The Markdown documents under `manuscript/nature_first/`, including the
14 September 2026 V6.27 / SI V5.19 files, are unchanged historical analysis-era
snapshots. Their author placeholders and earlier text do not describe the
current review manuscript. Consult [the snapshot index](manuscript/nature_first/README.md)
for their roles.

## Stored-result verification

Use the declared project environment (environment.yml and win-64 lock); on the
author Windows installation run through scripts/Invoke-ProjectPython.ps1. For a
separately prepared compatible environment, the portable entry is:

    python reproducibility/verify_stored_results.py --bundle-root . --report-dir ../verification

Seven main result modules plus three supporting checks are implemented. The
supporting checks recalculate flexible-calibration and repeated matched-scalar
contrasts from saved individual probabilities, averaging losses within person
across repeats. Their bootstrap intervals remain reported summaries: the frozen
59,391,693-byte index archive is retained locally and is not supplied. No new
bootstrap is generated. Substitution checks use primary participant probabilities
and 32 saved Fay-BRR replicate estimates; replicate predictions/refits are not
provided. Reports name the actual interval layer separately for each module.
These are arithmetic checks, not full source-to-model reconstruction.

## Fixed-input figure entry points

    python analysis/plot_nhanes_individual_agegap_contributions_extended_data_v0_2.py
    python analysis/plot_v6_22_main_figure_assembly_v0_1.py --validate-supplied-only
    python reproducibility/plot_submission_figure1_ed2.py --mode fixed-source --output-dir ../figure_redraw

ED4 reads the three original-path frozen small tables now provided; it does not
choose new display participants. Fig.1/ED2 fixed-source mode tests numerical
artists, intervals, labels and source preservation. Its output is a redraw for
inspection, not a replacement of the supplied publication images. To require
the original environment's historical pixel identity, use --mode
historical-pixel-audit instead. A pixel failure is reported as such, without
changing the supplied numerical evidence or bypassing numerical checks.

## Methods and provenance

CORE_NHANES_REPRODUCTION.md describes the original preparation/model entries.
The ordered 50/25-variable dictionary is reproducibility/ORDERED_VARIABLE_DICTIONARY.tsv.
V6_26_FROZEN_INPUTS.tsv records newly packaged input identities checked against
their unmodified frozen manifests in provenance/v6_26_input_manifests/.
The historical P2 direction-lock identity exception remains unresolved. P2 here
is an analysis stage, not NHANES III Phase 2. Original production scripts and
manifests remain provenance; model refitting and public-source reconstruction
have not been executed by this delivery repair.

The preserved V6.25 README is provenance/README_v6_25.md. This repair does not add
models, tuning, seeds, participant selection, raw-data downloads or a public release.


## Dependency closure, 14 September 2026

Ten previously omitted frozen dependencies are now supplied at their original
paths. See reproducibility/DEPENDENCY_CLOSURE_20260914.md for exact scope,
identity checks and the read-only preflight. This does not certify training
or public-source-to-model reconstruction.

The inventory for this navigation update is
[reproducibility/ARCHIVE_MEMBERS_20260922.tsv](reproducibility/ARCHIVE_MEMBERS_20260922.tsv).
It lists every member of this software archive except itself.
`reproducibility/V6_27_ARCHIVE_MEMBERS.tsv` and earlier inventories are retained
unchanged as historical snapshots, not as inventories of the revised navigation.
Frozen scientific inputs, production scripts, manuscript snapshot text and all
historical manifest payloads are unchanged. This update adds no analysis,
refitting, new scientific manuscript version or public reuse licence.
