## Supplementary_Software_1 for "Inferential boundaries of age prediction: why prediction does not establish biological age measurement": REPRODUCIBILITY_MANIFEST_SUBMISSION_20260831.md

# Reproducibility manifest — submission snapshot 20260831

Scientific pair:

- `MANUSCRIPT_NATURE_IDENTIFICATION_FIRST_v6_13_20260831.md`
- `SUPPLEMENTARY_INFORMATION_IDENTIFICATION_FIRST_v5_5_20260831.md`

This submission snapshot inherits the validated v6.12/v5.4 software archive and adds HUM-08 without refitting an age decoder or endpoint model. HUM-08 uses one frozen continuous-NHANES cross-fitted age-decoder pipeline, fold-local conditional centring and outcome-blind matching to show that near-identical conditionally centred gaps can arise from dissimilar measurement-contribution profiles.

The submission contains four main figures and five Extended Data figures. Extended Data Figure 4 is the HUM-08 contribution-profile display. The frozen acquisition resource previously assigned Extended Data Figure 4 is renumbered only as Extended Data Figure 5.

Included HUM-08 materials comprise the frozen specification, analysis and independent-audit scripts, fit-free plot script, aggregate result summaries, validation reports, artifact manifests, module contract, evidence registry, v6.13 resource indices and governance records. Participant-level derived tables, full contribution matrices, selected-person tables, fold coefficient tables and fitted-model binaries are excluded from the submission archive. Their internal persistence and reload-parity records are indexed by the project evidence manifest.

The public-use NHANES inputs remain available from the National Center for Health Statistics under their source terms. All Python workflows use the pinned `lhstate` environment and the project wrappers included here.
