## Supplementary_Software_1 for "Inferential boundaries of age prediction: why prediction does not establish biological age measurement": REVIEW_ACCESS_NOTICE.md

# Editorial and peer review access

The authors supply the accompanying code and derivative data for confidential editorial evaluation and peer review of “Chronological-age decoding does not identify biological age”. Editors and reviewers may inspect the files and run the supplied verification procedures for that purpose.

No public reuse licence is granted by this submission package. Public deposition, persistent identifiers and reuse licences remain to be finalized before publication. The package has not been publicly uploaded.

Third-party data, software and quoted source material retain their original terms and attribution requirements. This notice does not restrict rights already provided by those sources, and does not grant rights that the authors do not hold. Original NHANES and transplantation data should be obtained from the source repositories listed in the manuscript and accession inventories.

The stored-result verification layer is not a new analysis or an assurance that every source-to-model workflow has been reproduced on a clean machine. Its executable report states exactly which calculations were checked and which inputs were supplied.
