## Supplementary figures and images for "Inferential boundaries of age prediction: why prediction does not establish biological age measurement"

### Extended_Data_Figure_2.png

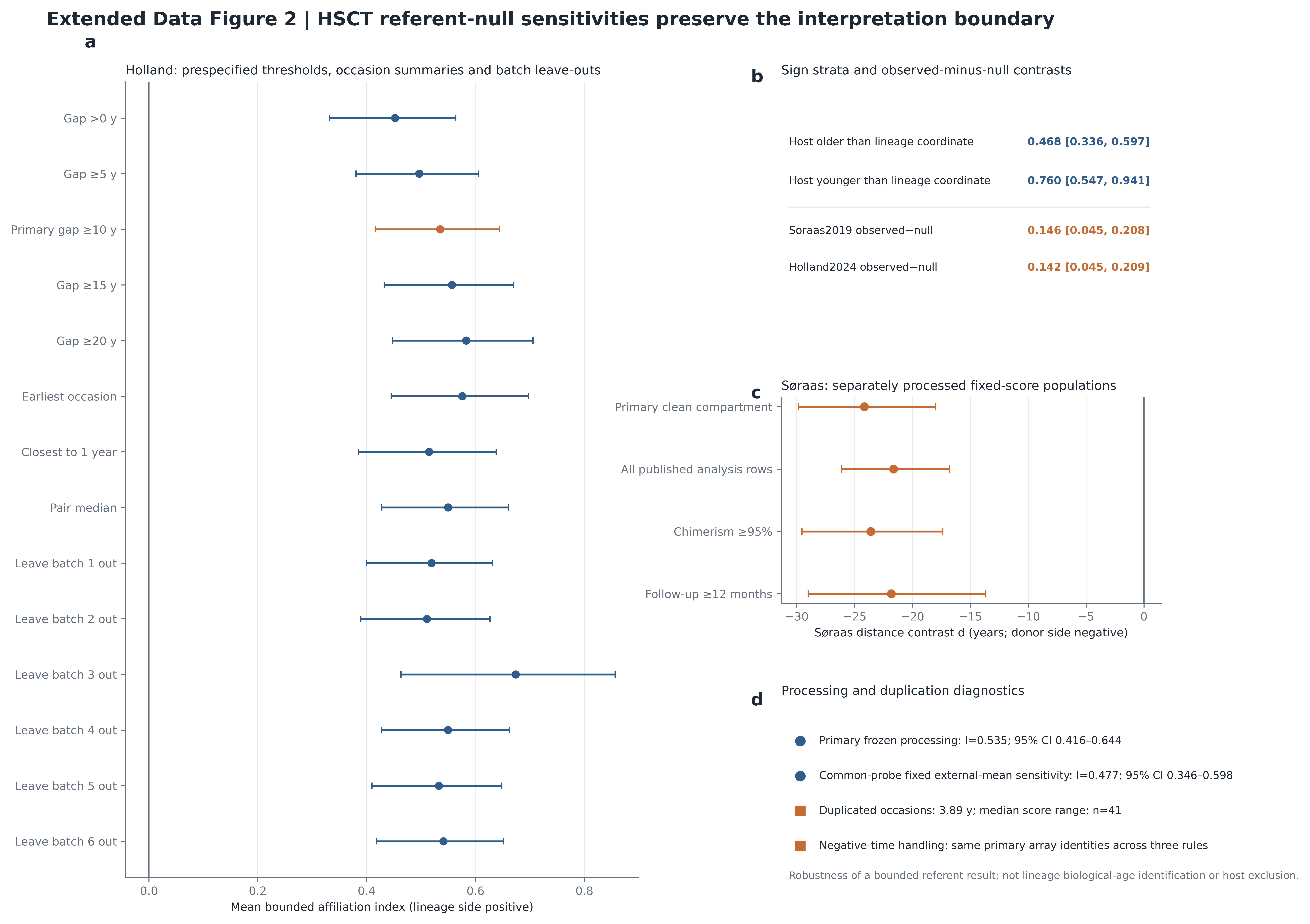

### Extended_Data_Figure_2_hsct_referent_sensitivities.png

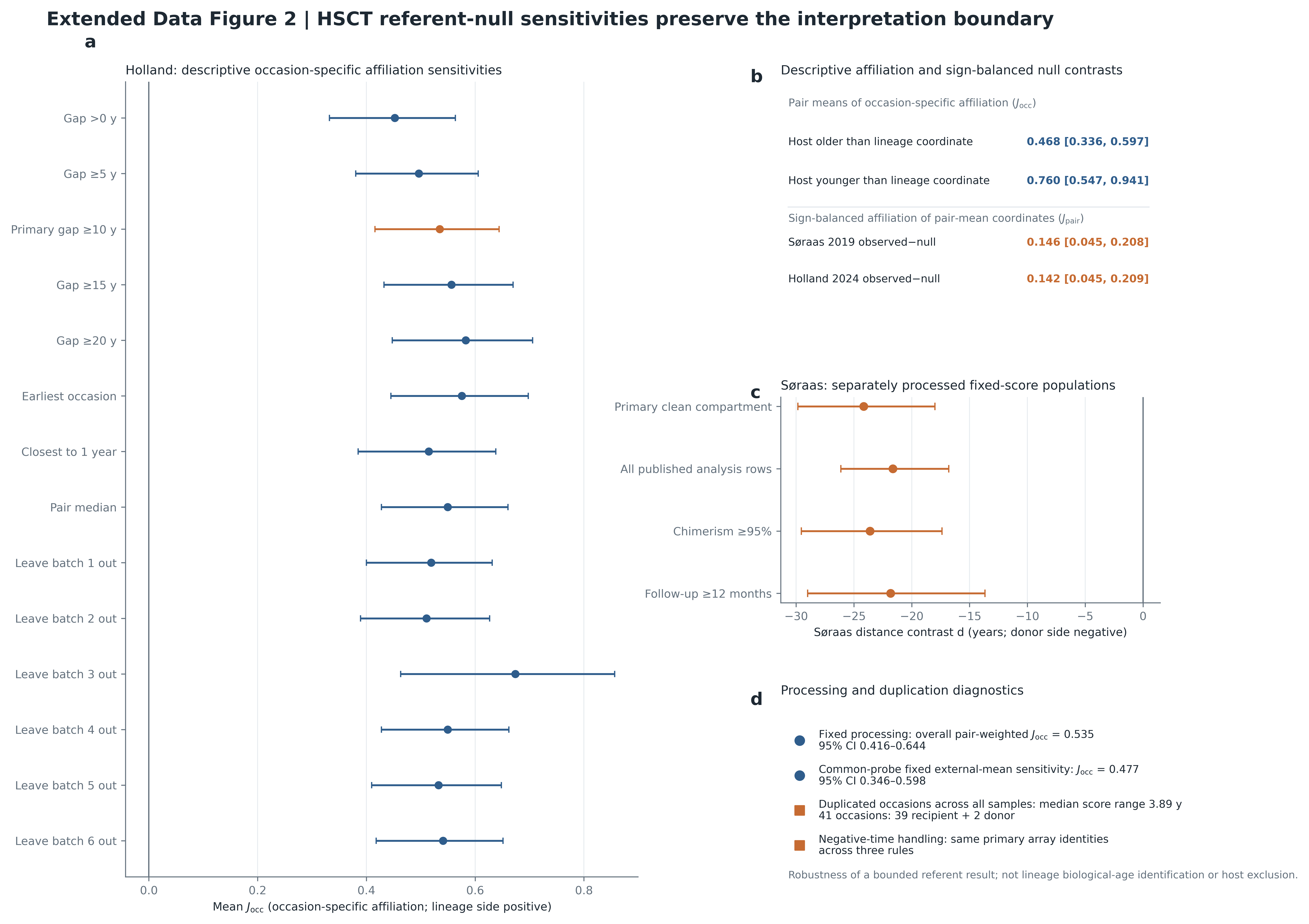

### Figure_1.png

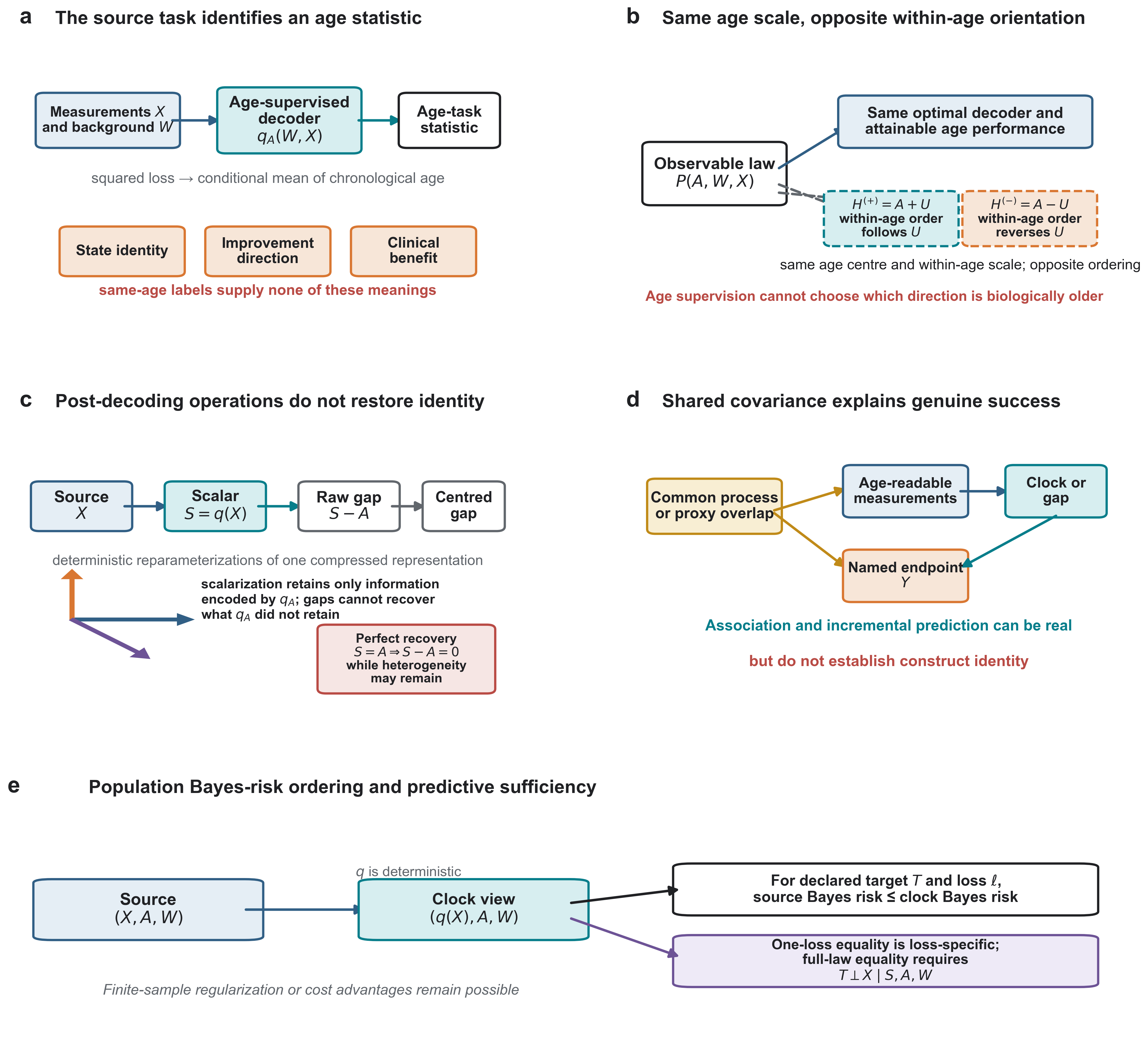

### Figure_1_age_decoding_identifies_an_age_task_not_biological_age.pdf

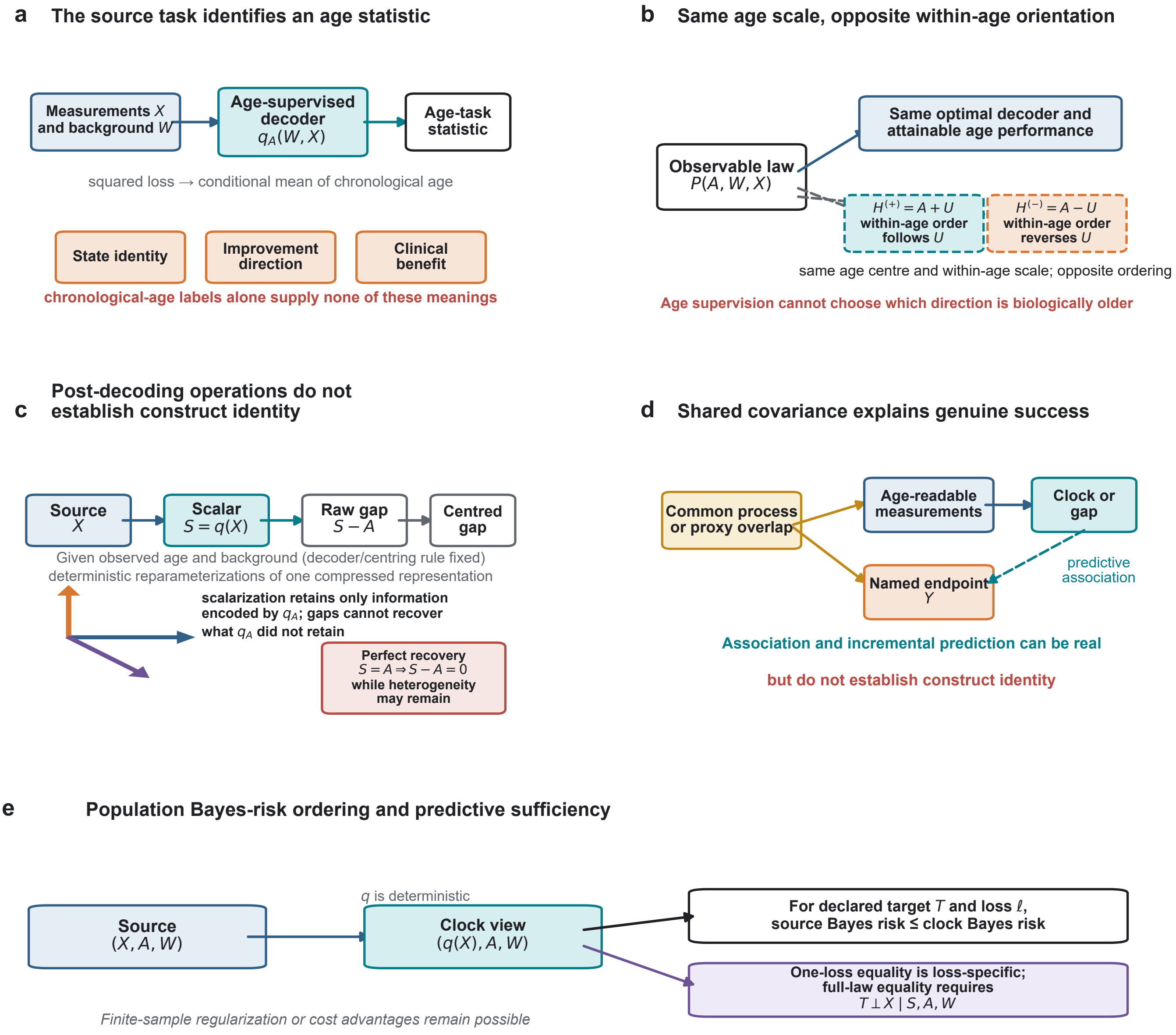

### Figure_1_age_decoding_identifies_an_age_task_not_biological_age.png

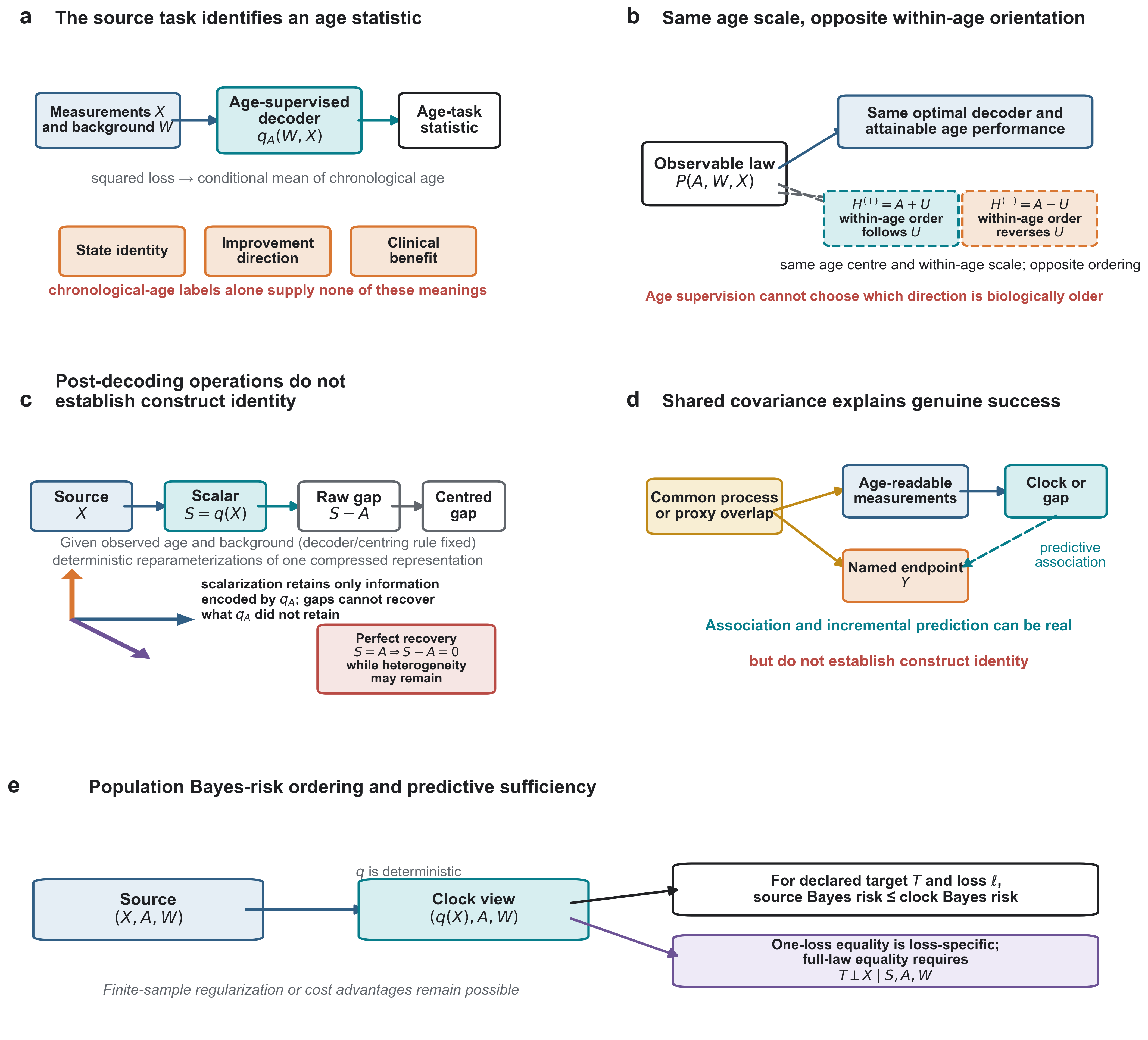

### Figure_3_NHANES_source_representation_and_supervision_target.pdf

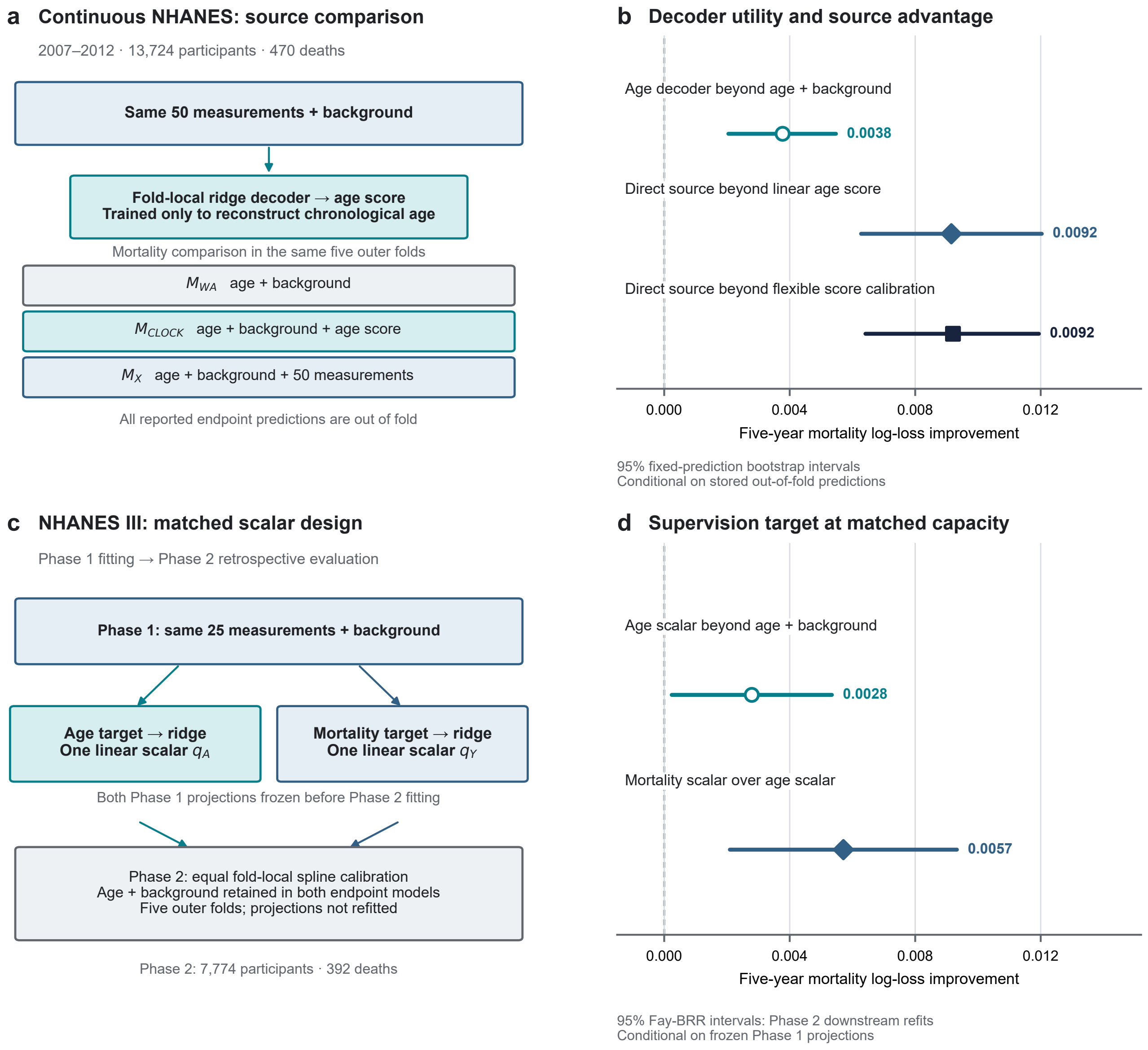

### Figure_3_NHANES_source_representation_and_supervision_target.png

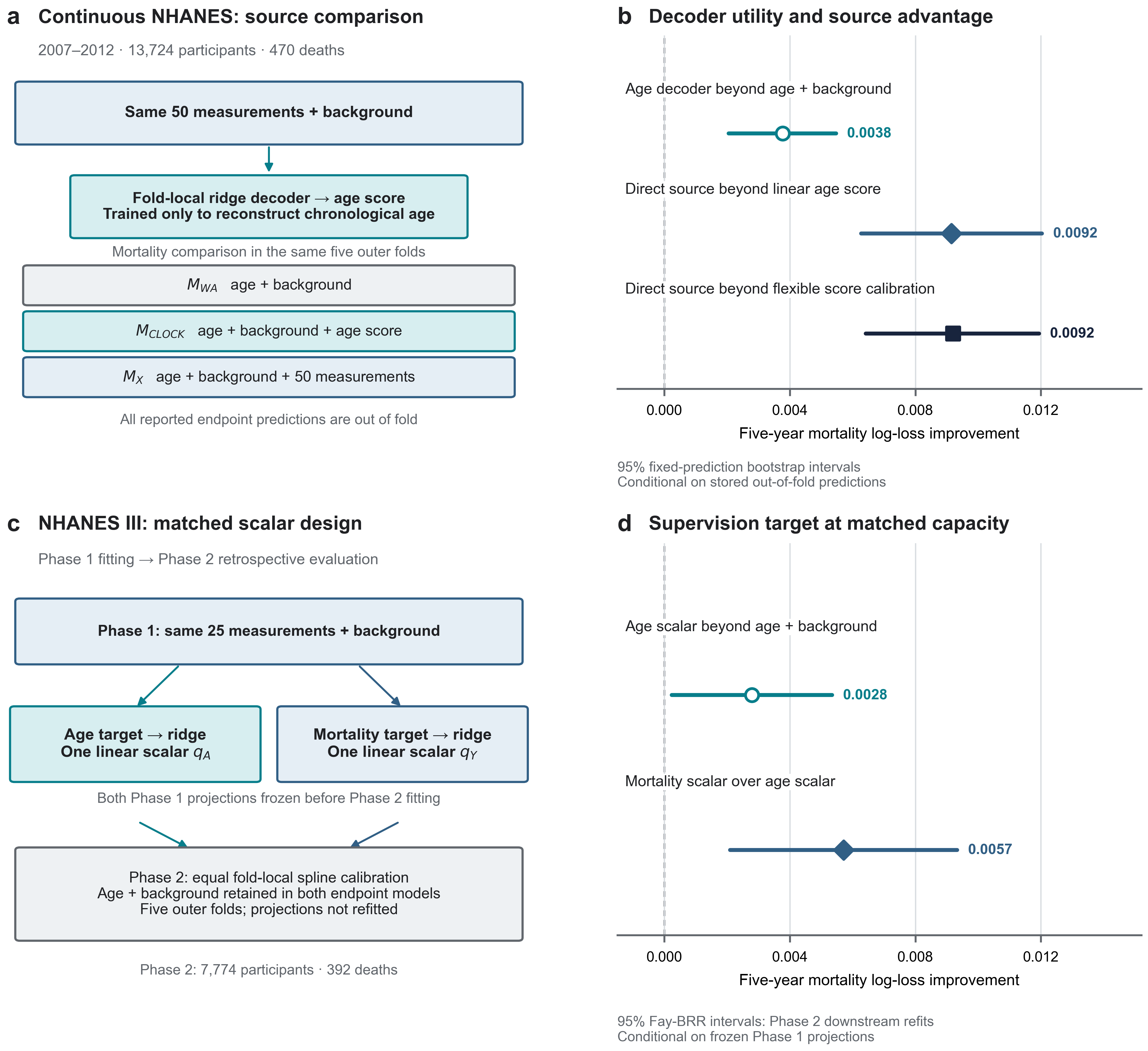

### Figure_4_target_first_measurement_and_evidence.png

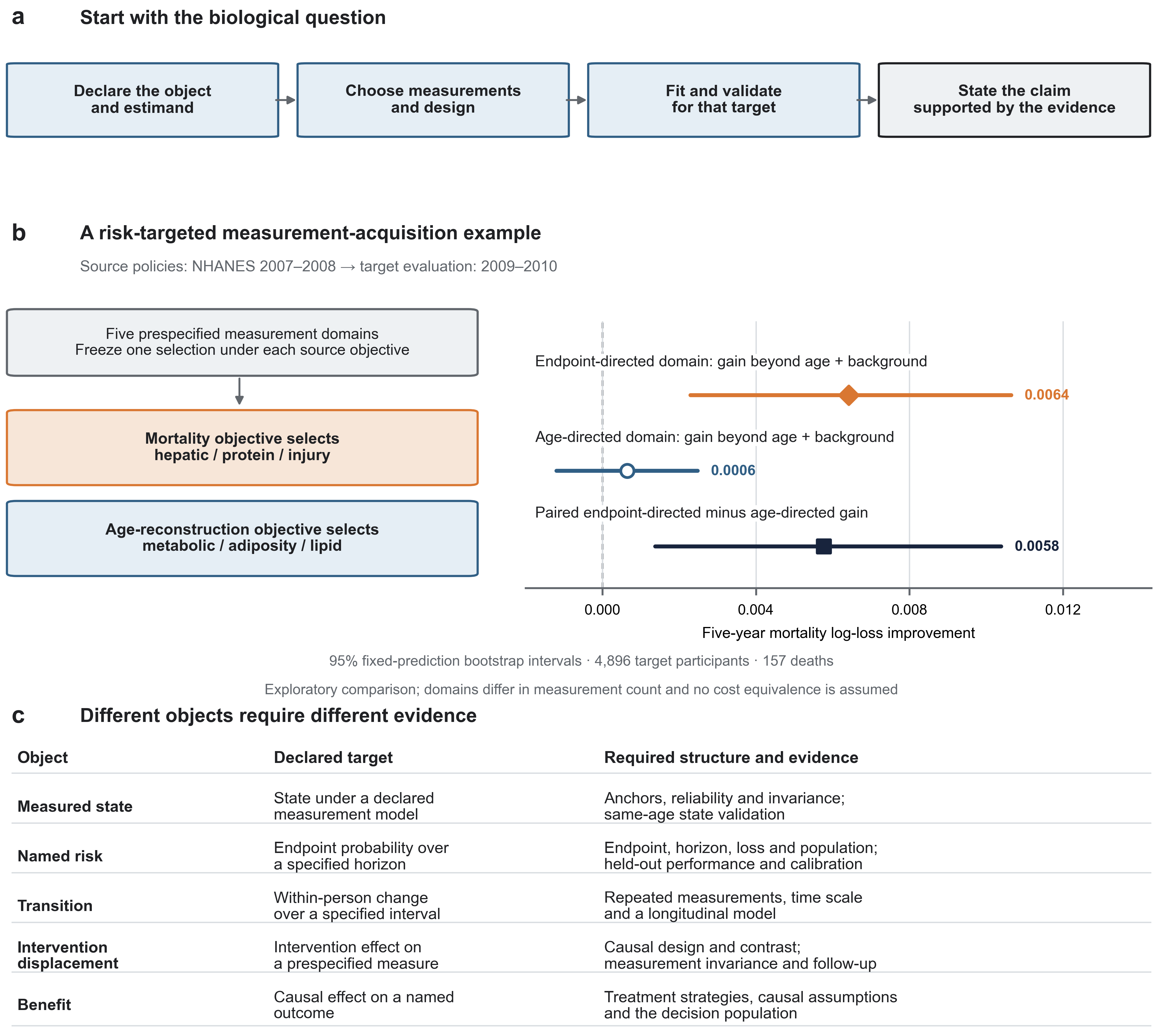

### Figure_4_target_first_measurement_and_evidence.png

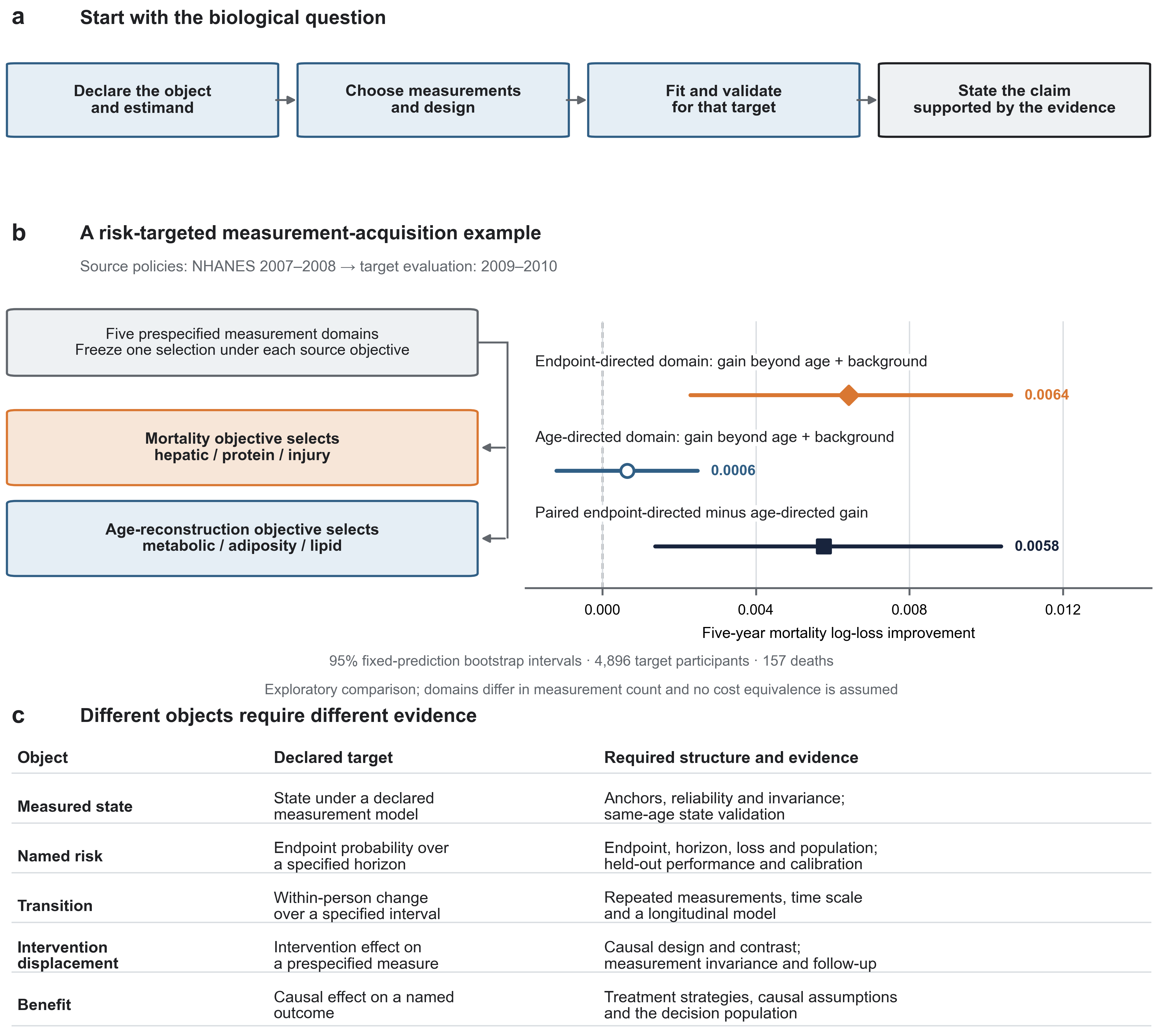
